# Upregulation of TNFR2 Precedes TOX Expression by Exhausted T cells and Restricts Antitumor and Antiviral Immunity

**DOI:** 10.1101/2024.07.12.603311

**Authors:** Alexandra Hoyt-Miggelbrink, Jessica Waibl Polania, Luke Wachsmuth, Selena Lorrey, Aditya Mohan, Andrew Hardigan, Emily Blandford, Emily Lerner, Daniel Wilkinson, Kelly M. Hotchkiss, Sarah Cook, Saskia Hemmers, Anoop Patel, Katayoun Ayasoufi, Peter Fecci

**Affiliations:** Duke Brain Tumor Immunotherapy Program, Duke University Medical Center, Durham, North Carolina; Department of Pathology, Duke University Medical Center, Durham, North Carolina; Duke University School of Medicine, Durham, North Carolina; Department of Integrative Immunobiology, Duke University Medical Center, Durham, North Carolina; Department of Neurosurgery, Duke University Medical Center, Durham, North Carolina; Department of Biomedical Engineering, Duke University Medical Center, Durham, North Carolina

## Abstract

Exhaustion represents a collection of programmed T cell differentiation states and an important mode of T cell dysfunction. T cell progression from progenitor to terminal exhaustion is associated with upregulation of the transcription factor TOX and expression of TIM3. Our understanding of factors regulating TOX expression and the transition from progenitor to terminal exhaustion, however, remains incomplete. We reveal here that T cell upregulation of tumor necrosis factor receptor type II (TNFR2) coincides with the gain of phenotypic markers and functions reflective of terminal exhaustion. Meanwhile, knocking out TNFR2 affords a novel population of T cells that express TIM3 but possess diminished TOX levels and functional characteristics of both progenitor and terminally exhausted cells. TIM3^+^ TNFR2 KO T cells exhibit reduced exhaustion transcriptional programs and enhanced AP1 pathway signatures. Finally, TNFR2 KO mice demonstrate improved T cell-dependent control of tumor and chronic lymphocytic choriomeningitis (cLCMV) infection, while pharmacologic antagonism of TNFR2 licenses responses to checkpoint blockade in multiple subcutaneous and intracranial tumor models.

## Introduction

T cell exhaustion is a known barrier to immunotherapeutic success^1,2^. The compromised function characterizing exhausted CD8^+^ T cells^3–6^ ultimately hinders their capacity to effect sustained tumor and virus control^7–10^. In addition to the loss of effector functions, exhausted T cells are marked by the upregulation of various co-inhibitory receptors, including programmed cell death protein 1 (PD-1)^11,12^ and cytotoxic T-lymphocyte associated protein 4 (CTLA4)^13^. Identification and targeting of these receptors spawned the advent of immune checkpoint inhibitors (ICI) that aim to restore T cell effector functions. While ICIs have been successful against a number of solid cancers, such as melanoma^14–16^, they have largely been ineffective against more notoriously treatment-resistant tumors, including glioblastoma (GBM)^17^. Particularly severe T cell exhaustion may be a significant contributor to treatment failures in these cancers^1,18^, creating a need to better understand the upstream regulators mediating exhaustion progression.

Multiple groups have identified two stages of exhaustion existing concomitantly amongst CD8^+^ T cells: progenitor T cell exhaustion (Tex_prog) and terminal T cell exhaustion (Tex_term)^19–21^. The Tex_prog population is characterized by expression of the transcription factor T cell factor 1 (TCF1) and the surface molecule SLAMF6. This population retains polyfunctional capacity, is self-renewing, and is responsive to ICI-treatment^19^. In contrast, Tex_term is marked by TIM3 on the cell surface. While this population exhibits a greater cytotoxic capacity, it loses polyfunctionality, the ability to self-renew, and proves unresponsive to ICI^19^. The transcription factor thymocyte selection-associated HMG BOX (TOX) is highly expressed in the Tex_term population and serves in its identification^22–24^. Expression of TOX has been shown to epigenetically reprogram exhausted CD8^+^ T cells, increasing their exhaustion signature and decreasing the effector program. Furthermore, overexpression of TOX is sufficient to skew CD8^+^ T cell differentiation towards exhaustion and away from effector and memory pathways^22^. While TOX appears to be required for CD8^+^ T cells to persist in a chronic antigen environment, reduced expression of TOX has been shown to increase anti-tumor effector functions^22^.

While it has been established that the NFAT^25^ and NR4A^24^ transcription factors are involved in the regulation of TOX expression, our understanding of the events leading to the upregulation of TOX remains incomplete. We hypothesized that tumor necrosis factor (TNF) might play a role in mediating the upregulation of TOX, as TNF has been shown previously to contribute to an exhausted T cell phenotype in various viral and parasitic infection models^26,27,28^. TNF signals through two receptors: TNF receptor 1 (TNFR1) and TNF receptor 2 (TNFR2). Whereas TNFR1 is expressed on most mammalian cells and induces a pro-inflammatory gene program, TNFR2 has a more restricted expression profile, including CD8^+^ T cells, and initiates immune-modulatory programming^29^. Therefore, we investigated the role of TNFR2 on CD8^+^ T cells in promoting progression of the exhaustion differentiation pathway.

We found that TNFR2, but not TNFR1, is expressed on CD8^+^ T cells in multiple murine models of cancer and chronic infection, including models of GBM, breast cancer, melanoma, and chronic lymphocytic choriomeningitis (cLCMV) infection. T cell expression levels of TNFR2 correlate with expression of canonical markers of exhaustion. Furthermore, TNFR2 upregulation heralds the loss of T cell function typically observed with exhaustion progression. Forced loss of TNFR2 from T cells uncouples TOX and TIM3 expression, creating a novel PD-1^+^TIM3^+^TOX^lo^ population that exhibits little of the exhaustion transcriptional program and possesses both enhanced cytotoxic capacity and polyfunctional features of Tex_prog. TNFR2 knockout (KO) mice display enhanced CD8^+^ effector function, decreased spleen weights following cLCMV infection, and improved CD8^+^ T cell-dependent tumor control. Finally, pharmacologic antagonism of TNFR2 synergizes with anti-PD1 in restricting tumor growth in multiple models. Our data place TNFR2 signaling as a potential upstream regulator of TOX expression in T cells and propose TNFR2 antagonism as a novel immunotherapeutic strategy.

## Results

### TNFR2 upregulation heralds the phenotypic transition of CD8^+^ T cells from progenitor to terminal exhaustion

Intrigued by the role TNF receptors might play in T cell exhaustion, we first investigated the expression patterns of TNFR1 and TNFR2 on T cells in various subcutaneous (sc) and intracranial (ic) models of breast cancer (E0771), glioblastoma (GBM) (CT2A), and melanoma (Yummer1.7) by flow cytometry **(Fig 1A-B** and **supplemental Fig 1A)**. Receptor levels were assessed on CD8^+^ T cells harvested at multiple time points from tumor, spleen, and tumor-draining lymph nodes (tdLN). While TNFR1 was generally not found on T cells from any tumor model, location, or time point **(Fig 1A)**, TNFR2 was highly expressed on CD8^+^ tumor-infiltrating lymphocytes (TIL) across models **(Fig 1B)**. Neither receptor demonstrated appreciable expression on T cells in the tdLN or the spleen of tumor-bearing (TB) mice **(Fig 1A-B)**. As TNFR2 was found highly expressed on CD8^+^ TILs from subcutaneously and intracranially implanted breast cancer, glioma, and melanoma models, its levels appeared to be independent of either tumor histology or anatomic compartment.

**Fig 1:**
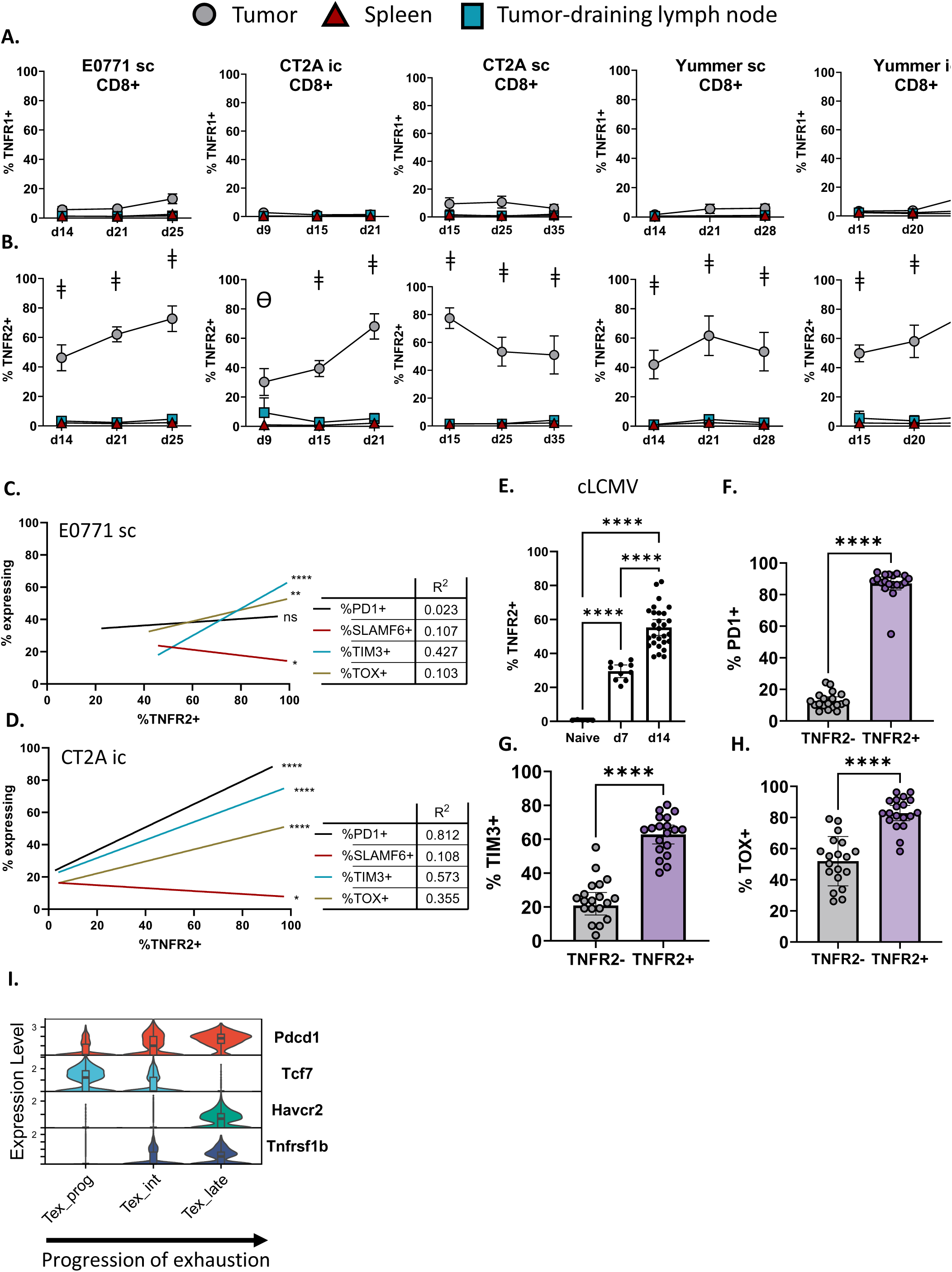
TNFR2 expression on CD8^+^ T cells is correlated with canonical markers of exhaustion. (A and B) TNFR1 (A) and TNFR2 (B) expression on CD8^+^ T cells harvested from the tumor, spleen, or tumor-draining lymph nodes from tumor-bearing mice over time, across tumor types and locations. N = 10-35 per time point/per tissue. ǂ represents p < 0.0001 between tumor and spleen and between tumor and tumor-draining lymph node. Ɵ represents p < 0.01 between tumor and tumor-draining lymph node and p < 0.0001 between tumor and spleen. (C and D) Correlation of TNFR2 expression on tumor-infiltrating CD8^+^ T cells with expression of canonical markers of exhaustion (PD1, SLAMF6, TIM3, and TOX) in subcutaneous E0771 (C) and intracranial CT2A (D). Tumors were harvested at similar timepoints as displayed in Figure 1A and B. A simple linear regression model was performed to determine significance and R^2^ values. (E) Expression of TNFR2 on CD8^+^ T cells in the spleen of mice infected with chronic LCMV at day 7 and 14 post-infection compared to naïve spleen. One-way ANOVA was performed. (F-H) Expression of PD1 (F), TIM3 (F), and TOX (H) on TNFR2- and TNFR2+ CD8^+^ T cells in the spleen of mice infected with chronic viral infection at peak viral titers (D14). Paired t-test was performed. (I) Single cell RNA sequencing was performed on CD3^+^ T cells sorted from intracranial CT2A at day 13 and day 19 post tumor implantation. Following QC, CD8^+^ T cells clustered into 6 populations. Trajectory analysis was performed on the clusters. Using the pathway identified, expression of *Pdcd1* (gene encoding PD1), *Tcf7* (gene encoding progenitor-associated transcription factor TCF1), *Havcr2* (gene encoding TIM3), and *Tnfrsf1b* (gene encoding TNFR2) was determined in the clusters Tex_prog, Tex_int, and Tex_late. Data represented as mean ± 95% confidence interval. ns not significant, * p < 0.05, ** p < 0.01, **** p < 0.0001

Given the above findings, we subsequently focused our attention on TNFR2. The predominant emergence of TNFR2 on T cells within the tumor microenvironment (TME) was reminiscent of the expression patterns of the canonical markers of Tex_term^18,19,30^. As TNF has a described role in promoting T cell exhaustion^26^, this prompted us to assess how TNFR2 levels may vary alongside those of PD-1 and TIM3. We used simple linear regression models to determine the correlation between TNFR2 and other exhaustion markers. Our analysis revealed a strong positive correlation between the expression of TNFR2 and the established terminal exhaustion markers PD-1, TIM3, and TOX ^22,24,31^ **(Fig 1C-D**, **supplemental Fig 1B-H**). We also examined the relationship with SLAMF6, which instead reflects the progenitor exhaustion (Tex_prog) differentiation state^19,32,33^ **(supplemental Fig 1H)**. While there was a significant negative correlation between SLAMF6 and TNFR2 levels in the sc breast cancer and ic GBM models **(Fig 1C-D)**, no apparent correlation was detected in the sc GBM and melanoma models **(supplemental Fig 1E-H)**. Interestingly, a slight positive correlation between SLAMF6 and TNFR2 was detected in the intracranial melanoma model **(supplemental Fig 1G)**.

To next examine whether TNFR2 expression in exhausted T cells was tumor-specific, we examined TNFR2 expression in a model of chronic viral infection. Similar to tumor-infiltrating CD8^+^ T cells, CD8^+^ T cells from the spleens of mice infected with chronic LCMV clone 13 (cLCMV) upregulated TNFR2 **(Fig 1E)**. At peak viral infection (D14)^9^, TNFR2^+^ CD8^+^ T cells expressed significantly higher levels of PD-1, TIM3, and TOX than their TNFR2^-^ counterparts **(Fig 1F-H)**. Conversely, SLAMF6 expression was significantly lower on splenic TNFR2^+^ CD8^+^ T cells **(supplementary Fig 1I)**.

To better understand the timing of TNFR2 upregulation on T cells along the progression from Tex_prog to Tex_term, we interrogated a scRNAseq dataset that we have previously established from murine CD8^+^ T cells infiltrating ic CT2A gliomas (Immunity in press)^34^. This dataset comprises of sorted CD8^+^ T cells from both an early (D13) and a late (D19) timepoint along tumor progression to evaluate the impact of time on exhaustion evolution. Ultimately, six phenotypic clusters were identified and subsequently annotated according to differentially expressed genes. Four of these clusters exhibited exhaustion phenotypes (Immunity in press)^34^. Pseudotime analysis was performed to infer temporal relationships between identified clusters. From this analysis, we established a trajectory from Tex_prog, through an intermediate population (Tex_int), and terminating in Tex_term (Tex_term clusters varied according to whether they were analyzed at the early or late timepoint) (Immunity in press)^34^. As expected, *Pdcd1* (gene encoding PD-1) expression increased along this trajectory and over time, while *Tcf7* (the transcription factor associated with SLAMF6 and Tex_prog) disappeared over the same period **(Fig 1I)**. *Havcr2* (encoding TIM3), in turn, appeared almost exclusively within Tex_term (late). Concomitantly, we examined the expression of *Tnfrsf1b* (encoding TNFR2) along the same trajectory. *Tnfrsf1b* first appeared within the Tex_int population, increasing further as *Tcf7* disappeared, but ultimately appearing prior to the acquisition of *Havcr2* **(Fig 1I)**. Thus, TNFR2 gene expression appears to arise as T cells are transitioning between Tex_prog and Tex_term.

### Acquisition of TNFR2 by T cells coincides with their loss of polyfunctionality (IFNγ^+^TNF^+^) and gain in cytotoxic capacity in the context of both tumor and chronic viral infection

These findings prompted us to examine the functional effects of TNFR2 upregulation on exhausted CD8^+^ T cells in models of both tumor and cLCMV infection. TNFR2^-^ and TNFR2^+^ CD8^+^ cells were sorted from E0771 (sc), CT2A (ic), and Yummer (sc, ic) tumors, and from spleens from cLCMV-infected mice. To ensure only antigen-experienced T cells were compared in subsequent functional assays, all sorted T cells were PD-1^+^ (i.e. CD8^+^PD-1^+^TNFR2^+^ and CD8^+^PD-1^+^TNFR2^-^) **(Fig 2A)**. Sorted samples were stimulated with PMA and ionomycin and stained intracellularly for TNF, IFNγ, interleukin 2 (IL-2), and granzyme B **(Fig 2B)**. Similar to T cells transitioning away from a progenitor exhausted state, TNFR2^+^CD8^+^ T cells from nearly all models displayed significantly lower polyfunctional capacity (IFNγ^+^TNF^+^, as described by Miller et al.^19^) **(Fig 2C)** and produced significantly less IL-2 **(Fig 2D)**. By contrast, these cells displayed functions more representative of Tex_term^19,35,36^, including mono-production of IFNγ without concomitant TNF **(Fig 2E)** as well as increased cytotoxic capacity (Granzyme B production) **(Fig 2F)**.

**Fig 2:**
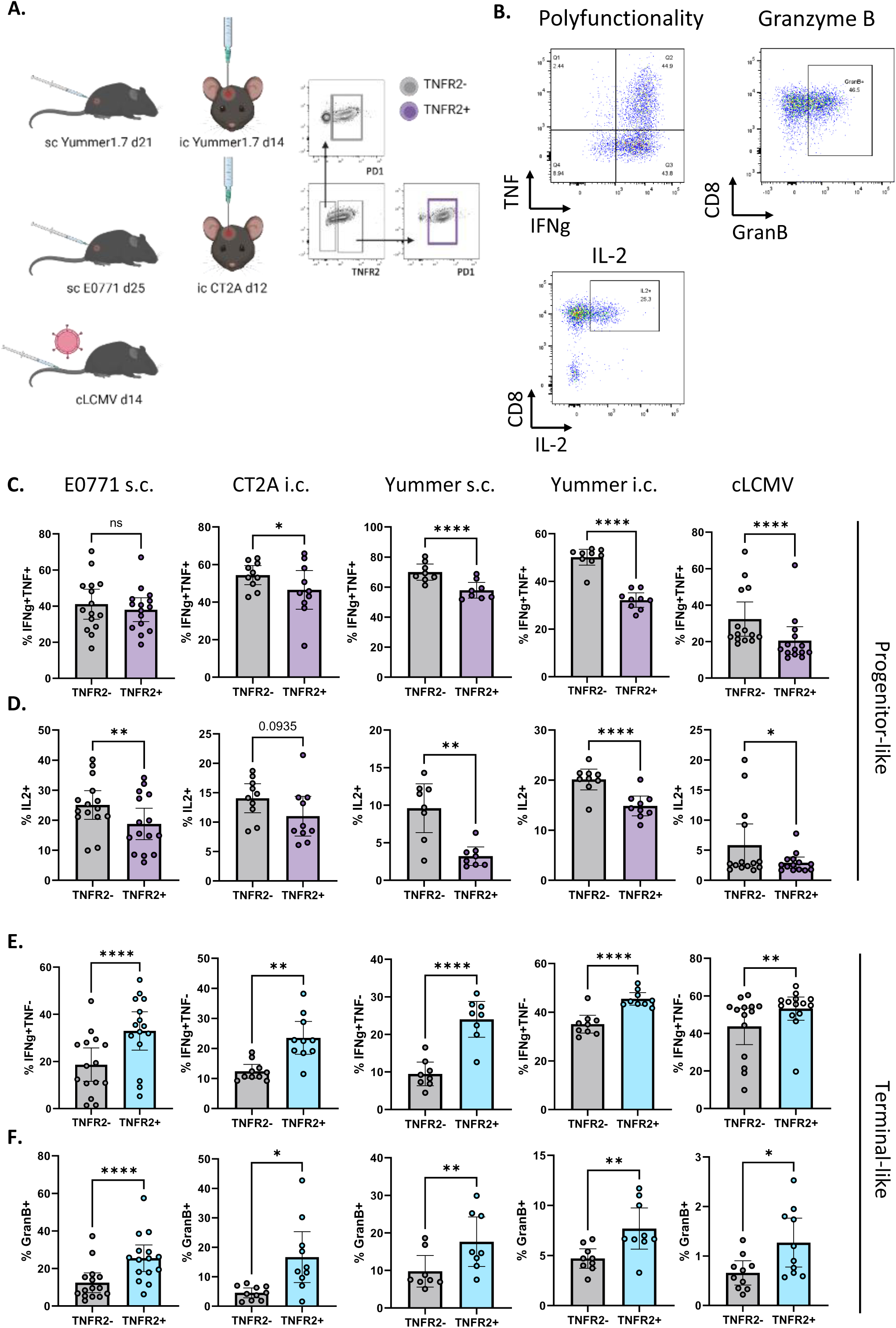
Upregulation of TNFR2 is associated with functional changes resembling a more terminally exhausted phenotype. (A) Experimental design to assess cytokine production in antigen-experienced (PD1^+^) TNFR2^-^ and TNFR2^+^ CD8^+^ T cells. Each tumor was harvested at a time point that produced sufficient TNFR2^-^ and TNFR2^+^ CD8^+^ T cells to allow sorting of each population. (B) Representative flow plots of IFNγ^+^ TNF^+^ (polyfunctional), IL-2, and granzyme B production of PD1^+^ CD8^+^ T cells. (C-D) Assessment of the progenitor-like functions polyfunctionality (IFNγ^+^ TNF^+^) (C) and IL-2 expression (D) in TNFR2^-^ and TNFR2^+^ CD8^+^ T cells harvested from tumors at time points displayed in (A) following stimulation with PMA, Ionomycin, GolgiStop, and GolgiPlug for 4 hours at 37°C. Paired t-test was performed. (E-F) Assessment of the terminal-like functions IFNγ^+^ single positive (TNF^-^) (E) and granzyme B (F) in TNFR2^-^ and TNFR2^+^ CD8^+^ T cells harvested from tumors at time points displayed in (A) following stimulation with PMA, Ionomycin, GolgiStop, and GolgiPlug for 4 hours at 37°C. Paired t-test was performed. Data represented as mean ± 95% confidence interval. ns not significant, * p < 0.05, ** p < 0.01, **** p < 0.0001

### TNFR2 deficiency leads to diminished TOX levels in chronically stimulated PD1^+^TIM3^+^ T cells

Given the association between TNFR2 upregulation and emergence of the Tex_term phenotype, we investigated the direct impact of TNFR2 loss on T cell exhaustion in tumor and cLCMV models using a TNFR2 knockout (KO) mouse^37^. Importantly, to begin, loss of TNFR2 did not result in a compensatory upregulation of TNFR1 on CD8^+^ T cells in any setting **(supplementary Fig 2A)**. The frequencies of memory precursor effector (MPEC), short-lived effector (SLEC), effector, and central memory T cell populations were also similar between wild type (WT) and TNFR2 KO mice across all models tested **(supplementary Fig 2B-E)**.

With TNFR2 loss demonstrating minimal impact on T cell memory differentiation states, we next turned to its influence on T cell differentiation along the exhaustion pathway. Once again examining various models of tumor and of LCMV, we found no consistent impact of TNFR2 KO on the frequency of TIM3 (or SLAMF6) expression on CD8^+^PD1^+^ T cells, and thus no consistent impact on the apparent phenotypic progenitor exhaustion : terminal exhaustion ratio (PETER)^34^. **(Fig 3A, supplementary Fig 3A, 3B)**. Consistent declines were seen, however, in expression levels of TOX within the PD-1^+^ TIM3^+^ (terminally exhausted) population **(Fig 3B)**. Amidst all models tested (tumors and cLCMV), global loss of TNFR2 resulted in a significant decrease of TOX expression in CD8^+^ T cells **(Fig 3C)**, without a concomitant decrease in the number that were PD-1^+^TIM3^+^ **(Fig 3A** and **supplementary Fig 3A)**. This led to the emergence of a novel population of CD8^+^PD-1^+^TIM3^+^TOX^lo^ Tex cells. Such a population is unique, as CD8^+^PD-1^+^TIM3^+^ Tex_term cells are typically TOX^hi^, given the established role of TOX in mediating exhaustion progression^22^. Therefore, we investigated whether it was TNFR2 loss in T cells, specifically, that led to the concomitant decline in TOX expression.

**Fig 3:**
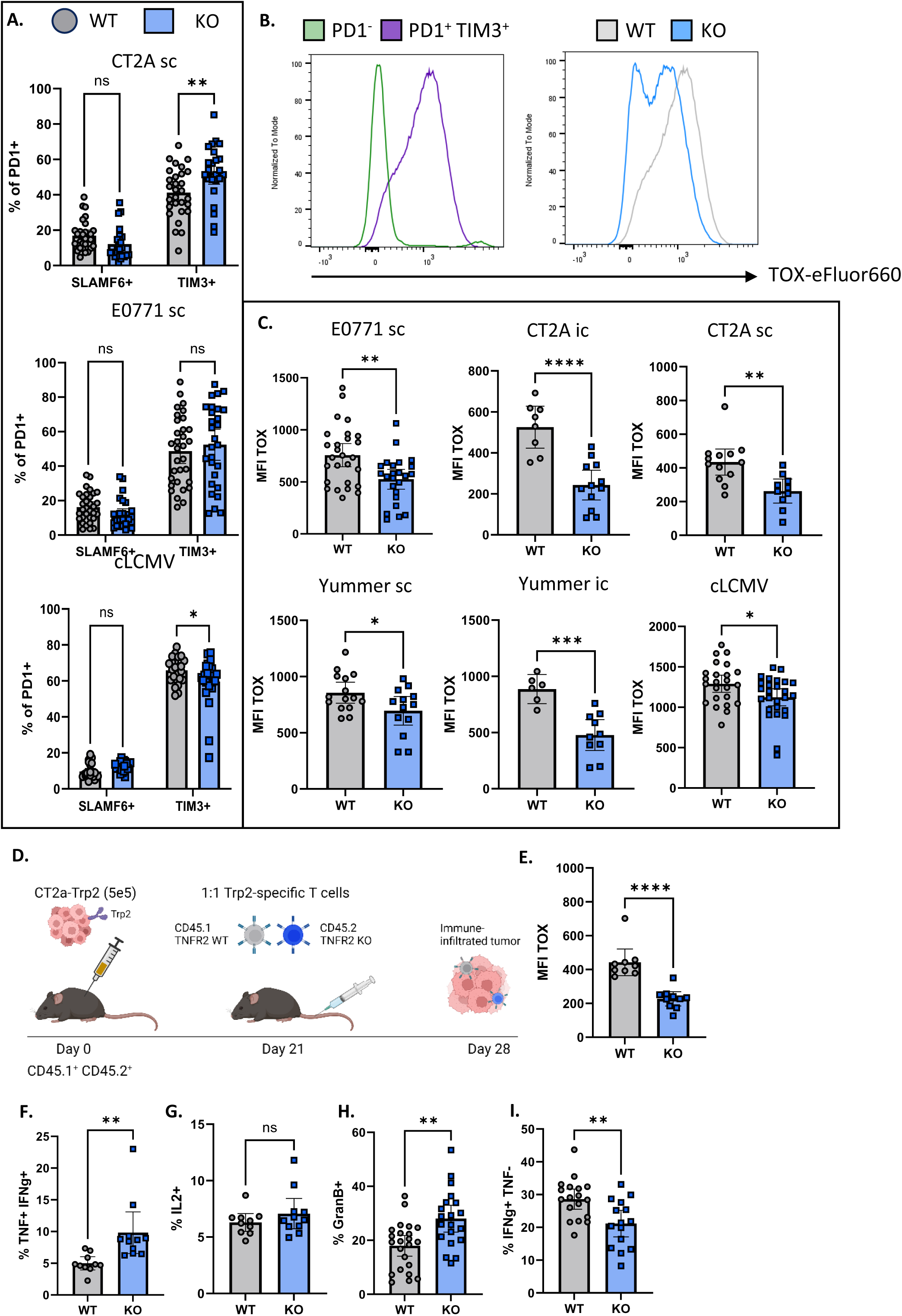
Loss of TNFR2 results in decreased TOX expression without a concomitant decrease in TIM3. (A) Frequency of progenitor (SLAMF6^+^TIM3^-^) and terminally exhausted (SLAMF6^-^TIM3^+^) CD8^+^ TILs from wild type (WT) and TNFR2 knock out (KO) in subcutaneous models of CT2A (D35, top), E0771 (D25, middle), and cLCMV (D14, bottom). Two-way repeated measures ANOVA was performed. (B) Representative flow plot of TOX expression in PD1^-^ (green) and PD1^+^TIM3^+^ (purple) in CD8^+^ TILs (left) and a histogram comparing TOX expression in WT (gray) and TNFR2 KO (blue) in PD1^+^TIM3^+^ CD8^+^ TILs (right) (C) Mean fluorescence intensity (MFI) of TOX in WT and TNFR2 KO TIM3^+^ CD8^+^ T cells harvested from tumor for sc E0771 (D25), ic CT2A (D21), sc CT2A (D35), sc Yummer (D28), ic Yummer (D25), and spleen for cLCMV (D14). Unpaired t-test was performed. (D-E) Experimental design to assess T cell intrinsic requirement of TNFR2 (D). CT2A-Trp2 was implanted subcutaneously into the flank of CD45.1^+^CD45.2^+^ mice. After 21 days, Trp2-specific T cells generated from WT (CD45.1) and TNFR2 KO (CD45.2) mice were injected intravenously at a 1:1 ratio. Tumors were harvested on day 28 and transferred Trp2-specific WT (CD45.1) and TNFR2 KO (CD45.2) T cells were assessed for TOX expression (E). Paired t-test was performed. (F-I) Polyfunctionality (IFNγ^+^ TNF^+^) (F), IL-2 (G), Granzyme B (H), and IFNγ^+^ single positive (I) was assessed on CD8^+^ T cells from splenocytes of WT and TNFR2 KO mice following chronic LCMV infection (D14) following stimulation with PMA, Ionomycin, GolgiStop, and GolgiPlug for 4 hours at 37°C. Unpaired t-test was performed. Data represented as mean ± 95% confidence interval. ns not significant, * p < 0.05, ** p < 0.01, **** p < 0.0001

To this end, CT2A gliomas previously engineered to express the tumor-associated antigen TRP2 (CT2A- TRP2) were implanted into the flanks of WT mice. On Day 21, a mixture of CD45.1 WT and CD45.2 TNFR2 KO TRP2 T cell receptor-transgenic (TRP2 TCR-tg) T cells were adoptively transferred into tumor-bearing mice at a 1:1 ratio **(Fig 3D)**. WT and KO T cells transduced at a comparable efficiency and were PD-1^-^ at the time of transfer **(supplementary Fig 3C-D)**. After 7 days, similar numbers of CD45.1 and CD45.2 transferred T cells could be recovered from the tumor **(supplementary Fig 3E-F)**, suggesting that TNFR2 KO T cells maintain normal trafficking functions. However, TNFR2 KO T cells recovered from tumor harbored significantly lower levels of TOX than their WT counterparts **(Fig 3E)**, suggesting that TNFR2 loss from T cells is indeed sufficient to mediate the accompanying decline in TOX.

As decreased expression of TOX is typically associated with improved T cell function^22^, we evaluated the functional capacities of TNFR2 KO CD8^+^ T cells. Cytokine expression profiles were examined in CD8^+^ T cells in the spleen from WT and TNFR2 KO mice in the cLCMV model. T cells were harvested at a time point corresponding to peak viral titers (D14). TNFR2 loss resulted in increased polyfunctional capacity **(Fig 3F)** reflective of a Tox^lo^ Tex_prog-type functional profile. In contrast, however, IL-2 production was not affected in mice lacking TNFR2 **(Fig 3G)**. Intriguingly, however, loss of TNFR2 also resulted in increased granzyme B expression **(Fig 3H),** but a decline in the frequency of IFNγ mono-producing cells **(Fig 3I)**. The net effect then, was a gain in functions reflecting typical features of both Tex_prog and Tex_term populations. Most notable was the simultaneous and atypical gain in both polyfunctionality and cytotoxicity, despite the expression of TIM3. While the above was observed in splenocytes from cLCMV-infected mice, similar findings were obtained across WT and TNFR2 KO CD8^+^ T cells harvested from tumor-bearing mice (sc CT2A and E0771). **(supplementary Fig 4A-E)**.

### TIM3+ TNFR2 KO T cells exhibit diminished exhaustion transcriptional programs and enhanced AP1 pathway signatures

To better understand the transcriptional profiles of TNFR2 KO T cells, we sorted PD-1^+^TIM3^+^CD8^+^ T cells from WT and TNFR2 KO LCMV-infected mice (D14) and performed bulk RNA-sequencing analyses on 6 WT and 6 KO samples **(Fig 4A)**. Principal component analysis showed clustering of genotypes, indicating a segregation of WT and TNFR2 KO samples **(Fig 4B)**. A total of 6584 genes were differentially expressed between WT and TNFR2 KO TIM3^+^ CD8^+^ T cells, of which 3010 were upregulated in the TNFR2 KO samples **(supplementary table 1)**. Among notable downregulated genes were various exhaustion-associated genes, including *Tox*, *Tox2*, *Tox4*, *Tbx21*, *Foxo1*, *Nfatc1*, *Nfatc2*, *Nfatc3*, and *Nfat5* **(Fig 4C)**. Interestingly, AP1 transcription factors normally associated with an effector phenotype were among the upregulated genes, including *Akt1*, *Jun*, and *Fos*^25^. Genes associated with memory and effector programs, such as *Batf3*^38^ and *Stat5a*^39^, were also upregulated in TNFR2 KO T cells, as were genes associated with increased cytolytic effector function, including *Klf4*^40^**(Fig 4C)**. To assess the transcriptional network in a more unbiased manner, we performed pathway analyses using the gene expression modules enlisted by Nanostring^41^ **(supplementary table 2)**. A significant decrease in pathways associated with exhaustion, immune checkpoints, PD-1 signaling, NF-κB signaling, and TNF signaling was observed for TNFR2 KO T cells **(Fig 4D)**. As TNFR2 signaling is upstream of the NF-κB pathway, we investigated genes related to NF-κB signaling: Most genes associated with this pathway were significantly downregulated, while *Nfkbia*, an inhibitor of NF-κB activation, was significantly increased in TNFR2 KO T cells **(supplementary Fig 5A)**.

**Fig 4:**
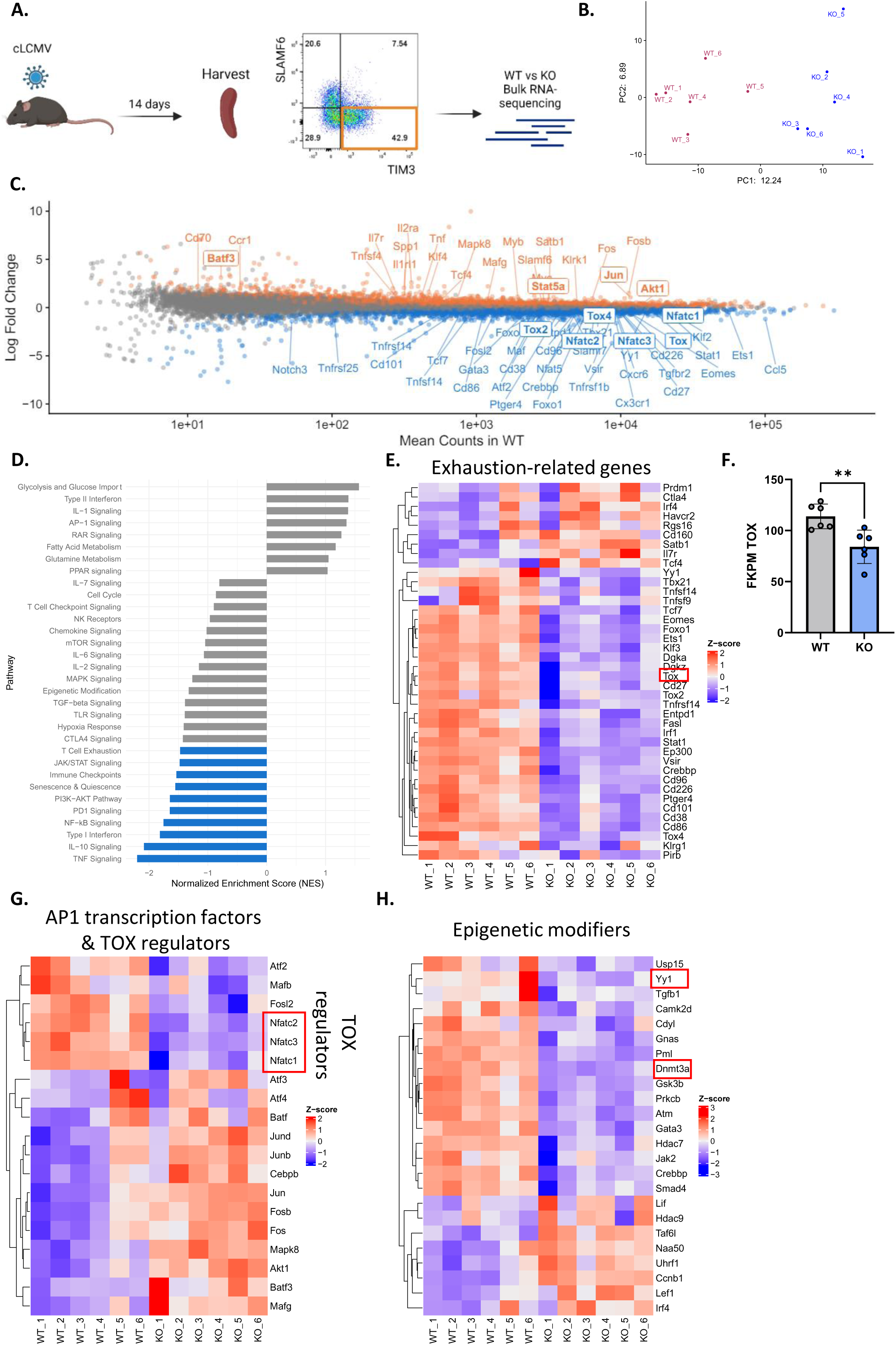
TNFR2 KO T cells display a diminished transcriptional exhaustion program. (A) Experimental design to assess transcriptional program of WT and TNFR2 KO CD8^+^ TIM3^+^ T cells. CD8^+^ PD1^+^ SLAMF6^-^ TIM3^+^ splenocytes were sorted from WT and TNFR2 KO mice 14 days post infection with cLCMV. (B) PCA plot representing distribution of WT and TNFR2 KO CD8^+^ TIM3^+^ T cells based on gene expression profile. (C) MA-plot highlighting significantly downregulated (blue) and significantly upregulated genes in TNFR2 KO CD8^+^ TIM3^+^ T cells. (D) Gene module enrichment analysis was performed based on the mouse NanoString nCounter^®^ Immune Exhaustion Panel **(genes listed in supplementary table 2)**. Pathways highlighted in blue were statistically downregulated using a corrected p-value <0.1. (E) Heatmaps displaying z-scores for exhausted-related genes. (F) Gene expression levels for *Tox* in WT and TNFR2 KO CD8^+^ TIM3^+^ T cells. Unpaired t-test was performed. (G-H) Heatmaps displaying z-scores for AP1-transcription factors and TOX regulators (G), and epigenetic modifiers (H) for each WT and TNFR2 KO CD8^+^ TIM3^+^ sample. * p < 0.05, ** p < 0.01

The downregulation of exhaustion and immune checkpoint pathways led us to examine the expression patterns of specific genes within these clusters. Examination of the genes downstream of immune checkpoint signaling and anergy pathways revealed a divergent pattern between WT and TNFR2 KO T cells **(supplementary Fig 5B-C)**. Within the exhaustion-related transcripts, some genes, including *Ctla4* and *Prdm1*, were upregulated in TNFR2 KO T cells. However, predominantly a significant downregulation of exhaustion-related genes, including *Eomes*, *Tbx21*, *Foxo1*, *Vsir*, and *Ptger4,* was more generally observed. Furthermore, affirming the downregulation of TOX seen earlier at the protein level in TNFR2 KO mice **(Fig 3C)**, transcript levels for *Tox* was also significantly lower **(Fig 4E-F)**.

Given the reduction in TOX expression **(Fig 4C)**, we next focused on genes associated with TOX upregulation. Nuclear factor of activated T cells (NFAT) transcription factors have been established as being directly upstream of TOX^23^. Traditionally, expression of NFAT in conjunction with AP1 transcription factors is thought to result in T cell effector differentiation, whereas NFAT expression in the absence of AP1 transcription factors leads to an exhaustion phenotype^25^. Interestingly, TNFR2 KO T cells displayed decreased expression of the *Nfatc1*, *Nfatc2*, and *Nfact3* genes and increased expression of AP1 transcription factors, including *Jund*, *Junb*, *Jun*, *Fosb*, *Fos*, *Akt1*, and *Batf3* **(Fig 4G)**. In contrast, AP1 transcription factors in the *Maf* family were mostly downregulated in the TNFR2 KO T cells **(Fig 4G)**. Considering the important role of TOX in the epigenetic reprogramming of exhausted T cells^22^, we subsequently investigated various epigenetic regulators. TNFR2 KO T cells displayed a vastly different epigenetic regulator profile than WT T cells **(Fig 4H)**. Some prominent downregulated epigenetic regulators, such as *Dnmt3a* and *Yy1*, have been directly associated with T cell exhaustion^42^ and upregulation of co-inhibitory markers^43^, respectively.

### TNFR2 deficiency elicits improved CD8^+^ T cell-dependent control of both tumor and chronic viral infection

Given the changes observed in TNFR2 KO T cells, we investigated the role of TNFR2 in restricting anti-tumor immunity. WT or TNFR2 KO mice were implanted in the flank with E0771 or CT2A tumors. A significant decrease in tumor growth rate and tumor burden was observed in the TNFR2 KO animals implanted with both E0771 **(Fig 5A and supplementary Fig 6A)** and CT2A **(Fig 5B and supplementary 6B)**. Interestingly, tumor growth inhibition was not observed in the sc melanoma model **(supplementary Fig 6C)**. Tumor control against sc CT2A in TNFR2 KO mice was entirely abolished by CD8 depletion **(Fig 5C-D and supplementary Fig 6D)**, suggesting that TNFR2 may play a role in limiting the antitumor immune response. We subsequently investigated whether TNFR2 KO mice exhibited an enhanced ability to control cLCMV infection. TNFR2 KO mice demonstrated significantly lower spleen weights at D14 than WT infected mice **(supplementary Fig 6E)**, as well as overall trends toward decreased viral titers **(supplementary Fig 6F)**.

**Fig 5:**
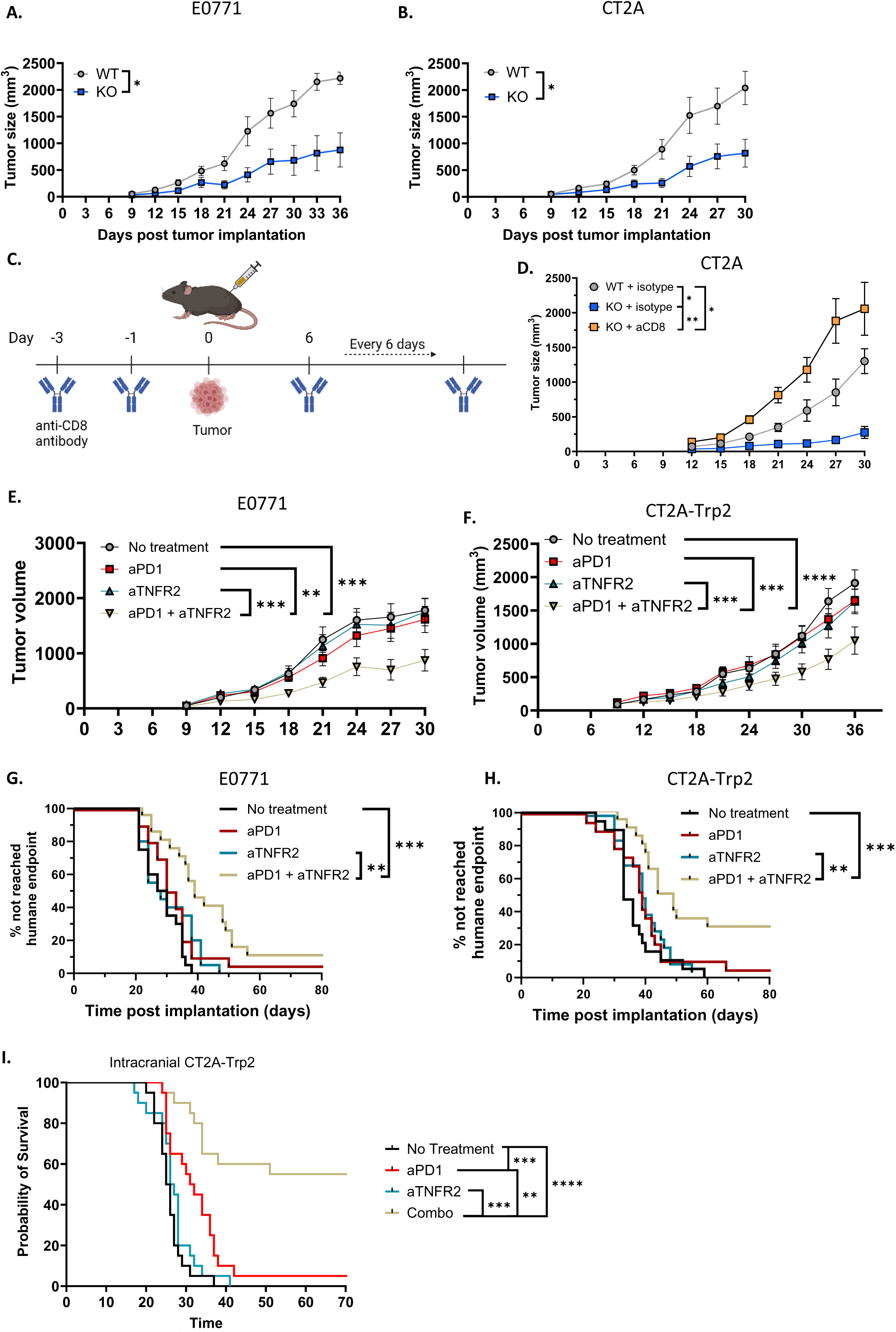
TNFR2 deficiency improves CD8^+^ T cell-dependent tumor control. (A-B) Tumor growth curves for WT and TNFR2 KO mice following subcutaneous implantation of E0771 (A) and CT2A (B). Tumors were measured every 3 days starting on day 9. Two-way repeated ANOVA was performed. (C) Experimental design to determine the role of CD8 T cell depletion on tumor control in TNFR2 KO mice. CD8 T cells were depleted with a depletion antibody day -3, day -1, and every 6 days post tumor implantation. CT2A was implanted subcutaneously on day 0. (D) Tumor growth curves for WT, TNFR2 KO, and CD8-depleted TNFR2 KO mice following subcutaneous implantation of CT2A. Tumors were measured every 3 days starting on day 9. Two-way repeated ANOVA was performed. (E-F) Tumor growth for WT mice dosed with anti-PD1, anti-TNFR2, or a combination of anti-PD1 and anti-TNFR2 following subcutaneous implantation of E0771 (E) or CT2A-Trp2 (F). Mice were injected with anti-PD1 (200ug), anti-TNFR2 (250ug), or both every 3 days starting on day 9. Tumors were measured every 3 days. Two-way repeated ANOVA was performed. N=20 per group. (G-H) Time to humane endpoint for mice displayed in E and F following subcutaneous implantation of E0771 (G) and CT2A-Trp2 (H). Log-rank test was performed between each group for a total of 6 comparisons. A corrected p-value of less than 0.0083 (p < 0.05/6 comparisons) was used as a cutoff for significance. Mice were followed for 80 days. If no tumor was detected at day 80, the mouse was considered a long-term survivor. N=19-20 per group. (I) Survival of mice post intracranial implantation of CT2A-Trp2. Mice were treated with 5 doses of anti-PD1, anti-TNFR2, or a combination (aPD1 + aTNFR2) starting at day 9. . A corrected p-value of less than 0.0083 (p < 0.05/6 comparisons) was used as a cutoff for significance. N=20 per group. Data represented as mean ± SEM p < 0.05, ** p < 0.01, *** p<0.001, **** p < 0.0001

Given the favorable results seen in sc tumors, we evaluated a more translatable approach using a TNFR2 antagonist. Treatment with a TNFR2 antagonist alone was not sufficient to decrease tumor growth in either CT2A-TRP2 or E0771 **(Fig 5E-F)**. However, the combination of TNFR2 antagonism and PD-1-blockade significantly reduced tumor volume **(Fig 5E-F)** and tumor burden **(Supplementary Fig 6G-H)** in both models. Six (6) out of 20 CT2A-TRP2-bearing mice dosed with the combination treatment achieved long-term survival **(Fig 5G-H)**. These survival benefits extended to an intracranial model of GBM, where 60% long-term survival was achieved in the combination treatment group **(Fig 5I)**. Ultimately, the combination of TNFR2 antagonism and PD-1-blockade either restricted tumor growth or extended survival regardless of the tumor model or compartment tested **(Supplementary Fig 6I)**.

## Discussion

T cell exhaustion plays a pivotal role in our understanding of immune responses to both chronic viral infection and solid tumors. With regard to the latter, the success of immunotherapeutic approaches, such as checkpoint blockade, has been recently tied to the persistence and/or the refurbishing of Tex_prog within tumors^44–47^, while response failures have instead been linked to a loss of T cell “stemness” and a predominant shift toward Tex_term^1^, marked often by the upregulation of both TOX and TIM3^19^. Increasingly, TOX has been identified as a critical transcription factor for establishing the unique epigenetic landscape observed amongst exhausted T cells^22,23^. A deeper understanding of the mechanisms underlying the transition between progenitor and terminal T cell exhaustion, as well as of the factors associated with the upregulation of TOX within T cells, has therefore become crucial to improving immunotherapeutic responses. Herein, we uncover that the TNFR2 receptor appears to play roles upstream of TOX in shepherding T cells from Tex_prog to Tex_term. TNFR2 emerges on T cells following the loss of SLAMF6 but before TIM3 appears during the transition to Tex_term. Its upregulation coincides with the gain of Tex_term phenotypic markers and functions, while knocking out of the receptor affords a novel population of cells that express TIM3 but have diminished TOX levels and functional characteristics of both Tex_prog and Tex_term. TIM3^+^ TNFR2 KO T cells exhibit diminished exhaustion transcriptional programs and enhanced AP1 pathway signatures. Furthermore, TNFR2 KO mice demonstrate improved T cell-dependent control of tumor and chronic infection with LCMV, while pharmacologic antagonism of TNFR2 licenses responses to checkpoint blockade in multiple peripheral tumor models.

Our investigation into TNFR2 followed from prior work suggesting roles for TNF in advancing T cell exhaustion. Exposure to TNF, for instance, is known to increase PD1 expression on T cells following viral infections. Furthermore, neutralization of TNF in LCMV is sufficient to increase the number and function of antigen-specific T cells, leading to diminished viral titers^26^. TNF signaling may occur through either TNFR1 or TNFR2. Whereas TNFR1 signaling is thought mainly to result in the induction of NF-κB and a pro-inflammatory signature, TNFR2 may elicit either pro-inflammatory or anti-inflammatory functions^48,49^. Cytotoxicity, however, is linked exclusively to activation of TNFR1^50^. TNFR2 agonists have demonstrated efficacy against autoimmune diseases supporting an anti-inflammatory role for TNFR2^51^. The pro- exhaustive functions of TNF and the seemingly immune-restrictive sequelae of TNFR2 signaling led us to examine the role of TNFR2 specifically in mediating the progression of T cell exhaustion in both viral infection and tumors. Our data revealed a selective upregulation of TNFR2 in CD8^+^ T cells in tumor microenvironments and in the spleens of mice infected with cLCMV, in a manner mimicking the expression patterns of canonical exhaustion markers^18,19,30^.

A curious finding was that T cells from TNFR2 KO mice appear to possess functional properties reflective of both progenitor and terminally exhausted T cells. In particular, the maintenance of both polyfunctionality (suggestive of Tex_prog) and cytotoxicity (suggestive of Tex_term), as well as the decreased levels of TOX observed, might imply the absence of exhaustion and more of an effector / memory phenotype. These T cells, however, still express high levels of PD-1 and TIM-3. Furthermore, the absence of traditional memory markers, such as KLRG1 and CD127 (reflecting SLECs and MPECs, respectively^52,53^), do not support a memory differentiation capacity for TNFR2 KO CD8^+^ T cells. Accordingly, chronic antigen stimulation, as these T cells experienced, is generally thought to prevent the acquisition of a memory phenotype capable of surviving without continuous antigen stimulation^54–58^. We therefore still assigned an “exhausted” label to CD8^+^PD1^+^TOX^lo^TIM3^+^TNFR2 KO T cells, even if their function and phenotype were somewhat inconclusive. As such, we conclude that TNFR2 signaling is not required for cells to acquire a terminally exhausted “phenotype.”

However, we did indeed observe a significant decrease in the expression of TOX in the “terminally exhausted” (TIM3^+^) population of TNFR2 KO T cells found amidst both tumors and chronic LCMV infection. While TOX has been demonstrated to be both sufficient and necessary to drive the terminal exhaustion pathway^22^, these data suggest that TOX and TIM3 are differentially regulated and that lack of TNFR2 signaling uncouples their expression. Furthermore, as TNF signaling is known to regulate the translocation of NFAT to the nucleus^28^ and NFAT is a known inducer of TOX^23,24,59^, this suggests that TNFR2 signaling is upstream of TOX, but not of TIM3. We have thus described a novel population, one that expresses high levels of TIM3, but that does not exhibit concomitantly high TOX expression.

Interestingly, TOX is not the only exhaustion-associated transcription factor whose expression waned in TNFR2 KO T cells. Amidst TIM3^+^ T cells from cLCMV-infected TNFR2 KO mice, we also observed significant downregulation of *Eomes*^60^ and *Entpd1* (encodes CD39)^61^, among others. Conversely, transcription factors associated with roles countering exhaustion, including *Irf4* and *Batf3*^62^, were significantly upregulated in TNFR2 KO TIM3^+^ T cells. Also diminished was the expression of genes directly upstream of the TOX program: *Nfatc1*, *Nfactc2*, and *Nfactc3*. This suggests that TNFR2 signaling is upstream of the currently established TOX-initiated exhaustion program. Additionally, TNFR2 KO TIM3^+^ T cells displayed significant upregulation of various AP1 transcription factors, including family members of *Jun* and *Fos*. NFAT expression alongside that of AP1 transcription factors is known to elicit a T cell “effector” program, whereas lack of AP1 transcription factors precipitates an exhaustion phenotype^25^. Therefore, our data also suggest that TNFR2 KO T cells possess an “effector-like phenotype.” This is further reflected in the increased levels of *Stat5a* we observed, as *Stat5a* has recently been associated with a durable effector state^39^.

As TOX is involved in the epigenetic reprogramming of exhausted T cells, we also evaluated epigenetic modifiers in TNFR2 KO T cells. Distinct epigenetic regulators were utilized by WT and TNFR2 KO T cells. TNFR2 KO T cells displayed a significant downregulation of various exhaustion-inducing genes, including *Yy1*^43^, *Dnmt3a*^42^, and *Gsk3b*^63^. Furthermore, TNFR2 KO T cells displayed significant upregulation of *Lef1*, a gene known to drive central memory programs in NKT cells^64^. A more direct assessment of the epigenetic landscape will be required to determine whether loss of TNFR2 is sufficient to reverse or prevent epigenetic changes commonly associated with TOX^22^. Finally, as the epigenetic stability of exhausted CD8^+^ T cells has been described as a key contributor to diminished long-term efficacy with PD-1 blockade^2^, TNFR2 antagonism might provide a novel strategy to increase the success of ICI. Further support for this notion is found in our observation of improved tumor control in mice lacking TNFR2, as well of signs of improved viral control in mice infected with cLCMV. More direct evidence yet is seen with the improved tumor control we obtained with the combination of a TNFR2 antagonist and anti-PD1. While the observed benefits proved to be CD8-dependent, future investigations into the detailed mechanisms of tumor control afforded by TNFR2 antagonism are warranted.

## Funding sources

Dr. Fecci is funded by a Cancer Research Institute (CRI) Arash Ferdowsi Lloyd J. Old STAR Award.

## Conflict of interest

The authors declare to have no conflicts of interest.

## Methods

### Animals

Wild-type C57BL/6J (Jackson Laboratory, cat #000664), B6.129S2-Tnfrsf1btm1Mwm/J (TNFR2 KO, Jackson Laboratory, cat # 002620), and B6.SJL-Ptprca Pepcb/BoyJ (CD45.1, Jackson Laboratory, cat# 002014) were used between 7 and 12 weeks of age. CD45.1 and C57BL/6J were crossed to generate recipients for adoptive transfer experiments (CD45.1 CD45.2 double-positive). All experiments were gender-matched and age-matched. As we observed significant tumor rejection in female mice, only male mice were used for experiments with the Yummer1.7 cell line. Mice were maintained under specific-free conditions and were bred in our animal facility at the Cancer Center Isolation Facility at the Duke University Medical Center. All experimental procedures were approved by the Institutional Animal Care and Use Committee.

### Cell lines and tumor implantation

All cell lines are syngeneic in C57BL/6 mice and were cultured at 37°C at 5% CO2. Cells were negative for Mycoplasma spp. The murine CT2A malignant glioma and Yummer1.7 melanoma cell lines were kindly provided by Robert L. Martuza (Massachusetts General Hospital, MA, USA) (47) and Kris Woods (Duke University, NC, USA), respectively. E0771 was purchased from ATCC (cat # CRL-3461). CT2A-TRP2 was generated by overexpressing a TRP2-containing lentiviral plasmid. CT2A and CT2A-Trp2 were expanded in vitro in DMEM containing 2mM 1-glutamine and 4.5mg/mL glucose (Sigma Aldrich, cat # D6429), supplemented with 10% heat-inactivated FBS (Sigma Aldrich, cat # F4135). E0771 was cultured in DMEM containing 2mM 1-glutamine and 4.5mg/mL glucose, supplemented with 10% heat-inactivated FBS, and Hepes (Gibco, cat # 15630-080). Yummer1.7 was cultured in F12 media (Gibco, cat # 11330-032), supplemented with non-essential amino acids (Gibco, cat # 11140-050) and 10% FBS. All cell lines were passaged at least one time after thawing prior to harvesting the cells in the logarithmic growth phase for implantation. Tumors were harvested at various time points as indicated per figure.

For intracranial injections, cells were harvested and resuspended in PBS (Sigma Aldrich, cat # D8537). Cells were then mixed in with 3% methylcellulose (Gibco, cat # 25030-081) and loaded into a 250μL Gastight syringe (Hamilton, cat #81120) attached to a 25G needle (BD, cat #305125). Mice were anesthetized using Isoflurane (Covetrus, cat # 11695-6777-2) and a midline incision was performed, exposing the skull. Using a stereotactic frame, the needle was positioned 2mm to the right of the bregma, and inserted 5mm below the surface of the skull before withdrawing 1mm, for an injection depth of 4mm. At this time, 1x10^4^ CT2A (2x10^6^/mL), 5x10^4^ Yummer1.7 (1x10^7^/mL), or 1x10^3^ (2x10^5^/mL) E0771 cells were delivered in a total volume of 5μL per mouse. The needle was slowly withdrawn over a period of 2 minutes to minimize leakage of cells out of the needle track. Mice received 100μL (0.5mg/mL) Meloxicam (Covetrus, cat # 11695-6936-1) injected subcutaneously and local administration of 3 drops of 0.25% Bupivacaine (Henry Schein, cat # 1411640) prior to wound closure using 7mm wound clips. Mice were monitored and sacrificed in accordance with Duke University Institutional Animal Care and Use Committee guidelines.

For subcutaneous injections, cells were harvested and resuspended in PBS at a concentration of 2.5x10^6^/mL for CT2A and Yummer1.7 and 1.25x10^6^/mL for E0771. Cells were injected in the subcutaneous space in the left flank of a mouse in a total volume of 200μL (5x10^5^ and 2.5x10^5^ cells per injection, respectively).

### Chronic LCMV infection

Chronic LCMV aliquots were kindly provided by Saskia Hemmers (Duke University, NC, USA). Mice were infected by intravenous injection of 2x10^6^ PFU LCMV clone 13. Mice were analyzed on various time points as indicated per figure.

### Tumor harvesting and tissue processing

#### Intracranial tumors

Mice were perfused by infusing 10mL PBS + 5% heparin (Fresenius Kabi, cat # C504730). The tumor-bearing hemisphere was removed and weighed. The tissue was transferred to a 7mL dounce tissue homogenizer (VWR, cat # 62400-620) and 3mL of HBSS+/+ (ThermoFisher, cat # 14025092) containing 0.5mg/mL Liberase DL (Sigma-Aldrich, cat #5401160001), 0.5mg/mL Liberase TL (Sigma-Aldrich, cat #5401020001), and 2mg/mL DNaseI (Sigma-Aldrich, cat #10104159001) was added. The tissue was gently homogenized using a loose pestle. The homogenized tissue was transferred to a 50mL conical, and the homogenizer was rinsed with 2mL HBSS with enzymes. After all tumor-bearing brains had been harvested, the samples were incubated in a 37°C water bath for 15 minutes. Following digestion, PBS was added to each tube to terminate the activity of the enzymes. After centrifugation, the tissues were resuspended in 2mL of 1x RBC lysis (Invitrogen, cat # 00-4300-54) for 2 minutes at room temperature. PBS was added to inactivate the reaction and samples were filtered through a 70µm cell strainer (Corning, cat # 431751). Samples were centrifuged and resuspended in 5mL 30% Percoll (Sigma-Aldrich, cat # P1644) to remove the myelin layer (48). Tissues were centrifuged at 500G for 20 minutes at 18°C, with low acceleration and brake. The myelin and remaining supernatant were carefully removed. The samples were resuspended in PBS or RPMI (Sigma-Aldrich, cat # R8758) depending on whether the cells were immediately stained, or *ex vivo* stimulated, respectively. Because of the limited cell number and T cell infiltration, the whole sample was stained for a single panel.

#### Subcutaneous tumors

The tumor was dissected out of the subcutaneous space. 10mL HBSS with enzymes was added to each tumor. The tumor plus digestion enzymes were transferred into a stomacher bag (Seward, cat # BA6040 and BA6040/CLR). The bags were sealed and placed inside the Stomacher 80 Biomaster Lab Blender (Seward, cat # 030010019) at 37°C for 30 minutes to one hour depending on the size of the tumor. Following digestion, the samples were transferred to a 50mL conical and centrifuged. Tissues were resuspended in 3mL of 1x RBC lysis for 3 minutes at room temperature. PBS was added to inactivate the reaction and samples were filtered through a 70µm cell strainer. Samples were centrifuged and resuspended in PBS or RPMI. Cells were counted using the Countess (ThermoFisher, cat # AMQAX2000) and 5 million cells were stained or stimulated per panel.

#### Secondary lymphoid organs

Spleen and tumor-draining lymph nodes were harvested from tumor-bearing and naïve mice. Organs were harvested in RPMI supplemented with 10% FBS. Spleens were meshed through a 70µm cell strainer to generate a single-cell suspension. After centrifugation, cells were resuspended in 3mL of 1x RBC lysis for 3 minutes at room temperature. PBS was added to inactivate the reaction and samples were filtered through a 70µm cell strainer. Samples were centrifuged and resuspended in PBS or RPMI. Cells were counted using the Countess and 5 million cells were stained or stimulated per panel. Tumor-draining lymph nodes were meshed through a 70µm cell strainer to generate a single-cell suspension. Cells were centrifuged and resuspended in PBS or RPMI. Because of the limited cell number, the entire lymph node was stained for a single panel.

### *Ex vivo* stimulation for cytokine detection

If cells were analyzed for cytokine production, samples were resuspended in 500μL RPMI supplemented with 10% FBS, HEPES, NEAA (Gibco, cat # 11140-050), L-glutamine (ThermoFisher, cat # 25030081), sodium pyruvate (ThermoFisher, cat # 11360070), and BME (Sigma Aldrich, cat # M3148) and transferred to a 48- well plate. Samples were then re-stimulated *ex vivo* with 50ng/mL PMA (Sigma-Aldrich, cat # P8139), 1µg/mL Ionomycin (Sigma-Aldrich, cat # I0634), 1/1500 GolgiStop (BD Biosciences, cat # 554724), and 1/1000 GolgiPlug (BD Biosciences, cat # 555029) for 4 hours at 37°C.

### Flow cytometry

The antibodies used are displayed in **Table 1**. All antibodies were anti-mouse unless specifically stated. All antibodies were diluted in FACS buffer (PBS + 1% FBS), except for live/dead staining which was performed in PBS. After samples had been centrifuged, cells were stained with 100µL of the diluted viability dye Zombie Aqua at room temperature for 15 minutes in the dark. Following the stain, cells were then resuspended in 50µL 1/500 FcBlock (Biolegend, cat # 101320) for 5-10 minutes at room temperature in the dark to prevent non-specific binding of antibodies. After the initial incubation with FcBlock, 50µL of a 2x antibody suspension was added to each sample, creating a final concentration of 1x. Samples were stained for 30 minutes at 4°C in the dark. If biotin antibodies were used, each sample was stained with 100µL of 1/100 diluted streptavidin for 30 min at 4°C in the dark. Tissues were fixed with the FoxP3/transcription factor kit according to manufacturer’s instructions (eBioscience, cat # 00-5523-00), unless the sample was stained for IL-2 or CCL5. In that case, the intracellular fixation and permeabilization Buffer (eBioscience, cat # 88-8824-00) was used according to manufacturer’s instructions. Samples fixed using the FoxP3 kit were resuspended in 100µL 1x fixation buffer and incubated for 30 minutes at 4°C in the dark. For the intracellular fixation kit, samples were resuspended in 100µL of the fixation buffer and incubated for 20 minutes at room temperature in the dark. For both methods, after fixation had been completed, cells were washed with 50µL 1x permeabilization buffer and centrifuged. Samples were resuspended in 200µL 1x permeabilization buffer until intracellular staining was performed.

**Table 1:** Antibody table.

| *Antibody* | *Color* | *Clone* | *Dilution* | *Source* | *Cat. number* |
| --- | --- | --- | --- | --- | --- |
| *Zombie Aqua™*<br>*Fixable Viability*<br>*Kit* | BV510 |  | 1/500 | BioLegend | 423102 |
| *CD120a (TNFR1)* | APC | 55R-286 | 1/100 | BioLegend | 113006 |
| *CD120a (TNFR1)* | Biotin |  | 1/100 |  |  |
| *CD120b (TNFR2)* | PE | TR75-89 | 1/100 | BioLegend | 113406 |
| *CD3* | PE-Cy5 | 145-2C11 | 1/200 | BioLegend | 100310 |
| *CD39* | PE-eFluor610 | 24DMS1 | 1/100 | ThermoFisher | 61-0391-82 |
| *CD39* | PE-Cy7 | 24DMS1 | 1/100 | ThermoFisher | 25-0391-82 |
| *CD44* | PE-CF594 | IM7 | 1/500 | BD Biosciences | 562464 |
| *CD44* | BV785 | IM7 | 1/200 | BioLegend | 103059 |
| *CD45* | APC-Cy7 | 30-F11 | 1/500 | BioLegend | 103116 |
| *CD45.1* | PE-Cy7 | A20 | 1/100 | BioLegend | 110730 |
| *CD45.2* | BV605 | 104 | 1/100 | BD Biosciences | 563051 |
| *CD8a* | BV605 | 53-6.7 | 1/100 | BioLegend |  |
| *CD8a* | BV650 | 53-6.7 | 1/100 | BioLegend | 100742 |
| *Anti-human*<br>*Granzyme B* | PE | GB12 | 1/200 | ThermoFisher | MHGB04 |
| *IFNγ* | APC-Cy7 | XMG1.2 | 1/200 | BioLegend | 505850 |
| *IFNγ* | AF488 | XMG1.2 | 1/200 | ThermoFisher | 53-7311-82 |
| *IL2* | BV785 | JES6-5H4 | 1/100 | BioLegend | 503843 |
| *IL2* | PE/Dazzle594 | JES6-5H4 | 1/100 | BioLegend | 503840 |
| *Ki67* | BV786 | B56 | 1/100 | BD Biosciences | 563756 |
| *Ki67* | Alexa Fluor 647 | B56 | 1/100 | BD Biosciences | 561126 |
| *Zombie Aqua* |  |  | 1/500 | BioLegend | 423102 |
| *PD-1* | BV711 | 29F.1A12 | 1/100 | BioLegend | 135231 |
| *PD-1* | PE/Dazzle594 | RMP1-30 | 1/100 | BioLegend | 109116 |
| *SLAMF6* | BV786 | 13G3 | 1/100 | BD Biosciences | 741030 |
| *SLAMF6* | BV650 | 13G3 | 1/100 | BD Biosciences | 740628 |
| *TIM3* | BV421 | RMT3-23 | 1/100 | BioLegend | 119723 |
| *TNF-α* | BV785 | MP6-XT22 | 1/100 | BioLegend | 506341 |
| *TNF-α* | PE/Dazzle594 | MP6-XT22 | 1/100 | BioLegend | 506346 |
| *TOX* | eFluor 660 | TXRX10 | 1/50 | ThermoFisher | 50-6502-82 |

Intracellular staining was performed on the day the samples were analyzed on the flow cytometer. Samples were stained with 80µL 1x permeabilization buffer with diluted antibodies. Samples were incubated for 30-45 minutes at room temperature in the dark. Samples were then resuspended in PBS containing 1/1000 dilution DNaseI (ThermoFisher, cat # 89836) and 2mM EDTA (ThermoFisher, cat # AM9269G). Samples were filtered using a FACS tube with a 35µM strainer cap. If counting beads (ThermoFisher, cat # C36950) were used to determine absolute counts, 10μL was added to each sample. Cells were collected on a BD Fortessa and analyzed using FlowJo version 10.10.

### Cell sorting

In experiments where populations were sorted, tissues were harvested as described above. To increase the sorting efficiency, populations of interest were enriched following the generation of a single cell suspension. For subcutaneous tumor samples, the sample was enriched for CD45+ cells using the murine CD45 (TIL) MicroBeads cocktail (Miltenyi, cat # 130-110-618) according to manufacturer’s protocol. Enrichment was achieved using the “possel” program on the Miltenyi autoMACS. In spleens from cLCMV- infected mice, a negative selection for CD8 T cells (Miltenyi, cat # 130-104-075) was performed according to manufacturer’s protocol. Enrichment was achieved using the “deplete” program on the Miltenyi autoMACS. The cells containing the enriched populations were then stained with the appropriate antibodies and were sorted using the Astrios Cell Sorter at the Duke Cancer Institute (DCI), BD FACSymphony S6, or BD FACSAria II at the Duke Human Vaccine Institute (DHVI) Research Flow Cytometry Shared Resource Facility (Durham, NC).

### *Ex* vivo generation of Trp2-specific T cells and adoptive lymphocyte transfer (ALT)

Trp2-specific T cells recognize the TRP2180-188 epitope in the context of H-2Kb (49). To generate Trp2- specific T cells, spleens were harvested from WT CD45.1 and TNFR2 KO CD45.2 mice. Splenocytes were resuspended at 2x10^6^ cells/mL in RPMI containing 10% FBS, 50 U/ml human IL-2 (Iovance), 2 μg/ml Concanavalin A (Sigma-Aldrich, cat # C5275), NEAA, sodium pyruvate, L-glutamine, Pen/Strep (Gibco, cat # 15140-122), and 50 µM BME. After 48 hours of culture, activated splenocytes were retrovirally transduced with the pMX-TRP2-TCRβ-2A-α vector (a kind gift from Dr. Schumacher, Netherlands Cancer Institute) (50). To generate the transduction supernatant, HEK293 T cells were transfected with pCL-ECO (Addgene, cat # 12371) and the TRP2 vector with lipofectamine 2000 (ThermoFisher, cat # 11668019) for 48-hours. This viral supernatant was combined with the activated splenocytes in Retronectin-coated (Takara bio, cat # T100B, 25μg/mL) non-tissue culture treated 24-well plates. The plates were centrifuged for 1.5h at 2000rpm with medium acceleration and brake. Complete RPMI with IL-2 was added, and cells were cultured for 2 days post-transduction, splitting daily. 21 days after implantation of CT2A-TRP2 in the subcutaneous space of CD45.1 CD45.2 recipients, 2.5x10^6^ WT (CD45.1) and 2.5 x10^6^ TNFR2 KO (CD45.2) Trp2-TCR transgenic cells were adoptively transferred into the same recipient via tail vein injection. Seven days post-transfer, tumors were harvested as described above, and WT and TNFR2 KO cells were investigated for exhaustion phenotypes.

### Bulk RNA-sequencing preparation

To investigated transcriptional differences in WT and TNFR2 KO CD8^+^ T cells, terminally (TIM3^+^) exhausted T cells were sorted from cLCMV-infected spleens. Cells were lysed using the TRI reagent (Zymo, cat # R2050-1-50) and RNA was isolated using the Direct-Zol MinPrep kit from Zymo (Cat # R2050), including the optional DNaseI treatment step. RNA was eluted in 25µL RNase- and DNase-free water. mRNA was purified using the NEBNext Poly(A) mRNA Magnetic Isolation Module (NEB, cat # E7490). Samples were processed using the NEBNext Ultra-II RNA library prep kit from New England Biolabs (cat # E7775S) and the NEBNext Multiplex Oligos for Illumina (96 Unique Dual Index Primer pairs) (NEB, cat # E6440S) according to manufacturer’s protocol.

### Bulk RNA-sequencing analysis

RNA-Seq data was processed and analyzed using R version 4.3.1. Initial preprocessing included mapping Ensembl gene IDs to gene symbols using the org.Mm.eg.db library. Data was aggregated by gene symbol, and count values were rounded to the nearest integer. Differential expression analysis was performed using the DESeq2 package. Genes with low counts were filtered out, and the dataset was analyzed to identify differentially expressed genes between wild-type (WT) and TNFR2 knockout (KO) samples. All analyses were adjusted for multiple testing using the Benjamini-Hochberg procedure to control the false discovery rate.

Principal Component Analysis (PCA) was conducted using the prcomp function on the VST-transformed data. The first two principal components were plotted to visualize the clustering of samples, with colors indicating different sample groups (WT and KO). Heatmaps were generated using the ComplexHeatmap package. Variance-stabilizing transformation (VST) was applied to the count data using vst from the DESeq2 package.

To perform gene module enrichment, pathways of interest were defined from a custom GMT file based on signatures taken from the mouse NanoString nCounter^®^ Immune Exhaustion Panel (lists of genes for each module are provided in the Supplemental Materials). The fgsea package was used to identify significant gene modules, and the results were summarized and visualized.

### Tumor growth assessment

Tumor sizes were measured every 3 days starting at 9 days post-implantation. Tumor volumes were calculated using the formula L * W * W/2. Mice were sacrificed if they reached the following humane endpoints: a total tumor volume of 2000mm^3^, a measurement of 20mm in any direction, or an ulceration. If mice were sacrificed due to volume or size endpoints, the volume recorded on the day of sacrifice was included in any subsequent time points. If a mouse was sacrificed due to an ulceration, the volumes were recorded until the time of sacrifice and the mouse was censored for any subsequent time points. To determine significant differences in tumor volume, a two-way ANOVA was employed. Because these decisions influence statistical approaches, time to reach humane endpoint was also evaluated. Finally, we devised a tumor burden output. It utilizes the area under the curve (AUC) of the tumor volume of each individual mouse throughout the experiment and is normalized to the number of days until humane endpoint.

### *In vivo* antibodies

To determine the role of CD8^+^ T cells in the tumor growth phenotype observed in TNFR2 KO mice, CD8^+^ T cells were depleted through intraperitoneal injections of 200µg anti-CD8a (BioXCell, clone 2.43) on days - 3, -1, and every 6 days post tumor implantation. To assess the effect of TNFR2 antagonism, with and without immune checkpoint blockade, 250μg TNFR2 antagonist (BioXCell, cat # BE0247) and 200μg anti- PD-1 (BioXCell, cat # BE0146) was injected intraperitoneally starting at day 9, and every 3 days thereafter.

### cLCMV viral titer aPCR

Spleens of cLCMV-infected mice were harvested at day 14 post-infection. After a single cell suspension was generated and ACK lysis was performed (see above), samples were counted. For each sample, 2 million cells were lysed using the TRI reagent. RNA was isolated using the Direct-Zol MinPrep kit from Zymo, including the optional DNaseI treatment step. RNA was eluted in 25µL RNase- and DNase-free water. cDNA was generated using the iScript cDNA synthesis kit (Biorad, cat # 1708891) according to manufacturer’s instructions. For each sample, 100ng RNA template was utilized. Following cDNA synthesis, qPCR was performed to determine viral titers as described previously (51). Briefly, 10μL SYBR green (Biorad, cat # 1725271), 2µL forward primer (5’ CATTCACCTGGACTTTGTCAGACTC), 2µL reverse primer (5’ GCAACTGCTGTGTTCCCGAAAC), and 2µL water/standard/sample were added. The standard curve was a kind gift from Dr. Hemmers (Duke University). A total of 8 points were included in the standard curve. The first point on the standard curve was undiluted. This point corresponded to 1ng/μL, which equates to 2.19x10^9^ virus particles. Each subsequent point was a ten-fold dilution. Each sample was run in duplicate. Cycling parameters: 3 minutes at 95C followed by 40 cycles of 15 seconds at 95C and 30 seconds at 60C. Finally, a melting curve test was performed.

### Statistical analysis

All data are representative of at least 2 separate experiments. Statistical analyses were performed using GraphPad Prism software version 10.0.0. Data are depicted as the mean ± standard error of the mean (SEM). When two groups were compared, a paired or unpaired t-test was utilized. When multiple groups were assessed, a one-way or two-way repeated measures ANOVA was applied. To compute correlations, simple linear regression was performed using the two-tailed p-value assessment and a 95% confidence interval.

For the time to humane endpoint analysis, a Kaplan Meier survival analysis with the Log-rank test was performed. The number of comparisons were determined for each experiment to dictate the corrected p-value: α corrected = 0.05/number of comparisons. In the experiment assessing the role of CD8 T cells, 2 comparisons were made: WT isotype vs KO isotype and KO isotype vs KO αCD8. The survival following treatment was considered significantly different when p < 0.025. For the tumor growth following TNFR2 antagonism with and without immune checkpoint blockade, the following 6 comparisons were made: untreated vs anti-PD-1, untreated vs anti-TNFR2, untreated vs combo treatment, anti-PD-1 vs anti-TNFR2, anti-PD-1 vs combo treatment, and anti-TNFR2 vs combo treatment. Survival was considered significantly different when p < 0.00833. For all other experiments, p-values < 0.05 were considered statistically significant.

---

**Supplementary Fig 1:**
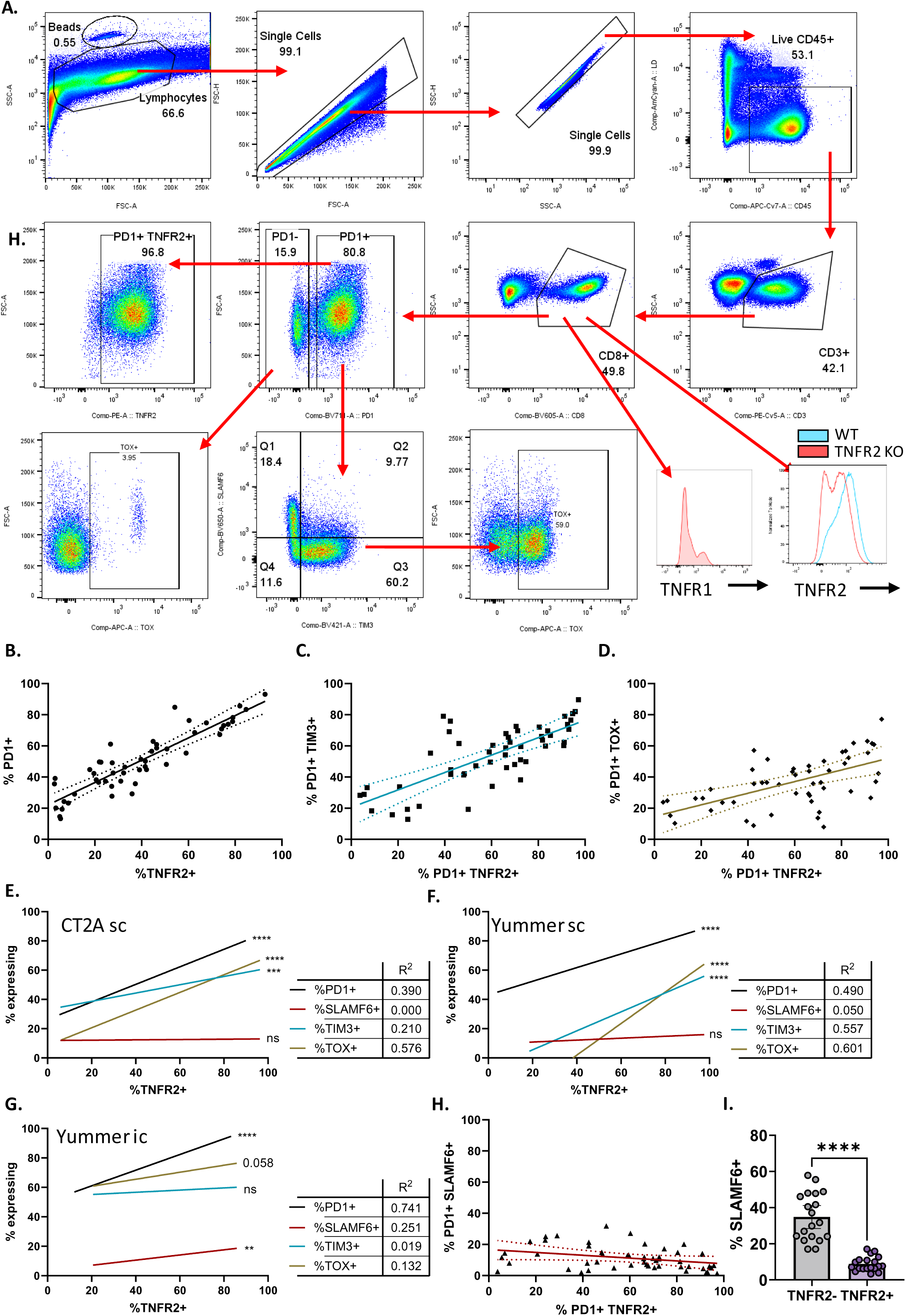
TNFR2 is correlated with canonical markers of exhaustion in multiple tumor models. (A) Gating strategy for assessing exhaustion markers. (B-D) Individual data points comparing frequency of TNFR2 to PD1 (B), TIM3 (C), and TOX (D) on tumor-infiltrating CD8^+^ T cells in intracranial CT2A. Simple linear regression was performed. Dotted line represents the 95% confidence interval. (E-G) Correlation of TNFR2 expression on tumor-infiltrating CD8^+^ T cells with expression of canonical markers of exhaustion (PD1, SLAMF6, TIM3, and TOX) in subcutaneous CT2A (E), subcutaneous Yummer1.7 (F), and intracranial Yummer1.7 (G). Tumors were harvested at similar timepoints as for Fig 1A and B. A simple linear regression model was performed to determine significance and R^2^ values. (H) Individual data points comparing frequency of TNFR2 to SLAMF6 on tumor-infiltrating CD8^+^ T cells in intracranial CT2A. Simple linear regression was performed. Dotted line represents the 95% confidence interval. (I) Frequency of SLAMF6+ cells in TNFR2- and TNFR2+ CD8^+^ T cells in the spleen of mice infected with chronic viral infection at peak viral titers (D14). Paired t-test was performed. Data represented as mean ± 95% confidence interval. ** p < 0.01, *** p<0.001, **** p < 0.0001

**Supplementary Fig 2:**
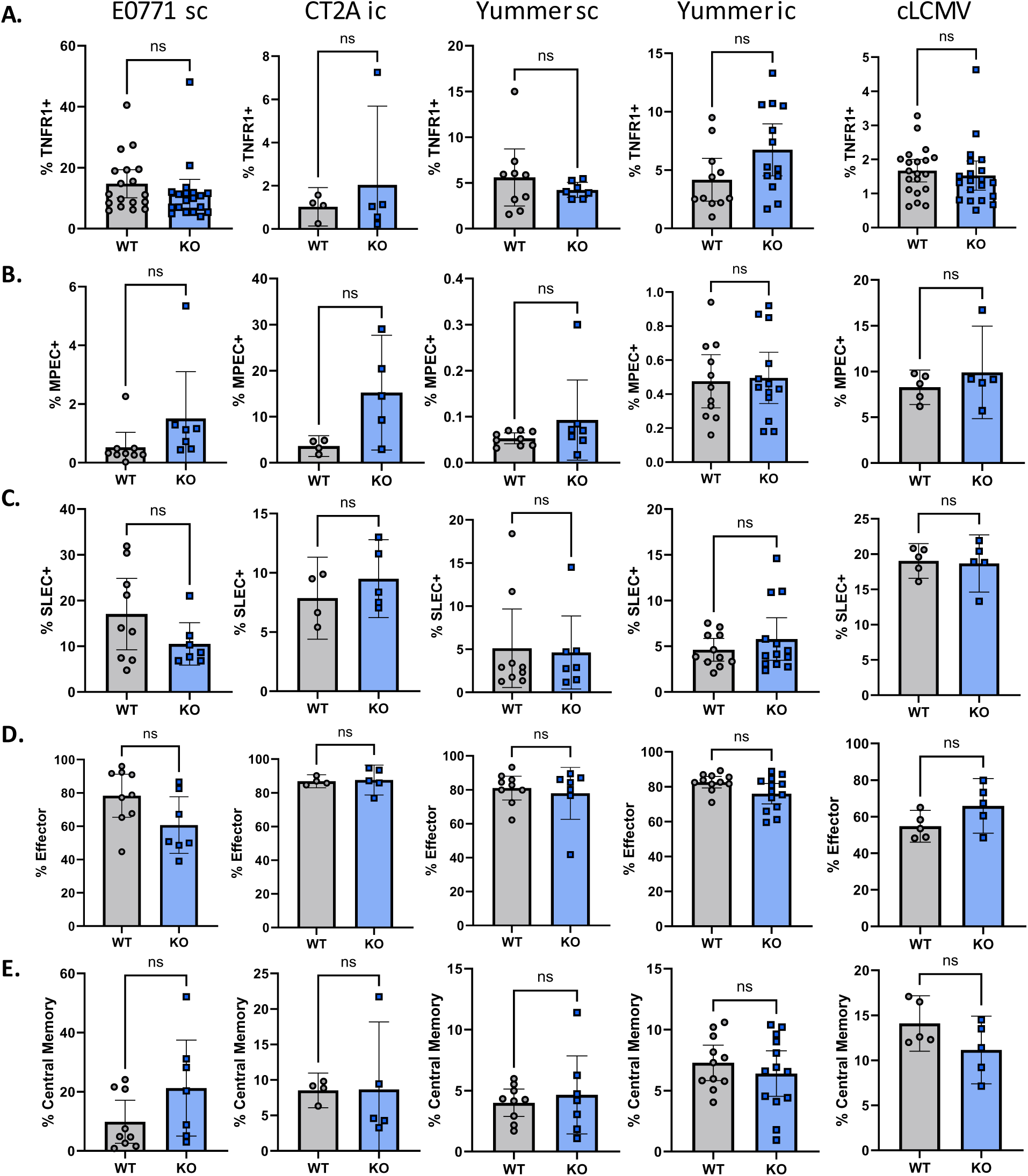
Loss of TNFR2 is not associated with increased expression of TNFR1 or improved memory phenotype acquisition. (A) Frequency of TNFR1+ on WT and TNFR2 KO CD8^+^ T cells harvested from tumor of mice implanted with sc E0771 (D25), ic CT2A (D21), sc Yummer (D28), ic Yummer (D25), or spleen for cLCMV (D14). Unpaired t-test was performed. (B-E) Frequency of memory precursor effector cells (MPECs; KLRG1^-^CD127^+^) (B), short-lived effector cells (SLECs; KLRG1^+^CD127^-^) (C), effector (CD62L^-^CD44^+^) (D), and central memory (CD62L^+^CD44^+^) (E) on WT and TNFR2 KO CD8^+^ T cells harvested from tumor of mice implanted with sc E0771 (D25), ic CT2A (D21), sc Yummer (D28), ic Yummer (D25), or spleen for cLCMV (D14). Unpaired t-test was performed. Data represented as mean ± 95% confidence interval. Ns not significant

**Supplementary Fig 3:**
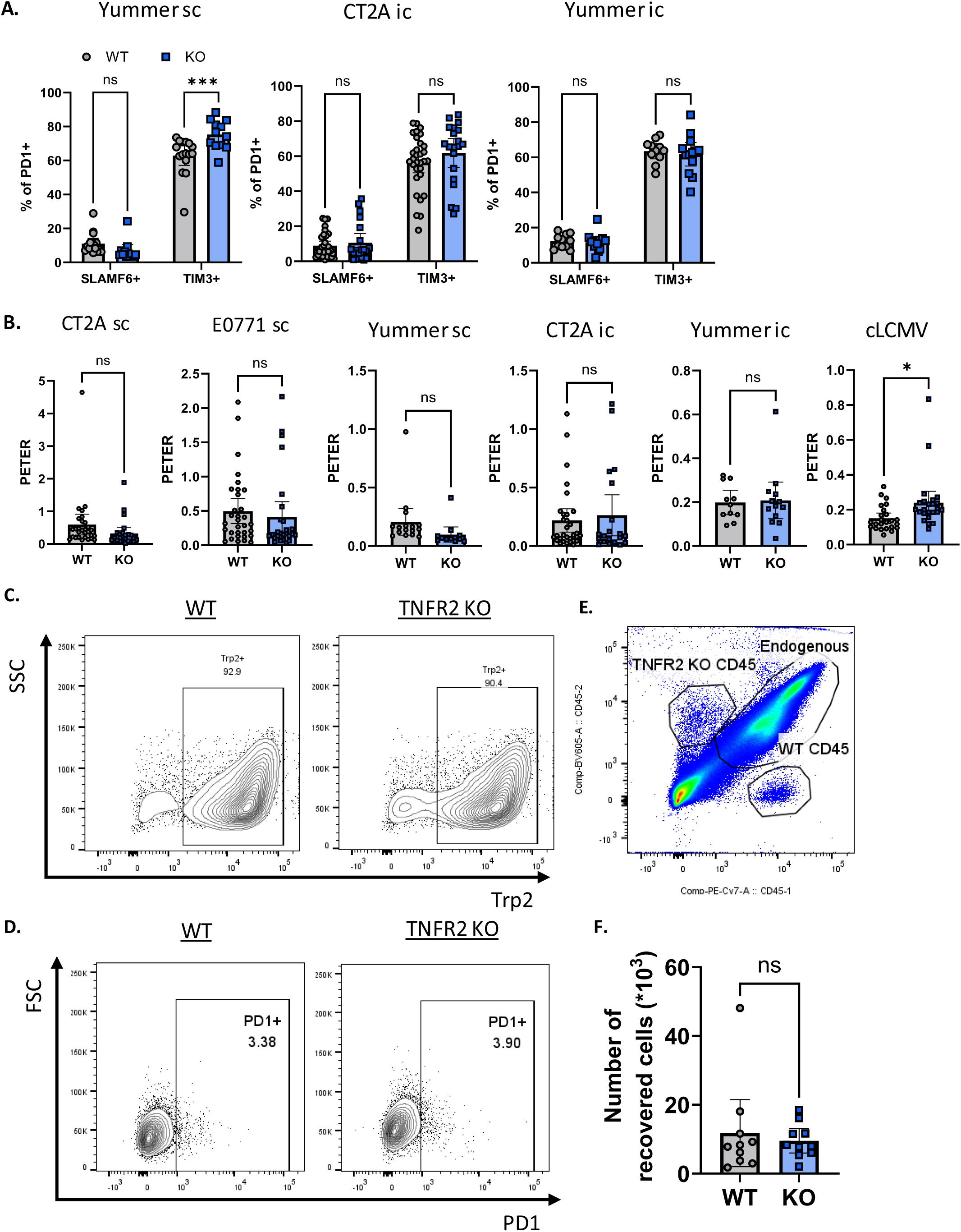
TNFR2 KO T cells display similar levels of progenitor and terminally exhausted phenotypes. (A) Frequency of progenitor (SLAMF6^+^TIM3^-^) and terminally exhausted (SLAMF6^-^TIM3^+^) CD8^+^ TILs from wild type (WT) and TNFR2 KO in sc Yummer1.7 (D28), ic CT2A (D21), ic Yummer1.7 (D25). Two-way repeated measures ANOVA was performed. (B) Progenitor to terminal ratio (PETER) on CD8^+^ TILs from wild type (WT) and TNFR2 KO in sc CT2A (D35), sc E0771 (D25), sc Yummer1.7 (D28), ic CT2A (D21), ic Yummer1.7 (D25), and CD8^+^T cells in the spleen of cLCMV-infected mice (D14). (C) Trp2 transduction efficiency of WT and TNFR2 KO splenocytes. (D) Pre-transfer frequency of PD1+ cells of Trp2-specific T cells. (E) Representative flow plot of CD45.1 (WT), CD45.2 (TNFR2 KO), and CD45.1 CD45.2 (endogenous T cell infiltration) harvested from CT2A-Trp2 tumors 7 days after adoptive transfer of Trp2-specific T cells. (F) Number Trp2-specific T cells recovered 7 days post transfer from a subcutaneous CT2A-trp2 tumor. Paired t-test was performed. Data represented as mean ± 95% confidence interval. Ns not significant, * p<0.05

**Supplementary Fig 4:**
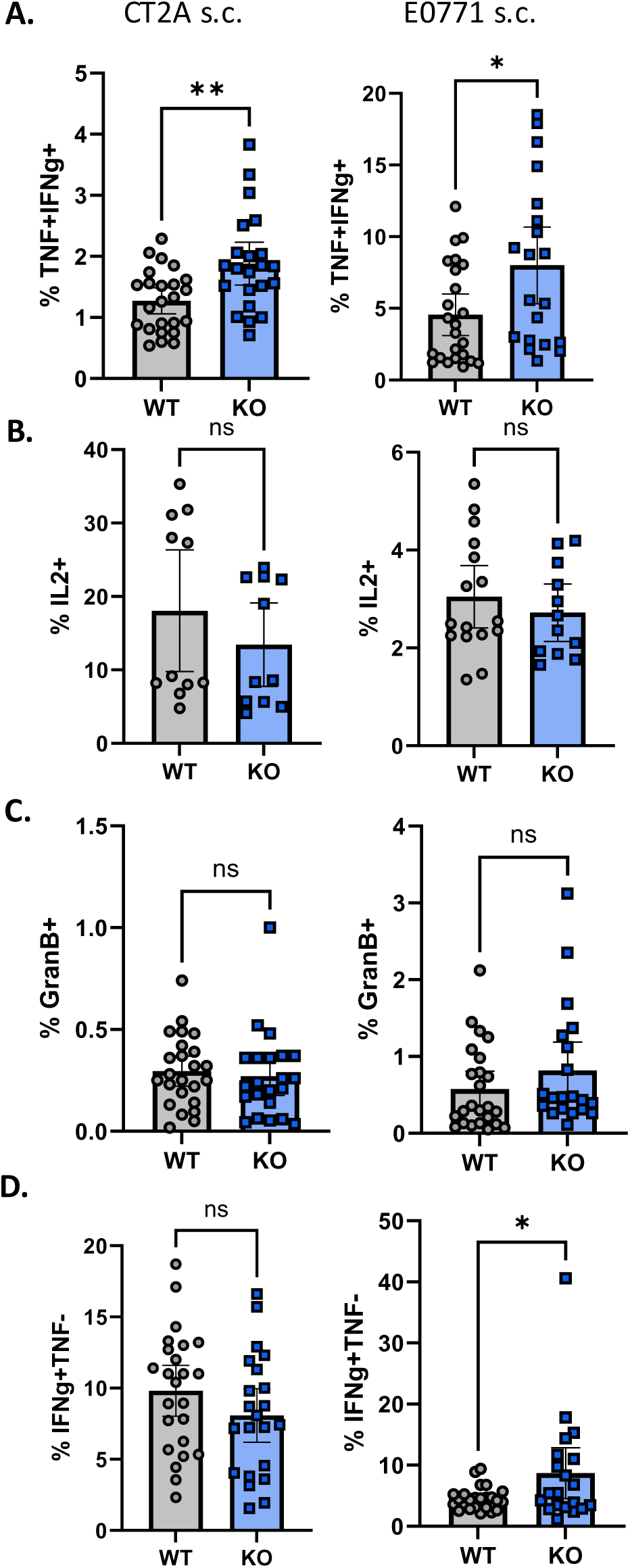
TNFR2 KO mice demonstrate improved CD8^+^ T cell functions in tumor-draining lymph nodes. (A-E) CD8^+^ T cells from WT and TNFR2 KO mice from tumor-draining lymph nodes were stimulated with PMA, Ionomycin, GolgiStop, and GolgiPlug for 4 hours at 37°C. Polyfunctionality (IFNγ^+^TNF^+^) (A), IL-2 (B), granzyme B (C), and IFNγ single-producing CD8^+^ T cells (D) was evaluated. Data represented as mean ± 95% confidence interval. ns not significant, * p < 0.05, ** p < 0.01, **** p < 0.0001

**Supplementary Fig 5:**
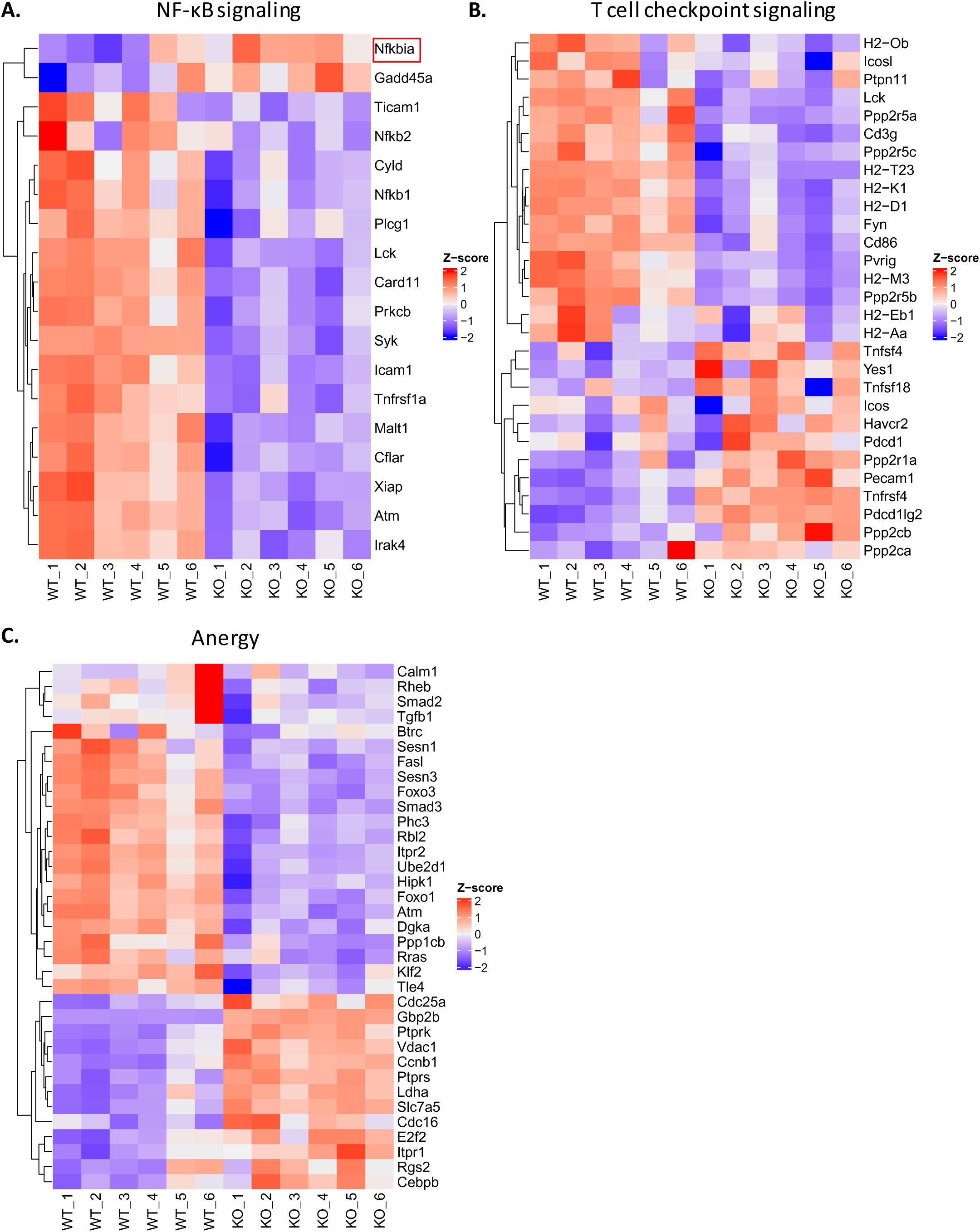
TNFR2 KO CD8^+^ T cells show decreased NF-κB activity and altered activation of inhibitory and anergy pathways. (A-C) Heatmaps displaying z-scores for NF-κB signaling (A), T cell checkpoint signaling (B), and anergy (C) each WT and TNFR2 KO CD8^+^ TIM3^+^ sample as defined the mouse NanoString nCounter^®^ Immune Exhaustion Panel.

**Supplementary Fig 6:**
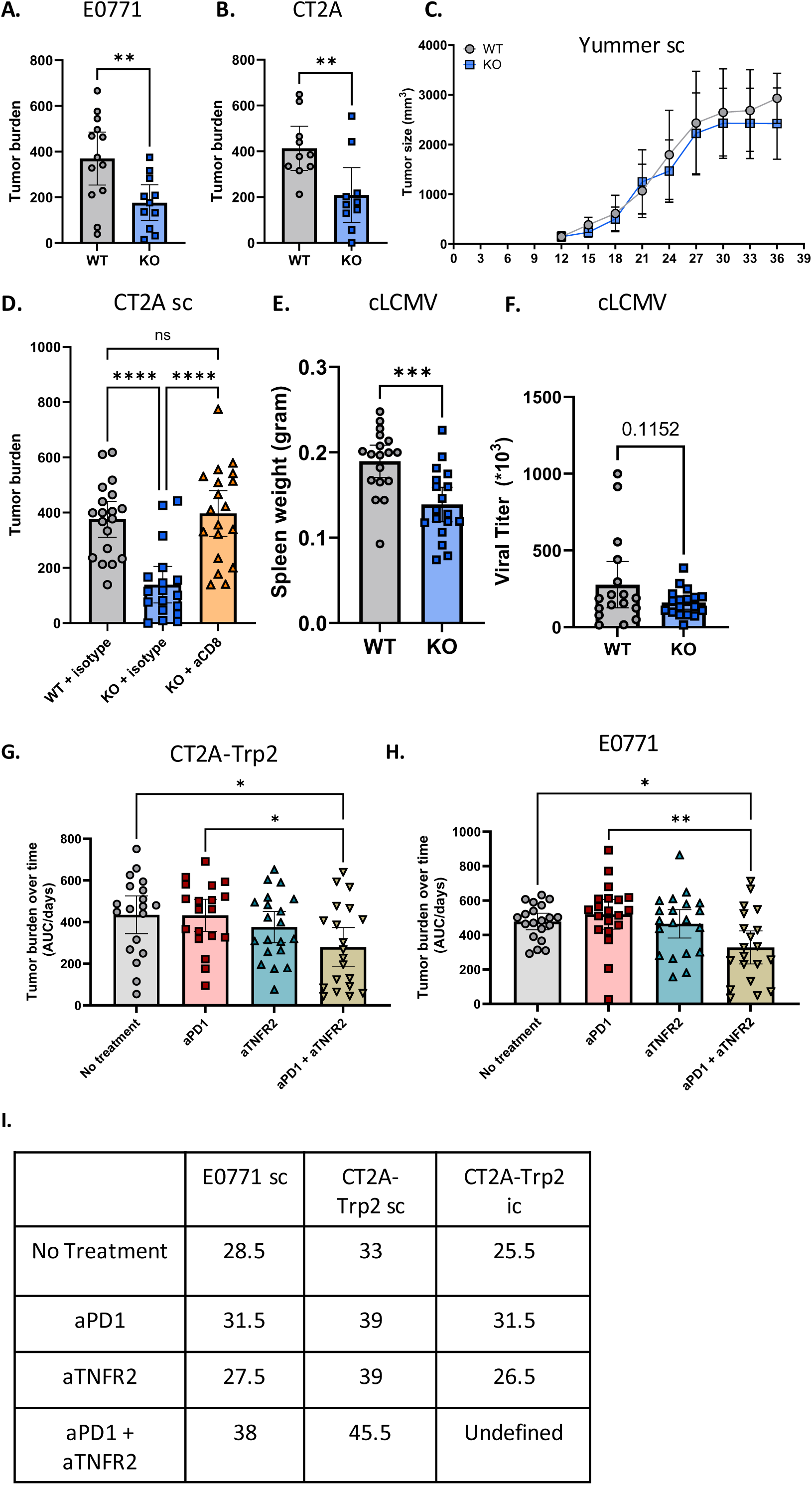
TNFR2 deficiency results in model-specific effects on tumor outcome and viral control. (A-B) Tumor burden in WT and TNFR2 KO mice implanted with sc E0771 (A) or sc CT2A (B). Tumor burden is defined as the area under the curve (AUC) of the tumor growth for each mouse over time divided by the number of days until reaching humane endpoint. This metric allows for the inclusion of ulcerated tumors and more adequately displays fast growing tumors. Unpaired t-test was performed. (C) Tumor volumes in WT and TNFR2 KO mice implanted with sc Yummer1.7 (D) Tumor burden in WT, TNFR2 KO, and TNFR2 KO mice dosed with an anti-CD8 depletion antibody following implantation of sc CT2A. One-way ANOVA was performed. (E-F) Spleen weights (E) and viral titers (F) in WT and TNFR2 KO mice on day 14 post infection. Unpaired t-test was performed. (G-H) Tumor burden in WT mice treated with anti-PD1, anti-TNFR2, or combination of anti-PD1 and anti-TNFR2 after sc implantation with CT2A-Trp2 (G) or E0771 (H). One-way ANOVA was performed. Mice were injected with anti-PD1 (200ug), anti-TNFR2 (250ug), or both every 3 days starting on day 9. Tumors were measured every 3 days. (I) Median time to humane endpoint for subcutaneous implantation of E0771 and CT2A-Trp2 or intracranial implantation of CT2A-Trp2. Median survival was not reached in the aTNFR2/aPD1 combination group following intracranial CT2A-Trp2 as 12 out of 20 survived long-term (more than 50 days). Data represented as mean ± 95% confidence interval. Ns not significant, * p<0.05, **p<0.01, *** p <0.001, **** p<0.0001

## Notes

### Competing Interest Statement

The authors have declared no competing interest.

### Summary of Updates

We have updated the paper to include anti-TNFR2/anti-PD1 survival data in an intracranial model of GBM

