## Supplemental Table 1 for "Upregulation of TNFR2 Precedes TOX Expression by Exhausted T cells and Restricts Antitumor and Antiviral Immunity"

| symbol | ensembl_II | User_ID | KO-WT_log | KO-WT_adj | Signifcant | Significant | Upregulated/Downreg |
|---|---|---|---|---|---|---|---|
| Eef1a1 | ENSMUSG( | ENSMUSG( | 0.229202 | 0.001071 | YES | NO | NA |
| Actb | ENSMUSG( | ENSMUSG( | -0.31063 | 4.10E-08 | YES | NO | NA |
| mt-Co1 | ENSMUSG( | ENSMUSG( | 0.165861 | 0.032878 | YES | NO | NA |
| Eef2 | ENSMUSG( | ENSMUSG( | 0.23427 | 6.70E-05 | YES | NO | NA |
| Actg1 | ENSMUSG( | ENSMUSG( | -0.07276 | 0.148544 | NO | NO | NA |
| Myh9 | ENSMUSG( | ENSMUSG( | -0.27014 | 3.09E-06 | YES | NO | NA |
| Tmsb4x | ENSMUSG( | ENSMUSG( | -0.24513 | 2.23E-05 | YES | NO | NA |
| Ahnak | ENSMUSG( | ENSMUSG( | -0.49425 | 2.97E-09 | YES | NO | NA |
| H2-K1 | ENSMUSG( | ENSMUSG( | -0.66628 | 1.43E-10 | YES | NO | NA |
| Ccl5 | ENSMUSG( | ENSMUSG( | -1.19021 | 3.82E-08 | YES | YES | Downregulated |
| Gapdh | ENSMUSG( | ENSMUSG( | 0.484085 | 8.42E-07 | YES | NO | NA |
| Ptma | ENSMUSG( | ENSMUSG( | 0.242592 | 0.33381 | NO | NO | NA |
| Flna | ENSMUSG( | ENSMUSG( | -0.17236 | 0.001964 | YES | NO | NA |
| Hspa8 | ENSMUSG( | ENSMUSG( | -0.31306 | 1.11E-05 | YES | NO | NA |
| Rn18s-rs5 | ENSMUSG( | ENSMUSG( | 0.65529 | 0.35551 | NO | NO | NA |
| Rplp0 | ENSMUSG( | ENSMUSG( | 0.390706 | 3.81E-08 | YES | NO | NA |
| Vim | ENSMUSG( | ENSMUSG( | 0.598086 | 9.05E-07 | YES | NO | NA |
| H2-D1 | ENSMUSG( | ENSMUSG( | -0.4587 | 8.68E-11 | YES | NO | NA |
| Pfn1 | ENSMUSG( | ENSMUSG( | -0.12031 | 0.001368 | YES | NO | NA |
| Hsp90ab1 | ENSMUSG( | ENSMUSG( | 0.15441 | 0.094999 | NO | NO | NA |
| Ptprc | ENSMUSG( | ENSMUSG( | -0.39788 | 5.81E-13 | YES | NO | NA |
| Lcp1 | ENSMUSG( | ENSMUSG( | 0.003915 | 0.951017 | NO | NO | NA |
| Rpsa | ENSMUSG( | ENSMUSG( | 0.408851 | 2.40E-08 | YES | NO | NA |
| Pabpc1 | ENSMUSG( | ENSMUSG( | 0.053856 | 0.558407 | NO | NO | NA |
| Tpt1 | ENSMUSG( | ENSMUSG( | -0.05625 | 0.468029 | NO | NO | NA |
| Rpl13a | ENSMUSG( | ENSMUSG( | -0.02768 | 0.688341 | NO | NO | NA |
| Coro1a | ENSMUSG( | ENSMUSG( | 0.10764 | 0.019259 | YES | NO | NA |
| B2m | ENSMUSG( | ENSMUSG( | -0.50156 | 3.40E-09 | YES | NO | NA |
| Gm43305 | ENSMUSG( | ENSMUSG( | 2.897809 | 1.44E-15 | YES | YES | Upregulated |
| Rpl4 | ENSMUSG( | ENSMUSG( | 0.344473 | 7.69E-09 | YES | NO | NA |
| Cfl1 | ENSMUSG( | ENSMUSG( | 0.092268 | 0.06932 | NO | NO | NA |
| H2-Q7 | ENSMUSG( | ENSMUSG( | -0.79219 | 7.64E-10 | YES | NO | NA |
| Rps2 | ENSMUSG( | ENSMUSG( | 0.630756 | 4.12E-09 | YES | NO | NA |
| Laptm5 | ENSMUSG( | ENSMUSG( | -0.53485 | 1.63E-13 | YES | NO | NA |
| Malat1 | ENSMUSG( | ENSMUSG( | -1.29177 | 1.31E-23 | YES | YES | Downregulated |
| Mki67 | ENSMUSG( | ENSMUSG( | 0.064378 | 0.60722 | NO | NO | NA |
| Hnrnpa2b1 | ENSMUSG( | ENSMUSG( | 0.079036 | 0.487278 | NO | NO | NA |
| Itga4 | ENSMUSG( | ENSMUSG( | -0.49608 | 4.70E-07 | YES | NO | NA |
| Pkm | ENSMUSG( | ENSMUSG( | 0.473083 | 1.17E-05 | YES | NO | NA |
| Itgal | ENSMUSG( | ENSMUSG( | -0.5149 | 4.26E-28 | YES | NO | NA |
| mt-Co2 | ENSMUSG( | ENSMUSG( | 0.117114 | 0.699119 | NO | NO | NA |
| Tubb5 | ENSMUSG( | ENSMUSG( | 0.150401 | 0.000958 | YES | NO | NA |
| Prex1 | ENSMUSG( | ENSMUSG( | -0.6532 | 5.45E-13 | YES | NO | NA |
| Hmgb2 | ENSMUSG( | ENSMUSG( | 0.223814 | 0.031357 | YES | NO | NA |
| Rpl3 | ENSMUSG( | ENSMUSG( | 0.410328 | 4.84E-11 | YES | NO | NA |
| Hnrnpa3 | ENSMUSG( | ENSMUSG( | 0.165006 | 0.104551 | NO | NO | NA |

| | | | | | | | |
|---|---|---|---|---|---|---|---|
| Gzmb | ENSMUSG( | ENSMUSG( | -0.09004 | 0.510027 | NO | NO | NA |
| Ddx5 | ENSMUSG( | ENSMUSG( | -0.16848 | 0.029026 | YES | NO | NA |
| Tln1 | ENSMUSG( | ENSMUSG( | -0.25868 | 9.09E-13 | YES | NO | NA |
| mt-Atp6 | ENSMUSG( | ENSMUSG( | 0.328882 | 0.008309 | YES | NO | NA |
| Rbm3 | ENSMUSG( | ENSMUSG( | 0.809647 | 4.19E-08 | YES | NO | NA |
| Actr3 | ENSMUSG( | ENSMUSG( | -0.14687 | 0.001238 | YES | NO | NA |
| mt-Cytb | ENSMUSG( | ENSMUSG( | 0.034785 | 0.693338 | NO | NO | NA |
| Macf1 | ENSMUSG( | ENSMUSG( | -0.7644 | 1.16E-20 | YES | NO | NA |
| mt-Nd1 | ENSMUSG( | ENSMUSG( | 0.275365 | 0.000633 | YES | NO | NA |
| Ets1 | ENSMUSG( | ENSMUSG( | -0.49307 | 1.54E-09 | YES | NO | NA |
| Anxa6 | ENSMUSG( | ENSMUSG( | -0.21031 | 1.29E-07 | YES | NO | NA |
| Rps6 | ENSMUSG( | ENSMUSG( | 0.370128 | 7.74E-08 | YES | NO | NA |
| Il2rb | ENSMUSG( | ENSMUSG( | -0.21396 | 0.000571 | YES | NO | NA |
| Rps3a1 | ENSMUSG( | ENSMUSG( | 0.344167 | 4.28E-07 | YES | NO | NA |
| Spn | ENSMUSG( | ENSMUSG( | -0.40273 | 5.12E-10 | YES | NO | NA |
| Xist | ENSMUSG( | ENSMUSG( | -0.36349 | 4.35E-07 | YES | NO | NA |
| Msn | ENSMUSG( | ENSMUSG( | -0.23898 | 3.34E-07 | YES | NO | NA |
| Eno1 | ENSMUSG( | ENSMUSG( | 0.513307 | 3.51E-05 | YES | NO | NA |
| mt-Nd5 | ENSMUSG( | ENSMUSG( | 0.145048 | 0.231138 | NO | NO | NA |
| Ppia | ENSMUSG( | ENSMUSG( | 0.20297 | 0.011743 | YES | NO | NA |
| Cd8a | ENSMUSG( | ENSMUSG( | -0.2926 | 1.25E-09 | YES | NO | NA |
| Esyt1 | ENSMUSG( | ENSMUSG( | -0.09487 | 0.153535 | NO | NO | NA |
| Rpl8 | ENSMUSG( | ENSMUSG( | 0.488801 | 1.49E-09 | YES | NO | NA |
| Rps3 | ENSMUSG( | ENSMUSG( | 0.26066 | 4.25E-07 | YES | NO | NA |
| Thy1 | ENSMUSG( | ENSMUSG( | -0.17758 | 0.000292 | YES | NO | NA |
| Rack1 | ENSMUSG( | ENSMUSG( | 0.317783 | 4.64E-07 | YES | NO | NA |
| Ubb | ENSMUSG( | ENSMUSG( | -0.14469 | 0.113903 | NO | NO | NA |
| Hnrnpu | ENSMUSG( | ENSMUSG( | 0.062617 | 0.570158 | NO | NO | NA |
| Lgals1 | ENSMUSG( | ENSMUSG( | -0.04191 | 0.677288 | NO | NO | NA |
| Ncl | ENSMUSG( | ENSMUSG( | 0.435387 | 0.001024 | YES | NO | NA |
| Arhgef1 | ENSMUSG( | ENSMUSG( | -0.21296 | 1.24E-05 | YES | NO | NA |
| mt-Nd4 | ENSMUSG( | ENSMUSG( | 0.034788 | 0.717688 | NO | NO | NA |
| Ldha | ENSMUSG( | ENSMUSG( | 0.614844 | 2.10E-06 | YES | NO | NA |
| Zfp36l2 | ENSMUSG( | ENSMUSG( | 0.187785 | 0.280449 | NO | NO | NA |
| Tmsb10 | ENSMUSG( | ENSMUSG( | 0.134965 | 0.231335 | NO | NO | NA |
| Arhgap45 | ENSMUSG( | ENSMUSG( | -0.49914 | 3.06E-11 | YES | NO | NA |
| Tpm3 | ENSMUSG( | ENSMUSG( | 0.087585 | 0.155123 | NO | NO | NA |
| Gnai2 | ENSMUSG( | ENSMUSG( | -0.14818 | 0.025189 | YES | NO | NA |
| Rpl18a | ENSMUSG( | ENSMUSG( | 0.168007 | 0.048504 | YES | NO | NA |
| Tuba1b | ENSMUSG( | ENSMUSG( | 0.301526 | 3.67E-08 | YES | NO | NA |
| Mbnl1 | ENSMUSG( | ENSMUSG( | -0.27335 | 0.001196 | YES | NO | NA |
| Eif4g2 | ENSMUSG( | ENSMUSG( | 0.039615 | 0.552587 | NO | NO | NA |
| Calm1 | ENSMUSG( | ENSMUSG( | -0.05869 | 0.438698 | NO | NO | NA |
| Fmnl1 | ENSMUSG( | ENSMUSG( | -0.29312 | 1.44E-08 | YES | NO | NA |
| Itgb2 | ENSMUSG( | ENSMUSG( | -0.51899 | 3.73E-23 | YES | NO | NA |
| Gm28661 | ENSMUSG( | ENSMUSG( | 0.14154 | 0.443843 | NO | NO | NA |
| Tnfaip3 | ENSMUSG( | ENSMUSG( | 0.003045 | 0.992393 | NO | NO | NA |

| | | | | | | | |
|---|---|---|---|---|---|---|---|
| Lsp1 | ENSMUSG( | ENSMUSG( | -0.35417 | 0.000124 | YES | NO | NA |
| H3f3b | ENSMUSG( | ENSMUSG( | -0.2679 | 0.001151 | YES | NO | NA |
| Ikzf1 | ENSMUSG( | ENSMUSG( | -0.07877 | 0.382461 | NO | NO | NA |
| Ctsd | ENSMUSG( | ENSMUSG( | -0.44232 | 4.20E-15 | YES | NO | NA |
| Srrm2 | ENSMUSG( | ENSMUSG( | -0.30514 | 2.45E-10 | YES | NO | NA |
| Rpl6 | ENSMUSG( | ENSMUSG( | 0.367816 | 8.36E-09 | YES | NO | NA |
| Gimap4 | ENSMUSG( | ENSMUSG( | -0.49656 | 6.21E-12 | YES | NO | NA |
| Gm42047 | ENSMUSG( | ENSMUSG( | 0.895617 | 0.000855 | YES | NO | NA |
| Hnrnpf | ENSMUSG( | ENSMUSG( | -0.00163 | 0.985073 | NO | NO | NA |
| Rplp1 | ENSMUSG( | ENSMUSG( | 0.396541 | 1.66E-09 | YES | NO | NA |
| Rpl13 | ENSMUSG( | ENSMUSG( | 0.349814 | 3.77E-10 | YES | NO | NA |
| Rps4x | ENSMUSG( | ENSMUSG( | 0.31208 | 4.52E-06 | YES | NO | NA |
| Rpl5 | ENSMUSG( | ENSMUSG( | 0.303034 | 1.43E-07 | YES | NO | NA |
| Rpl41 | ENSMUSG( | ENSMUSG( | 0.636115 | 4.89E-09 | YES | NO | NA |
| mt-Rnr2 | ENSMUSG( | ENSMUSG( | -0.4188 | 2.82E-05 | YES | NO | NA |
| Rac2 | ENSMUSG( | ENSMUSG( | -0.30253 | 2.68E-08 | YES | NO | NA |
| Iqgap1 | ENSMUSG( | ENSMUSG( | -0.14617 | 0.008037 | YES | NO | NA |
| Rpl7a | ENSMUSG( | ENSMUSG( | 0.333565 | 0.00111 | YES | NO | NA |
| Rps9 | ENSMUSG( | ENSMUSG( | 0.332569 | 5.81E-10 | YES | NO | NA |
| Rps8 | ENSMUSG( | ENSMUSG( | 0.448226 | 1.48E-09 | YES | NO | NA |
| Ms4a4b | ENSMUSG( | ENSMUSG( | -0.44657 | 1.86E-08 | YES | NO | NA |
| Hnrnpa1 | ENSMUSG( | ENSMUSG( | 0.338146 | 0.009519 | YES | NO | NA |
| Ywhaz | ENSMUSG( | ENSMUSG( | -0.1195 | 0.063705 | NO | NO | NA |
| Cap1 | ENSMUSG( | ENSMUSG( | -0.27924 | 1.97E-06 | YES | NO | NA |
| Rpl18 | ENSMUSG( | ENSMUSG( | 0.331566 | 1.11E-07 | YES | NO | NA |
| Arhgdib | ENSMUSG( | ENSMUSG( | -0.16474 | 0.005896 | YES | NO | NA |
| Atp5b | ENSMUSG( | ENSMUSG( | 0.372445 | 1.10E-07 | YES | NO | NA |
| Plec | ENSMUSG( | ENSMUSG( | -0.44258 | 7.16E-09 | YES | NO | NA |
| Gm28437 | ENSMUSG( | ENSMUSG( | 0.072447 | 0.670593 | NO | NO | NA |
| Dock2 | ENSMUSG( | ENSMUSG( | -0.42957 | 1.15E-11 | YES | NO | NA |
| Hnrnpab | ENSMUSG( | ENSMUSG( | 0.296996 | 0.002313 | YES | NO | NA |
| Akna | ENSMUSG( | ENSMUSG( | -0.48003 | 2.91E-11 | YES | NO | NA |
| Rps16 | ENSMUSG( | ENSMUSG( | 0.177276 | 0.044155 | YES | NO | NA |
| Rpl28 | ENSMUSG( | ENSMUSG( | 0.456713 | 2.05E-05 | YES | NO | NA |
| Nkg7 | ENSMUSG( | ENSMUSG( | -0.67079 | 2.90E-20 | YES | NO | NA |
| Ybx1 | ENSMUSG( | ENSMUSG( | 0.382332 | 2.71E-05 | YES | NO | NA |
| H2az1 | ENSMUSG( | ENSMUSG( | 0.397521 | 2.46E-09 | YES | NO | NA |
| Rps7 | ENSMUSG( | ENSMUSG( | 0.156538 | 0.014842 | YES | NO | NA |
| Arhgdia | ENSMUSG( | ENSMUSG( | 0.046539 | 0.490566 | NO | NO | NA |
| Rpl11 | ENSMUSG( | ENSMUSG( | 0.334644 | 1.51E-09 | YES | NO | NA |
| Shisa5 | ENSMUSG( | ENSMUSG( | -0.54263 | 2.43E-13 | YES | NO | NA |
| Rpl12 | ENSMUSG( | ENSMUSG( | 0.344865 | 2.42E-07 | YES | NO | NA |
| Rpl32 | ENSMUSG( | ENSMUSG( | 0.356145 | 4.59E-07 | YES | NO | NA |
| Rps23 | ENSMUSG( | ENSMUSG( | 0.309037 | 5.70E-06 | YES | NO | NA |
| Rps11 | ENSMUSG( | ENSMUSG( | 0.167735 | 0.076929 | NO | NO | NA |
| mt-Co3 | ENSMUSG( | ENSMUSG( | 0.232089 | 0.058345 | NO | NO | NA |
| Top2a | ENSMUSG( | ENSMUSG( | 0.006101 | 0.950219 | NO | NO | NA |

| | | | | | | | |
|---|---|---|---|---|---|---|---|
| Fyn | ENSMUSG( | ENSMUSG( | -0.66212 | 4.10E-08 | YES | NO | NA |
| Atp5a1 | ENSMUSG( | ENSMUSG( | 0.417294 | 2.53E-08 | YES | NO | NA |
| Lck | ENSMUSG( | ENSMUSG( | -0.32107 | 9.32E-10 | YES | NO | NA |
| Serbp1 | ENSMUSG( | ENSMUSG( | 0.255366 | 0.022027 | YES | NO | NA |
| Prrc2b | ENSMUSG( | ENSMUSG( | -0.52086 | 2.11E-12 | YES | NO | NA |
| Rnf213 | ENSMUSG( | ENSMUSG( | -1.00981 | 2.94E-19 | YES | YES | Downregulated |
| Cnn2 | ENSMUSG( | ENSMUSG( | -0.20603 | 3.22E-05 | YES | NO | NA |
| Gm47283 | ENSMUSG( | ENSMUSG( | 0.097272 | 0.854133 | NO | NO | NA |
| Rps18 | ENSMUSG( | ENSMUSG( | 0.458916 | 1.36E-10 | YES | NO | NA |
| Itgb7 | ENSMUSG( | ENSMUSG( | -0.14499 | 0.007969 | YES | NO | NA |
| Rpl17 | ENSMUSG( | ENSMUSG( | 0.145025 | 0.091511 | NO | NO | NA |
| Nbeal2 | ENSMUSG( | ENSMUSG( | -0.48746 | 1.45E-12 | YES | NO | NA |
| Rpl7 | ENSMUSG( | ENSMUSG( | 0.277843 | 3.87E-06 | YES | NO | NA |
| Actr2 | ENSMUSG( | ENSMUSG( | -0.08795 | 0.094505 | NO | NO | NA |
| Kif21b | ENSMUSG( | ENSMUSG( | -0.43843 | 3.73E-10 | YES | NO | NA |
| Hmgb1 | ENSMUSG( | ENSMUSG( | 0.22582 | 0.022629 | YES | NO | NA |
| Rpl23a | ENSMUSG( | ENSMUSG( | 0.175506 | 0.02064 | YES | NO | NA |
| Ikzf3 | ENSMUSG( | ENSMUSG( | -0.54993 | 2.91E-11 | YES | NO | NA |
| Cdc42 | ENSMUSG( | ENSMUSG( | -0.27239 | 1.62E-08 | YES | NO | NA |
| Grk2 | ENSMUSG( | ENSMUSG( | -0.19049 | 0.000141 | YES | NO | NA |
| Jak1 | ENSMUSG( | ENSMUSG( | -0.60996 | 9.73E-12 | YES | NO | NA |
| Gvin1 | ENSMUSG( | ENSMUSG( | -0.84514 | 6.39E-13 | YES | NO | NA |
| Sf3b1 | ENSMUSG( | ENSMUSG( | -0.14899 | 0.010471 | YES | NO | NA |
| Gnas | ENSMUSG( | ENSMUSG( | -0.23212 | 5.50E-06 | YES | NO | NA |
| Rpl19 | ENSMUSG( | ENSMUSG( | 0.141255 | 0.001494 | YES | NO | NA |
| Npm1 | ENSMUSG( | ENSMUSG( | 0.693071 | 1.66E-08 | YES | NO | NA |
| Rps14 | ENSMUSG( | ENSMUSG( | 0.167714 | 0.013071 | YES | NO | NA |
| Arpc1b | ENSMUSG( | ENSMUSG( | -0.10152 | 0.006357 | YES | NO | NA |
| Arpc2 | ENSMUSG( | ENSMUSG( | -0.07103 | 0.268553 | NO | NO | NA |
| Sptan1 | ENSMUSG( | ENSMUSG( | -0.44134 | 3.43E-09 | YES | NO | NA |
| Nup210 | ENSMUSG( | ENSMUSG( | -0.10916 | 0.017881 | YES | NO | NA |
| Hnrnpk | ENSMUSG( | ENSMUSG( | 0.090969 | 0.087885 | NO | NO | NA |
| Rpl21 | ENSMUSG( | ENSMUSG( | 0.22194 | 0.004682 | YES | NO | NA |
| Jund | ENSMUSG( | ENSMUSG( | 0.240389 | 0.015774 | YES | NO | NA |
| Wnk1 | ENSMUSG( | ENSMUSG( | -0.45812 | 3.44E-14 | YES | NO | NA |
| Prrc2a | ENSMUSG( | ENSMUSG( | -0.14867 | 0.000622 | YES | NO | NA |
| Uba1 | ENSMUSG( | ENSMUSG( | 0.083168 | 0.148146 | NO | NO | NA |
| Adam19 | ENSMUSG( | ENSMUSG( | -0.17781 | 0.040573 | YES | NO | NA |
| Lmnb1 | ENSMUSG( | ENSMUSG( | 0.246813 | 0.00408 | YES | NO | NA |
| Rps12 | ENSMUSG( | ENSMUSG( | 0.489015 | 2.00E-09 | YES | NO | NA |
| Ly6e | ENSMUSG( | ENSMUSG( | 0.302439 | 0.001136 | YES | NO | NA |
| Rpl10 | ENSMUSG( | ENSMUSG( | 0.406884 | 3.32E-07 | YES | NO | NA |
| Myo1g | ENSMUSG( | ENSMUSG( | -0.26381 | 3.32E-08 | YES | NO | NA |
| Rpl10a | ENSMUSG( | ENSMUSG( | 0.43097 | 5.86E-06 | YES | NO | NA |
| Hspa5 | ENSMUSG( | ENSMUSG( | -0.04966 | 0.625461 | NO | NO | NA |
| Hsp90b1 | ENSMUSG( | ENSMUSG( | -0.16749 | 0.098458 | NO | NO | NA |
| Eif5a | ENSMUSG( | ENSMUSG( | 0.667647 | 1.21E-08 | YES | NO | NA |

| | | | | | | | |
|---|---|---|---|---|---|---|---|
| Kmt2d | ENSMUSG( | ENSMUSG( | -0.73933 | 1.00E-12 | YES | NO | NA |
| Uba52 | ENSMUSG( | ENSMUSG( | 0.179538 | 0.000211 | YES | NO | NA |
| Rps5 | ENSMUSG( | ENSMUSG( | 0.332123 | 1.15E-08 | YES | NO | NA |
| Pik3cd | ENSMUSG( | ENSMUSG( | -0.51206 | 2.10E-09 | YES | NO | NA |
| Eif3a | ENSMUSG( | ENSMUSG( | 0.16819 | 0.211315 | NO | NO | NA |
| Rpl23 | ENSMUSG( | ENSMUSG( | 0.342324 | 6.61E-07 | YES | NO | NA |
| Ezr | ENSMUSG( | ENSMUSG( | 0.165011 | 0.039177 | YES | NO | NA |
| Capzb | ENSMUSG( | ENSMUSG( | -0.11041 | 0.046892 | YES | NO | NA |
| Atp1a1 | ENSMUSG( | ENSMUSG( | -0.15353 | 0.004155 | YES | NO | NA |
| Calr | ENSMUSG( | ENSMUSG( | -0.12802 | 0.236987 | NO | NO | NA |
| Chd4 | ENSMUSG( | ENSMUSG( | 0.084472 | 0.25241 | NO | NO | NA |
| Hnrnpa0 | ENSMUSG( | ENSMUSG( | 0.30576 | 2.90E-05 | YES | NO | NA |
| Eif4a1 | ENSMUSG( | ENSMUSG( | 0.328795 | 0.000267 | YES | NO | NA |
| Rasa3 | ENSMUSG( | ENSMUSG( | -0.58016 | 1.95E-11 | YES | NO | NA |
| B4galnt1 | ENSMUSG( | ENSMUSG( | -0.0202 | 0.829819 | NO | NO | NA |
| Nlrc5 | ENSMUSG( | ENSMUSG( | -0.57847 | 3.58E-11 | YES | NO | NA |
| Rps24 | ENSMUSG( | ENSMUSG( | 0.058399 | 0.544716 | NO | NO | NA |
| Ptbp1 | ENSMUSG( | ENSMUSG( | 0.221898 | 0.000266 | YES | NO | NA |
| Plek | ENSMUSG( | ENSMUSG( | -0.27744 | 0.001331 | YES | NO | NA |
| Hnrnpl | ENSMUSG( | ENSMUSG( | 0.060194 | 0.447068 | NO | NO | NA |
| H2ac24 | ENSMUSG( | ENSMUSG( | 0.124319 | 0.365388 | NO | NO | NA |
| Son | ENSMUSG( | ENSMUSG( | -0.39447 | 2.97E-10 | YES | NO | NA |
| Neat1 | ENSMUSG( | ENSMUSG( | -0.78605 | 3.17E-22 | YES | NO | NA |
| Hsp90aa1 | ENSMUSG( | ENSMUSG( | 0.046891 | 0.722177 | NO | NO | NA |
| Gvin-ps7 | ENSMUSG( | ENSMUSG( | -0.72746 | 5.72E-17 | YES | NO | NA |
| Dnmt1 | ENSMUSG( | ENSMUSG( | -0.07125 | 0.415446 | NO | NO | NA |
| Rps15 | ENSMUSG( | ENSMUSG( | 0.181591 | 0.057009 | NO | NO | NA |
| Ucp2 | ENSMUSG( | ENSMUSG( | -0.16197 | 0.022345 | YES | NO | NA |
| Bhlhe40 | ENSMUSG( | ENSMUSG( | 0.349334 | 0.153498 | NO | NO | NA |
| Gimap3 | ENSMUSG( | ENSMUSG( | -0.71834 | 7.93E-18 | YES | NO | NA |
| Eef1g | ENSMUSG( | ENSMUSG( | 0.474553 | 3.62E-13 | YES | NO | NA |
| Oaz1 | ENSMUSG( | ENSMUSG( | 0.03939 | 0.618667 | NO | NO | NA |
| Fau | ENSMUSG( | ENSMUSG( | 0.079852 | 0.374662 | NO | NO | NA |
| Rpl14 | ENSMUSG( | ENSMUSG( | 0.477312 | 1.28E-08 | YES | NO | NA |
| Pgk1 | ENSMUSG( | ENSMUSG( | 0.728633 | 2.19E-07 | YES | NO | NA |
| Aldoa ENS | ENSMUSG( | ENSMUSG( | 0.437862 | 0.00092 | YES | NO | NA |
| Septin9 | ENSMUSG( | ENSMUSG( | -0.17297 | 0.000755 | YES | NO | NA |
| Atp2a3 | ENSMUSG( | ENSMUSG( | -0.54352 | 2.59E-16 | YES | NO | NA |
| Hnrnpm | ENSMUSG( | ENSMUSG( | 0.127809 | 0.233451 | NO | NO | NA |
| Rps26 | ENSMUSG( | ENSMUSG( | 0.546158 | 1.60E-06 | YES | NO | NA |
| Akap13 | ENSMUSG( | ENSMUSG( | -0.33342 | 8.38E-07 | YES | NO | NA |
| Rpl9 | ENSMUSG( | ENSMUSG( | 0.167679 | 0.000831 | YES | NO | NA |
| Snrnp70 | ENSMUSG( | ENSMUSG( | 0.056802 | 0.608748 | NO | NO | NA |
| Hnrnpul1 | ENSMUSG( | ENSMUSG( | 0.076728 | 0.38644 | NO | NO | NA |
| Rpl27 | ENSMUSG( | ENSMUSG( | 0.157344 | 0.017891 | YES | NO | NA |
| Prkar1a | ENSMUSG( | ENSMUSG( | -0.07375 | 0.213885 | NO | NO | NA |
| Fus | ENSMUSG( | ENSMUSG( | 0.226154 | 0.145745 | NO | NO | NA |

| | | | | | | | |
|---|---|---|---|---|---|---|---|
| Rpl15 | ENSMUSG( | ENSMUSG( | 0.41974 | 4.21E-05 | YES | NO | NA |
| Gm10925 | ENSMUSG( | ENSMUSG( | 0.09452 | 0.594913 | NO | NO | NA |
| Rpl27a | ENSMUSG( | ENSMUSG( | 0.214634 | 0.021736 | YES | NO | NA |
| Sp100 | ENSMUSG( | ENSMUSG( | -0.61033 | 3.77E-14 | YES | NO | NA |
| Arhgap30 | ENSMUSG( | ENSMUSG( | -0.31116 | 2.28E-05 | YES | NO | NA |
| Selplg | ENSMUSG( | ENSMUSG( | -0.32268 | 2.32E-06 | YES | NO | NA |
| Fam107b | ENSMUSG( | ENSMUSG( | -0.08328 | 0.368134 | NO | NO | NA |
| Chsy1 | ENSMUSG( | ENSMUSG( | -0.85322 | 1.62E-19 | YES | NO | NA |
| Eif4g1 | ENSMUSG( | ENSMUSG( | 0.155261 | 0.030763 | YES | NO | NA |
| Gpi1 | ENSMUSG( | ENSMUSG( | 0.157107 | 0.026748 | YES | NO | NA |
| Sh2d2a | ENSMUSG( | ENSMUSG( | -0.13615 | 0.171311 | NO | NO | NA |
| Ppp1ca | ENSMUSG( | ENSMUSG( | -0.03654 | 0.350226 | NO | NO | NA |
| Set | ENSMUSG( | ENSMUSG( | 0.226847 | 0.013228 | YES | NO | NA |
| Tmpo | ENSMUSG( | ENSMUSG( | 0.106829 | 0.156375 | NO | NO | NA |
| Cyfip2 | ENSMUSG( | ENSMUSG( | -0.08951 | 0.014188 | YES | NO | NA |
| Sfpq | ENSMUSG( | ENSMUSG( | -0.05988 | 0.400548 | NO | NO | NA |
| Bin2 | ENSMUSG( | ENSMUSG( | -0.51335 | 4.99E-09 | YES | NO | NA |
| Zyx | ENSMUSG( | ENSMUSG( | -0.47753 | 5.86E-10 | YES | NO | NA |
| Rps20 | ENSMUSG( | ENSMUSG( | 0.562501 | 1.59E-11 | YES | NO | NA |
| Trbc2 | ENSMUSG( | ENSMUSG( | -0.19348 | 0.050463 | NO | NO | NA |
| H2-Q6 | ENSMUSG( | ENSMUSG( | -0.40947 | 6.26E-05 | YES | NO | NA |
| Hmgn2 | ENSMUSG( | ENSMUSG( | 0.273704 | 0.003642 | YES | NO | NA |
| Anp32b | ENSMUSG( | ENSMUSG( | 0.193489 | 0.267916 | NO | NO | NA |
| Gvin2 | ENSMUSG( | ENSMUSG( | -0.86863 | 1.90E-29 | YES | NO | NA |
| Prpf8 | ENSMUSG( | ENSMUSG( | -0.04409 | 0.380648 | NO | NO | NA |
| Myo1f | ENSMUSG( | ENSMUSG( | -0.67001 | 3.17E-22 | YES | NO | NA |
| Rap1b | ENSMUSG( | ENSMUSG( | -0.34429 | 9.88E-08 | YES | NO | NA |
| Actn4 | ENSMUSG( | ENSMUSG( | 0.060143 | 0.439816 | NO | NO | NA |
| Rps10 | ENSMUSG( | ENSMUSG( | 0.235315 | 6.44E-05 | YES | NO | NA |
| Slc25a5 | ENSMUSG( | ENSMUSG( | 0.513266 | 2.89E-08 | YES | NO | NA |
| Dgkz | ENSMUSG( | ENSMUSG( | -0.21079 | 0.00013 | YES | NO | NA |
| Arpc5 | ENSMUSG( | ENSMUSG( | -0.04918 | 0.500632 | NO | NO | NA |
| mt-Nd2 | ENSMUSG( | ENSMUSG( | -0.01057 | 0.918982 | NO | NO | NA |
| Anxa2 | ENSMUSG( | ENSMUSG( | 0.253987 | 0.0391 | YES | NO | NA |
| Cd8b1 | ENSMUSG( | ENSMUSG( | -0.45537 | 4.35E-12 | YES | NO | NA |
| Arid1a | ENSMUSG( | ENSMUSG( | -0.53591 | 1.07E-16 | YES | NO | NA |
| Klf2 | ENSMUSG( | ENSMUSG( | -0.4841 | 3.20E-06 | YES | NO | NA |
| Pdia3 | ENSMUSG( | ENSMUSG( | -0.01784 | 0.829991 | NO | NO | NA |
| Sub1 | ENSMUSG( | ENSMUSG( | -0.20625 | 0.188406 | NO | NO | NA |
| Rpl31 | ENSMUSG( | ENSMUSG( | 0.457525 | 5.63E-08 | YES | NO | NA |
| Prrc2c | ENSMUSG( | ENSMUSG( | -0.31041 | 2.90E-06 | YES | NO | NA |
| Tmbim6 | ENSMUSG( | ENSMUSG( | -0.23204 | 0.000103 | YES | NO | NA |
| Hnrnpul2 | ENSMUSG( | ENSMUSG( | -0.12286 | 0.271399 | NO | NO | NA |
| Grk6 | ENSMUSG( | ENSMUSG( | 0.116876 | 0.272132 | NO | NO | NA |
| Adgre5 | ENSMUSG( | ENSMUSG( | -0.34004 | 5.20E-13 | YES | NO | NA |
| Hnrnph1 | ENSMUSG( | ENSMUSG( | 0.094033 | 0.322221 | NO | NO | NA |
| Tap1 | ENSMUSG( | ENSMUSG( | -0.23114 | 0.000795 | YES | NO | NA |

| | | | | | | | |
|---|---|---|---|---|---|---|---|
| Dync1h1 | ENSMUSG( | ENSMUSG( | -0.38284 | 1.86E-13 | YES | NO | NA |
| Septin1 | ENSMUSG( | ENSMUSG( | -0.18204 | 9.21E-05 | YES | NO | NA |
| Serinc3 | ENSMUSG( | ENSMUSG( | -0.02365 | 0.831335 | NO | NO | NA |
| Btg1 | ENSMUSG( | ENSMUSG( | -0.44962 | 0.000524 | YES | NO | NA |
| Tle5 | ENSMUSG( | ENSMUSG( | -0.16946 | 0.011358 | YES | NO | NA |
| Kpnb1 | ENSMUSG( | ENSMUSG( | 0.295092 | 9.39E-06 | YES | NO | NA |
| Rps13 | ENSMUSG( | ENSMUSG( | 0.121849 | 0.13582 | NO | NO | NA |
| Itm2b | ENSMUSG( | ENSMUSG( | -0.2643 | 0.000199 | YES | NO | NA |
| Ablim1 | ENSMUSG( | ENSMUSG( | -0.43129 | 1.14E-08 | YES | NO | NA |
| Myl6 | ENSMUSG( | ENSMUSG( | -0.20988 | 0.001916 | YES | NO | NA |
| Arl6ip1 | ENSMUSG( | ENSMUSG( | -0.05977 | 0.436468 | NO | NO | NA |
| Rad21 | ENSMUSG( | ENSMUSG( | -0.02635 | 0.739482 | NO | NO | NA |
| Stk10 | ENSMUSG( | ENSMUSG( | -0.5787 | 1.34E-13 | YES | NO | NA |
| Mcm6 | ENSMUSG( | ENSMUSG( | 0.027578 | 0.664861 | NO | NO | NA |
| Nap1l1 | ENSMUSG( | ENSMUSG( | 0.236549 | 0.003308 | YES | NO | NA |
| Tpr | ENSMUSG( | ENSMUSG( | -0.07413 | 0.410802 | NO | NO | NA |
| Dek | ENSMUSG( | ENSMUSG( | 0.191238 | 0.047397 | YES | NO | NA |
| Ran | ENSMUSG( | ENSMUSG( | 0.293917 | 5.56E-05 | YES | NO | NA |
| Ankrd11 | ENSMUSG( | ENSMUSG( | -0.36596 | 8.19E-05 | YES | NO | NA |
| Samhd1 | ENSMUSG( | ENSMUSG( | -0.49115 | 3.80E-14 | YES | NO | NA |
| Kmt2a | ENSMUSG( | ENSMUSG( | -0.55765 | 1.66E-10 | YES | NO | NA |
| Ddx17 | ENSMUSG( | ENSMUSG( | -0.44304 | 8.21E-11 | YES | NO | NA |
| Sema4d | ENSMUSG( | ENSMUSG( | -0.09521 | 0.082945 | NO | NO | NA |
| Ripor2 | ENSMUSG( | ENSMUSG( | -0.75536 | 1.61E-11 | YES | NO | NA |
| Fosb | ENSMUSG( | ENSMUSG( | 1.479962 | 0.000875 | YES | YES | Upregulated |
| Wdr1 | ENSMUSG( | ENSMUSG( | -0.14286 | 0.000655 | YES | NO | NA |
| Ywhah | ENSMUSG( | ENSMUSG( | -0.10584 | 0.34315 | NO | NO | NA |
| Cotl1 | ENSMUSG( | ENSMUSG( | -0.31251 | 1.29E-06 | YES | NO | NA |
| Rasgrp2 | ENSMUSG( | ENSMUSG( | -0.34494 | 3.54E-05 | YES | NO | NA |
| Ncor1 | ENSMUSG( | ENSMUSG( | -0.38701 | 1.31E-08 | YES | NO | NA |
| Rplp2 | ENSMUSG( | ENSMUSG( | 0.272526 | 3.31E-06 | YES | NO | NA |
| Cd3e | ENSMUSG( | ENSMUSG( | -0.51024 | 2.14E-13 | YES | NO | NA |
| Grap2 | ENSMUSG( | ENSMUSG( | -0.46478 | 1.78E-14 | YES | NO | NA |
| Thrap3 | ENSMUSG( | ENSMUSG( | 0.118441 | 0.191677 | NO | NO | NA |
| Nono | ENSMUSG( | ENSMUSG( | 0.022361 | 0.77617 | NO | NO | NA |
| Rps15a | ENSMUSG( | ENSMUSG( | 0.109613 | 0.178072 | NO | NO | NA |
| Fnbp1 | ENSMUSG( | ENSMUSG( | -0.30623 | 3.46E-08 | YES | NO | NA |
| Capza1 | ENSMUSG( | ENSMUSG( | -0.25059 | 4.19E-09 | YES | NO | NA |
| Srsf2 | ENSMUSG( | ENSMUSG( | 0.301671 | 9.39E-06 | YES | NO | NA |
| Srsf5 | ENSMUSG( | ENSMUSG( | 0.088339 | 0.509559 | NO | NO | NA |
| Polr2a | ENSMUSG( | ENSMUSG( | -0.2711 | 1.28E-06 | YES | NO | NA |
| Trac | ENSMUSG( | ENSMUSG( | -0.17516 | 0.068933 | NO | NO | NA |
| Vasp | ENSMUSG( | ENSMUSG( | -0.31717 | 1.33E-12 | YES | NO | NA |
| Pfkp | ENSMUSG( | ENSMUSG( | 0.041226 | 0.582357 | NO | NO | NA |
| Cntrl | ENSMUSG( | ENSMUSG( | 0.023488 | 0.800931 | NO | NO | NA |
| Prkch | ENSMUSG( | ENSMUSG( | -0.22016 | 0.00795 | YES | NO | NA |
| Hnrnpd | ENSMUSG( | ENSMUSG( | 0.232386 | 0.177177 | NO | NO | NA |

| | | | | | | | |
|---|---|---|---|---|---|---|---|
| Itgb1 | ENSMUSG( | ENSMUSG( | -0.31348 | 0.002304 | YES | NO | NA |
| Il18r1 | ENSMUSG( | ENSMUSG( | 0.480629 | 0.001463 | YES | NO | NA |
| Eef1b2 | ENSMUSG( | ENSMUSG( | 0.54986 | 6.79E-11 | YES | NO | NA |
| Sh3kbp1 | ENSMUSG( | ENSMUSG( | -0.3287 | 2.75E-08 | YES | NO | NA |
| Dock10 | ENSMUSG( | ENSMUSG( | -0.34629 | 2.97E-09 | YES | NO | NA |
| Ncapd2 | ENSMUSG( | ENSMUSG( | 0.271239 | 0.000355 | YES | NO | NA |
| H2-Q4 | ENSMUSG( | ENSMUSG( | -0.26286 | 0.011366 | YES | NO | NA |
| Wipf1 | ENSMUSG( | ENSMUSG( | -0.35722 | 6.14E-06 | YES | NO | NA |
| Arsb | ENSMUSG( | ENSMUSG( | 0.176268 | 0.157988 | NO | NO | NA |
| Cd3g | ENSMUSG( | ENSMUSG( | -0.35981 | 1.25E-06 | YES | NO | NA |
| Rpl30 | ENSMUSG( | ENSMUSG( | 0.07222 | 0.6007 | NO | NO | NA |
| Rpl35 | ENSMUSG( | ENSMUSG( | 0.514107 | 4.21E-08 | YES | NO | NA |
| Rpl29 | ENSMUSG( | ENSMUSG( | 0.382477 | 1.95E-05 | YES | NO | NA |
| Huwe1 | ENSMUSG( | ENSMUSG( | -0.1696 | 1.77E-06 | YES | NO | NA |
| Sf3b2 | ENSMUSG( | ENSMUSG( | 0.064212 | 0.419671 | NO | NO | NA |
| Naca | ENSMUSG( | ENSMUSG( | 0.206146 | 0.00988 | YES | NO | NA |
| Rps17 | ENSMUSG( | ENSMUSG( | 0.622587 | 5.74E-08 | YES | NO | NA |
| Psap | ENSMUSG( | ENSMUSG( | -0.37132 | 2.18E-10 | YES | NO | NA |
| Rpl24 | ENSMUSG( | ENSMUSG( | 0.250806 | 4.44E-06 | YES | NO | NA |
| Sipa1 | ENSMUSG( | ENSMUSG( | -0.37943 | 2.05E-07 | YES | NO | NA |
| Gzma | ENSMUSG( | ENSMUSG( | -0.48091 | 0.518069 | NO | NO | NA |
| Ptp4a2 | ENSMUSG( | ENSMUSG( | 0.045499 | 0.599614 | NO | NO | NA |
| Caprin1 | ENSMUSG( | ENSMUSG( | 0.125449 | 0.024489 | YES | NO | NA |
| Srsf3 | ENSMUSG( | ENSMUSG( | 0.120104 | 0.12658 | NO | NO | NA |
| Ms4a6b | ENSMUSG( | ENSMUSG( | -0.40462 | 2.95E-10 | YES | NO | NA |
| Rps19 | ENSMUSG( | ENSMUSG( | 0.333435 | 6.73E-08 | YES | NO | NA |
| Acin1 | ENSMUSG( | ENSMUSG( | -0.19309 | 0.093322 | NO | NO | NA |
| Stat1 | ENSMUSG( | ENSMUSG( | -0.60299 | 5.89E-12 | YES | NO | NA |
| AI506816 | ENSMUSG( | ENSMUSG( | 0.35691 | 0.000126 | YES | NO | NA |
| Lrrfip1 | ENSMUSG( | ENSMUSG( | -0.26042 | 1.69E-05 | YES | NO | NA |
| Gimap6 | ENSMUSG( | ENSMUSG( | -0.37422 | 2.79E-06 | YES | NO | NA |
| Rps25 | ENSMUSG( | ENSMUSG( | 0.52232 | 1.25E-06 | YES | NO | NA |
| Efhd2 | ENSMUSG( | ENSMUSG( | 0.049631 | 0.523326 | NO | NO | NA |
| Fryl | ENSMUSG( | ENSMUSG( | -0.83658 | 1.45E-12 | YES | NO | NA |
| Gdi2 | ENSMUSG( | ENSMUSG( | 0.161739 | 0.006015 | YES | NO | NA |
| Psmb8 | ENSMUSG( | ENSMUSG( | 0.004382 | 0.912407 | NO | NO | NA |
| Ewsr1 | ENSMUSG( | ENSMUSG( | 0.034413 | 0.783319 | NO | NO | NA |
| Ptbp3 | ENSMUSG( | ENSMUSG( | -0.23497 | 0.000698 | YES | NO | NA |
| Jun | ENSMUSG( | ENSMUSG( | 1.059687 | 0.001254 | YES | YES | Upregulated |
| Btf3 | ENSMUSG( | ENSMUSG( | 0.226798 | 0.006963 | YES | NO | NA |
| Baz1b | ENSMUSG( | ENSMUSG( | -0.08716 | 0.240619 | NO | NO | NA |
| Kcnab2 | ENSMUSG( | ENSMUSG( | -0.13216 | 0.005509 | YES | NO | NA |
| Sf1 | ENSMUSG( | ENSMUSG( | -0.04809 | 0.437106 | NO | NO | NA |
| Stk17b | ENSMUSG( | ENSMUSG( | -0.35756 | 0.000198 | YES | NO | NA |
| Luc7l2 | ENSMUSG( | ENSMUSG( | -0.07822 | 0.430295 | NO | NO | NA |
| Stmn1 | ENSMUSG( | ENSMUSG( | 0.20995 | 0.004505 | YES | NO | NA |
| Hip1r | ENSMUSG( | ENSMUSG( | 0.023657 | 0.810346 | NO | NO | NA |

| | | | | | | | |
|---|---|---|---|---|---|---|---|
| Septin11 | ENSMUSG( | ENSMUSG( | -0.24595 | 0.000146 | YES | NO | NA |
| Birc6 | ENSMUSG( | ENSMUSG( | -0.45379 | 2.55E-13 | YES | NO | NA |
| Baiap3 | ENSMUSG( | ENSMUSG( | -0.79943 | 7.79E-12 | YES | NO | NA |
| Junb | ENSMUSG( | ENSMUSG( | 0.408289 | 0.018418 | YES | NO | NA |
| Vcp | ENSMUSG( | ENSMUSG( | 0.203808 | 6.69E-05 | YES | NO | NA |
| S100a4 | ENSMUSG( | ENSMUSG( | -0.30757 | 0.003211 | YES | NO | NA |
| Lbr | ENSMUSG( | ENSMUSG( | -0.02099 | 0.775027 | NO | NO | NA |
| Ppp1cc | ENSMUSG( | ENSMUSG( | 0.15826 | 0.015289 | YES | NO | NA |
| Nop53 | ENSMUSG( | ENSMUSG( | 0.019912 | 0.878599 | NO | NO | NA |
| Ube2i | ENSMUSG( | ENSMUSG( | -0.09247 | 0.07691 | NO | NO | NA |
| Srsf1 | ENSMUSG( | ENSMUSG( | 0.122988 | 0.04794 | YES | NO | NA |
| Nsmaf | ENSMUSG( | ENSMUSG( | -0.37017 | 6.66E-06 | YES | NO | NA |
| Ywhae | ENSMUSG( | ENSMUSG( | 0.217322 | 0.006794 | YES | NO | NA |
| Smc4 | ENSMUSG( | ENSMUSG( | -0.27476 | 0.295575 | NO | NO | NA |
| Ube2d3 | ENSMUSG( | ENSMUSG( | -0.1193 | 0.056166 | NO | NO | NA |
| Tpm4 | ENSMUSG( | ENSMUSG( | -0.02035 | 0.835725 | NO | NO | NA |
| Ogt | ENSMUSG( | ENSMUSG( | -0.62661 | 8.11E-09 | YES | NO | NA |
| Mcm5 | ENSMUSG( | ENSMUSG( | 0.189771 | 0.017669 | YES | NO | NA |
| Sptbn1 | ENSMUSG( | ENSMUSG( | 0.061084 | 0.543023 | NO | NO | NA |
| Clic1 | ENSMUSG( | ENSMUSG( | 0.009624 | 0.909884 | NO | NO | NA |
| Anp32e | ENSMUSG( | ENSMUSG( | 0.315452 | 0.009486 | YES | NO | NA |
| Calm3 | ENSMUSG( | ENSMUSG( | -0.02774 | 0.736727 | NO | NO | NA |
| Hcls1 | ENSMUSG( | ENSMUSG( | -0.24697 | 0.000633 | YES | NO | NA |
| Eif4b | ENSMUSG( | ENSMUSG( | 0.106057 | 0.131154 | NO | NO | NA |
| Jak3 | ENSMUSG( | ENSMUSG( | -0.14684 | 0.019722 | YES | NO | NA |
| Ddx3x | ENSMUSG( | ENSMUSG( | 0.358764 | 1.26E-05 | YES | NO | NA |
| Diaph1 | ENSMUSG( | ENSMUSG( | -0.19032 | 2.98E-05 | YES | NO | NA |
| Pkn1 | ENSMUSG( | ENSMUSG( | -0.21354 | 0.000201 | YES | NO | NA |
| Lasp1 | ENSMUSG( | ENSMUSG( | -0.49084 | 1.45E-14 | YES | NO | NA |
| Ifi203 | ENSMUSG( | ENSMUSG( | -0.73386 | 4.39E-15 | YES | NO | NA |
| Rps27a | ENSMUSG( | ENSMUSG( | 0.330121 | 5.64E-09 | YES | NO | NA |
| Tnfrsf1b | ENSMUSG( | ENSMUSG( | -3.70149 | 0 | YES | YES | Downregulated |
| Cd82 | ENSMUSG( | ENSMUSG( | -0.06469 | 0.539291 | NO | NO | NA |
| Ccdc88c | ENSMUSG( | ENSMUSG( | -0.45402 | 2.27E-08 | YES | NO | NA |
| Elf4 | ENSMUSG( | ENSMUSG( | -0.52981 | 1.05E-13 | YES | NO | NA |
| Trip12 | ENSMUSG( | ENSMUSG( | -0.3541 | 1.51E-10 | YES | NO | NA |
| Atp1b3 | ENSMUSG( | ENSMUSG( | -0.35464 | 2.18E-08 | YES | NO | NA |
| Rpl26 | ENSMUSG( | ENSMUSG( | 0.33133 | 2.55E-07 | YES | NO | NA |
| Mllt6 | ENSMUSG( | ENSMUSG( | -0.33367 | 1.45E-06 | YES | NO | NA |
| Ubr4 | ENSMUSG( | ENSMUSG( | -0.45888 | 6.64E-12 | YES | NO | NA |
| Hcfc1 | ENSMUSG( | ENSMUSG( | -0.24488 | 1.78E-08 | YES | NO | NA |
| Ly6c2 | ENSMUSG( | ENSMUSG( | -0.04536 | 0.757268 | NO | NO | NA |
| U2af2 | ENSMUSG( | ENSMUSG( | 0.13983 | 0.046378 | YES | NO | NA |
| Rhoa | ENSMUSG( | ENSMUSG( | -0.22185 | 0.000195 | YES | NO | NA |
| Pim1 | ENSMUSG( | ENSMUSG( | -0.18379 | 4.99E-05 | YES | NO | NA |
| Klf6 | ENSMUSG( | ENSMUSG( | -0.00975 | 0.973881 | NO | NO | NA |
| Cnot1 | ENSMUSG( | ENSMUSG( | -0.06718 | 0.209547 | NO | NO | NA |

| | | | | | | | |
|---|---|---|---|---|---|---|---|
| Mknk2 | ENSMUSG( | ENSMUSG( | -0.27971 | 5.44E-08 | YES | NO | NA |
| Ppp1r18 | ENSMUSG( | ENSMUSG( | -0.40844 | 7.64E-09 | YES | NO | NA |
| Ankrd44 | ENSMUSG( | ENSMUSG( | -0.50632 | 2.54E-19 | YES | NO | NA |
| Slc25a3 | ENSMUSG( | ENSMUSG( | 0.225301 | 0.00142 | YES | NO | NA |
| Cdc42se1 | ENSMUSG( | ENSMUSG( | -0.21975 | 4.71E-10 | YES | NO | NA |
| Zdhhc18 | ENSMUSG( | ENSMUSG( | -0.38268 | 7.74E-08 | YES | NO | NA |
| Eif4h | ENSMUSG( | ENSMUSG( | 0.158859 | 0.000281 | YES | NO | NA |
| Ppp1r12a | ENSMUSG( | ENSMUSG( | -0.72027 | 3.23E-14 | YES | NO | NA |
| Lfng | ENSMUSG( | ENSMUSG( | -0.85884 | 6.30E-15 | YES | NO | NA |
| Lnpep | ENSMUSG( | ENSMUSG( | -0.46302 | 4.27E-09 | YES | NO | NA |
| Ptpn22 | ENSMUSG( | ENSMUSG( | -0.26878 | 0.000867 | YES | NO | NA |
| Dock8 | ENSMUSG( | ENSMUSG( | -0.3606 | 4.90E-09 | YES | NO | NA |
| Rpl37a | ENSMUSG( | ENSMUSG( | 0.078133 | 0.389469 | NO | NO | NA |
| Bzw1 | ENSMUSG( | ENSMUSG( | 0.127724 | 0.034705 | YES | NO | NA |
| Zc3hav1 | ENSMUSG( | ENSMUSG( | -0.35042 | 7.13E-09 | YES | NO | NA |
| Eif1 | ENSMUSG( | ENSMUSG( | -0.20931 | 0.001366 | YES | NO | NA |
| Tubb4b | ENSMUSG( | ENSMUSG( | 0.28221 | 0.000166 | YES | NO | NA |
| Il18rap | ENSMUSG( | ENSMUSG( | 0.301665 | 0.041555 | YES | NO | NA |
| Rbm39 | ENSMUSG( | ENSMUSG( | -0.25786 | 6.30E-05 | YES | NO | NA |
| Fam120a | ENSMUSG( | ENSMUSG( | 0.037914 | 0.579415 | NO | NO | NA |
| Ifngr1 | ENSMUSG( | ENSMUSG( | 0.388126 | 0.009653 | YES | NO | NA |
| H2-T23 | ENSMUSG( | ENSMUSG( | -0.56825 | 8.37E-17 | YES | NO | NA |
| Cd52 | ENSMUSG( | ENSMUSG( | -0.02112 | 0.827284 | NO | NO | NA |
| Celf2 | ENSMUSG( | ENSMUSG( | 0.028066 | 0.755183 | NO | NO | NA |
| Hp1bp3 | ENSMUSG( | ENSMUSG( | -0.16426 | 0.027885 | YES | NO | NA |
| Kmt2e | ENSMUSG( | ENSMUSG( | -0.48004 | 1.08E-07 | YES | NO | NA |
| Adcy7 | ENSMUSG( | ENSMUSG( | -0.39296 | 6.30E-10 | YES | NO | NA |
| Tagln2 | ENSMUSG( | ENSMUSG( | 0.326559 | 0.001135 | YES | NO | NA |
| Rapgef1 | ENSMUSG( | ENSMUSG( | -0.23058 | 0.000247 | YES | NO | NA |
| Smg1 | ENSMUSG( | ENSMUSG( | -0.45936 | 9.13E-11 | YES | NO | NA |
| Mta2 | ENSMUSG( | ENSMUSG( | 0.004684 | 0.939136 | NO | NO | NA |
| Eif3c | ENSMUSG( | ENSMUSG( | 0.203467 | 0.018083 | YES | NO | NA |
| Pdcd1 | ENSMUSG( | ENSMUSG( | 0.119055 | 0.44084 | NO | NO | NA |
| Tpi1 | ENSMUSG( | ENSMUSG( | 0.960576 | 0.000234 | YES | NO | NA |
| Dazap2 | ENSMUSG( | ENSMUSG( | -0.02714 | 0.790471 | NO | NO | NA |
| Dhx9 | ENSMUSG( | ENSMUSG( | 0.113127 | 0.134433 | NO | NO | NA |
| Canx | ENSMUSG( | ENSMUSG( | 0.043376 | 0.680639 | NO | NO | NA |
| Gabpb2 | ENSMUSG( | ENSMUSG( | -0.12848 | 0.121345 | NO | NO | NA |
| Septin7 | ENSMUSG( | ENSMUSG( | 0.00515 | 0.963858 | NO | NO | NA |
| Anapc5 | ENSMUSG( | ENSMUSG( | 0.308317 | 0.000298 | YES | NO | NA |
| Arhgap9 | ENSMUSG( | ENSMUSG( | -0.12048 | 0.057485 | NO | NO | NA |
| Rps27 | ENSMUSG( | ENSMUSG( | -0.07452 | 0.567401 | NO | NO | NA |
| Smc1a | ENSMUSG( | ENSMUSG( | 0.086748 | 0.484356 | NO | NO | NA |
| Rasal3 | ENSMUSG( | ENSMUSG( | -0.279 | 1.09E-05 | YES | NO | NA |
| Osbpl3 | ENSMUSG( | ENSMUSG( | -0.38541 | 4.24E-11 | YES | NO | NA |
| Ssrp1 | ENSMUSG( | ENSMUSG( | 0.227467 | 0.006133 | YES | NO | NA |
| S100a6 | ENSMUSG( | ENSMUSG( | -0.28323 | 0.000785 | YES | NO | NA |

| | | | | | | | |
|---|---|---|---|---|---|---|---|
| Fxyd5 | ENSMUSG( | ENSMUSG( | -0.11102 | 0.170248 | NO | NO | NA |
| Casp3 | ENSMUSG( | ENSMUSG( | 0.342424 | 0.001362 | YES | NO | NA |
| Tap2 | ENSMUSG( | ENSMUSG( | -0.15992 | 1.75E-05 | YES | NO | NA |
| mt-Rnr1 | ENSMUSG( | ENSMUSG( | -0.56376 | 4.18E-07 | YES | NO | NA |
| Tox | ENSMUSG( | ENSMUSG( | -0.50343 | 0.000483 | YES | NO | NA |
| Maz | ENSMUSG( | ENSMUSG( | -0.07618 | 0.261706 | NO | NO | NA |
| Limd2 | ENSMUSG( | ENSMUSG( | -0.27629 | 3.95E-05 | YES | NO | NA |
| Fkbp5 | ENSMUSG( | ENSMUSG( | 0.33073 | 9.92E-05 | YES | NO | NA |
| Nfatc1 | ENSMUSG( | ENSMUSG( | -0.49686 | 2.99E-07 | YES | NO | NA |
| Prelid1 | ENSMUSG( | ENSMUSG( | 0.334493 | 0.000215 | YES | NO | NA |
| Ddx39b | ENSMUSG( | ENSMUSG( | 0.070295 | 0.143737 | NO | NO | NA |
| Rab8b | ENSMUSG( | ENSMUSG( | -0.27732 | 7.79E-05 | YES | NO | NA |
| Pcbp2 | ENSMUSG( | ENSMUSG( | 0.02986 | 0.593156 | NO | NO | NA |
| Eif3h | ENSMUSG( | ENSMUSG( | 0.159835 | 0.043927 | YES | NO | NA |
| Rinl | ENSMUSG( | ENSMUSG( | -0.34291 | 6.39E-09 | YES | NO | NA |
| Sh3bgrl3 | ENSMUSG( | ENSMUSG( | 0.091068 | 0.06603 | NO | NO | NA |
| Copa | ENSMUSG( | ENSMUSG( | -0.13391 | 0.004869 | YES | NO | NA |
| Pcna | ENSMUSG( | ENSMUSG( | 0.1522 | 0.005637 | YES | NO | NA |
| Rrm1 | ENSMUSG( | ENSMUSG( | 0.415988 | 2.73E-07 | YES | NO | NA |
| Slc44a2 | ENSMUSG( | ENSMUSG( | -0.26391 | 9.68E-08 | YES | NO | NA |
| Ubc | ENSMUSG( | ENSMUSG( | -0.38567 | 6.32E-08 | YES | NO | NA |
| Cct2 | ENSMUSG( | ENSMUSG( | 0.295773 | 7.81E-07 | YES | NO | NA |
| Rpl34 | ENSMUSG( | ENSMUSG( | -0.55233 | 5.03E-16 | YES | NO | NA |
| Rps28 | ENSMUSG( | ENSMUSG( | 0.47006 | 3.95E-08 | YES | NO | NA |
| Pdcd6ip | ENSMUSG( | ENSMUSG( | -0.24607 | 3.39E-05 | YES | NO | NA |
| Stk24 | ENSMUSG( | ENSMUSG( | -0.16727 | 0.019845 | YES | NO | NA |
| Eif3b | ENSMUSG( | ENSMUSG( | 0.429015 | 1.08E-05 | YES | NO | NA |
| Hip1 | ENSMUSG( | ENSMUSG( | -0.37009 | 4.81E-06 | YES | NO | NA |
| ENSMUSG | ENSMUSG( | ENSMUSG( | 0.481697 | 2.52E-05 | YES | NO | NA |
| S100a10 | ENSMUSG( | ENSMUSG( | -0.19947 | 0.000213 | YES | NO | NA |
| Rps29 | ENSMUSG( | ENSMUSG( | 0.376416 | 6.19E-06 | YES | NO | NA |
| Stk4 | ENSMUSG( | ENSMUSG( | -0.30371 | 3.45E-08 | YES | NO | NA |
| Brd2 | ENSMUSG( | ENSMUSG( | -0.13056 | 0.00517 | YES | NO | NA |
| Igf2r | ENSMUSG( | ENSMUSG( | -0.33905 | 1.49E-06 | YES | NO | NA |
| Ptprcap | ENSMUSG( | ENSMUSG( | -0.21506 | 0.000235 | YES | NO | NA |
| Pcbp1 | ENSMUSG( | ENSMUSG( | 0.135836 | 0.088085 | NO | NO | NA |
| Tapbp | ENSMUSG( | ENSMUSG( | -0.26612 | 1.31E-07 | YES | NO | NA |
| Mcm7 | ENSMUSG( | ENSMUSG( | 0.387732 | 9.45E-07 | YES | NO | NA |
| Ybx3 | ENSMUSG( | ENSMUSG( | 0.469978 | 9.21E-07 | YES | NO | NA |
| Zap70 | ENSMUSG( | ENSMUSG( | -0.11688 | 0.036576 | YES | NO | NA |
| Gnptab | ENSMUSG( | ENSMUSG( | 0.120748 | 0.229509 | NO | NO | NA |
| Ddb1 | ENSMUSG( | ENSMUSG( | 0.200443 | 0.00319 | YES | NO | NA |
| Ddx6 | ENSMUSG( | ENSMUSG( | -0.32039 | 7.51E-06 | YES | NO | NA |
| Lag3 | ENSMUSG( | ENSMUSG( | -0.13904 | 0.447057 | NO | NO | NA |
| Raly | ENSMUSG( | ENSMUSG( | 0.015033 | 0.861571 | NO | NO | NA |
| Runx3 | ENSMUSG( | ENSMUSG( | -0.44096 | 3.81E-09 | YES | NO | NA |
| Rac1 | ENSMUSG( | ENSMUSG( | 0.127374 | 0.049242 | YES | NO | NA |

| | | | | | | | |
|---|---|---|---|---|---|---|---|
| Helz2 | ENSMUSG( | ENSMUSG( | -0.82749 | 6.87E-21 | YES | NO | NA |
| Gnb1 | ENSMUSG( | ENSMUSG( | -0.11183 | 0.034467 | YES | NO | NA |
| Gimap8 | ENSMUSG( | ENSMUSG( | -0.62337 | 1.85E-13 | YES | NO | NA |
| Pds5a | ENSMUSG( | ENSMUSG( | -0.0304 | 0.649088 | NO | NO | NA |
| Mcm3 | ENSMUSG( | ENSMUSG( | 0.240505 | 0.00222 | YES | NO | NA |
| Cd2 | ENSMUSG( | ENSMUSG( | -0.31253 | 0.000279 | YES | NO | NA |
| Sorl1 | ENSMUSG( | ENSMUSG( | -0.61306 | 7.46E-12 | YES | NO | NA |
| Vars | ENSMUSG( | ENSMUSG( | 0.521026 | 8.54E-10 | YES | NO | NA |
| Csk | ENSMUSG( | ENSMUSG( | -0.24734 | 5.37E-07 | YES | NO | NA |
| Numa1 | ENSMUSG( | ENSMUSG( | -0.36217 | 4.44E-09 | YES | NO | NA |
| Tkt | ENSMUSG( | ENSMUSG( | 0.401355 | 4.89E-06 | YES | NO | NA |
| Eif3f | ENSMUSG( | ENSMUSG( | 0.103113 | 0.251772 | NO | NO | NA |
| Atxn2l | ENSMUSG( | ENSMUSG( | -0.20648 | 1.91E-06 | YES | NO | NA |
| Lat | ENSMUSG( | ENSMUSG( | -0.22052 | 0.001759 | YES | NO | NA |
| Arf6 | ENSMUSG( | ENSMUSG( | -0.15337 | 0.006784 | YES | NO | NA |
| Fyb | ENSMUSG( | ENSMUSG( | -0.43769 | 5.38E-14 | YES | NO | NA |
| Srrt | ENSMUSG( | ENSMUSG( | 0.090134 | 0.378862 | NO | NO | NA |
| Apobec3 | ENSMUSG( | ENSMUSG( | 0.044568 | 0.639481 | NO | NO | NA |
| Ptk2b | ENSMUSG( | ENSMUSG( | -0.3695 | 1.18E-06 | YES | NO | NA |
| Prpf38b | ENSMUSG( | ENSMUSG( | -0.20145 | 0.235139 | NO | NO | NA |
| Ep400 | ENSMUSG( | ENSMUSG( | -0.14745 | 0.021603 | YES | NO | NA |
| Atrx | ENSMUSG( | ENSMUSG( | -0.19086 | 0.010969 | YES | NO | NA |
| Acap1 | ENSMUSG( | ENSMUSG( | -0.14405 | 0.008858 | YES | NO | NA |
| Nfatc3 | ENSMUSG( | ENSMUSG( | -0.48243 | 1.27E-10 | YES | NO | NA |
| Il2rg | ENSMUSG( | ENSMUSG( | -0.49673 | 1.13E-19 | YES | NO | NA |
| Nckap1l | ENSMUSG( | ENSMUSG( | -0.24393 | 2.46E-06 | YES | NO | NA |
| Wdr26 | ENSMUSG( | ENSMUSG( | -0.15464 | 0.126385 | NO | NO | NA |
| Add1 | ENSMUSG( | ENSMUSG( | 0.020451 | 0.818021 | NO | NO | NA |
| Hdgf | ENSMUSG( | ENSMUSG( | 0.334295 | 2.09E-05 | YES | NO | NA |
| Lrp10 | ENSMUSG( | ENSMUSG( | -0.08642 | 0.309026 | NO | NO | NA |
| Rpl35a | ENSMUSG( | ENSMUSG( | 0.249194 | 0.00673 | YES | NO | NA |
| G3bp1 | ENSMUSG( | ENSMUSG( | 0.196187 | 0.005916 | YES | NO | NA |
| Cblb | ENSMUSG( | ENSMUSG( | -0.15394 | 0.124783 | NO | NO | NA |
| Racgap1 | ENSMUSG( | ENSMUSG( | 0.052553 | 0.562518 | NO | NO | NA |
| Aebp2 | ENSMUSG( | ENSMUSG( | 0.075547 | 0.478161 | NO | NO | NA |
| Itk | ENSMUSG( | ENSMUSG( | -0.40491 | 2.27E-08 | YES | NO | NA |
| Cdk6 | ENSMUSG( | ENSMUSG( | -0.35569 | 1.44E-05 | YES | NO | NA |
| Hjurp | ENSMUSG( | ENSMUSG( | 0.144006 | 0.057757 | NO | NO | NA |
| Larp1 | ENSMUSG( | ENSMUSG( | 0.16138 | 0.005205 | YES | NO | NA |
| Lgals3 | ENSMUSG( | ENSMUSG( | 0.024013 | 0.868273 | NO | NO | NA |
| Srsf6 | ENSMUSG( | ENSMUSG( | 0.167337 | 0.034788 | YES | NO | NA |
| Arf1 | ENSMUSG( | ENSMUSG( | 0.009078 | 0.868932 | NO | NO | NA |
| Tle3 | ENSMUSG( | ENSMUSG( | -0.60739 | 1.33E-16 | YES | NO | NA |
| Tacc1 | ENSMUSG( | ENSMUSG( | -0.15319 | 0.047037 | YES | NO | NA |
| Csde1 | ENSMUSG( | ENSMUSG( | -0.07908 | 0.189591 | NO | NO | NA |
| AW11201( | ENSMUSG( | ENSMUSG( | -0.31076 | 0.004587 | YES | NO | NA |
| Akt1 | ENSMUSG( | ENSMUSG( | 0.218826 | 0.000232 | YES | NO | NA |

| | | | | | | | |
|---|---|---|---|---|---|---|---|
| Ubr5 | ENSMUSG( | ENSMUSG( | -0.03348 | 0.746228 | NO | NO | NA |
| Snrnp200 | ENSMUSG( | ENSMUSG( | -0.01157 | 0.874639 | NO | NO | NA |
| Rps21 | ENSMUSG( | ENSMUSG( | 0.30041 | 2.28E-09 | YES | NO | NA |
| Ddx21 | ENSMUSG( | ENSMUSG( | 0.301956 | 0.008379 | YES | NO | NA |
| Stat3 | ENSMUSG( | ENSMUSG( | -0.06429 | 0.532939 | NO | NO | NA |
| Trbc1 | ENSMUSG( | ENSMUSG( | -0.38546 | 0.000636 | YES | NO | NA |
| Csnk1a1 | ENSMUSG( | ENSMUSG( | -0.00262 | 0.972742 | NO | NO | NA |
| Morf4l1 | ENSMUSG( | ENSMUSG( | -0.0141 | 0.846798 | NO | NO | NA |
| Hdac7 | ENSMUSG( | ENSMUSG( | -0.45386 | 6.28E-05 | YES | NO | NA |
| Rbl2 | ENSMUSG( | ENSMUSG( | -0.44649 | 2.82E-06 | YES | NO | NA |
| Mob1a | ENSMUSG( | ENSMUSG( | -0.323 | 3.47E-07 | YES | NO | NA |
| Rsrp1 | ENSMUSG( | ENSMUSG( | -0.3488 | 0.00036 | YES | NO | NA |
| Mbd2 | ENSMUSG( | ENSMUSG( | 0.052397 | 0.55484 | NO | NO | NA |
| Incenp | ENSMUSG( | ENSMUSG( | 0.072924 | 0.636419 | NO | NO | NA |
| Mcl1 | ENSMUSG( | ENSMUSG( | -0.30903 | 1.87E-06 | YES | NO | NA |
| Acly | ENSMUSG( | ENSMUSG( | -0.08332 | 0.030067 | YES | NO | NA |
| Sbf1 | ENSMUSG( | ENSMUSG( | -0.29954 | 1.39E-06 | YES | NO | NA |
| Dusp1 | ENSMUSG( | ENSMUSG( | 0.329597 | 0.349528 | NO | NO | NA |
| Setx | ENSMUSG( | ENSMUSG( | -0.17078 | 0.015816 | YES | NO | NA |
| Eif4a2 | ENSMUSG( | ENSMUSG( | -0.33555 | 1.15E-07 | YES | NO | NA |
| Rrm2 | ENSMUSG( | ENSMUSG( | 0.40018 | 1.58E-06 | YES | NO | NA |
| Cbx3 | ENSMUSG( | ENSMUSG( | 0.087074 | 0.216296 | NO | NO | NA |
| Prf1 | ENSMUSG( | ENSMUSG( | -0.18714 | 0.065467 | NO | NO | NA |
| Brd4 | ENSMUSG( | ENSMUSG( | -0.24254 | 0.000111 | YES | NO | NA |
| Abca2 | ENSMUSG( | ENSMUSG( | -0.20688 | 0.002158 | YES | NO | NA |
| Nab1 | ENSMUSG( | ENSMUSG( | -0.0825 | 0.484833 | NO | NO | NA |
| Cyth4 | ENSMUSG( | ENSMUSG( | -0.33391 | 2.96E-06 | YES | NO | NA |
| Dnaja1 | ENSMUSG( | ENSMUSG( | -0.74448 | 2.83E-11 | YES | NO | NA |
| Ppp6r1 | ENSMUSG( | ENSMUSG( | -0.13882 | 0.001489 | YES | NO | NA |
| Pclaf | ENSMUSG( | ENSMUSG( | 0.395254 | 0.000139 | YES | NO | NA |
| Hnrnpc | ENSMUSG( | ENSMUSG( | -0.01588 | 0.812176 | NO | NO | NA |
| Sun2 | ENSMUSG( | ENSMUSG( | -0.29136 | 8.28E-07 | YES | NO | NA |
| Cab39 | ENSMUSG( | ENSMUSG( | 0.019161 | 0.803783 | NO | NO | NA |
| Cep250 | ENSMUSG( | ENSMUSG( | -0.09397 | 0.283805 | NO | NO | NA |
| Cnbp | ENSMUSG( | ENSMUSG( | 0.23212 | 9.87E-06 | YES | NO | NA |
| Myo9b | ENSMUSG( | ENSMUSG( | -0.28523 | 1.94E-06 | YES | NO | NA |
| Xdh | ENSMUSG( | ENSMUSG( | -0.24488 | 0.001175 | YES | NO | NA |
| Eif4g3 | ENSMUSG( | ENSMUSG( | -0.18801 | 0.005091 | YES | NO | NA |
| Rpa1 | ENSMUSG( | ENSMUSG( | -0.01134 | 0.818283 | NO | NO | NA |
| Fth1 | ENSMUSG( | ENSMUSG( | 0.03222 | 0.729736 | NO | NO | NA |
| Cd164 | ENSMUSG( | ENSMUSG( | -0.44526 | 2.57E-09 | YES | NO | NA |
| Ubtf | ENSMUSG( | ENSMUSG( | -0.03857 | 0.633181 | NO | NO | NA |
| Baz1a | ENSMUSG( | ENSMUSG( | -0.16139 | 0.110389 | NO | NO | NA |
| Arhgef18 | ENSMUSG( | ENSMUSG( | -0.60533 | 1.30E-14 | YES | NO | NA |
| Cx3cr1 | ENSMUSG( | ENSMUSG( | -1.6153 | 9.28E-22 | YES | YES | Downregulated |
| Hif1a | ENSMUSG( | ENSMUSG( | 0.334247 | 0.007158 | YES | NO | NA |
| Srrm1 | ENSMUSG( | ENSMUSG( | 0.044248 | 0.648619 | NO | NO | NA |

| | | | | | | | |
|---|---|---|---|---|---|---|---|
| Sema4a | ENSMUSG( | ENSMUSG( | -0.16446 | 0.047273 | YES | NO | NA |
| Hectd1 | ENSMUSG( | ENSMUSG( | -0.10387 | 0.022599 | YES | NO | NA |
| Srcap | ENSMUSG( | ENSMUSG( | -0.41663 | 4.07E-12 | YES | NO | NA |
| Serpinb9 | ENSMUSG( | ENSMUSG( | 0.485255 | 2.22E-05 | YES | NO | NA |
| Sla | ENSMUSG( | ENSMUSG( | -0.17702 | 0.105617 | NO | NO | NA |
| Ctsb | ENSMUSG( | ENSMUSG( | -0.06635 | 0.443092 | NO | NO | NA |
| Dhx15 | ENSMUSG( | ENSMUSG( | 0.076026 | 0.198986 | NO | NO | NA |
| Cbfb | ENSMUSG( | ENSMUSG( | -0.01138 | 0.886978 | NO | NO | NA |
| Dgka | ENSMUSG( | ENSMUSG( | -0.44864 | 1.65E-07 | YES | NO | NA |
| Mcm2 | ENSMUSG( | ENSMUSG( | 0.092573 | 0.316602 | NO | NO | NA |
| Pum2 | ENSMUSG( | ENSMUSG( | -0.12279 | 0.050239 | NO | NO | NA |
| Atp2a2 | ENSMUSG( | ENSMUSG( | 0.220142 | 0.000811 | YES | NO | NA |
| Rbm25 | ENSMUSG( | ENSMUSG( | -0.10201 | 0.49711 | NO | NO | NA |
| Cd47 | ENSMUSG( | ENSMUSG( | -0.06152 | 0.456391 | NO | NO | NA |
| Epsti1 | ENSMUSG( | ENSMUSG( | -0.31399 | 0.000159 | YES | NO | NA |
| Fasn | ENSMUSG( | ENSMUSG( | 0.436977 | 0.000187 | YES | NO | NA |
| Lcp2 | ENSMUSG( | ENSMUSG( | -0.19176 | 0.00201 | YES | NO | NA |
| H3f3a | ENSMUSG( | ENSMUSG( | 0.096256 | 0.364935 | NO | NO | NA |
| Smarca4 | ENSMUSG( | ENSMUSG( | 0.002549 | 0.972271 | NO | NO | NA |
| Rpl39 | ENSMUSG( | ENSMUSG( | 0.167947 | 0.003012 | YES | NO | NA |
| Nktr | ENSMUSG( | ENSMUSG( | -0.29178 | 4.19E-06 | YES | NO | NA |
| Lig1 | ENSMUSG( | ENSMUSG( | 0.204745 | 0.013363 | YES | NO | NA |
| Serf2 | ENSMUSG( | ENSMUSG( | -0.3843 | 0.000675 | YES | NO | NA |
| Gnb2 | ENSMUSG( | ENSMUSG( | -0.05306 | 0.224015 | NO | NO | NA |
| Mcm4 | ENSMUSG( | ENSMUSG( | 0.046065 | 0.558244 | NO | NO | NA |
| Dnm2 | ENSMUSG( | ENSMUSG( | -0.16232 | 0.011233 | YES | NO | NA |
| Rps6ka1 | ENSMUSG( | ENSMUSG( | -0.37828 | 5.03E-11 | YES | NO | NA |
| Aak1 | ENSMUSG( | ENSMUSG( | -0.45228 | 7.27E-07 | YES | NO | NA |
| Ctbp1 | ENSMUSG( | ENSMUSG( | -0.01844 | 0.732555 | NO | NO | NA |
| Fermt3 | ENSMUSG( | ENSMUSG( | 0.043963 | 0.459449 | NO | NO | NA |
| Rcc2 | ENSMUSG( | ENSMUSG( | 0.445189 | 2.88E-08 | YES | NO | NA |
| Ap2m1 | ENSMUSG( | ENSMUSG( | 0.002858 | 0.961255 | NO | NO | NA |
| Tcp1 | ENSMUSG( | ENSMUSG( | 0.378676 | 5.72E-11 | YES | NO | NA |
| Spata13 | ENSMUSG( | ENSMUSG( | -0.55839 | 4.01E-13 | YES | NO | NA |
| Cdc25b | ENSMUSG( | ENSMUSG( | -0.49183 | 2.67E-08 | YES | NO | NA |
| Tra2b | ENSMUSG( | ENSMUSG( | 0.032582 | 0.756346 | NO | NO | NA |
| Topbp1 | ENSMUSG( | ENSMUSG( | 0.163319 | 0.003693 | YES | NO | NA |
| Crip1 | ENSMUSG( | ENSMUSG( | -0.13442 | 0.192792 | NO | NO | NA |
| Rpl36 | ENSMUSG( | ENSMUSG( | 0.363766 | 1.08E-05 | YES | NO | NA |
| Itpkb | ENSMUSG( | ENSMUSG( | -0.62729 | 7.57E-12 | YES | NO | NA |
| Nsd2 | ENSMUSG( | ENSMUSG( | -0.02838 | 0.645299 | NO | NO | NA |
| Mycbp2 | ENSMUSG( | ENSMUSG( | -0.52577 | 6.00E-13 | YES | NO | NA |
| Oga | ENSMUSG( | ENSMUSG( | -0.33256 | 2.64E-06 | YES | NO | NA |
| Gm49909 | ENSMUSG( | ENSMUSG( | -0.11996 | 0.364742 | NO | NO | NA |
| Abhd2 | ENSMUSG( | ENSMUSG( | -0.24424 | 0.00253 | YES | NO | NA |
| Ccdc88b | ENSMUSG( | ENSMUSG( | -0.20126 | 0.000168 | YES | NO | NA |
| Bptf | ENSMUSG( | ENSMUSG( | -0.33373 | 7.66E-06 | YES | NO | NA |

| | | | | | | | |
|---|---|---|---|---|---|---|---|
| Dusp2 | ENSMUSG( | ENSMUSG( | -0.0555 | 0.732349 | NO | NO | NA |
| Sash3 | ENSMUSG( | ENSMUSG( | -0.50609 | 4.71E-10 | YES | NO | NA |
| Hsph1 | ENSMUSG( | ENSMUSG( | -1.21793 | 5.63E-16 | YES | YES | Downregulated |
| Slc1a5 | ENSMUSG( | ENSMUSG( | -0.11478 | 0.05121 | NO | NO | NA |
| Flii | ENSMUSG( | ENSMUSG( | 0.02843 | 0.62793 | NO | NO | NA |
| Smchd1 | ENSMUSG( | ENSMUSG( | -0.31735 | 3.98E-08 | YES | NO | NA |
| Notch1 | ENSMUSG( | ENSMUSG( | -0.38807 | 3.70E-06 | YES | NO | NA |
| Eomes | ENSMUSG( | ENSMUSG( | -0.68153 | 8.03E-07 | YES | NO | NA |
| Cytip | ENSMUSG( | ENSMUSG( | 0.024756 | 0.81886 | NO | NO | NA |
| Uhrf1 | ENSMUSG( | ENSMUSG( | 0.337078 | 0.00123 | YES | NO | NA |
| Pnn | ENSMUSG( | ENSMUSG( | -0.10323 | 0.505843 | NO | NO | NA |
| Gimap7 | ENSMUSG( | ENSMUSG( | -0.70206 | 4.89E-15 | YES | NO | NA |
| Rgs16 | ENSMUSG( | ENSMUSG( | 0.036902 | 0.927457 | NO | NO | NA |
| Glud1 | ENSMUSG( | ENSMUSG( | -0.09775 | 0.216296 | NO | NO | NA |
| Cdc37 | ENSMUSG( | ENSMUSG( | 0.119355 | 0.083438 | NO | NO | NA |
| Smarcc2 | ENSMUSG( | ENSMUSG( | -0.15141 | 0.01292 | YES | NO | NA |
| Tm9sf3 | ENSMUSG( | ENSMUSG( | 0.004453 | 0.951939 | NO | NO | NA |
| Arpc4 | ENSMUSG( | ENSMUSG( | -0.1026 | 0.022875 | YES | NO | NA |
| Cd48 | ENSMUSG( | ENSMUSG( | -0.43101 | 9.38E-13 | YES | NO | NA |
| Mtpn | ENSMUSG( | ENSMUSG( | -0.09242 | 0.334781 | NO | NO | NA |
| Psmd2 | ENSMUSG( | ENSMUSG( | 0.162278 | 0.001343 | YES | NO | NA |
| Chd3 | ENSMUSG( | ENSMUSG( | -0.69989 | 6.56E-13 | YES | NO | NA |
| Smarca5 | ENSMUSG( | ENSMUSG( | 0.239482 | 0.002072 | YES | NO | NA |
| E2f2 | ENSMUSG( | ENSMUSG( | 0.298351 | 0.001279 | YES | NO | NA |
| Agap2 | ENSMUSG( | ENSMUSG( | 0.026535 | 0.857484 | NO | NO | NA |
| Zfp36 | ENSMUSG( | ENSMUSG( | 0.536114 | 0.001855 | YES | NO | NA |
| Pak2 | ENSMUSG( | ENSMUSG( | -0.22632 | 3.68E-05 | YES | NO | NA |
| Cd226 | ENSMUSG( | ENSMUSG( | -0.67038 | 5.71E-10 | YES | NO | NA |
| Tardbp | ENSMUSG( | ENSMUSG( | 0.101688 | 0.181363 | NO | NO | NA |
| P4hb | ENSMUSG( | ENSMUSG( | 0.117981 | 0.157925 | NO | NO | NA |
| Macroh2a | ENSMUSG( | ENSMUSG( | 0.053851 | 0.248136 | NO | NO | NA |
| Id2 | ENSMUSG( | ENSMUSG( | -0.18545 | 0.06781 | NO | NO | NA |
| Map4k2 | ENSMUSG( | ENSMUSG( | -0.2912 | 1.43E-05 | YES | NO | NA |
| Lgals3bp | ENSMUSG( | ENSMUSG( | -0.56054 | 1.35E-16 | YES | NO | NA |
| Add3 | ENSMUSG( | ENSMUSG( | -0.07254 | 0.449445 | NO | NO | NA |
| Rangap1 | ENSMUSG( | ENSMUSG( | -0.05706 | 0.506091 | NO | NO | NA |
| Wasf2 | ENSMUSG( | ENSMUSG( | -0.14935 | 0.028191 | YES | NO | NA |
| Smc2 | ENSMUSG( | ENSMUSG( | 0.111766 | 0.169199 | NO | NO | NA |
| Sumo2 | ENSMUSG( | ENSMUSG( | -0.07154 | 0.174022 | NO | NO | NA |
| Ccnd2 | ENSMUSG( | ENSMUSG( | -0.57947 | 5.50E-08 | YES | NO | NA |
| Rpl36a | ENSMUSG( | ENSMUSG( | 0.482132 | 6.73E-06 | YES | NO | NA |
| Ccna2 | ENSMUSG( | ENSMUSG( | 0.303841 | 0.000289 | YES | NO | NA |
| Ehmt2 | ENSMUSG( | ENSMUSG( | -0.01006 | 0.891155 | NO | NO | NA |
| Leng8 | ENSMUSG( | ENSMUSG( | -0.22076 | 0.003458 | YES | NO | NA |
| Cd6 | ENSMUSG( | ENSMUSG( | -0.15463 | 0.274415 | NO | NO | NA |
| Pik3ap1 | ENSMUSG( | ENSMUSG( | -0.51889 | 3.10E-09 | YES | NO | NA |
| Os9 | ENSMUSG( | ENSMUSG( | -0.21237 | 0.003944 | YES | NO | NA |

| | | | | | | | |
|---|---|---|---|---|---|---|---|
| Suz12 | ENSMUSG( | ENSMUSG( | 0.111501 | 0.211315 | NO | NO | NA |
| Pstpip1 | ENSMUSG( | ENSMUSG( | -0.17836 | 0.021854 | YES | NO | NA |
| Supt5 | ENSMUSG( | ENSMUSG( | -0.06052 | 0.204563 | NO | NO | NA |
| 2610507B: | ENSMUSG( | ENSMUSG( | 0.011384 | 0.876094 | NO | NO | NA |
| Cast | ENSMUSG( | ENSMUSG( | -0.18283 | 0.041187 | YES | NO | NA |
| M6pr | ENSMUSG( | ENSMUSG( | 0.137806 | 0.013718 | YES | NO | NA |
| Khsrp | ENSMUSG( | ENSMUSG( | 0.078599 | 0.135768 | NO | NO | NA |
| Rpl37 | ENSMUSG( | ENSMUSG( | 0.260379 | 0.000484 | YES | NO | NA |
| Gtf2i | ENSMUSG( | ENSMUSG( | -0.39393 | 2.35E-07 | YES | NO | NA |
| Arhgap4 | ENSMUSG( | ENSMUSG( | -0.51082 | 7.81E-16 | YES | NO | NA |
| Purb | ENSMUSG( | ENSMUSG( | -0.18 | 0.003219 | YES | NO | NA |
| Ppp2ca | ENSMUSG( | ENSMUSG( | 0.031643 | 0.640936 | NO | NO | NA |
| Bcl11b | ENSMUSG( | ENSMUSG( | -0.57495 | 3.12E-11 | YES | NO | NA |
| Trrap | ENSMUSG( | ENSMUSG( | -0.35989 | 5.76E-11 | YES | NO | NA |
| Madd | ENSMUSG( | ENSMUSG( | -0.33121 | 0.000146 | YES | NO | NA |
| Nsd3 | ENSMUSG( | ENSMUSG( | -0.48285 | 3.43E-11 | YES | NO | NA |
| Ctcf | ENSMUSG( | ENSMUSG( | -0.10843 | 0.231179 | NO | NO | NA |
| Prkcq | ENSMUSG( | ENSMUSG( | -0.41967 | 3.55E-08 | YES | NO | NA |
| Ddx46 | ENSMUSG( | ENSMUSG( | 0.049219 | 0.672948 | NO | NO | NA |
| Rab27a | ENSMUSG( | ENSMUSG( | 0.15198 | 0.127985 | NO | NO | NA |
| Smc3 | ENSMUSG( | ENSMUSG( | 0.099448 | 0.458022 | NO | NO | NA |
| Srgn | ENSMUSG( | ENSMUSG( | -0.00909 | 0.943802 | NO | NO | NA |
| Rbbp7 | ENSMUSG( | ENSMUSG( | 0.196361 | 0.007216 | YES | NO | NA |
| Ndfip1 | ENSMUSG( | ENSMUSG( | -0.04175 | 0.731835 | NO | NO | NA |
| Syncrip | ENSMUSG( | ENSMUSG( | 0.301333 | 0.000259 | YES | NO | NA |
| Cic | ENSMUSG( | ENSMUSG( | -0.42202 | 1.25E-11 | YES | NO | NA |
| Kat6a | ENSMUSG( | ENSMUSG( | -0.62573 | 6.12E-14 | YES | NO | NA |
| Ywhab | ENSMUSG( | ENSMUSG( | -0.21586 | 3.16E-07 | YES | NO | NA |
| BC005537 | ENSMUSG( | ENSMUSG( | -0.14545 | 0.012065 | YES | NO | NA |
| Pik3r5 | ENSMUSG( | ENSMUSG( | -0.56226 | 9.91E-11 | YES | NO | NA |
| Usp48 | ENSMUSG( | ENSMUSG( | -0.23231 | 0.009311 | YES | NO | NA |
| Themis | ENSMUSG( | ENSMUSG( | -0.18072 | 0.017436 | YES | NO | NA |
| Pafah1b1 | ENSMUSG( | ENSMUSG( | -0.30783 | 5.35E-11 | YES | NO | NA |
| Psme1 | ENSMUSG( | ENSMUSG( | -0.08814 | 0.104442 | NO | NO | NA |
| Map4 | ENSMUSG( | ENSMUSG( | -0.15824 | 0.002198 | YES | NO | NA |
| Elf1 | ENSMUSG( | ENSMUSG( | -0.38181 | 5.39E-08 | YES | NO | NA |
| Lyst | ENSMUSG( | ENSMUSG( | -0.33951 | 0.008288 | YES | NO | NA |
| Cct7 | ENSMUSG( | ENSMUSG( | 0.226511 | 6.15E-06 | YES | NO | NA |
| Fcho1 | ENSMUSG( | ENSMUSG( | -0.03674 | 0.540096 | NO | NO | NA |
| Fam168b | ENSMUSG( | ENSMUSG( | -0.24248 | 8.96E-06 | YES | NO | NA |
| Ppm1g | ENSMUSG( | ENSMUSG( | 0.199242 | 0.0353 | YES | NO | NA |
| Plekha2 | ENSMUSG( | ENSMUSG( | -0.20451 | 0.005554 | YES | NO | NA |
| Hipk1 | ENSMUSG( | ENSMUSG( | -0.37471 | 1.08E-06 | YES | NO | NA |
| Safb | ENSMUSG( | ENSMUSG( | -0.10171 | 0.432065 | NO | NO | NA |
| Ubap2l | ENSMUSG( | ENSMUSG( | -0.06821 | 0.342191 | NO | NO | NA |
| Ifng | ENSMUSG( | ENSMUSG( | 0.157201 | 0.438698 | NO | NO | NA |
| Mndal | ENSMUSG( | ENSMUSG( | -0.60285 | 3.62E-12 | YES | NO | NA |

| | | | | | | | |
|---|---|---|---|---|---|---|---|
| Psd4 | ENSMUSG( | ENSMUSG( | -0.3859 | 7.52E-16 | YES | NO | NA |
| Usp25 | ENSMUSG( | ENSMUSG( | -0.31741 | 1.97E-07 | YES | NO | NA |
| Cd3d | ENSMUSG( | ENSMUSG( | -0.23361 | 6.15E-06 | YES | NO | NA |
| Cd7 | ENSMUSG( | ENSMUSG( | -0.90397 | 2.39E-14 | YES | NO | NA |
| Klf13 | ENSMUSG( | ENSMUSG( | -0.29397 | 0.000513 | YES | NO | NA |
| Septin6 | ENSMUSG( | ENSMUSG( | -0.38091 | 6.23E-13 | YES | NO | NA |
| Tmed2 | ENSMUSG( | ENSMUSG( | 0.103438 | 0.042344 | YES | NO | NA |
| Itsn2 | ENSMUSG( | ENSMUSG( | -0.37588 | 1.95E-08 | YES | NO | NA |
| Cct5 | ENSMUSG( | ENSMUSG( | 0.209959 | 0.001447 | YES | NO | NA |
| Cox4i1 | ENSMUSG( | ENSMUSG( | 0.201326 | 0.005635 | YES | NO | NA |
| Calm2 | ENSMUSG( | ENSMUSG( | -0.14816 | 0.106491 | NO | NO | NA |
| Cct8 | ENSMUSG( | ENSMUSG( | 0.293107 | 2.47E-05 | YES | NO | NA |
| Ccni | ENSMUSG( | ENSMUSG( | -0.07769 | 0.43299 | NO | NO | NA |
| Mical1 | ENSMUSG( | ENSMUSG( | -0.30266 | 5.80E-08 | YES | NO | NA |
| Hspd1 | ENSMUSG( | ENSMUSG( | 0.281768 | 0.017051 | YES | NO | NA |
| Rpl22 | ENSMUSG( | ENSMUSG( | 0.327349 | 5.29E-07 | YES | NO | NA |
| Picalm | ENSMUSG( | ENSMUSG( | 0.027467 | 0.749922 | NO | NO | NA |
| Sf3b3 | ENSMUSG( | ENSMUSG( | 0.233791 | 7.64E-05 | YES | NO | NA |
| Kif11 | ENSMUSG( | ENSMUSG( | 0.02196 | 0.783213 | NO | NO | NA |
| Erdr1 | ENSMUSG( | ENSMUSG( | -0.16783 | 0.63996 | NO | NO | NA |
| Abr | ENSMUSG( | ENSMUSG( | -0.30241 | 4.43E-06 | YES | NO | NA |
| Gimap1 | ENSMUSG( | ENSMUSG( | -0.16167 | 0.004886 | YES | NO | NA |
| Eif2s3x | ENSMUSG( | ENSMUSG( | 0.135805 | 0.01018 | YES | NO | NA |
| Eif5 | ENSMUSG( | ENSMUSG( | 0.020499 | 0.758528 | NO | NO | NA |
| Pitpna | ENSMUSG( | ENSMUSG( | -0.15265 | 0.007035 | YES | NO | NA |
| Top2b | ENSMUSG( | ENSMUSG( | -0.19171 | 0.012081 | YES | NO | NA |
| Cltc | ENSMUSG( | ENSMUSG( | -0.06311 | 0.304777 | NO | NO | NA |
| Hspa4 | ENSMUSG( | ENSMUSG( | 0.219944 | 0.022383 | YES | NO | NA |
| Txn1 | ENSMUSG( | ENSMUSG( | 0.241178 | 0.000326 | YES | NO | NA |
| Saraf | ENSMUSG( | ENSMUSG( | -0.45175 | 2.35E-16 | YES | NO | NA |
| Cs | ENSMUSG( | ENSMUSG( | 0.192554 | 0.005914 | YES | NO | NA |
| Snx5 | ENSMUSG( | ENSMUSG( | 0.153329 | 0.145108 | NO | NO | NA |
| Cd247 | ENSMUSG( | ENSMUSG( | -0.29181 | 0.001095 | YES | NO | NA |
| Atp8b2 | ENSMUSG( | ENSMUSG( | -0.2913 | 3.81E-05 | YES | NO | NA |
| Ppp1r9b | ENSMUSG( | ENSMUSG( | -0.45773 | 1.70E-07 | YES | NO | NA |
| Ankrd13a | ENSMUSG( | ENSMUSG( | 0.018882 | 0.757131 | NO | NO | NA |
| Myl12a | ENSMUSG( | ENSMUSG( | -0.03924 | 0.467168 | NO | NO | NA |
| Nucb1 | ENSMUSG( | ENSMUSG( | -0.06158 | 0.449198 | NO | NO | NA |
| Anp32a | ENSMUSG( | ENSMUSG( | -0.47695 | 0.000769 | YES | NO | NA |
| Rbbp4 | ENSMUSG( | ENSMUSG( | 0.005259 | 0.914757 | NO | NO | NA |
| Herc1 | ENSMUSG( | ENSMUSG( | -0.45203 | 1.70E-11 | YES | NO | NA |
| Atf7ip | ENSMUSG( | ENSMUSG( | -0.83261 | 1.15E-23 | YES | NO | NA |
| Dnajc9 | ENSMUSG( | ENSMUSG( | -0.35374 | 6.55E-07 | YES | NO | NA |
| Srsf11 | ENSMUSG( | ENSMUSG( | -0.0019 | 0.984133 | NO | NO | NA |
| Mlec | ENSMUSG( | ENSMUSG( | -0.16495 | 0.013679 | YES | NO | NA |
| Xrn2 | ENSMUSG( | ENSMUSG( | 0.058738 | 0.475513 | NO | NO | NA |
| Chd7 | ENSMUSG( | ENSMUSG( | 0.022011 | 0.835507 | NO | NO | NA |

| | | | | | | | |
|---|---|---|---|---|---|---|---|
| Hnrnpdl | ENSMUSG( | ENSMUSG( | 0.201389 | 0.044218 | YES | NO | NA |
| Plcg1 | ENSMUSG( | ENSMUSG( | -0.27524 | 0.0014 | YES | NO | NA |
| Def6 | ENSMUSG( | ENSMUSG( | 0.043679 | 0.511138 | NO | NO | NA |
| Kpna2 | ENSMUSG( | ENSMUSG( | 0.164846 | 0.03173 | YES | NO | NA |
| Ywhaq | ENSMUSG( | ENSMUSG( | -0.11708 | 0.005024 | YES | NO | NA |
| Ogdh | ENSMUSG( | ENSMUSG( | -0.19944 | 5.08E-07 | YES | NO | NA |
| Ccnd3 | ENSMUSG( | ENSMUSG( | -0.42179 | 1.89E-06 | YES | NO | NA |
| Mdm4 | ENSMUSG( | ENSMUSG( | -0.30876 | 6.65E-05 | YES | NO | NA |
| Ide | ENSMUSG( | ENSMUSG( | -0.71327 | 2.25E-07 | YES | NO | NA |
| Gramd1a | ENSMUSG( | ENSMUSG( | -0.17339 | 0.001642 | YES | NO | NA |
| Fam102a | ENSMUSG( | ENSMUSG( | -0.31278 | 0.002216 | YES | NO | NA |
| Ago2 | ENSMUSG( | ENSMUSG( | -0.30258 | 8.65E-06 | YES | NO | NA |
| Rgs1 | ENSMUSG( | ENSMUSG( | 0.363894 | 0.158927 | NO | NO | NA |
| Lpin1 | ENSMUSG( | ENSMUSG( | -1.25274 | 1.59E-07 | YES | YES | Downregulated |
| Slc38a1 | ENSMUSG( | ENSMUSG( | 0.093375 | 0.068476 | NO | NO | NA |
| Rassf5 | ENSMUSG( | ENSMUSG( | -0.27186 | 0.000182 | YES | NO | NA |
| Utrn | ENSMUSG( | ENSMUSG( | -0.69274 | 1.34E-14 | YES | NO | NA |
| Ifi209 | ENSMUSG( | ENSMUSG( | -1.13076 | 1.40E-14 | YES | YES | Downregulated |
| S100a11 | ENSMUSG( | ENSMUSG( | -0.27361 | 0.000506 | YES | NO | NA |
| Tbc1d10c | ENSMUSG( | ENSMUSG( | -0.32235 | 6.14E-07 | YES | NO | NA |
| Ppp2r5c | ENSMUSG( | ENSMUSG( | -0.26621 | 0.000132 | YES | NO | NA |
| Ssb | ENSMUSG( | ENSMUSG( | 0.177824 | 0.03717 | YES | NO | NA |
| Agpat3 | ENSMUSG( | ENSMUSG( | -0.46842 | 2.59E-07 | YES | NO | NA |
| Pa2g4 | ENSMUSG( | ENSMUSG( | 0.581189 | 8.89E-08 | YES | NO | NA |
| Eif4ebp2 | ENSMUSG( | ENSMUSG( | -0.15917 | 0.009649 | YES | NO | NA |
| Rap1gap2 | ENSMUSG( | ENSMUSG( | -0.87263 | 6.23E-19 | YES | NO | NA |
| Hdlbp | ENSMUSG( | ENSMUSG( | 0.105467 | 0.120089 | NO | NO | NA |
| Trim28 | ENSMUSG( | ENSMUSG( | 0.25144 | 2.87E-07 | YES | NO | NA |
| Hdac1 | ENSMUSG( | ENSMUSG( | 0.013096 | 0.820172 | NO | NO | NA |
| Ankrd52 | ENSMUSG( | ENSMUSG( | 0.123825 | 0.127774 | NO | NO | NA |
| Mark2 | ENSMUSG( | ENSMUSG( | -0.34734 | 3.93E-08 | YES | NO | NA |
| Rrbp1 | ENSMUSG( | ENSMUSG( | -0.12645 | 0.491395 | NO | NO | NA |
| Tuba1c | ENSMUSG( | ENSMUSG( | 0.076537 | 0.197655 | NO | NO | NA |
| Zmiz1 | ENSMUSG( | ENSMUSG( | -0.25539 | 9.21E-05 | YES | NO | NA |
| Anxa11 | ENSMUSG( | ENSMUSG( | -0.13392 | 0.028549 | YES | NO | NA |
| Apbb1ip | ENSMUSG( | ENSMUSG( | -0.5517 | 1.48E-13 | YES | NO | NA |
| Ttc39b | ENSMUSG( | ENSMUSG( | -0.38314 | 9.04E-05 | YES | NO | NA |
| Baz2a | ENSMUSG( | ENSMUSG( | -0.40943 | 7.69E-07 | YES | NO | NA |
| Cep85 | ENSMUSG( | ENSMUSG( | 1.044233 | 3.73E-12 | YES | YES | Upregulated |
| Ivns1abp | ENSMUSG( | ENSMUSG( | 0.078746 | 0.175573 | NO | NO | NA |
| Safb2 | ENSMUSG( | ENSMUSG( | -0.18339 | 0.184777 | NO | NO | NA |
| Hk1 | ENSMUSG( | ENSMUSG( | -0.21131 | 1.63E-10 | YES | NO | NA |
| Papola | ENSMUSG( | ENSMUSG( | -0.18881 | 0.004274 | YES | NO | NA |
| 4932438A | ENSMUSG( | ENSMUSG( | -0.35885 | 3.26E-06 | YES | NO | NA |
| Clint1 | ENSMUSG( | ENSMUSG( | -0.18647 | 0.049129 | YES | NO | NA |
| Herc2 | ENSMUSG( | ENSMUSG( | -0.53268 | 1.10E-15 | YES | NO | NA |
| Sytl2 | ENSMUSG( | ENSMUSG( | -0.49699 | 4.71E-05 | YES | NO | NA |

| | | | | | | | |
|---|---|---|---|---|---|---|---|
| Khdrbs1 | ENSMUSG( | ENSMUSG( | 0.05486 | 0.322348 | NO | NO | NA |
| Ipcef1 | ENSMUSG( | ENSMUSG( | -0.45775 | 9.45E-07 | YES | NO | NA |
| Xpo1 | ENSMUSG( | ENSMUSG( | -0.09528 | 0.290698 | NO | NO | NA |
| Atp5g3 | ENSMUSG( | ENSMUSG( | 0.377141 | 6.61E-08 | YES | NO | NA |
| Rgs3 | ENSMUSG( | ENSMUSG( | -0.54695 | 3.53E-12 | YES | NO | NA |
| Unc13d | ENSMUSG( | ENSMUSG( | -0.29684 | 1.43E-10 | YES | NO | NA |
| Ppp2r1a | ENSMUSG( | ENSMUSG( | 0.130292 | 0.019374 | YES | NO | NA |
| Il10ra | ENSMUSG( | ENSMUSG( | -0.28372 | 0.04437 | YES | NO | NA |
| Akr1a1 | ENSMUSG( | ENSMUSG( | 0.222577 | 0.000306 | YES | NO | NA |
| Ptpn11 | ENSMUSG( | ENSMUSG( | -0.09047 | 0.152533 | NO | NO | NA |
| Eif3e | ENSMUSG( | ENSMUSG( | 0.238142 | 0.000711 | YES | NO | NA |
| Abcf1 | ENSMUSG( | ENSMUSG( | 0.07855 | 0.413484 | NO | NO | NA |
| Got2 | ENSMUSG( | ENSMUSG( | 0.417402 | 3.74E-05 | YES | NO | NA |
| Rpn2 | ENSMUSG( | ENSMUSG( | 0.17831 | 0.000556 | YES | NO | NA |
| Kif5b | ENSMUSG( | ENSMUSG( | -0.11064 | 0.05163 | NO | NO | NA |
| Sart1 | ENSMUSG( | ENSMUSG( | 0.034537 | 0.773408 | NO | NO | NA |
| Ipo5 | ENSMUSG( | ENSMUSG( | 0.4863 | 3.16E-08 | YES | NO | NA |
| Mapre1 | ENSMUSG( | ENSMUSG( | 0.079002 | 0.085539 | NO | NO | NA |
| Fkbp8 | ENSMUSG( | ENSMUSG( | 0.011238 | 0.862668 | NO | NO | NA |
| Srsf7 | ENSMUSG( | ENSMUSG( | 0.043039 | 0.574026 | NO | NO | NA |
| Tnks2 | ENSMUSG( | ENSMUSG( | -0.28029 | 1.72E-06 | YES | NO | NA |
| Prkacb | ENSMUSG( | ENSMUSG( | -0.3286 | 1.08E-05 | YES | NO | NA |
| Ep300 | ENSMUSG( | ENSMUSG( | -0.60362 | 5.99E-21 | YES | NO | NA |
| Cds2 | ENSMUSG( | ENSMUSG( | -0.35643 | 3.48E-08 | YES | NO | NA |
| Ostf1 | ENSMUSG( | ENSMUSG( | 0.155403 | 0.044011 | YES | NO | NA |
| Mapk1 | ENSMUSG( | ENSMUSG( | -0.20672 | 9.59E-06 | YES | NO | NA |
| Ftl1 | ENSMUSG( | ENSMUSG( | -0.05989 | 0.33387 | NO | NO | NA |
| Gls | ENSMUSG( | ENSMUSG( | -0.00841 | 0.897767 | NO | NO | NA |
| ENSMUSG | ENSMUSG( | ENSMUSG( | 0.078606 | 0.523831 | NO | NO | NA |
| Serpinb6b | ENSMUSG( | ENSMUSG( | 0.346096 | 0.000701 | YES | NO | NA |
| Psip1 | ENSMUSG( | ENSMUSG( | -0.21219 | 0.044554 | YES | NO | NA |
| Acsl5 | ENSMUSG( | ENSMUSG( | 0.023716 | 0.715825 | NO | NO | NA |
| Esyt2 | ENSMUSG( | ENSMUSG( | -0.66509 | 7.01E-15 | YES | NO | NA |
| Sdha | ENSMUSG( | ENSMUSG( | 0.127252 | 0.021736 | YES | NO | NA |
| Odc1 | ENSMUSG( | ENSMUSG( | 0.549608 | 4.16E-05 | YES | NO | NA |
| Ptpn7 | ENSMUSG( | ENSMUSG( | 0.143256 | 0.006003 | YES | NO | NA |
| Lrrc8c | ENSMUSG( | ENSMUSG( | -0.39964 | 1.54E-09 | YES | NO | NA |
| Myl12b | ENSMUSG( | ENSMUSG( | -0.20672 | 0.123848 | NO | NO | NA |
| Tsc22d4 | ENSMUSG( | ENSMUSG( | -0.37401 | 4.71E-11 | YES | NO | NA |
| Atxn1 | ENSMUSG( | ENSMUSG( | -0.55364 | 6.13E-09 | YES | NO | NA |
| Ranbp2 | ENSMUSG( | ENSMUSG( | -0.0834 | 0.168138 | NO | NO | NA |
| Ctsw | ENSMUSG( | ENSMUSG( | -0.31359 | 5.20E-09 | YES | NO | NA |
| Irf1 | ENSMUSG( | ENSMUSG( | -0.3377 | 1.78E-08 | YES | NO | NA |
| Atp8b4 | ENSMUSG( | ENSMUSG( | 0.313253 | 0.002964 | YES | NO | NA |
| Rab8a | ENSMUSG( | ENSMUSG( | -0.12069 | 0.055793 | NO | NO | NA |
| H2-T22 | ENSMUSG( | ENSMUSG( | -0.61772 | 9.79E-11 | YES | NO | NA |
| Irag2 | ENSMUSG( | ENSMUSG( | -0.14315 | 0.02644 | YES | NO | NA |

| | | | | | | | |
|---|---|---|---|---|---|---|---|
| Jpt1 | ENSMUSG( | ENSMUSG( | -0.14928 | 0.075876 | NO | NO | NA |
| Bag6 | ENSMUSG( | ENSMUSG( | -0.01838 | 0.735803 | NO | NO | NA |
| Cbx5 | ENSMUSG( | ENSMUSG( | 0.000132 | 0.99875 | NO | NO | NA |
| Galnt2 | ENSMUSG( | ENSMUSG( | -0.41066 | 1.39E-06 | YES | NO | NA |
| Gm42427 | ENSMUSG( | ENSMUSG( | 1.325143 | 7.72E-41 | YES | YES | Upregulated |
| Slc20a1 | ENSMUSG( | ENSMUSG( | -0.49928 | 1.62E-08 | YES | NO | NA |
| Rock1 | ENSMUSG( | ENSMUSG( | -0.16162 | 0.054676 | NO | NO | NA |
| Arf3 | ENSMUSG( | ENSMUSG( | -0.22059 | 0.000118 | YES | NO | NA |
| Mob3a | ENSMUSG( | ENSMUSG( | -0.24271 | 2.58E-07 | YES | NO | NA |
| Ly6a | ENSMUSG( | ENSMUSG( | -0.222 | 0.053309 | NO | NO | NA |
| Fos | ENSMUSG( | ENSMUSG( | 0.943604 | 0.015569 | YES | NO | NA |
| Gbp7 | ENSMUSG( | ENSMUSG( | -0.11592 | 0.222199 | NO | NO | NA |
| Asxl1 | ENSMUSG( | ENSMUSG( | -0.10782 | 0.181275 | NO | NO | NA |
| Taf15 | ENSMUSG( | ENSMUSG( | -0.38534 | 0.0056 | YES | NO | NA |
| Dock11 | ENSMUSG( | ENSMUSG( | -0.3582 | 4.50E-10 | YES | NO | NA |
| Sec61a1 | ENSMUSG( | ENSMUSG( | 0.058728 | 0.226058 | NO | NO | NA |
| Metap2 | ENSMUSG( | ENSMUSG( | 0.358011 | 0.000161 | YES | NO | NA |
| Dbnl | ENSMUSG( | ENSMUSG( | -0.15578 | 0.000764 | YES | NO | NA |
| Lbh | ENSMUSG( | ENSMUSG( | -0.61773 | 4.28E-08 | YES | NO | NA |
| Kmt2b | ENSMUSG( | ENSMUSG( | -0.32556 | 1.17E-09 | YES | NO | NA |
| mt-Nd6 | ENSMUSG( | ENSMUSG( | 0.185189 | 0.11807 | NO | NO | NA |
| Arcn1 | ENSMUSG( | ENSMUSG( | -0.00829 | 0.878391 | NO | NO | NA |
| Erbin | ENSMUSG( | ENSMUSG( | -0.36735 | 3.20E-05 | YES | NO | NA |
| Smap2 | ENSMUSG( | ENSMUSG( | -0.2059 | 0.003445 | YES | NO | NA |
| Ctsa | ENSMUSG( | ENSMUSG( | -0.3889 | 7.23E-06 | YES | NO | NA |
| Unc119b | ENSMUSG( | ENSMUSG( | -0.09026 | 0.435067 | NO | NO | NA |
| Tcf25 | ENSMUSG( | ENSMUSG( | -0.04485 | 0.527421 | NO | NO | NA |
| Tpx2 | ENSMUSG( | ENSMUSG( | 0.16842 | 0.115398 | NO | NO | NA |
| Parp14 | ENSMUSG( | ENSMUSG( | -0.62732 | 2.89E-13 | YES | NO | NA |
| B4galt5 | ENSMUSG( | ENSMUSG( | -0.32903 | 4.84E-11 | YES | NO | NA |
| Ttc7 | ENSMUSG( | ENSMUSG( | -0.30829 | 2.04E-07 | YES | NO | NA |
| Ccnl2 | ENSMUSG( | ENSMUSG( | -0.2448 | 6.01E-05 | YES | NO | NA |
| Capn2 | ENSMUSG( | ENSMUSG( | -0.52734 | 6.98E-20 | YES | NO | NA |
| Mast3 | ENSMUSG( | ENSMUSG( | -0.29989 | 9.80E-06 | YES | NO | NA |
| Arglu1 | ENSMUSG( | ENSMUSG( | -0.09165 | 0.483498 | NO | NO | NA |
| Ctnnb1 | ENSMUSG( | ENSMUSG( | -0.00185 | 0.966013 | NO | NO | NA |
| Kars | ENSMUSG( | ENSMUSG( | 0.320922 | 1.48E-05 | YES | NO | NA |
| Chmp4b | ENSMUSG( | ENSMUSG( | 0.011441 | 0.871655 | NO | NO | NA |
| Gatad2a | ENSMUSG( | ENSMUSG( | -0.04301 | 0.481538 | NO | NO | NA |
| Hspa9 | ENSMUSG( | ENSMUSG( | 0.689037 | 1.72E-07 | YES | NO | NA |
| Dazap1 | ENSMUSG( | ENSMUSG( | 0.196511 | 0.076362 | NO | NO | NA |
| Mdh2 | ENSMUSG( | ENSMUSG( | 0.271402 | 1.97E-06 | YES | NO | NA |
| Zfp207 | ENSMUSG( | ENSMUSG( | -0.01072 | 0.910368 | NO | NO | NA |
| Tgfbr2 | ENSMUSG( | ENSMUSG( | -0.36365 | 0.002944 | YES | NO | NA |
| Chtop | ENSMUSG( | ENSMUSG( | -0.23355 | 0.000953 | YES | NO | NA |
| Chd8 | ENSMUSG( | ENSMUSG( | -0.19851 | 3.57E-05 | YES | NO | NA |
| Med12 | ENSMUSG( | ENSMUSG( | -0.45422 | 6.99E-15 | YES | NO | NA |

| | | | | | | | |
|---|---|---|---|---|---|---|---|
| Gzmk | ENSMUSG( | ENSMUSG( | -0.28275 | 0.014115 | YES | NO | NA |
| Ppib | ENSMUSG( | ENSMUSG( | 0.104936 | 0.025988 | YES | NO | NA |
| Raf1 | ENSMUSG( | ENSMUSG( | -0.06292 | 0.653824 | NO | NO | NA |
| Ldb1 | ENSMUSG( | ENSMUSG( | -0.10686 | 0.238639 | NO | NO | NA |
| Snrpb | ENSMUSG( | ENSMUSG( | 0.250707 | 5.46E-05 | YES | NO | NA |
| Capn1 | ENSMUSG( | ENSMUSG( | -0.36676 | 3.59E-18 | YES | NO | NA |
| Cenpa | ENSMUSG( | ENSMUSG( | 0.183778 | 0.01094 | YES | NO | NA |
| Arhgap11a | ENSMUSG( | ENSMUSG( | -0.01082 | 0.945161 | NO | NO | NA |
| Supt16 | ENSMUSG( | ENSMUSG( | 0.210796 | 0.006963 | YES | NO | NA |
| Hmga1 | ENSMUSG( | ENSMUSG( | 0.236507 | 1.06E-05 | YES | NO | NA |
| Flnb | ENSMUSG( | ENSMUSG( | -0.20546 | 0.042273 | YES | NO | NA |
| Ptms | ENSMUSG( | ENSMUSG( | -0.33446 | 0.000749 | YES | NO | NA |
| Tcof1 | ENSMUSG( | ENSMUSG( | -0.0688 | 0.36873 | NO | NO | NA |
| Arl6ip5 | ENSMUSG( | ENSMUSG( | -0.24775 | 0.000319 | YES | NO | NA |
| Tab2 | ENSMUSG( | ENSMUSG( | -0.22356 | 0.003604 | YES | NO | NA |
| Ppp2r5a | ENSMUSG( | ENSMUSG( | -0.25638 | 1.72E-06 | YES | NO | NA |
| Ganab | ENSMUSG( | ENSMUSG( | -0.18856 | 1.99E-06 | YES | NO | NA |
| S1pr4 | ENSMUSG( | ENSMUSG( | -0.40232 | 3.64E-08 | YES | NO | NA |
| Capns1 | ENSMUSG( | ENSMUSG( | 0.015362 | 0.824518 | NO | NO | NA |
| Eftud2 | ENSMUSG( | ENSMUSG( | 0.273013 | 5.96E-06 | YES | NO | NA |
| Mef2d | ENSMUSG( | ENSMUSG( | -0.32855 | 0.000472 | YES | NO | NA |
| Usp7 | ENSMUSG( | ENSMUSG( | -0.00909 | 0.886329 | NO | NO | NA |
| Scaf11 | ENSMUSG( | ENSMUSG( | 0.085657 | 0.301508 | NO | NO | NA |
| Cyba | ENSMUSG( | ENSMUSG( | -0.41939 | 2.37E-09 | YES | NO | NA |
| Nsa2 | ENSMUSG( | ENSMUSG( | 0.159724 | 0.140955 | NO | NO | NA |
| Rab7 | ENSMUSG( | ENSMUSG( | -0.12637 | 0.037644 | YES | NO | NA |
| Pnisr | ENSMUSG( | ENSMUSG( | -0.31573 | 0.000827 | YES | NO | NA |
| Mybbp1a | ENSMUSG( | ENSMUSG( | 0.616976 | 1.08E-06 | YES | NO | NA |
| Cbl | ENSMUSG( | ENSMUSG( | -0.23699 | 4.43E-05 | YES | NO | NA |
| AU020206 | ENSMUSG( | ENSMUSG( | -0.42882 | 0.000504 | YES | NO | NA |
| Stip1 | ENSMUSG( | ENSMUSG( | -0.15072 | 0.145608 | NO | NO | NA |
| Nsd1 | ENSMUSG( | ENSMUSG( | -0.34738 | 1.29E-06 | YES | NO | NA |
| Arhgef2 | ENSMUSG( | ENSMUSG( | -0.35154 | 8.05E-11 | YES | NO | NA |
| Usp1 | ENSMUSG( | ENSMUSG( | 0.165586 | 0.060251 | NO | NO | NA |
| Sec24c | ENSMUSG( | ENSMUSG( | -0.25053 | 1.06E-13 | YES | NO | NA |
| Bclaf1 | ENSMUSG( | ENSMUSG( | 0.115403 | 0.201619 | NO | NO | NA |
| Psmb4 | ENSMUSG( | ENSMUSG( | 0.271423 | 6.41E-06 | YES | NO | NA |
| Ptpn1 | ENSMUSG( | ENSMUSG( | -0.07809 | 0.293832 | NO | NO | NA |
| Phgdh | ENSMUSG( | ENSMUSG( | 0.681176 | 3.82E-10 | YES | NO | NA |
| Coro2a | ENSMUSG( | ENSMUSG( | -0.0636 | 0.277583 | NO | NO | NA |
| Coro7 | ENSMUSG( | ENSMUSG( | 0.156175 | 0.086269 | NO | NO | NA |
| Srebf2 | ENSMUSG( | ENSMUSG( | 0.210166 | 0.00817 | YES | NO | NA |
| Ankrd17 | ENSMUSG( | ENSMUSG( | -0.23568 | 0.000217 | YES | NO | NA |
| Nxf1 | ENSMUSG( | ENSMUSG( | -0.11966 | 0.128766 | NO | NO | NA |
| Dgkd | ENSMUSG( | ENSMUSG( | -0.5006 | 5.15E-15 | YES | NO | NA |
| Ahcy | ENSMUSG( | ENSMUSG( | 0.633305 | 1.21E-09 | YES | NO | NA |
| Rnf10 | ENSMUSG( | ENSMUSG( | 0.100622 | 0.186976 | NO | NO | NA |

| | | | | | | | |
|---|---|---|---|---|---|---|---|
| Dusp11 | ENSMUSG( | ENSMUSG( | -0.15207 | 0.030282 | YES | NO | NA |
| Raver1 | ENSMUSG( | ENSMUSG( | 0.009729 | 0.83101 | NO | NO | NA |
| Cd53 | ENSMUSG( | ENSMUSG( | -0.14235 | 0.170195 | NO | NO | NA |
| Stag2 | ENSMUSG( | ENSMUSG( | -0.1348 | 0.063933 | NO | NO | NA |
| Cct6a | ENSMUSG( | ENSMUSG( | 0.400053 | 1.77E-05 | YES | NO | NA |
| Arpc3 | ENSMUSG( | ENSMUSG( | -0.08976 | 0.086923 | NO | NO | NA |
| Scd2 | ENSMUSG( | ENSMUSG( | 0.805795 | 4.48E-07 | YES | NO | NA |
| Samd9l | ENSMUSG( | ENSMUSG( | -0.77472 | 1.07E-17 | YES | NO | NA |
| Lman2 | ENSMUSG( | ENSMUSG( | 0.097821 | 0.017188 | YES | NO | NA |
| Pgam1 | ENSMUSG( | ENSMUSG( | 0.632721 | 6.40E-06 | YES | NO | NA |
| Prpf4b | ENSMUSG( | ENSMUSG( | -0.13072 | 0.073383 | NO | NO | NA |
| Gnai3 | ENSMUSG( | ENSMUSG( | 0.000749 | 0.991711 | NO | NO | NA |
| Cdv3 | ENSMUSG( | ENSMUSG( | 0.252707 | 0.002283 | YES | NO | NA |
| Atp13a3 | ENSMUSG( | ENSMUSG( | 0.164576 | 0.006855 | YES | NO | NA |
| Coro1b | ENSMUSG( | ENSMUSG( | -0.23258 | 2.12E-11 | YES | NO | NA |
| Sars | ENSMUSG( | ENSMUSG( | 0.059513 | 0.505653 | NO | NO | NA |
| Atp11b | ENSMUSG( | ENSMUSG( | -0.20007 | 0.011111 | YES | NO | NA |
| Tgoln1 | ENSMUSG( | ENSMUSG( | -0.4131 | 3.26E-08 | YES | NO | NA |
| Dut | ENSMUSG( | ENSMUSG( | 0.326784 | 4.55E-07 | YES | NO | NA |
| Ap2b1 | ENSMUSG( | ENSMUSG( | -0.32269 | 1.77E-12 | YES | NO | NA |
| Gmfg | ENSMUSG( | ENSMUSG( | 0.081692 | 0.559743 | NO | NO | NA |
| Nucks1 | ENSMUSG( | ENSMUSG( | 0.210844 | 0.009993 | YES | NO | NA |
| Grb2 | ENSMUSG( | ENSMUSG( | -0.16837 | 0.025777 | YES | NO | NA |
| Zmiz2 | ENSMUSG( | ENSMUSG( | 0.024488 | 0.714178 | NO | NO | NA |
| Slc25a4 | ENSMUSG( | ENSMUSG( | 0.484437 | 2.20E-06 | YES | NO | NA |
| G3bp2 | ENSMUSG( | ENSMUSG( | 0.000867 | 0.987528 | NO | NO | NA |
| Cd27 | ENSMUSG( | ENSMUSG( | -0.36746 | 3.29E-05 | YES | NO | NA |
| Ncaph2 | ENSMUSG( | ENSMUSG( | 0.168779 | 0.001099 | YES | NO | NA |
| Ppp1r15b | ENSMUSG( | ENSMUSG( | -0.2799 | 1.48E-08 | YES | NO | NA |
| Ube2l3 | ENSMUSG( | ENSMUSG( | 0.110254 | 0.2748 | NO | NO | NA |
| Cenpe | ENSMUSG( | ENSMUSG( | -0.11828 | 0.343845 | NO | NO | NA |
| Agfg1 | ENSMUSG( | ENSMUSG( | -0.353 | 1.12E-07 | YES | NO | NA |
| Niban1 | ENSMUSG( | ENSMUSG( | -0.71444 | 2.19E-12 | YES | NO | NA |
| Slc9a3r1 | ENSMUSG( | ENSMUSG( | -0.40461 | 1.62E-08 | YES | NO | NA |
| Atp2b4 | ENSMUSG( | ENSMUSG( | -0.33888 | 2.55E-08 | YES | NO | NA |
| Surf4 | ENSMUSG( | ENSMUSG( | -0.02701 | 0.638931 | NO | NO | NA |
| Dennd1c | ENSMUSG( | ENSMUSG( | -0.39258 | 7.32E-09 | YES | NO | NA |
| Ddit4 | ENSMUSG( | ENSMUSG( | 0.93453 | 3.44E-10 | YES | NO | NA |
| Uhmk1 | ENSMUSG( | ENSMUSG( | -0.16216 | 0.02382 | YES | NO | NA |
| Eml3 | ENSMUSG( | ENSMUSG( | -0.22937 | 1.09E-06 | YES | NO | NA |
| Atf4 | ENSMUSG( | ENSMUSG( | -0.04449 | 0.636498 | NO | NO | NA |
| Vsir | ENSMUSG( | ENSMUSG( | -0.69551 | 4.11E-13 | YES | NO | NA |
| Ube2v1 | ENSMUSG( | ENSMUSG( | -0.01575 | 0.852038 | NO | NO | NA |
| Spen | ENSMUSG( | ENSMUSG( | -0.52825 | 6.06E-13 | YES | NO | NA |
| Mink1 | ENSMUSG( | ENSMUSG( | -0.0524 | 0.534005 | NO | NO | NA |
| Puf60 | ENSMUSG( | ENSMUSG( | -0.10024 | 0.07542 | NO | NO | NA |
| Zc3h4 | ENSMUSG( | ENSMUSG( | -0.07114 | 0.417925 | NO | NO | NA |

| | | | | | | | |
|---|---|---|---|---|---|---|---|
| Epb41 | ENSMUSG( | ENSMUSG( | -0.34347 | 2.57E-05 | YES | NO | NA |
| Rbck1 | ENSMUSG( | ENSMUSG( | -0.15394 | 0.002587 | YES | NO | NA |
| Nfkbia | ENSMUSG( | ENSMUSG( | 0.493008 | 0.006004 | YES | NO | NA |
| Arrb2 | ENSMUSG( | ENSMUSG( | -0.05668 | 0.341079 | NO | NO | NA |
| Galnt1 | ENSMUSG( | ENSMUSG( | 0.027938 | 0.643873 | NO | NO | NA |
| Sh2d3c | ENSMUSG( | ENSMUSG( | -0.58841 | 1.00E-12 | YES | NO | NA |
| Slk | ENSMUSG( | ENSMUSG( | -0.04958 | 0.606953 | NO | NO | NA |
| Norad | ENSMUSG( | ENSMUSG( | 0.006558 | 0.92992 | NO | NO | NA |
| Mif | ENSMUSG( | ENSMUSG( | 0.737745 | 2.31E-07 | YES | NO | NA |
| Rnf44 | ENSMUSG( | ENSMUSG( | -0.44652 | 3.40E-10 | YES | NO | NA |
| Ptpa | ENSMUSG( | ENSMUSG( | 0.110889 | 0.047542 | YES | NO | NA |
| Rpl36al | ENSMUSG( | ENSMUSG( | 0.177226 | 0.002338 | YES | NO | NA |
| Zfp91 | ENSMUSG( | ENSMUSG( | -0.03447 | 0.706841 | NO | NO | NA |
| Tfdp1 | ENSMUSG( | ENSMUSG( | 0.347251 | 2.24E-06 | YES | NO | NA |
| Tmem50a | ENSMUSG( | ENSMUSG( | -0.10117 | 0.33719 | NO | NO | NA |
| Map4k1 | ENSMUSG( | ENSMUSG( | -0.10368 | 0.066692 | NO | NO | NA |
| Yy1 | ENSMUSG( | ENSMUSG( | -0.12091 | 0.041403 | YES | NO | NA |
| Etf1 | ENSMUSG( | ENSMUSG( | 0.414456 | 1.48E-07 | YES | NO | NA |
| Rpl38 | ENSMUSG( | ENSMUSG( | 0.209024 | 0.004832 | YES | NO | NA |
| Cxcr6 | ENSMUSG( | ENSMUSG( | -0.45706 | 4.96E-05 | YES | NO | NA |
| Cyrib | ENSMUSG( | ENSMUSG( | -0.04831 | 0.518497 | NO | NO | NA |
| Cnot6 | ENSMUSG( | ENSMUSG( | -0.08493 | 0.347686 | NO | NO | NA |
| Dcaf7 | ENSMUSG( | ENSMUSG( | -0.16265 | 0.00396 | YES | NO | NA |
| Aars | ENSMUSG( | ENSMUSG( | 0.12432 | 0.026496 | YES | NO | NA |
| Psme2 | ENSMUSG( | ENSMUSG( | -0.1141 | 0.006527 | YES | NO | NA |
| Psmb9 | ENSMUSG( | ENSMUSG( | -0.11021 | 0.034536 | YES | NO | NA |
| Nipbl | ENSMUSG( | ENSMUSG( | -0.33909 | 2.96E-06 | YES | NO | NA |
| ENSMUSG | ENSMUSG( | ENSMUSG( | -0.00722 | 0.974234 | NO | NO | NA |
| Gak | ENSMUSG( | ENSMUSG( | -0.08361 | 0.256547 | NO | NO | NA |
| Amfr | ENSMUSG( | ENSMUSG( | -0.12481 | 0.037644 | YES | NO | NA |
| Arap2 | ENSMUSG( | ENSMUSG( | -0.19773 | 0.022452 | YES | NO | NA |
| Nlrc3 | ENSMUSG( | ENSMUSG( | -0.30689 | 0.000151 | YES | NO | NA |
| Cep350 | ENSMUSG( | ENSMUSG( | -0.4658 | 1.21E-10 | YES | NO | NA |
| Lax1 | ENSMUSG( | ENSMUSG( | -0.65269 | 2.20E-06 | YES | NO | NA |
| Sbno1 | ENSMUSG( | ENSMUSG( | -0.16873 | 0.000192 | YES | NO | NA |
| Cxcr3 | ENSMUSG( | ENSMUSG( | -0.15881 | 0.11815 | NO | NO | NA |
| Srsf4 | ENSMUSG( | ENSMUSG( | 0.027833 | 0.729736 | NO | NO | NA |
| Rab5c | ENSMUSG( | ENSMUSG( | -0.12887 | 0.054978 | NO | NO | NA |
| Nedd9 | ENSMUSG( | ENSMUSG( | -0.5516 | 3.19E-09 | YES | NO | NA |
| Usp34 | ENSMUSG( | ENSMUSG( | -0.25175 | 7.88E-06 | YES | NO | NA |
| Rrp1 | ENSMUSG( | ENSMUSG( | 0.275083 | 0.000385 | YES | NO | NA |
| Sdf4 | ENSMUSG( | ENSMUSG( | 0.067315 | 0.25474 | NO | NO | NA |
| Smc6 | ENSMUSG( | ENSMUSG( | -0.00819 | 0.930577 | NO | NO | NA |
| Nipal3 | ENSMUSG( | ENSMUSG( | -0.64168 | 1.17E-15 | YES | NO | NA |
| Mir142hg | ENSMUSG( | ENSMUSG( | -0.8045 | 2.29E-14 | YES | NO | NA |
| Cad | ENSMUSG( | ENSMUSG( | 0.546744 | 1.92E-07 | YES | NO | NA |
| Ckap5 | ENSMUSG( | ENSMUSG( | -0.08837 | 0.239113 | NO | NO | NA |

| | | | | | | | |
|---|---|---|---|---|---|---|---|
| Nrd1 | ENSMUSG( | ENSMUSG( | -0.15869 | 0.014014 | YES | NO | NA |
| Supt6 | ENSMUSG( | ENSMUSG( | -0.11961 | 0.054304 | NO | NO | NA |
| Clta | ENSMUSG( | ENSMUSG( | 0.000383 | 0.995418 | NO | NO | NA |
| Rab10 | ENSMUSG( | ENSMUSG( | -0.10129 | 0.137221 | NO | NO | NA |
| Kdm4a | ENSMUSG( | ENSMUSG( | -0.47283 | 6.87E-09 | YES | NO | NA |
| Pitpnm1 | ENSMUSG( | ENSMUSG( | -0.15764 | 0.051369 | NO | NO | NA |
| Capza2 | ENSMUSG( | ENSMUSG( | 0.096727 | 0.246461 | NO | NO | NA |
| Znrf2 | ENSMUSG( | ENSMUSG( | -0.39975 | 9.58E-10 | YES | NO | NA |
| Carmil2 | ENSMUSG( | ENSMUSG( | -0.08887 | 0.216788 | NO | NO | NA |
| Ppp4r3a | ENSMUSG( | ENSMUSG( | -0.14222 | 0.027444 | YES | NO | NA |
| Tmod3 | ENSMUSG( | ENSMUSG( | -0.25776 | 0.000109 | YES | NO | NA |
| Map4k4 | ENSMUSG( | ENSMUSG( | -0.34031 | 1.21E-06 | YES | NO | NA |
| ENSMUSG | ENSMUSG( | ENSMUSG( | -0.31069 | 0.05234 | NO | NO | NA |
| Nup153 | ENSMUSG( | ENSMUSG( | -0.24746 | 0.000267 | YES | NO | NA |
| Rap1a | ENSMUSG( | ENSMUSG( | 0.001774 | 0.985756 | NO | NO | NA |
| Ezh2 | ENSMUSG( | ENSMUSG( | 0.117687 | 0.159401 | NO | NO | NA |
| Sp110 | ENSMUSG( | ENSMUSG( | -0.36309 | 5.35E-08 | YES | NO | NA |
| Tnk2 | ENSMUSG( | ENSMUSG( | -0.15174 | 0.010079 | YES | NO | NA |
| Ccnb2 | ENSMUSG( | ENSMUSG( | 0.50572 | 1.52E-07 | YES | NO | NA |
| Rpl34-ps1 | ENSMUSG( | ENSMUSG( | 7.100482 | 0 | YES | YES | Upregulated |
| Glg1 | ENSMUSG( | ENSMUSG( | -0.2376 | 1.73E-05 | YES | NO | NA |
| Prr13 | ENSMUSG( | ENSMUSG( | -0.44104 | 4.90E-09 | YES | NO | NA |
| Ppp1cb | ENSMUSG( | ENSMUSG( | -0.18233 | 0.00123 | YES | NO | NA |
| Zc3h18 | ENSMUSG( | ENSMUSG( | -0.08644 | 0.377002 | NO | NO | NA |
| Arih1 | ENSMUSG( | ENSMUSG( | -0.1524 | 0.016464 | YES | NO | NA |
| Khnyn | ENSMUSG( | ENSMUSG( | -0.61493 | 4.07E-16 | YES | NO | NA |
| Cenpf | ENSMUSG( | ENSMUSG( | 0.054724 | 0.700004 | NO | NO | NA |
| Tut7 | ENSMUSG( | ENSMUSG( | -0.31348 | 4.90E-05 | YES | NO | NA |
| Rdx | ENSMUSG( | ENSMUSG( | -0.06035 | 0.474574 | NO | NO | NA |
| Tigit | ENSMUSG( | ENSMUSG( | -0.04511 | 0.78099 | NO | NO | NA |
| Ifi206 | ENSMUSG( | ENSMUSG( | -1.14126 | 3.11E-19 | YES | YES | Downregulated |
| Rfwd3 | ENSMUSG( | ENSMUSG( | 0.081501 | 0.244982 | NO | NO | NA |
| Rbms1 | ENSMUSG( | ENSMUSG( | -0.0217 | 0.768227 | NO | NO | NA |
| Csnk1d | ENSMUSG( | ENSMUSG( | -0.15502 | 0.00349 | YES | NO | NA |
| Glipr2 | ENSMUSG( | ENSMUSG( | -0.2256 | 0.000646 | YES | NO | NA |
| Sod1 | ENSMUSG( | ENSMUSG( | 0.254933 | 0.000589 | YES | NO | NA |
| Ints1 | ENSMUSG( | ENSMUSG( | 0.002881 | 0.973166 | NO | NO | NA |
| Tmem30a | ENSMUSG( | ENSMUSG( | -0.14392 | 0.023606 | YES | NO | NA |
| Upf1 | ENSMUSG( | ENSMUSG( | -0.11388 | 0.050469 | NO | NO | NA |
| Cnot6l | ENSMUSG( | ENSMUSG( | -0.19953 | 0.043563 | YES | NO | NA |
| Gna13 | ENSMUSG( | ENSMUSG( | -0.25003 | 1.38E-05 | YES | NO | NA |
| Havcr2 | ENSMUSG( | ENSMUSG( | 0.111844 | 0.270046 | NO | NO | NA |
| Luc7l3 | ENSMUSG( | ENSMUSG( | -0.0306 | 0.775658 | NO | NO | NA |
| Ddx39a | ENSMUSG( | ENSMUSG( | 0.449138 | 2.26E-12 | YES | NO | NA |
| Tram1 | ENSMUSG( | ENSMUSG( | -0.22657 | 7.14E-08 | YES | NO | NA |
| Tnpo1 | ENSMUSG( | ENSMUSG( | 0.146605 | 0.00654 | YES | NO | NA |
| Mtch1 | ENSMUSG( | ENSMUSG( | 0.417184 | 1.83E-08 | YES | NO | NA |

| | | | | | | | |
|---|---|---|---|---|---|---|---|
| Lsm14a | ENSMUSG( | ENSMUSG( | -0.15092 | 0.00533 | YES | NO | NA |
| Tpst2 | ENSMUSG( | ENSMUSG( | -0.29374 | 1.08E-14 | YES | NO | NA |
| Zbtb7a | ENSMUSG( | ENSMUSG( | -0.29082 | 5.99E-07 | YES | NO | NA |
| Eif3l | ENSMUSG( | ENSMUSG( | 0.390771 | 2.51E-08 | YES | NO | NA |
| Sytl3 | ENSMUSG( | ENSMUSG( | 0.254476 | 0.001813 | YES | NO | NA |
| Eif5b | ENSMUSG( | ENSMUSG( | 0.118358 | 0.476489 | NO | NO | NA |
| Tnrc6a | ENSMUSG( | ENSMUSG( | -0.18938 | 0.025795 | YES | NO | NA |
| Plac8 | ENSMUSG( | ENSMUSG( | 0.285176 | 0.022628 | YES | NO | NA |
| Ranbp1 | ENSMUSG( | ENSMUSG( | 0.399318 | 0.007467 | YES | NO | NA |
| Gas5 | ENSMUSG( | ENSMUSG( | 0.33585 | 0.002798 | YES | NO | NA |
| Pnrc1 | ENSMUSG( | ENSMUSG( | -0.17432 | 0.256595 | NO | NO | NA |
| Kpna4 | ENSMUSG( | ENSMUSG( | -0.29543 | 3.39E-06 | YES | NO | NA |
| Anapc1 | ENSMUSG( | ENSMUSG( | 0.072779 | 0.109539 | NO | NO | NA |
| Gse1 | ENSMUSG( | ENSMUSG( | -0.82246 | 1.53E-12 | YES | NO | NA |
| Ehbp1l1 | ENSMUSG( | ENSMUSG( | -0.35059 | 9.66E-09 | YES | NO | NA |
| Nap1l4 | ENSMUSG( | ENSMUSG( | 0.051685 | 0.495773 | NO | NO | NA |
| Cd5 | ENSMUSG( | ENSMUSG( | -0.48413 | 4.56E-08 | YES | NO | NA |
| Nisch | ENSMUSG( | ENSMUSG( | -0.18477 | 0.003998 | YES | NO | NA |
| Atp6v0a2 | ENSMUSG( | ENSMUSG( | -0.26654 | 4.28E-09 | YES | NO | NA |
| Ifi27l2a | ENSMUSG( | ENSMUSG( | -1.34484 | 1.05E-09 | YES | YES | Downregulated |
| Ube2s | ENSMUSG( | ENSMUSG( | 0.092975 | 0.313701 | NO | NO | NA |
| Man2b1 | ENSMUSG( | ENSMUSG( | -0.35907 | 8.13E-12 | YES | NO | NA |
| Slbp | ENSMUSG( | ENSMUSG( | 0.024988 | 0.736231 | NO | NO | NA |
| Dock5 | ENSMUSG( | ENSMUSG( | -1.02946 | 1.85E-22 | YES | YES | Downregulated |
| Slc38a2 | ENSMUSG( | ENSMUSG( | -0.1659 | 0.002627 | YES | NO | NA |
| Rab11fip4 | ENSMUSG( | ENSMUSG( | -0.54608 | 8.36E-09 | YES | NO | NA |
| mt-Atp8 | ENSMUSG( | ENSMUSG( | 0.207625 | 0.022915 | YES | NO | NA |
| Phb2 | ENSMUSG( | ENSMUSG( | 0.474513 | 1.64E-08 | YES | NO | NA |
| Sltm | ENSMUSG( | ENSMUSG( | -0.01319 | 0.912025 | NO | NO | NA |
| Tax1bp1 | ENSMUSG( | ENSMUSG( | -0.10396 | 0.304215 | NO | NO | NA |
| Rtn3 | ENSMUSG( | ENSMUSG( | -0.2693 | 5.95E-08 | YES | NO | NA |
| Ptp4a3 | ENSMUSG( | ENSMUSG( | -0.53425 | 1.06E-12 | YES | NO | NA |
| Adar | ENSMUSG( | ENSMUSG( | -0.46914 | 1.88E-13 | YES | NO | NA |
| Clk1 | ENSMUSG( | ENSMUSG( | 0.012178 | 0.933155 | NO | NO | NA |
| Ddx54 | ENSMUSG( | ENSMUSG( | 0.001467 | 0.982791 | NO | NO | NA |
| Ccl3 | ENSMUSG( | ENSMUSG( | 0.099205 | 0.609095 | NO | NO | NA |
| Nin | ENSMUSG( | ENSMUSG( | -0.05335 | 0.563645 | NO | NO | NA |
| Uba2 | ENSMUSG( | ENSMUSG( | 0.221821 | 0.000855 | YES | NO | NA |
| Ssr3 | ENSMUSG( | ENSMUSG( | 0.122962 | 0.037244 | YES | NO | NA |
| Atad2 | ENSMUSG( | ENSMUSG( | -0.07557 | 0.353941 | NO | NO | NA |
| Ugcg | ENSMUSG( | ENSMUSG( | -0.28152 | 0.001032 | YES | NO | NA |
| Dnaja2 | ENSMUSG( | ENSMUSG( | 0.159488 | 0.038375 | YES | NO | NA |
| Gars | ENSMUSG( | ENSMUSG( | 0.184571 | 6.77E-05 | YES | NO | NA |
| Ier2 | ENSMUSG( | ENSMUSG( | 0.026415 | 0.858038 | NO | NO | NA |
| Zc3h7a | ENSMUSG( | ENSMUSG( | -0.08872 | 0.344904 | NO | NO | NA |
| Rnf166 | ENSMUSG( | ENSMUSG( | -0.30555 | 0.00121 | YES | NO | NA |
| Zfp106 | ENSMUSG( | ENSMUSG( | 0.244862 | 7.17E-05 | YES | NO | NA |

| | | | | | | | |
|---|---|---|---|---|---|---|---|
| Atp5pb | ENSMUSG( | ENSMUSG( | 0.125944 | 0.017246 | YES | NO | NA |
| Skiv2l | ENSMUSG( | ENSMUSG( | -0.17896 | 0.000155 | YES | NO | NA |
| Card11 | ENSMUSG( | ENSMUSG( | -0.85424 | 3.26E-14 | YES | NO | NA |
| Akap8 | ENSMUSG( | ENSMUSG( | -0.09823 | 0.231925 | NO | NO | NA |
| Bub1b | ENSMUSG( | ENSMUSG( | 0.271063 | 0.006807 | YES | NO | NA |
| Pcnt | ENSMUSG( | ENSMUSG( | -0.03765 | 0.675653 | NO | NO | NA |
| Arl4c | ENSMUSG( | ENSMUSG( | -0.54373 | 8.75E-09 | YES | NO | NA |
| Adamts10 | ENSMUSG( | ENSMUSG( | -0.20951 | 0.000567 | YES | NO | NA |
| Map7d1 | ENSMUSG( | ENSMUSG( | -0.15616 | 0.093836 | NO | NO | NA |
| Azin1 | ENSMUSG( | ENSMUSG( | -0.20608 | 0.000555 | YES | NO | NA |
| Pml | ENSMUSG( | ENSMUSG( | -0.52228 | 1.15E-19 | YES | NO | NA |
| Dnajb6 | ENSMUSG( | ENSMUSG( | -0.05054 | 0.560142 | NO | NO | NA |
| Sri | ENSMUSG( | ENSMUSG( | -0.04118 | 0.658376 | NO | NO | NA |
| Odf2 | ENSMUSG( | ENSMUSG( | 0.004249 | 0.950039 | NO | NO | NA |
| Eef1d | ENSMUSG( | ENSMUSG( | 0.177765 | 0.015854 | YES | NO | NA |
| Chd2 | ENSMUSG( | ENSMUSG( | -0.33622 | 0.00036 | YES | NO | NA |
| Phf20l1 | ENSMUSG( | ENSMUSG( | -0.36116 | 1.51E-06 | YES | NO | NA |
| Ppp3cb | ENSMUSG( | ENSMUSG( | -0.03416 | 0.645299 | NO | NO | NA |
| Matr3 | ENSMUSG( | ENSMUSG( | 0.063632 | 0.311172 | NO | NO | NA |
| Hadha | ENSMUSG( | ENSMUSG( | 0.102114 | 0.090643 | NO | NO | NA |
| Rab1b | ENSMUSG( | ENSMUSG( | 0.108735 | 0.178852 | NO | NO | NA |
| Crybg1 | ENSMUSG( | ENSMUSG( | -0.10647 | 0.172523 | NO | NO | NA |
| Otud4 | ENSMUSG( | ENSMUSG( | -0.14162 | 0.003097 | YES | NO | NA |
| Txnip | ENSMUSG( | ENSMUSG( | -0.27286 | 0.050875 | NO | NO | NA |
| Snd1 | ENSMUSG( | ENSMUSG( | 0.118325 | 0.10042 | NO | NO | NA |
| Pitpnc1 | ENSMUSG( | ENSMUSG( | -0.56871 | 7.84E-09 | YES | NO | NA |
| Oip5os1 | ENSMUSG( | ENSMUSG( | -0.43123 | 1.25E-05 | YES | NO | NA |
| Ipo7 | ENSMUSG( | ENSMUSG( | 0.218114 | 0.000655 | YES | NO | NA |
| Cct3 | ENSMUSG( | ENSMUSG( | 0.363022 | 1.05E-06 | YES | NO | NA |
| H2ax | ENSMUSG( | ENSMUSG( | 0.022714 | 0.859911 | NO | NO | NA |
| Dnajc5 | ENSMUSG( | ENSMUSG( | -0.22507 | 0.000849 | YES | NO | NA |
| Top1 | ENSMUSG( | ENSMUSG( | -0.16303 | 0.176712 | NO | NO | NA |
| Stk39 | ENSMUSG( | ENSMUSG( | -0.07162 | 0.302183 | NO | NO | NA |
| Paics | ENSMUSG( | ENSMUSG( | 0.334268 | 6.73E-06 | YES | NO | NA |
| Cox5b | ENSMUSG( | ENSMUSG( | 0.22419 | 0.139742 | NO | NO | NA |
| Hnrnpr | ENSMUSG( | ENSMUSG( | 0.056417 | 0.561165 | NO | NO | NA |
| Rora | ENSMUSG( | ENSMUSG( | -0.3432 | 0.03088 | YES | NO | NA |
| Bcl2 | ENSMUSG( | ENSMUSG( | -0.62642 | 8.07E-05 | YES | NO | NA |
| Psme3 | ENSMUSG( | ENSMUSG( | 0.184031 | 0.026261 | YES | NO | NA |
| Irf2 | ENSMUSG( | ENSMUSG( | -0.53976 | 6.07E-07 | YES | NO | NA |
| Nfkb1 | ENSMUSG( | ENSMUSG( | -0.28964 | 0.000233 | YES | NO | NA |
| AV099323 | ENSMUSG( | ENSMUSG( | 0.133854 | 0.516761 | NO | NO | NA |
| Eprs | ENSMUSG( | ENSMUSG( | 0.379525 | 1.91E-05 | YES | NO | NA |
| Rpa2 | ENSMUSG( | ENSMUSG( | 0.127621 | 0.040983 | YES | NO | NA |
| Api5 | ENSMUSG( | ENSMUSG( | 0.123442 | 0.054866 | NO | NO | NA |
| Stat5b | ENSMUSG( | ENSMUSG( | -0.04153 | 0.529609 | NO | NO | NA |
| Prdx1 | ENSMUSG( | ENSMUSG( | 0.049521 | 0.413266 | NO | NO | NA |

| | | | | | | | |
|---|---|---|---|---|---|---|---|
| Csnk2a1 | ENSMUSG( | ENSMUSG( | -0.18891 | 0.001847 | YES | NO | NA |
| Kdm5c | ENSMUSG( | ENSMUSG( | -0.35356 | 2.65E-12 | YES | NO | NA |
| Sf3a1 | ENSMUSG( | ENSMUSG( | 0.022001 | 0.733562 | NO | NO | NA |
| Nrgn | ENSMUSG( | ENSMUSG( | 1.607853 | 1.26E-07 | YES | YES | Upregulated |
| Vav1 | ENSMUSG( | ENSMUSG( | -0.08869 | 0.069686 | NO | NO | NA |
| Pold1 | ENSMUSG( | ENSMUSG( | 0.186863 | 0.04489 | YES | NO | NA |
| Edem1 | ENSMUSG( | ENSMUSG( | -0.09431 | 0.083596 | NO | NO | NA |
| Stk38 | ENSMUSG( | ENSMUSG( | -0.34773 | 5.67E-08 | YES | NO | NA |
| Gltp | ENSMUSG( | ENSMUSG( | -0.06707 | 0.26277 | NO | NO | NA |
| Xpo6 | ENSMUSG( | ENSMUSG( | -0.04934 | 0.252281 | NO | NO | NA |
| Sympk | ENSMUSG( | ENSMUSG( | -0.08047 | 0.043227 | YES | NO | NA |
| Nop56 | ENSMUSG( | ENSMUSG( | 0.401311 | 0.000156 | YES | NO | NA |
| Psmc3 | ENSMUSG( | ENSMUSG( | 0.162114 | 0.032527 | YES | NO | NA |
| Tuba1a | ENSMUSG( | ENSMUSG( | 0.06701 | 0.43629 | NO | NO | NA |
| Txnrd1 | ENSMUSG( | ENSMUSG( | -0.06068 | 0.275621 | NO | NO | NA |
| Sidt2 | ENSMUSG( | ENSMUSG( | -0.26466 | 0.000566 | YES | NO | NA |
| Ppp6r3 | ENSMUSG( | ENSMUSG( | -0.10017 | 0.11748 | NO | NO | NA |
| Rnps1 | ENSMUSG( | ENSMUSG( | 0.225199 | 0.00097 | YES | NO | NA |
| Ppp1r16b | ENSMUSG( | ENSMUSG( | -0.26052 | 0.006919 | YES | NO | NA |
| Adam10 | ENSMUSG( | ENSMUSG( | -0.19349 | 0.003734 | YES | NO | NA |
| Abca7 | ENSMUSG( | ENSMUSG( | -0.32569 | 8.37E-08 | YES | NO | NA |
| Ccnl1 | ENSMUSG( | ENSMUSG( | -0.19692 | 0.001662 | YES | NO | NA |
| Tmed9 | ENSMUSG( | ENSMUSG( | 0.13258 | 0.221922 | NO | NO | NA |
| Med13 | ENSMUSG( | ENSMUSG( | -0.44641 | 1.08E-08 | YES | NO | NA |
| Ppm1h | ENSMUSG( | ENSMUSG( | -0.19425 | 0.003991 | YES | NO | NA |
| 9930111J2 | ENSMUSG( | ENSMUSG( | -0.82771 | 9.49E-15 | YES | NO | NA |
| Clspn | ENSMUSG( | ENSMUSG( | 0.354912 | 0.000109 | YES | NO | NA |
| Psmd8 | ENSMUSG( | ENSMUSG( | 0.216752 | 0.001203 | YES | NO | NA |
| Xrn1 | ENSMUSG( | ENSMUSG( | -0.34051 | 3.24E-06 | YES | NO | NA |
| Sdcbp | ENSMUSG( | ENSMUSG( | -0.13722 | 0.160355 | NO | NO | NA |
| Copb2 | ENSMUSG( | ENSMUSG( | 0.057757 | 0.465973 | NO | NO | NA |
| Rcsd1 | ENSMUSG( | ENSMUSG( | 0.233339 | 0.010574 | YES | NO | NA |
| Fubp1 | ENSMUSG( | ENSMUSG( | -0.04574 | 0.512859 | NO | NO | NA |
| Fip1l1 | ENSMUSG( | ENSMUSG( | -0.27768 | 0.000821 | YES | NO | NA |
| Ncapg2 | ENSMUSG( | ENSMUSG( | 0.380004 | 1.39E-08 | YES | NO | NA |
| Cd37 | ENSMUSG( | ENSMUSG( | -0.39693 | 1.04E-10 | YES | NO | NA |
| Rasgrp1 | ENSMUSG( | ENSMUSG( | -0.5829 | 5.47E-09 | YES | NO | NA |
| Arhgef6 | ENSMUSG( | ENSMUSG( | -0.14782 | 0.018872 | YES | NO | NA |
| Klf3 | ENSMUSG( | ENSMUSG( | -1.54164 | 9.60E-08 | YES | YES | Downregulated |
| Il17ra | ENSMUSG( | ENSMUSG( | -0.50979 | 7.99E-11 | YES | NO | NA |
| Aco2 | ENSMUSG( | ENSMUSG( | 0.116109 | 0.013041 | YES | NO | NA |
| Pabpc4 | ENSMUSG( | ENSMUSG( | 0.478526 | 1.78E-06 | YES | NO | NA |
| Lime1 | ENSMUSG( | ENSMUSG( | -0.0445 | 0.743424 | NO | NO | NA |
| Nacc1 | ENSMUSG( | ENSMUSG( | -0.15628 | 0.000172 | YES | NO | NA |
| Naa15 | ENSMUSG( | ENSMUSG( | 0.222763 | 0.024247 | YES | NO | NA |
| S1pr1 | ENSMUSG( | ENSMUSG( | -0.43592 | 0.000269 | YES | NO | NA |
| Setd5 | ENSMUSG( | ENSMUSG( | -0.27482 | 5.42E-06 | YES | NO | NA |

| | | | | | | | |
|---|---|---|---|---|---|---|---|
| Rnf187 | ENSMUSG( | ENSMUSG( | -0.04633 | 0.492946 | NO | NO | NA |
| ENSMUSG( | ENSMUSG( | ENSMUSG( | 0.157581 | 0.130425 | NO | NO | NA |
| Cpne1 | ENSMUSG( | ENSMUSG( | -0.0605 | 0.473939 | NO | NO | NA |
| Ncor2 | ENSMUSG( | ENSMUSG( | 0.069077 | 0.384056 | NO | NO | NA |
| Dctn1 | ENSMUSG( | ENSMUSG( | -0.0847 | 0.35416 | NO | NO | NA |
| Trim30a | ENSMUSG( | ENSMUSG( | -1.00896 | 6.99E-15 | YES | YES | Downregulated |
| Runx2 | ENSMUSG( | ENSMUSG( | -0.09867 | 0.438825 | NO | NO | NA |
| Cct4 | ENSMUSG( | ENSMUSG( | 0.13225 | 0.024146 | YES | NO | NA |
| Tmem123 | ENSMUSG( | ENSMUSG( | -0.17375 | 0.013636 | YES | NO | NA |
| Rbbp6 | ENSMUSG( | ENSMUSG( | -0.00891 | 0.932859 | NO | NO | NA |
| Fgfr1op2 | ENSMUSG( | ENSMUSG( | -0.1903 | 0.000302 | YES | NO | NA |
| Ccar1 | ENSMUSG( | ENSMUSG( | 0.203867 | 0.021303 | YES | NO | NA |
| Csnk1g2 | ENSMUSG( | ENSMUSG( | -0.00459 | 0.955452 | NO | NO | NA |
| Klrc1 | ENSMUSG( | ENSMUSG( | -0.11882 | 0.071248 | NO | NO | NA |
| Fnbp4 | ENSMUSG( | ENSMUSG( | -0.0429 | 0.646058 | NO | NO | NA |
| Ppig | ENSMUSG( | ENSMUSG( | -0.06399 | 0.739912 | NO | NO | NA |
| Ctla4 | ENSMUSG( | ENSMUSG( | 0.394843 | 0.34 | NO | NO | NA |
| Prc1 | ENSMUSG( | ENSMUSG( | -0.04031 | 0.691178 | NO | NO | NA |
| Nr4a2 | ENSMUSG( | ENSMUSG( | -0.12327 | 0.612152 | NO | NO | NA |
| Vps37b | ENSMUSG( | ENSMUSG( | -0.2401 | 0.038194 | YES | NO | NA |
| Gspt1 | ENSMUSG( | ENSMUSG( | 0.275296 | 3.94E-05 | YES | NO | NA |
| Pdcd4 | ENSMUSG( | ENSMUSG( | -0.46897 | 5.33E-06 | YES | NO | NA |
| Dtx1 | ENSMUSG( | ENSMUSG( | -2.00416 | 5.06E-07 | YES | YES | Downregulated |
| Gtpbp2 | ENSMUSG( | ENSMUSG( | -0.22667 | 3.59E-05 | YES | NO | NA |
| Smarcd2 | ENSMUSG( | ENSMUSG( | -0.13725 | 0.001341 | YES | NO | NA |
| Dnajc7 | ENSMUSG( | ENSMUSG( | -0.04261 | 0.561709 | NO | NO | NA |
| Man2a1 | ENSMUSG( | ENSMUSG( | -0.04221 | 0.658781 | NO | NO | NA |
| Gbp6 | ENSMUSG( | ENSMUSG( | -0.70066 | 1.23E-08 | YES | NO | NA |
| Lars2 | ENSMUSG( | ENSMUSG( | 0.867736 | 0.021071 | YES | NO | NA |
| Psmd3 | ENSMUSG( | ENSMUSG( | 0.277362 | 9.56E-05 | YES | NO | NA |
| Pitpnm2 | ENSMUSG( | ENSMUSG( | -0.36014 | 8.19E-05 | YES | NO | NA |
| Wdr82 | ENSMUSG( | ENSMUSG( | -0.15966 | 4.45E-05 | YES | NO | NA |
| Rpn1 | ENSMUSG( | ENSMUSG( | 0.275785 | 0.000943 | YES | NO | NA |
| Ubr2 | ENSMUSG( | ENSMUSG( | -0.37268 | 4.66E-11 | YES | NO | NA |
| Oxsr1 | ENSMUSG( | ENSMUSG( | 0.245736 | 0.015465 | YES | NO | NA |
| Pxn | ENSMUSG( | ENSMUSG( | -0.10837 | 0.15672 | NO | NO | NA |
| Srpk1 | ENSMUSG( | ENSMUSG( | -0.07204 | 0.320809 | NO | NO | NA |
| Was | ENSMUSG( | ENSMUSG( | -0.06283 | 0.294292 | NO | NO | NA |
| Pdia6 | ENSMUSG( | ENSMUSG( | -0.00718 | 0.963858 | NO | NO | NA |
| Ddost | ENSMUSG( | ENSMUSG( | 0.181089 | 0.008394 | YES | NO | NA |
| Leprotl1 | ENSMUSG( | ENSMUSG( | -0.33075 | 4.59E-07 | YES | NO | NA |
| Prpf40a | ENSMUSG( | ENSMUSG( | 0.058799 | 0.515887 | NO | NO | NA |
| Rbm5 | ENSMUSG( | ENSMUSG( | -0.29484 | 3.71E-05 | YES | NO | NA |
| Celf1 | ENSMUSG( | ENSMUSG( | -0.31916 | 1.16E-06 | YES | NO | NA |
| Npc2 | ENSMUSG( | ENSMUSG( | -0.37568 | 6.73E-07 | YES | NO | NA |
| Ppp4c | ENSMUSG( | ENSMUSG( | -0.12117 | 0.083716 | NO | NO | NA |
| Phf3 | ENSMUSG( | ENSMUSG( | -0.30911 | 0.000513 | YES | NO | NA |

| | | | | | | | |
|---|---|---|---|---|---|---|---|
| Trpv2 | ENSMUSG( | ENSMUSG( | 0.226369 | 0.035226 | YES | NO | NA |
| Hbb-bs | ENSMUSG( | ENSMUSG( | 0.411534 | 0.625041 | NO | NO | NA |
| Ssr1 | ENSMUSG( | ENSMUSG( | 0.244406 | 2.56E-05 | YES | NO | NA |
| Sp3 | ENSMUSG( | ENSMUSG( | -0.2333 | 0.000342 | YES | NO | NA |
| Copg1 | ENSMUSG( | ENSMUSG( | -0.07607 | 0.07405 | NO | NO | NA |
| Slc27a4 | ENSMUSG( | ENSMUSG( | -1.10277 | 2.80E-17 | YES | YES | Downregulated |
| Slfn2 | ENSMUSG( | ENSMUSG( | -0.55192 | 8.99E-10 | YES | NO | NA |
| Larp4b | ENSMUSG( | ENSMUSG( | 0.008361 | 0.869303 | NO | NO | NA |
| Etnk1 | ENSMUSG( | ENSMUSG( | -0.36126 | 3.23E-06 | YES | NO | NA |
| Tsc22d3 | ENSMUSG( | ENSMUSG( | 0.3339 | 0.103506 | NO | NO | NA |
| Akt2 | ENSMUSG( | ENSMUSG( | -0.18298 | 7.21E-07 | YES | NO | NA |
| Gbp4 | ENSMUSG( | ENSMUSG( | -0.44548 | 1.48E-05 | YES | NO | NA |
| Tes | ENSMUSG( | ENSMUSG( | -0.24184 | 4.41E-05 | YES | NO | NA |
| Ppp4r3b | ENSMUSG( | ENSMUSG( | -0.01175 | 0.898037 | NO | NO | NA |
| Dtx3l | ENSMUSG( | ENSMUSG( | -0.62522 | 3.16E-17 | YES | NO | NA |
| Eif4a3 | ENSMUSG( | ENSMUSG( | 0.249323 | 3.01E-06 | YES | NO | NA |
| Alyref | ENSMUSG( | ENSMUSG( | 0.265414 | 0.002332 | YES | NO | NA |
| Eif3d | ENSMUSG( | ENSMUSG( | 0.152736 | 0.007201 | YES | NO | NA |
| Ubn1 | ENSMUSG( | ENSMUSG( | -0.25063 | 0.000102 | YES | NO | NA |
| Wapl | ENSMUSG( | ENSMUSG( | -0.06868 | 0.220868 | NO | NO | NA |
| Mapre2 | ENSMUSG( | ENSMUSG( | -0.13642 | 0.016419 | YES | NO | NA |
| Psmc1 | ENSMUSG( | ENSMUSG( | 0.380201 | 0.002077 | YES | NO | NA |
| Bak1 | ENSMUSG( | ENSMUSG( | 0.012161 | 0.89271 | NO | NO | NA |
| Erap1 | ENSMUSG( | ENSMUSG( | 0.020038 | 0.758872 | NO | NO | NA |
| Itpr3 | ENSMUSG( | ENSMUSG( | -0.47715 | 6.47E-08 | YES | NO | NA |
| Fkbp1a | ENSMUSG( | ENSMUSG( | -0.02745 | 0.723011 | NO | NO | NA |
| Gcn1 | ENSMUSG( | ENSMUSG( | -0.02854 | 0.609095 | NO | NO | NA |
| Pten | ENSMUSG( | ENSMUSG( | -0.16132 | 0.006384 | YES | NO | NA |
| Polr2m | ENSMUSG( | ENSMUSG( | -0.04771 | 0.475251 | NO | NO | NA |
| Abi1 | ENSMUSG( | ENSMUSG( | -0.30717 | 0.000283 | YES | NO | NA |
| Eif2s2 | ENSMUSG( | ENSMUSG( | 0.369369 | 0.000472 | YES | NO | NA |
| Ube2h | ENSMUSG( | ENSMUSG( | -0.63464 | 6.82E-12 | YES | NO | NA |
| Il21r | ENSMUSG( | ENSMUSG( | -0.35731 | 5.17E-08 | YES | NO | NA |
| F2r | ENSMUSG( | ENSMUSG( | -0.62943 | 3.82E-10 | YES | NO | NA |
| Gsk3a | ENSMUSG( | ENSMUSG( | -0.16799 | 0.000464 | YES | NO | NA |
| Gramd1b | ENSMUSG( | ENSMUSG( | -0.25245 | 0.011533 | YES | NO | NA |
| Tgfb1 | ENSMUSG( | ENSMUSG( | -0.10024 | 0.193795 | NO | NO | NA |
| Stt3a | ENSMUSG( | ENSMUSG( | 0.008167 | 0.926943 | NO | NO | NA |
| Tcf3 | ENSMUSG( | ENSMUSG( | 0.163346 | 0.007717 | YES | NO | NA |
| Arhgef3 | ENSMUSG( | ENSMUSG( | -0.62077 | 9.32E-11 | YES | NO | NA |
| Ddx23 | ENSMUSG( | ENSMUSG( | -0.05471 | 0.588985 | NO | NO | NA |
| Cabin1 | ENSMUSG( | ENSMUSG( | -0.07249 | 0.406115 | NO | NO | NA |
| Vmp1 | ENSMUSG( | ENSMUSG( | -0.39907 | 5.07E-05 | YES | NO | NA |
| Ilrun | ENSMUSG( | ENSMUSG( | -0.17556 | 4.79E-05 | YES | NO | NA |
| Gbp9 | ENSMUSG( | ENSMUSG( | -0.6818 | 1.25E-08 | YES | NO | NA |
| Tnrc6b | ENSMUSG( | ENSMUSG( | -0.64485 | 1.42E-13 | YES | NO | NA |
| Pik3r1 | ENSMUSG( | ENSMUSG( | -0.4015 | 5.38E-08 | YES | NO | NA |

| | | | | | | | |
|---|---|---|---|---|---|---|---|
| Itgax | ENSMUSG( | ENSMUSG( | -1.29644 | 1.97E-12 | YES | YES | Downregulated |
| Actr1a | ENSMUSG( | ENSMUSG( | 0.023845 | 0.700801 | NO | NO | NA |
| Psma2 | ENSMUSG( | ENSMUSG( | 0.168185 | 0.03323 | YES | NO | NA |
| Galnt6 | ENSMUSG( | ENSMUSG( | -0.39983 | 9.18E-06 | YES | NO | NA |
| Pbrm1 | ENSMUSG( | ENSMUSG( | -0.06569 | 0.286451 | NO | NO | NA |
| Trim56 | ENSMUSG( | ENSMUSG( | -0.62417 | 5.01E-16 | YES | NO | NA |
| Ctdsp1 | ENSMUSG( | ENSMUSG( | -0.26049 | 3.67E-06 | YES | NO | NA |
| Ythdc1 | ENSMUSG( | ENSMUSG( | -0.19748 | 0.027819 | YES | NO | NA |
| Tnrc18 | ENSMUSG( | ENSMUSG( | -0.33037 | 1.92E-06 | YES | NO | NA |
| Usp19 | ENSMUSG( | ENSMUSG( | -0.29006 | 1.58E-10 | YES | NO | NA |
| Impdh2 | ENSMUSG( | ENSMUSG( | 0.678443 | 6.08E-10 | YES | NO | NA |
| Smarcc1 | ENSMUSG( | ENSMUSG( | 0.130524 | 0.126215 | NO | NO | NA |
| Adap1 | ENSMUSG( | ENSMUSG( | 0.070767 | 0.378632 | NO | NO | NA |
| Strap | ENSMUSG( | ENSMUSG( | 0.352376 | 1.50E-06 | YES | NO | NA |
| Vdac2 | ENSMUSG( | ENSMUSG( | 0.2655 | 7.44E-06 | YES | NO | NA |
| Heca | ENSMUSG( | ENSMUSG( | -0.5026 | 5.09E-06 | YES | NO | NA |
| Nsun2 | ENSMUSG( | ENSMUSG( | 0.562086 | 1.31E-06 | YES | NO | NA |
| Nfkbiz | ENSMUSG( | ENSMUSG( | 0.086703 | 0.667184 | NO | NO | NA |
| Ap3d1 | ENSMUSG( | ENSMUSG( | -0.28272 | 1.28E-05 | YES | NO | NA |
| Coro1c | ENSMUSG( | ENSMUSG( | -0.04868 | 0.583984 | NO | NO | NA |
| Atp5c1 | ENSMUSG( | ENSMUSG( | 0.156745 | 0.001354 | YES | NO | NA |
| Elmo1 | ENSMUSG( | ENSMUSG( | -0.25318 | 5.87E-06 | YES | NO | NA |
| Srp72 | ENSMUSG( | ENSMUSG( | 0.061818 | 0.510726 | NO | NO | NA |
| Parp1 | ENSMUSG( | ENSMUSG( | 0.237279 | 0.002681 | YES | NO | NA |
| Uqcrc1 | ENSMUSG( | ENSMUSG( | 0.217471 | 0.000256 | YES | NO | NA |
| Uggt1 | ENSMUSG( | ENSMUSG( | -0.17299 | 0.001231 | YES | NO | NA |
| Tbx21 | ENSMUSG( | ENSMUSG( | -0.51606 | 1.19E-07 | YES | NO | NA |
| Traf3ip3 | ENSMUSG( | ENSMUSG( | -0.25846 | 9.99E-05 | YES | NO | NA |
| Rad23b | ENSMUSG( | ENSMUSG( | 0.181571 | 0.110143 | NO | NO | NA |
| Atl3 | ENSMUSG( | ENSMUSG( | 0.028625 | 0.772679 | NO | NO | NA |
| Ociad1 | ENSMUSG( | ENSMUSG( | -0.26079 | 6.20E-06 | YES | NO | NA |
| Sae1 | ENSMUSG( | ENSMUSG( | 0.329617 | 5.60E-05 | YES | NO | NA |
| Ctsc | ENSMUSG( | ENSMUSG( | -0.48598 | 1.86E-08 | YES | NO | NA |
| Taok3 | ENSMUSG( | ENSMUSG( | -0.17353 | 0.020635 | YES | NO | NA |
| Cdk2ap1 | ENSMUSG( | ENSMUSG( | 0.526261 | 1.67E-09 | YES | NO | NA |
| Ddx42 | ENSMUSG( | ENSMUSG( | 0.051648 | 0.508329 | NO | NO | NA |
| Trp53i11 | ENSMUSG( | ENSMUSG( | 0.411131 | 0.000196 | YES | NO | NA |
| Rbm33 | ENSMUSG( | ENSMUSG( | -0.1182 | 0.21546 | NO | NO | NA |
| Srsf10 | ENSMUSG( | ENSMUSG( | 0.099626 | 0.280492 | NO | NO | NA |
| Becn1 | ENSMUSG( | ENSMUSG( | -0.07804 | 0.118954 | NO | NO | NA |
| Dhx16 | ENSMUSG( | ENSMUSG( | 0.111346 | 0.05506 | NO | NO | NA |
| Noc2l | ENSMUSG( | ENSMUSG( | 0.288223 | 0.006167 | YES | NO | NA |
| Ghitm | ENSMUSG( | ENSMUSG( | -0.09397 | 0.112582 | NO | NO | NA |
| Kdm3b | ENSMUSG( | ENSMUSG( | -0.33529 | 1.12E-10 | YES | NO | NA |
| Lrrk1 | ENSMUSG( | ENSMUSG( | -0.27603 | 4.78E-05 | YES | NO | NA |
| Park7 | ENSMUSG( | ENSMUSG( | 0.163038 | 0.212591 | NO | NO | NA |
| Piezo1 | ENSMUSG( | ENSMUSG( | -0.26591 | 5.92E-05 | YES | NO | NA |

| | | | | | | | |
|---|---|---|---|---|---|---|---|
| Cpsf6 | ENSMUSG( | ENSMUSG( | 0.150304 | 0.004221 | YES | NO | NA |
| Crmp1 | ENSMUSG( | ENSMUSG( | -0.09315 | 0.244411 | NO | NO | NA |
| Gvin3 | ENSMUSG( | ENSMUSG( | -0.9463 | 1.04E-14 | YES | NO | NA |
| Cdc20 | ENSMUSG( | ENSMUSG( | 0.454463 | 1.72E-06 | YES | NO | NA |
| Taok2 | ENSMUSG( | ENSMUSG( | -0.23461 | 0.000119 | YES | NO | NA |
| Mthfd1l | ENSMUSG( | ENSMUSG( | 0.339603 | 1.44E-05 | YES | NO | NA |
| Tmc6 | ENSMUSG( | ENSMUSG( | -0.38337 | 0.000232 | YES | NO | NA |
| Cmip | ENSMUSG( | ENSMUSG( | -0.3319 | 9.02E-06 | YES | NO | NA |
| Cdk13 | ENSMUSG( | ENSMUSG( | -0.30749 | 3.26E-06 | YES | NO | NA |
| AI662270 | ENSMUSG( | ENSMUSG( | 0.166605 | 0.050861 | NO | NO | NA |
| Zgpat | ENSMUSG( | ENSMUSG( | -0.38441 | 1.49E-10 | YES | NO | NA |
| Pcnx3 | ENSMUSG( | ENSMUSG( | -0.30391 | 2.32E-06 | YES | NO | NA |
| Qars | ENSMUSG( | ENSMUSG( | 0.095135 | 0.215971 | NO | NO | NA |
| Cox8a | ENSMUSG( | ENSMUSG( | 0.146526 | 0.004993 | YES | NO | NA |
| H13 | ENSMUSG( | ENSMUSG( | -0.02473 | 0.717542 | NO | NO | NA |
| Tacc3 | ENSMUSG( | ENSMUSG( | 0.322533 | 0.001421 | YES | NO | NA |
| Pde2a | ENSMUSG( | ENSMUSG( | -1.8024 | 1.04E-09 | YES | YES | Downregulated |
| Bub3 | ENSMUSG( | ENSMUSG( | 0.159036 | 0.001752 | YES | NO | NA |
| Ifi47 | ENSMUSG( | ENSMUSG( | -0.38745 | 1.18E-07 | YES | NO | NA |
| Psmb1 | ENSMUSG( | ENSMUSG( | 0.064704 | 0.324184 | NO | NO | NA |
| Tyk2 | ENSMUSG( | ENSMUSG( | -0.45995 | 6.21E-14 | YES | NO | NA |
| Kmt2c | ENSMUSG( | ENSMUSG( | -0.62122 | 1.46E-12 | YES | NO | NA |
| Peak1 | ENSMUSG( | ENSMUSG( | -0.82137 | 1.09E-17 | YES | NO | NA |
| Lrrc59 | ENSMUSG( | ENSMUSG( | 0.312366 | 0.000517 | YES | NO | NA |
| Clic4 | ENSMUSG( | ENSMUSG( | 0.032258 | 0.676977 | NO | NO | NA |
| Gvin-ps6 | ENSMUSG( | ENSMUSG( | -0.95948 | 6.43E-17 | YES | NO | NA |
| Ube2d2a | ENSMUSG( | ENSMUSG( | -0.04927 | 0.445369 | NO | NO | NA |
| Vps13d | ENSMUSG( | ENSMUSG( | -0.6335 | 1.97E-21 | YES | NO | NA |
| Nme2 | ENSMUSG( | ENSMUSG( | 0.58479 | 1.73E-08 | YES | NO | NA |
| Git2 | ENSMUSG( | ENSMUSG( | -0.28213 | 0.000147 | YES | NO | NA |
| Efr3a | ENSMUSG( | ENSMUSG( | -0.13975 | 0.052054 | NO | NO | NA |
| Tbc1d2b | ENSMUSG( | ENSMUSG( | -0.29786 | 4.97E-06 | YES | NO | NA |
| Sqstm1 | ENSMUSG( | ENSMUSG( | -0.229 | 0.001106 | YES | NO | NA |
| Med14 | ENSMUSG( | ENSMUSG( | 0.024256 | 0.790376 | NO | NO | NA |
| Dnajb11 | ENSMUSG( | ENSMUSG( | -0.07417 | 0.366592 | NO | NO | NA |
| Mtdh | ENSMUSG( | ENSMUSG( | 0.198999 | 0.001334 | YES | NO | NA |
| Psmd1 | ENSMUSG( | ENSMUSG( | 0.350889 | 0.000381 | YES | NO | NA |
| Hyou1 | ENSMUSG( | ENSMUSG( | -0.06745 | 0.531711 | NO | NO | NA |
| Mier1 | ENSMUSG( | ENSMUSG( | -0.15422 | 0.054188 | NO | NO | NA |
| Nup62 | ENSMUSG( | ENSMUSG( | 0.260475 | 0.000546 | YES | NO | NA |
| Lsm4 | ENSMUSG( | ENSMUSG( | 0.339314 | 0.000243 | YES | NO | NA |
| Tcerg1 | ENSMUSG( | ENSMUSG( | 0.20448 | 0.004671 | YES | NO | NA |
| Ikbkb | ENSMUSG( | ENSMUSG( | -0.06864 | 0.420952 | NO | NO | NA |
| Gtpbp1 | ENSMUSG( | ENSMUSG( | -0.095 | 0.03133 | YES | NO | NA |
| Pi4ka | ENSMUSG( | ENSMUSG( | -0.24053 | 5.36E-09 | YES | NO | NA |
| Stat4 | ENSMUSG( | ENSMUSG( | -0.21679 | 0.000781 | YES | NO | NA |
| Nars | ENSMUSG( | ENSMUSG( | 0.254027 | 0.006617 | YES | NO | NA |

| | | | | | | | |
|---|---|---|---|---|---|---|---|
| Hint1 | ENSMUSG( | ENSMUSG( | 0.493794 | 2.64E-09 | YES | NO | NA |
| E2f4 | ENSMUSG( | ENSMUSG( | 0.133002 | 0.008372 | YES | NO | NA |
| Tecpr1 | ENSMUSG( | ENSMUSG( | -0.13139 | 0.252745 | NO | NO | NA |
| Spcs2 | ENSMUSG( | ENSMUSG( | -0.10259 | 0.260097 | NO | NO | NA |
| Fbl | ENSMUSG( | ENSMUSG( | 0.53816 | 0.000645 | YES | NO | NA |
| Tmem259 | ENSMUSG( | ENSMUSG( | 0.095371 | 0.271084 | NO | NO | NA |
| Rab11b | ENSMUSG( | ENSMUSG( | -0.20344 | 0.001354 | YES | NO | NA |
| Fam193b | ENSMUSG( | ENSMUSG( | -0.25005 | 1.63E-05 | YES | NO | NA |
| Plekhb2 | ENSMUSG( | ENSMUSG( | -0.37898 | 5.14E-06 | YES | NO | NA |
| Ddx24 | ENSMUSG( | ENSMUSG( | -0.12075 | 0.034186 | YES | NO | NA |
| Ik | ENSMUSG( | ENSMUSG( | 0.02536 | 0.789378 | NO | NO | NA |
| Cul3 | ENSMUSG( | ENSMUSG( | 0.047685 | 0.495773 | NO | NO | NA |
| Ncoa2 | ENSMUSG( | ENSMUSG( | -0.34381 | 4.21E-06 | YES | NO | NA |
| Tm6sf1 | ENSMUSG( | ENSMUSG( | -0.77054 | 7.82E-12 | YES | NO | NA |
| Mdh1 | ENSMUSG( | ENSMUSG( | 0.085913 | 0.149086 | NO | NO | NA |
| Rif1 | ENSMUSG( | ENSMUSG( | -0.13341 | 0.006081 | YES | NO | NA |
| Nudc | ENSMUSG( | ENSMUSG( | 0.237296 | 0.140653 | NO | NO | NA |
| Cit | ENSMUSG( | ENSMUSG( | -0.0426 | 0.606841 | NO | NO | NA |
| Slc38a10 | ENSMUSG( | ENSMUSG( | -0.16401 | 7.26E-08 | YES | NO | NA |
| Ubash3b | ENSMUSG( | ENSMUSG( | -0.17259 | 0.111628 | NO | NO | NA |
| Cnot3 | ENSMUSG( | ENSMUSG( | -0.13234 | 0.031464 | YES | NO | NA |
| Lypla2 | ENSMUSG( | ENSMUSG( | -0.21075 | 0.001932 | YES | NO | NA |
| Cdk11b | ENSMUSG( | ENSMUSG( | -0.15903 | 0.209547 | NO | NO | NA |
| Gdi1 | ENSMUSG( | ENSMUSG( | -0.38823 | 7.28E-11 | YES | NO | NA |
| Dlst | ENSMUSG( | ENSMUSG( | 0.062519 | 0.24151 | NO | NO | NA |
| Epn1 | ENSMUSG( | ENSMUSG( | 0.034099 | 0.667108 | NO | NO | NA |
| Mlf2 | ENSMUSG( | ENSMUSG( | 0.095457 | 0.108512 | NO | NO | NA |
| Setd2 | ENSMUSG( | ENSMUSG( | -0.28939 | 4.93E-05 | YES | NO | NA |
| Ly9 | ENSMUSG( | ENSMUSG( | -0.34684 | 1.71E-11 | YES | NO | NA |
| Rbm38 | ENSMUSG( | ENSMUSG( | -0.12939 | 0.054794 | NO | NO | NA |
| Bsg | ENSMUSG( | ENSMUSG( | 0.110879 | 0.147915 | NO | NO | NA |
| Cd69 | ENSMUSG( | ENSMUSG( | 0.406402 | 0.268146 | NO | NO | NA |
| Adss | ENSMUSG( | ENSMUSG( | 0.178866 | 0.040678 | YES | NO | NA |
| Virma | ENSMUSG( | ENSMUSG( | -0.28376 | 3.28E-07 | YES | NO | NA |
| Hells | ENSMUSG( | ENSMUSG( | 0.255211 | 0.006963 | YES | NO | NA |
| Inpp5d | ENSMUSG( | ENSMUSG( | -0.41781 | 5.29E-09 | YES | NO | NA |
| Zswim8 | ENSMUSG( | ENSMUSG( | -0.47139 | 4.73E-08 | YES | NO | NA |
| Selenot | ENSMUSG( | ENSMUSG( | -0.11703 | 0.013655 | YES | NO | NA |
| Jak2 | ENSMUSG( | ENSMUSG( | -0.19269 | 0.035788 | YES | NO | NA |
| Alkbh5 | ENSMUSG( | ENSMUSG( | -0.21836 | 6.65E-08 | YES | NO | NA |
| Ppm1b | ENSMUSG( | ENSMUSG( | -0.43328 | 1.96E-09 | YES | NO | NA |
| Golga4 | ENSMUSG( | ENSMUSG( | -0.36572 | 2.10E-07 | YES | NO | NA |
| Ywhag | ENSMUSG( | ENSMUSG( | -0.32131 | 6.04E-15 | YES | NO | NA |
| Creb1 | ENSMUSG( | ENSMUSG( | -0.44503 | 2.82E-10 | YES | NO | NA |
| Fam78a | ENSMUSG( | ENSMUSG( | -0.291 | 0.000156 | YES | NO | NA |
| Otud5 | ENSMUSG( | ENSMUSG( | -0.3485 | 8.06E-08 | YES | NO | NA |
| Copb1 | ENSMUSG( | ENSMUSG( | -0.06804 | 0.297659 | NO | NO | NA |

| | | | | | | | |
|---|---|---|---|---|---|---|---|
| Ccdc50 | ENSMUSG( | ENSMUSG( | -0.16134 | 0.000128 | YES | NO | NA |
| Tma7 | ENSMUSG( | ENSMUSG( | -0.03877 | 0.487915 | NO | NO | NA |
| Atp5g2 | ENSMUSG( | ENSMUSG( | 0.321827 | 2.95E-10 | YES | NO | NA |
| Usp4 | ENSMUSG( | ENSMUSG( | 0.008087 | 0.86804 | NO | NO | NA |
| Cox6a1 | ENSMUSG( | ENSMUSG( | 0.178802 | 0.004264 | YES | NO | NA |
| Tmed10 | ENSMUSG( | ENSMUSG( | 0.109603 | 0.093668 | NO | NO | NA |
| Aqr | ENSMUSG( | ENSMUSG( | -0.15455 | 0.005921 | YES | NO | NA |
| Dnajc8 | ENSMUSG( | ENSMUSG( | -0.10754 | 0.359969 | NO | NO | NA |
| Irak1 | ENSMUSG( | ENSMUSG( | 0.048492 | 0.601687 | NO | NO | NA |
| Iqgap2 | ENSMUSG( | ENSMUSG( | -0.22563 | 0.001609 | YES | NO | NA |
| Wac | ENSMUSG( | ENSMUSG( | -0.20969 | 0.000528 | YES | NO | NA |
| Acadl | ENSMUSG( | ENSMUSG( | 0.126654 | 0.10945 | NO | NO | NA |
| Nasp | ENSMUSG( | ENSMUSG( | 0.480249 | 7.43E-08 | YES | NO | NA |
| Sh3glb1 | ENSMUSG( | ENSMUSG( | -0.27084 | 0.00106 | YES | NO | NA |
| Gm17092 | ENSMUSG( | ENSMUSG( | -0.03285 | 0.945161 | NO | NO | NA |
| Trir | ENSMUSG( | ENSMUSG( | -0.05286 | 0.502435 | NO | NO | NA |
| Akr1b3 | ENSMUSG( | ENSMUSG( | 0.055113 | 0.126454 | NO | NO | NA |
| Trpc4ap | ENSMUSG( | ENSMUSG( | -0.16231 | 0.024441 | YES | NO | NA |
| Mrfap1 | ENSMUSG( | ENSMUSG( | -0.15981 | 0.059498 | NO | NO | NA |
| Mapk14 | ENSMUSG( | ENSMUSG( | -0.13144 | 0.019101 | YES | NO | NA |
| Cd28 | ENSMUSG( | ENSMUSG( | -0.0579 | 0.754398 | NO | NO | NA |
| Pde3b | ENSMUSG( | ENSMUSG( | -0.22481 | 0.00115 | YES | NO | NA |
| Psmd13 | ENSMUSG( | ENSMUSG( | 0.300088 | 4.52E-05 | YES | NO | NA |
| Cyld | ENSMUSG( | ENSMUSG( | -0.2677 | 0.000484 | YES | NO | NA |
| Bcl2l11 | ENSMUSG( | ENSMUSG( | -0.25169 | 0.001177 | YES | NO | NA |
| Rnf40 | ENSMUSG( | ENSMUSG( | 0.023141 | 0.800013 | NO | NO | NA |
| Kdm5a | ENSMUSG( | ENSMUSG( | -0.25547 | 0.000185 | YES | NO | NA |
| Cdc42se2 | ENSMUSG( | ENSMUSG( | -0.19755 | 0.011448 | YES | NO | NA |
| Kif13b | ENSMUSG( | ENSMUSG( | -0.38201 | 6.13E-05 | YES | NO | NA |
| Klrk1 | ENSMUSG( | ENSMUSG( | 0.622503 | 6.46E-05 | YES | NO | NA |
| Arl5a | ENSMUSG( | ENSMUSG( | -0.42693 | 1.89E-07 | YES | NO | NA |
| Taldo1 | ENSMUSG( | ENSMUSG( | -0.05855 | 0.341079 | NO | NO | NA |
| Srpr | ENSMUSG( | ENSMUSG( | -0.06712 | 0.271018 | NO | NO | NA |
| Arfgef1 | ENSMUSG( | ENSMUSG( | -0.25895 | 0.001254 | YES | NO | NA |
| Reep5 | ENSMUSG( | ENSMUSG( | -0.10283 | 0.220077 | NO | NO | NA |
| Rnf123 | ENSMUSG( | ENSMUSG( | -0.08032 | 0.368792 | NO | NO | NA |
| Pld3 | ENSMUSG( | ENSMUSG( | -0.43611 | 4.60E-08 | YES | NO | NA |
| Zbed6 | ENSMUSG( | ENSMUSG( | -0.34944 | 0.000481 | YES | NO | NA |
| Tor1aip1 | ENSMUSG( | ENSMUSG( | -0.31671 | 8.60E-07 | YES | NO | NA |
| Kcnn4 | ENSMUSG( | ENSMUSG( | 0.103268 | 0.156638 | NO | NO | NA |
| Nup50 | ENSMUSG( | ENSMUSG( | -0.16244 | 0.0044 | YES | NO | NA |
| Drap1 | ENSMUSG( | ENSMUSG( | 0.264134 | 0.00524 | YES | NO | NA |
| C2cd3 | ENSMUSG( | ENSMUSG( | -0.20149 | 0.000279 | YES | NO | NA |
| Atic | ENSMUSG( | ENSMUSG( | 0.496761 | 2.86E-05 | YES | NO | NA |
| Tlk1 | ENSMUSG( | ENSMUSG( | -0.05971 | 0.529845 | NO | NO | NA |
| Msl1 | ENSMUSG( | ENSMUSG( | -0.21159 | 0.000729 | YES | NO | NA |
| Mta1 | ENSMUSG( | ENSMUSG( | 0.263861 | 0.000171 | YES | NO | NA |

| | | | | | | | |
|---|---|---|---|---|---|---|---|
| Mapkapk2 | ENSMUSG( | ENSMUSG( | 0.364305 | 1.25E-05 | YES | NO | NA |
| Cdk4 | ENSMUSG( | ENSMUSG( | 0.380343 | 4.51E-05 | YES | NO | NA |
| Zfp871 | ENSMUSG( | ENSMUSG( | -0.54202 | 1.68E-08 | YES | NO | NA |
| Rnf114 | ENSMUSG( | ENSMUSG( | -0.4378 | 2.84E-08 | YES | NO | NA |
| Pbxip1 | ENSMUSG( | ENSMUSG( | -0.58409 | 3.71E-09 | YES | NO | NA |
| St3gal1 | ENSMUSG( | ENSMUSG( | -0.02476 | 0.838501 | NO | NO | NA |
| Sh3bp1 | ENSMUSG( | ENSMUSG( | -0.14765 | 0.000653 | YES | NO | NA |
| Dusp5 | ENSMUSG( | ENSMUSG( | -0.02317 | 0.897257 | NO | NO | NA |
| Slamf7 | ENSMUSG( | ENSMUSG( | -0.51587 | 2.10E-09 | YES | NO | NA |
| Rapgef6 | ENSMUSG( | ENSMUSG( | -0.1616 | 0.065891 | NO | NO | NA |
| Ncaph | ENSMUSG( | ENSMUSG( | 0.252641 | 2.38E-05 | YES | NO | NA |
| Nabp1 | ENSMUSG( | ENSMUSG( | -0.40982 | 1.00E-08 | YES | NO | NA |
| Cbx4 | ENSMUSG( | ENSMUSG( | 0.034964 | 0.812558 | NO | NO | NA |
| Foxm1 | ENSMUSG( | ENSMUSG( | 0.040853 | 0.618911 | NO | NO | NA |
| Bin1 | ENSMUSG( | ENSMUSG( | -0.2311 | 1.17E-06 | YES | NO | NA |
| Chaf1a | ENSMUSG( | ENSMUSG( | 0.205406 | 0.094921 | NO | NO | NA |
| Kdm2a | ENSMUSG( | ENSMUSG( | -0.30456 | 2.23E-06 | YES | NO | NA |
| Stat6 | ENSMUSG( | ENSMUSG( | -0.29028 | 4.44E-09 | YES | NO | NA |
| St13 | ENSMUSG( | ENSMUSG( | -0.06116 | 0.56132 | NO | NO | NA |
| Zc3h11a | ENSMUSG( | ENSMUSG( | -0.29775 | 3.92E-05 | YES | NO | NA |
| Pabpn1 | ENSMUSG( | ENSMUSG( | 0.096226 | 0.25241 | NO | NO | NA |
| Nup205 | ENSMUSG( | ENSMUSG( | 0.231631 | 1.54E-05 | YES | NO | NA |
| Rbl1 | ENSMUSG( | ENSMUSG( | -0.12745 | 0.064898 | NO | NO | NA |
| Bms1 | ENSMUSG( | ENSMUSG( | -0.0091 | 0.933709 | NO | NO | NA |
| Psmb10 | ENSMUSG( | ENSMUSG( | -0.12534 | 0.006967 | YES | NO | NA |
| Tcea1 | ENSMUSG( | ENSMUSG( | 0.070272 | 0.43299 | NO | NO | NA |
| Cd2bp2 | ENSMUSG( | ENSMUSG( | 0.114775 | 0.123354 | NO | NO | NA |
| Mlxip | ENSMUSG( | ENSMUSG( | -0.25347 | 0.000606 | YES | NO | NA |
| Sp1 | ENSMUSG( | ENSMUSG( | -0.13797 | 0.004534 | YES | NO | NA |
| Tbc1d10b | ENSMUSG( | ENSMUSG( | -0.24216 | 9.85E-07 | YES | NO | NA |
| Hprt | ENSMUSG( | ENSMUSG( | 0.193638 | 0.002051 | YES | NO | NA |
| Kidins220 | ENSMUSG( | ENSMUSG( | -0.2798 | 0.000236 | YES | NO | NA |
| Ptpn13 | ENSMUSG( | ENSMUSG( | -0.41411 | 1.43E-05 | YES | NO | NA |
| Csrp1 | ENSMUSG( | ENSMUSG( | 0.079046 | 0.253891 | NO | NO | NA |
| Pdpk1 | ENSMUSG( | ENSMUSG( | -0.20287 | 0.000501 | YES | NO | NA |
| Lims1 | ENSMUSG( | ENSMUSG( | -0.15511 | 0.088028 | NO | NO | NA |
| Tsn | ENSMUSG( | ENSMUSG( | 0.168273 | 0.003553 | YES | NO | NA |
| Mapk1ip1l | ENSMUSG( | ENSMUSG( | -0.14628 | 0.001408 | YES | NO | NA |
| Ube2g2 | ENSMUSG( | ENSMUSG( | -0.23212 | 0.003271 | YES | NO | NA |
| Ifi213 | ENSMUSG( | ENSMUSG( | -1.31188 | 2.37E-18 | YES | YES | Downregulated |
| Trim25 | ENSMUSG( | ENSMUSG( | -0.60949 | 1.34E-22 | YES | NO | NA |
| Zc3h13 | ENSMUSG( | ENSMUSG( | -0.1352 | 0.266773 | NO | NO | NA |
| Ccdc82 | ENSMUSG( | ENSMUSG( | -0.24213 | 0.000538 | YES | NO | NA |
| Crkl | ENSMUSG( | ENSMUSG( | -0.27883 | 1.65E-11 | YES | NO | NA |
| Parvg | ENSMUSG( | ENSMUSG( | -0.08938 | 0.061761 | NO | NO | NA |
| Cdk17 | ENSMUSG( | ENSMUSG( | -0.28866 | 0.000273 | YES | NO | NA |
| Psmb3 | ENSMUSG( | ENSMUSG( | 0.110173 | 0.04995 | YES | NO | NA |

| | | | | | | | |
|---|---|---|---|---|---|---|---|
| Htt | ENSMUSG( | ENSMUSG( | -0.27611 | 1.84E-12 | YES | NO | NA |
| Sun1 | ENSMUSG( | ENSMUSG( | -0.01732 | 0.862668 | NO | NO | NA |
| Phrf1 | ENSMUSG( | ENSMUSG( | -0.18222 | 0.000506 | YES | NO | NA |
| Smarce1 | ENSMUSG( | ENSMUSG( | 0.106385 | 0.502358 | NO | NO | NA |
| Rsrc2 | ENSMUSG( | ENSMUSG( | -0.17441 | 0.158086 | NO | NO | NA |
| Usf2 | ENSMUSG( | ENSMUSG( | 0.182954 | 0.047463 | YES | NO | NA |
| Pole | ENSMUSG( | ENSMUSG( | 0.161237 | 0.005494 | YES | NO | NA |
| Dpp8 | ENSMUSG( | ENSMUSG( | -0.0271 | 0.721838 | NO | NO | NA |
| Sec11c | ENSMUSG( | ENSMUSG( | -0.14775 | 0.015009 | YES | NO | NA |
| B4galt1 | ENSMUSG( | ENSMUSG( | -0.18927 | 0.029547 | YES | NO | NA |
| Scaf1 | ENSMUSG( | ENSMUSG( | -0.091 | 0.001105 | YES | NO | NA |
| Gart | ENSMUSG( | ENSMUSG( | 0.713255 | 2.57E-09 | YES | NO | NA |
| Padi2 | ENSMUSG( | ENSMUSG( | -0.53031 | 2.82E-08 | YES | NO | NA |
| Idh3a | ENSMUSG( | ENSMUSG( | 0.099253 | 0.205816 | NO | NO | NA |
| Ap1b1 | ENSMUSG( | ENSMUSG( | -0.24944 | 2.94E-05 | YES | NO | NA |
| Il16 | ENSMUSG( | ENSMUSG( | -0.65198 | 5.79E-12 | YES | NO | NA |
| Mllt3 | ENSMUSG( | ENSMUSG( | -0.09889 | 0.30141 | NO | NO | NA |
| Cpsf1 | ENSMUSG( | ENSMUSG( | 0.173506 | 0.002323 | YES | NO | NA |
| Atp5h | ENSMUSG( | ENSMUSG( | -0.10455 | 0.042735 | YES | NO | NA |
| Cerkl | ENSMUSG( | ENSMUSG( | -0.34557 | 0.002752 | YES | NO | NA |
| Cox7a2l | ENSMUSG( | ENSMUSG( | -0.02108 | 0.827094 | NO | NO | NA |
| Kif20a | ENSMUSG( | ENSMUSG( | 0.413882 | 7.75E-06 | YES | NO | NA |
| Zzef1 | ENSMUSG( | ENSMUSG( | -0.27341 | 3.31E-05 | YES | NO | NA |
| Usp9x | ENSMUSG( | ENSMUSG( | -0.10624 | 0.072091 | NO | NO | NA |
| Brd1 | ENSMUSG( | ENSMUSG( | -0.17273 | 0.006026 | YES | NO | NA |
| Atxn7l3 | ENSMUSG( | ENSMUSG( | -0.02805 | 0.620842 | NO | NO | NA |
| Ube2z | ENSMUSG( | ENSMUSG( | 0.033656 | 0.617169 | NO | NO | NA |
| Inpp5b | ENSMUSG( | ENSMUSG( | -0.21817 | 0.000161 | YES | NO | NA |
| Slc28a2 | ENSMUSG( | ENSMUSG( | -0.53969 | 3.05E-11 | YES | NO | NA |
| Hivep2 | ENSMUSG( | ENSMUSG( | -0.67169 | 3.67E-09 | YES | NO | NA |
| Fbxl14 | ENSMUSG( | ENSMUSG( | -0.12738 | 0.007751 | YES | NO | NA |
| Atp5d | ENSMUSG( | ENSMUSG( | 0.142709 | 0.014477 | YES | NO | NA |
| Usp47 | ENSMUSG( | ENSMUSG( | -0.14662 | 0.001397 | YES | NO | NA |
| Ccr5 | ENSMUSG( | ENSMUSG( | -0.28821 | 0.013072 | YES | NO | NA |
| Rab1a | ENSMUSG( | ENSMUSG( | -0.09201 | 0.437356 | NO | NO | NA |
| Ptpn6 | ENSMUSG( | ENSMUSG( | -0.16355 | 0.087711 | NO | NO | NA |
| 1700017B( | ENSMUSG( | ENSMUSG( | 0.357345 | 0.000391 | YES | NO | NA |
| Pip4k2a | ENSMUSG( | ENSMUSG( | -0.04567 | 0.624649 | NO | NO | NA |
| Rnf20 | ENSMUSG( | ENSMUSG( | -0.22109 | 0.000696 | YES | NO | NA |
| Atf6 | ENSMUSG( | ENSMUSG( | 0.170326 | 0.289258 | NO | NO | NA |
| Txndc5 | ENSMUSG( | ENSMUSG( | -0.13597 | 0.022765 | YES | NO | NA |
| Lrch4 | ENSMUSG( | ENSMUSG( | -0.18984 | 0.000203 | YES | NO | NA |
| Rassf2 | ENSMUSG( | ENSMUSG( | -0.48393 | 1.07E-05 | YES | NO | NA |
| Msl2 | ENSMUSG( | ENSMUSG( | -0.37391 | 1.84E-11 | YES | NO | NA |
| Tug1 | ENSMUSG( | ENSMUSG( | -0.29412 | 0.000143 | YES | NO | NA |
| Smg5 | ENSMUSG( | ENSMUSG( | 0.000495 | 0.993171 | NO | NO | NA |
| Chchd2 | ENSMUSG( | ENSMUSG( | 0.095157 | 0.067101 | NO | NO | NA |

| | | | | | | | |
|---|---|---|---|---|---|---|---|
| Eif3i | ENSMUSG( | ENSMUSG( | 0.416728 | 4.54E-14 | YES | NO | NA |
| Ptpn4 | ENSMUSG( | ENSMUSG( | -1.13523 | 3.88E-14 | YES | YES | Downregulated |
| Poldip3 | ENSMUSG( | ENSMUSG( | -0.18575 | 2.21E-07 | YES | NO | NA |
| Pom121 | ENSMUSG( | ENSMUSG( | 0.037602 | 0.639774 | NO | NO | NA |
| Zfpm1 | ENSMUSG( | ENSMUSG( | 0.00557 | 0.958772 | NO | NO | NA |
| Bod1l | ENSMUSG( | ENSMUSG( | -0.20973 | 0.012065 | YES | NO | NA |
| Eif3j1 | ENSMUSG( | ENSMUSG( | 0.203451 | 0.154131 | NO | NO | NA |
| Ccng1 | ENSMUSG( | ENSMUSG( | -0.17768 | 0.097673 | NO | NO | NA |
| Kifc1 | ENSMUSG( | ENSMUSG( | 0.325114 | 2.54E-05 | YES | NO | NA |
| Plcl2 | ENSMUSG( | ENSMUSG( | -0.26248 | 5.92E-05 | YES | NO | NA |
| Uba7 | ENSMUSG( | ENSMUSG( | -0.38879 | 1.83E-05 | YES | NO | NA |
| Psma4 | ENSMUSG( | ENSMUSG( | 0.157275 | 0.018344 | YES | NO | NA |
| Aup1 | ENSMUSG( | ENSMUSG( | -0.0173 | 0.789871 | NO | NO | NA |
| Dot1l | ENSMUSG( | ENSMUSG( | -0.03834 | 0.481807 | NO | NO | NA |
| Ilk | ENSMUSG( | ENSMUSG( | -0.03836 | 0.547317 | NO | NO | NA |
| Pigt | ENSMUSG( | ENSMUSG( | 0.103382 | 0.157527 | NO | NO | NA |
| Malt1 | ENSMUSG( | ENSMUSG( | -0.36929 | 5.43E-08 | YES | NO | NA |
| Csnk2b | ENSMUSG( | ENSMUSG( | 0.032712 | 0.534067 | NO | NO | NA |
| Kmt5a | ENSMUSG( | ENSMUSG( | 0.088004 | 0.222199 | NO | NO | NA |
| Spcs3 | ENSMUSG( | ENSMUSG( | 0.149344 | 0.002126 | YES | NO | NA |
| Foxo1 | ENSMUSG( | ENSMUSG( | -0.69535 | 1.76E-12 | YES | NO | NA |
| Kif15 | ENSMUSG( | ENSMUSG( | 0.271634 | 0.006483 | YES | NO | NA |
| Frmd8 | ENSMUSG( | ENSMUSG( | 0.017982 | 0.71797 | NO | NO | NA |
| Sap18 | ENSMUSG( | ENSMUSG( | 0.123244 | 0.12163 | NO | NO | NA |
| Cdkn1b | ENSMUSG( | ENSMUSG( | -0.68326 | 3.94E-11 | YES | NO | NA |
| Ddx58 | ENSMUSG( | ENSMUSG( | -0.73211 | 2.57E-12 | YES | NO | NA |
| Tuba4a | ENSMUSG( | ENSMUSG( | 0.550415 | 3.32E-07 | YES | NO | NA |
| Abracl | ENSMUSG( | ENSMUSG( | -0.18097 | 0.015213 | YES | NO | NA |
| Impdh1 | ENSMUSG( | ENSMUSG( | 0.089361 | 0.272668 | NO | NO | NA |
| H2ac23 | ENSMUSG( | ENSMUSG( | 0.480445 | 5.80E-08 | YES | NO | NA |
| Lrba | ENSMUSG( | ENSMUSG( | -0.1957 | 0.004734 | YES | NO | NA |
| Kif23 | ENSMUSG( | ENSMUSG( | 0.139051 | 0.065696 | NO | NO | NA |
| Birc5 | ENSMUSG( | ENSMUSG( | 0.302526 | 0.000251 | YES | NO | NA |
| Atp13a1 | ENSMUSG( | ENSMUSG( | 0.103199 | 0.094261 | NO | NO | NA |
| Pip5k1c | ENSMUSG( | ENSMUSG( | -0.05346 | 0.555025 | NO | NO | NA |
| Ss18 | ENSMUSG( | ENSMUSG( | -0.19012 | 0.000328 | YES | NO | NA |
| Pde7a | ENSMUSG( | ENSMUSG( | -0.5025 | 3.52E-09 | YES | NO | NA |
| Smg7 | ENSMUSG( | ENSMUSG( | -0.37136 | 3.42E-11 | YES | NO | NA |
| Dap | ENSMUSG( | ENSMUSG( | -0.0578 | 0.645714 | NO | NO | NA |
| Gng2 | ENSMUSG( | ENSMUSG( | -0.26871 | 0.001351 | YES | NO | NA |
| Cmtr1 | ENSMUSG( | ENSMUSG( | -0.27093 | 3.80E-12 | YES | NO | NA |
| Mtmr1 | ENSMUSG( | ENSMUSG( | 0.047302 | 0.66539 | NO | NO | NA |
| Ube2q1 | ENSMUSG( | ENSMUSG( | -0.19332 | 0.001128 | YES | NO | NA |
| Prkd2 | ENSMUSG( | ENSMUSG( | -0.42246 | 1.29E-10 | YES | NO | NA |
| Fkbp4 | ENSMUSG( | ENSMUSG( | -0.20063 | 0.089518 | NO | NO | NA |
| Nek2 | ENSMUSG( | ENSMUSG( | 0.554693 | 3.96E-09 | YES | NO | NA |
| Prpf19 | ENSMUSG( | ENSMUSG( | 0.388082 | 3.44E-06 | YES | NO | NA |

| | | | | | | | |
|---|---|---|---|---|---|---|---|
| Hectd4 | ENSMUSG( | ENSMUSG( | -0.46337 | 2.09E-12 | YES | NO | NA |
| Cggbp1 | ENSMUSG( | ENSMUSG( | -0.29365 | 1.89E-08 | YES | NO | NA |
| Nde1 | ENSMUSG( | ENSMUSG( | -0.03877 | 0.468284 | NO | NO | NA |
| Gimap5 | ENSMUSG( | ENSMUSG( | -0.26322 | 0.000391 | YES | NO | NA |
| Thoc2 | ENSMUSG( | ENSMUSG( | -0.27793 | 4.47E-05 | YES | NO | NA |
| Taf1 | ENSMUSG( | ENSMUSG( | -0.01816 | 0.868358 | NO | NO | NA |
| U2surp | ENSMUSG( | ENSMUSG( | 0.043762 | 0.62337 | NO | NO | NA |
| Bag1 | ENSMUSG( | ENSMUSG( | 0.074199 | 0.249622 | NO | NO | NA |
| Rnf145 | ENSMUSG( | ENSMUSG( | -0.07466 | 0.400316 | NO | NO | NA |
| Szt2 | ENSMUSG( | ENSMUSG( | -0.2306 | 0.002239 | YES | NO | NA |
| Eif3m | ENSMUSG( | ENSMUSG( | 0.184672 | 0.004781 | YES | NO | NA |
| Kansl1 | ENSMUSG( | ENSMUSG( | -0.36269 | 1.41E-05 | YES | NO | NA |
| Cmtm6 | ENSMUSG( | ENSMUSG( | -0.14061 | 0.022077 | YES | NO | NA |
| Cycs | ENSMUSG( | ENSMUSG( | 0.300534 | 0.001528 | YES | NO | NA |
| 2510039O | ENSMUSG( | ENSMUSG( | -0.50228 | 4.45E-42 | YES | NO | NA |
| Arf5 | ENSMUSG( | ENSMUSG( | 0.009986 | 0.885674 | NO | NO | NA |
| Immt | ENSMUSG( | ENSMUSG( | 0.387959 | 1.25E-07 | YES | NO | NA |
| Phip | ENSMUSG( | ENSMUSG( | -0.34691 | 1.26E-05 | YES | NO | NA |
| Dhx38 | ENSMUSG( | ENSMUSG( | -0.03856 | 0.68012 | NO | NO | NA |
| Bcl9l | ENSMUSG( | ENSMUSG( | -0.62688 | 3.83E-08 | YES | NO | NA |
| Synrg | ENSMUSG( | ENSMUSG( | -0.23898 | 0.000825 | YES | NO | NA |
| Me2 | ENSMUSG( | ENSMUSG( | 0.189304 | 0.006301 | YES | NO | NA |
| Ndufs2 | ENSMUSG( | ENSMUSG( | 0.251327 | 0.000112 | YES | NO | NA |
| Cpsf7 | ENSMUSG( | ENSMUSG( | -0.13023 | 0.016984 | YES | NO | NA |
| Usp5 | ENSMUSG( | ENSMUSG( | 0.286869 | 9.85E-05 | YES | NO | NA |
| Psma7 | ENSMUSG( | ENSMUSG( | 0.287294 | 5.24E-05 | YES | NO | NA |
| Rtcb | ENSMUSG( | ENSMUSG( | 0.225085 | 0.005417 | YES | NO | NA |
| Pygb | ENSMUSG( | ENSMUSG( | -0.23605 | 7.68E-08 | YES | NO | NA |
| Dnajc3 | ENSMUSG( | ENSMUSG( | -0.09793 | 0.013788 | YES | NO | NA |
| Myo18a | ENSMUSG( | ENSMUSG( | -0.52337 | 8.72E-11 | YES | NO | NA |
| Nras | ENSMUSG( | ENSMUSG( | 0.026127 | 0.764031 | NO | NO | NA |
| Kif1c | ENSMUSG( | ENSMUSG( | -0.02361 | 0.729543 | NO | NO | NA |
| Nptn | ENSMUSG( | ENSMUSG( | 0.016394 | 0.908629 | NO | NO | NA |
| Nans | ENSMUSG( | ENSMUSG( | 0.082296 | 0.411959 | NO | NO | NA |
| Syt11 | ENSMUSG( | ENSMUSG( | -0.37691 | 2.69E-05 | YES | NO | NA |
| Psat1 | ENSMUSG( | ENSMUSG( | 0.379684 | 8.08E-05 | YES | NO | NA |
| Gmip | ENSMUSG( | ENSMUSG( | -0.14476 | 0.011187 | YES | NO | NA |
| Timeless | ENSMUSG( | ENSMUSG( | 0.322019 | 0.000116 | YES | NO | NA |
| Cers2 | ENSMUSG( | ENSMUSG( | -0.12351 | 0.080192 | NO | NO | NA |
| Ankhd1 | ENSMUSG( | ENSMUSG( | -0.43439 | 3.36E-13 | YES | NO | NA |
| Pdia4 | ENSMUSG( | ENSMUSG( | -0.10913 | 0.354958 | NO | NO | NA |
| Ube2n | ENSMUSG( | ENSMUSG( | 0.105008 | 0.10192 | NO | NO | NA |
| Mprip | ENSMUSG( | ENSMUSG( | -0.11768 | 0.162247 | NO | NO | NA |
| Traf7 | ENSMUSG( | ENSMUSG( | 0.096448 | 0.199322 | NO | NO | NA |
| Pcnp | ENSMUSG( | ENSMUSG( | -0.15621 | 0.112702 | NO | NO | NA |
| Prmt1 | ENSMUSG( | ENSMUSG( | 0.546316 | 1.26E-07 | YES | NO | NA |
| Espl1 | ENSMUSG( | ENSMUSG( | 0.07552 | 0.357929 | NO | NO | NA |

| | | | | | | | |
|---|---|---|---|---|---|---|---|
| Eif3k | ENSMUSG( | ENSMUSG( | 0.111409 | 0.055482 | NO | NO | NA |
| Rnaset2b | ENSMUSG( | ENSMUSG( | -0.45878 | 7.28E-11 | YES | NO | NA |
| Rexo1 | ENSMUSG( | ENSMUSG( | -0.06925 | 0.304777 | NO | NO | NA |
| Ankrd12 | ENSMUSG( | ENSMUSG( | -0.40448 | 1.43E-05 | YES | NO | NA |
| Ralbp1 | ENSMUSG( | ENSMUSG( | -0.14184 | 0.002707 | YES | NO | NA |
| Atp2b1 | ENSMUSG( | ENSMUSG( | -0.50172 | 2.38E-13 | YES | NO | NA |
| Yme1l1 | ENSMUSG( | ENSMUSG( | -0.08601 | 0.093115 | NO | NO | NA |
| Chd1 | ENSMUSG( | ENSMUSG( | -0.12513 | 0.173376 | NO | NO | NA |
| Dpp3 | ENSMUSG( | ENSMUSG( | -0.02741 | 0.576568 | NO | NO | NA |
| Vps54 | ENSMUSG( | ENSMUSG( | -0.18805 | 0.126447 | NO | NO | NA |
| Pcf11 | ENSMUSG( | ENSMUSG( | -0.23986 | 0.001163 | YES | NO | NA |
| Spag5 | ENSMUSG( | ENSMUSG( | 0.246016 | 0.000355 | YES | NO | NA |
| Uck2 | ENSMUSG( | ENSMUSG( | 0.450152 | 1.52E-06 | YES | NO | NA |
| Syne1 | ENSMUSG( | ENSMUSG( | -0.71228 | 1.04E-13 | YES | NO | NA |
| Trak1 | ENSMUSG( | ENSMUSG( | -0.09451 | 0.247899 | NO | NO | NA |
| Cdc5l | ENSMUSG( | ENSMUSG( | -0.06237 | 0.460412 | NO | NO | NA |
| Tmc8 | ENSMUSG( | ENSMUSG( | -0.71532 | 1.06E-12 | YES | NO | NA |
| Dnajb1 | ENSMUSG( | ENSMUSG( | -0.58896 | 5.36E-09 | YES | NO | NA |
| Parp4 | ENSMUSG( | ENSMUSG( | -0.48397 | 7.78E-10 | YES | NO | NA |
| Kif22 | ENSMUSG( | ENSMUSG( | 0.221272 | 0.012201 | YES | NO | NA |
| Pcif1 | ENSMUSG( | ENSMUSG( | -0.03233 | 0.611373 | NO | NO | NA |
| Zdhhc20 | ENSMUSG( | ENSMUSG( | -0.26961 | 0.008308 | YES | NO | NA |
| Skap1 | ENSMUSG( | ENSMUSG( | -0.15755 | 0.001691 | YES | NO | NA |
| Azi2 | ENSMUSG( | ENSMUSG( | -0.31807 | 1.17E-05 | YES | NO | NA |
| Wbp11 | ENSMUSG( | ENSMUSG( | 0.07228 | 0.203292 | NO | NO | NA |
| N4bp1 | ENSMUSG( | ENSMUSG( | -0.41155 | 5.08E-08 | YES | NO | NA |
| Plk1 | ENSMUSG( | ENSMUSG( | 0.184806 | 0.056553 | NO | NO | NA |
| Ubp1 | ENSMUSG( | ENSMUSG( | -0.1625 | 0.007102 | YES | NO | NA |
| Stim1 | ENSMUSG( | ENSMUSG( | -0.4383 | 2.13E-09 | YES | NO | NA |
| Gon4l | ENSMUSG( | ENSMUSG( | -0.23542 | 7.88E-06 | YES | NO | NA |
| Dnajc13 | ENSMUSG( | ENSMUSG( | -0.07407 | 0.384546 | NO | NO | NA |
| Eif4enif1 | ENSMUSG( | ENSMUSG( | -0.07533 | 0.211661 | NO | NO | NA |
| Cc2d1b | ENSMUSG( | ENSMUSG( | 0.191268 | 0.002561 | YES | NO | NA |
| Itpr1 | ENSMUSG( | ENSMUSG( | 0.409247 | 0.000583 | YES | NO | NA |
| Cep192 | ENSMUSG( | ENSMUSG( | -0.04044 | 0.422853 | NO | NO | NA |
| Mtmr3 | ENSMUSG( | ENSMUSG( | -0.21072 | 0.002022 | YES | NO | NA |
| Map3k1 | ENSMUSG( | ENSMUSG( | -0.33767 | 2.86E-06 | YES | NO | NA |
| Marchf6 | ENSMUSG( | ENSMUSG( | -0.22078 | 5.18E-05 | YES | NO | NA |
| Cdk9 | ENSMUSG( | ENSMUSG( | 0.085468 | 0.428872 | NO | NO | NA |
| Psma3 | ENSMUSG( | ENSMUSG( | 0.246742 | 7.14E-06 | YES | NO | NA |
| Slc3a2 | ENSMUSG( | ENSMUSG( | 0.13569 | 0.204861 | NO | NO | NA |
| Mad2l1 | ENSMUSG( | ENSMUSG( | 0.290663 | 0.000192 | YES | NO | NA |
| Senp6 | ENSMUSG( | ENSMUSG( | -0.03197 | 0.758117 | NO | NO | NA |
| Nolc1 | ENSMUSG( | ENSMUSG( | 0.569453 | 2.71E-05 | YES | NO | NA |
| Apaf1 | ENSMUSG( | ENSMUSG( | -0.29566 | 8.06E-08 | YES | NO | NA |
| Polr1d | ENSMUSG( | ENSMUSG( | 0.231785 | 0.000717 | YES | NO | NA |
| Phc3 | ENSMUSG( | ENSMUSG( | -0.29268 | 5.91E-06 | YES | NO | NA |

| | | | | | | | |
|---|---|---|---|---|---|---|---|
| Lamc1 | ENSMUSG( | ENSMUSG( | -0.08907 | 0.165825 | NO | NO | NA |
| Pip4k2b | ENSMUSG( | ENSMUSG( | -0.16652 | 0.002465 | YES | NO | NA |
| Emp3 | ENSMUSG( | ENSMUSG( | -0.10711 | 0.180499 | NO | NO | NA |
| Pik3cg | ENSMUSG( | ENSMUSG( | -0.19185 | 6.20E-05 | YES | NO | NA |
| Tpp2 | ENSMUSG( | ENSMUSG( | -0.21689 | 4.16E-05 | YES | NO | NA |
| Tulp4 | ENSMUSG( | ENSMUSG( | 0.162404 | 0.124188 | NO | NO | NA |
| Sirt7 | ENSMUSG( | ENSMUSG( | -0.18295 | 2.52E-05 | YES | NO | NA |
| Mia3 | ENSMUSG( | ENSMUSG( | -0.22729 | 0.001158 | YES | NO | NA |
| Cxxc1 | ENSMUSG( | ENSMUSG( | -0.27465 | 3.44E-06 | YES | NO | NA |
| Ralgapb | ENSMUSG( | ENSMUSG( | -0.29843 | 0.000118 | YES | NO | NA |
| Cherp | ENSMUSG( | ENSMUSG( | -0.03678 | 0.391549 | NO | NO | NA |
| Kbtbd11 | ENSMUSG( | ENSMUSG( | 0.368089 | 0.001371 | YES | NO | NA |
| Abce1 | ENSMUSG( | ENSMUSG( | 0.413814 | 6.09E-07 | YES | NO | NA |
| Ppp1r12c | ENSMUSG( | ENSMUSG( | 0.051246 | 0.448353 | NO | NO | NA |
| Plekhg2 | ENSMUSG( | ENSMUSG( | -0.46926 | 6.29E-06 | YES | NO | NA |
| Ppp3r1 | ENSMUSG( | ENSMUSG( | -0.11313 | 0.17233 | NO | NO | NA |
| Slc2a3 | ENSMUSG( | ENSMUSG( | 0.080298 | 0.417345 | NO | NO | NA |
| Hba-a2 | ENSMUSG( | ENSMUSG( | 0.394751 | 0.63825 | NO | NO | NA |
| Mkrn1 | ENSMUSG( | ENSMUSG( | -0.1576 | 0.108435 | NO | NO | NA |
| Zmynd11 | ENSMUSG( | ENSMUSG( | -0.12045 | 0.295427 | NO | NO | NA |
| Nfat5 | ENSMUSG( | ENSMUSG( | -0.69138 | 1.62E-10 | YES | NO | NA |
| Zc3h15 | ENSMUSG( | ENSMUSG( | 0.005532 | 0.950806 | NO | NO | NA |
| Hba-a1 | ENSMUSG( | ENSMUSG( | 0.64113 | 0.397233 | NO | NO | NA |
| Ubap2 | ENSMUSG( | ENSMUSG( | 0.12888 | 0.020717 | YES | NO | NA |
| Mrtfa | ENSMUSG( | ENSMUSG( | -0.51978 | 4.93E-12 | YES | NO | NA |
| Ykt6 | ENSMUSG( | ENSMUSG( | 0.033589 | 0.640929 | NO | NO | NA |
| Cdca8 | ENSMUSG( | ENSMUSG( | 0.180694 | 0.111828 | NO | NO | NA |
| Selenof | ENSMUSG( | ENSMUSG( | 0.056505 | 0.595777 | NO | NO | NA |
| Txlna | ENSMUSG( | ENSMUSG( | 0.169808 | 0.003334 | YES | NO | NA |
| Elk3 | ENSMUSG( | ENSMUSG( | -0.22051 | 0.000397 | YES | NO | NA |
| Rnaset2a | ENSMUSG( | ENSMUSG( | -0.41499 | 0.000454 | YES | NO | NA |
| Ppp1r15a | ENSMUSG( | ENSMUSG( | 0.178672 | 0.567401 | NO | NO | NA |
| Gpd2 | ENSMUSG( | ENSMUSG( | -0.26827 | 6.12E-05 | YES | NO | NA |
| Gfpt1 | ENSMUSG( | ENSMUSG( | -0.64276 | 2.62E-16 | YES | NO | NA |
| Glyr1 | ENSMUSG( | ENSMUSG( | -0.03177 | 0.604949 | NO | NO | NA |
| Resf1 | ENSMUSG( | ENSMUSG( | -0.37117 | 1.50E-05 | YES | NO | NA |
| Map1lc3b | ENSMUSG( | ENSMUSG( | -0.26635 | 0.005388 | YES | NO | NA |
| Ak2 | ENSMUSG( | ENSMUSG( | 0.175921 | 0.070426 | NO | NO | NA |
| Naa50 | ENSMUSG( | ENSMUSG( | 0.198764 | 0.042344 | YES | NO | NA |
| Zc3h7b | ENSMUSG( | ENSMUSG( | 0.112403 | 0.062828 | NO | NO | NA |
| Rab35 | ENSMUSG( | ENSMUSG( | -0.17673 | 0.002123 | YES | NO | NA |
| Vps4b | ENSMUSG( | ENSMUSG( | -0.16957 | 0.000589 | YES | NO | NA |
| Zfp646 | ENSMUSG( | ENSMUSG( | -0.41521 | 1.32E-08 | YES | NO | NA |
| Gpbp1 | ENSMUSG( | ENSMUSG( | -0.27602 | 6.74E-05 | YES | NO | NA |
| Cmtm7 | ENSMUSG( | ENSMUSG( | 0.738129 | 4.70E-08 | YES | NO | NA |
| Nol7 | ENSMUSG( | ENSMUSG( | 0.210299 | 0.016261 | YES | NO | NA |
| Gramd4 | ENSMUSG( | ENSMUSG( | -0.40436 | 4.32E-06 | YES | NO | NA |

| | | | | | | | |
|---|---|---|---|---|---|---|---|
| Ino80d | ENSMUSG( | ENSMUSG( | -0.5829 | 5.77E-13 | YES | NO | NA |
| Axin1 | ENSMUSG( | ENSMUSG( | -0.07907 | 0.32874 | NO | NO | NA |
| Clptm1 | ENSMUSG( | ENSMUSG( | -0.02835 | 0.57285 | NO | NO | NA |
| Ube2j1 | ENSMUSG( | ENSMUSG( | 0.109629 | 0.203609 | NO | NO | NA |
| Ggnbp2 | ENSMUSG( | ENSMUSG( | -0.22576 | 0.002366 | YES | NO | NA |
| Idh3b | ENSMUSG( | ENSMUSG( | 0.076836 | 0.121615 | NO | NO | NA |
| Rhog | ENSMUSG( | ENSMUSG( | -0.21983 | 3.68E-06 | YES | NO | NA |
| Ap3b1 | ENSMUSG( | ENSMUSG( | -0.25382 | 0.000469 | YES | NO | NA |
| Sec62 | ENSMUSG( | ENSMUSG( | -0.24846 | 0.008653 | YES | NO | NA |
| Paxip1 | ENSMUSG( | ENSMUSG( | 0.179451 | 0.011434 | YES | NO | NA |
| Cpsf2 | ENSMUSG( | ENSMUSG( | 0.119073 | 0.131525 | NO | NO | NA |
| Dennd1b | ENSMUSG( | ENSMUSG( | -0.31557 | 9.17E-05 | YES | NO | NA |
| Dnajc2 | ENSMUSG( | ENSMUSG( | 0.3296 | 0.002948 | YES | NO | NA |
| Nusap1 | ENSMUSG( | ENSMUSG( | 0.072625 | 0.491395 | NO | NO | NA |
| Sp4 | ENSMUSG( | ENSMUSG( | -0.56952 | 2.97E-09 | YES | NO | NA |
| Rbm17 | ENSMUSG( | ENSMUSG( | 0.230386 | 0.062285 | NO | NO | NA |
| Stk11 | ENSMUSG( | ENSMUSG( | -0.10676 | 0.085384 | NO | NO | NA |
| Coa5 | ENSMUSG( | ENSMUSG( | -0.09096 | 0.341004 | NO | NO | NA |
| Srpk2 | ENSMUSG( | ENSMUSG( | 0.027699 | 0.772919 | NO | NO | NA |
| Cyb5b | ENSMUSG( | ENSMUSG( | 0.121874 | 0.033621 | YES | NO | NA |
| Psma1 | ENSMUSG( | ENSMUSG( | 0.127488 | 0.051964 | NO | NO | NA |
| Ubash3a | ENSMUSG( | ENSMUSG( | -0.15075 | 0.014407 | YES | NO | NA |
| Tnpo3 | ENSMUSG( | ENSMUSG( | 0.190707 | 0.004721 | YES | NO | NA |
| Ncoa3 | ENSMUSG( | ENSMUSG( | -0.52004 | 4.02E-11 | YES | NO | NA |
| Cited2 | ENSMUSG( | ENSMUSG( | 0.03107 | 0.850856 | NO | NO | NA |
| Dlgap5 | ENSMUSG( | ENSMUSG( | 0.252514 | 0.009389 | YES | NO | NA |
| Nup214 | ENSMUSG( | ENSMUSG( | -0.05507 | 0.459697 | NO | NO | NA |
| Ube2m | ENSMUSG( | ENSMUSG( | 0.066282 | 0.474445 | NO | NO | NA |
| Mapk8ip3 | ENSMUSG( | ENSMUSG( | -0.21559 | 0.000192 | YES | NO | NA |
| Gapvd1 | ENSMUSG( | ENSMUSG( | -0.08013 | 0.163563 | NO | NO | NA |
| Aspm | ENSMUSG( | ENSMUSG( | -0.01521 | 0.87566 | NO | NO | NA |
| Psmb7 | ENSMUSG( | ENSMUSG( | 0.196709 | 0.02016 | YES | NO | NA |
| Btg2 | ENSMUSG( | ENSMUSG( | -0.12545 | 0.370722 | NO | NO | NA |
| Nek9 | ENSMUSG( | ENSMUSG( | -0.02791 | 0.652767 | NO | NO | NA |
| Gabpa | ENSMUSG( | ENSMUSG( | 0.019641 | 0.800548 | NO | NO | NA |
| Cnp | ENSMUSG( | ENSMUSG( | -0.61847 | 5.35E-13 | YES | NO | NA |
| Cd84 | ENSMUSG( | ENSMUSG( | -0.41629 | 7.58E-05 | YES | NO | NA |
| Pou2f2 | ENSMUSG( | ENSMUSG( | -1.23857 | 4.28E-07 | YES | YES | Downregulated |
| R3hdm4 | ENSMUSG( | ENSMUSG( | -0.36405 | 1.19E-11 | YES | NO | NA |
| Foxn3 | ENSMUSG( | ENSMUSG( | -0.3392 | 0.000102 | YES | NO | NA |
| Setd1b | ENSMUSG( | ENSMUSG( | -0.56917 | 1.83E-09 | YES | NO | NA |
| Chtf8 | ENSMUSG( | ENSMUSG( | -0.14965 | 0.000239 | YES | NO | NA |
| Mcmbp | ENSMUSG( | ENSMUSG( | 0.085783 | 0.127982 | NO | NO | NA |
| Pfkl | ENSMUSG( | ENSMUSG( | 0.486787 | 0.000913 | YES | NO | NA |
| Syf2 | ENSMUSG( | ENSMUSG( | -0.12815 | 0.240782 | NO | NO | NA |
| Peli1 | ENSMUSG( | ENSMUSG( | -0.34615 | 0.000571 | YES | NO | NA |
| Pycard | ENSMUSG( | ENSMUSG( | -0.31628 | 0.000372 | YES | NO | NA |

| | | | | | | | |
|---|---|---|---|---|---|---|---|
| Erh | ENSMUSG( | ENSMUSG( | 0.43494 | 7.24E-09 | YES | NO | NA |
| Atp5j | ENSMUSG( | ENSMUSG( | 0.070504 | 0.581457 | NO | NO | NA |
| Gtf2f1 | ENSMUSG( | ENSMUSG( | 0.281816 | 0.005713 | YES | NO | NA |
| Psmc2 | ENSMUSG( | ENSMUSG( | 0.166278 | 0.017462 | YES | NO | NA |
| Dido1 | ENSMUSG( | ENSMUSG( | -0.30282 | 8.00E-09 | YES | NO | NA |
| Eri1 | ENSMUSG( | ENSMUSG( | 0.117763 | 0.112037 | NO | NO | NA |
| Lilrb4b | ENSMUSG( | ENSMUSG( | -0.09191 | 0.63616 | NO | NO | NA |
| Eif2s1 | ENSMUSG( | ENSMUSG( | 0.280456 | 0.007195 | YES | NO | NA |
| Eif4e2 | ENSMUSG( | ENSMUSG( | -0.11416 | 0.101393 | NO | NO | NA |
| Cbx1 | ENSMUSG( | ENSMUSG( | -0.06731 | 0.558551 | NO | NO | NA |
| Rabgap1l | ENSMUSG( | ENSMUSG( | -0.11182 | 0.244982 | NO | NO | NA |
| Dnah8 | ENSMUSG( | ENSMUSG( | -1.25553 | 6.98E-13 | YES | YES | Downregulated |
| Phf2 | ENSMUSG( | ENSMUSG( | -0.24848 | 8.98E-06 | YES | NO | NA |
| Srebf1 | ENSMUSG( | ENSMUSG( | 0.089006 | 0.330787 | NO | NO | NA |
| Gpsm3 | ENSMUSG( | ENSMUSG( | -0.33471 | 1.54E-08 | YES | NO | NA |
| Tapbpl | ENSMUSG( | ENSMUSG( | -0.18317 | 0.014881 | YES | NO | NA |
| Zfp292 | ENSMUSG( | ENSMUSG( | -0.41964 | 1.02E-06 | YES | NO | NA |
| Hnrnpll | ENSMUSG( | ENSMUSG( | 0.299151 | 0.000499 | YES | NO | NA |
| Snrk | ENSMUSG( | ENSMUSG( | -0.42212 | 8.51E-08 | YES | NO | NA |
| Ncapg | ENSMUSG( | ENSMUSG( | 0.190587 | 0.069886 | NO | NO | NA |
| Med15 | ENSMUSG( | ENSMUSG( | -0.13059 | 0.060568 | NO | NO | NA |
| Chfr | ENSMUSG( | ENSMUSG( | -0.2006 | 0.001902 | YES | NO | NA |
| Manf | ENSMUSG( | ENSMUSG( | 0.020442 | 0.892865 | NO | NO | NA |
| Gtf3c2 | ENSMUSG( | ENSMUSG( | 0.031745 | 0.623052 | NO | NO | NA |
| C2cd5 | ENSMUSG( | ENSMUSG( | 0.078217 | 0.39556 | NO | NO | NA |
| H2aj | ENSMUSG( | ENSMUSG( | 0.468035 | 2.36E-05 | YES | NO | NA |
| Strn | ENSMUSG( | ENSMUSG( | -0.18582 | 0.004774 | YES | NO | NA |
| Emg1 | ENSMUSG( | ENSMUSG( | 0.087237 | 0.025229 | YES | NO | NA |
| Mdn1 | ENSMUSG( | ENSMUSG( | 0.172034 | 0.089611 | NO | NO | NA |
| Rsl1d1 | ENSMUSG( | ENSMUSG( | 0.486786 | 3.92E-07 | YES | NO | NA |
| Dynll1 | ENSMUSG( | ENSMUSG( | -0.25843 | 0.000176 | YES | NO | NA |
| Zfp445 | ENSMUSG( | ENSMUSG( | -0.26789 | 0.000941 | YES | NO | NA |
| Aurkb | ENSMUSG( | ENSMUSG( | 0.315509 | 2.03E-09 | YES | NO | NA |
| Slc7a1 | ENSMUSG( | ENSMUSG( | 0.334084 | 0.000191 | YES | NO | NA |
| Smc5 | ENSMUSG( | ENSMUSG( | 0.065188 | 0.435434 | NO | NO | NA |
| Nop58 | ENSMUSG( | ENSMUSG( | 0.506892 | 2.19E-05 | YES | NO | NA |
| Atp5o | ENSMUSG( | ENSMUSG( | 0.192587 | 0.012959 | YES | NO | NA |
| Dars | ENSMUSG( | ENSMUSG( | 0.188584 | 0.033234 | YES | NO | NA |
| Marf1 | ENSMUSG( | ENSMUSG( | -0.54462 | 4.54E-10 | YES | NO | NA |
| Cpt1a | ENSMUSG( | ENSMUSG( | -0.16765 | 0.214168 | NO | NO | NA |
| Dyrk1a | ENSMUSG( | ENSMUSG( | -0.18197 | 0.011015 | YES | NO | NA |
| Zbp1 | ENSMUSG( | ENSMUSG( | -0.78508 | 1.17E-10 | YES | NO | NA |
| Trafd1 | ENSMUSG( | ENSMUSG( | -0.42812 | 4.63E-10 | YES | NO | NA |
| Serp1 | ENSMUSG( | ENSMUSG( | -0.08382 | 0.209815 | NO | NO | NA |
| Uqcrc2 | ENSMUSG( | ENSMUSG( | 0.093227 | 0.166298 | NO | NO | NA |
| Znfx1 | ENSMUSG( | ENSMUSG( | -0.75993 | 6.76E-21 | YES | NO | NA |
| Rab21 | ENSMUSG( | ENSMUSG( | -0.25056 | 0.001186 | YES | NO | NA |

| | | | | | | | |
|---|---|---|---|---|---|---|---|
| Sik1 | ENSMUSG( | ENSMUSG( | 0.68946 | 0.080464 | NO | NO | NA |
| Edf1 | ENSMUSG( | ENSMUSG( | -0.09932 | 0.383894 | NO | NO | NA |
| Ccnb1 | ENSMUSG( | ENSMUSG( | 0.467707 | 3.44E-07 | YES | NO | NA |
| Ncbp1 | ENSMUSG( | ENSMUSG( | 0.113565 | 0.04187 | YES | NO | NA |
| Aip | ENSMUSG( | ENSMUSG( | -0.10517 | 0.2625 | NO | NO | NA |
| Rfx7 | ENSMUSG( | ENSMUSG( | -0.20528 | 0.000634 | YES | NO | NA |
| Inpp4b | ENSMUSG( | ENSMUSG( | 0.042043 | 0.654211 | NO | NO | NA |
| Gm20186 | ENSMUSG( | ENSMUSG( | 0.572506 | 0.000262 | YES | NO | NA |
| Chp1 | ENSMUSG( | ENSMUSG( | -0.17703 | 0.021823 | YES | NO | NA |
| Mat2a | ENSMUSG( | ENSMUSG( | -0.08086 | 0.184406 | NO | NO | NA |
| Rab14 | ENSMUSG( | ENSMUSG( | -0.22139 | 2.48E-05 | YES | NO | NA |
| Suds3 | ENSMUSG( | ENSMUSG( | -0.0231 | 0.736696 | NO | NO | NA |
| Vps13a | ENSMUSG( | ENSMUSG( | -0.20004 | 0.023462 | YES | NO | NA |
| Lta4h | ENSMUSG( | ENSMUSG( | -0.10135 | 0.036809 | YES | NO | NA |
| Fen1 | ENSMUSG( | ENSMUSG( | 0.267875 | 0.000681 | YES | NO | NA |
| Trp53 | ENSMUSG( | ENSMUSG( | 0.202688 | 0.000635 | YES | NO | NA |
| Gosr2 | ENSMUSG( | ENSMUSG( | -0.02619 | 0.758671 | NO | NO | NA |
| Sms | ENSMUSG( | ENSMUSG( | 0.151608 | 0.004178 | YES | NO | NA |
| Supt20 | ENSMUSG( | ENSMUSG( | -0.36161 | 4.25E-07 | YES | NO | NA |
| Tk1 | ENSMUSG( | ENSMUSG( | 0.505595 | 1.13E-16 | YES | NO | NA |
| Emb | ENSMUSG( | ENSMUSG( | 0.802709 | 2.94E-05 | YES | NO | NA |
| Gatad2b | ENSMUSG( | ENSMUSG( | -0.37497 | 2.25E-09 | YES | NO | NA |
| Stag1 | ENSMUSG( | ENSMUSG( | -0.10532 | 0.160654 | NO | NO | NA |
| Clptm1l | ENSMUSG( | ENSMUSG( | -0.02983 | 0.643741 | NO | NO | NA |
| Eea1 | ENSMUSG( | ENSMUSG( | 0.067382 | 0.63164 | NO | NO | NA |
| Ankfy1 | ENSMUSG( | ENSMUSG( | -0.30599 | 4.44E-09 | YES | NO | NA |
| Satb1 | ENSMUSG( | ENSMUSG( | 0.835623 | 5.24E-08 | YES | NO | NA |
| Prdm2 | ENSMUSG( | ENSMUSG( | -0.75446 | 5.27E-20 | YES | NO | NA |
| Micu2 | ENSMUSG( | ENSMUSG( | 0.178724 | 0.040977 | YES | NO | NA |
| Lmf2 | ENSMUSG( | ENSMUSG( | 0.008308 | 0.91235 | NO | NO | NA |
| Tnik | ENSMUSG( | ENSMUSG( | -0.60511 | 2.66E-09 | YES | NO | NA |
| Tcirg1 | ENSMUSG( | ENSMUSG( | 0.003653 | 0.970144 | NO | NO | NA |
| Sdhb | ENSMUSG( | ENSMUSG( | 0.198921 | 0.002067 | YES | NO | NA |
| Fmr1 | ENSMUSG( | ENSMUSG( | -0.13601 | 0.091116 | NO | NO | NA |
| Asf1b | ENSMUSG( | ENSMUSG( | 0.207501 | 0.000646 | YES | NO | NA |
| Prkcsh | ENSMUSG( | ENSMUSG( | 0.045309 | 0.476215 | NO | NO | NA |
| Ilf3 | ENSMUSG( | ENSMUSG( | 0.168975 | 0.004574 | YES | NO | NA |
| Prpf18 | ENSMUSG( | ENSMUSG( | -0.10915 | 0.129726 | NO | NO | NA |
| Ankib1 | ENSMUSG( | ENSMUSG( | -0.34044 | 3.30E-06 | YES | NO | NA |
| Ranbp9 | ENSMUSG( | ENSMUSG( | -0.17185 | 0.01957 | YES | NO | NA |
| Eif6 | ENSMUSG( | ENSMUSG( | 0.256311 | 0.000877 | YES | NO | NA |
| Crebzf | ENSMUSG( | ENSMUSG( | -0.19929 | 0.003238 | YES | NO | NA |
| Serpina3g | ENSMUSG( | ENSMUSG( | 0.314651 | 0.02687 | YES | NO | NA |
| Adam17 | ENSMUSG( | ENSMUSG( | -0.13341 | 0.050511 | NO | NO | NA |
| Pds5b | ENSMUSG( | ENSMUSG( | -0.06446 | 0.512691 | NO | NO | NA |
| Sf3a2 | ENSMUSG( | ENSMUSG( | 0.0902 | 0.212685 | NO | NO | NA |
| Snx3 | ENSMUSG( | ENSMUSG( | -0.14956 | 2.18E-05 | YES | NO | NA |

| | | | | | | | |
|---|---|---|---|---|---|---|---|
| Nudt21 | ENSMUSG( | ENSMUSG( | 0.21669 | 2.86E-06 | YES | NO | NA |
| Wtap | ENSMUSG( | ENSMUSG( | -0.0886 | 0.088135 | NO | NO | NA |
| Rps6ka3 | ENSMUSG( | ENSMUSG( | 0.035826 | 0.546443 | NO | NO | NA |
| Mdm2 | ENSMUSG( | ENSMUSG( | -0.26604 | 3.99E-05 | YES | NO | NA |
| Ubqln1 | ENSMUSG( | ENSMUSG( | -0.07898 | 0.185248 | NO | NO | NA |
| Ddx27 | ENSMUSG( | ENSMUSG( | 0.276244 | 0.00388 | YES | NO | NA |
| Phf12 | ENSMUSG( | ENSMUSG( | -0.20132 | 1.90E-08 | YES | NO | NA |
| Ranbp3 | ENSMUSG( | ENSMUSG( | -0.04941 | 0.334781 | NO | NO | NA |
| Samm50 | ENSMUSG( | ENSMUSG( | 0.302178 | 7.35E-05 | YES | NO | NA |
| Tprgl | ENSMUSG( | ENSMUSG( | -0.31568 | 0.000302 | YES | NO | NA |
| Ciz1 | ENSMUSG( | ENSMUSG( | -0.02235 | 0.815358 | NO | NO | NA |
| Rfc2 | ENSMUSG( | ENSMUSG( | 0.099333 | 0.10351 | NO | NO | NA |
| Far1 | ENSMUSG( | ENSMUSG( | -0.1793 | 0.005174 | YES | NO | NA |
| Ipo9 | ENSMUSG( | ENSMUSG( | -0.09922 | 0.091202 | NO | NO | NA |
| Calu | ENSMUSG( | ENSMUSG( | 0.013433 | 0.880153 | NO | NO | NA |
| Adipor1 | ENSMUSG( | ENSMUSG( | -0.1089 | 0.160027 | NO | NO | NA |
| Gtf3c1 | ENSMUSG( | ENSMUSG( | -0.16512 | 0.000701 | YES | NO | NA |
| Cflar | ENSMUSG( | ENSMUSG( | -0.44285 | 7.77E-08 | YES | NO | NA |
| Zfp740 | ENSMUSG( | ENSMUSG( | -0.05726 | 0.479173 | NO | NO | NA |
| Orai2 | ENSMUSG( | ENSMUSG( | -0.20911 | 0.109767 | NO | NO | NA |
| Vamp8 | ENSMUSG( | ENSMUSG( | -0.04722 | 0.712266 | NO | NO | NA |
| Ylpm1 | ENSMUSG( | ENSMUSG( | -0.24505 | 8.72E-07 | YES | NO | NA |
| Ahcyl1 | ENSMUSG( | ENSMUSG( | 0.061867 | 0.279361 | NO | NO | NA |
| Glcci1 | ENSMUSG( | ENSMUSG( | -0.46538 | 8.46E-05 | YES | NO | NA |
| Rnf4 | ENSMUSG( | ENSMUSG( | -0.09738 | 0.129434 | NO | NO | NA |
| Psmd4 | ENSMUSG( | ENSMUSG( | 0.092957 | 0.289622 | NO | NO | NA |
| Cxcr4 | ENSMUSG( | ENSMUSG( | 0.349481 | 0.05543 | NO | NO | NA |
| Letm1 | ENSMUSG( | ENSMUSG( | 0.147579 | 0.020489 | YES | NO | NA |
| Ighg2c | ENSMUSG( | ENSMUSG( | -0.75072 | 0.508487 | NO | NO | NA |
| Rsbn1l | ENSMUSG( | ENSMUSG( | -0.30449 | 0.000177 | YES | NO | NA |
| Rtf1 | ENSMUSG( | ENSMUSG( | -0.01947 | 0.880986 | NO | NO | NA |
| Xiap | ENSMUSG( | ENSMUSG( | -0.40258 | 3.79E-08 | YES | NO | NA |
| Etfb | ENSMUSG( | ENSMUSG( | 0.193272 | 0.007168 | YES | NO | NA |
| Helz | ENSMUSG( | ENSMUSG( | -0.37606 | 2.27E-17 | YES | NO | NA |
| Ube2c | ENSMUSG( | ENSMUSG( | 0.179451 | 0.009217 | YES | NO | NA |
| Vdac1 | ENSMUSG( | ENSMUSG( | 0.405934 | 2.14E-06 | YES | NO | NA |
| Mau2 | ENSMUSG( | ENSMUSG( | -0.301 | 5.42E-05 | YES | NO | NA |
| Tdg | ENSMUSG( | ENSMUSG( | 0.464481 | 1.13E-05 | YES | NO | NA |
| Pacs1 | ENSMUSG( | ENSMUSG( | -0.41845 | 1.38E-07 | YES | NO | NA |
| Lypla1 | ENSMUSG( | ENSMUSG( | -0.07788 | 0.481807 | NO | NO | NA |
| Polr2b | ENSMUSG( | ENSMUSG( | 0.114193 | 0.057569 | NO | NO | NA |
| Rhoh | ENSMUSG( | ENSMUSG( | -0.28878 | 0.000101 | YES | NO | NA |
| Itm2c | ENSMUSG( | ENSMUSG( | -0.22436 | 0.015641 | YES | NO | NA |
| Banf1 | ENSMUSG( | ENSMUSG( | 0.187018 | 0.016493 | YES | NO | NA |
| Tspo | ENSMUSG( | ENSMUSG( | -0.20809 | 0.009306 | YES | NO | NA |
| Gigyf2 | ENSMUSG( | ENSMUSG( | -0.32687 | 0.002927 | YES | NO | NA |
| Crebbp | ENSMUSG( | ENSMUSG( | -0.50985 | 4.92E-07 | YES | NO | NA |

| | | | | | | | |
|---|---|---|---|---|---|---|---|
| Fgl2 | ENSMUSG( | ENSMUSG( | -1.03632 | 5.82E-12 | YES | YES | Downregulated |
| Emc10 | ENSMUSG( | ENSMUSG( | -0.0405 | 0.590438 | NO | NO | NA |
| Dynll2 | ENSMUSG( | ENSMUSG( | 0.054709 | 0.527225 | NO | NO | NA |
| Sec16a | ENSMUSG( | ENSMUSG( | -0.39633 | 6.88E-12 | YES | NO | NA |
| Adipor2 | ENSMUSG( | ENSMUSG( | -0.04031 | 0.600449 | NO | NO | NA |
| Sfswap | ENSMUSG( | ENSMUSG( | 0.003075 | 0.977199 | NO | NO | NA |
| Cert1 | ENSMUSG( | ENSMUSG( | -0.14221 | 0.015758 | YES | NO | NA |
| Fli1 | ENSMUSG( | ENSMUSG( | -0.315 | 4.19E-05 | YES | NO | NA |
| Dnajc14 | ENSMUSG( | ENSMUSG( | -0.31495 | 1.22E-08 | YES | NO | NA |
| Abtb2 | ENSMUSG( | ENSMUSG( | -0.89346 | 2.10E-08 | YES | NO | NA |
| Ddx1 | ENSMUSG( | ENSMUSG( | 0.524649 | 2.56E-07 | YES | NO | NA |
| Mgat2 | ENSMUSG( | ENSMUSG( | -0.01316 | 0.770717 | NO | NO | NA |
| Irf9 | ENSMUSG( | ENSMUSG( | -0.62379 | 3.21E-10 | YES | NO | NA |
| Limd1 | ENSMUSG( | ENSMUSG( | -0.34718 | 3.07E-11 | YES | NO | NA |
| Hmgcr | ENSMUSG( | ENSMUSG( | 0.17138 | 0.021844 | YES | NO | NA |
| Foxk1 | ENSMUSG( | ENSMUSG( | -0.26757 | 4.59E-05 | YES | NO | NA |
| Tmem127 | ENSMUSG( | ENSMUSG( | -0.42403 | 1.25E-06 | YES | NO | NA |
| Scamp3 | ENSMUSG( | ENSMUSG( | -0.20658 | 0.002825 | YES | NO | NA |
| Mcm3ap | ENSMUSG( | ENSMUSG( | -0.09763 | 0.208913 | NO | NO | NA |
| Cenpb | ENSMUSG( | ENSMUSG( | 0.103676 | 0.132792 | NO | NO | NA |
| Slc7a5 | ENSMUSG( | ENSMUSG( | 1.077879 | 5.89E-12 | YES | YES | Upregulated |
| Tra2a | ENSMUSG( | ENSMUSG( | -0.02779 | 0.691355 | NO | NO | NA |
| Pebp1 | ENSMUSG( | ENSMUSG( | 0.42691 | 5.47E-08 | YES | NO | NA |
| Pomp | ENSMUSG( | ENSMUSG( | 0.108341 | 0.068974 | NO | NO | NA |
| Wrnip1 | ENSMUSG( | ENSMUSG( | 0.016593 | 0.820264 | NO | NO | NA |
| Tmem234 | ENSMUSG( | ENSMUSG( | 0.010643 | 0.901916 | NO | NO | NA |
| Eif3g | ENSMUSG( | ENSMUSG( | 0.260531 | 0.003267 | YES | NO | NA |
| Selenow | ENSMUSG( | ENSMUSG( | -0.3015 | 6.36E-05 | YES | NO | NA |
| Gramd3 | ENSMUSG( | ENSMUSG( | -0.07875 | 0.527158 | NO | NO | NA |
| Cand1 | ENSMUSG( | ENSMUSG( | 0.082697 | 0.042735 | YES | NO | NA |
| Taf1d | ENSMUSG( | ENSMUSG( | 0.483907 | 7.96E-06 | YES | NO | NA |
| Ccar2 | ENSMUSG( | ENSMUSG( | 0.081515 | 0.252166 | NO | NO | NA |
| Fam111a | ENSMUSG( | ENSMUSG( | -0.06784 | 0.453495 | NO | NO | NA |
| Rmnd5a | ENSMUSG( | ENSMUSG( | -0.2035 | 0.020181 | YES | NO | NA |
| Ddi2 | ENSMUSG( | ENSMUSG( | -0.31471 | 2.72E-09 | YES | NO | NA |
| Prkcb | ENSMUSG( | ENSMUSG( | -0.46362 | 1.06E-07 | YES | NO | NA |
| Trappc10 | ENSMUSG( | ENSMUSG( | -0.12561 | 0.095211 | NO | NO | NA |
| Stxbp2 | ENSMUSG( | ENSMUSG( | -0.03307 | 0.609081 | NO | NO | NA |
| Bcor | ENSMUSG( | ENSMUSG( | -0.30915 | 8.32E-06 | YES | NO | NA |
| Cox6b1 | ENSMUSG( | ENSMUSG( | 0.10355 | 0.279867 | NO | NO | NA |
| Cse1l | ENSMUSG( | ENSMUSG( | 0.24216 | 0.010935 | YES | NO | NA |
| Psma6 | ENSMUSG( | ENSMUSG( | 0.230108 | 0.001609 | YES | NO | NA |
| Gm2808 | ENSMUSG( | ENSMUSG( | 0.008903 | 0.955338 | NO | NO | NA |
| Mta3 | ENSMUSG( | ENSMUSG( | -0.3185 | 2.26E-05 | YES | NO | NA |
| Cmah | ENSMUSG( | ENSMUSG( | -0.06842 | 0.682598 | NO | NO | NA |
| Ap2a1 | ENSMUSG( | ENSMUSG( | 0.088754 | 0.282929 | NO | NO | NA |
| Atp5l | ENSMUSG( | ENSMUSG( | 0.116777 | 0.326398 | NO | NO | NA |

| | | | | | | | |
|---|---|---|---|---|---|---|---|
| Cd96 | ENSMUSG( | ENSMUSG( | -0.47668 | 4.34E-10 | YES | NO | NA |
| Anapc2 | ENSMUSG( | ENSMUSG( | -0.07947 | 0.061576 | NO | NO | NA |
| Tomm70a | ENSMUSG( | ENSMUSG( | 0.265202 | 0.000536 | YES | NO | NA |
| Psmb6 | ENSMUSG( | ENSMUSG( | 0.325919 | 1.35E-05 | YES | NO | NA |
| Ptpra | ENSMUSG( | ENSMUSG( | -0.09853 | 0.107039 | NO | NO | NA |
| Ap1m1 | ENSMUSG( | ENSMUSG( | 0.055396 | 0.452233 | NO | NO | NA |
| Casp8 | ENSMUSG( | ENSMUSG( | -0.42968 | 1.61E-11 | YES | NO | NA |
| Mbtps1 | ENSMUSG( | ENSMUSG( | -0.09064 | 0.136138 | NO | NO | NA |
| Adnp | ENSMUSG( | ENSMUSG( | -0.16177 | 0.060584 | NO | NO | NA |
| Cdca3 | ENSMUSG( | ENSMUSG( | 0.455094 | 9.48E-10 | YES | NO | NA |
| Shmt2 | ENSMUSG( | ENSMUSG( | 0.608679 | 1.87E-08 | YES | NO | NA |
| Tatdn2 | ENSMUSG( | ENSMUSG( | 0.146416 | 0.106485 | NO | NO | NA |
| Sipa1l1 | ENSMUSG( | ENSMUSG( | 0.085124 | 0.176009 | NO | NO | NA |
| D16Ertd47 | ENSMUSG( | ENSMUSG( | -0.10013 | 0.097109 | NO | NO | NA |
| Ppp3ca | ENSMUSG( | ENSMUSG( | -0.09845 | 0.154201 | NO | NO | NA |
| Dync1li2 | ENSMUSG( | ENSMUSG( | -0.16126 | 0.031407 | YES | NO | NA |
| Dcaf15 | ENSMUSG( | ENSMUSG( | 0.08888 | 0.083555 | NO | NO | NA |
| Wdhd1 | ENSMUSG( | ENSMUSG( | 0.22179 | 2.63E-05 | YES | NO | NA |
| Sirt2 | ENSMUSG( | ENSMUSG( | -0.16472 | 0.034157 | YES | NO | NA |
| Evi2b | ENSMUSG( | ENSMUSG( | -0.51227 | 3.29E-07 | YES | NO | NA |
| Psmd14 | ENSMUSG( | ENSMUSG( | 0.161357 | 0.00034 | YES | NO | NA |
| Ahsa1 | ENSMUSG( | ENSMUSG( | -0.14697 | 0.10484 | NO | NO | NA |
| Parp9 | ENSMUSG( | ENSMUSG( | -0.35683 | 1.26E-05 | YES | NO | NA |
| Pdap1 | ENSMUSG( | ENSMUSG( | 0.267102 | 0.059566 | NO | NO | NA |
| Ppil2 | ENSMUSG( | ENSMUSG( | -0.19625 | 0.002291 | YES | NO | NA |
| Mthfd1 | ENSMUSG( | ENSMUSG( | 0.682557 | 1.16E-05 | YES | NO | NA |
| Shc1 | ENSMUSG( | ENSMUSG( | -0.06574 | 0.358484 | NO | NO | NA |
| Ssbp3 | ENSMUSG( | ENSMUSG( | -0.07019 | 0.41526 | NO | NO | NA |
| Ncapd3 | ENSMUSG( | ENSMUSG( | 0.058287 | 0.41526 | NO | NO | NA |
| Rbmxl1 | ENSMUSG( | ENSMUSG( | 0.273576 | 0.000589 | YES | NO | NA |
| Dnm1l | ENSMUSG( | ENSMUSG( | -0.02591 | 0.741426 | NO | NO | NA |
| Pdha1 | ENSMUSG( | ENSMUSG( | 0.273364 | 4.59E-05 | YES | NO | NA |
| Ccr2 | ENSMUSG( | ENSMUSG( | 0.792637 | 0.007176 | YES | NO | NA |
| Vamp2 | ENSMUSG( | ENSMUSG( | -0.48659 | 5.25E-09 | YES | NO | NA |
| Nmt1 | ENSMUSG( | ENSMUSG( | 0.042639 | 0.451846 | NO | NO | NA |
| Setd3 | ENSMUSG( | ENSMUSG( | -0.09232 | 0.218464 | NO | NO | NA |
| Ccl4 | ENSMUSG( | ENSMUSG( | -0.2185 | 0.319818 | NO | NO | NA |
| Naa12 | ENSMUSG( | ENSMUSG( | 0.27899 | 9.28E-05 | YES | NO | NA |
| Rtn4 | ENSMUSG( | ENSMUSG( | -0.03184 | 0.722873 | NO | NO | NA |
| Vps13b | ENSMUSG( | ENSMUSG( | -0.34104 | 1.65E-05 | YES | NO | NA |
| C1galt1 | ENSMUSG( | ENSMUSG( | -0.12814 | 0.254124 | NO | NO | NA |
| Cd44 | ENSMUSG( | ENSMUSG( | -0.11162 | 0.155997 | NO | NO | NA |
| Mef2a | ENSMUSG( | ENSMUSG( | -0.04758 | 0.646606 | NO | NO | NA |
| Adrm1 | ENSMUSG( | ENSMUSG( | 0.128271 | 0.034997 | YES | NO | NA |
| Trpm7 | ENSMUSG( | ENSMUSG( | -0.07074 | 0.541551 | NO | NO | NA |
| Psmc4 | ENSMUSG( | ENSMUSG( | 0.26221 | 6.04E-06 | YES | NO | NA |
| Dhcr24 | ENSMUSG( | ENSMUSG( | 1.224689 | 2.52E-05 | YES | YES | Upregulated |

| | | | | | | | |
|---|---|---|---|---|---|---|---|
| Ifi208 | ENSMUSG( | ENSMUSG( | -1.2669 | 1.02E-14 | YES | YES | Downregulated |
| Adgrl1 | ENSMUSG( | ENSMUSG( | -0.20576 | 1.10E-06 | YES | NO | NA |
| Keap1 | ENSMUSG( | ENSMUSG( | 0.058194 | 0.435788 | NO | NO | NA |
| Dpp9 | ENSMUSG( | ENSMUSG( | -0.01884 | 0.737715 | NO | NO | NA |
| Ssr2 | ENSMUSG( | ENSMUSG( | 0.181076 | 0.145686 | NO | NO | NA |
| Cstf2 | ENSMUSG( | ENSMUSG( | 0.139988 | 0.025159 | YES | NO | NA |
| Hipk3 | ENSMUSG( | ENSMUSG( | -0.13114 | 0.055139 | NO | NO | NA |
| Psen1 | ENSMUSG( | ENSMUSG( | -0.15823 | 0.001731 | YES | NO | NA |
| Mapk6 | ENSMUSG( | ENSMUSG( | 0.090573 | 0.227436 | NO | NO | NA |
| Tmed5 | ENSMUSG( | ENSMUSG( | -0.17787 | 0.007853 | YES | NO | NA |
| Zfc3h1 | ENSMUSG( | ENSMUSG( | -0.41151 | 1.45E-09 | YES | NO | NA |
| Plekho2 | ENSMUSG( | ENSMUSG( | -0.19326 | 0.000806 | YES | NO | NA |
| Nr1h2 | ENSMUSG( | ENSMUSG( | -0.1376 | 0.058218 | NO | NO | NA |
| Twf2 | ENSMUSG( | ENSMUSG( | -0.1536 | 1.72E-05 | YES | NO | NA |
| Oas3 | ENSMUSG( | ENSMUSG( | -1.31393 | 3.73E-23 | YES | YES | Downregulated |
| Rfc1 | ENSMUSG( | ENSMUSG( | 0.251478 | 0.001926 | YES | NO | NA |
| Sumo3 | ENSMUSG( | ENSMUSG( | 0.255697 | 0.000917 | YES | NO | NA |
| Nek7 | ENSMUSG( | ENSMUSG( | -0.09528 | 0.25474 | NO | NO | NA |
| Trim26 | ENSMUSG( | ENSMUSG( | -0.4607 | 2.24E-13 | YES | NO | NA |
| Rbm42 | ENSMUSG( | ENSMUSG( | 0.043194 | 0.466943 | NO | NO | NA |
| Ech1 | ENSMUSG( | ENSMUSG( | -0.25357 | 1.91E-05 | YES | NO | NA |
| Ip6k1 | ENSMUSG( | ENSMUSG( | -0.31562 | 0.000293 | YES | NO | NA |
| Actn1 | ENSMUSG( | ENSMUSG( | -0.32376 | 3.07E-06 | YES | NO | NA |
| Vps26b | ENSMUSG( | ENSMUSG( | -0.20855 | 0.000669 | YES | NO | NA |
| Dicer1 | ENSMUSG( | ENSMUSG( | -0.09627 | 0.064125 | NO | NO | NA |
| Med13l | ENSMUSG( | ENSMUSG( | -0.29719 | 0.001439 | YES | NO | NA |
| Golm1 | ENSMUSG( | ENSMUSG( | -0.66313 | 6.64E-12 | YES | NO | NA |
| Wdr43 | ENSMUSG( | ENSMUSG( | 0.436201 | 4.51E-05 | YES | NO | NA |
| Cox5a | ENSMUSG( | ENSMUSG( | 0.384745 | 9.35E-07 | YES | NO | NA |
| Ahctf1 | ENSMUSG( | ENSMUSG( | 0.048836 | 0.482229 | NO | NO | NA |
| Ssbp4 | ENSMUSG( | ENSMUSG( | 0.137506 | 0.005439 | YES | NO | NA |
| Uhrf2 | ENSMUSG( | ENSMUSG( | -0.26771 | 7.38E-08 | YES | NO | NA |
| Sec14l1 | ENSMUSG( | ENSMUSG( | -0.30079 | 0.00045 | YES | NO | NA |
| Cactin | ENSMUSG( | ENSMUSG( | 0.125967 | 0.22157 | NO | NO | NA |
| Wbp1l | ENSMUSG( | ENSMUSG( | -0.31997 | 4.61E-05 | YES | NO | NA |
| Ier5 | ENSMUSG( | ENSMUSG( | -0.05672 | 0.565343 | NO | NO | NA |
| Slc6a6 | ENSMUSG( | ENSMUSG( | 0.162391 | 0.078906 | NO | NO | NA |
| Dck | ENSMUSG( | ENSMUSG( | -0.00084 | 0.990103 | NO | NO | NA |
| Usp15 | ENSMUSG( | ENSMUSG( | -0.11213 | 0.053225 | NO | NO | NA |
| Iars2 | ENSMUSG( | ENSMUSG( | 0.459417 | 3.49E-06 | YES | NO | NA |
| Arfgef2 | ENSMUSG( | ENSMUSG( | -0.0395 | 0.636349 | NO | NO | NA |
| Pum1 | ENSMUSG( | ENSMUSG( | -0.3271 | 7.70E-07 | YES | NO | NA |
| Ubxn4 | ENSMUSG( | ENSMUSG( | -0.13603 | 0.170545 | NO | NO | NA |
| Preb | ENSMUSG( | ENSMUSG( | -0.05349 | 0.548545 | NO | NO | NA |
| Usp24 | ENSMUSG( | ENSMUSG( | -0.13105 | 0.000371 | YES | NO | NA |
| Trim8 | ENSMUSG( | ENSMUSG( | -0.21142 | 0.011368 | YES | NO | NA |
| Osbpl9 | ENSMUSG( | ENSMUSG( | -0.13519 | 0.203473 | NO | NO | NA |

| | | | | | | | |
|---|---|---|---|---|---|---|---|
| Taf10 | ENSMUSG( | ENSMUSG( | -0.05094 | 0.477503 | NO | NO | NA |
| Jade2 | ENSMUSG( | ENSMUSG( | -0.44784 | 3.07E-10 | YES | NO | NA |
| Srp9 | ENSMUSG( | ENSMUSG( | 0.25363 | 0.00201 | YES | NO | NA |
| Adam8 | ENSMUSG( | ENSMUSG( | 0.468029 | 0.000455 | YES | NO | NA |
| Rbm10 | ENSMUSG( | ENSMUSG( | -0.22136 | 0.002779 | YES | NO | NA |
| Srf | ENSMUSG( | ENSMUSG( | -0.05697 | 0.083576 | NO | NO | NA |
| Fbrs | ENSMUSG( | ENSMUSG( | -0.34603 | 3.79E-05 | YES | NO | NA |
| Riok3 | ENSMUSG( | ENSMUSG( | -0.09131 | 0.305959 | NO | NO | NA |
| Maea | ENSMUSG( | ENSMUSG( | -0.03856 | 0.503722 | NO | NO | NA |
| Sec31a | ENSMUSG( | ENSMUSG( | -0.0117 | 0.851529 | NO | NO | NA |
| Szrd1 | ENSMUSG( | ENSMUSG( | -0.15084 | 0.02416 | YES | NO | NA |
| Osbpl8 | ENSMUSG( | ENSMUSG( | -0.37543 | 3.39E-05 | YES | NO | NA |
| Eloa | ENSMUSG( | ENSMUSG( | -0.25252 | 8.30E-06 | YES | NO | NA |
| Tmed7 | ENSMUSG( | ENSMUSG( | -0.10987 | 0.03795 | YES | NO | NA |
| Lgals9 | ENSMUSG( | ENSMUSG( | -0.41992 | 7.79E-12 | YES | NO | NA |
| Sin3a | ENSMUSG( | ENSMUSG( | -0.16146 | 0.00572 | YES | NO | NA |
| Trip11 | ENSMUSG( | ENSMUSG( | -0.32326 | 8.28E-05 | YES | NO | NA |
| Brd8 | ENSMUSG( | ENSMUSG( | -0.03301 | 0.653824 | NO | NO | NA |
| Nup98 | ENSMUSG( | ENSMUSG( | 0.030182 | 0.582781 | NO | NO | NA |
| Ubxn1 | ENSMUSG( | ENSMUSG( | -0.0122 | 0.908368 | NO | NO | NA |
| Msh2 | ENSMUSG( | ENSMUSG( | -0.01272 | 0.892196 | NO | NO | NA |
| Cwc15 | ENSMUSG( | ENSMUSG( | 0.095332 | 0.243669 | NO | NO | NA |
| Eps15 | ENSMUSG( | ENSMUSG( | -0.23115 | 0.000987 | YES | NO | NA |
| Gm4316 | ENSMUSG( | ENSMUSG( | 0.763988 | 6.98E-08 | YES | NO | NA |
| Bcap31 | ENSMUSG( | ENSMUSG( | -0.09171 | 0.373837 | NO | NO | NA |
| Brpf1 | ENSMUSG( | ENSMUSG( | -0.00269 | 0.960639 | NO | NO | NA |
| Larp7 | ENSMUSG( | ENSMUSG( | 0.152989 | 0.14585 | NO | NO | NA |
| Ecpas | ENSMUSG( | ENSMUSG( | 0.036396 | 0.606789 | NO | NO | NA |
| Adgrg1 | ENSMUSG( | ENSMUSG( | -0.67957 | 0.000845 | YES | NO | NA |
| Zc3h14 | ENSMUSG( | ENSMUSG( | 0.042953 | 0.576726 | NO | NO | NA |
| Snw1 | ENSMUSG( | ENSMUSG( | -0.05383 | 0.497112 | NO | NO | NA |
| Prdm1 | ENSMUSG( | ENSMUSG( | 0.136579 | 0.439866 | NO | NO | NA |
| Plp2 | ENSMUSG( | ENSMUSG( | 0.47447 | 2.96E-06 | YES | NO | NA |
| Dnmt3a | ENSMUSG( | ENSMUSG( | -0.29231 | 6.55E-11 | YES | NO | NA |
| Tbcd | ENSMUSG( | ENSMUSG( | 0.387697 | 0.000257 | YES | NO | NA |
| Ftsj3 | ENSMUSG( | ENSMUSG( | 0.435795 | 0.000337 | YES | NO | NA |
| Rasa1 | ENSMUSG( | ENSMUSG( | -0.37918 | 1.98E-05 | YES | NO | NA |
| Nub1 | ENSMUSG( | ENSMUSG( | -0.23469 | 0.000782 | YES | NO | NA |
| Setd1a | ENSMUSG( | ENSMUSG( | 0.041833 | 0.468029 | NO | NO | NA |
| E2f8 | ENSMUSG( | ENSMUSG( | 0.180738 | 0.024134 | YES | NO | NA |
| Tnrc6c | ENSMUSG( | ENSMUSG( | -0.31114 | 0.000357 | YES | NO | NA |
| Ttc14 | ENSMUSG( | ENSMUSG( | -0.2192 | 0.051189 | NO | NO | NA |
| Scap | ENSMUSG( | ENSMUSG( | 0.182105 | 0.009617 | YES | NO | NA |
| Gpr68 | ENSMUSG( | ENSMUSG( | -0.3274 | 5.04E-06 | YES | NO | NA |
| Sin3b | ENSMUSG( | ENSMUSG( | -0.0142 | 0.800548 | NO | NO | NA |
| Atp5j2 | ENSMUSG( | ENSMUSG( | 0.047545 | 0.474587 | NO | NO | NA |
| Bnip2 | ENSMUSG( | ENSMUSG( | -0.15867 | 0.008173 | YES | NO | NA |

| | | | | | | | |
|---|---|---|---|---|---|---|---|
| Dennd5a | ENSMUSG( | ENSMUSG( | -0.35188 | 4.76E-08 | YES | NO | NA |
| Svil | ENSMUSG( | ENSMUSG( | -0.38461 | 6.36E-09 | YES | NO | NA |
| Isy1 | ENSMUSG( | ENSMUSG( | 0.047103 | 0.630359 | NO | NO | NA |
| Zbtb38 | ENSMUSG( | ENSMUSG( | -0.36174 | 7.55E-06 | YES | NO | NA |
| Ogfr | ENSMUSG( | ENSMUSG( | -0.07604 | 0.417925 | NO | NO | NA |
| Cox6c | ENSMUSG( | ENSMUSG( | 0.122907 | 0.112963 | NO | NO | NA |
| Ankrd10 | ENSMUSG( | ENSMUSG( | -0.12245 | 0.093525 | NO | NO | NA |
| Stk26 | ENSMUSG( | ENSMUSG( | -0.10307 | 0.327267 | NO | NO | NA |
| Morc3 | ENSMUSG( | ENSMUSG( | -0.33201 | 2.44E-08 | YES | NO | NA |
| Aplp2 | ENSMUSG( | ENSMUSG( | -0.67406 | 7.82E-12 | YES | NO | NA |
| Map2k2 | ENSMUSG( | ENSMUSG( | -0.02891 | 0.505024 | NO | NO | NA |
| Bax | ENSMUSG( | ENSMUSG( | -0.00256 | 0.966814 | NO | NO | NA |
| Rin3 | ENSMUSG( | ENSMUSG( | -0.33164 | 2.01E-07 | YES | NO | NA |
| Syne3 | ENSMUSG( | ENSMUSG( | 0.233318 | 0.000451 | YES | NO | NA |
| Rap1gds1 | ENSMUSG( | ENSMUSG( | -0.1338 | 0.035296 | YES | NO | NA |
| Slc12a6 | ENSMUSG( | ENSMUSG( | -0.37023 | 0.000128 | YES | NO | NA |
| Cmpk1 | ENSMUSG( | ENSMUSG( | -0.01722 | 0.792398 | NO | NO | NA |
| Kat7 | ENSMUSG( | ENSMUSG( | -0.09406 | 0.031407 | YES | NO | NA |
| Egln1 | ENSMUSG( | ENSMUSG( | 0.036692 | 0.662264 | NO | NO | NA |
| Trim35 | ENSMUSG( | ENSMUSG( | 0.161165 | 0.002546 | YES | NO | NA |
| Srp68 | ENSMUSG( | ENSMUSG( | 0.042406 | 0.717666 | NO | NO | NA |
| Gm44175 | ENSMUSG( | ENSMUSG( | -0.74819 | 3.11E-13 | YES | NO | NA |
| Nod1 | ENSMUSG( | ENSMUSG( | -0.98474 | 3.65E-13 | YES | NO | NA |
| Nfx1 | ENSMUSG( | ENSMUSG( | -0.11387 | 0.043192 | YES | NO | NA |
| Zbtb1 | ENSMUSG( | ENSMUSG( | -0.32783 | 1.15E-07 | YES | NO | NA |
| Clip1 | ENSMUSG( | ENSMUSG( | 0.244953 | 0.011189 | YES | NO | NA |
| Asxl2 | ENSMUSG( | ENSMUSG( | -0.50062 | 1.38E-11 | YES | NO | NA |
| Prr12 | ENSMUSG( | ENSMUSG( | -0.51025 | 6.95E-11 | YES | NO | NA |
| Rbm8a | ENSMUSG( | ENSMUSG( | 0.247165 | 0.076878 | NO | NO | NA |
| Snrpa | ENSMUSG( | ENSMUSG( | 0.33722 | 0.000244 | YES | NO | NA |
| B230219D | ENSMUSG( | ENSMUSG( | -0.06815 | 0.519401 | NO | NO | NA |
| Cnppd1 | ENSMUSG( | ENSMUSG( | -0.09235 | 0.274672 | NO | NO | NA |
| Gmfb | ENSMUSG( | ENSMUSG( | -0.13411 | 0.03571 | YES | NO | NA |
| Sar1a | ENSMUSG( | ENSMUSG( | 0.00891 | 0.865833 | NO | NO | NA |
| Med1 | ENSMUSG( | ENSMUSG( | -0.21975 | 2.75E-07 | YES | NO | NA |
| Mbp | ENSMUSG( | ENSMUSG( | -0.28825 | 0.003878 | YES | NO | NA |
| Slc12a7 | ENSMUSG( | ENSMUSG( | -1.02016 | 2.30E-17 | YES | YES | Downregulated |
| Ddx19a | ENSMUSG( | ENSMUSG( | 0.276422 | 0.000105 | YES | NO | NA |
| Gmps | ENSMUSG( | ENSMUSG( | 0.313501 | 2.03E-06 | YES | NO | NA |
| Tcf20 | ENSMUSG( | ENSMUSG( | -0.22803 | 0.000303 | YES | NO | NA |
| Dcaf5 | ENSMUSG( | ENSMUSG( | -0.24702 | 0.000523 | YES | NO | NA |
| Arl2bp | ENSMUSG( | ENSMUSG( | -0.0246 | 0.848056 | NO | NO | NA |
| Alg10b | ENSMUSG( | ENSMUSG( | -0.04632 | 0.686967 | NO | NO | NA |
| Gsk3b | ENSMUSG( | ENSMUSG( | -0.4687 | 4.07E-09 | YES | NO | NA |
| Lrwd1 | ENSMUSG( | ENSMUSG( | 0.195395 | 0.009969 | YES | NO | NA |
| Fam177a | ENSMUSG( | ENSMUSG( | 0.130245 | 0.383904 | NO | NO | NA |
| Timm23 | ENSMUSG( | ENSMUSG( | 0.334551 | 0.000443 | YES | NO | NA |

| | | | | | | | |
|---|---|---|---|---|---|---|---|
| Rpl22l1 | ENSMUSG( | ENSMUSG( | 0.276427 | 0.000435 | YES | NO | NA |
| Cracr2a | ENSMUSG( | ENSMUSG( | -0.4945 | 3.50E-07 | YES | NO | NA |
| Triobp | ENSMUSG( | ENSMUSG( | -0.11935 | 0.011547 | YES | NO | NA |
| Tnfrsf9 | ENSMUSG( | ENSMUSG( | -0.05394 | 0.63678 | NO | NO | NA |
| Knl1 | ENSMUSG( | ENSMUSG( | 0.027559 | 0.793916 | NO | NO | NA |
| Rela | ENSMUSG( | ENSMUSG( | -0.12056 | 0.152586 | NO | NO | NA |
| Ifnar1 | ENSMUSG( | ENSMUSG( | -0.25361 | 0.004124 | YES | NO | NA |
| Tox4 | ENSMUSG( | ENSMUSG( | -0.12756 | 0.005164 | YES | NO | NA |
| Mybl2 | ENSMUSG( | ENSMUSG( | 0.477748 | 4.54E-08 | YES | NO | NA |
| Ap1g2 | ENSMUSG( | ENSMUSG( | -0.29726 | 2.64E-11 | YES | NO | NA |
| Atf1 | ENSMUSG( | ENSMUSG( | 0.159538 | 0.001012 | YES | NO | NA |
| Brd9 | ENSMUSG( | ENSMUSG( | 0.00235 | 0.981136 | NO | NO | NA |
| Rictor | ENSMUSG( | ENSMUSG( | -0.32161 | 0.000387 | YES | NO | NA |
| Stt3b | ENSMUSG( | ENSMUSG( | 0.171642 | 0.00131 | YES | NO | NA |
| Pmpca | ENSMUSG( | ENSMUSG( | 0.175742 | 0.005502 | YES | NO | NA |
| Pgd | ENSMUSG( | ENSMUSG( | 0.16728 | 0.018416 | YES | NO | NA |
| Apol7e | ENSMUSG( | ENSMUSG( | -0.46447 | 0.001362 | YES | NO | NA |
| Clk3 | ENSMUSG( | ENSMUSG( | -0.11545 | 0.030763 | YES | NO | NA |
| Trak2 | ENSMUSG( | ENSMUSG( | -0.1233 | 0.168416 | NO | NO | NA |
| Sbno2 | ENSMUSG( | ENSMUSG( | -0.12343 | 0.114669 | NO | NO | NA |
| Rhof | ENSMUSG( | ENSMUSG( | -0.53197 | 2.34E-10 | YES | NO | NA |
| Man2b2 | ENSMUSG( | ENSMUSG( | -0.16445 | 0.069372 | NO | NO | NA |
| Atp6v0c | ENSMUSG( | ENSMUSG( | -0.08966 | 0.093257 | NO | NO | NA |
| Gm20390 | ENSMUSG( | ENSMUSG( | 0.625644 | 2.32E-07 | YES | NO | NA |
| Arel1 | ENSMUSG( | ENSMUSG( | -0.34612 | 7.05E-05 | YES | NO | NA |
| Spop | ENSMUSG( | ENSMUSG( | -0.16969 | 0.010192 | YES | NO | NA |
| Sacs | ENSMUSG( | ENSMUSG( | -0.34003 | 4.29E-07 | YES | NO | NA |
| Nup93 | ENSMUSG( | ENSMUSG( | 0.401732 | 3.70E-06 | YES | NO | NA |
| Limk2 | ENSMUSG( | ENSMUSG( | -0.21633 | 0.000172 | YES | NO | NA |
| Sel1l | ENSMUSG( | ENSMUSG( | -0.10225 | 0.207572 | NO | NO | NA |
| Itpr2 | ENSMUSG( | ENSMUSG( | -0.72975 | 1.00E-10 | YES | NO | NA |
| Itgam | ENSMUSG( | ENSMUSG( | -2.01 | 3.90E-09 | YES | YES | Downregulated |
| Elob | ENSMUSG( | ENSMUSG( | 0.087303 | 0.350806 | NO | NO | NA |
| Chordc1 | ENSMUSG( | ENSMUSG( | -0.50514 | 7.26E-08 | YES | NO | NA |
| Tomm6 | ENSMUSG( | ENSMUSG( | 0.11476 | 0.128224 | NO | NO | NA |
| Nup188 | ENSMUSG( | ENSMUSG( | 0.095147 | 0.054903 | NO | NO | NA |
| Lpcat4 | ENSMUSG( | ENSMUSG( | -0.20763 | 0.041631 | YES | NO | NA |
| Klc1 | ENSMUSG( | ENSMUSG( | -0.03759 | 0.604671 | NO | NO | NA |
| Mbd3 | ENSMUSG( | ENSMUSG( | 0.013301 | 0.859322 | NO | NO | NA |
| Acp5 | ENSMUSG( | ENSMUSG( | -0.39381 | 8.30E-13 | YES | NO | NA |
| Smad3 | ENSMUSG( | ENSMUSG( | -0.58915 | 4.84E-16 | YES | NO | NA |
| Pacsin2 | ENSMUSG( | ENSMUSG( | 0.00025 | 0.997466 | NO | NO | NA |
| Scaf8 | ENSMUSG( | ENSMUSG( | -0.25472 | 1.89E-07 | YES | NO | NA |
| Aftph | ENSMUSG( | ENSMUSG( | -0.29924 | 0.000791 | YES | NO | NA |
| Sik3 | ENSMUSG( | ENSMUSG( | -0.2669 | 0.00111 | YES | NO | NA |
| Smap1 | ENSMUSG( | ENSMUSG( | -0.25275 | 0.000748 | YES | NO | NA |
| Trp53inp1 | ENSMUSG( | ENSMUSG( | -1.03486 | 4.95E-11 | YES | YES | Downregulated |

| | | | | | | | |
|---|---|---|---|---|---|---|---|
| Qrich1 | ENSMUSG( | ENSMUSG( | -0.06276 | 0.280628 | NO | NO | NA |
| Kif4 | ENSMUSG( | ENSMUSG( | 0.221653 | 0.002343 | YES | NO | NA |
| Sh2b1 | ENSMUSG( | ENSMUSG( | -0.25232 | 5.35E-06 | YES | NO | NA |
| Scrib | ENSMUSG( | ENSMUSG( | -0.26713 | 3.77E-05 | YES | NO | NA |
| Adprh | ENSMUSG( | ENSMUSG( | -0.02862 | 0.665437 | NO | NO | NA |
| Slfn1 | ENSMUSG( | ENSMUSG( | -0.8325 | 1.87E-14 | YES | NO | NA |
| Rhot2 | ENSMUSG( | ENSMUSG( | -0.01968 | 0.7956 | NO | NO | NA |
| Nfatc2 | ENSMUSG( | ENSMUSG( | -0.42438 | 4.04E-10 | YES | NO | NA |
| Tfrc | ENSMUSG( | ENSMUSG( | 0.20115 | 0.073612 | NO | NO | NA |
| Clk2 | ENSMUSG( | ENSMUSG( | -0.17994 | 0.035298 | YES | NO | NA |
| Igtp | ENSMUSG( | ENSMUSG( | -0.32762 | 0.000176 | YES | NO | NA |
| Atp5e | ENSMUSG( | ENSMUSG( | -0.05968 | 0.203863 | NO | NO | NA |
| Tars | ENSMUSG( | ENSMUSG( | 0.142674 | 0.141701 | NO | NO | NA |
| Sla2 | ENSMUSG( | ENSMUSG( | -0.34608 | 2.91E-05 | YES | NO | NA |
| Cox7c | ENSMUSG( | ENSMUSG( | 0.162453 | 0.031517 | YES | NO | NA |
| Gabarap | ENSMUSG( | ENSMUSG( | -0.25674 | 0.000198 | YES | NO | NA |
| Cip2a | ENSMUSG( | ENSMUSG( | 0.198324 | 0.032779 | YES | NO | NA |
| Fzr1 | ENSMUSG( | ENSMUSG( | 0.091239 | 0.061772 | NO | NO | NA |
| Irf8 | ENSMUSG( | ENSMUSG( | -0.00418 | 0.970155 | NO | NO | NA |
| Ubl3 | ENSMUSG( | ENSMUSG( | -0.10067 | 0.19671 | NO | NO | NA |
| Impa1 | ENSMUSG( | ENSMUSG( | -0.10338 | 0.052799 | NO | NO | NA |
| Afg3l1 | ENSMUSG( | ENSMUSG( | 0.181606 | 0.000352 | YES | NO | NA |
| Tomm20 | ENSMUSG( | ENSMUSG( | 0.30611 | 3.99E-06 | YES | NO | NA |
| Ist1 | ENSMUSG( | ENSMUSG( | -0.14518 | 0.078227 | NO | NO | NA |
| Mtrex | ENSMUSG( | ENSMUSG( | -0.03223 | 0.679095 | NO | NO | NA |
| Mfap1b | ENSMUSG( | ENSMUSG( | -0.30084 | 0.005018 | YES | NO | NA |
| Sike1 | ENSMUSG( | ENSMUSG( | -0.40424 | 2.75E-07 | YES | NO | NA |
| Pan3 | ENSMUSG( | ENSMUSG( | -0.29104 | 0.00013 | YES | NO | NA |
| Rilpl2 | ENSMUSG( | ENSMUSG( | 0.252017 | 0.026185 | YES | NO | NA |
| Senp1 | ENSMUSG( | ENSMUSG( | -0.04135 | 0.424867 | NO | NO | NA |
| Wwp2 | ENSMUSG( | ENSMUSG( | -0.01573 | 0.78461 | NO | NO | NA |
| Golph3 | ENSMUSG( | ENSMUSG( | -0.04951 | 0.592341 | NO | NO | NA |
| Ltn1 | ENSMUSG( | ENSMUSG( | -0.09106 | 0.202765 | NO | NO | NA |
| Ak3 | ENSMUSG( | ENSMUSG( | -0.41624 | 2.45E-10 | YES | NO | NA |
| Prkag1 | ENSMUSG( | ENSMUSG( | 0.187266 | 0.003152 | YES | NO | NA |
| Tut4 | ENSMUSG( | ENSMUSG( | -0.51787 | 3.32E-08 | YES | NO | NA |
| Ppp2cb | ENSMUSG( | ENSMUSG( | 0.124885 | 0.043475 | YES | NO | NA |
| Sbk1 | ENSMUSG( | ENSMUSG( | -0.53652 | 1.85E-05 | YES | NO | NA |
| Ndufv1 | ENSMUSG( | ENSMUSG( | 0.627231 | 3.96E-09 | YES | NO | NA |
| Dennd4b | ENSMUSG( | ENSMUSG( | -0.55911 | 2.06E-11 | YES | NO | NA |
| Hnrnph2 | ENSMUSG( | ENSMUSG( | -0.02472 | 0.838946 | NO | NO | NA |
| Acot7 | ENSMUSG( | ENSMUSG( | -0.42764 | 1.20E-06 | YES | NO | NA |
| Crlf2 | ENSMUSG( | ENSMUSG( | 0.226712 | 0.175984 | NO | NO | NA |
| Psmc5 | ENSMUSG( | ENSMUSG( | 0.29403 | 1.25E-05 | YES | NO | NA |
| Cdt1 | ENSMUSG( | ENSMUSG( | 0.168775 | 0.000132 | YES | NO | NA |
| Gbf1 | ENSMUSG( | ENSMUSG( | -0.28944 | 3.81E-07 | YES | NO | NA |
| Atp10a | ENSMUSG( | ENSMUSG( | 0.032213 | 0.677884 | NO | NO | NA |

| | | | | | | | |
|---|---|---|---|---|---|---|---|
| Ppp3cc | ENSMUSG( | ENSMUSG( | 0.089573 | 0.321293 | NO | NO | NA |
| Spag9 | ENSMUSG( | ENSMUSG( | -0.10438 | 0.229729 | NO | NO | NA |
| Dnajc1 | ENSMUSG( | ENSMUSG( | -0.20109 | 0.002546 | YES | NO | NA |
| Mxd4 | ENSMUSG( | ENSMUSG( | -0.69894 | 8.26E-06 | YES | NO | NA |
| Tm9sf2 | ENSMUSG( | ENSMUSG( | -0.04191 | 0.562687 | NO | NO | NA |
| Gm2682 | ENSMUSG( | ENSMUSG( | -1.32497 | 1.09E-17 | YES | YES | Downregulated |
| Dcaf8 | ENSMUSG( | ENSMUSG( | -0.03049 | 0.655535 | NO | NO | NA |
| Rcc1 | ENSMUSG( | ENSMUSG( | 0.559834 | 4.49E-10 | YES | NO | NA |
| Eif1ax | ENSMUSG( | ENSMUSG( | 0.312934 | 0.000983 | YES | NO | NA |
| Zdhhc3 | ENSMUSG( | ENSMUSG( | 0.124806 | 0.017691 | YES | NO | NA |
| Ptpn12 | ENSMUSG( | ENSMUSG( | -0.04685 | 0.553915 | NO | NO | NA |
| Irf7 | ENSMUSG( | ENSMUSG( | -0.85164 | 2.82E-17 | YES | NO | NA |
| Nup107 | ENSMUSG( | ENSMUSG( | 0.3615 | 1.06E-05 | YES | NO | NA |
| Lonp1 | ENSMUSG( | ENSMUSG( | 0.617754 | 9.54E-08 | YES | NO | NA |
| Gsr | ENSMUSG( | ENSMUSG( | 0.195731 | 0.005881 | YES | NO | NA |
| Arhgap1 | ENSMUSG( | ENSMUSG( | -0.12989 | 0.009422 | YES | NO | NA |
| Ltb | ENSMUSG( | ENSMUSG( | -0.27697 | 0.012837 | YES | NO | NA |
| Tnpo2 | ENSMUSG( | ENSMUSG( | -0.06148 | 0.422267 | NO | NO | NA |
| Arid4a | ENSMUSG( | ENSMUSG( | -0.39233 | 4.49E-05 | YES | NO | NA |
| Gga1 | ENSMUSG( | ENSMUSG( | -0.07055 | 0.217871 | NO | NO | NA |
| Kctd10 | ENSMUSG( | ENSMUSG( | -0.07107 | 0.458761 | NO | NO | NA |
| Stard7 | ENSMUSG( | ENSMUSG( | 0.208762 | 0.001241 | YES | NO | NA |
| Actl6a | ENSMUSG( | ENSMUSG( | 0.174344 | 0.021977 | YES | NO | NA |
| Zfp36l1 | ENSMUSG( | ENSMUSG( | -0.33138 | 0.015202 | YES | NO | NA |
| Cope | ENSMUSG( | ENSMUSG( | 0.113881 | 0.140403 | NO | NO | NA |
| Cd160 | ENSMUSG( | ENSMUSG( | 0.129809 | 0.211065 | NO | NO | NA |
| Dpy19l1 | ENSMUSG( | ENSMUSG( | 0.10335 | 0.257728 | NO | NO | NA |
| Il27ra | ENSMUSG( | ENSMUSG( | -0.48273 | 9.51E-24 | YES | NO | NA |
| Ppp4r2 | ENSMUSG( | ENSMUSG( | -0.23725 | 0.000102 | YES | NO | NA |
| Rtf2 | ENSMUSG( | ENSMUSG( | -0.00459 | 0.961317 | NO | NO | NA |
| Phb | ENSMUSG( | ENSMUSG( | 0.51228 | 6.66E-07 | YES | NO | NA |
| Nudt4 | ENSMUSG( | ENSMUSG( | 0.162037 | 0.000876 | YES | NO | NA |
| Endod1 | ENSMUSG( | ENSMUSG( | 0.100411 | 0.354681 | NO | NO | NA |
| Sec63 | ENSMUSG( | ENSMUSG( | 0.121829 | 0.309494 | NO | NO | NA |
| Dync1i2 | ENSMUSG( | ENSMUSG( | -0.23353 | 0.000825 | YES | NO | NA |
| Gabbr1 | ENSMUSG( | ENSMUSG( | -0.51478 | 6.10E-09 | YES | NO | NA |
| Arpp19 | ENSMUSG( | ENSMUSG( | 0.146979 | 0.337668 | NO | NO | NA |
| Hars | ENSMUSG( | ENSMUSG( | 0.204512 | 0.007016 | YES | NO | NA |
| Kdm6a | ENSMUSG( | ENSMUSG( | -0.19678 | 0.008372 | YES | NO | NA |
| Sidt1 | ENSMUSG( | ENSMUSG( | -0.74591 | 2.00E-06 | YES | NO | NA |
| Msi2 | ENSMUSG( | ENSMUSG( | -0.39216 | 0.000655 | YES | NO | NA |
| Pglyrp1 | ENSMUSG( | ENSMUSG( | -0.39016 | 6.58E-07 | YES | NO | NA |
| Nploc4 | ENSMUSG( | ENSMUSG( | 0.082802 | 0.133409 | NO | NO | NA |
| Plaa | ENSMUSG( | ENSMUSG( | 0.10498 | 0.159685 | NO | NO | NA |
| Nomo1 | ENSMUSG( | ENSMUSG( | 0.448456 | 2.63E-06 | YES | NO | NA |
| Crot | ENSMUSG( | ENSMUSG( | -0.15199 | 0.136714 | NO | NO | NA |
| Grap | ENSMUSG( | ENSMUSG( | -0.27217 | 0.0005 | YES | NO | NA |

| | | | | | | | |
|---|---|---|---|---|---|---|---|
| Rer1 | ENSMUSG( | ENSMUSG( | -0.0084 | 0.910261 | NO | NO | NA |
| Adora2a | ENSMUSG( | ENSMUSG( | -0.24989 | 3.99E-05 | YES | NO | NA |
| Vps16 | ENSMUSG( | ENSMUSG( | 0.085099 | 0.241726 | NO | NO | NA |
| Xpo7 | ENSMUSG( | ENSMUSG( | -0.00263 | 0.970413 | NO | NO | NA |
| Wars | ENSMUSG( | ENSMUSG( | 0.140878 | 0.12036 | NO | NO | NA |
| Chmp1a | ENSMUSG( | ENSMUSG( | 0.038961 | 0.705171 | NO | NO | NA |
| Nat10 | ENSMUSG( | ENSMUSG( | -0.04525 | 0.473659 | NO | NO | NA |
| Acap2 | ENSMUSG( | ENSMUSG( | -0.26634 | 0.000325 | YES | NO | NA |
| Man1a | ENSMUSG( | ENSMUSG( | -0.47375 | 2.30E-07 | YES | NO | NA |
| Brip1os | ENSMUSG( | ENSMUSG( | -0.07177 | 0.132494 | NO | NO | NA |
| Sh3bgrl | ENSMUSG( | ENSMUSG( | -0.01594 | 0.897257 | NO | NO | NA |
| Wbp2 | ENSMUSG( | ENSMUSG( | -0.58479 | 5.59E-12 | YES | NO | NA |
| Kras | ENSMUSG( | ENSMUSG( | -0.03572 | 0.613046 | NO | NO | NA |
| Man2a2 | ENSMUSG( | ENSMUSG( | 0.232064 | 3.14E-06 | YES | NO | NA |
| Slc2a1 | ENSMUSG( | ENSMUSG( | 0.156922 | 0.322745 | NO | NO | NA |
| Prpf6 | ENSMUSG( | ENSMUSG( | 0.039541 | 0.595771 | NO | NO | NA |
| Rad51 | ENSMUSG( | ENSMUSG( | 0.289947 | 0.00083 | YES | NO | NA |
| Serinc1 | ENSMUSG( | ENSMUSG( | -0.27299 | 0.001082 | YES | NO | NA |
| Dop1b | ENSMUSG( | ENSMUSG( | -0.5576 | 7.62E-10 | YES | NO | NA |
| Akap9 | ENSMUSG( | ENSMUSG( | -0.2451 | 0.003696 | YES | NO | NA |
| Midn | ENSMUSG( | ENSMUSG( | -0.23802 | 0.000398 | YES | NO | NA |
| Pola2 | ENSMUSG( | ENSMUSG( | 0.289892 | 8.65E-06 | YES | NO | NA |
| Ehd3 | ENSMUSG( | ENSMUSG( | -0.07695 | 0.499294 | NO | NO | NA |
| Slain2 | ENSMUSG( | ENSMUSG( | -0.09242 | 0.219148 | NO | NO | NA |
| Chaserr | ENSMUSG( | ENSMUSG( | -0.40137 | 3.70E-07 | YES | NO | NA |
| Map3k11 | ENSMUSG( | ENSMUSG( | -0.20735 | 6.57E-05 | YES | NO | NA |
| Pmaip1 | ENSMUSG( | ENSMUSG( | 0.066623 | 0.679922 | NO | NO | NA |
| Hdgfl2 | ENSMUSG( | ENSMUSG( | 0.018391 | 0.840237 | NO | NO | NA |
| Ndufb11 | ENSMUSG( | ENSMUSG( | 0.208348 | 0.001982 | YES | NO | NA |
| Cst7 | ENSMUSG( | ENSMUSG( | -0.59496 | 6.59E-09 | YES | NO | NA |
| Sart3 | ENSMUSG( | ENSMUSG( | -0.13449 | 0.094237 | NO | NO | NA |
| Prkca | ENSMUSG( | ENSMUSG( | -0.3915 | 1.18E-06 | YES | NO | NA |
| Ptges3 | ENSMUSG( | ENSMUSG( | -0.02457 | 0.773412 | NO | NO | NA |
| Prdx2 | ENSMUSG( | ENSMUSG( | 0.286533 | 1.31E-05 | YES | NO | NA |
| Hspa1b | ENSMUSG( | ENSMUSG( | -2.97986 | 0.001395 | YES | YES | Downregulated |
| Derl1 | ENSMUSG( | ENSMUSG( | -0.1109 | 0.036828 | YES | NO | NA |
| Atad1 | ENSMUSG( | ENSMUSG( | 0.031653 | 0.727172 | NO | NO | NA |
| Scaf4 | ENSMUSG( | ENSMUSG( | -0.14772 | 0.005316 | YES | NO | NA |
| Kri1 | ENSMUSG( | ENSMUSG( | 0.141095 | 0.185456 | NO | NO | NA |
| Rad54l2 | ENSMUSG( | ENSMUSG( | 0.050362 | 0.600449 | NO | NO | NA |
| Bnip3l | ENSMUSG( | ENSMUSG( | -0.28965 | 0.014023 | YES | NO | NA |
| Batf | ENSMUSG( | ENSMUSG( | 0.069227 | 0.579897 | NO | NO | NA |
| Tmem254 | ENSMUSG( | ENSMUSG( | -0.08447 | 0.424857 | NO | NO | NA |
| Acsbg1 | ENSMUSG( | ENSMUSG( | -0.62881 | 4.37E-07 | YES | NO | NA |
| Crlf3 | ENSMUSG( | ENSMUSG( | -0.31444 | 1.09E-08 | YES | NO | NA |
| Pdcd11 | ENSMUSG( | ENSMUSG( | 0.196931 | 0.040573 | YES | NO | NA |
| Nelfb | ENSMUSG( | ENSMUSG( | -0.23589 | 2.21E-06 | YES | NO | NA |

| | | | | | | | |
|---|---|---|---|---|---|---|---|
| Snrpd2 | ENSMUSG( | ENSMUSG( | 0.504557 | 3.53E-09 | YES | NO | NA |
| Pgrmc1 | ENSMUSG( | ENSMUSG( | 0.342961 | 4.24E-06 | YES | NO | NA |
| Cul1 | ENSMUSG( | ENSMUSG( | 0.214324 | 0.001534 | YES | NO | NA |
| Ccdc47 | ENSMUSG( | ENSMUSG( | 0.070583 | 0.223305 | NO | NO | NA |
| Tecr | ENSMUSG( | ENSMUSG( | 0.009655 | 0.878987 | NO | NO | NA |
| Cyb5r4 | ENSMUSG( | ENSMUSG( | 0.00996 | 0.92214 | NO | NO | NA |
| Blmh | ENSMUSG( | ENSMUSG( | -0.00959 | 0.871801 | NO | NO | NA |
| Sppl2a | ENSMUSG( | ENSMUSG( | -0.15228 | 0.012065 | YES | NO | NA |
| Msh6 | ENSMUSG( | ENSMUSG( | 0.141042 | 0.092712 | NO | NO | NA |
| Snx4 | ENSMUSG( | ENSMUSG( | 0.019305 | 0.825294 | NO | NO | NA |
| Pes1 | ENSMUSG( | ENSMUSG( | 0.405468 | 8.07E-05 | YES | NO | NA |
| Exoc2 | ENSMUSG( | ENSMUSG( | -0.39434 | 5.52E-07 | YES | NO | NA |
| U2af1 | ENSMUSG( | ENSMUSG( | 0.201501 | 0.026692 | YES | NO | NA |
| Sephs1 | ENSMUSG( | ENSMUSG( | 0.223301 | 0.00604 | YES | NO | NA |
| Smu1 | ENSMUSG( | ENSMUSG( | 0.050246 | 0.355386 | NO | NO | NA |
| Gm49336 | ENSMUSG( | ENSMUSG( | 0.022174 | 0.777177 | NO | NO | NA |
| Ptp4a1 EN | ENSMUSG( | ENSMUSG( | -0.14081 | 0.358484 | NO | NO | NA |
| Ppfia1 | ENSMUSG( | ENSMUSG( | -0.04059 | 0.540149 | NO | NO | NA |
| Arhgap26 | ENSMUSG( | ENSMUSG( | -0.06206 | 0.594123 | NO | NO | NA |
| Zdhhc5 | ENSMUSG( | ENSMUSG( | -0.15853 | 0.002077 | YES | NO | NA |
| Egr1 | ENSMUSG( | ENSMUSG( | 0.381689 | 0.295743 | NO | NO | NA |
| Usf3 | ENSMUSG( | ENSMUSG( | -0.07304 | 0.424867 | NO | NO | NA |
| Osbp | ENSMUSG( | ENSMUSG( | -0.11416 | 0.062979 | NO | NO | NA |
| Tmx3 | ENSMUSG( | ENSMUSG( | -0.25569 | 0.002164 | YES | NO | NA |
| Ing1 | ENSMUSG( | ENSMUSG( | -0.09045 | 0.308111 | NO | NO | NA |
| Rbpj | ENSMUSG( | ENSMUSG( | 0.51541 | 1.84E-08 | YES | NO | NA |
| Marchf7 | ENSMUSG( | ENSMUSG( | -0.07004 | 0.53935 | NO | NO | NA |
| Cyth1 | ENSMUSG( | ENSMUSG( | -0.36319 | 9.73E-06 | YES | NO | NA |
| H2az2 | ENSMUSG( | ENSMUSG( | 0.019796 | 0.840512 | NO | NO | NA |
| Zfp598 | ENSMUSG( | ENSMUSG( | 0.065531 | 0.135872 | NO | NO | NA |
| Copz1 | ENSMUSG( | ENSMUSG( | -0.06459 | 0.475251 | NO | NO | NA |
| Ensa | ENSMUSG( | ENSMUSG( | -0.13149 | 0.025279 | YES | NO | NA |
| Zfp706 | ENSMUSG( | ENSMUSG( | -0.0864 | 0.153535 | NO | NO | NA |
| Smpdl3b | ENSMUSG( | ENSMUSG( | 0.19137 | 0.233182 | NO | NO | NA |
| Cr1l | ENSMUSG( | ENSMUSG( | -0.1873 | 0.021565 | YES | NO | NA |
| Pced1b | ENSMUSG( | ENSMUSG( | -0.74887 | 1.01E-14 | YES | NO | NA |
| Arhgap25 | ENSMUSG( | ENSMUSG( | -0.37066 | 1.45E-05 | YES | NO | NA |
| Otulinl | ENSMUSG( | ENSMUSG( | -0.24587 | 1.65E-05 | YES | NO | NA |
| Ppp5c | ENSMUSG( | ENSMUSG( | 0.140277 | 0.032685 | YES | NO | NA |
| Kctd20 | ENSMUSG( | ENSMUSG( | -0.06251 | 0.094124 | NO | NO | NA |
| Cdk1 | ENSMUSG( | ENSMUSG( | 0.190752 | 2.62E-05 | YES | NO | NA |
| Bscl2 | ENSMUSG( | ENSMUSG( | -0.11912 | 0.160392 | NO | NO | NA |
| Fam3c | ENSMUSG( | ENSMUSG( | 0.160174 | 0.008235 | YES | NO | NA |
| Nr2c2 | ENSMUSG( | ENSMUSG( | -0.27137 | 0.00063 | YES | NO | NA |
| Tbc1d14 | ENSMUSG( | ENSMUSG( | -0.08904 | 0.510933 | NO | NO | NA |
| Psmb5 | ENSMUSG( | ENSMUSG( | 0.221543 | 0.000513 | YES | NO | NA |
| Cnot9 | ENSMUSG( | ENSMUSG( | 0.280756 | 3.62E-05 | YES | NO | NA |

| | | | | | | | |
|---|---|---|---|---|---|---|---|
| Luc7l | ENSMUSG( | ENSMUSG( | 0.002515 | 0.967691 | NO | NO | NA |
| Rbm22 | ENSMUSG( | ENSMUSG( | -0.0027 | 0.976785 | NO | NO | NA |
| Ndufb7 | ENSMUSG( | ENSMUSG( | 0.042981 | 0.645714 | NO | NO | NA |
| Mon2 | ENSMUSG( | ENSMUSG( | -0.28816 | 4.43E-06 | YES | NO | NA |
| Ddx47 | ENSMUSG( | ENSMUSG( | 0.076112 | 0.274032 | NO | NO | NA |
| Nrip1 | ENSMUSG( | ENSMUSG( | -0.33492 | 0.000811 | YES | NO | NA |
| Tial1 | ENSMUSG( | ENSMUSG( | -0.04926 | 0.25122 | NO | NO | NA |
| Tmem71 | ENSMUSG( | ENSMUSG( | -0.50394 | 3.26E-06 | YES | NO | NA |
| Trim33 | ENSMUSG( | ENSMUSG( | -0.20408 | 0.008236 | YES | NO | NA |
| Slmap | ENSMUSG( | ENSMUSG( | -0.26745 | 0.000222 | YES | NO | NA |
| Tmf1 | ENSMUSG( | ENSMUSG( | -0.18056 | 0.027328 | YES | NO | NA |
| Ythdf3 | ENSMUSG( | ENSMUSG( | -0.344 | 2.84E-05 | YES | NO | NA |
| Srgap2 | ENSMUSG( | ENSMUSG( | -0.29517 | 5.13E-05 | YES | NO | NA |
| Abcb9 | ENSMUSG( | ENSMUSG( | -0.39484 | 4.23E-05 | YES | NO | NA |
| Idh3g | ENSMUSG( | ENSMUSG( | 0.259598 | 0.00216 | YES | NO | NA |
| Xab2 | ENSMUSG( | ENSMUSG( | 0.037064 | 0.669471 | NO | NO | NA |
| 1110004F: | ENSMUSG( | ENSMUSG( | 0.246932 | 0.006349 | YES | NO | NA |
| Rnf220 | ENSMUSG( | ENSMUSG( | -0.01097 | 0.860489 | NO | NO | NA |
| Cyc1 | ENSMUSG( | ENSMUSG( | 0.332514 | 3.92E-07 | YES | NO | NA |
| Vps35 | ENSMUSG( | ENSMUSG( | 0.002659 | 0.972758 | NO | NO | NA |
| Snrnp40 | ENSMUSG( | ENSMUSG( | 0.307855 | 1.77E-06 | YES | NO | NA |
| Kpna3 | ENSMUSG( | ENSMUSG( | 0.114359 | 0.141586 | NO | NO | NA |
| Exosc10 | ENSMUSG( | ENSMUSG( | 0.168031 | 0.015273 | YES | NO | NA |
| Zcchc8 | ENSMUSG( | ENSMUSG( | 0.014846 | 0.83522 | NO | NO | NA |
| Tsnax | ENSMUSG( | ENSMUSG( | 0.261641 | 5.78E-05 | YES | NO | NA |
| Npepl1 | ENSMUSG( | ENSMUSG( | 0.269768 | 0.000475 | YES | NO | NA |
| Rad50 | ENSMUSG( | ENSMUSG( | 0.104427 | 0.285481 | NO | NO | NA |
| Uqcrfs1 | ENSMUSG( | ENSMUSG( | 0.318656 | 8.93E-05 | YES | NO | NA |
| Mtmr12 | ENSMUSG( | ENSMUSG( | -0.28346 | 0.000542 | YES | NO | NA |
| Ric8a | ENSMUSG( | ENSMUSG( | 0.018586 | 0.874608 | NO | NO | NA |
| Inpp1 | ENSMUSG( | ENSMUSG( | -0.5162 | 5.33E-08 | YES | NO | NA |
| ENSMUSG( | ENSMUSG( | ENSMUSG( | -0.58079 | 2.94E-07 | YES | NO | NA |
| Ilf2 | ENSMUSG( | ENSMUSG( | 0.154291 | 0.061851 | NO | NO | NA |
| Arf4 | ENSMUSG( | ENSMUSG( | -0.39847 | 2.28E-07 | YES | NO | NA |
| Kansl3 | ENSMUSG( | ENSMUSG( | -0.15647 | 0.011988 | YES | NO | NA |
| Ints7 | ENSMUSG( | ENSMUSG( | 0.038185 | 0.439554 | NO | NO | NA |
| Rnf2 | ENSMUSG( | ENSMUSG( | -0.83886 | 9.91E-07 | YES | NO | NA |
| Golga2 | ENSMUSG( | ENSMUSG( | -0.03926 | 0.624845 | NO | NO | NA |
| Ap2s1 | ENSMUSG( | ENSMUSG( | 0.074526 | 0.12348 | NO | NO | NA |
| Hmgb3 | ENSMUSG( | ENSMUSG( | 0.796675 | 9.81E-09 | YES | NO | NA |
| Slu7 | ENSMUSG( | ENSMUSG( | -0.15078 | 0.135252 | NO | NO | NA |
| Srp54a | ENSMUSG( | ENSMUSG( | -0.0016 | 0.987651 | NO | NO | NA |
| Slc39a7 | ENSMUSG( | ENSMUSG( | -0.06971 | 0.476823 | NO | NO | NA |
| Ptch1 | ENSMUSG( | ENSMUSG( | -0.07306 | 0.439866 | NO | NO | NA |
| Ctdsp2 | ENSMUSG( | ENSMUSG( | -0.43558 | 0.000191 | YES | NO | NA |
| Ehd1 | ENSMUSG( | ENSMUSG( | 0.038963 | 0.672019 | NO | NO | NA |
| Gpbp1l1 | ENSMUSG( | ENSMUSG( | -0.31728 | 0.000249 | YES | NO | NA |

| | | | | | | | |
|---|---|---|---|---|---|---|---|
| Casp8ap2 | ENSMUSG( | ENSMUSG( | -0.08247 | 0.364689 | NO | NO | NA |
| Psme4 | ENSMUSG( | ENSMUSG( | 0.014497 | 0.890997 | NO | NO | NA |
| Lpgat1 | ENSMUSG( | ENSMUSG( | 0.211112 | 0.017701 | YES | NO | NA |
| Tspyl1 | ENSMUSG( | ENSMUSG( | -0.2177 | 0.007473 | YES | NO | NA |
| Dennd4c | ENSMUSG( | ENSMUSG( | -0.56627 | 1.98E-14 | YES | NO | NA |
| Tshz1 | ENSMUSG( | ENSMUSG( | -0.08263 | 0.307355 | NO | NO | NA |
| Ppm1a | ENSMUSG( | ENSMUSG( | -0.01672 | 0.796198 | NO | NO | NA |
| Trim44 | ENSMUSG( | ENSMUSG( | 0.08412 | 0.377977 | NO | NO | NA |
| Slfn8 | ENSMUSG( | ENSMUSG( | -0.97685 | 3.07E-13 | YES | NO | NA |
| Sfxn1 | ENSMUSG( | ENSMUSG( | 0.532273 | 1.46E-09 | YES | NO | NA |
| Chst11 | ENSMUSG( | ENSMUSG( | -0.12994 | 0.212967 | NO | NO | NA |
| Sec23ip | ENSMUSG( | ENSMUSG( | -0.05088 | 0.216209 | NO | NO | NA |
| Hectd3 | ENSMUSG( | ENSMUSG( | -0.16331 | 0.015141 | YES | NO | NA |
| Aggf1 | ENSMUSG( | ENSMUSG( | -0.10059 | 0.14938 | NO | NO | NA |
| Vps36 | ENSMUSG( | ENSMUSG( | 0.030291 | 0.733043 | NO | NO | NA |
| Rbm6 | ENSMUSG( | ENSMUSG( | -0.31705 | 0.000402 | YES | NO | NA |
| Ppip5k2 | ENSMUSG( | ENSMUSG( | -0.07057 | 0.399916 | NO | NO | NA |
| Icam1 | ENSMUSG( | ENSMUSG( | -0.70082 | 1.01E-12 | YES | NO | NA |
| Ube2k | ENSMUSG( | ENSMUSG( | -0.0605 | 0.758216 | NO | NO | NA |
| Dbf4 | ENSMUSG( | ENSMUSG( | 0.133342 | 0.106121 | NO | NO | NA |
| Snrpd3 | ENSMUSG( | ENSMUSG( | 0.237092 | 0.016979 | YES | NO | NA |
| Zcchc2 | ENSMUSG( | ENSMUSG( | -0.59814 | 2.81E-15 | YES | NO | NA |
| Cops2 | ENSMUSG( | ENSMUSG( | 0.214035 | 0.014288 | YES | NO | NA |
| Nup85 | ENSMUSG( | ENSMUSG( | 0.37104 | 8.54E-07 | YES | NO | NA |
| Vps28 | ENSMUSG( | ENSMUSG( | -0.17286 | 8.10E-05 | YES | NO | NA |
| Nup88 | ENSMUSG( | ENSMUSG( | 0.078477 | 0.076023 | NO | NO | NA |
| Iqsec1 | ENSMUSG( | ENSMUSG( | -0.46892 | 1.71E-06 | YES | NO | NA |
| Nrbp1 | ENSMUSG( | ENSMUSG( | -0.05736 | 0.493188 | NO | NO | NA |
| Rars | ENSMUSG( | ENSMUSG( | 0.31194 | 0.004734 | YES | NO | NA |
| Dr1 | ENSMUSG( | ENSMUSG( | -0.10762 | 0.052467 | NO | NO | NA |
| Mfap1a | ENSMUSG( | ENSMUSG( | -0.21546 | 0.007236 | YES | NO | NA |
| H2-Q5 | ENSMUSG( | ENSMUSG( | -0.47962 | 9.42E-06 | YES | NO | NA |
| ENSMUSG( | ENSMUSG( | ENSMUSG( | 0.102827 | 0.45373 | NO | NO | NA |
| Ripk1 | ENSMUSG( | ENSMUSG( | -0.18769 | 0.012999 | YES | NO | NA |
| Edc4 | ENSMUSG( | ENSMUSG( | -0.0181 | 0.786782 | NO | NO | NA |
| Rnf31 | ENSMUSG( | ENSMUSG( | -0.43678 | 2.74E-15 | YES | NO | NA |
| Cbx6 | ENSMUSG( | ENSMUSG( | -0.24893 | 0.000341 | YES | NO | NA |
| R3hdm2 | ENSMUSG( | ENSMUSG( | -0.37836 | 6.34E-07 | YES | NO | NA |
| Dgcr8 | ENSMUSG( | ENSMUSG( | 0.113451 | 0.13319 | NO | NO | NA |
| Vcpip1 | ENSMUSG( | ENSMUSG( | -0.46759 | 7.58E-13 | YES | NO | NA |
| Dcp2 | ENSMUSG( | ENSMUSG( | -0.08211 | 0.219229 | NO | NO | NA |
| Secisbp2l | ENSMUSG( | ENSMUSG( | -0.36229 | 1.10E-07 | YES | NO | NA |
| Hmgn5 | ENSMUSG( | ENSMUSG( | 0.157814 | 0.483231 | NO | NO | NA |
| Gm20605 | ENSMUSG( | ENSMUSG( | -0.10373 | 0.293318 | NO | NO | NA |
| Elavl1 | ENSMUSG( | ENSMUSG( | -0.14569 | 0.001746 | YES | NO | NA |
| Gm20559 | ENSMUSG( | ENSMUSG( | -0.91981 | 4.22E-14 | YES | NO | NA |
| Mogs | ENSMUSG( | ENSMUSG( | 0.314108 | 0.000353 | YES | NO | NA |

| | | | | | | | |
|---|---|---|---|---|---|---|---|
| Nudt3 | ENSMUSG( | ENSMUSG( | -0.07088 | 0.367293 | NO | NO | NA |
| Rbm14 | ENSMUSG( | ENSMUSG( | -0.11816 | 0.110015 | NO | NO | NA |
| C1qbp | ENSMUSG( | ENSMUSG( | 0.79027 | 2.61E-08 | YES | NO | NA |
| Ndufa10 | ENSMUSG( | ENSMUSG( | 0.392833 | 1.55E-06 | YES | NO | NA |
| Dennd4a | ENSMUSG( | ENSMUSG( | -0.13964 | 0.025527 | YES | NO | NA |
| Golga7 | ENSMUSG( | ENSMUSG( | -0.06773 | 0.294794 | NO | NO | NA |
| R3hdm1 | ENSMUSG( | ENSMUSG( | -0.04701 | 0.406744 | NO | NO | NA |
| Ubl5 | ENSMUSG( | ENSMUSG( | -0.09147 | 0.269521 | NO | NO | NA |
| Zfp622 | ENSMUSG( | ENSMUSG( | -0.05835 | 0.555485 | NO | NO | NA |
| Tia1 | ENSMUSG( | ENSMUSG( | -0.00588 | 0.970155 | NO | NO | NA |
| Retreg3 | ENSMUSG( | ENSMUSG( | -0.35569 | 1.52E-07 | YES | NO | NA |
| Tbl1x | ENSMUSG( | ENSMUSG( | -0.12696 | 0.057166 | NO | NO | NA |
| Ikzf2 | ENSMUSG( | ENSMUSG( | -0.8616 | 0.052799 | NO | NO | NA |
| Pafah1b2 | ENSMUSG( | ENSMUSG( | -0.03254 | 0.640175 | NO | NO | NA |
| Aamp | ENSMUSG( | ENSMUSG( | -0.10377 | 0.051248 | NO | NO | NA |
| Rnf19b | ENSMUSG( | ENSMUSG( | -0.43547 | 1.26E-05 | YES | NO | NA |
| Dad1 | ENSMUSG( | ENSMUSG( | 0.110227 | 0.113852 | NO | NO | NA |
| Zfp638 | ENSMUSG( | ENSMUSG( | -0.47759 | 6.00E-11 | YES | NO | NA |
| Gxylt1 | ENSMUSG( | ENSMUSG( | -0.40337 | 3.26E-09 | YES | NO | NA |
| Rnf38 | ENSMUSG( | ENSMUSG( | -0.30179 | 1.94E-06 | YES | NO | NA |
| Ap1g1 | ENSMUSG( | ENSMUSG( | -0.36353 | 5.06E-09 | YES | NO | NA |
| Aopep | ENSMUSG( | ENSMUSG( | 0.021662 | 0.833942 | NO | NO | NA |
| Mia2 | ENSMUSG( | ENSMUSG( | 0.112104 | 0.22789 | NO | NO | NA |
| Nemf | ENSMUSG( | ENSMUSG( | 0.059277 | 0.568419 | NO | NO | NA |
| Kif20b | ENSMUSG( | ENSMUSG( | 0.009352 | 0.946243 | NO | NO | NA |
| Itpk1 | ENSMUSG( | ENSMUSG( | -0.38024 | 1.94E-08 | YES | NO | NA |
| Dnajc15 | ENSMUSG( | ENSMUSG( | 0.471199 | 0.000742 | YES | NO | NA |
| Pfdn5 | ENSMUSG( | ENSMUSG( | -0.11556 | 0.271399 | NO | NO | NA |
| Prim1 | ENSMUSG( | ENSMUSG( | 0.119209 | 0.088713 | NO | NO | NA |
| Eaf1 | ENSMUSG( | ENSMUSG( | -0.05241 | 0.531711 | NO | NO | NA |
| Ehd4 | ENSMUSG( | ENSMUSG( | -0.18304 | 0.023222 | YES | NO | NA |
| Cops6 | ENSMUSG( | ENSMUSG( | 0.162466 | 0.239491 | NO | NO | NA |
| Hadhb | ENSMUSG( | ENSMUSG( | 0.152639 | 0.01973 | YES | NO | NA |
| Nfe2l1 | ENSMUSG( | ENSMUSG( | -0.23345 | 0.001209 | YES | NO | NA |
| Ndufa13 | ENSMUSG( | ENSMUSG( | -0.12764 | 0.305266 | NO | NO | NA |
| Foxp4 | ENSMUSG( | ENSMUSG( | -0.34902 | 0.000318 | YES | NO | NA |
| Gorasp2 | ENSMUSG( | ENSMUSG( | 0.18398 | 0.004163 | YES | NO | NA |
| Cox17 | ENSMUSG( | ENSMUSG( | -0.07104 | 0.50712 | NO | NO | NA |
| Pogk | ENSMUSG( | ENSMUSG( | -0.52708 | 5.87E-08 | YES | NO | NA |
| Golgb1 | ENSMUSG( | ENSMUSG( | -0.11646 | 0.383678 | NO | NO | NA |
| Gm47015 | ENSMUSG( | ENSMUSG( | 1.470638 | 4.02E-11 | YES | YES | Upregulated |
| Elovl5 | ENSMUSG( | ENSMUSG( | -0.14564 | 0.006766 | YES | NO | NA |
| Casc3 | ENSMUSG( | ENSMUSG( | -0.27181 | 3.44E-05 | YES | NO | NA |
| Pdzd8 | ENSMUSG( | ENSMUSG( | 0.000553 | 0.994565 | NO | NO | NA |
| Cdk2 | ENSMUSG( | ENSMUSG( | 0.109842 | 0.046859 | YES | NO | NA |
| Aida | ENSMUSG( | ENSMUSG( | -0.19959 | 0.006271 | YES | NO | NA |
| Unc45a | ENSMUSG( | ENSMUSG( | -0.10399 | 0.060809 | NO | NO | NA |

| | | | | | | | |
|---|---|---|---|---|---|---|---|
| Tob2 | ENSMUSG( | ENSMUSG( | -0.09741 | 0.374936 | NO | NO | NA |
| Hmmr | ENSMUSG( | ENSMUSG( | 0.109819 | 0.288858 | NO | NO | NA |
| Gpatch8 | ENSMUSG( | ENSMUSG( | -0.26372 | 4.14E-05 | YES | NO | NA |
| Smarca2 | ENSMUSG( | ENSMUSG( | -0.24005 | 0.002029 | YES | NO | NA |
| Gfod1 | ENSMUSG( | ENSMUSG( | -0.40371 | 2.89E-05 | YES | NO | NA |
| Hspa14 | ENSMUSG( | ENSMUSG( | 0.120101 | 0.084325 | NO | NO | NA |
| Zfhx2 | ENSMUSG( | ENSMUSG( | -1.0936 | 3.86E-17 | YES | YES | Downregulated |
| Zfr | ENSMUSG( | ENSMUSG( | -0.06003 | 0.430386 | NO | NO | NA |
| Fam117b | ENSMUSG( | ENSMUSG( | -0.35536 | 3.25E-07 | YES | NO | NA |
| Snrpg | ENSMUSG( | ENSMUSG( | 0.275674 | 2.31E-06 | YES | NO | NA |
| Panx1 | ENSMUSG( | ENSMUSG( | -0.21983 | 0.000497 | YES | NO | NA |
| Gm49980 | ENSMUSG( | ENSMUSG( | 2.457405 | 1.39E-12 | YES | YES | Upregulated |
| Pdhb | ENSMUSG( | ENSMUSG( | 0.110377 | 0.184436 | NO | NO | NA |
| Cndp2 | ENSMUSG( | ENSMUSG( | 0.14635 | 0.01453 | YES | NO | NA |
| Dok2 | ENSMUSG( | ENSMUSG( | -0.17617 | 0.054642 | NO | NO | NA |
| Trim27 | ENSMUSG( | ENSMUSG( | 0.138573 | 0.001842 | YES | NO | NA |
| Tent4b | ENSMUSG( | ENSMUSG( | -0.09241 | 0.226504 | NO | NO | NA |
| Gbp3 | ENSMUSG( | ENSMUSG( | 0.477112 | 6.89E-15 | YES | NO | NA |
| Dctn2 | ENSMUSG( | ENSMUSG( | 0.332034 | 0.000264 | YES | NO | NA |
| Ncoa6 | ENSMUSG( | ENSMUSG( | -0.2358 | 0.002077 | YES | NO | NA |
| Nedd8 | ENSMUSG( | ENSMUSG( | 0.090932 | 0.357367 | NO | NO | NA |
| Smg6 | ENSMUSG( | ENSMUSG( | -0.06747 | 0.460586 | NO | NO | NA |
| Cdkn2aipn | ENSMUSG( | ENSMUSG( | -0.02851 | 0.688344 | NO | NO | NA |
| Apeh | ENSMUSG( | ENSMUSG( | -0.29228 | 1.19E-08 | YES | NO | NA |
| Ccdc6 | ENSMUSG( | ENSMUSG( | -0.25646 | 3.41E-05 | YES | NO | NA |
| Kmt5b | ENSMUSG( | ENSMUSG( | -0.53647 | 9.81E-11 | YES | NO | NA |
| Ebna1bp2 | ENSMUSG( | ENSMUSG( | 0.277556 | 0.00185 | YES | NO | NA |
| Rbm27 | ENSMUSG( | ENSMUSG( | -0.18601 | 0.063952 | NO | NO | NA |
| ENSMUSG | ENSMUSG( | ENSMUSG( | -0.26827 | 0.000209 | YES | NO | NA |
| Ncstn | ENSMUSG( | ENSMUSG( | -0.05905 | 0.535806 | NO | NO | NA |
| Apc | ENSMUSG( | ENSMUSG( | -0.3186 | 0.000529 | YES | NO | NA |
| Slc35e1 | ENSMUSG( | ENSMUSG( | -0.15776 | 0.029154 | YES | NO | NA |
| Psmd7 | ENSMUSG( | ENSMUSG( | 0.233804 | 0.064121 | NO | NO | NA |
| Acsl4 | ENSMUSG( | ENSMUSG( | -0.06096 | 0.458411 | NO | NO | NA |
| Trap1 | ENSMUSG( | ENSMUSG( | 0.313718 | 0.000332 | YES | NO | NA |
| Exosc8 | ENSMUSG( | ENSMUSG( | 0.113158 | 0.120341 | NO | NO | NA |
| Trim12c | ENSMUSG( | ENSMUSG( | -0.69201 | 3.13E-12 | YES | NO | NA |
| Pola1 | ENSMUSG( | ENSMUSG( | -0.02569 | 0.810189 | NO | NO | NA |
| Exoc5 | ENSMUSG( | ENSMUSG( | 0.036226 | 0.600183 | NO | NO | NA |
| Arhgap19 | ENSMUSG( | ENSMUSG( | 0.280112 | 0.000109 | YES | NO | NA |
| Mat2b | ENSMUSG( | ENSMUSG( | -0.17544 | 0.040619 | YES | NO | NA |
| Cdca4 | ENSMUSG( | ENSMUSG( | 0.138841 | 0.003309 | YES | NO | NA |
| Elk4 | ENSMUSG( | ENSMUSG( | -0.28323 | 0.001156 | YES | NO | NA |
| Ipo8 | ENSMUSG( | ENSMUSG( | -0.14848 | 1.88E-05 | YES | NO | NA |
| Arrdc1 | ENSMUSG( | ENSMUSG( | -0.14063 | 0.025187 | YES | NO | NA |
| Itgb3 | ENSMUSG( | ENSMUSG( | 0.35239 | 0.008802 | YES | NO | NA |
| Polr3c | ENSMUSG( | ENSMUSG( | -0.00257 | 0.972626 | NO | NO | NA |

| | | | | | | | |
|---|---|---|---|---|---|---|---|
| Max | ENSMUSG( | ENSMUSG( | -0.17732 | 0.134001 | NO | NO | NA |
| Gimap9 | ENSMUSG( | ENSMUSG( | -0.29509 | 0.000506 | YES | NO | NA |
| Sh3gl1 | ENSMUSG( | ENSMUSG( | -0.33237 | 1.89E-11 | YES | NO | NA |
| Dpf2 | ENSMUSG( | ENSMUSG( | -0.17906 | 0.000498 | YES | NO | NA |
| Fyco1 | ENSMUSG( | ENSMUSG( | -0.35674 | 0.000109 | YES | NO | NA |
| Ncoa1 | ENSMUSG( | ENSMUSG( | -0.23944 | 9.53E-05 | YES | NO | NA |
| Sec23b | ENSMUSG( | ENSMUSG( | -0.15164 | 0.001434 | YES | NO | NA |
| Myo5a | ENSMUSG( | ENSMUSG( | 0.142145 | 0.044089 | YES | NO | NA |
| Sec24b | ENSMUSG( | ENSMUSG( | -0.29229 | 9.80E-07 | YES | NO | NA |
| Sugt1 | ENSMUSG( | ENSMUSG( | -0.01715 | 0.801826 | NO | NO | NA |
| Sptlc2 | ENSMUSG( | ENSMUSG( | 0.002852 | 0.975721 | NO | NO | NA |
| Adamts14 | ENSMUSG( | ENSMUSG( | 0.007318 | 0.949938 | NO | NO | NA |
| Thap12 | ENSMUSG( | ENSMUSG( | 0.39167 | 4.07E-05 | YES | NO | NA |
| Ppil1 | ENSMUSG( | ENSMUSG( | 0.311095 | 0.000108 | YES | NO | NA |
| Ash1l | ENSMUSG( | ENSMUSG( | -0.46006 | 1.31E-07 | YES | NO | NA |
| Bicral | ENSMUSG( | ENSMUSG( | -0.52728 | 9.39E-12 | YES | NO | NA |
| Tspoap1 | ENSMUSG( | ENSMUSG( | 0.166469 | 0.396816 | NO | NO | NA |
| Map2k7 | ENSMUSG( | ENSMUSG( | -0.01424 | 0.881484 | NO | NO | NA |
| Stk32c | ENSMUSG( | ENSMUSG( | 0.254976 | 0.007279 | YES | NO | NA |
| Iws1 | ENSMUSG( | ENSMUSG( | -0.06068 | 0.495195 | NO | NO | NA |
| Rc3h1 | ENSMUSG( | ENSMUSG( | -0.45454 | 2.92E-08 | YES | NO | NA |
| Wdr33 | ENSMUSG( | ENSMUSG( | 0.040947 | 0.323716 | NO | NO | NA |
| Scp2 | ENSMUSG( | ENSMUSG( | -0.21912 | 0.020336 | YES | NO | NA |
| Cdkn2c | ENSMUSG( | ENSMUSG( | -0.00815 | 0.942273 | NO | NO | NA |
| Prelid3b | ENSMUSG( | ENSMUSG( | 0.373084 | 3.24E-07 | YES | NO | NA |
| Lars | ENSMUSG( | ENSMUSG( | 0.32349 | 0.000269 | YES | NO | NA |
| Etfa | ENSMUSG( | ENSMUSG( | 0.206512 | 0.03147 | YES | NO | NA |
| Mtmr6 | ENSMUSG( | ENSMUSG( | -0.16641 | 0.108578 | NO | NO | NA |
| Usp8 | ENSMUSG( | ENSMUSG( | -0.08904 | 0.415514 | NO | NO | NA |
| Arhgap27 | ENSMUSG( | ENSMUSG( | -0.38784 | 8.25E-09 | YES | NO | NA |
| Prr14 | ENSMUSG( | ENSMUSG( | -0.20464 | 0.000169 | YES | NO | NA |
| Maf1 | ENSMUSG( | ENSMUSG( | -0.25804 | 0.005039 | YES | NO | NA |
| Tbl1xr1 | ENSMUSG( | ENSMUSG( | -0.05994 | 0.580206 | NO | NO | NA |
| Sumo1 | ENSMUSG( | ENSMUSG( | 0.100254 | 0.107146 | NO | NO | NA |
| Itch | ENSMUSG( | ENSMUSG( | -0.23531 | 0.003449 | YES | NO | NA |
| Cdc34 | ENSMUSG( | ENSMUSG( | 0.233311 | 0.001528 | YES | NO | NA |
| Ildr1 | ENSMUSG( | ENSMUSG( | -0.80018 | 2.88E-08 | YES | NO | NA |
| Tnfaip8 | ENSMUSG( | ENSMUSG( | 0.041556 | 0.500516 | NO | NO | NA |
| Psmd11 | ENSMUSG( | ENSMUSG( | -0.11429 | 0.095652 | NO | NO | NA |
| Tfip11 | ENSMUSG( | ENSMUSG( | -0.16518 | 0.000194 | YES | NO | NA |
| Esd | ENSMUSG( | ENSMUSG( | 0.085201 | 0.260444 | NO | NO | NA |
| Ndufa4 | ENSMUSG( | ENSMUSG( | 0.037649 | 0.57964 | NO | NO | NA |
| Mif4gd | ENSMUSG( | ENSMUSG( | 0.181803 | 0.048822 | YES | NO | NA |
| Zfp148 | ENSMUSG( | ENSMUSG( | -0.25454 | 0.002777 | YES | NO | NA |
| Ctla2a | ENSMUSG( | ENSMUSG( | 0.381544 | 0.028189 | YES | NO | NA |
| Chuk | ENSMUSG( | ENSMUSG( | 0.048925 | 0.5778 | NO | NO | NA |
| Zfand5 | ENSMUSG( | ENSMUSG( | -0.10269 | 0.23514 | NO | NO | NA |

| | | | | | | | |
|---|---|---|---|---|---|---|---|
| Nf2 | ENSMUSG( | ENSMUSG( | -0.16851 | 0.002331 | YES | NO | NA |
| Naa60 | ENSMUSG( | ENSMUSG( | -0.3136 | 3.47E-05 | YES | NO | NA |
| Slc35a4 | ENSMUSG( | ENSMUSG( | 0.118799 | 0.030878 | YES | NO | NA |
| Fxr1 | ENSMUSG( | ENSMUSG( | -0.11403 | 0.250962 | NO | NO | NA |
| Bop1 | ENSMUSG( | ENSMUSG( | 0.357704 | 5.27E-06 | YES | NO | NA |
| AI504432 | ENSMUSG( | ENSMUSG( | -0.44783 | 0.000483 | YES | NO | NA |
| Pglyrp2 | ENSMUSG( | ENSMUSG( | -0.80872 | 5.97E-12 | YES | NO | NA |
| Ptger4 | ENSMUSG( | ENSMUSG( | -0.67884 | 1.08E-08 | YES | NO | NA |
| Ostm1 | ENSMUSG( | ENSMUSG( | -0.18904 | 0.039238 | YES | NO | NA |
| Utp3 | ENSMUSG( | ENSMUSG( | 0.129847 | 0.351405 | NO | NO | NA |
| Dynlrb1 | ENSMUSG( | ENSMUSG( | -0.13526 | 0.115093 | NO | NO | NA |
| B3gat3 | ENSMUSG( | ENSMUSG( | -0.27792 | 1.12E-10 | YES | NO | NA |
| Fam32a | ENSMUSG( | ENSMUSG( | -0.26886 | 0.000806 | YES | NO | NA |
| Mirt1 | ENSMUSG( | ENSMUSG( | -0.6138 | 7.09E-26 | YES | NO | NA |
| Septin2 EN | ENSMUSG( | ENSMUSG( | -0.05908 | 0.425871 | NO | NO | NA |
| Chmp2a | ENSMUSG( | ENSMUSG( | -0.02776 | 0.827588 | NO | NO | NA |
| Gm10222 | ENSMUSG( | ENSMUSG( | -0.20722 | 0.391098 | NO | NO | NA |
| Tet3 | ENSMUSG( | ENSMUSG( | -0.50649 | 7.13E-08 | YES | NO | NA |
| Nfrkb | ENSMUSG( | ENSMUSG( | -0.11114 | 0.218244 | NO | NO | NA |
| Nr3c1 | ENSMUSG( | ENSMUSG( | -0.38787 | 0.000141 | YES | NO | NA |
| Foxk2 | ENSMUSG( | ENSMUSG( | -0.00129 | 0.984529 | NO | NO | NA |
| Suco | ENSMUSG( | ENSMUSG( | -0.31666 | 0.000391 | YES | NO | NA |
| Zmynd8 | ENSMUSG( | ENSMUSG( | -0.47999 | 2.17E-13 | YES | NO | NA |
| Strn4 | ENSMUSG( | ENSMUSG( | -0.13907 | 0.000853 | YES | NO | NA |
| Actr1b | ENSMUSG( | ENSMUSG( | -0.01796 | 0.747282 | NO | NO | NA |
| Sntb2 | ENSMUSG( | ENSMUSG( | -0.132 | 0.209077 | NO | NO | NA |
| Trabd | ENSMUSG( | ENSMUSG( | -0.04018 | 0.473973 | NO | NO | NA |
| Lrrc47 | ENSMUSG( | ENSMUSG( | -0.00615 | 0.931584 | NO | NO | NA |
| Dgcr2 | ENSMUSG( | ENSMUSG( | -0.2808 | 1.04E-05 | YES | NO | NA |
| Larp4 | ENSMUSG( | ENSMUSG( | 0.191345 | 0.005889 | YES | NO | NA |
| Fis1 | ENSMUSG( | ENSMUSG( | -0.05556 | 0.475513 | NO | NO | NA |
| Ube4a | ENSMUSG( | ENSMUSG( | -0.09231 | 0.077981 | NO | NO | NA |
| Vgll4 | ENSMUSG( | ENSMUSG( | -0.73231 | 1.34E-12 | YES | NO | NA |
| Itih5 | ENSMUSG( | ENSMUSG( | 0.094743 | 0.273898 | NO | NO | NA |
| Ric1 | ENSMUSG( | ENSMUSG( | -0.38041 | 1.48E-05 | YES | NO | NA |
| 2900026A( | ENSMUSG( | ENSMUSG( | -1.09091 | 3.46E-07 | YES | YES | Downregulated |
| Psma5 | ENSMUSG( | ENSMUSG( | 0.260539 | 4.83E-06 | YES | NO | NA |
| Pip4k2c | ENSMUSG( | ENSMUSG( | -0.22963 | 3.68E-05 | YES | NO | NA |
| Dda1 | ENSMUSG( | ENSMUSG( | -0.08779 | 0.191679 | NO | NO | NA |
| Slc4a2 | ENSMUSG( | ENSMUSG( | -0.12988 | 0.010727 | YES | NO | NA |
| Pik3ca | ENSMUSG( | ENSMUSG( | -0.25546 | 0.000143 | YES | NO | NA |
| Brap | ENSMUSG( | ENSMUSG( | 0.039674 | 0.663478 | NO | NO | NA |
| Eif4e | ENSMUSG( | ENSMUSG( | 0.197851 | 0.018097 | YES | NO | NA |
| Arih2 | ENSMUSG( | ENSMUSG( | -0.01283 | 0.920392 | NO | NO | NA |
| Nfyb | ENSMUSG( | ENSMUSG( | -0.09931 | 0.328452 | NO | NO | NA |
| B3gnt2 | ENSMUSG( | ENSMUSG( | -0.1806 | 0.005591 | YES | NO | NA |
| Dtl | ENSMUSG( | ENSMUSG( | 0.321977 | 3.96E-06 | YES | NO | NA |

| | | | | | | | |
|---|---|---|---|---|---|---|---|
| Ncbp3 | ENSMUSG( | ENSMUSG( | 0.180511 | 4.32E-06 | YES | NO | NA |
| Ckap2l | ENSMUSG( | ENSMUSG( | 0.135201 | 0.141441 | NO | NO | NA |
| Arap1 | ENSMUSG( | ENSMUSG( | -0.43171 | 4.57E-12 | YES | NO | NA |
| Ppp2r2a | ENSMUSG( | ENSMUSG( | -0.13172 | 0.002769 | YES | NO | NA |
| Ndufs1 | ENSMUSG( | ENSMUSG( | 0.217759 | 0.000219 | YES | NO | NA |
| Wiz | ENSMUSG( | ENSMUSG( | 0.001616 | 0.981136 | NO | NO | NA |
| Sem1 | ENSMUSG( | ENSMUSG( | 0.110671 | 0.135458 | NO | NO | NA |
| Snrpf | ENSMUSG( | ENSMUSG( | 0.3876 | 0.000145 | YES | NO | NA |
| Zfp592 | ENSMUSG( | ENSMUSG( | -0.3475 | 1.76E-07 | YES | NO | NA |
| Notch2 | ENSMUSG( | ENSMUSG( | -0.65601 | 8.13E-15 | YES | NO | NA |
| Brk1 | ENSMUSG( | ENSMUSG( | -0.13603 | 0.012064 | YES | NO | NA |
| Bicd2 | ENSMUSG( | ENSMUSG( | -0.37131 | 1.47E-07 | YES | NO | NA |
| Slain1 | ENSMUSG( | ENSMUSG( | -0.11721 | 0.176034 | NO | NO | NA |
| Hif1an | ENSMUSG( | ENSMUSG( | -0.60464 | 6.30E-10 | YES | NO | NA |
| Uso1 | ENSMUSG( | ENSMUSG( | -0.11667 | 0.115746 | NO | NO | NA |
| Map3k7 | ENSMUSG( | ENSMUSG( | -0.01656 | 0.852826 | NO | NO | NA |
| Litaf | ENSMUSG( | ENSMUSG( | 0.098084 | 0.425984 | NO | NO | NA |
| Ndfip2 | ENSMUSG( | ENSMUSG( | -0.22947 | 0.006473 | YES | NO | NA |
| Mthfd2 | ENSMUSG( | ENSMUSG( | 0.392447 | 3.97E-06 | YES | NO | NA |
| Gstp1 | ENSMUSG( | ENSMUSG( | 0.136104 | 0.182925 | NO | NO | NA |
| Arid1b | ENSMUSG( | ENSMUSG( | -0.40242 | 1.60E-12 | YES | NO | NA |
| Afg3l2 | ENSMUSG( | ENSMUSG( | 0.304221 | 1.62E-05 | YES | NO | NA |
| Vdac3 | ENSMUSG( | ENSMUSG( | 0.111754 | 0.190366 | NO | NO | NA |
| Ankrd40 | ENSMUSG( | ENSMUSG( | -0.15206 | 0.004928 | YES | NO | NA |
| Brd7 | ENSMUSG( | ENSMUSG( | 0.04489 | 0.647827 | NO | NO | NA |
| Krit1 | ENSMUSG( | ENSMUSG( | -0.26628 | 0.000681 | YES | NO | NA |
| Tmx1 | ENSMUSG( | ENSMUSG( | -0.0792 | 0.105725 | NO | NO | NA |
| Snx17 | ENSMUSG( | ENSMUSG( | 0.047245 | 0.529934 | NO | NO | NA |
| Entpd1 | ENSMUSG( | ENSMUSG( | -0.56404 | 8.40E-05 | YES | NO | NA |
| Iars | ENSMUSG( | ENSMUSG( | 0.147979 | 0.062767 | NO | NO | NA |
| Nvl | ENSMUSG( | ENSMUSG( | 0.139729 | 0.159326 | NO | NO | NA |
| Suv39h1 | ENSMUSG( | ENSMUSG( | 0.352493 | 3.45E-05 | YES | NO | NA |
| Traf3 | ENSMUSG( | ENSMUSG( | -0.30603 | 0.000104 | YES | NO | NA |
| Paip2 | ENSMUSG( | ENSMUSG( | -0.16765 | 0.063933 | NO | NO | NA |
| Nufip2 | ENSMUSG( | ENSMUSG( | -0.44074 | 3.69E-08 | YES | NO | NA |
| Mfsd14a | ENSMUSG( | ENSMUSG( | -0.15196 | 0.074969 | NO | NO | NA |
| Araf | ENSMUSG( | ENSMUSG( | -0.10651 | 0.056008 | NO | NO | NA |
| Sf3b4 | ENSMUSG( | ENSMUSG( | 0.136411 | 0.00856 | YES | NO | NA |
| Zdhhc2 | ENSMUSG( | ENSMUSG( | -0.13519 | 0.027036 | YES | NO | NA |
| Ube4b | ENSMUSG( | ENSMUSG( | -0.19446 | 0.001136 | YES | NO | NA |
| Eml4 | ENSMUSG( | ENSMUSG( | -0.12633 | 0.008319 | YES | NO | NA |
| Ccm2 | ENSMUSG( | ENSMUSG( | -0.42387 | 3.15E-07 | YES | NO | NA |
| Otub1 | ENSMUSG( | ENSMUSG( | 0.029204 | 0.693448 | NO | NO | NA |
| Ccne1 | ENSMUSG( | ENSMUSG( | 0.444375 | 6.15E-09 | YES | NO | NA |
| Fam53b | ENSMUSG( | ENSMUSG( | -0.44922 | 3.96E-09 | YES | NO | NA |
| Tomm22 | ENSMUSG( | ENSMUSG( | -0.00017 | 0.998348 | NO | NO | NA |
| Psmd12 | ENSMUSG( | ENSMUSG( | 0.303223 | 0.00229 | YES | NO | NA |

| | | | | | | | |
|---|---|---|---|---|---|---|---|
| Fbxw2 | ENSMUSG( | ENSMUSG( | -0.15669 | 0.026929 | YES | NO | NA |
| Prr11 | ENSMUSG( | ENSMUSG( | 0.086371 | 0.390736 | NO | NO | NA |
| Ubald2 | ENSMUSG( | ENSMUSG( | 0.191108 | 0.060563 | NO | NO | NA |
| Rbm28 | ENSMUSG( | ENSMUSG( | -0.03648 | 0.645518 | NO | NO | NA |
| Nbr1 | ENSMUSG( | ENSMUSG( | -0.49545 | 2.83E-09 | YES | NO | NA |
| Plaat3 | ENSMUSG( | ENSMUSG( | -0.45795 | 7.36E-10 | YES | NO | NA |
| Knstrn | ENSMUSG( | ENSMUSG( | 0.327914 | 0.015951 | YES | NO | NA |
| Daxx | ENSMUSG( | ENSMUSG( | -0.42416 | 1.67E-10 | YES | NO | NA |
| Pdcd10 | ENSMUSG( | ENSMUSG( | -0.16022 | 0.06851 | NO | NO | NA |
| Rfc5 | ENSMUSG( | ENSMUSG( | 0.283884 | 2.56E-07 | YES | NO | NA |
| Rnf115 | ENSMUSG( | ENSMUSG( | -0.15537 | 0.05147 | NO | NO | NA |
| Prr14l | ENSMUSG( | ENSMUSG( | -0.38547 | 7.73E-08 | YES | NO | NA |
| Fbxl3 | ENSMUSG( | ENSMUSG( | -0.20089 | 0.012447 | YES | NO | NA |
| Eif2ak1 | ENSMUSG( | ENSMUSG( | 0.058646 | 0.377555 | NO | NO | NA |
| Eps15l1 | ENSMUSG( | ENSMUSG( | -0.39673 | 5.72E-11 | YES | NO | NA |
| Sap130 | ENSMUSG( | ENSMUSG( | -0.23535 | 3.08E-05 | YES | NO | NA |
| Stat2 | ENSMUSG( | ENSMUSG( | -0.41708 | 4.51E-11 | YES | NO | NA |
| Slc25a24 | ENSMUSG( | ENSMUSG( | -0.5028 | 8.34E-08 | YES | NO | NA |
| Ddx50 | ENSMUSG( | ENSMUSG( | 0.076696 | 0.419115 | NO | NO | NA |
| Atm | ENSMUSG( | ENSMUSG( | -0.37198 | 3.94E-10 | YES | NO | NA |
| Khdc4 | ENSMUSG( | ENSMUSG( | -0.15894 | 0.052605 | NO | NO | NA |
| Trim59 | ENSMUSG( | ENSMUSG( | -0.14353 | 0.04127 | YES | NO | NA |
| Anxa7 | ENSMUSG( | ENSMUSG( | -0.1297 | 0.017279 | YES | NO | NA |
| Snu13 | ENSMUSG( | ENSMUSG( | 0.246072 | 0.000194 | YES | NO | NA |
| Foxn2 | ENSMUSG( | ENSMUSG( | -0.29647 | 2.96E-06 | YES | NO | NA |
| Fkbp15 | ENSMUSG( | ENSMUSG( | -0.08418 | 0.190779 | NO | NO | NA |
| Setbp1 | ENSMUSG( | ENSMUSG( | -0.8065 | 1.57E-12 | YES | NO | NA |
| Agfg2 | ENSMUSG( | ENSMUSG( | -0.14019 | 0.076802 | NO | NO | NA |
| Angel2 | ENSMUSG( | ENSMUSG( | 0.054552 | 0.585511 | NO | NO | NA |
| Ube2b | ENSMUSG( | ENSMUSG( | -0.45042 | 1.43E-07 | YES | NO | NA |
| Lin7c | ENSMUSG( | ENSMUSG( | 0.012504 | 0.916397 | NO | NO | NA |
| Il12rb1 | ENSMUSG( | ENSMUSG( | -0.1584 | 0.015374 | YES | NO | NA |
| Med24 | ENSMUSG( | ENSMUSG( | 0.118383 | 0.209948 | NO | NO | NA |
| Slc9a1 | ENSMUSG( | ENSMUSG( | -0.42583 | 4.39E-09 | YES | NO | NA |
| Mars1 | ENSMUSG( | ENSMUSG( | 0.236726 | 0.001357 | YES | NO | NA |
| Opa1 | ENSMUSG( | ENSMUSG( | 0.110671 | 0.279361 | NO | NO | NA |
| Fxr2 | ENSMUSG( | ENSMUSG( | -0.06407 | 0.121345 | NO | NO | NA |
| Phf14 | ENSMUSG( | ENSMUSG( | -0.03406 | 0.734711 | NO | NO | NA |
| Ano6 | ENSMUSG( | ENSMUSG( | -0.26309 | 0.000102 | YES | NO | NA |
| Zfp384 | ENSMUSG( | ENSMUSG( | 0.053952 | 0.547577 | NO | NO | NA |
| Armc7 | ENSMUSG( | ENSMUSG( | -0.61633 | 1.25E-06 | YES | NO | NA |
| Cat | ENSMUSG( | ENSMUSG( | -0.24233 | 0.000449 | YES | NO | NA |
| Celsr1 | ENSMUSG( | ENSMUSG( | -0.5094 | 1.63E-06 | YES | NO | NA |
| Hira | ENSMUSG( | ENSMUSG( | 0.167151 | 0.003881 | YES | NO | NA |
| Kif2a | ENSMUSG( | ENSMUSG( | 0.155734 | 0.083518 | NO | NO | NA |
| Polg | ENSMUSG( | ENSMUSG( | 0.0218 | 0.80634 | NO | NO | NA |
| Naa40 | ENSMUSG( | ENSMUSG( | -0.01522 | 0.770192 | NO | NO | NA |

| | | | | | | | |
|---|---|---|---|---|---|---|---|
| 5830417I1 | ENSMUSG( | ENSMUSG( | -0.2973 | 1.05E-11 | YES | NO | NA |
| Snx1 | ENSMUSG( | ENSMUSG( | -0.12421 | 0.122158 | NO | NO | NA |
| Nubp1 | ENSMUSG( | ENSMUSG( | 0.306396 | 4.84E-05 | YES | NO | NA |
| Rcn1 | ENSMUSG( | ENSMUSG( | -0.4775 | 1.60E-09 | YES | NO | NA |
| Setd7 | ENSMUSG( | ENSMUSG( | -0.31435 | 6.92E-05 | YES | NO | NA |
| Sf3a3 | ENSMUSG( | ENSMUSG( | 0.118468 | 0.298727 | NO | NO | NA |
| Npm3 | ENSMUSG( | ENSMUSG( | 0.449538 | 8.26E-07 | YES | NO | NA |
| Mindy2 | ENSMUSG( | ENSMUSG( | -0.32115 | 5.30E-07 | YES | NO | NA |
| Zyg11b | ENSMUSG( | ENSMUSG( | 0.059898 | 0.629563 | NO | NO | NA |
| Cpsf3 | ENSMUSG( | ENSMUSG( | 0.129473 | 0.025583 | YES | NO | NA |
| Clasp1 | ENSMUSG( | ENSMUSG( | 0.064231 | 0.400137 | NO | NO | NA |
| Elf2 | ENSMUSG( | ENSMUSG( | -0.1482 | 0.051029 | NO | NO | NA |
| Tmem229I | ENSMUSG( | ENSMUSG( | -0.26506 | 7.73E-07 | YES | NO | NA |
| Usp38 | ENSMUSG( | ENSMUSG( | -0.2355 | 0.000195 | YES | NO | NA |
| Icos | ENSMUSG( | ENSMUSG( | -0.0138 | 0.949513 | NO | NO | NA |
| Nadk | ENSMUSG( | ENSMUSG( | 0.091248 | 0.093463 | NO | NO | NA |
| Ube2j2 | ENSMUSG( | ENSMUSG( | 0.078458 | 0.22364 | NO | NO | NA |
| Srgap3 | ENSMUSG( | ENSMUSG( | -0.73138 | 9.04E-09 | YES | NO | NA |
| Xpnpep1 | ENSMUSG( | ENSMUSG( | 0.252897 | 0.000255 | YES | NO | NA |
| Clcn4 | ENSMUSG( | ENSMUSG( | -0.16041 | 0.034857 | YES | NO | NA |
| 9530068E( | ENSMUSG( | ENSMUSG( | 0.032334 | 0.560369 | NO | NO | NA |
| Vps52 | ENSMUSG( | ENSMUSG( | -0.05436 | 0.391166 | NO | NO | NA |
| Mgat1 | ENSMUSG( | ENSMUSG( | -0.28439 | 2.49E-08 | YES | NO | NA |
| Myd88 | ENSMUSG( | ENSMUSG( | 0.152462 | 0.022804 | YES | NO | NA |
| Rnpep | ENSMUSG( | ENSMUSG( | 0.045414 | 0.567401 | NO | NO | NA |
| Herc6 | ENSMUSG( | ENSMUSG( | -0.80648 | 1.54E-17 | YES | NO | NA |
| Pwwp3a | ENSMUSG( | ENSMUSG( | 0.036094 | 0.48269 | NO | NO | NA |
| Rtraf | ENSMUSG( | ENSMUSG( | 0.241888 | 0.00515 | YES | NO | NA |
| Bcas2 | ENSMUSG( | ENSMUSG( | 0.044377 | 0.45469 | NO | NO | NA |
| Specc1l | ENSMUSG( | ENSMUSG( | 0.012885 | 0.859542 | NO | NO | NA |
| Trappc8 | ENSMUSG( | ENSMUSG( | -0.37856 | 7.85E-09 | YES | NO | NA |
| Cdk12 | ENSMUSG( | ENSMUSG( | -0.22432 | 0.000806 | YES | NO | NA |
| Kif14 | ENSMUSG( | ENSMUSG( | 0.128809 | 0.159529 | NO | NO | NA |
| Aldh18a1 | ENSMUSG( | ENSMUSG( | -0.10401 | 0.103235 | NO | NO | NA |
| Helb | ENSMUSG( | ENSMUSG( | -0.05926 | 0.465162 | NO | NO | NA |
| Ccnk | ENSMUSG( | ENSMUSG( | -0.14196 | 0.09763 | NO | NO | NA |
| Prpf39 | ENSMUSG( | ENSMUSG( | -0.18982 | 0.020928 | YES | NO | NA |
| Tagap | ENSMUSG( | ENSMUSG( | -0.45455 | 8.40E-05 | YES | NO | NA |
| Fignl1 | ENSMUSG( | ENSMUSG( | 0.220818 | 0.021137 | YES | NO | NA |
| Asnsd1 | ENSMUSG( | ENSMUSG( | 0.103943 | 0.148433 | NO | NO | NA |
| Heatr1 | ENSMUSG( | ENSMUSG( | 0.316143 | 0.000362 | YES | NO | NA |
| Glipr1 | ENSMUSG( | ENSMUSG( | 0.10154 | 0.347354 | NO | NO | NA |
| Tex261 | ENSMUSG( | ENSMUSG( | 0.102659 | 0.155527 | NO | NO | NA |
| Entr1 | ENSMUSG( | ENSMUSG( | -0.13288 | 0.00735 | YES | NO | NA |
| Kif18b | ENSMUSG( | ENSMUSG( | 0.081829 | 0.3311 | NO | NO | NA |
| Cirbp | ENSMUSG( | ENSMUSG( | 0.725464 | 1.94E-08 | YES | NO | NA |
| Ubac2 | ENSMUSG( | ENSMUSG( | -0.13425 | 0.009208 | YES | NO | NA |

| | | | | | | | |
|---|---|---|---|---|---|---|---|
| Hmgcs1 | ENSMUSG( | ENSMUSG( | 0.465602 | 0.000409 | YES | NO | NA |
| Hbs1l | ENSMUSG( | ENSMUSG( | 0.29868 | 4.18E-07 | YES | NO | NA |
| Taf5l | ENSMUSG( | ENSMUSG( | 0.157119 | 0.033754 | YES | NO | NA |
| Tet2 | ENSMUSG( | ENSMUSG( | -0.19939 | 0.07382 | NO | NO | NA |
| Rai1 | ENSMUSG( | ENSMUSG( | 0.156653 | 0.039522 | YES | NO | NA |
| Brpf3 | ENSMUSG( | ENSMUSG( | -0.2009 | 0.025054 | YES | NO | NA |
| Ostc | ENSMUSG( | ENSMUSG( | 0.084928 | 0.365952 | NO | NO | NA |
| Dkc1 | ENSMUSG( | ENSMUSG( | 0.513226 | 9.47E-06 | YES | NO | NA |
| Tbc1d20 | ENSMUSG( | ENSMUSG( | -0.35603 | 4.23E-06 | YES | NO | NA |
| Bdp1 | ENSMUSG( | ENSMUSG( | -0.23443 | 0.000549 | YES | NO | NA |
| Mark3 | ENSMUSG( | ENSMUSG( | -0.14955 | 0.017776 | YES | NO | NA |
| Tmem131l | ENSMUSG( | ENSMUSG( | -0.6756 | 2.76E-09 | YES | NO | NA |
| Ndufb10 | ENSMUSG( | ENSMUSG( | 0.297512 | 2.67E-05 | YES | NO | NA |
| Inpp5k | ENSMUSG( | ENSMUSG( | -0.12072 | 0.384056 | NO | NO | NA |
| Cep295 | ENSMUSG( | ENSMUSG( | 0.018554 | 0.855635 | NO | NO | NA |
| Srek1 | ENSMUSG( | ENSMUSG( | 0.032436 | 0.781929 | NO | NO | NA |
| Slc35c1 | ENSMUSG( | ENSMUSG( | -0.38785 | 7.58E-13 | YES | NO | NA |
| Hexb | ENSMUSG( | ENSMUSG( | -0.28173 | 0.000384 | YES | NO | NA |
| Camk2g | ENSMUSG( | ENSMUSG( | 0.052708 | 0.561096 | NO | NO | NA |
| Jakmip1 | ENSMUSG( | ENSMUSG( | -0.1954 | 0.004949 | YES | NO | NA |
| Fam149b | ENSMUSG( | ENSMUSG( | -0.6293 | 3.64E-08 | YES | NO | NA |
| Atp6v1f | ENSMUSG( | ENSMUSG( | -0.08147 | 0.478362 | NO | NO | NA |
| Ube3b | ENSMUSG( | ENSMUSG( | -0.14033 | 0.028709 | YES | NO | NA |
| Paf1 | ENSMUSG( | ENSMUSG( | 0.021399 | 0.838685 | NO | NO | NA |
| Mapkapk3 | ENSMUSG( | ENSMUSG( | -0.00054 | 0.995654 | NO | NO | NA |
| Itgav | ENSMUSG( | ENSMUSG( | -0.44528 | 2.79E-06 | YES | NO | NA |
| Entpd5 | ENSMUSG( | ENSMUSG( | 0.530494 | 1.17E-05 | YES | NO | NA |
| Retreg2 | ENSMUSG( | ENSMUSG( | -0.12417 | 0.13709 | NO | NO | NA |
| Rfx1 | ENSMUSG( | ENSMUSG( | -0.40381 | 1.90E-09 | YES | NO | NA |
| Sfr1 | ENSMUSG( | ENSMUSG( | 0.047922 | 0.683829 | NO | NO | NA |
| Gbp8 | ENSMUSG( | ENSMUSG( | -0.73197 | 1.09E-09 | YES | NO | NA |
| Ski | ENSMUSG( | ENSMUSG( | -0.44113 | 2.84E-06 | YES | NO | NA |
| Usp3 | ENSMUSG( | ENSMUSG( | -0.22986 | 0.001942 | YES | NO | NA |
| Snrpe | ENSMUSG( | ENSMUSG( | 0.290215 | 1.48E-08 | YES | NO | NA |
| Sephs2 | ENSMUSG( | ENSMUSG( | 0.265456 | 0.001215 | YES | NO | NA |
| Hpf1 | ENSMUSG( | ENSMUSG( | 0.024874 | 0.798718 | NO | NO | NA |
| Pnrc2 | ENSMUSG( | ENSMUSG( | -0.37718 | 1.16E-05 | YES | NO | NA |
| Atad5 | ENSMUSG( | ENSMUSG( | 0.228962 | 0.001598 | YES | NO | NA |
| Arl1 | ENSMUSG( | ENSMUSG( | 0.242921 | 0.001094 | YES | NO | NA |
| Degs1 | ENSMUSG( | ENSMUSG( | 0.015448 | 0.812809 | NO | NO | NA |
| Furin | ENSMUSG( | ENSMUSG( | 0.260244 | 0.001242 | YES | NO | NA |
| Map2k3 | ENSMUSG( | ENSMUSG( | 0.195975 | 0.000474 | YES | NO | NA |
| Taok1 | ENSMUSG( | ENSMUSG( | -0.35669 | 3.39E-05 | YES | NO | NA |
| Pank2 | ENSMUSG( | ENSMUSG( | -0.10366 | 0.223572 | NO | NO | NA |
| Nedd4 | ENSMUSG( | ENSMUSG( | -0.42873 | 0.000814 | YES | NO | NA |
| Wrn | ENSMUSG( | ENSMUSG( | -0.34308 | 2.13E-06 | YES | NO | NA |
| Dtymk | ENSMUSG( | ENSMUSG( | 0.460752 | 8.59E-09 | YES | NO | NA |

| | | | | | | | |
|---|---|---|---|---|---|---|---|
| Rap2c | ENSMUSG( | ENSMUSG( | -0.18975 | 0.00558 | YES | NO | NA |
| Oas1a | ENSMUSG( | ENSMUSG( | -1.34677 | 9.24E-14 | YES | YES | Downregulated |
| Slc4a7 | ENSMUSG( | ENSMUSG( | -0.47064 | 1.10E-05 | YES | NO | NA |
| Dlg1 | ENSMUSG( | ENSMUSG( | -0.3464 | 1.45E-05 | YES | NO | NA |
| Atp6ap1 | ENSMUSG( | ENSMUSG( | -0.29586 | 4.34E-05 | YES | NO | NA |
| Gm49751 | ENSMUSG( | ENSMUSG( | 0.153135 | 0.438825 | NO | NO | NA |
| Ccnt2 | ENSMUSG( | ENSMUSG( | -0.37813 | 1.10E-06 | YES | NO | NA |
| Dld | ENSMUSG( | ENSMUSG( | 0.28225 | 0.000204 | YES | NO | NA |
| Tor4a | ENSMUSG( | ENSMUSG( | -0.56954 | 1.05E-11 | YES | NO | NA |
| Mxd1 | ENSMUSG( | ENSMUSG( | -0.75985 | 4.85E-11 | YES | NO | NA |
| Ppp2r5d | ENSMUSG( | ENSMUSG( | 0.037824 | 0.492973 | NO | NO | NA |
| Phf8 | ENSMUSG( | ENSMUSG( | -0.41962 | 5.80E-08 | YES | NO | NA |
| Golga3 | ENSMUSG( | ENSMUSG( | -0.20505 | 0.00023 | YES | NO | NA |
| Ndc80 | ENSMUSG( | ENSMUSG( | 0.204833 | 0.036993 | YES | NO | NA |
| Nuf2 | ENSMUSG( | ENSMUSG( | 0.211995 | 0.025296 | YES | NO | NA |
| Ppp1r8 | ENSMUSG( | ENSMUSG( | 0.224782 | 0.002395 | YES | NO | NA |
| Irgm1 | ENSMUSG( | ENSMUSG( | -0.56379 | 1.12E-12 | YES | NO | NA |
| Cep57 | ENSMUSG( | ENSMUSG( | 0.057412 | 0.581023 | NO | NO | NA |
| Irf2bpl | ENSMUSG( | ENSMUSG( | -0.54172 | 2.97E-10 | YES | NO | NA |
| Bap1 | ENSMUSG( | ENSMUSG( | 0.009931 | 0.890997 | NO | NO | NA |
| Abhd8 | ENSMUSG( | ENSMUSG( | 0.184771 | 0.05772 | NO | NO | NA |
| Vezf1 | ENSMUSG( | ENSMUSG( | -0.43398 | 8.49E-08 | YES | NO | NA |
| Tgtp2 | ENSMUSG( | ENSMUSG( | -0.33509 | 0.017867 | YES | NO | NA |
| Gpr174 | ENSMUSG( | ENSMUSG( | -0.3228 | 0.001709 | YES | NO | NA |
| Susd6 | ENSMUSG( | ENSMUSG( | -0.67326 | 1.48E-13 | YES | NO | NA |
| Arid4b | ENSMUSG( | ENSMUSG( | -0.21619 | 0.036476 | YES | NO | NA |
| Slc29a1 | ENSMUSG( | ENSMUSG( | 0.437084 | 1.99E-07 | YES | NO | NA |
| Ulbp1 | ENSMUSG( | ENSMUSG( | -0.13748 | 0.327153 | NO | NO | NA |
| Elmo2 | ENSMUSG( | ENSMUSG( | -0.22417 | 1.00E-05 | YES | NO | NA |
| Rnh1 | ENSMUSG( | ENSMUSG( | -0.01463 | 0.814735 | NO | NO | NA |
| Gigyf1 | ENSMUSG( | ENSMUSG( | -0.1457 | 0.140198 | NO | NO | NA |
| Selenoh | ENSMUSG( | ENSMUSG( | 0.340463 | 1.43E-05 | YES | NO | NA |
| Akirin1 | ENSMUSG( | ENSMUSG( | -0.05882 | 0.542733 | NO | NO | NA |
| Ciapin1 | ENSMUSG( | ENSMUSG( | 0.220681 | 0.007881 | YES | NO | NA |
| Paxbp1 | ENSMUSG( | ENSMUSG( | -0.03877 | 0.515683 | NO | NO | NA |
| Rftn1 | ENSMUSG( | ENSMUSG( | 0.016618 | 0.78461 | NO | NO | NA |
| Cluh | ENSMUSG( | ENSMUSG( | 0.790855 | 1.72E-09 | YES | NO | NA |
| Zranb2 | ENSMUSG( | ENSMUSG( | -0.08676 | 0.280969 | NO | NO | NA |
| Get4 | ENSMUSG( | ENSMUSG( | 0.381808 | 1.73E-07 | YES | NO | NA |
| Supt4a | ENSMUSG( | ENSMUSG( | 0.014058 | 0.853511 | NO | NO | NA |
| Crtc2 | ENSMUSG( | ENSMUSG( | -0.1036 | 0.209948 | NO | NO | NA |
| Zfp236 | ENSMUSG( | ENSMUSG( | -0.41923 | 2.18E-06 | YES | NO | NA |
| Pkd1 | ENSMUSG( | ENSMUSG( | -0.45309 | 1.99E-06 | YES | NO | NA |
| Emsy | ENSMUSG( | ENSMUSG( | -0.35051 | 4.10E-08 | YES | NO | NA |
| Glrx3 | ENSMUSG( | ENSMUSG( | 0.326029 | 0.000286 | YES | NO | NA |
| Ash2l | ENSMUSG( | ENSMUSG( | 0.349981 | 3.27E-07 | YES | NO | NA |
| Arid2 | ENSMUSG( | ENSMUSG( | -0.05419 | 0.575254 | NO | NO | NA |

| | | | | | | | |
|---|---|---|---|---|---|---|---|
| Ago1 | ENSMUSG( | ENSMUSG( | -0.13444 | 0.102928 | NO | NO | NA |
| Mtf2 | ENSMUSG( | ENSMUSG( | 0.165588 | 0.028298 | YES | NO | NA |
| Trim39 | ENSMUSG( | ENSMUSG( | -0.16224 | 0.058323 | NO | NO | NA |
| Kcnk5 | ENSMUSG( | ENSMUSG( | -0.60308 | 5.12E-11 | YES | NO | NA |
| Herc4 | ENSMUSG( | ENSMUSG( | -0.09521 | 0.159533 | NO | NO | NA |
| Sec13 | ENSMUSG( | ENSMUSG( | 0.167725 | 0.002973 | YES | NO | NA |
| Anxa5 | ENSMUSG( | ENSMUSG( | -0.08187 | 0.478799 | NO | NO | NA |
| Art2b | ENSMUSG( | ENSMUSG( | -1.28784 | 1.01E-20 | YES | YES | Downregulated |
| Hltf | ENSMUSG( | ENSMUSG( | -0.27019 | 0.000168 | YES | NO | NA |
| Gem | ENSMUSG( | ENSMUSG( | -0.73132 | 1.50E-09 | YES | NO | NA |
| Cdipt | ENSMUSG( | ENSMUSG( | -0.29687 | 1.17E-05 | YES | NO | NA |
| Ubn2 | ENSMUSG( | ENSMUSG( | -0.46499 | 1.94E-09 | YES | NO | NA |
| Fam76b | ENSMUSG( | ENSMUSG( | -0.10625 | 0.30234 | NO | NO | NA |
| Dhx30 | ENSMUSG( | ENSMUSG( | 0.001815 | 0.975184 | NO | NO | NA |
| Sdf2l1 | ENSMUSG( | ENSMUSG( | 0.103813 | 0.442832 | NO | NO | NA |
| Nus1 | ENSMUSG( | ENSMUSG( | 0.154595 | 0.019704 | YES | NO | NA |
| Prmt5 | ENSMUSG( | ENSMUSG( | 0.372118 | 6.97E-05 | YES | NO | NA |
| Gpr55 | ENSMUSG( | ENSMUSG( | -0.52027 | 0.003991 | YES | NO | NA |
| Ccdc124 | ENSMUSG( | ENSMUSG( | 0.225247 | 0.049347 | YES | NO | NA |
| Acaca | ENSMUSG( | ENSMUSG( | 0.481429 | 2.29E-09 | YES | NO | NA |
| Glul | ENSMUSG( | ENSMUSG( | -0.32325 | 2.13E-06 | YES | NO | NA |
| Nudcd3 | ENSMUSG( | ENSMUSG( | 0.021897 | 0.819751 | NO | NO | NA |
| Rev3l | ENSMUSG( | ENSMUSG( | -0.25108 | 0.007585 | YES | NO | NA |
| Wdr76 | ENSMUSG( | ENSMUSG( | 0.006229 | 0.939403 | NO | NO | NA |
| Ppp2r5e | ENSMUSG( | ENSMUSG( | -0.18078 | 0.011599 | YES | NO | NA |
| Tbk1 | ENSMUSG( | ENSMUSG( | -0.14245 | 0.012449 | YES | NO | NA |
| Hexim1 | ENSMUSG( | ENSMUSG( | -0.00689 | 0.964687 | NO | NO | NA |
| Dennd2d | ENSMUSG( | ENSMUSG( | -0.09242 | 0.356061 | NO | NO | NA |
| Btbd1 | ENSMUSG( | ENSMUSG( | 0.178305 | 0.01101 | YES | NO | NA |
| Gm2a | ENSMUSG( | ENSMUSG( | 0.024598 | 0.808819 | NO | NO | NA |
| Cnot2 | ENSMUSG( | ENSMUSG( | -0.13174 | 0.187151 | NO | NO | NA |
| Runx1 | ENSMUSG( | ENSMUSG( | -0.52013 | 1.43E-09 | YES | NO | NA |
| Cox7b | ENSMUSG( | ENSMUSG( | 0.196909 | 0.035196 | YES | NO | NA |
| Faah | ENSMUSG( | ENSMUSG( | -0.18709 | 0.04053 | YES | NO | NA |
| Eed | ENSMUSG( | ENSMUSG( | 0.0573 | 0.501479 | NO | NO | NA |
| Dedd2 | ENSMUSG( | ENSMUSG( | -0.44888 | 2.61E-08 | YES | NO | NA |
| Trbv3 | ENSMUSG( | ENSMUSG( | -0.44561 | 0.059292 | NO | NO | NA |
| Ralgapa2 | ENSMUSG( | ENSMUSG( | -0.284 | 0.000891 | YES | NO | NA |
| Nt5c | ENSMUSG( | ENSMUSG( | 0.204237 | 0.029202 | YES | NO | NA |
| Zmat2 | ENSMUSG( | ENSMUSG( | -0.06402 | 0.581127 | NO | NO | NA |
| Flot2 | ENSMUSG( | ENSMUSG( | -0.02049 | 0.808313 | NO | NO | NA |
| Kdm3a | ENSMUSG( | ENSMUSG( | -0.20982 | 0.081891 | NO | NO | NA |
| Rtel1 | ENSMUSG( | ENSMUSG( | 0.124521 | 0.078072 | NO | NO | NA |
| Agps | ENSMUSG( | ENSMUSG( | -0.0087 | 0.847174 | NO | NO | NA |
| Nsmce4a | ENSMUSG( | ENSMUSG( | 0.052462 | 0.422059 | NO | NO | NA |
| Kdm1a | ENSMUSG( | ENSMUSG( | -0.02998 | 0.588955 | NO | NO | NA |
| Rbm15b | ENSMUSG( | ENSMUSG( | -0.13431 | 0.002399 | YES | NO | NA |

| | | | | | | | |
|---|---|---|---|---|---|---|---|
| Uqcrh | ENSMUSG( | ENSMUSG( | -0.09248 | 0.16902 | NO | NO | NA |
| Gm8369 | ENSMUSG( | ENSMUSG( | -0.04438 | 0.776895 | NO | NO | NA |
| Pkmyt1 | ENSMUSG( | ENSMUSG( | 0.184585 | 0.000444 | YES | NO | NA |
| Snrpd1 | ENSMUSG( | ENSMUSG( | 0.294314 | 0.068369 | NO | NO | NA |
| Nsfl1c | ENSMUSG( | ENSMUSG( | 0.080426 | 0.377352 | NO | NO | NA |
| Hat1 | ENSMUSG( | ENSMUSG( | 0.206666 | 0.005113 | YES | NO | NA |
| Cdca7l | ENSMUSG( | ENSMUSG( | -0.08744 | 0.036693 | YES | NO | NA |
| Nrf1 | ENSMUSG( | ENSMUSG( | -0.06093 | 0.418988 | NO | NO | NA |
| Rassf1 | ENSMUSG( | ENSMUSG( | -0.19767 | 0.001845 | YES | NO | NA |
| Ccng2 | ENSMUSG( | ENSMUSG( | -0.27859 | 0.001646 | YES | NO | NA |
| Pold3 | ENSMUSG( | ENSMUSG( | 0.004897 | 0.948511 | NO | NO | NA |
| Mbd6 | ENSMUSG( | ENSMUSG( | -0.41871 | 2.20E-12 | YES | NO | NA |
| Pan2 | ENSMUSG( | ENSMUSG( | 0.033585 | 0.778769 | NO | NO | NA |
| Ndst1 | ENSMUSG( | ENSMUSG( | 0.395143 | 0.000134 | YES | NO | NA |
| Llph | ENSMUSG( | ENSMUSG( | 0.306825 | 0.024177 | YES | NO | NA |
| Gm28438 | ENSMUSG( | ENSMUSG( | -0.08969 | 0.679983 | NO | NO | NA |
| Mms19 | ENSMUSG( | ENSMUSG( | -0.0574 | 0.593553 | NO | NO | NA |
| Psmb2 | ENSMUSG( | ENSMUSG( | 0.253127 | 2.08E-05 | YES | NO | NA |
| Aldh9a1 | ENSMUSG( | ENSMUSG( | 0.180963 | 0.003392 | YES | NO | NA |
| Fkbp3 | ENSMUSG( | ENSMUSG( | 0.167083 | 0.013787 | YES | NO | NA |
| Jmjd1c | ENSMUSG( | ENSMUSG( | -0.144 | 0.021565 | YES | NO | NA |
| Dctn5 | ENSMUSG( | ENSMUSG( | -0.11342 | 0.155265 | NO | NO | NA |
| Bpnt2 | ENSMUSG( | ENSMUSG( | -0.09221 | 0.351949 | NO | NO | NA |
| Rad23a | ENSMUSG( | ENSMUSG( | -0.06742 | 0.319847 | NO | NO | NA |
| Pnpla2 | ENSMUSG( | ENSMUSG( | -0.03916 | 0.669375 | NO | NO | NA |
| Rcor1 | ENSMUSG( | ENSMUSG( | -0.14653 | 0.015818 | YES | NO | NA |
| Nmi | ENSMUSG( | ENSMUSG( | -0.25099 | 0.004099 | YES | NO | NA |
| Mllt10 | ENSMUSG( | ENSMUSG( | -0.08289 | 0.375234 | NO | NO | NA |
| Ccnt1 | ENSMUSG( | ENSMUSG( | -0.20055 | 0.000142 | YES | NO | NA |
| Stat5a | ENSMUSG( | ENSMUSG( | 0.239129 | 0.000159 | YES | NO | NA |
| Ramac | ENSMUSG( | ENSMUSG( | -0.009 | 0.953489 | NO | NO | NA |
| Rnf126 | ENSMUSG( | ENSMUSG( | 0.264906 | 8.49E-05 | YES | NO | NA |
| Stxbp5 | ENSMUSG( | ENSMUSG( | -0.04766 | 0.52342 | NO | NO | NA |
| Reps1 | ENSMUSG( | ENSMUSG( | -0.26947 | 0.000332 | YES | NO | NA |
| Camta2 | ENSMUSG( | ENSMUSG( | -0.27963 | 0.001334 | YES | NO | NA |
| Ncoa4 | ENSMUSG( | ENSMUSG( | 0.014336 | 0.86261 | NO | NO | NA |
| Nol6 | ENSMUSG( | ENSMUSG( | 0.322992 | 0.000306 | YES | NO | NA |
| Traf1 | ENSMUSG( | ENSMUSG( | -0.25818 | 0.000447 | YES | NO | NA |
| Ighm | ENSMUSG( | ENSMUSG( | -0.05521 | 0.797347 | NO | NO | NA |
| Cep55 | ENSMUSG( | ENSMUSG( | 0.144092 | 0.131748 | NO | NO | NA |
| Fbxw8 | ENSMUSG( | ENSMUSG( | 0.248668 | 0.001165 | YES | NO | NA |
| Slx4 | ENSMUSG( | ENSMUSG( | -0.07543 | 0.288272 | NO | NO | NA |
| Sharpin | ENSMUSG( | ENSMUSG( | 0.012172 | 0.915251 | NO | NO | NA |
| Sdhd | ENSMUSG( | ENSMUSG( | 0.397839 | 1.00E-06 | YES | NO | NA |
| Ermp1 | ENSMUSG( | ENSMUSG( | -0.18902 | 0.001816 | YES | NO | NA |
| Lonp2 | ENSMUSG( | ENSMUSG( | -0.37361 | 6.92E-09 | YES | NO | NA |
| St14 | ENSMUSG( | ENSMUSG( | 0.547959 | 9.45E-07 | YES | NO | NA |

| | | | | | | | |
|---|---|---|---|---|---|---|---|
| Lef1 | ENSMUSG( | ENSMUSG( | 0.437733 | 0.003854 | YES | NO | NA |
| Senp2 | ENSMUSG( | ENSMUSG( | -0.21052 | 0.000233 | YES | NO | NA |
| Fry | ENSMUSG( | ENSMUSG( | -0.79633 | 5.15E-10 | YES | NO | NA |
| Spg21 | ENSMUSG( | ENSMUSG( | 0.361303 | 0.084092 | NO | NO | NA |
| Zfand6 | ENSMUSG( | ENSMUSG( | -0.3123 | 1.51E-05 | YES | NO | NA |
| Elp2 | ENSMUSG( | ENSMUSG( | 0.086808 | 0.07313 | NO | NO | NA |
| Arhgap15 | ENSMUSG( | ENSMUSG( | -0.09329 | 0.298442 | NO | NO | NA |
| Gm10252 | ENSMUSG( | ENSMUSG( | -0.07448 | 0.429185 | NO | NO | NA |
| Epc1 | ENSMUSG( | ENSMUSG( | -0.36626 | 9.53E-07 | YES | NO | NA |
| Nhlrc2 | ENSMUSG( | ENSMUSG( | -0.13359 | 0.111049 | NO | NO | NA |
| Tor1aip2 | ENSMUSG( | ENSMUSG( | -0.27805 | 2.16E-06 | YES | NO | NA |
| Cant1 | ENSMUSG( | ENSMUSG( | -0.31556 | 1.16E-11 | YES | NO | NA |
| Dera | ENSMUSG( | ENSMUSG( | 0.160854 | 0.044179 | YES | NO | NA |
| Oxct1 | ENSMUSG( | ENSMUSG( | 0.1798 | 0.131363 | NO | NO | NA |
| Reep4 | ENSMUSG( | ENSMUSG( | 0.119492 | 0.014751 | YES | NO | NA |
| Rock2 | ENSMUSG( | ENSMUSG( | -0.17094 | 0.011977 | YES | NO | NA |
| Cep120 | ENSMUSG( | ENSMUSG( | -0.19836 | 0.004195 | YES | NO | NA |
| Ttf2 | ENSMUSG( | ENSMUSG( | 0.12537 | 0.041588 | YES | NO | NA |
| 1810037I1 | ENSMUSG( | ENSMUSG( | -0.09162 | 0.410248 | NO | NO | NA |
| Parp10 | ENSMUSG( | ENSMUSG( | -0.36237 | 1.70E-05 | YES | NO | NA |
| Rnf6 | ENSMUSG( | ENSMUSG( | -0.16286 | 0.037948 | YES | NO | NA |
| Snrnp48 | ENSMUSG( | ENSMUSG( | -0.22018 | 8.49E-05 | YES | NO | NA |
| Ccdc12 | ENSMUSG( | ENSMUSG( | -0.11831 | 0.174954 | NO | NO | NA |
| Spty2d1 | ENSMUSG( | ENSMUSG( | -0.18773 | 0.05291 | NO | NO | NA |
| Tm9sf1 | ENSMUSG( | ENSMUSG( | -0.04509 | 0.354923 | NO | NO | NA |
| Ifi214 | ENSMUSG( | ENSMUSG( | -1.22987 | 1.36E-10 | YES | YES | Downregulated |
| Use1 | ENSMUSG( | ENSMUSG( | 0.02304 | 0.842144 | NO | NO | NA |
| Tnip1 | ENSMUSG( | ENSMUSG( | -0.2972 | 1.43E-05 | YES | NO | NA |
| Mast2 | ENSMUSG( | ENSMUSG( | -0.01349 | 0.854408 | NO | NO | NA |
| Arpc5l | ENSMUSG( | ENSMUSG( | 0.09008 | 0.157795 | NO | NO | NA |
| Riox2 | ENSMUSG( | ENSMUSG( | -0.11206 | 0.043941 | YES | NO | NA |
| Rb1 | ENSMUSG( | ENSMUSG( | -0.25004 | 0.018239 | YES | NO | NA |
| Dpp4 | ENSMUSG( | ENSMUSG( | -0.17811 | 0.00669 | YES | NO | NA |
| Phf11b | ENSMUSG( | ENSMUSG( | -0.46623 | 1.62E-13 | YES | NO | NA |
| L1cam | ENSMUSG( | ENSMUSG( | -0.56592 | 1.01E-14 | YES | NO | NA |
| Glmp | ENSMUSG( | ENSMUSG( | -0.42221 | 5.84E-11 | YES | NO | NA |
| Gm47761 | ENSMUSG( | ENSMUSG( | 0.866476 | 2.22E-05 | YES | NO | NA |
| Kdm2b | ENSMUSG( | ENSMUSG( | 0.005202 | 0.970155 | NO | NO | NA |
| Smarcb1 | ENSMUSG( | ENSMUSG( | 0.135155 | 0.011895 | YES | NO | NA |
| Trerf1 | ENSMUSG( | ENSMUSG( | -0.13199 | 0.374884 | NO | NO | NA |
| Ncald | ENSMUSG( | ENSMUSG( | -0.23832 | 2.83E-05 | YES | NO | NA |
| Tubgcp4 | ENSMUSG( | ENSMUSG( | -0.18335 | 0.00218 | YES | NO | NA |
| Sft2d2 | ENSMUSG( | ENSMUSG( | -0.18505 | 0.024014 | YES | NO | NA |
| Psmd6 | ENSMUSG( | ENSMUSG( | 0.255695 | 0.000691 | YES | NO | NA |
| Hsd11b1 | ENSMUSG( | ENSMUSG( | -0.29043 | 1.42E-05 | YES | NO | NA |
| Esco1 | ENSMUSG( | ENSMUSG( | -0.17012 | 0.063052 | NO | NO | NA |
| Ppp2r1b | ENSMUSG( | ENSMUSG( | -0.01655 | 0.807141 | NO | NO | NA |

| | | | | | | | |
|---|---|---|---|---|---|---|---|
| Dync1li1 | ENSMUSG( | ENSMUSG( | -0.14457 | 0.024941 | YES | NO | NA |
| Mrs2 | ENSMUSG( | ENSMUSG( | -0.11179 | 0.2254 | NO | NO | NA |
| Sfxn3 | ENSMUSG( | ENSMUSG( | -0.58189 | 3.43E-09 | YES | NO | NA |
| Sec61b | ENSMUSG( | ENSMUSG( | 0.037963 | 0.706841 | NO | NO | NA |
| Tbc1d22b | ENSMUSG( | ENSMUSG( | -0.24691 | 9.74E-07 | YES | NO | NA |
| Csnk2a2 | ENSMUSG( | ENSMUSG( | -0.10106 | 0.02687 | YES | NO | NA |
| Atp6v1b2 | ENSMUSG( | ENSMUSG( | -0.10432 | 0.126492 | NO | NO | NA |
| Zbtb32 | ENSMUSG( | ENSMUSG( | 0.169187 | 0.272959 | NO | NO | NA |
| Txnl1 | ENSMUSG( | ENSMUSG( | 0.220453 | 0.000377 | YES | NO | NA |
| Btaf1 | ENSMUSG( | ENSMUSG( | -0.35514 | 0.000341 | YES | NO | NA |
| Pkp3 | ENSMUSG( | ENSMUSG( | -0.0197 | 0.899573 | NO | NO | NA |
| Rps6kb1 | ENSMUSG( | ENSMUSG( | 0.047691 | 0.567556 | NO | NO | NA |
| Extl3 | ENSMUSG( | ENSMUSG( | -0.02116 | 0.783353 | NO | NO | NA |
| Wdr6 | ENSMUSG( | ENSMUSG( | 0.046269 | 0.593553 | NO | NO | NA |
| Nopchap1 | ENSMUSG( | ENSMUSG( | 0.166211 | 0.010467 | YES | NO | NA |
| Pmf1 | ENSMUSG( | ENSMUSG( | 0.294295 | 0.000107 | YES | NO | NA |
| Atp9b | ENSMUSG( | ENSMUSG( | -0.00741 | 0.949307 | NO | NO | NA |
| Cdk2ap2 | ENSMUSG( | ENSMUSG( | -0.13423 | 0.018023 | YES | NO | NA |
| Pum3 | ENSMUSG( | ENSMUSG( | 0.325574 | 0.001735 | YES | NO | NA |
| Uimc1 | ENSMUSG( | ENSMUSG( | -0.20032 | 0.007826 | YES | NO | NA |
| Gmnn | ENSMUSG( | ENSMUSG( | 0.499782 | 3.97E-09 | YES | NO | NA |
| Pfas | ENSMUSG( | ENSMUSG( | 0.484361 | 3.62E-07 | YES | NO | NA |
| Ppat | ENSMUSG( | ENSMUSG( | 0.250635 | 0.005218 | YES | NO | NA |
| Klf10 | ENSMUSG( | ENSMUSG( | -0.48483 | 6.42E-07 | YES | NO | NA |
| Patl1 | ENSMUSG( | ENSMUSG( | -0.19058 | 0.014247 | YES | NO | NA |
| Ngly1 | ENSMUSG( | ENSMUSG( | -0.16669 | 0.065378 | NO | NO | NA |
| Denr | ENSMUSG( | ENSMUSG( | 0.253407 | 0.025988 | YES | NO | NA |
| Cacybp | ENSMUSG( | ENSMUSG( | 0.088518 | 0.450269 | NO | NO | NA |
| Hirip3 | ENSMUSG( | ENSMUSG( | 0.545031 | 0.00349 | YES | NO | NA |
| Twsg1 | ENSMUSG( | ENSMUSG( | -0.11483 | 0.369627 | NO | NO | NA |
| Tcf19 | ENSMUSG( | ENSMUSG( | 0.212341 | 0.034593 | YES | NO | NA |
| Dhx8 | ENSMUSG( | ENSMUSG( | -0.13263 | 0.001339 | YES | NO | NA |
| Aph1a | ENSMUSG( | ENSMUSG( | 0.003867 | 0.961255 | NO | NO | NA |
| Prr5l | ENSMUSG( | ENSMUSG( | -1.55743 | 1.87E-06 | YES | YES | Downregulated |
| Kif2c | ENSMUSG( | ENSMUSG( | 0.328355 | 6.13E-05 | YES | NO | NA |
| Rnaseh2b | ENSMUSG( | ENSMUSG( | 0.383576 | 6.06E-08 | YES | NO | NA |
| Dip2b | ENSMUSG( | ENSMUSG( | -0.00196 | 0.984402 | NO | NO | NA |
| Ssh1 | ENSMUSG( | ENSMUSG( | -0.60338 | 7.17E-08 | YES | NO | NA |
| Taf4 | ENSMUSG( | ENSMUSG( | -0.35403 | 6.39E-09 | YES | NO | NA |
| Sec11a | ENSMUSG( | ENSMUSG( | -0.1138 | 0.005071 | YES | NO | NA |
| Slc25a11 | ENSMUSG( | ENSMUSG( | -0.15101 | 0.018241 | YES | NO | NA |
| Ola1 | ENSMUSG( | ENSMUSG( | 0.341717 | 0.001719 | YES | NO | NA |
| Plk4 | ENSMUSG( | ENSMUSG( | 0.231588 | 0.000499 | YES | NO | NA |
| Asap1 | ENSMUSG( | ENSMUSG( | -0.51872 | 4.73E-08 | YES | NO | NA |
| Cramp1 | ENSMUSG( | ENSMUSG( | -0.39251 | 1.83E-12 | YES | NO | NA |
| Ints3 | ENSMUSG( | ENSMUSG( | -0.24043 | 1.97E-06 | YES | NO | NA |
| Fgd3 | ENSMUSG( | ENSMUSG( | -0.62581 | 8.13E-12 | YES | NO | NA |

| | | | | | | | |
|---|---|---|---|---|---|---|---|
| Ice1 | ENSMUSG( | ENSMUSG( | -0.15076 | 0.020517 | YES | NO | NA |
| Camk4 | ENSMUSG( | ENSMUSG( | 0.11954 | 0.231542 | NO | NO | NA |
| Peds1 | ENSMUSG( | ENSMUSG( | -0.12637 | 0.083555 | NO | NO | NA |
| Pqlc3 | ENSMUSG( | ENSMUSG( | -0.33317 | 0.000428 | YES | NO | NA |
| Samd1 | ENSMUSG( | ENSMUSG( | 0.062586 | 0.670137 | NO | NO | NA |
| Ythdf1 | ENSMUSG( | ENSMUSG( | -0.04853 | 0.46692 | NO | NO | NA |
| Pigs | ENSMUSG( | ENSMUSG( | -0.184 | 0.002706 | YES | NO | NA |
| Kcmf1 | ENSMUSG( | ENSMUSG( | 0.04552 | 0.446235 | NO | NO | NA |
| Osbpl5 | ENSMUSG( | ENSMUSG( | -0.5174 | 1.18E-10 | YES | NO | NA |
| Mt1 | ENSMUSG( | ENSMUSG( | 1.883598 | 1.50E-07 | YES | YES | Upregulated |
| Lamp1 | ENSMUSG( | ENSMUSG( | 0.146761 | 0.130502 | NO | NO | NA |
| Tollip | ENSMUSG( | ENSMUSG( | -0.10208 | 0.230423 | NO | NO | NA |
| Gtpbp4 | ENSMUSG( | ENSMUSG( | 0.285213 | 1.89E-05 | YES | NO | NA |
| Diaph3 | ENSMUSG( | ENSMUSG( | 0.223299 | 0.025426 | YES | NO | NA |
| Fbxo5 | ENSMUSG( | ENSMUSG( | 0.093587 | 0.250201 | NO | NO | NA |
| 4833420G | ENSMUSG( | ENSMUSG( | -0.07201 | 0.44719 | NO | NO | NA |
| Syngr2 | ENSMUSG( | ENSMUSG( | 0.060798 | 0.286533 | NO | NO | NA |
| Mrpl9 | ENSMUSG( | ENSMUSG( | 0.13316 | 0.088135 | NO | NO | NA |
| Psmc6 | ENSMUSG( | ENSMUSG( | 0.188419 | 0.007081 | YES | NO | NA |
| Tbrg1 | ENSMUSG( | ENSMUSG( | -0.06988 | 0.369971 | NO | NO | NA |
| Dcaf1 | ENSMUSG( | ENSMUSG( | -0.18197 | 0.001439 | YES | NO | NA |
| Man1a2 | ENSMUSG( | ENSMUSG( | -0.15541 | 0.021703 | YES | NO | NA |
| Adh5 | ENSMUSG( | ENSMUSG( | 0.108985 | 0.084644 | NO | NO | NA |
| Cks1b | ENSMUSG( | ENSMUSG( | 0.313606 | 0.000116 | YES | NO | NA |
| Gripap1 | ENSMUSG( | ENSMUSG( | -0.30314 | 1.09E-05 | YES | NO | NA |
| Myef2 | ENSMUSG( | ENSMUSG( | -0.01293 | 0.860491 | NO | NO | NA |
| Prep | ENSMUSG( | ENSMUSG( | 0.40081 | 1.57E-05 | YES | NO | NA |
| Ncf1 | ENSMUSG( | ENSMUSG( | -0.56132 | 0.000152 | YES | NO | NA |
| Ndufv2 | ENSMUSG( | ENSMUSG( | 0.175558 | 0.126451 | NO | NO | NA |
| Magt1 | ENSMUSG( | ENSMUSG( | -0.16041 | 0.095935 | NO | NO | NA |
| Ak6 | ENSMUSG( | ENSMUSG( | 0.465327 | 1.16E-05 | YES | NO | NA |
| Ccser2 | ENSMUSG( | ENSMUSG( | -0.46142 | 4.08E-08 | YES | NO | NA |
| Uri1 | ENSMUSG( | ENSMUSG( | 0.30403 | 2.01E-05 | YES | NO | NA |
| Arfgap2 | ENSMUSG( | ENSMUSG( | -0.01497 | 0.840839 | NO | NO | NA |
| Tubgcp2 | ENSMUSG( | ENSMUSG( | 0.158745 | 0.006764 | YES | NO | NA |
| Atp6v1a | ENSMUSG( | ENSMUSG( | 0.011543 | 0.841559 | NO | NO | NA |
| Tmco4 | ENSMUSG( | ENSMUSG( | 0.116417 | 0.107078 | NO | NO | NA |
| Ddx52 | ENSMUSG( | ENSMUSG( | 0.294005 | 0.000261 | YES | NO | NA |
| Ifi27 | ENSMUSG( | ENSMUSG( | -0.22141 | 0.001196 | YES | NO | NA |
| Clk4 | ENSMUSG( | ENSMUSG( | -0.16647 | 0.00478 | YES | NO | NA |
| Zdhhc21 | ENSMUSG( | ENSMUSG( | -0.03175 | 0.726189 | NO | NO | NA |
| Npepps | ENSMUSG( | ENSMUSG( | -0.07588 | 0.268921 | NO | NO | NA |
| Cnot10 | ENSMUSG( | ENSMUSG( | 0.242524 | 3.58E-05 | YES | NO | NA |
| Nt5c3b | ENSMUSG( | ENSMUSG( | -0.24877 | 0.00027 | YES | NO | NA |
| Qk | ENSMUSG( | ENSMUSG( | 0.199799 | 0.001196 | YES | NO | NA |
| Isg20l2 | ENSMUSG( | ENSMUSG( | 0.145999 | 0.038311 | YES | NO | NA |
| Cog5 | ENSMUSG( | ENSMUSG( | -0.23056 | 0.001176 | YES | NO | NA |

| | | | | | | | |
|---|---|---|---|---|---|---|---|
| Prkdc | ENSMUSG( | ENSMUSG( | -0.44073 | 3.65E-07 | YES | NO | NA |
| Tubgcp3 | ENSMUSG( | ENSMUSG( | -0.0213 | 0.75156 | NO | NO | NA |
| Vopp1 | ENSMUSG( | ENSMUSG( | -0.34937 | 0.000187 | YES | NO | NA |
| Ndufa9 | ENSMUSG( | ENSMUSG( | 0.205767 | 0.023941 | YES | NO | NA |
| Tufm | ENSMUSG( | ENSMUSG( | 0.243783 | 0.00173 | YES | NO | NA |
| Dhx36 | ENSMUSG( | ENSMUSG( | -0.02135 | 0.805944 | NO | NO | NA |
| Slc25a12 | ENSMUSG( | ENSMUSG( | -0.0798 | 0.175013 | NO | NO | NA |
| Rrp7a | ENSMUSG( | ENSMUSG( | 0.187556 | 0.002136 | YES | NO | NA |
| Ccny | ENSMUSG( | ENSMUSG( | 0.015163 | 0.869437 | NO | NO | NA |
| Hmgn1 | ENSMUSG( | ENSMUSG( | 0.375478 | 0.000581 | YES | NO | NA |
| Ddx18 | ENSMUSG( | ENSMUSG( | 0.290237 | 0.000329 | YES | NO | NA |
| 4930523C( | ENSMUSG( | ENSMUSG( | -0.11423 | 0.157881 | NO | NO | NA |
| Rab2a | ENSMUSG( | ENSMUSG( | -0.09094 | 0.100129 | NO | NO | NA |
| Arpc1a | ENSMUSG( | ENSMUSG( | 0.221982 | 0.012272 | YES | NO | NA |
| Zmpste24 | ENSMUSG( | ENSMUSG( | -0.0246 | 0.75192 | NO | NO | NA |
| Sgk1 | ENSMUSG( | ENSMUSG( | -0.13054 | 0.173595 | NO | NO | NA |
| Tbc1d1 | ENSMUSG( | ENSMUSG( | -0.44452 | 9.27E-11 | YES | NO | NA |
| Rbm26 | ENSMUSG( | ENSMUSG( | -0.15735 | 0.015174 | YES | NO | NA |
| Abcf2 | ENSMUSG( | ENSMUSG( | 0.416292 | 4.36E-07 | YES | NO | NA |
| Ppid | ENSMUSG( | ENSMUSG( | 0.01074 | 0.930307 | NO | NO | NA |
| ENSMUSG | ENSMUSG( | ENSMUSG( | -0.40767 | 1.83E-09 | YES | NO | NA |
| Mga | ENSMUSG( | ENSMUSG( | -0.30845 | 1.36E-06 | YES | NO | NA |
| Eif2a | ENSMUSG( | ENSMUSG( | 0.362615 | 3.75E-05 | YES | NO | NA |
| Mmd | ENSMUSG( | ENSMUSG( | -0.02845 | 0.803915 | NO | NO | NA |
| Wdtc1 | ENSMUSG( | ENSMUSG( | -0.25706 | 0.00012 | YES | NO | NA |
| Swi5 | ENSMUSG( | ENSMUSG( | 0.278859 | 0.000634 | YES | NO | NA |
| Unkl | ENSMUSG( | ENSMUSG( | -0.20684 | 0.005128 | YES | NO | NA |
| Hdac2 | ENSMUSG( | ENSMUSG( | 0.224986 | 0.013163 | YES | NO | NA |
| Ksr1 | ENSMUSG( | ENSMUSG( | 0.197523 | 0.005063 | YES | NO | NA |
| Smdt1 | ENSMUSG( | ENSMUSG( | 0.044051 | 0.735159 | NO | NO | NA |
| Isyna1 | ENSMUSG( | ENSMUSG( | 0.346409 | 2.47E-05 | YES | NO | NA |
| Pcmt1 | ENSMUSG( | ENSMUSG( | 0.027436 | 0.696107 | NO | NO | NA |
| Micos10 | ENSMUSG( | ENSMUSG( | 0.215182 | 0.003282 | YES | NO | NA |
| Snhg1 | ENSMUSG( | ENSMUSG( | 0.585378 | 2.62E-05 | YES | NO | NA |
| Snx2 | ENSMUSG( | ENSMUSG( | -0.16133 | 0.016495 | YES | NO | NA |
| Tmem248 | ENSMUSG( | ENSMUSG( | -0.05627 | 0.465617 | NO | NO | NA |
| Irf3 | ENSMUSG( | ENSMUSG( | -0.17263 | 0.006171 | YES | NO | NA |
| Smndc1 | ENSMUSG( | ENSMUSG( | -0.11945 | 0.176012 | NO | NO | NA |
| Cdc27 | ENSMUSG( | ENSMUSG( | 0.099486 | 0.1501 | NO | NO | NA |
| Ubr7 | ENSMUSG( | ENSMUSG( | 0.153661 | 0.031883 | YES | NO | NA |
| Socs4 | ENSMUSG( | ENSMUSG( | -0.47424 | 4.51E-11 | YES | NO | NA |
| Ikbke | ENSMUSG( | ENSMUSG( | -0.43535 | 4.08E-08 | YES | NO | NA |
| Lmtk2 | ENSMUSG( | ENSMUSG( | -0.31837 | 8.27E-11 | YES | NO | NA |
| Pqbp1 | ENSMUSG( | ENSMUSG( | 0.023121 | 0.863904 | NO | NO | NA |
| Nhp2 | ENSMUSG( | ENSMUSG( | 0.433229 | 0.000461 | YES | NO | NA |
| Ttc13 | ENSMUSG( | ENSMUSG( | -0.03007 | 0.660302 | NO | NO | NA |
| Bub1 | ENSMUSG( | ENSMUSG( | 0.172626 | 0.099771 | NO | NO | NA |

| | | | | | | | |
|---|---|---|---|---|---|---|---|
| Map3k2 | ENSMUSG( | ENSMUSG( | -0.42217 | 1.02E-06 | YES | NO | NA |
| Sting1 | ENSMUSG( | ENSMUSG( | -0.05107 | 0.421949 | NO | NO | NA |
| Pacsin1 | ENSMUSG( | ENSMUSG( | -0.10041 | 0.471033 | NO | NO | NA |
| Spin1 | ENSMUSG( | ENSMUSG( | -0.17943 | 0.001508 | YES | NO | NA |
| Aff1 | ENSMUSG( | ENSMUSG( | -0.58844 | 5.91E-09 | YES | NO | NA |
| Ap2a2 | ENSMUSG( | ENSMUSG( | -0.05778 | 0.300349 | NO | NO | NA |
| Sesn3 | ENSMUSG( | ENSMUSG( | -0.94089 | 1.16E-14 | YES | NO | NA |
| AI467606 | ENSMUSG( | ENSMUSG( | -0.52799 | 5.12E-10 | YES | NO | NA |
| Shkbp1 | ENSMUSG( | ENSMUSG( | -0.10085 | 0.153206 | NO | NO | NA |
| Trmt112 | ENSMUSG( | ENSMUSG( | 0.138213 | 0.028193 | YES | NO | NA |
| Rprd2 | ENSMUSG( | ENSMUSG( | -0.20322 | 0.005133 | YES | NO | NA |
| Lrch1 | ENSMUSG( | ENSMUSG( | -0.01398 | 0.921433 | NO | NO | NA |
| Wsb1 | ENSMUSG( | ENSMUSG( | 0.084423 | 0.178024 | NO | NO | NA |
| Ascc2 | ENSMUSG( | ENSMUSG( | 0.070287 | 0.207666 | NO | NO | NA |
| Sacm1l | ENSMUSG( | ENSMUSG( | -0.20109 | 0.006648 | YES | NO | NA |
| Atxn10 | ENSMUSG( | ENSMUSG( | 0.094394 | 0.254111 | NO | NO | NA |
| Washc5 | ENSMUSG( | ENSMUSG( | 0.010294 | 0.91236 | NO | NO | NA |
| Fbxo38 | ENSMUSG( | ENSMUSG( | -0.25343 | 0.000297 | YES | NO | NA |
| Ibtk | ENSMUSG( | ENSMUSG( | 0.063668 | 0.502427 | NO | NO | NA |
| Txn2 | ENSMUSG( | ENSMUSG( | 0.128108 | 0.077761 | NO | NO | NA |
| Ino80e | ENSMUSG( | ENSMUSG( | 0.212064 | 0.002128 | YES | NO | NA |
| Kntc1 | ENSMUSG( | ENSMUSG( | 0.124644 | 0.28302 | NO | NO | NA |
| Gpr132 | ENSMUSG( | ENSMUSG( | -0.1529 | 0.054803 | NO | NO | NA |
| Utp6 | ENSMUSG( | ENSMUSG( | 0.182669 | 0.013397 | YES | NO | NA |
| Gosr1 | ENSMUSG( | ENSMUSG( | -0.38185 | 4.81E-10 | YES | NO | NA |
| Xpo5 | ENSMUSG( | ENSMUSG( | 0.08364 | 0.134333 | NO | NO | NA |
| Tyms | ENSMUSG( | ENSMUSG( | 0.293562 | 0.000728 | YES | NO | NA |
| Emd | ENSMUSG( | ENSMUSG( | 0.148167 | 0.083554 | NO | NO | NA |
| Ssh2 | ENSMUSG( | ENSMUSG( | -0.10738 | 0.179905 | NO | NO | NA |
| Mdc1 | ENSMUSG( | ENSMUSG( | -0.38329 | 1.68E-05 | YES | NO | NA |
| Dmtf1 | ENSMUSG( | ENSMUSG( | -0.36342 | 1.07E-05 | YES | NO | NA |
| Erp44 | ENSMUSG( | ENSMUSG( | -0.08298 | 0.181956 | NO | NO | NA |
| Alcam | ENSMUSG( | ENSMUSG( | 0.315215 | 0.008328 | YES | NO | NA |
| Atg2a | ENSMUSG( | ENSMUSG( | -0.30529 | 2.49E-07 | YES | NO | NA |
| Mrpl18 | ENSMUSG( | ENSMUSG( | 0.20258 | 0.006512 | YES | NO | NA |
| Cebpz | ENSMUSG( | ENSMUSG( | 0.170373 | 0.031234 | YES | NO | NA |
| Med25 | ENSMUSG( | ENSMUSG( | 0.216221 | 0.009159 | YES | NO | NA |
| Aff4 | ENSMUSG( | ENSMUSG( | -0.24431 | 0.010008 | YES | NO | NA |
| Mkrn2 | ENSMUSG( | ENSMUSG( | 0.033285 | 0.822108 | NO | NO | NA |
| Phtf1 | ENSMUSG( | ENSMUSG( | 0.019028 | 0.843977 | NO | NO | NA |
| Ambra1 | ENSMUSG( | ENSMUSG( | -0.30255 | 3.30E-06 | YES | NO | NA |
| Man1b1 | ENSMUSG( | ENSMUSG( | -0.15069 | 0.006449 | YES | NO | NA |
| Cbx7 | ENSMUSG( | ENSMUSG( | 0.02325 | 0.842943 | NO | NO | NA |
| Cdca2 | ENSMUSG( | ENSMUSG( | -0.04263 | 0.610654 | NO | NO | NA |
| Tmem167 | ENSMUSG( | ENSMUSG( | -0.0121 | 0.861347 | NO | NO | NA |
| Hdhd5 | ENSMUSG( | ENSMUSG( | -0.67288 | 4.12E-12 | YES | NO | NA |
| H1f2 | ENSMUSG( | ENSMUSG( | 0.009647 | 0.964642 | NO | NO | NA |

| | | | | | | | |
|---|---|---|---|---|---|---|---|
| Lrch3 | ENSMUSG( | ENSMUSG( | -0.45823 | 1.12E-05 | YES | NO | NA |
| Fam13b | ENSMUSG( | ENSMUSG( | -0.57293 | 1.24E-09 | YES | NO | NA |
| Ppil4 | ENSMUSG( | ENSMUSG( | -0.03199 | 0.72311 | NO | NO | NA |
| Zfp335 | ENSMUSG( | ENSMUSG( | -0.28777 | 9.26E-07 | YES | NO | NA |
| Cops5 | ENSMUSG( | ENSMUSG( | 0.312034 | 7.15E-06 | YES | NO | NA |
| Rab4b | ENSMUSG( | ENSMUSG( | -0.00754 | 0.94486 | NO | NO | NA |
| Pitpnb | ENSMUSG( | ENSMUSG( | 0.076723 | 0.466465 | NO | NO | NA |
| Gnptg | ENSMUSG( | ENSMUSG( | -0.49939 | 3.74E-14 | YES | NO | NA |
| Slamf6 | ENSMUSG( | ENSMUSG( | 0.273754 | 0.025982 | YES | NO | NA |
| Tmem164 | ENSMUSG( | ENSMUSG( | -0.08573 | 0.067664 | NO | NO | NA |
| Pknox1 | ENSMUSG( | ENSMUSG( | -0.10574 | 0.031418 | YES | NO | NA |
| Cdk19 | ENSMUSG( | ENSMUSG( | -0.09622 | 0.423041 | NO | NO | NA |
| Gpsm2 | ENSMUSG( | ENSMUSG( | 0.578215 | 2.67E-10 | YES | NO | NA |
| Sh2d1a | ENSMUSG( | ENSMUSG( | -0.34556 | 0.000487 | YES | NO | NA |
| Clcn3 | ENSMUSG( | ENSMUSG( | 0.0184 | 0.870698 | NO | NO | NA |
| Rpia | ENSMUSG( | ENSMUSG( | 0.086689 | 0.195893 | NO | NO | NA |
| Zadh2 | ENSMUSG( | ENSMUSG( | 0.147536 | 0.230686 | NO | NO | NA |
| Nelfcd | ENSMUSG( | ENSMUSG( | 0.18006 | 0.012467 | YES | NO | NA |
| Xpot | ENSMUSG( | ENSMUSG( | 0.261823 | 1.91E-06 | YES | NO | NA |
| Thumpd1 | ENSMUSG( | ENSMUSG( | 0.307583 | 0.001299 | YES | NO | NA |
| Prps2 | ENSMUSG( | ENSMUSG( | -0.17545 | 0.00032 | YES | NO | NA |
| Stard4 | ENSMUSG( | ENSMUSG( | 0.162285 | 0.032844 | YES | NO | NA |
| Snrpb2 | ENSMUSG( | ENSMUSG( | 0.205728 | 0.08249 | NO | NO | NA |
| Tox2 | ENSMUSG( | ENSMUSG( | -0.53335 | 0.002467 | YES | NO | NA |
| Rxrb | ENSMUSG( | ENSMUSG( | -0.12269 | 0.054076 | NO | NO | NA |
| Nme1 | ENSMUSG( | ENSMUSG( | 0.539265 | 1.60E-07 | YES | NO | NA |
| Ticrr | ENSMUSG( | ENSMUSG( | 0.072101 | 0.354025 | NO | NO | NA |
| Casp2 | ENSMUSG( | ENSMUSG( | -0.01523 | 0.793495 | NO | NO | NA |
| Bcl2a1b | ENSMUSG( | ENSMUSG( | 0.199464 | 0.072898 | NO | NO | NA |
| Tmem14c | ENSMUSG( | ENSMUSG( | 0.155758 | 0.059198 | NO | NO | NA |
| Ttll12 | ENSMUSG( | ENSMUSG( | 0.073395 | 0.407513 | NO | NO | NA |
| Farp1 | ENSMUSG( | ENSMUSG( | -0.01504 | 0.859322 | NO | NO | NA |
| Rsf1 | ENSMUSG( | ENSMUSG( | -0.33217 | 3.49E-07 | YES | NO | NA |
| Uchl5 | ENSMUSG( | ENSMUSG( | 0.542309 | 5.03E-07 | YES | NO | NA |
| Ccdc97 | ENSMUSG( | ENSMUSG( | -0.21988 | 0.000989 | YES | NO | NA |
| Plagl2 | ENSMUSG( | ENSMUSG( | -0.17974 | 0.000894 | YES | NO | NA |
| Rab18 | ENSMUSG( | ENSMUSG( | -0.05479 | 0.636386 | NO | NO | NA |
| Colgalt1 | ENSMUSG( | ENSMUSG( | 0.238531 | 0.000847 | YES | NO | NA |
| Gpr65 | ENSMUSG( | ENSMUSG( | -0.05498 | 0.639044 | NO | NO | NA |
| Rnf26 | ENSMUSG( | ENSMUSG( | 0.037047 | 0.460937 | NO | NO | NA |
| Ireb2 | ENSMUSG( | ENSMUSG( | -0.07565 | 0.397531 | NO | NO | NA |
| Tomm34 | ENSMUSG( | ENSMUSG( | -0.17719 | 0.00349 | YES | NO | NA |
| Chd6 | ENSMUSG( | ENSMUSG( | -0.19092 | 0.165562 | NO | NO | NA |
| Cdc6 | ENSMUSG( | ENSMUSG( | 0.226476 | 0.061207 | NO | NO | NA |
| Selenoi | ENSMUSG( | ENSMUSG( | 0.031552 | 0.729736 | NO | NO | NA |
| Pon2 | ENSMUSG( | ENSMUSG( | 0.007916 | 0.943843 | NO | NO | NA |
| Farsa | ENSMUSG( | ENSMUSG( | 0.236675 | 0.00043 | YES | NO | NA |

| | | | | | | | |
|---|---|---|---|---|---|---|---|
| Ndufb8 | ENSMUSG( | ENSMUSG( | 0.160707 | 0.044696 | YES | NO | NA |
| 1600014C: | ENSMUSG( | ENSMUSG( | -0.68155 | 4.07E-09 | YES | NO | NA |
| Ccnf | ENSMUSG( | ENSMUSG( | 0.063187 | 0.440252 | NO | NO | NA |
| Zc3h12d | ENSMUSG( | ENSMUSG( | -0.05838 | 0.604642 | NO | NO | NA |
| Tank | ENSMUSG( | ENSMUSG( | -0.24456 | 0.006802 | YES | NO | NA |
| Atxn7l3b | ENSMUSG( | ENSMUSG( | -0.14265 | 0.052691 | NO | NO | NA |
| Hivep1 | ENSMUSG( | ENSMUSG( | -0.03874 | 0.763959 | NO | NO | NA |
| Brf1 | ENSMUSG( | ENSMUSG( | -0.01972 | 0.796012 | NO | NO | NA |
| Stx11 | ENSMUSG( | ENSMUSG( | -0.30365 | 0.000399 | YES | NO | NA |
| Pggt1b | ENSMUSG( | ENSMUSG( | -0.02386 | 0.770537 | NO | NO | NA |
| Vac14 | ENSMUSG( | ENSMUSG( | 0.078889 | 0.418544 | NO | NO | NA |
| Tsr1 | ENSMUSG( | ENSMUSG( | 0.444155 | 8.40E-05 | YES | NO | NA |
| Crnkl1 | ENSMUSG( | ENSMUSG( | 0.057022 | 0.438715 | NO | NO | NA |
| Bspry | ENSMUSG( | ENSMUSG( | 0.470691 | 1.52E-05 | YES | NO | NA |
| Gps1 | ENSMUSG( | ENSMUSG( | 0.215668 | 1.38E-06 | YES | NO | NA |
| Fem1a | ENSMUSG( | ENSMUSG( | -0.11752 | 0.175573 | NO | NO | NA |
| Chmp1b | ENSMUSG( | ENSMUSG( | -0.22081 | 0.016234 | YES | NO | NA |
| Ythdf2 | ENSMUSG( | ENSMUSG( | -0.03032 | 0.636498 | NO | NO | NA |
| Tjp2 | ENSMUSG( | ENSMUSG( | -0.04354 | 0.770862 | NO | NO | NA |
| Brca1 | ENSMUSG( | ENSMUSG( | 0.177461 | 0.029586 | YES | NO | NA |
| Nsf | ENSMUSG( | ENSMUSG( | -0.1252 | 0.030902 | YES | NO | NA |
| Faf2 | ENSMUSG( | ENSMUSG( | -0.15263 | 0.007478 | YES | NO | NA |
| Etv3 | ENSMUSG( | ENSMUSG( | -0.3043 | 3.78E-06 | YES | NO | NA |
| Mfn2 | ENSMUSG( | ENSMUSG( | 0.0459 | 0.640823 | NO | NO | NA |
| Myb | ENSMUSG( | ENSMUSG( | 1.108282 | 6.21E-07 | YES | YES | Upregulated |
| Git1 | ENSMUSG( | ENSMUSG( | -0.04609 | 0.60154 | NO | NO | NA |
| Rgl2 | ENSMUSG( | ENSMUSG( | -0.33796 | 1.34E-12 | YES | NO | NA |
| Zbed4 | ENSMUSG( | ENSMUSG( | -0.02857 | 0.746387 | NO | NO | NA |
| Ccndbp1 | ENSMUSG( | ENSMUSG( | -0.41452 | 2.69E-05 | YES | NO | NA |
| Bcl2l13 | ENSMUSG( | ENSMUSG( | -0.02315 | 0.617833 | NO | NO | NA |
| Cdca7 | ENSMUSG( | ENSMUSG( | 0.33861 | 0.002472 | YES | NO | NA |
| Rbm15 | ENSMUSG( | ENSMUSG( | -0.14635 | 0.173128 | NO | NO | NA |
| Hspa4l | ENSMUSG( | ENSMUSG( | -0.34168 | 4.68E-05 | YES | NO | NA |
| Ctnnbl1 | ENSMUSG( | ENSMUSG( | 0.234203 | 0.006133 | YES | NO | NA |
| Limk1 | ENSMUSG( | ENSMUSG( | -0.46176 | 1.12E-13 | YES | NO | NA |
| Imp4 | ENSMUSG( | ENSMUSG( | 0.152225 | 0.002365 | YES | NO | NA |
| Spryd3 | ENSMUSG( | ENSMUSG( | -0.1238 | 0.05409 | NO | NO | NA |
| Prdx5 | ENSMUSG( | ENSMUSG( | -0.01456 | 0.817607 | NO | NO | NA |
| Uqcrq | ENSMUSG( | ENSMUSG( | 0.301835 | 0.00015 | YES | NO | NA |
| Pcsk7 | ENSMUSG( | ENSMUSG( | -0.14999 | 0.005699 | YES | NO | NA |
| Nemp2 | ENSMUSG( | ENSMUSG( | -0.05609 | 0.6068 | NO | NO | NA |
| Naa10 | ENSMUSG( | ENSMUSG( | 0.320226 | 3.79E-05 | YES | NO | NA |
| Senp3 | ENSMUSG( | ENSMUSG( | 0.078902 | 0.174594 | NO | NO | NA |
| Nabp2 | ENSMUSG( | ENSMUSG( | 0.081879 | 0.311741 | NO | NO | NA |
| Tmem131 | ENSMUSG( | ENSMUSG( | 0.034518 | 0.709483 | NO | NO | NA |
| Tomm40 | ENSMUSG( | ENSMUSG( | 0.697463 | 7.76E-11 | YES | NO | NA |
| Ctdspl2 | ENSMUSG( | ENSMUSG( | -0.11694 | 0.140859 | NO | NO | NA |

| | | | | | | | |
|---|---|---|---|---|---|---|---|
| Zfp865 | ENSMUSG( | ENSMUSG( | -0.05886 | 0.454653 | NO | NO | NA |
| Tcf12 | ENSMUSG( | ENSMUSG( | -0.16483 | 0.019866 | YES | NO | NA |
| Pcm1 | ENSMUSG( | ENSMUSG( | -0.13626 | 0.14973 | NO | NO | NA |
| Abhd17a | ENSMUSG( | ENSMUSG( | 0.11638 | 0.060604 | NO | NO | NA |
| Trappc14 | ENSMUSG( | ENSMUSG( | -0.57531 | 8.49E-09 | YES | NO | NA |
| Rnf167 | ENSMUSG( | ENSMUSG( | -0.67992 | 1.86E-11 | YES | NO | NA |
| Wdr3 | ENSMUSG( | ENSMUSG( | 0.122482 | 0.072544 | NO | NO | NA |
| Necap2 | ENSMUSG( | ENSMUSG( | -0.10653 | 0.165846 | NO | NO | NA |
| Lpxn | ENSMUSG( | ENSMUSG( | -0.02588 | 0.774615 | NO | NO | NA |
| Mdfic | ENSMUSG( | ENSMUSG( | -0.11885 | 0.082067 | NO | NO | NA |
| Sh3glb2 | ENSMUSG( | ENSMUSG( | -0.09356 | 0.211211 | NO | NO | NA |
| Ripk3 | ENSMUSG( | ENSMUSG( | -0.08476 | 0.209647 | NO | NO | NA |
| Snrpa1 | ENSMUSG( | ENSMUSG( | 0.247183 | 0.001102 | YES | NO | NA |
| Atxn7 | ENSMUSG( | ENSMUSG( | -0.41713 | 7.04E-05 | YES | NO | NA |
| Shoc2 | ENSMUSG( | ENSMUSG( | -0.21324 | 0.004708 | YES | NO | NA |
| Pole4 | ENSMUSG( | ENSMUSG( | -0.33261 | 2.41E-11 | YES | NO | NA |
| Pold2 | ENSMUSG( | ENSMUSG( | 0.34746 | 2.65E-05 | YES | NO | NA |
| Ctdnep1 | ENSMUSG( | ENSMUSG( | -0.06273 | 0.24867 | NO | NO | NA |
| Snrpc | ENSMUSG( | ENSMUSG( | 0.138957 | 0.231472 | NO | NO | NA |
| Ergic3 | ENSMUSG( | ENSMUSG( | 0.16343 | 0.020523 | YES | NO | NA |
| Rab22a | ENSMUSG( | ENSMUSG( | -0.19821 | 0.00261 | YES | NO | NA |
| Lyar | ENSMUSG( | ENSMUSG( | 0.442531 | 0.010053 | YES | NO | NA |
| N4bp2l2 | ENSMUSG( | ENSMUSG( | -0.15428 | 0.007908 | YES | NO | NA |
| Cmas | ENSMUSG( | ENSMUSG( | -0.21314 | 0.00096 | YES | NO | NA |
| Mbd1 | ENSMUSG( | ENSMUSG( | -0.22257 | 0.000253 | YES | NO | NA |
| Vapb | ENSMUSG( | ENSMUSG( | -0.26277 | 1.56E-06 | YES | NO | NA |
| Prpf31 | ENSMUSG( | ENSMUSG( | 0.366123 | 0.000423 | YES | NO | NA |
| Lclat1 | ENSMUSG( | ENSMUSG( | -0.58768 | 1.09E-05 | YES | NO | NA |
| AU040320 | ENSMUSG( | ENSMUSG( | -0.56355 | 1.33E-12 | YES | NO | NA |
| Mtarc2 | ENSMUSG( | ENSMUSG( | 0.163766 | 0.029179 | YES | NO | NA |
| Ap1s1 | ENSMUSG( | ENSMUSG( | 0.243263 | 2.94E-05 | YES | NO | NA |
| Hspe1 | ENSMUSG( | ENSMUSG( | 0.134552 | 0.374884 | NO | NO | NA |
| Adsl | ENSMUSG( | ENSMUSG( | 0.315717 | 0.000322 | YES | NO | NA |
| Vps35l | ENSMUSG( | ENSMUSG( | -0.15608 | 0.072252 | NO | NO | NA |
| Cd38 | ENSMUSG( | ENSMUSG( | -0.90856 | 4.90E-09 | YES | NO | NA |
| Appl1 | ENSMUSG( | ENSMUSG( | 0.006974 | 0.930568 | NO | NO | NA |
| Arid5a | ENSMUSG( | ENSMUSG( | -0.23257 | 0.004972 | YES | NO | NA |
| Mfsd1 | ENSMUSG( | ENSMUSG( | -0.0631 | 0.576974 | NO | NO | NA |
| Lsm6 | ENSMUSG( | ENSMUSG( | 0.310198 | 8.07E-05 | YES | NO | NA |
| Oxa1l | ENSMUSG( | ENSMUSG( | 0.084284 | 0.364097 | NO | NO | NA |
| Spg7 | ENSMUSG( | ENSMUSG( | 0.067051 | 0.42939 | NO | NO | NA |
| Ube3a | ENSMUSG( | ENSMUSG( | -0.23754 | 0.00221 | YES | NO | NA |
| Ly75 | ENSMUSG( | ENSMUSG( | -0.26939 | 0.014477 | YES | NO | NA |
| Klhdc2 | ENSMUSG( | ENSMUSG( | 0.40402 | 0.000343 | YES | NO | NA |
| Sgpp1 | ENSMUSG( | ENSMUSG( | -0.16319 | 0.008225 | YES | NO | NA |
| Aprt | ENSMUSG( | ENSMUSG( | 0.265234 | 0.000498 | YES | NO | NA |
| Gle1 | ENSMUSG( | ENSMUSG( | 0.087195 | 0.11748 | NO | NO | NA |

| | | | | | | | |
|---|---|---|---|---|---|---|---|
| Zfp217 | ENSMUSG( | ENSMUSG( | -0.40119 | 3.06E-06 | YES | NO | NA |
| Tmem109 | ENSMUSG( | ENSMUSG( | 0.517107 | 6.66E-06 | YES | NO | NA |
| Phc2 | ENSMUSG( | ENSMUSG( | -0.23996 | 0.001755 | YES | NO | NA |
| Recql | ENSMUSG( | ENSMUSG( | 0.054465 | 0.535468 | NO | NO | NA |
| Mrps23 | ENSMUSG( | ENSMUSG( | 0.220287 | 0.014703 | YES | NO | NA |
| Wdr48 | ENSMUSG( | ENSMUSG( | -0.14509 | 0.002239 | YES | NO | NA |
| Vrk3 | ENSMUSG( | ENSMUSG( | -0.42751 | 5.44E-16 | YES | NO | NA |
| Ncln | ENSMUSG( | ENSMUSG( | -0.32058 | 1.00E-06 | YES | NO | NA |
| Cers4 | ENSMUSG( | ENSMUSG( | -0.70353 | 1.12E-12 | YES | NO | NA |
| Mrpl4 | ENSMUSG( | ENSMUSG( | -0.07223 | 0.328689 | NO | NO | NA |
| Washc2 | ENSMUSG( | ENSMUSG( | -0.11389 | 0.322595 | NO | NO | NA |
| Babam1 | ENSMUSG( | ENSMUSG( | 0.028413 | 0.651195 | NO | NO | NA |
| Cybc1 | ENSMUSG( | ENSMUSG( | -0.37302 | 6.58E-09 | YES | NO | NA |
| Ypel3 | ENSMUSG( | ENSMUSG( | -0.87412 | 1.06E-08 | YES | NO | NA |
| Htatsf1 | ENSMUSG( | ENSMUSG( | 0.104663 | 0.325038 | NO | NO | NA |
| Ndufv3 | ENSMUSG( | ENSMUSG( | 0.233911 | 0.000258 | YES | NO | NA |
| Tmem94 | ENSMUSG( | ENSMUSG( | -0.19657 | 0.000361 | YES | NO | NA |
| Ect2 | ENSMUSG( | ENSMUSG( | 0.161917 | 0.040867 | YES | NO | NA |
| Jarid2 | ENSMUSG( | ENSMUSG( | -0.17776 | 0.083821 | NO | NO | NA |
| Per1 | ENSMUSG( | ENSMUSG( | 0.333345 | 0.076756 | NO | NO | NA |
| Pdlim2 | ENSMUSG( | ENSMUSG( | -0.2077 | 1.53E-05 | YES | NO | NA |
| Cc2d1a | ENSMUSG( | ENSMUSG( | -0.1886 | 0.001072 | YES | NO | NA |
| Plekha5 | ENSMUSG( | ENSMUSG( | -0.23981 | 8.65E-05 | YES | NO | NA |
| Ankrd13c | ENSMUSG( | ENSMUSG( | -0.1341 | 0.037913 | YES | NO | NA |
| Ptpn2 | ENSMUSG( | ENSMUSG( | 0.081331 | 0.481493 | NO | NO | NA |
| Aimp1 | ENSMUSG( | ENSMUSG( | 0.341977 | 2.87E-05 | YES | NO | NA |
| Otud7b | ENSMUSG( | ENSMUSG( | -0.08164 | 0.402155 | NO | NO | NA |
| Lilrb4a | ENSMUSG( | ENSMUSG( | -0.31704 | 0.01379 | YES | NO | NA |
| Mak16 | ENSMUSG( | ENSMUSG( | 0.141031 | 0.165584 | NO | NO | NA |
| Bicra | ENSMUSG( | ENSMUSG( | -0.43484 | 1.09E-08 | YES | NO | NA |
| Fh1 | ENSMUSG( | ENSMUSG( | 0.447388 | 4.37E-09 | YES | NO | NA |
| Vps53 | ENSMUSG( | ENSMUSG( | 0.048823 | 0.558403 | NO | NO | NA |
| Sf3b5 | ENSMUSG( | ENSMUSG( | 0.272219 | 9.71E-06 | YES | NO | NA |
| Abcd1 | ENSMUSG( | ENSMUSG( | -0.2147 | 7.60E-05 | YES | NO | NA |
| Rab43 | ENSMUSG( | ENSMUSG( | 0.196124 | 0.089051 | NO | NO | NA |
| Txk | ENSMUSG( | ENSMUSG( | -0.56883 | 2.20E-06 | YES | NO | NA |
| Traf6 | ENSMUSG( | ENSMUSG( | -0.01064 | 0.925265 | NO | NO | NA |
| Smcr8 | ENSMUSG( | ENSMUSG( | -0.424 | 8.49E-07 | YES | NO | NA |
| Mctp2 | ENSMUSG( | ENSMUSG( | -0.4053 | 5.77E-07 | YES | NO | NA |
| Tsc2 | ENSMUSG( | ENSMUSG( | -0.221 | 0.00246 | YES | NO | NA |
| Ap1s2 | ENSMUSG( | ENSMUSG( | -0.55562 | 2.03E-09 | YES | NO | NA |
| Yipf3 | ENSMUSG( | ENSMUSG( | -0.0119 | 0.867509 | NO | NO | NA |
| Rragc | ENSMUSG( | ENSMUSG( | -0.08471 | 0.362789 | NO | NO | NA |
| Ubr3 | ENSMUSG( | ENSMUSG( | -0.10513 | 0.30402 | NO | NO | NA |
| Parg | ENSMUSG( | ENSMUSG( | -0.17215 | 0.002123 | YES | NO | NA |
| Tmem245 | ENSMUSG( | ENSMUSG( | -0.36544 | 7.28E-05 | YES | NO | NA |
| Nup160 | ENSMUSG( | ENSMUSG( | 0.116741 | 0.206676 | NO | NO | NA |

| | | | | | | | |
|---|---|---|---|---|---|---|---|
| Wdr81 | ENSMUSG( | ENSMUSG( | -0.43178 | 1.91E-06 | YES | NO | NA |
| Entpd4 | ENSMUSG( | ENSMUSG( | 0.745149 | 1.11E-16 | YES | NO | NA |
| Prkar2a | ENSMUSG( | ENSMUSG( | 0.084555 | 0.218663 | NO | NO | NA |
| Dcaf13 | ENSMUSG( | ENSMUSG( | 0.209948 | 9.46E-05 | YES | NO | NA |
| Polr1a | ENSMUSG( | ENSMUSG( | 0.239801 | 0.014188 | YES | NO | NA |
| Exoc4 | ENSMUSG( | ENSMUSG( | -0.13866 | 0.016758 | YES | NO | NA |
| Ttyh3 | ENSMUSG( | ENSMUSG( | -0.08695 | 0.516992 | NO | NO | NA |
| Mcm10 | ENSMUSG( | ENSMUSG( | 0.329996 | 0.000155 | YES | NO | NA |
| Icmt | ENSMUSG( | ENSMUSG( | -0.09175 | 0.239113 | NO | NO | NA |
| Ptpn18 | ENSMUSG( | ENSMUSG( | -0.17516 | 0.071457 | NO | NO | NA |
| Nfya | ENSMUSG( | ENSMUSG( | -0.15093 | 0.000121 | YES | NO | NA |
| Mms22l | ENSMUSG( | ENSMUSG( | 0.115856 | 0.097803 | NO | NO | NA |
| Rbmx | ENSMUSG( | ENSMUSG( | 0.006851 | 0.920499 | NO | NO | NA |
| Synj1 | ENSMUSG( | ENSMUSG( | -0.35421 | 2.85E-05 | YES | NO | NA |
| Zer1 | ENSMUSG( | ENSMUSG( | -0.3652 | 0.000298 | YES | NO | NA |
| Zzz3 | ENSMUSG( | ENSMUSG( | -0.02914 | 0.71207 | NO | NO | NA |
| Pag1 | ENSMUSG( | ENSMUSG( | 0.34557 | 0.003171 | YES | NO | NA |
| Srsf9 | ENSMUSG( | ENSMUSG( | 0.178797 | 0.009092 | YES | NO | NA |
| Ifit1bl1 | ENSMUSG( | ENSMUSG( | -1.20874 | 2.93E-12 | YES | YES | Downregulated |
| Abl1 | ENSMUSG( | ENSMUSG( | -0.12996 | 0.097673 | NO | NO | NA |
| Smpd4 | ENSMUSG( | ENSMUSG( | 0.411972 | 6.58E-11 | YES | NO | NA |
| Atp6v0e | ENSMUSG( | ENSMUSG( | -0.18003 | 0.015111 | YES | NO | NA |
| Rps6ka4 | ENSMUSG( | ENSMUSG( | -0.11317 | 0.006104 | YES | NO | NA |
| Slc35a3 | ENSMUSG( | ENSMUSG( | -0.04263 | 0.609081 | NO | NO | NA |
| Mfsd14b | ENSMUSG( | ENSMUSG( | -0.26879 | 0.000554 | YES | NO | NA |
| Chic2 | ENSMUSG( | ENSMUSG( | -0.27576 | 0.000193 | YES | NO | NA |
| Sat1 | ENSMUSG( | ENSMUSG( | -0.24094 | 0.032736 | YES | NO | NA |
| Tbc1d9b | ENSMUSG( | ENSMUSG( | -0.28929 | 0.000198 | YES | NO | NA |
| Ddx10 | ENSMUSG( | ENSMUSG( | 0.215558 | 0.064799 | NO | NO | NA |
| Hdac3 | ENSMUSG( | ENSMUSG( | 0.056306 | 0.288187 | NO | NO | NA |
| Prkcd | ENSMUSG( | ENSMUSG( | -0.15669 | 0.03381 | YES | NO | NA |
| Irf4 | ENSMUSG( | ENSMUSG( | 0.242977 | 0.222904 | NO | NO | NA |
| Umps | ENSMUSG( | ENSMUSG( | 0.401808 | 0.000361 | YES | NO | NA |
| Ubr1 | ENSMUSG( | ENSMUSG( | -0.20105 | 0.020889 | YES | NO | NA |
| Ube3c | ENSMUSG( | ENSMUSG( | -0.0063 | 0.942632 | NO | NO | NA |
| Tipin | ENSMUSG( | ENSMUSG( | 0.473833 | 4.37E-06 | YES | NO | NA |
| Gpat4 | ENSMUSG( | ENSMUSG( | 0.120754 | 0.023694 | YES | NO | NA |
| Ubxn7 | ENSMUSG( | ENSMUSG( | -0.25516 | 0.000173 | YES | NO | NA |
| Drosha | ENSMUSG( | ENSMUSG( | 0.036946 | 0.667611 | NO | NO | NA |
| Glrx | ENSMUSG( | ENSMUSG( | -0.08252 | 0.480845 | NO | NO | NA |
| Lsm12 | ENSMUSG( | ENSMUSG( | 0.061668 | 0.459146 | NO | NO | NA |
| Nup155 | ENSMUSG( | ENSMUSG( | 0.064233 | 0.452531 | NO | NO | NA |
| Ruvbl2 | ENSMUSG( | ENSMUSG( | 0.391476 | 1.77E-05 | YES | NO | NA |
| Stx16 | ENSMUSG( | ENSMUSG( | -0.52141 | 3.43E-11 | YES | NO | NA |
| Rom1 | ENSMUSG( | ENSMUSG( | -0.19308 | 0.014393 | YES | NO | NA |
| Chd9 | ENSMUSG( | ENSMUSG( | -0.30639 | 0.000894 | YES | NO | NA |
| Tubgcp6 | ENSMUSG( | ENSMUSG( | -0.13568 | 0.011574 | YES | NO | NA |

| | | | | | | | |
|---|---|---|---|---|---|---|---|
| Stub1 | ENSMUSG( | ENSMUSG( | 0.009685 | 0.912113 | NO | NO | NA |
| Lpin2 | ENSMUSG( | ENSMUSG( | -0.18034 | 0.025019 | YES | NO | NA |
| Rbbp5 | ENSMUSG( | ENSMUSG( | -0.1848 | 0.006403 | YES | NO | NA |
| Map3k3 | ENSMUSG( | ENSMUSG( | -0.22302 | 0.002161 | YES | NO | NA |
| 9930111J2 | ENSMUSG( | ENSMUSG( | -0.80907 | 3.56E-16 | YES | NO | NA |
| L3mbtl2 | ENSMUSG( | ENSMUSG( | 0.12981 | 0.037218 | YES | NO | NA |
| Smpdl3a | ENSMUSG( | ENSMUSG( | -0.77667 | 5.76E-09 | YES | NO | NA |
| Smad4 | ENSMUSG( | ENSMUSG( | -0.25571 | 0.001351 | YES | NO | NA |
| Ckap2 | ENSMUSG( | ENSMUSG( | 0.293979 | 0.00022 | YES | NO | NA |
| Gpd1l | ENSMUSG( | ENSMUSG( | -0.01992 | 0.83183 | NO | NO | NA |
| Herpud2 | ENSMUSG( | ENSMUSG( | -0.2729 | 0.001646 | YES | NO | NA |
| Slc43a3 | ENSMUSG( | ENSMUSG( | 0.41233 | 1.99E-05 | YES | NO | NA |
| Galnt7 | ENSMUSG( | ENSMUSG( | -0.00171 | 0.978105 | NO | NO | NA |
| Morf4l2 | ENSMUSG( | ENSMUSG( | 0.252536 | 0.00118 | YES | NO | NA |
| Ubald1 | ENSMUSG( | ENSMUSG( | 0.041556 | 0.740295 | NO | NO | NA |
| Ranbp10 | ENSMUSG( | ENSMUSG( | -0.26247 | 0.001146 | YES | NO | NA |
| Rlf | ENSMUSG( | ENSMUSG( | -0.44386 | 0.00012 | YES | NO | NA |
| Rbx1 | ENSMUSG( | ENSMUSG( | -0.05183 | 0.33087 | NO | NO | NA |
| Mtor | ENSMUSG( | ENSMUSG( | 0.009394 | 0.916888 | NO | NO | NA |
| D8Ertd738 | ENSMUSG( | ENSMUSG( | 0.07517 | 0.425784 | NO | NO | NA |
| Anapc4 | ENSMUSG( | ENSMUSG( | 0.118257 | 0.065359 | NO | NO | NA |
| Crbn | ENSMUSG( | ENSMUSG( | -0.18531 | 0.022173 | YES | NO | NA |
| BE692007 | ENSMUSG( | ENSMUSG( | -0.91652 | 3.52E-09 | YES | NO | NA |
| Mepce | ENSMUSG( | ENSMUSG( | -0.21282 | 0.002472 | YES | NO | NA |
| Cers5 | ENSMUSG( | ENSMUSG( | -0.0775 | 0.159995 | NO | NO | NA |
| Fam172a | ENSMUSG( | ENSMUSG( | 0.009893 | 0.927591 | NO | NO | NA |
| Mtch2 | ENSMUSG( | ENSMUSG( | 0.227846 | 0.003302 | YES | NO | NA |
| Foxo3 | ENSMUSG( | ENSMUSG( | -1.10302 | 9.31E-14 | YES | YES | Downregulated |
| Zcchc7 | ENSMUSG( | ENSMUSG( | -0.15394 | 0.143121 | NO | NO | NA |
| Otulin | ENSMUSG( | ENSMUSG( | 0.078015 | 0.265756 | NO | NO | NA |
| Clasrp | ENSMUSG( | ENSMUSG( | -0.03269 | 0.600449 | NO | NO | NA |
| Sdad1 | ENSMUSG( | ENSMUSG( | 0.432087 | 9.79E-05 | YES | NO | NA |
| Actr10 | ENSMUSG( | ENSMUSG( | 0.051019 | 0.504438 | NO | NO | NA |
| Eif2ak2 | ENSMUSG( | ENSMUSG( | -0.44318 | 3.80E-10 | YES | NO | NA |
| Ruvbl1 | ENSMUSG( | ENSMUSG( | 0.622277 | 1.92E-07 | YES | NO | NA |
| Ints11 | ENSMUSG( | ENSMUSG( | -0.06573 | 0.514897 | NO | NO | NA |
| ENSMUSG( | ENSMUSG( | ENSMUSG( | 0.334658 | 0.000438 | YES | NO | NA |
| H2-T10 | ENSMUSG( | ENSMUSG( | -0.58256 | 6.82E-12 | YES | NO | NA |
| Ehmt1 | ENSMUSG( | ENSMUSG( | -0.08517 | 0.178813 | NO | NO | NA |
| Slc36a1 | ENSMUSG( | ENSMUSG( | -0.69956 | 3.86E-11 | YES | NO | NA |
| Rab5a | ENSMUSG( | ENSMUSG( | -0.19353 | 0.078786 | NO | NO | NA |
| P2ry10 | ENSMUSG( | ENSMUSG( | -0.16837 | 0.262034 | NO | NO | NA |
| Dnttip2 | ENSMUSG( | ENSMUSG( | -0.02315 | 0.85779 | NO | NO | NA |
| Wdr90 | ENSMUSG( | ENSMUSG( | 0.19187 | 0.015367 | YES | NO | NA |
| Gns | ENSMUSG( | ENSMUSG( | -1.03888 | 1.26E-08 | YES | YES | Downregulated |
| Chd1l | ENSMUSG( | ENSMUSG( | 0.159642 | 0.063932 | NO | NO | NA |
| Rabl6 | ENSMUSG( | ENSMUSG( | 0.096235 | 0.027048 | YES | NO | NA |

| | | | | | | | |
|---|---|---|---|---|---|---|---|
| Chst2 | ENSMUSG( | ENSMUSG( | -0.44072 | 0.001217 | YES | NO | NA |
| Prcc | ENSMUSG( | ENSMUSG( | -0.08824 | 0.192105 | NO | NO | NA |
| Kpna1 | ENSMUSG( | ENSMUSG( | 0.007896 | 0.89804 | NO | NO | NA |
| Mecp2 | ENSMUSG( | ENSMUSG( | -0.1621 | 0.015566 | YES | NO | NA |
| Tnfsf10 | ENSMUSG( | ENSMUSG( | -0.37928 | 2.48E-05 | YES | NO | NA |
| Vapa | ENSMUSG( | ENSMUSG( | -0.05062 | 0.409568 | NO | NO | NA |
| Upf3a | ENSMUSG( | ENSMUSG( | 0.057064 | 0.664861 | NO | NO | NA |
| Gpx4 | ENSMUSG( | ENSMUSG( | 0.182317 | 0.001018 | YES | NO | NA |
| Mx1 | ENSMUSG( | ENSMUSG( | -0.8949 | 4.31E-14 | YES | NO | NA |
| Napa | ENSMUSG( | ENSMUSG( | -0.06116 | 0.421121 | NO | NO | NA |
| Mapk3 | ENSMUSG( | ENSMUSG( | -0.13786 | 0.085539 | NO | NO | NA |
| Cdc14a | ENSMUSG( | ENSMUSG( | -0.15192 | 0.08249 | NO | NO | NA |
| Sppl3 | ENSMUSG( | ENSMUSG( | -0.20393 | 0.00237 | YES | NO | NA |
| Gtf3a | ENSMUSG( | ENSMUSG( | 0.194148 | 0.003797 | YES | NO | NA |
| Dlgap4 | ENSMUSG( | ENSMUSG( | -0.58507 | 4.39E-15 | YES | NO | NA |
| Eif1ad | ENSMUSG( | ENSMUSG( | 0.114134 | 0.044426 | YES | NO | NA |
| Map2k1 | ENSMUSG( | ENSMUSG( | -0.26055 | 1.43E-07 | YES | NO | NA |
| Pkp4 | ENSMUSG( | ENSMUSG( | -0.01562 | 0.87041 | NO | NO | NA |
| Usp39 | ENSMUSG( | ENSMUSG( | 0.140755 | 0.002509 | YES | NO | NA |
| Pfkfb3 | ENSMUSG( | ENSMUSG( | 0.019485 | 0.87566 | NO | NO | NA |
| Ralgapa1 | ENSMUSG( | ENSMUSG( | -0.24096 | 0.001981 | YES | NO | NA |
| Sec22b | ENSMUSG( | ENSMUSG( | -0.04087 | 0.536617 | NO | NO | NA |
| Rab3gap1 | ENSMUSG( | ENSMUSG( | -0.07694 | 0.319978 | NO | NO | NA |
| Chaf1b | ENSMUSG( | ENSMUSG( | 0.230655 | 0.000765 | YES | NO | NA |
| Stx4a | ENSMUSG( | ENSMUSG( | -0.23461 | 0.001635 | YES | NO | NA |
| Kdm7a | ENSMUSG( | ENSMUSG( | -0.67991 | 1.01E-06 | YES | NO | NA |
| Rbbp8 | ENSMUSG( | ENSMUSG( | 0.067525 | 0.513919 | NO | NO | NA |
| Gbp2 | ENSMUSG( | ENSMUSG( | -0.46591 | 5.50E-05 | YES | NO | NA |
| Ube2r2 | ENSMUSG( | ENSMUSG( | -0.13657 | 0.026869 | YES | NO | NA |
| Tasor2 | ENSMUSG( | ENSMUSG( | -0.3247 | 0.000671 | YES | NO | NA |
| Pink1 | ENSMUSG( | ENSMUSG( | -1.32569 | 2.62E-20 | YES | YES | Downregulated |
| Capn7 | ENSMUSG( | ENSMUSG( | -0.16103 | 0.054157 | NO | NO | NA |
| 1810030O | ENSMUSG( | ENSMUSG( | 0.041709 | 0.648008 | NO | NO | NA |
| Atp2c1 | ENSMUSG( | ENSMUSG( | -0.1644 | 0.006301 | YES | NO | NA |
| Gnl1 | ENSMUSG( | ENSMUSG( | 0.144077 | 0.017231 | YES | NO | NA |
| Mkln1 | ENSMUSG( | ENSMUSG( | -0.20295 | 0.002142 | YES | NO | NA |
| Ciao2a | ENSMUSG( | ENSMUSG( | 0.195315 | 0.005528 | YES | NO | NA |
| Phf6 | ENSMUSG( | ENSMUSG( | -0.12036 | 0.18308 | NO | NO | NA |
| Cdc73 | ENSMUSG( | ENSMUSG( | -0.11994 | 0.146854 | NO | NO | NA |
| Nosip | ENSMUSG( | ENSMUSG( | 0.189951 | 0.005589 | YES | NO | NA |
| Sdhc | ENSMUSG( | ENSMUSG( | 0.209062 | 0.000966 | YES | NO | NA |
| Ctr9 | ENSMUSG( | ENSMUSG( | -0.22202 | 0.000107 | YES | NO | NA |
| Gstp3 | ENSMUSG( | ENSMUSG( | 0.772313 | 3.40E-05 | YES | NO | NA |
| Dgkh | ENSMUSG( | ENSMUSG( | -0.29051 | 0.000216 | YES | NO | NA |
| Map3k4 | ENSMUSG( | ENSMUSG( | -0.12035 | 0.104719 | NO | NO | NA |
| Arl6ip4 | ENSMUSG( | ENSMUSG( | 0.287373 | 0.005869 | YES | NO | NA |
| Slc30a9 | ENSMUSG( | ENSMUSG( | -0.00967 | 0.897577 | NO | NO | NA |

| | | | | | | | |
|---|---|---|---|---|---|---|---|
| Ppp6c | ENSMUSG( | ENSMUSG( | -0.13798 | 0.115469 | NO | NO | NA |
| Dcaf12 | ENSMUSG( | ENSMUSG( | -0.21519 | 3.55E-06 | YES | NO | NA |
| Igkc | ENSMUSG( | ENSMUSG( | -0.49456 | 0.471322 | NO | NO | NA |
| Mfng | ENSMUSG( | ENSMUSG( | -0.23974 | 1.02E-05 | YES | NO | NA |
| Zw10 | ENSMUSG( | ENSMUSG( | -0.12637 | 0.027552 | YES | NO | NA |
| Foxj3 | ENSMUSG( | ENSMUSG( | -0.15961 | 0.021776 | YES | NO | NA |
| Ctdp1 | ENSMUSG( | ENSMUSG( | 0.046709 | 0.557045 | NO | NO | NA |
| Cdk5rap3 | ENSMUSG( | ENSMUSG( | 0.068573 | 0.476831 | NO | NO | NA |
| Dcun1d5 | ENSMUSG( | ENSMUSG( | 0.056273 | 0.534005 | NO | NO | NA |
| Sgta | ENSMUSG( | ENSMUSG( | 0.134568 | 0.03606 | YES | NO | NA |
| Pdcd6 | ENSMUSG( | ENSMUSG( | 0.175995 | 0.037958 | YES | NO | NA |
| Trim12a | ENSMUSG( | ENSMUSG( | -0.58133 | 9.13E-11 | YES | NO | NA |
| Acap3 | ENSMUSG( | ENSMUSG( | -0.16348 | 2.88E-05 | YES | NO | NA |
| Fam193a | ENSMUSG( | ENSMUSG( | -0.3666 | 3.99E-05 | YES | NO | NA |
| Snx25 | ENSMUSG( | ENSMUSG( | 0.096353 | 0.254163 | NO | NO | NA |
| Cpox | ENSMUSG( | ENSMUSG( | 0.213184 | 0.029079 | YES | NO | NA |
| Asb6 | ENSMUSG( | ENSMUSG( | -0.16997 | 0.00498 | YES | NO | NA |
| Washc1 | ENSMUSG( | ENSMUSG( | 0.09741 | 0.339176 | NO | NO | NA |
| Exoc1 | ENSMUSG( | ENSMUSG( | 0.025964 | 0.80971 | NO | NO | NA |
| Erp29 | ENSMUSG( | ENSMUSG( | 0.17717 | 0.003629 | YES | NO | NA |
| Pigm | ENSMUSG( | ENSMUSG( | -0.21609 | 0.005591 | YES | NO | NA |
| Arhgef7 | ENSMUSG( | ENSMUSG( | -0.09171 | 0.138116 | NO | NO | NA |
| Rnf111 | ENSMUSG( | ENSMUSG( | -0.46975 | 3.52E-06 | YES | NO | NA |
| Mllt1 | ENSMUSG( | ENSMUSG( | -0.24456 | 0.000219 | YES | NO | NA |
| Setdb1 | ENSMUSG( | ENSMUSG( | 0.011112 | 0.855874 | NO | NO | NA |
| Ap1ar | ENSMUSG( | ENSMUSG( | 0.053938 | 0.559018 | NO | NO | NA |
| Adcy3 | ENSMUSG( | ENSMUSG( | -0.42123 | 3.16E-11 | YES | NO | NA |
| Exosc3 | ENSMUSG( | ENSMUSG( | 0.191035 | 0.147833 | NO | NO | NA |
| Tinf2 | ENSMUSG( | ENSMUSG( | 0.03572 | 0.600183 | NO | NO | NA |
| Prkx | ENSMUSG( | ENSMUSG( | -0.55535 | 2.69E-09 | YES | NO | NA |
| Ifi35 | ENSMUSG( | ENSMUSG( | -0.4169 | 4.89E-08 | YES | NO | NA |
| Pfdn6 | ENSMUSG( | ENSMUSG( | 0.241773 | 0.106744 | NO | NO | NA |
| Vps39 | ENSMUSG( | ENSMUSG( | -0.13303 | 0.015853 | YES | NO | NA |
| Llgl1 | ENSMUSG( | ENSMUSG( | -0.11475 | 0.066001 | NO | NO | NA |
| Fam98b | ENSMUSG( | ENSMUSG( | 0.14286 | 0.102306 | NO | NO | NA |
| Carhsp1 | ENSMUSG( | ENSMUSG( | -0.22132 | 0.006048 | YES | NO | NA |
| Rnpepl1 | ENSMUSG( | ENSMUSG( | -0.23401 | 0.001571 | YES | NO | NA |
| Rnasel | ENSMUSG( | ENSMUSG( | -0.33952 | 0.000758 | YES | NO | NA |
| Sp2 | ENSMUSG( | ENSMUSG( | -0.16706 | 0.021584 | YES | NO | NA |
| Stk40 | ENSMUSG( | ENSMUSG( | -0.5016 | 1.14E-06 | YES | NO | NA |
| Ufc1 | ENSMUSG( | ENSMUSG( | 0.19785 | 0.013546 | YES | NO | NA |
| Cdan1 | ENSMUSG( | ENSMUSG( | -0.00353 | 0.953027 | NO | NO | NA |
| Brox | ENSMUSG( | ENSMUSG( | -0.1283 | 0.319151 | NO | NO | NA |
| Atad2b | ENSMUSG( | ENSMUSG( | -0.29925 | 1.09E-05 | YES | NO | NA |
| Amd1 | ENSMUSG( | ENSMUSG( | 0.45434 | 1.36E-07 | YES | NO | NA |
| Ubqln4 | ENSMUSG( | ENSMUSG( | 0.018384 | 0.791732 | NO | NO | NA |
| Agpat1 | ENSMUSG( | ENSMUSG( | -0.41336 | 1.43E-13 | YES | NO | NA |

| | | | | | | | |
|---|---|---|---|---|---|---|---|
| Ascc3 | ENSMUSG( | ENSMUSG( | -0.28924 | 4.52E-05 | YES | NO | NA |
| Rwdd1 | ENSMUSG( | ENSMUSG( | 0.097249 | 0.313018 | NO | NO | NA |
| Ppp4r1 | ENSMUSG( | ENSMUSG( | -0.39158 | 1.51E-09 | YES | NO | NA |
| Nfkb2 | ENSMUSG( | ENSMUSG( | -0.08976 | 0.134603 | NO | NO | NA |
| Cdc45 | ENSMUSG( | ENSMUSG( | 0.181202 | 0.045279 | YES | NO | NA |
| Gnl3l | ENSMUSG( | ENSMUSG( | 0.088637 | 0.190242 | NO | NO | NA |
| Gna15 | ENSMUSG( | ENSMUSG( | -0.12828 | 0.280969 | NO | NO | NA |
| Vbp1 | ENSMUSG( | ENSMUSG( | 0.156149 | 0.070162 | NO | NO | NA |
| Abcb7 | ENSMUSG( | ENSMUSG( | -0.05065 | 0.524985 | NO | NO | NA |
| Ttc9c | ENSMUSG( | ENSMUSG( | -0.1869 | 0.025908 | YES | NO | NA |
| Arl8b | ENSMUSG( | ENSMUSG( | 0.0245 | 0.702923 | NO | NO | NA |
| Med16 | ENSMUSG( | ENSMUSG( | 0.345415 | 3.70E-05 | YES | NO | NA |
| Myo1c | ENSMUSG( | ENSMUSG( | 0.011206 | 0.909942 | NO | NO | NA |
| Grina | ENSMUSG( | ENSMUSG( | -0.02053 | 0.878742 | NO | NO | NA |
| Fam126a | ENSMUSG( | ENSMUSG( | -0.39944 | 4.67E-07 | YES | NO | NA |
| Pym1 | ENSMUSG( | ENSMUSG( | 0.287712 | 9.03E-07 | YES | NO | NA |
| Gps2 | ENSMUSG( | ENSMUSG( | -0.00652 | 0.926979 | NO | NO | NA |
| Rgs14 | ENSMUSG( | ENSMUSG( | -0.34465 | 2.75E-07 | YES | NO | NA |
| Gm19585 | ENSMUSG( | ENSMUSG( | 0.08389 | 0.46475 | NO | NO | NA |
| Ndufa8 | ENSMUSG( | ENSMUSG( | 0.120572 | 0.162572 | NO | NO | NA |
| Rnf181 | ENSMUSG( | ENSMUSG( | -0.18298 | 0.003929 | YES | NO | NA |
| Ufd1 | ENSMUSG( | ENSMUSG( | 0.042829 | 0.660946 | NO | NO | NA |
| Slc25a46 | ENSMUSG( | ENSMUSG( | -0.01884 | 0.848177 | NO | NO | NA |
| Acbd3 | ENSMUSG( | ENSMUSG( | -0.174 | 0.025324 | YES | NO | NA |
| Myc | ENSMUSG( | ENSMUSG( | 0.268493 | 0.012764 | YES | NO | NA |
| Ppp6r2 | ENSMUSG( | ENSMUSG( | -0.07745 | 0.479158 | NO | NO | NA |
| Kpna6 | ENSMUSG( | ENSMUSG( | -0.14223 | 0.001646 | YES | NO | NA |
| Zfp142 | ENSMUSG( | ENSMUSG( | -0.2771 | 9.73E-05 | YES | NO | NA |
| Tlk2 | ENSMUSG( | ENSMUSG( | -0.12744 | 0.217858 | NO | NO | NA |
| Zfp652 | ENSMUSG( | ENSMUSG( | -0.50299 | 4.14E-07 | YES | NO | NA |
| Bysl | ENSMUSG( | ENSMUSG( | 0.102664 | 0.314293 | NO | NO | NA |
| Fam20b | ENSMUSG( | ENSMUSG( | -0.14634 | 0.030763 | YES | NO | NA |
| Haus6 | ENSMUSG( | ENSMUSG( | 0.15861 | 0.002712 | YES | NO | NA |
| Rgs2 | ENSMUSG( | ENSMUSG( | 0.453621 | 0.254163 | NO | NO | NA |
| Mphosph9 | ENSMUSG( | ENSMUSG( | 0.040276 | 0.587391 | NO | NO | NA |
| Slc25a51 | ENSMUSG( | ENSMUSG( | -0.10283 | 0.086754 | NO | NO | NA |
| Avl9 | ENSMUSG( | ENSMUSG( | -0.28469 | 0.001196 | YES | NO | NA |
| Dcaf11 | ENSMUSG( | ENSMUSG( | -0.26465 | 0.019231 | YES | NO | NA |
| Rab5b | ENSMUSG( | ENSMUSG( | -0.46773 | 9.63E-07 | YES | NO | NA |
| Itga6 | ENSMUSG( | ENSMUSG( | -0.30573 | 0.000126 | YES | NO | NA |
| Ankle2 | ENSMUSG( | ENSMUSG( | -0.20119 | 0.00106 | YES | NO | NA |
| Prdx6 | ENSMUSG( | ENSMUSG( | 0.208003 | 0.015063 | YES | NO | NA |
| Pkn2 | ENSMUSG( | ENSMUSG( | -0.31757 | 1.37E-06 | YES | NO | NA |
| Fhip2a | ENSMUSG( | ENSMUSG( | -0.48026 | 3.55E-09 | YES | NO | NA |
| Usp10 | ENSMUSG( | ENSMUSG( | 0.115732 | 0.209768 | NO | NO | NA |
| Mid1ip1 | ENSMUSG( | ENSMUSG( | 0.118372 | 0.234446 | NO | NO | NA |
| Acaa1a | ENSMUSG( | ENSMUSG( | -0.10235 | 0.069145 | NO | NO | NA |

| | | | | | | | |
|---|---|---|---|---|---|---|---|
| Wdr36 | ENSMUSG( | ENSMUSG( | 0.435714 | 4.62E-06 | YES | NO | NA |
| Ppp1r2 | ENSMUSG( | ENSMUSG( | 0.122993 | 0.108783 | NO | NO | NA |
| Tmem258 | ENSMUSG( | ENSMUSG( | 0.0196 | 0.930577 | NO | NO | NA |
| Aldh3a2 | ENSMUSG( | ENSMUSG( | -0.36252 | 2.56E-06 | YES | NO | NA |
| Grsf1 | ENSMUSG( | ENSMUSG( | 0.184145 | 0.01203 | YES | NO | NA |
| Yeats4 | ENSMUSG( | ENSMUSG( | 0.125711 | 0.176859 | NO | NO | NA |
| G6pdx | ENSMUSG( | ENSMUSG( | 0.231081 | 3.87E-05 | YES | NO | NA |
| Rae1 | ENSMUSG( | ENSMUSG( | 0.162393 | 0.014715 | YES | NO | NA |
| Aldh16a1 | ENSMUSG( | ENSMUSG( | 0.342686 | 0.000255 | YES | NO | NA |
| Rheb | ENSMUSG( | ENSMUSG( | -0.1144 | 0.114589 | NO | NO | NA |
| Tmem106l | ENSMUSG( | ENSMUSG( | -0.24334 | 0.065015 | NO | NO | NA |
| Tle4 | ENSMUSG( | ENSMUSG( | -0.372 | 0.000186 | YES | NO | NA |
| Rp9 | ENSMUSG( | ENSMUSG( | 0.185908 | 0.025263 | YES | NO | NA |
| Srp14 | ENSMUSG( | ENSMUSG( | -0.17538 | 0.0012 | YES | NO | NA |
| Cipc | ENSMUSG( | ENSMUSG( | -1.06434 | 2.52E-11 | YES | YES | Downregulated |
| Slc39a6 | ENSMUSG( | ENSMUSG( | -0.29652 | 0.000982 | YES | NO | NA |
| Ercc3 | ENSMUSG( | ENSMUSG( | 0.059846 | 0.499325 | NO | NO | NA |
| Igbp1 | ENSMUSG( | ENSMUSG( | 0.016559 | 0.844822 | NO | NO | NA |
| Spsb3 | ENSMUSG( | ENSMUSG( | -0.01217 | 0.913346 | NO | NO | NA |
| Selenok | ENSMUSG( | ENSMUSG( | -0.16954 | 0.071616 | NO | NO | NA |
| Ddx19b | ENSMUSG( | ENSMUSG( | 0.289 | 0.004734 | YES | NO | NA |
| Uhrf1bp1l | ENSMUSG( | ENSMUSG( | 0.420519 | 2.83E-05 | YES | NO | NA |
| Ap3m1 | ENSMUSG( | ENSMUSG( | 0.087014 | 0.243018 | NO | NO | NA |
| Ino80c | ENSMUSG( | ENSMUSG( | 0.02163 | 0.836189 | NO | NO | NA |
| Ankrd16 | ENSMUSG( | ENSMUSG( | 0.156017 | 0.148937 | NO | NO | NA |
| Ppp1r7 | ENSMUSG( | ENSMUSG( | 0.275178 | 7.65E-05 | YES | NO | NA |
| Bccip | ENSMUSG( | ENSMUSG( | 0.250814 | 0.017856 | YES | NO | NA |
| Elovl1 | ENSMUSG( | ENSMUSG( | -0.06405 | 0.298001 | NO | NO | NA |
| Gm46620 | ENSMUSG( | ENSMUSG( | 1.475098 | 6.51E-30 | YES | YES | Upregulated |
| Adgrg5 | ENSMUSG( | ENSMUSG( | 0.773558 | 0.001364 | YES | NO | NA |
| Eif2b5 | ENSMUSG( | ENSMUSG( | 0.313394 | 2.36E-05 | YES | NO | NA |
| Cdc123 | ENSMUSG( | ENSMUSG( | 0.105274 | 0.294351 | NO | NO | NA |
| Zscan29 | ENSMUSG( | ENSMUSG( | -0.44858 | 1.21E-05 | YES | NO | NA |
| Usf1 | ENSMUSG( | ENSMUSG( | -0.15811 | 0.014477 | YES | NO | NA |
| Gusb | ENSMUSG( | ENSMUSG( | 0.359892 | 1.86E-06 | YES | NO | NA |
| Slc16a6 | ENSMUSG( | ENSMUSG( | -0.06566 | 0.276792 | NO | NO | NA |
| Morc2a | ENSMUSG( | ENSMUSG( | 0.152681 | 0.083438 | NO | NO | NA |
| Fbxw11 | ENSMUSG( | ENSMUSG( | -0.08051 | 0.278735 | NO | NO | NA |
| Gm49759 | ENSMUSG( | ENSMUSG( | 0.529446 | 0.000323 | YES | NO | NA |
| Als2cl | ENSMUSG( | ENSMUSG( | -0.37727 | 0.014942 | YES | NO | NA |
| Noct | ENSMUSG( | ENSMUSG( | -0.07991 | 0.341314 | NO | NO | NA |
| Pigq | ENSMUSG( | ENSMUSG( | 0.055587 | 0.48942 | NO | NO | NA |
| Bri3bp | ENSMUSG( | ENSMUSG( | 0.891807 | 4.92E-11 | YES | NO | NA |
| Cops4 | ENSMUSG( | ENSMUSG( | 0.213686 | 0.007407 | YES | NO | NA |
| Tmem134 | ENSMUSG( | ENSMUSG( | -0.09349 | 0.257835 | NO | NO | NA |
| Usp16 | ENSMUSG( | ENSMUSG( | -0.02203 | 0.853757 | NO | NO | NA |
| Fbxo42 | ENSMUSG( | ENSMUSG( | -0.20716 | 0.00075 | YES | NO | NA |

| | | | | | | | |
|---|---|---|---|---|---|---|---|
| Upf2 | ENSMUSG( | ENSMUSG( | -0.03675 | 0.763512 | NO | NO | NA |
| Gpr171 | ENSMUSG( | ENSMUSG( | 0.077415 | 0.580673 | NO | NO | NA |
| Slc37a1 | ENSMUSG( | ENSMUSG( | -0.52232 | 2.03E-08 | YES | NO | NA |
| Exoc3 | ENSMUSG( | ENSMUSG( | -0.1905 | 0.000283 | YES | NO | NA |
| Skp1 | ENSMUSG( | ENSMUSG( | 0.010367 | 0.906987 | NO | NO | NA |
| Ncdn | ENSMUSG( | ENSMUSG( | 0.078192 | 0.229109 | NO | NO | NA |
| Haus4 | ENSMUSG( | ENSMUSG( | 0.333302 | 0.000163 | YES | NO | NA |
| Lats2 | ENSMUSG( | ENSMUSG( | -0.86589 | 2.24E-10 | YES | NO | NA |
| Tubg1 | ENSMUSG( | ENSMUSG( | 0.275931 | 0.000218 | YES | NO | NA |
| Eloc | ENSMUSG( | ENSMUSG( | 0.115297 | 0.33212 | NO | NO | NA |
| Zwint | ENSMUSG( | ENSMUSG( | 0.004932 | 0.959488 | NO | NO | NA |
| Uba3 | ENSMUSG( | ENSMUSG( | -0.19732 | 0.003816 | YES | NO | NA |
| Fbrsl1 | ENSMUSG( | ENSMUSG( | 0.010292 | 0.930465 | NO | NO | NA |
| Spc24 | ENSMUSG( | ENSMUSG( | 0.306431 | 0.000373 | YES | NO | NA |
| Mns1 | ENSMUSG( | ENSMUSG( | 0.159367 | 0.350628 | NO | NO | NA |
| Fam91a1 | ENSMUSG( | ENSMUSG( | 0.054763 | 0.580769 | NO | NO | NA |
| Pgm1 | ENSMUSG( | ENSMUSG( | 0.458504 | 6.22E-06 | YES | NO | NA |
| Zkscan17 | ENSMUSG( | ENSMUSG( | 0.160473 | 0.086921 | NO | NO | NA |
| Pspc1 | ENSMUSG( | ENSMUSG( | 0.20697 | 0.023524 | YES | NO | NA |
| Cdc7 | ENSMUSG( | ENSMUSG( | 0.243987 | 0.002136 | YES | NO | NA |
| Arl14ep | ENSMUSG( | ENSMUSG( | 0.640788 | 8.39E-08 | YES | NO | NA |
| Cst3 | ENSMUSG( | ENSMUSG( | 0.081821 | 0.335952 | NO | NO | NA |
| Fanca | ENSMUSG( | ENSMUSG( | 0.206059 | 0.000932 | YES | NO | NA |
| Clasp2 | ENSMUSG( | ENSMUSG( | 0.037391 | 0.60683 | NO | NO | NA |
| Mrps25 | ENSMUSG( | ENSMUSG( | -0.16107 | 0.02042 | YES | NO | NA |
| Riok1 | ENSMUSG( | ENSMUSG( | 0.15493 | 0.072765 | NO | NO | NA |
| Paqr4 | ENSMUSG( | ENSMUSG( | 0.035358 | 0.729736 | NO | NO | NA |
| Flt3l | ENSMUSG( | ENSMUSG( | -0.39742 | 9.97E-06 | YES | NO | NA |
| Slf2 | ENSMUSG( | ENSMUSG( | -0.12975 | 0.004777 | YES | NO | NA |
| Nfic | ENSMUSG( | ENSMUSG( | -0.10156 | 0.157988 | NO | NO | NA |
| Cebpg | ENSMUSG( | ENSMUSG( | -0.31152 | 6.11E-06 | YES | NO | NA |
| Cks2 | ENSMUSG( | ENSMUSG( | 0.296479 | 1.45E-06 | YES | NO | NA |
| Emc3 | ENSMUSG( | ENSMUSG( | 0.017454 | 0.821131 | NO | NO | NA |
| Atg3 | ENSMUSG( | ENSMUSG( | -0.21703 | 0.045793 | YES | NO | NA |
| Sugp1 | ENSMUSG( | ENSMUSG( | -0.01674 | 0.826879 | NO | NO | NA |
| Gba2 | ENSMUSG( | ENSMUSG( | -0.28025 | 0.000373 | YES | NO | NA |
| Dtx3 | ENSMUSG( | ENSMUSG( | 0.095736 | 0.369772 | NO | NO | NA |
| Rab6a | ENSMUSG( | ENSMUSG( | -0.27583 | 9.36E-06 | YES | NO | NA |
| Vamp3 | ENSMUSG( | ENSMUSG( | -0.09847 | 0.216398 | NO | NO | NA |
| Eif2b2 | ENSMUSG( | ENSMUSG( | 0.366008 | 6.80E-06 | YES | NO | NA |
| Mapk9 | ENSMUSG( | ENSMUSG( | -0.10433 | 0.306811 | NO | NO | NA |
| Dyrk2 | ENSMUSG( | ENSMUSG( | -0.32092 | 0.001029 | YES | NO | NA |
| Slc16a1 | ENSMUSG( | ENSMUSG( | 0.73043 | 1.78E-07 | YES | NO | NA |
| Ncf4 | ENSMUSG( | ENSMUSG( | -0.02033 | 0.841607 | NO | NO | NA |
| Sema7a | ENSMUSG( | ENSMUSG( | 0.594621 | 1.69E-05 | YES | NO | NA |
| Ndufs8 | ENSMUSG( | ENSMUSG( | 0.244359 | 0.001961 | YES | NO | NA |
| Rpl7l1 | ENSMUSG( | ENSMUSG( | 0.085187 | 0.3488 | NO | NO | NA |

| | | | | | | | |
|---|---|---|---|---|---|---|---|
| Polr2e | ENSMUSG( | ENSMUSG( | 0.179069 | 0.000198 | YES | NO | NA |
| Phkg2 | ENSMUSG( | ENSMUSG( | -0.0059 | 0.934291 | NO | NO | NA |
| Zdhhc7 | ENSMUSG( | ENSMUSG( | -0.03567 | 0.645311 | NO | NO | NA |
| Eya3 | ENSMUSG( | ENSMUSG( | -0.22074 | 3.00E-09 | YES | NO | NA |
| Atf6b | ENSMUSG( | ENSMUSG( | -0.23257 | 3.16E-06 | YES | NO | NA |
| Scyl3 | ENSMUSG( | ENSMUSG( | -0.07393 | 0.50518 | NO | NO | NA |
| Tbrg4 | ENSMUSG( | ENSMUSG( | 0.405709 | 1.19E-09 | YES | NO | NA |
| Gpx1 | ENSMUSG( | ENSMUSG( | 0.436816 | 2.07E-14 | YES | NO | NA |
| Snx10 | ENSMUSG( | ENSMUSG( | -0.3128 | 1.00E-06 | YES | NO | NA |
| Tonsl | ENSMUSG( | ENSMUSG( | -0.02578 | 0.772679 | NO | NO | NA |
| Ergic2 | ENSMUSG( | ENSMUSG( | -0.18775 | 0.018515 | YES | NO | NA |
| Csnk1g1 | ENSMUSG( | ENSMUSG( | -0.19931 | 0.010887 | YES | NO | NA |
| 4930453N | ENSMUSG( | ENSMUSG( | -0.25611 | 0.00022 | YES | NO | NA |
| Pear1 | ENSMUSG( | ENSMUSG( | -0.62582 | 4.05E-07 | YES | NO | NA |
| Leprot | ENSMUSG( | ENSMUSG( | -0.01496 | 0.887324 | NO | NO | NA |
| Ccpg1 | ENSMUSG( | ENSMUSG( | -0.73344 | 8.79E-16 | YES | NO | NA |
| Cacul1 | ENSMUSG( | ENSMUSG( | -0.07483 | 0.309928 | NO | NO | NA |
| Trmt2a | ENSMUSG( | ENSMUSG( | 0.10713 | 0.121286 | NO | NO | NA |
| Ttpal | ENSMUSG( | ENSMUSG( | -0.11341 | 0.048698 | YES | NO | NA |
| Cish | ENSMUSG( | ENSMUSG( | -0.77394 | 5.51E-05 | YES | NO | NA |
| Snf8 | ENSMUSG( | ENSMUSG( | 0.084621 | 0.492715 | NO | NO | NA |
| Vps13c | ENSMUSG( | ENSMUSG( | -0.65054 | 4.68E-11 | YES | NO | NA |
| Rlim | ENSMUSG( | ENSMUSG( | -0.11306 | 0.106118 | NO | NO | NA |
| Lemd2 | ENSMUSG( | ENSMUSG( | -0.22482 | 0.000184 | YES | NO | NA |
| Birc2 | ENSMUSG( | ENSMUSG( | -0.25141 | 0.003025 | YES | NO | NA |
| Rrs1 | ENSMUSG( | ENSMUSG( | 0.459088 | 1.21E-05 | YES | NO | NA |
| Ndufb9 | ENSMUSG( | ENSMUSG( | 0.305055 | 0.000116 | YES | NO | NA |
| Cyp51 | ENSMUSG( | ENSMUSG( | 0.774012 | 6.14E-06 | YES | NO | NA |
| Tedc1 | ENSMUSG( | ENSMUSG( | 0.169998 | 0.002713 | YES | NO | NA |
| Rexo4 | ENSMUSG( | ENSMUSG( | -0.06129 | 0.530804 | NO | NO | NA |
| Mnt | ENSMUSG( | ENSMUSG( | -0.15342 | 0.032736 | YES | NO | NA |
| Filip1 | ENSMUSG( | ENSMUSG( | 0.937627 | 1.49E-05 | YES | NO | NA |
| H1f0 | ENSMUSG( | ENSMUSG( | 1.134705 | 1.63E-06 | YES | YES | Upregulated |
| Dnajc11 | ENSMUSG( | ENSMUSG( | 0.281949 | 0.000159 | YES | NO | NA |
| Micall1 | ENSMUSG( | ENSMUSG( | -0.2688 | 2.39E-05 | YES | NO | NA |
| Taf2 | ENSMUSG( | ENSMUSG( | 0.005227 | 0.958652 | NO | NO | NA |
| Gnl3 | ENSMUSG( | ENSMUSG( | 0.525971 | 4.43E-06 | YES | NO | NA |
| Sf3b6 | ENSMUSG( | ENSMUSG( | 0.104361 | 0.290434 | NO | NO | NA |
| Abcb1a | ENSMUSG( | ENSMUSG( | -0.64173 | 2.10E-09 | YES | NO | NA |
| Heatr5b | ENSMUSG( | ENSMUSG( | -0.29439 | 1.50E-05 | YES | NO | NA |
| Rnf125 | ENSMUSG( | ENSMUSG( | -0.21244 | 0.067367 | NO | NO | NA |
| Gpcpd1 | ENSMUSG( | ENSMUSG( | -0.37218 | 0.003832 | YES | NO | NA |
| Nr4a1 | ENSMUSG( | ENSMUSG( | 0.303541 | 0.435507 | NO | NO | NA |
| Phf23 | ENSMUSG( | ENSMUSG( | -0.18903 | 0.063705 | NO | NO | NA |
| Ahdc1 | ENSMUSG( | ENSMUSG( | -0.45202 | 6.88E-06 | YES | NO | NA |
| Phf1 | ENSMUSG( | ENSMUSG( | -0.64255 | 6.70E-09 | YES | NO | NA |
| 2010016I1 | ENSMUSG( | ENSMUSG( | -0.78033 | 7.13E-10 | YES | NO | NA |

| | | | | | | | |
|---|---|---|---|---|---|---|---|
| Klhdc4 | ENSMUSG( | ENSMUSG( | 0.045673 | 0.663297 | NO | NO | NA |
| Kif3b | ENSMUSG( | ENSMUSG( | -0.22838 | 0.042873 | YES | NO | NA |
| Uck1 | ENSMUSG( | ENSMUSG( | 0.032176 | 0.721185 | NO | NO | NA |
| Ssu72 | ENSMUSG( | ENSMUSG( | 0.14969 | 0.006032 | YES | NO | NA |
| Snx27 | ENSMUSG( | ENSMUSG( | -0.2903 | 2.03E-09 | YES | NO | NA |
| Atg2b | ENSMUSG( | ENSMUSG( | -0.09859 | 0.097482 | NO | NO | NA |
| Il10rb | ENSMUSG( | ENSMUSG( | -0.3246 | 1.09E-05 | YES | NO | NA |
| Ei24 | ENSMUSG( | ENSMUSG( | -0.10843 | 0.181363 | NO | NO | NA |
| Prkaca | ENSMUSG( | ENSMUSG( | -0.02321 | 0.75279 | NO | NO | NA |
| Rrp1b | ENSMUSG( | ENSMUSG( | 0.204947 | 0.004926 | YES | NO | NA |
| Gar1 | ENSMUSG( | ENSMUSG( | 0.628888 | 0.002488 | YES | NO | NA |
| Anapc7 | ENSMUSG( | ENSMUSG( | 0.233807 | 0.002856 | YES | NO | NA |
| Ccz1 | ENSMUSG( | ENSMUSG( | 0.236793 | 0.003664 | YES | NO | NA |
| Med10 | ENSMUSG( | ENSMUSG( | 0.011946 | 0.881144 | NO | NO | NA |
| Tent5c | ENSMUSG( | ENSMUSG( | -0.13308 | 0.456867 | NO | NO | NA |
| Sinhcaf | ENSMUSG( | ENSMUSG( | 0.064734 | 0.47288 | NO | NO | NA |
| Bmp2k | ENSMUSG( | ENSMUSG( | 0.278083 | 0.001706 | YES | NO | NA |
| Ap3s1 | ENSMUSG( | ENSMUSG( | 0.070135 | 0.442109 | NO | NO | NA |
| Ctss | ENSMUSG( | ENSMUSG( | -0.27424 | 0.010119 | YES | NO | NA |
| Ngdn | ENSMUSG( | ENSMUSG( | -0.00061 | 0.996207 | NO | NO | NA |
| Rad54l | ENSMUSG( | ENSMUSG( | 0.218276 | 0.007735 | YES | NO | NA |
| Drg1 | ENSMUSG( | ENSMUSG( | 0.074386 | 0.270106 | NO | NO | NA |
| Xpo4 | ENSMUSG( | ENSMUSG( | 0.057067 | 0.408636 | NO | NO | NA |
| Ext2 | ENSMUSG( | ENSMUSG( | 0.158757 | 0.126166 | NO | NO | NA |
| Secisbp2 | ENSMUSG( | ENSMUSG( | -0.0525 | 0.295595 | NO | NO | NA |
| Desi1 | ENSMUSG( | ENSMUSG( | 0.155562 | 0.022792 | YES | NO | NA |
| Kansl2 | ENSMUSG( | ENSMUSG( | 0.221403 | 0.001626 | YES | NO | NA |
| Stk25 | ENSMUSG( | ENSMUSG( | -0.14683 | 0.062463 | NO | NO | NA |
| Plekhj1 | ENSMUSG( | ENSMUSG( | -0.24071 | 0.000919 | YES | NO | NA |
| Ndufc2 | ENSMUSG( | ENSMUSG( | 0.358707 | 0.002243 | YES | NO | NA |
| Stim2 | ENSMUSG( | ENSMUSG( | -0.41305 | 2.81E-05 | YES | NO | NA |
| Yars | ENSMUSG( | ENSMUSG( | 0.373602 | 5.70E-06 | YES | NO | NA |
| Nubp2 | ENSMUSG( | ENSMUSG( | 0.105753 | 0.089051 | NO | NO | NA |
| Fasl | ENSMUSG( | ENSMUSG( | -0.73757 | 1.67E-08 | YES | NO | NA |
| Rexo2 | ENSMUSG( | ENSMUSG( | 0.367237 | 0.000595 | YES | NO | NA |
| Zfp654 | ENSMUSG( | ENSMUSG( | -0.40048 | 1.60E-05 | YES | NO | NA |
| Sft2d1 | ENSMUSG( | ENSMUSG( | -0.22959 | 9.78E-06 | YES | NO | NA |
| Cep97 | ENSMUSG( | ENSMUSG( | -0.10517 | 0.361085 | NO | NO | NA |
| Atp5g1 | ENSMUSG( | ENSMUSG( | 0.473341 | 1.63E-08 | YES | NO | NA |
| Trim37 | ENSMUSG( | ENSMUSG( | -0.07321 | 0.379744 | NO | NO | NA |
| Hook3 | ENSMUSG( | ENSMUSG( | -0.34993 | 3.35E-06 | YES | NO | NA |
| Wipf2 | ENSMUSG( | ENSMUSG( | -0.36389 | 2.33E-07 | YES | NO | NA |
| Cnot7 | ENSMUSG( | ENSMUSG( | 0.02139 | 0.7985 | NO | NO | NA |
| Cwc22 | ENSMUSG( | ENSMUSG( | 0.214967 | 0.026496 | YES | NO | NA |
| Epb41l2 | ENSMUSG( | ENSMUSG( | -0.37975 | 0.000511 | YES | NO | NA |
| Txndc9 | ENSMUSG( | ENSMUSG( | -0.11846 | 0.031974 | YES | NO | NA |
| Zfp367 | ENSMUSG( | ENSMUSG( | 0.092812 | 0.330426 | NO | NO | NA |

| | | | | | | | |
|---|---|---|---|---|---|---|---|
| Gpalpp1 | ENSMUSG( | ENSMUSG( | 0.072802 | 0.548121 | NO | NO | NA |
| Atg4d | ENSMUSG( | ENSMUSG( | -0.28697 | 4.97E-07 | YES | NO | NA |
| Ccp110 | ENSMUSG( | ENSMUSG( | 0.098308 | 0.313789 | NO | NO | NA |
| Zfand3 | ENSMUSG( | ENSMUSG( | -0.37264 | 6.30E-07 | YES | NO | NA |
| Clns1a | ENSMUSG( | ENSMUSG( | 0.292281 | 4.10E-08 | YES | NO | NA |
| Ptdss1 | ENSMUSG( | ENSMUSG( | -0.16187 | 0.001683 | YES | NO | NA |
| Tpm1 | ENSMUSG( | ENSMUSG( | -0.08608 | 0.213378 | NO | NO | NA |
| Rab11fip1 | ENSMUSG( | ENSMUSG( | -0.38856 | 6.22E-06 | YES | NO | NA |
| Lamp2 | ENSMUSG( | ENSMUSG( | -0.15301 | 0.141594 | NO | NO | NA |
| Smarcd1 | ENSMUSG( | ENSMUSG( | 0.127999 | 0.185287 | NO | NO | NA |
| Vps25 | ENSMUSG( | ENSMUSG( | 0.087327 | 0.361047 | NO | NO | NA |
| Tmem165 | ENSMUSG( | ENSMUSG( | 0.000519 | 0.99494 | NO | NO | NA |
| Cog1 | ENSMUSG( | ENSMUSG( | -0.11151 | 0.021179 | YES | NO | NA |
| Polh | ENSMUSG( | ENSMUSG( | 0.030412 | 0.638705 | NO | NO | NA |
| Ifit2 | ENSMUSG( | ENSMUSG( | -0.44692 | 2.54E-07 | YES | NO | NA |
| Ppa1 | ENSMUSG( | ENSMUSG( | 0.480714 | 7.76E-05 | YES | NO | NA |
| Tspan14 | ENSMUSG( | ENSMUSG( | -0.35241 | 1.21E-06 | YES | NO | NA |
| Pak1ip1 | ENSMUSG( | ENSMUSG( | 0.157484 | 0.118283 | NO | NO | NA |
| Slc39a9 | ENSMUSG( | ENSMUSG( | -0.01398 | 0.833106 | NO | NO | NA |
| E130309D | ENSMUSG( | ENSMUSG( | 0.073886 | 0.133102 | NO | NO | NA |
| Rc3h2 | ENSMUSG( | ENSMUSG( | -0.43068 | 2.87E-05 | YES | NO | NA |
| Cnbd2 | ENSMUSG( | ENSMUSG( | 2.214164 | 6.41E-50 | YES | YES | Upregulated |
| Tfam | ENSMUSG( | ENSMUSG( | 0.072058 | 0.117453 | NO | NO | NA |
| Cd244a | ENSMUSG( | ENSMUSG( | -0.82189 | 8.87E-06 | YES | NO | NA |
| Emc7 | ENSMUSG( | ENSMUSG( | 0.133421 | 0.100716 | NO | NO | NA |
| Bud31 | ENSMUSG( | ENSMUSG( | -0.00464 | 0.96675 | NO | NO | NA |
| Got1 | ENSMUSG( | ENSMUSG( | 0.171532 | 0.036696 | YES | NO | NA |
| Anapc16 | ENSMUSG( | ENSMUSG( | 0.310728 | 4.66E-05 | YES | NO | NA |
| Bmi1 | ENSMUSG( | ENSMUSG( | 0.017947 | 0.85784 | NO | NO | NA |
| Ost4 | ENSMUSG( | ENSMUSG( | -0.20511 | 0.04128 | YES | NO | NA |
| Srp54c | ENSMUSG( | ENSMUSG( | -0.2042 | 0.043222 | YES | NO | NA |
| Flot1 | ENSMUSG( | ENSMUSG( | -0.11824 | 0.359777 | NO | NO | NA |
| Ddrgk1 | ENSMUSG( | ENSMUSG( | 0.070356 | 0.553738 | NO | NO | NA |
| Mrpl24 | ENSMUSG( | ENSMUSG( | -0.04438 | 0.586729 | NO | NO | NA |
| Fam53c | ENSMUSG( | ENSMUSG( | -0.26041 | 1.64E-05 | YES | NO | NA |
| Pura | ENSMUSG( | ENSMUSG( | -0.13225 | 0.048008 | YES | NO | NA |
| Tmem9b | ENSMUSG( | ENSMUSG( | -0.40916 | 4.49E-08 | YES | NO | NA |
| Zfx | ENSMUSG( | ENSMUSG( | -0.26042 | 3.28E-05 | YES | NO | NA |
| Mpdu1 | ENSMUSG( | ENSMUSG( | 0.155729 | 0.076122 | NO | NO | NA |
| Atxn7l1 | ENSMUSG( | ENSMUSG( | -0.40491 | 2.94E-05 | YES | NO | NA |
| Inf2 | ENSMUSG( | ENSMUSG( | -0.01235 | 0.923891 | NO | NO | NA |
| Dnajc10 | ENSMUSG( | ENSMUSG( | 0.184367 | 0.097109 | NO | NO | NA |
| Ccdc9 | ENSMUSG( | ENSMUSG( | 0.069408 | 0.626786 | NO | NO | NA |
| Tsix | ENSMUSG( | ENSMUSG( | -0.45101 | 0.000113 | YES | NO | NA |
| Top3b | ENSMUSG( | ENSMUSG( | -0.19272 | 0.003122 | YES | NO | NA |
| Gng10 | ENSMUSG( | ENSMUSG( | 0.11587 | 0.125831 | NO | NO | NA |
| Nutf2 | ENSMUSG( | ENSMUSG( | 0.309026 | 0.004375 | YES | NO | NA |

| | | | | | | | |
|---|---|---|---|---|---|---|---|
| Nfyc | ENSMUSG( | ENSMUSG( | -0.01326 | 0.888811 | NO | NO | NA |
| Brca2 | ENSMUSG( | ENSMUSG( | 0.34258 | 0.000502 | YES | NO | NA |
| Lztr1 | ENSMUSG( | ENSMUSG( | -0.03119 | 0.678821 | NO | NO | NA |
| Gnpda1 | ENSMUSG( | ENSMUSG( | -0.31582 | 1.72E-05 | YES | NO | NA |
| Carm1 | ENSMUSG( | ENSMUSG( | 0.479124 | 1.15E-07 | YES | NO | NA |
| Aaas | ENSMUSG( | ENSMUSG( | 0.130626 | 0.049202 | YES | NO | NA |
| Mapkap1 | ENSMUSG( | ENSMUSG( | 0.081156 | 0.379287 | NO | NO | NA |
| Ifih1 | ENSMUSG( | ENSMUSG( | -0.45579 | 2.66E-05 | YES | NO | NA |
| Lmnb2 | ENSMUSG( | ENSMUSG( | 0.442977 | 2.81E-06 | YES | NO | NA |
| Wdr5 | ENSMUSG( | ENSMUSG( | 0.15067 | 0.002562 | YES | NO | NA |
| Scyl1 | ENSMUSG( | ENSMUSG( | -0.10287 | 0.086825 | NO | NO | NA |
| Ilkap | ENSMUSG( | ENSMUSG( | -0.03367 | 0.61985 | NO | NO | NA |
| Ecd | ENSMUSG( | ENSMUSG( | 0.239003 | 1.43E-05 | YES | NO | NA |
| Trps1 | ENSMUSG( | ENSMUSG( | -0.2861 | 0.004062 | YES | NO | NA |
| Crk | ENSMUSG( | ENSMUSG( | -0.14726 | 0.001157 | YES | NO | NA |
| Tent2 | ENSMUSG( | ENSMUSG( | 0.122398 | 0.236987 | NO | NO | NA |
| Atp6v1d | ENSMUSG( | ENSMUSG( | -0.21111 | 0.012212 | YES | NO | NA |
| Chchd3 | ENSMUSG( | ENSMUSG( | 0.123129 | 0.098291 | NO | NO | NA |
| Srp54b | ENSMUSG( | ENSMUSG( | 0.294468 | 0.041848 | YES | NO | NA |
| 4833439L1 | ENSMUSG( | ENSMUSG( | -0.19242 | 0.001695 | YES | NO | NA |
| Dmxl1 | ENSMUSG( | ENSMUSG( | -0.28298 | 0.008791 | YES | NO | NA |
| Scyl2 | ENSMUSG( | ENSMUSG( | -0.27182 | 5.94E-05 | YES | NO | NA |
| Naa35 | ENSMUSG( | ENSMUSG( | -0.05183 | 0.424857 | NO | NO | NA |
| Seh1l | ENSMUSG( | ENSMUSG( | 0.242484 | 0.0036 | YES | NO | NA |
| Xaf1 | ENSMUSG( | ENSMUSG( | -0.79317 | 8.40E-16 | YES | NO | NA |
| Arnt | ENSMUSG( | ENSMUSG( | -0.27022 | 0.000188 | YES | NO | NA |
| Lrpprc | ENSMUSG( | ENSMUSG( | 0.299559 | 5.83E-05 | YES | NO | NA |
| Zfp746 | ENSMUSG( | ENSMUSG( | -0.12459 | 0.085087 | NO | NO | NA |
| Rnf14 | ENSMUSG( | ENSMUSG( | -0.18191 | 0.066828 | NO | NO | NA |
| Spg11 | ENSMUSG( | ENSMUSG( | -0.27456 | 2.88E-05 | YES | NO | NA |
| Hacd3 | ENSMUSG( | ENSMUSG( | -0.06667 | 0.467256 | NO | NO | NA |
| Cmklr1 | ENSMUSG( | ENSMUSG( | -1.00537 | 8.46E-13 | YES | YES | Downregulated |
| Arhgap31 | ENSMUSG( | ENSMUSG( | -0.28407 | 0.006815 | YES | NO | NA |
| Mov10 | ENSMUSG( | ENSMUSG( | -0.72725 | 2.03E-12 | YES | NO | NA |
| Derl2 | ENSMUSG( | ENSMUSG( | -0.1949 | 0.000284 | YES | NO | NA |
| Gabarapl2 | ENSMUSG( | ENSMUSG( | -0.43247 | 6.61E-08 | YES | NO | NA |
| Fbxo11 | ENSMUSG( | ENSMUSG( | -0.3709 | 0.000368 | YES | NO | NA |
| Bard1 | ENSMUSG( | ENSMUSG( | 0.002458 | 0.977763 | NO | NO | NA |
| Cox7a2 | ENSMUSG( | ENSMUSG( | 0.165983 | 0.018626 | YES | NO | NA |
| Ugdh | ENSMUSG( | ENSMUSG( | 0.080618 | 0.387568 | NO | NO | NA |
| Nol8 | ENSMUSG( | ENSMUSG( | -0.22795 | 0.005044 | YES | NO | NA |
| Dcp1a | ENSMUSG( | ENSMUSG( | -0.0606 | 0.397319 | NO | NO | NA |
| Abcg1 | ENSMUSG( | ENSMUSG( | -0.46361 | 0.003932 | YES | NO | NA |
| Epc2 | ENSMUSG( | ENSMUSG( | -0.28067 | 0.012568 | YES | NO | NA |
| Klhl20 | ENSMUSG( | ENSMUSG( | -0.23366 | 1.17E-05 | YES | NO | NA |
| Cdk5rap2 | ENSMUSG( | ENSMUSG( | -0.18306 | 0.015397 | YES | NO | NA |
| Krr1 | ENSMUSG( | ENSMUSG( | 0.058196 | 0.3473 | NO | NO | NA |

| | | | | | | | |
|---|---|---|---|---|---|---|---|
| Fam133b | ENSMUSG( | ENSMUSG( | 0.046505 | 0.712298 | NO | NO | NA |
| Mon1b | ENSMUSG( | ENSMUSG( | -0.41077 | 7.26E-13 | YES | NO | NA |
| Csrnp1 | ENSMUSG( | ENSMUSG( | 0.109741 | 0.603838 | NO | NO | NA |
| Scamp2 | ENSMUSG( | ENSMUSG( | -0.2409 | 0.00011 | YES | NO | NA |
| Zfp990 | ENSMUSG( | ENSMUSG( | 9.967392 | 4.97E-119 | YES | YES | Upregulated |
| Atp5k | ENSMUSG( | ENSMUSG( | 0.152007 | 0.083343 | NO | NO | NA |
| Atf2 | ENSMUSG( | ENSMUSG( | -0.24494 | 0.001342 | YES | NO | NA |
| Ints6 | ENSMUSG( | ENSMUSG( | -0.27747 | 0.000913 | YES | NO | NA |
| Edem3 | ENSMUSG( | ENSMUSG( | -0.24981 | 0.00034 | YES | NO | NA |
| Ero1a | ENSMUSG( | ENSMUSG( | 0.211563 | 0.025835 | YES | NO | NA |
| Mrpl58 | ENSMUSG( | ENSMUSG( | 0.096997 | 0.350951 | NO | NO | NA |
| Appbp2 | ENSMUSG( | ENSMUSG( | -0.04607 | 0.496053 | NO | NO | NA |
| Polq | ENSMUSG( | ENSMUSG( | 0.108506 | 0.308887 | NO | NO | NA |
| Cdc40 | ENSMUSG( | ENSMUSG( | 0.03384 | 0.646606 | NO | NO | NA |
| Brwd1 | ENSMUSG( | ENSMUSG( | -0.38899 | 1.69E-11 | YES | NO | NA |
| Bex3 | ENSMUSG( | ENSMUSG( | 0.041519 | 0.634637 | NO | NO | NA |
| Rabac1 | ENSMUSG( | ENSMUSG( | -0.01475 | 0.895322 | NO | NO | NA |
| Armc10 | ENSMUSG( | ENSMUSG( | 0.047483 | 0.597006 | NO | NO | NA |
| Rgs19 | ENSMUSG( | ENSMUSG( | -0.14227 | 0.006961 | YES | NO | NA |
| Casp1 | ENSMUSG( | ENSMUSG( | 0.062104 | 0.606989 | NO | NO | NA |
| Arid5b | ENSMUSG( | ENSMUSG( | -0.43608 | 0.004187 | YES | NO | NA |
| Dusp4 | ENSMUSG( | ENSMUSG( | 0.949382 | 1.25E-06 | YES | NO | NA |
| Aspscr1 | ENSMUSG( | ENSMUSG( | -0.08234 | 0.075938 | NO | NO | NA |
| Lsg1 | ENSMUSG( | ENSMUSG( | 0.223023 | 0.003238 | YES | NO | NA |
| Mfsd10 | ENSMUSG( | ENSMUSG( | 0.135932 | 0.084644 | NO | NO | NA |
| Cnot11 | ENSMUSG( | ENSMUSG( | -0.06684 | 0.232205 | NO | NO | NA |
| Acp1 | ENSMUSG( | ENSMUSG( | 0.379944 | 1.08E-06 | YES | NO | NA |
| Gsdmd | ENSMUSG( | ENSMUSG( | -0.39433 | 1.07E-12 | YES | NO | NA |
| Pank3 | ENSMUSG( | ENSMUSG( | -0.27432 | 0.000167 | YES | NO | NA |
| Lrrc8a | ENSMUSG( | ENSMUSG( | -0.60038 | 3.04E-08 | YES | NO | NA |
| Cpsf4 | ENSMUSG( | ENSMUSG( | 0.101431 | 0.167728 | NO | NO | NA |
| Rgp1 | ENSMUSG( | ENSMUSG( | -0.16234 | 0.085873 | NO | NO | NA |
| Kctd9 | ENSMUSG( | ENSMUSG( | -0.03787 | 0.723945 | NO | NO | NA |
| Golga5 | ENSMUSG( | ENSMUSG( | 0.005204 | 0.965501 | NO | NO | NA |
| Cdc23 | ENSMUSG( | ENSMUSG( | 0.088301 | 0.228525 | NO | NO | NA |
| Map1s | ENSMUSG( | ENSMUSG( | 0.002475 | 0.976793 | NO | NO | NA |
| Camk1d | ENSMUSG( | ENSMUSG( | -0.63814 | 5.38E-08 | YES | NO | NA |
| Gm42742 | ENSMUSG( | ENSMUSG( | -0.28725 | 0.002554 | YES | NO | NA |
| Pptc7 | ENSMUSG( | ENSMUSG( | -0.30387 | 7.73E-07 | YES | NO | NA |
| Heg1 | ENSMUSG( | ENSMUSG( | -0.08554 | 0.317944 | NO | NO | NA |
| Cdc37l1 | ENSMUSG( | ENSMUSG( | -0.388 | 2.68E-05 | YES | NO | NA |
| Trim21 | ENSMUSG( | ENSMUSG( | -0.54499 | 1.72E-11 | YES | NO | NA |
| Antkmt | ENSMUSG( | ENSMUSG( | -0.05258 | 0.512821 | NO | NO | NA |
| Ncoa5 | ENSMUSG( | ENSMUSG( | 0.057834 | 0.399347 | NO | NO | NA |
| Bcl2a1d | ENSMUSG( | ENSMUSG( | 0.247211 | 0.039753 | YES | NO | NA |
| Stau1 | ENSMUSG( | ENSMUSG( | -0.21492 | 0.000439 | YES | NO | NA |
| Inpp4a | ENSMUSG( | ENSMUSG( | -0.69716 | 1.18E-12 | YES | NO | NA |

| | | | | | | | |
|---|---|---|---|---|---|---|---|
| Aurka | ENSMUSG( | ENSMUSG( | 0.218756 | 0.000171 | YES | NO | NA |
| Phf5a | ENSMUSG( | ENSMUSG( | 0.280121 | 1.83E-05 | YES | NO | NA |
| Hk2 | ENSMUSG( | ENSMUSG( | 1.047582 | 2.53E-10 | YES | YES | Upregulated |
| Magoh | ENSMUSG( | ENSMUSG( | 0.112183 | 0.379341 | NO | NO | NA |
| Cpeb2 | ENSMUSG( | ENSMUSG( | -1.29078 | 1.34E-13 | YES | YES | Downregulated |
| Ercc6l2 | ENSMUSG( | ENSMUSG( | 0.059919 | 0.475214 | NO | NO | NA |
| Epas1 | ENSMUSG( | ENSMUSG( | 0.778297 | 8.99E-05 | YES | NO | NA |
| Slc9a8 | ENSMUSG( | ENSMUSG( | 0.014612 | 0.851755 | NO | NO | NA |
| Mre11a | ENSMUSG( | ENSMUSG( | 0.071155 | 0.332987 | NO | NO | NA |
| Ezh1 | ENSMUSG( | ENSMUSG( | -0.41011 | 1.38E-07 | YES | NO | NA |
| Utp20 | ENSMUSG( | ENSMUSG( | 0.513143 | 3.67E-07 | YES | NO | NA |
| Cdc25a | ENSMUSG( | ENSMUSG( | 0.223024 | 0.000353 | YES | NO | NA |
| ENSMUSG( | ENSMUSG( | ENSMUSG( | -0.97147 | 1.01E-10 | YES | NO | NA |
| Qsox1 | ENSMUSG( | ENSMUSG( | -0.10464 | 0.244442 | NO | NO | NA |
| Crybg3 | ENSMUSG( | ENSMUSG( | 0.061092 | 0.478091 | NO | NO | NA |
| Klhl6 | ENSMUSG( | ENSMUSG( | -0.09528 | 0.282716 | NO | NO | NA |
| Depdc1a | ENSMUSG( | ENSMUSG( | 0.011887 | 0.929502 | NO | NO | NA |
| Krcc1 | ENSMUSG( | ENSMUSG( | -0.13894 | 0.18152 | NO | NO | NA |
| Trmt6 | ENSMUSG( | ENSMUSG( | 0.446018 | 3.52E-09 | YES | NO | NA |
| Prpf4 | ENSMUSG( | ENSMUSG( | 0.108956 | 0.069404 | NO | NO | NA |
| Plgrkt | ENSMUSG( | ENSMUSG( | -0.29073 | 0.001196 | YES | NO | NA |
| E2f1 | ENSMUSG( | ENSMUSG( | 0.077352 | 0.321549 | NO | NO | NA |
| Ppp2r2d | ENSMUSG( | ENSMUSG( | 0.079162 | 0.190582 | NO | NO | NA |
| Traf5 | ENSMUSG( | ENSMUSG( | -0.30349 | 6.29E-05 | YES | NO | NA |
| Chmp5 | ENSMUSG( | ENSMUSG( | -0.35179 | 3.08E-06 | YES | NO | NA |
| Gm28439 | ENSMUSG( | ENSMUSG( | -0.38226 | 0.541669 | NO | NO | NA |
| Pde4d | ENSMUSG( | ENSMUSG( | -0.09835 | 0.5946 | NO | NO | NA |
| BC004004 | ENSMUSG( | ENSMUSG( | 0.249875 | 0.010461 | YES | NO | NA |
| Wbp4 | ENSMUSG( | ENSMUSG( | 0.015806 | 0.85784 | NO | NO | NA |
| Dbi | ENSMUSG( | ENSMUSG( | 0.383822 | 0.000498 | YES | NO | NA |
| Rcbtb1 | ENSMUSG( | ENSMUSG( | -0.11834 | 0.085539 | NO | NO | NA |
| Atpif1 | ENSMUSG( | ENSMUSG( | 0.387117 | 0.00079 | YES | NO | NA |
| Smad2 | ENSMUSG( | ENSMUSG( | -0.19827 | 0.058138 | NO | NO | NA |
| Mzt1 | ENSMUSG( | ENSMUSG( | 0.209541 | 0.053721 | NO | NO | NA |
| Themis2 | ENSMUSG( | ENSMUSG( | -0.70584 | 4.58E-09 | YES | NO | NA |
| Herc3 | ENSMUSG( | ENSMUSG( | -0.05434 | 0.658479 | NO | NO | NA |
| Ampd2 | ENSMUSG( | ENSMUSG( | 0.26855 | 0.003192 | YES | NO | NA |
| Tmem43 | ENSMUSG( | ENSMUSG( | -0.32541 | 2.05E-06 | YES | NO | NA |
| Mis12 | ENSMUSG( | ENSMUSG( | -0.18143 | 0.012235 | YES | NO | NA |
| Fbxl20 | ENSMUSG( | ENSMUSG( | -0.30987 | 0.0825 | NO | NO | NA |
| 1810013L2 | ENSMUSG( | ENSMUSG( | -0.31875 | 3.77E-07 | YES | NO | NA |
| Topors | ENSMUSG( | ENSMUSG( | -0.15237 | 0.068097 | NO | NO | NA |
| Lman1 | ENSMUSG( | ENSMUSG( | 0.114884 | 0.097841 | NO | NO | NA |
| Mrpl12 | ENSMUSG( | ENSMUSG( | 0.458262 | 2.53E-07 | YES | NO | NA |
| Ptprj | ENSMUSG( | ENSMUSG( | 0.043497 | 0.685397 | NO | NO | NA |
| Psme3ip1 | ENSMUSG( | ENSMUSG( | -0.08287 | 0.362321 | NO | NO | NA |
| Atp6v1c1 | ENSMUSG( | ENSMUSG( | -0.11272 | 0.160835 | NO | NO | NA |

| | | | | | | | |
|---|---|---|---|---|---|---|---|
| Gatad1 | ENSMUSG( | ENSMUSG( | -0.07733 | 0.13609 | NO | NO | NA |
| Mcrs1 | ENSMUSG( | ENSMUSG( | 0.136868 | 0.060251 | NO | NO | NA |
| Mid1 | ENSMUSG( | ENSMUSG( | 0.076466 | 0.923171 | NO | NO | NA |
| Cdca5 | ENSMUSG( | ENSMUSG( | 0.307325 | 3.27E-06 | YES | NO | NA |
| Maf | ENSMUSG( | ENSMUSG( | -0.60855 | 0.000447 | YES | NO | NA |
| Bst2 | ENSMUSG( | ENSMUSG( | -0.46991 | 1.96E-09 | YES | NO | NA |
| Marchf2 | ENSMUSG( | ENSMUSG( | -0.14025 | 0.239113 | NO | NO | NA |
| Prpf3 | ENSMUSG( | ENSMUSG( | 0.003231 | 0.977213 | NO | NO | NA |
| Ddx41 | ENSMUSG( | ENSMUSG( | 0.177232 | 0.066803 | NO | NO | NA |
| Ahcyl2 | ENSMUSG( | ENSMUSG( | -0.22444 | 0.001727 | YES | NO | NA |
| Smad7 | ENSMUSG( | ENSMUSG( | -0.40457 | 0.003739 | YES | NO | NA |
| Rbm7 | ENSMUSG( | ENSMUSG( | -0.00256 | 0.975272 | NO | NO | NA |
| Eif1a | ENSMUSG( | ENSMUSG( | 0.274347 | 0.006963 | YES | NO | NA |
| Stx5a | ENSMUSG( | ENSMUSG( | -0.1531 | 0.117208 | NO | NO | NA |
| Gm10123 | ENSMUSG( | ENSMUSG( | 0.202757 | 0.076279 | NO | NO | NA |
| Sec24a | ENSMUSG( | ENSMUSG( | -0.43284 | 4.03E-09 | YES | NO | NA |
| Lmbr1l | ENSMUSG( | ENSMUSG( | -0.19541 | 0.050613 | NO | NO | NA |
| S100pbp | ENSMUSG( | ENSMUSG( | -0.16804 | 0.072352 | NO | NO | NA |
| Ankrd28 | ENSMUSG( | ENSMUSG( | 0.361724 | 5.90E-05 | YES | NO | NA |
| Rrp12 | ENSMUSG( | ENSMUSG( | 0.730612 | 3.27E-06 | YES | NO | NA |
| Rmnd5b | ENSMUSG( | ENSMUSG( | 0.242374 | 0.007246 | YES | NO | NA |
| Xbp1 | ENSMUSG( | ENSMUSG( | -0.13974 | 0.174199 | NO | NO | NA |
| Zfp451 | ENSMUSG( | ENSMUSG( | -0.23967 | 8.36E-05 | YES | NO | NA |
| Mroh1 | ENSMUSG( | ENSMUSG( | -0.37439 | 3.83E-08 | YES | NO | NA |
| Rhno1 | ENSMUSG( | ENSMUSG( | 0.032808 | 0.536342 | NO | NO | NA |
| Gsap | ENSMUSG( | ENSMUSG( | -0.49634 | 2.98E-11 | YES | NO | NA |
| Ints13 | ENSMUSG( | ENSMUSG( | -0.11229 | 0.279867 | NO | NO | NA |
| Lnx2 | ENSMUSG( | ENSMUSG( | -0.3715 | 7.33E-06 | YES | NO | NA |
| Ptar1 | ENSMUSG( | ENSMUSG( | -0.1791 | 0.004035 | YES | NO | NA |
| Gtf2h1 | ENSMUSG( | ENSMUSG( | 0.142955 | 0.189707 | NO | NO | NA |
| Btbd7 | ENSMUSG( | ENSMUSG( | -0.31448 | 3.48E-08 | YES | NO | NA |
| Prps1 | ENSMUSG( | ENSMUSG( | 0.384551 | 1.50E-06 | YES | NO | NA |
| Evl | ENSMUSG( | ENSMUSG( | -0.33098 | 3.10E-08 | YES | NO | NA |
| Atp10d | ENSMUSG( | ENSMUSG( | -0.71671 | 4.07E-09 | YES | NO | NA |
| Commd3 | ENSMUSG( | ENSMUSG( | 0.016091 | 0.858854 | NO | NO | NA |
| Tmem183a | ENSMUSG( | ENSMUSG( | 0.21387 | 0.000272 | YES | NO | NA |
| Trim24 | ENSMUSG( | ENSMUSG( | 0.129817 | 0.203979 | NO | NO | NA |
| Irf2bp1 | ENSMUSG( | ENSMUSG( | -0.23521 | 8.16E-05 | YES | NO | NA |
| Pou6f1 | ENSMUSG( | ENSMUSG( | -1.04558 | 0.000747 | YES | YES | Downregulated |
| Samsn1 | ENSMUSG( | ENSMUSG( | -0.37105 | 0.001665 | YES | NO | NA |
| Zfp280b | ENSMUSG( | ENSMUSG( | -0.05979 | 0.299148 | NO | NO | NA |
| Rbm45 | ENSMUSG( | ENSMUSG( | -0.00837 | 0.9084 | NO | NO | NA |
| Gpr107 | ENSMUSG( | ENSMUSG( | -0.34214 | 2.43E-05 | YES | NO | NA |
| Gtse1 | ENSMUSG( | ENSMUSG( | 0.139283 | 0.216209 | NO | NO | NA |
| Nom1 | ENSMUSG( | ENSMUSG( | 0.274768 | 0.000614 | YES | NO | NA |
| Mysm1 | ENSMUSG( | ENSMUSG( | -0.2208 | 0.053551 | NO | NO | NA |
| Gnpat | ENSMUSG( | ENSMUSG( | -0.02727 | 0.707387 | NO | NO | NA |

| | | | | | | | |
|---|---|---|---|---|---|---|---|
| Rpap1 | ENSMUSG( | ENSMUSG( | 0.034298 | 0.617105 | NO | NO | NA |
| Synj2bp | ENSMUSG( | ENSMUSG( | -0.27195 | 6.88E-05 | YES | NO | NA |
| Ppp1r14b | ENSMUSG( | ENSMUSG( | 0.565786 | 8.89E-08 | YES | NO | NA |
| Vps11 | ENSMUSG( | ENSMUSG( | 0.046115 | 0.736541 | NO | NO | NA |
| Dctn4 | ENSMUSG( | ENSMUSG( | -0.05937 | 0.427669 | NO | NO | NA |
| Sptlc1 | ENSMUSG( | ENSMUSG( | 0.061016 | 0.531711 | NO | NO | NA |
| Spcs1 | ENSMUSG( | ENSMUSG( | 0.103468 | 0.346781 | NO | NO | NA |
| Farsb | ENSMUSG( | ENSMUSG( | 0.327075 | 3.24E-07 | YES | NO | NA |
| Usp36 | ENSMUSG( | ENSMUSG( | -0.00564 | 0.950071 | NO | NO | NA |
| Scand1 | ENSMUSG( | ENSMUSG( | 0.16375 | 0.013929 | YES | NO | NA |
| Pmm2 | ENSMUSG( | ENSMUSG( | -0.04013 | 0.638583 | NO | NO | NA |
| Arfgap1 | ENSMUSG( | ENSMUSG( | -0.05264 | 0.562672 | NO | NO | NA |
| Vps29 | ENSMUSG( | ENSMUSG( | -0.06548 | 0.446439 | NO | NO | NA |
| Chkb | ENSMUSG( | ENSMUSG( | -0.13721 | 0.063052 | NO | NO | NA |
| Ptprs | ENSMUSG( | ENSMUSG( | 0.715863 | 3.63E-12 | YES | NO | NA |
| Mff | ENSMUSG( | ENSMUSG( | -0.18694 | 0.014413 | YES | NO | NA |
| Neb | ENSMUSG( | ENSMUSG( | -0.76438 | 7.74E-09 | YES | NO | NA |
| Kbtbd2 | ENSMUSG( | ENSMUSG( | -0.35653 | 1.83E-07 | YES | NO | NA |
| Las1l | ENSMUSG( | ENSMUSG( | 0.217072 | 0.056765 | NO | NO | NA |
| Slamf1 | ENSMUSG( | ENSMUSG( | -0.35834 | 0.001831 | YES | NO | NA |
| Pde4dip | ENSMUSG( | ENSMUSG( | -0.30043 | 1.12E-05 | YES | NO | NA |
| Cisd2 | ENSMUSG( | ENSMUSG( | -0.3413 | 0.000117 | YES | NO | NA |
| Tor1b | ENSMUSG( | ENSMUSG( | -0.13989 | 0.023674 | YES | NO | NA |
| Vps4a | ENSMUSG( | ENSMUSG( | -0.20152 | 0.000112 | YES | NO | NA |
| Hvcn1 | ENSMUSG( | ENSMUSG( | -0.05891 | 0.572582 | NO | NO | NA |
| Ppp1r10 | ENSMUSG( | ENSMUSG( | -0.25498 | 2.66E-05 | YES | NO | NA |
| Men1 | ENSMUSG( | ENSMUSG( | -0.10575 | 0.246461 | NO | NO | NA |
| Heatr6 | ENSMUSG( | ENSMUSG( | -0.10197 | 0.088563 | NO | NO | NA |
| Cog2 | ENSMUSG( | ENSMUSG( | -0.04703 | 0.553112 | NO | NO | NA |
| Gclc | ENSMUSG( | ENSMUSG( | 0.140097 | 0.237132 | NO | NO | NA |
| Rnf216 | ENSMUSG( | ENSMUSG( | -0.26363 | 0.002502 | YES | NO | NA |
| Rtp4 | ENSMUSG( | ENSMUSG( | -1.22527 | 3.16E-17 | YES | YES | Downregulated |
| Chmp3 | ENSMUSG( | ENSMUSG( | -0.06268 | 0.430364 | NO | NO | NA |
| Mrps9 | ENSMUSG( | ENSMUSG( | 0.423116 | 4.69E-06 | YES | NO | NA |
| Pgp | ENSMUSG( | ENSMUSG( | 0.366263 | 7.75E-07 | YES | NO | NA |
| Atr | ENSMUSG( | ENSMUSG( | 0.086728 | 0.262034 | NO | NO | NA |
| Fbxl5 | ENSMUSG( | ENSMUSG( | 0.01832 | 0.856111 | NO | NO | NA |
| Aurkaip1 | ENSMUSG( | ENSMUSG( | 0.0948 | 0.049003 | YES | NO | NA |
| Mxi1 | ENSMUSG( | ENSMUSG( | -0.29058 | 0.016357 | YES | NO | NA |
| Ssr4 | ENSMUSG( | ENSMUSG( | 0.253952 | 0.000803 | YES | NO | NA |
| Vps41 | ENSMUSG( | ENSMUSG( | -0.14973 | 0.085149 | NO | NO | NA |
| Tmed8 | ENSMUSG( | ENSMUSG( | -0.38353 | 0.000111 | YES | NO | NA |
| Thoc1 | ENSMUSG( | ENSMUSG( | 0.313612 | 0.000102 | YES | NO | NA |
| Rps27l | ENSMUSG( | ENSMUSG( | 0.352307 | 0.000117 | YES | NO | NA |
| Plekhm1 | ENSMUSG( | ENSMUSG( | -0.42576 | 1.66E-09 | YES | NO | NA |
| Trappc11 | ENSMUSG( | ENSMUSG( | -0.10708 | 0.109767 | NO | NO | NA |
| Fam117a | ENSMUSG( | ENSMUSG( | -0.47517 | 3.30E-06 | YES | NO | NA |

| | | | | | | | |
|---|---|---|---|---|---|---|---|
| Shcbp1 | ENSMUSG( | ENSMUSG( | 0.246585 | 0.004869 | YES | NO | NA |
| Cdkn2d | ENSMUSG( | ENSMUSG( | -0.1034 | 0.308915 | NO | NO | NA |
| Eid1 | ENSMUSG( | ENSMUSG( | 0.164935 | 0.024199 | YES | NO | NA |
| Sucla2 | ENSMUSG( | ENSMUSG( | 0.203157 | 0.002213 | YES | NO | NA |
| Gnl2 | ENSMUSG( | ENSMUSG( | 0.381715 | 0.000105 | YES | NO | NA |
| Idi1 | ENSMUSG( | ENSMUSG( | 0.69375 | 1.81E-05 | YES | NO | NA |
| Trim11 | ENSMUSG( | ENSMUSG( | -0.26682 | 7.75E-06 | YES | NO | NA |
| Uqcr10 | ENSMUSG( | ENSMUSG( | 0.404608 | 1.38E-05 | YES | NO | NA |
| Stil | ENSMUSG( | ENSMUSG( | 0.120086 | 0.232063 | NO | NO | NA |
| Fuca2 | ENSMUSG( | ENSMUSG( | -0.08926 | 0.348227 | NO | NO | NA |
| Vti1b | ENSMUSG( | ENSMUSG( | 0.113192 | 0.102087 | NO | NO | NA |
| Usp12 | ENSMUSG( | ENSMUSG( | -0.33606 | 6.14E-06 | YES | NO | NA |
| Fuca1 | ENSMUSG( | ENSMUSG( | 0.232373 | 0.003761 | YES | NO | NA |
| Rsl24d1 | ENSMUSG( | ENSMUSG( | 0.194207 | 0.007244 | YES | NO | NA |
| Relch | ENSMUSG( | ENSMUSG( | -0.15616 | 0.198849 | NO | NO | NA |
| Cep152 | ENSMUSG( | ENSMUSG( | 0.151308 | 0.184715 | NO | NO | NA |
| Sp140 | ENSMUSG( | ENSMUSG( | -0.25786 | 0.001602 | YES | NO | NA |
| Nop9 | ENSMUSG( | ENSMUSG( | 0.304388 | 1.24E-05 | YES | NO | NA |
| Macir | ENSMUSG( | ENSMUSG( | -0.55404 | 1.35E-07 | YES | NO | NA |
| Fastk | ENSMUSG( | ENSMUSG( | -0.05376 | 0.214873 | NO | NO | NA |
| Atg16l2 | ENSMUSG( | ENSMUSG( | -0.14282 | 0.024476 | YES | NO | NA |
| Syne2 | ENSMUSG( | ENSMUSG( | -0.2649 | 1.23E-05 | YES | NO | NA |
| Nup133 | ENSMUSG( | ENSMUSG( | 0.481059 | 4.00E-07 | YES | NO | NA |
| Arfrp1 | ENSMUSG( | ENSMUSG( | -0.32351 | 1.65E-06 | YES | NO | NA |
| Bcap29 | ENSMUSG( | ENSMUSG( | 0.563118 | 5.11E-07 | YES | NO | NA |
| Ppan | ENSMUSG( | ENSMUSG( | 0.409213 | 0.000108 | YES | NO | NA |
| Lats1 | ENSMUSG( | ENSMUSG( | -0.15446 | 0.112117 | NO | NO | NA |
| Pecam1 | ENSMUSG( | ENSMUSG( | 0.675834 | 1.01E-05 | YES | NO | NA |
| Frg1 | ENSMUSG( | ENSMUSG( | -0.2609 | 0.032682 | YES | NO | NA |
| Ccdc117 | ENSMUSG( | ENSMUSG( | -0.79935 | 2.81E-15 | YES | NO | NA |
| Zfp251 | ENSMUSG( | ENSMUSG( | -0.29217 | 0.00111 | YES | NO | NA |
| Nup54 | ENSMUSG( | ENSMUSG( | 0.176953 | 0.062966 | NO | NO | NA |
| Agtpbp1 | ENSMUSG( | ENSMUSG( | -0.22873 | 4.30E-08 | YES | NO | NA |
| Minpp1 | ENSMUSG( | ENSMUSG( | 0.266029 | 1.57E-05 | YES | NO | NA |
| Dvl1 | ENSMUSG( | ENSMUSG( | -0.19822 | 0.000867 | YES | NO | NA |
| Snx19 | ENSMUSG( | ENSMUSG( | -0.19423 | 0.002575 | YES | NO | NA |
| Pacs2 | ENSMUSG( | ENSMUSG( | -0.02968 | 0.729933 | NO | NO | NA |
| Mrps26 | ENSMUSG( | ENSMUSG( | 0.27247 | 0.00042 | YES | NO | NA |
| Sys1 | ENSMUSG( | ENSMUSG( | 0.033996 | 0.717512 | NO | NO | NA |
| Socs7 | ENSMUSG( | ENSMUSG( | -0.21492 | 0.000706 | YES | NO | NA |
| Fem1c | ENSMUSG( | ENSMUSG( | -0.29219 | 1.95E-05 | YES | NO | NA |
| Faf1 | ENSMUSG( | ENSMUSG( | -0.05016 | 0.507817 | NO | NO | NA |
| Hsd17b12 | ENSMUSG( | ENSMUSG( | 0.297906 | 0.004221 | YES | NO | NA |
| Sptssa | ENSMUSG( | ENSMUSG( | 0.292572 | 9.56E-05 | YES | NO | NA |
| Nop2 | ENSMUSG( | ENSMUSG( | 0.541391 | 6.54E-06 | YES | NO | NA |
| Dus3l | ENSMUSG( | ENSMUSG( | 0.175361 | 0.006617 | YES | NO | NA |
| Cyth2 | ENSMUSG( | ENSMUSG( | 0.017849 | 0.859925 | NO | NO | NA |

| | | | | | | | |
|---|---|---|---|---|---|---|---|
| Ubxn8 | ENSMUSG( | ENSMUSG( | 0.227298 | 0.010266 | YES | NO | NA |
| Pias3 | ENSMUSG( | ENSMUSG( | -0.3562 | 9.82E-06 | YES | NO | NA |
| mt-Nd3 | ENSMUSG( | ENSMUSG( | 0.426657 | 0.287329 | NO | NO | NA |
| Slc30a5 | ENSMUSG( | ENSMUSG( | 0.105601 | 0.174694 | NO | NO | NA |
| Cog3 | ENSMUSG( | ENSMUSG( | -0.03102 | 0.667184 | NO | NO | NA |
| Usp37 | ENSMUSG( | ENSMUSG( | -0.22653 | 0.000115 | YES | NO | NA |
| Fam207a | ENSMUSG( | ENSMUSG( | 0.053434 | 0.456591 | NO | NO | NA |
| Tmem87a | ENSMUSG( | ENSMUSG( | -0.15691 | 0.006457 | YES | NO | NA |
| Tmem209 | ENSMUSG( | ENSMUSG( | 0.096282 | 0.253615 | NO | NO | NA |
| Zfp131 | ENSMUSG( | ENSMUSG( | -0.13281 | 0.094896 | NO | NO | NA |
| Marchf5 | ENSMUSG( | ENSMUSG( | -0.0675 | 0.516168 | NO | NO | NA |
| Mtr | ENSMUSG( | ENSMUSG( | -0.16249 | 0.023529 | YES | NO | NA |
| Senp5 | ENSMUSG( | ENSMUSG( | -0.13287 | 0.056896 | NO | NO | NA |
| Pik3c3 | ENSMUSG( | ENSMUSG( | -0.03954 | 0.622643 | NO | NO | NA |
| Prdm4 | ENSMUSG( | ENSMUSG( | 0.16679 | 0.04662 | YES | NO | NA |
| Ctc1 | ENSMUSG( | ENSMUSG( | 0.377602 | 2.93E-05 | YES | NO | NA |
| Rab3gap2 | ENSMUSG( | ENSMUSG( | -4.02E-05 | 0.999611 | NO | NO | NA |
| Ino80 | ENSMUSG( | ENSMUSG( | -0.14564 | 0.001638 | YES | NO | NA |
| Timm50 | ENSMUSG( | ENSMUSG( | 0.524319 | 6.33E-08 | YES | NO | NA |
| Fhod1 | ENSMUSG( | ENSMUSG( | -0.26524 | 9.32E-11 | YES | NO | NA |
| Tbc1d15 | ENSMUSG( | ENSMUSG( | 0.053678 | 0.609667 | NO | NO | NA |
| Aifm1 | ENSMUSG( | ENSMUSG( | 0.254474 | 0.002216 | YES | NO | NA |
| Gmeb2 | ENSMUSG( | ENSMUSG( | -0.03919 | 0.60683 | NO | NO | NA |
| Atf7 | ENSMUSG( | ENSMUSG( | -0.35805 | 1.62E-08 | YES | NO | NA |
| Ddx49 | ENSMUSG( | ENSMUSG( | 0.295146 | 0.004647 | YES | NO | NA |
| Fundc2 | ENSMUSG( | ENSMUSG( | 0.08676 | 0.138589 | NO | NO | NA |
| Usp14 | ENSMUSG( | ENSMUSG( | 0.19929 | 0.002176 | YES | NO | NA |
| Dap3 | ENSMUSG( | ENSMUSG( | 0.344984 | 4.19E-05 | YES | NO | NA |
| Flywch1 | ENSMUSG( | ENSMUSG( | 0.154361 | 0.195678 | NO | NO | NA |
| Brms1 | ENSMUSG( | ENSMUSG( | 0.149618 | 0.00127 | YES | NO | NA |
| Gm26740 | ENSMUSG( | ENSMUSG( | -0.60619 | 0.000159 | YES | NO | NA |
| Fam89b | ENSMUSG( | ENSMUSG( | 0.008149 | 0.930482 | NO | NO | NA |
| Akap11 | ENSMUSG( | ENSMUSG( | -0.1277 | 0.10548 | NO | NO | NA |
| Arhgap17 | ENSMUSG( | ENSMUSG( | -0.23658 | 1.49E-05 | YES | NO | NA |
| Sos2 | ENSMUSG( | ENSMUSG( | -0.04066 | 0.765827 | NO | NO | NA |
| Als2 | ENSMUSG( | ENSMUSG( | -0.74038 | 8.66E-16 | YES | NO | NA |
| Tbl2 | ENSMUSG( | ENSMUSG( | -0.20894 | 7.06E-07 | YES | NO | NA |
| Zdhhc8 | ENSMUSG( | ENSMUSG( | -0.0273 | 0.685592 | NO | NO | NA |
| Pdcl3 | ENSMUSG( | ENSMUSG( | -0.10807 | 0.449709 | NO | NO | NA |
| Smim15 | ENSMUSG( | ENSMUSG( | 0.050672 | 0.589716 | NO | NO | NA |
| Thoc5 | ENSMUSG( | ENSMUSG( | 0.102137 | 0.231177 | NO | NO | NA |
| Gmeb1 | ENSMUSG( | ENSMUSG( | -0.02391 | 0.603197 | NO | NO | NA |
| Dis3 | ENSMUSG( | ENSMUSG( | 0.421227 | 1.63E-07 | YES | NO | NA |
| Hopx | ENSMUSG( | ENSMUSG( | -0.34009 | 2.90E-05 | YES | NO | NA |
| Atmin | ENSMUSG( | ENSMUSG( | -0.11987 | 0.254587 | NO | NO | NA |
| Naa20 | ENSMUSG( | ENSMUSG( | -0.15436 | 0.008511 | YES | NO | NA |
| Tbcb | ENSMUSG( | ENSMUSG( | 0.187088 | 0.057487 | NO | NO | NA |

| | | | | | | | |
|---|---|---|---|---|---|---|---|
| Parl | ENSMUSG( | ENSMUSG( | 0.367114 | 9.32E-06 | YES | NO | NA |
| Haus3 | ENSMUSG( | ENSMUSG( | -0.2185 | 0.009908 | YES | NO | NA |
| Zfp362 | ENSMUSG( | ENSMUSG( | -0.11888 | 0.378631 | NO | NO | NA |
| Rnf19a | ENSMUSG( | ENSMUSG( | 0.008869 | 0.94162 | NO | NO | NA |
| Sarnp | ENSMUSG( | ENSMUSG( | -0.03095 | 0.810325 | NO | NO | NA |
| Kmt5c | ENSMUSG( | ENSMUSG( | 0.017715 | 0.802041 | NO | NO | NA |
| Tbc1d13 | ENSMUSG( | ENSMUSG( | -0.26044 | 0.000102 | YES | NO | NA |
| Gfm1 | ENSMUSG( | ENSMUSG( | 0.391581 | 6.51E-07 | YES | NO | NA |
| 0610030E2 | ENSMUSG( | ENSMUSG( | -0.25683 | 0.000104 | YES | NO | NA |
| Nsrp1 | ENSMUSG( | ENSMUSG( | -0.11974 | 0.524266 | NO | NO | NA |
| Dapk3 | ENSMUSG( | ENSMUSG( | -0.01492 | 0.854465 | NO | NO | NA |
| Trp53bp1 | ENSMUSG( | ENSMUSG( | -0.15526 | 0.019641 | YES | NO | NA |
| Polr3b | ENSMUSG( | ENSMUSG( | 0.00226 | 0.975732 | NO | NO | NA |
| Spast | ENSMUSG( | ENSMUSG( | -0.20567 | 0.003135 | YES | NO | NA |
| Sypl | ENSMUSG( | ENSMUSG( | 0.21235 | 0.014241 | YES | NO | NA |
| Oxr1 | ENSMUSG( | ENSMUSG( | -0.36289 | 2.64E-05 | YES | NO | NA |
| Zfp831 | ENSMUSG( | ENSMUSG( | -1.38034 | 3.11E-11 | YES | YES | Downregulated |
| Dus1l | ENSMUSG( | ENSMUSG( | 0.282214 | 8.59E-05 | YES | NO | NA |
| Tor2a | ENSMUSG( | ENSMUSG( | 0.209609 | 0.025184 | YES | NO | NA |
| Mib1 | ENSMUSG( | ENSMUSG( | -0.00442 | 0.966048 | NO | NO | NA |
| Ube2g1 | ENSMUSG( | ENSMUSG( | -0.03697 | 0.664677 | NO | NO | NA |
| Pygo2 | ENSMUSG( | ENSMUSG( | 0.020376 | 0.818237 | NO | NO | NA |
| Kat2a | ENSMUSG( | ENSMUSG( | 0.12921 | 0.037109 | YES | NO | NA |
| Hps5 | ENSMUSG( | ENSMUSG( | -0.25825 | 0.000584 | YES | NO | NA |
| Tmem135 | ENSMUSG( | ENSMUSG( | -0.6458 | 5.80E-08 | YES | NO | NA |
| Pgm2l1 | ENSMUSG( | ENSMUSG( | -0.96751 | 7.16E-15 | YES | NO | NA |
| Lamtor1 | ENSMUSG( | ENSMUSG( | 0.041883 | 0.706185 | NO | NO | NA |
| Thoc3 | ENSMUSG( | ENSMUSG( | 0.452903 | 3.13E-06 | YES | NO | NA |
| Mtmr2 | ENSMUSG( | ENSMUSG( | 0.062835 | 0.481958 | NO | NO | NA |
| Dpysl2 | ENSMUSG( | ENSMUSG( | -0.49035 | 3.51E-08 | YES | NO | NA |
| Metap1 | ENSMUSG( | ENSMUSG( | 0.252174 | 6.40E-06 | YES | NO | NA |
| Trim14 | ENSMUSG( | ENSMUSG( | -0.51279 | 2.37E-10 | YES | NO | NA |
| Por | ENSMUSG( | ENSMUSG( | 0.083952 | 0.192483 | NO | NO | NA |
| Foxp1 | ENSMUSG( | ENSMUSG( | -0.42717 | 1.47E-07 | YES | NO | NA |
| Pttg1ip | ENSMUSG( | ENSMUSG( | -0.22867 | 0.004617 | YES | NO | NA |
| Atp6v1e1 | ENSMUSG( | ENSMUSG( | -0.17178 | 0.004972 | YES | NO | NA |
| Traf2 | ENSMUSG( | ENSMUSG( | 0.020848 | 0.743006 | NO | NO | NA |
| Gga3 | ENSMUSG( | ENSMUSG( | -0.09435 | 0.130973 | NO | NO | NA |
| Chmp7 | ENSMUSG( | ENSMUSG( | -0.11088 | 0.042273 | YES | NO | NA |
| Ints14 | ENSMUSG( | ENSMUSG( | 0.207504 | 0.006718 | YES | NO | NA |
| Eif2ak3 | ENSMUSG( | ENSMUSG( | 0.301264 | 0.010035 | YES | NO | NA |
| Wipi2 | ENSMUSG( | ENSMUSG( | -0.09148 | 0.053109 | NO | NO | NA |
| Ubqln2 | ENSMUSG( | ENSMUSG( | -0.17432 | 0.002315 | YES | NO | NA |
| Lgalsl | ENSMUSG( | ENSMUSG( | -0.35502 | 0.001358 | YES | NO | NA |
| Acat1 | ENSMUSG( | ENSMUSG( | 0.519132 | 4.69E-06 | YES | NO | NA |
| Chtf18 | ENSMUSG( | ENSMUSG( | 0.37625 | 8.01E-06 | YES | NO | NA |
| Nefh | ENSMUSG( | ENSMUSG( | 0.486934 | 0.018198 | YES | NO | NA |

| | | | | | | | |
|---|---|---|---|---|---|---|---|
| Capg | ENSMUSG( | ENSMUSG( | 0.90885 | 2.71E-09 | YES | NO | NA |
| Depdc5 | ENSMUSG( | ENSMUSG( | -0.1103 | 0.058607 | NO | NO | NA |
| Atg16l1 | ENSMUSG( | ENSMUSG( | -0.19525 | 0.022205 | YES | NO | NA |
| Clcf1 | ENSMUSG( | ENSMUSG( | -0.66407 | 7.08E-07 | YES | NO | NA |
| Prkd3 | ENSMUSG( | ENSMUSG( | -0.07615 | 0.207706 | NO | NO | NA |
| Glrx5 | ENSMUSG( | ENSMUSG( | 0.352291 | 1.35E-05 | YES | NO | NA |
| Akap8l | ENSMUSG( | ENSMUSG( | -0.36571 | 6.29E-05 | YES | NO | NA |
| Prpf38a | ENSMUSG( | ENSMUSG( | 0.275461 | 1.34E-05 | YES | NO | NA |
| Rnf169 | ENSMUSG( | ENSMUSG( | -0.3682 | 0.00014 | YES | NO | NA |
| Zbtb11 | ENSMUSG( | ENSMUSG( | -0.26018 | 0.000622 | YES | NO | NA |
| Trappc12 | ENSMUSG( | ENSMUSG( | -0.19771 | 0.00148 | YES | NO | NA |
| Hccs | ENSMUSG( | ENSMUSG( | 0.184525 | 0.126645 | NO | NO | NA |
| Fbxo22 | ENSMUSG( | ENSMUSG( | -0.10658 | 0.048121 | YES | NO | NA |
| Raph1 | ENSMUSG( | ENSMUSG( | 0.031356 | 0.807853 | NO | NO | NA |
| Dvl3 | ENSMUSG( | ENSMUSG( | -0.20351 | 0.002014 | YES | NO | NA |
| Pxk | ENSMUSG( | ENSMUSG( | -0.2147 | 0.008308 | YES | NO | NA |
| Tbcel | ENSMUSG( | ENSMUSG( | -0.23875 | 0.000511 | YES | NO | NA |
| Kif1b | ENSMUSG( | ENSMUSG( | -0.95423 | 3.02E-18 | YES | NO | NA |
| Hgs | ENSMUSG( | ENSMUSG( | -0.2231 | 0.00233 | YES | NO | NA |
| Sprtn | ENSMUSG( | ENSMUSG( | -0.18775 | 0.03798 | YES | NO | NA |
| Tmem33 | ENSMUSG( | ENSMUSG( | -0.24131 | 0.000965 | YES | NO | NA |
| Sec24d | ENSMUSG( | ENSMUSG( | -0.06138 | 0.52971 | NO | NO | NA |
| Blm | ENSMUSG( | ENSMUSG( | 0.412595 | 7.82E-08 | YES | NO | NA |
| Tnks | ENSMUSG( | ENSMUSG( | -0.28353 | 0.002562 | YES | NO | NA |
| Mpnd | ENSMUSG( | ENSMUSG( | -0.05207 | 0.526309 | NO | NO | NA |
| Cdk8 | ENSMUSG( | ENSMUSG( | 0.14291 | 0.02575 | YES | NO | NA |
| Tasor | ENSMUSG( | ENSMUSG( | -0.497 | 5.61E-09 | YES | NO | NA |
| Klhl24 | ENSMUSG( | ENSMUSG( | -0.55034 | 3.62E-05 | YES | NO | NA |
| Mbnl3 | ENSMUSG( | ENSMUSG( | 0.294108 | 2.19E-05 | YES | NO | NA |
| Ndrg3 | ENSMUSG( | ENSMUSG( | -0.23936 | 3.33E-05 | YES | NO | NA |
| Pelp1 | ENSMUSG( | ENSMUSG( | 0.139656 | 0.129289 | NO | NO | NA |
| N4bp2 | ENSMUSG( | ENSMUSG( | 0.367386 | 1.95E-05 | YES | NO | NA |
| Terf2 | ENSMUSG( | ENSMUSG( | 0.032763 | 0.745859 | NO | NO | NA |
| Zbtb2 | ENSMUSG( | ENSMUSG( | -0.49068 | 4.10E-08 | YES | NO | NA |
| Ndufs3 | ENSMUSG( | ENSMUSG( | 0.200174 | 0.017051 | YES | NO | NA |
| Cops3 | ENSMUSG( | ENSMUSG( | 0.255758 | 0.001323 | YES | NO | NA |
| S100a13 | ENSMUSG( | ENSMUSG( | -0.32082 | 4.88E-06 | YES | NO | NA |
| Rps6kb2 | ENSMUSG( | ENSMUSG( | 0.034154 | 0.588985 | NO | NO | NA |
| Polr3k | ENSMUSG( | ENSMUSG( | 0.220634 | 0.011949 | YES | NO | NA |
| Dcun1d1 | ENSMUSG( | ENSMUSG( | -0.09721 | 0.327176 | NO | NO | NA |
| Pde12 | ENSMUSG( | ENSMUSG( | 0.11857 | 0.033027 | YES | NO | NA |
| Slc23a2 | ENSMUSG( | ENSMUSG( | -0.46328 | 3.27E-07 | YES | NO | NA |
| Stoml2 | ENSMUSG( | ENSMUSG( | 0.365362 | 0.000121 | YES | NO | NA |
| Tapt1 | ENSMUSG( | ENSMUSG( | -0.01483 | 0.88631 | NO | NO | NA |
| Rb1cc1 | ENSMUSG( | ENSMUSG( | -0.51478 | 8.28E-07 | YES | NO | NA |
| Fkbp2 | ENSMUSG( | ENSMUSG( | 0.126824 | 0.573192 | NO | NO | NA |
| Gsto1 | ENSMUSG( | ENSMUSG( | 1.041757 | 1.38E-08 | YES | YES | Upregulated |

| | | | | | | | |
|---|---|---|---|---|---|---|---|
| Cep135 | ENSMUSG( | ENSMUSG( | 0.023738 | 0.797391 | NO | NO | NA |
| Anapc11 | ENSMUSG( | ENSMUSG( | -0.09991 | 0.185287 | NO | NO | NA |
| Sgo2a | ENSMUSG( | ENSMUSG( | 0.077794 | 0.468472 | NO | NO | NA |
| Senp7 | ENSMUSG( | ENSMUSG( | -0.52971 | 1.18E-07 | YES | NO | NA |
| Rsbn1 | ENSMUSG( | ENSMUSG( | -0.17966 | 0.002726 | YES | NO | NA |
| Edc3 | ENSMUSG( | ENSMUSG( | -0.06412 | 0.455434 | NO | NO | NA |
| Usp20 | ENSMUSG( | ENSMUSG( | -0.24507 | 0.006257 | YES | NO | NA |
| Slc7a6 | ENSMUSG( | ENSMUSG( | 0.17574 | 0.027101 | YES | NO | NA |
| Rmc1 | ENSMUSG( | ENSMUSG( | -0.38083 | 2.18E-07 | YES | NO | NA |
| Hmgxb4 | ENSMUSG( | ENSMUSG( | -0.07866 | 0.351004 | NO | NO | NA |
| Mis18bp1 | ENSMUSG( | ENSMUSG( | 0.094574 | 0.431293 | NO | NO | NA |
| St8sia4 | ENSMUSG( | ENSMUSG( | -0.4916 | 8.93E-07 | YES | NO | NA |
| Ndufa12 | ENSMUSG( | ENSMUSG( | 0.37671 | 0.003445 | YES | NO | NA |
| Cyb5r3 | ENSMUSG( | ENSMUSG( | 0.048828 | 0.684279 | NO | NO | NA |
| Thoc7 | ENSMUSG( | ENSMUSG( | 0.147604 | 0.033629 | YES | NO | NA |
| Trib2 | ENSMUSG( | ENSMUSG( | 0.330774 | 0.022705 | YES | NO | NA |
| Timm13 | ENSMUSG( | ENSMUSG( | 0.251729 | 0.001477 | YES | NO | NA |
| Gata3 | ENSMUSG( | ENSMUSG( | -0.50582 | 8.96E-08 | YES | NO | NA |
| Pogz | ENSMUSG( | ENSMUSG( | -0.31068 | 1.31E-06 | YES | NO | NA |
| Sirt1 | ENSMUSG( | ENSMUSG( | -0.28267 | 4.04E-06 | YES | NO | NA |
| Map2k4 | ENSMUSG( | ENSMUSG( | -0.14635 | 0.190077 | NO | NO | NA |
| Pdlim5 | ENSMUSG( | ENSMUSG( | -0.56663 | 1.28E-16 | YES | NO | NA |
| Napg | ENSMUSG( | ENSMUSG( | -0.29871 | 0.000215 | YES | NO | NA |
| Foxj2 | ENSMUSG( | ENSMUSG( | -0.34746 | 0.00255 | YES | NO | NA |
| Gpkow | ENSMUSG( | ENSMUSG( | -0.19724 | 0.014477 | YES | NO | NA |
| Wdr37 | ENSMUSG( | ENSMUSG( | -0.09256 | 0.302959 | NO | NO | NA |
| Katnb1 | ENSMUSG( | ENSMUSG( | 0.043937 | 0.680543 | NO | NO | NA |
| Timm44 | ENSMUSG( | ENSMUSG( | 0.207619 | 0.003933 | YES | NO | NA |
| Cdk16 | ENSMUSG( | ENSMUSG( | -0.09365 | 0.153294 | NO | NO | NA |
| Utp18 | ENSMUSG( | ENSMUSG( | 0.288593 | 0.009438 | YES | NO | NA |
| Pgrmc2 | ENSMUSG( | ENSMUSG( | 0.015978 | 0.91236 | NO | NO | NA |
| Ift52 | ENSMUSG( | ENSMUSG( | 0.039592 | 0.724414 | NO | NO | NA |
| Exoc6 | ENSMUSG( | ENSMUSG( | -0.07802 | 0.446809 | NO | NO | NA |
| Zhx1 | ENSMUSG( | ENSMUSG( | -0.0609 | 0.568075 | NO | NO | NA |
| Csnk1g3 | ENSMUSG( | ENSMUSG( | -0.17505 | 0.021739 | YES | NO | NA |
| Chst12 | ENSMUSG( | ENSMUSG( | -0.33135 | 0.000281 | YES | NO | NA |
| Cfdp1 | ENSMUSG( | ENSMUSG( | -0.01148 | 0.93675 | NO | NO | NA |
| Syvn1 | ENSMUSG( | ENSMUSG( | -0.35471 | 1.19E-08 | YES | NO | NA |
| Mex3c | ENSMUSG( | ENSMUSG( | 0.081325 | 0.33719 | NO | NO | NA |
| Sugp2 | ENSMUSG( | ENSMUSG( | -0.04955 | 0.49528 | NO | NO | NA |
| Isca1 | ENSMUSG( | ENSMUSG( | 0.040874 | 0.69132 | NO | NO | NA |
| Gba | ENSMUSG( | ENSMUSG( | -0.27743 | 0.002751 | YES | NO | NA |
| Gprin3 | ENSMUSG( | ENSMUSG( | -0.52967 | 1.56E-05 | YES | NO | NA |
| Tmem160 | ENSMUSG( | ENSMUSG( | 0.220588 | 0.010324 | YES | NO | NA |
| Tbca | ENSMUSG( | ENSMUSG( | -0.11925 | 0.356141 | NO | NO | NA |
| Scarb2 | ENSMUSG( | ENSMUSG( | -0.43049 | 4.18E-07 | YES | NO | NA |
| Snx6 | ENSMUSG( | ENSMUSG( | -0.20279 | 0.00704 | YES | NO | NA |

| | | | | | | | |
|---|---|---|---|---|---|---|---|
| Cstf3 | ENSMUSG( | ENSMUSG( | 0.074276 | 0.436798 | NO | NO | NA |
| Vps51 | ENSMUSG( | ENSMUSG( | 0.016988 | 0.84357 | NO | NO | NA |
| Irf2bp2 | ENSMUSG( | ENSMUSG( | -0.19656 | 0.039979 | YES | NO | NA |
| Galk1 | ENSMUSG( | ENSMUSG( | 0.989909 | 6.61E-08 | YES | NO | NA |
| Rab5if | ENSMUSG( | ENSMUSG( | 0.134488 | 0.05073 | NO | NO | NA |
| Tiprl | ENSMUSG( | ENSMUSG( | -0.02809 | 0.794703 | NO | NO | NA |
| Skil | ENSMUSG( | ENSMUSG( | -0.42324 | 0.000112 | YES | NO | NA |
| BC005624 | ENSMUSG( | ENSMUSG( | -0.19945 | 0.050077 | NO | NO | NA |
| Creld2 | ENSMUSG( | ENSMUSG( | 0.117279 | 0.224995 | NO | NO | NA |
| Ap5m1 | ENSMUSG( | ENSMUSG( | -0.28173 | 0.000342 | YES | NO | NA |
| Exosc9 | ENSMUSG( | ENSMUSG( | 0.414612 | 4.41E-08 | YES | NO | NA |
| Fem1b | ENSMUSG( | ENSMUSG( | -0.0637 | 0.344138 | NO | NO | NA |
| Pi4kb | ENSMUSG( | ENSMUSG( | -0.22182 | 0.000103 | YES | NO | NA |
| Pcmtd1 | ENSMUSG( | ENSMUSG( | -0.60225 | 7.53E-08 | YES | NO | NA |
| Tmem214 | ENSMUSG( | ENSMUSG( | -0.05651 | 0.541333 | NO | NO | NA |
| Tspan32 | ENSMUSG( | ENSMUSG( | 0.242266 | 0.141753 | NO | NO | NA |
| Mrpl10 | ENSMUSG( | ENSMUSG( | 0.103388 | 0.037711 | YES | NO | NA |
| Ctps2 | ENSMUSG( | ENSMUSG( | -0.39286 | 1.82E-07 | YES | NO | NA |
| Upf3b | ENSMUSG( | ENSMUSG( | -0.01255 | 0.94245 | NO | NO | NA |
| Rprd1b | ENSMUSG( | ENSMUSG( | 0.091437 | 0.112587 | NO | NO | NA |
| Iffo1 | ENSMUSG( | ENSMUSG( | -0.05245 | 0.636296 | NO | NO | NA |
| 2310011J( | ENSMUSG( | ENSMUSG( | 0.111101 | 0.171547 | NO | NO | NA |
| Gtf2b | ENSMUSG( | ENSMUSG( | -0.06469 | 0.206733 | NO | NO | NA |
| Rfc3 | ENSMUSG( | ENSMUSG( | 0.356469 | 3.88E-08 | YES | NO | NA |
| Ttc7b | ENSMUSG( | ENSMUSG( | -0.33188 | 0.000263 | YES | NO | NA |
| 1500009L: | ENSMUSG( | ENSMUSG( | 0.896231 | 5.34E-08 | YES | NO | NA |
| Ctnna1 | ENSMUSG( | ENSMUSG( | -0.06385 | 0.598033 | NO | NO | NA |
| Jmjd6 | ENSMUSG( | ENSMUSG( | -0.00182 | 0.9833 | NO | NO | NA |
| Fancd2 | ENSMUSG( | ENSMUSG( | 0.215358 | 0.034587 | YES | NO | NA |
| Egln2 | ENSMUSG( | ENSMUSG( | -0.101 | 0.246634 | NO | NO | NA |
| Brd3 | ENSMUSG( | ENSMUSG( | -0.13098 | 0.055972 | NO | NO | NA |
| Ankrd27 | ENSMUSG( | ENSMUSG( | -0.09831 | 0.165584 | NO | NO | NA |
| Ktn1 | ENSMUSG( | ENSMUSG( | 0.228621 | 0.00404 | YES | NO | NA |
| Cplane1 | ENSMUSG( | ENSMUSG( | -0.13464 | 0.238332 | NO | NO | NA |
| Nrp1 | ENSMUSG( | ENSMUSG( | 0.106943 | 0.593553 | NO | NO | NA |
| Fhip2b | ENSMUSG( | ENSMUSG( | -0.21142 | 0.005847 | YES | NO | NA |
| Ankrd50 | ENSMUSG( | ENSMUSG( | -0.15047 | 0.171615 | NO | NO | NA |
| 2410002F: | ENSMUSG( | ENSMUSG( | 0.437645 | 8.14E-08 | YES | NO | NA |
| Narf | ENSMUSG( | ENSMUSG( | -0.24829 | 0.057144 | NO | NO | NA |
| Gopc | ENSMUSG( | ENSMUSG( | -0.24663 | 0.000894 | YES | NO | NA |
| Rcbtb2 | ENSMUSG( | ENSMUSG( | -0.11781 | 0.101829 | NO | NO | NA |
| Phf13 | ENSMUSG( | ENSMUSG( | -0.95469 | 6.18E-09 | YES | NO | NA |
| Acadvl | ENSMUSG( | ENSMUSG( | -0.04256 | 0.553112 | NO | NO | NA |
| Ppt1 | ENSMUSG( | ENSMUSG( | -0.11787 | 0.166586 | NO | NO | NA |
| Uspl1 | ENSMUSG( | ENSMUSG( | -0.36552 | 6.09E-06 | YES | NO | NA |
| E2f7 | ENSMUSG( | ENSMUSG( | 0.278382 | 5.78E-07 | YES | NO | NA |
| Ssna1 | ENSMUSG( | ENSMUSG( | 0.137413 | 0.037733 | YES | NO | NA |

| | | | | | | | |
|---|---|---|---|---|---|---|---|
| Prag1 | ENSMUSG( | ENSMUSG( | -0.98505 | 6.54E-06 | YES | NO | NA |
| Nol9 | ENSMUSG( | ENSMUSG( | 0.126531 | 0.031765 | YES | NO | NA |
| Rrn3 | ENSMUSG( | ENSMUSG( | 0.138856 | 0.034031 | YES | NO | NA |
| Smurf2 | ENSMUSG( | ENSMUSG( | -0.27221 | 0.000656 | YES | NO | NA |
| Bcl3 | ENSMUSG( | ENSMUSG( | -0.36303 | 8.79E-05 | YES | NO | NA |
| Cnot8 | ENSMUSG( | ENSMUSG( | -0.21786 | 0.000125 | YES | NO | NA |
| Pfdn2 | ENSMUSG( | ENSMUSG( | -0.11561 | 0.624821 | NO | NO | NA |
| Ugp2 | ENSMUSG( | ENSMUSG( | 0.017297 | 0.872834 | NO | NO | NA |
| Hook1 | ENSMUSG( | ENSMUSG( | 0.348925 | 0.004231 | YES | NO | NA |
| Fbxo28 | ENSMUSG( | ENSMUSG( | -0.37475 | 1.92E-15 | YES | NO | NA |
| Styx | ENSMUSG( | ENSMUSG( | -0.04174 | 0.722873 | NO | NO | NA |
| Atp6v1h | ENSMUSG( | ENSMUSG( | -0.08021 | 0.199081 | NO | NO | NA |
| Dcps | ENSMUSG( | ENSMUSG( | -0.05291 | 0.499614 | NO | NO | NA |
| Ass1 | ENSMUSG( | ENSMUSG( | 0.248055 | 0.118976 | NO | NO | NA |
| Tlnrd1 | ENSMUSG( | ENSMUSG( | -0.18078 | 0.000812 | YES | NO | NA |
| Rap2b | ENSMUSG( | ENSMUSG( | -0.30197 | 2.95E-05 | YES | NO | NA |
| Ern1 | ENSMUSG( | ENSMUSG( | -0.39095 | 4.43E-08 | YES | NO | NA |
| Katna1 | ENSMUSG( | ENSMUSG( | 0.003974 | 0.959325 | NO | NO | NA |
| Mvb12a | ENSMUSG( | ENSMUSG( | -0.21198 | 3.11E-05 | YES | NO | NA |
| Samd4b | ENSMUSG( | ENSMUSG( | -0.18131 | 0.049281 | YES | NO | NA |
| Rel | ENSMUSG( | ENSMUSG( | 0.064672 | 0.594085 | NO | NO | NA |
| Cep83 | ENSMUSG( | ENSMUSG( | 0.418867 | 0.000289 | YES | NO | NA |
| Ggt1 | ENSMUSG( | ENSMUSG( | -0.38122 | 0.025483 | YES | NO | NA |
| Herpud1 | ENSMUSG( | ENSMUSG( | -0.03413 | 0.702923 | NO | NO | NA |
| Cry1 | ENSMUSG( | ENSMUSG( | -0.19344 | 0.091258 | NO | NO | NA |
| Ubl7 | ENSMUSG( | ENSMUSG( | -0.04579 | 0.609667 | NO | NO | NA |
| Ndufa3 | ENSMUSG( | ENSMUSG( | -0.02813 | 0.772033 | NO | NO | NA |
| Cnih1 | ENSMUSG( | ENSMUSG( | 0.197394 | 0.025189 | YES | NO | NA |
| Ccdc25 | ENSMUSG( | ENSMUSG( | 0.05349 | 0.503792 | NO | NO | NA |
| Yipf1 | ENSMUSG( | ENSMUSG( | -0.05559 | 0.482543 | NO | NO | NA |
| Rapgef2 | ENSMUSG( | ENSMUSG( | -0.58545 | 2.95E-09 | YES | NO | NA |
| Sar1b | ENSMUSG( | ENSMUSG( | 0.074908 | 0.291658 | NO | NO | NA |
| St3gal6 | ENSMUSG( | ENSMUSG( | -0.43499 | 6.13E-06 | YES | NO | NA |
| Efcab14 | ENSMUSG( | ENSMUSG( | -0.28628 | 1.09E-05 | YES | NO | NA |
| Gfi1 | ENSMUSG( | ENSMUSG( | -0.20391 | 0.00874 | YES | NO | NA |
| Jade1 | ENSMUSG( | ENSMUSG( | -0.05408 | 0.565868 | NO | NO | NA |
| Lgals8 | ENSMUSG( | ENSMUSG( | -0.29796 | 0.003991 | YES | NO | NA |
| Dhfr | ENSMUSG( | ENSMUSG( | 0.341216 | 1.88E-06 | YES | NO | NA |
| Mettl14 | ENSMUSG( | ENSMUSG( | 0.125777 | 0.187394 | NO | NO | NA |
| Pif1 | ENSMUSG( | ENSMUSG( | 0.141727 | 0.08304 | NO | NO | NA |
| Agap3 | ENSMUSG( | ENSMUSG( | 0.010881 | 0.882272 | NO | NO | NA |
| Ttc5 | ENSMUSG( | ENSMUSG( | -0.23686 | 0.000446 | YES | NO | NA |
| Hgsnat | ENSMUSG( | ENSMUSG( | 0.170658 | 0.160976 | NO | NO | NA |
| Zfyve26 | ENSMUSG( | ENSMUSG( | -0.35867 | 4.28E-07 | YES | NO | NA |
| Klhdc3 | ENSMUSG( | ENSMUSG( | -0.0356 | 0.591676 | NO | NO | NA |
| Ints8 | ENSMUSG( | ENSMUSG( | -0.30008 | 1.00E-06 | YES | NO | NA |
| Mad1l1 | ENSMUSG( | ENSMUSG( | 0.113098 | 0.279176 | NO | NO | NA |

| | | | | | | | |
|---|---|---|---|---|---|---|---|
| Pnpla6 | ENSMUSG( | ENSMUSG( | -0.19303 | 0.016898 | YES | NO | NA |
| Cul4b | ENSMUSG( | ENSMUSG( | 0.03434 | 0.727413 | NO | NO | NA |
| Rnf138 | ENSMUSG( | ENSMUSG( | -0.08618 | 0.426095 | NO | NO | NA |
| Ccdc59 | ENSMUSG( | ENSMUSG( | -0.07489 | 0.584983 | NO | NO | NA |
| Proser1 | ENSMUSG( | ENSMUSG( | -0.1621 | 0.233115 | NO | NO | NA |
| Polr2c | ENSMUSG( | ENSMUSG( | 0.209165 | 0.000248 | YES | NO | NA |
| Zrsr2 | ENSMUSG( | ENSMUSG( | -0.14727 | 0.180503 | NO | NO | NA |
| Maml1 | ENSMUSG( | ENSMUSG( | -0.39871 | 0.000301 | YES | NO | NA |
| Prkaa1 | ENSMUSG( | ENSMUSG( | -0.22879 | 0.000205 | YES | NO | NA |
| Ptpn9 | ENSMUSG( | ENSMUSG( | 0.119422 | 0.173764 | NO | NO | NA |
| Cyria | ENSMUSG( | ENSMUSG( | -0.64599 | 9.05E-09 | YES | NO | NA |
| Msl3 | ENSMUSG( | ENSMUSG( | 0.085965 | 0.48942 | NO | NO | NA |
| Tep1 | ENSMUSG( | ENSMUSG( | -0.40936 | 6.89E-07 | YES | NO | NA |
| Hadh | ENSMUSG( | ENSMUSG( | 0.234045 | 0.035433 | YES | NO | NA |
| Laptm4a | ENSMUSG( | ENSMUSG( | -0.27759 | 1.74E-05 | YES | NO | NA |
| Pea15a | ENSMUSG( | ENSMUSG( | -0.55497 | 9.80E-15 | YES | NO | NA |
| Prkab1 | ENSMUSG( | ENSMUSG( | -0.09675 | 0.201631 | NO | NO | NA |
| Dhx29 | ENSMUSG( | ENSMUSG( | 0.092215 | 0.408317 | NO | NO | NA |
| Sgpl1 | ENSMUSG( | ENSMUSG( | -0.31175 | 2.80E-12 | YES | NO | NA |
| Armc1 | ENSMUSG( | ENSMUSG( | -0.10216 | 0.153826 | NO | NO | NA |
| Idnk | ENSMUSG( | ENSMUSG( | -0.25048 | 0.000136 | YES | NO | NA |
| Tradd | ENSMUSG( | ENSMUSG( | -0.17867 | 0.00444 | YES | NO | NA |
| Zfp664 | ENSMUSG( | ENSMUSG( | -0.20992 | 0.017056 | YES | NO | NA |
| Ndufs7 | ENSMUSG( | ENSMUSG( | 0.099766 | 0.149037 | NO | NO | NA |
| BC031181 | ENSMUSG( | ENSMUSG( | -0.22431 | 0.003453 | YES | NO | NA |
| Rnaseh2c | ENSMUSG( | ENSMUSG( | -0.08493 | 0.299432 | NO | NO | NA |
| Trappc6b | ENSMUSG( | ENSMUSG( | -0.25394 | 0.014284 | YES | NO | NA |
| Usp42 | ENSMUSG( | ENSMUSG( | -0.22672 | 5.43E-06 | YES | NO | NA |
| Mief1 | ENSMUSG( | ENSMUSG( | -0.10697 | 0.05163 | NO | NO | NA |
| Atp11a | ENSMUSG( | ENSMUSG( | -0.48527 | 7.97E-07 | YES | NO | NA |
| Get3 | ENSMUSG( | ENSMUSG( | 0.036926 | 0.624845 | NO | NO | NA |
| Cobll1 | ENSMUSG( | ENSMUSG( | -0.22445 | 0.001321 | YES | NO | NA |
| Fcho2 | ENSMUSG( | ENSMUSG( | -0.58949 | 1.22E-06 | YES | NO | NA |
| Cables2 | ENSMUSG( | ENSMUSG( | -0.16714 | 0.022083 | YES | NO | NA |
| Vps33a | ENSMUSG( | ENSMUSG( | -0.01223 | 0.894638 | NO | NO | NA |
| Esco2 | ENSMUSG( | ENSMUSG( | 0.024979 | 0.83522 | NO | NO | NA |
| Nelfe | ENSMUSG( | ENSMUSG( | 0.185816 | 0.05711 | NO | NO | NA |
| Plpbp | ENSMUSG( | ENSMUSG( | -0.05486 | 0.579897 | NO | NO | NA |
| Sapcd2 | ENSMUSG( | ENSMUSG( | 0.423082 | 1.24E-06 | YES | NO | NA |
| Gpr108 | ENSMUSG( | ENSMUSG( | -0.19738 | 0.004889 | YES | NO | NA |
| Mideas | ENSMUSG( | ENSMUSG( | -0.53269 | 9.02E-09 | YES | NO | NA |
| Sap30bp | ENSMUSG( | ENSMUSG( | 0.167213 | 0.018532 | YES | NO | NA |
| Neil3 | ENSMUSG( | ENSMUSG( | 0.162593 | 0.041332 | YES | NO | NA |
| Orc3 | ENSMUSG( | ENSMUSG( | 0.093177 | 0.247496 | NO | NO | NA |
| Tmem59 | ENSMUSG( | ENSMUSG( | -0.42143 | 1.08E-09 | YES | NO | NA |
| Fbxo3 | ENSMUSG( | ENSMUSG( | -0.12578 | 0.058623 | NO | NO | NA |
| Bbx | ENSMUSG( | ENSMUSG( | -0.54738 | 1.42E-08 | YES | NO | NA |

| | | | | | | | |
|---|---|---|---|---|---|---|---|
| Ppdpf | ENSMUSG( | ENSMUSG( | -0.25843 | 0.000113 | YES | NO | NA |
| Slc35b1 | ENSMUSG( | ENSMUSG( | 0.145809 | 0.129265 | NO | NO | NA |
| Chn2 | ENSMUSG( | ENSMUSG( | -0.97904 | 1.55E-11 | YES | NO | NA |
| Ikbkg | ENSMUSG( | ENSMUSG( | -0.14735 | 0.2254 | NO | NO | NA |
| Ebp | ENSMUSG( | ENSMUSG( | -0.03357 | 0.615925 | NO | NO | NA |
| Zbtb4 | ENSMUSG( | ENSMUSG( | -0.82096 | 1.79E-09 | YES | NO | NA |
| Pdk1 | ENSMUSG( | ENSMUSG( | 0.622147 | 0.000291 | YES | NO | NA |
| Nrros | ENSMUSG( | ENSMUSG( | -0.11854 | 0.141155 | NO | NO | NA |
| Rpf1 | ENSMUSG( | ENSMUSG( | 0.047439 | 0.729736 | NO | NO | NA |
| Acox3 | ENSMUSG( | ENSMUSG( | -0.33474 | 0.000246 | YES | NO | NA |
| Vps72 | ENSMUSG( | ENSMUSG( | -0.07678 | 0.289283 | NO | NO | NA |
| Nfkbib | ENSMUSG( | ENSMUSG( | 0.039917 | 0.664536 | NO | NO | NA |
| Spock2 | ENSMUSG( | ENSMUSG( | 0.507827 | 5.86E-05 | YES | NO | NA |
| Mphosph8 | ENSMUSG( | ENSMUSG( | -0.21993 | 0.087536 | NO | NO | NA |
| Srp19 | ENSMUSG( | ENSMUSG( | 0.044345 | 0.60191 | NO | NO | NA |
| Trim41 | ENSMUSG( | ENSMUSG( | -0.14949 | 0.084388 | NO | NO | NA |
| Hnrnph3 | ENSMUSG( | ENSMUSG( | -0.22024 | 0.03906 | YES | NO | NA |
| Nck1 | ENSMUSG( | ENSMUSG( | -0.5523 | 8.79E-07 | YES | NO | NA |
| Snx18 | ENSMUSG( | ENSMUSG( | -0.55779 | 1.38E-10 | YES | NO | NA |
| Pycr2 | ENSMUSG( | ENSMUSG( | 0.321217 | 1.23E-05 | YES | NO | NA |
| Mfsd6 | ENSMUSG( | ENSMUSG( | -0.10172 | 0.364915 | NO | NO | NA |
| Kat14 | ENSMUSG( | ENSMUSG( | 0.107691 | 0.167728 | NO | NO | NA |
| Pals2 | ENSMUSG( | ENSMUSG( | 0.560339 | 1.63E-08 | YES | NO | NA |
| Impa2 | ENSMUSG( | ENSMUSG( | 0.390991 | 1.03E-05 | YES | NO | NA |
| Tbc1d22a | ENSMUSG( | ENSMUSG( | 0.098526 | 0.35234 | NO | NO | NA |
| 4930503L1 | ENSMUSG( | ENSMUSG( | -0.169 | 0.045607 | YES | NO | NA |
| Nudt5 | ENSMUSG( | ENSMUSG( | 0.173649 | 0.05543 | NO | NO | NA |
| Mtap | ENSMUSG( | ENSMUSG( | 0.410327 | 2.98E-05 | YES | NO | NA |
| Pnkp | ENSMUSG( | ENSMUSG( | -0.16035 | 0.020009 | YES | NO | NA |
| Dna2 | ENSMUSG( | ENSMUSG( | 0.366743 | 7.26E-08 | YES | NO | NA |
| Zeb2 | ENSMUSG( | ENSMUSG( | -1.05674 | 2.14E-13 | YES | YES | Downregulated |
| Psenen | ENSMUSG( | ENSMUSG( | -0.1206 | 0.071904 | NO | NO | NA |
| Rere | ENSMUSG( | ENSMUSG( | -0.63462 | 5.50E-07 | YES | NO | NA |
| Rabep2 | ENSMUSG( | ENSMUSG( | -0.32412 | 7.46E-05 | YES | NO | NA |
| Tars2 | ENSMUSG( | ENSMUSG( | 0.119535 | 0.157446 | NO | NO | NA |
| Dock9 | ENSMUSG( | ENSMUSG( | 0.112758 | 0.188844 | NO | NO | NA |
| Rpe | ENSMUSG( | ENSMUSG( | -0.25948 | 2.00E-05 | YES | NO | NA |
| Slc25a37 | ENSMUSG( | ENSMUSG( | 0.129179 | 0.214439 | NO | NO | NA |
| Etv6 | ENSMUSG( | ENSMUSG( | 0.069955 | 0.383295 | NO | NO | NA |
| Ank | ENSMUSG( | ENSMUSG( | -0.50241 | 0.000146 | YES | NO | NA |
| Hsd17b4 | ENSMUSG( | ENSMUSG( | -0.0407 | 0.737669 | NO | NO | NA |
| Rbm12 | ENSMUSG( | ENSMUSG( | -0.2019 | 0.004723 | YES | NO | NA |
| Pigo | ENSMUSG( | ENSMUSG( | -0.00082 | 0.993026 | NO | NO | NA |
| Ndufb5 | ENSMUSG( | ENSMUSG( | 0.118213 | 0.15151 | NO | NO | NA |
| Lpcat1 | ENSMUSG( | ENSMUSG( | 0.165845 | 0.053951 | NO | NO | NA |
| Pdpr | ENSMUSG( | ENSMUSG( | -0.01727 | 0.897157 | NO | NO | NA |
| Garre1 | ENSMUSG( | ENSMUSG( | -0.34151 | 0.000207 | YES | NO | NA |

| | | | | | | | |
|---|---|---|---|---|---|---|---|
| Nab2 | ENSMUSG( | ENSMUSG( | -0.18873 | 0.233451 | NO | NO | NA |
| Pias1 | ENSMUSG( | ENSMUSG( | -0.38098 | 6.36E-07 | YES | NO | NA |
| Desi2 | ENSMUSG( | ENSMUSG( | 0.205848 | 0.002355 | YES | NO | NA |
| Crtc3 | ENSMUSG( | ENSMUSG( | -0.02902 | 0.827284 | NO | NO | NA |
| Ptpn23 | ENSMUSG( | ENSMUSG( | -0.2826 | 5.84E-07 | YES | NO | NA |
| Atxn1l | ENSMUSG( | ENSMUSG( | -0.09011 | 0.305015 | NO | NO | NA |
| Sdhaf2 | ENSMUSG( | ENSMUSG( | -0.16924 | 0.016318 | YES | NO | NA |
| Krtcap2 | ENSMUSG( | ENSMUSG( | 0.046194 | 0.639533 | NO | NO | NA |
| Fam189b | ENSMUSG( | ENSMUSG( | -0.24263 | 0.049838 | YES | NO | NA |
| Pdcd5 | ENSMUSG( | ENSMUSG( | 0.204051 | 0.013132 | YES | NO | NA |
| Mbnl2 | ENSMUSG( | ENSMUSG( | -0.40674 | 0.000387 | YES | NO | NA |
| Cuta | ENSMUSG( | ENSMUSG( | -0.10317 | 0.037192 | YES | NO | NA |
| Uvrag | ENSMUSG( | ENSMUSG( | -0.16792 | 0.134954 | NO | NO | NA |
| Ndor1 ENS | ENSMUSG( | ENSMUSG( | 0.093218 | 0.360063 | NO | NO | NA |
| Birc3 | ENSMUSG( | ENSMUSG( | 0.046346 | 0.661917 | NO | NO | NA |
| Ints5 | ENSMUSG( | ENSMUSG( | -0.00799 | 0.914675 | NO | NO | NA |
| Shprh | ENSMUSG( | ENSMUSG( | -0.33476 | 6.05E-07 | YES | NO | NA |
| Gm10076 | ENSMUSG( | ENSMUSG( | 0.185338 | 0.33087 | NO | NO | NA |
| Grcc10 | ENSMUSG( | ENSMUSG( | -0.06932 | 0.417632 | NO | NO | NA |
| Prpsap1 | ENSMUSG( | ENSMUSG( | 0.10147 | 0.162531 | NO | NO | NA |
| Pidd1 | ENSMUSG( | ENSMUSG( | 0.171874 | 0.007843 | YES | NO | NA |
| Ntan1 | ENSMUSG( | ENSMUSG( | -0.08037 | 0.216335 | NO | NO | NA |
| Cops8 | ENSMUSG( | ENSMUSG( | 0.143509 | 0.106491 | NO | NO | NA |
| Arfip1 | ENSMUSG( | ENSMUSG( | -0.00088 | 0.993583 | NO | NO | NA |
| Asrgl1 | ENSMUSG( | ENSMUSG( | 0.29017 | 4.16E-05 | YES | NO | NA |
| Mrpl51 | ENSMUSG( | ENSMUSG( | 0.203322 | 0.01471 | YES | NO | NA |
| Gins4 | ENSMUSG( | ENSMUSG( | 0.164751 | 0.001323 | YES | NO | NA |
| Insig1 | ENSMUSG( | ENSMUSG( | 0.706451 | 3.33E-05 | YES | NO | NA |
| Nedd1 | ENSMUSG( | ENSMUSG( | -0.00308 | 0.973203 | NO | NO | NA |
| Apmap | ENSMUSG( | ENSMUSG( | 0.041696 | 0.593861 | NO | NO | NA |
| Nmt2 | ENSMUSG( | ENSMUSG( | -0.057 | 0.466448 | NO | NO | NA |
| Slc39a1 | ENSMUSG( | ENSMUSG( | 0.129225 | 0.161841 | NO | NO | NA |
| Swap70 | ENSMUSG( | ENSMUSG( | -0.77309 | 3.02E-20 | YES | NO | NA |
| Rmi1 | ENSMUSG( | ENSMUSG( | -0.26155 | 0.000785 | YES | NO | NA |
| Orc2 | ENSMUSG( | ENSMUSG( | 0.372146 | 5.57E-07 | YES | NO | NA |
| Ercc6 | ENSMUSG( | ENSMUSG( | -0.23306 | 0.000196 | YES | NO | NA |
| Ints10 | ENSMUSG( | ENSMUSG( | -0.02648 | 0.717607 | NO | NO | NA |
| Zpr1 | ENSMUSG( | ENSMUSG( | 0.212689 | 0.022448 | YES | NO | NA |
| Med17 | ENSMUSG( | ENSMUSG( | 0.039867 | 0.616241 | NO | NO | NA |
| Prrc1 | ENSMUSG( | ENSMUSG( | -0.05574 | 0.484233 | NO | NO | NA |
| Fam120b | ENSMUSG( | ENSMUSG( | -0.25807 | 0.003179 | YES | NO | NA |
| Exoc7 | ENSMUSG( | ENSMUSG( | -0.05358 | 0.63321 | NO | NO | NA |
| Abcc4 | ENSMUSG( | ENSMUSG( | -0.03116 | 0.80545 | NO | NO | NA |
| Tsc22d2 | ENSMUSG( | ENSMUSG( | -0.40168 | 1.19E-05 | YES | NO | NA |
| Cenpc1 | ENSMUSG( | ENSMUSG( | 0.018131 | 0.847349 | NO | NO | NA |
| Ifrd1 | ENSMUSG( | ENSMUSG( | 0.523105 | 0.000104 | YES | NO | NA |
| Tfg | ENSMUSG( | ENSMUSG( | 0.031261 | 0.763686 | NO | NO | NA |

| | | | | | | | |
|---|---|---|---|---|---|---|---|
| Map3k5 | ENSMUSG( | ENSMUSG( | -0.29986 | 0.000148 | YES | NO | NA |
| Hmox2 | ENSMUSG( | ENSMUSG( | -0.14015 | 0.219159 | NO | NO | NA |
| Trmt1 | ENSMUSG( | ENSMUSG( | 0.50951 | 5.48E-08 | YES | NO | NA |
| Etfdh | ENSMUSG( | ENSMUSG( | 0.123395 | 0.241243 | NO | NO | NA |
| Zfp512b | ENSMUSG( | ENSMUSG( | 0.389389 | 0.000552 | YES | NO | NA |
| Pbdc1 | ENSMUSG( | ENSMUSG( | 0.248809 | 0.056195 | NO | NO | NA |
| Camkk2 | ENSMUSG( | ENSMUSG( | 0.031352 | 0.644893 | NO | NO | NA |
| Utp15 | ENSMUSG( | ENSMUSG( | 0.272958 | 0.000508 | YES | NO | NA |
| Surf6 | ENSMUSG( | ENSMUSG( | 0.15022 | 0.068651 | NO | NO | NA |
| Atp5md | ENSMUSG( | ENSMUSG( | 0.132279 | 0.132657 | NO | NO | NA |
| Zfp62 | ENSMUSG( | ENSMUSG( | -0.36107 | 0.000243 | YES | NO | NA |
| Cul4a | ENSMUSG( | ENSMUSG( | -0.12472 | 0.048124 | YES | NO | NA |
| Cyhr1 | ENSMUSG( | ENSMUSG( | -0.09641 | 0.278397 | NO | NO | NA |
| Unc119 | ENSMUSG( | ENSMUSG( | 0.065344 | 0.480832 | NO | NO | NA |
| Cog8 | ENSMUSG( | ENSMUSG( | -0.2129 | 1.31E-05 | YES | NO | NA |
| Rad9a | ENSMUSG( | ENSMUSG( | -0.31808 | 1.84E-09 | YES | NO | NA |
| Orc6 | ENSMUSG( | ENSMUSG( | 0.289888 | 0.001848 | YES | NO | NA |
| Grpel1 | ENSMUSG( | ENSMUSG( | 0.250665 | 0.017332 | YES | NO | NA |
| Kctd2 | ENSMUSG( | ENSMUSG( | -0.0932 | 0.374463 | NO | NO | NA |
| Isoc1 | ENSMUSG( | ENSMUSG( | 0.292603 | 7.44E-09 | YES | NO | NA |
| Slc4a1 | ENSMUSG( | ENSMUSG( | 1.293851 | 0.073376 | NO | NO | NA |
| Tbl3 | ENSMUSG( | ENSMUSG( | 0.488056 | 2.50E-05 | YES | NO | NA |
| Gpatch2l | ENSMUSG( | ENSMUSG( | -0.21731 | 0.001468 | YES | NO | NA |
| Ctps | ENSMUSG( | ENSMUSG( | 0.48282 | 5.36E-06 | YES | NO | NA |
| Rchy1 | ENSMUSG( | ENSMUSG( | -0.15202 | 0.081214 | NO | NO | NA |
| Bcl7c | ENSMUSG( | ENSMUSG( | 0.411278 | 1.27E-07 | YES | NO | NA |
| Pias4 | ENSMUSG( | ENSMUSG( | -0.12864 | 0.067478 | NO | NO | NA |
| Rnf7 | ENSMUSG( | ENSMUSG( | 0.227051 | 0.002623 | YES | NO | NA |
| Nampt | ENSMUSG( | ENSMUSG( | -0.04633 | 0.611454 | NO | NO | NA |
| Reep3 | ENSMUSG( | ENSMUSG( | -0.04832 | 0.640929 | NO | NO | NA |
| Carnmt1 | ENSMUSG( | ENSMUSG( | 0.323721 | 0.000125 | YES | NO | NA |
| Mrpl3 | ENSMUSG( | ENSMUSG( | 0.405342 | 7.51E-08 | YES | NO | NA |
| Ifnar2 | ENSMUSG( | ENSMUSG( | -0.95136 | 2.84E-11 | YES | NO | NA |
| Dele1 | ENSMUSG( | ENSMUSG( | -0.02101 | 0.776671 | NO | NO | NA |
| Cmpk2 | ENSMUSG( | ENSMUSG( | -0.50032 | 4.35E-12 | YES | NO | NA |
| Nol11 | ENSMUSG( | ENSMUSG( | 0.40196 | 2.82E-07 | YES | NO | NA |
| Ldlrap1 | ENSMUSG( | ENSMUSG( | -0.502 | 0.000373 | YES | NO | NA |
| Pigk | ENSMUSG( | ENSMUSG( | 0.075693 | 0.397319 | NO | NO | NA |
| Dlat | ENSMUSG( | ENSMUSG( | 0.259315 | 0.00106 | YES | NO | NA |
| Atg12 | ENSMUSG( | ENSMUSG( | -0.3429 | 8.05E-05 | YES | NO | NA |
| Slc35b4 | ENSMUSG( | ENSMUSG( | -0.16115 | 0.045044 | YES | NO | NA |
| Utp4 | ENSMUSG( | ENSMUSG( | 0.284745 | 0.011886 | YES | NO | NA |
| Rtca | ENSMUSG( | ENSMUSG( | 0.452596 | 9.51E-05 | YES | NO | NA |
| Plrg1 | ENSMUSG( | ENSMUSG( | 0.227758 | 0.001153 | YES | NO | NA |
| Gcc2 | ENSMUSG( | ENSMUSG( | -0.07069 | 0.431957 | NO | NO | NA |
| Yeats2 | ENSMUSG( | ENSMUSG( | -0.13794 | 0.029044 | YES | NO | NA |
| Kdelr1 | ENSMUSG( | ENSMUSG( | -0.16394 | 0.014119 | YES | NO | NA |

| | | | | | | | |
|---|---|---|---|---|---|---|---|
| Dhrs7 | ENSMUSG( | ENSMUSG( | -0.20883 | 0.018521 | YES | NO | NA |
| Brix1 | ENSMUSG( | ENSMUSG( | 0.373759 | 0.000571 | YES | NO | NA |
| Fanci | ENSMUSG( | ENSMUSG( | 0.199991 | 0.012259 | YES | NO | NA |
| Dctn6 | ENSMUSG( | ENSMUSG( | -0.10523 | 0.136506 | NO | NO | NA |
| Pip5k1a | ENSMUSG( | ENSMUSG( | -0.23027 | 0.004304 | YES | NO | NA |
| Zfp410 | ENSMUSG( | ENSMUSG( | -0.00471 | 0.954162 | NO | NO | NA |
| Gins2 | ENSMUSG( | ENSMUSG( | 0.021654 | 0.787164 | NO | NO | NA |
| St3gal4 | ENSMUSG( | ENSMUSG( | -0.26111 | 5.46E-08 | YES | NO | NA |
| Atl2 | ENSMUSG( | ENSMUSG( | -0.02165 | 0.775742 | NO | NO | NA |
| Cul5 | ENSMUSG( | ENSMUSG( | 0.040042 | 0.6204 | NO | NO | NA |
| Mboat7 | ENSMUSG( | ENSMUSG( | -0.02197 | 0.768026 | NO | NO | NA |
| Eif2d | ENSMUSG( | ENSMUSG( | -0.02325 | 0.741426 | NO | NO | NA |
| Xpr1 | ENSMUSG( | ENSMUSG( | -0.00115 | 0.991623 | NO | NO | NA |
| Tomm7 | ENSMUSG( | ENSMUSG( | 0.032815 | 0.753653 | NO | NO | NA |
| Klhl9 | ENSMUSG( | ENSMUSG( | -0.20592 | 0.0011 | YES | NO | NA |
| Acaa2 | ENSMUSG( | ENSMUSG( | 0.372112 | 9.74E-05 | YES | NO | NA |
| Fam169b | ENSMUSG( | ENSMUSG( | -0.15482 | 0.07668 | NO | NO | NA |
| Mrto4 | ENSMUSG( | ENSMUSG( | 0.640109 | 1.50E-06 | YES | NO | NA |
| Fut8 | ENSMUSG( | ENSMUSG( | -0.30626 | 5.28E-06 | YES | NO | NA |
| Ttc3 | ENSMUSG( | ENSMUSG( | -0.24896 | 0.059602 | NO | NO | NA |
| Ppme1 | ENSMUSG( | ENSMUSG( | -0.23236 | 5.58E-05 | YES | NO | NA |
| Lamtor3 | ENSMUSG( | ENSMUSG( | -0.02984 | 0.600449 | NO | NO | NA |
| Rfk | ENSMUSG( | ENSMUSG( | -0.02888 | 0.633271 | NO | NO | NA |
| Pim2 | ENSMUSG( | ENSMUSG( | -0.31751 | 0.000954 | YES | NO | NA |
| Pphln1 | ENSMUSG( | ENSMUSG( | -0.04129 | 0.650518 | NO | NO | NA |
| Ulk2 | ENSMUSG( | ENSMUSG( | -0.39834 | 4.70E-06 | YES | NO | NA |
| Usp18 | ENSMUSG( | ENSMUSG( | -1.07171 | 2.24E-10 | YES | YES | Downregulated |
| Rptor | ENSMUSG( | ENSMUSG( | -0.34769 | 3.89E-05 | YES | NO | NA |
| Gdf11 | ENSMUSG( | ENSMUSG( | -1.00109 | 3.31E-07 | YES | YES | Downregulated |
| Utp14a | ENSMUSG( | ENSMUSG( | 0.359505 | 0.000544 | YES | NO | NA |
| Fbh1 | ENSMUSG( | ENSMUSG( | -0.27132 | 5.13E-05 | YES | NO | NA |
| Msh3 | ENSMUSG( | ENSMUSG( | -0.01982 | 0.856543 | NO | NO | NA |
| Memo1 | ENSMUSG( | ENSMUSG( | 0.226018 | 0.003289 | YES | NO | NA |
| Dnajb12 | ENSMUSG( | ENSMUSG( | 0.114953 | 0.270824 | NO | NO | NA |
| Oser1 | ENSMUSG( | ENSMUSG( | -0.11385 | 0.323191 | NO | NO | NA |
| Crebrf | ENSMUSG( | ENSMUSG( | -0.72945 | 2.97E-07 | YES | NO | NA |
| Scly | ENSMUSG( | ENSMUSG( | -0.46306 | 1.68E-08 | YES | NO | NA |
| Ccdc186 | ENSMUSG( | ENSMUSG( | -0.38208 | 1.96E-06 | YES | NO | NA |
| Aagab | ENSMUSG( | ENSMUSG( | -0.13088 | 0.040274 | YES | NO | NA |
| Fnta | ENSMUSG( | ENSMUSG( | 0.095194 | 0.45373 | NO | NO | NA |
| Xndc1 | ENSMUSG( | ENSMUSG( | -0.15965 | 0.012433 | YES | NO | NA |
| B4galt7 | ENSMUSG( | ENSMUSG( | -0.25019 | 7.75E-05 | YES | NO | NA |
| Dclk2 | ENSMUSG( | ENSMUSG( | -0.26008 | 0.008425 | YES | NO | NA |
| Rasa2 | ENSMUSG( | ENSMUSG( | -0.08542 | 0.121286 | NO | NO | NA |
| Commd7 | ENSMUSG( | ENSMUSG( | -0.13076 | 0.089784 | NO | NO | NA |
| Capn15 | ENSMUSG( | ENSMUSG( | -0.27874 | 5.19E-06 | YES | NO | NA |
| Togaram1 | ENSMUSG( | ENSMUSG( | -0.0555 | 0.521137 | NO | NO | NA |

| | | | | | | | |
|---|---|---|---|---|---|---|---|
| Taf12 | ENSMUSG( | ENSMUSG( | -0.12649 | 0.181278 | NO | NO | NA |
| Suclg1 | ENSMUSG( | ENSMUSG( | 0.288152 | 1.37E-05 | YES | NO | NA |
| Ifit3 | ENSMUSG( | ENSMUSG( | -1.19462 | 1.66E-10 | YES | YES | Downregulated |
| Zfyve27 | ENSMUSG( | ENSMUSG( | -0.21702 | 0.004916 | YES | NO | NA |
| Hdac5 | ENSMUSG( | ENSMUSG( | -0.40987 | 2.53E-11 | YES | NO | NA |
| Snhg8 | ENSMUSG( | ENSMUSG( | 0.346408 | 0.020803 | YES | NO | NA |
| Gabpb1 | ENSMUSG( | ENSMUSG( | 0.001599 | 0.984402 | NO | NO | NA |
| 2410006H | ENSMUSG( | ENSMUSG( | -0.03744 | 0.784462 | NO | NO | NA |
| Naa16 | ENSMUSG( | ENSMUSG( | 0.110949 | 0.222429 | NO | NO | NA |
| Abtb1 | ENSMUSG( | ENSMUSG( | 0.017441 | 0.911696 | NO | NO | NA |
| Guf1 | ENSMUSG( | ENSMUSG( | 0.331286 | 0.007753 | YES | NO | NA |
| Kdm4c | ENSMUSG( | ENSMUSG( | -0.36092 | 2.23E-07 | YES | NO | NA |
| Cetn2 | ENSMUSG( | ENSMUSG( | -0.16921 | 0.087146 | NO | NO | NA |
| Slc25a36 | ENSMUSG( | ENSMUSG( | -0.36998 | 0.000987 | YES | NO | NA |
| Traf3ip2 | ENSMUSG( | ENSMUSG( | -0.34258 | 7.80E-09 | YES | NO | NA |
| Gm4204 | ENSMUSG( | ENSMUSG( | 0.167893 | 0.249092 | NO | NO | NA |
| Ttc17 | ENSMUSG( | ENSMUSG( | -0.34944 | 0.000363 | YES | NO | NA |
| Tmco1 | ENSMUSG( | ENSMUSG( | 0.027316 | 0.771854 | NO | NO | NA |
| Chpf | ENSMUSG( | ENSMUSG( | 0.004349 | 0.958772 | NO | NO | NA |
| Syap1 | ENSMUSG( | ENSMUSG( | -0.2842 | 0.026929 | YES | NO | NA |
| Bcl10 | ENSMUSG( | ENSMUSG( | -0.03997 | 0.50675 | NO | NO | NA |
| Zbtb17 | ENSMUSG( | ENSMUSG( | 0.052663 | 0.451496 | NO | NO | NA |
| Ndel1 | ENSMUSG( | ENSMUSG( | -0.10107 | 0.262914 | NO | NO | NA |
| Mdp1 | ENSMUSG( | ENSMUSG( | -0.16706 | 0.005715 | YES | NO | NA |
| Phf19 | ENSMUSG( | ENSMUSG( | 0.346685 | 2.17E-06 | YES | NO | NA |
| Med28 | ENSMUSG( | ENSMUSG( | -0.4097 | 1.77E-11 | YES | NO | NA |
| Itgb1bp1 | ENSMUSG( | ENSMUSG( | -0.04381 | 0.631052 | NO | NO | NA |
| Pole3 | ENSMUSG( | ENSMUSG( | 0.505752 | 1.48E-05 | YES | NO | NA |
| Mrps30 | ENSMUSG( | ENSMUSG( | 0.284859 | 0.001237 | YES | NO | NA |
| Smpd1 | ENSMUSG( | ENSMUSG( | -0.06974 | 0.546992 | NO | NO | NA |
| Ttf1 | ENSMUSG( | ENSMUSG( | -0.20353 | 0.002495 | YES | NO | NA |
| Rabep1 | ENSMUSG( | ENSMUSG( | -0.13432 | 0.047595 | YES | NO | NA |
| Atp6v0d1 | ENSMUSG( | ENSMUSG( | -0.24677 | 0.001927 | YES | NO | NA |
| Slc25a17 | ENSMUSG( | ENSMUSG( | 0.29876 | 7.05E-05 | YES | NO | NA |
| Pou2f1 | ENSMUSG( | ENSMUSG( | -0.1293 | 0.109241 | NO | NO | NA |
| Hps3 | ENSMUSG( | ENSMUSG( | -0.16464 | 0.030763 | YES | NO | NA |
| Copg2 | ENSMUSG( | ENSMUSG( | 0.00581 | 0.949082 | NO | NO | NA |
| Ubxn2a | ENSMUSG( | ENSMUSG( | -0.04748 | 0.63545 | NO | NO | NA |
| Pdgfb | ENSMUSG( | ENSMUSG( | -1.14741 | 5.46E-08 | YES | YES | Downregulated |
| Mrpl33 | ENSMUSG( | ENSMUSG( | 0.111705 | 0.071689 | NO | NO | NA |
| Kcna3 | ENSMUSG( | ENSMUSG( | -0.40039 | 6.48E-05 | YES | NO | NA |
| Ints6l | ENSMUSG( | ENSMUSG( | -0.12432 | 0.364408 | NO | NO | NA |
| Trmt1l | ENSMUSG( | ENSMUSG( | -0.45369 | 1.76E-08 | YES | NO | NA |
| Fcf1 | ENSMUSG( | ENSMUSG( | 0.280484 | 0.154699 | NO | NO | NA |
| Ufm1 | ENSMUSG( | ENSMUSG( | 0.073246 | 0.499294 | NO | NO | NA |
| Zfp330 | ENSMUSG( | ENSMUSG( | 0.197075 | 0.037898 | YES | NO | NA |
| Slc26a2 | ENSMUSG( | ENSMUSG( | -0.22837 | 0.005207 | YES | NO | NA |

| | | | | | | | |
|---|---|---|---|---|---|---|---|
| Hmg20a | ENSMUSG( | ENSMUSG( | -0.45508 | 2.35E-09 | YES | NO | NA |
| Gid8 | ENSMUSG( | ENSMUSG( | -0.13467 | 0.050511 | NO | NO | NA |
| Mrpl42 | ENSMUSG( | ENSMUSG( | 0.18456 | 0.038024 | YES | NO | NA |
| Zfp263 | ENSMUSG( | ENSMUSG( | -0.11497 | 0.175017 | NO | NO | NA |
| Taf8 | ENSMUSG( | ENSMUSG( | -0.21362 | 0.016677 | YES | NO | NA |
| Scamp4 | ENSMUSG( | ENSMUSG( | -0.09317 | 0.288847 | NO | NO | NA |
| Mindy3 | ENSMUSG( | ENSMUSG( | -0.05281 | 0.393625 | NO | NO | NA |
| Prps1l3 | ENSMUSG( | ENSMUSG( | 0.020191 | 0.760732 | NO | NO | NA |
| Ncbp2 | ENSMUSG( | ENSMUSG( | 0.077328 | 0.354826 | NO | NO | NA |
| Tpp1 | ENSMUSG( | ENSMUSG( | -0.35839 | 0.001087 | YES | NO | NA |
| Pgs1 | ENSMUSG( | ENSMUSG( | -0.20599 | 0.001611 | YES | NO | NA |
| Pprc1 | ENSMUSG( | ENSMUSG( | 0.210343 | 0.044079 | YES | NO | NA |
| Hmgn2-ps | ENSMUSG( | ENSMUSG( | 0.276203 | 0.047595 | YES | NO | NA |
| Tacc2 | ENSMUSG( | ENSMUSG( | -0.18959 | 0.271956 | NO | NO | NA |
| Sgo1 | ENSMUSG( | ENSMUSG( | 0.18375 | 0.168709 | NO | NO | NA |
| Zfp512 | ENSMUSG( | ENSMUSG( | 0.082599 | 0.393625 | NO | NO | NA |
| Pdk3 | ENSMUSG( | ENSMUSG( | 0.114944 | 0.137221 | NO | NO | NA |
| Ints9 | ENSMUSG( | ENSMUSG( | 0.156754 | 0.009519 | YES | NO | NA |
| Trgc2 | ENSMUSG( | ENSMUSG( | -0.30285 | 0.402496 | NO | NO | NA |
| Med22 | ENSMUSG( | ENSMUSG( | 0.187694 | 0.067288 | NO | NO | NA |
| Zcchc17 | ENSMUSG( | ENSMUSG( | 0.02406 | 0.80634 | NO | NO | NA |
| Naa30 | ENSMUSG( | ENSMUSG( | -0.30421 | 0.000706 | YES | NO | NA |
| Atg4b | ENSMUSG( | ENSMUSG( | -0.10757 | 0.110176 | NO | NO | NA |
| Txndc17 | ENSMUSG( | ENSMUSG( | 0.042284 | 0.696353 | NO | NO | NA |
| Nop14 | ENSMUSG( | ENSMUSG( | 0.316011 | 0.001752 | YES | NO | NA |
| Bckdk | ENSMUSG( | ENSMUSG( | 0.296172 | 0.001845 | YES | NO | NA |
| Ndufa11 | ENSMUSG( | ENSMUSG( | 0.292589 | 3.39E-05 | YES | NO | NA |
| Cuedc2 | ENSMUSG( | ENSMUSG( | 0.550293 | 1.68E-06 | YES | NO | NA |
| Ccdc93 | ENSMUSG( | ENSMUSG( | -0.45961 | 7.38E-10 | YES | NO | NA |
| Trip4 | ENSMUSG( | ENSMUSG( | -0.60566 | 1.47E-10 | YES | NO | NA |
| Atg7 | ENSMUSG( | ENSMUSG( | 0.153805 | 0.00728 | YES | NO | NA |
| Mtfr1l | ENSMUSG( | ENSMUSG( | 0.027661 | 0.847167 | NO | NO | NA |
| Emc8 | ENSMUSG( | ENSMUSG( | 0.194075 | 0.014801 | YES | NO | NA |
| Dnajc21 | ENSMUSG( | ENSMUSG( | -0.07293 | 0.579976 | NO | NO | NA |
| Nipa2 | ENSMUSG( | ENSMUSG( | 0.003778 | 0.967434 | NO | NO | NA |
| Rap2a | ENSMUSG( | ENSMUSG( | -0.26827 | 0.029648 | YES | NO | NA |
| Hsbp1 | ENSMUSG( | ENSMUSG( | -0.21662 | 0.004631 | YES | NO | NA |
| Tmem163 | ENSMUSG( | ENSMUSG( | -0.02935 | 0.782858 | NO | NO | NA |
| Rad18 | ENSMUSG( | ENSMUSG( | 0.317288 | 1.26E-05 | YES | NO | NA |
| Stx6 | ENSMUSG( | ENSMUSG( | -0.06914 | 0.283331 | NO | NO | NA |
| Mmadhc | ENSMUSG( | ENSMUSG( | 0.018359 | 0.835507 | NO | NO | NA |
| Plekhg3 | ENSMUSG( | ENSMUSG( | -0.10642 | 0.446431 | NO | NO | NA |
| Man2c1 | ENSMUSG( | ENSMUSG( | 0.343467 | 6.23E-07 | YES | NO | NA |
| Lemd3 | ENSMUSG( | ENSMUSG( | -0.14226 | 0.105893 | NO | NO | NA |
| Fancm | ENSMUSG( | ENSMUSG( | 0.115562 | 0.077981 | NO | NO | NA |
| Sigirr | ENSMUSG( | ENSMUSG( | -0.01951 | 0.842522 | NO | NO | NA |
| Map3k12 | ENSMUSG( | ENSMUSG( | 0.091484 | 0.299432 | NO | NO | NA |

| | | | | | | | |
|---|---|---|---|---|---|---|---|
| Eci2 | ENSMUSG( | ENSMUSG( | 0.409957 | 0.000113 | YES | NO | NA |
| Atp6ap2 | ENSMUSG( | ENSMUSG( | 0.011653 | 0.897686 | NO | NO | NA |
| Rraga | ENSMUSG( | ENSMUSG( | -0.01106 | 0.899457 | NO | NO | NA |
| Txndc11 | ENSMUSG( | ENSMUSG( | -0.10726 | 0.125709 | NO | NO | NA |
| Micos13 | ENSMUSG( | ENSMUSG( | 0.093513 | 0.375343 | NO | NO | NA |
| Fam98a | ENSMUSG( | ENSMUSG( | 0.125805 | 0.181278 | NO | NO | NA |
| Shfl | ENSMUSG( | ENSMUSG( | -0.35556 | 4.10E-05 | YES | NO | NA |
| Fam76a | ENSMUSG( | ENSMUSG( | 0.112692 | 0.353933 | NO | NO | NA |
| Sfmbt1 | ENSMUSG( | ENSMUSG( | 0.067801 | 0.535118 | NO | NO | NA |
| Dnaaf5 | ENSMUSG( | ENSMUSG( | -0.17216 | 0.086841 | NO | NO | NA |
| Nmd3 | ENSMUSG( | ENSMUSG( | 0.328819 | 7.56E-07 | YES | NO | NA |
| Rabggtb | ENSMUSG( | ENSMUSG( | 0.270438 | 0.001344 | YES | NO | NA |
| Toe1 | ENSMUSG( | ENSMUSG( | 0.260654 | 0.001465 | YES | NO | NA |
| 2210016F1 | ENSMUSG( | ENSMUSG( | 0.115862 | 0.165778 | NO | NO | NA |
| Inip | ENSMUSG( | ENSMUSG( | 0.041467 | 0.729736 | NO | NO | NA |
| Nt5dc1 | ENSMUSG( | ENSMUSG( | 0.085474 | 0.402132 | NO | NO | NA |
| Zmynd19 | ENSMUSG( | ENSMUSG( | 0.363296 | 2.94E-05 | YES | NO | NA |
| Sra1 | ENSMUSG( | ENSMUSG( | -0.03249 | 0.754118 | NO | NO | NA |
| Zbtb14 | ENSMUSG( | ENSMUSG( | -0.24005 | 7.10E-05 | YES | NO | NA |
| Ddx11 | ENSMUSG( | ENSMUSG( | -0.15226 | 0.148023 | NO | NO | NA |
| Xrcc6 | ENSMUSG( | ENSMUSG( | 0.186404 | 0.13277 | NO | NO | NA |
| Pcyt2 | ENSMUSG( | ENSMUSG( | 0.230186 | 0.007339 | YES | NO | NA |
| Cyfip1 | ENSMUSG( | ENSMUSG( | -0.10552 | 0.216756 | NO | NO | NA |
| Lsm8 | ENSMUSG( | ENSMUSG( | 0.053189 | 0.611454 | NO | NO | NA |
| Med30 | ENSMUSG( | ENSMUSG( | -0.12511 | 0.102149 | NO | NO | NA |
| Mppe1 | ENSMUSG( | ENSMUSG( | -0.33985 | 1.59E-05 | YES | NO | NA |
| Tsg101 | ENSMUSG( | ENSMUSG( | -0.11099 | 0.066828 | NO | NO | NA |
| Mrps16 | ENSMUSG( | ENSMUSG( | -0.04448 | 0.538512 | NO | NO | NA |
| Selenos | ENSMUSG( | ENSMUSG( | 0.101329 | 0.243074 | NO | NO | NA |
| Ndufa2 | ENSMUSG( | ENSMUSG( | 0.17773 | 0.001071 | YES | NO | NA |
| Eml2 | ENSMUSG( | ENSMUSG( | -0.18197 | 0.009216 | YES | NO | NA |
| Stk11ip | ENSMUSG( | ENSMUSG( | 0.103725 | 0.433694 | NO | NO | NA |
| Slc16a3 | ENSMUSG( | ENSMUSG( | 0.376208 | 0.181671 | NO | NO | NA |
| Pgam5 | ENSMUSG( | ENSMUSG( | 0.205808 | 0.009127 | YES | NO | NA |
| Mtbp | ENSMUSG( | ENSMUSG( | 0.227584 | 0.006526 | YES | NO | NA |
| Nifk | ENSMUSG( | ENSMUSG( | 0.3668 | 0.000183 | YES | NO | NA |
| Zfp260 | ENSMUSG( | ENSMUSG( | -0.57114 | 3.79E-08 | YES | NO | NA |
| Snx12 | ENSMUSG( | ENSMUSG( | -0.17765 | 0.001951 | YES | NO | NA |
| Noc4l | ENSMUSG( | ENSMUSG( | 0.438306 | 8.73E-07 | YES | NO | NA |
| Parp2 | ENSMUSG( | ENSMUSG( | 0.263878 | 0.002162 | YES | NO | NA |
| Pik3r4 | ENSMUSG( | ENSMUSG( | -0.21385 | 0.008811 | YES | NO | NA |
| Metrn | ENSMUSG( | ENSMUSG( | 0.345537 | 1.82E-07 | YES | NO | NA |
| Ppp1r21 | ENSMUSG( | ENSMUSG( | -0.2065 | 0.000729 | YES | NO | NA |
| Aim2 | ENSMUSG( | ENSMUSG( | 0.055874 | 0.665416 | NO | NO | NA |
| Gm20716 | ENSMUSG( | ENSMUSG( | -0.18097 | 0.476823 | NO | NO | NA |
| Exosc2 | ENSMUSG( | ENSMUSG( | 0.373964 | 8.28E-05 | YES | NO | NA |
| Otud6b | ENSMUSG( | ENSMUSG( | 0.492312 | 0.000528 | YES | NO | NA |

| | | | | | | | |
|---|---|---|---|---|---|---|---|
| Hsf1 | ENSMUSG( | ENSMUSG( | -0.12159 | 0.027222 | YES | NO | NA |
| Spata2 | ENSMUSG( | ENSMUSG( | -0.09958 | 0.428302 | NO | NO | NA |
| Stk35 | ENSMUSG( | ENSMUSG( | -0.12748 | 0.062639 | NO | NO | NA |
| Gpaa1 | ENSMUSG( | ENSMUSG( | 0.018459 | 0.777177 | NO | NO | NA |
| Meaf6 | ENSMUSG( | ENSMUSG( | 0.023075 | 0.786278 | NO | NO | NA |
| Polr2f | ENSMUSG( | ENSMUSG( | 0.421704 | 8.07E-05 | YES | NO | NA |
| Wdr75 | ENSMUSG( | ENSMUSG( | 0.427243 | 4.50E-06 | YES | NO | NA |
| Elac2 | ENSMUSG( | ENSMUSG( | 0.384406 | 1.01E-06 | YES | NO | NA |
| Mlx | ENSMUSG( | ENSMUSG( | -0.20495 | 0.006604 | YES | NO | NA |
| Rexo5 | ENSMUSG( | ENSMUSG( | -0.60788 | 7.19E-08 | YES | NO | NA |
| Rabif | ENSMUSG( | ENSMUSG( | -0.19203 | 0.001501 | YES | NO | NA |
| Pgls | ENSMUSG( | ENSMUSG( | 0.222207 | 0.004563 | YES | NO | NA |
| Nudt16l1 | ENSMUSG( | ENSMUSG( | -0.0669 | 0.35495 | NO | NO | NA |
| Ddhd1 | ENSMUSG( | ENSMUSG( | -0.46732 | 1.98E-09 | YES | NO | NA |
| Fbxl6 | ENSMUSG( | ENSMUSG( | 0.097312 | 0.338282 | NO | NO | NA |
| Zdhhc6 | ENSMUSG( | ENSMUSG( | 0.176313 | 0.049393 | YES | NO | NA |
| Jpt2 | ENSMUSG( | ENSMUSG( | 0.546937 | 9.88E-08 | YES | NO | NA |
| Strip1 | ENSMUSG( | ENSMUSG( | -0.03079 | 0.678924 | NO | NO | NA |
| Taf6 | ENSMUSG( | ENSMUSG( | -0.05022 | 0.569187 | NO | NO | NA |
| Hmg20b | ENSMUSG( | ENSMUSG( | 0.136026 | 0.074096 | NO | NO | NA |
| Mettl23 | ENSMUSG( | ENSMUSG( | -0.28565 | 5.34E-05 | YES | NO | NA |
| Ints4 | ENSMUSG( | ENSMUSG( | -0.10035 | 0.032036 | YES | NO | NA |
| Pofut1 | ENSMUSG( | ENSMUSG( | -0.02276 | 0.801815 | NO | NO | NA |
| Plscr1 | ENSMUSG( | ENSMUSG( | 0.210766 | 0.172742 | NO | NO | NA |
| ENSMUSG( | ENSMUSG( | ENSMUSG( | 0.593207 | 3.12E-11 | YES | NO | NA |
| Klrd1 | ENSMUSG( | ENSMUSG( | -0.19162 | 0.218523 | NO | NO | NA |
| Abcb1b | ENSMUSG( | ENSMUSG( | 0.616678 | 1.35E-09 | YES | NO | NA |
| 0610010K: | ENSMUSG( | ENSMUSG( | 0.112512 | 0.092989 | NO | NO | NA |
| Slc25a28 | ENSMUSG( | ENSMUSG( | 0.075244 | 0.501496 | NO | NO | NA |
| Trappc3 | ENSMUSG( | ENSMUSG( | -0.01763 | 0.851437 | NO | NO | NA |
| Bin3 | ENSMUSG( | ENSMUSG( | 0.024757 | 0.803298 | NO | NO | NA |
| Atp13a2 | ENSMUSG( | ENSMUSG( | 0.131217 | 0.172478 | NO | NO | NA |
| Slc35g1 | ENSMUSG( | ENSMUSG( | 0.507536 | 4.15E-07 | YES | NO | NA |
| Phax | ENSMUSG( | ENSMUSG( | 0.075165 | 0.480077 | NO | NO | NA |
| Ppm1m | ENSMUSG( | ENSMUSG( | -0.3294 | 5.38E-06 | YES | NO | NA |
| Trbv4 | ENSMUSG( | ENSMUSG( | -0.72761 | 0.001897 | YES | NO | NA |
| Gm44174 | ENSMUSG( | ENSMUSG( | -0.83945 | 9.90E-16 | YES | NO | NA |
| Tpcn1 | ENSMUSG( | ENSMUSG( | 0.268598 | 0.001572 | YES | NO | NA |
| Surf1 | ENSMUSG( | ENSMUSG( | 0.04927 | 0.52654 | NO | NO | NA |
| Vta1 | ENSMUSG( | ENSMUSG( | 0.041128 | 0.626634 | NO | NO | NA |
| Tm9sf4 | ENSMUSG( | ENSMUSG( | -0.09388 | 0.119346 | NO | NO | NA |
| Npat | ENSMUSG( | ENSMUSG( | -0.12727 | 0.015477 | YES | NO | NA |
| Rnf168 | ENSMUSG( | ENSMUSG( | -0.05909 | 0.323055 | NO | NO | NA |
| Mospd3 | ENSMUSG( | ENSMUSG( | -0.11273 | 0.203991 | NO | NO | NA |
| Leo1 | ENSMUSG( | ENSMUSG( | -0.08671 | 0.413253 | NO | NO | NA |
| 6030458C: | ENSMUSG( | ENSMUSG( | -0.01454 | 0.844265 | NO | NO | NA |
| Znrf1 | ENSMUSG( | ENSMUSG( | -0.14075 | 0.206006 | NO | NO | NA |

| | | | | | | | |
|---|---|---|---|---|---|---|---|
| 6530402F1 | ENSMUSG0 | ENSMUSG0 | -0.3342 | 3.02E-07 | YES | NO | NA |
| Rab11a | ENSMUSG0 | ENSMUSG0 | -0.12819 | 0.054094 | NO | NO | NA |
| Shmt1 | ENSMUSG0 | ENSMUSG0 | 0.81262 | 1.31E-07 | YES | NO | NA |
| Prim2 | ENSMUSG0 | ENSMUSG0 | 0.313339 | 2.88E-08 | YES | NO | NA |
| Timm17a | ENSMUSG0 | ENSMUSG0 | 0.222264 | 0.007088 | YES | NO | NA |
| Bahd1 | ENSMUSG0 | ENSMUSG0 | -0.37451 | 4.94E-08 | YES | NO | NA |
| Fbxo31 | ENSMUSG0 | ENSMUSG0 | -0.1501 | 0.059498 | NO | NO | NA |
| Ube2e1 | ENSMUSG0 | ENSMUSG0 | 0.134469 | 0.102306 | NO | NO | NA |
| Ece1 | ENSMUSG0 | ENSMUSG0 | 0.31219 | 0.000213 | YES | NO | NA |
| Dpy30 | ENSMUSG0 | ENSMUSG0 | 0.173496 | 0.050239 | NO | NO | NA |
| Arl8a | ENSMUSG0 | ENSMUSG0 | -0.35899 | 2.87E-07 | YES | NO | NA |
| Ulk3 | ENSMUSG0 | ENSMUSG0 | -0.37417 | 6.05E-07 | YES | NO | NA |
| mt-Nd4l | ENSMUSG0 | ENSMUSG0 | 0.25107 | 0.296699 | NO | NO | NA |
| Arhgap18 | ENSMUSG0 | ENSMUSG0 | -0.22918 | 0.012693 | YES | NO | NA |
| Mafk | ENSMUSG0 | ENSMUSG0 | -0.18212 | 0.164525 | NO | NO | NA |
| Bzw2 | ENSMUSG0 | ENSMUSG0 | 0.690679 | 1.09E-08 | YES | NO | NA |
| Sec61g | ENSMUSG0 | ENSMUSG0 | 0.133408 | 0.024824 | YES | NO | NA |
| Mrpl15 | ENSMUSG0 | ENSMUSG0 | 0.235823 | 0.006913 | YES | NO | NA |
| Tyw1 | ENSMUSG0 | ENSMUSG0 | 0.017475 | 0.826558 | NO | NO | NA |
| Tgfbrap1 | ENSMUSG0 | ENSMUSG0 | -0.24644 | 0.000706 | YES | NO | NA |
| Aven | ENSMUSG0 | ENSMUSG0 | -0.01189 | 0.894092 | NO | NO | NA |
| Zmym5 | ENSMUSG0 | ENSMUSG0 | -0.39606 | 9.68E-05 | YES | NO | NA |
| Cop1 | ENSMUSG0 | ENSMUSG0 | -0.02325 | 0.758707 | NO | NO | NA |
| Ap3s2 | ENSMUSG0 | ENSMUSG0 | 0.155063 | 0.038915 | YES | NO | NA |
| Acbd5 | ENSMUSG0 | ENSMUSG0 | -0.13032 | 0.177188 | NO | NO | NA |
| Esm1 | ENSMUSG0 | ENSMUSG0 | 1.146375 | 3.27E-07 | YES | YES | Upregulated |
| Braf | ENSMUSG0 | ENSMUSG0 | -0.20163 | 0.00699 | YES | NO | NA |
| Baiap2 | ENSMUSG0 | ENSMUSG0 | -0.16232 | 0.006617 | YES | NO | NA |
| Zfp52 | ENSMUSG0 | ENSMUSG0 | -0.04471 | 0.769274 | NO | NO | NA |
| Pih1d1 | ENSMUSG0 | ENSMUSG0 | 0.035323 | 0.655006 | NO | NO | NA |
| Nob1 | ENSMUSG0 | ENSMUSG0 | 0.292469 | 0.00154 | YES | NO | NA |
| Serpinb6a | ENSMUSG0 | ENSMUSG0 | 0.273199 | 0.00916 | YES | NO | NA |
| Xrcc5 | ENSMUSG0 | ENSMUSG0 | 0.335677 | 5.88E-06 | YES | NO | NA |
| Hid1 | ENSMUSG0 | ENSMUSG0 | -0.30393 | 0.071195 | NO | NO | NA |
| Casp7 | ENSMUSG0 | ENSMUSG0 | -0.39466 | 7.56E-08 | YES | NO | NA |
| Srprb | ENSMUSG0 | ENSMUSG0 | -0.30087 | 1.31E-08 | YES | NO | NA |
| Nol10 | ENSMUSG0 | ENSMUSG0 | 0.211871 | 0.0056 | YES | NO | NA |
| Kifc5b | ENSMUSG0 | ENSMUSG0 | 0.225804 | 0.007328 | YES | NO | NA |
| Gimap1os | ENSMUSG0 | ENSMUSG0 | -0.54634 | 7.59E-07 | YES | NO | NA |
| Net1 | ENSMUSG0 | ENSMUSG0 | 0.017408 | 0.833737 | NO | NO | NA |
| AW54987 | ENSMUSG0 | ENSMUSG0 | -0.44998 | 5.13E-05 | YES | NO | NA |
| Rab19 | ENSMUSG0 | ENSMUSG0 | -0.06114 | 0.616526 | NO | NO | NA |
| Msto1 | ENSMUSG0 | ENSMUSG0 | 0.391239 | 6.21E-07 | YES | NO | NA |
| Zfp414 | ENSMUSG0 | ENSMUSG0 | 0.345645 | 0.002449 | YES | NO | NA |
| Hrh2 | ENSMUSG0 | ENSMUSG0 | -0.27734 | 0.041403 | YES | NO | NA |
| Brcc3 | ENSMUSG0 | ENSMUSG0 | 0.058693 | 0.547231 | NO | NO | NA |
| Gm28036 | ENSMUSG0 | ENSMUSG0 | 0.03579 | 0.818259 | NO | NO | NA |

| | | | | | | | |
|---|---|---|---|---|---|---|---|
| Mapkapk5 | ENSMUSG( | ENSMUSG( | -0.01901 | 0.84081 | NO | NO | NA |
| Atp11c | ENSMUSG( | ENSMUSG( | -0.2503 | 0.006856 | YES | NO | NA |
| Socs3 | ENSMUSG( | ENSMUSG( | 0.289967 | 0.053551 | NO | NO | NA |
| Patz1 | ENSMUSG( | ENSMUSG( | 0.023651 | 0.749636 | NO | NO | NA |
| Miip | ENSMUSG( | ENSMUSG( | -0.42373 | 3.30E-05 | YES | NO | NA |
| Zbtb8os | ENSMUSG( | ENSMUSG( | -0.01621 | 0.814223 | NO | NO | NA |
| Ska2 | ENSMUSG( | ENSMUSG( | -0.08109 | 0.421068 | NO | NO | NA |
| Gtf2a1 | ENSMUSG( | ENSMUSG( | -0.27652 | 0.000666 | YES | NO | NA |
| Sqle | ENSMUSG( | ENSMUSG( | 0.955174 | 3.61E-08 | YES | NO | NA |
| Cyth3 | ENSMUSG( | ENSMUSG( | 0.168219 | 0.389673 | NO | NO | NA |
| Ltv1 | ENSMUSG( | ENSMUSG( | 0.483672 | 3.57E-05 | YES | NO | NA |
| Nipsnap2 | ENSMUSG( | ENSMUSG( | 0.253033 | 0.019355 | YES | NO | NA |
| Pitrm1 | ENSMUSG( | ENSMUSG( | 0.317391 | 0.000278 | YES | NO | NA |
| Errfi1 | ENSMUSG( | ENSMUSG( | -0.58789 | 3.93E-05 | YES | NO | NA |
| Polr3a | ENSMUSG( | ENSMUSG( | 0.051904 | 0.528583 | NO | NO | NA |
| Trim34a | ENSMUSG( | ENSMUSG( | -0.8707 | 9.31E-14 | YES | NO | NA |
| Imp3 | ENSMUSG( | ENSMUSG( | 0.25083 | 0.000641 | YES | NO | NA |
| Podnl1 | ENSMUSG( | ENSMUSG( | 0.38321 | 0.005538 | YES | NO | NA |
| Med23 | ENSMUSG( | ENSMUSG( | -0.08105 | 0.441436 | NO | NO | NA |
| Mrpl37 | ENSMUSG( | ENSMUSG( | 0.191208 | 0.024006 | YES | NO | NA |
| Psmf1 | ENSMUSG( | ENSMUSG( | 0.079254 | 0.450533 | NO | NO | NA |
| Dnttip1 | ENSMUSG( | ENSMUSG( | 0.020406 | 0.862377 | NO | NO | NA |
| Ndufa1 | ENSMUSG( | ENSMUSG( | 0.079576 | 0.465617 | NO | NO | NA |
| Mob2 | ENSMUSG( | ENSMUSG( | 0.195393 | 0.087541 | NO | NO | NA |
| Bfar | ENSMUSG( | ENSMUSG( | -0.03946 | 0.653348 | NO | NO | NA |
| Nelfa | ENSMUSG( | ENSMUSG( | -0.00317 | 0.970155 | NO | NO | NA |
| Vti1a | ENSMUSG( | ENSMUSG( | 0.298645 | 0.005714 | YES | NO | NA |
| Zswim6 | ENSMUSG( | ENSMUSG( | -0.66639 | 3.84E-12 | YES | NO | NA |
| Sod2 | ENSMUSG( | ENSMUSG( | 0.144773 | 0.03596 | YES | NO | NA |
| Fbxo33 | ENSMUSG( | ENSMUSG( | -0.17189 | 0.092239 | NO | NO | NA |
| Neurl4 | ENSMUSG( | ENSMUSG( | -0.14226 | 0.01246 | YES | NO | NA |
| Steep1 | ENSMUSG( | ENSMUSG( | -0.29577 | 0.000171 | YES | NO | NA |
| Plekha3 | ENSMUSG( | ENSMUSG( | -0.31767 | 1.97E-05 | YES | NO | NA |
| D5Ertd579 | ENSMUSG( | ENSMUSG( | -0.27974 | 0.001211 | YES | NO | NA |
| Vps26c | ENSMUSG( | ENSMUSG( | -0.11376 | 0.091364 | NO | NO | NA |
| Bola2 | ENSMUSG( | ENSMUSG( | 0.394688 | 4.63E-10 | YES | NO | NA |
| E4f1 | ENSMUSG( | ENSMUSG( | -0.13391 | 0.145481 | NO | NO | NA |
| Itfg1 | ENSMUSG( | ENSMUSG( | -0.17187 | 0.185833 | NO | NO | NA |
| Tti1 | ENSMUSG( | ENSMUSG( | -0.13487 | 0.020248 | YES | NO | NA |
| Inpp5e | ENSMUSG( | ENSMUSG( | -0.22463 | 0.024932 | YES | NO | NA |
| Snap29 | ENSMUSG( | ENSMUSG( | -0.36684 | 7.41E-10 | YES | NO | NA |
| Snx9 | ENSMUSG( | ENSMUSG( | 0.750676 | 3.96E-06 | YES | NO | NA |
| Mvp | ENSMUSG( | ENSMUSG( | -0.15471 | 0.0063 | YES | NO | NA |
| Snip1 | ENSMUSG( | ENSMUSG( | 0.033582 | 0.594461 | NO | NO | NA |
| Tbpl1 | ENSMUSG( | ENSMUSG( | -0.20739 | 0.001279 | YES | NO | NA |
| Itpa | ENSMUSG( | ENSMUSG( | 0.308347 | 1.09E-05 | YES | NO | NA |
| Uqcrb | ENSMUSG( | ENSMUSG( | -0.10525 | 0.403857 | NO | NO | NA |

| | | | | | | | |
|---|---|---|---|---|---|---|---|
| Gcc1 | ENSMUSG( | ENSMUSG( | -0.30622 | 1.09E-06 | YES | NO | NA |
| Tent4a | ENSMUSG( | ENSMUSG( | -0.1826 | 0.007443 | YES | NO | NA |
| Ttll4 | ENSMUSG( | ENSMUSG( | 0.030015 | 0.791854 | NO | NO | NA |
| Lrif1 | ENSMUSG( | ENSMUSG( | -0.11151 | 0.498479 | NO | NO | NA |
| Ndufs6 | ENSMUSG( | ENSMUSG( | 0.153515 | 0.250124 | NO | NO | NA |
| Fam71b | ENSMUSG( | ENSMUSG( | -0.50282 | 4.86E-05 | YES | NO | NA |
| Spns1 | ENSMUSG( | ENSMUSG( | -0.17636 | 0.00077 | YES | NO | NA |
| Hbb-bt | ENSMUSG( | ENSMUSG( | 0.4962 | 0.510027 | NO | NO | NA |
| Prdx3 | ENSMUSG( | ENSMUSG( | 0.220057 | 0.000619 | YES | NO | NA |
| Gng5 | ENSMUSG( | ENSMUSG( | -0.26394 | 8.07E-05 | YES | NO | NA |
| Dhx58 | ENSMUSG( | ENSMUSG( | -0.75626 | 3.98E-10 | YES | NO | NA |
| Slc25a30 | ENSMUSG( | ENSMUSG( | -0.14695 | 0.027691 | YES | NO | NA |
| Zmym2 | ENSMUSG( | ENSMUSG( | -0.39259 | 8.32E-07 | YES | NO | NA |
| Mrtfb | ENSMUSG( | ENSMUSG( | -0.19628 | 0.011456 | YES | NO | NA |
| Dhx37 | ENSMUSG( | ENSMUSG( | 0.11314 | 0.115539 | NO | NO | NA |
| Sde2 | ENSMUSG( | ENSMUSG( | -0.08503 | 0.379311 | NO | NO | NA |
| Elp1 | ENSMUSG( | ENSMUSG( | 0.325226 | 0.001196 | YES | NO | NA |
| Oat | ENSMUSG( | ENSMUSG( | 0.11932 | 0.066782 | NO | NO | NA |
| Rnf146 | ENSMUSG( | ENSMUSG( | -0.28491 | 0.000109 | YES | NO | NA |
| Gemin5 | ENSMUSG( | ENSMUSG( | 0.313116 | 0.00062 | YES | NO | NA |
| Mrpl57 | ENSMUSG( | ENSMUSG( | 0.289013 | 1.36E-07 | YES | NO | NA |
| St6galnac4 | ENSMUSG( | ENSMUSG( | 0.066558 | 0.517147 | NO | NO | NA |
| Fbxw5 | ENSMUSG( | ENSMUSG( | 0.025362 | 0.838627 | NO | NO | NA |
| Abhd13 | ENSMUSG( | ENSMUSG( | -0.37368 | 1.71E-06 | YES | NO | NA |
| Slc25a22 | ENSMUSG( | ENSMUSG( | -0.06216 | 0.335564 | NO | NO | NA |
| Scfd1 | ENSMUSG( | ENSMUSG( | -0.0417 | 0.620189 | NO | NO | NA |
| Hps1 | ENSMUSG( | ENSMUSG( | -0.12281 | 0.186591 | NO | NO | NA |
| Polr2g | ENSMUSG( | ENSMUSG( | -0.04125 | 0.537952 | NO | NO | NA |
| Fam114a2 | ENSMUSG( | ENSMUSG( | -0.21685 | 0.006673 | YES | NO | NA |
| Slc4a1ap | ENSMUSG( | ENSMUSG( | -0.15475 | 0.097535 | NO | NO | NA |
| Tex10 | ENSMUSG( | ENSMUSG( | 0.00345 | 0.968865 | NO | NO | NA |
| Pofut2 | ENSMUSG( | ENSMUSG( | 0.11191 | 0.204408 | NO | NO | NA |
| Kdm6b | ENSMUSG( | ENSMUSG( | -0.07687 | 0.558028 | NO | NO | NA |
| Ppm1d | ENSMUSG( | ENSMUSG( | -0.1576 | 0.009489 | YES | NO | NA |
| G2e3 | ENSMUSG( | ENSMUSG( | -0.30479 | 0.001071 | YES | NO | NA |
| Atp6v1g1 | ENSMUSG( | ENSMUSG( | -0.1248 | 0.118757 | NO | NO | NA |
| Wdr46 | ENSMUSG( | ENSMUSG( | 0.531971 | 5.46E-05 | YES | NO | NA |
| Bicdl1 | ENSMUSG( | ENSMUSG( | -0.00316 | 0.981021 | NO | NO | NA |
| Mtmr9 | ENSMUSG( | ENSMUSG( | -0.05483 | 0.552976 | NO | NO | NA |
| Elp3 | ENSMUSG( | ENSMUSG( | 0.178645 | 0.009409 | YES | NO | NA |
| Pus1 | ENSMUSG( | ENSMUSG( | 0.320113 | 0.015696 | YES | NO | NA |
| Zfp507 | ENSMUSG( | ENSMUSG( | -0.22353 | 8.73E-05 | YES | NO | NA |
| Wdr62 | ENSMUSG( | ENSMUSG( | 0.231166 | 0.022752 | YES | NO | NA |
| Sos1 | ENSMUSG( | ENSMUSG( | -0.53527 | 1.95E-11 | YES | NO | NA |
| Wsb2 | ENSMUSG( | ENSMUSG( | -0.10609 | 0.341852 | NO | NO | NA |
| Acot9 | ENSMUSG( | ENSMUSG( | 0.118322 | 0.298442 | NO | NO | NA |
| Haus5 | ENSMUSG( | ENSMUSG( | 0.516558 | 2.20E-10 | YES | NO | NA |

| | | | | | | | |
|---|---|---|---|---|---|---|---|
| 8030462N | ENSMUSG( | ENSMUSG( | -0.19036 | 0.001552 | YES | NO | NA |
| Mcrip1 | ENSMUSG( | ENSMUSG( | 0.052738 | 0.675764 | NO | NO | NA |
| Myg1 | ENSMUSG( | ENSMUSG( | 0.091533 | 0.112117 | NO | NO | NA |
| Emc1 | ENSMUSG( | ENSMUSG( | 0.127736 | 0.062463 | NO | NO | NA |
| Slc25a39 | ENSMUSG( | ENSMUSG( | 0.237583 | 0.006617 | YES | NO | NA |
| Phactr2 | ENSMUSG( | ENSMUSG( | -0.46049 | 0.001071 | YES | NO | NA |
| Bcl7b | ENSMUSG( | ENSMUSG( | -0.15456 | 0.039349 | YES | NO | NA |
| Cetn3 | ENSMUSG( | ENSMUSG( | 0.104054 | 0.431229 | NO | NO | NA |
| Pdcl | ENSMUSG( | ENSMUSG( | -0.2341 | 0.000149 | YES | NO | NA |
| Gbp2b | ENSMUSG( | ENSMUSG( | 8.344507 | 1.70E-163 | YES | YES | Upregulated |
| Zbtb33 | ENSMUSG( | ENSMUSG( | -0.14872 | 0.034718 | YES | NO | NA |
| Taf3 | ENSMUSG( | ENSMUSG( | -0.00858 | 0.945713 | NO | NO | NA |
| Hmbs | ENSMUSG( | ENSMUSG( | 0.508116 | 6.91E-08 | YES | NO | NA |
| Zbtb44 | ENSMUSG( | ENSMUSG( | -0.29604 | 0.000425 | YES | NO | NA |
| Chpf2 | ENSMUSG( | ENSMUSG( | -0.25855 | 0.004118 | YES | NO | NA |
| Tmem128 | ENSMUSG( | ENSMUSG( | -0.01987 | 0.888811 | NO | NO | NA |
| Mtf1 | ENSMUSG( | ENSMUSG( | -0.26716 | 1.98E-05 | YES | NO | NA |
| Thap3 | ENSMUSG( | ENSMUSG( | -0.68065 | 1.66E-09 | YES | NO | NA |
| Urb2 | ENSMUSG( | ENSMUSG( | 0.327824 | 0.000915 | YES | NO | NA |
| Sdr39u1 | ENSMUSG( | ENSMUSG( | -0.05516 | 0.579782 | NO | NO | NA |
| Tex2 | ENSMUSG( | ENSMUSG( | 0.358636 | 0.001375 | YES | NO | NA |
| Donson | ENSMUSG( | ENSMUSG( | -0.10814 | 0.145063 | NO | NO | NA |
| Golim4 | ENSMUSG( | ENSMUSG( | -0.16107 | 0.060594 | NO | NO | NA |
| Bcl2l1 | ENSMUSG( | ENSMUSG( | -0.28011 | 0.009199 | YES | NO | NA |
| Dapk2 | ENSMUSG( | ENSMUSG( | -1.45443 | 1.65E-13 | YES | YES | Downregulated |
| Tceal9 | ENSMUSG( | ENSMUSG( | 0.049347 | 0.676261 | NO | NO | NA |
| Mien1 | ENSMUSG( | ENSMUSG( | -0.26293 | 0.000332 | YES | NO | NA |
| Zfp869 | ENSMUSG( | ENSMUSG( | -0.18658 | 0.017278 | YES | NO | NA |
| Cntrob | ENSMUSG( | ENSMUSG( | -0.1853 | 0.051246 | NO | NO | NA |
| Kxd1 | ENSMUSG( | ENSMUSG( | 0.077178 | 0.432927 | NO | NO | NA |
| Dctn3 | ENSMUSG( | ENSMUSG( | -0.0768 | 0.389594 | NO | NO | NA |
| Trip13 | ENSMUSG( | ENSMUSG( | 0.622012 | 6.93E-08 | YES | NO | NA |
| Gmcl1 | ENSMUSG( | ENSMUSG( | -0.14192 | 0.034705 | YES | NO | NA |
| Tsc1 | ENSMUSG( | ENSMUSG( | -0.10069 | 0.255296 | NO | NO | NA |
| Llgl2 | ENSMUSG( | ENSMUSG( | -0.6074 | 3.53E-09 | YES | NO | NA |
| Them6 | ENSMUSG( | ENSMUSG( | 0.368034 | 0.000301 | YES | NO | NA |
| H2-Q10 | ENSMUSG( | ENSMUSG( | -0.2137 | 0.084106 | NO | NO | NA |
| Dhdds | ENSMUSG( | ENSMUSG( | 0.074937 | 0.238544 | NO | NO | NA |
| Zfas1 | ENSMUSG( | ENSMUSG( | 0.483095 | 1.85E-05 | YES | NO | NA |
| Glo1 | ENSMUSG( | ENSMUSG( | 0.178905 | 0.031979 | YES | NO | NA |
| Gm42031 | ENSMUSG( | ENSMUSG( | 1.506241 | 1.65E-14 | YES | YES | Upregulated |
| Pdxdc1 | ENSMUSG( | ENSMUSG( | 0.156069 | 0.052571 | NO | NO | NA |
| Tdp2 | ENSMUSG( | ENSMUSG( | 0.127005 | 0.101811 | NO | NO | NA |
| Eny2 | ENSMUSG( | ENSMUSG( | 0.202793 | 0.016012 | YES | NO | NA |
| Dleu2 | ENSMUSG( | ENSMUSG( | -0.4964 | 1.05E-05 | YES | NO | NA |
| Agpat4 | ENSMUSG( | ENSMUSG( | -0.06384 | 0.656664 | NO | NO | NA |
| Dhx57 | ENSMUSG( | ENSMUSG( | -0.26299 | 0.000945 | YES | NO | NA |

| | | | | | | | |
|---|---|---|---|---|---|---|---|
| Abhd12 | ENSMUSG( | ENSMUSG( | -0.02642 | 0.80969 | NO | NO | NA |
| Soat1 | ENSMUSG( | ENSMUSG( | -0.33951 | 3.43E-09 | YES | NO | NA |
| Kat5 | ENSMUSG( | ENSMUSG( | 0.103795 | 0.073653 | NO | NO | NA |
| Gfm2 | ENSMUSG( | ENSMUSG( | -0.15924 | 0.094161 | NO | NO | NA |
| Spag7 | ENSMUSG( | ENSMUSG( | -0.10117 | 0.384021 | NO | NO | NA |
| Plk3 | ENSMUSG( | ENSMUSG( | 0.067709 | 0.703822 | NO | NO | NA |
| Acbd6 | ENSMUSG( | ENSMUSG( | 0.195136 | 0.005743 | YES | NO | NA |
| Ing5 | ENSMUSG( | ENSMUSG( | 0.02888 | 0.687926 | NO | NO | NA |
| H2-M3 | ENSMUSG( | ENSMUSG( | -0.66773 | 3.23E-09 | YES | NO | NA |
| Trnt1 | ENSMUSG( | ENSMUSG( | 0.110047 | 0.244262 | NO | NO | NA |
| Psen2 | ENSMUSG( | ENSMUSG( | -0.18277 | 0.020214 | YES | NO | NA |
| Atg5 | ENSMUSG( | ENSMUSG( | -0.04451 | 0.599709 | NO | NO | NA |
| Eif4e3 | ENSMUSG( | ENSMUSG( | 0.130058 | 0.279303 | NO | NO | NA |
| Rngtt | ENSMUSG( | ENSMUSG( | -0.11651 | 0.169272 | NO | NO | NA |
| Opa3 | ENSMUSG( | ENSMUSG( | 0.002714 | 0.976434 | NO | NO | NA |
| Slc50a1 | ENSMUSG( | ENSMUSG( | -0.10475 | 0.327762 | NO | NO | NA |
| Ubap1 | ENSMUSG( | ENSMUSG( | -0.28424 | 0.000107 | YES | NO | NA |
| Tnfrsf18 | ENSMUSG( | ENSMUSG( | 0.121178 | 0.187457 | NO | NO | NA |
| Cpeb4 | ENSMUSG( | ENSMUSG( | -0.52598 | 9.93E-07 | YES | NO | NA |
| Srbd1 | ENSMUSG( | ENSMUSG( | -0.01495 | 0.900065 | NO | NO | NA |
| Lrrfip2 | ENSMUSG( | ENSMUSG( | -0.29058 | 3.00E-06 | YES | NO | NA |
| Pmpcb | ENSMUSG( | ENSMUSG( | 0.269357 | 0.000657 | YES | NO | NA |
| Ccdc71l | ENSMUSG( | ENSMUSG( | -0.255 | 0.000436 | YES | NO | NA |
| Rwdd4a | ENSMUSG( | ENSMUSG( | 0.108188 | 0.052273 | NO | NO | NA |
| Maml2 | ENSMUSG( | ENSMUSG( | -0.5477 | 9.84E-06 | YES | NO | NA |
| Twf1 | ENSMUSG( | ENSMUSG( | 0.139347 | 0.173351 | NO | NO | NA |
| Smim14 | ENSMUSG( | ENSMUSG( | -0.28475 | 0.002352 | YES | NO | NA |
| Bora | ENSMUSG( | ENSMUSG( | 0.314815 | 2.16E-06 | YES | NO | NA |
| Dusp6 | ENSMUSG( | ENSMUSG( | 0.229664 | 0.086401 | NO | NO | NA |
| Sdf2 | ENSMUSG( | ENSMUSG( | -0.33964 | 6.61E-08 | YES | NO | NA |
| Armc5 | ENSMUSG( | ENSMUSG( | -0.14066 | 0.114735 | NO | NO | NA |
| Stk16 | ENSMUSG( | ENSMUSG( | 0.154547 | 0.043201 | YES | NO | NA |
| Haspin | ENSMUSG( | ENSMUSG( | -0.09498 | 0.158805 | NO | NO | NA |
| Pttg1 | ENSMUSG( | ENSMUSG( | -0.74304 | 1.75E-20 | YES | NO | NA |
| Timm29 | ENSMUSG( | ENSMUSG( | 0.048615 | 0.601436 | NO | NO | NA |
| Irgm2 | ENSMUSG( | ENSMUSG( | -0.68199 | 4.35E-12 | YES | NO | NA |
| Zranb1 | ENSMUSG( | ENSMUSG( | -0.02903 | 0.784854 | NO | NO | NA |
| Melk | ENSMUSG( | ENSMUSG( | 0.377878 | 1.25E-06 | YES | NO | NA |
| Ndufb3 | ENSMUSG( | ENSMUSG( | 0.066111 | 0.671878 | NO | NO | NA |
| Gadd45gip | ENSMUSG( | ENSMUSG( | 0.281664 | 0.004442 | YES | NO | NA |
| Ube2o | ENSMUSG( | ENSMUSG( | 0.219382 | 0.014119 | YES | NO | NA |
| Fabp5 | ENSMUSG( | ENSMUSG( | 0.852071 | 8.81E-08 | YES | NO | NA |
| Mob4 | ENSMUSG( | ENSMUSG( | -0.23242 | 0.007857 | YES | NO | NA |
| Ptcd3 | ENSMUSG( | ENSMUSG( | 0.127937 | 0.120598 | NO | NO | NA |
| Jdp2 | ENSMUSG( | ENSMUSG( | -0.16845 | 0.172742 | NO | NO | NA |
| Cchcr1 | ENSMUSG( | ENSMUSG( | 0.005995 | 0.957086 | NO | NO | NA |
| Nfs1 | ENSMUSG( | ENSMUSG( | 0.198094 | 0.002346 | YES | NO | NA |

| | | | | | | | |
|---|---|---|---|---|---|---|---|
| Edem2 | ENSMUSG( | ENSMUSG( | -0.08375 | 0.171785 | NO | NO | NA |
| Foxo4 | ENSMUSG( | ENSMUSG( | -0.41844 | 2.96E-06 | YES | NO | NA |
| Irak4 | ENSMUSG( | ENSMUSG( | -0.36197 | 7.81E-06 | YES | NO | NA |
| Snx20 | ENSMUSG( | ENSMUSG( | -0.358 | 2.67E-06 | YES | NO | NA |
| Unk | ENSMUSG( | ENSMUSG( | -0.08417 | 0.297993 | NO | NO | NA |
| Trim5 | ENSMUSG( | ENSMUSG( | -0.48871 | 1.79E-07 | YES | NO | NA |
| Lrrc58 | ENSMUSG( | ENSMUSG( | 0.008493 | 0.936625 | NO | NO | NA |
| Mfap3 | ENSMUSG( | ENSMUSG( | -0.15889 | 0.033871 | YES | NO | NA |
| Ergic1 | ENSMUSG( | ENSMUSG( | 0.269342 | 1.04E-05 | YES | NO | NA |
| Tgs1 | ENSMUSG( | ENSMUSG( | -0.09676 | 0.303467 | NO | NO | NA |
| Slc48a1 | ENSMUSG( | ENSMUSG( | 0.303867 | 0.010623 | YES | NO | NA |
| Eif1b | ENSMUSG( | ENSMUSG( | -0.13031 | 0.046216 | YES | NO | NA |
| Nol12 | ENSMUSG( | ENSMUSG( | 0.258714 | 0.001515 | YES | NO | NA |
| Slc7a6os | ENSMUSG( | ENSMUSG( | -0.01085 | 0.861045 | NO | NO | NA |
| Stard10 | ENSMUSG( | ENSMUSG( | -0.67489 | 4.89E-09 | YES | NO | NA |
| Rala | ENSMUSG( | ENSMUSG( | -0.01616 | 0.925689 | NO | NO | NA |
| Pcnx | ENSMUSG( | ENSMUSG( | -0.08735 | 0.406889 | NO | NO | NA |
| Zmym3 | ENSMUSG( | ENSMUSG( | -0.07901 | 0.305056 | NO | NO | NA |
| Mxd3 | ENSMUSG( | ENSMUSG( | 0.049733 | 0.666266 | NO | NO | NA |
| Dop1a | ENSMUSG( | ENSMUSG( | -0.06039 | 0.532755 | NO | NO | NA |
| Nae1 | ENSMUSG( | ENSMUSG( | 0.03057 | 0.771401 | NO | NO | NA |
| Sbds | ENSMUSG( | ENSMUSG( | 0.07578 | 0.346178 | NO | NO | NA |
| Dynlt1f | ENSMUSG( | ENSMUSG( | 0.01078 | 0.923109 | NO | NO | NA |
| Lsm14b | ENSMUSG( | ENSMUSG( | -0.39545 | 6.50E-08 | YES | NO | NA |
| Commd4 | ENSMUSG( | ENSMUSG( | -0.12744 | 0.316602 | NO | NO | NA |
| Zfp282 | ENSMUSG( | ENSMUSG( | 0.066454 | 0.400316 | NO | NO | NA |
| Lipa | ENSMUSG( | ENSMUSG( | -0.33315 | 1.49E-07 | YES | NO | NA |
| Cdc16 | ENSMUSG( | ENSMUSG( | 0.213478 | 0.004217 | YES | NO | NA |
| Fbxo45 | ENSMUSG( | ENSMUSG( | -0.04274 | 0.562171 | NO | NO | NA |
| Elof1 | ENSMUSG( | ENSMUSG( | 0.334804 | 3.48E-08 | YES | NO | NA |
| Pex5 | ENSMUSG( | ENSMUSG( | -0.11581 | 0.286804 | NO | NO | NA |
| Tbce | ENSMUSG( | ENSMUSG( | 0.085015 | 0.454653 | NO | NO | NA |
| Deaf1 | ENSMUSG( | ENSMUSG( | 0.044112 | 0.600536 | NO | NO | NA |
| Car5b | ENSMUSG( | ENSMUSG( | 0.175605 | 0.31092 | NO | NO | NA |
| Pin1 | ENSMUSG( | ENSMUSG( | 0.164613 | 0.028425 | YES | NO | NA |
| Cspp1 | ENSMUSG( | ENSMUSG( | -0.19248 | 0.08615 | NO | NO | NA |
| Cir1 | ENSMUSG( | ENSMUSG( | -0.15881 | 0.225444 | NO | NO | NA |
| Zcrb1 | ENSMUSG( | ENSMUSG( | -0.03211 | 0.71797 | NO | NO | NA |
| Alg5 | ENSMUSG( | ENSMUSG( | 0.223322 | 0.013718 | YES | NO | NA |
| Xylt2 | ENSMUSG( | ENSMUSG( | -0.06255 | 0.58144 | NO | NO | NA |
| C2cd2l | ENSMUSG( | ENSMUSG( | 0.040893 | 0.601067 | NO | NO | NA |
| Trbv29 | ENSMUSG( | ENSMUSG( | -0.02085 | 0.970413 | NO | NO | NA |
| Polr2j | ENSMUSG( | ENSMUSG( | 0.252052 | 0.022562 | YES | NO | NA |
| Champ1 | ENSMUSG( | ENSMUSG( | 0.00117 | 0.986386 | NO | NO | NA |
| Brip1 | ENSMUSG( | ENSMUSG( | 0.05351 | 0.655215 | NO | NO | NA |
| Wscd2 | ENSMUSG( | ENSMUSG( | -0.45389 | 0.000123 | YES | NO | NA |
| Yipf4 | ENSMUSG( | ENSMUSG( | -0.18307 | 0.015868 | YES | NO | NA |

| | | | | | | | |
|---|---|---|---|---|---|---|---|
| Rbm4 | ENSMUSG( | ENSMUSG( | -0.29374 | 1.82E-05 | YES | NO | NA |
| Cab39l | ENSMUSG( | ENSMUSG( | -0.1824 | 0.003204 | YES | NO | NA |
| Zfp687 | ENSMUSG( | ENSMUSG( | -0.10739 | 0.231157 | NO | NO | NA |
| Ap1s3 | ENSMUSG( | ENSMUSG( | -0.17087 | 0.02961 | YES | NO | NA |
| Fkrp | ENSMUSG( | ENSMUSG( | -0.21933 | 0.006944 | YES | NO | NA |
| Trbv16 | ENSMUSG( | ENSMUSG( | 0.33711 | 0.33087 | NO | NO | NA |
| Atg13 | ENSMUSG( | ENSMUSG( | -0.21054 | 0.019585 | YES | NO | NA |
| Phf20 | ENSMUSG( | ENSMUSG( | 0.008114 | 0.930011 | NO | NO | NA |
| Anln | ENSMUSG( | ENSMUSG( | -0.02143 | 0.844349 | NO | NO | NA |
| Pms2 | ENSMUSG( | ENSMUSG( | 0.118509 | 0.066938 | NO | NO | NA |
| Faim | ENSMUSG( | ENSMUSG( | -0.01001 | 0.921065 | NO | NO | NA |
| Dhcr7 | ENSMUSG( | ENSMUSG( | -0.03636 | 0.517784 | NO | NO | NA |
| Thumpd3 | ENSMUSG( | ENSMUSG( | 0.001075 | 0.989942 | NO | NO | NA |
| Zfp68 | ENSMUSG( | ENSMUSG( | -0.2419 | 0.002871 | YES | NO | NA |
| Zfp319 | ENSMUSG( | ENSMUSG( | -0.48645 | 6.61E-08 | YES | NO | NA |
| Tusc3 | ENSMUSG( | ENSMUSG( | 0.187116 | 0.113719 | NO | NO | NA |
| Hipk2 | ENSMUSG( | ENSMUSG( | -0.33507 | 0.00068 | YES | NO | NA |
| Dbr1 | ENSMUSG( | ENSMUSG( | 0.125156 | 0.21965 | NO | NO | NA |
| Abcc5 | ENSMUSG( | ENSMUSG( | -0.25329 | 0.016857 | YES | NO | NA |
| Zfp800 | ENSMUSG( | ENSMUSG( | -0.30049 | 0.000364 | YES | NO | NA |
| Mknk1 | ENSMUSG( | ENSMUSG( | -0.25074 | 0.004219 | YES | NO | NA |
| Prr5 | ENSMUSG( | ENSMUSG( | 1.197084 | 6.73E-08 | YES | YES | Upregulated |
| Lnpk | ENSMUSG( | ENSMUSG( | 0.188351 | 0.006403 | YES | NO | NA |
| Tspyl3 | ENSMUSG( | ENSMUSG( | -0.20829 | 0.007511 | YES | NO | NA |
| Rhot1 | ENSMUSG( | ENSMUSG( | -0.0131 | 0.86993 | NO | NO | NA |
| Tmem167l | ENSMUSG( | ENSMUSG( | -0.2196 | 0.013858 | YES | NO | NA |
| Gen1 | ENSMUSG( | ENSMUSG( | 0.134964 | 0.201052 | NO | NO | NA |
| Npc1 | ENSMUSG( | ENSMUSG( | -0.18447 | 0.044064 | YES | NO | NA |
| Ccdc34 | ENSMUSG( | ENSMUSG( | 0.281979 | 0.062492 | NO | NO | NA |
| Ndufa7 | ENSMUSG( | ENSMUSG( | 0.156948 | 0.035788 | YES | NO | NA |
| Esf1 | ENSMUSG( | ENSMUSG( | 0.163815 | 0.202451 | NO | NO | NA |
| Polr3f | ENSMUSG( | ENSMUSG( | -0.02341 | 0.816357 | NO | NO | NA |
| Klhl28 | ENSMUSG( | ENSMUSG( | -0.60123 | 4.01E-08 | YES | NO | NA |
| Tmem184l | ENSMUSG( | ENSMUSG( | -0.53687 | 3.54E-11 | YES | NO | NA |
| Cstf2t | ENSMUSG( | ENSMUSG( | -0.15865 | 0.047442 | YES | NO | NA |
| Armc8 | ENSMUSG( | ENSMUSG( | -0.16882 | 0.002376 | YES | NO | NA |
| Zdhhc16 | ENSMUSG( | ENSMUSG( | 0.067918 | 0.324564 | NO | NO | NA |
| Ing4 | ENSMUSG( | ENSMUSG( | -0.32502 | 0.000713 | YES | NO | NA |
| Akirin2 | ENSMUSG( | ENSMUSG( | -0.23039 | 0.002228 | YES | NO | NA |
| Gm42715 | ENSMUSG( | ENSMUSG( | 0.071973 | 0.616932 | NO | NO | NA |
| Mrpl43 | ENSMUSG( | ENSMUSG( | 0.092781 | 0.236838 | NO | NO | NA |
| Tada1 | ENSMUSG( | ENSMUSG( | -0.15163 | 0.003439 | YES | NO | NA |
| Pts | ENSMUSG( | ENSMUSG( | 0.010932 | 0.953526 | NO | NO | NA |
| Mrps7 | ENSMUSG( | ENSMUSG( | 0.229069 | 0.001554 | YES | NO | NA |
| Pwwp2a | ENSMUSG( | ENSMUSG( | -0.29588 | 3.00E-05 | YES | NO | NA |
| Aen | ENSMUSG( | ENSMUSG( | 0.187907 | 0.004991 | YES | NO | NA |
| Ddx55 | ENSMUSG( | ENSMUSG( | -0.13855 | 0.079173 | NO | NO | NA |

| | | | | | | | |
|---|---|---|---|---|---|---|---|
| Mis18a | ENSMUSG( | ENSMUSG( | -0.07078 | 0.26752 | NO | NO | NA |
| Hmgxb3 | ENSMUSG( | ENSMUSG( | -0.20528 | 0.004452 | YES | NO | NA |
| Spry2 | ENSMUSG( | ENSMUSG( | 0.012251 | 0.961918 | NO | NO | NA |
| Ppcdc | ENSMUSG( | ENSMUSG( | -0.08375 | 0.366356 | NO | NO | NA |
| Knop1 | ENSMUSG( | ENSMUSG( | 0.182915 | 0.00332 | YES | NO | NA |
| Emc2 | ENSMUSG( | ENSMUSG( | 0.114936 | 0.312678 | NO | NO | NA |
| Comt | ENSMUSG( | ENSMUSG( | 0.385027 | 6.31E-05 | YES | NO | NA |
| Tmx2 | ENSMUSG( | ENSMUSG( | -0.04901 | 0.508373 | NO | NO | NA |
| Mob1b | ENSMUSG( | ENSMUSG( | -0.16986 | 0.079037 | NO | NO | NA |
| Gtf2h5 | ENSMUSG( | ENSMUSG( | 0.313521 | 0.001563 | YES | NO | NA |
| Auh | ENSMUSG( | ENSMUSG( | -0.19929 | 0.005789 | YES | NO | NA |
| Mettl9 | ENSMUSG( | ENSMUSG( | 0.331464 | 0.000131 | YES | NO | NA |
| Fmnl3 | ENSMUSG( | ENSMUSG( | 0.264189 | 0.001868 | YES | NO | NA |
| Pex19 | ENSMUSG( | ENSMUSG( | 0.04226 | 0.610637 | NO | NO | NA |
| ENSMUSG( | ENSMUSG( | ENSMUSG( | -0.59264 | 9.99E-07 | YES | NO | NA |
| Xrcc1 | ENSMUSG( | ENSMUSG( | -0.07405 | 0.307123 | NO | NO | NA |
| Hdac4 | ENSMUSG( | ENSMUSG( | -0.29303 | 0.002453 | YES | NO | NA |
| Letmd1 | ENSMUSG( | ENSMUSG( | -0.28044 | 8.07E-05 | YES | NO | NA |
| Fdft1 | ENSMUSG( | ENSMUSG( | 0.35457 | 0.001686 | YES | NO | NA |
| Cux1 | ENSMUSG( | ENSMUSG( | -0.02173 | 0.815712 | NO | NO | NA |
| Hpcal1 | ENSMUSG( | ENSMUSG( | -0.20304 | 0.041083 | YES | NO | NA |
| Urod | ENSMUSG( | ENSMUSG( | 0.155262 | 0.034548 | YES | NO | NA |
| Osbpl7 | ENSMUSG( | ENSMUSG( | -0.14418 | 0.037584 | YES | NO | NA |
| Ano8 | ENSMUSG( | ENSMUSG( | -0.2846 | 0.00311 | YES | NO | NA |
| Kat2b | ENSMUSG( | ENSMUSG( | -0.31708 | 0.000147 | YES | NO | NA |
| Mad2l2 | ENSMUSG( | ENSMUSG( | 1.136588 | 3.94E-23 | YES | YES | Upregulated |
| Utp11 | ENSMUSG( | ENSMUSG( | 0.082572 | 0.49485 | NO | NO | NA |
| Bcr | ENSMUSG( | ENSMUSG( | -0.14262 | 0.013832 | YES | NO | NA |
| Dhrs1 | ENSMUSG( | ENSMUSG( | -0.03965 | 0.706458 | NO | NO | NA |
| Rbm19 | ENSMUSG( | ENSMUSG( | 0.380532 | 0.000827 | YES | NO | NA |
| Troap | ENSMUSG( | ENSMUSG( | 0.294608 | 0.002766 | YES | NO | NA |
| Zfp266 | ENSMUSG( | ENSMUSG( | -0.18356 | 0.004105 | YES | NO | NA |
| Sufu | ENSMUSG( | ENSMUSG( | -0.08221 | 0.224995 | NO | NO | NA |
| Ypel5 | ENSMUSG( | ENSMUSG( | -0.29993 | 0.023797 | YES | NO | NA |
| Parn | ENSMUSG( | ENSMUSG( | 0.185503 | 0.026368 | YES | NO | NA |
| Whamm | ENSMUSG( | ENSMUSG( | -0.26528 | 0.000266 | YES | NO | NA |
| Cdc42ep3 | ENSMUSG( | ENSMUSG( | -0.61381 | 6.16E-18 | YES | NO | NA |
| Bsdc1 | ENSMUSG( | ENSMUSG( | -0.44481 | 4.36E-09 | YES | NO | NA |
| Acd | ENSMUSG( | ENSMUSG( | 0.130437 | 0.053236 | NO | NO | NA |
| Frmd4b | ENSMUSG( | ENSMUSG( | -0.28009 | 0.060243 | NO | NO | NA |
| Traip | ENSMUSG( | ENSMUSG( | 0.7778 | 2.73E-13 | YES | NO | NA |
| Abcc1 | ENSMUSG( | ENSMUSG( | 0.219622 | 0.000377 | YES | NO | NA |
| Kctd18 | ENSMUSG( | ENSMUSG( | -0.12398 | 0.3727 | NO | NO | NA |
| Ccdc71 | ENSMUSG( | ENSMUSG( | -0.17752 | 0.002137 | YES | NO | NA |
| Clstn1 | ENSMUSG( | ENSMUSG( | -0.22894 | 0.00041 | YES | NO | NA |
| Cox15 | ENSMUSG( | ENSMUSG( | 0.022158 | 0.784434 | NO | NO | NA |
| Msrb1 | ENSMUSG( | ENSMUSG( | 0.229722 | 0.02414 | YES | NO | NA |

| | | | | | | | |
|---|---|---|---|---|---|---|---|
| Ing3 | ENSMUSG( | ENSMUSG( | -0.02129 | 0.857106 | NO | NO | NA |
| Hras | ENSMUSG( | ENSMUSG( | 0.118653 | 0.016898 | YES | NO | NA |
| Krba1 | ENSMUSG( | ENSMUSG( | 0.258688 | 0.054682 | NO | NO | NA |
| Plcxd2 | ENSMUSG( | ENSMUSG( | -0.41176 | 4.35E-05 | YES | NO | NA |
| Ormdl1 | ENSMUSG( | ENSMUSG( | -0.42316 | 6.61E-07 | YES | NO | NA |
| Chmp6 | ENSMUSG( | ENSMUSG( | 0.090574 | 0.419047 | NO | NO | NA |
| Pcmtd2 | ENSMUSG( | ENSMUSG( | -0.50635 | 4.67E-06 | YES | NO | NA |
| Polr1c | ENSMUSG( | ENSMUSG( | 0.201735 | 0.026436 | YES | NO | NA |
| Smarcad1 | ENSMUSG( | ENSMUSG( | 0.037738 | 0.620441 | NO | NO | NA |
| Zfp24 | ENSMUSG( | ENSMUSG( | 0.007059 | 0.948326 | NO | NO | NA |
| Mark4 | ENSMUSG( | ENSMUSG( | -0.31208 | 0.001004 | YES | NO | NA |
| Esrra | ENSMUSG( | ENSMUSG( | -0.42056 | 1.66E-09 | YES | NO | NA |
| Cnst | ENSMUSG( | ENSMUSG( | -0.32254 | 6.48E-08 | YES | NO | NA |
| A630001G | ENSMUSG( | ENSMUSG( | -0.08125 | 0.583866 | NO | NO | NA |
| Strbp | ENSMUSG( | ENSMUSG( | 0.227957 | 0.086033 | NO | NO | NA |
| Fam50a | ENSMUSG( | ENSMUSG( | -0.08565 | 0.547207 | NO | NO | NA |
| Tmem201 | ENSMUSG( | ENSMUSG( | 0.079764 | 0.411282 | NO | NO | NA |
| Afmid | ENSMUSG( | ENSMUSG( | -0.31336 | 9.60E-05 | YES | NO | NA |
| Sfn | ENSMUSG( | ENSMUSG( | 0.023402 | 0.862571 | NO | NO | NA |
| Ppox | ENSMUSG( | ENSMUSG( | -0.10641 | 0.262918 | NO | NO | NA |
| Papss1 | ENSMUSG( | ENSMUSG( | 0.070563 | 0.329729 | NO | NO | NA |
| Ndufa6 | ENSMUSG( | ENSMUSG( | 0.011531 | 0.905904 | NO | NO | NA |
| Dynlt1b | ENSMUSG( | ENSMUSG( | 0.741667 | 1.90E-29 | YES | NO | NA |
| Pdrg1 | ENSMUSG( | ENSMUSG( | -0.25001 | 0.001617 | YES | NO | NA |
| Trim30d | ENSMUSG( | ENSMUSG( | -1.16596 | 7.49E-13 | YES | YES | Downregulated |
| Mphosph1 | ENSMUSG( | ENSMUSG( | 0.178058 | 0.286895 | NO | NO | NA |
| Spopl | ENSMUSG( | ENSMUSG( | 0.638205 | 3.24E-32 | YES | NO | NA |
| Epg5 | ENSMUSG( | ENSMUSG( | -0.26947 | 0.001402 | YES | NO | NA |
| Clcn6 | ENSMUSG( | ENSMUSG( | 0.896422 | 3.23E-32 | YES | NO | NA |
| Usp11 | ENSMUSG( | ENSMUSG( | -0.18605 | 0.065923 | NO | NO | NA |
| Ung | ENSMUSG( | ENSMUSG( | 0.474649 | 1.88E-06 | YES | NO | NA |
| Klf16 | ENSMUSG( | ENSMUSG( | -0.11225 | 0.286533 | NO | NO | NA |
| Babam2 | ENSMUSG( | ENSMUSG( | 0.064984 | 0.470463 | NO | NO | NA |
| Ndc1 | ENSMUSG( | ENSMUSG( | 0.406056 | 9.05E-06 | YES | NO | NA |
| Ddb2 | ENSMUSG( | ENSMUSG( | -0.13451 | 0.17992 | NO | NO | NA |
| Gm15500 | ENSMUSG( | ENSMUSG( | 0.584374 | 0.013063 | YES | NO | NA |
| Rab24 | ENSMUSG( | ENSMUSG( | -0.00585 | 0.949307 | NO | NO | NA |
| Smim7 | ENSMUSG( | ENSMUSG( | -0.03662 | 0.665935 | NO | NO | NA |
| Gyg | ENSMUSG( | ENSMUSG( | -0.0536 | 0.476823 | NO | NO | NA |
| Gucd1 | ENSMUSG( | ENSMUSG( | -0.28307 | 0.000815 | YES | NO | NA |
| Ttk | ENSMUSG( | ENSMUSG( | 0.117775 | 0.22488 | NO | NO | NA |
| Fdps | ENSMUSG( | ENSMUSG( | 0.722042 | 1.87E-06 | YES | NO | NA |
| Spdl1 | ENSMUSG( | ENSMUSG( | 0.346143 | 0.003552 | YES | NO | NA |
| Dok1 | ENSMUSG( | ENSMUSG( | -0.07584 | 0.474169 | NO | NO | NA |
| AI837181 | ENSMUSG( | ENSMUSG( | -0.29829 | 2.54E-05 | YES | NO | NA |
| Tarbp2 | ENSMUSG( | ENSMUSG( | 0.129685 | 0.009853 | YES | NO | NA |
| Gbp5 | ENSMUSG( | ENSMUSG( | -0.46204 | 4.36E-05 | YES | NO | NA |

| | | | | | | | |
|---|---|---|---|---|---|---|---|
| Cstf1 | ENSMUSG( | ENSMUSG( | 0.04873 | 0.415514 | NO | NO | NA |
| Cisd3 | ENSMUSG( | ENSMUSG( | 0.285433 | 2.13E-06 | YES | NO | NA |
| Arhgap33 | ENSMUSG( | ENSMUSG( | 0.382054 | 0.000212 | YES | NO | NA |
| Itprip | ENSMUSG( | ENSMUSG( | -0.12844 | 0.063735 | NO | NO | NA |
| Lamtor5 | ENSMUSG( | ENSMUSG( | -0.00155 | 0.982586 | NO | NO | NA |
| Vps26a | ENSMUSG( | ENSMUSG( | -0.19308 | 0.034488 | YES | NO | NA |
| Slc43a2 | ENSMUSG( | ENSMUSG( | -0.4646 | 4.26E-07 | YES | NO | NA |
| Orc4 | ENSMUSG( | ENSMUSG( | -0.04668 | 0.527544 | NO | NO | NA |
| Sipa1l3 | ENSMUSG( | ENSMUSG( | -0.31562 | 7.55E-06 | YES | NO | NA |
| Tmem168 | ENSMUSG( | ENSMUSG( | -0.08626 | 0.328216 | NO | NO | NA |
| Mrps5 | ENSMUSG( | ENSMUSG( | 0.352694 | 0.003127 | YES | NO | NA |
| Ids | ENSMUSG( | ENSMUSG( | -0.17572 | 0.187457 | NO | NO | NA |
| Mcur1 | ENSMUSG( | ENSMUSG( | 0.141048 | 0.149758 | NO | NO | NA |
| C1d | ENSMUSG( | ENSMUSG( | 0.246175 | 0.006963 | YES | NO | NA |
| Ppie | ENSMUSG( | ENSMUSG( | 0.288664 | 0.001784 | YES | NO | NA |
| Tspan31 | ENSMUSG( | ENSMUSG( | 0.889427 | 1.93E-07 | YES | NO | NA |
| Hcst | ENSMUSG( | ENSMUSG( | -0.2572 | 0.031908 | YES | NO | NA |
| Mrpl30 | ENSMUSG( | ENSMUSG( | -0.00819 | 0.934051 | NO | NO | NA |
| Nxpe3 | ENSMUSG( | ENSMUSG( | -0.26655 | 1.46E-08 | YES | NO | NA |
| Wdr77 | ENSMUSG( | ENSMUSG( | -0.03731 | 0.575381 | NO | NO | NA |
| Mfsd5 | ENSMUSG( | ENSMUSG( | -0.1225 | 0.001907 | YES | NO | NA |
| Sh3bp2 | ENSMUSG( | ENSMUSG( | 0.088735 | 0.512637 | NO | NO | NA |
| Nop10 | ENSMUSG( | ENSMUSG( | 0.364943 | 0.000887 | YES | NO | NA |
| Ago3 | ENSMUSG( | ENSMUSG( | -0.39526 | 8.46E-05 | YES | NO | NA |
| Cln3 | ENSMUSG( | ENSMUSG( | 0.106791 | 0.372606 | NO | NO | NA |
| Rnmt | ENSMUSG( | ENSMUSG( | 0.173997 | 0.007988 | YES | NO | NA |
| Rusc1 | ENSMUSG( | ENSMUSG( | -0.13691 | 0.037487 | YES | NO | NA |
| Ubfd1 | ENSMUSG( | ENSMUSG( | -0.28416 | 2.98E-06 | YES | NO | NA |
| Hspa1a | ENSMUSG( | ENSMUSG( | -2.67039 | 0.007052 | YES | YES | Downregulated |
| Kctd17 | ENSMUSG( | ENSMUSG( | 0.404861 | 0.002395 | YES | NO | NA |
| Depdc1b | ENSMUSG( | ENSMUSG( | 0.101815 | 0.13582 | NO | NO | NA |
| Trub2 | ENSMUSG( | ENSMUSG( | -0.21148 | 0.06989 | NO | NO | NA |
| Cnot4 | ENSMUSG( | ENSMUSG( | -0.41906 | 5.10E-06 | YES | NO | NA |
| Eapp | ENSMUSG( | ENSMUSG( | -0.11394 | 0.145292 | NO | NO | NA |
| Ess2 | ENSMUSG( | ENSMUSG( | 0.074094 | 0.443843 | NO | NO | NA |
| Stard3nl | ENSMUSG( | ENSMUSG( | -0.21099 | 0.006219 | YES | NO | NA |
| Tdp1 | ENSMUSG( | ENSMUSG( | 0.109199 | 0.383665 | NO | NO | NA |
| Ahsa2 | ENSMUSG( | ENSMUSG( | -0.28332 | 0.000123 | YES | NO | NA |
| Phlda1 | ENSMUSG( | ENSMUSG( | 0.261053 | 0.213047 | NO | NO | NA |
| Septin2 EN | ENSMUSG( | ENSMUSG( | 0.084441 | 0.726446 | NO | NO | NA |
| Ddhd2 | ENSMUSG( | ENSMUSG( | 0.060941 | 0.446235 | NO | NO | NA |
| ENSMUSG | ENSMUSG( | ENSMUSG( | 0.322029 | 0.04391 | YES | NO | NA |
| Pex6 | ENSMUSG( | ENSMUSG( | -0.58492 | 9.88E-07 | YES | NO | NA |
| AB124611 | ENSMUSG( | ENSMUSG( | -0.14872 | 0.243906 | NO | NO | NA |
| Pcgf5 | ENSMUSG( | ENSMUSG( | -0.11793 | 0.011907 | YES | NO | NA |
| 2310022A | ENSMUSG( | ENSMUSG( | 0.041168 | 0.545331 | NO | NO | NA |
| Abcf3 | ENSMUSG( | ENSMUSG( | -0.00095 | 0.991918 | NO | NO | NA |

| | | | | | | | |
|---|---|---|---|---|---|---|---|
| Hmces | ENSMUSG( | ENSMUSG( | -0.02356 | 0.84357 | NO | NO | NA |
| Papolg | ENSMUSG( | ENSMUSG( | -0.44692 | 3.35E-07 | YES | NO | NA |
| Rufy2 | ENSMUSG( | ENSMUSG( | -0.22563 | 0.07543 | NO | NO | NA |
| Med8 | ENSMUSG( | ENSMUSG( | -0.01216 | 0.906481 | NO | NO | NA |
| Rbm43 | ENSMUSG( | ENSMUSG( | -0.15551 | 0.039582 | YES | NO | NA |
| Smg9 | ENSMUSG( | ENSMUSG( | 0.046765 | 0.475214 | NO | NO | NA |
| Pop4 | ENSMUSG( | ENSMUSG( | 0.180982 | 0.008662 | YES | NO | NA |
| Asb1 | ENSMUSG( | ENSMUSG( | -0.27748 | 0.000761 | YES | NO | NA |
| Slc4a8 | ENSMUSG( | ENSMUSG( | -0.34348 | 3.22E-05 | YES | NO | NA |
| Apobr | ENSMUSG( | ENSMUSG( | 0.137247 | 0.575033 | NO | NO | NA |
| Gna11 | ENSMUSG( | ENSMUSG( | -0.36532 | 2.13E-07 | YES | NO | NA |
| Slc41a1 | ENSMUSG( | ENSMUSG( | -0.48017 | 0.000102 | YES | NO | NA |
| Lamtor4 | ENSMUSG( | ENSMUSG( | -0.0773 | 0.484647 | NO | NO | NA |
| Cog4 | ENSMUSG( | ENSMUSG( | -0.10576 | 0.120341 | NO | NO | NA |
| Slc25a45 | ENSMUSG( | ENSMUSG( | -0.49161 | 3.27E-07 | YES | NO | NA |
| Zgrf1 | ENSMUSG( | ENSMUSG( | 0.198091 | 0.076213 | NO | NO | NA |
| Tbp | ENSMUSG( | ENSMUSG( | -0.02927 | 0.713566 | NO | NO | NA |
| Dnpep | ENSMUSG( | ENSMUSG( | 0.006326 | 0.961546 | NO | NO | NA |
| Lsm3 | ENSMUSG( | ENSMUSG( | 0.153606 | 0.085856 | NO | NO | NA |
| Lrp6 | ENSMUSG( | ENSMUSG( | -0.11782 | 0.105785 | NO | NO | NA |
| Rfc4 | ENSMUSG( | ENSMUSG( | 0.38228 | 0.000169 | YES | NO | NA |
| Romo1 | ENSMUSG( | ENSMUSG( | 0.566886 | 0.000101 | YES | NO | NA |
| Mios | ENSMUSG( | ENSMUSG( | 0.221349 | 0.01943 | YES | NO | NA |
| Cep164 | ENSMUSG( | ENSMUSG( | 0.016734 | 0.91234 | NO | NO | NA |
| Zfp281 | ENSMUSG( | ENSMUSG( | -0.29781 | 4.35E-05 | YES | NO | NA |
| Lin54 | ENSMUSG( | ENSMUSG( | 0.013826 | 0.874442 | NO | NO | NA |
| Kdm1b | ENSMUSG( | ENSMUSG( | -0.28618 | 0.000728 | YES | NO | NA |
| Plekhf2 | ENSMUSG( | ENSMUSG( | -0.13973 | 0.184585 | NO | NO | NA |
| Mea1 | ENSMUSG( | ENSMUSG( | -0.17267 | 0.011796 | YES | NO | NA |
| Phlpp1 | ENSMUSG( | ENSMUSG( | 0.239656 | 0.020103 | YES | NO | NA |
| Eif2b4 | ENSMUSG( | ENSMUSG( | 0.111397 | 0.135294 | NO | NO | NA |
| Rhbdf2 | ENSMUSG( | ENSMUSG( | 0.407633 | 8.07E-05 | YES | NO | NA |
| Vps37a | ENSMUSG( | ENSMUSG( | -0.34802 | 7.05E-05 | YES | NO | NA |
| Nck2 | ENSMUSG( | ENSMUSG( | -0.24509 | 0.002715 | YES | NO | NA |
| Cib1 | ENSMUSG( | ENSMUSG( | -0.01713 | 0.896865 | NO | NO | NA |
| Dhx40 | ENSMUSG( | ENSMUSG( | 0.122875 | 0.133858 | NO | NO | NA |
| Tcp11l2 | ENSMUSG( | ENSMUSG( | -0.89854 | 2.97E-09 | YES | NO | NA |
| Ercc4 | ENSMUSG( | ENSMUSG( | -0.06605 | 0.484899 | NO | NO | NA |
| Pno1 | ENSMUSG( | ENSMUSG( | 0.651283 | 2.00E-06 | YES | NO | NA |
| Psmd9 | ENSMUSG( | ENSMUSG( | 0.382005 | 3.67E-07 | YES | NO | NA |
| Zbtb37 | ENSMUSG( | ENSMUSG( | -0.62831 | 8.21E-07 | YES | NO | NA |
| Paqr7 | ENSMUSG( | ENSMUSG( | -0.01672 | 0.910986 | NO | NO | NA |
| Ccdc115 | ENSMUSG( | ENSMUSG( | 0.342224 | 7.53E-05 | YES | NO | NA |
| Psmd5 | ENSMUSG( | ENSMUSG( | -0.00351 | 0.965837 | NO | NO | NA |
| Mettl16 | ENSMUSG( | ENSMUSG( | 0.189085 | 0.074581 | NO | NO | NA |
| Ubxn6 | ENSMUSG( | ENSMUSG( | -0.16089 | 0.178813 | NO | NO | NA |
| Polm | ENSMUSG( | ENSMUSG( | -0.00709 | 0.9574 | NO | NO | NA |

| | | | | | | | |
|---|---|---|---|---|---|---|---|
| Nfatc2ip | ENSMUSG( | ENSMUSG( | -0.05307 | 0.452849 | NO | NO | NA |
| Nt5c3 | ENSMUSG( | ENSMUSG( | -0.25686 | 0.000437 | YES | NO | NA |
| Gm39323 | ENSMUSG( | ENSMUSG( | -0.38032 | 0.001739 | YES | NO | NA |
| Sertad2 | ENSMUSG( | ENSMUSG( | -0.6718 | 1.16E-08 | YES | NO | NA |
| Trbv13-1 | ENSMUSG( | ENSMUSG( | -0.38228 | 0.434211 | NO | NO | NA |
| Txndc12 | ENSMUSG( | ENSMUSG( | -0.0881 | 0.307944 | NO | NO | NA |
| ENSMUSG( | ENSMUSG( | ENSMUSG( | -0.39768 | 8.17E-07 | YES | NO | NA |
| Zfp26 | ENSMUSG( | ENSMUSG( | -0.16169 | 0.101725 | NO | NO | NA |
| Mrpl19 | ENSMUSG( | ENSMUSG( | 0.258159 | 0.000786 | YES | NO | NA |
| Phf11c | ENSMUSG( | ENSMUSG( | -0.53822 | 1.17E-07 | YES | NO | NA |
| Kctd5 | ENSMUSG( | ENSMUSG( | 0.067932 | 0.310431 | NO | NO | NA |
| Tfeb | ENSMUSG( | ENSMUSG( | 0.071867 | 0.435436 | NO | NO | NA |
| H2-T24 | ENSMUSG( | ENSMUSG( | -1.27634 | 9.88E-28 | YES | YES | Downregulated |
| Dnajb9 | ENSMUSG( | ENSMUSG( | -0.53857 | 0.00038 | YES | NO | NA |
| Gipc1 | ENSMUSG( | ENSMUSG( | 0.209437 | 0.031407 | YES | NO | NA |
| Pbx2 | ENSMUSG( | ENSMUSG( | -0.30895 | 4.67E-07 | YES | NO | NA |
| Cops9 | ENSMUSG( | ENSMUSG( | -0.02737 | 0.803298 | NO | NO | NA |
| Pxylp1 | ENSMUSG( | ENSMUSG( | 0.055454 | 0.640929 | NO | NO | NA |
| Xlr4b | ENSMUSG( | ENSMUSG( | -0.08891 | 0.548849 | NO | NO | NA |
| Ubac1 | ENSMUSG( | ENSMUSG( | 0.126158 | 0.022036 | YES | NO | NA |
| 1500011B( | ENSMUSG( | ENSMUSG( | 0.401992 | 9.37E-05 | YES | NO | NA |
| Zbtb22 | ENSMUSG( | ENSMUSG( | 0.064441 | 0.429171 | NO | NO | NA |
| Flad1 | ENSMUSG( | ENSMUSG( | -0.21399 | 0.007975 | YES | NO | NA |
| Htatip2 | ENSMUSG( | ENSMUSG( | 0.124162 | 0.233033 | NO | NO | NA |
| Uqcr11 | ENSMUSG( | ENSMUSG( | 0.362182 | 0.000682 | YES | NO | NA |
| Pank4 | ENSMUSG( | ENSMUSG( | 0.020544 | 0.770606 | NO | NO | NA |
| Ufl1 | ENSMUSG( | ENSMUSG( | 0.012664 | 0.89964 | NO | NO | NA |
| Tnks1bp1 | ENSMUSG( | ENSMUSG( | -0.23923 | 0.011225 | YES | NO | NA |
| Anapc15 | ENSMUSG( | ENSMUSG( | 0.450386 | 7.82E-08 | YES | NO | NA |
| Myadm | ENSMUSG( | ENSMUSG( | -0.74811 | 1.07E-11 | YES | NO | NA |
| Uba5 | ENSMUSG( | ENSMUSG( | -0.17807 | 0.0105 | YES | NO | NA |
| Trappc9 | ENSMUSG( | ENSMUSG( | 0.027995 | 0.801851 | NO | NO | NA |
| Gtf3c5 | ENSMUSG( | ENSMUSG( | 0.171633 | 0.012201 | YES | NO | NA |
| Slc25a44 | ENSMUSG( | ENSMUSG( | -0.20594 | 0.006622 | YES | NO | NA |
| Mrpl20 | ENSMUSG( | ENSMUSG( | 0.498184 | 1.42E-08 | YES | NO | NA |
| Rras2 | ENSMUSG( | ENSMUSG( | -0.20386 | 0.270825 | NO | NO | NA |
| Paip2b | ENSMUSG( | ENSMUSG( | -0.03044 | 0.689596 | NO | NO | NA |
| Mrpl13 | ENSMUSG( | ENSMUSG( | 0.243304 | 0.004018 | YES | NO | NA |
| Trappc4 | ENSMUSG( | ENSMUSG( | -0.04423 | 0.637771 | NO | NO | NA |
| Morf4l1-ps | ENSMUSG( | ENSMUSG( | -0.02809 | 0.801815 | NO | NO | NA |
| Kif16b | ENSMUSG( | ENSMUSG( | -0.01181 | 0.915968 | NO | NO | NA |
| Med20 | ENSMUSG( | ENSMUSG( | -0.07116 | 0.284137 | NO | NO | NA |
| Clcc1 | ENSMUSG( | ENSMUSG( | 0.091996 | 0.366592 | NO | NO | NA |
| Itpripl1 | ENSMUSG( | ENSMUSG( | -0.03117 | 0.624402 | NO | NO | NA |
| Etaa1 | ENSMUSG( | ENSMUSG( | -0.13999 | 0.219649 | NO | NO | NA |
| Gne | ENSMUSG( | ENSMUSG( | 0.039002 | 0.531711 | NO | NO | NA |
| Aarsd1 | ENSMUSG( | ENSMUSG( | 0.284652 | 0.003117 | YES | NO | NA |

| | | | | | | | |
|---|---|---|---|---|---|---|---|
| Echs1 | ENSMUSG( | ENSMUSG( | 0.330699 | 0.00025 | YES | NO | NA |
| Zfp41 | ENSMUSG( | ENSMUSG( | -0.54485 | 8.72E-07 | YES | NO | NA |
| Pcid2 | ENSMUSG( | ENSMUSG( | -0.14975 | 0.003077 | YES | NO | NA |
| Uchl3 | ENSMUSG( | ENSMUSG( | 0.251564 | 0.032303 | YES | NO | NA |
| Dcaf10 | ENSMUSG( | ENSMUSG( | -0.16333 | 0.003773 | YES | NO | NA |
| Mrpl54 | ENSMUSG( | ENSMUSG( | 0.538871 | 1.53E-05 | YES | NO | NA |
| Tmem63a | ENSMUSG( | ENSMUSG( | -0.24841 | 0.000253 | YES | NO | NA |
| Hinfp | ENSMUSG( | ENSMUSG( | -0.07658 | 0.344904 | NO | NO | NA |
| Slc22a15 | ENSMUSG( | ENSMUSG( | -0.06436 | 0.6204 | NO | NO | NA |
| Mcoln1 | ENSMUSG( | ENSMUSG( | -0.12603 | 0.398144 | NO | NO | NA |
| Ttc37 | ENSMUSG( | ENSMUSG( | 0.124119 | 0.103215 | NO | NO | NA |
| Mtif2 | ENSMUSG( | ENSMUSG( | 0.070709 | 0.518102 | NO | NO | NA |
| Maco1 | ENSMUSG( | ENSMUSG( | 0.06565 | 0.646928 | NO | NO | NA |
| Mettl3 | ENSMUSG( | ENSMUSG( | -0.22656 | 0.001272 | YES | NO | NA |
| Fto | ENSMUSG( | ENSMUSG( | 0.028694 | 0.732447 | NO | NO | NA |
| Uba6 | ENSMUSG( | ENSMUSG( | -0.2361 | 1.24E-05 | YES | NO | NA |
| Wdr83os | ENSMUSG( | ENSMUSG( | 0.140584 | 0.158327 | NO | NO | NA |
| Uvssa | ENSMUSG( | ENSMUSG( | 0.415131 | 0.001372 | YES | NO | NA |
| Polr1f | ENSMUSG( | ENSMUSG( | 0.074222 | 0.391427 | NO | NO | NA |
| Pnpt1 | ENSMUSG( | ENSMUSG( | 0.069214 | 0.51558 | NO | NO | NA |
| Mbtd1 | ENSMUSG( | ENSMUSG( | -0.01482 | 0.884746 | NO | NO | NA |
| Cfap20 | ENSMUSG( | ENSMUSG( | 0.478383 | 4.81E-07 | YES | NO | NA |
| Galnt3 | ENSMUSG( | ENSMUSG( | 0.619741 | 1.25E-06 | YES | NO | NA |
| Cyb561d1 | ENSMUSG( | ENSMUSG( | -0.43041 | 1.26E-05 | YES | NO | NA |
| Anks1 | ENSMUSG( | ENSMUSG( | -0.19141 | 0.048686 | YES | NO | NA |
| Eif2b1 | ENSMUSG( | ENSMUSG( | 0.349071 | 3.58E-05 | YES | NO | NA |
| Uap1 | ENSMUSG( | ENSMUSG( | -0.00871 | 0.889318 | NO | NO | NA |
| Marcksl1 | ENSMUSG( | ENSMUSG( | 1.19191 | 3.38E-10 | YES | YES | Upregulated |
| Mfn1 | ENSMUSG( | ENSMUSG( | 0.213936 | 0.032389 | YES | NO | NA |
| Atp6v0b | ENSMUSG( | ENSMUSG( | -0.17378 | 0.022342 | YES | NO | NA |
| Plxnc1 | ENSMUSG( | ENSMUSG( | 0.505034 | 0.002395 | YES | NO | NA |
| Flcn | ENSMUSG( | ENSMUSG( | -0.392 | 1.30E-05 | YES | NO | NA |
| Slc30a7 | ENSMUSG( | ENSMUSG( | -0.34813 | 1.37E-07 | YES | NO | NA |
| Nlk | ENSMUSG( | ENSMUSG( | -0.34909 | 3.50E-06 | YES | NO | NA |
| Dsn1 | ENSMUSG( | ENSMUSG( | 0.193458 | 0.089784 | NO | NO | NA |
| Ttc1 | ENSMUSG( | ENSMUSG( | 0.092382 | 0.33848 | NO | NO | NA |
| Ppm1f | ENSMUSG( | ENSMUSG( | 0.12612 | 0.077328 | NO | NO | NA |
| Smurf1 | ENSMUSG( | ENSMUSG( | -0.17491 | 0.176807 | NO | NO | NA |
| Akap10 | ENSMUSG( | ENSMUSG( | -0.24516 | 0.000874 | YES | NO | NA |
| Nudcd2 | ENSMUSG( | ENSMUSG( | 0.521549 | 6.10E-08 | YES | NO | NA |
| Camk2n1 | ENSMUSG( | ENSMUSG( | -1.05941 | 1.78E-09 | YES | YES | Downregulated |
| Spc25 | ENSMUSG( | ENSMUSG( | 0.018648 | 0.860844 | NO | NO | NA |
| Gm17334 | ENSMUSG( | ENSMUSG( | 0.164206 | 0.384723 | NO | NO | NA |
| Mrps18c | ENSMUSG( | ENSMUSG( | 0.125083 | 0.401609 | NO | NO | NA |
| Zup1 | ENSMUSG( | ENSMUSG( | -0.52698 | 7.26E-11 | YES | NO | NA |
| Gm50230 | ENSMUSG( | ENSMUSG( | -0.34437 | 0.01269 | YES | NO | NA |
| Ikzf5 | ENSMUSG( | ENSMUSG( | -0.25836 | 0.000724 | YES | NO | NA |

| | | | | | | | |
|---|---|---|---|---|---|---|---|
| Ctu2 | ENSMUSG( | ENSMUSG( | 0.116552 | 0.119248 | NO | NO | NA |
| Snapin | ENSMUSG( | ENSMUSG( | -0.14343 | 0.320524 | NO | NO | NA |
| Mrpl27 | ENSMUSG( | ENSMUSG( | 0.517611 | 2.73E-07 | YES | NO | NA |
| Klhl25 | ENSMUSG( | ENSMUSG( | -0.40509 | 6.54E-06 | YES | NO | NA |
| Cep128 | ENSMUSG( | ENSMUSG( | 0.116975 | 0.392719 | NO | NO | NA |
| Fech | ENSMUSG( | ENSMUSG( | 0.65255 | 3.87E-06 | YES | NO | NA |
| Snapc4 | ENSMUSG( | ENSMUSG( | 0.079553 | 0.298258 | NO | NO | NA |
| Nfil3 | ENSMUSG( | ENSMUSG( | 0.049405 | 0.803364 | NO | NO | NA |
| Mt2 | ENSMUSG( | ENSMUSG( | 2.20134 | 3.56E-10 | YES | YES | Upregulated |
| Ythdc2 | ENSMUSG( | ENSMUSG( | -0.24275 | 0.001847 | YES | NO | NA |
| Zfp110 | ENSMUSG( | ENSMUSG( | -0.23862 | 0.007104 | YES | NO | NA |
| Rrp9 | ENSMUSG( | ENSMUSG( | 0.273713 | 0.005847 | YES | NO | NA |
| Glp1r | ENSMUSG( | ENSMUSG( | -1.10357 | 0.002533 | YES | YES | Downregulated |
| Cemip2 | ENSMUSG( | ENSMUSG( | -0.27992 | 0.001785 | YES | NO | NA |
| Tnfrsf1a | ENSMUSG( | ENSMUSG( | -0.50786 | 1.91E-06 | YES | NO | NA |
| Map3k14 | ENSMUSG( | ENSMUSG( | -0.50718 | 4.00E-07 | YES | NO | NA |
| Rab10os | ENSMUSG( | ENSMUSG( | -0.12749 | 0.123095 | NO | NO | NA |
| Zwilch | ENSMUSG( | ENSMUSG( | 0.430869 | 4.06E-06 | YES | NO | NA |
| Irak2 | ENSMUSG( | ENSMUSG( | -0.4953 | 3.75E-05 | YES | NO | NA |
| Ip6k2 | ENSMUSG( | ENSMUSG( | -0.21815 | 0.0082 | YES | NO | NA |
| Ube2v2 | ENSMUSG( | ENSMUSG( | 0.13687 | 0.274188 | NO | NO | NA |
| Vwa8 | ENSMUSG( | ENSMUSG( | 0.324237 | 2.44E-05 | YES | NO | NA |
| Cul2 | ENSMUSG( | ENSMUSG( | 0.192897 | 0.013631 | YES | NO | NA |
| Sc5d | ENSMUSG( | ENSMUSG( | 0.286988 | 2.21E-05 | YES | NO | NA |
| Scpep1 | ENSMUSG( | ENSMUSG( | 0.488638 | 0.003179 | YES | NO | NA |
| Pwp2 | ENSMUSG( | ENSMUSG( | 0.261361 | 0.008222 | YES | NO | NA |
| Trbv14 | ENSMUSG( | ENSMUSG( | -0.82007 | 0.100329 | NO | NO | NA |
| Zkscan3 | ENSMUSG( | ENSMUSG( | -0.30371 | 2.59E-06 | YES | NO | NA |
| Noa1 | ENSMUSG( | ENSMUSG( | 0.214428 | 0.017067 | YES | NO | NA |
| Dffa | ENSMUSG( | ENSMUSG( | -0.03374 | 0.703762 | NO | NO | NA |
| Enpp1 | ENSMUSG( | ENSMUSG( | -0.5608 | 2.07E-08 | YES | NO | NA |
| Rspry1 | ENSMUSG( | ENSMUSG( | -0.10642 | 0.086155 | NO | NO | NA |
| Ccnc | ENSMUSG( | ENSMUSG( | -0.07287 | 0.309752 | NO | NO | NA |
| Pwp1 | ENSMUSG( | ENSMUSG( | 0.349597 | 7.02E-06 | YES | NO | NA |
| Aco1 | ENSMUSG( | ENSMUSG( | -0.10957 | 0.268831 | NO | NO | NA |
| Dgke | ENSMUSG( | ENSMUSG( | 0.040785 | 0.749342 | NO | NO | NA |
| Btbd10 | ENSMUSG( | ENSMUSG( | -0.21886 | 0.012937 | YES | NO | NA |
| Nsmf | ENSMUSG( | ENSMUSG( | -0.08523 | 0.49793 | NO | NO | NA |
| Tmx4 | ENSMUSG( | ENSMUSG( | 0.08885 | 0.539291 | NO | NO | NA |
| Eipr1 | ENSMUSG( | ENSMUSG( | -0.10895 | 0.279222 | NO | NO | NA |
| Cept1 | ENSMUSG( | ENSMUSG( | -0.18795 | 0.005508 | YES | NO | NA |
| Clpx | ENSMUSG( | ENSMUSG( | 0.225752 | 0.002651 | YES | NO | NA |
| Cyb561a3 | ENSMUSG( | ENSMUSG( | -0.4023 | 2.56E-06 | YES | NO | NA |
| Atad3a | ENSMUSG( | ENSMUSG( | 0.622989 | 8.35E-06 | YES | NO | NA |
| Dennd6a | ENSMUSG( | ENSMUSG( | -0.18261 | 0.063704 | NO | NO | NA |
| Ncoa7 | ENSMUSG( | ENSMUSG( | -0.28743 | 0.00149 | YES | NO | NA |
| Tysnd1 | ENSMUSG( | ENSMUSG( | 0.111657 | 0.128864 | NO | NO | NA |

| | | | | | | | |
|---|---|---|---|---|---|---|---|
| Pithd1 | ENSMUSG( | ENSMUSG( | 0.249968 | 2.42E-06 | YES | NO | NA |
| Rnasek | ENSMUSG( | ENSMUSG( | -0.33475 | 0.000822 | YES | NO | NA |
| Pnp | ENSMUSG( | ENSMUSG( | -0.04551 | 0.433475 | NO | NO | NA |
| Mrps21 | ENSMUSG( | ENSMUSG( | -0.04045 | 0.659108 | NO | NO | NA |
| Mrpl40 | ENSMUSG( | ENSMUSG( | 0.141755 | 0.070047 | NO | NO | NA |
| Fam104a | ENSMUSG( | ENSMUSG( | -0.11383 | 0.186757 | NO | NO | NA |
| Clcn7 | ENSMUSG( | ENSMUSG( | -0.20811 | 0.006529 | YES | NO | NA |
| Cars | ENSMUSG( | ENSMUSG( | 0.374082 | 1.27E-05 | YES | NO | NA |
| Plbd2 | ENSMUSG( | ENSMUSG( | 0.185468 | 0.02418 | YES | NO | NA |
| Wdr7 | ENSMUSG( | ENSMUSG( | -0.12028 | 0.105884 | NO | NO | NA |
| Osbpl11 | ENSMUSG( | ENSMUSG( | -0.16708 | 0.008851 | YES | NO | NA |
| Jaml | ENSMUSG( | ENSMUSG( | 0.075628 | 0.733043 | NO | NO | NA |
| Ptpre | ENSMUSG( | ENSMUSG( | -0.54013 | 8.19E-10 | YES | NO | NA |
| Fndc3a | ENSMUSG( | ENSMUSG( | -0.20669 | 0.067915 | NO | NO | NA |
| Atxn3 | ENSMUSG( | ENSMUSG( | -0.12557 | 0.136424 | NO | NO | NA |
| N4bp2l1 | ENSMUSG( | ENSMUSG( | -0.3052 | 0.028724 | YES | NO | NA |
| Nemp1 | ENSMUSG( | ENSMUSG( | 0.231226 | 0.013492 | YES | NO | NA |
| Abl2 | ENSMUSG( | ENSMUSG( | -0.20273 | 0.021709 | YES | NO | NA |
| Zbtb45 | ENSMUSG( | ENSMUSG( | -0.0453 | 0.46777 | NO | NO | NA |
| Kdelr2 | ENSMUSG( | ENSMUSG( | 0.038819 | 0.63616 | NO | NO | NA |
| Paip1 | ENSMUSG( | ENSMUSG( | -0.23413 | 7.33E-06 | YES | NO | NA |
| Lrpap1 | ENSMUSG( | ENSMUSG( | -0.07004 | 0.471485 | NO | NO | NA |
| Naa25 | ENSMUSG( | ENSMUSG( | 0.288662 | 0.017834 | YES | NO | NA |
| Ino80b | ENSMUSG( | ENSMUSG( | -0.03724 | 0.741786 | NO | NO | NA |
| Il3ra | ENSMUSG( | ENSMUSG( | 0.236456 | 0.060981 | NO | NO | NA |
| Cdkn2aip | ENSMUSG( | ENSMUSG( | -0.03638 | 0.749795 | NO | NO | NA |
| Slc25a19 | ENSMUSG( | ENSMUSG( | 0.104184 | 0.166928 | NO | NO | NA |
| Mrps24 | ENSMUSG( | ENSMUSG( | 0.150623 | 0.022697 | YES | NO | NA |
| Galk2 | ENSMUSG( | ENSMUSG( | -0.12992 | 0.067443 | NO | NO | NA |
| Nudt9 | ENSMUSG( | ENSMUSG( | 0.166651 | 0.047086 | YES | NO | NA |
| Rest | ENSMUSG( | ENSMUSG( | -0.13639 | 0.098564 | NO | NO | NA |
| Zfp143 | ENSMUSG( | ENSMUSG( | -0.26543 | 1.64E-05 | YES | NO | NA |
| H6pd | ENSMUSG( | ENSMUSG( | -0.26736 | 0.002788 | YES | NO | NA |
| Arhgap10 | ENSMUSG( | ENSMUSG( | -0.39917 | 1.37E-08 | YES | NO | NA |
| 2810004N | ENSMUSG( | ENSMUSG( | 0.174989 | 0.187178 | NO | NO | NA |
| Nfu1 | ENSMUSG( | ENSMUSG( | 0.182308 | 0.120491 | NO | NO | NA |
| Mrpl28 | ENSMUSG( | ENSMUSG( | 0.351055 | 0.002002 | YES | NO | NA |
| Ndufb6 | ENSMUSG( | ENSMUSG( | 0.354146 | 0.000152 | YES | NO | NA |
| Clec2i | ENSMUSG( | ENSMUSG( | -0.23227 | 0.01075 | YES | NO | NA |
| Smarcal1 | ENSMUSG( | ENSMUSG( | -0.05637 | 0.41526 | NO | NO | NA |
| Zfp7 | ENSMUSG( | ENSMUSG( | -0.29472 | 7.55E-05 | YES | NO | NA |
| Hsd17b10 | ENSMUSG( | ENSMUSG( | 0.099178 | 0.40141 | NO | NO | NA |
| Fam168a | ENSMUSG( | ENSMUSG( | -0.40449 | 4.28E-05 | YES | NO | NA |
| Neurl3 | ENSMUSG( | ENSMUSG( | -0.25825 | 0.030535 | YES | NO | NA |
| Mrpl48 | ENSMUSG( | ENSMUSG( | 0.238082 | 0.002477 | YES | NO | NA |
| Lrrc8b | ENSMUSG( | ENSMUSG( | -0.03997 | 0.663492 | NO | NO | NA |
| Rnf139 | ENSMUSG( | ENSMUSG( | -0.18076 | 0.016576 | YES | NO | NA |

| | | | | | | | |
|---|---|---|---|---|---|---|---|
| Glod4 | ENSMUSG( | ENSMUSG( | 0.126026 | 0.282568 | NO | NO | NA |
| Wdr18 | ENSMUSG( | ENSMUSG( | 0.436972 | 2.53E-07 | YES | NO | NA |
| Wee1 | ENSMUSG( | ENSMUSG( | -0.0552 | 0.531711 | NO | NO | NA |
| Slc35b3 | ENSMUSG( | ENSMUSG( | -0.06609 | 0.653854 | NO | NO | NA |
| Clp1 | ENSMUSG( | ENSMUSG( | 0.185577 | 0.000259 | YES | NO | NA |
| Med4 | ENSMUSG( | ENSMUSG( | 0.164704 | 0.063406 | NO | NO | NA |
| Taf1b | ENSMUSG( | ENSMUSG( | 0.017283 | 0.843013 | NO | NO | NA |
| Tmem260 | ENSMUSG( | ENSMUSG( | -0.14929 | 0.235016 | NO | NO | NA |
| Commd8 | ENSMUSG( | ENSMUSG( | -0.13737 | 0.095652 | NO | NO | NA |
| Ilvbl | ENSMUSG( | ENSMUSG( | -0.05002 | 0.600045 | NO | NO | NA |
| Plekhm2 | ENSMUSG( | ENSMUSG( | -0.23815 | 0.00388 | YES | NO | NA |
| Anapc13 | ENSMUSG( | ENSMUSG( | 0.053177 | 0.739792 | NO | NO | NA |
| Trbv13-2 | ENSMUSG( | ENSMUSG( | -0.48208 | 0.069438 | NO | NO | NA |
| Rsrc1 | ENSMUSG( | ENSMUSG( | 0.278131 | 0.039057 | YES | NO | NA |
| Poldip2 | ENSMUSG( | ENSMUSG( | 0.465543 | 2.62E-05 | YES | NO | NA |
| Cwc25 | ENSMUSG( | ENSMUSG( | -0.05027 | 0.471033 | NO | NO | NA |
| Heatr3 | ENSMUSG( | ENSMUSG( | 0.648469 | 3.48E-08 | YES | NO | NA |
| Pdik1l | ENSMUSG( | ENSMUSG( | -0.20991 | 0.001918 | YES | NO | NA |
| Tbc1d31 | ENSMUSG( | ENSMUSG( | -0.01009 | 0.944621 | NO | NO | NA |
| Nrde2 | ENSMUSG( | ENSMUSG( | 0.241078 | 0.007592 | YES | NO | NA |
| Snx11 | ENSMUSG( | ENSMUSG( | -0.10775 | 0.30141 | NO | NO | NA |
| Tor1a | ENSMUSG( | ENSMUSG( | 0.081079 | 0.209935 | NO | NO | NA |
| Nmral1 | ENSMUSG( | ENSMUSG( | 0.694034 | 1.17E-15 | YES | NO | NA |
| Clec2d | ENSMUSG( | ENSMUSG( | -0.77932 | 1.31E-07 | YES | NO | NA |
| Tada3 | ENSMUSG( | ENSMUSG( | -0.12008 | 0.179885 | NO | NO | NA |
| Alad | ENSMUSG( | ENSMUSG( | 0.320532 | 0.000528 | YES | NO | NA |
| Rabgap1 | ENSMUSG( | ENSMUSG( | -0.14751 | 0.101026 | NO | NO | NA |
| Wdfy1 | ENSMUSG( | ENSMUSG( | 0.17324 | 0.333824 | NO | NO | NA |
| Lamtor2 | ENSMUSG( | ENSMUSG( | -0.19713 | 0.032845 | YES | NO | NA |
| Galnt12 | ENSMUSG( | ENSMUSG( | 0.114095 | 0.158522 | NO | NO | NA |
| Ate1 | ENSMUSG( | ENSMUSG( | -0.1516 | 0.050077 | NO | NO | NA |
| Ska3 | ENSMUSG( | ENSMUSG( | 0.173085 | 0.034186 | YES | NO | NA |
| Wdr45b | ENSMUSG( | ENSMUSG( | 0.166117 | 0.007589 | YES | NO | NA |
| Parpbp | ENSMUSG( | ENSMUSG( | 0.105665 | 0.289824 | NO | NO | NA |
| Inppl1 | ENSMUSG( | ENSMUSG( | -0.30035 | 1.22E-06 | YES | NO | NA |
| Pfdn1 | ENSMUSG( | ENSMUSG( | 0.2317 | 0.008363 | YES | NO | NA |
| Cd55 | ENSMUSG( | ENSMUSG( | -0.79815 | 4.83E-06 | YES | NO | NA |
| Bend4 | ENSMUSG( | ENSMUSG( | -0.74602 | 4.76E-05 | YES | NO | NA |
| Def8 | ENSMUSG( | ENSMUSG( | 0.326057 | 0.004623 | YES | NO | NA |
| Gde1 | ENSMUSG( | ENSMUSG( | 0.354928 | 1.25E-05 | YES | NO | NA |
| Usp32 | ENSMUSG( | ENSMUSG( | -0.26274 | 0.004394 | YES | NO | NA |
| Ndufs4 | ENSMUSG( | ENSMUSG( | -0.07135 | 0.431466 | NO | NO | NA |
| Suclg2 | ENSMUSG( | ENSMUSG( | 0.251132 | 0.000527 | YES | NO | NA |
| Tstd2 | ENSMUSG( | ENSMUSG( | -0.13527 | 0.009669 | YES | NO | NA |
| Pmm1 | ENSMUSG( | ENSMUSG( | 0.423444 | 0.003383 | YES | NO | NA |
| Ighg1 | ENSMUSG( | ENSMUSG( | 0.420131 | 0.758402 | NO | NO | NA |
| Micu1 | ENSMUSG( | ENSMUSG( | 0.067968 | 0.276551 | NO | NO | NA |

| | | | | | | | |
|---|---|---|---|---|---|---|---|
| Atp5mpl | ENSMUSG( | ENSMUSG( | 0.220479 | 0.025388 | YES | NO | NA |
| Flvcr1 | ENSMUSG( | ENSMUSG( | 0.296182 | 0.006399 | YES | NO | NA |
| Sars2 | ENSMUSG( | ENSMUSG( | 0.410594 | 5.15E-05 | YES | NO | NA |
| Emc4 | ENSMUSG( | ENSMUSG( | 0.190522 | 0.178499 | NO | NO | NA |
| Zeb1 | ENSMUSG( | ENSMUSG( | 0.030397 | 0.764892 | NO | NO | NA |
| Higd2a | ENSMUSG( | ENSMUSG( | 0.040083 | 0.726112 | NO | NO | NA |
| Dnaja3 | ENSMUSG( | ENSMUSG( | 0.208887 | 0.01207 | YES | NO | NA |
| Ggta1 | ENSMUSG( | ENSMUSG( | -0.02367 | 0.765557 | NO | NO | NA |
| Gan | ENSMUSG( | ENSMUSG( | 0.067227 | 0.300699 | NO | NO | NA |
| Erg28 | ENSMUSG( | ENSMUSG( | 0.037918 | 0.63321 | NO | NO | NA |
| ENSMUSG | ENSMUSG( | ENSMUSG( | -0.1303 | 0.44719 | NO | NO | NA |
| Arntl | ENSMUSG( | ENSMUSG( | -0.47472 | 1.23E-07 | YES | NO | NA |
| Rybp | ENSMUSG( | ENSMUSG( | -0.40648 | 1.39E-07 | YES | NO | NA |
| Pi4k2a | ENSMUSG( | ENSMUSG( | -0.0383 | 0.647827 | NO | NO | NA |
| Tmub2 | ENSMUSG( | ENSMUSG( | -0.14161 | 0.125291 | NO | NO | NA |
| Lcor | ENSMUSG( | ENSMUSG( | -0.38812 | 3.81E-06 | YES | NO | NA |
| Clpb | ENSMUSG( | ENSMUSG( | 0.275934 | 0.000896 | YES | NO | NA |
| Stap1 | ENSMUSG( | ENSMUSG( | -0.18498 | 0.008177 | YES | NO | NA |
| ENSMUSG | ENSMUSG( | ENSMUSG( | -0.63759 | 0.008691 | YES | NO | NA |
| Sit1 | ENSMUSG( | ENSMUSG( | 0.416033 | 0.000187 | YES | NO | NA |
| Zfp574 | ENSMUSG( | ENSMUSG( | -0.27326 | 5.98E-09 | YES | NO | NA |
| Tjap1 | ENSMUSG( | ENSMUSG( | -0.14803 | 0.007028 | YES | NO | NA |
| Gclm | ENSMUSG( | ENSMUSG( | 0.36761 | 0.001926 | YES | NO | NA |
| Ssbp1 | ENSMUSG( | ENSMUSG( | 0.31038 | 3.78E-06 | YES | NO | NA |
| Qdpr | ENSMUSG( | ENSMUSG( | 0.185693 | 0.028549 | YES | NO | NA |
| Mtmr14 | ENSMUSG( | ENSMUSG( | -0.01186 | 0.914861 | NO | NO | NA |
| Ppp1r37 | ENSMUSG( | ENSMUSG( | -0.03915 | 0.601396 | NO | NO | NA |
| Crcp | ENSMUSG( | ENSMUSG( | 0.198783 | 0.065109 | NO | NO | NA |
| Fbxo9 | ENSMUSG( | ENSMUSG( | -0.50702 | 8.71E-12 | YES | NO | NA |
| Srm | ENSMUSG( | ENSMUSG( | 0.632774 | 0.000151 | YES | NO | NA |
| Pign | ENSMUSG( | ENSMUSG( | -0.27166 | 0.0106 | YES | NO | NA |
| Taf9b | ENSMUSG( | ENSMUSG( | -0.32895 | 0.009024 | YES | NO | NA |
| B3galnt2 | ENSMUSG( | ENSMUSG( | 0.099459 | 0.259399 | NO | NO | NA |
| Cltb | ENSMUSG( | ENSMUSG( | 0.31093 | 0.002693 | YES | NO | NA |
| Cers6 | ENSMUSG( | ENSMUSG( | -0.31328 | 0.000833 | YES | NO | NA |
| Golt1b | ENSMUSG( | ENSMUSG( | -0.04906 | 0.606888 | NO | NO | NA |
| Mcu | ENSMUSG( | ENSMUSG( | -0.33941 | 0.000177 | YES | NO | NA |
| Ndufab1 | ENSMUSG( | ENSMUSG( | 0.252248 | 0.000943 | YES | NO | NA |
| Dis3l | ENSMUSG( | ENSMUSG( | -0.08368 | 0.223803 | NO | NO | NA |
| Rbm34 | ENSMUSG( | ENSMUSG( | 0.266796 | 0.007792 | YES | NO | NA |
| Ubl4a | ENSMUSG( | ENSMUSG( | -0.1802 | 0.009486 | YES | NO | NA |
| Wdr70 | ENSMUSG( | ENSMUSG( | 0.080506 | 0.492717 | NO | NO | NA |
| Fbxw4 | ENSMUSG( | ENSMUSG( | -0.44002 | 9.90E-06 | YES | NO | NA |
| Acss1 | ENSMUSG( | ENSMUSG( | -0.65906 | 1.92E-07 | YES | NO | NA |
| Nkap | ENSMUSG( | ENSMUSG( | -0.15719 | 0.12565 | NO | NO | NA |
| Phc1 | ENSMUSG( | ENSMUSG( | 0.038738 | 0.785998 | NO | NO | NA |
| Ercc2 | ENSMUSG( | ENSMUSG( | 0.108428 | 0.299093 | NO | NO | NA |

| | | | | | | | |
|---|---|---|---|---|---|---|---|
| Mfsd4a | ENSMUSG( | ENSMUSG( | -0.26332 | 0.002225 | YES | NO | NA |
| Thoc6 | ENSMUSG( | ENSMUSG( | 0.204337 | 0.046136 | YES | NO | NA |
| P4ha1 | ENSMUSG( | ENSMUSG( | -0.50584 | 7.52E-16 | YES | NO | NA |
| Nt5c2 | ENSMUSG( | ENSMUSG( | 0.43864 | 5.78E-07 | YES | NO | NA |
| Lmbrd2 | ENSMUSG( | ENSMUSG( | -0.21604 | 0.019354 | YES | NO | NA |
| Mlst8 | ENSMUSG( | ENSMUSG( | 0.089277 | 0.406768 | NO | NO | NA |
| Tmem184 | ENSMUSG( | ENSMUSG( | -0.01568 | 0.878742 | NO | NO | NA |
| Ier3ip1 | ENSMUSG( | ENSMUSG( | -0.14564 | 0.119971 | NO | NO | NA |
| Cldnd1 | ENSMUSG( | ENSMUSG( | 0.04625 | 0.706298 | NO | NO | NA |
| E2f3 | ENSMUSG( | ENSMUSG( | 0.061218 | 0.501178 | NO | NO | NA |
| Vps33b | ENSMUSG( | ENSMUSG( | 0.053507 | 0.476831 | NO | NO | NA |
| Zfp318 | ENSMUSG( | ENSMUSG( | -0.16837 | 0.163347 | NO | NO | NA |
| Slc35b2 | ENSMUSG( | ENSMUSG( | -0.06862 | 0.244414 | NO | NO | NA |
| Rad51ap1 | ENSMUSG( | ENSMUSG( | 0.320958 | 0.001077 | YES | NO | NA |
| Terf2ip | ENSMUSG( | ENSMUSG( | -0.14296 | 0.101738 | NO | NO | NA |
| Stk19 | ENSMUSG( | ENSMUSG( | 0.257543 | 0.020005 | YES | NO | NA |
| Mydgf | ENSMUSG( | ENSMUSG( | 0.12192 | 0.16766 | NO | NO | NA |
| 1810009A | ENSMUSG( | ENSMUSG( | -0.08004 | 0.514713 | NO | NO | NA |
| Mlh1 | ENSMUSG( | ENSMUSG( | 0.307352 | 9.82E-05 | YES | NO | NA |
| Ccdc134 | ENSMUSG( | ENSMUSG( | -0.06959 | 0.354347 | NO | NO | NA |
| Fhip1b | ENSMUSG( | ENSMUSG( | -0.54987 | 4.85E-07 | YES | NO | NA |
| Ddx56 | ENSMUSG( | ENSMUSG( | 0.403172 | 3.50E-06 | YES | NO | NA |
| Helq | ENSMUSG( | ENSMUSG( | 0.020254 | 0.795988 | NO | NO | NA |
| Acad9 | ENSMUSG( | ENSMUSG( | 0.178791 | 0.112409 | NO | NO | NA |
| Ttc4 | ENSMUSG( | ENSMUSG( | 0.086636 | 0.326072 | NO | NO | NA |
| Iscu | ENSMUSG( | ENSMUSG( | 0.150159 | 0.054271 | NO | NO | NA |
| Hspbp1 | ENSMUSG( | ENSMUSG( | 0.514351 | 6.83E-06 | YES | NO | NA |
| Urb1 | ENSMUSG( | ENSMUSG( | 0.449813 | 8.07E-05 | YES | NO | NA |
| Zscan26 | ENSMUSG( | ENSMUSG( | 0.103534 | 0.430226 | NO | NO | NA |
| Stam2 | ENSMUSG( | ENSMUSG( | -0.08737 | 0.327902 | NO | NO | NA |
| Hes6 | ENSMUSG( | ENSMUSG( | -0.10026 | 0.225087 | NO | NO | NA |
| Filip1l | ENSMUSG( | ENSMUSG( | 0.067351 | 0.732409 | NO | NO | NA |
| L3mbtl3 | ENSMUSG( | ENSMUSG( | -0.84551 | 3.83E-12 | YES | NO | NA |
| Klrg1 | ENSMUSG( | ENSMUSG( | -0.36772 | 0.118234 | NO | NO | NA |
| Trmt2b | ENSMUSG( | ENSMUSG( | -0.05797 | 0.517788 | NO | NO | NA |
| Pde1b | ENSMUSG( | ENSMUSG( | 0.066856 | 0.664439 | NO | NO | NA |
| Klhl5 | ENSMUSG( | ENSMUSG( | 0.503898 | 2.20E-06 | YES | NO | NA |
| Slfn9 | ENSMUSG( | ENSMUSG( | -0.20302 | 0.074552 | NO | NO | NA |
| Fam204a | ENSMUSG( | ENSMUSG( | 0.231255 | 0.001045 | YES | NO | NA |
| Zmym4 | ENSMUSG( | ENSMUSG( | 0.090899 | 0.305959 | NO | NO | NA |
| Washc4 | ENSMUSG( | ENSMUSG( | -0.20693 | 0.009811 | YES | NO | NA |
| Cmtm4 | ENSMUSG( | ENSMUSG( | -0.17415 | 0.061444 | NO | NO | NA |
| Ppp1r13b | ENSMUSG( | ENSMUSG( | -0.76477 | 4.04E-11 | YES | NO | NA |
| Ints2 | ENSMUSG( | ENSMUSG( | 0.198806 | 0.042893 | YES | NO | NA |
| Cd80 | ENSMUSG( | ENSMUSG( | 0.215919 | 0.168759 | NO | NO | NA |
| Mri1 | ENSMUSG( | ENSMUSG( | 0.094009 | 0.296568 | NO | NO | NA |
| Kti12 | ENSMUSG( | ENSMUSG( | 0.248541 | 0.000392 | YES | NO | NA |

| | | | | | | | |
|---|---|---|---|---|---|---|---|
| Tmem170l | ENSMUSG( | ENSMUSG( | -0.27921 | 0.006598 | YES | NO | NA |
| Phf10 | ENSMUSG( | ENSMUSG( | 0.240234 | 0.009892 | YES | NO | NA |
| Gtf3c6 | ENSMUSG( | ENSMUSG( | -0.02644 | 0.872677 | NO | NO | NA |
| Bcas3 | ENSMUSG( | ENSMUSG( | -0.01921 | 0.897257 | NO | NO | NA |
| Sap30l | ENSMUSG( | ENSMUSG( | 0.020128 | 0.829321 | NO | NO | NA |
| Tubgcp5 | ENSMUSG( | ENSMUSG( | -0.04923 | 0.656785 | NO | NO | NA |
| Tmem19 | ENSMUSG( | ENSMUSG( | 0.048886 | 0.725258 | NO | NO | NA |
| Ptbp2 | ENSMUSG( | ENSMUSG( | -0.05825 | 0.664323 | NO | NO | NA |
| Dhx32 | ENSMUSG( | ENSMUSG( | 0.113288 | 0.134262 | NO | NO | NA |
| Lsm5 | ENSMUSG( | ENSMUSG( | 0.381427 | 0.001025 | YES | NO | NA |
| Vamp4 | ENSMUSG( | ENSMUSG( | -0.29345 | 2.49E-06 | YES | NO | NA |
| Zfp398 | ENSMUSG( | ENSMUSG( | -0.03695 | 0.758872 | NO | NO | NA |
| Ciao2b | ENSMUSG( | ENSMUSG( | -0.20451 | 0.029938 | YES | NO | NA |
| Polr3e | ENSMUSG( | ENSMUSG( | 0.29584 | 0.001752 | YES | NO | NA |
| Ankrd54 | ENSMUSG( | ENSMUSG( | 0.036128 | 0.695741 | NO | NO | NA |
| Pja1 | ENSMUSG( | ENSMUSG( | -0.17431 | 0.043273 | YES | NO | NA |
| Ipo4 | ENSMUSG( | ENSMUSG( | 0.680995 | 2.55E-06 | YES | NO | NA |
| Adpgk | ENSMUSG( | ENSMUSG( | 0.288487 | 0.000928 | YES | NO | NA |
| Tmem222 | ENSMUSG( | ENSMUSG( | 0.153664 | 0.126819 | NO | NO | NA |
| Bud23 | ENSMUSG( | ENSMUSG( | 0.101099 | 0.201773 | NO | NO | NA |
| Lzic | ENSMUSG( | ENSMUSG( | 0.073301 | 0.406658 | NO | NO | NA |
| Coq9 | ENSMUSG( | ENSMUSG( | 0.196145 | 0.037868 | YES | NO | NA |
| Naxe | ENSMUSG( | ENSMUSG( | 0.096189 | 0.26454 | NO | NO | NA |
| Polr2l | ENSMUSG( | ENSMUSG( | 0.37661 | 0.000179 | YES | NO | NA |
| Yipf5 | ENSMUSG( | ENSMUSG( | -0.04498 | 0.662185 | NO | NO | NA |
| Lap3 | ENSMUSG( | ENSMUSG( | 0.523568 | 0.000735 | YES | NO | NA |
| Zfp180 | ENSMUSG( | ENSMUSG( | -0.22874 | 0.00141 | YES | NO | NA |
| Hsdl1 | ENSMUSG( | ENSMUSG( | -0.36065 | 9.29E-09 | YES | NO | NA |
| Josd1 | ENSMUSG( | ENSMUSG( | 0.255187 | 4.67E-05 | YES | NO | NA |
| Ctso | ENSMUSG( | ENSMUSG( | -0.2463 | 0.030461 | YES | NO | NA |
| Lysmd3 | ENSMUSG( | ENSMUSG( | -0.3232 | 3.46E-06 | YES | NO | NA |
| Lig3 | ENSMUSG( | ENSMUSG( | 0.181015 | 0.020632 | YES | NO | NA |
| Arid3b | ENSMUSG( | ENSMUSG( | -0.51984 | 5.68E-08 | YES | NO | NA |
| Serpine2 | ENSMUSG( | ENSMUSG( | 1.201707 | 9.96E-10 | YES | YES | Upregulated |
| Tmem181a | ENSMUSG( | ENSMUSG( | 0.159365 | 0.081426 | NO | NO | NA |
| Acot11 | ENSMUSG( | ENSMUSG( | -0.14689 | 0.330787 | NO | NO | NA |
| Gpatch4 | ENSMUSG( | ENSMUSG( | 0.818829 | 0.00029 | YES | NO | NA |
| Fiz1 | ENSMUSG( | ENSMUSG( | -0.09771 | 0.23928 | NO | NO | NA |
| F730043M | ENSMUSG( | ENSMUSG( | -0.65093 | 1.98E-09 | YES | NO | NA |
| Mtmr4 | ENSMUSG( | ENSMUSG( | 0.053308 | 0.610282 | NO | NO | NA |
| Lactb | ENSMUSG( | ENSMUSG( | -0.02309 | 0.841324 | NO | NO | NA |
| Prmt7 | ENSMUSG( | ENSMUSG( | 0.540328 | 4.55E-07 | YES | NO | NA |
| Zfyve1 | ENSMUSG( | ENSMUSG( | -0.26706 | 1.96E-05 | YES | NO | NA |
| Tprkb | ENSMUSG( | ENSMUSG( | 0.102942 | 0.149383 | NO | NO | NA |
| Zfp609 | ENSMUSG( | ENSMUSG( | -0.0965 | 0.35495 | NO | NO | NA |
| Relt | ENSMUSG( | ENSMUSG( | -0.16698 | 0.026013 | YES | NO | NA |
| Ell | ENSMUSG( | ENSMUSG( | -0.08801 | 0.410282 | NO | NO | NA |

| | | | | | | | |
|---|---|---|---|---|---|---|---|
| Ppp1r11 | ENSMUSG( | ENSMUSG( | -0.12771 | 0.179362 | NO | NO | NA |
| Tmem41b | ENSMUSG( | ENSMUSG( | 0.076537 | 0.328993 | NO | NO | NA |
| Trappc5 | ENSMUSG( | ENSMUSG( | 0.044655 | 0.604896 | NO | NO | NA |
| Mitd1 | ENSMUSG( | ENSMUSG( | -0.26415 | 0.000858 | YES | NO | NA |
| Ccnh | ENSMUSG( | ENSMUSG( | 0.182887 | 0.038073 | YES | NO | NA |
| Ccdc66 | ENSMUSG( | ENSMUSG( | -0.2904 | 0.002643 | YES | NO | NA |
| Kdm4b | ENSMUSG( | ENSMUSG( | -0.29691 | 0.00041 | YES | NO | NA |
| Zfp644 | ENSMUSG( | ENSMUSG( | -0.19153 | 0.058739 | NO | NO | NA |
| Ccdc92 | ENSMUSG( | ENSMUSG( | 0.21614 | 0.034575 | YES | NO | NA |
| Mettl4 | ENSMUSG( | ENSMUSG( | -0.23996 | 0.000143 | YES | NO | NA |
| Zfp280c | ENSMUSG( | ENSMUSG( | 0.155649 | 0.285795 | NO | NO | NA |
| Pgm2 | ENSMUSG( | ENSMUSG( | -0.41045 | 2.33E-07 | YES | NO | NA |
| Fbxo21 | ENSMUSG( | ENSMUSG( | -0.00771 | 0.902274 | NO | NO | NA |
| Napsa | ENSMUSG( | ENSMUSG( | 0.372677 | 0.027392 | YES | NO | NA |
| Slc37a3 | ENSMUSG( | ENSMUSG( | -0.17502 | 0.063766 | NO | NO | NA |
| Ldah | ENSMUSG( | ENSMUSG( | -0.09052 | 0.291361 | NO | NO | NA |
| Lsm7 | ENSMUSG( | ENSMUSG( | 0.433139 | 0.001537 | YES | NO | NA |
| Plekha1 | ENSMUSG( | ENSMUSG( | -0.82955 | 7.86E-10 | YES | NO | NA |
| Nip7 | ENSMUSG( | ENSMUSG( | 0.244592 | 0.002313 | YES | NO | NA |
| Gstcd | ENSMUSG( | ENSMUSG( | 0.276796 | 0.002592 | YES | NO | NA |
| Rnf34 | ENSMUSG( | ENSMUSG( | -0.1918 | 0.006786 | YES | NO | NA |
| Slc35e2 | ENSMUSG( | ENSMUSG( | -0.16903 | 0.046105 | YES | NO | NA |
| Trim65 | ENSMUSG( | ENSMUSG( | -0.86561 | 2.39E-18 | YES | NO | NA |
| Nsmce2 | ENSMUSG( | ENSMUSG( | 0.372497 | 7.86E-07 | YES | NO | NA |
| Rhob | ENSMUSG( | ENSMUSG( | 0.445613 | 0.108867 | NO | NO | NA |
| Mrpl2 | ENSMUSG( | ENSMUSG( | 0.663858 | 2.15E-07 | YES | NO | NA |
| Ifit1 | ENSMUSG( | ENSMUSG( | -1.39021 | 1.49E-13 | YES | YES | Downregulated |
| Cenpo | ENSMUSG( | ENSMUSG( | 0.081873 | 0.278173 | NO | NO | NA |
| Adprs | ENSMUSG( | ENSMUSG( | 0.053237 | 0.417268 | NO | NO | NA |
| Cerk | ENSMUSG( | ENSMUSG( | -0.07212 | 0.471208 | NO | NO | NA |
| Naip2 | ENSMUSG( | ENSMUSG( | -0.70402 | 5.18E-12 | YES | NO | NA |
| Ccdc69 | ENSMUSG( | ENSMUSG( | -0.39671 | 1.73E-07 | YES | NO | NA |
| Slc25a40 | ENSMUSG( | ENSMUSG( | -0.36309 | 0.005898 | YES | NO | NA |
| 2610318N | ENSMUSG( | ENSMUSG( | 0.359949 | 0.000347 | YES | NO | NA |
| Cep104 | ENSMUSG( | ENSMUSG( | -0.07973 | 0.450894 | NO | NO | NA |
| Yaf2 | ENSMUSG( | ENSMUSG( | -0.34376 | 1.68E-05 | YES | NO | NA |
| Strada | ENSMUSG( | ENSMUSG( | 0.11833 | 0.168084 | NO | NO | NA |
| Slc10a7 | ENSMUSG( | ENSMUSG( | -0.14815 | 0.009147 | YES | NO | NA |
| Btbd11 | ENSMUSG( | ENSMUSG( | -0.36886 | 0.03453 | YES | NO | NA |
| Exosc5 | ENSMUSG( | ENSMUSG( | 0.224841 | 0.000681 | YES | NO | NA |
| Ipmk | ENSMUSG( | ENSMUSG( | 0.059815 | 0.528583 | NO | NO | NA |
| Polr1b | ENSMUSG( | ENSMUSG( | 0.470176 | 0.00022 | YES | NO | NA |
| Vars2 | ENSMUSG( | ENSMUSG( | 0.168795 | 0.006332 | YES | NO | NA |
| Trp53bp2 | ENSMUSG( | ENSMUSG( | -0.19964 | 0.030869 | YES | NO | NA |
| Wdr61 | ENSMUSG( | ENSMUSG( | 0.090141 | 0.254094 | NO | NO | NA |
| Txlng | ENSMUSG( | ENSMUSG( | -0.23212 | 0.003396 | YES | NO | NA |
| Tomm5 | ENSMUSG( | ENSMUSG( | 0.246045 | 0.006895 | YES | NO | NA |

| | | | | | | | |
|---|---|---|---|---|---|---|---|
| H2-Ke6 | ENSMUSG( | ENSMUSG( | -0.10848 | 0.095947 | NO | NO | NA |
| Mphosph6 | ENSMUSG( | ENSMUSG( | 0.195368 | 0.114903 | NO | NO | NA |
| Cdon | ENSMUSG( | ENSMUSG( | -0.22549 | 0.028584 | YES | NO | NA |
| Ipo11 | ENSMUSG( | ENSMUSG( | 0.410685 | 9.54E-08 | YES | NO | NA |
| 1110051M | ENSMUSG( | ENSMUSG( | 0.135183 | 0.218945 | NO | NO | NA |
| Rnf113a2 | ENSMUSG( | ENSMUSG( | -0.00538 | 0.975344 | NO | NO | NA |
| Dpy19l3 | ENSMUSG( | ENSMUSG( | -0.34734 | 0.000585 | YES | NO | NA |
| Fchsd2 | ENSMUSG( | ENSMUSG( | -0.44206 | 0.00013 | YES | NO | NA |
| Pim3 | ENSMUSG( | ENSMUSG( | 0.174957 | 0.15403 | NO | NO | NA |
| Stard3 | ENSMUSG( | ENSMUSG( | -0.24818 | 0.0044 | YES | NO | NA |
| Sfi1 | ENSMUSG( | ENSMUSG( | -0.23694 | 0.032043 | YES | NO | NA |
| Zfp628 | ENSMUSG( | ENSMUSG( | -0.13279 | 0.233019 | NO | NO | NA |
| Mfhas1 | ENSMUSG( | ENSMUSG( | 0.04673 | 0.700795 | NO | NO | NA |
| Mrps15 | ENSMUSG( | ENSMUSG( | 0.038842 | 0.772033 | NO | NO | NA |
| Ulk1 | ENSMUSG( | ENSMUSG( | -0.21183 | 0.074356 | NO | NO | NA |
| Dvl2 | ENSMUSG( | ENSMUSG( | -0.07679 | 0.248308 | NO | NO | NA |
| Smn1 | ENSMUSG( | ENSMUSG( | 0.406011 | 0.000443 | YES | NO | NA |
| Zdhhc4 | ENSMUSG( | ENSMUSG( | 0.318829 | 0.002097 | YES | NO | NA |
| Phf21a | ENSMUSG( | ENSMUSG( | -0.45942 | 2.11E-05 | YES | NO | NA |
| Ammecr1 | ENSMUSG( | ENSMUSG( | -0.04045 | 0.752616 | NO | NO | NA |
| Arid3a | ENSMUSG( | ENSMUSG( | -0.28793 | 4.79E-05 | YES | NO | NA |
| Mrpl39 | ENSMUSG( | ENSMUSG( | 0.035808 | 0.734883 | NO | NO | NA |
| Dym | ENSMUSG( | ENSMUSG( | -0.04587 | 0.634607 | NO | NO | NA |
| Hivep3 | ENSMUSG( | ENSMUSG( | -0.28042 | 0.034186 | YES | NO | NA |
| Hace1 | ENSMUSG( | ENSMUSG( | -0.2303 | 0.001919 | YES | NO | NA |
| Erf | ENSMUSG( | ENSMUSG( | -0.05725 | 0.685344 | NO | NO | NA |
| Ift140 | ENSMUSG( | ENSMUSG( | 0.068139 | 0.495773 | NO | NO | NA |
| Osbpl2 | ENSMUSG( | ENSMUSG( | -0.22435 | 0.001436 | YES | NO | NA |
| Abhd16a | ENSMUSG( | ENSMUSG( | -0.0172 | 0.851755 | NO | NO | NA |
| Palm | ENSMUSG( | ENSMUSG( | 0.254954 | 0.057228 | NO | NO | NA |
| Niban2 | ENSMUSG( | ENSMUSG( | 0.858587 | 6.13E-05 | YES | NO | NA |
| Pgap6 | ENSMUSG( | ENSMUSG( | -0.35278 | 2.22E-05 | YES | NO | NA |
| Hic1 | ENSMUSG( | ENSMUSG( | -0.71527 | 0.000443 | YES | NO | NA |
| Cd274 | ENSMUSG( | ENSMUSG( | -0.23523 | 0.105707 | NO | NO | NA |
| Cep43 | ENSMUSG( | ENSMUSG( | 0.44422 | 0.000176 | YES | NO | NA |
| Zbtb25 | ENSMUSG( | ENSMUSG( | -0.00949 | 0.911186 | NO | NO | NA |
| Bcat2 | ENSMUSG( | ENSMUSG( | 0.21913 | 0.005698 | YES | NO | NA |
| Mrps14 | ENSMUSG( | ENSMUSG( | 0.198326 | 0.001686 | YES | NO | NA |
| Brwd3 | ENSMUSG( | ENSMUSG( | -0.34271 | 0.000111 | YES | NO | NA |
| Crtam | ENSMUSG( | ENSMUSG( | 0.744212 | 5.92E-10 | YES | NO | NA |
| Frrs1 | ENSMUSG( | ENSMUSG( | -0.28161 | 1.97E-05 | YES | NO | NA |
| Sav1 | ENSMUSG( | ENSMUSG( | -0.03783 | 0.63825 | NO | NO | NA |
| Recql4 | ENSMUSG( | ENSMUSG( | 0.545629 | 3.03E-05 | YES | NO | NA |
| Acads | ENSMUSG( | ENSMUSG( | 0.081249 | 0.419662 | NO | NO | NA |
| 4933434E2 | ENSMUSG( | ENSMUSG( | -0.17166 | 0.011513 | YES | NO | NA |
| Higd1a | ENSMUSG( | ENSMUSG( | 0.042631 | 0.677734 | NO | NO | NA |
| Fbxo46 | ENSMUSG( | ENSMUSG( | -0.29509 | 0.000136 | YES | NO | NA |

| | | | | | | | |
|---|---|---|---|---|---|---|---|
| Mrgbp | ENSMUSG( | ENSMUSG( | -0.05054 | 0.399916 | NO | NO | NA |
| Rcan3 | ENSMUSG( | ENSMUSG( | 0.007408 | 0.932872 | NO | NO | NA |
| Ankrd26 | ENSMUSG( | ENSMUSG( | 0.08865 | 0.484558 | NO | NO | NA |
| Banp | ENSMUSG( | ENSMUSG( | -0.34643 | 9.48E-10 | YES | NO | NA |
| Dhps | ENSMUSG( | ENSMUSG( | 0.113277 | 0.02941 | YES | NO | NA |
| Cep68 | ENSMUSG( | ENSMUSG( | -0.14658 | 0.042877 | YES | NO | NA |
| Snap23 | ENSMUSG( | ENSMUSG( | -0.18727 | 0.01229 | YES | NO | NA |
| Snx14 | ENSMUSG( | ENSMUSG( | -0.36179 | 1.20E-06 | YES | NO | NA |
| Pskh1 | ENSMUSG( | ENSMUSG( | -0.31677 | 0.019491 | YES | NO | NA |
| Scai | ENSMUSG( | ENSMUSG( | -0.3891 | 0.001971 | YES | NO | NA |
| Nit1 | ENSMUSG( | ENSMUSG( | -0.18234 | 0.001506 | YES | NO | NA |
| Vamp1 | ENSMUSG( | ENSMUSG( | -0.53384 | 6.07E-05 | YES | NO | NA |
| Polrmt | ENSMUSG( | ENSMUSG( | 0.100213 | 0.225137 | NO | NO | NA |
| 5031425E2 | ENSMUSG( | ENSMUSG( | -0.1796 | 0.130505 | NO | NO | NA |
| Tmem97 | ENSMUSG( | ENSMUSG( | 0.970915 | 5.93E-08 | YES | NO | NA |
| Chek1 | ENSMUSG( | ENSMUSG( | 0.032957 | 0.796294 | NO | NO | NA |
| Exosc7 | ENSMUSG( | ENSMUSG( | 0.244678 | 0.004036 | YES | NO | NA |
| 1500004A | ENSMUSG( | ENSMUSG( | -0.28119 | 0.005714 | YES | NO | NA |
| Ppif | ENSMUSG( | ENSMUSG( | 0.146978 | 0.026344 | YES | NO | NA |
| Ube2e3 | ENSMUSG( | ENSMUSG( | 0.354962 | 5.26E-07 | YES | NO | NA |
| Mrpl45 | ENSMUSG( | ENSMUSG( | 0.443388 | 9.05E-11 | YES | NO | NA |
| Mcm9 | ENSMUSG( | ENSMUSG( | 0.088893 | 0.211211 | NO | NO | NA |
| Klhdc10 | ENSMUSG( | ENSMUSG( | 0.222237 | 0.028191 | YES | NO | NA |
| Tab1 | ENSMUSG( | ENSMUSG( | 0.485171 | 2.11E-06 | YES | NO | NA |
| Gna12 | ENSMUSG( | ENSMUSG( | -0.06075 | 0.590627 | NO | NO | NA |
| Atox1 | ENSMUSG( | ENSMUSG( | 0.090326 | 0.374636 | NO | NO | NA |
| Riok2 | ENSMUSG( | ENSMUSG( | 0.253562 | 0.003364 | YES | NO | NA |
| Zcchc9 | ENSMUSG( | ENSMUSG( | -0.00089 | 0.991454 | NO | NO | NA |
| Polr2d | ENSMUSG( | ENSMUSG( | 0.265413 | 0.000891 | YES | NO | NA |
| Dtnb | ENSMUSG( | ENSMUSG( | -0.27976 | 0.009017 | YES | NO | NA |
| Fbxw7 | ENSMUSG( | ENSMUSG( | -0.26656 | 0.000992 | YES | NO | NA |
| Actr8 | ENSMUSG( | ENSMUSG( | -0.02114 | 0.794099 | NO | NO | NA |
| Camk2d | ENSMUSG( | ENSMUSG( | -0.24847 | 0.000596 | YES | NO | NA |
| Retreg1 | ENSMUSG( | ENSMUSG( | -0.43709 | 4.02E-11 | YES | NO | NA |
| Adnp2 | ENSMUSG( | ENSMUSG( | -0.14651 | 0.010009 | YES | NO | NA |
| Zfp276 | ENSMUSG( | ENSMUSG( | -0.41436 | 3.25E-09 | YES | NO | NA |
| Il6st | ENSMUSG( | ENSMUSG( | -0.26258 | 0.013783 | YES | NO | NA |
| Armt1 | ENSMUSG( | ENSMUSG( | 0.036226 | 0.584088 | NO | NO | NA |
| Setdb2 | ENSMUSG( | ENSMUSG( | -0.19878 | 0.000758 | YES | NO | NA |
| Mtpap | ENSMUSG( | ENSMUSG( | 0.032414 | 0.725834 | NO | NO | NA |
| Zfp518a | ENSMUSG( | ENSMUSG( | -0.35619 | 0.000337 | YES | NO | NA |
| Exoc8 | ENSMUSG( | ENSMUSG( | -0.31582 | 6.13E-05 | YES | NO | NA |
| Ercc5 | ENSMUSG( | ENSMUSG( | -0.03502 | 0.772033 | NO | NO | NA |
| Ube2d1 | ENSMUSG( | ENSMUSG( | -0.39663 | 7.65E-06 | YES | NO | NA |
| Cbll1 | ENSMUSG( | ENSMUSG( | -0.27043 | 5.29E-05 | YES | NO | NA |
| Cfap97 | ENSMUSG( | ENSMUSG( | -0.13713 | 0.111463 | NO | NO | NA |
| Atrn | ENSMUSG( | ENSMUSG( | -0.09769 | 0.389321 | NO | NO | NA |

| | | | | | | | |
|---|---|---|---|---|---|---|---|
| Sec22c | ENSMUSG( | ENSMUSG( | -0.11903 | 0.251677 | NO | NO | NA |
| Fam53a | ENSMUSG( | ENSMUSG( | -0.02472 | 0.784577 | NO | NO | NA |
| Rex1bd | ENSMUSG( | ENSMUSG( | 0.210408 | 0.02414 | YES | NO | NA |
| Emc6 | ENSMUSG( | ENSMUSG( | 0.208376 | 0.001121 | YES | NO | NA |
| Drg2 | ENSMUSG( | ENSMUSG( | 0.215716 | 0.003913 | YES | NO | NA |
| Snhg5 | ENSMUSG( | ENSMUSG( | 0.614128 | 2.13E-06 | YES | NO | NA |
| Zfp280d | ENSMUSG( | ENSMUSG( | 0.008045 | 0.909568 | NO | NO | NA |
| Ddx28 | ENSMUSG( | ENSMUSG( | -0.3844 | 2.56E-13 | YES | NO | NA |
| Pop5 | ENSMUSG( | ENSMUSG( | 0.225633 | 0.006919 | YES | NO | NA |
| Tmem263 | ENSMUSG( | ENSMUSG( | -0.23246 | 0.000181 | YES | NO | NA |
| Lss | ENSMUSG( | ENSMUSG( | 0.473644 | 2.78E-06 | YES | NO | NA |
| Zfp317 | ENSMUSG( | ENSMUSG( | -0.31332 | 0.00019 | YES | NO | NA |
| Skap2 | ENSMUSG( | ENSMUSG( | 0.269706 | 0.014453 | YES | NO | NA |
| Ube2a | ENSMUSG( | ENSMUSG( | -0.06607 | 0.45456 | NO | NO | NA |
| Cenpn | ENSMUSG( | ENSMUSG( | 0.438828 | 9.41E-14 | YES | NO | NA |
| Pot1b | ENSMUSG( | ENSMUSG( | -0.42169 | 6.06E-05 | YES | NO | NA |
| Btf3l4 | ENSMUSG( | ENSMUSG( | 0.395518 | 0.000691 | YES | NO | NA |
| Cdc42bpg | ENSMUSG( | ENSMUSG( | -0.17061 | 0.107146 | NO | NO | NA |
| Wdr44 | ENSMUSG( | ENSMUSG( | -0.35585 | 1.62E-05 | YES | NO | NA |
| Tmem242 | ENSMUSG( | ENSMUSG( | 0.33406 | 9.85E-05 | YES | NO | NA |
| Pus10 | ENSMUSG( | ENSMUSG( | 0.191337 | 0.043421 | YES | NO | NA |
| Dpagt1 | ENSMUSG( | ENSMUSG( | 0.261767 | 0.000846 | YES | NO | NA |
| Mrpl17 | ENSMUSG( | ENSMUSG( | 0.415833 | 1.74E-07 | YES | NO | NA |
| Ormdl3 | ENSMUSG( | ENSMUSG( | -0.32633 | 0.004108 | YES | NO | NA |
| Nbas | ENSMUSG( | ENSMUSG( | 0.044698 | 0.609667 | NO | NO | NA |
| Mdm1 | ENSMUSG( | ENSMUSG( | 0.086964 | 0.491132 | NO | NO | NA |
| Ddx20 | ENSMUSG( | ENSMUSG( | 0.181434 | 0.000429 | YES | NO | NA |
| Zfp777 | ENSMUSG( | ENSMUSG( | 0.163467 | 0.07313 | NO | NO | NA |
| Snapc1 | ENSMUSG( | ENSMUSG( | -0.06311 | 0.573474 | NO | NO | NA |
| Dalrd3 | ENSMUSG( | ENSMUSG( | -0.04932 | 0.658479 | NO | NO | NA |
| Ipo13 | ENSMUSG( | ENSMUSG( | -0.03949 | 0.600373 | NO | NO | NA |
| Cnnm3 | ENSMUSG( | ENSMUSG( | -0.29382 | 0.000645 | YES | NO | NA |
| Vamp5 | ENSMUSG( | ENSMUSG( | -0.33391 | 0.000447 | YES | NO | NA |
| Tent5a | ENSMUSG( | ENSMUSG( | -0.2667 | 0.010575 | YES | NO | NA |
| Chm | ENSMUSG( | ENSMUSG( | -0.14404 | 0.105925 | NO | NO | NA |
| Mrpl52 | ENSMUSG( | ENSMUSG( | 0.348169 | 6.01E-07 | YES | NO | NA |
| Arl5c | ENSMUSG( | ENSMUSG( | -0.99996 | 4.55E-07 | YES | NO | NA |
| Bag4 | ENSMUSG( | ENSMUSG( | 0.07855 | 0.307944 | NO | NO | NA |
| Suox | ENSMUSG( | ENSMUSG( | -0.4863 | 1.97E-06 | YES | NO | NA |
| Gm28043 | ENSMUSG( | ENSMUSG( | -0.14183 | 0.23247 | NO | NO | NA |
| Zbtb7b | ENSMUSG( | ENSMUSG( | -0.51902 | 5.20E-08 | YES | NO | NA |
| Gtf2f2 | ENSMUSG( | ENSMUSG( | 0.305067 | 2.48E-05 | YES | NO | NA |
| Snapc5 | ENSMUSG( | ENSMUSG( | 0.032158 | 0.844004 | NO | NO | NA |
| Wdr74 | ENSMUSG( | ENSMUSG( | 0.502641 | 2.30E-07 | YES | NO | NA |
| Dcp1b | ENSMUSG( | ENSMUSG( | -0.3562 | 0.000275 | YES | NO | NA |
| Prdm10 | ENSMUSG( | ENSMUSG( | -0.39086 | 3.90E-06 | YES | NO | NA |
| Alms1 | ENSMUSG( | ENSMUSG( | -0.0635 | 0.378248 | NO | NO | NA |

| | | | | | | | |
|---|---|---|---|---|---|---|---|
| Epm2aip1 | ENSMUSG( | ENSMUSG( | -0.07 | 0.412994 | NO | NO | NA |
| Kin | ENSMUSG( | ENSMUSG( | -0.13548 | 0.354025 | NO | NO | NA |
| Rbsn | ENSMUSG( | ENSMUSG( | 0.031532 | 0.695107 | NO | NO | NA |
| Ccr9 | ENSMUSG( | ENSMUSG( | 0.308791 | 0.477841 | NO | NO | NA |
| Tex264 | ENSMUSG( | ENSMUSG( | -0.37251 | 1.28E-08 | YES | NO | NA |
| Zfp212 | ENSMUSG( | ENSMUSG( | -0.188 | 0.027785 | YES | NO | NA |
| Phka2 | ENSMUSG( | ENSMUSG( | -0.08455 | 0.438501 | NO | NO | NA |
| Uckl1 | ENSMUSG( | ENSMUSG( | -0.29971 | 0.005832 | YES | NO | NA |
| Rbfa | ENSMUSG( | ENSMUSG( | 0.334412 | 0.000475 | YES | NO | NA |
| Ap5z1 | ENSMUSG( | ENSMUSG( | -0.40225 | 2.98E-05 | YES | NO | NA |
| Ppih | ENSMUSG( | ENSMUSG( | 0.45073 | 4.35E-07 | YES | NO | NA |
| Mrps33 | ENSMUSG( | ENSMUSG( | 0.315437 | 0.000358 | YES | NO | NA |
| Zfp655 | ENSMUSG( | ENSMUSG( | -0.34349 | 5.50E-05 | YES | NO | NA |
| Fdxr | ENSMUSG( | ENSMUSG( | -0.19214 | 0.012777 | YES | NO | NA |
| Smyd5 | ENSMUSG( | ENSMUSG( | 0.819301 | 3.12E-10 | YES | NO | NA |
| Sass6 | ENSMUSG( | ENSMUSG( | 0.111618 | 0.352248 | NO | NO | NA |
| Fyttd1 | ENSMUSG( | ENSMUSG( | 0.228889 | 0.000561 | YES | NO | NA |
| Rpa3 | ENSMUSG( | ENSMUSG( | 0.142777 | 0.066281 | NO | NO | NA |
| Blvra | ENSMUSG( | ENSMUSG( | 0.134458 | 0.08852 | NO | NO | NA |
| Cenpj | ENSMUSG( | ENSMUSG( | -0.2761 | 0.001813 | YES | NO | NA |
| Adam9 | ENSMUSG( | ENSMUSG( | 0.25524 | 0.017966 | YES | NO | NA |
| Ggps1 | ENSMUSG( | ENSMUSG( | -0.13676 | 0.032532 | YES | NO | NA |
| Diablo | ENSMUSG( | ENSMUSG( | -0.0627 | 0.517571 | NO | NO | NA |
| Atp8a1 | ENSMUSG( | ENSMUSG( | -0.39923 | 0.000344 | YES | NO | NA |
| Slc37a2 | ENSMUSG( | ENSMUSG( | -0.56546 | 0.002093 | YES | NO | NA |
| Focad | ENSMUSG( | ENSMUSG( | 0.463653 | 8.26E-06 | YES | NO | NA |
| Ap4e1 | ENSMUSG( | ENSMUSG( | -0.00139 | 0.986203 | NO | NO | NA |
| Cox10 | ENSMUSG( | ENSMUSG( | 0.470865 | 4.93E-07 | YES | NO | NA |
| Dis3l2 | ENSMUSG( | ENSMUSG( | 0.23897 | 0.023876 | YES | NO | NA |
| Mrpl16 | ENSMUSG( | ENSMUSG( | 0.404493 | 9.53E-08 | YES | NO | NA |
| Ppp1r35 | ENSMUSG( | ENSMUSG( | -0.11671 | 0.194586 | NO | NO | NA |
| Lin9 | ENSMUSG( | ENSMUSG( | -0.04094 | 0.688983 | NO | NO | NA |
| Gas7 | ENSMUSG( | ENSMUSG( | -1.00914 | 2.81E-07 | YES | YES | Downregulated |
| Klhl18 | ENSMUSG( | ENSMUSG( | -0.19872 | 0.002501 | YES | NO | NA |
| Rnf214 | ENSMUSG( | ENSMUSG( | -0.32452 | 1.53E-05 | YES | NO | NA |
| Arfip2 | ENSMUSG( | ENSMUSG( | 0.19436 | 0.002658 | YES | NO | NA |
| Tmem192 | ENSMUSG( | ENSMUSG( | 0.212253 | 0.054431 | NO | NO | NA |
| Efl1 | ENSMUSG( | ENSMUSG( | 0.120504 | 0.225969 | NO | NO | NA |
| Mylip | ENSMUSG( | ENSMUSG( | -0.81542 | 5.15E-06 | YES | NO | NA |
| Hspa13 | ENSMUSG( | ENSMUSG( | -0.19676 | 0.031839 | YES | NO | NA |
| Smyd2 | ENSMUSG( | ENSMUSG( | 0.990989 | 1.06E-09 | YES | NO | NA |
| Tspan3 | ENSMUSG( | ENSMUSG( | -0.06164 | 0.683724 | NO | NO | NA |
| Phlpp2 | ENSMUSG( | ENSMUSG( | -0.0702 | 0.34275 | NO | NO | NA |
| Rfxap | ENSMUSG( | ENSMUSG( | -0.0099 | 0.942764 | NO | NO | NA |
| Mturn | ENSMUSG( | ENSMUSG( | -1.04204 | 4.13E-07 | YES | YES | Downregulated |
| Golph3l | ENSMUSG( | ENSMUSG( | -0.26193 | 0.004216 | YES | NO | NA |
| Mtx1 | ENSMUSG( | ENSMUSG( | 0.442149 | 1.12E-06 | YES | NO | NA |

| | | | | | | | |
|---|---|---|---|---|---|---|---|
| Cars2 | ENSMUSG( | ENSMUSG( | -0.14323 | 0.016104 | YES | NO | NA |
| Spata5 | ENSMUSG( | ENSMUSG( | -0.11694 | 0.152285 | NO | NO | NA |
| Gmppa | ENSMUSG( | ENSMUSG( | -0.14659 | 0.017519 | YES | NO | NA |
| Ccdc174 | ENSMUSG( | ENSMUSG( | 0.071834 | 0.473525 | NO | NO | NA |
| Stxbp3 | ENSMUSG( | ENSMUSG( | -0.36492 | 6.19E-07 | YES | NO | NA |
| Odr4 | ENSMUSG( | ENSMUSG( | 0.004987 | 0.974234 | NO | NO | NA |
| Cacfd1 | ENSMUSG( | ENSMUSG( | -0.18342 | 0.024018 | YES | NO | NA |
| Parp11 | ENSMUSG( | ENSMUSG( | -0.49789 | 5.93E-09 | YES | NO | NA |
| Sell | ENSMUSG( | ENSMUSG( | -0.07094 | 0.897193 | NO | NO | NA |
| Gm45140 | ENSMUSG( | ENSMUSG( | -0.40387 | 0.437949 | NO | NO | NA |
| Strn3 | ENSMUSG( | ENSMUSG( | -0.17082 | 0.027741 | YES | NO | NA |
| Vma21 | ENSMUSG( | ENSMUSG( | 0.126643 | 0.211198 | NO | NO | NA |
| Mtfmt | ENSMUSG( | ENSMUSG( | 0.033461 | 0.759182 | NO | NO | NA |
| Mavs | ENSMUSG( | ENSMUSG( | -0.45395 | 1.19E-08 | YES | NO | NA |
| Slf1 | ENSMUSG( | ENSMUSG( | 0.157259 | 0.153706 | NO | NO | NA |
| Ofd1 | ENSMUSG( | ENSMUSG( | 0.132884 | 0.158897 | NO | NO | NA |
| Siah2 | ENSMUSG( | ENSMUSG( | 0.104538 | 0.476831 | NO | NO | NA |
| Aatf | ENSMUSG( | ENSMUSG( | 0.355629 | 0.004303 | YES | NO | NA |
| Cenpl | ENSMUSG( | ENSMUSG( | 0.081225 | 0.322605 | NO | NO | NA |
| R3hcc1l | ENSMUSG( | ENSMUSG( | -0.08454 | 0.320474 | NO | NO | NA |
| Fbxl17 | ENSMUSG( | ENSMUSG( | -0.09114 | 0.509593 | NO | NO | NA |
| Dgat1 | ENSMUSG( | ENSMUSG( | -0.14124 | 0.126934 | NO | NO | NA |
| Mpc1 | ENSMUSG( | ENSMUSG( | 0.522757 | 1.12E-05 | YES | NO | NA |
| Nrarp | ENSMUSG( | ENSMUSG( | -0.72181 | 1.96E-08 | YES | NO | NA |
| Rabggta | ENSMUSG( | ENSMUSG( | -0.09114 | 0.251475 | NO | NO | NA |
| Abhd17b | ENSMUSG( | ENSMUSG( | -0.14613 | 0.126912 | NO | NO | NA |
| Phykpl | ENSMUSG( | ENSMUSG( | 0.300311 | 0.001556 | YES | NO | NA |
| Tfdp2 | ENSMUSG( | ENSMUSG( | 0.265736 | 0.101419 | NO | NO | NA |
| Ccdc127 | ENSMUSG( | ENSMUSG( | -0.09693 | 0.234786 | NO | NO | NA |
| Tm7sf3 | ENSMUSG( | ENSMUSG( | 0.100691 | 0.475214 | NO | NO | NA |
| Ccdc167 | ENSMUSG( | ENSMUSG( | -0.20739 | 0.008689 | YES | NO | NA |
| Tbc1d8b | ENSMUSG( | ENSMUSG( | -0.48141 | 7.40E-09 | YES | NO | NA |
| Bloc1s6 | ENSMUSG( | ENSMUSG( | -0.10123 | 0.38454 | NO | NO | NA |
| Ggct | ENSMUSG( | ENSMUSG( | -0.03741 | 0.755908 | NO | NO | NA |
| Ciao1 | ENSMUSG( | ENSMUSG( | -0.19991 | 0.000881 | YES | NO | NA |
| Lias | ENSMUSG( | ENSMUSG( | 0.384134 | 0.001242 | YES | NO | NA |
| Haus8 | ENSMUSG( | ENSMUSG( | -0.22478 | 0.00794 | YES | NO | NA |
| Pigx | ENSMUSG( | ENSMUSG( | -0.05612 | 0.603517 | NO | NO | NA |
| Jtb | ENSMUSG( | ENSMUSG( | 0.134722 | 0.113368 | NO | NO | NA |
| A930004J1 | ENSMUSG( | ENSMUSG( | -0.02234 | 0.862699 | NO | NO | NA |
| Ogfod1 | ENSMUSG( | ENSMUSG( | -0.19068 | 0.002365 | YES | NO | NA |
| Ankle1 | ENSMUSG( | ENSMUSG( | -0.03964 | 0.826557 | NO | NO | NA |
| Tut1 | ENSMUSG( | ENSMUSG( | 0.043858 | 0.638822 | NO | NO | NA |
| Vrk1 | ENSMUSG( | ENSMUSG( | 0.351697 | 0.004798 | YES | NO | NA |
| Rnf13 | ENSMUSG( | ENSMUSG( | -0.32955 | 1.96E-05 | YES | NO | NA |
| Tsen54 | ENSMUSG( | ENSMUSG( | 0.144443 | 0.10351 | NO | NO | NA |
| Tmem154 | ENSMUSG( | ENSMUSG( | 0.252871 | 0.043391 | YES | NO | NA |

| | | | | | | | |
|---|---|---|---|---|---|---|---|
| Htra2 | ENSMUSG( | ENSMUSG( | 0.400997 | 7.56E-05 | YES | NO | NA |
| Ccdc61 | ENSMUSG( | ENSMUSG( | 0.011881 | 0.927531 | NO | NO | NA |
| Cdc26 | ENSMUSG( | ENSMUSG( | -0.01745 | 0.87566 | NO | NO | NA |
| Rrp8 | ENSMUSG( | ENSMUSG( | 0.031881 | 0.738195 | NO | NO | NA |
| Cyb5a | ENSMUSG( | ENSMUSG( | 0.081989 | 0.299612 | NO | NO | NA |
| Nsl1 | ENSMUSG( | ENSMUSG( | 0.258197 | 0.001584 | YES | NO | NA |
| Myo1e | ENSMUSG( | ENSMUSG( | -0.53075 | 1.42E-05 | YES | NO | NA |
| Btg3 | ENSMUSG( | ENSMUSG( | -0.25771 | 0.007155 | YES | NO | NA |
| Tsr3 | ENSMUSG( | ENSMUSG( | -0.09662 | 0.240847 | NO | NO | NA |
| Csgalnact2 | ENSMUSG( | ENSMUSG( | 0.122557 | 0.442731 | NO | NO | NA |
| Cnep1r1 | ENSMUSG( | ENSMUSG( | -0.05943 | 0.554806 | NO | NO | NA |
| Foxred1 | ENSMUSG( | ENSMUSG( | 0.037569 | 0.645518 | NO | NO | NA |
| Timm17b | ENSMUSG( | ENSMUSG( | -0.01883 | 0.797821 | NO | NO | NA |
| Cep170 | ENSMUSG( | ENSMUSG( | -0.13939 | 0.078786 | NO | NO | NA |
| Gm16867 | ENSMUSG( | ENSMUSG( | 0.947597 | 8.84E-06 | YES | NO | NA |
| Neu1 | ENSMUSG( | ENSMUSG( | 0.081914 | 0.511559 | NO | NO | NA |
| Noc3l | ENSMUSG( | ENSMUSG( | 0.271727 | 0.047099 | YES | NO | NA |
| Gatd3a | ENSMUSG( | ENSMUSG( | -0.059 | 0.502427 | NO | NO | NA |
| Bloc1s2 | ENSMUSG( | ENSMUSG( | 0.192825 | 0.055507 | NO | NO | NA |
| Psph | ENSMUSG( | ENSMUSG( | 0.053296 | 0.509559 | NO | NO | NA |
| Tchp | ENSMUSG( | ENSMUSG( | -0.2393 | 0.069886 | NO | NO | NA |
| Tmem115 | ENSMUSG( | ENSMUSG( | -0.20328 | 7.66E-05 | YES | NO | NA |
| Erlec1 | ENSMUSG( | ENSMUSG( | 0.131784 | 0.343744 | NO | NO | NA |
| Il4ra | ENSMUSG( | ENSMUSG( | -0.05541 | 0.33098 | NO | NO | NA |
| Usp45 | ENSMUSG( | ENSMUSG( | 0.05953 | 0.584807 | NO | NO | NA |
| Tceanc2 | ENSMUSG( | ENSMUSG( | -0.2174 | 0.009269 | YES | NO | NA |
| Cisd1 | ENSMUSG( | ENSMUSG( | 0.322804 | 0.001638 | YES | NO | NA |
| Dohh | ENSMUSG( | ENSMUSG( | 0.406644 | 7.37E-07 | YES | NO | NA |
| Trappc13 | ENSMUSG( | ENSMUSG( | -0.32521 | 2.98E-06 | YES | NO | NA |
| Ptcd2 | ENSMUSG( | ENSMUSG( | 0.124768 | 0.127533 | NO | NO | NA |
| Slc35c2 | ENSMUSG( | ENSMUSG( | 0.069636 | 0.442109 | NO | NO | NA |
| Dubr | ENSMUSG( | ENSMUSG( | -0.56058 | 1.23E-11 | YES | NO | NA |
| Ift20 | ENSMUSG( | ENSMUSG( | 0.062469 | 0.527709 | NO | NO | NA |
| Trex1 | ENSMUSG( | ENSMUSG( | -0.69283 | 1.33E-12 | YES | NO | NA |
| Nrm | ENSMUSG( | ENSMUSG( | 0.166911 | 0.018507 | YES | NO | NA |
| Terf1 | ENSMUSG( | ENSMUSG( | -0.23899 | 0.021827 | YES | NO | NA |
| Iqce | ENSMUSG( | ENSMUSG( | -0.19001 | 0.095173 | NO | NO | NA |
| Glb1 | ENSMUSG( | ENSMUSG( | -0.23076 | 0.020977 | YES | NO | NA |
| Rpf2 | ENSMUSG( | ENSMUSG( | 0.595062 | 0.000159 | YES | NO | NA |
| Tesk1 | ENSMUSG( | ENSMUSG( | -0.35666 | 0.00141 | YES | NO | NA |
| Eefsec | ENSMUSG( | ENSMUSG( | 0.277972 | 0.01081 | YES | NO | NA |
| Mrpl49 | ENSMUSG( | ENSMUSG( | 0.035267 | 0.58825 | NO | NO | NA |
| Spint2 | ENSMUSG( | ENSMUSG( | 0.108195 | 0.294113 | NO | NO | NA |
| Mrps36 | ENSMUSG( | ENSMUSG( | 0.192979 | 0.034705 | YES | NO | NA |
| 1810058I2 | ENSMUSG( | ENSMUSG( | -0.12889 | 0.243638 | NO | NO | NA |
| Gm49342 | ENSMUSG( | ENSMUSG( | -0.06596 | 0.677417 | NO | NO | NA |
| Pask | ENSMUSG( | ENSMUSG( | 0.202447 | 0.005092 | YES | NO | NA |

| | | | | | | | |
|---|---|---|---|---|---|---|---|
| Lztfl1 | ENSMUSG( | ENSMUSG( | 0.686971 | 0.00185 | YES | NO | NA |
| Scml4 | ENSMUSG( | ENSMUSG( | -1.10639 | 6.13E-10 | YES | YES | Downregulated |
| Klrc2 | ENSMUSG( | ENSMUSG( | -0.54587 | 9.16E-06 | YES | NO | NA |
| Rnf8 | ENSMUSG( | ENSMUSG( | 0.04723 | 0.710988 | NO | NO | NA |
| H4c9 | ENSMUSG( | ENSMUSG( | 0.823688 | 9.87E-07 | YES | NO | NA |
| Tab3 | ENSMUSG( | ENSMUSG( | 0.032943 | 0.826335 | NO | NO | NA |
| Pole2 | ENSMUSG( | ENSMUSG( | 0.074963 | 0.397233 | NO | NO | NA |
| Slco4a1 | ENSMUSG( | ENSMUSG( | 0.747718 | 3.91E-07 | YES | NO | NA |
| Rnf5 | ENSMUSG( | ENSMUSG( | -0.29304 | 0.000288 | YES | NO | NA |
| Cep76 | ENSMUSG( | ENSMUSG( | 0.149651 | 0.08615 | NO | NO | NA |
| Btbd9 | ENSMUSG( | ENSMUSG( | -0.26396 | 6.66E-07 | YES | NO | NA |
| Abraxas2 | ENSMUSG( | ENSMUSG( | -0.13726 | 0.028516 | YES | NO | NA |
| Pde4b | ENSMUSG( | ENSMUSG( | -0.37255 | 0.000808 | YES | NO | NA |
| Stx12 | ENSMUSG( | ENSMUSG( | -0.14703 | 0.040567 | YES | NO | NA |
| Rnft1 | ENSMUSG( | ENSMUSG( | -0.1005 | 0.310617 | NO | NO | NA |
| Bola3 | ENSMUSG( | ENSMUSG( | 0.324617 | 0.0013 | YES | NO | NA |
| Rrp15 | ENSMUSG( | ENSMUSG( | 0.730297 | 5.38E-06 | YES | NO | NA |
| Car2 | ENSMUSG( | ENSMUSG( | 0.331753 | 0.532803 | NO | NO | NA |
| 9930021J( | ENSMUSG( | ENSMUSG( | -0.17967 | 0.034705 | YES | NO | NA |
| Primpol | ENSMUSG( | ENSMUSG( | -0.40489 | 4.14E-06 | YES | NO | NA |
| Gskip | ENSMUSG( | ENSMUSG( | -0.06619 | 0.513868 | NO | NO | NA |
| Wdr11 | ENSMUSG( | ENSMUSG( | -0.00311 | 0.976785 | NO | NO | NA |
| Rnf121 | ENSMUSG( | ENSMUSG( | -0.31268 | 1.91E-05 | YES | NO | NA |
| Mynn | ENSMUSG( | ENSMUSG( | -0.22639 | 0.02669 | YES | NO | NA |
| Ppp2r3c | ENSMUSG( | ENSMUSG( | 0.110043 | 0.25172 | NO | NO | NA |
| Med7 | ENSMUSG( | ENSMUSG( | -0.03358 | 0.829598 | NO | NO | NA |
| Ap3m2 | ENSMUSG( | ENSMUSG( | -0.08912 | 0.522804 | NO | NO | NA |
| Hdgfl3 | ENSMUSG( | ENSMUSG( | -0.18871 | 0.157988 | NO | NO | NA |
| Cep89 | ENSMUSG( | ENSMUSG( | 0.433438 | 0.000472 | YES | NO | NA |
| ENSMUSG( | ENSMUSG( | ENSMUSG( | 0.438877 | 0.027823 | YES | NO | NA |
| Pik3c2a | ENSMUSG( | ENSMUSG( | -0.19259 | 0.124547 | NO | NO | NA |
| Psmd10 | ENSMUSG( | ENSMUSG( | -0.02042 | 0.859542 | NO | NO | NA |
| Pnpla7 | ENSMUSG( | ENSMUSG( | -0.28349 | 0.048394 | YES | NO | NA |
| Poc1b | ENSMUSG( | ENSMUSG( | 0.08469 | 0.357007 | NO | NO | NA |
| Fut11 | ENSMUSG( | ENSMUSG( | -0.26789 | 2.04E-07 | YES | NO | NA |
| Gm12840 | ENSMUSG( | ENSMUSG( | 0.365004 | 0.071089 | NO | NO | NA |
| Mbtps2 | ENSMUSG( | ENSMUSG( | 0.020403 | 0.863447 | NO | NO | NA |
| Gm2000 | ENSMUSG( | ENSMUSG( | 0.653501 | 7.88E-06 | YES | NO | NA |
| Uqcc2 | ENSMUSG( | ENSMUSG( | 0.250095 | 0.000135 | YES | NO | NA |
| Syk | ENSMUSG( | ENSMUSG( | -0.95039 | 2.49E-17 | YES | NO | NA |
| Frs2 | ENSMUSG( | ENSMUSG( | -0.31688 | 0.003953 | YES | NO | NA |
| Slc12a4 | ENSMUSG( | ENSMUSG( | -0.19493 | 0.002108 | YES | NO | NA |
| Hars2 | ENSMUSG( | ENSMUSG( | -0.04831 | 0.545216 | NO | NO | NA |
| Ddx60 | ENSMUSG( | ENSMUSG( | -1.11025 | 2.44E-14 | YES | YES | Downregulated |
| Ganc | ENSMUSG( | ENSMUSG( | -0.17363 | 0.05121 | NO | NO | NA |
| Snrnp27 | ENSMUSG( | ENSMUSG( | -0.07759 | 0.613606 | NO | NO | NA |
| Dph3 | ENSMUSG( | ENSMUSG( | 0.185087 | 0.000876 | YES | NO | NA |

| | | | | | | | |
|---|---|---|---|---|---|---|---|
| B3gnt5 | ENSMUSG( | ENSMUSG( | -0.59795 | 5.92E-06 | YES | NO | NA |
| Slco3a1 | ENSMUSG( | ENSMUSG( | -0.21632 | 0.006149 | YES | NO | NA |
| Rnf149 | ENSMUSG( | ENSMUSG( | -0.13784 | 0.065051 | NO | NO | NA |
| Bdh1 | ENSMUSG( | ENSMUSG( | 0.285478 | 0.013647 | YES | NO | NA |
| Pimreg | ENSMUSG( | ENSMUSG( | 0.356889 | 0.001791 | YES | NO | NA |
| Taf1c | ENSMUSG( | ENSMUSG( | -0.34678 | 8.34E-06 | YES | NO | NA |
| Ptgr2 | ENSMUSG( | ENSMUSG( | 0.042584 | 0.699813 | NO | NO | NA |
| Urm1 | ENSMUSG( | ENSMUSG( | 0.339591 | 7.08E-05 | YES | NO | NA |
| Actr5 | ENSMUSG( | ENSMUSG( | 0.081114 | 0.226058 | NO | NO | NA |
| Lancl1 | ENSMUSG( | ENSMUSG( | 0.281792 | 0.021238 | YES | NO | NA |
| Oard1 | ENSMUSG( | ENSMUSG( | -0.06726 | 0.465954 | NO | NO | NA |
| Rad54b | ENSMUSG( | ENSMUSG( | 0.095881 | 0.248793 | NO | NO | NA |
| Rab29 | ENSMUSG( | ENSMUSG( | -0.33857 | 0.000152 | YES | NO | NA |
| Abitram | ENSMUSG( | ENSMUSG( | 0.21588 | 0.016752 | YES | NO | NA |
| Zfp513 | ENSMUSG( | ENSMUSG( | -0.33001 | 2.10E-05 | YES | NO | NA |
| Ube2q2 | ENSMUSG( | ENSMUSG( | 0.069309 | 0.483451 | NO | NO | NA |
| Sppl2b | ENSMUSG( | ENSMUSG( | 0.100202 | 0.496187 | NO | NO | NA |
| Tpgs1 | ENSMUSG( | ENSMUSG( | 0.19471 | 0.075277 | NO | NO | NA |
| Tomm40l | ENSMUSG( | ENSMUSG( | 0.218345 | 0.000275 | YES | NO | NA |
| Exo1 | ENSMUSG( | ENSMUSG( | 0.372806 | 0.000352 | YES | NO | NA |
| Prmt3 | ENSMUSG( | ENSMUSG( | 0.174788 | 0.026882 | YES | NO | NA |
| Clec16a | ENSMUSG( | ENSMUSG( | -0.29118 | 0.000125 | YES | NO | NA |
| Sntb1 | ENSMUSG( | ENSMUSG( | -0.83868 | 3.79E-27 | YES | NO | NA |
| Top3a | ENSMUSG( | ENSMUSG( | 0.129424 | 0.010751 | YES | NO | NA |
| Slc25a20 | ENSMUSG( | ENSMUSG( | -0.10441 | 0.435434 | NO | NO | NA |
| Slc25a38 | ENSMUSG( | ENSMUSG( | 0.003695 | 0.978564 | NO | NO | NA |
| Fosl2 | ENSMUSG( | ENSMUSG( | -0.48387 | 9.31E-05 | YES | NO | NA |
| Cog6 | ENSMUSG( | ENSMUSG( | -0.19235 | 0.018666 | YES | NO | NA |
| Vps37c | ENSMUSG( | ENSMUSG( | -0.06626 | 0.408456 | NO | NO | NA |
| Mgst2 | ENSMUSG( | ENSMUSG( | -0.37982 | 0.001433 | YES | NO | NA |
| Cd2ap | ENSMUSG( | ENSMUSG( | 0.743573 | 4.19E-08 | YES | NO | NA |
| Pcbp4 | ENSMUSG( | ENSMUSG( | -0.18082 | 0.004094 | YES | NO | NA |
| Rnaseh2a | ENSMUSG( | ENSMUSG( | 0.171563 | 0.029851 | YES | NO | NA |
| Apex1 | ENSMUSG( | ENSMUSG( | 0.939783 | 4.87E-09 | YES | NO | NA |
| Rpap3 | ENSMUSG( | ENSMUSG( | 0.490031 | 5.07E-06 | YES | NO | NA |
| Rbm18 | ENSMUSG( | ENSMUSG( | -0.22467 | 0.001152 | YES | NO | NA |
| Zfp275 | ENSMUSG( | ENSMUSG( | -0.51198 | 2.34E-07 | YES | NO | NA |
| Cwc27 | ENSMUSG( | ENSMUSG( | 0.318524 | 0.013895 | YES | NO | NA |
| Ankra2 | ENSMUSG( | ENSMUSG( | -0.17722 | 0.112361 | NO | NO | NA |
| Ctsz | ENSMUSG( | ENSMUSG( | 0.538216 | 0.000123 | YES | NO | NA |
| Dedd | ENSMUSG( | ENSMUSG( | -0.08882 | 0.182865 | NO | NO | NA |
| Tbc1d10a | ENSMUSG( | ENSMUSG( | -0.10815 | 0.221618 | NO | NO | NA |
| Tmem161; | ENSMUSG( | ENSMUSG( | -0.05469 | 0.515031 | NO | NO | NA |
| Gpatch2 | ENSMUSG( | ENSMUSG( | -0.30449 | 0.001146 | YES | NO | NA |
| Klf7 | ENSMUSG( | ENSMUSG( | 0.043905 | 0.661917 | NO | NO | NA |
| Axl | ENSMUSG( | ENSMUSG( | 0.915319 | 2.09E-05 | YES | NO | NA |
| Bid | ENSMUSG( | ENSMUSG( | -0.09718 | 0.311391 | NO | NO | NA |

| | | | | | | | |
|---|---|---|---|---|---|---|---|
| Zfp407 | ENSMUSG( | ENSMUSG( | -0.27901 | 0.000896 | YES | NO | NA |
| Phactr4 | ENSMUSG( | ENSMUSG( | -0.11394 | 0.308752 | NO | NO | NA |
| Rhbdd1 | ENSMUSG( | ENSMUSG( | -0.11675 | 0.116764 | NO | NO | NA |
| Commd10 | ENSMUSG( | ENSMUSG( | 0.069589 | 0.314912 | NO | NO | NA |
| Zfp639 | ENSMUSG( | ENSMUSG( | 0.138067 | 0.05753 | NO | NO | NA |
| Exd2 | ENSMUSG( | ENSMUSG( | -0.18612 | 0.000858 | YES | NO | NA |
| Prmt6 | ENSMUSG( | ENSMUSG( | 0.164732 | 0.088785 | NO | NO | NA |
| 3830406C: | ENSMUSG( | ENSMUSG( | 0.056884 | 0.567511 | NO | NO | NA |
| Gtf2a2 | ENSMUSG( | ENSMUSG( | 0.181081 | 0.039829 | YES | NO | NA |
| Nkapd1 | ENSMUSG( | ENSMUSG( | 0.173391 | 0.088004 | NO | NO | NA |
| Nxt1 | ENSMUSG( | ENSMUSG( | 0.144736 | 0.004791 | YES | NO | NA |
| Hax1 | ENSMUSG( | ENSMUSG( | 0.11055 | 0.282001 | NO | NO | NA |
| Snapc3 | ENSMUSG( | ENSMUSG( | -0.2507 | 0.026344 | YES | NO | NA |
| Dtnbp1 | ENSMUSG( | ENSMUSG( | 0.127615 | 0.281448 | NO | NO | NA |
| Slc39a14 | ENSMUSG( | ENSMUSG( | 0.464671 | 5.20E-06 | YES | NO | NA |
| Vipas39 | ENSMUSG( | ENSMUSG( | -0.05599 | 0.587391 | NO | NO | NA |
| Ttc38 | ENSMUSG( | ENSMUSG( | -0.43682 | 4.51E-05 | YES | NO | NA |
| Uxs1 | ENSMUSG( | ENSMUSG( | 0.631423 | 1.17E-08 | YES | NO | NA |
| Cenph | ENSMUSG( | ENSMUSG( | 0.536693 | 2.08E-05 | YES | NO | NA |
| Mrps17 | ENSMUSG( | ENSMUSG( | 0.178487 | 0.082419 | NO | NO | NA |
| Ppm1j | ENSMUSG( | ENSMUSG( | -0.4648 | 0.000171 | YES | NO | NA |
| Wasl | ENSMUSG( | ENSMUSG( | -0.4 | 0.001225 | YES | NO | NA |
| 1700123O | ENSMUSG( | ENSMUSG( | -0.01483 | 0.822566 | NO | NO | NA |
| Cgas | ENSMUSG( | ENSMUSG( | -0.2148 | 0.094428 | NO | NO | NA |
| Ccdc43 | ENSMUSG( | ENSMUSG( | 0.18142 | 0.041983 | YES | NO | NA |
| Acsf2 | ENSMUSG( | ENSMUSG( | -0.3116 | 0.000343 | YES | NO | NA |
| Camsap1 | ENSMUSG( | ENSMUSG( | -0.14971 | 0.12664 | NO | NO | NA |
| Ppwd1 | ENSMUSG( | ENSMUSG( | -0.2286 | 0.01434 | YES | NO | NA |
| Clock | ENSMUSG( | ENSMUSG( | -0.0505 | 0.610095 | NO | NO | NA |
| Parp3 | ENSMUSG( | ENSMUSG( | -0.78754 | 1.45E-12 | YES | NO | NA |
| Ercc6l | ENSMUSG( | ENSMUSG( | 0.074282 | 0.551123 | NO | NO | NA |
| Tpd52l2 | ENSMUSG( | ENSMUSG( | -0.03356 | 0.699123 | NO | NO | NA |
| Fam234a | ENSMUSG( | ENSMUSG( | -0.03334 | 0.865953 | NO | NO | NA |
| Thop1 | ENSMUSG( | ENSMUSG( | 0.920814 | 6.41E-07 | YES | NO | NA |
| Dnah14 | ENSMUSG( | ENSMUSG( | 0.105273 | 0.374748 | NO | NO | NA |
| Abt1 | ENSMUSG( | ENSMUSG( | 0.008839 | 0.944823 | NO | NO | NA |
| Cep20 | ENSMUSG( | ENSMUSG( | 0.004579 | 0.970155 | NO | NO | NA |
| Btbd19 | ENSMUSG( | ENSMUSG( | 0.204229 | 0.074313 | NO | NO | NA |
| 4833407H | ENSMUSG( | ENSMUSG( | -0.47605 | 3.28E-07 | YES | NO | NA |
| Marveld1 | ENSMUSG( | ENSMUSG( | -0.49672 | 3.90E-05 | YES | NO | NA |
| Eri2 | ENSMUSG( | ENSMUSG( | 0.077529 | 0.423196 | NO | NO | NA |
| Coq2 | ENSMUSG( | ENSMUSG( | 0.161262 | 0.070464 | NO | NO | NA |
| Oaz2 | ENSMUSG( | ENSMUSG( | -0.23321 | 0.010973 | YES | NO | NA |
| Wdr4 | ENSMUSG( | ENSMUSG( | 0.611552 | 8.51E-08 | YES | NO | NA |
| Bbip1 | ENSMUSG( | ENSMUSG( | -0.33249 | 0.002388 | YES | NO | NA |
| A430093F: | ENSMUSG( | ENSMUSG( | -0.25113 | 0.006615 | YES | NO | NA |
| Thada | ENSMUSG( | ENSMUSG( | -0.04225 | 0.572582 | NO | NO | NA |

| | | | | | | | |
|---|---|---|---|---|---|---|---|
| Osgep | ENSMUSG( | ENSMUSG( | 0.248828 | 0.002071 | YES | NO | NA |
| Dctpp1 | ENSMUSG( | ENSMUSG( | 0.483309 | 5.99E-05 | YES | NO | NA |
| Isg15 | ENSMUSG( | ENSMUSG( | -1.07609 | 1.27E-09 | YES | YES | Downregulated |
| Dapp1 | ENSMUSG( | ENSMUSG( | -0.45859 | 2.45E-09 | YES | NO | NA |
| Hlcs | ENSMUSG( | ENSMUSG( | 0.051494 | 0.586519 | NO | NO | NA |
| Zfp397 | ENSMUSG( | ENSMUSG( | -0.26052 | 0.020415 | YES | NO | NA |
| Hsdl2 | ENSMUSG( | ENSMUSG( | 0.06898 | 0.491395 | NO | NO | NA |
| Fbxl19 | ENSMUSG( | ENSMUSG( | 0.009191 | 0.928155 | NO | NO | NA |
| Poc1a | ENSMUSG( | ENSMUSG( | 0.498222 | 2.61E-06 | YES | NO | NA |
| Ankrd46 | ENSMUSG( | ENSMUSG( | -0.09727 | 0.352514 | NO | NO | NA |
| Rassf3 | ENSMUSG( | ENSMUSG( | -0.23239 | 0.01355 | YES | NO | NA |
| Tpra1 | ENSMUSG( | ENSMUSG( | -0.33548 | 3.59E-05 | YES | NO | NA |
| Cep162 | ENSMUSG( | ENSMUSG( | -0.24552 | 0.02605 | YES | NO | NA |
| Arhgef12 | ENSMUSG( | ENSMUSG( | 0.309733 | 0.112826 | NO | NO | NA |
| Lrrc40 | ENSMUSG( | ENSMUSG( | 0.563364 | 1.37E-07 | YES | NO | NA |
| Ryk | ENSMUSG( | ENSMUSG( | -0.0399 | 0.734336 | NO | NO | NA |
| Zfp53 | ENSMUSG( | ENSMUSG( | -0.16405 | 0.250943 | NO | NO | NA |
| Dpm1 | ENSMUSG( | ENSMUSG( | -0.07782 | 0.445886 | NO | NO | NA |
| Eme1 | ENSMUSG( | ENSMUSG( | 0.23786 | 0.019231 | YES | NO | NA |
| Fubp3 | ENSMUSG( | ENSMUSG( | 0.458816 | 8.28E-05 | YES | NO | NA |
| Mocs2 | ENSMUSG( | ENSMUSG( | 0.152803 | 0.108608 | NO | NO | NA |
| Sdccag8 | ENSMUSG( | ENSMUSG( | -0.04971 | 0.704289 | NO | NO | NA |
| Gfus | ENSMUSG( | ENSMUSG( | 0.0577 | 0.444093 | NO | NO | NA |
| Map6 | ENSMUSG( | ENSMUSG( | -0.7748 | 1.49E-08 | YES | NO | NA |
| Lrrcc1 | ENSMUSG( | ENSMUSG( | -0.18423 | 0.029316 | YES | NO | NA |
| Gvin-ps4 | ENSMUSG( | ENSMUSG( | -0.676 | 5.57E-05 | YES | NO | NA |
| Yrdc | ENSMUSG( | ENSMUSG( | 0.016592 | 0.869991 | NO | NO | NA |
| Ttc39c | ENSMUSG( | ENSMUSG( | -0.37589 | 0.004414 | YES | NO | NA |
| Rab3ip | ENSMUSG( | ENSMUSG( | -0.42299 | 9.53E-06 | YES | NO | NA |
| Atn1 | ENSMUSG( | ENSMUSG( | -0.14149 | 0.154197 | NO | NO | NA |
| Rpgrip1 | ENSMUSG( | ENSMUSG( | 0.226732 | 0.005372 | YES | NO | NA |
| Acadm | ENSMUSG( | ENSMUSG( | 0.147618 | 0.115839 | NO | NO | NA |
| Vmn2r97 | ENSMUSG( | ENSMUSG( | -0.80418 | 9.61E-13 | YES | NO | NA |
| Tssc4 | ENSMUSG( | ENSMUSG( | 0.272975 | 0.001355 | YES | NO | NA |
| Sdcbp2 | ENSMUSG( | ENSMUSG( | -0.72348 | 1.42E-06 | YES | NO | NA |
| Rmdn3 | ENSMUSG( | ENSMUSG( | 0.188451 | 0.026783 | YES | NO | NA |
| Gpr89 | ENSMUSG( | ENSMUSG( | 0.013042 | 0.892196 | NO | NO | NA |
| Ube2f | ENSMUSG( | ENSMUSG( | -0.26864 | 0.003074 | YES | NO | NA |
| Rtn4rl1 | ENSMUSG( | ENSMUSG( | -1.26477 | 2.95E-09 | YES | YES | Downregulated |
| Acox1 | ENSMUSG( | ENSMUSG( | -0.16899 | 0.224118 | NO | NO | NA |
| Anxa1 | ENSMUSG( | ENSMUSG( | 1.600418 | 1.22E-08 | YES | YES | Upregulated |
| Entpd4b | ENSMUSG( | ENSMUSG( | 0.353758 | 0.211633 | NO | NO | NA |
| Usb1 | ENSMUSG( | ENSMUSG( | -0.31396 | 9.96E-08 | YES | NO | NA |
| Pja2 | ENSMUSG( | ENSMUSG( | -0.06534 | 0.609081 | NO | NO | NA |
| Mrpl21 | ENSMUSG( | ENSMUSG( | 0.199523 | 0.010517 | YES | NO | NA |
| Zfp84 | ENSMUSG( | ENSMUSG( | -0.36709 | 4.11E-06 | YES | NO | NA |
| Ndufs5 | ENSMUSG( | ENSMUSG( | 0.139131 | 0.177382 | NO | NO | NA |

| | | | | | | | |
|---|---|---|---|---|---|---|---|
| Cwf19l2 | ENSMUSG( | ENSMUSG( | -0.05088 | 0.624377 | NO | NO | NA |
| Gldc | ENSMUSG( | ENSMUSG( | 1.560571 | 5.82E-12 | YES | YES | Upregulated |
| Snai3 | ENSMUSG( | ENSMUSG( | -0.40734 | 8.48E-07 | YES | NO | NA |
| Trbv15 | ENSMUSG( | ENSMUSG( | -0.49435 | 0.481807 | NO | NO | NA |
| Serhl | ENSMUSG( | ENSMUSG( | 0.065389 | 0.638601 | NO | NO | NA |
| Ndst2 | ENSMUSG( | ENSMUSG( | -0.1416 | 0.247715 | NO | NO | NA |
| Ano10 | ENSMUSG( | ENSMUSG( | 0.461557 | 0.003858 | YES | NO | NA |
| Dusp10 | ENSMUSG( | ENSMUSG( | -0.31663 | 5.13E-05 | YES | NO | NA |
| Cnpy4 | ENSMUSG( | ENSMUSG( | -0.16004 | 0.208027 | NO | NO | NA |
| Dcun1d4 | ENSMUSG( | ENSMUSG( | -0.77139 | 1.37E-06 | YES | NO | NA |
| Surf2 | ENSMUSG( | ENSMUSG( | 0.221012 | 0.018453 | YES | NO | NA |
| Rundc1 | ENSMUSG( | ENSMUSG( | -0.14207 | 0.055365 | NO | NO | NA |
| Cenpt | ENSMUSG( | ENSMUSG( | 0.133701 | 0.081419 | NO | NO | NA |
| Usp33 | ENSMUSG( | ENSMUSG( | 0.214636 | 0.005607 | YES | NO | NA |
| Zcchc18 | ENSMUSG( | ENSMUSG( | -0.92225 | 6.32E-12 | YES | NO | NA |
| Klhl21 | ENSMUSG( | ENSMUSG( | -0.33551 | 0.000287 | YES | NO | NA |
| Dhx34 | ENSMUSG( | ENSMUSG( | -0.37598 | 3.43E-08 | YES | NO | NA |
| Nrbf2 | ENSMUSG( | ENSMUSG( | -0.17908 | 0.006171 | YES | NO | NA |
| Tbck | ENSMUSG( | ENSMUSG( | -0.46918 | 2.54E-08 | YES | NO | NA |
| 1110038F: | ENSMUSG( | ENSMUSG( | 0.204046 | 0.003373 | YES | NO | NA |
| Pop1 | ENSMUSG( | ENSMUSG( | 0.220601 | 0.034341 | YES | NO | NA |
| Dipk2a | ENSMUSG( | ENSMUSG( | 0.088469 | 0.309644 | NO | NO | NA |
| Crocc | ENSMUSG( | ENSMUSG( | 0.148354 | 0.166736 | NO | NO | NA |
| Mettl6 | ENSMUSG( | ENSMUSG( | 0.106207 | 0.202542 | NO | NO | NA |
| Ppt2 | ENSMUSG( | ENSMUSG( | -0.00909 | 0.950071 | NO | NO | NA |
| Zdhhc15 | ENSMUSG( | ENSMUSG( | -0.55932 | 5.38E-07 | YES | NO | NA |
| Abcb8 | ENSMUSG( | ENSMUSG( | 0.632858 | 9.75E-05 | YES | NO | NA |
| Plin3 | ENSMUSG( | ENSMUSG( | 0.83274 | 5.27E-07 | YES | NO | NA |
| Taf11 | ENSMUSG( | ENSMUSG( | 0.226662 | 0.044559 | YES | NO | NA |
| Lsm2 | ENSMUSG( | ENSMUSG( | 0.183647 | 0.082363 | NO | NO | NA |
| Pdcd7 | ENSMUSG( | ENSMUSG( | -0.05633 | 0.589529 | NO | NO | NA |
| Gm7964 | ENSMUSG( | ENSMUSG( | -0.00731 | 0.961918 | NO | NO | NA |
| Rbis | ENSMUSG( | ENSMUSG( | 0.259998 | 0.123367 | NO | NO | NA |
| Larp1b | ENSMUSG( | ENSMUSG( | 0.382323 | 0.000362 | YES | NO | NA |
| Cd24a | ENSMUSG( | ENSMUSG( | 1.255252 | 1.84E-05 | YES | YES | Upregulated |
| Snrnp25 | ENSMUSG( | ENSMUSG( | 0.388659 | 1.52E-05 | YES | NO | NA |
| Cep170b | ENSMUSG( | ENSMUSG( | -0.11376 | 0.077477 | NO | NO | NA |
| Adk | ENSMUSG( | ENSMUSG( | 0.966754 | 1.98E-09 | YES | NO | NA |
| Taf13 | ENSMUSG( | ENSMUSG( | -0.05901 | 0.712283 | NO | NO | NA |
| Rrm2b | ENSMUSG( | ENSMUSG( | -0.31472 | 8.90E-05 | YES | NO | NA |
| Orc5 | ENSMUSG( | ENSMUSG( | 0.109147 | 0.285194 | NO | NO | NA |
| Zbtb41 | ENSMUSG( | ENSMUSG( | -0.24892 | 0.003056 | YES | NO | NA |
| Recql5 | ENSMUSG( | ENSMUSG( | -0.14572 | 0.225036 | NO | NO | NA |
| Cops7a | ENSMUSG( | ENSMUSG( | 0.124583 | 0.086218 | NO | NO | NA |
| Tfe3 | ENSMUSG( | ENSMUSG( | 0.003734 | 0.957155 | NO | NO | NA |
| Rttn | ENSMUSG( | ENSMUSG( | 0.045511 | 0.711222 | NO | NO | NA |
| Gm36723 | ENSMUSG( | ENSMUSG( | -0.26658 | 0.223588 | NO | NO | NA |

| | | | | | | | |
|---|---|---|---|---|---|---|---|
| AI413582 | ENSMUSG( | ENSMUSG( | -0.10409 | 0.444352 | NO | NO | NA |
| Cebpb | ENSMUSG( | ENSMUSG( | 0.332315 | 0.008494 | YES | NO | NA |
| Cep63 | ENSMUSG( | ENSMUSG( | -0.02775 | 0.771808 | NO | NO | NA |
| Ptger1 | ENSMUSG( | ENSMUSG( | -0.11414 | 0.361843 | NO | NO | NA |
| Atg9a | ENSMUSG( | ENSMUSG( | -0.03717 | 0.81143 | NO | NO | NA |
| Fibp | ENSMUSG( | ENSMUSG( | -0.24405 | 0.025988 | YES | NO | NA |
| Pias2 | ENSMUSG( | ENSMUSG( | 0.155354 | 0.163732 | NO | NO | NA |
| Stx7 | ENSMUSG( | ENSMUSG( | -0.32369 | 0.001877 | YES | NO | NA |
| Appl2 | ENSMUSG( | ENSMUSG( | -0.12996 | 0.210595 | NO | NO | NA |
| Cep95 | ENSMUSG( | ENSMUSG( | -0.27107 | 0.004642 | YES | NO | NA |
| Gm16042 | ENSMUSG( | ENSMUSG( | -0.39743 | 0.011882 | YES | NO | NA |
| Commd6 | ENSMUSG( | ENSMUSG( | -0.09608 | 0.398195 | NO | NO | NA |
| Tmed4 | ENSMUSG( | ENSMUSG( | 0.01473 | 0.888811 | NO | NO | NA |
| Gtpbp6 | ENSMUSG( | ENSMUSG( | -0.16669 | 0.096024 | NO | NO | NA |
| Impact | ENSMUSG( | ENSMUSG( | -0.14016 | 0.235641 | NO | NO | NA |
| Lrrc45 | ENSMUSG( | ENSMUSG( | -0.15015 | 0.215273 | NO | NO | NA |
| Mtfr1 | ENSMUSG( | ENSMUSG( | -0.04024 | 0.552587 | NO | NO | NA |
| Thyn1 | ENSMUSG( | ENSMUSG( | 0.428037 | 2.84E-06 | YES | NO | NA |
| Thtpa | ENSMUSG( | ENSMUSG( | -0.12134 | 0.284468 | NO | NO | NA |
| Fbxo34 | ENSMUSG( | ENSMUSG( | -0.03772 | 0.701067 | NO | NO | NA |
| Mafg | ENSMUSG( | ENSMUSG( | 0.230939 | 0.002877 | YES | NO | NA |
| Gtf2e1 | ENSMUSG( | ENSMUSG( | 0.052778 | 0.55111 | NO | NO | NA |
| Ap4b1 | ENSMUSG( | ENSMUSG( | 0.064795 | 0.489506 | NO | NO | NA |
| Amdhd2 | ENSMUSG( | ENSMUSG( | -0.20547 | 0.003881 | YES | NO | NA |
| Cdyl | ENSMUSG( | ENSMUSG( | -0.21566 | 0.011419 | YES | NO | NA |
| Zbed3 | ENSMUSG( | ENSMUSG( | 0.331589 | 0.003348 | YES | NO | NA |
| Cd200r4 | ENSMUSG( | ENSMUSG( | -1.43666 | 6.84E-09 | YES | YES | Downregulated |
| Nipsnap3b | ENSMUSG( | ENSMUSG( | -0.11735 | 0.320056 | NO | NO | NA |
| Dzip3 | ENSMUSG( | ENSMUSG( | -0.66676 | 1.82E-08 | YES | NO | NA |
| Edrf1 | ENSMUSG( | ENSMUSG( | -0.16649 | 0.034679 | YES | NO | NA |
| Rp2 | ENSMUSG( | ENSMUSG( | -0.12864 | 0.18855 | NO | NO | NA |
| Lmbrd1 | ENSMUSG( | ENSMUSG( | -0.20459 | 0.058632 | NO | NO | NA |
| Ankmy2 | ENSMUSG( | ENSMUSG( | 0.00448 | 0.955293 | NO | NO | NA |
| Nhlrc3 | ENSMUSG( | ENSMUSG( | -0.46035 | 3.52E-06 | YES | NO | NA |
| Agbl5 | ENSMUSG( | ENSMUSG( | -0.15932 | 0.150945 | NO | NO | NA |
| Piga | ENSMUSG( | ENSMUSG( | 0.276901 | 0.025997 | YES | NO | NA |
| Gm9821 | ENSMUSG( | ENSMUSG( | -0.24939 | 0.030472 | YES | NO | NA |
| Dipk1a | ENSMUSG( | ENSMUSG( | -0.22374 | 0.025966 | YES | NO | NA |
| Aimp2 | ENSMUSG( | ENSMUSG( | 0.329021 | 0.006029 | YES | NO | NA |
| 2310061I0 | ENSMUSG( | ENSMUSG( | 0.250647 | 0.003962 | YES | NO | NA |
| Sh3bp5l | ENSMUSG( | ENSMUSG( | -0.11989 | 0.053541 | NO | NO | NA |
| Cdkn1a | ENSMUSG( | ENSMUSG( | 0.917867 | 1.14E-09 | YES | NO | NA |
| Klhl17 | ENSMUSG( | ENSMUSG( | -0.35341 | 5.87E-05 | YES | NO | NA |
| Usp6nl | ENSMUSG( | ENSMUSG( | -0.05021 | 0.572967 | NO | NO | NA |
| Heatr5a | ENSMUSG( | ENSMUSG( | 0.195015 | 0.002168 | YES | NO | NA |
| Hmbox1 | ENSMUSG( | ENSMUSG( | -0.26199 | 0.003069 | YES | NO | NA |
| Hdac6 | ENSMUSG( | ENSMUSG( | 0.411685 | 4.74E-07 | YES | NO | NA |

| | | | | | | | |
|---|---|---|---|---|---|---|---|
| Creb3 | ENSMUSG( | ENSMUSG( | -0.09001 | 0.365952 | NO | NO | NA |
| Pcyt1a | ENSMUSG( | ENSMUSG( | 0.189098 | 0.193795 | NO | NO | NA |
| Urgcp | ENSMUSG( | ENSMUSG( | -0.08755 | 0.197173 | NO | NO | NA |
| Rnf11 | ENSMUSG( | ENSMUSG( | -0.29547 | 0.012678 | YES | NO | NA |
| Mrpl32 | ENSMUSG( | ENSMUSG( | -0.04343 | 0.585964 | NO | NO | NA |
| Ecsit | ENSMUSG( | ENSMUSG( | 0.265906 | 0.006827 | YES | NO | NA |
| Gnpda2 | ENSMUSG( | ENSMUSG( | 0.101012 | 0.532011 | NO | NO | NA |
| Bloc1s1 | ENSMUSG( | ENSMUSG( | -0.12431 | 0.357003 | NO | NO | NA |
| 2610002M | ENSMUSG( | ENSMUSG( | -0.29514 | 0.001916 | YES | NO | NA |
| Bhlhb9 | ENSMUSG( | ENSMUSG( | 0.277636 | 0.004274 | YES | NO | NA |
| Tspan13 | ENSMUSG( | ENSMUSG( | -0.12034 | 0.256454 | NO | NO | NA |
| Kat6b | ENSMUSG( | ENSMUSG( | -0.11207 | 0.156279 | NO | NO | NA |
| Snhg17 | ENSMUSG( | ENSMUSG( | 0.241369 | 0.005294 | YES | NO | NA |
| Dpy19l4 | ENSMUSG( | ENSMUSG( | 0.015296 | 0.908506 | NO | NO | NA |
| Rcl1 | ENSMUSG( | ENSMUSG( | 0.740965 | 1.35E-07 | YES | NO | NA |
| Trmt10c | ENSMUSG( | ENSMUSG( | 0.06511 | 0.444377 | NO | NO | NA |
| Alkbh1 | ENSMUSG( | ENSMUSG( | 0.040361 | 0.591125 | NO | NO | NA |
| Pcgf3 | ENSMUSG( | ENSMUSG( | -0.26843 | 0.001305 | YES | NO | NA |
| Dxo | ENSMUSG( | ENSMUSG( | 0.028586 | 0.790471 | NO | NO | NA |
| Dpm2 | ENSMUSG( | ENSMUSG( | -0.11835 | 0.255342 | NO | NO | NA |
| Acat2 | ENSMUSG( | ENSMUSG( | 0.089903 | 0.418988 | NO | NO | NA |
| Kremen1 | ENSMUSG( | ENSMUSG( | 0.121027 | 0.360474 | NO | NO | NA |
| Spg20 | ENSMUSG( | ENSMUSG( | -0.04834 | 0.584439 | NO | NO | NA |
| Fig4 | ENSMUSG( | ENSMUSG( | 0.020985 | 0.879395 | NO | NO | NA |
| Cnpy2 | ENSMUSG( | ENSMUSG( | 0.227214 | 0.025862 | YES | NO | NA |
| Ap4m1 | ENSMUSG( | ENSMUSG( | -0.07276 | 0.394648 | NO | NO | NA |
| Arl4a | ENSMUSG( | ENSMUSG( | -0.07801 | 0.414083 | NO | NO | NA |
| Rsad2 | ENSMUSG( | ENSMUSG( | -1.3136 | 6.19E-11 | YES | YES | Downregulated |
| Usp22 | ENSMUSG( | ENSMUSG( | -0.04579 | 0.62909 | NO | NO | NA |
| E130307A | ENSMUSG( | ENSMUSG( | 0.03867 | 0.711964 | NO | NO | NA |
| Tbc1d5 | ENSMUSG( | ENSMUSG( | -0.26884 | 1.42E-05 | YES | NO | NA |
| Dnaaf9 | ENSMUSG( | ENSMUSG( | -0.45786 | 7.82E-06 | YES | NO | NA |
| Cenpx | ENSMUSG( | ENSMUSG( | 0.149681 | 0.081399 | NO | NO | NA |
| Zfp346 | ENSMUSG( | ENSMUSG( | -0.06694 | 0.43991 | NO | NO | NA |
| Wdr55 | ENSMUSG( | ENSMUSG( | -0.02612 | 0.862649 | NO | NO | NA |
| Arrb1 | ENSMUSG( | ENSMUSG( | -0.22543 | 0.16583 | NO | NO | NA |
| Mrpl36 | ENSMUSG( | ENSMUSG( | 0.233925 | 0.018388 | YES | NO | NA |
| Zfp146 | ENSMUSG( | ENSMUSG( | 0.045778 | 0.573204 | NO | NO | NA |
| Mapk7 | ENSMUSG( | ENSMUSG( | -0.12925 | 0.082121 | NO | NO | NA |
| Churc1 | ENSMUSG( | ENSMUSG( | 0.073296 | 0.479388 | NO | NO | NA |
| Selenoo | ENSMUSG( | ENSMUSG( | -0.12592 | 0.274524 | NO | NO | NA |
| Gdap2 | ENSMUSG( | ENSMUSG( | -0.03246 | 0.689625 | NO | NO | NA |
| Fancg | ENSMUSG( | ENSMUSG( | -0.06417 | 0.320156 | NO | NO | NA |
| Apobec2 | ENSMUSG( | ENSMUSG( | -0.7649 | 0.000339 | YES | NO | NA |
| Pced1a | ENSMUSG( | ENSMUSG( | -0.10701 | 0.202772 | NO | NO | NA |
| Wdr12 | ENSMUSG( | ENSMUSG( | 0.589343 | 1.92E-08 | YES | NO | NA |
| Slc25a32 | ENSMUSG( | ENSMUSG( | 0.15853 | 0.042448 | YES | NO | NA |

| | | | | | | | |
|---|---|---|---|---|---|---|---|
| Vasn | ENSMUSG( | ENSMUSG( | 0.076581 | 0.639073 | NO | NO | NA |
| Prmt9 | ENSMUSG( | ENSMUSG( | -0.12559 | 0.100739 | NO | NO | NA |
| Med11 | ENSMUSG( | ENSMUSG( | 0.24105 | 0.001231 | YES | NO | NA |
| Tigd2 | ENSMUSG( | ENSMUSG( | -0.0923 | 0.412712 | NO | NO | NA |
| Rbms2 | ENSMUSG( | ENSMUSG( | -0.54123 | 1.83E-08 | YES | NO | NA |
| Dhrs4 | ENSMUSG( | ENSMUSG( | 0.380739 | 0.000791 | YES | NO | NA |
| Slc12a2 | ENSMUSG( | ENSMUSG( | 0.672758 | 3.57E-08 | YES | NO | NA |
| Gm12250 | ENSMUSG( | ENSMUSG( | -0.55073 | 5.11E-09 | YES | NO | NA |
| Tmbim4 | ENSMUSG( | ENSMUSG( | 0.322121 | 5.34E-05 | YES | NO | NA |
| Zbtb24 | ENSMUSG( | ENSMUSG( | -0.37246 | 7.88E-06 | YES | NO | NA |
| Cmc2 | ENSMUSG( | ENSMUSG( | 0.027254 | 0.732447 | NO | NO | NA |
| Kif18a | ENSMUSG( | ENSMUSG( | 0.10612 | 0.312553 | NO | NO | NA |
| Hikeshi | ENSMUSG( | ENSMUSG( | -0.13084 | 0.142439 | NO | NO | NA |
| Wdr24 | ENSMUSG( | ENSMUSG( | -0.11523 | 0.103235 | NO | NO | NA |
| Zfp715 | ENSMUSG( | ENSMUSG( | -0.38152 | 5.56E-05 | YES | NO | NA |
| Polk | ENSMUSG( | ENSMUSG( | -0.38205 | 1.71E-06 | YES | NO | NA |
| Ifi30 | ENSMUSG( | ENSMUSG( | 0.388626 | 4.43E-06 | YES | NO | NA |
| Sphk2 | ENSMUSG( | ENSMUSG( | -0.14661 | 0.104008 | NO | NO | NA |
| Gabarapl1 | ENSMUSG( | ENSMUSG( | 0.180635 | 0.12565 | NO | NO | NA |
| Tada2b | ENSMUSG( | ENSMUSG( | -0.06226 | 0.455954 | NO | NO | NA |
| Srd5a3 | ENSMUSG( | ENSMUSG( | -0.19734 | 0.031439 | YES | NO | NA |
| Haao | ENSMUSG( | ENSMUSG( | -0.1949 | 0.05601 | NO | NO | NA |
| Nfkbie | ENSMUSG( | ENSMUSG( | -0.53951 | 3.12E-10 | YES | NO | NA |
| Ddx51 | ENSMUSG( | ENSMUSG( | 0.386416 | 4.07E-06 | YES | NO | NA |
| Pip4p1 | ENSMUSG( | ENSMUSG( | 0.015041 | 0.901555 | NO | NO | NA |
| Zfp809 | ENSMUSG( | ENSMUSG( | -0.46192 | 1.74E-11 | YES | NO | NA |
| Zbtb40 | ENSMUSG( | ENSMUSG( | -0.31819 | 2.06E-08 | YES | NO | NA |
| Pigu | ENSMUSG( | ENSMUSG( | 0.082175 | 0.284292 | NO | NO | NA |
| Zhx3 | ENSMUSG( | ENSMUSG( | -0.19726 | 0.013072 | YES | NO | NA |
| Unc50 | ENSMUSG( | ENSMUSG( | 0.062157 | 0.505687 | NO | NO | NA |
| Cln5 | ENSMUSG( | ENSMUSG( | -0.208 | 0.000618 | YES | NO | NA |
| Nudt16 | ENSMUSG( | ENSMUSG( | 0.214805 | 0.162484 | NO | NO | NA |
| Atg101 | ENSMUSG( | ENSMUSG( | -0.07738 | 0.324777 | NO | NO | NA |
| Manba | ENSMUSG( | ENSMUSG( | -0.43014 | 6.32E-05 | YES | NO | NA |
| Miga1 | ENSMUSG( | ENSMUSG( | 0.253692 | 0.007053 | YES | NO | NA |
| Mrpl41 | ENSMUSG( | ENSMUSG( | 0.1172 | 0.1374 | NO | NO | NA |
| Bcl7a | ENSMUSG( | ENSMUSG( | -0.24762 | 0.011252 | YES | NO | NA |
| Cript | ENSMUSG( | ENSMUSG( | -0.33607 | 0.000183 | YES | NO | NA |
| Tafazzin | ENSMUSG( | ENSMUSG( | -0.23336 | 0.014358 | YES | NO | NA |
| Coa8 | ENSMUSG( | ENSMUSG( | 0.360618 | 0.000862 | YES | NO | NA |
| Vps8 | ENSMUSG( | ENSMUSG( | -0.19505 | 0.001282 | YES | NO | NA |
| Rad17 | ENSMUSG( | ENSMUSG( | -0.08314 | 0.127357 | NO | NO | NA |
| Dhx35 | ENSMUSG( | ENSMUSG( | 0.018237 | 0.855008 | NO | NO | NA |
| Ccne2 | ENSMUSG( | ENSMUSG( | 0.327406 | 0.001493 | YES | NO | NA |
| Zfand2a | ENSMUSG( | ENSMUSG( | -0.3332 | 0.002567 | YES | NO | NA |
| Tex9 | ENSMUSG( | ENSMUSG( | 0.301709 | 0.004529 | YES | NO | NA |
| Wrap73 | ENSMUSG( | ENSMUSG( | 0.008806 | 0.914861 | NO | NO | NA |

| | | | | | | | |
|---|---|---|---|---|---|---|---|
| Rbm4b | ENSMUSG( | ENSMUSG( | -0.36807 | 2.17E-07 | YES | NO | NA |
| Hcfc2 | ENSMUSG( | ENSMUSG( | -0.22643 | 0.001902 | YES | NO | NA |
| Asah1 | ENSMUSG( | ENSMUSG( | 0.142453 | 0.231343 | NO | NO | NA |
| Slc38a9 | ENSMUSG( | ENSMUSG( | -0.05081 | 0.624845 | NO | NO | NA |
| Lpcat3 | ENSMUSG( | ENSMUSG( | 0.036107 | 0.771003 | NO | NO | NA |
| Tex30 | ENSMUSG( | ENSMUSG( | 0.301002 | 0.000472 | YES | NO | NA |
| Tmco6 | ENSMUSG( | ENSMUSG( | -0.1686 | 0.143603 | NO | NO | NA |
| Ptcd1 | ENSMUSG( | ENSMUSG( | -0.15262 | 0.082352 | NO | NO | NA |
| Gpr18 | ENSMUSG( | ENSMUSG( | 0.013935 | 0.919297 | NO | NO | NA |
| Commd1 | ENSMUSG( | ENSMUSG( | 0.17346 | 0.043014 | YES | NO | NA |
| F2rl1 | ENSMUSG( | ENSMUSG( | -1.04499 | 1.14E-09 | YES | YES | Downregulated |
| Gtf2e2 | ENSMUSG( | ENSMUSG( | 0.186352 | 0.034857 | YES | NO | NA |
| Nfkbid | ENSMUSG( | ENSMUSG( | -0.22655 | 0.061462 | NO | NO | NA |
| Slc37a4 | ENSMUSG( | ENSMUSG( | 0.250283 | 0.002793 | YES | NO | NA |
| Zfand4 | ENSMUSG( | ENSMUSG( | 0.202606 | 0.072213 | NO | NO | NA |
| Gtpbp3 | ENSMUSG( | ENSMUSG( | 0.016998 | 0.812414 | NO | NO | NA |
| E2f6 | ENSMUSG( | ENSMUSG( | -0.08328 | 0.197008 | NO | NO | NA |
| Tstd3 | ENSMUSG( | ENSMUSG( | 0.027079 | 0.793871 | NO | NO | NA |
| Nsmce3 | ENSMUSG( | ENSMUSG( | 0.030034 | 0.781396 | NO | NO | NA |
| Bag5 | ENSMUSG( | ENSMUSG( | 0.066946 | 0.476775 | NO | NO | NA |
| Gpr146 | ENSMUSG( | ENSMUSG( | 0.127867 | 0.437342 | NO | NO | NA |
| Zfp120 | ENSMUSG( | ENSMUSG( | -0.45035 | 0.000222 | YES | NO | NA |
| Yipf6 | ENSMUSG( | ENSMUSG( | -0.21251 | 0.006807 | YES | NO | NA |
| Gtf3c4 | ENSMUSG( | ENSMUSG( | -0.11997 | 0.154904 | NO | NO | NA |
| Irgq | ENSMUSG( | ENSMUSG( | -0.28828 | 0.005141 | YES | NO | NA |
| Nudt19 | ENSMUSG( | ENSMUSG( | 0.241319 | 0.034186 | YES | NO | NA |
| Thap11 | ENSMUSG( | ENSMUSG( | -0.07449 | 0.229509 | NO | NO | NA |
| Lockd | ENSMUSG( | ENSMUSG( | 0.020385 | 0.852826 | NO | NO | NA |
| Gpn3 | ENSMUSG( | ENSMUSG( | 0.190468 | 0.024163 | YES | NO | NA |
| Gadd45b | ENSMUSG( | ENSMUSG( | 0.561266 | 0.000305 | YES | NO | NA |
| Pigv | ENSMUSG( | ENSMUSG( | -0.30518 | 0.006617 | YES | NO | NA |
| Smim10l1 | ENSMUSG( | ENSMUSG( | 0.196706 | 0.086036 | NO | NO | NA |
| Plxdc1 | ENSMUSG( | ENSMUSG( | -0.177 | 0.073894 | NO | NO | NA |
| Pef1 | ENSMUSG( | ENSMUSG( | -0.06594 | 0.414728 | NO | NO | NA |
| Dstn | ENSMUSG( | ENSMUSG( | 0.118797 | 0.19959 | NO | NO | NA |
| Casd1 | ENSMUSG( | ENSMUSG( | -0.34774 | 4.35E-05 | YES | NO | NA |
| Timm8a1 | ENSMUSG( | ENSMUSG( | 0.697791 | 1.05E-06 | YES | NO | NA |
| Ints12 | ENSMUSG( | ENSMUSG( | 0.064036 | 0.503631 | NO | NO | NA |
| Fam98c | ENSMUSG( | ENSMUSG( | 0.055164 | 0.617881 | NO | NO | NA |
| Asb13 | ENSMUSG( | ENSMUSG( | -0.65768 | 3.38E-13 | YES | NO | NA |
| Fam214a | ENSMUSG( | ENSMUSG( | -0.57791 | 1.90E-05 | YES | NO | NA |
| Commd2 | ENSMUSG( | ENSMUSG( | 0.102242 | 0.157181 | NO | NO | NA |
| Mapkbp1 | ENSMUSG( | ENSMUSG( | -0.22183 | 0.063195 | NO | NO | NA |
| Eps8l1 | ENSMUSG( | ENSMUSG( | -0.56928 | 0.323894 | NO | NO | NA |
| Dhx33 | ENSMUSG( | ENSMUSG( | 0.182688 | 0.02382 | YES | NO | NA |
| ENSMUSG | ENSMUSG( | ENSMUSG( | -0.41668 | 4.34E-06 | YES | NO | NA |
| 2210016L2 | ENSMUSG( | ENSMUSG( | -0.09039 | 0.170241 | NO | NO | NA |

| | | | | | | | |
|---|---|---|---|---|---|---|---|
| Kcnq1ot1 | ENSMUSG( | ENSMUSG( | 0.13192 | 0.584088 | NO | NO | NA |
| Rubcn | ENSMUSG( | ENSMUSG( | -0.19463 | 0.002996 | YES | NO | NA |
| Zfp64 | ENSMUSG( | ENSMUSG( | 0.190652 | 0.079299 | NO | NO | NA |
| Abi2 | ENSMUSG( | ENSMUSG( | 0.444926 | 8.34E-06 | YES | NO | NA |
| Gm33858 | ENSMUSG( | ENSMUSG( | 1.066134 | 0.012004 | YES | YES | Upregulated |
| Pin4 | ENSMUSG( | ENSMUSG( | 0.058844 | 0.801026 | NO | NO | NA |
| Atpaf2 | ENSMUSG( | ENSMUSG( | 0.034726 | 0.746794 | NO | NO | NA |
| Rdh11 | ENSMUSG( | ENSMUSG( | -0.08021 | 0.502435 | NO | NO | NA |
| Gmpr2 | ENSMUSG( | ENSMUSG( | 0.20181 | 0.01128 | YES | NO | NA |
| Cercam | ENSMUSG( | ENSMUSG( | -0.27899 | 0.000241 | YES | NO | NA |
| Rabepk | ENSMUSG( | ENSMUSG( | 0.21768 | 0.000508 | YES | NO | NA |
| Ammecr1l | ENSMUSG( | ENSMUSG( | -0.35876 | 2.00E-07 | YES | NO | NA |
| Slc45a4 | ENSMUSG( | ENSMUSG( | -0.44321 | 0.001332 | YES | NO | NA |
| Stambpl1 | ENSMUSG( | ENSMUSG( | -0.03911 | 0.679852 | NO | NO | NA |
| Saal1 | ENSMUSG( | ENSMUSG( | 0.167748 | 0.034658 | YES | NO | NA |
| Cbfa2t2 | ENSMUSG( | ENSMUSG( | -0.0514 | 0.580155 | NO | NO | NA |
| Klf11 | ENSMUSG( | ENSMUSG( | 0.044684 | 0.843184 | NO | NO | NA |
| Zfp422 | ENSMUSG( | ENSMUSG( | 0.396557 | 0.000473 | YES | NO | NA |
| Slc29a3 | ENSMUSG( | ENSMUSG( | -0.15669 | 0.194787 | NO | NO | NA |
| Pgap2 | ENSMUSG( | ENSMUSG( | -0.02682 | 0.772331 | NO | NO | NA |
| Ndufb4 | ENSMUSG( | ENSMUSG( | 0.258756 | 0.009129 | YES | NO | NA |
| Zfp597 | ENSMUSG( | ENSMUSG( | -0.45839 | 3.79E-05 | YES | NO | NA |
| Cd200r1 | ENSMUSG( | ENSMUSG( | -0.75505 | 0.005241 | YES | NO | NA |
| Spata5l1 | ENSMUSG( | ENSMUSG( | -0.13604 | 0.049895 | YES | NO | NA |
| Vhl | ENSMUSG( | ENSMUSG( | 0.024464 | 0.840265 | NO | NO | NA |
| Gbp10 | ENSMUSG( | ENSMUSG( | -1.34913 | 4.67E-07 | YES | YES | Downregulated |
| Fgf13 | ENSMUSG( | ENSMUSG( | -0.67088 | 8.80E-07 | YES | NO | NA |
| Chmp2b | ENSMUSG( | ENSMUSG( | 0.01665 | 0.900004 | NO | NO | NA |
| Sec23a | ENSMUSG( | ENSMUSG( | -0.4452 | 2.12E-05 | YES | NO | NA |
| Cklf | ENSMUSG( | ENSMUSG( | -0.19895 | 0.005091 | YES | NO | NA |
| Mettl17 | ENSMUSG( | ENSMUSG( | -0.08902 | 0.235458 | NO | NO | NA |
| Gsn | ENSMUSG( | ENSMUSG( | -1.13497 | 8.72E-10 | YES | YES | Downregulated |
| Mastl | ENSMUSG( | ENSMUSG( | 0.392006 | 2.90E-05 | YES | NO | NA |
| Cibar1 | ENSMUSG( | ENSMUSG( | 0.257251 | 0.024965 | YES | NO | NA |
| Miat | ENSMUSG( | ENSMUSG( | -0.98072 | 9.40E-07 | YES | NO | NA |
| Tbkbp1 | ENSMUSG( | ENSMUSG( | -0.62396 | 9.21E-06 | YES | NO | NA |
| Ifrd2 | ENSMUSG( | ENSMUSG( | 0.793658 | 1.19E-08 | YES | NO | NA |
| Zc3h6 | ENSMUSG( | ENSMUSG( | 0.643805 | 0.001431 | YES | NO | NA |
| Cnpy3 | ENSMUSG( | ENSMUSG( | 0.107311 | 0.167253 | NO | NO | NA |
| Zc3h12a | ENSMUSG( | ENSMUSG( | 0.023735 | 0.894092 | NO | NO | NA |
| 9430015G | ENSMUSG( | ENSMUSG( | -0.01916 | 0.857648 | NO | NO | NA |
| Evi2a | ENSMUSG( | ENSMUSG( | 0.017898 | 0.885851 | NO | NO | NA |
| Cenpq | ENSMUSG( | ENSMUSG( | 0.09152 | 0.41526 | NO | NO | NA |
| Grwd1 | ENSMUSG( | ENSMUSG( | 0.6026 | 3.79E-05 | YES | NO | NA |
| Mettl7a1 | ENSMUSG( | ENSMUSG( | -0.00368 | 0.980563 | NO | NO | NA |
| Ppfibp1 | ENSMUSG( | ENSMUSG( | 0.314077 | 0.006414 | YES | NO | NA |
| Scfd2 | ENSMUSG( | ENSMUSG( | -0.29941 | 8.21E-06 | YES | NO | NA |

| | | | | | | | |
|---|---|---|---|---|---|---|---|
| Cdk7 | ENSMUSG( | ENSMUSG( | 0.185433 | 0.004631 | YES | NO | NA |
| Plxnd1 | ENSMUSG( | ENSMUSG( | 0.739965 | 1.68E-09 | YES | NO | NA |
| Mpv17l2 | ENSMUSG( | ENSMUSG( | 0.264829 | 0.011793 | YES | NO | NA |
| Alkbh8 | ENSMUSG( | ENSMUSG( | -0.09806 | 0.296812 | NO | NO | NA |
| Rbm12b2 | ENSMUSG( | ENSMUSG( | -0.66736 | 2.48E-07 | YES | NO | NA |
| Pex2 | ENSMUSG( | ENSMUSG( | 0.214341 | 0.152029 | NO | NO | NA |
| Slc25a16 | ENSMUSG( | ENSMUSG( | 0.309758 | 0.034341 | YES | NO | NA |
| Phkb | ENSMUSG( | ENSMUSG( | -0.15364 | 0.192584 | NO | NO | NA |
| Slc19a1 | ENSMUSG( | ENSMUSG( | 0.637964 | 0.000139 | YES | NO | NA |
| Ggh | ENSMUSG( | ENSMUSG( | 0.29675 | 0.033623 | YES | NO | NA |
| 4933427D | ENSMUSG( | ENSMUSG( | -0.04263 | 0.734883 | NO | NO | NA |
| Pot1a | ENSMUSG( | ENSMUSG( | -0.27719 | 0.000523 | YES | NO | NA |
| Tgtp1 | ENSMUSG( | ENSMUSG( | 0.008446 | 0.975934 | NO | NO | NA |
| Ndufb1 | ENSMUSG( | ENSMUSG( | -0.03044 | 0.781352 | NO | NO | NA |
| St7l | ENSMUSG( | ENSMUSG( | -0.07477 | 0.48992 | NO | NO | NA |
| Tob1 | ENSMUSG( | ENSMUSG( | -0.30528 | 0.029547 | YES | NO | NA |
| Pstk | ENSMUSG( | ENSMUSG( | -0.11231 | 0.234366 | NO | NO | NA |
| Mrps34 | ENSMUSG( | ENSMUSG( | 0.384599 | 0.001909 | YES | NO | NA |
| Wwp1 | ENSMUSG( | ENSMUSG( | 0.049314 | 0.763649 | NO | NO | NA |
| Mrpl23 | ENSMUSG( | ENSMUSG( | 0.565475 | 1.55E-09 | YES | NO | NA |
| Pdcd2 | ENSMUSG( | ENSMUSG( | 0.011806 | 0.901146 | NO | NO | NA |
| Slc14a1 | ENSMUSG( | ENSMUSG( | -0.27847 | 0.139978 | NO | NO | NA |
| Pbk | ENSMUSG( | ENSMUSG( | -0.16462 | 0.034807 | YES | NO | NA |
| Asl | ENSMUSG( | ENSMUSG( | 0.149596 | 0.021293 | YES | NO | NA |
| Zmat3 | ENSMUSG( | ENSMUSG( | 0.128726 | 0.351132 | NO | NO | NA |
| Zfp326 | ENSMUSG( | ENSMUSG( | 0.379365 | 0.001223 | YES | NO | NA |
| Thap4 | ENSMUSG( | ENSMUSG( | -0.01938 | 0.828296 | NO | NO | NA |
| Cd86 | ENSMUSG( | ENSMUSG( | -0.72632 | 1.47E-10 | YES | NO | NA |
| Spr | ENSMUSG( | ENSMUSG( | 0.267976 | 0.001029 | YES | NO | NA |
| Sema4f | ENSMUSG( | ENSMUSG( | -0.22785 | 0.070538 | NO | NO | NA |
| Zfp511 | ENSMUSG( | ENSMUSG( | -0.02772 | 0.761001 | NO | NO | NA |
| Zc3hc1 | ENSMUSG( | ENSMUSG( | 0.463741 | 1.33E-09 | YES | NO | NA |
| Wdr95 | ENSMUSG( | ENSMUSG( | -0.24998 | 0.320477 | NO | NO | NA |
| Fam122b | ENSMUSG( | ENSMUSG( | -0.04622 | 0.679095 | NO | NO | NA |
| Dmac2 | ENSMUSG( | ENSMUSG( | -0.14199 | 0.210086 | NO | NO | NA |
| Mrps35 | ENSMUSG( | ENSMUSG( | 0.127602 | 0.129614 | NO | NO | NA |
| Nfkbil1 | ENSMUSG( | ENSMUSG( | 0.492745 | 0.005108 | YES | NO | NA |
| Usp31 | ENSMUSG( | ENSMUSG( | -0.15827 | 0.045917 | YES | NO | NA |
| Rrnad1 | ENSMUSG( | ENSMUSG( | -0.09435 | 0.323577 | NO | NO | NA |
| Entpd6 | ENSMUSG( | ENSMUSG( | -0.02822 | 0.722243 | NO | NO | NA |
| Car1 | ENSMUSG( | ENSMUSG( | 1.082506 | 0.160171 | NO | NO | NA |
| Smad5 | ENSMUSG( | ENSMUSG( | -0.3203 | 0.002638 | YES | NO | NA |
| Gm45902 | ENSMUSG( | ENSMUSG( | 0.130928 | 0.471129 | NO | NO | NA |
| Nqo2 | ENSMUSG( | ENSMUSG( | -0.33985 | 1.17E-05 | YES | NO | NA |
| Fchsd1 | ENSMUSG( | ENSMUSG( | -0.49533 | 3.05E-11 | YES | NO | NA |
| Ing2 | ENSMUSG( | ENSMUSG( | 0.065989 | 0.717706 | NO | NO | NA |
| Heatr9 | ENSMUSG( | ENSMUSG( | -0.74682 | 1.51E-10 | YES | NO | NA |

| | | | | | | | |
|---|---|---|---|---|---|---|---|
| Zfp426 | ENSMUSG( | ENSMUSG( | -0.33323 | 0.000806 | YES | NO | NA |
| Mmgt1 | ENSMUSG( | ENSMUSG( | 0.039196 | 0.748181 | NO | NO | NA |
| Uqcc3 | ENSMUSG( | ENSMUSG( | 0.196674 | 0.045837 | YES | NO | NA |
| Gpatch11 | ENSMUSG( | ENSMUSG( | -0.22623 | 0.026503 | YES | NO | NA |
| Ccsap | ENSMUSG( | ENSMUSG( | -0.04806 | 0.696194 | NO | NO | NA |
| ENSMUSG( | ENSMUSG( | ENSMUSG( | -0.153 | 0.096178 | NO | NO | NA |
| Mrpl38 | ENSMUSG( | ENSMUSG( | 0.294697 | 0.000455 | YES | NO | NA |
| Uap1l1 | ENSMUSG( | ENSMUSG( | -0.07556 | 0.483813 | NO | NO | NA |
| Rad51d | ENSMUSG( | ENSMUSG( | -0.12757 | 0.13709 | NO | NO | NA |
| Xlr4c | ENSMUSG( | ENSMUSG( | -0.24832 | 0.337098 | NO | NO | NA |
| Smtn | ENSMUSG( | ENSMUSG( | -0.20632 | 0.03918 | YES | NO | NA |
| Ttll5 | ENSMUSG( | ENSMUSG( | -0.02312 | 0.873694 | NO | NO | NA |
| Tnfaip8l2 | ENSMUSG( | ENSMUSG( | -0.35493 | 3.17E-05 | YES | NO | NA |
| Mrps2 | ENSMUSG( | ENSMUSG( | 0.282078 | 0.000837 | YES | NO | NA |
| Mad2l1bp | ENSMUSG( | ENSMUSG( | 0.00452 | 0.949367 | NO | NO | NA |
| Smim30 | ENSMUSG( | ENSMUSG( | 0.053978 | 0.682083 | NO | NO | NA |
| Mical2 | ENSMUSG( | ENSMUSG( | -0.84535 | 7.61E-10 | YES | NO | NA |
| Ppip5k1 | ENSMUSG( | ENSMUSG( | -0.07023 | 0.456421 | NO | NO | NA |
| Rell1 | ENSMUSG( | ENSMUSG( | -0.54436 | 4.93E-12 | YES | NO | NA |
| Nop16 | ENSMUSG( | ENSMUSG( | 0.727882 | 5.57E-07 | YES | NO | NA |
| Zbtb9 | ENSMUSG( | ENSMUSG( | -0.42807 | 8.58E-07 | YES | NO | NA |
| Zfp942 | ENSMUSG( | ENSMUSG( | -0.33942 | 0.00795 | YES | NO | NA |
| Alkbh4 | ENSMUSG( | ENSMUSG( | -0.24397 | 0.005137 | YES | NO | NA |
| Anks3 | ENSMUSG( | ENSMUSG( | -0.08907 | 0.333591 | NO | NO | NA |
| Alas1 | ENSMUSG( | ENSMUSG( | 0.364747 | 1.26E-05 | YES | NO | NA |
| Cops7b | ENSMUSG( | ENSMUSG( | 0.170781 | 0.03424 | YES | NO | NA |
| Kctd3 | ENSMUSG( | ENSMUSG( | -0.04045 | 0.714272 | NO | NO | NA |
| Tigar | ENSMUSG( | ENSMUSG( | -0.02238 | 0.808208 | NO | NO | NA |
| Fhl2 | ENSMUSG( | ENSMUSG( | -0.28425 | 0.000106 | YES | NO | NA |
| 4921524J1 | ENSMUSG( | ENSMUSG( | -0.11445 | 0.308111 | NO | NO | NA |
| Gtf2h2 | ENSMUSG( | ENSMUSG( | 0.061852 | 0.552734 | NO | NO | NA |
| Sumf1 | ENSMUSG( | ENSMUSG( | 0.049984 | 0.73361 | NO | NO | NA |
| 9130401M | ENSMUSG( | ENSMUSG( | 0.352585 | 0.002213 | YES | NO | NA |
| Rabgef1 | ENSMUSG( | ENSMUSG( | -0.39519 | 2.45E-11 | YES | NO | NA |
| Nsmce1 | ENSMUSG( | ENSMUSG( | 0.235158 | 0.021773 | YES | NO | NA |
| Prdm15 | ENSMUSG( | ENSMUSG( | -0.25522 | 0.000383 | YES | NO | NA |
| Ttc21b | ENSMUSG( | ENSMUSG( | -0.11135 | 0.29696 | NO | NO | NA |
| Haus2 | ENSMUSG( | ENSMUSG( | 0.25141 | 0.018616 | YES | NO | NA |
| Numb | ENSMUSG( | ENSMUSG( | -0.11563 | 0.476797 | NO | NO | NA |
| Usp30 | ENSMUSG( | ENSMUSG( | 0.04791 | 0.690509 | NO | NO | NA |
| Rrp36 | ENSMUSG( | ENSMUSG( | 0.050366 | 0.5314 | NO | NO | NA |
| Cdc25c | ENSMUSG( | ENSMUSG( | 0.418978 | 7.96E-07 | YES | NO | NA |
| Antxr2 | ENSMUSG( | ENSMUSG( | -0.75369 | 4.39E-10 | YES | NO | NA |
| Map2k5 | ENSMUSG( | ENSMUSG( | 0.037232 | 0.728736 | NO | NO | NA |
| Dnajc19 | ENSMUSG( | ENSMUSG( | 0.182535 | 0.180747 | NO | NO | NA |
| Orai1 | ENSMUSG( | ENSMUSG( | -0.13537 | 0.385531 | NO | NO | NA |
| Cinp | ENSMUSG( | ENSMUSG( | 0.388221 | 0.000276 | YES | NO | NA |

| | | | | | | | |
|---|---|---|---|---|---|---|---|
| Zc3h10 | ENSMUSG( | ENSMUSG( | -0.20934 | 5.46E-05 | YES | NO | NA |
| Traf4 | ENSMUSG( | ENSMUSG( | -0.15105 | 0.043586 | YES | NO | NA |
| Ndufa5 | ENSMUSG( | ENSMUSG( | 0.235236 | 0.052939 | NO | NO | NA |
| Gm37194 | ENSMUSG( | ENSMUSG( | 0.577864 | 0.48389 | NO | NO | NA |
| Plpp6 | ENSMUSG( | ENSMUSG( | 0.001555 | 0.993521 | NO | NO | NA |
| Bclaf3 | ENSMUSG( | ENSMUSG( | -0.09406 | 0.424383 | NO | NO | NA |
| Acot8 | ENSMUSG( | ENSMUSG( | -0.18683 | 0.002291 | YES | NO | NA |
| Zfp668 | ENSMUSG( | ENSMUSG( | -0.01592 | 0.838522 | NO | NO | NA |
| Tarbp1 | ENSMUSG( | ENSMUSG( | 0.319058 | 5.83E-06 | YES | NO | NA |
| Dguok | ENSMUSG( | ENSMUSG( | -0.03836 | 0.763512 | NO | NO | NA |
| 2310033P( | ENSMUSG( | ENSMUSG( | 0.052424 | 0.626178 | NO | NO | NA |
| BC029722 | ENSMUSG( | ENSMUSG( | 0.274763 | 0.001852 | YES | NO | NA |
| Rhobtb2 | ENSMUSG( | ENSMUSG( | -0.12951 | 0.065315 | NO | NO | NA |
| Zfp984 | ENSMUSG( | ENSMUSG( | -1.23857 | 7.50E-34 | YES | YES | Downregulated |
| Sigmar1 | ENSMUSG( | ENSMUSG( | 0.514343 | 0.000273 | YES | NO | NA |
| Pus7 | ENSMUSG( | ENSMUSG( | 0.892696 | 2.84E-06 | YES | NO | NA |
| Kbtbd4 | ENSMUSG( | ENSMUSG( | -0.24535 | 0.002257 | YES | NO | NA |
| Ginm1 | ENSMUSG( | ENSMUSG( | 0.270126 | 0.067101 | NO | NO | NA |
| Plekhf1 | ENSMUSG( | ENSMUSG( | -0.78248 | 2.10E-09 | YES | NO | NA |
| 4930579G | ENSMUSG( | ENSMUSG( | 0.159689 | 0.010027 | YES | NO | NA |
| Crtc1 | ENSMUSG( | ENSMUSG( | 0.198529 | 0.073 | NO | NO | NA |
| Hus1 | ENSMUSG( | ENSMUSG( | 0.063575 | 0.410461 | NO | NO | NA |
| Prpsap2 | ENSMUSG( | ENSMUSG( | -0.21716 | 0.015662 | YES | NO | NA |
| Tmem50b | ENSMUSG( | ENSMUSG( | 0.216257 | 0.269371 | NO | NO | NA |
| ENSMUSG( | ENSMUSG( | ENSMUSG( | 0.154717 | 0.405313 | NO | NO | NA |
| Mpst | ENSMUSG( | ENSMUSG( | 0.069991 | 0.552744 | NO | NO | NA |
| Usp21 | ENSMUSG( | ENSMUSG( | -0.25477 | 3.37E-05 | YES | NO | NA |
| Sgsh | ENSMUSG( | ENSMUSG( | -0.12723 | 0.388544 | NO | NO | NA |
| Mrps18a | ENSMUSG( | ENSMUSG( | 0.216479 | 0.014188 | YES | NO | NA |
| Wdr20 | ENSMUSG( | ENSMUSG( | -0.35604 | 0.000375 | YES | NO | NA |
| Mrpl44 | ENSMUSG( | ENSMUSG( | 0.220582 | 0.012239 | YES | NO | NA |
| Mbd4 | ENSMUSG( | ENSMUSG( | -0.0768 | 0.406864 | NO | NO | NA |
| Vps50 | ENSMUSG( | ENSMUSG( | 0.217417 | 0.040091 | YES | NO | NA |
| Pms1 | ENSMUSG( | ENSMUSG( | 0.1384 | 0.160836 | NO | NO | NA |
| Sanbr | ENSMUSG( | ENSMUSG( | 0.512945 | 0.001501 | YES | NO | NA |
| Ero1b | ENSMUSG( | ENSMUSG( | -0.09188 | 0.306823 | NO | NO | NA |
| Rpap2 | ENSMUSG( | ENSMUSG( | -0.15399 | 0.149026 | NO | NO | NA |
| Adamts6 | ENSMUSG( | ENSMUSG( | -0.20424 | 0.152586 | NO | NO | NA |
| Csnk1e | ENSMUSG( | ENSMUSG( | -0.38389 | 0.000913 | YES | NO | NA |
| Wdr91 | ENSMUSG( | ENSMUSG( | -0.06368 | 0.562509 | NO | NO | NA |
| Ring1 | ENSMUSG( | ENSMUSG( | 0.016473 | 0.874667 | NO | NO | NA |
| D2hgdh | ENSMUSG( | ENSMUSG( | -0.46652 | 0.000785 | YES | NO | NA |
| Cdip1 | ENSMUSG( | ENSMUSG( | 0.244568 | 0.002534 | YES | NO | NA |
| Siva1 | ENSMUSG( | ENSMUSG( | 0.494731 | 4.66E-05 | YES | NO | NA |
| Rusf1 | ENSMUSG( | ENSMUSG( | 0.110898 | 0.261706 | NO | NO | NA |
| Asb8 | ENSMUSG( | ENSMUSG( | 0.060975 | 0.432246 | NO | NO | NA |
| Gtf2h3 | ENSMUSG( | ENSMUSG( | 0.092849 | 0.42412 | NO | NO | NA |

| | | | | | | | |
|---|---|---|---|---|---|---|---|
| Ric8b | ENSMUSG( | ENSMUSG( | -0.24005 | 0.013585 | YES | NO | NA |
| Gm7336 | ENSMUSG( | ENSMUSG( | 0.545931 | 0.000903 | YES | NO | NA |
| Serpini1 | ENSMUSG( | ENSMUSG( | -1.31526 | 8.62E-14 | YES | YES | Downregulated |
| Naxd | ENSMUSG( | ENSMUSG( | -0.07 | 0.365272 | NO | NO | NA |
| H2-T-ps | ENSMUSG( | ENSMUSG( | -0.408 | 1.69E-07 | YES | NO | NA |
| Naa38 | ENSMUSG( | ENSMUSG( | 0.132872 | 0.213047 | NO | NO | NA |
| Mesd | ENSMUSG( | ENSMUSG( | 0.019372 | 0.825135 | NO | NO | NA |
| ENSMUSG( | ENSMUSG( | ENSMUSG( | -0.16419 | 0.095435 | NO | NO | NA |
| Pramel12 | ENSMUSG( | ENSMUSG( | -0.43798 | 9.54E-08 | YES | NO | NA |
| Arl6ip6 | ENSMUSG( | ENSMUSG( | 0.006566 | 0.951501 | NO | NO | NA |
| Gcfc2 | ENSMUSG( | ENSMUSG( | 0.332005 | 1.02E-05 | YES | NO | NA |
| Zc3h3 | ENSMUSG( | ENSMUSG( | -0.23151 | 0.085849 | NO | NO | NA |
| Tram2 | ENSMUSG( | ENSMUSG( | -0.38839 | 5.64E-06 | YES | NO | NA |
| Nadk2 | ENSMUSG( | ENSMUSG( | 0.044843 | 0.680543 | NO | NO | NA |
| Vps9d1 | ENSMUSG( | ENSMUSG( | -0.17141 | 0.081671 | NO | NO | NA |
| Manbal | ENSMUSG( | ENSMUSG( | 0.203111 | 0.26162 | NO | NO | NA |
| 1700097N | ENSMUSG( | ENSMUSG( | 0.479503 | 0.002433 | YES | NO | NA |
| Nxn | ENSMUSG( | ENSMUSG( | 0.021776 | 0.824786 | NO | NO | NA |
| Riox1 | ENSMUSG( | ENSMUSG( | 0.099708 | 0.25335 | NO | NO | NA |
| Rarg | ENSMUSG( | ENSMUSG( | -0.57973 | 1.08E-07 | YES | NO | NA |
| Mtrr | ENSMUSG( | ENSMUSG( | 0.083253 | 0.323498 | NO | NO | NA |
| Gmppb | ENSMUSG( | ENSMUSG( | -0.05849 | 0.544325 | NO | NO | NA |
| Borcs8 | ENSMUSG( | ENSMUSG( | -0.2694 | 0.000422 | YES | NO | NA |
| Yif1b | ENSMUSG( | ENSMUSG( | 0.424068 | 0.000184 | YES | NO | NA |
| Gm44226 | ENSMUSG( | ENSMUSG( | -0.27367 | 0.049756 | YES | NO | NA |
| Ppp2r3d | ENSMUSG( | ENSMUSG( | 0.192419 | 0.104897 | NO | NO | NA |
| Nprl2 | ENSMUSG( | ENSMUSG( | -0.33201 | 9.46E-07 | YES | NO | NA |
| Mrps10 | ENSMUSG( | ENSMUSG( | 0.496364 | 1.74E-07 | YES | NO | NA |
| Tmem175 | ENSMUSG( | ENSMUSG( | -0.11497 | 0.41934 | NO | NO | NA |
| Slc26a10 | ENSMUSG( | ENSMUSG( | 0.12112 | 0.60057 | NO | NO | NA |
| Slc25a25 | ENSMUSG( | ENSMUSG( | -0.07719 | 0.402703 | NO | NO | NA |
| Frat2 | ENSMUSG( | ENSMUSG( | -0.54087 | 0.000647 | YES | NO | NA |
| Gins1 | ENSMUSG( | ENSMUSG( | 0.396306 | 8.95E-07 | YES | NO | NA |
| Wdr13 | ENSMUSG( | ENSMUSG( | -0.11278 | 0.362561 | NO | NO | NA |
| Lcmt1 | ENSMUSG( | ENSMUSG( | 0.066946 | 0.552691 | NO | NO | NA |
| Dcun1d2 | ENSMUSG( | ENSMUSG( | -0.30661 | 4.46E-05 | YES | NO | NA |
| Ahi1 | ENSMUSG( | ENSMUSG( | 0.10753 | 0.416217 | NO | NO | NA |
| Slc8b1 | ENSMUSG( | ENSMUSG( | -0.29594 | 0.00174 | YES | NO | NA |
| Arhgap27c | ENSMUSG( | ENSMUSG( | -0.23563 | 0.03314 | YES | NO | NA |
| Elp5 | ENSMUSG( | ENSMUSG( | 0.177929 | 0.036666 | YES | NO | NA |
| Wbp1 | ENSMUSG( | ENSMUSG( | -0.27133 | 0.001437 | YES | NO | NA |
| Slc25a10 | ENSMUSG( | ENSMUSG( | 0.378901 | 5.71E-06 | YES | NO | NA |
| Fzd5 | ENSMUSG( | ENSMUSG( | -0.42968 | 1.73E-06 | YES | NO | NA |
| Mrps12 | ENSMUSG( | ENSMUSG( | 0.118018 | 0.182925 | NO | NO | NA |
| Btbd2 | ENSMUSG( | ENSMUSG( | -0.22538 | 0.007246 | YES | NO | NA |
| Wrap53 | ENSMUSG( | ENSMUSG( | 0.129866 | 0.224696 | NO | NO | NA |
| Gpank1 | ENSMUSG( | ENSMUSG( | -0.02955 | 0.705171 | NO | NO | NA |

| | | | | | | | |
|---|---|---|---|---|---|---|---|
| Tmem147 | ENSMUSG( | ENSMUSG( | 0.485655 | 6.26E-05 | YES | NO | NA |
| Mpp1 | ENSMUSG( | ENSMUSG( | 0.058729 | 0.652373 | NO | NO | NA |
| Wdr83 | ENSMUSG( | ENSMUSG( | 0.178588 | 0.022039 | YES | NO | NA |
| Psme2b | ENSMUSG( | ENSMUSG( | -0.30157 | 9.80E-07 | YES | NO | NA |
| Tmem18 | ENSMUSG( | ENSMUSG( | -0.10876 | 0.352248 | NO | NO | NA |
| Pycrl | ENSMUSG( | ENSMUSG( | 0.292275 | 0.000117 | YES | NO | NA |
| Dclre1c | ENSMUSG( | ENSMUSG( | -0.07597 | 0.55751 | NO | NO | NA |
| Gid4 | ENSMUSG( | ENSMUSG( | -0.01812 | 0.83474 | NO | NO | NA |
| Cdkn3 | ENSMUSG( | ENSMUSG( | 0.473898 | 1.17E-05 | YES | NO | NA |
| Scnm1 | ENSMUSG( | ENSMUSG( | -0.17102 | 0.088193 | NO | NO | NA |
| Pex14 | ENSMUSG( | ENSMUSG( | -0.03008 | 0.796109 | NO | NO | NA |
| ENSMUSG | ENSMUSG( | ENSMUSG( | 1.308183 | 5.00E-08 | YES | YES | Upregulated |
| Fbxo7 | ENSMUSG( | ENSMUSG( | -0.07886 | 0.423238 | NO | NO | NA |
| Rab33b | ENSMUSG( | ENSMUSG( | -0.41363 | 4.41E-05 | YES | NO | NA |
| Rab37 | ENSMUSG( | ENSMUSG( | -0.24761 | 0.203151 | NO | NO | NA |
| Fads1 | ENSMUSG( | ENSMUSG( | -0.07192 | 0.484488 | NO | NO | NA |
| Chrac1 | ENSMUSG( | ENSMUSG( | 0.106585 | 0.480215 | NO | NO | NA |
| Rara | ENSMUSG( | ENSMUSG( | -1.24472 | 1.88E-12 | YES | YES | Downregulated |
| Lactb2 | ENSMUSG( | ENSMUSG( | -0.14327 | 0.042521 | YES | NO | NA |
| Ankzf1 | ENSMUSG( | ENSMUSG( | -0.19416 | 0.025988 | YES | NO | NA |
| Mus81 | ENSMUSG( | ENSMUSG( | -0.28201 | 0.002461 | YES | NO | NA |
| Ebag9 | ENSMUSG( | ENSMUSG( | -0.06501 | 0.63395 | NO | NO | NA |
| Kdsr | ENSMUSG( | ENSMUSG( | 0.154252 | 0.114114 | NO | NO | NA |
| Katnip | ENSMUSG( | ENSMUSG( | -0.06866 | 0.444657 | NO | NO | NA |
| Trim23 | ENSMUSG( | ENSMUSG( | -0.04219 | 0.757906 | NO | NO | NA |
| BC147527 | ENSMUSG( | ENSMUSG( | -1.24421 | 8.04E-12 | YES | YES | Downregulated |
| Cdkn2a | ENSMUSG( | ENSMUSG( | 0.433322 | 0.018074 | YES | NO | NA |
| Eno3 | ENSMUSG( | ENSMUSG( | 0.597692 | 2.57E-07 | YES | NO | NA |
| Eva1b | ENSMUSG( | ENSMUSG( | -0.52446 | 4.61E-07 | YES | NO | NA |
| Il15ra | ENSMUSG( | ENSMUSG( | -0.15113 | 0.227594 | NO | NO | NA |
| Ivd | ENSMUSG( | ENSMUSG( | 0.369082 | 0.01151 | YES | NO | NA |
| Emp1 | ENSMUSG( | ENSMUSG( | 0.379266 | 0.060434 | NO | NO | NA |
| Gm29719 | ENSMUSG( | ENSMUSG( | 0.150335 | 0.516168 | NO | NO | NA |
| Znhit1 | ENSMUSG( | ENSMUSG( | 0.149064 | 0.254163 | NO | NO | NA |
| Spice1 | ENSMUSG( | ENSMUSG( | -0.71534 | 7.90E-08 | YES | NO | NA |
| Eme2 | ENSMUSG( | ENSMUSG( | 0.049256 | 0.650641 | NO | NO | NA |
| Nfe2l2 | ENSMUSG( | ENSMUSG( | -0.47508 | 1.35E-07 | YES | NO | NA |
| Habp4 | ENSMUSG( | ENSMUSG( | -0.10703 | 0.374279 | NO | NO | NA |
| Hbp1 | ENSMUSG( | ENSMUSG( | -0.52543 | 7.72E-06 | YES | NO | NA |
| Mars2 | ENSMUSG( | ENSMUSG( | 0.019518 | 0.808232 | NO | NO | NA |
| Hsh2d | ENSMUSG( | ENSMUSG( | -0.45767 | 2.59E-06 | YES | NO | NA |
| Bet1l | ENSMUSG( | ENSMUSG( | -0.03335 | 0.677082 | NO | NO | NA |
| R3hcc1 | ENSMUSG( | ENSMUSG( | 0.155413 | 0.253615 | NO | NO | NA |
| Epop | ENSMUSG( | ENSMUSG( | 0.623655 | 0.000127 | YES | NO | NA |
| Eif5a2 | ENSMUSG( | ENSMUSG( | 0.392218 | 0.00777 | YES | NO | NA |
| Iqgap3 | ENSMUSG( | ENSMUSG( | 0.206182 | 0.037218 | YES | NO | NA |
| Vkorc1l1 | ENSMUSG( | ENSMUSG( | 0.205662 | 0.046624 | YES | NO | NA |

| | | | | | | | |
|---|---|---|---|---|---|---|---|
| Fbxl12 | ENSMUSG( | ENSMUSG( | -0.55406 | 1.65E-07 | YES | NO | NA |
| Plcb2 | ENSMUSG( | ENSMUSG( | -1.06563 | 8.11E-08 | YES | YES | Downregulated |
| Tmem268 | ENSMUSG( | ENSMUSG( | -0.12097 | 0.204125 | NO | NO | NA |
| Pex7 | ENSMUSG( | ENSMUSG( | 0.25159 | 0.004972 | YES | NO | NA |
| Cdadc1 | ENSMUSG( | ENSMUSG( | 0.105667 | 0.35722 | NO | NO | NA |
| Lmtk3 | ENSMUSG( | ENSMUSG( | -0.12022 | 0.54169 | NO | NO | NA |
| Msantd4 | ENSMUSG( | ENSMUSG( | 0.001355 | 0.987646 | NO | NO | NA |
| Ttc33 | ENSMUSG( | ENSMUSG( | -0.29394 | 2.02E-05 | YES | NO | NA |
| Ube2l6 | ENSMUSG( | ENSMUSG( | -0.70992 | 2.17E-07 | YES | NO | NA |
| Cln6 | ENSMUSG( | ENSMUSG( | 0.079816 | 0.568223 | NO | NO | NA |
| Itfg2 | ENSMUSG( | ENSMUSG( | 0.090083 | 0.307552 | NO | NO | NA |
| Rps6-ps4 | ENSMUSG( | ENSMUSG( | 0.501548 | 7.88E-06 | YES | NO | NA |
| Firre | ENSMUSG( | ENSMUSG( | 0.041278 | 0.820264 | NO | NO | NA |
| Bpgm | ENSMUSG( | ENSMUSG( | 0.059521 | 0.693256 | NO | NO | NA |
| Smg8 | ENSMUSG( | ENSMUSG( | -0.3687 | 6.15E-08 | YES | NO | NA |
| Setd6 | ENSMUSG( | ENSMUSG( | 0.211027 | 0.047921 | YES | NO | NA |
| Cox20 | ENSMUSG( | ENSMUSG( | -0.20056 | 0.053804 | NO | NO | NA |
| Tspyl4 | ENSMUSG( | ENSMUSG( | -0.54411 | 4.36E-06 | YES | NO | NA |
| Dtd1 | ENSMUSG( | ENSMUSG( | 0.293242 | 0.001351 | YES | NO | NA |
| Ipp | ENSMUSG( | ENSMUSG( | 0.150483 | 0.083333 | NO | NO | NA |
| Xylt1 | ENSMUSG( | ENSMUSG( | -0.10173 | 0.487541 | NO | NO | NA |
| Dmac1 | ENSMUSG( | ENSMUSG( | 0.08873 | 0.267015 | NO | NO | NA |
| Calcoco1 | ENSMUSG( | ENSMUSG( | -0.47086 | 0.000139 | YES | NO | NA |
| Tmed3 | ENSMUSG( | ENSMUSG( | 0.108823 | 0.216788 | NO | NO | NA |
| Prune1 | ENSMUSG( | ENSMUSG( | -0.22647 | 0.006538 | YES | NO | NA |
| Zdhhc17 | ENSMUSG( | ENSMUSG( | -0.07834 | 0.537122 | NO | NO | NA |
| Ccdc102a | ENSMUSG( | ENSMUSG( | 1.134802 | 2.43E-06 | YES | YES | Upregulated |
| 0610012G | ENSMUSG( | ENSMUSG( | 0.276895 | 0.006021 | YES | NO | NA |
| Card6 | ENSMUSG( | ENSMUSG( | -0.72027 | 9.60E-09 | YES | NO | NA |
| Pelo | ENSMUSG( | ENSMUSG( | 0.007042 | 0.954655 | NO | NO | NA |
| Gm20431 | ENSMUSG( | ENSMUSG( | 0.232273 | 0.679132 | NO | NO | NA |
| Sytl1 | ENSMUSG( | ENSMUSG( | 0.022628 | 0.851529 | NO | NO | NA |
| Tbxa2r | ENSMUSG( | ENSMUSG( | -0.12271 | 0.230403 | NO | NO | NA |
| Relb | ENSMUSG( | ENSMUSG( | -0.36039 | 0.00154 | YES | NO | NA |
| Ado | ENSMUSG( | ENSMUSG( | -0.18827 | 0.044059 | YES | NO | NA |
| Eogt | ENSMUSG( | ENSMUSG( | -0.18316 | 0.044559 | YES | NO | NA |
| Snhg12 | ENSMUSG( | ENSMUSG( | 0.884481 | 3.33E-09 | YES | NO | NA |
| Prdm11 | ENSMUSG( | ENSMUSG( | -0.34355 | 2.81E-06 | YES | NO | NA |
| Pgghg | ENSMUSG( | ENSMUSG( | -0.27461 | 0.005127 | YES | NO | NA |
| Poc5 | ENSMUSG( | ENSMUSG( | 0.169758 | 0.065928 | NO | NO | NA |
| Prkag2 | ENSMUSG( | ENSMUSG( | -0.11326 | 0.351916 | NO | NO | NA |
| Gnb4 | ENSMUSG( | ENSMUSG( | -0.7421 | 2.80E-12 | YES | NO | NA |
| Spred2 | ENSMUSG( | ENSMUSG( | -0.06427 | 0.637795 | NO | NO | NA |
| Morrbid | ENSMUSG( | ENSMUSG( | -0.04755 | 0.580861 | NO | NO | NA |
| Dpcd | ENSMUSG( | ENSMUSG( | 0.099168 | 0.449445 | NO | NO | NA |
| Snx13 | ENSMUSG( | ENSMUSG( | -0.02765 | 0.832243 | NO | NO | NA |
| Tbc1d17 | ENSMUSG( | ENSMUSG( | -0.36805 | 9.55E-06 | YES | NO | NA |

| | | | | | | | |
|---|---|---|---|---|---|---|---|
| Klhl12 | ENSMUSG( | ENSMUSG( | 0.122477 | 0.183659 | NO | NO | NA |
| Pdzd11 | ENSMUSG( | ENSMUSG( | -0.03115 | 0.820924 | NO | NO | NA |
| Nle1 | ENSMUSG( | ENSMUSG( | 0.79248 | 2.70E-07 | YES | NO | NA |
| Gm27177 | ENSMUSG( | ENSMUSG( | 0.652129 | 0.000202 | YES | NO | NA |
| Lrrc8d | ENSMUSG( | ENSMUSG( | 0.016482 | 0.843266 | NO | NO | NA |
| Klf4 | ENSMUSG( | ENSMUSG( | 1.638276 | 2.35E-06 | YES | YES | Upregulated |
| Tor3a | ENSMUSG( | ENSMUSG( | -0.19867 | 0.070132 | NO | NO | NA |
| Fam219b | ENSMUSG( | ENSMUSG( | -0.23324 | 0.001924 | YES | NO | NA |
| Rnf41 | ENSMUSG( | ENSMUSG( | -0.19877 | 0.017708 | YES | NO | NA |
| ENSMUSG( | ENSMUSG( | ENSMUSG( | -0.60077 | 4.15E-15 | YES | NO | NA |
| Phf11a | ENSMUSG( | ENSMUSG( | -0.66952 | 7.73E-11 | YES | NO | NA |
| Ppard | ENSMUSG( | ENSMUSG( | -0.11048 | 0.112328 | NO | NO | NA |
| Pigb | ENSMUSG( | ENSMUSG( | -0.10884 | 0.13205 | NO | NO | NA |
| Abca3 | ENSMUSG( | ENSMUSG( | -0.28009 | 0.106362 | NO | NO | NA |
| Rsu1 | ENSMUSG( | ENSMUSG( | 0.245017 | 0.003436 | YES | NO | NA |
| Mcph1 | ENSMUSG( | ENSMUSG( | 0.127018 | 0.166103 | NO | NO | NA |
| Fbxo6 | ENSMUSG( | ENSMUSG( | 0.306702 | 0.020714 | YES | NO | NA |
| Rdm1 | ENSMUSG( | ENSMUSG( | -0.43098 | 0.000224 | YES | NO | NA |
| Washc3 | ENSMUSG( | ENSMUSG( | -0.08165 | 0.275571 | NO | NO | NA |
| Pdlim1 | ENSMUSG( | ENSMUSG( | 0.096965 | 0.469551 | NO | NO | NA |
| Gm15441 | ENSMUSG( | ENSMUSG( | -0.30963 | 0.036652 | YES | NO | NA |
| Tmem11 | ENSMUSG( | ENSMUSG( | 0.123531 | 0.189919 | NO | NO | NA |
| Hibadh | ENSMUSG( | ENSMUSG( | -0.08614 | 0.340311 | NO | NO | NA |
| Gm11508 | ENSMUSG( | ENSMUSG( | -0.05897 | 0.768418 | NO | NO | NA |
| Eef1e1 | ENSMUSG( | ENSMUSG( | 0.061004 | 0.624649 | NO | NO | NA |
| Pnpla8 | ENSMUSG( | ENSMUSG( | -0.06596 | 0.605421 | NO | NO | NA |
| Kif3c | ENSMUSG( | ENSMUSG( | -0.47158 | 3.67E-14 | YES | NO | NA |
| Rce1 | ENSMUSG( | ENSMUSG( | 0.023165 | 0.824786 | NO | NO | NA |
| D030056L: | ENSMUSG( | ENSMUSG( | 0.120864 | 0.344724 | NO | NO | NA |
| Smim36 | ENSMUSG( | ENSMUSG( | 0.752774 | 1.56E-06 | YES | NO | NA |
| Nudcd1 | ENSMUSG( | ENSMUSG( | 0.336437 | 0.000648 | YES | NO | NA |
| Dtx2 | ENSMUSG( | ENSMUSG( | -0.33461 | 0.000198 | YES | NO | NA |
| Tsfm | ENSMUSG( | ENSMUSG( | 0.364316 | 0.000442 | YES | NO | NA |
| ENSMUSG | ENSMUSG( | ENSMUSG( | 0.038277 | 0.656664 | NO | NO | NA |
| Trbv19 | ENSMUSG( | ENSMUSG( | -0.58759 | 0.008886 | YES | NO | NA |
| Mmgt2 | ENSMUSG( | ENSMUSG( | -0.07144 | 0.450816 | NO | NO | NA |
| Nufip1 | ENSMUSG( | ENSMUSG( | 0.390519 | 4.11E-06 | YES | NO | NA |
| Hscb | ENSMUSG( | ENSMUSG( | 0.059206 | 0.626663 | NO | NO | NA |
| Atg4a | ENSMUSG( | ENSMUSG( | 0.061148 | 0.648432 | NO | NO | NA |
| Csf1 | ENSMUSG( | ENSMUSG( | 0.406845 | 0.018671 | YES | NO | NA |
| Gmds | ENSMUSG( | ENSMUSG( | 0.374697 | 4.28E-05 | YES | NO | NA |
| Ssh3 | ENSMUSG( | ENSMUSG( | 0.060694 | 0.665935 | NO | NO | NA |
| Taf5 | ENSMUSG( | ENSMUSG( | 0.378246 | 7.03E-09 | YES | NO | NA |
| Sgf29 | ENSMUSG( | ENSMUSG( | 0.420773 | 0.000205 | YES | NO | NA |
| Thg1l | ENSMUSG( | ENSMUSG( | 0.694134 | 2.95E-08 | YES | NO | NA |
| Hrob | ENSMUSG( | ENSMUSG( | 0.22197 | 0.013134 | YES | NO | NA |
| Cd1d1 | ENSMUSG( | ENSMUSG( | -0.52799 | 1.64E-05 | YES | NO | NA |

| | | | | | | | |
|---|---|---|---|---|---|---|---|
| Pcdhgc3 | ENSMUSG( | ENSMUSG( | -0.47662 | 4.35E-05 | YES | NO | NA |
| Mon1a | ENSMUSG( | ENSMUSG( | -0.17121 | 0.040764 | YES | NO | NA |
| Akap12 | ENSMUSG( | ENSMUSG( | -0.15267 | 0.21335 | NO | NO | NA |
| Frmd4a | ENSMUSG( | ENSMUSG( | 0.201664 | 0.027048 | YES | NO | NA |
| Nln | ENSMUSG( | ENSMUSG( | 0.248479 | 0.001046 | YES | NO | NA |
| Zbtb43 | ENSMUSG( | ENSMUSG( | -0.24025 | 0.000803 | YES | NO | NA |
| Zfp386 | ENSMUSG( | ENSMUSG( | 0.025293 | 0.854593 | NO | NO | NA |
| Ndufb2 | ENSMUSG( | ENSMUSG( | 0.187337 | 0.061919 | NO | NO | NA |
| Zfp395 | ENSMUSG( | ENSMUSG( | -0.9402 | 6.98E-08 | YES | NO | NA |
| Stimate | ENSMUSG( | ENSMUSG( | 0.193852 | 0.078381 | NO | NO | NA |
| Polr1e | ENSMUSG( | ENSMUSG( | 0.24871 | 0.02161 | YES | NO | NA |
| Fnip1 | ENSMUSG( | ENSMUSG( | -0.01881 | 0.899231 | NO | NO | NA |
| Tfb2m | ENSMUSG( | ENSMUSG( | 0.413207 | 0.000352 | YES | NO | NA |
| Stoml1 | ENSMUSG( | ENSMUSG( | 0.425131 | 0.003764 | YES | NO | NA |
| Itpripl2 | ENSMUSG( | ENSMUSG( | -0.37883 | 0.012467 | YES | NO | NA |
| Mcts1 | ENSMUSG( | ENSMUSG( | 0.132258 | 0.091511 | NO | NO | NA |
| Chst10 | ENSMUSG( | ENSMUSG( | -0.74095 | 7.36E-10 | YES | NO | NA |
| Dcun1d3 | ENSMUSG( | ENSMUSG( | -0.43036 | 6.29E-05 | YES | NO | NA |
| 4930581F2 | ENSMUSG( | ENSMUSG( | -0.27715 | 0.000981 | YES | NO | NA |
| A430005L1 | ENSMUSG( | ENSMUSG( | 0.164283 | 0.11068 | NO | NO | NA |
| Prr33 | ENSMUSG( | ENSMUSG( | -0.52868 | 1.42E-09 | YES | NO | NA |
| Tagap1 | ENSMUSG( | ENSMUSG( | 0.071255 | 0.518523 | NO | NO | NA |
| Rfng | ENSMUSG( | ENSMUSG( | 0.043054 | 0.682283 | NO | NO | NA |
| Asns | ENSMUSG( | ENSMUSG( | 1.016909 | 1.70E-05 | YES | YES | Upregulated |
| Ermard | ENSMUSG( | ENSMUSG( | -0.13904 | 0.324184 | NO | NO | NA |
| Arhgap21 | ENSMUSG( | ENSMUSG( | 0.747913 | 2.96E-05 | YES | NO | NA |
| Rnpc3 | ENSMUSG( | ENSMUSG( | -0.04044 | 0.751588 | NO | NO | NA |
| Srfbp1 | ENSMUSG( | ENSMUSG( | 0.020213 | 0.903965 | NO | NO | NA |
| Zfp182 | ENSMUSG( | ENSMUSG( | -0.70982 | 1.00E-07 | YES | NO | NA |
| Gypc | ENSMUSG( | ENSMUSG( | 0.135269 | 0.333753 | NO | NO | NA |
| Cnnm4 | ENSMUSG( | ENSMUSG( | -0.15286 | 0.005134 | YES | NO | NA |
| Rhoq | ENSMUSG( | ENSMUSG( | -0.39571 | 0.015566 | YES | NO | NA |
| Glrx2 | ENSMUSG( | ENSMUSG( | -0.03722 | 0.751327 | NO | NO | NA |
| Dennd1a | ENSMUSG( | ENSMUSG( | -0.12046 | 0.276646 | NO | NO | NA |
| Tmem104 | ENSMUSG( | ENSMUSG( | -0.37285 | 0.000174 | YES | NO | NA |
| Nceh1 | ENSMUSG( | ENSMUSG( | -0.02392 | 0.829364 | NO | NO | NA |
| Accs | ENSMUSG( | ENSMUSG( | -0.02373 | 0.847785 | NO | NO | NA |
| Tspan2 | ENSMUSG( | ENSMUSG( | 0.245966 | 0.219116 | NO | NO | NA |
| Zswim4 | ENSMUSG( | ENSMUSG( | 0.053131 | 0.703089 | NO | NO | NA |
| Scaper | ENSMUSG( | ENSMUSG( | -0.0556 | 0.700622 | NO | NO | NA |
| Cysltr2 | ENSMUSG( | ENSMUSG( | -0.18832 | 0.07833 | NO | NO | NA |
| Igsf8 | ENSMUSG( | ENSMUSG( | 0.100137 | 0.297374 | NO | NO | NA |
| Mlkl | ENSMUSG( | ENSMUSG( | 0.009574 | 0.937826 | NO | NO | NA |
| Adprm | ENSMUSG( | ENSMUSG( | -0.39275 | 3.88E-07 | YES | NO | NA |
| Gm10524 | ENSMUSG( | ENSMUSG( | 0.261201 | 0.30875 | NO | NO | NA |
| Rbmx2 | ENSMUSG( | ENSMUSG( | 0.000308 | 0.99843 | NO | NO | NA |
| Sirt6 | ENSMUSG( | ENSMUSG( | -0.14542 | 0.059756 | NO | NO | NA |

| | | | | | | | |
|---|---|---|---|---|---|---|---|
| Eif2ak4 | ENSMUSG( | ENSMUSG( | 0.159483 | 0.09841 | NO | NO | NA |
| Ocel1 | ENSMUSG( | ENSMUSG( | -0.25724 | 0.002034 | YES | NO | NA |
| Tmem243 | ENSMUSG( | ENSMUSG( | 0.206667 | 0.008076 | YES | NO | NA |
| Erlin2 | ENSMUSG( | ENSMUSG( | -0.0973 | 0.451427 | NO | NO | NA |
| Cenpi | ENSMUSG( | ENSMUSG( | 0.274793 | 0.004375 | YES | NO | NA |
| Siae | ENSMUSG( | ENSMUSG( | -0.19733 | 0.042633 | YES | NO | NA |
| Cyb561d2 | ENSMUSG( | ENSMUSG( | -0.15479 | 0.071094 | NO | NO | NA |
| Il1rap | ENSMUSG( | ENSMUSG( | -0.3371 | 1.84E-06 | YES | NO | NA |
| Icam2 | ENSMUSG( | ENSMUSG( | -0.4318 | 7.22E-07 | YES | NO | NA |
| Zfp866 | ENSMUSG( | ENSMUSG( | -0.06751 | 0.601138 | NO | NO | NA |
| Chpt1 | ENSMUSG( | ENSMUSG( | -0.2265 | 0.054334 | NO | NO | NA |
| Dclre1b | ENSMUSG( | ENSMUSG( | 0.187788 | 0.014952 | YES | NO | NA |
| Nkiras2 | ENSMUSG( | ENSMUSG( | 0.037838 | 0.724938 | NO | NO | NA |
| Yod1 | ENSMUSG( | ENSMUSG( | -0.22548 | 0.009565 | YES | NO | NA |
| Mtfr2 | ENSMUSG( | ENSMUSG( | 0.290675 | 2.81E-06 | YES | NO | NA |
| Bud13 | ENSMUSG( | ENSMUSG( | 0.040096 | 0.667475 | NO | NO | NA |
| Gm37691 | ENSMUSG( | ENSMUSG( | -0.99235 | 2.16E-12 | YES | NO | NA |
| Arhgap12 | ENSMUSG( | ENSMUSG( | -0.16709 | 0.120955 | NO | NO | NA |
| Paxx | ENSMUSG( | ENSMUSG( | -0.16057 | 0.032169 | YES | NO | NA |
| Baz2b | ENSMUSG( | ENSMUSG( | -0.45487 | 0.000193 | YES | NO | NA |
| Telo2 | ENSMUSG( | ENSMUSG( | 0.426583 | 1.36E-05 | YES | NO | NA |
| Cenpm | ENSMUSG( | ENSMUSG( | 0.467783 | 0.00045 | YES | NO | NA |
| Galns | ENSMUSG( | ENSMUSG( | -0.01517 | 0.852776 | NO | NO | NA |
| Pccb | ENSMUSG( | ENSMUSG( | 0.775525 | 1.01E-06 | YES | NO | NA |
| Zcchc4 | ENSMUSG( | ENSMUSG( | 0.16235 | 0.090084 | NO | NO | NA |
| Rnaseh1 | ENSMUSG( | ENSMUSG( | 0.292304 | 0.001993 | YES | NO | NA |
| Nagpa | ENSMUSG( | ENSMUSG( | 0.007943 | 0.951017 | NO | NO | NA |
| Zfp329 | ENSMUSG( | ENSMUSG( | -0.50873 | 4.03E-05 | YES | NO | NA |
| Faap20 | ENSMUSG( | ENSMUSG( | 0.025268 | 0.754962 | NO | NO | NA |
| Fbxl8 | ENSMUSG( | ENSMUSG( | -0.22893 | 0.000837 | YES | NO | NA |
| Ubxn11 | ENSMUSG( | ENSMUSG( | -0.00793 | 0.951721 | NO | NO | NA |
| Cdk5 | ENSMUSG( | ENSMUSG( | -0.17517 | 0.104424 | NO | NO | NA |
| Coa3 | ENSMUSG( | ENSMUSG( | -0.08429 | 0.464432 | NO | NO | NA |
| Dars2 | ENSMUSG( | ENSMUSG( | -0.10009 | 0.563166 | NO | NO | NA |
| Bc1 | ENSMUSG( | ENSMUSG( | -0.36954 | NA | NO | NO | NA |
| Rint1 | ENSMUSG( | ENSMUSG( | 0.092031 | 0.303529 | NO | NO | NA |
| Tmem191( | ENSMUSG( | ENSMUSG( | 0.189985 | 0.17406 | NO | NO | NA |
| Wnt10b | ENSMUSG( | ENSMUSG( | 2.213497 | 1.55E-10 | YES | YES | Upregulated |
| Elmod2 | ENSMUSG( | ENSMUSG( | -0.35897 | 4.10E-05 | YES | NO | NA |
| Clgn | ENSMUSG( | ENSMUSG( | 0.326754 | 0.050161 | NO | NO | NA |
| Vat1 | ENSMUSG( | ENSMUSG( | 0.210641 | 0.004808 | YES | NO | NA |
| Ldlr | ENSMUSG( | ENSMUSG( | 1.166437 | 5.78E-07 | YES | YES | Upregulated |
| Ar | ENSMUSG( | ENSMUSG( | 0.042052 | 0.78906 | NO | NO | NA |
| Dnlz | ENSMUSG( | ENSMUSG( | -0.03109 | 0.789532 | NO | NO | NA |
| Rpusd4 | ENSMUSG( | ENSMUSG( | -0.05687 | 0.452392 | NO | NO | NA |
| Ptger2 | ENSMUSG( | ENSMUSG( | -0.60262 | 3.09E-05 | YES | NO | NA |
| Mplkip | ENSMUSG( | ENSMUSG( | 0.017324 | 0.887796 | NO | NO | NA |

| | | | | | | | |
|---|---|---|---|---|---|---|---|
| Tcf7 | ENSMUSG( | ENSMUSG( | -0.84066 | 1.49E-07 | YES | NO | NA |
| Zfand2b | ENSMUSG( | ENSMUSG( | 0.108899 | 0.340554 | NO | NO | NA |
| Dync2h1 | ENSMUSG( | ENSMUSG( | 0.426565 | 0.010583 | YES | NO | NA |
| Zfp408 | ENSMUSG( | ENSMUSG( | -0.08386 | 0.361227 | NO | NO | NA |
| Slc39a10 | ENSMUSG( | ENSMUSG( | -0.23033 | 0.127714 | NO | NO | NA |
| Lamb3 | ENSMUSG( | ENSMUSG( | -0.14335 | 0.350029 | NO | NO | NA |
| Mfsd11 | ENSMUSG( | ENSMUSG( | -0.37362 | 1.54E-05 | YES | NO | NA |
| Tmtc4 | ENSMUSG( | ENSMUSG( | -0.46115 | 3.52E-07 | YES | NO | NA |
| Zxdb | ENSMUSG( | ENSMUSG( | -0.40344 | 4.14E-05 | YES | NO | NA |
| Mrpl11 | ENSMUSG( | ENSMUSG( | 0.477262 | 7.64E-08 | YES | NO | NA |
| Txnl4a | ENSMUSG( | ENSMUSG( | -0.16903 | 0.084942 | NO | NO | NA |
| Nup37 | ENSMUSG( | ENSMUSG( | 0.454597 | 1.17E-08 | YES | NO | NA |
| Swt1 | ENSMUSG( | ENSMUSG( | -0.20991 | 0.024789 | YES | NO | NA |
| Gpn2 | ENSMUSG( | ENSMUSG( | 0.290366 | 0.000103 | YES | NO | NA |
| Nprl3 | ENSMUSG( | ENSMUSG( | 0.27979 | 0.018868 | YES | NO | NA |
| Zfp322a | ENSMUSG( | ENSMUSG( | -0.27324 | 0.000223 | YES | NO | NA |
| Cwf19l1 | ENSMUSG( | ENSMUSG( | 0.173204 | 0.004476 | YES | NO | NA |
| Orai3 | ENSMUSG( | ENSMUSG( | -0.26309 | 0.004537 | YES | NO | NA |
| Dffb | ENSMUSG( | ENSMUSG( | 0.156427 | 0.126385 | NO | NO | NA |
| Prrt1 | ENSMUSG( | ENSMUSG( | 0.296908 | 0.00394 | YES | NO | NA |
| Txndc16 | ENSMUSG( | ENSMUSG( | -0.35318 | 0.000193 | YES | NO | NA |
| Socs6 | ENSMUSG( | ENSMUSG( | 0.060533 | 0.566196 | NO | NO | NA |
| Gm12895 | ENSMUSG( | ENSMUSG( | 1.258912 | 0.645714 | NO | NO | NA |
| Serinc5 | ENSMUSG( | ENSMUSG( | -0.33784 | 0.010733 | YES | NO | NA |
| Gm12992 | ENSMUSG( | ENSMUSG( | 0.231235 | 0.015872 | YES | NO | NA |
| Slc33a1 | ENSMUSG( | ENSMUSG( | 0.159632 | 0.05662 | NO | NO | NA |
| Ripor1 | ENSMUSG( | ENSMUSG( | -0.47461 | 1.43E-06 | YES | NO | NA |
| Tmem70 | ENSMUSG( | ENSMUSG( | 0.216487 | 0.005947 | YES | NO | NA |
| Slc15a2 | ENSMUSG( | ENSMUSG( | -1.17225 | 0.161751 | NO | NO | NA |
| Camk2b | ENSMUSG( | ENSMUSG( | 3.022021 | 7.28E-41 | YES | YES | Upregulated |
| Tmem39a | ENSMUSG( | ENSMUSG( | 0.18641 | 0.025344 | YES | NO | NA |
| Itga3 | ENSMUSG( | ENSMUSG( | 0.393254 | 0.004483 | YES | NO | NA |
| Lrrc61 | ENSMUSG( | ENSMUSG( | -0.56422 | 1.21E-10 | YES | NO | NA |
| Tbc1d23 | ENSMUSG( | ENSMUSG( | -0.04739 | 0.593657 | NO | NO | NA |
| ENSMUSG( | ENSMUSG( | ENSMUSG( | -0.81369 | 1.15E-09 | YES | NO | NA |
| Zfp523 | ENSMUSG( | ENSMUSG( | 0.410167 | 0.00219 | YES | NO | NA |
| B4galt3 | ENSMUSG( | ENSMUSG( | -0.19212 | 0.054928 | NO | NO | NA |
| Trnau1ap | ENSMUSG( | ENSMUSG( | 0.101723 | 0.282122 | NO | NO | NA |
| Gatd1 | ENSMUSG( | ENSMUSG( | -0.10084 | 0.327932 | NO | NO | NA |
| Calhm2 | ENSMUSG( | ENSMUSG( | -0.58959 | 1.47E-09 | YES | NO | NA |
| Slc9a5 | ENSMUSG( | ENSMUSG( | 0.770942 | 7.03E-08 | YES | NO | NA |
| Fndc9 | ENSMUSG( | ENSMUSG( | 0.09208 | 0.554294 | NO | NO | NA |
| Repin1 | ENSMUSG( | ENSMUSG( | 0.51507 | 4.06E-06 | YES | NO | NA |
| Cstb | ENSMUSG( | ENSMUSG( | 0.121014 | 0.27189 | NO | NO | NA |
| Tk2 | ENSMUSG( | ENSMUSG( | -0.24509 | 0.05358 | NO | NO | NA |
| Dgkq | ENSMUSG( | ENSMUSG( | -0.21655 | 0.020489 | YES | NO | NA |
| Hyls1 | ENSMUSG( | ENSMUSG( | 0.00457 | 0.968966 | NO | NO | NA |

| | | | | | | | |
|---|---|---|---|---|---|---|---|
| Ap5b1 | ENSMUSG( | ENSMUSG( | -0.65804 | 6.14E-07 | YES | NO | NA |
| 1700037H | ENSMUSG( | ENSMUSG( | 0.276712 | 0.006837 | YES | NO | NA |
| Rfx5 | ENSMUSG( | ENSMUSG( | -0.30758 | 0.013146 | YES | NO | NA |
| Xlr4a | ENSMUSG( | ENSMUSG( | -0.31742 | 0.021177 | YES | NO | NA |
| Rnf170 | ENSMUSG( | ENSMUSG( | -0.03241 | 0.843184 | NO | NO | NA |
| Ercc1 | ENSMUSG( | ENSMUSG( | 0.113731 | 0.338529 | NO | NO | NA |
| Inpp5f | ENSMUSG( | ENSMUSG( | -0.11719 | 0.120917 | NO | NO | NA |
| Ifitm10 | ENSMUSG( | ENSMUSG( | -0.74263 | 1.28E-05 | YES | NO | NA |
| Gm38244 | ENSMUSG( | ENSMUSG( | -1.10981 | 2.34E-12 | YES | YES | Downregulated |
| Pikfyve | ENSMUSG( | ENSMUSG( | -0.08753 | 0.244262 | NO | NO | NA |
| Ift27 | ENSMUSG( | ENSMUSG( | 0.262254 | 0.002979 | YES | NO | NA |
| Agk | ENSMUSG( | ENSMUSG( | 0.264074 | 0.000117 | YES | NO | NA |
| Atrip | ENSMUSG( | ENSMUSG( | 0.158932 | 0.04163 | YES | NO | NA |
| Ighmbp2 | ENSMUSG( | ENSMUSG( | -0.09921 | 0.232794 | NO | NO | NA |
| Tprn | ENSMUSG( | ENSMUSG( | 0.124305 | 0.203392 | NO | NO | NA |
| Lrp8 | ENSMUSG( | ENSMUSG( | 0.60259 | 5.59E-07 | YES | NO | NA |
| Rccd1 | ENSMUSG( | ENSMUSG( | 0.122741 | 0.315652 | NO | NO | NA |
| Klhl2 | ENSMUSG( | ENSMUSG( | -0.06816 | 0.397671 | NO | NO | NA |
| Nr2c2ap | ENSMUSG( | ENSMUSG( | 0.206684 | 0.076071 | NO | NO | NA |
| Dbt | ENSMUSG( | ENSMUSG( | 0.111922 | 0.407211 | NO | NO | NA |
| Rbm47 | ENSMUSG( | ENSMUSG( | 0.41564 | 6.98E-06 | YES | NO | NA |
| Nepro | ENSMUSG( | ENSMUSG( | 0.157606 | 0.155609 | NO | NO | NA |
| Lin37 | ENSMUSG( | ENSMUSG( | -0.25174 | 0.002424 | YES | NO | NA |
| Rfxank | ENSMUSG( | ENSMUSG( | -0.08572 | 0.464811 | NO | NO | NA |
| Pcgf6 | ENSMUSG( | ENSMUSG( | 0.041024 | 0.629327 | NO | NO | NA |
| Sik2 | ENSMUSG( | ENSMUSG( | -0.32252 | 0.00015 | YES | NO | NA |
| Manea | ENSMUSG( | ENSMUSG( | 0.04824 | 0.740776 | NO | NO | NA |
| Vrk2 | ENSMUSG( | ENSMUSG( | -0.18435 | 0.071504 | NO | NO | NA |
| Mrpl46 | ENSMUSG( | ENSMUSG( | 0.312936 | 0.00049 | YES | NO | NA |
| Polr3gl | ENSMUSG( | ENSMUSG( | -0.00104 | 0.993761 | NO | NO | NA |
| Vps18 | ENSMUSG( | ENSMUSG( | -0.00675 | 0.945479 | NO | NO | NA |
| U2af1l4 | ENSMUSG( | ENSMUSG( | 0.04482 | 0.676357 | NO | NO | NA |
| Gins3 | ENSMUSG( | ENSMUSG( | 0.107926 | 0.25122 | NO | NO | NA |
| Marveld2 | ENSMUSG( | ENSMUSG( | 0.377043 | 0.001225 | YES | NO | NA |
| Cfap36 | ENSMUSG( | ENSMUSG( | 0.445347 | 0.001459 | YES | NO | NA |
| Ska1 | ENSMUSG( | ENSMUSG( | 0.219938 | 0.111612 | NO | NO | NA |
| Dennd10 | ENSMUSG( | ENSMUSG( | -0.27946 | 0.000837 | YES | NO | NA |
| Ift80 | ENSMUSG( | ENSMUSG( | -0.58445 | 0.006348 | YES | NO | NA |
| Steap3 | ENSMUSG( | ENSMUSG( | 0.235035 | 0.162412 | NO | NO | NA |
| Bach1 | ENSMUSG( | ENSMUSG( | -0.64257 | 7.61E-12 | YES | NO | NA |
| Zfp653 | ENSMUSG( | ENSMUSG( | 0.082366 | 0.542176 | NO | NO | NA |
| Lsm1 | ENSMUSG( | ENSMUSG( | 0.062229 | 0.577933 | NO | NO | NA |
| Slc31a1 | ENSMUSG( | ENSMUSG( | 0.588638 | 1.07E-05 | YES | NO | NA |
| Abcd3 | ENSMUSG( | ENSMUSG( | 0.127458 | 0.292762 | NO | NO | NA |
| Zfpl1 | ENSMUSG( | ENSMUSG( | -0.06093 | 0.562282 | NO | NO | NA |
| Dus2 | ENSMUSG( | ENSMUSG( | -0.38027 | 0.000377 | YES | NO | NA |
| Endov | ENSMUSG( | ENSMUSG( | -0.13057 | 0.35193 | NO | NO | NA |

| | | | | | | | |
|---|---|---|---|---|---|---|---|
| Pyroxd1 | ENSMUSG( | ENSMUSG( | 0.049102 | 0.715024 | NO | NO | NA |
| Exosc1 | ENSMUSG( | ENSMUSG( | 0.290219 | 0.001397 | YES | NO | NA |
| Tspan5 | ENSMUSG( | ENSMUSG( | -0.66364 | 1.71E-07 | YES | NO | NA |
| Tango2 | ENSMUSG( | ENSMUSG( | -0.19615 | 0.011533 | YES | NO | NA |
| Tctn1 | ENSMUSG( | ENSMUSG( | 0.141266 | 0.188133 | NO | NO | NA |
| Kifbp | ENSMUSG( | ENSMUSG( | -0.17796 | 0.105946 | NO | NO | NA |
| Sil1 | ENSMUSG( | ENSMUSG( | 0.707104 | 5.60E-05 | YES | NO | NA |
| Med9 | ENSMUSG( | ENSMUSG( | 0.407053 | 8.59E-05 | YES | NO | NA |
| Mipep | ENSMUSG( | ENSMUSG( | 0.52046 | 2.86E-06 | YES | NO | NA |
| Foxd2os | ENSMUSG( | ENSMUSG( | 0.154642 | 0.02941 | YES | NO | NA |
| Cyp20a1 | ENSMUSG( | ENSMUSG( | 0.295767 | 0.003745 | YES | NO | NA |
| Lpar5 | ENSMUSG( | ENSMUSG( | -0.56534 | 2.30E-07 | YES | NO | NA |
| Zfp12 | ENSMUSG( | ENSMUSG( | -0.31407 | 0.010936 | YES | NO | NA |
| Gcdh | ENSMUSG( | ENSMUSG( | -0.10825 | 0.324996 | NO | NO | NA |
| Myo1h | ENSMUSG( | ENSMUSG( | -0.06789 | 0.731835 | NO | NO | NA |
| Msantd2 | ENSMUSG( | ENSMUSG( | -0.04108 | 0.722103 | NO | NO | NA |
| Mrps18b | ENSMUSG( | ENSMUSG( | 0.614147 | 2.43E-09 | YES | NO | NA |
| Tnfaip8l1 | ENSMUSG( | ENSMUSG( | -0.05328 | 0.694142 | NO | NO | NA |
| Fbxo8 | ENSMUSG( | ENSMUSG( | -0.1506 | 0.067307 | NO | NO | NA |
| Zfp827 | ENSMUSG( | ENSMUSG( | -0.24865 | 0.024146 | YES | NO | NA |
| Slc12a9 | ENSMUSG( | ENSMUSG( | -0.2969 | 0.001585 | YES | NO | NA |
| BC055324 | ENSMUSG( | ENSMUSG( | 0.429407 | 1.08E-08 | YES | NO | NA |
| Gpatch1 | ENSMUSG( | ENSMUSG( | -0.05123 | 0.690509 | NO | NO | NA |
| Brat1 | ENSMUSG( | ENSMUSG( | -0.21421 | 0.000331 | YES | NO | NA |
| Dmap1 | ENSMUSG( | ENSMUSG( | 0.091908 | 0.395362 | NO | NO | NA |
| Arf2 | ENSMUSG( | ENSMUSG( | 0.191776 | 0.211222 | NO | NO | NA |
| Dynlt3 | ENSMUSG( | ENSMUSG( | -0.00866 | 0.946459 | NO | NO | NA |
| 2700049A( | ENSMUSG( | ENSMUSG( | -0.09438 | 0.304247 | NO | NO | NA |
| Adck2 | ENSMUSG( | ENSMUSG( | -0.26586 | 0.000196 | YES | NO | NA |
| Cmc1 | ENSMUSG( | ENSMUSG( | 0.007856 | 0.938698 | NO | NO | NA |
| Tnfaip1 | ENSMUSG( | ENSMUSG( | 0.404448 | 0.000163 | YES | NO | NA |
| Lat2 | ENSMUSG( | ENSMUSG( | -0.19061 | 0.194578 | NO | NO | NA |
| Aacs | ENSMUSG( | ENSMUSG( | 0.544106 | 0.000536 | YES | NO | NA |
| Zxdc | ENSMUSG( | ENSMUSG( | -0.0301 | 0.757248 | NO | NO | NA |
| Vcpkmt | ENSMUSG( | ENSMUSG( | -0.00832 | 0.950631 | NO | NO | NA |
| Zmat5 | ENSMUSG( | ENSMUSG( | -0.05478 | 0.576298 | NO | NO | NA |
| Mrpl34 | ENSMUSG( | ENSMUSG( | 0.232154 | 0.005637 | YES | NO | NA |
| Mrpl50 | ENSMUSG( | ENSMUSG( | 0.201188 | 0.008372 | YES | NO | NA |
| Atraid | ENSMUSG( | ENSMUSG( | 0.114773 | 0.467096 | NO | NO | NA |
| Ccdc77 | ENSMUSG( | ENSMUSG( | 0.011577 | 0.914757 | NO | NO | NA |
| Pcgf2 | ENSMUSG( | ENSMUSG( | -0.04814 | 0.714029 | NO | NO | NA |
| Psmg2 | ENSMUSG( | ENSMUSG( | 0.258341 | 0.010304 | YES | NO | NA |
| Smim20 | ENSMUSG( | ENSMUSG( | 0.040602 | 0.715727 | NO | NO | NA |
| LTO1 | ENSMUSG( | ENSMUSG( | -0.26706 | 0.004378 | YES | NO | NA |
| Zfp830 | ENSMUSG( | ENSMUSG( | -0.0756 | 0.462135 | NO | NO | NA |
| Slc52a3 | ENSMUSG( | ENSMUSG( | -0.15709 | 0.292026 | NO | NO | NA |
| Kptn | ENSMUSG( | ENSMUSG( | -0.0964 | 0.364181 | NO | NO | NA |

| | | | | | | | |
|---|---|---|---|---|---|---|---|
| Slc30a4 | ENSMUSG( | ENSMUSG( | 1.310047 | 8.95E-11 | YES | YES | Upregulated |
| Trappc1 | ENSMUSG( | ENSMUSG( | -0.01949 | 0.803364 | NO | NO | NA |
| Utp25 | ENSMUSG( | ENSMUSG( | 0.204295 | 0.01128 | YES | NO | NA |
| Ascc1 | ENSMUSG( | ENSMUSG( | 0.066866 | 0.574941 | NO | NO | NA |
| Zfp553 | ENSMUSG( | ENSMUSG( | -0.20769 | 0.001024 | YES | NO | NA |
| Abhd17c | ENSMUSG( | ENSMUSG( | 0.404236 | 2.18E-07 | YES | NO | NA |
| Ube2t | ENSMUSG( | ENSMUSG( | -0.01772 | 0.868932 | NO | NO | NA |
| Eif2b3 | ENSMUSG( | ENSMUSG( | 0.599336 | 6.15E-06 | YES | NO | NA |
| Ubxn2b | ENSMUSG( | ENSMUSG( | -0.05177 | 0.647663 | NO | NO | NA |
| Pcbd2 | ENSMUSG( | ENSMUSG( | -0.14665 | 0.414988 | NO | NO | NA |
| Klc2 | ENSMUSG( | ENSMUSG( | -0.09265 | 0.414327 | NO | NO | NA |
| Gemin7 | ENSMUSG( | ENSMUSG( | -0.19442 | 0.027698 | YES | NO | NA |
| Scarb1 | ENSMUSG( | ENSMUSG( | 0.847512 | 1.01E-12 | YES | NO | NA |
| Polr3h | ENSMUSG( | ENSMUSG( | 0.761336 | 4.79E-09 | YES | NO | NA |
| Parp6 | ENSMUSG( | ENSMUSG( | -0.39282 | 1.18E-06 | YES | NO | NA |
| Yju2 | ENSMUSG( | ENSMUSG( | 0.017591 | 0.895822 | NO | NO | NA |
| Ccrl2 | ENSMUSG( | ENSMUSG( | -0.45453 | 0.002851 | YES | NO | NA |
| Ubiad1 | ENSMUSG( | ENSMUSG( | 0.752578 | 8.70E-27 | YES | NO | NA |
| Gm29721 | ENSMUSG( | ENSMUSG( | 0.251628 | 0.06 | NO | NO | NA |
| Stx17 | ENSMUSG( | ENSMUSG( | 0.110378 | 0.388409 | NO | NO | NA |
| E130308A | ENSMUSG( | ENSMUSG( | -0.1616 | 0.252204 | NO | NO | NA |
| Mier2 | ENSMUSG( | ENSMUSG( | -0.06985 | 0.435507 | NO | NO | NA |
| Tmem120a | ENSMUSG( | ENSMUSG( | 0.026204 | 0.845649 | NO | NO | NA |
| Tmem62 | ENSMUSG( | ENSMUSG( | -0.15335 | 0.141422 | NO | NO | NA |
| Bcat1 | ENSMUSG( | ENSMUSG( | 0.995419 | 5.78E-06 | YES | NO | NA |
| Cfap298 | ENSMUSG( | ENSMUSG( | 0.499757 | 7.22E-05 | YES | NO | NA |
| Dnaaf2 | ENSMUSG( | ENSMUSG( | 0.023146 | 0.854133 | NO | NO | NA |
| Gm14236 | ENSMUSG( | ENSMUSG( | 0.743338 | 0.401302 | NO | NO | NA |
| Gas2 | ENSMUSG( | ENSMUSG( | 0.683909 | 0.000226 | YES | NO | NA |
| Coq5 | ENSMUSG( | ENSMUSG( | 0.263827 | 0.000112 | YES | NO | NA |
| Tbc1d2 | ENSMUSG( | ENSMUSG( | 0.437415 | 0.000497 | YES | NO | NA |
| Lrrc42 | ENSMUSG( | ENSMUSG( | -0.20216 | 0.04355 | YES | NO | NA |
| Znhit2 | ENSMUSG( | ENSMUSG( | 0.219628 | 0.036005 | YES | NO | NA |
| Tmem126a | ENSMUSG( | ENSMUSG( | 0.079944 | 0.476489 | NO | NO | NA |
| Zfyve19 | ENSMUSG( | ENSMUSG( | 0.10327 | 0.131087 | NO | NO | NA |
| Zfp277 | ENSMUSG( | ENSMUSG( | 0.111645 | 0.266125 | NO | NO | NA |
| Mrps11 | ENSMUSG( | ENSMUSG( | 0.264387 | 0.006617 | YES | NO | NA |
| Tgfbr1 | ENSMUSG( | ENSMUSG( | -0.41167 | 3.90E-05 | YES | NO | NA |
| Mrpl35 | ENSMUSG( | ENSMUSG( | 0.422073 | 1.03E-07 | YES | NO | NA |
| Slc11a2 | ENSMUSG( | ENSMUSG( | 0.414961 | 1.04E-05 | YES | NO | NA |
| Umad1 | ENSMUSG( | ENSMUSG( | -0.06956 | 0.491395 | NO | NO | NA |
| Amz2 | ENSMUSG( | ENSMUSG( | 0.317254 | 0.036524 | YES | NO | NA |
| Rufy1 | ENSMUSG( | ENSMUSG( | -0.16301 | 0.11648 | NO | NO | NA |
| Sgcb | ENSMUSG( | ENSMUSG( | 0.132854 | 0.259206 | NO | NO | NA |
| Lrig2 | ENSMUSG( | ENSMUSG( | -0.30956 | 0.004397 | YES | NO | NA |
| Zfp141 | ENSMUSG( | ENSMUSG( | -0.09933 | 0.488317 | NO | NO | NA |
| Ephb6 | ENSMUSG( | ENSMUSG( | -0.31274 | 0.07265 | NO | NO | NA |

| | | | | | | | |
|---|---|---|---|---|---|---|---|
| Ttc27 | ENSMUSG( | ENSMUSG( | 0.683183 | 5.00E-08 | YES | NO | NA |
| Pak4 | ENSMUSG( | ENSMUSG( | -0.22332 | 0.048671 | YES | NO | NA |
| Naga | ENSMUSG( | ENSMUSG( | 0.039711 | 0.792778 | NO | NO | NA |
| Qrsl1 | ENSMUSG( | ENSMUSG( | 0.277739 | 0.00654 | YES | NO | NA |
| 2410004B: | ENSMUSG( | ENSMUSG( | -0.10314 | 0.328456 | NO | NO | NA |
| Mmut | ENSMUSG( | ENSMUSG( | 0.081662 | 0.637281 | NO | NO | NA |
| Isca2 | ENSMUSG( | ENSMUSG( | 0.051107 | 0.539846 | NO | NO | NA |
| Lancl2 | ENSMUSG( | ENSMUSG( | 0.298 | 0.004487 | YES | NO | NA |
| Bloc1s4 | ENSMUSG( | ENSMUSG( | -0.08194 | 0.429242 | NO | NO | NA |
| Mrps6 | ENSMUSG( | ENSMUSG( | 0.508254 | 7.06E-09 | YES | NO | NA |
| Gm37401 | ENSMUSG( | ENSMUSG( | -0.06136 | 0.625461 | NO | NO | NA |
| Prss16 | ENSMUSG( | ENSMUSG( | 0.491738 | 0.015705 | YES | NO | NA |
| 2310001H | ENSMUSG( | ENSMUSG( | -0.33888 | 0.011125 | YES | NO | NA |
| Faap24 | ENSMUSG( | ENSMUSG( | 0.125784 | 0.128813 | NO | NO | NA |
| Gprasp1 | ENSMUSG( | ENSMUSG( | -0.44256 | 1.56E-05 | YES | NO | NA |
| Pmvk | ENSMUSG( | ENSMUSG( | 0.281842 | 0.009928 | YES | NO | NA |
| Txndc15 | ENSMUSG( | ENSMUSG( | 0.027483 | 0.722171 | NO | NO | NA |
| F2rl3 | ENSMUSG( | ENSMUSG( | 0.314451 | 0.000433 | YES | NO | NA |
| Atxn2 | ENSMUSG( | ENSMUSG( | -0.12883 | 0.196467 | NO | NO | NA |
| 5730455P: | ENSMUSG( | ENSMUSG( | 0.162937 | 0.087695 | NO | NO | NA |
| Tiam1 | ENSMUSG( | ENSMUSG( | 0.769726 | 1.94E-06 | YES | NO | NA |
| Nsun5 | ENSMUSG( | ENSMUSG( | 0.346703 | 0.000338 | YES | NO | NA |
| Fam102b | ENSMUSG( | ENSMUSG( | -0.46517 | 4.94E-07 | YES | NO | NA |
| G6pc3 | ENSMUSG( | ENSMUSG( | 0.342667 | 0.001996 | YES | NO | NA |
| Itprid2 | ENSMUSG( | ENSMUSG( | -0.12774 | 0.229669 | NO | NO | NA |
| Tmem87b | ENSMUSG( | ENSMUSG( | -0.34778 | 0.000322 | YES | NO | NA |
| Wdyhv1 | ENSMUSG( | ENSMUSG( | -0.54346 | 8.66E-17 | YES | NO | NA |
| Gvin-ps2 | ENSMUSG( | ENSMUSG( | -0.32931 | 0.066807 | NO | NO | NA |
| Lrrc41 | ENSMUSG( | ENSMUSG( | -0.14472 | 0.105284 | NO | NO | NA |
| Ten1 | ENSMUSG( | ENSMUSG( | 0.205148 | 0.009775 | YES | NO | NA |
| Serpinb9b | ENSMUSG( | ENSMUSG( | 1.023227 | 0.004956 | YES | YES | Upregulated |
| Gpn1 | ENSMUSG( | ENSMUSG( | 0.174657 | 0.021006 | YES | NO | NA |
| Tmem208 | ENSMUSG( | ENSMUSG( | 0.010663 | 0.917001 | NO | NO | NA |
| Lsm11 | ENSMUSG( | ENSMUSG( | 0.131013 | 0.202451 | NO | NO | NA |
| Tmem179l | ENSMUSG( | ENSMUSG( | -0.09801 | 0.340753 | NO | NO | NA |
| Brms1l | ENSMUSG( | ENSMUSG( | 0.121602 | 0.341395 | NO | NO | NA |
| Stx2 | ENSMUSG( | ENSMUSG( | -0.79057 | 1.10E-08 | YES | NO | NA |
| Trbv12-2 | ENSMUSG( | ENSMUSG( | -0.12889 | 0.799508 | NO | NO | NA |
| St6galnac6 | ENSMUSG( | ENSMUSG( | 0.90558 | 1.64E-12 | YES | NO | NA |
| Alkbh3 | ENSMUSG( | ENSMUSG( | 0.228797 | 0.04771 | YES | NO | NA |
| Hypk | ENSMUSG( | ENSMUSG( | 0.024381 | 0.816297 | NO | NO | NA |
| Prr3 | ENSMUSG( | ENSMUSG( | -0.00082 | 0.994565 | NO | NO | NA |
| Trp53rka | ENSMUSG( | ENSMUSG( | -0.10004 | 0.235139 | NO | NO | NA |
| Gm49602 | ENSMUSG( | ENSMUSG( | -0.31472 | 0.001483 | YES | NO | NA |
| Nlrx1 | ENSMUSG( | ENSMUSG( | 0.134367 | 0.336657 | NO | NO | NA |
| Fxn | ENSMUSG( | ENSMUSG( | 0.3202 | 6.38E-05 | YES | NO | NA |
| Ranbp6 | ENSMUSG( | ENSMUSG( | -0.24787 | 0.011933 | YES | NO | NA |

| | | | | | | | |
|---|---|---|---|---|---|---|---|
| Bbln | ENSMUSG( | ENSMUSG( | -0.16623 | 0.145762 | NO | NO | NA |
| Timm10b | ENSMUSG( | ENSMUSG( | 0.118671 | 0.393625 | NO | NO | NA |
| Mpc2 | ENSMUSG( | ENSMUSG( | 0.082043 | 0.373516 | NO | NO | NA |
| Grpel2 | ENSMUSG( | ENSMUSG( | -0.08927 | 0.304919 | NO | NO | NA |
| B3glct | ENSMUSG( | ENSMUSG( | -0.00932 | 0.92372 | NO | NO | NA |
| Gnpnat1 | ENSMUSG( | ENSMUSG( | 0.063424 | 0.46978 | NO | NO | NA |
| Katnbl1 | ENSMUSG( | ENSMUSG( | 0.05705 | 0.504719 | NO | NO | NA |
| Mul1 | ENSMUSG( | ENSMUSG( | -0.00403 | 0.970238 | NO | NO | NA |
| Maskbp3 | ENSMUSG( | ENSMUSG( | -0.25329 | 0.031122 | YES | NO | NA |
| Sclt1 | ENSMUSG( | ENSMUSG( | 0.144304 | 0.236426 | NO | NO | NA |
| Dennd11 | ENSMUSG( | ENSMUSG( | 0.190941 | 0.137859 | NO | NO | NA |
| Fam185a | ENSMUSG( | ENSMUSG( | 0.086298 | 0.350235 | NO | NO | NA |
| Rcn2 | ENSMUSG( | ENSMUSG( | 0.328303 | 0.000274 | YES | NO | NA |
| Agpat5 | ENSMUSG( | ENSMUSG( | 0.379682 | 0.00022 | YES | NO | NA |
| Gpr183 | ENSMUSG( | ENSMUSG( | 0.813412 | 0.000707 | YES | NO | NA |
| Lpp | ENSMUSG( | ENSMUSG( | -0.35359 | 0.003315 | YES | NO | NA |
| Dcaf17 | ENSMUSG( | ENSMUSG( | -0.19305 | 0.030902 | YES | NO | NA |
| Apbb3 | ENSMUSG( | ENSMUSG( | -0.08691 | 0.465633 | NO | NO | NA |
| ENSMUSG( | ENSMUSG( | ENSMUSG( | 0.266173 | 0.077007 | NO | NO | NA |
| Nipsnap1 | ENSMUSG( | ENSMUSG( | 0.11858 | 0.079237 | NO | NO | NA |
| N4bp3 | ENSMUSG( | ENSMUSG( | -0.44157 | 0.001981 | YES | NO | NA |
| Stx18 | ENSMUSG( | ENSMUSG( | 0.195713 | 0.096024 | NO | NO | NA |
| Gas2l3 | ENSMUSG( | ENSMUSG( | -0.11022 | 0.415514 | NO | NO | NA |
| Zfp526 | ENSMUSG( | ENSMUSG( | -0.51825 | 2.03E-08 | YES | NO | NA |
| Natd1 | ENSMUSG( | ENSMUSG( | -0.56219 | 2.64E-05 | YES | NO | NA |
| Cenpw | ENSMUSG( | ENSMUSG( | 0.328476 | 0.000811 | YES | NO | NA |
| Apex2 | ENSMUSG( | ENSMUSG( | -0.22216 | 0.010193 | YES | NO | NA |
| Tmem161l | ENSMUSG( | ENSMUSG( | -0.02956 | 0.79108 | NO | NO | NA |
| Dnaja4 | ENSMUSG( | ENSMUSG( | -0.51872 | 0.000101 | YES | NO | NA |
| Obi1 | ENSMUSG( | ENSMUSG( | -0.02471 | 0.796351 | NO | NO | NA |
| Tepsin | ENSMUSG( | ENSMUSG( | -0.48503 | 8.60E-09 | YES | NO | NA |
| BC003965 | ENSMUSG( | ENSMUSG( | -0.02915 | 0.770991 | NO | NO | NA |
| Ets2 | ENSMUSG( | ENSMUSG( | -0.32624 | 0.017322 | YES | NO | NA |
| Slc25a53 | ENSMUSG( | ENSMUSG( | -0.45674 | 1.05E-06 | YES | NO | NA |
| Phaf1 | ENSMUSG( | ENSMUSG( | -0.02898 | 0.790471 | NO | NO | NA |
| Hacd2 | ENSMUSG( | ENSMUSG( | -0.062 | 0.5876 | NO | NO | NA |
| Mpg | ENSMUSG( | ENSMUSG( | 0.293806 | 0.023841 | YES | NO | NA |
| Thap7 | ENSMUSG( | ENSMUSG( | 0.034238 | 0.675887 | NO | NO | NA |
| ENSMUSG | ENSMUSG( | ENSMUSG( | -0.2515 | 0.01307 | YES | NO | NA |
| Zfp369 | ENSMUSG( | ENSMUSG( | -0.51361 | 9.63E-06 | YES | NO | NA |
| Mgat4a | ENSMUSG( | ENSMUSG( | -0.13991 | 0.147915 | NO | NO | NA |
| Slc36a4 | ENSMUSG( | ENSMUSG( | 0.317861 | 0.010851 | YES | NO | NA |
| Acss2 | ENSMUSG( | ENSMUSG( | -0.41263 | 0.021967 | YES | NO | NA |
| Nup35 | ENSMUSG( | ENSMUSG( | -0.17171 | 0.049638 | YES | NO | NA |
| Faap100 | ENSMUSG( | ENSMUSG( | -0.06845 | 0.549983 | NO | NO | NA |
| Ciao3 | ENSMUSG( | ENSMUSG( | 0.069566 | 0.488115 | NO | NO | NA |
| Timm22 | ENSMUSG( | ENSMUSG( | 0.25603 | 0.001703 | YES | NO | NA |

| | | | | | | | |
|---|---|---|---|---|---|---|---|
| Gm44878 | ENSMUSG( | ENSMUSG( | -0.13478 | 0.442957 | NO | NO | NA |
| Rtl8b | ENSMUSG( | ENSMUSG( | 0.02083 | 0.851968 | NO | NO | NA |
| Tnfsf14 | ENSMUSG( | ENSMUSG( | -0.33488 | 0.009244 | YES | NO | NA |
| Zfp568 | ENSMUSG( | ENSMUSG( | 0.078649 | 0.388409 | NO | NO | NA |
| Cenps | ENSMUSG( | ENSMUSG( | 0.412293 | 0.002484 | YES | NO | NA |
| Svbp | ENSMUSG( | ENSMUSG( | -0.01055 | 0.930482 | NO | NO | NA |
| Lims2 | ENSMUSG( | ENSMUSG( | 0.55069 | 0.006418 | YES | NO | NA |
| Il2ra | ENSMUSG( | ENSMUSG( | 2.592388 | 8.23E-16 | YES | YES | Upregulated |
| Smim3 | ENSMUSG( | ENSMUSG( | 0.388506 | 0.000683 | YES | NO | NA |
| Mterf3 | ENSMUSG( | ENSMUSG( | -0.2807 | 9.61E-06 | YES | NO | NA |
| Wdfy2 | ENSMUSG( | ENSMUSG( | 0.247669 | 0.042059 | YES | NO | NA |
| Exosc4 | ENSMUSG( | ENSMUSG( | 0.032804 | 0.723821 | NO | NO | NA |
| E530011L2 | ENSMUSG( | ENSMUSG( | -0.09358 | 0.466218 | NO | NO | NA |
| Zfp703 | ENSMUSG( | ENSMUSG( | -0.09602 | 0.309567 | NO | NO | NA |
| Tamm41 | ENSMUSG( | ENSMUSG( | 0.558007 | 2.61E-06 | YES | NO | NA |
| Phospho2 | ENSMUSG( | ENSMUSG( | 0.017153 | 0.853406 | NO | NO | NA |
| Atpaf1 | ENSMUSG( | ENSMUSG( | 0.018378 | 0.846519 | NO | NO | NA |
| Mex3b | ENSMUSG( | ENSMUSG( | -0.27999 | 0.058546 | NO | NO | NA |
| Orc1 | ENSMUSG( | ENSMUSG( | 0.400547 | 1.68E-05 | YES | NO | NA |
| Hmgcl | ENSMUSG( | ENSMUSG( | 0.139156 | 0.065066 | NO | NO | NA |
| Gstt2 | ENSMUSG( | ENSMUSG( | 0.51116 | 0.000327 | YES | NO | NA |
| Slc15a4 | ENSMUSG( | ENSMUSG( | -0.32892 | 0.00012 | YES | NO | NA |
| AI597479 | ENSMUSG( | ENSMUSG( | -0.19435 | 0.026845 | YES | NO | NA |
| Agtrap | ENSMUSG( | ENSMUSG( | -0.44226 | 2.05E-06 | YES | NO | NA |
| Trbv13-3 | ENSMUSG( | ENSMUSG( | -0.35629 | 0.247168 | NO | NO | NA |
| Tada2a | ENSMUSG( | ENSMUSG( | 0.424961 | 0.000309 | YES | NO | NA |
| Fancf | ENSMUSG( | ENSMUSG( | 0.195928 | 0.034676 | YES | NO | NA |
| Tarsl2 | ENSMUSG( | ENSMUSG( | 0.487322 | 6.78E-05 | YES | NO | NA |
| Enpp4 | ENSMUSG( | ENSMUSG( | -0.5037 | 5.90E-05 | YES | NO | NA |
| Uevld | ENSMUSG( | ENSMUSG( | 0.021783 | 0.773278 | NO | NO | NA |
| Il1rl1 | ENSMUSG( | ENSMUSG( | 2.039087 | 6.16E-18 | YES | YES | Upregulated |
| Triap1 | ENSMUSG( | ENSMUSG( | 0.034716 | 0.751196 | NO | NO | NA |
| ENSMUSG( | ENSMUSG( | ENSMUSG( | -0.07911 | 0.49287 | NO | NO | NA |
| Itgad | ENSMUSG( | ENSMUSG( | -1.77463 | 8.08E-08 | YES | YES | Downregulated |
| Cyb5r1 | ENSMUSG( | ENSMUSG( | 0.177256 | 0.069438 | NO | NO | NA |
| Gdpd5 | ENSMUSG( | ENSMUSG( | 0.610877 | 6.76E-05 | YES | NO | NA |
| Wdr45 | ENSMUSG( | ENSMUSG( | -0.19914 | 0.049393 | YES | NO | NA |
| Zfp956 | ENSMUSG( | ENSMUSG( | 0.096869 | 0.484233 | NO | NO | NA |
| Enkd1 | ENSMUSG( | ENSMUSG( | 0.558799 | 4.18E-08 | YES | NO | NA |
| Polr2h | ENSMUSG( | ENSMUSG( | 0.475352 | 1.59E-05 | YES | NO | NA |
| Polr2k | ENSMUSG( | ENSMUSG( | 0.101194 | 0.216296 | NO | NO | NA |
| Trappc6a | ENSMUSG( | ENSMUSG( | 0.267798 | 0.065334 | NO | NO | NA |
| Ublcp1 | ENSMUSG( | ENSMUSG( | 0.251566 | 0.094848 | NO | NO | NA |
| Slc20a2 | ENSMUSG( | ENSMUSG( | -0.00588 | 0.963858 | NO | NO | NA |
| Grn | ENSMUSG( | ENSMUSG( | -0.14521 | 0.264275 | NO | NO | NA |
| Nif3l1 | ENSMUSG( | ENSMUSG( | 0.283343 | 0.003889 | YES | NO | NA |
| Bsn | ENSMUSG( | ENSMUSG( | -0.28813 | 0.007645 | YES | NO | NA |

| | | | | | | | |
|---|---|---|---|---|---|---|---|
| Ighg2b | ENSMUSG( | ENSMUSG( | -3.43293 | 0.000276 | YES | YES | Downregulated |
| Ufsp2 | ENSMUSG( | ENSMUSG( | 0.126314 | 0.277353 | NO | NO | NA |
| Bnip1 | ENSMUSG( | ENSMUSG( | 0.118283 | 0.287275 | NO | NO | NA |
| Ppp1r12b | ENSMUSG( | ENSMUSG( | -0.16964 | 0.041864 | YES | NO | NA |
| Haghl | ENSMUSG( | ENSMUSG( | 0.447252 | 0.000484 | YES | NO | NA |
| Fance | ENSMUSG( | ENSMUSG( | 0.05964 | 0.57351 | NO | NO | NA |
| Stx8 | ENSMUSG( | ENSMUSG( | -0.15592 | 0.13474 | NO | NO | NA |
| Ptrh2 | ENSMUSG( | ENSMUSG( | 0.255144 | 0.000511 | YES | NO | NA |
| Ppil3 | ENSMUSG( | ENSMUSG( | 0.03357 | 0.710704 | NO | NO | NA |
| Phpt1 | ENSMUSG( | ENSMUSG( | 0.659602 | 7.25E-06 | YES | NO | NA |
| Gm14029 | ENSMUSG( | ENSMUSG( | -0.90911 | 4.67E-09 | YES | NO | NA |
| Zkscan8 | ENSMUSG( | ENSMUSG( | 0.146035 | 0.336818 | NO | NO | NA |
| Zfp219 | ENSMUSG( | ENSMUSG( | 0.405794 | 0.000622 | YES | NO | NA |
| Ngrn | ENSMUSG( | ENSMUSG( | 0.079842 | 0.470703 | NO | NO | NA |
| Galt | ENSMUSG( | ENSMUSG( | 0.003115 | 0.982598 | NO | NO | NA |
| Ccnq | ENSMUSG( | ENSMUSG( | 0.139971 | 0.187728 | NO | NO | NA |
| Eri3 | ENSMUSG( | ENSMUSG( | 0.269663 | 0.029984 | YES | NO | NA |
| Btla | ENSMUSG( | ENSMUSG( | -0.17658 | 0.229121 | NO | NO | NA |
| Anxa4 | ENSMUSG( | ENSMUSG( | -0.18846 | 0.103888 | NO | NO | NA |
| Gm33104 | ENSMUSG( | ENSMUSG( | 0.206243 | 0.141155 | NO | NO | NA |
| Coasy | ENSMUSG( | ENSMUSG( | 0.207825 | 0.065031 | NO | NO | NA |
| Rpp14 | ENSMUSG( | ENSMUSG( | 0.143766 | 0.040251 | YES | NO | NA |
| Ccdc85b | ENSMUSG( | ENSMUSG( | -0.07436 | 0.534082 | NO | NO | NA |
| Golga1 | ENSMUSG( | ENSMUSG( | -0.15866 | 0.22187 | NO | NO | NA |
| Polb | ENSMUSG( | ENSMUSG( | -0.20763 | 0.006807 | YES | NO | NA |
| Mier3 | ENSMUSG( | ENSMUSG( | -0.27334 | 0.002088 | YES | NO | NA |
| Tgif1 | ENSMUSG( | ENSMUSG( | 0.101435 | 0.584439 | NO | NO | NA |
| Naf1 | ENSMUSG( | ENSMUSG( | 0.649557 | 1.87E-08 | YES | NO | NA |
| Itga1 | ENSMUSG( | ENSMUSG( | 0.06072 | 0.840237 | NO | NO | NA |
| Fbxl18 | ENSMUSG( | ENSMUSG( | -0.14218 | 0.166103 | NO | NO | NA |
| Dip2a | ENSMUSG( | ENSMUSG( | -0.4165 | 1.05E-06 | YES | NO | NA |
| Mosmo | ENSMUSG( | ENSMUSG( | -0.28317 | 0.00149 | YES | NO | NA |
| Tmem129 | ENSMUSG( | ENSMUSG( | -0.25653 | 0.001051 | YES | NO | NA |
| Zfp707 | ENSMUSG( | ENSMUSG( | -0.29191 | 0.000332 | YES | NO | NA |
| 2310057M | ENSMUSG( | ENSMUSG( | -0.03493 | 0.752107 | NO | NO | NA |
| 2510002D | ENSMUSG( | ENSMUSG( | -0.08368 | 0.272668 | NO | NO | NA |
| Insig2 | ENSMUSG( | ENSMUSG( | -0.12527 | 0.390646 | NO | NO | NA |
| Slc52a2 | ENSMUSG( | ENSMUSG( | 0.075565 | 0.461006 | NO | NO | NA |
| Usp49 | ENSMUSG( | ENSMUSG( | 0.029645 | 0.832038 | NO | NO | NA |
| Pop7 | ENSMUSG( | ENSMUSG( | 0.098105 | 0.474071 | NO | NO | NA |
| Galnt4 | ENSMUSG( | ENSMUSG( | -0.04772 | 0.729544 | NO | NO | NA |
| Phyh | ENSMUSG( | ENSMUSG( | -0.2346 | 0.028209 | YES | NO | NA |
| Med19 | ENSMUSG( | ENSMUSG( | 0.015308 | 0.888811 | NO | NO | NA |
| Crim1 | ENSMUSG( | ENSMUSG( | -0.66836 | 0.002758 | YES | NO | NA |
| Fam110a | ENSMUSG( | ENSMUSG( | 0.01917 | 0.874667 | NO | NO | NA |
| Lage3 | ENSMUSG( | ENSMUSG( | 0.15742 | 0.04695 | YES | NO | NA |
| Mgat5 | ENSMUSG( | ENSMUSG( | -0.30941 | 0.001115 | YES | NO | NA |

| | | | | | | | |
|---|---|---|---|---|---|---|---|
| Dnajc18 | ENSMUSG( | ENSMUSG( | -0.32788 | 0.000102 | YES | NO | NA |
| Caap1 | ENSMUSG( | ENSMUSG( | -0.04551 | 0.729948 | NO | NO | NA |
| Ovca2 | ENSMUSG( | ENSMUSG( | -0.1426 | 0.203986 | NO | NO | NA |
| Isg20 | ENSMUSG( | ENSMUSG( | -0.31288 | 0.015693 | YES | NO | NA |
| Ppp1r16a | ENSMUSG( | ENSMUSG( | 0.331042 | 0.00856 | YES | NO | NA |
| Rn7s1 | ENSMUSG( | ENSMUSG( | 0.423545 | 0.846207 | NO | NO | NA |
| Slirp | ENSMUSG( | ENSMUSG( | 0.288095 | 0.009757 | YES | NO | NA |
| Rab3d | ENSMUSG( | ENSMUSG( | -0.00432 | 0.978798 | NO | NO | NA |
| Coq10b | ENSMUSG( | ENSMUSG( | 0.054878 | 0.724666 | NO | NO | NA |
| Foxred2 | ENSMUSG( | ENSMUSG( | 0.743524 | 1.08E-06 | YES | NO | NA |
| Tubb3 | ENSMUSG( | ENSMUSG( | 0.749571 | 2.92E-06 | YES | NO | NA |
| Tec | ENSMUSG( | ENSMUSG( | -0.49372 | 3.24E-06 | YES | NO | NA |
| Pepd | ENSMUSG( | ENSMUSG( | 0.00741 | 0.944823 | NO | NO | NA |
| Slc35d2 | ENSMUSG( | ENSMUSG( | -0.04003 | 0.812809 | NO | NO | NA |
| Nsdhl | ENSMUSG( | ENSMUSG( | -0.10578 | 0.405994 | NO | NO | NA |
| 1600012H | ENSMUSG( | ENSMUSG( | -0.1922 | 0.005428 | YES | NO | NA |
| Ccdc86 | ENSMUSG( | ENSMUSG( | 0.378324 | 0.002634 | YES | NO | NA |
| Fam210a | ENSMUSG( | ENSMUSG( | -0.1089 | 0.25195 | NO | NO | NA |
| AW01173 | ENSMUSG( | ENSMUSG( | -1.29738 | 3.19E-09 | YES | YES | Downregulated |
| Cpped1 | ENSMUSG( | ENSMUSG( | -0.03853 | 0.773862 | NO | NO | NA |
| Tgds | ENSMUSG( | ENSMUSG( | 0.023097 | 0.840237 | NO | NO | NA |
| Snhg15 | ENSMUSG( | ENSMUSG( | 0.531651 | 0.001038 | YES | NO | NA |
| Ppm1k | ENSMUSG( | ENSMUSG( | -0.26785 | 0.004417 | YES | NO | NA |
| Ppa2 | ENSMUSG( | ENSMUSG( | 0.211739 | 0.013107 | YES | NO | NA |
| Xpc | ENSMUSG( | ENSMUSG( | -0.15491 | 0.309961 | NO | NO | NA |
| Tmem250 | ENSMUSG( | ENSMUSG( | 0.307202 | 0.005542 | YES | NO | NA |
| Nsun3 | ENSMUSG( | ENSMUSG( | 0.218278 | 0.049241 | YES | NO | NA |
| Bri3 | ENSMUSG( | ENSMUSG( | 0.216208 | 0.034255 | YES | NO | NA |
| Pxmp4 | ENSMUSG( | ENSMUSG( | -0.11373 | 0.421379 | NO | NO | NA |
| Akip1 | ENSMUSG( | ENSMUSG( | 0.197465 | 0.023512 | YES | NO | NA |
| Rcc1l | ENSMUSG( | ENSMUSG( | 0.898761 | 9.41E-09 | YES | NO | NA |
| Klk8 | ENSMUSG( | ENSMUSG( | -0.17233 | 0.092443 | NO | NO | NA |
| Echdc1 | ENSMUSG( | ENSMUSG( | -0.06756 | 0.625598 | NO | NO | NA |
| Slc39a3 | ENSMUSG( | ENSMUSG( | -0.08733 | 0.369399 | NO | NO | NA |
| Jkamp | ENSMUSG( | ENSMUSG( | -0.22229 | 0.024338 | YES | NO | NA |
| Xrcc3 | ENSMUSG( | ENSMUSG( | 0.176027 | 0.202809 | NO | NO | NA |
| Gm19765 | ENSMUSG( | ENSMUSG( | -0.7748 | 0.000355 | YES | NO | NA |
| 1110038B | ENSMUSG( | ENSMUSG( | 1.11034 | 3.29E-09 | YES | YES | Upregulated |
| Clpp | ENSMUSG( | ENSMUSG( | 0.365471 | 5.96E-05 | YES | NO | NA |
| Aar2 | ENSMUSG( | ENSMUSG( | -0.04716 | 0.71535 | NO | NO | NA |
| Aplp1 | ENSMUSG( | ENSMUSG( | -0.88822 | 1.54E-08 | YES | NO | NA |
| Slc39a4 | ENSMUSG( | ENSMUSG( | 0.171943 | 0.029316 | YES | NO | NA |
| Stam | ENSMUSG( | ENSMUSG( | -0.00186 | 0.987807 | NO | NO | NA |
| Zfp51 | ENSMUSG( | ENSMUSG( | -0.19989 | 0.10893 | NO | NO | NA |
| Ankrd39 | ENSMUSG( | ENSMUSG( | 0.131275 | 0.060885 | NO | NO | NA |
| Plin2 | ENSMUSG( | ENSMUSG( | -0.65234 | 1.85E-07 | YES | NO | NA |
| Tti2 | ENSMUSG( | ENSMUSG( | -0.03755 | 0.626685 | NO | NO | NA |

| | | | | | | | |
|---|---|---|---|---|---|---|---|
| Oxsm | ENSMUSG( | ENSMUSG( | -0.26688 | 0.005004 | YES | NO | NA |
| Mcat | ENSMUSG( | ENSMUSG( | 0.283575 | 0.004786 | YES | NO | NA |
| Prdx4 | ENSMUSG( | ENSMUSG( | 1.2691 | 5.08E-08 | YES | YES | Upregulated |
| Cox16 | ENSMUSG( | ENSMUSG( | 0.020883 | 0.797995 | NO | NO | NA |
| Alg8 | ENSMUSG( | ENSMUSG( | 0.142188 | 0.262914 | NO | NO | NA |
| Slc35a2 | ENSMUSG( | ENSMUSG( | -0.29596 | 7.06E-06 | YES | NO | NA |
| Zfp868 | ENSMUSG( | ENSMUSG( | -0.13169 | 0.296203 | NO | NO | NA |
| Ccdc22 | ENSMUSG( | ENSMUSG( | -0.14418 | 0.194576 | NO | NO | NA |
| Bckdha | ENSMUSG( | ENSMUSG( | 0.174717 | 0.293315 | NO | NO | NA |
| Zfp287 | ENSMUSG( | ENSMUSG( | -0.16757 | 0.087098 | NO | NO | NA |
| Xpa | ENSMUSG( | ENSMUSG( | 0.004289 | 0.973542 | NO | NO | NA |
| Itm2a | ENSMUSG( | ENSMUSG( | -0.08397 | 0.643741 | NO | NO | NA |
| Zfp945 | ENSMUSG( | ENSMUSG( | -0.22889 | 0.187394 | NO | NO | NA |
| Dnajc16 | ENSMUSG( | ENSMUSG( | -0.13061 | 0.30358 | NO | NO | NA |
| Tvp23b | ENSMUSG( | ENSMUSG( | -0.20027 | 0.022839 | YES | NO | NA |
| Vcl | ENSMUSG( | ENSMUSG( | 0.510916 | 0.002128 | YES | NO | NA |
| Mecr | ENSMUSG( | ENSMUSG( | 0.909484 | 4.90E-08 | YES | NO | NA |
| Zfp382 | ENSMUSG( | ENSMUSG( | -0.34978 | 0.001212 | YES | NO | NA |
| Guk1 | ENSMUSG( | ENSMUSG( | 0.286685 | 0.004673 | YES | NO | NA |
| Tmprss13 | ENSMUSG( | ENSMUSG( | -1.30865 | 1.43E-07 | YES | YES | Downregulated |
| Kat8 | ENSMUSG( | ENSMUSG( | -0.04647 | 0.650518 | NO | NO | NA |
| Borcs6 | ENSMUSG( | ENSMUSG( | -0.17304 | 0.067929 | NO | NO | NA |
| Enoph1 | ENSMUSG( | ENSMUSG( | 0.364164 | 5.82E-05 | YES | NO | NA |
| Nccrp1 | ENSMUSG( | ENSMUSG( | -0.60504 | 0.00086 | YES | NO | NA |
| Gm30054 | ENSMUSG( | ENSMUSG( | -0.80487 | 1.13E-06 | YES | NO | NA |
| Pus3 | ENSMUSG( | ENSMUSG( | -0.19373 | 0.097159 | NO | NO | NA |
| Ndufaf7 | ENSMUSG( | ENSMUSG( | 0.115816 | 0.302365 | NO | NO | NA |
| Ddt | ENSMUSG( | ENSMUSG( | 0.296629 | 0.033916 | YES | NO | NA |
| Rab28 | ENSMUSG( | ENSMUSG( | 0.155669 | 0.267015 | NO | NO | NA |
| Hsd17b7 | ENSMUSG( | ENSMUSG( | 0.625008 | 6.49E-05 | YES | NO | NA |
| Cdr2 | ENSMUSG( | ENSMUSG( | 0.00547 | 0.978405 | NO | NO | NA |
| Parp12 | ENSMUSG( | ENSMUSG( | -0.86715 | 1.38E-07 | YES | NO | NA |
| Snhg20 | ENSMUSG( | ENSMUSG( | 0.07964 | 0.512304 | NO | NO | NA |
| Cd101 | ENSMUSG( | ENSMUSG( | -0.86212 | 3.92E-05 | YES | NO | NA |
| Golm2 | ENSMUSG( | ENSMUSG( | 0.501201 | 8.55E-06 | YES | NO | NA |
| Adrb1 | ENSMUSG( | ENSMUSG( | -0.52067 | 6.79E-05 | YES | NO | NA |
| 5430416N | ENSMUSG( | ENSMUSG( | 0.180878 | 0.382542 | NO | NO | NA |
| Zfp787 | ENSMUSG( | ENSMUSG( | -0.11602 | 0.302496 | NO | NO | NA |
| Fbxw17 | ENSMUSG( | ENSMUSG( | 0.078089 | 0.645945 | NO | NO | NA |
| Snhg6 | ENSMUSG( | ENSMUSG( | 0.255733 | 0.005443 | YES | NO | NA |
| Hdhd2 | ENSMUSG( | ENSMUSG( | 0.01125 | 0.921939 | NO | NO | NA |
| Tnf | ENSMUSG( | ENSMUSG( | 1.340866 | 9.91E-05 | YES | YES | Upregulated |
| 4833438C( | ENSMUSG( | ENSMUSG( | 0.224667 | 0.18641 | NO | NO | NA |
| Nudt1 | ENSMUSG( | ENSMUSG( | 0.261972 | 0.042165 | YES | NO | NA |
| Sgms1 | ENSMUSG( | ENSMUSG( | -0.07515 | 0.698535 | NO | NO | NA |
| Spata6 | ENSMUSG( | ENSMUSG( | 0.379782 | 0.02876 | YES | NO | NA |
| Zfp213 | ENSMUSG( | ENSMUSG( | 0.210498 | 0.094428 | NO | NO | NA |

| | | | | | | | |
|---|---|---|---|---|---|---|---|
| Osgin2 | ENSMUSG( | ENSMUSG( | -0.01133 | 0.92009 | NO | NO | NA |
| Hddc2 | ENSMUSG( | ENSMUSG( | 0.858221 | 3.07E-07 | YES | NO | NA |
| Mvd | ENSMUSG( | ENSMUSG( | -0.17159 | 0.111142 | NO | NO | NA |
| Spidr | ENSMUSG( | ENSMUSG( | -0.02044 | 0.897193 | NO | NO | NA |
| Tpgs2 | ENSMUSG( | ENSMUSG( | 0.022214 | 0.874407 | NO | NO | NA |
| Zdhhc13 | ENSMUSG( | ENSMUSG( | -0.01353 | 0.86099 | NO | NO | NA |
| D6Wsu163 | ENSMUSG( | ENSMUSG( | -0.0673 | 0.418988 | NO | NO | NA |
| Fam8a1 | ENSMUSG( | ENSMUSG( | -0.38776 | 0.001177 | YES | NO | NA |
| Gtf3c3 | ENSMUSG( | ENSMUSG( | 0.027362 | 0.832892 | NO | NO | NA |
| Trp53i13 | ENSMUSG( | ENSMUSG( | -0.09319 | 0.277974 | NO | NO | NA |
| Zfp729a | ENSMUSG( | ENSMUSG( | -0.31696 | 0.004348 | YES | NO | NA |
| Spout1 | ENSMUSG( | ENSMUSG( | 0.190032 | 0.028701 | YES | NO | NA |
| Traf3ip1 | ENSMUSG( | ENSMUSG( | -0.51474 | 4.36E-05 | YES | NO | NA |
| Timmdc1 | ENSMUSG( | ENSMUSG( | 0.064217 | 0.590346 | NO | NO | NA |
| Fundc1 | ENSMUSG( | ENSMUSG( | -0.06023 | 0.557764 | NO | NO | NA |
| Casp9 | ENSMUSG( | ENSMUSG( | -0.22142 | 0.018095 | YES | NO | NA |
| Timm8b | ENSMUSG( | ENSMUSG( | 0.356102 | 0.000151 | YES | NO | NA |
| Pde4a | ENSMUSG( | ENSMUSG( | -0.1574 | 0.224475 | NO | NO | NA |
| Dolpp1 | ENSMUSG( | ENSMUSG( | -0.05416 | 0.518349 | NO | NO | NA |
| Galnt11 | ENSMUSG( | ENSMUSG( | -0.25205 | 0.007988 | YES | NO | NA |
| Zkscan1 | ENSMUSG( | ENSMUSG( | -0.25113 | 0.00346 | YES | NO | NA |
| Rcor3 | ENSMUSG( | ENSMUSG( | 0.063735 | 0.466856 | NO | NO | NA |
| Trgv2 | ENSMUSG( | ENSMUSG( | -0.34634 | 0.231542 | NO | NO | NA |
| Mettl26 | ENSMUSG( | ENSMUSG( | 0.216899 | 0.043563 | YES | NO | NA |
| Zfp444 | ENSMUSG( | ENSMUSG( | -0.10773 | 0.125956 | NO | NO | NA |
| Pex16 | ENSMUSG( | ENSMUSG( | 0.063411 | 0.544336 | NO | NO | NA |
| Supt7l | ENSMUSG( | ENSMUSG( | -0.00822 | 0.92158 | NO | NO | NA |
| Synj2 | ENSMUSG( | ENSMUSG( | 0.144634 | 0.391549 | NO | NO | NA |
| Mthfsl | ENSMUSG( | ENSMUSG( | -0.0279 | 0.844265 | NO | NO | NA |
| Paox | ENSMUSG( | ENSMUSG( | 0.933069 | 6.58E-07 | YES | NO | NA |
| Mapk8 | ENSMUSG( | ENSMUSG( | 0.436459 | 9.18E-08 | YES | NO | NA |
| Arl5b | ENSMUSG( | ENSMUSG( | -0.24265 | 0.0032 | YES | NO | NA |
| Rars2 | ENSMUSG( | ENSMUSG( | 0.137703 | 0.232063 | NO | NO | NA |
| Rpp21 | ENSMUSG( | ENSMUSG( | -0.04155 | 0.658025 | NO | NO | NA |
| Klhl42 | ENSMUSG( | ENSMUSG( | -0.37554 | 2.01E-05 | YES | NO | NA |
| Card19 | ENSMUSG( | ENSMUSG( | -0.05251 | 0.661866 | NO | NO | NA |
| Nbeal1 | ENSMUSG( | ENSMUSG( | -0.32163 | 0.006104 | YES | NO | NA |
| Cnih4 | ENSMUSG( | ENSMUSG( | 0.261626 | 0.008069 | YES | NO | NA |
| Gpam | ENSMUSG( | ENSMUSG( | 0.373859 | 7.38E-05 | YES | NO | NA |
| Med27 | ENSMUSG( | ENSMUSG( | 0.059546 | 0.581864 | NO | NO | NA |
| Asf1a | ENSMUSG( | ENSMUSG( | 0.283621 | 0.026635 | YES | NO | NA |
| Rgs8 | ENSMUSG( | ENSMUSG( | 1.15639 | 3.19E-05 | YES | YES | Upregulated |
| Vps45 | ENSMUSG( | ENSMUSG( | -0.07297 | 0.616427 | NO | NO | NA |
| Rsph3b | ENSMUSG( | ENSMUSG( | 0.327079 | 0.004262 | YES | NO | NA |
| Dcaf6 | ENSMUSG( | ENSMUSG( | -0.01585 | 0.901248 | NO | NO | NA |
| Mcoln2 | ENSMUSG( | ENSMUSG( | 0.424393 | 0.000106 | YES | NO | NA |
| Cdc14b | ENSMUSG( | ENSMUSG( | -0.58868 | 4.36E-08 | YES | NO | NA |

| | | | | | | | |
|---|---|---|---|---|---|---|---|
| Mtif3 | ENSMUSG( | ENSMUSG( | -0.07424 | 0.58248 | NO | NO | NA |
| Tmem65 | ENSMUSG( | ENSMUSG( | -0.08223 | 0.496489 | NO | NO | NA |
| Nudt18 | ENSMUSG( | ENSMUSG( | -0.39688 | 0.000633 | YES | NO | NA |
| Supv3l1 | ENSMUSG( | ENSMUSG( | 0.301301 | 0.00011 | YES | NO | NA |
| Haus7 | ENSMUSG( | ENSMUSG( | 0.37348 | 0.00047 | YES | NO | NA |
| Oma1 | ENSMUSG( | ENSMUSG( | -0.13976 | 0.203595 | NO | NO | NA |
| 9330175E: | ENSMUSG( | ENSMUSG( | -0.15968 | 0.396285 | NO | NO | NA |
| Tbcc | ENSMUSG( | ENSMUSG( | -0.07346 | 0.480215 | NO | NO | NA |
| Ptprk | ENSMUSG( | ENSMUSG( | 1.304079 | 1.15E-08 | YES | YES | Upregulated |
| Tom1l2 | ENSMUSG( | ENSMUSG( | -0.01083 | 0.950071 | NO | NO | NA |
| Fra10ac1 | ENSMUSG( | ENSMUSG( | 0.142322 | 0.219559 | NO | NO | NA |
| Polr1h | ENSMUSG( | ENSMUSG( | -0.00276 | 0.976454 | NO | NO | NA |
| Mtx2 | ENSMUSG( | ENSMUSG( | 0.418959 | 3.68E-06 | YES | NO | NA |
| Cluap1 | ENSMUSG( | ENSMUSG( | 0.089291 | 0.349117 | NO | NO | NA |
| Poli | ENSMUSG( | ENSMUSG( | -0.01941 | 0.867801 | NO | NO | NA |
| Plekhm3 | ENSMUSG( | ENSMUSG( | -0.29584 | 0.000464 | YES | NO | NA |
| Slc9a6 | ENSMUSG( | ENSMUSG( | -0.02714 | 0.840374 | NO | NO | NA |
| Mfge8 | ENSMUSG( | ENSMUSG( | -0.06174 | 0.534652 | NO | NO | NA |
| Abhd5 | ENSMUSG( | ENSMUSG( | -0.14172 | 0.079036 | NO | NO | NA |
| Gm49077 | ENSMUSG( | ENSMUSG( | -0.1452 | 0.472886 | NO | NO | NA |
| Pex1 | ENSMUSG( | ENSMUSG( | 0.197671 | 0.158522 | NO | NO | NA |
| Pck2 | ENSMUSG( | ENSMUSG( | -0.11278 | 0.36325 | NO | NO | NA |
| 2610008E: | ENSMUSG( | ENSMUSG( | -0.30745 | 0.059554 | NO | NO | NA |
| Wdr47 | ENSMUSG( | ENSMUSG( | -0.1433 | 0.220243 | NO | NO | NA |
| Uqcc1 | ENSMUSG( | ENSMUSG( | 0.077511 | 0.352248 | NO | NO | NA |
| Cenpv | ENSMUSG( | ENSMUSG( | 0.403232 | 0.005317 | YES | NO | NA |
| Rab12 | ENSMUSG( | ENSMUSG( | -0.20109 | 0.035937 | YES | NO | NA |
| Hps6 | ENSMUSG( | ENSMUSG( | -0.36629 | 1.18E-06 | YES | NO | NA |
| Ssx2ip | ENSMUSG( | ENSMUSG( | 0.394448 | 0.003567 | YES | NO | NA |
| Rit1 | ENSMUSG( | ENSMUSG( | -0.16367 | 0.151526 | NO | NO | NA |
| Rbm12b1 | ENSMUSG( | ENSMUSG( | -0.55425 | 1.99E-07 | YES | NO | NA |
| Iah1 | ENSMUSG( | ENSMUSG( | -0.10928 | 0.263012 | NO | NO | NA |
| Tmem64 | ENSMUSG( | ENSMUSG( | 0.260389 | 0.008037 | YES | NO | NA |
| Eef1aknmt | ENSMUSG( | ENSMUSG( | 0.599965 | 6.73E-07 | YES | NO | NA |
| Ndufc1 | ENSMUSG( | ENSMUSG( | 0.288178 | 0.002754 | YES | NO | NA |
| Ccs | ENSMUSG( | ENSMUSG( | 0.134343 | 0.222474 | NO | NO | NA |
| Btbd6 | ENSMUSG( | ENSMUSG( | -0.14758 | 0.149643 | NO | NO | NA |
| Clcn5 | ENSMUSG( | ENSMUSG( | -0.0128 | 0.903228 | NO | NO | NA |
| Apba3 | ENSMUSG( | ENSMUSG( | -0.1162 | 0.219273 | NO | NO | NA |
| Tmem186 | ENSMUSG( | ENSMUSG( | 0.207268 | 0.010743 | YES | NO | NA |
| Itga2 | ENSMUSG( | ENSMUSG( | -1.03533 | 1.04E-07 | YES | YES | Downregulated |
| Zfp950 | ENSMUSG( | ENSMUSG( | -0.31443 | 0.030122 | YES | NO | NA |
| Josd2 | ENSMUSG( | ENSMUSG( | -0.34236 | 2.90E-05 | YES | NO | NA |
| Pik3ip1 | ENSMUSG( | ENSMUSG( | 0.495914 | 0.02409 | YES | NO | NA |
| Rnf141 | ENSMUSG( | ENSMUSG( | -0.3696 | 0.000276 | YES | NO | NA |
| Dusp7 | ENSMUSG( | ENSMUSG( | -0.25546 | 0.056569 | NO | NO | NA |
| Ntng2 | ENSMUSG( | ENSMUSG( | -0.17162 | 0.226904 | NO | NO | NA |

| | | | | | | | |
|---|---|---|---|---|---|---|---|
| Samd10 | ENSMUSG( | ENSMUSG( | 0.328959 | 0.008791 | YES | NO | NA |
| 2610021A( | ENSMUSG( | ENSMUSG( | -0.17498 | 0.216926 | NO | NO | NA |
| Wdr73 | ENSMUSG( | ENSMUSG( | 0.237198 | 0.010219 | YES | NO | NA |
| Rprd1a | ENSMUSG( | ENSMUSG( | -0.13415 | 0.297078 | NO | NO | NA |
| Slc9a9 | ENSMUSG( | ENSMUSG( | -0.47974 | 0.000146 | YES | NO | NA |
| Mettl2 | ENSMUSG( | ENSMUSG( | 0.186305 | 0.215985 | NO | NO | NA |
| Akt3 | ENSMUSG( | ENSMUSG( | -0.35788 | 0.001413 | YES | NO | NA |
| Pick1 | ENSMUSG( | ENSMUSG( | 0.109427 | 0.360749 | NO | NO | NA |
| Ephx1 | ENSMUSG( | ENSMUSG( | -0.44024 | 0.003998 | YES | NO | NA |
| Gtf2ird1 | ENSMUSG( | ENSMUSG( | 0.947429 | 5.47E-07 | YES | NO | NA |
| Hgh1 | ENSMUSG( | ENSMUSG( | 0.415741 | 2.06E-09 | YES | NO | NA |
| Palb2 | ENSMUSG( | ENSMUSG( | 0.17502 | 0.052915 | NO | NO | NA |
| Sptb | ENSMUSG( | ENSMUSG( | 0.983663 | 0.017674 | YES | NO | NA |
| Cox14 | ENSMUSG( | ENSMUSG( | 0.040219 | 0.636129 | NO | NO | NA |
| Xrcc2 | ENSMUSG( | ENSMUSG( | 0.196575 | 0.022772 | YES | NO | NA |
| Rpp25l | ENSMUSG( | ENSMUSG( | -0.16994 | 0.055969 | NO | NO | NA |
| Med26 | ENSMUSG( | ENSMUSG( | -0.16887 | 0.129145 | NO | NO | NA |
| Slc25a1 | ENSMUSG( | ENSMUSG( | 0.239526 | 0.026194 | YES | NO | NA |
| Gcsh | ENSMUSG( | ENSMUSG( | 0.806156 | 1.06E-06 | YES | NO | NA |
| Blzf1 | ENSMUSG( | ENSMUSG( | 0.139395 | 0.222453 | NO | NO | NA |
| Oas1b | ENSMUSG( | ENSMUSG( | -0.72213 | 3.25E-10 | YES | NO | NA |
| Utp23 | ENSMUSG( | ENSMUSG( | -0.34299 | 4.58E-07 | YES | NO | NA |
| Tkfc | ENSMUSG( | ENSMUSG( | 0.050969 | 0.62774 | NO | NO | NA |
| Gch1 | ENSMUSG( | ENSMUSG( | -0.24252 | 0.071497 | NO | NO | NA |
| Mt3 | ENSMUSG( | ENSMUSG( | 1.097071 | 4.39E-10 | YES | YES | Upregulated |
| ENSMUSG( | ENSMUSG( | ENSMUSG( | -0.51088 | 1.79E-05 | YES | NO | NA |
| Tmem68 | ENSMUSG( | ENSMUSG( | -0.22684 | 0.016571 | YES | NO | NA |
| Tnip2 | ENSMUSG( | ENSMUSG( | 0.211778 | 0.162256 | NO | NO | NA |
| Zfp692 | ENSMUSG( | ENSMUSG( | 0.063412 | 0.476831 | NO | NO | NA |
| Sema4b | ENSMUSG( | ENSMUSG( | 0.100333 | 0.471485 | NO | NO | NA |
| Adi1 | ENSMUSG( | ENSMUSG( | 0.290052 | 0.00066 | YES | NO | NA |
| Gm49391 | ENSMUSG( | ENSMUSG( | 0.533168 | 0.011053 | YES | NO | NA |
| Ccdc125 | ENSMUSG( | ENSMUSG( | 0.535699 | 3.87E-06 | YES | NO | NA |
| Alg1 | ENSMUSG( | ENSMUSG( | -0.06287 | 0.62015 | NO | NO | NA |
| Unc93b1 | ENSMUSG( | ENSMUSG( | -0.07179 | 0.699819 | NO | NO | NA |
| Tmem185l | ENSMUSG( | ENSMUSG( | -0.11732 | 0.319022 | NO | NO | NA |
| Polr2i | ENSMUSG( | ENSMUSG( | 0.300871 | 0.004274 | YES | NO | NA |
| Fam136a | ENSMUSG( | ENSMUSG( | 0.306641 | 0.057187 | NO | NO | NA |
| Hibch | ENSMUSG( | ENSMUSG( | 0.157347 | 0.090892 | NO | NO | NA |
| Ift46 | ENSMUSG( | ENSMUSG( | 0.258383 | 0.001371 | YES | NO | NA |
| Taf7 | ENSMUSG( | ENSMUSG( | -0.40095 | 0.000107 | YES | NO | NA |
| Synpo2 | ENSMUSG( | ENSMUSG( | 1.064835 | 7.33E-05 | YES | YES | Upregulated |
| Pdss1 | ENSMUSG( | ENSMUSG( | 0.719324 | 2.02E-09 | YES | NO | NA |
| Lcorl | ENSMUSG( | ENSMUSG( | 0.046169 | 0.65894 | NO | NO | NA |
| Gm12715 | ENSMUSG( | ENSMUSG( | -0.09139 | 0.637997 | NO | NO | NA |
| Arhgap39 | ENSMUSG( | ENSMUSG( | -0.62186 | 2.35E-06 | YES | NO | NA |
| Fam220a E | ENSMUSG( | ENSMUSG( | -0.24695 | 0.197407 | NO | NO | NA |

| | | | | | | | |
|---|---|---|---|---|---|---|---|
| Gm7072 | ENSMUSG( | ENSMUSG( | -0.06614 | 0.696353 | NO | NO | NA |
| Tpcn2 | ENSMUSG( | ENSMUSG( | -0.02966 | 0.790471 | NO | NO | NA |
| Nos1 | ENSMUSG( | ENSMUSG( | 2.147459 | 1.38E-12 | YES | YES | Upregulated |
| Gtf2ird2 | ENSMUSG( | ENSMUSG( | -0.12307 | 0.171384 | NO | NO | NA |
| Snhg16 | ENSMUSG( | ENSMUSG( | 0.544603 | 8.96E-07 | YES | NO | NA |
| Trp73 | ENSMUSG( | ENSMUSG( | 1.015136 | 7.31E-08 | YES | YES | Upregulated |
| Grhpr | ENSMUSG( | ENSMUSG( | 0.430927 | 0.031289 | YES | NO | NA |
| Nt5dc3 | ENSMUSG( | ENSMUSG( | -0.02821 | 0.78461 | NO | NO | NA |
| Rffl | ENSMUSG( | ENSMUSG( | 0.462717 | 3.57E-05 | YES | NO | NA |
| Pisd | ENSMUSG( | ENSMUSG( | 0.112341 | 0.212815 | NO | NO | NA |
| Elac1 | ENSMUSG( | ENSMUSG( | 0.384598 | 0.002575 | YES | NO | NA |
| ENSMUSG( | ENSMUSG( | ENSMUSG( | 0.236433 | 0.147209 | NO | NO | NA |
| Ccdc84 | ENSMUSG( | ENSMUSG( | -0.14315 | 0.267916 | NO | NO | NA |
| Trav8n-2 | ENSMUSG( | ENSMUSG( | -0.56034 | 0.587391 | NO | NO | NA |
| Skp2 | ENSMUSG( | ENSMUSG( | 0.369792 | 2.23E-06 | YES | NO | NA |
| Akr1b10 | ENSMUSG( | ENSMUSG( | -0.27341 | 0.034128 | YES | NO | NA |
| Prkrip1 | ENSMUSG( | ENSMUSG( | 0.135298 | 0.2223 | NO | NO | NA |
| Zbtb6 | ENSMUSG( | ENSMUSG( | -0.41257 | 1.02E-05 | YES | NO | NA |
| Mrps27 | ENSMUSG( | ENSMUSG( | 0.590768 | 1.10E-06 | YES | NO | NA |
| Fars2 | ENSMUSG( | ENSMUSG( | 0.406963 | 1.29E-05 | YES | NO | NA |
| Zxda | ENSMUSG( | ENSMUSG( | -0.3456 | 0.025232 | YES | NO | NA |
| Magohb | ENSMUSG( | ENSMUSG( | 0.387067 | 0.008751 | YES | NO | NA |
| Smpd2 | ENSMUSG( | ENSMUSG( | -0.14561 | 0.216908 | NO | NO | NA |
| D630039A | ENSMUSG( | ENSMUSG( | -0.1372 | 0.171349 | NO | NO | NA |
| Cep72 | ENSMUSG( | ENSMUSG( | 0.424736 | 4.57E-05 | YES | NO | NA |
| Dscc1 | ENSMUSG( | ENSMUSG( | 0.262079 | 0.068843 | NO | NO | NA |
| Map2 | ENSMUSG( | ENSMUSG( | -0.71302 | 7.68E-10 | YES | NO | NA |
| Ccdc32 | ENSMUSG( | ENSMUSG( | 0.089257 | 0.378259 | NO | NO | NA |
| Rmi2 | ENSMUSG( | ENSMUSG( | -0.01843 | 0.922432 | NO | NO | NA |
| Glmn | ENSMUSG( | ENSMUSG( | -0.03628 | 0.724216 | NO | NO | NA |
| Poll | ENSMUSG( | ENSMUSG( | 0.009248 | 0.950039 | NO | NO | NA |
| Mnat1 | ENSMUSG( | ENSMUSG( | 0.358053 | 0.0149 | YES | NO | NA |
| Prxl2c | ENSMUSG( | ENSMUSG( | -0.15777 | 0.094529 | NO | NO | NA |
| Zfp593 | ENSMUSG( | ENSMUSG( | 0.471021 | 1.25E-05 | YES | NO | NA |
| Mthfsd | ENSMUSG( | ENSMUSG( | -0.12494 | 0.086784 | NO | NO | NA |
| Mtrf1l | ENSMUSG( | ENSMUSG( | -0.17104 | 0.164596 | NO | NO | NA |
| Stn1 | ENSMUSG( | ENSMUSG( | -0.18236 | 0.044863 | YES | NO | NA |
| Chchd1 | ENSMUSG( | ENSMUSG( | 0.217966 | 0.088325 | NO | NO | NA |
| Tube1 | ENSMUSG( | ENSMUSG( | 0.242264 | 0.056195 | NO | NO | NA |
| Asb3 | ENSMUSG( | ENSMUSG( | -0.01979 | 0.817551 | NO | NO | NA |
| Lrrc57 | ENSMUSG( | ENSMUSG( | -0.20828 | 0.037349 | YES | NO | NA |
| Pnpo | ENSMUSG( | ENSMUSG( | 0.191921 | 0.078843 | NO | NO | NA |
| Axin2 | ENSMUSG( | ENSMUSG( | -0.98049 | 0.000671 | YES | NO | NA |
| Aldh7a1 | ENSMUSG( | ENSMUSG( | 0.44097 | 1.22E-05 | YES | NO | NA |
| Ppp2r5b | ENSMUSG( | ENSMUSG( | -0.53349 | 5.34E-09 | YES | NO | NA |
| Rnf215 | ENSMUSG( | ENSMUSG( | -0.1575 | 0.260276 | NO | NO | NA |
| Gpsm1 | ENSMUSG( | ENSMUSG( | -0.52152 | 0.000187 | YES | NO | NA |

| | | | | | | | |
|---|---|---|---|---|---|---|---|
| Tjp3 | ENSMUSG( | ENSMUSG( | -0.44263 | 1.05E-05 | YES | NO | NA |
| Pinx1 | ENSMUSG( | ENSMUSG( | 0.514191 | 2.71E-05 | YES | NO | NA |
| Poglut1 | ENSMUSG( | ENSMUSG( | 0.217807 | 0.002533 | YES | NO | NA |
| Fbxo25 | ENSMUSG( | ENSMUSG( | -0.38279 | 0.000149 | YES | NO | NA |
| Mib2 | ENSMUSG( | ENSMUSG( | -0.64795 | 1.97E-07 | YES | NO | NA |
| Armh3 | ENSMUSG( | ENSMUSG( | -0.00133 | 0.991511 | NO | NO | NA |
| Ryr1 | ENSMUSG( | ENSMUSG( | -0.17101 | 0.259107 | NO | NO | NA |
| Tmem256 | ENSMUSG( | ENSMUSG( | 0.136951 | 0.143881 | NO | NO | NA |
| Gm37416 | ENSMUSG( | ENSMUSG( | 0.728614 | 2.79E-05 | YES | NO | NA |
| Coil | ENSMUSG( | ENSMUSG( | -0.00945 | 0.929258 | NO | NO | NA |
| Cd68 | ENSMUSG( | ENSMUSG( | -0.56422 | 0.003647 | YES | NO | NA |
| Selenon | ENSMUSG( | ENSMUSG( | -0.83412 | 1.13E-09 | YES | NO | NA |
| Hps4 | ENSMUSG( | ENSMUSG( | -0.12079 | 0.199403 | NO | NO | NA |
| Hs2st1 | ENSMUSG( | ENSMUSG( | 0.20863 | 0.04063 | YES | NO | NA |
| Tm2d2 | ENSMUSG( | ENSMUSG( | 0.242076 | 0.020578 | YES | NO | NA |
| Cox18 | ENSMUSG( | ENSMUSG( | 0.090213 | 0.386356 | NO | NO | NA |
| Dpm3 | ENSMUSG( | ENSMUSG( | 0.29337 | 0.013228 | YES | NO | NA |
| Arrdc3 | ENSMUSG( | ENSMUSG( | -0.90299 | 6.17E-06 | YES | NO | NA |
| B930095G | ENSMUSG( | ENSMUSG( | -0.16063 | 0.047841 | YES | NO | NA |
| Tef | ENSMUSG( | ENSMUSG( | 0.09299 | 0.679852 | NO | NO | NA |
| Sap30 | ENSMUSG( | ENSMUSG( | 0.084918 | 0.580155 | NO | NO | NA |
| Gale | ENSMUSG( | ENSMUSG( | -0.15244 | 0.127357 | NO | NO | NA |
| Zfp35 | ENSMUSG( | ENSMUSG( | -0.12274 | 0.127868 | NO | NO | NA |
| Slc25a13 | ENSMUSG( | ENSMUSG( | 0.705805 | 4.19E-08 | YES | NO | NA |
| ENSMUSG | ENSMUSG( | ENSMUSG( | 0.928656 | 4.47E-07 | YES | NO | NA |
| Vax2 | ENSMUSG( | ENSMUSG( | 0.363714 | 0.055677 | NO | NO | NA |
| Dhodh | ENSMUSG( | ENSMUSG( | 0.445226 | 0.000425 | YES | NO | NA |
| Tmem230 | ENSMUSG( | ENSMUSG( | -0.42662 | 7.45E-06 | YES | NO | NA |
| Dph2 | ENSMUSG( | ENSMUSG( | 0.077408 | 0.421169 | NO | NO | NA |
| Sap25 | ENSMUSG( | ENSMUSG( | -0.34234 | 0.023824 | YES | NO | NA |
| Pold4 | ENSMUSG( | ENSMUSG( | -0.10806 | 0.454063 | NO | NO | NA |
| Zfp46 | ENSMUSG( | ENSMUSG( | -0.24751 | 0.009378 | YES | NO | NA |
| Il12rb2 | ENSMUSG( | ENSMUSG( | 1.127316 | 3.64E-07 | YES | YES | Upregulated |
| Fuz | ENSMUSG( | ENSMUSG( | -0.1841 | 0.077368 | NO | NO | NA |
| Polr3d | ENSMUSG( | ENSMUSG( | 0.20111 | 0.07265 | NO | NO | NA |
| Ftsj1 | ENSMUSG( | ENSMUSG( | 0.318645 | 4.50E-05 | YES | NO | NA |
| Gm48274 | ENSMUSG( | ENSMUSG( | 1.252718 | 1.05E-07 | YES | YES | Upregulated |
| Scart2 | ENSMUSG( | ENSMUSG( | 2.621664 | 0.000978 | YES | YES | Upregulated |
| Pomgnt1 | ENSMUSG( | ENSMUSG( | -0.07496 | 0.511623 | NO | NO | NA |
| Haus1 | ENSMUSG( | ENSMUSG( | 0.522944 | 0.000312 | YES | NO | NA |
| Cep78 | ENSMUSG( | ENSMUSG( | 0.410368 | 1.16E-06 | YES | NO | NA |
| Gm11131 | ENSMUSG( | ENSMUSG( | -0.60166 | 2.07E-06 | YES | NO | NA |
| Mrpl1 | ENSMUSG( | ENSMUSG( | 0.34209 | 0.000935 | YES | NO | NA |
| Phtf2 | ENSMUSG( | ENSMUSG( | 0.139284 | 0.118033 | NO | NO | NA |
| Nckap1 | ENSMUSG( | ENSMUSG( | 0.554622 | 0.004425 | YES | NO | NA |
| Rwdd2b | ENSMUSG( | ENSMUSG( | 0.057054 | 0.536617 | NO | NO | NA |
| Fbxo4 | ENSMUSG( | ENSMUSG( | 0.304361 | 0.000269 | YES | NO | NA |

| | | | | | | | |
|---|---|---|---|---|---|---|---|
| Osm | ENSMUSG( | ENSMUSG( | 0.346553 | 0.001048 | YES | NO | NA |
| 1300002E: | ENSMUSG( | ENSMUSG( | -0.47633 | 1.12E-06 | YES | NO | NA |
| Izumo4 | ENSMUSG( | ENSMUSG( | -0.624 | 6.60E-08 | YES | NO | NA |
| AI987944 | ENSMUSG( | ENSMUSG( | -0.2439 | 0.073612 | NO | NO | NA |
| D430020J( | ENSMUSG( | ENSMUSG( | 0.147514 | 0.410561 | NO | NO | NA |
| Cd72 | ENSMUSG( | ENSMUSG( | -0.484 | 0.004696 | YES | NO | NA |
| Mrps22 | ENSMUSG( | ENSMUSG( | 0.419725 | 0.001022 | YES | NO | NA |
| Spindoc | ENSMUSG( | ENSMUSG( | -0.16557 | 0.32583 | NO | NO | NA |
| Slc16a10 | ENSMUSG( | ENSMUSG( | -0.82578 | 2.51E-06 | YES | NO | NA |
| Gtpbp10 | ENSMUSG( | ENSMUSG( | 0.356682 | 0.000111 | YES | NO | NA |
| Rsph3a | ENSMUSG( | ENSMUSG( | 0.28304 | 0.011277 | YES | NO | NA |
| Amn1 | ENSMUSG( | ENSMUSG( | 0.080448 | 0.551005 | NO | NO | NA |
| Abcb10 | ENSMUSG( | ENSMUSG( | 0.233516 | 0.068988 | NO | NO | NA |
| Trbv17 | ENSMUSG( | ENSMUSG( | 0.139125 | 0.689372 | NO | NO | NA |
| Tfpt | ENSMUSG( | ENSMUSG( | 0.101936 | 0.465954 | NO | NO | NA |
| Smug1 | ENSMUSG( | ENSMUSG( | -0.43402 | 0.001368 | YES | NO | NA |
| Gfer | ENSMUSG( | ENSMUSG( | 0.305087 | 0.009675 | YES | NO | NA |
| Stambp | ENSMUSG( | ENSMUSG( | 0.042766 | 0.706403 | NO | NO | NA |
| Trmt13 | ENSMUSG( | ENSMUSG( | 0.035001 | 0.792583 | NO | NO | NA |
| Prickle3 | ENSMUSG( | ENSMUSG( | -0.22842 | 0.118798 | NO | NO | NA |
| Invs | ENSMUSG( | ENSMUSG( | -0.4729 | 4.67E-07 | YES | NO | NA |
| Serpina3f | ENSMUSG( | ENSMUSG( | 0.783383 | 0.000177 | YES | NO | NA |
| Ebpl | ENSMUSG( | ENSMUSG( | 0.058782 | 0.722171 | NO | NO | NA |
| Fam126b | ENSMUSG( | ENSMUSG( | -0.01381 | 0.928272 | NO | NO | NA |
| Gm33989 | ENSMUSG( | ENSMUSG( | -0.54129 | 0.000191 | YES | NO | NA |
| F8a | ENSMUSG( | ENSMUSG( | 0.16355 | 0.136624 | NO | NO | NA |
| Daglb | ENSMUSG( | ENSMUSG( | -0.04124 | 0.782763 | NO | NO | NA |
| Czib | ENSMUSG( | ENSMUSG( | -0.08037 | 0.463112 | NO | NO | NA |
| Pex11b | ENSMUSG( | ENSMUSG( | -0.01286 | 0.895322 | NO | NO | NA |
| Ddias | ENSMUSG( | ENSMUSG( | 0.216711 | 0.082784 | NO | NO | NA |
| Mvb12b | ENSMUSG( | ENSMUSG( | 0.607483 | 0.0021 | YES | NO | NA |
| Rbm48 | ENSMUSG( | ENSMUSG( | -0.09388 | 0.499325 | NO | NO | NA |
| Gm49396 | ENSMUSG( | ENSMUSG( | -0.07338 | 0.553112 | NO | NO | NA |
| Lpar6 | ENSMUSG( | ENSMUSG( | -0.79248 | 2.02E-07 | YES | NO | NA |
| Plekhn1 | ENSMUSG( | ENSMUSG( | -0.73498 | 1.42E-08 | YES | NO | NA |
| Nkrf | ENSMUSG( | ENSMUSG( | 0.489807 | 3.03E-05 | YES | NO | NA |
| Pgm3 | ENSMUSG( | ENSMUSG( | 0.276876 | 0.001109 | YES | NO | NA |
| Zranb3 | ENSMUSG( | ENSMUSG( | -0.19687 | 0.172945 | NO | NO | NA |
| Mtss1 | ENSMUSG( | ENSMUSG( | 1.200811 | 4.18E-12 | YES | YES | Upregulated |
| Trbv12-1 | ENSMUSG( | ENSMUSG( | -1.36141 | 1.25E-05 | YES | YES | Downregulated |
| Snhg4 ENS | ENSMUSG( | ENSMUSG( | 0.991003 | 9.33E-13 | YES | NO | NA |
| Rev1 | ENSMUSG( | ENSMUSG( | 0.37526 | 5.67E-06 | YES | NO | NA |
| Abcg3 | ENSMUSG( | ENSMUSG( | -1.21464 | 3.62E-12 | YES | YES | Downregulated |
| Il7r | ENSMUSG( | ENSMUSG( | 1.49819 | 0.000253 | YES | YES | Upregulated |
| Sco2 | ENSMUSG( | ENSMUSG( | 0.393632 | 9.96E-05 | YES | NO | NA |
| Pex13 | ENSMUSG( | ENSMUSG( | -0.09017 | 0.428766 | NO | NO | NA |
| Tfap4 | ENSMUSG( | ENSMUSG( | 0.870464 | 2.84E-09 | YES | NO | NA |

| | | | | | | | |
|---|---|---|---|---|---|---|---|
| Atxn7l2 | ENSMUSG( | ENSMUSG( | -0.02716 | 0.810987 | NO | NO | NA |
| Ctla2b | ENSMUSG( | ENSMUSG( | 0.166738 | 0.436017 | NO | NO | NA |
| Tiparp | ENSMUSG( | ENSMUSG( | -0.12348 | 0.366033 | NO | NO | NA |
| Jagn1 | ENSMUSG( | ENSMUSG( | 0.263773 | 0.012201 | YES | NO | NA |
| Trbv5 | ENSMUSG( | ENSMUSG( | 0.377161 | 0.597166 | NO | NO | NA |
| Elk1 | ENSMUSG( | ENSMUSG( | -0.24497 | 0.021977 | YES | NO | NA |
| Sepsecs | ENSMUSG( | ENSMUSG( | 0.002412 | 0.98477 | NO | NO | NA |
| Rbak | ENSMUSG( | ENSMUSG( | 0.03205 | 0.868704 | NO | NO | NA |
| Gss | ENSMUSG( | ENSMUSG( | 0.599177 | 3.47E-05 | YES | NO | NA |
| Rnf103 | ENSMUSG( | ENSMUSG( | -0.08469 | 0.574263 | NO | NO | NA |
| Chka | ENSMUSG( | ENSMUSG( | 0.2064 | 0.091723 | NO | NO | NA |
| Mfsd8 | ENSMUSG( | ENSMUSG( | -0.16206 | 0.140252 | NO | NO | NA |
| Nfxl1 | ENSMUSG( | ENSMUSG( | 0.022893 | 0.846339 | NO | NO | NA |
| Rpp30 | ENSMUSG( | ENSMUSG( | 0.401763 | 8.04E-05 | YES | NO | NA |
| Wwox | ENSMUSG( | ENSMUSG( | 0.347882 | 0.002588 | YES | NO | NA |
| Gemin2 | ENSMUSG( | ENSMUSG( | 0.102802 | 0.275121 | NO | NO | NA |
| Nek8 | ENSMUSG( | ENSMUSG( | 0.407264 | 0.000934 | YES | NO | NA |
| Btrc | ENSMUSG( | ENSMUSG( | -0.12879 | 0.312213 | NO | NO | NA |
| Zbtb20 | ENSMUSG( | ENSMUSG( | 0.196853 | 0.512691 | NO | NO | NA |
| Msmo1 | ENSMUSG( | ENSMUSG( | 0.531013 | 3.92E-07 | YES | NO | NA |
| Nr4a3 | ENSMUSG( | ENSMUSG( | 0.408901 | 0.072893 | NO | NO | NA |
| Tmco3 | ENSMUSG( | ENSMUSG( | -0.53875 | 1.16E-08 | YES | NO | NA |
| Bcl2l12 | ENSMUSG( | ENSMUSG( | -0.03045 | 0.725782 | NO | NO | NA |
| Tmem39b | ENSMUSG( | ENSMUSG( | -0.08558 | 0.353715 | NO | NO | NA |
| ENSMUSG( | ENSMUSG( | ENSMUSG( | -0.62767 | 1.02E-07 | YES | NO | NA |
| Kctd13 | ENSMUSG( | ENSMUSG( | -0.00872 | 0.931291 | NO | NO | NA |
| Perp | ENSMUSG( | ENSMUSG( | -0.76128 | 4.42E-06 | YES | NO | NA |
| Ttc19 | ENSMUSG( | ENSMUSG( | -0.06086 | 0.637041 | NO | NO | NA |
| Syce2 | ENSMUSG( | ENSMUSG( | 0.49837 | 0.002477 | YES | NO | NA |
| Psmc3ip | ENSMUSG( | ENSMUSG( | 0.258103 | 0.008857 | YES | NO | NA |
| Ormdl2 | ENSMUSG( | ENSMUSG( | -0.15777 | 0.045813 | YES | NO | NA |
| Sfxn2 | ENSMUSG( | ENSMUSG( | 0.103731 | 0.32459 | NO | NO | NA |
| Zfp160 | ENSMUSG( | ENSMUSG( | -0.33579 | 0.001985 | YES | NO | NA |
| S1pr2 | ENSMUSG( | ENSMUSG( | -0.66874 | 3.97E-05 | YES | NO | NA |
| Odf2l | ENSMUSG( | ENSMUSG( | 0.386056 | 2.12E-06 | YES | NO | NA |
| Casp4 | ENSMUSG( | ENSMUSG( | 0.097476 | 0.595836 | NO | NO | NA |
| Sdhaf1 | ENSMUSG( | ENSMUSG( | 0.685614 | 3.15E-06 | YES | NO | NA |
| Rnf157 | ENSMUSG( | ENSMUSG( | 0.944572 | 3.94E-10 | YES | NO | NA |
| Fam199x | ENSMUSG( | ENSMUSG( | 0.201491 | 0.150183 | NO | NO | NA |
| Gm12481 | ENSMUSG( | ENSMUSG( | -0.12561 | 0.860652 | NO | NO | NA |
| Rab9 | ENSMUSG( | ENSMUSG( | -0.01113 | 0.917747 | NO | NO | NA |
| Zfp579 | ENSMUSG( | ENSMUSG( | 0.081271 | 0.63596 | NO | NO | NA |
| Elovl6 | ENSMUSG( | ENSMUSG( | 0.949101 | 4.38E-17 | YES | NO | NA |
| Septin4 | ENSMUSG( | ENSMUSG( | -1.20901 | 3.48E-08 | YES | YES | Downregulated |
| S1pr5 | ENSMUSG( | ENSMUSG( | -2.35125 | 9.48E-08 | YES | YES | Downregulated |
| Rad51c | ENSMUSG( | ENSMUSG( | -0.12639 | 0.302975 | NO | NO | NA |
| Cenpk | ENSMUSG( | ENSMUSG( | 0.357184 | 0.019672 | YES | NO | NA |

| | | | | | | | |
|---|---|---|---|---|---|---|---|
| Agl | ENSMUSG( | ENSMUSG( | 0.385643 | 0.000778 | YES | NO | NA |
| Nbn | ENSMUSG( | ENSMUSG( | 0.546308 | 4.35E-06 | YES | NO | NA |
| Smyd1 | ENSMUSG( | ENSMUSG( | 0.285752 | 0.000706 | YES | NO | NA |
| Trav6n-7 | ENSMUSG( | ENSMUSG( | -0.02255 | 0.93838 | NO | NO | NA |
| Galnt10 | ENSMUSG( | ENSMUSG( | -1.04514 | 1.15E-09 | YES | YES | Downregulated |
| Mllt11 | ENSMUSG( | ENSMUSG( | -0.46192 | 0.001548 | YES | NO | NA |
| Znhit6 | ENSMUSG( | ENSMUSG( | 0.236269 | 0.237115 | NO | NO | NA |
| Psmg1 | ENSMUSG( | ENSMUSG( | 0.592102 | 0.000171 | YES | NO | NA |
| Snapc2 | ENSMUSG( | ENSMUSG( | 0.059705 | 0.613303 | NO | NO | NA |
| Pfdn4 | ENSMUSG( | ENSMUSG( | 0.163113 | 0.201819 | NO | NO | NA |
| Fam221b | ENSMUSG( | ENSMUSG( | -0.11336 | 0.488115 | NO | NO | NA |
| Slc35a5 | ENSMUSG( | ENSMUSG( | -0.52074 | 0.000106 | YES | NO | NA |
| Cd9 | ENSMUSG( | ENSMUSG( | -0.31216 | 0.306075 | NO | NO | NA |
| 1700025G | ENSMUSG( | ENSMUSG( | -0.9417 | 3.28E-05 | YES | NO | NA |
| Klrb1c | ENSMUSG( | ENSMUSG( | 0.934536 | 0.030441 | YES | NO | NA |
| Inafm2 | ENSMUSG( | ENSMUSG( | 0.513733 | 2.66E-07 | YES | NO | NA |
| Zbtb5 | ENSMUSG( | ENSMUSG( | -0.11912 | 0.30875 | NO | NO | NA |
| Pkdcc | ENSMUSG( | ENSMUSG( | -0.75813 | 0.000117 | YES | NO | NA |
| Zfp961 | ENSMUSG( | ENSMUSG( | -0.23939 | 0.004702 | YES | NO | NA |
| Gin1 | ENSMUSG( | ENSMUSG( | -0.08724 | 0.515346 | NO | NO | NA |
| Map3k10 | ENSMUSG( | ENSMUSG( | 0.062863 | 0.703614 | NO | NO | NA |
| Npnt | ENSMUSG( | ENSMUSG( | 1.331112 | 1.41E-08 | YES | YES | Upregulated |
| Rpl17-ps8 | ENSMUSG( | ENSMUSG( | 0.099138 | 0.421127 | NO | NO | NA |
| Mto1 | ENSMUSG( | ENSMUSG( | 0.21145 | 0.044672 | YES | NO | NA |
| Platr25 | ENSMUSG( | ENSMUSG( | -0.35666 | 0.000224 | YES | NO | NA |
| Pdp2 | ENSMUSG( | ENSMUSG( | -0.05124 | 0.573921 | NO | NO | NA |
| Ier3 | ENSMUSG( | ENSMUSG( | -0.50349 | 0.004109 | YES | NO | NA |
| 2810013P( | ENSMUSG( | ENSMUSG( | -0.05369 | 0.731009 | NO | NO | NA |
| Car12 | ENSMUSG( | ENSMUSG( | 1.607606 | 1.07E-11 | YES | YES | Upregulated |
| Slfn3 | ENSMUSG( | ENSMUSG( | 0.1249 | 0.355996 | NO | NO | NA |
| Pam16 | ENSMUSG( | ENSMUSG( | 0.175489 | 0.086667 | NO | NO | NA |
| Akap1 | ENSMUSG( | ENSMUSG( | 1.259586 | 9.94E-08 | YES | YES | Upregulated |
| Fam174a | ENSMUSG( | ENSMUSG( | -0.0302 | 0.838501 | NO | NO | NA |
| Gbe1 | ENSMUSG( | ENSMUSG( | 0.190495 | 0.041505 | YES | NO | NA |
| Zfand1 | ENSMUSG( | ENSMUSG( | 0.361561 | 0.004414 | YES | NO | NA |
| Lyrm9 | ENSMUSG( | ENSMUSG( | 0.213218 | 0.029648 | YES | NO | NA |
| Pdxp | ENSMUSG( | ENSMUSG( | -0.03584 | 0.758867 | NO | NO | NA |
| Cyp4v3 | ENSMUSG( | ENSMUSG( | -0.19442 | 0.30973 | NO | NO | NA |
| Dnmt3b | ENSMUSG( | ENSMUSG( | -0.05364 | 0.665302 | NO | NO | NA |
| Acsf3 | ENSMUSG( | ENSMUSG( | 0.088266 | 0.406876 | NO | NO | NA |
| Zfp729b | ENSMUSG( | ENSMUSG( | -0.39452 | 0.003734 | YES | NO | NA |
| Dph5 | ENSMUSG( | ENSMUSG( | 0.284941 | 0.007959 | YES | NO | NA |
| Zfp954 | ENSMUSG( | ENSMUSG( | -0.1553 | 0.302839 | NO | NO | NA |
| Pigc | ENSMUSG( | ENSMUSG( | -0.35657 | 3.77E-06 | YES | NO | NA |
| Pdlim7 | ENSMUSG( | ENSMUSG( | -0.18504 | 0.003755 | YES | NO | NA |
| Gm48942 | ENSMUSG( | ENSMUSG( | 0.314699 | 0.389704 | NO | NO | NA |
| Gm43802 | ENSMUSG( | ENSMUSG( | 0.114422 | 0.394547 | NO | NO | NA |

| | | | | | | | |
|---|---|---|---|---|---|---|---|
| Mlycd | ENSMUSG( | ENSMUSG( | -0.28636 | 0.000281 | YES | NO | NA |
| Trpm1 | ENSMUSG( | ENSMUSG( | -0.00699 | 0.975934 | NO | NO | NA |
| Bpnt1 | ENSMUSG( | ENSMUSG( | 0.058518 | 0.534809 | NO | NO | NA |
| Eci1 | ENSMUSG( | ENSMUSG( | -0.0612 | 0.606907 | NO | NO | NA |
| Ankrd49 | ENSMUSG( | ENSMUSG( | 0.236732 | 0.030535 | YES | NO | NA |
| Fam83d | ENSMUSG( | ENSMUSG( | 0.216172 | 0.050469 | NO | NO | NA |
| Gm13610 | ENSMUSG( | ENSMUSG( | -0.6541 | 7.59E-06 | YES | NO | NA |
| Zbtb42 | ENSMUSG( | ENSMUSG( | -0.35527 | 0.002722 | YES | NO | NA |
| Pdp1 | ENSMUSG( | ENSMUSG( | 0.177063 | 0.062346 | NO | NO | NA |
| Gm49339 | ENSMUSG( | ENSMUSG( | -0.30769 | 0.042715 | YES | NO | NA |
| Soat2 | ENSMUSG( | ENSMUSG( | 0.009247 | 0.96437 | NO | NO | NA |
| Fpgs | ENSMUSG( | ENSMUSG( | 0.794849 | 1.18E-07 | YES | NO | NA |
| Gm4631 | ENSMUSG( | ENSMUSG( | -0.04216 | 0.787243 | NO | NO | NA |
| Prkci | ENSMUSG( | ENSMUSG( | 0.37025 | 0.001419 | YES | NO | NA |
| Zkscan6 | ENSMUSG( | ENSMUSG( | -0.22295 | 0.002319 | YES | NO | NA |
| Abcd4 | ENSMUSG( | ENSMUSG( | -0.21747 | 0.044902 | YES | NO | NA |
| Cox19 | ENSMUSG( | ENSMUSG( | -0.07394 | 0.473873 | NO | NO | NA |
| Atcay | ENSMUSG( | ENSMUSG( | -0.08486 | 0.575279 | NO | NO | NA |
| Mtg2 | ENSMUSG( | ENSMUSG( | -0.24274 | 0.002175 | YES | NO | NA |
| Gtf2h4 | ENSMUSG( | ENSMUSG( | 0.481646 | 1.74E-07 | YES | NO | NA |
| Ndufaf4 | ENSMUSG( | ENSMUSG( | 0.380866 | 8.56E-05 | YES | NO | NA |
| Eef1akmt1 | ENSMUSG( | ENSMUSG( | -0.01449 | 0.914757 | NO | NO | NA |
| Trav6d-7 | ENSMUSG( | ENSMUSG( | 0.022894 | 0.934193 | NO | NO | NA |
| Tma16 | ENSMUSG( | ENSMUSG( | 0.799141 | 9.89E-08 | YES | NO | NA |
| Tnfrsf4 | ENSMUSG( | ENSMUSG( | 1.433694 | 3.23E-13 | YES | YES | Upregulated |
| Acot2 | ENSMUSG( | ENSMUSG( | -0.09368 | 0.43166 | NO | NO | NA |
| Isoc2a | ENSMUSG( | ENSMUSG( | 0.230703 | 0.043925 | YES | NO | NA |
| Rbm44 | ENSMUSG( | ENSMUSG( | 1.438326 | 1.06E-07 | YES | YES | Upregulated |
| Ralgds | ENSMUSG( | ENSMUSG( | -0.28503 | 0.006484 | YES | NO | NA |
| Zfp943 | ENSMUSG( | ENSMUSG( | -0.54112 | 3.21E-05 | YES | NO | NA |
| Gphn | ENSMUSG( | ENSMUSG( | 0.196285 | 0.021363 | YES | NO | NA |
| B9d2 | ENSMUSG( | ENSMUSG( | 0.370251 | 0.002135 | YES | NO | NA |
| Gcnt1 | ENSMUSG( | ENSMUSG( | -0.02333 | 0.916437 | NO | NO | NA |
| Fam162a | ENSMUSG( | ENSMUSG( | 0.282063 | 0.037708 | YES | NO | NA |
| Abi3 | ENSMUSG( | ENSMUSG( | -0.83298 | 0.000565 | YES | NO | NA |
| 4933404O | ENSMUSG( | ENSMUSG( | 0.404607 | 0.003835 | YES | NO | NA |
| Tax1bp3 | ENSMUSG( | ENSMUSG( | -0.46154 | 8.04E-06 | YES | NO | NA |
| Borcs5 | ENSMUSG( | ENSMUSG( | 0.022686 | 0.888656 | NO | NO | NA |
| Zfp169 | ENSMUSG( | ENSMUSG( | -0.8758 | 2.55E-10 | YES | NO | NA |
| Gm45133 | ENSMUSG( | ENSMUSG( | -0.31605 | 0.070622 | NO | NO | NA |
| Mroh2a | ENSMUSG( | ENSMUSG( | -0.13076 | 0.280628 | NO | NO | NA |
| Socs1 | ENSMUSG( | ENSMUSG( | 0.03768 | 0.783471 | NO | NO | NA |
| N6amt1 | ENSMUSG( | ENSMUSG( | 0.297746 | 0.000524 | YES | NO | NA |
| C1qtnf6 | ENSMUSG( | ENSMUSG( | -0.3796 | 0.004709 | YES | NO | NA |
| 9330162G | ENSMUSG( | ENSMUSG( | 0.14604 | 0.640929 | NO | NO | NA |
| Faxc | ENSMUSG( | ENSMUSG( | -0.42919 | 0.001465 | YES | NO | NA |
| Lrrc14 | ENSMUSG( | ENSMUSG( | -0.17872 | 0.074422 | NO | NO | NA |

| | | | | | | | |
|---|---|---|---|---|---|---|---|
| Lysmd2 | ENSMUSG( | ENSMUSG( | -0.50106 | 0.004296 | YES | NO | NA |
| Nf1 | ENSMUSG( | ENSMUSG( | -0.00028 | 0.997489 | NO | NO | NA |
| Rmnd1 | ENSMUSG( | ENSMUSG( | -0.13805 | 0.203489 | NO | NO | NA |
| Calhm6 | ENSMUSG( | ENSMUSG( | -0.10908 | 0.243801 | NO | NO | NA |
| Ap4s1 | ENSMUSG( | ENSMUSG( | -0.02663 | 0.83079 | NO | NO | NA |
| Snx15 | ENSMUSG( | ENSMUSG( | 0.082568 | 0.449445 | NO | NO | NA |
| Mocs1 | ENSMUSG( | ENSMUSG( | -0.0336 | 0.806496 | NO | NO | NA |
| Qpctl | ENSMUSG( | ENSMUSG( | 0.297766 | 0.015382 | YES | NO | NA |
| Tsga10 | ENSMUSG( | ENSMUSG( | -0.41905 | 0.00424 | YES | NO | NA |
| ENSMUSG | ENSMUSG( | ENSMUSG( | 0.368175 | 0.018865 | YES | NO | NA |
| Fbxo32 | ENSMUSG( | ENSMUSG( | -0.50048 | 0.000977 | YES | NO | NA |
| Pdxk | ENSMUSG( | ENSMUSG( | 0.613261 | 8.35E-06 | YES | NO | NA |
| Pld2 | ENSMUSG( | ENSMUSG( | -0.29216 | 0.00875 | YES | NO | NA |
| Fancc | ENSMUSG( | ENSMUSG( | 0.095345 | 0.436272 | NO | NO | NA |
| Zbtb21 | ENSMUSG( | ENSMUSG( | -0.00432 | 0.96972 | NO | NO | NA |
| Coq10a | ENSMUSG( | ENSMUSG( | 0.135126 | 0.349653 | NO | NO | NA |
| Bet1 | ENSMUSG( | ENSMUSG( | 0.093451 | 0.501479 | NO | NO | NA |
| Zfp626 | ENSMUSG( | ENSMUSG( | -0.32266 | 0.011443 | YES | NO | NA |
| Fam234b | ENSMUSG( | ENSMUSG( | -0.03063 | 0.826879 | NO | NO | NA |
| Tmem80 | ENSMUSG( | ENSMUSG( | -0.05428 | 0.683173 | NO | NO | NA |
| Snx33 | ENSMUSG( | ENSMUSG( | -0.40077 | 0.023136 | YES | NO | NA |
| ENSMUSG( | ENSMUSG( | ENSMUSG( | -0.89164 | 0.007514 | YES | NO | NA |
| Zscan20 | ENSMUSG( | ENSMUSG( | -0.62062 | 2.07E-07 | YES | NO | NA |
| Qtrt1 | ENSMUSG( | ENSMUSG( | 0.400616 | 0.001171 | YES | NO | NA |
| Exo5 | ENSMUSG( | ENSMUSG( | -0.16843 | 0.078158 | NO | NO | NA |
| Cdh24 | ENSMUSG( | ENSMUSG( | -0.41557 | 9.91E-05 | YES | NO | NA |
| Mccc1 | ENSMUSG( | ENSMUSG( | 0.258967 | 0.00458 | YES | NO | NA |
| Casp6 | ENSMUSG( | ENSMUSG( | -0.15292 | 0.241993 | NO | NO | NA |
| Jmjd4 | ENSMUSG( | ENSMUSG( | -0.27209 | 0.000367 | YES | NO | NA |
| Gm4673 | ENSMUSG( | ENSMUSG( | 0.197335 | 0.05352 | NO | NO | NA |
| Eral1 | ENSMUSG( | ENSMUSG( | 0.140552 | 0.14275 | NO | NO | NA |
| 1600020E( | ENSMUSG( | ENSMUSG( | 0.138611 | 0.469756 | NO | NO | NA |
| Exoc6b | ENSMUSG( | ENSMUSG( | -0.40294 | 1.75E-05 | YES | NO | NA |
| Erc1 | ENSMUSG( | ENSMUSG( | 0.146316 | 0.342258 | NO | NO | NA |
| Cd3eap | ENSMUSG( | ENSMUSG( | 0.677466 | 3.30E-06 | YES | NO | NA |
| Klhl36 | ENSMUSG( | ENSMUSG( | -0.19519 | 0.11748 | NO | NO | NA |
| Gm48161 | ENSMUSG( | ENSMUSG( | 0.020457 | 0.928272 | NO | NO | NA |
| Neurl1b | ENSMUSG( | ENSMUSG( | -0.24589 | 0.001234 | YES | NO | NA |
| Ctse | ENSMUSG( | ENSMUSG( | 0.109339 | 0.614449 | NO | NO | NA |
| Dcaf4 | ENSMUSG( | ENSMUSG( | -0.01138 | 0.93409 | NO | NO | NA |
| Adck5 | ENSMUSG( | ENSMUSG( | 0.493391 | 1.27E-05 | YES | NO | NA |
| Bmt2 | ENSMUSG( | ENSMUSG( | -0.13105 | 0.300735 | NO | NO | NA |
| Aptx | ENSMUSG( | ENSMUSG( | 0.164083 | 0.250201 | NO | NO | NA |
| Mpp2 | ENSMUSG( | ENSMUSG( | 0.281164 | 0.120832 | NO | NO | NA |
| Slc5a3 | ENSMUSG( | ENSMUSG( | -0.25029 | 0.101651 | NO | NO | NA |
| Tdrd3 | ENSMUSG( | ENSMUSG( | 0.066091 | 0.58553 | NO | NO | NA |
| St7 | ENSMUSG( | ENSMUSG( | 0.550554 | 1.99E-05 | YES | NO | NA |

| | | | | | | | |
|---|---|---|---|---|---|---|---|
| AI480526 | ENSMUSG( | ENSMUSG( | -0.32526 | 0.034211 | YES | NO | NA |
| Mcub | ENSMUSG( | ENSMUSG( | -4.65048 | 2.78E-166 | YES | YES | Downregulated |
| Timm21 | ENSMUSG( | ENSMUSG( | 0.161291 | 0.251026 | NO | NO | NA |
| Dnajb4 | ENSMUSG( | ENSMUSG( | -0.40429 | 0.001573 | YES | NO | NA |
| Rps6ka5 | ENSMUSG( | ENSMUSG( | -0.05598 | 0.703822 | NO | NO | NA |
| Hcfc1r1 | ENSMUSG( | ENSMUSG( | -0.08704 | 0.485963 | NO | NO | NA |
| 1110059E: | ENSMUSG( | ENSMUSG( | 0.153363 | 0.121345 | NO | NO | NA |
| Slc35d1 | ENSMUSG( | ENSMUSG( | -0.05502 | 0.6204 | NO | NO | NA |
| Slc6a20b | ENSMUSG( | ENSMUSG( | -0.02409 | 0.856147 | NO | NO | NA |
| Ank1 | ENSMUSG( | ENSMUSG( | 0.983595 | 0.066667 | NO | NO | NA |
| Tbccd1 | ENSMUSG( | ENSMUSG( | -0.05743 | 0.567556 | NO | NO | NA |
| Nup43 | ENSMUSG( | ENSMUSG( | 0.465288 | 2.50E-05 | YES | NO | NA |
| Alg9 | ENSMUSG( | ENSMUSG( | 0.218568 | 0.015641 | YES | NO | NA |
| Poglut3 | ENSMUSG( | ENSMUSG( | 0.470002 | 0.001248 | YES | NO | NA |
| Gm13456 | ENSMUSG( | ENSMUSG( | 0.2977 | 0.110814 | NO | NO | NA |
| Idua | ENSMUSG( | ENSMUSG( | -0.60941 | 2.18E-05 | YES | NO | NA |
| Spred1 | ENSMUSG( | ENSMUSG( | 0.362345 | 0.000359 | YES | NO | NA |
| Arap3 | ENSMUSG( | ENSMUSG( | -1.62326 | 0.000182 | YES | YES | Downregulated |
| 1110059G | ENSMUSG( | ENSMUSG( | -0.06943 | 0.624649 | NO | NO | NA |
| Cep41 | ENSMUSG( | ENSMUSG( | 0.05895 | 0.667184 | NO | NO | NA |
| Cd74 | ENSMUSG( | ENSMUSG( | 0.253234 | 0.502992 | NO | NO | NA |
| Zscan21 | ENSMUSG( | ENSMUSG( | 0.00163 | 0.991711 | NO | NO | NA |
| Txnrd2 | ENSMUSG( | ENSMUSG( | 0.467476 | 0.000179 | YES | NO | NA |
| Scoc | ENSMUSG( | ENSMUSG( | -0.15514 | 0.084384 | NO | NO | NA |
| Slc2a8 | ENSMUSG( | ENSMUSG( | 0.01305 | 0.939793 | NO | NO | NA |
| Zdhhc9 | ENSMUSG( | ENSMUSG( | -0.54369 | 9.49E-09 | YES | NO | NA |
| Caml | ENSMUSG( | ENSMUSG( | -0.14231 | 0.159583 | NO | NO | NA |
| Mast4 | ENSMUSG( | ENSMUSG( | 0.007467 | 0.964378 | NO | NO | NA |
| Pnkd | ENSMUSG( | ENSMUSG( | 0.152214 | 0.076751 | NO | NO | NA |
| Metap1d | ENSMUSG( | ENSMUSG( | 0.787003 | 7.32E-11 | YES | NO | NA |
| Eef1akmt2 | ENSMUSG( | ENSMUSG( | 0.013533 | 0.901916 | NO | NO | NA |
| Mrpl14 | ENSMUSG( | ENSMUSG( | -0.03007 | 0.784434 | NO | NO | NA |
| Unc5cl | ENSMUSG( | ENSMUSG( | -0.71576 | 0.001735 | YES | NO | NA |
| Rnf144a | ENSMUSG( | ENSMUSG( | -0.64184 | 9.12E-09 | YES | NO | NA |
| Ice2 | ENSMUSG( | ENSMUSG( | 0.124974 | 0.239269 | NO | NO | NA |
| Ripk2 | ENSMUSG( | ENSMUSG( | -0.17684 | 0.076535 | NO | NO | NA |
| Tmem237 | ENSMUSG( | ENSMUSG( | -0.01755 | 0.894744 | NO | NO | NA |
| Gm17300 | ENSMUSG( | ENSMUSG( | 0.648096 | 7.80E-05 | YES | NO | NA |
| Coq7 | ENSMUSG( | ENSMUSG( | 0.752468 | 1.08E-09 | YES | NO | NA |
| Dag1 | ENSMUSG( | ENSMUSG( | 0.297247 | 0.040689 | YES | NO | NA |
| Pex3 | ENSMUSG( | ENSMUSG( | 0.530691 | 2.73E-06 | YES | NO | NA |
| Cep19 | ENSMUSG( | ENSMUSG( | 0.393323 | 0.000145 | YES | NO | NA |
| Aktip | ENSMUSG( | ENSMUSG( | 0.042017 | 0.732555 | NO | NO | NA |
| Zfp39 | ENSMUSG( | ENSMUSG( | -0.37454 | 0.000283 | YES | NO | NA |
| Rab11fip2 | ENSMUSG( | ENSMUSG( | -0.09968 | 0.584747 | NO | NO | NA |
| Dhrs7b | ENSMUSG( | ENSMUSG( | 0.063548 | 0.545055 | NO | NO | NA |
| Ogfod2 | ENSMUSG( | ENSMUSG( | 0.104904 | 0.276373 | NO | NO | NA |

| | | | | | | | |
|---|---|---|---|---|---|---|---|
| Pde6d | ENSMUSG( | ENSMUSG( | -0.414 | 1.49E-06 | YES | NO | NA |
| Fadd | ENSMUSG( | ENSMUSG( | 0.27173 | 0.011742 | YES | NO | NA |
| Qrfp | ENSMUSG( | ENSMUSG( | 1.499578 | 4.15E-11 | YES | YES | Upregulated |
| Acp2 | ENSMUSG( | ENSMUSG( | 0.073232 | 0.633247 | NO | NO | NA |
| Slc35a1 | ENSMUSG( | ENSMUSG( | 0.090709 | 0.516453 | NO | NO | NA |
| 9030025P: | ENSMUSG( | ENSMUSG( | -0.20172 | 0.280824 | NO | NO | NA |
| E330020D | ENSMUSG( | ENSMUSG( | -0.0994 | 0.626538 | NO | NO | NA |
| Pgap3 | ENSMUSG( | ENSMUSG( | -0.00061 | 0.995654 | NO | NO | NA |
| Fbf1 | ENSMUSG( | ENSMUSG( | -0.08194 | 0.600992 | NO | NO | NA |
| Samd8 | ENSMUSG( | ENSMUSG( | -0.09599 | 0.453495 | NO | NO | NA |
| 2610005L( | ENSMUSG( | ENSMUSG( | 0.117022 | 0.580585 | NO | NO | NA |
| Rnf185 | ENSMUSG( | ENSMUSG( | -0.02007 | 0.859851 | NO | NO | NA |
| Slx4ip | ENSMUSG( | ENSMUSG( | -0.36745 | 0.008603 | YES | NO | NA |
| Stard5 | ENSMUSG( | ENSMUSG( | -0.20402 | 0.013326 | YES | NO | NA |
| Btd | ENSMUSG( | ENSMUSG( | -0.21251 | 0.002982 | YES | NO | NA |
| Immp1l | ENSMUSG( | ENSMUSG( | -0.27045 | 0.003361 | YES | NO | NA |
| Mcm8 | ENSMUSG( | ENSMUSG( | 0.249729 | 0.078883 | NO | NO | NA |
| Mutyh | ENSMUSG( | ENSMUSG( | 0.227703 | 0.072796 | NO | NO | NA |
| Fntb | ENSMUSG( | ENSMUSG( | 0.138489 | 0.206239 | NO | NO | NA |
| Sertad1 | ENSMUSG( | ENSMUSG( | -0.06198 | 0.664181 | NO | NO | NA |
| Grb7 | ENSMUSG( | ENSMUSG( | 0.26325 | 0.187474 | NO | NO | NA |
| Rcn3 | ENSMUSG( | ENSMUSG( | -0.60392 | 0.000128 | YES | NO | NA |
| Gm10130 | ENSMUSG( | ENSMUSG( | -0.80278 | 2.72E-05 | YES | NO | NA |
| Cep83os | ENSMUSG( | ENSMUSG( | 0.474685 | 0.011581 | YES | NO | NA |
| Reck | ENSMUSG( | ENSMUSG( | 0.117607 | 0.288771 | NO | NO | NA |
| Vav3 | ENSMUSG( | ENSMUSG( | -0.29254 | 0.04111 | YES | NO | NA |
| Fancl | ENSMUSG( | ENSMUSG( | 0.099253 | 0.288861 | NO | NO | NA |
| Aqp1 | ENSMUSG( | ENSMUSG( | 0.971659 | 0.126934 | NO | NO | NA |
| Gm10182 | ENSMUSG( | ENSMUSG( | 0.405232 | 0.006963 | YES | NO | NA |
| Akt1s1 | ENSMUSG( | ENSMUSG( | 0.190043 | 0.052956 | NO | NO | NA |
| Zfyve16 | ENSMUSG( | ENSMUSG( | -0.35227 | 0.000373 | YES | NO | NA |
| Dcakd | ENSMUSG( | ENSMUSG( | -0.05341 | 0.596956 | NO | NO | NA |
| Acad8 | ENSMUSG( | ENSMUSG( | 0.155377 | 0.353555 | NO | NO | NA |
| Rab40c | ENSMUSG( | ENSMUSG( | -0.2729 | 0.001196 | YES | NO | NA |
| Armcx5 | ENSMUSG( | ENSMUSG( | 0.163541 | 0.374748 | NO | NO | NA |
| Gm17134 | ENSMUSG( | ENSMUSG( | 0.105004 | 0.352248 | NO | NO | NA |
| Ddit3 | ENSMUSG( | ENSMUSG( | -0.22703 | 0.191184 | NO | NO | NA |
| Rbm41 | ENSMUSG( | ENSMUSG( | -0.26993 | 0.012657 | YES | NO | NA |
| Ccdc137 | ENSMUSG( | ENSMUSG( | -0.19026 | 0.054263 | NO | NO | NA |
| Arl3 | ENSMUSG( | ENSMUSG( | 0.155244 | 0.407673 | NO | NO | NA |
| Rad1 | ENSMUSG( | ENSMUSG( | -0.04643 | 0.628563 | NO | NO | NA |
| Slc46a3 | ENSMUSG( | ENSMUSG( | -0.3421 | 0.022491 | YES | NO | NA |
| Cd99l2 | ENSMUSG( | ENSMUSG( | -0.17437 | 0.096745 | NO | NO | NA |
| Penk | ENSMUSG( | ENSMUSG( | 2.00982 | 7.99E-13 | YES | YES | Upregulated |
| Ddx31 | ENSMUSG( | ENSMUSG( | -0.06544 | 0.518438 | NO | NO | NA |
| Fam118b | ENSMUSG( | ENSMUSG( | -0.11804 | 0.176575 | NO | NO | NA |
| Zmym1 | ENSMUSG( | ENSMUSG( | 0.090744 | 0.403656 | NO | NO | NA |

| | | | | | | | |
|---|---|---|---|---|---|---|---|
| Fez2 | ENSMUSG( | ENSMUSG( | 0.004761 | 0.970821 | NO | NO | NA |
| Clip2 | ENSMUSG( | ENSMUSG( | -0.3232 | 0.00097 | YES | NO | NA |
| Trav16 | ENSMUSG( | ENSMUSG( | -0.75246 | 0.05844 | NO | NO | NA |
| Cep44 | ENSMUSG( | ENSMUSG( | 0.045994 | 0.706841 | NO | NO | NA |
| Xpnpep3 | ENSMUSG( | ENSMUSG( | -0.05305 | 0.613051 | NO | NO | NA |
| Dph6 | ENSMUSG( | ENSMUSG( | 0.148411 | 0.173244 | NO | NO | NA |
| Gk | ENSMUSG( | ENSMUSG( | 0.402878 | 0.000219 | YES | NO | NA |
| Bbs9 | ENSMUSG( | ENSMUSG( | 0.220581 | 0.352352 | NO | NO | NA |
| Tmem251 | ENSMUSG( | ENSMUSG( | 0.305149 | 0.00269 | YES | NO | NA |
| Tsr2 | ENSMUSG( | ENSMUSG( | 0.349809 | 0.00502 | YES | NO | NA |
| Chchd4 | ENSMUSG( | ENSMUSG( | 0.520021 | 1.38E-05 | YES | NO | NA |
| Zswim1 | ENSMUSG( | ENSMUSG( | 0.052664 | 0.561653 | NO | NO | NA |
| Necap1 | ENSMUSG( | ENSMUSG( | -0.29252 | 0.018944 | YES | NO | NA |
| Zscan25 | ENSMUSG( | ENSMUSG( | 0.084822 | 0.454706 | NO | NO | NA |
| Asb7 | ENSMUSG( | ENSMUSG( | -0.03911 | 0.805041 | NO | NO | NA |
| Ticam1 | ENSMUSG( | ENSMUSG( | -0.16555 | 0.064027 | NO | NO | NA |
| Tbc1d25 | ENSMUSG( | ENSMUSG( | -0.20447 | 0.002306 | YES | NO | NA |
| Kbtbd7 | ENSMUSG( | ENSMUSG( | -0.15746 | 0.329531 | NO | NO | NA |
| Dse | ENSMUSG( | ENSMUSG( | 0.36511 | 0.023396 | YES | NO | NA |
| Bcorl1 | ENSMUSG( | ENSMUSG( | -0.82948 | 1.18E-12 | YES | NO | NA |
| Atpsckmt | ENSMUSG( | ENSMUSG( | -0.03429 | 0.802185 | NO | NO | NA |
| Taf6l | ENSMUSG( | ENSMUSG( | 0.358493 | 0.000207 | YES | NO | NA |
| Gm28875 | ENSMUSG( | ENSMUSG( | 0.066184 | 0.624805 | NO | NO | NA |
| Capn10 | ENSMUSG( | ENSMUSG( | 0.237336 | 0.071853 | NO | NO | NA |
| Nanos1 | ENSMUSG( | ENSMUSG( | -0.23055 | 0.094921 | NO | NO | NA |
| Pbx3 | ENSMUSG( | ENSMUSG( | 0.04538 | 0.709959 | NO | NO | NA |
| Zfp758 | ENSMUSG( | ENSMUSG( | -0.51119 | 0.000254 | YES | NO | NA |
| Tmem185a | ENSMUSG( | ENSMUSG( | 0.015713 | 0.912074 | NO | NO | NA |
| Oasl1 | ENSMUSG( | ENSMUSG( | -1.06323 | 4.57E-07 | YES | YES | Downregulated |
| Alg12 | ENSMUSG( | ENSMUSG( | -0.34308 | 7.86E-05 | YES | NO | NA |
| Ptpmt1 | ENSMUSG( | ENSMUSG( | 0.683024 | 1.25E-06 | YES | NO | NA |
| Zfp738 | ENSMUSG( | ENSMUSG( | -0.24429 | 0.047379 | YES | NO | NA |
| Myo9a | ENSMUSG( | ENSMUSG( | -0.27513 | 0.004054 | YES | NO | NA |
| Gm26885 | ENSMUSG( | ENSMUSG( | 0.455349 | 0.019048 | YES | NO | NA |
| Ndrg1 | ENSMUSG( | ENSMUSG( | -0.06246 | 0.70182 | NO | NO | NA |
| Tnfrsf14 | ENSMUSG( | ENSMUSG( | -0.4805 | 0.000247 | YES | NO | NA |
| Dennd6b | ENSMUSG( | ENSMUSG( | 0.342426 | 0.000175 | YES | NO | NA |
| Zfp87 | ENSMUSG( | ENSMUSG( | -0.5415 | 0.000103 | YES | NO | NA |
| H2-DMa | ENSMUSG( | ENSMUSG( | 0.032335 | 0.810257 | NO | NO | NA |
| Zfp101 | ENSMUSG( | ENSMUSG( | 0.045053 | 0.686039 | NO | NO | NA |
| Bend3 | ENSMUSG( | ENSMUSG( | 0.354708 | 0.013079 | YES | NO | NA |
| Ippk | ENSMUSG( | ENSMUSG( | 0.069443 | 0.619035 | NO | NO | NA |
| Smim11 | ENSMUSG( | ENSMUSG( | -0.01193 | 0.939814 | NO | NO | NA |
| Nme3 | ENSMUSG( | ENSMUSG( | -0.14667 | 0.155729 | NO | NO | NA |
| Dclre1a | ENSMUSG( | ENSMUSG( | 0.095797 | 0.460937 | NO | NO | NA |
| Serp2 | ENSMUSG( | ENSMUSG( | -0.40618 | 0.028701 | YES | NO | NA |
| Xxylt1 | ENSMUSG( | ENSMUSG( | 0.458487 | 0.00018 | YES | NO | NA |

| | | | | | | | |
|---|---|---|---|---|---|---|---|
| Pafah1b3 | ENSMUSG( | ENSMUSG( | 0.329227 | 0.003429 | YES | NO | NA |
| Dtd2 | ENSMUSG( | ENSMUSG( | 0.483458 | 2.99E-06 | YES | NO | NA |
| Eml5 | ENSMUSG( | ENSMUSG( | 0.089391 | 0.463646 | NO | NO | NA |
| Entpd7 | ENSMUSG( | ENSMUSG( | 0.068916 | 0.422216 | NO | NO | NA |
| Erbb3 | ENSMUSG( | ENSMUSG( | -0.00371 | 0.990647 | NO | NO | NA |
| Mbd5 | ENSMUSG( | ENSMUSG( | -0.20659 | 0.117621 | NO | NO | NA |
| Bad | ENSMUSG( | ENSMUSG( | 0.147061 | 0.226749 | NO | NO | NA |
| Timp2 | ENSMUSG( | ENSMUSG( | -0.44223 | 0.003302 | YES | NO | NA |
| Coq6 | ENSMUSG( | ENSMUSG( | 0.133544 | 0.137186 | NO | NO | NA |
| Pvr | ENSMUSG( | ENSMUSG( | 0.082438 | 0.587432 | NO | NO | NA |
| Zmym6 | ENSMUSG( | ENSMUSG( | -0.05395 | 0.733315 | NO | NO | NA |
| Wdr41 | ENSMUSG( | ENSMUSG( | 0.181237 | 0.207559 | NO | NO | NA |
| Mettl21a | ENSMUSG( | ENSMUSG( | 0.129168 | 0.442367 | NO | NO | NA |
| Sesn1 | ENSMUSG( | ENSMUSG( | -0.36496 | 0.002956 | YES | NO | NA |
| Rfx3 | ENSMUSG( | ENSMUSG( | -0.41522 | 6.39E-06 | YES | NO | NA |
| Coq8b | ENSMUSG( | ENSMUSG( | 0.010537 | 0.931584 | NO | NO | NA |
| Nmrk1 | ENSMUSG( | ENSMUSG( | 0.610741 | 4.52E-05 | YES | NO | NA |
| Alg2 | ENSMUSG( | ENSMUSG( | -0.40656 | 0.000364 | YES | NO | NA |
| F2rl2 | ENSMUSG( | ENSMUSG( | -0.79705 | 1.00E-08 | YES | NO | NA |
| Cep70 | ENSMUSG( | ENSMUSG( | 0.085415 | 0.520986 | NO | NO | NA |
| Gm14569 | ENSMUSG( | ENSMUSG( | 0.713577 | 0.014751 | YES | NO | NA |
| Abhd11 | ENSMUSG( | ENSMUSG( | 0.241764 | 0.061382 | NO | NO | NA |
| Exog | ENSMUSG( | ENSMUSG( | -0.23021 | 0.031651 | YES | NO | NA |
| Diaph2 | ENSMUSG( | ENSMUSG( | 0.206532 | 0.219544 | NO | NO | NA |
| Gadd45g | ENSMUSG( | ENSMUSG( | 0.939874 | 3.56E-08 | YES | NO | NA |
| Alg3 | ENSMUSG( | ENSMUSG( | 0.107377 | 0.484318 | NO | NO | NA |
| Tusc2 | ENSMUSG( | ENSMUSG( | -0.15493 | 0.184246 | NO | NO | NA |
| Gon7 | ENSMUSG( | ENSMUSG( | 0.003684 | 0.976657 | NO | NO | NA |
| Tmcc1 | ENSMUSG( | ENSMUSG( | 0.186098 | 0.122009 | NO | NO | NA |
| Prepl | ENSMUSG( | ENSMUSG( | -0.33751 | 0.004954 | YES | NO | NA |
| ENSMUSG( | ENSMUSG( | ENSMUSG( | 0.363778 | 0.005306 | YES | NO | NA |
| Chid1 | ENSMUSG( | ENSMUSG( | -0.16637 | 0.268944 | NO | NO | NA |
| Gtdc1 | ENSMUSG( | ENSMUSG( | -0.20252 | 0.183146 | NO | NO | NA |
| Znrd2 | ENSMUSG( | ENSMUSG( | 0.510399 | 0.000953 | YES | NO | NA |
| Kdm8 | ENSMUSG( | ENSMUSG( | 0.279271 | 0.019753 | YES | NO | NA |
| Fam219a | ENSMUSG( | ENSMUSG( | -0.17671 | 0.356564 | NO | NO | NA |
| Bag3 | ENSMUSG( | ENSMUSG( | -0.19018 | 0.147081 | NO | NO | NA |
| Abhd18 | ENSMUSG( | ENSMUSG( | -0.11128 | 0.362672 | NO | NO | NA |
| Mboat1 | ENSMUSG( | ENSMUSG( | 0.316772 | 0.000554 | YES | NO | NA |
| Arhgef4 | ENSMUSG( | ENSMUSG( | -0.28847 | 0.020108 | YES | NO | NA |
| Ttll3 | ENSMUSG( | ENSMUSG( | -0.43817 | 0.000127 | YES | NO | NA |
| Fam118a | ENSMUSG( | ENSMUSG( | -0.11412 | 0.332677 | NO | NO | NA |
| Mrps28 | ENSMUSG( | ENSMUSG( | 0.577535 | 0.000153 | YES | NO | NA |
| Apool | ENSMUSG( | ENSMUSG( | 0.022416 | 0.882467 | NO | NO | NA |
| Kansl1l | ENSMUSG( | ENSMUSG( | -0.337 | 0.034985 | YES | NO | NA |
| Fdx2 | ENSMUSG( | ENSMUSG( | 0.642883 | 3.85E-06 | YES | NO | NA |
| Pi4k2b | ENSMUSG( | ENSMUSG( | 0.390263 | 0.003863 | YES | NO | NA |

| | | | | | | | |
|---|---|---|---|---|---|---|---|
| Cdin1 | ENSMUSG( | ENSMUSG( | -0.13705 | 0.294351 | NO | NO | NA |
| Chek2 | ENSMUSG( | ENSMUSG( | 0.057092 | 0.643308 | NO | NO | NA |
| Psmg4 | ENSMUSG( | ENSMUSG( | 0.35082 | 0.000439 | YES | NO | NA |
| Dnaaf10 | ENSMUSG( | ENSMUSG( | 0.130654 | 0.291794 | NO | NO | NA |
| Mpzl3 | ENSMUSG( | ENSMUSG( | 0.363738 | 0.009925 | YES | NO | NA |
| Med21 | ENSMUSG( | ENSMUSG( | -0.00103 | 0.992674 | NO | NO | NA |
| Cox11 | ENSMUSG( | ENSMUSG( | 0.115154 | 0.323235 | NO | NO | NA |
| Zbtb48 | ENSMUSG( | ENSMUSG( | -0.13181 | 0.279303 | NO | NO | NA |
| Dram2 | ENSMUSG( | ENSMUSG( | -0.24058 | 0.046421 | YES | NO | NA |
| Nme7 | ENSMUSG( | ENSMUSG( | 0.331428 | 0.001802 | YES | NO | NA |
| Sdc3 | ENSMUSG( | ENSMUSG( | -1.59324 | 7.07E-12 | YES | YES | Downregulated |
| Arhgap35 | ENSMUSG( | ENSMUSG( | -0.00212 | 0.987651 | NO | NO | NA |
| Slc39a11 | ENSMUSG( | ENSMUSG( | 0.452252 | 0.007102 | YES | NO | NA |
| Thoc2l | ENSMUSG( | ENSMUSG( | 0.019047 | 0.819148 | NO | NO | NA |
| Mrpl47 | ENSMUSG( | ENSMUSG( | 0.24773 | 0.03373 | YES | NO | NA |
| Carns1 | ENSMUSG( | ENSMUSG( | 0.020529 | 0.894999 | NO | NO | NA |
| Tmem199 | ENSMUSG( | ENSMUSG( | 0.0228 | 0.832038 | NO | NO | NA |
| B3gntl1 | ENSMUSG( | ENSMUSG( | -0.10234 | 0.432927 | NO | NO | NA |
| Stk38l | ENSMUSG( | ENSMUSG( | 0.268639 | 3.47E-05 | YES | NO | NA |
| Zfp672 | ENSMUSG( | ENSMUSG( | -0.29297 | 0.000169 | YES | NO | NA |
| Phf7 | ENSMUSG( | ENSMUSG( | 0.354864 | 0.006454 | YES | NO | NA |
| Zfp790 | ENSMUSG( | ENSMUSG( | -0.30938 | 0.00522 | YES | NO | NA |
| Ccdc14 | ENSMUSG( | ENSMUSG( | 0.444619 | 0.000272 | YES | NO | NA |
| Tcf4 | ENSMUSG( | ENSMUSG( | 0.588537 | 0.000824 | YES | NO | NA |
| Naip5 | ENSMUSG( | ENSMUSG( | -0.55232 | 0.001532 | YES | NO | NA |
| Kcnk6 | ENSMUSG( | ENSMUSG( | -0.49751 | 7.86E-07 | YES | NO | NA |
| 9930104L( | ENSMUSG( | ENSMUSG( | -0.16262 | 0.261847 | NO | NO | NA |
| Tmem223 | ENSMUSG( | ENSMUSG( | 0.143704 | 0.222797 | NO | NO | NA |
| Zfp472 | ENSMUSG( | ENSMUSG( | -0.15816 | 0.162335 | NO | NO | NA |
| Sac3d1 | ENSMUSG( | ENSMUSG( | 0.198453 | 0.028609 | YES | NO | NA |
| Klhdc1 | ENSMUSG( | ENSMUSG( | -0.26843 | 0.106197 | NO | NO | NA |
| Fbxl2 | ENSMUSG( | ENSMUSG( | -0.49162 | 1.50E-06 | YES | NO | NA |
| Cd81 | ENSMUSG( | ENSMUSG( | 1.011784 | 0.000107 | YES | YES | Upregulated |
| Thap2 | ENSMUSG( | ENSMUSG( | -0.62777 | 3.71E-05 | YES | NO | NA |
| Srrd | ENSMUSG( | ENSMUSG( | -0.08011 | 0.430927 | NO | NO | NA |
| Gm45221 | ENSMUSG( | ENSMUSG( | -0.24662 | 0.2351 | NO | NO | NA |
| Rft1 | ENSMUSG( | ENSMUSG( | 0.176522 | 0.185287 | NO | NO | NA |
| AA467197 | ENSMUSG( | ENSMUSG( | 1.416565 | 5.36E-07 | YES | YES | Upregulated |
| Ccdc18 | ENSMUSG( | ENSMUSG( | 0.233275 | 0.309567 | NO | NO | NA |
| Gm53012 | ENSMUSG( | ENSMUSG( | -0.17948 | 0.18584 | NO | NO | NA |
| Fut7 | ENSMUSG( | ENSMUSG( | 0.79432 | 2.81E-06 | YES | NO | NA |
| Gm26782 | ENSMUSG( | ENSMUSG( | 0.076142 | 0.554579 | NO | NO | NA |
| ENSMUSG( | ENSMUSG( | ENSMUSG( | 0.141936 | 0.143834 | NO | NO | NA |
| Slc35f6 | ENSMUSG( | ENSMUSG( | 0.014812 | 0.903282 | NO | NO | NA |
| Gm46430 | ENSMUSG( | ENSMUSG( | 0.100876 | 0.387184 | NO | NO | NA |
| 9830144P: | ENSMUSG( | ENSMUSG( | -0.55747 | 2.19E-06 | YES | NO | NA |
| Twnk | ENSMUSG( | ENSMUSG( | 0.064979 | 0.637146 | NO | NO | NA |

| | | | | | | | |
|---|---|---|---|---|---|---|---|
| Mindy1 | ENSMUSG( | ENSMUSG( | -0.35212 | 0.000568 | YES | NO | NA |
| Sco1 | ENSMUSG( | ENSMUSG( | 0.06922 | 0.487655 | NO | NO | NA |
| Cables1 | ENSMUSG( | ENSMUSG( | 0.868001 | 0.000517 | YES | NO | NA |
| Cmss1 | ENSMUSG( | ENSMUSG( | 0.890717 | 9.44E-11 | YES | NO | NA |
| Tmppe | ENSMUSG( | ENSMUSG( | -0.33829 | 0.011026 | YES | NO | NA |
| Spats2 | ENSMUSG( | ENSMUSG( | 0.491952 | 3.28E-05 | YES | NO | NA |
| Casz1 | ENSMUSG( | ENSMUSG( | -0.49184 | 0.004034 | YES | NO | NA |
| Lrrc20 | ENSMUSG( | ENSMUSG( | 0.125387 | 0.22328 | NO | NO | NA |
| AU022252 | ENSMUSG( | ENSMUSG( | 0.000901 | 0.992393 | NO | NO | NA |
| Lman2l | ENSMUSG( | ENSMUSG( | -0.29589 | 0.054786 | NO | NO | NA |
| Mief2 | ENSMUSG( | ENSMUSG( | -0.10651 | 0.461948 | NO | NO | NA |
| Shld2 | ENSMUSG( | ENSMUSG( | 0.081599 | 0.585511 | NO | NO | NA |
| Aars2 | ENSMUSG( | ENSMUSG( | 0.330787 | 0.006674 | YES | NO | NA |
| Nek1 | ENSMUSG( | ENSMUSG( | 0.313397 | 0.050939 | NO | NO | NA |
| Srek1ip1 | ENSMUSG( | ENSMUSG( | 0.014199 | 0.921962 | NO | NO | NA |
| Mettl1 | ENSMUSG( | ENSMUSG( | 0.877933 | 1.66E-10 | YES | NO | NA |
| Lmf1 | ENSMUSG( | ENSMUSG( | -0.27044 | 0.002528 | YES | NO | NA |
| Yif1a | ENSMUSG( | ENSMUSG( | 0.117377 | 0.402567 | NO | NO | NA |
| Mgme1 | ENSMUSG( | ENSMUSG( | 0.15335 | 0.167959 | NO | NO | NA |
| Smim19 | ENSMUSG( | ENSMUSG( | -0.15861 | 0.18346 | NO | NO | NA |
| Prkcz | ENSMUSG( | ENSMUSG( | -0.90912 | 2.37E-06 | YES | NO | NA |
| ENSMUSG( | ENSMUSG( | ENSMUSG( | -0.49937 | 0.881796 | NO | NO | NA |
| Gm17745 | ENSMUSG( | ENSMUSG( | 0.900936 | 0.000129 | YES | NO | NA |
| Sumf2 | ENSMUSG( | ENSMUSG( | 0.097293 | 0.353524 | NO | NO | NA |
| Dph7 | ENSMUSG( | ENSMUSG( | 0.17142 | 0.1501 | NO | NO | NA |
| Slc19a2 | ENSMUSG( | ENSMUSG( | 0.698392 | 5.11E-09 | YES | NO | NA |
| Cdk10 | ENSMUSG( | ENSMUSG( | -0.27151 | 0.017178 | YES | NO | NA |
| Ulk4 | ENSMUSG( | ENSMUSG( | 0.096066 | 0.610282 | NO | NO | NA |
| Dnajc17 | ENSMUSG( | ENSMUSG( | 0.273855 | 0.020347 | YES | NO | NA |
| Cbx8 | ENSMUSG( | ENSMUSG( | -0.23579 | 0.070951 | NO | NO | NA |
| Oas1c | ENSMUSG( | ENSMUSG( | -0.36851 | 0.008803 | YES | NO | NA |
| Lrp5 | ENSMUSG( | ENSMUSG( | 0.972885 | 1.99E-08 | YES | NO | NA |
| Vezt | ENSMUSG( | ENSMUSG( | 0.031544 | 0.828882 | NO | NO | NA |
| Smyd4 | ENSMUSG( | ENSMUSG( | -0.15276 | 0.053541 | NO | NO | NA |
| Fastkd2 | ENSMUSG( | ENSMUSG( | 0.307068 | 0.002991 | YES | NO | NA |
| Trav16n | ENSMUSG( | ENSMUSG( | -0.2565 | 0.607428 | NO | NO | NA |
| Klrb1b | ENSMUSG( | ENSMUSG( | 1.32695 | 5.21E-07 | YES | YES | Upregulated |
| Mthfs | ENSMUSG( | ENSMUSG( | -0.22242 | 0.081641 | NO | NO | NA |
| Gm50394 | ENSMUSG( | ENSMUSG( | 0.026076 | 0.896584 | NO | NO | NA |
| Gatc | ENSMUSG( | ENSMUSG( | 0.024357 | 0.824442 | NO | NO | NA |
| Slc16a11 | ENSMUSG( | ENSMUSG( | 0.847226 | 1.62E-06 | YES | NO | NA |
| Ighg3 | ENSMUSG( | ENSMUSG( | -2.15937 | 0.150531 | NO | NO | NA |
| Rps19bp1 | ENSMUSG( | ENSMUSG( | 0.310122 | 0.002347 | YES | NO | NA |
| ENSMUSG | ENSMUSG( | ENSMUSG( | -0.9064 | 1.69E-18 | YES | NO | NA |
| Mettl5 | ENSMUSG( | ENSMUSG( | 0.053721 | 0.614855 | NO | NO | NA |
| Gemin4 | ENSMUSG( | ENSMUSG( | 0.265723 | 0.171026 | NO | NO | NA |
| Arfgap3 | ENSMUSG( | ENSMUSG( | -0.32077 | 0.00051 | YES | NO | NA |

| | | | | | | | |
|---|---|---|---|---|---|---|---|
| Pus7l | ENSMUSG( | ENSMUSG( | 0.456113 | 1.10E-05 | YES | NO | NA |
| Gaa | ENSMUSG( | ENSMUSG( | 0.175653 | 0.115387 | NO | NO | NA |
| Crtap | ENSMUSG( | ENSMUSG( | 0.311331 | 0.007246 | YES | NO | NA |
| Jrkl | ENSMUSG( | ENSMUSG( | -0.27782 | 0.017601 | YES | NO | NA |
| Chchd10 | ENSMUSG( | ENSMUSG( | 0.537158 | 8.85E-05 | YES | NO | NA |
| Prkce | ENSMUSG( | ENSMUSG( | 0.272469 | 0.070757 | NO | NO | NA |
| Trib1 | ENSMUSG( | ENSMUSG( | 0.245454 | 0.228978 | NO | NO | NA |
| Rundc3b | ENSMUSG( | ENSMUSG( | -0.04408 | 0.74914 | NO | NO | NA |
| Gpc1 | ENSMUSG( | ENSMUSG( | 0.519536 | 0.000971 | YES | NO | NA |
| Pla2g6 | ENSMUSG( | ENSMUSG( | 0.077707 | 0.512053 | NO | NO | NA |
| Klhl11 | ENSMUSG( | ENSMUSG( | -0.20871 | 0.071198 | NO | NO | NA |
| Fastkd3 | ENSMUSG( | ENSMUSG( | 0.106559 | 0.336818 | NO | NO | NA |
| Gm49553 | ENSMUSG( | ENSMUSG( | -0.57316 | 2.56E-07 | YES | NO | NA |
| Olfr60 | ENSMUSG( | ENSMUSG( | 1.5573 | 0.002061 | YES | YES | Upregulated |
| Gtpbp8 | ENSMUSG( | ENSMUSG( | 0.029521 | 0.819022 | NO | NO | NA |
| Pvt1 | ENSMUSG( | ENSMUSG( | 0.03606 | 0.772214 | NO | NO | NA |
| Susd3 | ENSMUSG( | ENSMUSG( | -0.27553 | 0.016495 | YES | NO | NA |
| Pcca | ENSMUSG( | ENSMUSG( | 0.60037 | 9.46E-05 | YES | NO | NA |
| Mpp7 | ENSMUSG( | ENSMUSG( | 0.027032 | 0.858038 | NO | NO | NA |
| Rab23 | ENSMUSG( | ENSMUSG( | -0.02048 | 0.863539 | NO | NO | NA |
| St3gal2 | ENSMUSG( | ENSMUSG( | 0.002707 | 0.983029 | NO | NO | NA |
| Tecpr2 | ENSMUSG( | ENSMUSG( | 0.160289 | 0.057689 | NO | NO | NA |
| Gm20604 | ENSMUSG( | ENSMUSG( | 0.268445 | 0.014898 | YES | NO | NA |
| Phldb3 | ENSMUSG( | ENSMUSG( | -0.67888 | 5.58E-09 | YES | NO | NA |
| Ocrl | ENSMUSG( | ENSMUSG( | 0.445277 | 7.62E-07 | YES | NO | NA |
| Mtg1 | ENSMUSG( | ENSMUSG( | 0.423483 | 0.000432 | YES | NO | NA |
| Zfp952 | ENSMUSG( | ENSMUSG( | -0.40038 | 0.002336 | YES | NO | NA |
| Jmy | ENSMUSG( | ENSMUSG( | 0.066721 | 0.580769 | NO | NO | NA |
| Ankrd23 | ENSMUSG( | ENSMUSG( | -0.01919 | 0.861496 | NO | NO | NA |
| Gm46223 | ENSMUSG( | ENSMUSG( | 1.207059 | 2.88E-06 | YES | YES | Upregulated |
| Ftx | ENSMUSG( | ENSMUSG( | 0.058333 | 0.617485 | NO | NO | NA |
| Spring1 | ENSMUSG( | ENSMUSG( | -0.05534 | 0.712884 | NO | NO | NA |
| Abhd4 | ENSMUSG( | ENSMUSG( | 0.603257 | 0.001686 | YES | NO | NA |
| Ggact | ENSMUSG( | ENSMUSG( | 0.503591 | 0.00031 | YES | NO | NA |
| Rnf25 | ENSMUSG( | ENSMUSG( | -0.36432 | 0.003199 | YES | NO | NA |
| Gm47469 | ENSMUSG( | ENSMUSG( | -0.28137 | 0.051002 | NO | NO | NA |
| Prr7 | ENSMUSG( | ENSMUSG( | 0.232005 | 0.131129 | NO | NO | NA |
| Slc22a5 | ENSMUSG( | ENSMUSG( | 0.064054 | 0.672518 | NO | NO | NA |
| Naip6 | ENSMUSG( | ENSMUSG( | -0.32942 | 0.037168 | YES | NO | NA |
| Apol11b | ENSMUSG( | ENSMUSG( | 0.956001 | 0.068295 | NO | NO | NA |
| 9130023H | ENSMUSG( | ENSMUSG( | -0.26917 | 0.00772 | YES | NO | NA |
| Ndufaf3 | ENSMUSG( | ENSMUSG( | -0.11337 | 0.328293 | NO | NO | NA |
| Kif5c | ENSMUSG( | ENSMUSG( | -0.59224 | 0.000355 | YES | NO | NA |
| Mob3c | ENSMUSG( | ENSMUSG( | -0.16191 | 0.309679 | NO | NO | NA |
| Fas | ENSMUSG( | ENSMUSG( | 0.022597 | 0.901916 | NO | NO | NA |
| Smim24 | ENSMUSG( | ENSMUSG( | -0.86549 | 0.000373 | YES | NO | NA |
| Pdhx | ENSMUSG( | ENSMUSG( | 0.417669 | 5.21E-05 | YES | NO | NA |

| | | | | | | | |
|---|---|---|---|---|---|---|---|
| Cptp | ENSMUSG( | ENSMUSG( | 0.044525 | 0.665553 | NO | NO | NA |
| Scrn3 | ENSMUSG( | ENSMUSG( | -0.12033 | 0.159012 | NO | NO | NA |
| Nnt ENSM | ENSMUSG( | ENSMUSG( | -0.15681 | 0.926979 | NO | NO | NA |
| Mrpl22 | ENSMUSG( | ENSMUSG( | 0.295436 | 0.001977 | YES | NO | NA |
| ENSMUSG( | ENSMUSG( | ENSMUSG( | 0.463719 | 1.55E-05 | YES | NO | NA |
| H2bc8 | ENSMUSG( | ENSMUSG( | 0.43818 | 0.010048 | YES | NO | NA |
| Ccnyl1 | ENSMUSG( | ENSMUSG( | 0.221787 | 0.087247 | NO | NO | NA |
| Arl15 | ENSMUSG( | ENSMUSG( | -0.35417 | 0.00041 | YES | NO | NA |
| Mrpl53 | ENSMUSG( | ENSMUSG( | -0.07255 | 0.555728 | NO | NO | NA |
| Smim12 | ENSMUSG( | ENSMUSG( | 0.062698 | 0.572312 | NO | NO | NA |
| Disp1 | ENSMUSG( | ENSMUSG( | -0.22202 | 0.01513 | YES | NO | NA |
| Tm2d3 | ENSMUSG( | ENSMUSG( | -0.06992 | 0.453757 | NO | NO | NA |
| Hs6st1 | ENSMUSG( | ENSMUSG( | -0.12053 | 0.188866 | NO | NO | NA |
| Plod3 | ENSMUSG( | ENSMUSG( | 0.114482 | 0.360428 | NO | NO | NA |
| Wdr89 | ENSMUSG( | ENSMUSG( | 0.267259 | 0.015368 | YES | NO | NA |
| 5530601H | ENSMUSG( | ENSMUSG( | -0.05996 | 0.623853 | NO | NO | NA |
| Elp4 | ENSMUSG( | ENSMUSG( | 0.232092 | 0.007735 | YES | NO | NA |
| Egr2 | ENSMUSG( | ENSMUSG( | 0.574694 | 0.00876 | YES | NO | NA |
| Maip1 | ENSMUSG( | ENSMUSG( | 0.268287 | 0.007016 | YES | NO | NA |
| Pet100 | ENSMUSG( | ENSMUSG( | -0.0343 | 0.785685 | NO | NO | NA |
| Vsig10l | ENSMUSG( | ENSMUSG( | -0.5005 | 0.000207 | YES | NO | NA |
| Acbd4 | ENSMUSG( | ENSMUSG( | -0.04506 | 0.735932 | NO | NO | NA |
| Gm45552 | ENSMUSG( | ENSMUSG( | -0.44953 | 0.002906 | YES | NO | NA |
| Palm3 | ENSMUSG( | ENSMUSG( | -0.29542 | 0.022763 | YES | NO | NA |
| Pip4p2 | ENSMUSG( | ENSMUSG( | 0.154389 | 0.407389 | NO | NO | NA |
| Zbtb34 | ENSMUSG( | ENSMUSG( | -0.42681 | 4.71E-07 | YES | NO | NA |
| Pcdhgc4 | ENSMUSG( | ENSMUSG( | -0.40787 | 0.009565 | YES | NO | NA |
| Fam216a | ENSMUSG( | ENSMUSG( | 0.582924 | 6.56E-05 | YES | NO | NA |
| Polg2 | ENSMUSG( | ENSMUSG( | 0.213414 | 0.027164 | YES | NO | NA |
| Gzf1 | ENSMUSG( | ENSMUSG( | -0.15148 | 0.147564 | NO | NO | NA |
| ENSMUSG | ENSMUSG( | ENSMUSG( | -0.23327 | 0.047925 | YES | NO | NA |
| Gucy1a1 | ENSMUSG( | ENSMUSG( | 1.117482 | 1.68E-07 | YES | YES | Upregulated |
| Agpat2 | ENSMUSG( | ENSMUSG( | -0.13532 | 0.197429 | NO | NO | NA |
| Tmbim1 | ENSMUSG( | ENSMUSG( | 0.020709 | 0.869914 | NO | NO | NA |
| Slc35e3 | ENSMUSG( | ENSMUSG( | -0.22397 | 0.032968 | YES | NO | NA |
| Mtx3 | ENSMUSG( | ENSMUSG( | 0.332993 | 0.028292 | YES | NO | NA |
| Irf5 | ENSMUSG( | ENSMUSG( | 0.189966 | 0.092357 | NO | NO | NA |
| Fdx1 | ENSMUSG( | ENSMUSG( | 0.321302 | 0.000103 | YES | NO | NA |
| Gpr160 | ENSMUSG( | ENSMUSG( | 0.298711 | 0.013847 | YES | NO | NA |
| Engase | ENSMUSG( | ENSMUSG( | -0.03335 | 0.761446 | NO | NO | NA |
| Mtmr10 | ENSMUSG( | ENSMUSG( | 0.049146 | 0.802479 | NO | NO | NA |
| Usp46 | ENSMUSG( | ENSMUSG( | 0.123164 | 0.319978 | NO | NO | NA |
| Kremen2 | ENSMUSG( | ENSMUSG( | -0.40946 | 0.019055 | YES | NO | NA |
| Asmt | ENSMUSG( | ENSMUSG( | 0.393962 | 0.05785 | NO | NO | NA |
| Hsf2 | ENSMUSG( | ENSMUSG( | 0.213862 | 0.027722 | YES | NO | NA |
| Erlin1 | ENSMUSG( | ENSMUSG( | 0.415453 | 7.55E-05 | YES | NO | NA |
| Tns1 | ENSMUSG( | ENSMUSG( | 0.822939 | 3.24E-10 | YES | NO | NA |

| | | | | | | | |
|---|---|---|---|---|---|---|---|
| Cep85l | ENSMUSG( | ENSMUSG( | 0.017587 | 0.899157 | NO | NO | NA |
| Klf8 | ENSMUSG( | ENSMUSG( | -0.13518 | 0.522612 | NO | NO | NA |
| Zfp969 | ENSMUSG( | ENSMUSG( | -0.46545 | 0.09287 | NO | NO | NA |
| Naaa | ENSMUSG( | ENSMUSG( | -0.27014 | 0.00231 | YES | NO | NA |
| Thap1 | ENSMUSG( | ENSMUSG( | -0.2738 | 0.003054 | YES | NO | NA |
| Crebl2 | ENSMUSG( | ENSMUSG( | -0.62308 | 1.43E-06 | YES | NO | NA |
| Mgat4b | ENSMUSG( | ENSMUSG( | 0.126971 | 0.290434 | NO | NO | NA |
| Zfp157 | ENSMUSG( | ENSMUSG( | -0.48807 | 2.22E-06 | YES | NO | NA |
| Tmem9 | ENSMUSG( | ENSMUSG( | 0.3182 | 0.003872 | YES | NO | NA |
| Plcb3 | ENSMUSG( | ENSMUSG( | -0.40519 | 0.001198 | YES | NO | NA |
| Atp2b2 | ENSMUSG( | ENSMUSG( | 0.58511 | 0.010086 | YES | NO | NA |
| Iffo2 | ENSMUSG( | ENSMUSG( | -0.55776 | 2.50E-05 | YES | NO | NA |
| Gm43197 | ENSMUSG( | ENSMUSG( | -1.00782 | 6.09E-07 | YES | YES | Downregulated |
| Zfp935 | ENSMUSG( | ENSMUSG( | -0.47012 | 0.000211 | YES | NO | NA |
| Qtrt2 | ENSMUSG( | ENSMUSG( | 0.432762 | 0.002883 | YES | NO | NA |
| D130020L( | ENSMUSG( | ENSMUSG( | -0.17995 | 0.608374 | NO | NO | NA |
| Ctnnbip1 | ENSMUSG( | ENSMUSG( | -0.06727 | 0.596362 | NO | NO | NA |
| Rhbdd2 | ENSMUSG( | ENSMUSG( | 0.01632 | 0.90812 | NO | NO | NA |
| Aasdhppt | ENSMUSG( | ENSMUSG( | -0.00358 | 0.976537 | NO | NO | NA |
| Zfp710 | ENSMUSG( | ENSMUSG( | -0.25592 | 0.047714 | YES | NO | NA |
| Timm9 | ENSMUSG( | ENSMUSG( | 0.444519 | 3.38E-05 | YES | NO | NA |
| Inafm1 | ENSMUSG( | ENSMUSG( | -0.01297 | 0.91728 | NO | NO | NA |
| Armcx4 | ENSMUSG( | ENSMUSG( | 0.084359 | 0.505568 | NO | NO | NA |
| Rabl3 | ENSMUSG( | ENSMUSG( | 0.201499 | 0.060521 | NO | NO | NA |
| Tm2d1 | ENSMUSG( | ENSMUSG( | -0.22023 | 0.10029 | NO | NO | NA |
| 9630013D | ENSMUSG( | ENSMUSG( | 0.057107 | 0.716059 | NO | NO | NA |
| D11Wsu47 | ENSMUSG( | ENSMUSG( | -0.26955 | 0.002128 | YES | NO | NA |
| Otud1 | ENSMUSG( | ENSMUSG( | -0.33946 | 6.38E-05 | YES | NO | NA |
| Msra | ENSMUSG( | ENSMUSG( | 0.312238 | 0.007856 | YES | NO | NA |
| Nanp | ENSMUSG( | ENSMUSG( | 0.105546 | 0.352904 | NO | NO | NA |
| 9530082P | ENSMUSG( | ENSMUSG( | -0.22487 | 0.146854 | NO | NO | NA |
| Mzt2 | ENSMUSG( | ENSMUSG( | 0.948554 | 2.89E-07 | YES | NO | NA |
| Leng1 | ENSMUSG( | ENSMUSG( | -0.24111 | 0.048345 | YES | NO | NA |
| Frmd6 | ENSMUSG( | ENSMUSG( | -0.29173 | 0.030807 | YES | NO | NA |
| Jchain | ENSMUSG( | ENSMUSG( | -1.59927 | 0.032169 | YES | YES | Downregulated |
| Sgsm2 | ENSMUSG( | ENSMUSG( | 0.242427 | 0.088758 | NO | NO | NA |
| Zkscan14 | ENSMUSG( | ENSMUSG( | -0.54001 | 0.00114 | YES | NO | NA |
| Abhd10 | ENSMUSG( | ENSMUSG( | 0.08688 | 0.465815 | NO | NO | NA |
| Fam222b | ENSMUSG( | ENSMUSG( | -0.21937 | 0.015654 | YES | NO | NA |
| Cpt2 | ENSMUSG( | ENSMUSG( | -0.02241 | 0.863183 | NO | NO | NA |
| Lgals4 | ENSMUSG( | ENSMUSG( | -0.00262 | 0.988803 | NO | NO | NA |
| Asap2 | ENSMUSG( | ENSMUSG( | -0.83254 | 3.97E-07 | YES | NO | NA |
| Ociad2 | ENSMUSG( | ENSMUSG( | -0.81352 | 0.000657 | YES | NO | NA |
| Alkbh6 | ENSMUSG( | ENSMUSG( | 0.005722 | 0.966048 | NO | NO | NA |
| Slc15a3 | ENSMUSG( | ENSMUSG( | 0.313613 | 0.079299 | NO | NO | NA |
| Zfp524 | ENSMUSG( | ENSMUSG( | -0.19861 | 0.034718 | YES | NO | NA |
| Dnajc25 | ENSMUSG( | ENSMUSG( | 0.158133 | 0.19548 | NO | NO | NA |

| | | | | | | | |
|---|---|---|---|---|---|---|---|
| Det1 | ENSMUSG( | ENSMUSG( | 0.219446 | 0.003975 | YES | NO | NA |
| Sirt3 | ENSMUSG( | ENSMUSG( | 0.311993 | 0.00111 | YES | NO | NA |
| 1700066N | ENSMUSG( | ENSMUSG( | -0.13068 | 0.366763 | NO | NO | NA |
| Cby1 | ENSMUSG( | ENSMUSG( | 0.473766 | 0.002127 | YES | NO | NA |
| Snx32 | ENSMUSG( | ENSMUSG( | -0.2646 | 0.009514 | YES | NO | NA |
| Tsen34 | ENSMUSG( | ENSMUSG( | 0.054754 | 0.692132 | NO | NO | NA |
| Gm15491 | ENSMUSG( | ENSMUSG( | -0.06505 | 0.729989 | NO | NO | NA |
| Arl13b | ENSMUSG( | ENSMUSG( | -0.17567 | 0.130858 | NO | NO | NA |
| Yars2 | ENSMUSG( | ENSMUSG( | 0.209557 | 0.048809 | YES | NO | NA |
| Rai14 | ENSMUSG( | ENSMUSG( | 1.039734 | 4.16E-06 | YES | YES | Upregulated |
| Zfp688 | ENSMUSG( | ENSMUSG( | -0.03059 | 0.823568 | NO | NO | NA |
| Mospd2 | ENSMUSG( | ENSMUSG( | 0.043776 | 0.659678 | NO | NO | NA |
| Dusp22 | ENSMUSG( | ENSMUSG( | 0.144034 | 0.263914 | NO | NO | NA |
| H2bc11 | ENSMUSG( | ENSMUSG( | 0.648939 | 2.56E-07 | YES | NO | NA |
| A530088E( | ENSMUSG( | ENSMUSG( | -0.23721 | 0.152575 | NO | NO | NA |
| Ptdss2 | ENSMUSG( | ENSMUSG( | 0.140763 | 0.11648 | NO | NO | NA |
| Prmt2 | ENSMUSG( | ENSMUSG( | -0.6687 | 7.46E-06 | YES | NO | NA |
| Tmem203 | ENSMUSG( | ENSMUSG( | 0.220677 | 0.201812 | NO | NO | NA |
| Spp1 | ENSMUSG( | ENSMUSG( | 1.205239 | 0.001971 | YES | YES | Upregulated |
| Scart1 | ENSMUSG( | ENSMUSG( | 1.196494 | 0.000907 | YES | YES | Upregulated |
| ENSMUSG | ENSMUSG( | ENSMUSG( | 0.05955 | 0.756154 | NO | NO | NA |
| Zfp61 | ENSMUSG( | ENSMUSG( | 0.254135 | 0.038079 | YES | NO | NA |
| Hexa | ENSMUSG( | ENSMUSG( | -0.07823 | 0.653047 | NO | NO | NA |
| Phf21b | ENSMUSG( | ENSMUSG( | -0.26548 | 0.001791 | YES | NO | NA |
| 4833403J1 | ENSMUSG( | ENSMUSG( | -0.08058 | 0.594452 | NO | NO | NA |
| Slc30a6 | ENSMUSG( | ENSMUSG( | -0.04546 | 0.660508 | NO | NO | NA |
| Elp6 | ENSMUSG( | ENSMUSG( | 0.361639 | 0.001273 | YES | NO | NA |
| Med6 | ENSMUSG( | ENSMUSG( | -0.10832 | 0.612079 | NO | NO | NA |
| Mbip | ENSMUSG( | ENSMUSG( | 0.056197 | 0.68133 | NO | NO | NA |
| D17H6S53 | ENSMUSG( | ENSMUSG( | -0.3202 | 0.013129 | YES | NO | NA |
| 2810001G | ENSMUSG( | ENSMUSG( | 0.66468 | 0.001603 | YES | NO | NA |
| Ndufaf1 | ENSMUSG( | ENSMUSG( | 0.167918 | 0.332415 | NO | NO | NA |
| Gm12940 | ENSMUSG( | ENSMUSG( | 0.018385 | 0.915968 | NO | NO | NA |
| Fam241a | ENSMUSG( | ENSMUSG( | 0.834951 | 2.25E-06 | YES | NO | NA |
| Tango6 | ENSMUSG( | ENSMUSG( | 0.29209 | 0.020001 | YES | NO | NA |
| Tpd52 | ENSMUSG( | ENSMUSG( | -0.13594 | 0.286533 | NO | NO | NA |
| C87436 | ENSMUSG( | ENSMUSG( | -0.3049 | 0.002668 | YES | NO | NA |
| Ddx59 | ENSMUSG( | ENSMUSG( | 0.202794 | 0.097085 | NO | NO | NA |
| Tedc2 | ENSMUSG( | ENSMUSG( | 0.105093 | 0.446859 | NO | NO | NA |
| Zfp933 | ENSMUSG( | ENSMUSG( | 0.783956 | 1.11E-08 | YES | NO | NA |
| Bbc3 | ENSMUSG( | ENSMUSG( | 0.537745 | 0.053582 | NO | NO | NA |
| Iqcc | ENSMUSG( | ENSMUSG( | -0.00053 | 0.996207 | NO | NO | NA |
| Zc3hav1l | ENSMUSG( | ENSMUSG( | 0.230066 | 0.062964 | NO | NO | NA |
| Ube2w | ENSMUSG( | ENSMUSG( | -0.31378 | 0.00025 | YES | NO | NA |
| Avpr2 | ENSMUSG( | ENSMUSG( | -0.31218 | 0.059775 | NO | NO | NA |
| Rgs12 | ENSMUSG( | ENSMUSG( | 0.021008 | 0.887901 | NO | NO | NA |
| Grik5 | ENSMUSG( | ENSMUSG( | 0.843307 | 3.27E-05 | YES | NO | NA |

| | | | | | | | |
|---|---|---|---|---|---|---|---|
| Gm5547 | ENSMUSG( | ENSMUSG( | -0.26522 | 0.057412 | NO | NO | NA |
| Plod2 | ENSMUSG( | ENSMUSG( | -0.0054 | 0.970413 | NO | NO | NA |
| Tspyl2 | ENSMUSG( | ENSMUSG( | -0.19302 | 0.041691 | YES | NO | NA |
| Gpr180 | ENSMUSG( | ENSMUSG( | 0.260427 | 0.086967 | NO | NO | NA |
| Smyd3 | ENSMUSG( | ENSMUSG( | 0.110493 | 0.305132 | NO | NO | NA |
| Erich1 | ENSMUSG( | ENSMUSG( | 0.044963 | 0.730207 | NO | NO | NA |
| Kdm5b | ENSMUSG( | ENSMUSG( | -0.63974 | 0.000598 | YES | NO | NA |
| Ppcs | ENSMUSG( | ENSMUSG( | -0.09055 | 0.350628 | NO | NO | NA |
| Oip5 | ENSMUSG( | ENSMUSG( | 0.311338 | 0.00986 | YES | NO | NA |
| Kctd12 | ENSMUSG( | ENSMUSG( | 0.431195 | 0.022452 | YES | NO | NA |
| Vwa5a | ENSMUSG( | ENSMUSG( | -0.50563 | 0.002903 | YES | NO | NA |
| Cog7 | ENSMUSG( | ENSMUSG( | 0.311949 | 0.000416 | YES | NO | NA |
| Ptpn3 | ENSMUSG( | ENSMUSG( | 0.303942 | 0.064549 | NO | NO | NA |
| Zfp617 | ENSMUSG( | ENSMUSG( | -0.23968 | 0.122149 | NO | NO | NA |
| Mocs3 | ENSMUSG( | ENSMUSG( | -0.0769 | 0.44358 | NO | NO | NA |
| Rpusd1 | ENSMUSG( | ENSMUSG( | 0.213984 | 0.07159 | NO | NO | NA |
| Acot13 | ENSMUSG( | ENSMUSG( | 0.083511 | 0.475251 | NO | NO | NA |
| Cmtm3 | ENSMUSG( | ENSMUSG( | -0.57249 | 1.48E-09 | YES | NO | NA |
| Aga | ENSMUSG( | ENSMUSG( | -0.11929 | 0.328598 | NO | NO | NA |
| Dimt1 | ENSMUSG( | ENSMUSG( | 0.172048 | 0.171335 | NO | NO | NA |
| Gm10509 | ENSMUSG( | ENSMUSG( | -0.26814 | 0.042533 | YES | NO | NA |
| Mybl1 | ENSMUSG( | ENSMUSG( | -0.02973 | 0.867506 | NO | NO | NA |
| Lcmt2 | ENSMUSG( | ENSMUSG( | -0.1097 | 0.173362 | NO | NO | NA |
| Crls1 | ENSMUSG( | ENSMUSG( | 0.167216 | 0.12233 | NO | NO | NA |
| Pusl1 | ENSMUSG( | ENSMUSG( | 0.55777 | 8.85E-05 | YES | NO | NA |
| Zfp748 | ENSMUSG( | ENSMUSG( | -0.44132 | 0.001267 | YES | NO | NA |
| Ttl | ENSMUSG( | ENSMUSG( | 0.25889 | 0.049056 | YES | NO | NA |
| Wdr59 | ENSMUSG( | ENSMUSG( | 0.094773 | 0.390676 | NO | NO | NA |
| Pradc1 | ENSMUSG( | ENSMUSG( | 0.190576 | 0.187457 | NO | NO | NA |
| Fastkd1 | ENSMUSG( | ENSMUSG( | 0.080122 | 0.524288 | NO | NO | NA |
| Mr1 | ENSMUSG( | ENSMUSG( | -0.21237 | 0.078611 | NO | NO | NA |
| Cryzl1 | ENSMUSG( | ENSMUSG( | -0.18319 | 0.172805 | NO | NO | NA |
| Aasdh | ENSMUSG( | ENSMUSG( | -0.00173 | 0.992366 | NO | NO | NA |
| Cdkal1 | ENSMUSG( | ENSMUSG( | -0.19293 | 0.074829 | NO | NO | NA |
| Gm17491 | ENSMUSG( | ENSMUSG( | -0.07995 | 0.624063 | NO | NO | NA |
| Tmem29 | ENSMUSG( | ENSMUSG( | -0.20407 | 0.22157 | NO | NO | NA |
| Afdn | ENSMUSG( | ENSMUSG( | 0.803187 | 0.000671 | YES | NO | NA |
| Fam122a | ENSMUSG( | ENSMUSG( | 0.049721 | 0.73959 | NO | NO | NA |
| ENSMUSG | ENSMUSG( | ENSMUSG( | -0.43086 | 0.004003 | YES | NO | NA |
| Gm10521 | ENSMUSG( | ENSMUSG( | -0.64228 | 3.66E-06 | YES | NO | NA |
| Slc35f2 | ENSMUSG( | ENSMUSG( | 0.08194 | 0.454706 | NO | NO | NA |
| Plppr3 | ENSMUSG( | ENSMUSG( | -0.01031 | 0.942331 | NO | NO | NA |
| Pgap1 | ENSMUSG( | ENSMUSG( | -0.42366 | 0.00149 | YES | NO | NA |
| Homer1 | ENSMUSG( | ENSMUSG( | -0.03843 | 0.758216 | NO | NO | NA |
| Chml | ENSMUSG( | ENSMUSG( | 0.194732 | 0.210086 | NO | NO | NA |
| Dnajc30 | ENSMUSG( | ENSMUSG( | 0.09251 | 0.577104 | NO | NO | NA |
| Crem | ENSMUSG( | ENSMUSG( | -0.05534 | 0.711222 | NO | NO | NA |

| | | | | | | | |
|---|---|---|---|---|---|---|---|
| Ccdc107 | ENSMUSG( | ENSMUSG( | 0.516808 | 0.000932 | YES | NO | NA |
| Ccdc166 | ENSMUSG( | ENSMUSG( | -0.15676 | 0.164747 | NO | NO | NA |
| Akr1e1 | ENSMUSG( | ENSMUSG( | 0.042268 | 0.740889 | NO | NO | NA |
| Med12l | ENSMUSG( | ENSMUSG( | 0.682898 | 5.15E-06 | YES | NO | NA |
| Kif24 | ENSMUSG( | ENSMUSG( | 0.152311 | 0.292273 | NO | NO | NA |
| Jmjd7 | ENSMUSG( | ENSMUSG( | 0.125879 | 0.276671 | NO | NO | NA |
| Tg | ENSMUSG( | ENSMUSG( | 0.722258 | 2.13E-05 | YES | NO | NA |
| Plekhg5 | ENSMUSG( | ENSMUSG( | -0.3841 | 2.00E-05 | YES | NO | NA |
| Gm50387 | ENSMUSG( | ENSMUSG( | 0.144969 | 0.562509 | NO | NO | NA |
| Acsl1 | ENSMUSG( | ENSMUSG( | 0.816845 | 6.45E-06 | YES | NO | NA |
| Tmem219 | ENSMUSG( | ENSMUSG( | 0.070651 | 0.499294 | NO | NO | NA |
| St3gal3 | ENSMUSG( | ENSMUSG( | -0.2995 | 0.025862 | YES | NO | NA |
| Nudt13 | ENSMUSG( | ENSMUSG( | -0.04519 | 0.734883 | NO | NO | NA |
| ENSMUSG( | ENSMUSG( | ENSMUSG( | 1.317025 | 4.23E-09 | YES | YES | Upregulated |
| 3110040N | ENSMUSG( | ENSMUSG( | -0.10493 | 0.569381 | NO | NO | NA |
| Aldh4a1 | ENSMUSG( | ENSMUSG( | -0.40988 | 0.001223 | YES | NO | NA |
| Ydjc | ENSMUSG( | ENSMUSG( | 0.70815 | 5.08E-05 | YES | NO | NA |
| Abcc10 | ENSMUSG( | ENSMUSG( | -0.18714 | 0.398411 | NO | NO | NA |
| Znf41-ps | ENSMUSG( | ENSMUSG( | -3.57425 | 1.37E-24 | YES | YES | Downregulated |
| Ppic | ENSMUSG( | ENSMUSG( | 0.317412 | 0.096295 | NO | NO | NA |
| Pnma8c | ENSMUSG( | ENSMUSG( | -0.55384 | 0.00111 | YES | NO | NA |
| Fan1 | ENSMUSG( | ENSMUSG( | 0.445696 | 0.000197 | YES | NO | NA |
| Bcdin3d | ENSMUSG( | ENSMUSG( | 0.2206 | 0.179797 | NO | NO | NA |
| Gm15459 | ENSMUSG( | ENSMUSG( | -0.2946 | 0.052691 | NO | NO | NA |
| Aunip | ENSMUSG( | ENSMUSG( | 0.335466 | 0.05358 | NO | NO | NA |
| Dnajb5 | ENSMUSG( | ENSMUSG( | 0.188021 | 0.218534 | NO | NO | NA |
| Zfp780b | ENSMUSG( | ENSMUSG( | -0.35989 | 0.001293 | YES | NO | NA |
| Bloc1s3 | ENSMUSG( | ENSMUSG( | -0.19403 | 0.108217 | NO | NO | NA |
| Taco1 | ENSMUSG( | ENSMUSG( | 0.024768 | 0.835507 | NO | NO | NA |
| Taf1a | ENSMUSG( | ENSMUSG( | -0.00725 | 0.953534 | NO | NO | NA |
| Fam3a | ENSMUSG( | ENSMUSG( | -0.0094 | 0.940462 | NO | NO | NA |
| Abraxas1 | ENSMUSG( | ENSMUSG( | 0.173288 | 0.129211 | NO | NO | NA |
| Neto2 | ENSMUSG( | ENSMUSG( | 0.446402 | 0.000724 | YES | NO | NA |
| Acp6 | ENSMUSG( | ENSMUSG( | -0.07356 | 0.616503 | NO | NO | NA |
| Mfsd4b4 | ENSMUSG( | ENSMUSG( | 0.39654 | 0.044613 | YES | NO | NA |
| Zfp930 | ENSMUSG( | ENSMUSG( | -0.06139 | 0.647956 | NO | NO | NA |
| Uxt | ENSMUSG( | ENSMUSG( | 0.290517 | 0.026013 | YES | NO | NA |
| Apobec1 | ENSMUSG( | ENSMUSG( | -0.37514 | 0.02914 | YES | NO | NA |
| Usp40 | ENSMUSG( | ENSMUSG( | -0.08871 | 0.611892 | NO | NO | NA |
| Ypel1 | ENSMUSG( | ENSMUSG( | -0.41323 | 0.004327 | YES | NO | NA |
| Klra3 | ENSMUSG( | ENSMUSG( | -0.46318 | 0.077134 | NO | NO | NA |
| Tmcc3 | ENSMUSG( | ENSMUSG( | -0.34886 | 0.113846 | NO | NO | NA |
| D330041H | ENSMUSG( | ENSMUSG( | -0.36437 | 0.00021 | YES | NO | NA |
| Lxn | ENSMUSG( | ENSMUSG( | -0.50423 | 0.000821 | YES | NO | NA |
| Lrrc28 | ENSMUSG( | ENSMUSG( | 0.172806 | 0.22611 | NO | NO | NA |
| ENSMUSG | ENSMUSG( | ENSMUSG( | 0.088675 | 0.531711 | NO | NO | NA |
| Neu3 | ENSMUSG( | ENSMUSG( | -0.08759 | 0.575669 | NO | NO | NA |

| | | | | | | | |
|---|---|---|---|---|---|---|---|
| Ptp4a1 EN | ENSMUSG( | ENSMUSG( | 0.297721 | 0.217601 | NO | NO | NA |
| Tmem106 | ENSMUSG( | ENSMUSG( | 0.237484 | 0.022629 | YES | NO | NA |
| Naif1 | ENSMUSG( | ENSMUSG( | 0.067384 | 0.572582 | NO | NO | NA |
| Adal | ENSMUSG( | ENSMUSG( | -0.06082 | 0.618922 | NO | NO | NA |
| Pla2g12a | ENSMUSG( | ENSMUSG( | 0.890165 | 8.53E-07 | YES | NO | NA |
| Gm42428 | ENSMUSG( | ENSMUSG( | 0.959721 | 6.64E-05 | YES | NO | NA |
| Enox2 | ENSMUSG( | ENSMUSG( | -0.47991 | 6.18E-08 | YES | NO | NA |
| Ccdc85c | ENSMUSG( | ENSMUSG( | 0.489481 | 0.000332 | YES | NO | NA |
| Zfp775 | ENSMUSG( | ENSMUSG( | 0.357984 | 0.010943 | YES | NO | NA |
| Cbx2 | ENSMUSG( | ENSMUSG( | -0.13871 | 0.365472 | NO | NO | NA |
| Tmub1 | ENSMUSG( | ENSMUSG( | -0.07017 | 0.596362 | NO | NO | NA |
| Ankrd37 | ENSMUSG( | ENSMUSG( | 0.742362 | 0.0077 | YES | NO | NA |
| Zfp677 | ENSMUSG( | ENSMUSG( | -0.37436 | 9.45E-07 | YES | NO | NA |
| Nt5m | ENSMUSG( | ENSMUSG( | 0.251196 | 0.028848 | YES | NO | NA |
| Rhoc | ENSMUSG( | ENSMUSG( | -0.66861 | 0.000686 | YES | NO | NA |
| B3galt4 | ENSMUSG( | ENSMUSG( | -0.60111 | 1.60E-09 | YES | NO | NA |
| Slc10a3 | ENSMUSG( | ENSMUSG( | -0.14496 | 0.325766 | NO | NO | NA |
| Pigl | ENSMUSG( | ENSMUSG( | -0.09809 | 0.440958 | NO | NO | NA |
| Gm12216 | ENSMUSG( | ENSMUSG( | -0.32401 | 0.000665 | YES | NO | NA |
| Tasp1 | ENSMUSG( | ENSMUSG( | 0.270078 | 0.027926 | YES | NO | NA |
| Suv39h2 | ENSMUSG( | ENSMUSG( | 0.168961 | 0.233033 | NO | NO | NA |
| Coa7 | ENSMUSG( | ENSMUSG( | 0.563578 | 0.000208 | YES | NO | NA |
| Ank3 | ENSMUSG( | ENSMUSG( | 0.364911 | 0.121286 | NO | NO | NA |
| Cfl2 | ENSMUSG( | ENSMUSG( | -0.30023 | 0.002093 | YES | NO | NA |
| Rtn4ip1 | ENSMUSG( | ENSMUSG( | -0.04076 | 0.760754 | NO | NO | NA |
| Alg6 | ENSMUSG( | ENSMUSG( | -0.1181 | 0.242187 | NO | NO | NA |
| Tmem101 | ENSMUSG( | ENSMUSG( | -0.05564 | 0.684963 | NO | NO | NA |
| Malsu1 | ENSMUSG( | ENSMUSG( | 0.2344 | 0.028131 | YES | NO | NA |
| ENSMUSG | ENSMUSG( | ENSMUSG( | -0.27739 | 0.015406 | YES | NO | NA |
| Ccdc181 | ENSMUSG( | ENSMUSG( | 0.282258 | 0.079114 | NO | NO | NA |
| Mrpl55 | ENSMUSG( | ENSMUSG( | 0.281929 | 0.093677 | NO | NO | NA |
| Pfkm | ENSMUSG( | ENSMUSG( | 0.262504 | 0.024751 | YES | NO | NA |
| Plcl1 | ENSMUSG( | ENSMUSG( | 0.814916 | 6.29E-05 | YES | NO | NA |
| Slc30a1 | ENSMUSG( | ENSMUSG( | -0.24119 | 0.01314 | YES | NO | NA |
| Alg13 | ENSMUSG( | ENSMUSG( | -0.03845 | 0.736727 | NO | NO | NA |
| Mpo | ENSMUSG( | ENSMUSG( | 1.13521 | 0.114866 | NO | NO | NA |
| Prcp | ENSMUSG( | ENSMUSG( | 0.256993 | 0.010727 | YES | NO | NA |
| Gm6548 | ENSMUSG( | ENSMUSG( | -0.33729 | 0.010236 | YES | NO | NA |
| Gm44170 | ENSMUSG( | ENSMUSG( | 0.164944 | 0.13687 | NO | NO | NA |
| Tpst1 | ENSMUSG( | ENSMUSG( | 0.739086 | 6.69E-08 | YES | NO | NA |
| Praf2 | ENSMUSG( | ENSMUSG( | -0.32036 | 0.000306 | YES | NO | NA |
| Insrr | ENSMUSG( | ENSMUSG( | 0.223409 | 0.350235 | NO | NO | NA |
| Zfp810 | ENSMUSG( | ENSMUSG( | -0.13019 | 0.456206 | NO | NO | NA |
| Rab11fip3 | ENSMUSG( | ENSMUSG( | -0.40252 | 3.19E-06 | YES | NO | NA |
| Hs3st3b1 | ENSMUSG( | ENSMUSG( | 0.469157 | 0.005056 | YES | NO | NA |
| Hdac8 | ENSMUSG( | ENSMUSG( | -0.09345 | 0.571086 | NO | NO | NA |
| Dnajc27 | ENSMUSG( | ENSMUSG( | 0.425275 | 0.000201 | YES | NO | NA |

| | | | | | | | |
|---|---|---|---|---|---|---|---|
| Gm49369 | ENSMUSG( | ENSMUSG( | 0.221938 | 0.814576 | NO | NO | NA |
| Ntmt1 | ENSMUSG( | ENSMUSG( | 0.223826 | 0.03662 | YES | NO | NA |
| Nadsyn1 | ENSMUSG( | ENSMUSG( | -0.34102 | 0.000983 | YES | NO | NA |
| Zdhhc12 | ENSMUSG( | ENSMUSG( | -0.15421 | 0.323141 | NO | NO | NA |
| Tlcd3a | ENSMUSG( | ENSMUSG( | 0.022528 | 0.865915 | NO | NO | NA |
| Nrp2 | ENSMUSG( | ENSMUSG( | -1.62605 | 0.00058 | YES | YES | Downregulated |
| Fgr | ENSMUSG( | ENSMUSG( | -0.23991 | 0.044728 | YES | NO | NA |
| Gm48283 | ENSMUSG( | ENSMUSG( | -0.11454 | 0.588548 | NO | NO | NA |
| 3110082I1 | ENSMUSG( | ENSMUSG( | 1.503294 | 2.86E-08 | YES | YES | Upregulated |
| Snhg3 | ENSMUSG( | ENSMUSG( | 0.943349 | 0.00049 | YES | NO | NA |
| B230307C: | ENSMUSG( | ENSMUSG( | -0.15074 | 0.166188 | NO | NO | NA |
| Prorsd1 | ENSMUSG( | ENSMUSG( | -0.12428 | 0.344361 | NO | NO | NA |
| Trpm4 | ENSMUSG( | ENSMUSG( | 0.148184 | 0.331156 | NO | NO | NA |
| Cenpu | ENSMUSG( | ENSMUSG( | 0.318237 | 0.013548 | YES | NO | NA |
| Lair1 | ENSMUSG( | ENSMUSG( | 0.084054 | 0.600449 | NO | NO | NA |
| Zfp65 | ENSMUSG( | ENSMUSG( | -0.2146 | 0.04646 | YES | NO | NA |
| Popdc2 | ENSMUSG( | ENSMUSG( | 0.141573 | 0.444377 | NO | NO | NA |
| Chchd7 | ENSMUSG( | ENSMUSG( | 0.272195 | 0.019719 | YES | NO | NA |
| Alyref2 | ENSMUSG( | ENSMUSG( | -0.13114 | 0.429043 | NO | NO | NA |
| Zfp268 | ENSMUSG( | ENSMUSG( | -0.27768 | 0.051104 | NO | NO | NA |
| Ttc16 | ENSMUSG( | ENSMUSG( | -0.11561 | 0.398638 | NO | NO | NA |
| Srr | ENSMUSG( | ENSMUSG( | 0.059579 | 0.746048 | NO | NO | NA |
| Emc9 | ENSMUSG( | ENSMUSG( | -0.40361 | 0.0018 | YES | NO | NA |
| Commd5 | ENSMUSG( | ENSMUSG( | 0.047 | 0.62092 | NO | NO | NA |
| Trmt61a | ENSMUSG( | ENSMUSG( | 0.579543 | 1.92E-05 | YES | NO | NA |
| Mrm1 | ENSMUSG( | ENSMUSG( | 0.260871 | 0.029448 | YES | NO | NA |
| Cpne3 | ENSMUSG( | ENSMUSG( | 0.042546 | 0.747282 | NO | NO | NA |
| A630072N | ENSMUSG( | ENSMUSG( | -0.10697 | 0.388544 | NO | NO | NA |
| Spin4 | ENSMUSG( | ENSMUSG( | 0.865292 | 2.27E-06 | YES | NO | NA |
| Epdr1 | ENSMUSG( | ENSMUSG( | 0.401781 | 0.047781 | YES | NO | NA |
| Ubox5 | ENSMUSG( | ENSMUSG( | 0.248027 | 0.088306 | NO | NO | NA |
| 1110002L( | ENSMUSG( | ENSMUSG( | -0.40228 | 0.000264 | YES | NO | NA |
| Leng9 | ENSMUSG( | ENSMUSG( | -0.04132 | 0.753216 | NO | NO | NA |
| Hyi | ENSMUSG( | ENSMUSG( | -0.02807 | 0.843643 | NO | NO | NA |
| Klc3 | ENSMUSG( | ENSMUSG( | -0.38852 | 0.011344 | YES | NO | NA |
| Map3k8 | ENSMUSG( | ENSMUSG( | 0.406729 | 0.00072 | YES | NO | NA |
| Man1c1 | ENSMUSG( | ENSMUSG( | -0.214 | 0.434731 | NO | NO | NA |
| Arhgef9 | ENSMUSG( | ENSMUSG( | 1.572215 | 7.68E-08 | YES | YES | Upregulated |
| Zfp949 | ENSMUSG( | ENSMUSG( | -0.42547 | 0.001691 | YES | NO | NA |
| Psrc1 | ENSMUSG( | ENSMUSG( | 0.220471 | 0.31005 | NO | NO | NA |
| Rtl8a | ENSMUSG( | ENSMUSG( | 0.131657 | 0.432522 | NO | NO | NA |
| Rab39b | ENSMUSG( | ENSMUSG( | 0.197119 | 0.308111 | NO | NO | NA |
| Mrps31 | ENSMUSG( | ENSMUSG( | 0.102994 | 0.472399 | NO | NO | NA |
| Ell2 | ENSMUSG( | ENSMUSG( | 0.118987 | 0.279507 | NO | NO | NA |
| Arsa | ENSMUSG( | ENSMUSG( | -0.40847 | 1.35E-05 | YES | NO | NA |
| Nkiras1 | ENSMUSG( | ENSMUSG( | -0.18484 | 0.160921 | NO | NO | NA |
| Trmt10a | ENSMUSG( | ENSMUSG( | 0.16531 | 0.177617 | NO | NO | NA |

| | | | | | | | |
|---|---|---|---|---|---|---|---|
| Rreb1 | ENSMUSG( | ENSMUSG( | -0.26215 | 0.030902 | YES | NO | NA |
| Tmem38b | ENSMUSG( | ENSMUSG( | -0.15577 | 0.160836 | NO | NO | NA |
| Coq3 | ENSMUSG( | ENSMUSG( | 0.263522 | 0.06282 | NO | NO | NA |
| Lpar2 | ENSMUSG( | ENSMUSG( | -0.16182 | 0.205258 | NO | NO | NA |
| Gm8013 | ENSMUSG( | ENSMUSG( | 6.019698 | 1.60E-86 | YES | YES | Upregulated |
| Gm28901 | ENSMUSG( | ENSMUSG( | -0.41832 | 0.153294 | NO | NO | NA |
| Ogfrl1 | ENSMUSG( | ENSMUSG( | 0.080229 | 0.672948 | NO | NO | NA |
| Bach2 | ENSMUSG( | ENSMUSG( | -0.63358 | 0.000875 | YES | NO | NA |
| Jade3 | ENSMUSG( | ENSMUSG( | 0.066564 | 0.569366 | NO | NO | NA |
| Pigh | ENSMUSG( | ENSMUSG( | -0.12862 | 0.360734 | NO | NO | NA |
| Klc4 | ENSMUSG( | ENSMUSG( | -0.22145 | 0.263079 | NO | NO | NA |
| Cebpzos | ENSMUSG( | ENSMUSG( | 0.160926 | 0.319967 | NO | NO | NA |
| Gm48275 | ENSMUSG( | ENSMUSG( | 1.930946 | 0.312909 | NO | NO | NA |
| Rdh10 | ENSMUSG( | ENSMUSG( | -0.0931 | 0.511078 | NO | NO | NA |
| Zfp932 | ENSMUSG( | ENSMUSG( | -0.16894 | 0.226504 | NO | NO | NA |
| Wdr53 | ENSMUSG( | ENSMUSG( | -0.1621 | 0.138073 | NO | NO | NA |
| Ms4a6d | ENSMUSG( | ENSMUSG( | -0.35567 | 0.000639 | YES | NO | NA |
| Dusp14 | ENSMUSG( | ENSMUSG( | 0.30883 | 0.010882 | YES | NO | NA |
| Coa6 | ENSMUSG( | ENSMUSG( | 0.128195 | 0.209547 | NO | NO | NA |
| Iigp1 | ENSMUSG( | ENSMUSG( | 0.257849 | 0.355036 | NO | NO | NA |
| 2310022B( | ENSMUSG( | ENSMUSG( | 0.210992 | 0.231157 | NO | NO | NA |
| Dhrs11 | ENSMUSG( | ENSMUSG( | 0.16553 | 0.302707 | NO | NO | NA |
| Dnajb13 | ENSMUSG( | ENSMUSG( | -0.113 | 0.592899 | NO | NO | NA |
| Tmem126l | ENSMUSG( | ENSMUSG( | -0.23042 | 0.022972 | YES | NO | NA |
| Plpp5 | ENSMUSG( | ENSMUSG( | 0.028918 | 0.786202 | NO | NO | NA |
| Aig1 | ENSMUSG( | ENSMUSG( | -0.06355 | 0.683724 | NO | NO | NA |
| Spo11 | ENSMUSG( | ENSMUSG( | 0.106555 | 0.677554 | NO | NO | NA |
| 5830408C: | ENSMUSG( | ENSMUSG( | -0.2219 | 0.074318 | NO | NO | NA |
| Fpgt | ENSMUSG( | ENSMUSG( | -0.37759 | 0.002799 | YES | NO | NA |
| Ptges2 | ENSMUSG( | ENSMUSG( | 0.465452 | 5.10E-06 | YES | NO | NA |
| Smim8 | ENSMUSG( | ENSMUSG( | 0.157719 | 0.370029 | NO | NO | NA |
| Trappc2l | ENSMUSG( | ENSMUSG( | -0.31638 | 0.004364 | YES | NO | NA |
| Trmt5 | ENSMUSG( | ENSMUSG( | 0.107298 | 0.343905 | NO | NO | NA |
| Ccdc28b | ENSMUSG( | ENSMUSG( | 0.485311 | 0.023008 | YES | NO | NA |
| Map2k6 | ENSMUSG( | ENSMUSG( | -0.57992 | 4.16E-05 | YES | NO | NA |
| Dyrk1b | ENSMUSG( | ENSMUSG( | -0.64149 | 0.00047 | YES | NO | NA |
| Camsap2 | ENSMUSG( | ENSMUSG( | 1.008604 | 3.15E-05 | YES | YES | Upregulated |
| Ust | ENSMUSG( | ENSMUSG( | -0.21323 | 0.044793 | YES | NO | NA |
| Cela1 | ENSMUSG( | ENSMUSG( | -0.89226 | 0.000109 | YES | NO | NA |
| C2cd2 | ENSMUSG( | ENSMUSG( | -0.55358 | 1.95E-10 | YES | NO | NA |
| Hook2 | ENSMUSG( | ENSMUSG( | 0.340106 | 0.00111 | YES | NO | NA |
| Zfp955a | ENSMUSG( | ENSMUSG( | -0.08914 | 0.535949 | NO | NO | NA |
| Snhg14 | ENSMUSG( | ENSMUSG( | -0.12836 | 0.322298 | NO | NO | NA |
| Dph1 | ENSMUSG( | ENSMUSG( | 0.651861 | 1.27E-06 | YES | NO | NA |
| Hacl1 | ENSMUSG( | ENSMUSG( | 0.330446 | 0.155091 | NO | NO | NA |
| 2700097O | ENSMUSG( | ENSMUSG( | 0.230384 | 0.047707 | YES | NO | NA |
| Egr3 | ENSMUSG( | ENSMUSG( | 1.010964 | 0.001084 | YES | YES | Upregulated |

| | | | | | | | |
|---|---|---|---|---|---|---|---|
| Gm26616 | ENSMUSG( | ENSMUSG( | -0.26891 | 0.02914 | YES | NO | NA |
| 4930417O | ENSMUSG( | ENSMUSG( | -0.64322 | 0.001622 | YES | NO | NA |
| Med29 | ENSMUSG( | ENSMUSG( | 0.155814 | 0.294199 | NO | NO | NA |
| Kics2 | ENSMUSG( | ENSMUSG( | 0.242656 | 0.077981 | NO | NO | NA |
| Gm16675 | ENSMUSG( | ENSMUSG( | -0.10549 | 0.510118 | NO | NO | NA |
| Zfta | ENSMUSG( | ENSMUSG( | -0.12785 | 0.418466 | NO | NO | NA |
| Megf9 | ENSMUSG( | ENSMUSG( | 0.296216 | 0.016289 | YES | NO | NA |
| Gnaq | ENSMUSG( | ENSMUSG( | 0.366404 | 0.000536 | YES | NO | NA |
| Pvrig | ENSMUSG( | ENSMUSG( | -1.22989 | 3.34E-08 | YES | YES | Downregulated |
| Calcrl | ENSMUSG( | ENSMUSG( | -0.51041 | 0.0002 | YES | NO | NA |
| Gm43041 | ENSMUSG( | ENSMUSG( | 0.585884 | 0.000313 | YES | NO | NA |
| Adck1 | ENSMUSG( | ENSMUSG( | 0.165047 | 0.297301 | NO | NO | NA |
| Trp53inp2 | ENSMUSG( | ENSMUSG( | -0.52171 | 0.027116 | YES | NO | NA |
| Rdh13 | ENSMUSG( | ENSMUSG( | 0.827941 | 1.74E-08 | YES | NO | NA |
| Sesn2 | ENSMUSG( | ENSMUSG( | -0.19076 | 0.204053 | NO | NO | NA |
| ENSMUSG | ENSMUSG( | ENSMUSG( | 0.349024 | 0.010511 | YES | NO | NA |
| 5830432E( | ENSMUSG( | ENSMUSG( | -0.4549 | 0.000601 | YES | NO | NA |
| Zfp821 | ENSMUSG( | ENSMUSG( | -0.04449 | 0.745666 | NO | NO | NA |
| Aste1 | ENSMUSG( | ENSMUSG( | 0.028244 | 0.792881 | NO | NO | NA |
| Bace1 | ENSMUSG( | ENSMUSG( | -0.18812 | 0.106841 | NO | NO | NA |
| Ptprv | ENSMUSG( | ENSMUSG( | -0.00284 | 0.986635 | NO | NO | NA |
| Cep57l1 | ENSMUSG( | ENSMUSG( | -0.20871 | 0.071778 | NO | NO | NA |
| Mfsd12 | ENSMUSG( | ENSMUSG( | 0.578418 | 5.60E-06 | YES | NO | NA |
| Gm21982 | ENSMUSG( | ENSMUSG( | 0.740807 | 0.001926 | YES | NO | NA |
| Decr1 | ENSMUSG( | ENSMUSG( | 0.166772 | 0.077511 | NO | NO | NA |
| Tom1 | ENSMUSG( | ENSMUSG( | 0.014017 | 0.92214 | NO | NO | NA |
| Trdmt1 | ENSMUSG( | ENSMUSG( | 0.315975 | 0.010222 | YES | NO | NA |
| Pla2g15 | ENSMUSG( | ENSMUSG( | -0.21153 | 0.092061 | NO | NO | NA |
| Zfp429 | ENSMUSG( | ENSMUSG( | -0.62246 | 1.35E-05 | YES | NO | NA |
| Zcchc10 | ENSMUSG( | ENSMUSG( | 0.05959 | 0.651522 | NO | NO | NA |
| Mrgpre | ENSMUSG( | ENSMUSG( | 0.179459 | 0.211213 | NO | NO | NA |
| Naglu | ENSMUSG( | ENSMUSG( | -0.19077 | 0.090097 | NO | NO | NA |
| Xlr3b | ENSMUSG( | ENSMUSG( | 0.643116 | 0.496053 | NO | NO | NA |
| Cbr4 | ENSMUSG( | ENSMUSG( | 0.461981 | 0.005869 | YES | NO | NA |
| Ddx43 | ENSMUSG( | ENSMUSG( | -0.91213 | 4.74E-10 | YES | NO | NA |
| Trbv1 | ENSMUSG( | ENSMUSG( | -0.20105 | 0.61345 | NO | NO | NA |
| Psmg3 | ENSMUSG( | ENSMUSG( | 0.092887 | 0.481555 | NO | NO | NA |
| Ccdc51 | ENSMUSG( | ENSMUSG( | -0.15319 | 0.173088 | NO | NO | NA |
| Jazf1 | ENSMUSG( | ENSMUSG( | -0.0545 | 0.718224 | NO | NO | NA |
| Amigo3 | ENSMUSG( | ENSMUSG( | 0.075007 | 0.602691 | NO | NO | NA |
| Trmt11 | ENSMUSG( | ENSMUSG( | 0.309132 | 0.012968 | YES | NO | NA |
| Nr1d2 | ENSMUSG( | ENSMUSG( | -0.17107 | 0.428235 | NO | NO | NA |
| Magee1 | ENSMUSG( | ENSMUSG( | -0.30898 | 0.000442 | YES | NO | NA |
| Trim68 | ENSMUSG( | ENSMUSG( | -0.17523 | 0.393625 | NO | NO | NA |
| Lrr1 | ENSMUSG( | ENSMUSG( | 0.281838 | 0.036286 | YES | NO | NA |
| Arl2 | ENSMUSG( | ENSMUSG( | 0.341673 | 0.000493 | YES | NO | NA |
| Dexi | ENSMUSG( | ENSMUSG( | -0.14538 | 0.382946 | NO | NO | NA |

| | | | | | | | |
|---|---|---|---|---|---|---|---|
| Tpt1-ps3 | ENSMUSG( | ENSMUSG( | 0.007968 | 0.968865 | NO | NO | NA |
| Rpain | ENSMUSG( | ENSMUSG( | 0.009207 | 0.95124 | NO | NO | NA |
| Zfp607a | ENSMUSG( | ENSMUSG( | -0.60964 | 1.10E-05 | YES | NO | NA |
| A430088P | ENSMUSG( | ENSMUSG( | -0.2167 | 0.384 | NO | NO | NA |
| Cyren | ENSMUSG( | ENSMUSG( | -0.40392 | 0.00037 | YES | NO | NA |
| Stradb | ENSMUSG( | ENSMUSG( | -0.16145 | 0.413797 | NO | NO | NA |
| Zfp850 | ENSMUSG( | ENSMUSG( | 0.052016 | 0.690509 | NO | NO | NA |
| Btbd8 | ENSMUSG( | ENSMUSG( | -0.20879 | 0.156284 | NO | NO | NA |
| Zfp944 | ENSMUSG( | ENSMUSG( | -0.13532 | 0.232231 | NO | NO | NA |
| 4933412E: | ENSMUSG( | ENSMUSG( | -0.22508 | 0.054068 | NO | NO | NA |
| Fbxo48 | ENSMUSG( | ENSMUSG( | -0.07944 | 0.545532 | NO | NO | NA |
| Prxl2a | ENSMUSG( | ENSMUSG( | 1.353314 | 2.81E-11 | YES | YES | Upregulated |
| 4932422M | ENSMUSG( | ENSMUSG( | -0.07509 | 0.699791 | NO | NO | NA |
| Trbv20 | ENSMUSG( | ENSMUSG( | -0.81567 | 0.021341 | YES | NO | NA |
| Adrb2 | ENSMUSG( | ENSMUSG( | -0.21583 | 0.209513 | NO | NO | NA |
| Ccdc130 | ENSMUSG( | ENSMUSG( | -0.19412 | 0.121618 | NO | NO | NA |
| Hdac10 | ENSMUSG( | ENSMUSG( | -0.28693 | 0.003734 | YES | NO | NA |
| Gm6563 | ENSMUSG( | ENSMUSG( | -0.01719 | 0.92214 | NO | NO | NA |
| Dusp19 | ENSMUSG( | ENSMUSG( | -0.24513 | 0.034362 | YES | NO | NA |
| Rad52 | ENSMUSG( | ENSMUSG( | -0.32526 | 0.004442 | YES | NO | NA |
| Pdcd2l | ENSMUSG( | ENSMUSG( | 0.039894 | 0.700926 | NO | NO | NA |
| Neil1 | ENSMUSG( | ENSMUSG( | -0.28712 | 0.012777 | YES | NO | NA |
| Gm16973 | ENSMUSG( | ENSMUSG( | -0.0195 | 0.875932 | NO | NO | NA |
| Slc26a11 | ENSMUSG( | ENSMUSG( | 0.503277 | 0.016997 | YES | NO | NA |
| Zfp595 | ENSMUSG( | ENSMUSG( | -0.54553 | 0.000593 | YES | NO | NA |
| Pcyox1 | ENSMUSG( | ENSMUSG( | 0.229241 | 0.043032 | YES | NO | NA |
| Iglv1 | ENSMUSG( | ENSMUSG( | -1.17838 | 0.414655 | NO | NO | NA |
| Cacna1i | ENSMUSG( | ENSMUSG( | 0.518419 | 0.14014 | NO | NO | NA |
| Sft2d3 | ENSMUSG( | ENSMUSG( | 0.189986 | 0.209122 | NO | NO | NA |
| Mrc2 | ENSMUSG( | ENSMUSG( | -0.63739 | 0.247168 | NO | NO | NA |
| Mpv17 | ENSMUSG( | ENSMUSG( | -0.16807 | 0.201985 | NO | NO | NA |
| 2010315B( | ENSMUSG( | ENSMUSG( | -0.11808 | 0.515 | NO | NO | NA |
| Pomt2 | ENSMUSG( | ENSMUSG( | 0.310661 | 0.054372 | NO | NO | NA |
| Gm10143 | ENSMUSG( | ENSMUSG( | 0.252171 | 0.072572 | NO | NO | NA |
| Ndufaf2 | ENSMUSG( | ENSMUSG( | 0.28995 | 0.204053 | NO | NO | NA |
| Caskin2 | ENSMUSG( | ENSMUSG( | -0.37967 | 0.039686 | YES | NO | NA |
| Zbtb39 | ENSMUSG( | ENSMUSG( | 0.106575 | 0.573474 | NO | NO | NA |
| Amigo1 | ENSMUSG( | ENSMUSG( | -0.27184 | 0.134349 | NO | NO | NA |
| Tmem241 | ENSMUSG( | ENSMUSG( | -0.00732 | 0.966706 | NO | NO | NA |
| Xrcc4 | ENSMUSG( | ENSMUSG( | -0.29557 | 0.000651 | YES | NO | NA |
| Camta1 | ENSMUSG( | ENSMUSG( | 0.072792 | 0.615844 | NO | NO | NA |
| Rxra | ENSMUSG( | ENSMUSG( | 0.289866 | 0.125824 | NO | NO | NA |
| Acoxl | ENSMUSG( | ENSMUSG( | 0.532689 | 0.00464 | YES | NO | NA |
| BC002059 | ENSMUSG( | ENSMUSG( | -0.31861 | 0.006104 | YES | NO | NA |
| Trim32 | ENSMUSG( | ENSMUSG( | -0.14857 | 0.216209 | NO | NO | NA |
| Hagh | ENSMUSG( | ENSMUSG( | 0.042629 | 0.828365 | NO | NO | NA |
| Cutc | ENSMUSG( | ENSMUSG( | -0.02525 | 0.869827 | NO | NO | NA |

| | | | | | | | |
|---|---|---|---|---|---|---|---|
| B130046B: | ENSMUSG( | ENSMUSG( | -0.05704 | 0.726544 | NO | NO | NA |
| H2aw | ENSMUSG( | ENSMUSG( | 0.139431 | 0.559175 | NO | NO | NA |
| Nckap5l | ENSMUSG( | ENSMUSG( | -0.87381 | 6.39E-08 | YES | NO | NA |
| Polr3g | ENSMUSG( | ENSMUSG( | 0.023098 | 0.881357 | NO | NO | NA |
| Alg11 | ENSMUSG( | ENSMUSG( | -0.21837 | 0.044962 | YES | NO | NA |
| BC035044 | ENSMUSG( | ENSMUSG( | 0.72482 | 0.000222 | YES | NO | NA |
| Ifi211 | ENSMUSG( | ENSMUSG( | 0.744317 | 1.29E-05 | YES | NO | NA |
| Tubb4a | ENSMUSG( | ENSMUSG( | 0.334126 | 0.044473 | YES | NO | NA |
| Gla | ENSMUSG( | ENSMUSG( | -0.01828 | 0.915023 | NO | NO | NA |
| Rxylt1 | ENSMUSG( | ENSMUSG( | 0.600582 | 0.000256 | YES | NO | NA |
| Myo19 | ENSMUSG( | ENSMUSG( | 0.252293 | 0.065311 | NO | NO | NA |
| Usp53 | ENSMUSG( | ENSMUSG( | 0.296544 | 0.012549 | YES | NO | NA |
| 1810049J1 | ENSMUSG( | ENSMUSG( | 0.206342 | 0.492717 | NO | NO | NA |
| Idh2 | ENSMUSG( | ENSMUSG( | 0.004003 | 0.980947 | NO | NO | NA |
| Rps6kc1 | ENSMUSG( | ENSMUSG( | -0.13655 | 0.185449 | NO | NO | NA |
| Fam214b | ENSMUSG( | ENSMUSG( | -0.76174 | 6.18E-06 | YES | NO | NA |
| Cyb5d1 | ENSMUSG( | ENSMUSG( | 0.012611 | 0.923681 | NO | NO | NA |
| Tyro3 | ENSMUSG( | ENSMUSG( | -0.50913 | 0.001477 | YES | NO | NA |
| Evi2 | ENSMUSG( | ENSMUSG( | -0.65629 | 7.47E-05 | YES | NO | NA |
| Exph5 | ENSMUSG( | ENSMUSG( | 1.413063 | 1.72E-09 | YES | YES | Upregulated |
| Ggcx | ENSMUSG( | ENSMUSG( | -0.31314 | 0.009893 | YES | NO | NA |
| Nfia | ENSMUSG( | ENSMUSG( | 0.522454 | 0.1614 | NO | NO | NA |
| Rtl6 | ENSMUSG( | ENSMUSG( | -0.02179 | 0.923834 | NO | NO | NA |
| Cdpf1 | ENSMUSG( | ENSMUSG( | 0.054756 | 0.633647 | NO | NO | NA |
| Wdcp | ENSMUSG( | ENSMUSG( | -0.2192 | 0.107774 | NO | NO | NA |
| Fam81a | ENSMUSG( | ENSMUSG( | 0.119601 | 0.701752 | NO | NO | NA |
| Gm49383 | ENSMUSG( | ENSMUSG( | 0.257298 | 0.09432 | NO | NO | NA |
| Trit1 | ENSMUSG( | ENSMUSG( | -0.00507 | 0.968945 | NO | NO | NA |
| Iba57 | ENSMUSG( | ENSMUSG( | 0.100837 | 0.468404 | NO | NO | NA |
| Dscaml1 | ENSMUSG( | ENSMUSG( | 3.453941 | 9.88E-54 | YES | YES | Upregulated |
| Nckipsd | ENSMUSG( | ENSMUSG( | -0.06124 | 0.655006 | NO | NO | NA |
| Dst | ENSMUSG( | ENSMUSG( | -0.27107 | 0.124971 | NO | NO | NA |
| Gm9888 | ENSMUSG( | ENSMUSG( | 0.782251 | 0.000105 | YES | NO | NA |
| Tfb1m | ENSMUSG( | ENSMUSG( | 0.139423 | 0.37124 | NO | NO | NA |
| Cbr1 | ENSMUSG( | ENSMUSG( | 0.200306 | 0.147015 | NO | NO | NA |
| Nupl2 | ENSMUSG( | ENSMUSG( | 0.56544 | 0.000716 | YES | NO | NA |
| Trub1 | ENSMUSG( | ENSMUSG( | 0.519213 | 0.000216 | YES | NO | NA |
| Blcap | ENSMUSG( | ENSMUSG( | -0.25631 | 0.092091 | NO | NO | NA |
| Socs2 | ENSMUSG( | ENSMUSG( | 1.049416 | 4.12E-07 | YES | YES | Upregulated |
| Slc17a9 | ENSMUSG( | ENSMUSG( | -0.41222 | 0.000821 | YES | NO | NA |
| Pigf | ENSMUSG( | ENSMUSG( | 0.156497 | 0.065114 | NO | NO | NA |
| Ctns | ENSMUSG( | ENSMUSG( | -0.25729 | 0.199708 | NO | NO | NA |
| Klhl7 | ENSMUSG( | ENSMUSG( | 0.403595 | 0.017856 | YES | NO | NA |
| Rufy3 | ENSMUSG( | ENSMUSG( | -0.12846 | 0.331544 | NO | NO | NA |
| Wls | ENSMUSG( | ENSMUSG( | -0.27234 | 0.147209 | NO | NO | NA |
| Zfp747 | ENSMUSG( | ENSMUSG( | -0.11904 | 0.39701 | NO | NO | NA |
| 1700020L: | ENSMUSG( | ENSMUSG( | -0.43231 | 0.005427 | YES | NO | NA |

| | | | | | | | |
|---|---|---|---|---|---|---|---|
| Zfp955b | ENSMUSG( | ENSMUSG( | -0.14816 | 0.339449 | NO | NO | NA |
| Zfp959 | ENSMUSG( | ENSMUSG( | -0.14139 | 0.366421 | NO | NO | NA |
| Fmc1 | ENSMUSG( | ENSMUSG( | 0.704219 | 2.75E-06 | YES | NO | NA |
| ENSMUSG( | ENSMUSG( | ENSMUSG( | -0.05085 | 0.874667 | NO | NO | NA |
| Pik3r2 | ENSMUSG( | ENSMUSG( | 0.674112 | 4.32E-05 | YES | NO | NA |
| Znhit3 | ENSMUSG( | ENSMUSG( | 0.502825 | 2.45E-05 | YES | NO | NA |
| Homez | ENSMUSG( | ENSMUSG( | 0.166483 | 0.254124 | NO | NO | NA |
| Tmem138 | ENSMUSG( | ENSMUSG( | 0.224654 | 0.012624 | YES | NO | NA |
| Tnip3 | ENSMUSG( | ENSMUSG( | 0.324118 | 0.16152 | NO | NO | NA |
| Trbv2 | ENSMUSG( | ENSMUSG( | -0.57633 | 0.172768 | NO | NO | NA |
| Cpd | ENSMUSG( | ENSMUSG( | 1.294085 | 9.35E-06 | YES | YES | Upregulated |
| Mef2c | ENSMUSG( | ENSMUSG( | 0.281354 | 0.156149 | NO | NO | NA |
| Sorbs1 | ENSMUSG( | ENSMUSG( | -0.59192 | 6.06E-05 | YES | NO | NA |
| Cul9 | ENSMUSG( | ENSMUSG( | 0.439703 | 0.052624 | NO | NO | NA |
| Siah1a | ENSMUSG( | ENSMUSG( | -0.42542 | 1.35E-05 | YES | NO | NA |
| Slc2a9 | ENSMUSG( | ENSMUSG( | 0.316289 | 0.092956 | NO | NO | NA |
| Gm13134 | ENSMUSG( | ENSMUSG( | 10.62338 | 2.05E-32 | YES | YES | Upregulated |
| Nsun6 | ENSMUSG( | ENSMUSG( | -0.37265 | 0.001126 | YES | NO | NA |
| Zfp968 | ENSMUSG( | ENSMUSG( | -0.29793 | 0.818237 | NO | NO | NA |
| Mcfd2 | ENSMUSG( | ENSMUSG( | 0.009569 | 0.94245 | NO | NO | NA |
| Gm28042 | ENSMUSG( | ENSMUSG( | -0.7914 | 4.98E-05 | YES | NO | NA |
| Ptpn21 | ENSMUSG( | ENSMUSG( | -0.05645 | 0.775658 | NO | NO | NA |
| Arrdc2 | ENSMUSG( | ENSMUSG( | -0.08094 | 0.607904 | NO | NO | NA |
| Snx30 | ENSMUSG( | ENSMUSG( | 0.206816 | 0.1248 | NO | NO | NA |
| Cand2 | ENSMUSG( | ENSMUSG( | 0.045377 | 0.759918 | NO | NO | NA |
| Cilk1 | ENSMUSG( | ENSMUSG( | 0.150569 | 0.392392 | NO | NO | NA |
| Mfsd13a | ENSMUSG( | ENSMUSG( | 0.904674 | 5.51E-09 | YES | NO | NA |
| Mpi | ENSMUSG( | ENSMUSG( | 0.068607 | 0.666136 | NO | NO | NA |
| Gm32856 | ENSMUSG( | ENSMUSG( | -0.06106 | 0.646606 | NO | NO | NA |
| Zbtb49 | ENSMUSG( | ENSMUSG( | -0.09975 | 0.548323 | NO | NO | NA |
| Gm16933 | ENSMUSG( | ENSMUSG( | -0.05777 | 0.820001 | NO | NO | NA |
| Ptrh1 | ENSMUSG( | ENSMUSG( | 0.611145 | 0.000508 | YES | NO | NA |
| Rpl3-ps1 | ENSMUSG( | ENSMUSG( | 0.495692 | 0.001084 | YES | NO | NA |
| ENSMUSG | ENSMUSG( | ENSMUSG( | -0.01321 | 0.952073 | NO | NO | NA |
| Zfp770 | ENSMUSG( | ENSMUSG( | -0.06568 | 0.646643 | NO | NO | NA |
| Zfp113 | ENSMUSG( | ENSMUSG( | -0.01959 | 0.858038 | NO | NO | NA |
| 2310043P: | ENSMUSG( | ENSMUSG( | -0.62604 | 0.00073 | YES | NO | NA |
| Zfp983 | ENSMUSG( | ENSMUSG( | -0.66962 | 2.30E-06 | YES | NO | NA |
| Poglut2 | ENSMUSG( | ENSMUSG( | 0.635371 | 8.55E-05 | YES | NO | NA |
| Tbc1d7 | ENSMUSG( | ENSMUSG( | 0.404748 | 0.002995 | YES | NO | NA |
| Nek6 | ENSMUSG( | ENSMUSG( | 0.436833 | 0.019215 | YES | NO | NA |
| Lin52 | ENSMUSG( | ENSMUSG( | -0.25117 | 0.003893 | YES | NO | NA |
| Izumo1r | ENSMUSG( | ENSMUSG( | -0.19292 | 0.415514 | NO | NO | NA |
| Matk | ENSMUSG( | ENSMUSG( | 0.605495 | 0.004234 | YES | NO | NA |
| Taf9 | ENSMUSG( | ENSMUSG( | 0.880376 | 5.98E-05 | YES | NO | NA |
| Xkr5 | ENSMUSG( | ENSMUSG( | 0.868054 | 2.61E-06 | YES | NO | NA |
| Gm43737 | ENSMUSG( | ENSMUSG( | 0.328462 | 0.089518 | NO | NO | NA |

| | | | | | | | |
|---|---|---|---|---|---|---|---|
| Metrnl | ENSMUSG( | ENSMUSG( | -0.76604 | 2.77E-06 | YES | NO | NA |
| Ogg1 | ENSMUSG( | ENSMUSG( | -0.08032 | 0.529506 | NO | NO | NA |
| C030006K: | ENSMUSG( | ENSMUSG( | 0.033403 | 0.870435 | NO | NO | NA |
| Trim62 | ENSMUSG( | ENSMUSG( | -0.27742 | 0.057805 | NO | NO | NA |
| Mycbp | ENSMUSG( | ENSMUSG( | 0.279206 | 0.004375 | YES | NO | NA |
| Mterf1a | ENSMUSG( | ENSMUSG( | -0.2538 | 0.102099 | NO | NO | NA |
| Ermap | ENSMUSG( | ENSMUSG( | 0.893035 | 0.054053 | NO | NO | NA |
| Gm2792 | ENSMUSG( | ENSMUSG( | 0.453672 | 0.307005 | NO | NO | NA |
| Pcyox1l | ENSMUSG( | ENSMUSG( | -0.32476 | 0.085518 | NO | NO | NA |
| Hint2 | ENSMUSG( | ENSMUSG( | 0.420767 | 0.010337 | YES | NO | NA |
| Kcnq5 | ENSMUSG( | ENSMUSG( | 0.266747 | 0.078588 | NO | NO | NA |
| Bcl2l2 | ENSMUSG( | ENSMUSG( | 0.266892 | 0.22489 | NO | NO | NA |
| Slx1b | ENSMUSG( | ENSMUSG( | 0.18561 | 0.164527 | NO | NO | NA |
| Mtcp1 | ENSMUSG( | ENSMUSG( | -0.32485 | 0.005082 | YES | NO | NA |
| ENSMUSG | ENSMUSG( | ENSMUSG( | 0.650125 | 0.000263 | YES | NO | NA |
| Trmu | ENSMUSG( | ENSMUSG( | 0.298982 | 0.021709 | YES | NO | NA |
| Selenop | ENSMUSG( | ENSMUSG( | -2.24926 | 8.34E-17 | YES | YES | Downregulated |
| Ret | ENSMUSG( | ENSMUSG( | -1.24753 | 8.04E-06 | YES | YES | Downregulated |
| Alas2 | ENSMUSG( | ENSMUSG( | 0.296502 | 0.636769 | NO | NO | NA |
| Gpatch3 | ENSMUSG( | ENSMUSG( | 0.170915 | 0.377415 | NO | NO | NA |
| Cbwd1 | ENSMUSG( | ENSMUSG( | 0.198391 | 0.029154 | YES | NO | NA |
| Hspa2 | ENSMUSG( | ENSMUSG( | 0.377485 | 0.020179 | YES | NO | NA |
| Fes | ENSMUSG( | ENSMUSG( | 0.493371 | 0.041187 | YES | NO | NA |
| Mlh3 | ENSMUSG( | ENSMUSG( | 0.074951 | 0.631052 | NO | NO | NA |
| Rasal1 | ENSMUSG( | ENSMUSG( | -0.79313 | 0.000187 | YES | NO | NA |
| Nr2f6 | ENSMUSG( | ENSMUSG( | -6.32E-06 | 0.999952 | NO | NO | NA |
| Arl16 | ENSMUSG( | ENSMUSG( | -0.00961 | 0.945161 | NO | NO | NA |
| Fhl3 | ENSMUSG( | ENSMUSG( | -0.08037 | 0.65086 | NO | NO | NA |
| Tnfrsf13c | ENSMUSG( | ENSMUSG( | -0.07869 | 0.559552 | NO | NO | NA |
| Gm37352 | ENSMUSG( | ENSMUSG( | 0.995739 | 0.018254 | YES | NO | NA |
| Dnase1l1 | ENSMUSG( | ENSMUSG( | -0.05371 | 0.672518 | NO | NO | NA |
| 4933439C: | ENSMUSG( | ENSMUSG( | 0.149235 | 0.175573 | NO | NO | NA |
| Epb41l4ao | ENSMUSG( | ENSMUSG( | 0.523764 | 1.14E-06 | YES | NO | NA |
| Zfp839 | ENSMUSG( | ENSMUSG( | 0.235258 | 0.228181 | NO | NO | NA |
| AW55491ł | ENSMUSG( | ENSMUSG( | -0.03677 | 0.759335 | NO | NO | NA |
| Rbks | ENSMUSG( | ENSMUSG( | 0.472257 | 0.041138 | YES | NO | NA |
| Dnase2a | ENSMUSG( | ENSMUSG( | -0.08887 | 0.421429 | NO | NO | NA |
| Ltbp4 | ENSMUSG( | ENSMUSG( | 0.482723 | 0.108218 | NO | NO | NA |
| Gm6637 | ENSMUSG( | ENSMUSG( | -0.1657 | 0.386118 | NO | NO | NA |
| Tnfrsf8 | ENSMUSG( | ENSMUSG( | 0.294841 | 0.099073 | NO | NO | NA |
| 9330133O | ENSMUSG( | ENSMUSG( | 0.159007 | 0.263743 | NO | NO | NA |
| Usp28 | ENSMUSG( | ENSMUSG( | 0.095502 | 0.526309 | NO | NO | NA |
| Zdhhc24 | ENSMUSG( | ENSMUSG( | -0.20123 | 0.114849 | NO | NO | NA |
| Ndufaf5 | ENSMUSG( | ENSMUSG( | 0.089216 | 0.517146 | NO | NO | NA |
| Fam78b | ENSMUSG( | ENSMUSG( | -0.56639 | 0.024093 | YES | NO | NA |
| Lig4 | ENSMUSG( | ENSMUSG( | -0.35777 | 0.000143 | YES | NO | NA |
| Supt3 | ENSMUSG( | ENSMUSG( | 0.049105 | 0.780696 | NO | NO | NA |

| | | | | | | | |
|---|---|---|---|---|---|---|---|
| Slc9a7 | ENSMUSG( | ENSMUSG( | -1.0634 | 1.86E-13 | YES | YES | Downregulated |
| Llph-ps2 | ENSMUSG( | ENSMUSG( | 0.179477 | 0.555025 | NO | NO | NA |
| Acpp | ENSMUSG( | ENSMUSG( | 0.093734 | 0.567401 | NO | NO | NA |
| Ficd | ENSMUSG( | ENSMUSG( | -0.26618 | 0.012004 | YES | NO | NA |
| Rragb | ENSMUSG( | ENSMUSG( | 0.113533 | 0.276671 | NO | NO | NA |
| Gm47210 | ENSMUSG( | ENSMUSG( | 0.561447 | 0.002583 | YES | NO | NA |
| Zfp953 | ENSMUSG( | ENSMUSG( | -0.51917 | 0.000148 | YES | NO | NA |
| Zfp324 | ENSMUSG( | ENSMUSG( | -0.18455 | 0.102405 | NO | NO | NA |
| Klhl26 | ENSMUSG( | ENSMUSG( | -0.47811 | 2.19E-05 | YES | NO | NA |
| Cmtr2 | ENSMUSG( | ENSMUSG( | 0.202558 | 0.151427 | NO | NO | NA |
| Dnajc24 | ENSMUSG( | ENSMUSG( | 0.050206 | 0.682397 | NO | NO | NA |
| Tubd1 | ENSMUSG( | ENSMUSG( | -0.03715 | 0.827445 | NO | NO | NA |
| Galc | ENSMUSG( | ENSMUSG( | 0.485882 | 3.22E-05 | YES | NO | NA |
| Creg1 | ENSMUSG( | ENSMUSG( | 0.169265 | 0.365236 | NO | NO | NA |
| Map1a | ENSMUSG( | ENSMUSG( | 0.404283 | 0.014336 | YES | NO | NA |
| Lbhd1 | ENSMUSG( | ENSMUSG( | 0.408689 | 0.005108 | YES | NO | NA |
| Tmem60 | ENSMUSG( | ENSMUSG( | 0.097897 | 0.351901 | NO | NO | NA |
| Gpld1 | ENSMUSG( | ENSMUSG( | 0.440147 | 0.032953 | YES | NO | NA |
| Acadsb | ENSMUSG( | ENSMUSG( | -0.19723 | 0.280897 | NO | NO | NA |
| Chit1 | ENSMUSG( | ENSMUSG( | -0.38194 | 0.134672 | NO | NO | NA |
| Ercc8 | ENSMUSG( | ENSMUSG( | 0.354431 | 0.000535 | YES | NO | NA |
| Borcs7 | ENSMUSG( | ENSMUSG( | -0.34911 | 0.003069 | YES | NO | NA |
| Gas8 | ENSMUSG( | ENSMUSG( | 0.718111 | 1.92E-05 | YES | NO | NA |
| Lsm10 | ENSMUSG( | ENSMUSG( | -0.45813 | 0.000167 | YES | NO | NA |
| Cdk5rap1 | ENSMUSG( | ENSMUSG( | 0.0853 | 0.542905 | NO | NO | NA |
| Chchd5 | ENSMUSG( | ENSMUSG( | 0.147663 | 0.269544 | NO | NO | NA |
| Tirap | ENSMUSG( | ENSMUSG( | 0.277642 | 0.004105 | YES | NO | NA |
| Wdr5b | ENSMUSG( | ENSMUSG( | -0.085 | 0.6068 | NO | NO | NA |
| Tnfsf4 | ENSMUSG( | ENSMUSG( | 0.54181 | 0.009021 | YES | NO | NA |
| Tmem143 | ENSMUSG( | ENSMUSG( | 0.149017 | 0.330503 | NO | NO | NA |
| Trappc2 | ENSMUSG( | ENSMUSG( | -0.32631 | 0.015592 | YES | NO | NA |
| Gstz1 | ENSMUSG( | ENSMUSG( | 0.306432 | 0.036314 | YES | NO | NA |
| Kiz | ENSMUSG( | ENSMUSG( | -0.05825 | 0.759406 | NO | NO | NA |
| Hsf4 | ENSMUSG( | ENSMUSG( | -0.34912 | 0.036776 | YES | NO | NA |
| Zc3h12c | ENSMUSG( | ENSMUSG( | 1.281109 | 1.21E-08 | YES | YES | Upregulated |
| Amer1 | ENSMUSG( | ENSMUSG( | -0.31349 | 0.013749 | YES | NO | NA |
| Zc3h8 | ENSMUSG( | ENSMUSG( | 0.226588 | 0.115582 | NO | NO | NA |
| ENSMUSG | ENSMUSG( | ENSMUSG( | 0.253236 | 0.150749 | NO | NO | NA |
| Glt8d1 | ENSMUSG( | ENSMUSG( | -0.2942 | 0.121345 | NO | NO | NA |
| Mmab | ENSMUSG( | ENSMUSG( | 0.461237 | 0.000863 | YES | NO | NA |
| 5430405H | ENSMUSG( | ENSMUSG( | -0.00784 | 0.960063 | NO | NO | NA |
| Gga2 | ENSMUSG( | ENSMUSG( | 0.206121 | 0.220257 | NO | NO | NA |
| Zfp60 | ENSMUSG( | ENSMUSG( | -0.47775 | 5.52E-06 | YES | NO | NA |
| Iqcg | ENSMUSG( | ENSMUSG( | -0.20956 | 0.267916 | NO | NO | NA |
| Nat2 | ENSMUSG( | ENSMUSG( | -0.04795 | 0.763382 | NO | NO | NA |
| Pomt1 | ENSMUSG( | ENSMUSG( | 0.263764 | 0.018028 | YES | NO | NA |
| Mrrf | ENSMUSG( | ENSMUSG( | 0.025426 | 0.860115 | NO | NO | NA |

| | | | | | | | |
|---|---|---|---|---|---|---|---|
| Tmem221 | ENSMUSG( | ENSMUSG( | -0.64123 | 1.86E-05 | YES | NO | NA |
| Naa80 | ENSMUSG( | ENSMUSG( | -0.00457 | 0.976024 | NO | NO | NA |
| Gm45867 | ENSMUSG( | ENSMUSG( | -0.7305 | 0.003122 | YES | NO | NA |
| Tuft1 | ENSMUSG( | ENSMUSG( | 0.054268 | 0.701965 | NO | NO | NA |
| Taf4b | ENSMUSG( | ENSMUSG( | 0.81834 | 2.47E-07 | YES | NO | NA |
| 5830428M | ENSMUSG( | ENSMUSG( | -0.08558 | 0.642562 | NO | NO | NA |
| Trmt44 | ENSMUSG( | ENSMUSG( | -0.02052 | 0.877074 | NO | NO | NA |
| Gm16568 | ENSMUSG( | ENSMUSG( | 0.128668 | 0.523483 | NO | NO | NA |
| Smco4 | ENSMUSG( | ENSMUSG( | -0.10267 | 0.373608 | NO | NO | NA |
| Gm44729 | ENSMUSG( | ENSMUSG( | 0.147101 | 0.591064 | NO | NO | NA |
| Pcyt1b | ENSMUSG( | ENSMUSG( | 0.089188 | 0.688983 | NO | NO | NA |
| Eola1 | ENSMUSG( | ENSMUSG( | -0.0673 | 0.739663 | NO | NO | NA |
| Tmem86b | ENSMUSG( | ENSMUSG( | 0.185567 | 0.184927 | NO | NO | NA |
| Trav7-4 | ENSMUSG( | ENSMUSG( | -0.1339 | 0.805716 | NO | NO | NA |
| Zfp40 | ENSMUSG( | ENSMUSG( | -0.26709 | 0.063705 | NO | NO | NA |
| ENSMUSG( | ENSMUSG( | ENSMUSG( | -0.35284 | 0.001137 | YES | NO | NA |
| Fbxo30 | ENSMUSG( | ENSMUSG( | -0.05992 | 0.650468 | NO | NO | NA |
| 2610044O | ENSMUSG( | ENSMUSG( | -0.14097 | 0.423134 | NO | NO | NA |
| H2bc4 | ENSMUSG( | ENSMUSG( | 0.399127 | 0.077341 | NO | NO | NA |
| Peg13 | ENSMUSG( | ENSMUSG( | -0.65176 | 3.67E-05 | YES | NO | NA |
| Smim13 | ENSMUSG( | ENSMUSG( | 0.004908 | 0.974457 | NO | NO | NA |
| Ralgps1 | ENSMUSG( | ENSMUSG( | -0.88033 | 2.12E-06 | YES | NO | NA |
| Plcg2 | ENSMUSG( | ENSMUSG( | 0.119352 | 0.500004 | NO | NO | NA |
| Stom | ENSMUSG( | ENSMUSG( | 0.517071 | 0.001766 | YES | NO | NA |
| Brf2 | ENSMUSG( | ENSMUSG( | 0.070371 | 0.536004 | NO | NO | NA |
| Lysmd1 | ENSMUSG( | ENSMUSG( | -0.26324 | 0.004928 | YES | NO | NA |
| Trp53rkb | ENSMUSG( | ENSMUSG( | 0.042577 | 0.71246 | NO | NO | NA |
| Gm10366 | ENSMUSG( | ENSMUSG( | 0.018464 | 0.915707 | NO | NO | NA |
| Ighv1-82 | ENSMUSG( | ENSMUSG( | 0.262787 | 0.895165 | NO | NO | NA |
| St6galnac3 | ENSMUSG( | ENSMUSG( | 0.05046 | 0.830613 | NO | NO | NA |
| Ccdc58 | ENSMUSG( | ENSMUSG( | 0.498781 | 9.99E-05 | YES | NO | NA |
| Pigyl | ENSMUSG( | ENSMUSG( | 0.026943 | 0.836456 | NO | NO | NA |
| Gm11454 | ENSMUSG( | ENSMUSG( | -0.5867 | 0.025835 | YES | NO | NA |
| Pik3cb | ENSMUSG( | ENSMUSG( | 0.01788 | 0.89271 | NO | NO | NA |
| Cdyl2 | ENSMUSG( | ENSMUSG( | -0.04671 | 0.72481 | NO | NO | NA |
| Acer3 | ENSMUSG( | ENSMUSG( | -0.02481 | 0.868688 | NO | NO | NA |
| Rida | ENSMUSG( | ENSMUSG( | -0.04219 | 0.788575 | NO | NO | NA |
| B4gat1 | ENSMUSG( | ENSMUSG( | 0.711524 | 0.000337 | YES | NO | NA |
| Fcsk | ENSMUSG( | ENSMUSG( | -0.30568 | 0.000588 | YES | NO | NA |
| Gm29216 | ENSMUSG( | ENSMUSG( | 0.188082 | 0.513766 | NO | NO | NA |
| Cenpp | ENSMUSG( | ENSMUSG( | 0.731327 | 2.70E-07 | YES | NO | NA |
| Ndufaf8 | ENSMUSG( | ENSMUSG( | 0.073613 | 0.635315 | NO | NO | NA |
| Tsen15 | ENSMUSG( | ENSMUSG( | 0.029302 | 0.844653 | NO | NO | NA |
| Cybb | ENSMUSG( | ENSMUSG( | -1.48132 | 0.00057 | YES | YES | Downregulated |
| Pcgf1 | ENSMUSG( | ENSMUSG( | 0.07672 | 0.663944 | NO | NO | NA |
| Nsun4 | ENSMUSG( | ENSMUSG( | -0.03509 | 0.812696 | NO | NO | NA |
| Micu3 | ENSMUSG( | ENSMUSG( | 0.085798 | 0.616442 | NO | NO | NA |

| | | | | | | | |
|---|---|---|---|---|---|---|---|
| Ccdc15 | ENSMUSG( | ENSMUSG( | 0.03245 | 0.832943 | NO | NO | NA |
| Ppp2r3a | ENSMUSG( | ENSMUSG( | 0.378899 | 0.003832 | YES | NO | NA |
| Nudt15 | ENSMUSG( | ENSMUSG( | 0.120028 | 0.327932 | NO | NO | NA |
| Yipf2 | ENSMUSG( | ENSMUSG( | -0.14869 | 0.294351 | NO | NO | NA |
| Dhrs3 | ENSMUSG( | ENSMUSG( | 1.72077 | 2.73E-37 | YES | YES | Upregulated |
| E430024I0 | ENSMUSG( | ENSMUSG( | -0.04522 | 0.798662 | NO | NO | NA |
| Rfx2 | ENSMUSG( | ENSMUSG( | 0.099411 | 0.628823 | NO | NO | NA |
| Magi3 | ENSMUSG( | ENSMUSG( | 0.338465 | 0.044696 | YES | NO | NA |
| Oxnad1 | ENSMUSG( | ENSMUSG( | 0.111184 | 0.421499 | NO | NO | NA |
| Lyrm2 | ENSMUSG( | ENSMUSG( | 0.197587 | 0.30141 | NO | NO | NA |
| Ptpn5 | ENSMUSG( | ENSMUSG( | -0.06084 | 0.797332 | NO | NO | NA |
| Gm20498 | ENSMUSG( | ENSMUSG( | -0.07287 | 0.511663 | NO | NO | NA |
| Lrrc1 | ENSMUSG( | ENSMUSG( | -0.33396 | 0.010402 | YES | NO | NA |
| Rasgef1a | ENSMUSG( | ENSMUSG( | -0.5004 | 0.00036 | YES | NO | NA |
| Tmem63b | ENSMUSG( | ENSMUSG( | -0.29564 | 0.001119 | YES | NO | NA |
| 2810021J2 | ENSMUSG( | ENSMUSG( | -0.45102 | 0.001286 | YES | NO | NA |
| Slfn5 | ENSMUSG( | ENSMUSG( | -0.65825 | 0.000345 | YES | NO | NA |
| Gdpgp1 | ENSMUSG( | ENSMUSG( | -0.2618 | 0.067715 | NO | NO | NA |
| Snrnp35 | ENSMUSG( | ENSMUSG( | 0.072863 | 0.75842 | NO | NO | NA |
| Zfp958 | ENSMUSG( | ENSMUSG( | -0.30161 | 0.050504 | NO | NO | NA |
| Zfp74 | ENSMUSG( | ENSMUSG( | -0.10789 | 0.459618 | NO | NO | NA |
| Hhat | ENSMUSG( | ENSMUSG( | 1.83288 | 1.55E-07 | YES | YES | Upregulated |
| Rab6b | ENSMUSG( | ENSMUSG( | -5.0428 | 1.35E-30 | YES | YES | Downregulated |
| Qsox2 | ENSMUSG( | ENSMUSG( | 0.367947 | 0.003943 | YES | NO | NA |
| Tyw5 | ENSMUSG( | ENSMUSG( | -0.10656 | 0.450067 | NO | NO | NA |
| Mvk | ENSMUSG( | ENSMUSG( | 0.818296 | 7.01E-09 | YES | NO | NA |
| Gm9887 | ENSMUSG( | ENSMUSG( | -0.15625 | 0.257658 | NO | NO | NA |
| Lca5 | ENSMUSG( | ENSMUSG( | 0.0241 | 0.909337 | NO | NO | NA |
| Trav3-3 | ENSMUSG( | ENSMUSG( | 0.165049 | 0.566624 | NO | NO | NA |
| Rn7s2 | ENSMUSG( | ENSMUSG( | 2.159001 | 0.566716 | NO | NO | NA |
| Gm12896 | ENSMUSG( | ENSMUSG( | 0.333162 | 0.67946 | NO | NO | NA |
| Zfp81 | ENSMUSG( | ENSMUSG( | -0.12227 | 0.379251 | NO | NO | NA |
| B230354K | ENSMUSG( | ENSMUSG( | -0.18336 | 0.230313 | NO | NO | NA |
| Ms4a4c | ENSMUSG( | ENSMUSG( | -0.27068 | 0.069591 | NO | NO | NA |
| Rassf7 | ENSMUSG( | ENSMUSG( | 0.037007 | 0.797332 | NO | NO | NA |
| Gm4117 | ENSMUSG( | ENSMUSG( | 0.612391 | 0.5021 | NO | NO | NA |
| Ttn | ENSMUSG( | ENSMUSG( | 0.690051 | 0.001609 | YES | NO | NA |
| Usp54 | ENSMUSG( | ENSMUSG( | 0.159339 | 0.278533 | NO | NO | NA |
| Pigg | ENSMUSG( | ENSMUSG( | 0.172669 | 0.241792 | NO | NO | NA |
| Nmb | ENSMUSG( | ENSMUSG( | -0.32714 | 0.006606 | YES | NO | NA |
| Slc38a6 | ENSMUSG( | ENSMUSG( | 0.033145 | 0.836247 | NO | NO | NA |
| Gemin6 | ENSMUSG( | ENSMUSG( | 0.463301 | 0.00053 | YES | NO | NA |
| Gm15446 | ENSMUSG( | ENSMUSG( | -0.24564 | 0.027822 | YES | NO | NA |
| Galm | ENSMUSG( | ENSMUSG( | -0.71983 | 1.98E-08 | YES | NO | NA |
| Exoc3l | ENSMUSG( | ENSMUSG( | 0.708652 | 0.00249 | YES | NO | NA |
| Zfp771 | ENSMUSG( | ENSMUSG( | 0.380894 | 0.003784 | YES | NO | NA |
| Zfp788 | ENSMUSG( | ENSMUSG( | 0.010672 | 0.933709 | NO | NO | NA |

| | | | | | | | |
|---|---|---|---|---|---|---|---|
| Zfp90 | ENSMUSG( | ENSMUSG( | -0.3217 | 0.003025 | YES | NO | NA |
| C1galt1c1 | ENSMUSG( | ENSMUSG( | -0.23117 | 0.150945 | NO | NO | NA |
| 2810429I0 | ENSMUSG( | ENSMUSG( | 0.720555 | 8.55E-08 | YES | NO | NA |
| Apoo | ENSMUSG( | ENSMUSG( | 0.565025 | 3.17E-06 | YES | NO | NA |
| Trav14d-1 | ENSMUSG( | ENSMUSG( | 0.438187 | 0.798585 | NO | NO | NA |
| 2700062C( | ENSMUSG( | ENSMUSG( | 0.389632 | 0.011104 | YES | NO | NA |
| Rgs11 | ENSMUSG( | ENSMUSG( | -0.04051 | 0.852829 | NO | NO | NA |
| Pex12 | ENSMUSG( | ENSMUSG( | 0.128908 | 0.237913 | NO | NO | NA |
| As3mt | ENSMUSG( | ENSMUSG( | -1.07453 | 0.002424 | YES | YES | Downregulated |
| ENSMUSG | ENSMUSG( | ENSMUSG( | -0.40142 | 0.000971 | YES | NO | NA |
| Bod1 | ENSMUSG( | ENSMUSG( | 0.086298 | 0.582481 | NO | NO | NA |
| Eef2kmt | ENSMUSG( | ENSMUSG( | 0.116333 | 0.427102 | NO | NO | NA |
| Gab3 | ENSMUSG( | ENSMUSG( | -0.60907 | 3.46E-05 | YES | NO | NA |
| P2ry10b | ENSMUSG( | ENSMUSG( | -0.43809 | 0.002834 | YES | NO | NA |
| Glb1l | ENSMUSG( | ENSMUSG( | 0.213478 | 0.203811 | NO | NO | NA |
| ENSMUSG | ENSMUSG( | ENSMUSG( | 0.318412 | 0.104815 | NO | NO | NA |
| H1f4 | ENSMUSG( | ENSMUSG( | 0.022765 | 0.910843 | NO | NO | NA |
| Rnf43 | ENSMUSG( | ENSMUSG( | -0.63959 | 0.017114 | YES | NO | NA |
| Gm49325 | ENSMUSG( | ENSMUSG( | 0.242011 | 0.846414 | NO | NO | NA |
| Trav1 | ENSMUSG( | ENSMUSG( | 0.049256 | 0.971034 | NO | NO | NA |
| Elmod3 | ENSMUSG( | ENSMUSG( | 0.092296 | 0.58481 | NO | NO | NA |
| Decr2 | ENSMUSG( | ENSMUSG( | -0.06909 | 0.614865 | NO | NO | NA |
| Pex26 | ENSMUSG( | ENSMUSG( | -0.48894 | 6.24E-06 | YES | NO | NA |
| Agrn | ENSMUSG( | ENSMUSG( | -0.89721 | 9.16E-06 | YES | NO | NA |
| Ston2 | ENSMUSG( | ENSMUSG( | 1.248929 | 1.31E-07 | YES | YES | Upregulated |
| Pyurf | ENSMUSG( | ENSMUSG( | 0.548741 | 0.00118 | YES | NO | NA |
| H1f3 | ENSMUSG( | ENSMUSG( | 0.068704 | 0.785803 | NO | NO | NA |
| Zfp683 | ENSMUSG( | ENSMUSG( | 1.605379 | 3.02E-06 | YES | YES | Upregulated |
| H2ac19 | ENSMUSG( | ENSMUSG( | 0.50502 | 0.00848 | YES | NO | NA |
| D3Ertd751 | ENSMUSG( | ENSMUSG( | -0.29224 | 0.050727 | NO | NO | NA |
| Gm14391 | ENSMUSG( | ENSMUSG( | -0.12435 | 0.65219 | NO | NO | NA |
| Ermn | ENSMUSG( | ENSMUSG( | 1.346065 | 7.31E-09 | YES | YES | Upregulated |
| Pibf1 | ENSMUSG( | ENSMUSG( | -0.00876 | 0.96061 | NO | NO | NA |
| 2310039H | ENSMUSG( | ENSMUSG( | 0.802906 | 0.000136 | YES | NO | NA |
| Gm38253 | ENSMUSG( | ENSMUSG( | -0.13893 | 0.485239 | NO | NO | NA |
| Evi5l | ENSMUSG( | ENSMUSG( | -0.05848 | 0.729331 | NO | NO | NA |
| Qprt | ENSMUSG( | ENSMUSG( | -0.69351 | 3.27E-09 | YES | NO | NA |
| Zhx2 | ENSMUSG( | ENSMUSG( | -0.40392 | 0.005951 | YES | NO | NA |
| Sh2b3 | ENSMUSG( | ENSMUSG( | 0.266021 | 0.116692 | NO | NO | NA |
| Tanc2 | ENSMUSG( | ENSMUSG( | -0.43746 | 0.000159 | YES | NO | NA |
| Zfp235 | ENSMUSG( | ENSMUSG( | -0.40463 | 0.002623 | YES | NO | NA |
| Snap47 | ENSMUSG( | ENSMUSG( | -0.23177 | 0.133821 | NO | NO | NA |
| Aplf | ENSMUSG( | ENSMUSG( | 0.023868 | 0.865515 | NO | NO | NA |
| Abcb6 | ENSMUSG( | ENSMUSG( | 0.609336 | 0.000142 | YES | NO | NA |
| Zfp456 | ENSMUSG( | ENSMUSG( | -0.72993 | 0.000385 | YES | NO | NA |
| Rpl35a-ps2 | ENSMUSG( | ENSMUSG( | 0.224942 | 0.898856 | NO | NO | NA |
| Itsn1 | ENSMUSG( | ENSMUSG( | 0.561198 | 0.002134 | YES | NO | NA |

| | | | | | | | |
|---|---|---|---|---|---|---|---|
| 9130221H | ENSMUSG( | ENSMUSG( | -0.28612 | 0.013783 | YES | NO | NA |
| Rnf165 | ENSMUSG( | ENSMUSG( | -0.53303 | 0.128838 | NO | NO | NA |
| Wars2 | ENSMUSG( | ENSMUSG( | 0.01789 | 0.901489 | NO | NO | NA |
| Art2a | ENSMUSG( | ENSMUSG( | -0.93722 | 0.000153 | YES | NO | NA |
| Glce | ENSMUSG( | ENSMUSG( | 0.73725 | 6.46E-08 | YES | NO | NA |
| Akr7a5 | ENSMUSG( | ENSMUSG( | 0.564301 | 0.000416 | YES | NO | NA |
| Slc39a13 | ENSMUSG( | ENSMUSG( | 0.106164 | 0.534573 | NO | NO | NA |
| Endog | ENSMUSG( | ENSMUSG( | 0.804365 | 5.82E-06 | YES | NO | NA |
| Zfp874a | ENSMUSG( | ENSMUSG( | -0.82783 | 1.40E-06 | YES | NO | NA |
| Fam210b | ENSMUSG( | ENSMUSG( | -0.08329 | 0.665321 | NO | NO | NA |
| Pdf | ENSMUSG( | ENSMUSG( | 0.305258 | 0.075582 | NO | NO | NA |
| Hint3 | ENSMUSG( | ENSMUSG( | -0.26109 | 0.109893 | NO | NO | NA |
| Ccdc163 | ENSMUSG( | ENSMUSG( | 0.147233 | 0.204125 | NO | NO | NA |
| Sh3rf1 | ENSMUSG( | ENSMUSG( | 0.040184 | 0.812047 | NO | NO | NA |
| Gm1976 | ENSMUSG( | ENSMUSG( | 0.293579 | 0.076751 | NO | NO | NA |
| Klhl15 | ENSMUSG( | ENSMUSG( | -0.01782 | 0.919414 | NO | NO | NA |
| Zkscan7 | ENSMUSG( | ENSMUSG( | -0.24222 | 0.146822 | NO | NO | NA |
| Lrrc75b | ENSMUSG( | ENSMUSG( | -1.03431 | 3.16E-06 | YES | YES | Downregulated |
| Parp8 | ENSMUSG( | ENSMUSG( | 0.049103 | 0.801947 | NO | NO | NA |
| Lmna | ENSMUSG( | ENSMUSG( | 0.634439 | 0.000454 | YES | NO | NA |
| Sertad3 | ENSMUSG( | ENSMUSG( | -0.3469 | 0.004169 | YES | NO | NA |
| Atf5 | ENSMUSG( | ENSMUSG( | -0.1436 | 0.364915 | NO | NO | NA |
| Optn | ENSMUSG( | ENSMUSG( | 0.033147 | 0.841547 | NO | NO | NA |
| Ntpcr | ENSMUSG( | ENSMUSG( | 0.241556 | 0.061444 | NO | NO | NA |
| Dlg4 | ENSMUSG( | ENSMUSG( | 0.262933 | 0.560273 | NO | NO | NA |
| Sergef | ENSMUSG( | ENSMUSG( | 0.32276 | 0.000507 | YES | NO | NA |
| Gm12758 | ENSMUSG( | ENSMUSG( | -0.11478 | 0.317999 | NO | NO | NA |
| Spire1 | ENSMUSG( | ENSMUSG( | 1.340536 | 1.19E-06 | YES | YES | Upregulated |
| Slc25a15 | ENSMUSG( | ENSMUSG( | 0.635667 | 4.23E-05 | YES | NO | NA |
| Zfp874b | ENSMUSG( | ENSMUSG( | -0.88342 | 1.23E-07 | YES | NO | NA |
| Gm11427 | ENSMUSG( | ENSMUSG( | -0.16934 | 0.369755 | NO | NO | NA |
| Tubb2a | ENSMUSG( | ENSMUSG( | 0.265268 | 0.18742 | NO | NO | NA |
| Mterf4 | ENSMUSG( | ENSMUSG( | -0.22435 | 0.131395 | NO | NO | NA |
| Tatdn1 | ENSMUSG( | ENSMUSG( | -0.06156 | 0.687679 | NO | NO | NA |
| Gm37233 | ENSMUSG( | ENSMUSG( | -0.27394 | 0.298484 | NO | NO | NA |
| 5730480H | ENSMUSG( | ENSMUSG( | 0.194777 | 0.038697 | YES | NO | NA |
| Slc27a1 | ENSMUSG( | ENSMUSG( | 0.505998 | 0.00039 | YES | NO | NA |
| Fam120c | ENSMUSG( | ENSMUSG( | -0.32026 | 0.027968 | YES | NO | NA |
| Gm4617 | ENSMUSG( | ENSMUSG( | 0.453271 | 0.004924 | YES | NO | NA |
| Siah1b | ENSMUSG( | ENSMUSG( | 0.462317 | 0.004212 | YES | NO | NA |
| Gm38431 | ENSMUSG( | ENSMUSG( | 0.061389 | 0.863456 | NO | NO | NA |
| Klre1 | ENSMUSG( | ENSMUSG( | -0.72495 | 0.209972 | NO | NO | NA |
| Plk2 | ENSMUSG( | ENSMUSG( | 0.619541 | 0.029249 | YES | NO | NA |
| 1700030K( | ENSMUSG( | ENSMUSG( | -0.00362 | 0.980226 | NO | NO | NA |
| H1f10 | ENSMUSG( | ENSMUSG( | 1.881681 | 0.000142 | YES | YES | Upregulated |
| Atp7a | ENSMUSG( | ENSMUSG( | -0.62755 | 2.93E-06 | YES | NO | NA |
| 1700008J( | ENSMUSG( | ENSMUSG( | -0.35184 | 0.04106 | YES | NO | NA |

| | | | | | | | |
|---|---|---|---|---|---|---|---|
| Slc38a7 | ENSMUSG( | ENSMUSG( | -0.45808 | 0.020449 | YES | NO | NA |
| Klhl22 | ENSMUSG( | ENSMUSG( | -0.44243 | 7.56E-05 | YES | NO | NA |
| Zfp3 | ENSMUSG( | ENSMUSG( | -0.179 | 0.326393 | NO | NO | NA |
| Mapk1ip1 | ENSMUSG( | ENSMUSG( | -0.47521 | 0.001071 | YES | NO | NA |
| Trav8d-2 | ENSMUSG( | ENSMUSG( | -1.54397 | 0.113352 | NO | NO | NA |
| Ppargc1b | ENSMUSG( | ENSMUSG( | 0.154307 | 0.462691 | NO | NO | NA |
| Zswim9 | ENSMUSG( | ENSMUSG( | -0.34846 | 0.007335 | YES | NO | NA |
| ENSMUSG( | ENSMUSG( | ENSMUSG( | 0.52463 | 0.001458 | YES | NO | NA |
| Insl6 | ENSMUSG( | ENSMUSG( | -0.03434 | 0.771401 | NO | NO | NA |
| Tesk2 | ENSMUSG( | ENSMUSG( | -0.41149 | 0.001045 | YES | NO | NA |
| Zfp948 | ENSMUSG( | ENSMUSG( | 0.116573 | 0.493174 | NO | NO | NA |
| Creb3l2 | ENSMUSG( | ENSMUSG( | -0.30527 | 0.012752 | YES | NO | NA |
| Osgepl1 | ENSMUSG( | ENSMUSG( | -0.00496 | 0.983133 | NO | NO | NA |
| Prdm16 | ENSMUSG( | ENSMUSG( | -0.30655 | 0.352248 | NO | NO | NA |
| Zfp709 | ENSMUSG( | ENSMUSG( | -0.09138 | 0.490349 | NO | NO | NA |
| Trav5-4 | ENSMUSG( | ENSMUSG( | -0.72552 | 0.300133 | NO | NO | NA |
| Vmn2r76 | ENSMUSG( | ENSMUSG( | 0.574225 | 0.628823 | NO | NO | NA |
| Dglucy | ENSMUSG( | ENSMUSG( | 0.458409 | 0.067934 | NO | NO | NA |
| Trav14n-1 | ENSMUSG( | ENSMUSG( | -0.60335 | 0.574263 | NO | NO | NA |
| Gm20008 | ENSMUSG( | ENSMUSG( | -0.09479 | 0.616746 | NO | NO | NA |
| Srxn1 | ENSMUSG( | ENSMUSG( | 0.151802 | 0.484899 | NO | NO | NA |
| Cep290 | ENSMUSG( | ENSMUSG( | 0.919605 | 0.00057 | YES | NO | NA |
| Myef2l | ENSMUSG( | ENSMUSG( | 0.024767 | 0.915152 | NO | NO | NA |
| Prkar2b | ENSMUSG( | ENSMUSG( | 0.158811 | 0.493751 | NO | NO | NA |
| Zfp27 | ENSMUSG( | ENSMUSG( | -0.06473 | 0.726446 | NO | NO | NA |
| Vegfb | ENSMUSG( | ENSMUSG( | 0.107546 | 0.378429 | NO | NO | NA |
| Mettl8 | ENSMUSG( | ENSMUSG( | 0.010374 | 0.952204 | NO | NO | NA |
| Gm46519 | ENSMUSG( | ENSMUSG( | -0.86845 | 0.000229 | YES | NO | NA |
| Angel1 | ENSMUSG( | ENSMUSG( | -0.2082 | 0.149599 | NO | NO | NA |
| Ctu1 | ENSMUSG( | ENSMUSG( | -0.18663 | 0.240544 | NO | NO | NA |
| Mypopos | ENSMUSG( | ENSMUSG( | -0.50533 | 2.58E-05 | YES | NO | NA |
| Gm47079 | ENSMUSG( | ENSMUSG( | -0.42347 | 0.009596 | YES | NO | NA |
| Lef1os1 | ENSMUSG( | ENSMUSG( | 0.528218 | 0.064125 | NO | NO | NA |
| Apip | ENSMUSG( | ENSMUSG( | 0.343264 | 0.017842 | YES | NO | NA |
| Il20rb | ENSMUSG( | ENSMUSG( | 0.78987 | 0.002509 | YES | NO | NA |
| Lratd2 | ENSMUSG( | ENSMUSG( | -0.21102 | 0.131992 | NO | NO | NA |
| Zfp69 | ENSMUSG( | ENSMUSG( | -0.37452 | 0.003819 | YES | NO | NA |
| Dsp | ENSMUSG( | ENSMUSG( | -1.3027 | 7.59E-05 | YES | YES | Downregulated |
| Pak6 | ENSMUSG( | ENSMUSG( | -0.98396 | 2.81E-05 | YES | NO | NA |
| Gt(ROSA)2 | ENSMUSG( | ENSMUSG( | 0.241639 | 0.054911 | NO | NO | NA |
| Has3 | ENSMUSG( | ENSMUSG( | -0.14839 | 0.181203 | NO | NO | NA |
| Isoc2b | ENSMUSG( | ENSMUSG( | 0.189478 | 0.373608 | NO | NO | NA |
| Igf2bp3 | ENSMUSG( | ENSMUSG( | 0.079369 | 0.68133 | NO | NO | NA |
| 4931406C( | ENSMUSG( | ENSMUSG( | 0.527035 | 5.88E-05 | YES | NO | NA |
| Smim4 | ENSMUSG( | ENSMUSG( | 0.489452 | 0.000232 | YES | NO | NA |
| Myh10 | ENSMUSG( | ENSMUSG( | 0.865194 | 0.000632 | YES | NO | NA |
| 2810002D | ENSMUSG( | ENSMUSG( | -0.71267 | 1.65E-07 | YES | NO | NA |

| | | | | | | | |
|---|---|---|---|---|---|---|---|
| Gm17494 | ENSMUSG( | ENSMUSG( | 0.145097 | 0.494513 | NO | NO | NA |
| Trim36 | ENSMUSG( | ENSMUSG( | -0.19615 | 0.056991 | NO | NO | NA |
| Slc9b2 | ENSMUSG( | ENSMUSG( | 2.478939 | 9.80E-25 | YES | YES | Upregulated |
| B130055N | ENSMUSG( | ENSMUSG( | -0.48022 | 2.79E-06 | YES | NO | NA |
| Commd9 | ENSMUSG( | ENSMUSG( | 0.245424 | 0.112037 | NO | NO | NA |
| Gm10603 | ENSMUSG( | ENSMUSG( | -0.11002 | 0.6937 | NO | NO | NA |
| Gm9752 | ENSMUSG( | ENSMUSG( | -0.08215 | 0.700632 | NO | NO | NA |
| Slc25a33 | ENSMUSG( | ENSMUSG( | -0.42482 | 0.003283 | YES | NO | NA |
| Tmem140 | ENSMUSG( | ENSMUSG( | -0.76391 | 3.59E-06 | YES | NO | NA |
| Tamalin | ENSMUSG( | ENSMUSG( | 0.243419 | 0.062635 | NO | NO | NA |
| Ift74 | ENSMUSG( | ENSMUSG( | 1.273778 | 6.02E-05 | YES | YES | Upregulated |
| Gm49322 | ENSMUSG( | ENSMUSG( | 0.151548 | 0.522926 | NO | NO | NA |
| Ifit3b | ENSMUSG( | ENSMUSG( | -1.47809 | 1.11E-11 | YES | YES | Downregulated |
| 4931414P: | ENSMUSG( | ENSMUSG( | -0.27171 | 0.011096 | YES | NO | NA |
| Rdh14 | ENSMUSG( | ENSMUSG( | -0.12782 | 0.321717 | NO | NO | NA |
| ENSMUSG | ENSMUSG( | ENSMUSG( | -0.39813 | 0.007215 | YES | NO | NA |
| Actr6 | ENSMUSG( | ENSMUSG( | 0.0485 | 0.691734 | NO | NO | NA |
| Meak7 | ENSMUSG( | ENSMUSG( | 0.066094 | 0.628904 | NO | NO | NA |
| Zfp940 | ENSMUSG( | ENSMUSG( | 0.167995 | 0.355386 | NO | NO | NA |
| 6330403K( | ENSMUSG( | ENSMUSG( | -0.96137 | 1.45E-07 | YES | NO | NA |
| Tanc1 | ENSMUSG( | ENSMUSG( | -0.25887 | 0.054682 | NO | NO | NA |
| Zkscan5 | ENSMUSG( | ENSMUSG( | -0.03235 | 0.85452 | NO | NO | NA |
| Nek4 | ENSMUSG( | ENSMUSG( | 0.692696 | 8.39E-06 | YES | NO | NA |
| ENSMUSG( | ENSMUSG( | ENSMUSG( | 0.180526 | 0.422508 | NO | NO | NA |
| ENSMUSG( | ENSMUSG( | ENSMUSG( | 0.16686 | 0.665493 | NO | NO | NA |
| Yae1d1 | ENSMUSG( | ENSMUSG( | 0.107761 | 0.352248 | NO | NO | NA |
| Tmem81 | ENSMUSG( | ENSMUSG( | -0.41046 | 6.37E-05 | YES | NO | NA |
| Ankrd13d | ENSMUSG( | ENSMUSG( | -0.12928 | 0.59843 | NO | NO | NA |
| Tln2 | ENSMUSG( | ENSMUSG( | 1.303381 | 1.24E-08 | YES | YES | Upregulated |
| Gatb | ENSMUSG( | ENSMUSG( | 0.287392 | 0.099093 | NO | NO | NA |
| Gm16001 | ENSMUSG( | ENSMUSG( | 0.746727 | 0.000399 | YES | NO | NA |
| 1700019D | ENSMUSG( | ENSMUSG( | 0.619112 | 0.005833 | YES | NO | NA |
| Ankrd13b | ENSMUSG( | ENSMUSG( | 1.46833 | 4.98E-10 | YES | YES | Upregulated |
| Pltp | ENSMUSG( | ENSMUSG( | -0.90301 | 1.66E-11 | YES | NO | NA |
| Zfp316 | ENSMUSG( | ENSMUSG( | -0.0507 | 0.736541 | NO | NO | NA |
| Zfp420 | ENSMUSG( | ENSMUSG( | -0.35223 | 0.018225 | YES | NO | NA |
| Gpr137 | ENSMUSG( | ENSMUSG( | -0.20694 | 0.168084 | NO | NO | NA |
| Hilpda | ENSMUSG( | ENSMUSG( | 0.507578 | 0.00185 | YES | NO | NA |
| Daam1 | ENSMUSG( | ENSMUSG( | 0.543726 | 0.035616 | YES | NO | NA |
| Hspb6 | ENSMUSG( | ENSMUSG( | 0.572729 | 0.001382 | YES | NO | NA |
| Slc26a6 | ENSMUSG( | ENSMUSG( | -0.26476 | 0.012503 | YES | NO | NA |
| Gm49190 | ENSMUSG( | ENSMUSG( | -0.07218 | 0.583918 | NO | NO | NA |
| Trbv31 | ENSMUSG( | ENSMUSG( | -0.52186 | 0.310173 | NO | NO | NA |
| Crat | ENSMUSG( | ENSMUSG( | -0.30159 | 0.035454 | YES | NO | NA |
| Ankrd35 | ENSMUSG( | ENSMUSG( | 0.795475 | 0.001971 | YES | NO | NA |
| Armc6 | ENSMUSG( | ENSMUSG( | 0.20477 | 0.183847 | NO | NO | NA |
| Ccnj | ENSMUSG( | ENSMUSG( | -0.15791 | 0.259206 | NO | NO | NA |

| | | | | | | | |
|---|---|---|---|---|---|---|---|
| ENSMUSG | ENSMUSG( | ENSMUSG( | 0.78533 | 6.25E-09 | YES | NO | NA |
| Arl4d | ENSMUSG( | ENSMUSG( | 0.072753 | 0.629973 | NO | NO | NA |
| Tcea2 | ENSMUSG( | ENSMUSG( | 0.239354 | 0.041695 | YES | NO | NA |
| Zfp365 | ENSMUSG( | ENSMUSG( | -0.27194 | 0.038375 | YES | NO | NA |
| Igha | ENSMUSG( | ENSMUSG( | -1.90241 | 0.386118 | NO | NO | NA |
| Mir17hg | ENSMUSG( | ENSMUSG( | 0.327743 | 0.000645 | YES | NO | NA |
| Rnf135 | ENSMUSG( | ENSMUSG( | -0.35669 | 0.00135 | YES | NO | NA |
| Cdkl3 | ENSMUSG( | ENSMUSG( | 0.604469 | 0.000608 | YES | NO | NA |
| A930005H | ENSMUSG( | ENSMUSG( | 0.020128 | 0.888289 | NO | NO | NA |
| Pigp | ENSMUSG( | ENSMUSG( | -0.01699 | 0.901746 | NO | NO | NA |
| Bcs1l | ENSMUSG( | ENSMUSG( | 0.053497 | 0.701427 | NO | NO | NA |
| Gm17251 | ENSMUSG( | ENSMUSG( | 0.127739 | 0.427013 | NO | NO | NA |
| A930015D | ENSMUSG( | ENSMUSG( | -0.4291 | 0.001802 | YES | NO | NA |
| Ypel2 | ENSMUSG( | ENSMUSG( | -0.12415 | 0.616138 | NO | NO | NA |
| Tesc | ENSMUSG( | ENSMUSG( | 0.087007 | 0.660272 | NO | NO | NA |
| Zfp764 | ENSMUSG( | ENSMUSG( | 0.015786 | 0.92009 | NO | NO | NA |
| Otud3 | ENSMUSG( | ENSMUSG( | -0.26729 | 0.09763 | NO | NO | NA |
| Gucy1b1 | ENSMUSG( | ENSMUSG( | 0.450289 | 0.015374 | YES | NO | NA |
| Uros | ENSMUSG( | ENSMUSG( | -0.35592 | 0.000208 | YES | NO | NA |
| Rsad1 | ENSMUSG( | ENSMUSG( | 0.676901 | 0.005294 | YES | NO | NA |
| Zfp82 | ENSMUSG( | ENSMUSG( | -0.45438 | 0.000327 | YES | NO | NA |
| Cd109 | ENSMUSG( | ENSMUSG( | 0.764741 | 0.014923 | YES | NO | NA |
| Gm19605 | ENSMUSG( | ENSMUSG( | -0.33718 | 0.019972 | YES | NO | NA |
| Rasal2 | ENSMUSG( | ENSMUSG( | 1.000263 | 2.62E-05 | YES | YES | Upregulated |
| Rpl9-ps6 | ENSMUSG( | ENSMUSG( | 0.282471 | 0.026743 | YES | NO | NA |
| Il1rl2 | ENSMUSG( | ENSMUSG( | 0.923887 | 3.86E-08 | YES | NO | NA |
| Ica1 | ENSMUSG( | ENSMUSG( | 0.288325 | 0.279785 | NO | NO | NA |
| Dnmbp | ENSMUSG( | ENSMUSG( | -0.36769 | 0.047201 | YES | NO | NA |
| Cd151 | ENSMUSG( | ENSMUSG( | -0.22538 | 0.158691 | NO | NO | NA |
| Numbl | ENSMUSG( | ENSMUSG( | -0.10277 | 0.614295 | NO | NO | NA |
| Fancb | ENSMUSG( | ENSMUSG( | 0.182869 | 0.158534 | NO | NO | NA |
| 4930500M | ENSMUSG( | ENSMUSG( | -0.04822 | 0.784601 | NO | NO | NA |
| Gm4739 | ENSMUSG( | ENSMUSG( | 0.326203 | 0.040091 | YES | NO | NA |
| Zbtb8a | ENSMUSG( | ENSMUSG( | -0.00837 | 0.946361 | NO | NO | NA |
| Anapc10 | ENSMUSG( | ENSMUSG( | -0.20861 | 0.27261 | NO | NO | NA |
| 2610507I0 | ENSMUSG( | ENSMUSG( | -0.00806 | 0.945736 | NO | NO | NA |
| Lancl3 | ENSMUSG( | ENSMUSG( | 0.201254 | 0.173088 | NO | NO | NA |
| Nars2 | ENSMUSG( | ENSMUSG( | 0.38628 | 0.00031 | YES | NO | NA |
| Nbdy | ENSMUSG( | ENSMUSG( | 0.128744 | 0.468476 | NO | NO | NA |
| Slc35d3 | ENSMUSG( | ENSMUSG( | 0.942227 | 0.00067 | YES | NO | NA |
| Trav12-2 | ENSMUSG( | ENSMUSG( | -0.52403 | 0.377938 | NO | NO | NA |
| Ethe1 | ENSMUSG( | ENSMUSG( | -0.4764 | 2.01E-06 | YES | NO | NA |
| Tubb6 | ENSMUSG( | ENSMUSG( | 1.188542 | 2.62E-10 | YES | YES | Upregulated |
| Dusp12 | ENSMUSG( | ENSMUSG( | 0.324854 | 0.005675 | YES | NO | NA |
| ENSMUSG( | ENSMUSG( | ENSMUSG( | 1.471577 | 0.002827 | YES | YES | Upregulated |
| Bloc1s5 | ENSMUSG( | ENSMUSG( | 0.27592 | 0.034193 | YES | NO | NA |
| Mex3d | ENSMUSG( | ENSMUSG( | 0.095867 | 0.536477 | NO | NO | NA |

| | | | | | | | |
|---|---|---|---|---|---|---|---|
| Atp6v1g2 | ENSMUSG( | ENSMUSG( | 1.06374 | 0.000137 | YES | YES | Upregulated |
| Rpl35a-ps( | ENSMUSG( | ENSMUSG( | 0.545366 | 0.791759 | NO | NO | NA |
| Trav9-4 | ENSMUSG( | ENSMUSG( | 0.485496 | 0.5354 | NO | NO | NA |
| Smpd5 | ENSMUSG( | ENSMUSG( | -0.36553 | 0.044653 | YES | NO | NA |
| Chrnb2 | ENSMUSG( | ENSMUSG( | -0.19266 | 0.183414 | NO | NO | NA |
| Tmem238 | ENSMUSG( | ENSMUSG( | 0.499428 | 0.000373 | YES | NO | NA |
| Socs5 | ENSMUSG( | ENSMUSG( | -0.44409 | 0.004723 | YES | NO | NA |
| Plscr3 | ENSMUSG( | ENSMUSG( | 0.221653 | 0.2625 | NO | NO | NA |
| ENSMUSG | ENSMUSG( | ENSMUSG( | -0.49512 | 0.000872 | YES | NO | NA |
| Tcf7l1 | ENSMUSG( | ENSMUSG( | -0.12543 | 0.590734 | NO | NO | NA |
| Meis3 | ENSMUSG( | ENSMUSG( | -0.4728 | 4.96E-06 | YES | NO | NA |
| Rrad | ENSMUSG( | ENSMUSG( | -0.21812 | 0.117079 | NO | NO | NA |
| Cfap410 | ENSMUSG( | ENSMUSG( | 0.492071 | 0.020035 | YES | NO | NA |
| Unc13a | ENSMUSG( | ENSMUSG( | 0.424699 | 0.034681 | YES | NO | NA |
| Itpkc | ENSMUSG( | ENSMUSG( | -0.31972 | 0.042521 | YES | NO | NA |
| Slc29a4 | ENSMUSG( | ENSMUSG( | -0.48736 | 0.000654 | YES | NO | NA |
| Gm21988 | ENSMUSG( | ENSMUSG( | -0.22179 | 0.157054 | NO | NO | NA |
| Gm37676 | ENSMUSG( | ENSMUSG( | 0.222479 | 0.085835 | NO | NO | NA |
| Miga2 | ENSMUSG( | ENSMUSG( | -0.11153 | 0.319978 | NO | NO | NA |
| Gm47260 | ENSMUSG( | ENSMUSG( | -0.22143 | 0.307303 | NO | NO | NA |
| Dnajc4 | ENSMUSG( | ENSMUSG( | 0.080391 | 0.61095 | NO | NO | NA |
| Rpsa-ps10 | ENSMUSG( | ENSMUSG( | 0.560882 | 0.011087 | YES | NO | NA |
| Cldn12 | ENSMUSG( | ENSMUSG( | -0.06026 | 0.775565 | NO | NO | NA |
| Parp16 | ENSMUSG( | ENSMUSG( | -0.30186 | 0.152532 | NO | NO | NA |
| Nit2 | ENSMUSG( | ENSMUSG( | 0.488679 | 6.80E-05 | YES | NO | NA |
| Tbc1d4 | ENSMUSG( | ENSMUSG( | -0.36937 | 0.341357 | NO | NO | NA |
| Stard9 | ENSMUSG( | ENSMUSG( | 0.62078 | 1.01E-08 | YES | NO | NA |
| Hmgb1-ps | ENSMUSG( | ENSMUSG( | -0.49765 | 0.002728 | YES | NO | NA |
| Gm47798 | ENSMUSG( | ENSMUSG( | -0.04961 | 0.801288 | NO | NO | NA |
| Gm45193 | ENSMUSG( | ENSMUSG( | -0.50732 | 0.012486 | YES | NO | NA |
| Tnfsf13b | ENSMUSG( | ENSMUSG( | 0.179249 | 0.343885 | NO | NO | NA |
| Gm37917 | ENSMUSG( | ENSMUSG( | 0.732861 | 1.39E-05 | YES | NO | NA |
| Ptrhd1 | ENSMUSG( | ENSMUSG( | 0.201415 | 0.05206 | NO | NO | NA |
| Ghdc | ENSMUSG( | ENSMUSG( | 0.083151 | 0.562501 | NO | NO | NA |
| Sord | ENSMUSG( | ENSMUSG( | -0.21033 | 0.051947 | NO | NO | NA |
| Xcl1 | ENSMUSG( | ENSMUSG( | 0.982632 | 0.000127 | YES | NO | NA |
| Rapgef5 | ENSMUSG( | ENSMUSG( | -0.13242 | 0.422853 | NO | NO | NA |
| Bivm | ENSMUSG( | ENSMUSG( | 0.342501 | 0.018023 | YES | NO | NA |
| Ccdc142 | ENSMUSG( | ENSMUSG( | 0.0052 | 0.979835 | NO | NO | NA |
| Snrpert | ENSMUSG( | ENSMUSG( | 0.335155 | 0.001572 | YES | NO | NA |
| Fah | ENSMUSG( | ENSMUSG( | 0.057474 | 0.731835 | NO | NO | NA |
| Nnt ENSM | ENSMUSG( | ENSMUSG( | -0.27883 | 0.518917 | NO | NO | NA |
| Gm11110 | ENSMUSG( | ENSMUSG( | -0.08074 | 0.714548 | NO | NO | NA |
| Hmox1 | ENSMUSG( | ENSMUSG( | -0.40672 | 0.012005 | YES | NO | NA |
| Trav7-6 | ENSMUSG( | ENSMUSG( | -1.00511 | 0.230439 | NO | NO | NA |
| Pde8a | ENSMUSG( | ENSMUSG( | 0.151402 | 0.456867 | NO | NO | NA |
| Rpusd2 | ENSMUSG( | ENSMUSG( | 0.371356 | 0.00011 | YES | NO | NA |

| | | | | | | | |
|---|---|---|---|---|---|---|---|
| Zfp994 | ENSMUSG( | ENSMUSG( | -0.22125 | 0.153972 | NO | NO | NA |
| Gm43302 | ENSMUSG( | ENSMUSG( | -9.86896 | 0.000318 | YES | YES | Downregulated |
| 3110056K( | ENSMUSG( | ENSMUSG( | -0.35244 | 0.006717 | YES | NO | NA |
| 1810055G | ENSMUSG( | ENSMUSG( | 0.249565 | 0.091827 | NO | NO | NA |
| Setd4 | ENSMUSG( | ENSMUSG( | 0.349649 | 0.000373 | YES | NO | NA |
| E430024P1 | ENSMUSG( | ENSMUSG( | 0.400105 | 0.036253 | YES | NO | NA |
| Ndnf | ENSMUSG( | ENSMUSG( | -0.45265 | 0.009919 | YES | NO | NA |
| A130010J1 | ENSMUSG( | ENSMUSG( | 0.212267 | 0.104551 | NO | NO | NA |
| Rtkn | ENSMUSG( | ENSMUSG( | 0.699335 | 0.011977 | YES | NO | NA |
| Pls1 | ENSMUSG( | ENSMUSG( | 1.59067 | 4.24E-05 | YES | YES | Upregulated |
| Sgms2 | ENSMUSG( | ENSMUSG( | 1.889732 | 0.051125 | NO | NO | NA |
| Hlf | ENSMUSG( | ENSMUSG( | 0.679301 | 0.000447 | YES | NO | NA |
| Rpgrip1l | ENSMUSG( | ENSMUSG( | -0.05705 | 0.781484 | NO | NO | NA |
| Gm10033 | ENSMUSG( | ENSMUSG( | -0.13527 | 0.454181 | NO | NO | NA |
| H1f5 | ENSMUSG( | ENSMUSG( | 0.177983 | 0.263297 | NO | NO | NA |
| Gm45837 | ENSMUSG( | ENSMUSG( | -1.28072 | 0.669294 | NO | NO | NA |
| Zfp605 | ENSMUSG( | ENSMUSG( | -0.27449 | 0.125021 | NO | NO | NA |
| Slc25a42 | ENSMUSG( | ENSMUSG( | 0.066842 | 0.662767 | NO | NO | NA |
| Rhebl1 | ENSMUSG( | ENSMUSG( | 0.261299 | 0.175573 | NO | NO | NA |
| Mtrfr | ENSMUSG( | ENSMUSG( | -0.19626 | 0.255342 | NO | NO | NA |
| Pik3c2b | ENSMUSG( | ENSMUSG( | 0.241649 | 0.251042 | NO | NO | NA |
| Trav9-2 | ENSMUSG( | ENSMUSG( | 0.499907 | 0.38454 | NO | NO | NA |
| Gm11346 | ENSMUSG( | ENSMUSG( | -1.25398 | 2.70E-05 | YES | YES | Downregulated |
| I730030J2 | ENSMUSG( | ENSMUSG( | 0.25585 | 0.319978 | NO | NO | NA |
| ENSMUSG | ENSMUSG( | ENSMUSG( | 0.29477 | 0.257899 | NO | NO | NA |
| Sowahc | ENSMUSG( | ENSMUSG( | 1.087929 | 1.76E-06 | YES | YES | Upregulated |
| Cradd | ENSMUSG( | ENSMUSG( | 0.034839 | 0.849155 | NO | NO | NA |
| Alkbh7 | ENSMUSG( | ENSMUSG( | 0.076008 | 0.633815 | NO | NO | NA |
| 2810402E1 | ENSMUSG( | ENSMUSG( | 0.4253 | 0.048922 | YES | NO | NA |
| Gm26533 | ENSMUSG( | ENSMUSG( | -0.26187 | 0.047007 | YES | NO | NA |
| Rcan1 | ENSMUSG( | ENSMUSG( | 0.097524 | 0.472454 | NO | NO | NA |
| Spta1 | ENSMUSG( | ENSMUSG( | 0.929119 | 0.092991 | NO | NO | NA |
| Phtf1os | ENSMUSG( | ENSMUSG( | -0.22174 | 0.107955 | NO | NO | NA |
| Atg14 | ENSMUSG( | ENSMUSG( | 0.045783 | 0.819148 | NO | NO | NA |
| Pfkfb2 | ENSMUSG( | ENSMUSG( | -0.25807 | 0.095513 | NO | NO | NA |
| ENSMUSG | ENSMUSG( | ENSMUSG( | 0.277741 | 0.12252 | NO | NO | NA |
| Rab2b | ENSMUSG( | ENSMUSG( | -0.46846 | 0.003532 | YES | NO | NA |
| Kantr | ENSMUSG( | ENSMUSG( | -0.6032 | 2.43E-06 | YES | NO | NA |
| Luzp1 | ENSMUSG( | ENSMUSG( | 0.392476 | 0.05785 | NO | NO | NA |
| Gm37305 | ENSMUSG( | ENSMUSG( | 0.443817 | 0.094611 | NO | NO | NA |
| Bbs12 | ENSMUSG( | ENSMUSG( | 0.045984 | 0.799499 | NO | NO | NA |
| Pars2 | ENSMUSG( | ENSMUSG( | 0.103313 | 0.455451 | NO | NO | NA |
| Eldr | ENSMUSG( | ENSMUSG( | -0.25488 | 0.083102 | NO | NO | NA |
| Nt5dc2 | ENSMUSG( | ENSMUSG( | 0.056983 | 0.819022 | NO | NO | NA |
| Fam72a | ENSMUSG( | ENSMUSG( | 0.173868 | 0.227099 | NO | NO | NA |
| Rpl35a-ps4 | ENSMUSG( | ENSMUSG( | 1.071473 | 0.582357 | NO | NO | NA |
| Caprin2 | ENSMUSG( | ENSMUSG( | -0.34425 | 0.05543 | NO | NO | NA |

| | | | | | | | |
|---|---|---|---|---|---|---|---|
| Trmt12 | ENSMUSG( | ENSMUSG( | 0.11198 | 0.45087 | NO | NO | NA |
| Ndor1 ENS | ENSMUSG( | ENSMUSG( | -0.05873 | 0.816153 | NO | NO | NA |
| Uprt | ENSMUSG( | ENSMUSG( | -0.39046 | 0.000511 | YES | NO | NA |
| Frat1 | ENSMUSG( | ENSMUSG( | -0.12493 | 0.464388 | NO | NO | NA |
| Scrn2 | ENSMUSG( | ENSMUSG( | 0.287799 | 0.04408 | YES | NO | NA |
| Slc17a5 | ENSMUSG( | ENSMUSG( | -0.07427 | 0.638532 | NO | NO | NA |
| Gm21811 | ENSMUSG( | ENSMUSG( | -0.01926 | 0.930403 | NO | NO | NA |
| Igsf9b | ENSMUSG( | ENSMUSG( | -0.0813 | 0.826005 | NO | NO | NA |
| Blvrb | ENSMUSG( | ENSMUSG( | 0.5528 | 0.083438 | NO | NO | NA |
| Trmt61b | ENSMUSG( | ENSMUSG( | 0.224503 | 0.072067 | NO | NO | NA |
| Tpm3-rs7 | ENSMUSG( | ENSMUSG( | 0.220051 | 0.142318 | NO | NO | NA |
| Rsl1 | ENSMUSG( | ENSMUSG( | -0.51154 | 0.000773 | YES | NO | NA |
| Ebf1 | ENSMUSG( | ENSMUSG( | 0.834467 | 0.280619 | NO | NO | NA |
| Gm14440 | ENSMUSG( | ENSMUSG( | -0.2231 | 0.141657 | NO | NO | NA |
| Gm42547 | ENSMUSG( | ENSMUSG( | -0.7643 | 9.02E-06 | YES | NO | NA |
| Zfp446 | ENSMUSG( | ENSMUSG( | 0.284472 | 0.063735 | NO | NO | NA |
| Ehhadh | ENSMUSG( | ENSMUSG( | -0.19751 | 0.254615 | NO | NO | NA |
| Gpr155 | ENSMUSG( | ENSMUSG( | -0.57979 | 0.000241 | YES | NO | NA |
| Hemgn | ENSMUSG( | ENSMUSG( | 0.767694 | 0.250576 | NO | NO | NA |
| Rhbdd3 | ENSMUSG( | ENSMUSG( | 0.573741 | 7.32E-07 | YES | NO | NA |
| Rn7sk | ENSMUSG( | ENSMUSG( | 0.864153 | 0.140105 | NO | NO | NA |
| Dolk | ENSMUSG( | ENSMUSG( | -0.02687 | 0.843786 | NO | NO | NA |
| Derpc | ENSMUSG( | ENSMUSG( | -0.27063 | 0.180932 | NO | NO | NA |
| Trpc1 | ENSMUSG( | ENSMUSG( | 0.407853 | 0.256634 | NO | NO | NA |
| Gm45799 | ENSMUSG( | ENSMUSG( | -0.19787 | 0.487209 | NO | NO | NA |
| Uhrf1bp1 | ENSMUSG( | ENSMUSG( | 0.221389 | 0.065232 | NO | NO | NA |
| Fastkd5 | ENSMUSG( | ENSMUSG( | -0.01723 | 0.925531 | NO | NO | NA |
| Shq1 | ENSMUSG( | ENSMUSG( | 0.358637 | 0.093565 | NO | NO | NA |
| Xkr8 | ENSMUSG( | ENSMUSG( | -0.42052 | 0.000761 | YES | NO | NA |
| Get1 | ENSMUSG( | ENSMUSG( | 0.421827 | 0.011833 | YES | NO | NA |
| Patj | ENSMUSG( | ENSMUSG( | 0.561259 | 0.00374 | YES | NO | NA |
| Zfp712 | ENSMUSG( | ENSMUSG( | -0.46449 | 0.000251 | YES | NO | NA |
| Celsr2 | ENSMUSG( | ENSMUSG( | -0.35274 | 0.112821 | NO | NO | NA |
| Dctd | ENSMUSG( | ENSMUSG( | 0.744775 | 5.05E-05 | YES | NO | NA |
| D930048N | ENSMUSG( | ENSMUSG( | -0.44356 | 1.20E-05 | YES | NO | NA |
| Zfp606 | ENSMUSG( | ENSMUSG( | -0.19383 | 0.159746 | NO | NO | NA |
| Sdhaf4 | ENSMUSG( | ENSMUSG( | -0.35995 | 0.002037 | YES | NO | NA |
| Zfp808 | ENSMUSG( | ENSMUSG( | -0.49042 | 1.14E-05 | YES | NO | NA |
| Ankrd29 | ENSMUSG( | ENSMUSG( | 0.140816 | 0.530191 | NO | NO | NA |
| Gm20683 | ENSMUSG( | ENSMUSG( | -0.15739 | 0.582542 | NO | NO | NA |
| B3galt6 | ENSMUSG( | ENSMUSG( | 0.106944 | 0.474914 | NO | NO | NA |
| Acy1 | ENSMUSG( | ENSMUSG( | 0.202386 | 0.221443 | NO | NO | NA |
| Ext1 | ENSMUSG( | ENSMUSG( | -0.00412 | 0.97899 | NO | NO | NA |
| Coq8a | ENSMUSG( | ENSMUSG( | -0.1217 | 0.595991 | NO | NO | NA |
| Cxcl10 | ENSMUSG( | ENSMUSG( | -0.38935 | 0.192831 | NO | NO | NA |
| Dnal4 | ENSMUSG( | ENSMUSG( | 0.432176 | 0.050442 | NO | NO | NA |
| Gm6712 | ENSMUSG( | ENSMUSG( | -0.26384 | 0.0825 | NO | NO | NA |

| | | | | | | | |
|---|---|---|---|---|---|---|---|
| Pacc1 | ENSMUSG( | ENSMUSG( | -0.0083 | 0.954374 | NO | NO | NA |
| 2310009B | ENSMUSG( | ENSMUSG( | 0.491208 | 0.014136 | YES | NO | NA |
| Acacb | ENSMUSG( | ENSMUSG( | -0.19434 | 0.315494 | NO | NO | NA |
| Cry2 | ENSMUSG( | ENSMUSG( | -0.12495 | 0.423332 | NO | NO | NA |
| H2bc21 | ENSMUSG( | ENSMUSG( | 0.250702 | 0.223494 | NO | NO | NA |
| Il10 | ENSMUSG( | ENSMUSG( | 1.12999 | 0.000673 | YES | YES | Upregulated |
| Nrn1 | ENSMUSG( | ENSMUSG( | 1.388241 | 6.02E-05 | YES | YES | Upregulated |
| Zfp689 | ENSMUSG( | ENSMUSG( | -0.02333 | 0.856625 | NO | NO | NA |
| Nrxn1 | ENSMUSG( | ENSMUSG( | 0.773564 | 0.513114 | NO | NO | NA |
| Lrrc49 | ENSMUSG( | ENSMUSG( | 1.266071 | 0.000217 | YES | YES | Upregulated |
| Lmo4 | ENSMUSG( | ENSMUSG( | -0.28017 | 0.054149 | NO | NO | NA |
| Fdxacb1 | ENSMUSG( | ENSMUSG( | 0.267966 | 0.107933 | NO | NO | NA |
| Pctp | ENSMUSG( | ENSMUSG( | 0.269637 | 0.150925 | NO | NO | NA |
| Tnfrsf13b | ENSMUSG( | ENSMUSG( | -0.26486 | 0.166298 | NO | NO | NA |
| Sestd1 | ENSMUSG( | ENSMUSG( | 1.332705 | 7.20E-08 | YES | YES | Upregulated |
| Pomk | ENSMUSG( | ENSMUSG( | 0.036436 | 0.809587 | NO | NO | NA |
| Mks1 | ENSMUSG( | ENSMUSG( | 0.349087 | 0.041787 | YES | NO | NA |
| Tcaim | ENSMUSG( | ENSMUSG( | -0.18042 | 0.359812 | NO | NO | NA |
| Specc1 | ENSMUSG( | ENSMUSG( | 0.16268 | 0.394041 | NO | NO | NA |
| Nebl | ENSMUSG( | ENSMUSG( | 1.520663 | 2.18E-07 | YES | YES | Upregulated |
| Ms4a6c | ENSMUSG( | ENSMUSG( | -0.581 | 1.53E-05 | YES | NO | NA |
| Sec22a | ENSMUSG( | ENSMUSG( | 0.109472 | 0.478634 | NO | NO | NA |
| Gm43118 | ENSMUSG( | ENSMUSG( | 0.333964 | 0.071198 | NO | NO | NA |
| Cpm | ENSMUSG( | ENSMUSG( | -0.10175 | 0.684648 | NO | NO | NA |
| Nagk | ENSMUSG( | ENSMUSG( | -0.18397 | 0.274152 | NO | NO | NA |
| Cbarp | ENSMUSG( | ENSMUSG( | -0.13235 | 0.481807 | NO | NO | NA |
| Gm9917 | ENSMUSG( | ENSMUSG( | 0.001374 | 0.992393 | NO | NO | NA |
| Edaradd | ENSMUSG( | ENSMUSG( | 0.589993 | 0.02416 | YES | NO | NA |
| Zrsr1 | ENSMUSG( | ENSMUSG( | -0.44463 | 0.024441 | YES | NO | NA |
| Tmem265 | ENSMUSG( | ENSMUSG( | -0.02543 | 0.9296 | NO | NO | NA |
| Abhd6 | ENSMUSG( | ENSMUSG( | -0.18422 | 0.367207 | NO | NO | NA |
| Csf2ra | ENSMUSG( | ENSMUSG( | 0.972458 | 6.80E-08 | YES | NO | NA |
| Gm49364 | ENSMUSG( | ENSMUSG( | -0.45397 | 0.18403 | NO | NO | NA |
| Sult2b1 | ENSMUSG( | ENSMUSG( | 0.135727 | 0.392394 | NO | NO | NA |
| Trmo | ENSMUSG( | ENSMUSG( | -0.00129 | 0.992533 | NO | NO | NA |
| ENSMUSG( | ENSMUSG( | ENSMUSG( | -0.13017 | 0.450534 | NO | NO | NA |
| Angptl4 | ENSMUSG( | ENSMUSG( | 0.745581 | 1.15E-06 | YES | NO | NA |
| Fam174b | ENSMUSG( | ENSMUSG( | -0.67979 | 0.004871 | YES | NO | NA |
| BC024978 | ENSMUSG( | ENSMUSG( | -0.22486 | 0.113352 | NO | NO | NA |
| 2900076A | ENSMUSG( | ENSMUSG( | -0.36216 | 0.104897 | NO | NO | NA |
| Zfp825 | ENSMUSG( | ENSMUSG( | -0.32002 | 0.023284 | YES | NO | NA |
| Rfesd | ENSMUSG( | ENSMUSG( | -0.08529 | 0.585511 | NO | NO | NA |
| Ikzf4 | ENSMUSG( | ENSMUSG( | -0.26265 | 0.501066 | NO | NO | NA |
| Meg3 | ENSMUSG( | ENSMUSG( | 0.595316 | 0.642462 | NO | NO | NA |
| Dlg3 | ENSMUSG( | ENSMUSG( | -0.72857 | 0.000574 | YES | NO | NA |
| Plxdc2 | ENSMUSG( | ENSMUSG( | 1.296668 | 1.11E-14 | YES | YES | Upregulated |
| Cpne2 | ENSMUSG( | ENSMUSG( | 0.572986 | 0.003688 | YES | NO | NA |

| | | | | | | | |
|---|---|---|---|---|---|---|---|
| Gm37422 | ENSMUSG( | ENSMUSG( | 1.00065 | 3.10E-05 | YES | YES | Upregulated |
| 1110065P: | ENSMUSG( | ENSMUSG( | 0.345122 | 0.019215 | YES | NO | NA |
| A130071D | ENSMUSG( | ENSMUSG( | 0.06263 | 0.854133 | NO | NO | NA |
| Gm49599 | ENSMUSG( | ENSMUSG( | -0.17567 | 0.297659 | NO | NO | NA |
| Ss18l2 | ENSMUSG( | ENSMUSG( | 0.152044 | 0.343011 | NO | NO | NA |
| Mgrn1 | ENSMUSG( | ENSMUSG( | 0.385228 | 0.012161 | YES | NO | NA |
| Etohd2 | ENSMUSG( | ENSMUSG( | -0.29406 | 0.100152 | NO | NO | NA |
| Commd1b | ENSMUSG( | ENSMUSG( | 0.463987 | 0.70818 | NO | NO | NA |
| Nfib | ENSMUSG( | ENSMUSG( | 0.657541 | 0.582357 | NO | NO | NA |
| Gm12166 | ENSMUSG( | ENSMUSG( | -0.29468 | 0.025269 | YES | NO | NA |
| Dynlt1a | ENSMUSG( | ENSMUSG( | 0.136189 | 0.336832 | NO | NO | NA |
| Xntrpc | ENSMUSG( | ENSMUSG( | 0.146544 | 0.449643 | NO | NO | NA |
| Gm42748 | ENSMUSG( | ENSMUSG( | 0.106658 | 0.584094 | NO | NO | NA |
| Gm10167 | ENSMUSG( | ENSMUSG( | 0.311126 | 0.094593 | NO | NO | NA |
| Plpp2 | ENSMUSG( | ENSMUSG( | 0.181654 | 0.247188 | NO | NO | NA |
| Etfrf1 | ENSMUSG( | ENSMUSG( | -0.19573 | 0.372394 | NO | NO | NA |
| Cryl1 | ENSMUSG( | ENSMUSG( | 0.646429 | 0.000307 | YES | NO | NA |
| Slc44a1 | ENSMUSG( | ENSMUSG( | 0.626736 | 2.01E-06 | YES | NO | NA |
| Serpinb1a | ENSMUSG( | ENSMUSG( | 1.074232 | 0.002474 | YES | YES | Upregulated |
| Nav2 | ENSMUSG( | ENSMUSG( | 0.499391 | 0.488166 | NO | NO | NA |
| B3gat2 | ENSMUSG( | ENSMUSG( | -0.31093 | 0.106919 | NO | NO | NA |
| Rpph1 | ENSMUSG( | ENSMUSG( | 0.835627 | 0.004522 | YES | NO | NA |
| Gm38287 | ENSMUSG( | ENSMUSG( | -0.65628 | 0.004707 | YES | NO | NA |
| Comtd1 | ENSMUSG( | ENSMUSG( | 0.091103 | 0.504561 | NO | NO | NA |
| Zfp799 | ENSMUSG( | ENSMUSG( | -0.39702 | 0.011875 | YES | NO | NA |
| Znrf3 | ENSMUSG( | ENSMUSG( | 0.097009 | 0.550072 | NO | NO | NA |
| Ttc32 | ENSMUSG( | ENSMUSG( | 0.127797 | 0.428872 | NO | NO | NA |
| Nlgn2 | ENSMUSG( | ENSMUSG( | 0.443062 | 0.004082 | YES | NO | NA |
| Wdr19 | ENSMUSG( | ENSMUSG( | 0.265326 | 0.506979 | NO | NO | NA |
| Top1mt | ENSMUSG( | ENSMUSG( | 0.716078 | 2.26E-05 | YES | NO | NA |
| Akap17b | ENSMUSG( | ENSMUSG( | 0.1214 | 0.430091 | NO | NO | NA |
| Pals1 | ENSMUSG( | ENSMUSG( | 0.182203 | 0.175992 | NO | NO | NA |
| Nradd | ENSMUSG( | ENSMUSG( | -0.23113 | 0.284217 | NO | NO | NA |
| Atrnl1 | ENSMUSG( | ENSMUSG( | 0.591091 | 0.001902 | YES | NO | NA |
| Timm10 | ENSMUSG( | ENSMUSG( | 0.344868 | 0.023932 | YES | NO | NA |
| Letm2 | ENSMUSG( | ENSMUSG( | -0.18965 | 0.150037 | NO | NO | NA |
| Gm43766 | ENSMUSG( | ENSMUSG( | 0.073739 | 0.62014 | NO | NO | NA |
| Acyp1 | ENSMUSG( | ENSMUSG( | -0.35816 | 0.00453 | YES | NO | NA |
| Marchf8 | ENSMUSG( | ENSMUSG( | 0.759291 | 0.000802 | YES | NO | NA |
| Gm21781 | ENSMUSG( | ENSMUSG( | -0.47791 | 0.011533 | YES | NO | NA |
| Bola1 | ENSMUSG( | ENSMUSG( | 0.823151 | 1.05E-05 | YES | NO | NA |
| Lsr | ENSMUSG( | ENSMUSG( | 0.311313 | 0.159326 | NO | NO | NA |
| Ccdc40 | ENSMUSG( | ENSMUSG( | 1.254814 | 1.16E-07 | YES | YES | Upregulated |
| Gm45606 | ENSMUSG( | ENSMUSG( | -0.51612 | 3.14E-05 | YES | NO | NA |
| Gm10676 | ENSMUSG( | ENSMUSG( | 0.272415 | 0.004737 | YES | NO | NA |
| ENSMUSG( | ENSMUSG( | ENSMUSG( | -0.17211 | 0.350484 | NO | NO | NA |
| Gm44164 | ENSMUSG( | ENSMUSG( | -0.38825 | 0.03176 | YES | NO | NA |

| | | | | | | | |
|---|---|---|---|---|---|---|---|
| Myo10 | ENSMUSG( | ENSMUSG( | 1.128082 | 9.35E-07 | YES | YES | Upregulated |
| Lipe | ENSMUSG( | ENSMUSG( | 0.12382 | 0.431995 | NO | NO | NA |
| Ttbk2 | ENSMUSG( | ENSMUSG( | 0.219951 | 0.364342 | NO | NO | NA |
| Gm49204 | ENSMUSG( | ENSMUSG( | -0.42178 | 0.006622 | YES | NO | NA |
| Map3k15 | ENSMUSG( | ENSMUSG( | -0.131 | 0.545964 | NO | NO | NA |
| 4930430F( | ENSMUSG( | ENSMUSG( | 0.082559 | 0.586288 | NO | NO | NA |
| Gm38118 | ENSMUSG( | ENSMUSG( | -0.8917 | 7.87E-09 | YES | NO | NA |
| Tmem177 | ENSMUSG( | ENSMUSG( | -0.0565 | 0.751111 | NO | NO | NA |
| Etfbkmt | ENSMUSG( | ENSMUSG( | -0.0664 | 0.713191 | NO | NO | NA |
| Add2 | ENSMUSG( | ENSMUSG( | 1.012424 | 0.093525 | NO | NO | NA |
| Gm8186 | ENSMUSG( | ENSMUSG( | 0.288722 | 0.120777 | NO | NO | NA |
| Trav12-3 | ENSMUSG( | ENSMUSG( | -0.23067 | 0.779851 | NO | NO | NA |
| Zfat | ENSMUSG( | ENSMUSG( | -0.42753 | 0.000869 | YES | NO | NA |
| Septin8 | ENSMUSG( | ENSMUSG( | 0.84157 | 0.049807 | YES | NO | NA |
| Gm20536 | ENSMUSG( | ENSMUSG( | -0.3478 | 0.007876 | YES | NO | NA |
| Cntd1 | ENSMUSG( | ENSMUSG( | 0.201392 | 0.428302 | NO | NO | NA |
| Gm49368 | ENSMUSG( | ENSMUSG( | -0.779 | 0.19471 | NO | NO | NA |
| Zfp58 | ENSMUSG( | ENSMUSG( | -0.21422 | 0.085625 | NO | NO | NA |
| C130046K: | ENSMUSG( | ENSMUSG( | -0.29423 | 0.005585 | YES | NO | NA |
| Rpusd3 | ENSMUSG( | ENSMUSG( | 0.140976 | 0.424383 | NO | NO | NA |
| Shld1 | ENSMUSG( | ENSMUSG( | -0.20184 | 0.072222 | NO | NO | NA |
| Zfp1 | ENSMUSG( | ENSMUSG( | -0.1294 | 0.368428 | NO | NO | NA |
| Trav9n-4 | ENSMUSG( | ENSMUSG( | -0.71624 | 0.218909 | NO | NO | NA |
| Ighv8-12 | ENSMUSG( | ENSMUSG( | 0.271432 | 0.885851 | NO | NO | NA |
| L2hgdh | ENSMUSG( | ENSMUSG( | 0.656151 | 2.42E-05 | YES | NO | NA |
| Gm26982 | ENSMUSG( | ENSMUSG( | 0.176608 | 0.322523 | NO | NO | NA |
| Fam220a E | ENSMUSG( | ENSMUSG( | -0.42909 | 0.606624 | NO | NO | NA |
| Ap5s1 | ENSMUSG( | ENSMUSG( | -0.17345 | 0.312475 | NO | NO | NA |
| Zfp473 | ENSMUSG( | ENSMUSG( | -0.08838 | 0.597779 | NO | NO | NA |
| Tceanc | ENSMUSG( | ENSMUSG( | 0.19906 | 0.250934 | NO | NO | NA |
| Nmnat2 | ENSMUSG( | ENSMUSG( | -0.01516 | 0.932809 | NO | NO | NA |
| Mettl25 | ENSMUSG( | ENSMUSG( | -0.04739 | 0.76235 | NO | NO | NA |
| Mrm3 | ENSMUSG( | ENSMUSG( | 0.695892 | 4.35E-05 | YES | NO | NA |
| Tmem198 | ENSMUSG( | ENSMUSG( | -0.02552 | 0.884247 | NO | NO | NA |
| Tsen2 | ENSMUSG( | ENSMUSG( | 0.706594 | 0.00048 | YES | NO | NA |
| Slc66a2 | ENSMUSG( | ENSMUSG( | 0.246104 | 0.154419 | NO | NO | NA |
| Proz | ENSMUSG( | ENSMUSG( | -0.11882 | 0.484832 | NO | NO | NA |
| Ddr1 | ENSMUSG( | ENSMUSG( | -0.16834 | 0.562405 | NO | NO | NA |
| Tbc1d30 | ENSMUSG( | ENSMUSG( | -0.46606 | 0.004641 | YES | NO | NA |
| Crip2 | ENSMUSG( | ENSMUSG( | -0.19488 | 0.446966 | NO | NO | NA |
| Pax6 | ENSMUSG( | ENSMUSG( | 0.622641 | 0.401017 | NO | NO | NA |
| Glyctk | ENSMUSG( | ENSMUSG( | -0.2133 | 0.176062 | NO | NO | NA |
| Lins1 | ENSMUSG( | ENSMUSG( | 0.092272 | 0.599709 | NO | NO | NA |
| Mir6236 | ENSMUSG( | ENSMUSG( | 0.986997 | 0.044621 | YES | NO | NA |
| Adat2 | ENSMUSG( | ENSMUSG( | 0.290402 | 0.094611 | NO | NO | NA |
| Krtcap3 | ENSMUSG( | ENSMUSG( | -0.04604 | 0.794623 | NO | NO | NA |
| Gm51425 | ENSMUSG( | ENSMUSG( | -0.2982 | 0.017427 | YES | NO | NA |

| | | | | | | | |
|---|---|---|---|---|---|---|---|
| Prorp | ENSMUSG( | ENSMUSG( | 0.340635 | 0.022036 | YES | NO | NA |
| Zfp97 | ENSMUSG( | ENSMUSG( | -0.31336 | 0.093869 | NO | NO | NA |
| Dcbld2 | ENSMUSG( | ENSMUSG( | 0.457196 | 0.010323 | YES | NO | NA |
| Nudt6 | ENSMUSG( | ENSMUSG( | 0.247561 | 0.129908 | NO | NO | NA |
| BC043934 | ENSMUSG( | ENSMUSG( | -0.55158 | 8.31E-05 | YES | NO | NA |
| Gm29585 | ENSMUSG( | ENSMUSG( | -0.17948 | 0.350806 | NO | NO | NA |
| Tctn3 | ENSMUSG( | ENSMUSG( | 0.081781 | 0.660508 | NO | NO | NA |
| Nxt2 | ENSMUSG( | ENSMUSG( | -0.23091 | 0.091968 | NO | NO | NA |
| Ntrk3 | ENSMUSG( | ENSMUSG( | 0.108493 | 0.819022 | NO | NO | NA |
| Swsap1 | ENSMUSG( | ENSMUSG( | 0.007023 | 0.976269 | NO | NO | NA |
| Ttc28 | ENSMUSG( | ENSMUSG( | 0.394627 | 0.012202 | YES | NO | NA |
| Apbb1 | ENSMUSG( | ENSMUSG( | -0.58336 | 0.000109 | YES | NO | NA |
| Gm11585 | ENSMUSG( | ENSMUSG( | 0.402889 | 0.015724 | YES | NO | NA |
| Gm20427 | ENSMUSG( | ENSMUSG( | -0.26276 | 0.374884 | NO | NO | NA |
| Snupn | ENSMUSG( | ENSMUSG( | 0.064472 | 0.722194 | NO | NO | NA |
| Gm27232 | ENSMUSG( | ENSMUSG( | -0.03264 | 0.842144 | NO | NO | NA |
| Gm28053 | ENSMUSG( | ENSMUSG( | -0.22865 | 0.127867 | NO | NO | NA |
| Gm49874 | ENSMUSG( | ENSMUSG( | -0.00344 | 0.985983 | NO | NO | NA |
| Ccdc138 | ENSMUSG( | ENSMUSG( | 0.266799 | 0.019536 | YES | NO | NA |
| Sh2d5 | ENSMUSG( | ENSMUSG( | 0.636511 | 0.001744 | YES | NO | NA |
| Skida1 | ENSMUSG( | ENSMUSG( | 0.063751 | 0.733152 | NO | NO | NA |
| Castor1 | ENSMUSG( | ENSMUSG( | 0.031094 | 0.83633 | NO | NO | NA |
| Cdk14 | ENSMUSG( | ENSMUSG( | -0.46948 | 0.05774 | NO | NO | NA |
| Cln8 | ENSMUSG( | ENSMUSG( | 0.242071 | 0.111049 | NO | NO | NA |
| Nudt14 | ENSMUSG( | ENSMUSG( | -0.15731 | 0.293903 | NO | NO | NA |
| Crybg2 | ENSMUSG( | ENSMUSG( | 1.123505 | 3.88E-07 | YES | YES | Upregulated |
| Moap1 | ENSMUSG( | ENSMUSG( | 0.553692 | 0.000189 | YES | NO | NA |
| Gm15337 | ENSMUSG( | ENSMUSG( | 0.026488 | 0.918145 | NO | NO | NA |
| Trav6d-3 | ENSMUSG( | ENSMUSG( | 0.93242 | 0.548076 | NO | NO | NA |
| Enc1 | ENSMUSG( | ENSMUSG( | 0.137074 | 0.301318 | NO | NO | NA |
| Mtres1 | ENSMUSG( | ENSMUSG( | 0.000897 | 0.996207 | NO | NO | NA |
| Zan | ENSMUSG( | ENSMUSG( | -0.13923 | 0.619095 | NO | NO | NA |
| Myl4 | ENSMUSG( | ENSMUSG( | 0.717424 | 8.03E-09 | YES | NO | NA |
| Mterf2 | ENSMUSG( | ENSMUSG( | -0.14984 | 0.435713 | NO | NO | NA |
| Gm8226 | ENSMUSG( | ENSMUSG( | 0.429298 | 0.000384 | YES | NO | NA |
| Spata24 | ENSMUSG( | ENSMUSG( | 0.776225 | 8.68E-09 | YES | NO | NA |
| Gcat ENSN | ENSMUSG( | ENSMUSG( | 1.2587 | 5.72E-05 | YES | YES | Upregulated |
| Kbtbd8 | ENSMUSG( | ENSMUSG( | -0.06054 | 0.762738 | NO | NO | NA |
| Cecr2 | ENSMUSG( | ENSMUSG( | 0.680624 | 0.307944 | NO | NO | NA |
| Ptgfrn | ENSMUSG( | ENSMUSG( | 1.358518 | 2.85E-06 | YES | YES | Upregulated |
| Sirt4 | ENSMUSG( | ENSMUSG( | -0.0136 | 0.933725 | NO | NO | NA |
| Vsig8 | ENSMUSG( | ENSMUSG( | 1.262137 | 0.019012 | YES | YES | Upregulated |
| ENSMUSG | ENSMUSG( | ENSMUSG( | -0.07458 | 0.784854 | NO | NO | NA |
| Oas2 | ENSMUSG( | ENSMUSG( | -1.85597 | 6.09E-10 | YES | YES | Downregulated |
| Mfsd2a | ENSMUSG( | ENSMUSG( | 0.5054 | 0.005409 | YES | NO | NA |
| Gm49783 | ENSMUSG( | ENSMUSG( | 0.196315 | 0.370814 | NO | NO | NA |
| 9430038I0 | ENSMUSG( | ENSMUSG( | 0.65394 | 3.31E-08 | YES | NO | NA |

| | | | | | | | |
|---|---|---|---|---|---|---|---|
| Gm32687 | ENSMUSG( | ENSMUSG( | -0.0014 | 0.995252 | NO | NO | NA |
| Sgsm3 | ENSMUSG( | ENSMUSG( | 0.325095 | 0.02423 | YES | NO | NA |
| Nmnat3 | ENSMUSG( | ENSMUSG( | 0.638032 | 0.000466 | YES | NO | NA |
| Tmem67 | ENSMUSG( | ENSMUSG( | 0.288027 | 0.254163 | NO | NO | NA |
| Airn | ENSMUSG( | ENSMUSG( | -0.73509 | 8.10E-05 | YES | NO | NA |
| Grk5 | ENSMUSG( | ENSMUSG( | -0.33473 | 0.112285 | NO | NO | NA |
| Ybey | ENSMUSG( | ENSMUSG( | 0.142994 | 0.57996 | NO | NO | NA |
| Zfp341 | ENSMUSG( | ENSMUSG( | -0.25345 | 0.005559 | YES | NO | NA |
| Spata2l | ENSMUSG( | ENSMUSG( | -0.30514 | 0.053715 | NO | NO | NA |
| Septin10 | ENSMUSG( | ENSMUSG( | 0.818081 | 0.000138 | YES | NO | NA |
| Slc25a14 | ENSMUSG( | ENSMUSG( | 0.097359 | 0.53935 | NO | NO | NA |
| Trav14-1 | ENSMUSG( | ENSMUSG( | -0.41013 | 0.544688 | NO | NO | NA |
| Hspbap1 | ENSMUSG( | ENSMUSG( | -0.29902 | 0.051113 | NO | NO | NA |
| 1700001O | ENSMUSG( | ENSMUSG( | -1.26514 | 0.001257 | YES | YES | Downregulated |
| Smarcd3 | ENSMUSG( | ENSMUSG( | -0.10166 | 0.540932 | NO | NO | NA |
| Pald1 | ENSMUSG( | ENSMUSG( | -0.47012 | 0.002634 | YES | NO | NA |
| Trav10 | ENSMUSG( | ENSMUSG( | -0.40645 | 0.503809 | NO | NO | NA |
| Auts2 | ENSMUSG( | ENSMUSG( | 0.478407 | 0.665475 | NO | NO | NA |
| Gm45885 | ENSMUSG( | ENSMUSG( | 6.232557 | NA | NO | NO | NA |
| Arsk | ENSMUSG( | ENSMUSG( | -0.26093 | 0.152569 | NO | NO | NA |
| Nav1 | ENSMUSG( | ENSMUSG( | 0.640476 | 0.512293 | NO | NO | NA |
| Gm43863 | ENSMUSG( | ENSMUSG( | -0.19114 | 0.28741 | NO | NO | NA |
| Tfcp2 | ENSMUSG( | ENSMUSG( | 0.133347 | 0.460383 | NO | NO | NA |
| Acad11 | ENSMUSG( | ENSMUSG( | 0.017384 | 0.926556 | NO | NO | NA |
| Gm9800 | ENSMUSG( | ENSMUSG( | 0.371217 | 0.140741 | NO | NO | NA |
| Tnfsf9 | ENSMUSG( | ENSMUSG( | -0.19926 | 0.390459 | NO | NO | NA |
| Gys1 | ENSMUSG( | ENSMUSG( | 0.040511 | 0.817665 | NO | NO | NA |
| Spryd7 | ENSMUSG( | ENSMUSG( | 0.028352 | 0.8557 | NO | NO | NA |
| Trav11d | ENSMUSG( | ENSMUSG( | -1.90449 | 0.541019 | NO | NO | NA |
| Dync2i2 | ENSMUSG( | ENSMUSG( | 1.703224 | 1.63E-06 | YES | YES | Upregulated |
| Nipal1 | ENSMUSG( | ENSMUSG( | -0.43063 | 0.01404 | YES | NO | NA |
| Gadd45a | ENSMUSG( | ENSMUSG( | 0.309384 | 0.116016 | NO | NO | NA |
| Saysd1 | ENSMUSG( | ENSMUSG( | -0.30894 | 0.006963 | YES | NO | NA |
| Nyap1 | ENSMUSG( | ENSMUSG( | -0.43378 | 0.014505 | YES | NO | NA |
| Gm36445 | ENSMUSG( | ENSMUSG( | 0.310753 | 0.037252 | YES | NO | NA |
| Fcgrt | ENSMUSG( | ENSMUSG( | -0.47873 | 0.000606 | YES | NO | NA |
| Trmt10b | ENSMUSG( | ENSMUSG( | -0.08136 | 0.534088 | NO | NO | NA |
| Tmod1 | ENSMUSG( | ENSMUSG( | 0.175058 | 0.245332 | NO | NO | NA |
| Arhgef39 | ENSMUSG( | ENSMUSG( | 0.39326 | 0.032887 | YES | NO | NA |
| Arl6 | ENSMUSG( | ENSMUSG( | 0.90985 | 0.0011 | YES | NO | NA |
| Dqx1 | ENSMUSG( | ENSMUSG( | -0.46771 | 0.000207 | YES | NO | NA |
| Gorab | ENSMUSG( | ENSMUSG( | -0.12209 | 0.501497 | NO | NO | NA |
| Lyrm4 | ENSMUSG( | ENSMUSG( | 0.217379 | 0.195955 | NO | NO | NA |
| 5033430I1 | ENSMUSG( | ENSMUSG( | -0.03347 | 0.840292 | NO | NO | NA |
| Dzip1 | ENSMUSG( | ENSMUSG( | 0.151821 | 0.40295 | NO | NO | NA |
| Spag1 | ENSMUSG( | ENSMUSG( | -0.42374 | 0.038356 | YES | NO | NA |
| Smox | ENSMUSG( | ENSMUSG( | -0.21669 | 0.296719 | NO | NO | NA |

| | | | | | | | |
|---|---|---|---|---|---|---|---|
| Pdss2 | ENSMUSG( | ENSMUSG( | 0.403092 | 0.015591 | YES | NO | NA |
| Dmwd | ENSMUSG( | ENSMUSG( | 1.449867 | 1.45E-05 | YES | YES | Upregulated |
| Atp23 | ENSMUSG( | ENSMUSG( | 0.194841 | 0.148287 | NO | NO | NA |
| Fahd2a | ENSMUSG( | ENSMUSG( | 0.293117 | 0.119447 | NO | NO | NA |
| Nat8f4 | ENSMUSG( | ENSMUSG( | -0.23119 | 0.20405 | NO | NO | NA |
| Slc25a35 | ENSMUSG( | ENSMUSG( | -0.26032 | 0.171936 | NO | NO | NA |
| 4933440N | ENSMUSG( | ENSMUSG( | 0.102922 | 0.693868 | NO | NO | NA |
| Klhl3 | ENSMUSG( | ENSMUSG( | -0.26403 | 0.040233 | YES | NO | NA |
| Slc16a13 | ENSMUSG( | ENSMUSG( | -0.01756 | 0.91013 | NO | NO | NA |
| 1700052K: | ENSMUSG( | ENSMUSG( | 0.139387 | 0.349117 | NO | NO | NA |
| Carf | ENSMUSG( | ENSMUSG( | 0.310608 | 0.121871 | NO | NO | NA |
| Gm20342 | ENSMUSG( | ENSMUSG( | -0.51244 | 0.005408 | YES | NO | NA |
| Mmaa | ENSMUSG( | ENSMUSG( | -0.3404 | 0.009489 | YES | NO | NA |
| Dennd3 | ENSMUSG( | ENSMUSG( | -0.06278 | 0.742607 | NO | NO | NA |
| Gm50322 | ENSMUSG( | ENSMUSG( | 0.181327 | 0.365426 | NO | NO | NA |
| Igf1r | ENSMUSG( | ENSMUSG( | 0.410003 | 0.492717 | NO | NO | NA |
| Sox11 | ENSMUSG( | ENSMUSG( | 0.616084 | 0.625664 | NO | NO | NA |
| Thumpd2 | ENSMUSG( | ENSMUSG( | 0.046085 | 0.728736 | NO | NO | NA |
| Tmem156 | ENSMUSG( | ENSMUSG( | -0.1945 | 0.134397 | NO | NO | NA |
| Ift57 | ENSMUSG( | ENSMUSG( | 1.181094 | 4.69E-06 | YES | YES | Upregulated |
| Reep2 | ENSMUSG( | ENSMUSG( | 0.375857 | 0.053197 | NO | NO | NA |
| Tle1 | ENSMUSG( | ENSMUSG( | 1.398978 | 7.58E-07 | YES | YES | Upregulated |
| Csrnp2 | ENSMUSG( | ENSMUSG( | -0.14139 | 0.596394 | NO | NO | NA |
| 4933407K: | ENSMUSG( | ENSMUSG( | -0.05633 | 0.80032 | NO | NO | NA |
| Cyp4f13 | ENSMUSG( | ENSMUSG( | -0.15078 | 0.467968 | NO | NO | NA |
| Eef2k | ENSMUSG( | ENSMUSG( | 0.441507 | 0.039766 | YES | NO | NA |
| ENSMUSG | ENSMUSG( | ENSMUSG( | -0.78371 | 0.002883 | YES | NO | NA |
| Dnm3 | ENSMUSG( | ENSMUSG( | 0.30064 | 0.494189 | NO | NO | NA |
| Fn3krp | ENSMUSG( | ENSMUSG( | -0.05441 | 0.732349 | NO | NO | NA |
| Lysmd4 | ENSMUSG( | ENSMUSG( | -0.38175 | 0.005408 | YES | NO | NA |
| Mblac2 | ENSMUSG( | ENSMUSG( | -0.08091 | 0.618809 | NO | NO | NA |
| Hexdc | ENSMUSG( | ENSMUSG( | 0.667064 | 1.91E-05 | YES | NO | NA |
| Zik1 | ENSMUSG( | ENSMUSG( | 0.361246 | 0.011696 | YES | NO | NA |
| Zscan22 | ENSMUSG( | ENSMUSG( | -0.04443 | 0.792835 | NO | NO | NA |
| Nudt22 | ENSMUSG( | ENSMUSG( | -0.04696 | 0.784646 | NO | NO | NA |
| Rasl11b | ENSMUSG( | ENSMUSG( | -0.52124 | 0.034312 | YES | NO | NA |
| Gm48362 | ENSMUSG( | ENSMUSG( | 0.209943 | 0.729736 | NO | NO | NA |
| Idh1 | ENSMUSG( | ENSMUSG( | 0.125318 | 0.664907 | NO | NO | NA |
| AW20949: | ENSMUSG( | ENSMUSG( | 0.00104 | 0.994565 | NO | NO | NA |
| Aamdc | ENSMUSG( | ENSMUSG( | 0.185245 | 0.326972 | NO | NO | NA |
| Trav9d-3 | ENSMUSG( | ENSMUSG( | -1.46009 | 0.676249 | NO | NO | NA |
| Cyb5d2 | ENSMUSG( | ENSMUSG( | -0.22604 | 0.223369 | NO | NO | NA |
| Ccdc112 | ENSMUSG( | ENSMUSG( | -0.03007 | 0.876368 | NO | NO | NA |
| Gorasp1 | ENSMUSG( | ENSMUSG( | -0.32266 | 0.011375 | YES | NO | NA |
| Tcta | ENSMUSG( | ENSMUSG( | -0.23944 | 0.300607 | NO | NO | NA |
| Gm43980 | ENSMUSG( | ENSMUSG( | 0.147747 | 0.406581 | NO | NO | NA |
| AW14615 | ENSMUSG( | ENSMUSG( | -0.42467 | 0.011853 | YES | NO | NA |

| | | | | | | | |
|---|---|---|---|---|---|---|---|
| Gm15601 | ENSMUSG( | ENSMUSG( | -0.09541 | 0.593913 | NO | NO | NA |
| Rnf128 | ENSMUSG( | ENSMUSG( | 0.687207 | 0.000358 | YES | NO | NA |
| Fam89a | ENSMUSG( | ENSMUSG( | -0.64751 | 0.000394 | YES | NO | NA |
| Proser3 | ENSMUSG( | ENSMUSG( | -0.43341 | 0.000348 | YES | NO | NA |
| 0610009B: | ENSMUSG( | ENSMUSG( | 0.140915 | 0.450269 | NO | NO | NA |
| Gm14326 | ENSMUSG( | ENSMUSG( | 0.259625 | 0.220077 | NO | NO | NA |
| Sgk3 | ENSMUSG( | ENSMUSG( | -0.48817 | 0.036085 | YES | NO | NA |
| Zfp580 | ENSMUSG( | ENSMUSG( | 1.460524 | 4.89E-06 | YES | YES | Upregulated |
| Xk | ENSMUSG( | ENSMUSG( | 0.206945 | 0.129107 | NO | NO | NA |
| Gm14295 | ENSMUSG( | ENSMUSG( | 0.628547 | 0.002434 | YES | NO | NA |
| Igkv10-96 | ENSMUSG( | ENSMUSG( | -1.07595 | 0.457817 | NO | NO | NA |
| Gm11478 | ENSMUSG( | ENSMUSG( | 0.115328 | 0.649088 | NO | NO | NA |
| Fads2 | ENSMUSG( | ENSMUSG( | 2.746574 | 0.000277 | YES | YES | Upregulated |
| Tmem159 | ENSMUSG( | ENSMUSG( | 0.009343 | 0.966064 | NO | NO | NA |
| Gm16196 | ENSMUSG( | ENSMUSG( | 0.424423 | 0.457961 | NO | NO | NA |
| Enpp5 | ENSMUSG( | ENSMUSG( | -0.88382 | 2.11E-05 | YES | NO | NA |
| Cul7 | ENSMUSG( | ENSMUSG( | 0.192283 | 0.291809 | NO | NO | NA |
| Nrxn2 | ENSMUSG( | ENSMUSG( | 0.165328 | 0.612241 | NO | NO | NA |
| Nrbp2 | ENSMUSG( | ENSMUSG( | -0.23882 | 0.254302 | NO | NO | NA |
| D030028A | ENSMUSG( | ENSMUSG( | 0.063729 | 0.721186 | NO | NO | NA |
| Gm15427 | ENSMUSG( | ENSMUSG( | 0.458581 | 0.002575 | YES | NO | NA |
| Dynlt1c | ENSMUSG( | ENSMUSG( | -0.19527 | 0.333497 | NO | NO | NA |
| Adat3 | ENSMUSG( | ENSMUSG( | 0.205404 | 0.457961 | NO | NO | NA |
| H2-Ab1 | ENSMUSG( | ENSMUSG( | 0.197983 | 0.628823 | NO | NO | NA |
| Gm10138 | ENSMUSG( | ENSMUSG( | 0.180073 | 0.555025 | NO | NO | NA |
| Gm42664 | ENSMUSG( | ENSMUSG( | -0.34348 | 0.070503 | NO | NO | NA |
| Rims3 | ENSMUSG( | ENSMUSG( | 0.577061 | 7.04E-05 | YES | NO | NA |
| H3c3 | ENSMUSG( | ENSMUSG( | 0.36863 | 0.103206 | NO | NO | NA |
| Zbtb26 | ENSMUSG( | ENSMUSG( | -0.30102 | 0.275501 | NO | NO | NA |
| Zbtb11os1 | ENSMUSG( | ENSMUSG( | 0.101409 | 0.659678 | NO | NO | NA |
| H2bu2 | ENSMUSG( | ENSMUSG( | 0.212747 | 0.41702 | NO | NO | NA |
| Trav6-3 | ENSMUSG( | ENSMUSG( | -0.18514 | 0.796087 | NO | NO | NA |
| Gm43481 | ENSMUSG( | ENSMUSG( | -0.1814 | 0.30206 | NO | NO | NA |
| Mospd1 | ENSMUSG( | ENSMUSG( | -0.03532 | 0.806706 | NO | NO | NA |
| Tmcc2 | ENSMUSG( | ENSMUSG( | 0.704397 | 0.244677 | NO | NO | NA |
| Gemin8 | ENSMUSG( | ENSMUSG( | 0.352347 | 0.034857 | YES | NO | NA |
| Txnl4b | ENSMUSG( | ENSMUSG( | -0.09876 | 0.510869 | NO | NO | NA |
| Gm43065 | ENSMUSG( | ENSMUSG( | -0.51379 | 0.001422 | YES | NO | NA |
| 1700037C: | ENSMUSG( | ENSMUSG( | 0.027267 | 0.841312 | NO | NO | NA |
| Ctbs | ENSMUSG( | ENSMUSG( | 0.205445 | 0.301687 | NO | NO | NA |
| Adam15 | ENSMUSG( | ENSMUSG( | 0.257131 | 0.07256 | NO | NO | NA |
| Katnal1 | ENSMUSG( | ENSMUSG( | -0.04307 | 0.808877 | NO | NO | NA |
| Gm38077 | ENSMUSG( | ENSMUSG( | -0.08982 | 0.7116 | NO | NO | NA |
| Eif4ebp1 | ENSMUSG( | ENSMUSG( | 0.358441 | 0.086923 | NO | NO | NA |
| Gm12185 | ENSMUSG( | ENSMUSG( | -0.65989 | 4.17E-05 | YES | NO | NA |
| ENSMUSG | ENSMUSG( | ENSMUSG( | 0.168838 | 0.268652 | NO | NO | NA |
| Tnfaip2 | ENSMUSG( | ENSMUSG( | 0.759507 | 0.150749 | NO | NO | NA |

| | | | | | | | |
|---|---|---|---|---|---|---|---|
| Dgcr6 | ENSMUSG( | ENSMUSG( | 1.286979 | 1.48E-05 | YES | YES | Upregulated |
| Khk | ENSMUSG( | ENSMUSG( | 0.23357 | 0.236195 | NO | NO | NA |
| Sirt5 | ENSMUSG( | ENSMUSG( | 0.389451 | 0.025918 | YES | NO | NA |
| Mettl22 | ENSMUSG( | ENSMUSG( | 0.416894 | 0.002212 | YES | NO | NA |
| Tnfsf8 | ENSMUSG( | ENSMUSG( | -0.25461 | 0.454653 | NO | NO | NA |
| Ceacam16 | ENSMUSG( | ENSMUSG( | 0.520122 | 0.030171 | YES | NO | NA |
| ENSMUSG | ENSMUSG( | ENSMUSG( | 0.436075 | 0.001332 | YES | NO | NA |
| Ncam1 | ENSMUSG( | ENSMUSG( | 0.711626 | 0.042303 | YES | NO | NA |
| Zfp934 | ENSMUSG( | ENSMUSG( | -0.39636 | 0.001355 | YES | NO | NA |
| Etv5 | ENSMUSG( | ENSMUSG( | 0.248808 | 0.029892 | YES | NO | NA |
| Zswim3 | ENSMUSG( | ENSMUSG( | -0.58833 | 1.38E-05 | YES | NO | NA |
| Gm11808 | ENSMUSG( | ENSMUSG( | 0.191194 | 0.30875 | NO | NO | NA |
| Ndufb4c | ENSMUSG( | ENSMUSG( | -0.7608 | 0.660508 | NO | NO | NA |
| St6gal1 | ENSMUSG( | ENSMUSG( | 0.633827 | 0.2714 | NO | NO | NA |
| Icosl | ENSMUSG( | ENSMUSG( | -0.45914 | 0.110475 | NO | NO | NA |
| Zcwpw1 | ENSMUSG( | ENSMUSG( | 0.075425 | 0.73356 | NO | NO | NA |
| Oplah | ENSMUSG( | ENSMUSG( | -0.66519 | 3.58E-06 | YES | NO | NA |
| Pygm | ENSMUSG( | ENSMUSG( | 0.023691 | 0.903102 | NO | NO | NA |
| Kif7 | ENSMUSG( | ENSMUSG( | 0.730348 | 9.46E-05 | YES | NO | NA |
| Mrnip | ENSMUSG( | ENSMUSG( | 0.517789 | 0.000594 | YES | NO | NA |
| E030022I1 | ENSMUSG( | ENSMUSG( | -0.75493 | 9.88E-07 | YES | NO | NA |
| C1qtnf12 | ENSMUSG( | ENSMUSG( | 0.64053 | 0.085409 | NO | NO | NA |
| Rasgef1b | ENSMUSG( | ENSMUSG( | -0.29305 | 0.560577 | NO | NO | NA |
| Samd3 | ENSMUSG( | ENSMUSG( | -1.19016 | 1.25E-05 | YES | YES | Downregulated |
| Kcnc1 | ENSMUSG( | ENSMUSG( | -1.15762 | 2.42E-09 | YES | YES | Downregulated |
| Zfp719 | ENSMUSG( | ENSMUSG( | -0.2032 | 0.178208 | NO | NO | NA |
| Gm28035 | ENSMUSG( | ENSMUSG( | -0.85108 | 1.81E-05 | YES | NO | NA |
| Gm21092 | ENSMUSG( | ENSMUSG( | 0.389284 | 0.032824 | YES | NO | NA |
| Cep131 | ENSMUSG( | ENSMUSG( | -0.061 | 0.785998 | NO | NO | NA |
| Tasl | ENSMUSG( | ENSMUSG( | -0.17946 | 0.299905 | NO | NO | NA |
| Cmc4 | ENSMUSG( | ENSMUSG( | 0.235291 | 0.161056 | NO | NO | NA |
| Pigbos1 | ENSMUSG( | ENSMUSG( | -0.11631 | 0.414065 | NO | NO | NA |
| Mthfr | ENSMUSG( | ENSMUSG( | -0.36717 | 0.079144 | NO | NO | NA |
| Ccdc30 | ENSMUSG( | ENSMUSG( | 0.029758 | 0.869416 | NO | NO | NA |
| Enpp6 | ENSMUSG( | ENSMUSG( | -0.12521 | 0.649781 | NO | NO | NA |
| Fam174c | ENSMUSG( | ENSMUSG( | -0.20095 | 0.169824 | NO | NO | NA |
| Lrsam1 | ENSMUSG( | ENSMUSG( | -0.46767 | 0.006032 | YES | NO | NA |
| Mypop | ENSMUSG( | ENSMUSG( | -0.24344 | 0.130256 | NO | NO | NA |
| Fbxl15 | ENSMUSG( | ENSMUSG( | 0.153859 | 0.201381 | NO | NO | NA |
| Harbi1 | ENSMUSG( | ENSMUSG( | 0.049916 | 0.784991 | NO | NO | NA |
| Dusp28 | ENSMUSG( | ENSMUSG( | 0.388688 | 0.082584 | NO | NO | NA |
| Trav7-3 | ENSMUSG( | ENSMUSG( | -0.61267 | 0.402033 | NO | NO | NA |
| Tns2 | ENSMUSG( | ENSMUSG( | 0.710185 | 7.91E-06 | YES | NO | NA |
| A930024E( | ENSMUSG( | ENSMUSG( | 0.014496 | 0.947496 | NO | NO | NA |
| Bicd1 | ENSMUSG( | ENSMUSG( | 1.148491 | 2.89E-07 | YES | YES | Upregulated |
| Gm20687 | ENSMUSG( | ENSMUSG( | -1.16694 | 5.98E-07 | YES | YES | Downregulated |
| Ss18l1 | ENSMUSG( | ENSMUSG( | -0.36378 | 0.010768 | YES | NO | NA |

| | | | | | | | |
|---|---|---|---|---|---|---|---|
| Gm49760 | ENSMUSG( | ENSMUSG( | -0.16449 | 0.348662 | NO | NO | NA |
| Oasl2 | ENSMUSG( | ENSMUSG( | -0.17456 | 0.567401 | NO | NO | NA |
| Zscan2 | ENSMUSG( | ENSMUSG( | -0.22429 | 0.159078 | NO | NO | NA |
| Dipk1b | ENSMUSG( | ENSMUSG( | 0.546895 | 0.000314 | YES | NO | NA |
| Ank2 | ENSMUSG( | ENSMUSG( | 0.562348 | 0.650468 | NO | NO | NA |
| Hpse | ENSMUSG( | ENSMUSG( | -0.42727 | 0.00164 | YES | NO | NA |
| Dtwd1 | ENSMUSG( | ENSMUSG( | 0.45798 | 0.004112 | YES | NO | NA |
| Dpp7 | ENSMUSG( | ENSMUSG( | -0.21088 | 0.198038 | NO | NO | NA |
| Gm44231 | ENSMUSG( | ENSMUSG( | 0.797416 | 0.312909 | NO | NO | NA |
| Slc5a6 | ENSMUSG( | ENSMUSG( | -0.01896 | 0.892287 | NO | NO | NA |
| 5830444B( | ENSMUSG( | ENSMUSG( | -1.13058 | 0.172449 | NO | NO | NA |
| Cd200 | ENSMUSG( | ENSMUSG( | -0.2075 | 0.534652 | NO | NO | NA |
| P2rx4 | ENSMUSG( | ENSMUSG( | 0.316337 | 0.190511 | NO | NO | NA |
| Pfkfb1 | ENSMUSG( | ENSMUSG( | 0.070947 | 0.747414 | NO | NO | NA |
| D630045J1 | ENSMUSG( | ENSMUSG( | 1.367958 | 2.95E-11 | YES | YES | Upregulated |
| Hoxb4 | ENSMUSG( | ENSMUSG( | -0.31128 | 0.01114 | YES | NO | NA |
| Mx2 | ENSMUSG( | ENSMUSG( | -1.61225 | 9.32E-12 | YES | YES | Downregulated |
| Mccc2 | ENSMUSG( | ENSMUSG( | 0.830757 | 9.87E-06 | YES | NO | NA |
| Zfp54 | ENSMUSG( | ENSMUSG( | -0.33339 | 0.043955 | YES | NO | NA |
| Med31 | ENSMUSG( | ENSMUSG( | -0.09845 | 0.574311 | NO | NO | NA |
| Gm3055 | ENSMUSG( | ENSMUSG( | -0.46093 | 0.067556 | NO | NO | NA |
| Gm13835 | ENSMUSG( | ENSMUSG( | 0.35946 | 0.007073 | YES | NO | NA |
| Klrh1 | ENSMUSG( | ENSMUSG( | 1.020807 | 0.209948 | NO | NO | NA |
| Arv1 | ENSMUSG( | ENSMUSG( | 0.04318 | 0.791732 | NO | NO | NA |
| Gm48529 | ENSMUSG( | ENSMUSG( | -0.22031 | 0.328452 | NO | NO | NA |
| 4930404A | ENSMUSG( | ENSMUSG( | -0.29702 | 0.010881 | YES | NO | NA |
| Far2 | ENSMUSG( | ENSMUSG( | -0.34534 | 0.108402 | NO | NO | NA |
| Cd93 | ENSMUSG( | ENSMUSG( | 0.973822 | 0.000891 | YES | NO | NA |
| Bmf | ENSMUSG( | ENSMUSG( | -1.22433 | 0.003328 | YES | YES | Downregulated |
| Rras | ENSMUSG( | ENSMUSG( | -0.46881 | 0.002336 | YES | NO | NA |
| Gm8752 | ENSMUSG( | ENSMUSG( | -0.03423 | 0.930376 | NO | NO | NA |
| Zfp560 | ENSMUSG( | ENSMUSG( | -0.03212 | 0.85174 | NO | NO | NA |
| Fktn | ENSMUSG( | ENSMUSG( | -0.17451 | 0.240422 | NO | NO | NA |
| Gm21992 | ENSMUSG( | ENSMUSG( | 0.13823 | 0.570134 | NO | NO | NA |
| Gm48348 | ENSMUSG( | ENSMUSG( | 0.843172 | 0.000502 | YES | NO | NA |
| Gm36043 | ENSMUSG( | ENSMUSG( | -0.01014 | 0.974434 | NO | NO | NA |
| Cgrrf1 | ENSMUSG( | ENSMUSG( | 0.045744 | 0.712875 | NO | NO | NA |
| Aoc2 | ENSMUSG( | ENSMUSG( | 0.277014 | 0.116413 | NO | NO | NA |
| Zfp458 | ENSMUSG( | ENSMUSG( | -0.18029 | 0.374815 | NO | NO | NA |
| A430033K( | ENSMUSG( | ENSMUSG( | -0.07192 | 0.664907 | NO | NO | NA |
| Grin2b | ENSMUSG( | ENSMUSG( | 0.815095 | 0.513114 | NO | NO | NA |
| Trav6d-6 | ENSMUSG( | ENSMUSG( | 0.134894 | 0.972271 | NO | NO | NA |
| Gm38102 | ENSMUSG( | ENSMUSG( | -0.58653 | 0.000869 | YES | NO | NA |
| Sema4c | ENSMUSG( | ENSMUSG( | 0.692968 | 0.000982 | YES | NO | NA |
| Gm45705 | ENSMUSG( | ENSMUSG( | -0.05181 | 0.810767 | NO | NO | NA |
| Pex11g | ENSMUSG( | ENSMUSG( | 0.042448 | 0.795085 | NO | NO | NA |
| 4632404H | ENSMUSG( | ENSMUSG( | -0.02488 | 0.86989 | NO | NO | NA |

| | | | | | | | |
|---|---|---|---|---|---|---|---|
| Pkig | ENSMUSG( | ENSMUSG( | 0.867718 | 0.003046 | YES | NO | NA |
| Nudt7 | ENSMUSG( | ENSMUSG( | -0.30273 | 0.013666 | YES | NO | NA |
| Bbs4 | ENSMUSG( | ENSMUSG( | 0.45499 | 4.74E-05 | YES | NO | NA |
| ENSMUSG( | ENSMUSG( | ENSMUSG( | -0.3462 | 0.023057 | YES | NO | NA |
| 4931428F( | ENSMUSG( | ENSMUSG( | -0.31687 | 0.096211 | NO | NO | NA |
| Pard6a | ENSMUSG( | ENSMUSG( | 0.202422 | 0.139079 | NO | NO | NA |
| Fam20a | ENSMUSG( | ENSMUSG( | 1.230315 | 0.000602 | YES | YES | Upregulated |
| Susd2 | ENSMUSG( | ENSMUSG( | -0.86806 | 0.001425 | YES | NO | NA |
| Sec61a2 | ENSMUSG( | ENSMUSG( | 0.03428 | 0.880464 | NO | NO | NA |
| Gm43300 | ENSMUSG( | ENSMUSG( | 0.093169 | 0.72046 | NO | NO | NA |
| Abhd14a | ENSMUSG( | ENSMUSG( | 0.64318 | 0.000843 | YES | NO | NA |
| A630089N | ENSMUSG( | ENSMUSG( | -0.43714 | 3.76E-05 | YES | NO | NA |
| Castor2 | ENSMUSG( | ENSMUSG( | -0.04868 | 0.832943 | NO | NO | NA |
| Pcnx4 | ENSMUSG( | ENSMUSG( | -0.06888 | 0.697041 | NO | NO | NA |
| Dtwd2 | ENSMUSG( | ENSMUSG( | 0.220585 | 0.313391 | NO | NO | NA |
| D730005E | ENSMUSG( | ENSMUSG( | -0.11593 | 0.493365 | NO | NO | NA |
| Gm15232 | ENSMUSG( | ENSMUSG( | -0.05396 | 0.739465 | NO | NO | NA |
| 1110032A( | ENSMUSG( | ENSMUSG( | 0.643478 | 0.000138 | YES | NO | NA |
| Rgs10 | ENSMUSG( | ENSMUSG( | 0.287942 | 0.13582 | NO | NO | NA |
| Igsf3 | ENSMUSG( | ENSMUSG( | 0.323706 | 0.556141 | NO | NO | NA |
| Extl2 | ENSMUSG( | ENSMUSG( | 0.320756 | 0.032815 | YES | NO | NA |
| ENSMUSG | ENSMUSG( | ENSMUSG( | 0.288481 | 0.239263 | NO | NO | NA |
| Kctd7 | ENSMUSG( | ENSMUSG( | -0.12108 | 0.501479 | NO | NO | NA |
| Gm29609 | ENSMUSG( | ENSMUSG( | -0.54455 | 0.231297 | NO | NO | NA |
| Ighv1-66 | ENSMUSG( | ENSMUSG( | -0.60662 | 0.824067 | NO | NO | NA |
| ENSMUSG | ENSMUSG( | ENSMUSG( | 0.051445 | 0.769794 | NO | NO | NA |
| Rpl35a-ps3 | ENSMUSG( | ENSMUSG( | -0.48486 | 0.831409 | NO | NO | NA |
| Kank3 | ENSMUSG( | ENSMUSG( | 0.714918 | 0.001382 | YES | NO | NA |
| Med18 | ENSMUSG( | ENSMUSG( | 0.256873 | 0.344724 | NO | NO | NA |
| 2300009A( | ENSMUSG( | ENSMUSG( | 0.390502 | 0.003985 | YES | NO | NA |
| Shld3 | ENSMUSG( | ENSMUSG( | -0.03855 | 0.854348 | NO | NO | NA |
| Gm38190 | ENSMUSG( | ENSMUSG( | 0.118606 | 0.505315 | NO | NO | NA |
| Rpl10-ps3 | ENSMUSG( | ENSMUSG( | 0.531071 | 0.006367 | YES | NO | NA |
| Nmnat1 | ENSMUSG( | ENSMUSG( | 0.108766 | 0.470684 | NO | NO | NA |
| Rps23-ps1 | ENSMUSG( | ENSMUSG( | 0.524347 | 0.000143 | YES | NO | NA |
| Mrm2 | ENSMUSG( | ENSMUSG( | 0.011628 | 0.938452 | NO | NO | NA |
| Rbms3 | ENSMUSG( | ENSMUSG( | 0.786443 | 0.502435 | NO | NO | NA |
| Gm5165 | ENSMUSG( | ENSMUSG( | -0.29436 | 0.128778 | NO | NO | NA |
| Jpx | ENSMUSG( | ENSMUSG( | 0.058677 | 0.745283 | NO | NO | NA |
| Pcx | ENSMUSG( | ENSMUSG( | 0.922248 | 0.001958 | YES | NO | NA |
| Tmed1 | ENSMUSG( | ENSMUSG( | 0.437607 | 0.002964 | YES | NO | NA |
| Prx | ENSMUSG( | ENSMUSG( | 0.19247 | 0.347976 | NO | NO | NA |
| 2010008C1 | ENSMUSG( | ENSMUSG( | -0.5917 | 0.001733 | YES | NO | NA |
| ENSMUSG | ENSMUSG( | ENSMUSG( | 0.45182 | 0.000242 | YES | NO | NA |
| Ssbp2 | ENSMUSG( | ENSMUSG( | 1.055788 | 7.35E-05 | YES | YES | Upregulated |
| Dmtn | ENSMUSG( | ENSMUSG( | 0.595602 | 0.300876 | NO | NO | NA |
| Sv2c | ENSMUSG( | ENSMUSG( | 0.619302 | 7.56E-06 | YES | NO | NA |

| | | | | | | | |
|---|---|---|---|---|---|---|---|
| Gm20707 | ENSMUSG( | ENSMUSG( | -0.97061 | 3.00E-09 | YES | NO | NA |
| Rnf122 | ENSMUSG( | ENSMUSG( | 0.397545 | 0.119978 | NO | NO | NA |
| AI847159 | ENSMUSG( | ENSMUSG( | -0.84614 | 1.43E-05 | YES | NO | NA |
| Jmjd8 | ENSMUSG( | ENSMUSG( | -0.09871 | 0.623689 | NO | NO | NA |
| Gm26648 | ENSMUSG( | ENSMUSG( | 0.165051 | 0.530158 | NO | NO | NA |
| Ears2 | ENSMUSG( | ENSMUSG( | 0.186934 | 0.281672 | NO | NO | NA |
| ENSMUSG | ENSMUSG( | ENSMUSG( | -0.4957 | 0.022599 | YES | NO | NA |
| Ino80dos | ENSMUSG( | ENSMUSG( | 0.067908 | 0.706675 | NO | NO | NA |
| Klf12 | ENSMUSG( | ENSMUSG( | -0.63162 | 0.001193 | YES | NO | NA |
| ENSMUSG( | ENSMUSG( | ENSMUSG( | -0.24762 | 0.065766 | NO | NO | NA |
| Mocos | ENSMUSG( | ENSMUSG( | -1.06118 | 1.35E-08 | YES | YES | Downregulated |
| Prdm9 | ENSMUSG( | ENSMUSG( | 0.406316 | 0.00344 | YES | NO | NA |
| Kndc1 | ENSMUSG( | ENSMUSG( | 0.530052 | 0.002251 | YES | NO | NA |
| Gsdme | ENSMUSG( | ENSMUSG( | 0.393911 | 0.067503 | NO | NO | NA |
| Gm9843 | ENSMUSG( | ENSMUSG( | 0.200994 | 0.34493 | NO | NO | NA |
| H2ac11 | ENSMUSG( | ENSMUSG( | 0.202916 | 0.278855 | NO | NO | NA |
| Tlcd1 | ENSMUSG( | ENSMUSG( | 0.466428 | 0.001074 | YES | NO | NA |
| Gm31597 | ENSMUSG( | ENSMUSG( | 0.181221 | 0.324751 | NO | NO | NA |
| Milr1 | ENSMUSG( | ENSMUSG( | 0.523696 | 0.009255 | YES | NO | NA |
| Vmn2r102 | ENSMUSG( | ENSMUSG( | 0.640234 | 0.610287 | NO | NO | NA |
| Ldlrad4 | ENSMUSG( | ENSMUSG( | -1.01298 | 7.37E-07 | YES | YES | Downregulated |
| Gm29361 | ENSMUSG( | ENSMUSG( | 0.053898 | 0.863696 | NO | NO | NA |
| Trav9n-3 | ENSMUSG( | ENSMUSG( | 0.179051 | 0.914204 | NO | NO | NA |
| Plod1 | ENSMUSG( | ENSMUSG( | -1.61164 | 2.11E-15 | YES | YES | Downregulated |
| Olfr56 | ENSMUSG( | ENSMUSG( | 0.795644 | 0.004015 | YES | NO | NA |
| Ticam2 | ENSMUSG( | ENSMUSG( | 0.649603 | 0.001835 | YES | NO | NA |
| Vmac | ENSMUSG( | ENSMUSG( | 0.07535 | 0.786739 | NO | NO | NA |
| Ccdc157 | ENSMUSG( | ENSMUSG( | -0.11925 | 0.647539 | NO | NO | NA |
| Osgin1 | ENSMUSG( | ENSMUSG( | -0.61604 | 0.260726 | NO | NO | NA |
| Zfp189 | ENSMUSG( | ENSMUSG( | -0.38632 | 0.001707 | YES | NO | NA |
| Egln3 | ENSMUSG( | ENSMUSG( | 1.257283 | 0.000286 | YES | YES | Upregulated |
| Fbxo44 | ENSMUSG( | ENSMUSG( | -0.99643 | 2.14E-19 | YES | NO | NA |
| Gm44423 | ENSMUSG( | ENSMUSG( | -0.30016 | 0.178208 | NO | NO | NA |
| Pbx4 | ENSMUSG( | ENSMUSG( | 0.315589 | 0.043827 | YES | NO | NA |
| 4930577N | ENSMUSG( | ENSMUSG( | 0.102851 | 0.727172 | NO | NO | NA |
| Gm45412 | ENSMUSG( | ENSMUSG( | -1.5938 | NA | NO | NO | NA |
| Csad | ENSMUSG( | ENSMUSG( | -0.16944 | 0.296465 | NO | NO | NA |
| Gm42659 | ENSMUSG( | ENSMUSG( | -0.29202 | 0.11386 | NO | NO | NA |
| Trav7d-2 | ENSMUSG( | ENSMUSG( | 0.730569 | 0.597508 | NO | NO | NA |
| Ralb | ENSMUSG( | ENSMUSG( | 0.258429 | 0.020578 | YES | NO | NA |
| Kif3a | ENSMUSG( | ENSMUSG( | 0.64875 | 0.024721 | YES | NO | NA |
| Trav14n-3 | ENSMUSG( | ENSMUSG( | 0.748375 | 0.23712 | NO | NO | NA |
| Rhag | ENSMUSG( | ENSMUSG( | 1.40243 | 0.024355 | YES | YES | Upregulated |
| Mtm1 | ENSMUSG( | ENSMUSG( | -0.00154 | 0.991511 | NO | NO | NA |
| 1810034E: | ENSMUSG( | ENSMUSG( | 0.036022 | 0.820973 | NO | NO | NA |
| ENSMUSG( | ENSMUSG( | ENSMUSG( | 0.540882 | 6.78E-05 | YES | NO | NA |
| Ccdc91 | ENSMUSG( | ENSMUSG( | 0.158709 | 0.475513 | NO | NO | NA |

| | | | | | | | |
|---|---|---|---|---|---|---|---|
| C330018D | ENSMUSG( | ENSMUSG( | -0.45517 | 0.010494 | YES | NO | NA |
| Gm45713 | ENSMUSG( | ENSMUSG( | -0.15161 | 0.779344 | NO | NO | NA |
| Sipa1l2 | ENSMUSG( | ENSMUSG( | -0.43439 | 0.023398 | YES | NO | NA |
| Kifc2 | ENSMUSG( | ENSMUSG( | -0.20082 | 0.193979 | NO | NO | NA |
| Coa4 | ENSMUSG( | ENSMUSG( | -0.15818 | 0.342272 | NO | NO | NA |
| Simc1 | ENSMUSG( | ENSMUSG( | -0.10524 | 0.442832 | NO | NO | NA |
| Zfp760 | ENSMUSG( | ENSMUSG( | -0.60821 | 0.003991 | YES | NO | NA |
| Kcnk7 | ENSMUSG( | ENSMUSG( | 0.542124 | 0.010511 | YES | NO | NA |
| Bmpr2 | ENSMUSG( | ENSMUSG( | 0.100937 | 0.588526 | NO | NO | NA |
| Fbxl4 | ENSMUSG( | ENSMUSG( | -0.09597 | 0.602085 | NO | NO | NA |
| Bag2 | ENSMUSG( | ENSMUSG( | 0.385284 | 0.016947 | YES | NO | NA |
| Agap1 | ENSMUSG( | ENSMUSG( | 0.522071 | 0.473607 | NO | NO | NA |
| Abcb4 | ENSMUSG( | ENSMUSG( | 0.961098 | 0.052605 | NO | NO | NA |
| Trim3 | ENSMUSG( | ENSMUSG( | 0.305563 | 0.029592 | YES | NO | NA |
| Gm43672 | ENSMUSG( | ENSMUSG( | 0.644393 | 0.040579 | YES | NO | NA |
| Nat9 | ENSMUSG( | ENSMUSG( | 0.065658 | 0.768418 | NO | NO | NA |
| Pagr1a | ENSMUSG( | ENSMUSG( | 0.618892 | 0.450504 | NO | NO | NA |
| Tafa3 | ENSMUSG( | ENSMUSG( | -0.46029 | 0.022055 | YES | NO | NA |
| Tgfbr3 | ENSMUSG( | ENSMUSG( | 1.456751 | 9.78E-12 | YES | YES | Upregulated |
| 1700019L1 | ENSMUSG( | ENSMUSG( | 0.520601 | 0.36939 | NO | NO | NA |
| Mcoln3 | ENSMUSG( | ENSMUSG( | 0.247108 | 0.366015 | NO | NO | NA |
| 2610020C( | ENSMUSG( | ENSMUSG( | 0.185231 | 0.202851 | NO | NO | NA |
| ENSMUSG | ENSMUSG( | ENSMUSG( | 0.049484 | 0.799056 | NO | NO | NA |
| Slpi | ENSMUSG( | ENSMUSG( | -0.52452 | 0.663248 | NO | NO | NA |
| Trav6-7-dv | ENSMUSG( | ENSMUSG( | -0.50567 | 0.05362 | NO | NO | NA |
| Tal1 | ENSMUSG( | ENSMUSG( | 1.084943 | 0.009409 | YES | YES | Upregulated |
| Dsel | ENSMUSG( | ENSMUSG( | 0.424551 | 0.098577 | NO | NO | NA |
| Dkkl1 | ENSMUSG( | ENSMUSG( | 0.921249 | 9.49E-10 | YES | NO | NA |
| Spryd4 | ENSMUSG( | ENSMUSG( | 0.067651 | 0.611854 | NO | NO | NA |
| Lhx4 | ENSMUSG( | ENSMUSG( | 0.905515 | 0.439353 | NO | NO | NA |
| ENSMUSG | ENSMUSG( | ENSMUSG( | 0.345904 | 0.15116 | NO | NO | NA |
| ENSMUSG( | ENSMUSG( | ENSMUSG( | 0.144982 | 0.45456 | NO | NO | NA |
| Ttll1 | ENSMUSG( | ENSMUSG( | -0.21261 | 0.172805 | NO | NO | NA |
| Ccdc171 | ENSMUSG( | ENSMUSG( | -0.31856 | 0.062083 | NO | NO | NA |
| ENSMUSG( | ENSMUSG( | ENSMUSG( | -0.33465 | 0.006878 | YES | NO | NA |
| Klf1 | ENSMUSG( | ENSMUSG( | 1.047299 | 0.058516 | NO | NO | NA |
| Gm49521 | ENSMUSG( | ENSMUSG( | -0.22023 | 0.495773 | NO | NO | NA |
| Pla2g4b | ENSMUSG( | ENSMUSG( | -0.30704 | 0.220257 | NO | NO | NA |
| Cetn4 | ENSMUSG( | ENSMUSG( | 0.985294 | 5.70E-07 | YES | NO | NA |
| Epb41l5 | ENSMUSG( | ENSMUSG( | 0.550577 | 0.179798 | NO | NO | NA |
| Gm47071 | ENSMUSG( | ENSMUSG( | -0.43373 | 0.000706 | YES | NO | NA |
| Gm10288 | ENSMUSG( | ENSMUSG( | 0.322633 | 0.176932 | NO | NO | NA |
| Rogdi | ENSMUSG( | ENSMUSG( | 0.316742 | 0.111642 | NO | NO | NA |
| Atp1b1 | ENSMUSG( | ENSMUSG( | -0.44319 | 0.383463 | NO | NO | NA |
| Trav7-5 | ENSMUSG( | ENSMUSG( | -0.36841 | 0.615815 | NO | NO | NA |
| Csf2rb | ENSMUSG( | ENSMUSG( | 0.964826 | 0.000846 | YES | NO | NA |
| Zfp296 | ENSMUSG( | ENSMUSG( | 0.629469 | 0.002216 | YES | NO | NA |

| | | | | | | | |
|---|---|---|---|---|---|---|---|
| Snx21 | ENSMUSG( | ENSMUSG( | -0.6512 | 0.000537 | YES | NO | NA |
| Klf9 | ENSMUSG( | ENSMUSG( | 0.403113 | 0.007149 | YES | NO | NA |
| 5031439G | ENSMUSG( | ENSMUSG( | -0.08953 | 0.625461 | NO | NO | NA |
| Zfp995 | ENSMUSG( | ENSMUSG( | 0.243001 | 0.264799 | NO | NO | NA |
| Tead3 | ENSMUSG( | ENSMUSG( | 0.079873 | 0.78478 | NO | NO | NA |
| Rmdn2 | ENSMUSG( | ENSMUSG( | -0.31808 | 0.028874 | YES | NO | NA |
| ENSMUSG( | ENSMUSG( | ENSMUSG( | 0.358134 | 0.018868 | YES | NO | NA |
| Ciart | ENSMUSG( | ENSMUSG( | 0.362597 | 0.01673 | YES | NO | NA |
| Stx3 | ENSMUSG( | ENSMUSG( | 0.655844 | 0.021844 | YES | NO | NA |
| Pex11a | ENSMUSG( | ENSMUSG( | -0.0136 | 0.959488 | NO | NO | NA |
| Gm2007 | ENSMUSG( | ENSMUSG( | -0.57949 | 0.714063 | NO | NO | NA |
| Adssl1 | ENSMUSG( | ENSMUSG( | 0.515352 | 0.00665 | YES | NO | NA |
| Zfp846 | ENSMUSG( | ENSMUSG( | -0.02375 | 0.901337 | NO | NO | NA |
| Dram1 | ENSMUSG( | ENSMUSG( | -0.49354 | 0.013156 | YES | NO | NA |
| Tespa1 | ENSMUSG( | ENSMUSG( | -0.29622 | 0.328452 | NO | NO | NA |
| Zfhx3 | ENSMUSG( | ENSMUSG( | 0.564586 | 0.482306 | NO | NO | NA |
| 3010003L2 | ENSMUSG( | ENSMUSG( | 0.541737 | 0.023828 | YES | NO | NA |
| Zfp267 | ENSMUSG( | ENSMUSG( | 0.450047 | 0.003734 | YES | NO | NA |
| 4732487G | ENSMUSG( | ENSMUSG( | 0.009414 | 0.963475 | NO | NO | NA |
| Aifm2 | ENSMUSG( | ENSMUSG( | 0.396917 | 0.176825 | NO | NO | NA |
| Gm45871 | ENSMUSG( | ENSMUSG( | -0.27276 | 0.131089 | NO | NO | NA |
| Myo6 | ENSMUSG( | ENSMUSG( | -0.31911 | 0.249552 | NO | NO | NA |
| Gm38220 | ENSMUSG( | ENSMUSG( | 0.154784 | 0.732447 | NO | NO | NA |
| Yy2 | ENSMUSG( | ENSMUSG( | 0.201633 | 0.419854 | NO | NO | NA |
| Sema6b | ENSMUSG( | ENSMUSG( | 0.660273 | 0.012134 | YES | NO | NA |
| Ccdc28a | ENSMUSG( | ENSMUSG( | 0.082548 | 0.646058 | NO | NO | NA |
| 9230112E( | ENSMUSG( | ENSMUSG( | -0.01755 | 0.928853 | NO | NO | NA |
| Ccl25 | ENSMUSG( | ENSMUSG( | -0.05164 | 0.832793 | NO | NO | NA |
| Clybl | ENSMUSG( | ENSMUSG( | 1.260696 | 0.000279 | YES | YES | Upregulated |
| Ogfod3 | ENSMUSG( | ENSMUSG( | 0.585162 | 0.000277 | YES | NO | NA |
| Gm47694 | ENSMUSG( | ENSMUSG( | -0.14804 | 0.489587 | NO | NO | NA |
| Serpinc1 | ENSMUSG( | ENSMUSG( | 0.155973 | 0.624776 | NO | NO | NA |
| Meis1 | ENSMUSG( | ENSMUSG( | 0.787783 | 0.288272 | NO | NO | NA |
| Olfr524 | ENSMUSG( | ENSMUSG( | 1.407102 | 0.007955 | YES | YES | Upregulated |
| Coq4 | ENSMUSG( | ENSMUSG( | -0.42611 | 0.000673 | YES | NO | NA |
| Rpl19-ps1 | ENSMUSG( | ENSMUSG( | 0.214812 | 0.338522 | NO | NO | NA |
| Kctd6 | ENSMUSG( | ENSMUSG( | 0.079496 | 0.531711 | NO | NO | NA |
| Ces2g | ENSMUSG( | ENSMUSG( | 1.052395 | 0.121179 | NO | NO | NA |
| Fnip2 | ENSMUSG( | ENSMUSG( | 0.017848 | 0.914861 | NO | NO | NA |
| G530011O | ENSMUSG( | ENSMUSG( | 0.761394 | 0.074597 | NO | NO | NA |
| Gxylt2 | ENSMUSG( | ENSMUSG( | 0.797354 | 0.534553 | NO | NO | NA |
| Rdh12 | ENSMUSG( | ENSMUSG( | -0.56254 | 0.008175 | YES | NO | NA |
| Msl3l2 | ENSMUSG( | ENSMUSG( | 0.782523 | 0.008554 | YES | NO | NA |
| ENSMUSG( | ENSMUSG( | ENSMUSG( | 0.147874 | 0.39796 | NO | NO | NA |
| Gm12184 | ENSMUSG( | ENSMUSG( | 0.670544 | 0.001558 | YES | NO | NA |
| Kalrn | ENSMUSG( | ENSMUSG( | 0.67309 | 0.399941 | NO | NO | NA |
| Ankrd9 | ENSMUSG( | ENSMUSG( | 0.208758 | 0.11693 | NO | NO | NA |

| | | | | | | | |
|---|---|---|---|---|---|---|---|
| Tspan33 | ENSMUSG( | ENSMUSG( | 0.956979 | 0.113545 | NO | NO | NA |
| Zfp658 | ENSMUSG( | ENSMUSG( | -0.28314 | 0.070398 | NO | NO | NA |
| ENSMUSG | ENSMUSG( | ENSMUSG( | -0.30182 | 0.149053 | NO | NO | NA |
| BC025920 | ENSMUSG( | ENSMUSG( | -0.09264 | 0.583116 | NO | NO | NA |
| Fkbp10 | ENSMUSG( | ENSMUSG( | -2.88155 | 5.87E-08 | YES | YES | Downregulated |
| Zcchc3 | ENSMUSG( | ENSMUSG( | 0.686762 | 0.000203 | YES | NO | NA |
| Gm17552 | ENSMUSG( | ENSMUSG( | 0.052396 | 0.810919 | NO | NO | NA |
| Zfp992 | ENSMUSG( | ENSMUSG( | 1.338155 | 5.85E-11 | YES | YES | Upregulated |
| Gm50166 | ENSMUSG( | ENSMUSG( | 0.739415 | 0.421281 | NO | NO | NA |
| Ppp1r3e | ENSMUSG( | ENSMUSG( | 0.226608 | 0.103122 | NO | NO | NA |
| Mtrf1 | ENSMUSG( | ENSMUSG( | 0.33115 | 0.034255 | YES | NO | NA |
| Gm14305 | ENSMUSG( | ENSMUSG( | 0.174098 | 0.432526 | NO | NO | NA |
| Slc66a1 | ENSMUSG( | ENSMUSG( | -0.30958 | 0.060434 | NO | NO | NA |
| Gm9522 | ENSMUSG( | ENSMUSG( | 1.271876 | 0.475251 | NO | NO | NA |
| Gm10052 | ENSMUSG( | ENSMUSG( | 0.475001 | 8.67E-05 | YES | NO | NA |
| Il18bp | ENSMUSG( | ENSMUSG( | -0.54728 | 0.000691 | YES | NO | NA |
| Mir155hg | ENSMUSG( | ENSMUSG( | -0.59942 | 0.011147 | YES | NO | NA |
| Ift88 | ENSMUSG( | ENSMUSG( | 1.098484 | 0.000203 | YES | YES | Upregulated |
| Ltk | ENSMUSG( | ENSMUSG( | -0.55023 | 0.012148 | YES | NO | NA |
| Mtmr7 | ENSMUSG( | ENSMUSG( | -1.42027 | 2.29E-10 | YES | YES | Downregulated |
| Trav6-6 | ENSMUSG( | ENSMUSG( | -0.68827 | 0.729736 | NO | NO | NA |
| Arhgef10 | ENSMUSG( | ENSMUSG( | 0.328815 | 0.007443 | YES | NO | NA |
| 9330151L1 | ENSMUSG( | ENSMUSG( | -0.18847 | 0.477095 | NO | NO | NA |
| Rnf130 | ENSMUSG( | ENSMUSG( | 0.071495 | 0.688954 | NO | NO | NA |
| Pkd1l3 | ENSMUSG( | ENSMUSG( | 0.826217 | 0.001161 | YES | NO | NA |
| Gm10762 | ENSMUSG( | ENSMUSG( | 0.251654 | 0.113475 | NO | NO | NA |
| Lyn | ENSMUSG( | ENSMUSG( | 0.23174 | 0.42412 | NO | NO | NA |
| ENSMUSG | ENSMUSG( | ENSMUSG( | 0.259367 | 0.16788 | NO | NO | NA |
| Ralgps2 | ENSMUSG( | ENSMUSG( | -0.06548 | 0.735159 | NO | NO | NA |
| Zfp125 | ENSMUSG( | ENSMUSG( | -0.62836 | 0.000801 | YES | NO | NA |
| Kbtbd6 | ENSMUSG( | ENSMUSG( | 0.226873 | 0.186566 | NO | NO | NA |
| Iglc1 | ENSMUSG( | ENSMUSG( | -0.76945 | 0.590865 | NO | NO | NA |
| Trav4-4-dv | ENSMUSG( | ENSMUSG( | 0.064504 | 0.929293 | NO | NO | NA |
| Sox4 | ENSMUSG( | ENSMUSG( | 0.825452 | 0.030476 | YES | NO | NA |
| Tmem267 | ENSMUSG( | ENSMUSG( | -0.26951 | 0.224595 | NO | NO | NA |
| Tmem106a | ENSMUSG( | ENSMUSG( | -0.54472 | 5.14E-05 | YES | NO | NA |
| Tdrd7 | ENSMUSG( | ENSMUSG( | -0.54595 | 0.001509 | YES | NO | NA |
| Lzts1 | ENSMUSG( | ENSMUSG( | 0.839335 | 0.000281 | YES | NO | NA |
| 4930513N | ENSMUSG( | ENSMUSG( | 0.217417 | 0.4519 | NO | NO | NA |
| Tpk1 | ENSMUSG( | ENSMUSG( | -0.12591 | 0.409568 | NO | NO | NA |
| Ro60 | ENSMUSG( | ENSMUSG( | 0.013548 | 0.946237 | NO | NO | NA |
| Tmem41a | ENSMUSG( | ENSMUSG( | 0.554829 | 0.00637 | YES | NO | NA |
| Trim46 | ENSMUSG( | ENSMUSG( | -0.25039 | 0.10042 | NO | NO | NA |
| Napepld | ENSMUSG( | ENSMUSG( | -0.1312 | 0.468284 | NO | NO | NA |
| Wfikkn2 | ENSMUSG( | ENSMUSG( | -0.70419 | 0.009552 | YES | NO | NA |
| 4930522L1 | ENSMUSG( | ENSMUSG( | -0.03162 | 0.863904 | NO | NO | NA |
| Olfr525 | ENSMUSG( | ENSMUSG( | 1.7725 | 0.000161 | YES | YES | Upregulated |

| | | | | | | | |
|---|---|---|---|---|---|---|---|
| Gm20300 | ENSMUSG( | ENSMUSG( | -0.23159 | 0.211455 | NO | NO | NA |
| Kyat1 | ENSMUSG( | ENSMUSG( | -0.31541 | 0.102846 | NO | NO | NA |
| Tmem69 | ENSMUSG( | ENSMUSG( | -0.04032 | 0.83839 | NO | NO | NA |
| Zfp763 | ENSMUSG( | ENSMUSG( | -0.30407 | 0.162825 | NO | NO | NA |
| Dhrs13 | ENSMUSG( | ENSMUSG( | 0.382185 | 0.030571 | YES | NO | NA |
| ENSMUSG | ENSMUSG( | ENSMUSG( | 0.320049 | 0.431435 | NO | NO | NA |
| Mkks | ENSMUSG( | ENSMUSG( | -0.45456 | 0.022793 | YES | NO | NA |
| Gm35558 | ENSMUSG( | ENSMUSG( | -0.52544 | 0.01218 | YES | NO | NA |
| Phkg1 | ENSMUSG( | ENSMUSG( | 0.718257 | 0.361115 | NO | NO | NA |
| Igkv12-44 | ENSMUSG( | ENSMUSG( | 0.783715 | 0.702923 | NO | NO | NA |
| Ighv8-8 | ENSMUSG( | ENSMUSG( | 0.066479 | 0.976454 | NO | NO | NA |
| Gm7846 | ENSMUSG( | ENSMUSG( | 0.295086 | 0.238227 | NO | NO | NA |
| Gm4735 | ENSMUSG( | ENSMUSG( | 0.523672 | 0.045813 | YES | NO | NA |
| Gm17173 | ENSMUSG( | ENSMUSG( | -0.53387 | 0.007857 | YES | NO | NA |
| Gm4924 | ENSMUSG( | ENSMUSG( | -0.12496 | 0.397264 | NO | NO | NA |
| Gm49797 | ENSMUSG( | ENSMUSG( | 0.037082 | 0.854072 | NO | NO | NA |
| Dhdh | ENSMUSG( | ENSMUSG( | 0.674175 | 0.006502 | YES | NO | NA |
| Cd22 | ENSMUSG( | ENSMUSG( | 0.989927 | 0.003895 | YES | NO | NA |
| Wdr35 | ENSMUSG( | ENSMUSG( | 1.56703 | 3.16E-06 | YES | YES | Upregulated |
| Gm6545 | ENSMUSG( | ENSMUSG( | -1.44087 | 1.51E-09 | YES | YES | Downregulated |
| Particl | ENSMUSG( | ENSMUSG( | -0.02977 | 0.896339 | NO | NO | NA |
| Gca | ENSMUSG( | ENSMUSG( | 0.176695 | 0.374035 | NO | NO | NA |
| Dlg2 | ENSMUSG( | ENSMUSG( | 1.126225 | 2.17E-09 | YES | YES | Upregulated |
| Nhej1 | ENSMUSG( | ENSMUSG( | 0.104618 | 0.70411 | NO | NO | NA |
| Rad9b | ENSMUSG( | ENSMUSG( | 0.139247 | 0.379041 | NO | NO | NA |
| Pawr | ENSMUSG( | ENSMUSG( | 0.084098 | 0.625261 | NO | NO | NA |
| Psd | ENSMUSG( | ENSMUSG( | 0.439016 | 0.015752 | YES | NO | NA |
| Dscam | ENSMUSG( | ENSMUSG( | 1.636134 | 7.34E-11 | YES | YES | Upregulated |
| Dock7 | ENSMUSG( | ENSMUSG( | -0.19528 | 0.477841 | NO | NO | NA |
| Gng3 | ENSMUSG( | ENSMUSG( | -0.34364 | 0.212284 | NO | NO | NA |
| Gm21596 | ENSMUSG( | ENSMUSG( | 0.435199 | 0.006529 | YES | NO | NA |
| Myo3b | ENSMUSG( | ENSMUSG( | -0.07626 | 0.734564 | NO | NO | NA |
| Dus4l | ENSMUSG( | ENSMUSG( | 0.739765 | 0.000148 | YES | NO | NA |
| Fbxo27 | ENSMUSG( | ENSMUSG( | -0.43891 | 0.037662 | YES | NO | NA |
| Gm9856 | ENSMUSG( | ENSMUSG( | -0.03059 | 0.89535 | NO | NO | NA |
| Snx16 | ENSMUSG( | ENSMUSG( | -0.14714 | 0.364958 | NO | NO | NA |
| Tbx3 | ENSMUSG( | ENSMUSG( | 0.795599 | 0.519345 | NO | NO | NA |
| Gm17344 | ENSMUSG( | ENSMUSG( | -0.14771 | 0.474914 | NO | NO | NA |
| Gm8730 | ENSMUSG( | ENSMUSG( | 0.36951 | 0.175013 | NO | NO | NA |
| Lyrm1 | ENSMUSG( | ENSMUSG( | 0.473593 | 7.71E-05 | YES | NO | NA |
| Prelid2 | ENSMUSG( | ENSMUSG( | 0.616555 | 0.00028 | YES | NO | NA |
| 5830405F( | ENSMUSG( | ENSMUSG( | -0.90499 | 0.003985 | YES | NO | NA |
| Gkap1 | ENSMUSG( | ENSMUSG( | 0.738094 | 0.00146 | YES | NO | NA |
| Gm7536 | ENSMUSG( | ENSMUSG( | 0.235694 | 0.254114 | NO | NO | NA |
| Dnajb2 | ENSMUSG( | ENSMUSG( | 0.303348 | 0.034255 | YES | NO | NA |
| Kifap3 | ENSMUSG( | ENSMUSG( | 1.000073 | 0.005443 | YES | YES | Upregulated |
| Zdhhc23 | ENSMUSG( | ENSMUSG( | 0.130809 | 0.501479 | NO | NO | NA |

| | | | | | | | |
|---|---|---|---|---|---|---|---|
| Myo7a | ENSMUSG( | ENSMUSG( | -0.36115 | 0.004622 | YES | NO | NA |
| Hykk | ENSMUSG( | ENSMUSG( | 0.86412 | 0.199499 | NO | NO | NA |
| Igkv19-93 | ENSMUSG( | ENSMUSG( | -0.98249 | 0.764367 | NO | NO | NA |
| Ccdc141 | ENSMUSG( | ENSMUSG( | -0.12935 | 0.520088 | NO | NO | NA |
| Gm42141 | ENSMUSG( | ENSMUSG( | -0.71008 | 0.000169 | YES | NO | NA |
| Tmem218 | ENSMUSG( | ENSMUSG( | 0.211796 | 0.3724 | NO | NO | NA |
| Mlf1 | ENSMUSG( | ENSMUSG( | 0.659662 | 0.019845 | YES | NO | NA |
| Gm47484 | ENSMUSG( | ENSMUSG( | -0.05888 | 0.781352 | NO | NO | NA |
| Ptov1 | ENSMUSG( | ENSMUSG( | 0.383937 | 0.104473 | NO | NO | NA |
| E2f5 | ENSMUSG( | ENSMUSG( | 0.038622 | 0.778559 | NO | NO | NA |
| Phlda3 | ENSMUSG( | ENSMUSG( | -0.19876 | 0.419387 | NO | NO | NA |
| Cdk5r1 | ENSMUSG( | ENSMUSG( | -0.01864 | 0.933155 | NO | NO | NA |
| Zfp248 | ENSMUSG( | ENSMUSG( | 0.176798 | 0.461948 | NO | NO | NA |
| Gm48226 | ENSMUSG( | ENSMUSG( | 0.260849 | 0.091758 | NO | NO | NA |
| Il21 | ENSMUSG( | ENSMUSG( | 0.356584 | 0.387041 | NO | NO | NA |
| Zfp873 | ENSMUSG( | ENSMUSG( | -0.24014 | 0.11977 | NO | NO | NA |
| Jup | ENSMUSG( | ENSMUSG( | -1.47856 | 2.62E-05 | YES | YES | Downregulated |
| Atp9a | ENSMUSG( | ENSMUSG( | 0.282299 | 0.291985 | NO | NO | NA |
| Gm3435 | ENSMUSG( | ENSMUSG( | -0.07615 | 0.872995 | NO | NO | NA |
| Zfp536 | ENSMUSG( | ENSMUSG( | 0.681335 | 0.428944 | NO | NO | NA |
| Rpl10a-ps1 | ENSMUSG( | ENSMUSG( | 0.519975 | 0.010562 | YES | NO | NA |
| Trav6-1 | ENSMUSG( | ENSMUSG( | -0.59005 | 0.518069 | NO | NO | NA |
| Gm12359 | ENSMUSG( | ENSMUSG( | -0.32251 | 0.075635 | NO | NO | NA |
| Mtus2 | ENSMUSG( | ENSMUSG( | 0.280509 | 0.569187 | NO | NO | NA |
| Armcx2 | ENSMUSG( | ENSMUSG( | 0.02645 | 0.928071 | NO | NO | NA |
| Nek3 | ENSMUSG( | ENSMUSG( | -0.43106 | 0.00207 | YES | NO | NA |
| ENSMUSG( | ENSMUSG( | ENSMUSG( | -0.45564 | 0.009389 | YES | NO | NA |
| Zdhhc1 | ENSMUSG( | ENSMUSG( | 0.708987 | 0.000351 | YES | NO | NA |
| Mtln | ENSMUSG( | ENSMUSG( | 0.774372 | 0.000162 | YES | NO | NA |
| Zfp637 | ENSMUSG( | ENSMUSG( | -0.06192 | 0.772141 | NO | NO | NA |
| Gm14296 | ENSMUSG( | ENSMUSG( | 0.096082 | 0.874011 | NO | NO | NA |
| Zfp433 | ENSMUSG( | ENSMUSG( | -0.09828 | 0.731835 | NO | NO | NA |
| Gm49338 | ENSMUSG( | ENSMUSG( | -0.32505 | 0.102099 | NO | NO | NA |
| Tspan4 | ENSMUSG( | ENSMUSG( | 0.462841 | 0.023524 | YES | NO | NA |
| Gm49037 | ENSMUSG( | ENSMUSG( | 0.206622 | 0.664472 | NO | NO | NA |
| Vav2 | ENSMUSG( | ENSMUSG( | 0.307343 | 0.164377 | NO | NO | NA |
| Plcxd1 | ENSMUSG( | ENSMUSG( | -0.00962 | 0.962708 | NO | NO | NA |
| Vmn2r53 | ENSMUSG( | ENSMUSG( | 0.576662 | 0.613662 | NO | NO | NA |
| Cpne7 | ENSMUSG( | ENSMUSG( | -1.01797 | 0.001355 | YES | YES | Downregulated |
| Ikbip | ENSMUSG( | ENSMUSG( | 0.232524 | 0.391281 | NO | NO | NA |
| Spef1 | ENSMUSG( | ENSMUSG( | -0.12355 | 0.73702 | NO | NO | NA |
| Eid2 | ENSMUSG( | ENSMUSG( | 1.150251 | 6.38E-05 | YES | YES | Upregulated |
| Ninj1 | ENSMUSG( | ENSMUSG( | 0.551685 | 0.005017 | YES | NO | NA |
| P4htm | ENSMUSG( | ENSMUSG( | 0.46682 | 0.001595 | YES | NO | NA |
| H2bc12 | ENSMUSG( | ENSMUSG( | 0.723622 | 0.000201 | YES | NO | NA |
| Gm37254 | ENSMUSG( | ENSMUSG( | -0.32447 | 0.115108 | NO | NO | NA |
| Prrx1 | ENSMUSG( | ENSMUSG( | 0.982937 | 0.23664 | NO | NO | NA |

| | | | | | | | |
|---|---|---|---|---|---|---|---|
| Gm30373 | ENSMUSG( | ENSMUSG( | 0.506991 | 0.006808 | YES | NO | NA |
| Slc35e4 | ENSMUSG( | ENSMUSG( | -0.6749 | 2.60E-06 | YES | NO | NA |
| Tmem170 | ENSMUSG( | ENSMUSG( | -0.11391 | 0.562748 | NO | NO | NA |
| Slc31a2 | ENSMUSG( | ENSMUSG( | -0.32486 | 0.012463 | YES | NO | NA |
| Paqr3 | ENSMUSG( | ENSMUSG( | 0.316463 | 0.05358 | NO | NO | NA |
| Hemk1 | ENSMUSG( | ENSMUSG( | -0.10535 | 0.664907 | NO | NO | NA |
| Rundc3a | ENSMUSG( | ENSMUSG( | 0.433521 | 0.209647 | NO | NO | NA |
| 1700030C: | ENSMUSG( | ENSMUSG( | 1.304136 | 0.054673 | NO | NO | NA |
| Rangrf | ENSMUSG( | ENSMUSG( | 1.07018 | 9.04E-05 | YES | YES | Upregulated |
| Tcn2 | ENSMUSG( | ENSMUSG( | -0.93728 | 2.01E-05 | YES | NO | NA |
| Trav7d-4 | ENSMUSG( | ENSMUSG( | -0.02602 | 0.979373 | NO | NO | NA |
| H4c17 | ENSMUSG( | ENSMUSG( | 0.485336 | 0.022853 | YES | NO | NA |
| Star | ENSMUSG( | ENSMUSG( | 0.19761 | 0.381121 | NO | NO | NA |
| Igf2bp1 | ENSMUSG( | ENSMUSG( | 0.632643 | 0.616516 | NO | NO | NA |
| Acrbp | ENSMUSG( | ENSMUSG( | -0.33632 | 0.045859 | YES | NO | NA |
| Rubcnl | ENSMUSG( | ENSMUSG( | 0.715119 | 3.26E-07 | YES | NO | NA |
| Gm12744 | ENSMUSG( | ENSMUSG( | 1.155436 | 2.89E-05 | YES | YES | Upregulated |
| ENSMUSG | ENSMUSG( | ENSMUSG( | -0.16376 | 0.415723 | NO | NO | NA |
| 2500002B: | ENSMUSG( | ENSMUSG( | 0.008556 | 0.976785 | NO | NO | NA |
| Gnb1l | ENSMUSG( | ENSMUSG( | 0.044335 | 0.797821 | NO | NO | NA |
| Creb5 | ENSMUSG( | ENSMUSG( | 0.708817 | 0.574941 | NO | NO | NA |
| H2ac6 | ENSMUSG( | ENSMUSG( | 0.591815 | 5.93E-05 | YES | NO | NA |
| Sall2 | ENSMUSG( | ENSMUSG( | 0.539224 | 0.001047 | YES | NO | NA |
| Trav12d-2 | ENSMUSG( | ENSMUSG( | -0.71602 | 0.774247 | NO | NO | NA |
| Zfp691 | ENSMUSG( | ENSMUSG( | -0.30119 | 0.079589 | NO | NO | NA |
| Tff1 | ENSMUSG( | ENSMUSG( | 1.203048 | 0.014081 | YES | YES | Upregulated |
| A730011C | ENSMUSG( | ENSMUSG( | 0.056754 | 0.749566 | NO | NO | NA |
| Rps27rt | ENSMUSG( | ENSMUSG( | -0.08826 | 0.641748 | NO | NO | NA |
| Dand5 | ENSMUSG( | ENSMUSG( | -0.51565 | 0.007818 | YES | NO | NA |
| Tyw3 | ENSMUSG( | ENSMUSG( | 0.229948 | 0.167592 | NO | NO | NA |
| Gm49747 | ENSMUSG( | ENSMUSG( | -0.26686 | 0.250943 | NO | NO | NA |
| Prkab2 | ENSMUSG( | ENSMUSG( | -0.40879 | 0.000936 | YES | NO | NA |
| Tgif2 | ENSMUSG( | ENSMUSG( | 0.072996 | 0.685592 | NO | NO | NA |
| Snx8 | ENSMUSG( | ENSMUSG( | 0.393055 | 0.019845 | YES | NO | NA |
| Cfap418 | ENSMUSG( | ENSMUSG( | 0.615138 | 0.009686 | YES | NO | NA |
| Apold1 | ENSMUSG( | ENSMUSG( | 0.626735 | 0.008379 | YES | NO | NA |
| Pgpep1 | ENSMUSG( | ENSMUSG( | 0.108348 | 0.668038 | NO | NO | NA |
| Marchf9 | ENSMUSG( | ENSMUSG( | -0.08431 | 0.672518 | NO | NO | NA |
| Gm38365 | ENSMUSG( | ENSMUSG( | 0.574915 | 0.415514 | NO | NO | NA |
| Pkn3 | ENSMUSG( | ENSMUSG( | -0.11414 | 0.616241 | NO | NO | NA |
| E130317F: | ENSMUSG( | ENSMUSG( | 0.206728 | 0.3276 | NO | NO | NA |
| Stk3 | ENSMUSG( | ENSMUSG( | -0.07314 | 0.732555 | NO | NO | NA |
| Zmat1 | ENSMUSG( | ENSMUSG( | -0.44777 | 0.044387 | YES | NO | NA |
| Gm47483 | ENSMUSG( | ENSMUSG( | -0.6205 | 0.004146 | YES | NO | NA |
| Gm37033 | ENSMUSG( | ENSMUSG( | -0.3332 | 0.385362 | NO | NO | NA |
| Ddah2 | ENSMUSG( | ENSMUSG( | 0.15286 | 0.390053 | NO | NO | NA |
| Mical3 | ENSMUSG( | ENSMUSG( | 0.614523 | 0.105237 | NO | NO | NA |

| | | | | | | | |
|---|---|---|---|---|---|---|---|
| Gm37760 | ENSMUSG( | ENSMUSG( | 0.169634 | 0.692132 | NO | NO | NA |
| Fcer1g | ENSMUSG( | ENSMUSG( | 0.756809 | 0.165658 | NO | NO | NA |
| Trav3d-3 | ENSMUSG( | ENSMUSG( | -0.37356 | 0.860006 | NO | NO | NA |
| Gm14434 | ENSMUSG( | ENSMUSG( | 0.711062 | 0.412416 | NO | NO | NA |
| Nudt2 | ENSMUSG( | ENSMUSG( | 0.667007 | 0.000267 | YES | NO | NA |
| Nrxn3 | ENSMUSG( | ENSMUSG( | 0.565684 | 0.567007 | NO | NO | NA |
| B3gnt9 | ENSMUSG( | ENSMUSG( | -0.19002 | 0.195893 | NO | NO | NA |
| Gm8818 | ENSMUSG( | ENSMUSG( | 0.803193 | 0.008184 | YES | NO | NA |
| Gm2788 | ENSMUSG( | ENSMUSG( | -0.19607 | 0.560998 | NO | NO | NA |
| 9930012K: | ENSMUSG( | ENSMUSG( | -0.37499 | 0.0388 | YES | NO | NA |
| Abcg2 | ENSMUSG( | ENSMUSG( | 0.572269 | 0.001397 | YES | NO | NA |
| Ccdc90b | ENSMUSG( | ENSMUSG( | 0.517442 | 0.004552 | YES | NO | NA |
| 4632415L( | ENSMUSG( | ENSMUSG( | -0.00275 | 0.989686 | NO | NO | NA |
| 6720489N | ENSMUSG( | ENSMUSG( | -0.38964 | 0.067592 | NO | NO | NA |
| Trgc1 | ENSMUSG( | ENSMUSG( | -1.29984 | 0.005848 | YES | YES | Downregulated |
| C130036L: | ENSMUSG( | ENSMUSG( | 0.292486 | 0.171026 | NO | NO | NA |
| Gm45774 | ENSMUSG( | ENSMUSG( | 1.42706 | 0.000858 | YES | YES | Upregulated |
| 1700109H | ENSMUSG( | ENSMUSG( | -0.18014 | 0.291953 | NO | NO | NA |
| Igip | ENSMUSG( | ENSMUSG( | -0.26403 | 0.088713 | NO | NO | NA |
| Sfxn5 | ENSMUSG( | ENSMUSG( | 0.644602 | 0.149455 | NO | NO | NA |
| 6230400D | ENSMUSG( | ENSMUSG( | -0.05608 | 0.790468 | NO | NO | NA |
| Smim26 | ENSMUSG( | ENSMUSG( | 0.081049 | 0.688851 | NO | NO | NA |
| Gm26708 | ENSMUSG( | ENSMUSG( | 5.103194 | NA | NO | NO | NA |
| Trip6 | ENSMUSG( | ENSMUSG( | -0.28246 | 0.160317 | NO | NO | NA |
| Eif3j2 | ENSMUSG( | ENSMUSG( | 0.355813 | 0.244423 | NO | NO | NA |
| Rps7-ps3 | ENSMUSG( | ENSMUSG( | 0.641359 | 0.576784 | NO | NO | NA |
| Dnph1 | ENSMUSG( | ENSMUSG( | 1.647392 | 6.95E-08 | YES | YES | Upregulated |
| Enah | ENSMUSG( | ENSMUSG( | 0.630017 | 0.526045 | NO | NO | NA |
| Gm17066 | ENSMUSG( | ENSMUSG( | -0.01042 | 0.954102 | NO | NO | NA |
| Cacng2 | ENSMUSG( | ENSMUSG( | 0.363513 | 0.684648 | NO | NO | NA |
| Rnf227 | ENSMUSG( | ENSMUSG( | -0.07271 | 0.684536 | NO | NO | NA |
| Mettl15 | ENSMUSG( | ENSMUSG( | 0.048827 | 0.803948 | NO | NO | NA |
| Nxnl1 | ENSMUSG( | ENSMUSG( | -0.33619 | 0.05601 | NO | NO | NA |
| 4930486L: | ENSMUSG( | ENSMUSG( | 0.312016 | 0.362173 | NO | NO | NA |
| Ablim2 | ENSMUSG( | ENSMUSG( | -0.42848 | 0.00565 | YES | NO | NA |
| Gm39326 | ENSMUSG( | ENSMUSG( | -0.28562 | 0.519025 | NO | NO | NA |
| Kctd1 | ENSMUSG( | ENSMUSG( | -0.22509 | 0.146071 | NO | NO | NA |
| Armcx3 | ENSMUSG( | ENSMUSG( | -0.84151 | 0.007469 | YES | NO | NA |
| Tet1 | ENSMUSG( | ENSMUSG( | 0.389425 | 0.424258 | NO | NO | NA |
| Ctnnal1 | ENSMUSG( | ENSMUSG( | 0.962611 | 9.53E-07 | YES | NO | NA |
| Nubpl | ENSMUSG( | ENSMUSG( | 0.737337 | 0.002147 | YES | NO | NA |
| Rps3a2 | ENSMUSG( | ENSMUSG( | 0.528537 | 0.007245 | YES | NO | NA |
| Gnal | ENSMUSG( | ENSMUSG( | -0.07894 | 0.715997 | NO | NO | NA |
| Bcl11a | ENSMUSG( | ENSMUSG( | 0.87733 | 0.077337 | NO | NO | NA |
| Rnf113a1 | ENSMUSG( | ENSMUSG( | 0.174025 | 0.425984 | NO | NO | NA |
| Myl6b | ENSMUSG( | ENSMUSG( | 0.160039 | 0.522588 | NO | NO | NA |
| Syce1 | ENSMUSG( | ENSMUSG( | 2.306731 | 0.377048 | NO | NO | NA |

| | | | | | | | |
|---|---|---|---|---|---|---|---|
| Anxa9 | ENSMUSG( | ENSMUSG( | 0.899281 | 0.002572 | YES | NO | NA |
| Elavl3 | ENSMUSG( | ENSMUSG( | 0.434018 | 0.614865 | NO | NO | NA |
| Ada | ENSMUSG( | ENSMUSG( | 0.220745 | 0.227436 | NO | NO | NA |
| Pcbp3 | ENSMUSG( | ENSMUSG( | 0.848667 | 0.027124 | YES | NO | NA |
| Gm42653 | ENSMUSG( | ENSMUSG( | 0.703578 | 0.22411 | NO | NO | NA |
| Adgrl3 | ENSMUSG( | ENSMUSG( | 0.65658 | 0.609219 | NO | NO | NA |
| Gm26581 | ENSMUSG( | ENSMUSG( | -0.46884 | 0.147596 | NO | NO | NA |
| Rps4l | ENSMUSG( | ENSMUSG( | 0.531902 | 0.426371 | NO | NO | NA |
| Atp1b2 | ENSMUSG( | ENSMUSG( | 0.920789 | 0.220641 | NO | NO | NA |
| ENSMUSG | ENSMUSG( | ENSMUSG( | -0.39448 | 0.003972 | YES | NO | NA |
| 1700096K: | ENSMUSG( | ENSMUSG( | 0.379721 | 0.168155 | NO | NO | NA |
| C130073E: | ENSMUSG( | ENSMUSG( | 0.559254 | 0.646606 | NO | NO | NA |
| Foxp2 | ENSMUSG( | ENSMUSG( | 0.702062 | 0.473607 | NO | NO | NA |
| Gm12918 | ENSMUSG( | ENSMUSG( | 0.49319 | 0.034531 | YES | NO | NA |
| Zfp563 | ENSMUSG( | ENSMUSG( | -0.60581 | 0.012957 | YES | NO | NA |
| Gm42549 | ENSMUSG( | ENSMUSG( | -0.46668 | 0.004026 | YES | NO | NA |
| Plekho1 | ENSMUSG( | ENSMUSG( | 0.213107 | 0.339968 | NO | NO | NA |
| Rian | ENSMUSG( | ENSMUSG( | 0.989611 | 0.004099 | YES | NO | NA |
| Trbv26 | ENSMUSG( | ENSMUSG( | 0.04362 | 0.924763 | NO | NO | NA |
| Gm26850 | ENSMUSG( | ENSMUSG( | -0.04331 | 0.773948 | NO | NO | NA |
| Sdhaf3 | ENSMUSG( | ENSMUSG( | 0.74587 | 0.000122 | YES | NO | NA |
| Celf4 | ENSMUSG( | ENSMUSG( | 0.775384 | 0.042681 | YES | NO | NA |
| Gm47662 | ENSMUSG( | ENSMUSG( | 1.55076 | 0.000346 | YES | YES | Upregulated |
| Clip3 | ENSMUSG( | ENSMUSG( | 0.778519 | 0.003952 | YES | NO | NA |
| Wdr25 | ENSMUSG( | ENSMUSG( | 0.282211 | 0.131806 | NO | NO | NA |
| 2810408I1 | ENSMUSG( | ENSMUSG( | 0.757469 | 1.45E-05 | YES | NO | NA |
| Map1lc3a | ENSMUSG( | ENSMUSG( | 0.075589 | 0.727172 | NO | NO | NA |
| Dnajc12 | ENSMUSG( | ENSMUSG( | 0.272779 | 0.31893 | NO | NO | NA |
| Zfp973 | ENSMUSG( | ENSMUSG( | -0.3405 | 0.627411 | NO | NO | NA |
| Dock4 | ENSMUSG( | ENSMUSG( | 0.613654 | 0.422503 | NO | NO | NA |
| ENSMUSG( | ENSMUSG( | ENSMUSG( | 0.196879 | 0.527914 | NO | NO | NA |
| Hsd17b11 | ENSMUSG( | ENSMUSG( | -0.74383 | 0.007565 | YES | NO | NA |
| Hand2os1 | ENSMUSG( | ENSMUSG( | 0.911161 | 0.24151 | NO | NO | NA |
| Gm6477 | ENSMUSG( | ENSMUSG( | 0.63171 | 0.01203 | YES | NO | NA |
| Rpl15-ps3 | ENSMUSG( | ENSMUSG( | 0.528524 | 0.063915 | NO | NO | NA |
| Gm11175 | ENSMUSG( | ENSMUSG( | 0.304136 | 0.136138 | NO | NO | NA |
| Vsig10 | ENSMUSG( | ENSMUSG( | -0.51621 | 0.318148 | NO | NO | NA |
| Eid2b | ENSMUSG( | ENSMUSG( | 0.195627 | 0.292632 | NO | NO | NA |
| Armc3 | ENSMUSG( | ENSMUSG( | -0.58389 | 0.00946 | YES | NO | NA |
| Tmem216 | ENSMUSG( | ENSMUSG( | 0.631536 | 0.005552 | YES | NO | NA |
| Gm10557 | ENSMUSG( | ENSMUSG( | 0.152124 | 0.611373 | NO | NO | NA |
| Cryz | ENSMUSG( | ENSMUSG( | 0.451252 | 0.004949 | YES | NO | NA |
| ENSMUSG | ENSMUSG( | ENSMUSG( | 0.089812 | 0.825486 | NO | NO | NA |
| Trav10n | ENSMUSG( | ENSMUSG( | 0.524016 | 0.529107 | NO | NO | NA |
| 1700028E: | ENSMUSG( | ENSMUSG( | 0.465034 | 0.001809 | YES | NO | NA |
| Zfp518b | ENSMUSG( | ENSMUSG( | 0.664698 | 0.000142 | YES | NO | NA |
| ENSMUSG( | ENSMUSG( | ENSMUSG( | 0.275282 | 0.149155 | NO | NO | NA |

| | | | | | | | |
|---|---|---|---|---|---|---|---|
| Ggt7 | ENSMUSG( | ENSMUSG( | -0.12374 | 0.450569 | NO | NO | NA |
| 4930556H | ENSMUSG( | ENSMUSG( | -0.50252 | 0.021104 | YES | NO | NA |
| ENSMUSG( | ENSMUSG( | ENSMUSG( | 0.581584 | 0.00076 | YES | NO | NA |
| ENSMUSG( | ENSMUSG( | ENSMUSG( | -0.38741 | 0.390053 | NO | NO | NA |
| Fndc10 | ENSMUSG( | ENSMUSG( | 0.629289 | 0.048708 | YES | NO | NA |
| ENSMUSG | ENSMUSG( | ENSMUSG( | 0.18519 | 0.618809 | NO | NO | NA |
| Mnd1 | ENSMUSG( | ENSMUSG( | 0.882739 | 0.002564 | YES | NO | NA |
| Ppp1r3f | ENSMUSG( | ENSMUSG( | -0.06221 | 0.821271 | NO | NO | NA |
| Cyb561 | ENSMUSG( | ENSMUSG( | 1.014635 | 0.016828 | YES | YES | Upregulated |
| Zfp951 | ENSMUSG( | ENSMUSG( | 0.46926 | 0.051649 | NO | NO | NA |
| A930037H | ENSMUSG( | ENSMUSG( | -0.44955 | 0.000622 | YES | NO | NA |
| Gm49384 | ENSMUSG( | ENSMUSG( | 0.438432 | 0.087381 | NO | NO | NA |
| ENSMUSG | ENSMUSG( | ENSMUSG( | -0.83045 | 2.62E-05 | YES | NO | NA |
| Cyp4f17 | ENSMUSG( | ENSMUSG( | -0.21554 | 0.352768 | NO | NO | NA |
| Gm15545 | ENSMUSG( | ENSMUSG( | 0.076032 | 0.660302 | NO | NO | NA |
| Smim1 | ENSMUSG( | ENSMUSG( | 0.063792 | 0.861865 | NO | NO | NA |
| P2rx7 | ENSMUSG( | ENSMUSG( | -0.28173 | 0.509685 | NO | NO | NA |
| Arxes2 | ENSMUSG( | ENSMUSG( | 0.603168 | 0.001857 | YES | NO | NA |
| Gm14966 | ENSMUSG( | ENSMUSG( | -0.07286 | 0.758117 | NO | NO | NA |
| Rps16-ps2 | ENSMUSG( | ENSMUSG( | 0.325746 | 0.053263 | NO | NO | NA |
| Vmn2r96 | ENSMUSG( | ENSMUSG( | -0.41423 | 0.107665 | NO | NO | NA |
| Insr | ENSMUSG( | ENSMUSG( | 0.409686 | 0.171936 | NO | NO | NA |
| Snhg4 ENS | ENSMUSG( | ENSMUSG( | 1.399008 | 0.004363 | YES | YES | Upregulated |
| Igflr1 | ENSMUSG( | ENSMUSG( | 0.277043 | 0.281481 | NO | NO | NA |
| Trav8-1 | ENSMUSG( | ENSMUSG( | 0.990118 | 0.318088 | NO | NO | NA |
| Plxna1 | ENSMUSG( | ENSMUSG( | 0.442493 | 0.038365 | YES | NO | NA |
| Gm42429 | ENSMUSG( | ENSMUSG( | 9.525248 | 1.41E-07 | YES | YES | Upregulated |
| Syngr3 | ENSMUSG( | ENSMUSG( | -0.19784 | 0.40575 | NO | NO | NA |
| Nfe2 | ENSMUSG( | ENSMUSG( | 0.59582 | 0.000258 | YES | NO | NA |
| Runx1t1 | ENSMUSG( | ENSMUSG( | 0.73163 | 0.465954 | NO | NO | NA |
| Trav19 | ENSMUSG( | ENSMUSG( | 1.097346 | 0.250934 | NO | NO | NA |
| Tert | ENSMUSG( | ENSMUSG( | -0.22306 | 0.476831 | NO | NO | NA |
| B130024G | ENSMUSG( | ENSMUSG( | 0.533876 | 0.663671 | NO | NO | NA |
| Pklr | ENSMUSG( | ENSMUSG( | 0.806523 | 0.089446 | NO | NO | NA |
| Gm43679 | ENSMUSG( | ENSMUSG( | 0.67276 | 0.005514 | YES | NO | NA |
| Cyb5rl | ENSMUSG( | ENSMUSG( | 0.639808 | 0.00097 | YES | NO | NA |
| Zfp85 | ENSMUSG( | ENSMUSG( | -0.64838 | 0.003076 | YES | NO | NA |
| Ift172 | ENSMUSG( | ENSMUSG( | 0.234638 | 0.319438 | NO | NO | NA |
| Cfap91 | ENSMUSG( | ENSMUSG( | 2.094907 | 2.06E-11 | YES | YES | Upregulated |
| Zfp599 | ENSMUSG( | ENSMUSG( | -0.16272 | 0.41964 | NO | NO | NA |
| ENSMUSG( | ENSMUSG( | ENSMUSG( | 1.058954 | 0.042027 | YES | YES | Upregulated |
| Mylpf | ENSMUSG( | ENSMUSG( | 0.150462 | 0.395625 | NO | NO | NA |
| Gm28373 | ENSMUSG( | ENSMUSG( | -0.6703 | 0.000955 | YES | NO | NA |
| Ltf | ENSMUSG( | ENSMUSG( | 0.033187 | 0.9589 | NO | NO | NA |
| Rpl14-ps1 | ENSMUSG( | ENSMUSG( | 0.635342 | 0.001621 | YES | NO | NA |
| Dancr | ENSMUSG( | ENSMUSG( | 0.078439 | 0.738771 | NO | NO | NA |
| Col11a2 | ENSMUSG( | ENSMUSG( | -0.20674 | 0.202765 | NO | NO | NA |

| | | | | | | | |
|---|---|---|---|---|---|---|---|
| Vmn2r113 | ENSMUSG( | ENSMUSG( | 0.387826 | 0.760265 | NO | NO | NA |
| Gm42658 | ENSMUSG( | ENSMUSG( | -0.19611 | 0.358421 | NO | NO | NA |
| Fv1 | ENSMUSG( | ENSMUSG( | -0.25522 | 0.118516 | NO | NO | NA |
| Gm29666 | ENSMUSG( | ENSMUSG( | 0.502318 | 0.04237 | YES | NO | NA |
| Fitm2 | ENSMUSG( | ENSMUSG( | 0.125369 | 0.590846 | NO | NO | NA |
| Gpm6b | ENSMUSG( | ENSMUSG( | 1.130169 | 2.81E-08 | YES | YES | Upregulated |
| Mttp | ENSMUSG( | ENSMUSG( | 1.190799 | 4.25E-07 | YES | YES | Upregulated |
| 4933421O | ENSMUSG( | ENSMUSG( | 0.454285 | 0.143834 | NO | NO | NA |
| Gm4366 | ENSMUSG( | ENSMUSG( | 0.505781 | 2.02E-05 | YES | NO | NA |
| Ranbp17 | ENSMUSG( | ENSMUSG( | 0.246976 | 0.244982 | NO | NO | NA |
| Gm26510 | ENSMUSG( | ENSMUSG( | 0.218833 | 0.459449 | NO | NO | NA |
| Prss30 | ENSMUSG( | ENSMUSG( | 1.669827 | 9.88E-05 | YES | YES | Upregulated |
| ENSMUSG | ENSMUSG( | ENSMUSG( | 0.238921 | 0.199054 | NO | NO | NA |
| Gm49774 | ENSMUSG( | ENSMUSG( | 0.517056 | 0.278855 | NO | NO | NA |
| Cdh17 | ENSMUSG( | ENSMUSG( | 0.301446 | 0.429671 | NO | NO | NA |
| Wdsub1 | ENSMUSG( | ENSMUSG( | -0.0937 | 0.72269 | NO | NO | NA |
| Gm49883 | ENSMUSG( | ENSMUSG( | 0.117525 | 0.804412 | NO | NO | NA |
| 9130019O | ENSMUSG( | ENSMUSG( | -0.08273 | 0.642775 | NO | NO | NA |
| Gm45062 | ENSMUSG( | ENSMUSG( | -2.76243 | 0.508329 | NO | NO | NA |
| Deptor | ENSMUSG( | ENSMUSG( | 0.709614 | 0.243373 | NO | NO | NA |
| ENSMUSG( | ENSMUSG( | ENSMUSG( | -0.18674 | 0.44719 | NO | NO | NA |
| Gm7008 | ENSMUSG( | ENSMUSG( | 0.312901 | 0.306821 | NO | NO | NA |
| Ccdc191 | ENSMUSG( | ENSMUSG( | 0.128101 | 0.5876 | NO | NO | NA |
| 1500015A( | ENSMUSG( | ENSMUSG( | 0.47628 | 0.006607 | YES | NO | NA |
| Zfp239 | ENSMUSG( | ENSMUSG( | 0.288791 | 0.06047 | NO | NO | NA |
| Lacc1 | ENSMUSG( | ENSMUSG( | 0.947813 | 1.85E-05 | YES | NO | NA |
| Gm49392 | ENSMUSG( | ENSMUSG( | 0.428141 | 0.377109 | NO | NO | NA |
| Mcc | ENSMUSG( | ENSMUSG( | 0.587553 | 0.464937 | NO | NO | NA |
| Chchd6 | ENSMUSG( | ENSMUSG( | 1.405272 | 2.95E-05 | YES | YES | Upregulated |
| Gm10642 | ENSMUSG( | ENSMUSG( | -0.07467 | 0.770717 | NO | NO | NA |
| D7Bwg082 | ENSMUSG( | ENSMUSG( | 0.371286 | 0.007908 | YES | NO | NA |
| Tfcp2l1 | ENSMUSG( | ENSMUSG( | 0.930911 | 3.06E-05 | YES | NO | NA |
| Igkv1-117 | ENSMUSG( | ENSMUSG( | -0.74959 | 0.661684 | NO | NO | NA |
| 4930431P( | ENSMUSG( | ENSMUSG( | -0.18082 | 0.628805 | NO | NO | NA |
| Srl | ENSMUSG( | ENSMUSG( | 0.927899 | 0.001171 | YES | NO | NA |
| ENSMUSG | ENSMUSG( | ENSMUSG( | -1.37289 | 0.001284 | YES | YES | Downregulated |
| Gm38372 | ENSMUSG( | ENSMUSG( | -0.89283 | 1.75E-05 | YES | NO | NA |
| Vangl2 | ENSMUSG( | ENSMUSG( | 0.724568 | 0.352179 | NO | NO | NA |
| Gm9725 | ENSMUSG( | ENSMUSG( | -0.22138 | 0.602529 | NO | NO | NA |
| ENSMUSG( | ENSMUSG( | ENSMUSG( | 0.283671 | 0.108318 | NO | NO | NA |
| Gm29340 | ENSMUSG( | ENSMUSG( | 0.932233 | 0.108576 | NO | NO | NA |
| Zpbp | ENSMUSG( | ENSMUSG( | 0.247394 | 0.266593 | NO | NO | NA |
| Msh5 | ENSMUSG( | ENSMUSG( | -0.04043 | 0.862831 | NO | NO | NA |
| Serpinb1b | ENSMUSG( | ENSMUSG( | 0.474967 | 0.016037 | YES | NO | NA |
| Itgb3bp | ENSMUSG( | ENSMUSG( | 0.079461 | 0.644442 | NO | NO | NA |
| A930007I1 | ENSMUSG( | ENSMUSG( | -0.22156 | 0.343775 | NO | NO | NA |
| Map3k20 | ENSMUSG( | ENSMUSG( | 0.831749 | 0.065688 | NO | NO | NA |

| | | | | | | | |
|---|---|---|---|---|---|---|---|
| Gm31374 | ENSMUSG( | ENSMUSG( | -0.60858 | 0.04319 | YES | NO | NA |
| Mettl18 | ENSMUSG( | ENSMUSG( | -0.20395 | 0.17233 | NO | NO | NA |
| Tmtc3 | ENSMUSG( | ENSMUSG( | 1.312097 | 0.000155 | YES | YES | Upregulated |
| Ngp | ENSMUSG( | ENSMUSG( | 2.364103 | 0.18787 | NO | NO | NA |
| Tha1 | ENSMUSG( | ENSMUSG( | 0.498263 | 0.052697 | NO | NO | NA |
| Acvr1b | ENSMUSG( | ENSMUSG( | 0.332443 | 0.138443 | NO | NO | NA |
| Jrk | ENSMUSG( | ENSMUSG( | -0.08259 | 0.741426 | NO | NO | NA |
| Ift22 | ENSMUSG( | ENSMUSG( | 0.789058 | 0.004599 | YES | NO | NA |
| Gramd1c | ENSMUSG( | ENSMUSG( | -0.20924 | 0.305175 | NO | NO | NA |
| Dlgap1 | ENSMUSG( | ENSMUSG( | 0.614449 | 0.527421 | NO | NO | NA |
| Asah2 | ENSMUSG( | ENSMUSG( | 0.235717 | 0.439866 | NO | NO | NA |
| Gpt | ENSMUSG( | ENSMUSG( | -0.09375 | 0.541781 | NO | NO | NA |
| Gm16536 | ENSMUSG( | ENSMUSG( | -0.33337 | 0.05347 | NO | NO | NA |
| Trav3n-3 | ENSMUSG( | ENSMUSG( | 0.245109 | 0.801288 | NO | NO | NA |
| Trim71 | ENSMUSG( | ENSMUSG( | 0.62085 | 0.49424 | NO | NO | NA |
| Mthfd2l | ENSMUSG( | ENSMUSG( | 0.208716 | 0.170189 | NO | NO | NA |
| Il20ra | ENSMUSG( | ENSMUSG( | 0.849869 | 0.003705 | YES | NO | NA |
| Gm12258 | ENSMUSG( | ENSMUSG( | -0.31396 | 0.152477 | NO | NO | NA |
| Armc9 | ENSMUSG( | ENSMUSG( | 0.479875 | 0.008168 | YES | NO | NA |
| Tnnt1 | ENSMUSG( | ENSMUSG( | 0.895963 | 3.67E-05 | YES | NO | NA |
| Fgf2 | ENSMUSG( | ENSMUSG( | 0.240908 | 0.299077 | NO | NO | NA |
| ENSMUSG | ENSMUSG( | ENSMUSG( | 0.300392 | 0.185193 | NO | NO | NA |
| Kazn | ENSMUSG( | ENSMUSG( | 0.679096 | 0.433549 | NO | NO | NA |
| Hopxos | ENSMUSG( | ENSMUSG( | 0.124236 | 0.519025 | NO | NO | NA |
| Hcn2 | ENSMUSG( | ENSMUSG( | 0.097983 | 0.640013 | NO | NO | NA |
| A730017L: | ENSMUSG( | ENSMUSG( | 0.097906 | 0.617169 | NO | NO | NA |
| Gm37169 | ENSMUSG( | ENSMUSG( | -0.15847 | 0.537526 | NO | NO | NA |
| Tex15 | ENSMUSG( | ENSMUSG( | 0.027822 | 0.894092 | NO | NO | NA |
| Trat1 | ENSMUSG( | ENSMUSG( | 0.239659 | 0.472671 | NO | NO | NA |
| 9130230N | ENSMUSG( | ENSMUSG( | 0.051629 | 0.860871 | NO | NO | NA |
| Rpgr | ENSMUSG( | ENSMUSG( | 0.121055 | 0.642983 | NO | NO | NA |
| Zfp882 | ENSMUSG( | ENSMUSG( | -0.59451 | 0.005665 | YES | NO | NA |
| Plxna3 | ENSMUSG( | ENSMUSG( | 0.997387 | 1.34E-05 | YES | NO | NA |
| Bckdhb | ENSMUSG( | ENSMUSG( | 0.719913 | 0.000571 | YES | NO | NA |
| C130013H | ENSMUSG( | ENSMUSG( | 0.216935 | 0.314547 | NO | NO | NA |
| Gm26759 | ENSMUSG( | ENSMUSG( | -0.02508 | 0.911399 | NO | NO | NA |
| Dnajb14 | ENSMUSG( | ENSMUSG( | -0.46114 | 0.102696 | NO | NO | NA |
| Gm37985 | ENSMUSG( | ENSMUSG( | 0.557314 | 0.64643 | NO | NO | NA |
| Samd14 | ENSMUSG( | ENSMUSG( | 1.088582 | 0.017974 | YES | YES | Upregulated |
| Zfp772 | ENSMUSG( | ENSMUSG( | 0.032841 | 0.841559 | NO | NO | NA |
| Adra2b | ENSMUSG( | ENSMUSG( | 0.71227 | 0.399941 | NO | NO | NA |
| Zfp991 | ENSMUSG( | ENSMUSG( | -1.10159 | 1.35E-05 | YES | YES | Downregulated |
| Lad1 | ENSMUSG( | ENSMUSG( | 2.733471 | 1.14E-07 | YES | YES | Upregulated |
| Zfp938 | ENSMUSG( | ENSMUSG( | -0.21933 | 0.194359 | NO | NO | NA |
| Slc24a5 | ENSMUSG( | ENSMUSG( | 0.490512 | 0.022003 | YES | NO | NA |
| Dcxr | ENSMUSG( | ENSMUSG( | 0.005364 | 0.978798 | NO | NO | NA |
| Gm3095 | ENSMUSG( | ENSMUSG( | -0.21606 | 0.435788 | NO | NO | NA |

| | | | | | | | |
|---|---|---|---|---|---|---|---|
| Eif4a-ps4 | ENSMUSG( | ENSMUSG( | 0.510699 | 0.010823 | YES | NO | NA |
| Gm10125 | ENSMUSG( | ENSMUSG( | 0.482273 | 0.501256 | NO | NO | NA |
| Gm12319 | ENSMUSG( | ENSMUSG( | 0.012098 | 0.956656 | NO | NO | NA |
| Gm43379 | ENSMUSG( | ENSMUSG( | 0.339706 | 0.214549 | NO | NO | NA |
| ENSMUSG( | ENSMUSG( | ENSMUSG( | 0.345557 | 0.341767 | NO | NO | NA |
| Gm14410 | ENSMUSG( | ENSMUSG( | -0.12262 | 0.687659 | NO | NO | NA |
| Gm11847 | ENSMUSG( | ENSMUSG( | 0.240788 | 0.309567 | NO | NO | NA |
| Nfam1 | ENSMUSG( | ENSMUSG( | 0.515268 | 0.563166 | NO | NO | NA |
| Gm5141 | ENSMUSG( | ENSMUSG( | -0.38561 | 0.038022 | YES | NO | NA |
| Atg10 | ENSMUSG( | ENSMUSG( | 0.261791 | 0.096212 | NO | NO | NA |
| ENSMUSG | ENSMUSG( | ENSMUSG( | -0.68216 | 1.11E-06 | YES | NO | NA |
| Slc8a1 | ENSMUSG( | ENSMUSG( | 0.662735 | 0.475267 | NO | NO | NA |
| Enpp2 | ENSMUSG( | ENSMUSG( | 0.414458 | 0.34303 | NO | NO | NA |
| Gm15675 | ENSMUSG( | ENSMUSG( | -1.16041 | 3.10E-06 | YES | YES | Downregulated |
| Bbs2 | ENSMUSG( | ENSMUSG( | 0.610419 | 0.060778 | NO | NO | NA |
| Gm43474 | ENSMUSG( | ENSMUSG( | -0.31191 | 0.184306 | NO | NO | NA |
| Tcf7l2 | ENSMUSG( | ENSMUSG( | 0.537359 | 0.502908 | NO | NO | NA |
| Msc | ENSMUSG( | ENSMUSG( | 1.372601 | 0.001241 | YES | YES | Upregulated |
| Hspb11 | ENSMUSG( | ENSMUSG( | -0.48806 | 0.001902 | YES | NO | NA |
| 1700048O | ENSMUSG( | ENSMUSG( | 1.238122 | 1.65E-07 | YES | YES | Upregulated |
| Ptprd | ENSMUSG( | ENSMUSG( | 0.627744 | 0.619035 | NO | NO | NA |
| Dusp16 | ENSMUSG( | ENSMUSG( | 0.580801 | 0.001626 | YES | NO | NA |
| Acat3 | ENSMUSG( | ENSMUSG( | 0.786164 | 0.079105 | NO | NO | NA |
| 1190007I0 | ENSMUSG( | ENSMUSG( | 0.205465 | 0.318298 | NO | NO | NA |
| Afg1l | ENSMUSG( | ENSMUSG( | -0.61208 | 7.63E-05 | YES | NO | NA |
| Igkv14-111 | ENSMUSG( | ENSMUSG( | -0.76025 | 0.702778 | NO | NO | NA |
| Tfpi | ENSMUSG( | ENSMUSG( | 0.338434 | 0.477503 | NO | NO | NA |
| Cdkl2 | ENSMUSG( | ENSMUSG( | -0.05241 | 0.848306 | NO | NO | NA |
| Gm38357 | ENSMUSG( | ENSMUSG( | -0.89989 | 5.13E-05 | YES | NO | NA |
| Gm47640 | ENSMUSG( | ENSMUSG( | 0.4807 | 0.01018 | YES | NO | NA |
| C130026I2 | ENSMUSG( | ENSMUSG( | 0.927664 | 0.000293 | YES | NO | NA |
| Onecut2 | ENSMUSG( | ENSMUSG( | 0.787636 | 0.288274 | NO | NO | NA |
| Rasa4 | ENSMUSG( | ENSMUSG( | 0.404548 | 0.06657 | NO | NO | NA |
| Scn8a | ENSMUSG( | ENSMUSG( | 0.606722 | 0.482306 | NO | NO | NA |
| Fgfbp3 | ENSMUSG( | ENSMUSG( | -0.06743 | 0.875932 | NO | NO | NA |
| Zbtb12 | ENSMUSG( | ENSMUSG( | 0.411544 | 0.028558 | YES | NO | NA |
| Pxmp2 | ENSMUSG( | ENSMUSG( | 0.053323 | 0.75148 | NO | NO | NA |
| Drp2 | ENSMUSG( | ENSMUSG( | 0.634848 | 0.006874 | YES | NO | NA |
| Cadm1 | ENSMUSG( | ENSMUSG( | 0.490345 | 0.471485 | NO | NO | NA |
| 5_8S_rRN/ | ENSMUSG( | ENSMUSG( | 1.288892 | 0.086255 | NO | NO | NA |
| Stxbp1 | ENSMUSG( | ENSMUSG( | 0.695981 | 0.001459 | YES | NO | NA |
| Apbb2 | ENSMUSG( | ENSMUSG( | 0.74108 | 0.164329 | NO | NO | NA |
| Gm10451 | ENSMUSG( | ENSMUSG( | -1.04608 | 5.21E-05 | YES | YES | Downregulated |
| Zfp773 | ENSMUSG( | ENSMUSG( | -0.39506 | 0.150073 | NO | NO | NA |
| Raver2 | ENSMUSG( | ENSMUSG( | 0.935296 | 0.001214 | YES | NO | NA |
| Cuedc1 | ENSMUSG( | ENSMUSG( | -0.31422 | 0.111065 | NO | NO | NA |
| Alg14 | ENSMUSG( | ENSMUSG( | -0.07144 | 0.673953 | NO | NO | NA |

| | | | | | | | |
|---|---|---|---|---|---|---|---|
| Gm49388 | ENSMUSG( | ENSMUSG( | -0.29641 | 0.443659 | NO | NO | NA |
| Rnf24 | ENSMUSG( | ENSMUSG( | 0.147876 | 0.584443 | NO | NO | NA |
| Dnah11 | ENSMUSG( | ENSMUSG( | -0.73338 | 0.005959 | YES | NO | NA |
| Lmntd2 | ENSMUSG( | ENSMUSG( | -0.58216 | 0.000332 | YES | NO | NA |
| Cracdl | ENSMUSG( | ENSMUSG( | 0.340508 | 0.133854 | NO | NO | NA |
| H2ac8 | ENSMUSG( | ENSMUSG( | 0.328897 | 0.102238 | NO | NO | NA |
| 5_8S_rRN/ | ENSMUSG( | ENSMUSG( | 0.956414 | 0.12658 | NO | NO | NA |
| Mmachc | ENSMUSG( | ENSMUSG( | 0.569737 | 0.001672 | YES | NO | NA |
| Lcn4 | ENSMUSG( | ENSMUSG( | 0.499274 | 0.070215 | NO | NO | NA |
| Zfp358 | ENSMUSG( | ENSMUSG( | -0.06859 | 0.792835 | NO | NO | NA |
| Adat1 | ENSMUSG( | ENSMUSG( | -0.08657 | 0.672778 | NO | NO | NA |
| St3gal5 | ENSMUSG( | ENSMUSG( | 0.350527 | 0.231083 | NO | NO | NA |
| Olfr1328 | ENSMUSG( | ENSMUSG( | 1.269074 | 0.205123 | NO | NO | NA |
| Eef1a1-ps: | ENSMUSG( | ENSMUSG( | 0.278891 | 0.127786 | NO | NO | NA |
| Gm45222 | ENSMUSG( | ENSMUSG( | -0.26966 | 0.110475 | NO | NO | NA |
| Mro | ENSMUSG( | ENSMUSG( | 0.728322 | 0.443203 | NO | NO | NA |
| ENSMUSG | ENSMUSG( | ENSMUSG( | 0.590048 | 0.006529 | YES | NO | NA |
| 2700038G | ENSMUSG( | ENSMUSG( | 0.245246 | 0.150945 | NO | NO | NA |
| Rbfox3 | ENSMUSG( | ENSMUSG( | 0.465144 | 0.567472 | NO | NO | NA |
| Tgfb3 | ENSMUSG( | ENSMUSG( | 0.60388 | 0.13609 | NO | NO | NA |
| Zfp59 | ENSMUSG( | ENSMUSG( | -0.14135 | 0.459146 | NO | NO | NA |
| Ryr3 | ENSMUSG( | ENSMUSG( | -0.06156 | 0.781691 | NO | NO | NA |
| Fign | ENSMUSG( | ENSMUSG( | 0.665646 | 0.508487 | NO | NO | NA |
| Gm6277 | ENSMUSG( | ENSMUSG( | 0.853153 | 0.013941 | YES | NO | NA |
| Alkbh2 | ENSMUSG( | ENSMUSG( | -0.32842 | 0.035714 | YES | NO | NA |
| Tnfrsf11a | ENSMUSG( | ENSMUSG( | 0.857826 | 0.424153 | NO | NO | NA |
| Nhsl2 | ENSMUSG( | ENSMUSG( | 0.314713 | 0.420117 | NO | NO | NA |
| Gm43328 | ENSMUSG( | ENSMUSG( | -0.35069 | 0.049537 | YES | NO | NA |
| 2410022M | ENSMUSG( | ENSMUSG( | 0.766021 | 0.001932 | YES | NO | NA |
| Ahr | ENSMUSG( | ENSMUSG( | 0.016889 | 0.955707 | NO | NO | NA |
| Kcnb1 | ENSMUSG( | ENSMUSG( | 0.293892 | 0.46303 | NO | NO | NA |
| H2bc23 | ENSMUSG( | ENSMUSG( | 0.685888 | 0.115469 | NO | NO | NA |
| Creld1 | ENSMUSG( | ENSMUSG( | -0.13416 | 0.601079 | NO | NO | NA |
| Lrrc56 | ENSMUSG( | ENSMUSG( | 0.339679 | 0.15317 | NO | NO | NA |
| E130311K: | ENSMUSG( | ENSMUSG( | 0.203019 | 0.428872 | NO | NO | NA |
| Rpp40 | ENSMUSG( | ENSMUSG( | 0.868962 | 1.14E-05 | YES | NO | NA |
| Mfsd2b | ENSMUSG( | ENSMUSG( | 0.689151 | 0.071089 | NO | NO | NA |
| Gm45223 | ENSMUSG( | ENSMUSG( | -0.08549 | 0.839726 | NO | NO | NA |
| Gm34794 | ENSMUSG( | ENSMUSG( | -0.63731 | 0.001371 | YES | NO | NA |
| Gm2026 | ENSMUSG( | ENSMUSG( | -0.05897 | 0.906059 | NO | NO | NA |
| Lamc2 | ENSMUSG( | ENSMUSG( | 0.980589 | 0.000148 | YES | NO | NA |
| Gm10778 | ENSMUSG( | ENSMUSG( | -0.26024 | 0.481807 | NO | NO | NA |
| H4c18 | ENSMUSG( | ENSMUSG( | 0.407377 | 0.406887 | NO | NO | NA |
| Gm44505 | ENSMUSG( | ENSMUSG( | 0.205348 | 0.899465 | NO | NO | NA |
| Trav12n-2 | ENSMUSG( | ENSMUSG( | 0.846564 | 0.741426 | NO | NO | NA |
| Xylb | ENSMUSG( | ENSMUSG( | 0.754639 | 0.000124 | YES | NO | NA |
| E230034D | ENSMUSG( | ENSMUSG( | 0.117663 | 0.607114 | NO | NO | NA |

| | | | | | | | |
|---|---|---|---|---|---|---|---|
| Tsc22d1 | ENSMUSG( | ENSMUSG( | -0.06991 | 0.720662 | NO | NO | NA |
| Tmsb15l | ENSMUSG( | ENSMUSG( | -0.21723 | 0.424448 | NO | NO | NA |
| Gm45464 | ENSMUSG( | ENSMUSG( | -0.19902 | 0.2395 | NO | NO | NA |
| Styx-ps | ENSMUSG( | ENSMUSG( | 0.056924 | 0.853406 | NO | NO | NA |
| Cbfa2t3 | ENSMUSG( | ENSMUSG( | 0.300145 | 0.536182 | NO | NO | NA |
| Sulf2 | ENSMUSG( | ENSMUSG( | 2.182404 | 1.13E-05 | YES | YES | Upregulated |
| Trim13 | ENSMUSG( | ENSMUSG( | -0.07491 | 0.70225 | NO | NO | NA |
| Map4k5 | ENSMUSG( | ENSMUSG( | 0.293249 | 0.22942 | NO | NO | NA |
| Gm49794 | ENSMUSG( | ENSMUSG( | 0.752198 | 0.127413 | NO | NO | NA |
| ENSMUSG | ENSMUSG( | ENSMUSG( | -0.58287 | 0.009073 | YES | NO | NA |
| Stxbp4 | ENSMUSG( | ENSMUSG( | -0.00307 | 0.989102 | NO | NO | NA |
| Gypa | ENSMUSG( | ENSMUSG( | 0.635812 | 0.254949 | NO | NO | NA |
| 2310015A | ENSMUSG( | ENSMUSG( | 0.048606 | 0.857932 | NO | NO | NA |
| H2ac10 | ENSMUSG( | ENSMUSG( | 0.597757 | 0.478075 | NO | NO | NA |
| H2bc3 | ENSMUSG( | ENSMUSG( | 0.812304 | 0.00029 | YES | NO | NA |
| Prox2 | ENSMUSG( | ENSMUSG( | 0.46968 | 0.000264 | YES | NO | NA |
| 4933440M | ENSMUSG( | ENSMUSG( | -1.84239 | 0.000117 | YES | YES | Downregulated |
| Slc1a2 | ENSMUSG( | ENSMUSG( | 0.537026 | 0.4913 | NO | NO | NA |
| Gm10382 | ENSMUSG( | ENSMUSG( | -0.15778 | 0.545164 | NO | NO | NA |
| Rpl28-ps1 | ENSMUSG( | ENSMUSG( | 0.718438 | 0.00338 | YES | NO | NA |
| Gm9794 | ENSMUSG( | ENSMUSG( | 0.433774 | 0.008144 | YES | NO | NA |
| Shpk | ENSMUSG( | ENSMUSG( | -0.26108 | 0.147357 | NO | NO | NA |
| Tmem42 | ENSMUSG( | ENSMUSG( | 0.116168 | 0.466606 | NO | NO | NA |
| Gpr19 | ENSMUSG( | ENSMUSG( | -0.16651 | 0.2874 | NO | NO | NA |
| Gm6225 | ENSMUSG( | ENSMUSG( | 0.281166 | 0.431957 | NO | NO | NA |
| Gm14288 | ENSMUSG( | ENSMUSG( | 0.223676 | 0.80634 | NO | NO | NA |
| Susd1 | ENSMUSG( | ENSMUSG( | -0.16005 | 0.370688 | NO | NO | NA |
| Gucy2e | ENSMUSG( | ENSMUSG( | 2.670842 | 1.71E-19 | YES | YES | Upregulated |
| ENSMUSG | ENSMUSG( | ENSMUSG( | 0.275965 | 0.130018 | NO | NO | NA |
| Pex10 | ENSMUSG( | ENSMUSG( | -0.06597 | 0.740948 | NO | NO | NA |
| Adarb1 | ENSMUSG( | ENSMUSG( | 0.652582 | 0.044901 | YES | NO | NA |
| Mid2 | ENSMUSG( | ENSMUSG( | 2.222203 | 1.79E-06 | YES | YES | Upregulated |
| Pabpc1l | ENSMUSG( | ENSMUSG( | -0.42554 | 0.01826 | YES | NO | NA |
| Gm37893 | ENSMUSG( | ENSMUSG( | -0.36775 | 0.0815 | NO | NO | NA |
| Gcat ENSM | ENSMUSG( | ENSMUSG( | 0.81291 | 0.612079 | NO | NO | NA |
| Pigw | ENSMUSG( | ENSMUSG( | 0.085631 | 0.681348 | NO | NO | NA |
| Sh3bp5 | ENSMUSG( | ENSMUSG( | -0.021 | 0.914446 | NO | NO | NA |
| Gpr137b | ENSMUSG( | ENSMUSG( | -0.69158 | 0.001373 | YES | NO | NA |
| Tbc1d19 | ENSMUSG( | ENSMUSG( | 1.282391 | 8.07E-05 | YES | YES | Upregulated |
| Bnip3 | ENSMUSG( | ENSMUSG( | 0.798115 | 0.10268 | NO | NO | NA |
| Gm28935 | ENSMUSG( | ENSMUSG( | 0.022668 | 0.921962 | NO | NO | NA |
| Gm10505 | ENSMUSG( | ENSMUSG( | -0.16159 | 0.444552 | NO | NO | NA |
| Gzmf | ENSMUSG( | ENSMUSG( | 5.14629 | 1.78E-12 | YES | YES | Upregulated |
| Rph3al | ENSMUSG( | ENSMUSG( | -0.58064 | 0.000881 | YES | NO | NA |
| Hdnr | ENSMUSG( | ENSMUSG( | 0.959248 | 0.354681 | NO | NO | NA |
| Gm37206 | ENSMUSG( | ENSMUSG( | -0.2546 | 0.355589 | NO | NO | NA |
| Zfp11 | ENSMUSG( | ENSMUSG( | -0.06522 | 0.747571 | NO | NO | NA |

| | | | | | | | |
|---|---|---|---|---|---|---|---|
| Apoe | ENSMUSG( | ENSMUSG( | 0.264506 | 0.221304 | NO | NO | NA |
| Tppp3 | ENSMUSG( | ENSMUSG( | -0.36712 | 0.204887 | NO | NO | NA |
| Vangl1 | ENSMUSG( | ENSMUSG( | 0.797252 | 0.199413 | NO | NO | NA |
| 2500004C( | ENSMUSG( | ENSMUSG( | -0.19336 | 0.467463 | NO | NO | NA |
| Gm20517 | ENSMUSG( | ENSMUSG( | 0.620568 | 0.259399 | NO | NO | NA |
| Nme6 | ENSMUSG( | ENSMUSG( | 0.498474 | 0.009279 | YES | NO | NA |
| Cacna1a | ENSMUSG( | ENSMUSG( | 0.342592 | 0.065617 | NO | NO | NA |
| Dip2c | ENSMUSG( | ENSMUSG( | 1.135677 | 6.08E-05 | YES | YES | Upregulated |
| Dock6 | ENSMUSG( | ENSMUSG( | 0.608532 | 0.035433 | YES | NO | NA |
| Slc25a26 | ENSMUSG( | ENSMUSG( | 0.454425 | 0.00153 | YES | NO | NA |
| Galnt17 | ENSMUSG( | ENSMUSG( | 0.577651 | 0.574311 | NO | NO | NA |
| Ccdc116 | ENSMUSG( | ENSMUSG( | 0.821376 | 2.83E-05 | YES | NO | NA |
| Gm6560 | ENSMUSG( | ENSMUSG( | 0.558985 | 0.015839 | YES | NO | NA |
| Gm20544 | ENSMUSG( | ENSMUSG( | 0.303658 | 0.450269 | NO | NO | NA |
| Cerox1 | ENSMUSG( | ENSMUSG( | -0.70933 | 0.000123 | YES | NO | NA |
| Gm17017 | ENSMUSG( | ENSMUSG( | -0.15638 | 0.449568 | NO | NO | NA |
| Atat1 | ENSMUSG( | ENSMUSG( | -0.52646 | 0.000881 | YES | NO | NA |
| C030034I2 | ENSMUSG( | ENSMUSG( | -0.04649 | 0.808364 | NO | NO | NA |
| Gpnmb | ENSMUSG( | ENSMUSG( | 0.873751 | 1.22E-05 | YES | NO | NA |
| Gm10335 | ENSMUSG( | ENSMUSG( | 0.099817 | 0.771858 | NO | NO | NA |
| Rhd | ENSMUSG( | ENSMUSG( | 0.746431 | 0.280628 | NO | NO | NA |
| Prtg | ENSMUSG( | ENSMUSG( | 0.86608 | 0.39039 | NO | NO | NA |
| Atp1a2 | ENSMUSG( | ENSMUSG( | 0.763728 | 0.033158 | YES | NO | NA |
| Gm48604 | ENSMUSG( | ENSMUSG( | -0.1962 | 0.451447 | NO | NO | NA |
| Gm16364 | ENSMUSG( | ENSMUSG( | -0.79294 | 5.67E-05 | YES | NO | NA |
| Nrg4 | ENSMUSG( | ENSMUSG( | -0.0831 | 0.655576 | NO | NO | NA |
| Actn2 | ENSMUSG( | ENSMUSG( | -1.11458 | 1.70E-05 | YES | YES | Downregulated |
| Slc1a3 | ENSMUSG( | ENSMUSG( | 1.154022 | 0.234753 | NO | NO | NA |
| Spsb2 | ENSMUSG( | ENSMUSG( | -0.65158 | 0.001359 | YES | NO | NA |
| Agbl3 | ENSMUSG( | ENSMUSG( | 0.842062 | 0.02221 | YES | NO | NA |
| Sdk1 | ENSMUSG( | ENSMUSG( | 0.766143 | 0.049655 | YES | NO | NA |
| Flt1 | ENSMUSG( | ENSMUSG( | 0.216366 | 0.584546 | NO | NO | NA |
| Gm19412 | ENSMUSG( | ENSMUSG( | 0.076515 | 0.779065 | NO | NO | NA |
| Sccpdh | ENSMUSG( | ENSMUSG( | 0.710717 | 0.003503 | YES | NO | NA |
| Sdr42e1 | ENSMUSG( | ENSMUSG( | 0.13258 | 0.656122 | NO | NO | NA |
| Gngt2 | ENSMUSG( | ENSMUSG( | -0.64711 | 0.004886 | YES | NO | NA |
| Ctnnd2 | ENSMUSG( | ENSMUSG( | 0.72249 | 0.153498 | NO | NO | NA |
| Nfix | ENSMUSG( | ENSMUSG( | 0.29467 | 0.406718 | NO | NO | NA |
| Tomt | ENSMUSG( | ENSMUSG( | 0.678158 | 0.014511 | YES | NO | NA |
| Plekha8 | ENSMUSG( | ENSMUSG( | 0.29625 | 0.05601 | NO | NO | NA |
| Kctd11 | ENSMUSG( | ENSMUSG( | -0.2769 | 0.239483 | NO | NO | NA |
| Gm43813 | ENSMUSG( | ENSMUSG( | 0.354719 | 0.508226 | NO | NO | NA |
| Gm44597 | ENSMUSG( | ENSMUSG( | -0.10016 | 0.678687 | NO | NO | NA |
| Bmp1 | ENSMUSG( | ENSMUSG( | 0.47525 | 0.56202 | NO | NO | NA |
| Bcl9 | ENSMUSG( | ENSMUSG( | 0.474026 | 0.177666 | NO | NO | NA |
| Ccdc126 | ENSMUSG( | ENSMUSG( | 0.069841 | 0.794266 | NO | NO | NA |
| Rps12-ps3 | ENSMUSG( | ENSMUSG( | 0.88312 | 1.10E-06 | YES | NO | NA |

| | | | | | | | |
|---|---|---|---|---|---|---|---|
| Trav4n-4 | ENSMUSG( | ENSMUSG( | -1.09547 | 0.756944 | NO | NO | NA |
| 5430403G | ENSMUSG( | ENSMUSG( | -0.12996 | 0.627562 | NO | NO | NA |
| Gm20743 | ENSMUSG( | ENSMUSG( | 2.167379 | 6.61E-07 | YES | YES | Upregulated |
| Hic2 | ENSMUSG( | ENSMUSG( | 0.299258 | 0.083139 | NO | NO | NA |
| Cnnm2 | ENSMUSG( | ENSMUSG( | -0.20506 | 0.374444 | NO | NO | NA |
| 3110009E: | ENSMUSG( | ENSMUSG( | 0.111575 | 0.601079 | NO | NO | NA |
| Tm7sf2 | ENSMUSG( | ENSMUSG( | 0.058937 | 0.723449 | NO | NO | NA |
| E230001N | ENSMUSG( | ENSMUSG( | 0.317587 | 0.128446 | NO | NO | NA |
| Nbea | ENSMUSG( | ENSMUSG( | 0.763503 | 0.072352 | NO | NO | NA |
| ENSMUSG | ENSMUSG( | ENSMUSG( | 0.472428 | 0.100901 | NO | NO | NA |
| Gm11613 | ENSMUSG( | ENSMUSG( | -0.10837 | 0.674118 | NO | NO | NA |
| Iqcb1 | ENSMUSG( | ENSMUSG( | 0.188896 | 0.384148 | NO | NO | NA |
| Adcy6 | ENSMUSG( | ENSMUSG( | 1.127571 | 0.012404 | YES | YES | Upregulated |
| Gm16091 | ENSMUSG( | ENSMUSG( | -0.0914 | 0.751088 | NO | NO | NA |
| Nectin1 | ENSMUSG( | ENSMUSG( | 0.369195 | 0.620616 | NO | NO | NA |
| Lncpint | ENSMUSG( | ENSMUSG( | -0.08071 | 0.768848 | NO | NO | NA |
| Fbxw9 | ENSMUSG( | ENSMUSG( | 0.537936 | 0.03917 | YES | NO | NA |
| 1700056E: | ENSMUSG( | ENSMUSG( | 0.490795 | 0.006903 | YES | NO | NA |
| Foxd2 | ENSMUSG( | ENSMUSG( | -0.41252 | 0.011883 | YES | NO | NA |
| Vash1 | ENSMUSG( | ENSMUSG( | 0.217585 | 0.29931 | NO | NO | NA |
| Gli3 | ENSMUSG( | ENSMUSG( | 0.92617 | 0.386947 | NO | NO | NA |
| Muc20 | ENSMUSG( | ENSMUSG( | -0.93167 | 0.000254 | YES | NO | NA |
| Gstm4 | ENSMUSG( | ENSMUSG( | 0.490539 | 0.019613 | YES | NO | NA |
| Xlr | ENSMUSG( | ENSMUSG( | 1.637743 | 0.025759 | YES | YES | Upregulated |
| Rab32 | ENSMUSG( | ENSMUSG( | 0.300997 | 0.260999 | NO | NO | NA |
| Uchl1 | ENSMUSG( | ENSMUSG( | 0.524685 | 0.1755 | NO | NO | NA |
| Nt5e | ENSMUSG( | ENSMUSG( | 0.22412 | 0.498883 | NO | NO | NA |
| Epb41l3 | ENSMUSG( | ENSMUSG( | 0.216746 | 0.74342 | NO | NO | NA |
| Arhgef11 | ENSMUSG( | ENSMUSG( | 0.211421 | 0.220888 | NO | NO | NA |
| Gm38248 | ENSMUSG( | ENSMUSG( | 0.033878 | 0.869914 | NO | NO | NA |
| Trgc4 | ENSMUSG( | ENSMUSG( | -1.00034 | 0.075796 | NO | NO | NA |
| Gm1043 | ENSMUSG( | ENSMUSG( | -0.386 | 0.017957 | YES | NO | NA |
| Cspg5 | ENSMUSG( | ENSMUSG( | 0.245034 | 0.73647 | NO | NO | NA |
| Mterf1b | ENSMUSG( | ENSMUSG( | -0.40782 | 0.042634 | YES | NO | NA |
| Bcl6 | ENSMUSG( | ENSMUSG( | -0.10808 | 0.745508 | NO | NO | NA |
| 4632427E: | ENSMUSG( | ENSMUSG( | 0.222132 | 0.515606 | NO | NO | NA |
| Arhgef25 | ENSMUSG( | ENSMUSG( | 0.257711 | 0.428834 | NO | NO | NA |
| Tigd3 | ENSMUSG( | ENSMUSG( | 0.193631 | 0.371398 | NO | NO | NA |
| Fgl1 | ENSMUSG( | ENSMUSG( | 0.791421 | 0.449709 | NO | NO | NA |
| Gm14325 | ENSMUSG( | ENSMUSG( | 0.069819 | 0.74345 | NO | NO | NA |
| Gm42939 | ENSMUSG( | ENSMUSG( | 1.036647 | 0.315494 | NO | NO | NA |
| Tlr12 | ENSMUSG( | ENSMUSG( | -0.31744 | 0.265275 | NO | NO | NA |
| Gm35019 | ENSMUSG( | ENSMUSG( | 0.120999 | 0.679453 | NO | NO | NA |
| Zfp566 | ENSMUSG( | ENSMUSG( | -0.12748 | 0.431435 | NO | NO | NA |
| ENSMUSG( | ENSMUSG( | ENSMUSG( | 0.04404 | 0.866454 | NO | NO | NA |
| Rcor2 | ENSMUSG( | ENSMUSG( | -1.23331 | 3.51E-05 | YES | YES | Downregulated |
| Tmem171 | ENSMUSG( | ENSMUSG( | -0.00559 | 0.985797 | NO | NO | NA |

| | | | | | | | |
|---|---|---|---|---|---|---|---|
| Niban3 | ENSMUSG( | ENSMUSG( | -0.45227 | 0.017134 | YES | NO | NA |
| Smad1 | ENSMUSG( | ENSMUSG( | 0.156104 | 0.391447 | NO | NO | NA |
| Usp29 | ENSMUSG( | ENSMUSG( | 0.677974 | 0.509593 | NO | NO | NA |
| Svip | ENSMUSG( | ENSMUSG( | 1.048856 | 0.000289 | YES | YES | Upregulated |
| Pet117 | ENSMUSG( | ENSMUSG( | -0.26699 | 0.871067 | NO | NO | NA |
| Msrb2 | ENSMUSG( | ENSMUSG( | -0.248 | 0.22757 | NO | NO | NA |
| Srsf12 | ENSMUSG( | ENSMUSG( | 0.084303 | 0.750821 | NO | NO | NA |
| Mcts2 | ENSMUSG( | ENSMUSG( | -0.10177 | 0.697751 | NO | NO | NA |
| Endou | ENSMUSG( | ENSMUSG( | 1.590536 | 6.87E-09 | YES | YES | Upregulated |
| A830035O | ENSMUSG( | ENSMUSG( | -0.44263 | 0.045988 | YES | NO | NA |
| Eya2 | ENSMUSG( | ENSMUSG( | 1.405608 | 4.49E-05 | YES | YES | Upregulated |
| Tmem198l | ENSMUSG( | ENSMUSG( | 0.040841 | 0.86261 | NO | NO | NA |
| ENSMUSG | ENSMUSG( | ENSMUSG( | -0.12075 | 0.538693 | NO | NO | NA |
| Csf1r | ENSMUSG( | ENSMUSG( | -0.55831 | 0.075002 | NO | NO | NA |
| Glis1 | ENSMUSG( | ENSMUSG( | 1.280335 | 8.13E-10 | YES | YES | Upregulated |
| Dstyk | ENSMUSG( | ENSMUSG( | 0.571023 | 0.002951 | YES | NO | NA |
| Rtkn2 | ENSMUSG( | ENSMUSG( | 0.237755 | 0.417158 | NO | NO | NA |
| ENSMUSG( | ENSMUSG( | ENSMUSG( | 0.183113 | 0.366285 | NO | NO | NA |
| Lipo3 | ENSMUSG( | ENSMUSG( | -0.39221 | 0.083025 | NO | NO | NA |
| Asph | ENSMUSG( | ENSMUSG( | 1.13327 | 0.000539 | YES | YES | Upregulated |
| Ugt1a6a | ENSMUSG( | ENSMUSG( | 0.239299 | 0.746599 | NO | NO | NA |
| Gm43634 | ENSMUSG( | ENSMUSG( | -0.47279 | 0.087671 | NO | NO | NA |
| Khdc3 | ENSMUSG( | ENSMUSG( | 0.17426 | 0.551005 | NO | NO | NA |
| Gm26586 | ENSMUSG( | ENSMUSG( | -0.05934 | 0.711913 | NO | NO | NA |
| Rpl23a-ps3 | ENSMUSG( | ENSMUSG( | 0.318677 | 0.319978 | NO | NO | NA |
| Pkd2l2 | ENSMUSG( | ENSMUSG( | -0.2118 | 0.348614 | NO | NO | NA |
| Nr6a1 | ENSMUSG( | ENSMUSG( | 0.498193 | 0.342573 | NO | NO | NA |
| Gm44777 | ENSMUSG( | ENSMUSG( | -0.02846 | 0.911471 | NO | NO | NA |
| Gm20402 | ENSMUSG( | ENSMUSG( | 0.038981 | 0.914861 | NO | NO | NA |
| AA414768 | ENSMUSG( | ENSMUSG( | -0.12984 | 0.491449 | NO | NO | NA |
| P2ry14 | ENSMUSG( | ENSMUSG( | 0.931268 | 0.002623 | YES | NO | NA |
| Ighv1-55 | ENSMUSG( | ENSMUSG( | -5.04801 | 0.00161 | YES | YES | Downregulated |
| Gm5537 | ENSMUSG( | ENSMUSG( | 0.29979 | 0.851772 | NO | NO | NA |
| Spata7 | ENSMUSG( | ENSMUSG( | 0.724194 | 0.083888 | NO | NO | NA |
| Zfp946 | ENSMUSG( | ENSMUSG( | 0.153119 | 0.632955 | NO | NO | NA |
| Zfp870 | ENSMUSG( | ENSMUSG( | -0.76233 | 0.000815 | YES | NO | NA |
| ENSMUSG( | ENSMUSG( | ENSMUSG( | 0.603856 | 0.158927 | NO | NO | NA |
| Gm37010 | ENSMUSG( | ENSMUSG( | -0.60414 | 0.00564 | YES | NO | NA |
| Zscan12 | ENSMUSG( | ENSMUSG( | 0.891238 | 6.57E-06 | YES | NO | NA |
| Inka2 | ENSMUSG( | ENSMUSG( | 0.156915 | 0.481807 | NO | NO | NA |
| ENSMUSG | ENSMUSG( | ENSMUSG( | -0.61448 | 0.072544 | NO | NO | NA |
| Grk4 | ENSMUSG( | ENSMUSG( | 0.003399 | 0.987646 | NO | NO | NA |
| Gm36995 | ENSMUSG( | ENSMUSG( | -0.21181 | 0.533073 | NO | NO | NA |
| Gm6136 | ENSMUSG( | ENSMUSG( | 0.259339 | 0.159563 | NO | NO | NA |
| H3c13 | ENSMUSG( | ENSMUSG( | 0.330597 | 0.19541 | NO | NO | NA |
| Ccdc62 | ENSMUSG( | ENSMUSG( | 0.44798 | 0.003552 | YES | NO | NA |
| A630052C | ENSMUSG( | ENSMUSG( | 0.185855 | 0.505568 | NO | NO | NA |

| | | | | | | | |
|---|---|---|---|---|---|---|---|
| Mpeg1 | ENSMUSG( | ENSMUSG( | -1.45813 | 0.000645 | YES | YES | Downregulated |
| Gm26770 | ENSMUSG( | ENSMUSG( | 0.103333 | 0.607026 | NO | NO | NA |
| Zfp784 | ENSMUSG( | ENSMUSG( | 0.131988 | 0.655215 | NO | NO | NA |
| Clcn2 | ENSMUSG( | ENSMUSG( | 0.732401 | 0.01355 | YES | NO | NA |
| Insl3 | ENSMUSG( | ENSMUSG( | 0.781298 | 0.084775 | NO | NO | NA |
| Trav4d-4 | ENSMUSG( | ENSMUSG( | -0.19841 | 0.948515 | NO | NO | NA |
| Tnfrsf21 | ENSMUSG( | ENSMUSG( | 0.213168 | 0.472454 | NO | NO | NA |
| Aldh5a1 | ENSMUSG( | ENSMUSG( | 0.714485 | 0.004431 | YES | NO | NA |
| Hcn1 | ENSMUSG( | ENSMUSG( | 0.752364 | 0.432797 | NO | NO | NA |
| Zfp462 | ENSMUSG( | ENSMUSG( | 0.6477 | 0.476511 | NO | NO | NA |
| Ttbk1 | ENSMUSG( | ENSMUSG( | 0.135372 | 0.83556 | NO | NO | NA |
| AI854703 | ENSMUSG( | ENSMUSG( | 1.248362 | 1.13E-05 | YES | YES | Upregulated |
| Macrod1 | ENSMUSG( | ENSMUSG( | 1.080592 | 0.000487 | YES | YES | Upregulated |
| Ighv1-7 | ENSMUSG( | ENSMUSG( | -5.26528 | NA | NO | NO | NA |
| Arc | ENSMUSG( | ENSMUSG( | -0.19762 | 0.389321 | NO | NO | NA |
| Aldoc | ENSMUSG( | ENSMUSG( | 0.676432 | 0.10947 | NO | NO | NA |
| Trim45 | ENSMUSG( | ENSMUSG( | 0.598357 | 0.006617 | YES | NO | NA |
| Dnase1 | ENSMUSG( | ENSMUSG( | -0.07626 | 0.702792 | NO | NO | NA |
| P2ry12 | ENSMUSG( | ENSMUSG( | 1.217452 | 0.000527 | YES | YES | Upregulated |
| Eno1b | ENSMUSG( | ENSMUSG( | 0.606738 | 0.01825 | YES | NO | NA |
| Gm42669 | ENSMUSG( | ENSMUSG( | -0.55181 | 0.49117 | NO | NO | NA |
| Mtfp1 | ENSMUSG( | ENSMUSG( | 0.683412 | 0.000342 | YES | NO | NA |
| Tbc1d12 | ENSMUSG( | ENSMUSG( | 0.140465 | 0.475161 | NO | NO | NA |
| Card10 | ENSMUSG( | ENSMUSG( | -0.00772 | 0.976023 | NO | NO | NA |
| Gm16754 | ENSMUSG( | ENSMUSG( | -0.32432 | 0.049159 | YES | NO | NA |
| Zfp459 | ENSMUSG( | ENSMUSG( | -0.68989 | 0.001293 | YES | NO | NA |
| Chac2 | ENSMUSG( | ENSMUSG( | 0.00921 | 0.976417 | NO | NO | NA |
| Gm28370 | ENSMUSG( | ENSMUSG( | -0.77322 | 0.000548 | YES | NO | NA |
| 8430429K( | ENSMUSG( | ENSMUSG( | 0.533278 | 0.001622 | YES | NO | NA |
| Gm20667 | ENSMUSG( | ENSMUSG( | -0.15066 | 0.53045 | NO | NO | NA |
| B3gnt3 | ENSMUSG( | ENSMUSG( | -0.04543 | 0.839726 | NO | NO | NA |
| Mettl27 | ENSMUSG( | ENSMUSG( | 0.29975 | 0.259199 | NO | NO | NA |
| Gm35576 | ENSMUSG( | ENSMUSG( | 0.807198 | 0.453331 | NO | NO | NA |
| D930028M | ENSMUSG( | ENSMUSG( | -0.03659 | 0.899457 | NO | NO | NA |
| Ifitm3 | ENSMUSG( | ENSMUSG( | 0.186568 | 0.649152 | NO | NO | NA |
| Mex3a | ENSMUSG( | ENSMUSG( | -0.15176 | 0.506019 | NO | NO | NA |
| Trav4-3 | ENSMUSG( | ENSMUSG( | 0.198264 | 0.746814 | NO | NO | NA |
| C77080 | ENSMUSG( | ENSMUSG( | 0.126742 | 0.63907 | NO | NO | NA |
| Six4 | ENSMUSG( | ENSMUSG( | 0.648535 | 0.539424 | NO | NO | NA |
| Slc39a8 | ENSMUSG( | ENSMUSG( | 1.283066 | 2.00E-05 | YES | YES | Upregulated |
| Mir22hg | ENSMUSG( | ENSMUSG( | -0.1376 | 0.468862 | NO | NO | NA |
| D830025C | ENSMUSG( | ENSMUSG( | 0.342541 | 0.471631 | NO | NO | NA |
| Gm3604 | ENSMUSG( | ENSMUSG( | -0.26876 | 0.162247 | NO | NO | NA |
| Cacna1d | ENSMUSG( | ENSMUSG( | -0.50962 | 0.243059 | NO | NO | NA |
| 5031434O | ENSMUSG( | ENSMUSG( | 0.361686 | 0.226265 | NO | NO | NA |
| Odad1 | ENSMUSG( | ENSMUSG( | 0.332222 | 0.515507 | NO | NO | NA |
| Gm21985 | ENSMUSG( | ENSMUSG( | -2.44352 | 0.522888 | NO | NO | NA |

| | | | | | | | |
|---|---|---|---|---|---|---|---|
| Shank1 | ENSMUSG( | ENSMUSG( | 0.691811 | 0.400572 | NO | NO | NA |
| Elmo3 | ENSMUSG( | ENSMUSG( | -0.38586 | 0.022527 | YES | NO | NA |
| Zfp963 | ENSMUSG( | ENSMUSG( | 0.254332 | 0.254003 | NO | NO | NA |
| Kyat3 | ENSMUSG( | ENSMUSG( | 0.976396 | 9.54E-06 | YES | NO | NA |
| Rorb | ENSMUSG( | ENSMUSG( | 0.778 | 0.215985 | NO | NO | NA |
| Dpysl3 | ENSMUSG( | ENSMUSG( | 0.140923 | 0.64011 | NO | NO | NA |
| B930094E( | ENSMUSG( | ENSMUSG( | 0.399941 | 0.307775 | NO | NO | NA |
| Akr1c13 | ENSMUSG( | ENSMUSG( | -0.21834 | 0.328488 | NO | NO | NA |
| Nedd4l | ENSMUSG( | ENSMUSG( | 0.281695 | 0.469377 | NO | NO | NA |
| Vmn2r130 | ENSMUSG( | ENSMUSG( | 0.731226 | 0.494121 | NO | NO | NA |
| Gm7160 | ENSMUSG( | ENSMUSG( | -0.0335 | 0.89298 | NO | NO | NA |
| Hyal2 | ENSMUSG( | ENSMUSG( | -0.11607 | 0.580206 | NO | NO | NA |
| Zfp965 | ENSMUSG( | ENSMUSG( | -0.04705 | 0.963475 | NO | NO | NA |
| Tmem107 | ENSMUSG( | ENSMUSG( | 0.366867 | 0.069193 | NO | NO | NA |
| Stx1a | ENSMUSG( | ENSMUSG( | 0.425823 | 0.013651 | YES | NO | NA |
| Mfsd4b5 | ENSMUSG( | ENSMUSG( | -1.60534 | 1.69E-05 | YES | YES | Downregulated |
| Gm42727 | ENSMUSG( | ENSMUSG( | -0.31671 | 0.111599 | NO | NO | NA |
| Cnga1 | ENSMUSG( | ENSMUSG( | -0.91784 | 0.000146 | YES | NO | NA |
| App | ENSMUSG( | ENSMUSG( | 0.456774 | 0.123913 | NO | NO | NA |
| Arhgap23 | ENSMUSG( | ENSMUSG( | 0.653456 | 0.259323 | NO | NO | NA |
| Ptgr1 | ENSMUSG( | ENSMUSG( | 0.039001 | 0.892518 | NO | NO | NA |
| Gm43597 | ENSMUSG( | ENSMUSG( | 0.175027 | 0.507817 | NO | NO | NA |
| Mapt | ENSMUSG( | ENSMUSG( | 0.743927 | 0.418294 | NO | NO | NA |
| Ifi44 | ENSMUSG( | ENSMUSG( | -0.90335 | 0.000588 | YES | NO | NA |
| Arvcf | ENSMUSG( | ENSMUSG( | 0.264312 | 0.187802 | NO | NO | NA |
| Per3 | ENSMUSG( | ENSMUSG( | 0.114587 | 0.732456 | NO | NO | NA |
| Cpt1b | ENSMUSG( | ENSMUSG( | 0.491658 | 0.019482 | YES | NO | NA |
| Nenf | ENSMUSG( | ENSMUSG( | 0.095155 | 0.624843 | NO | NO | NA |
| Naprt | ENSMUSG( | ENSMUSG( | -0.47092 | 0.198254 | NO | NO | NA |
| Gm35339 | ENSMUSG( | ENSMUSG( | -0.47314 | 0.185594 | NO | NO | NA |
| Ciita | ENSMUSG( | ENSMUSG( | 0.332566 | 0.58481 | NO | NO | NA |
| ENSMUSG( | ENSMUSG( | ENSMUSG( | 0.023405 | 0.927553 | NO | NO | NA |
| Emilin1 | ENSMUSG( | ENSMUSG( | -0.09041 | 0.681501 | NO | NO | NA |
| ENSMUSG( | ENSMUSG( | ENSMUSG( | 0.091869 | 0.837445 | NO | NO | NA |
| Ubtd1 | ENSMUSG( | ENSMUSG( | -0.91486 | 4.35E-05 | YES | NO | NA |
| Gas2l1 | ENSMUSG( | ENSMUSG( | -0.28547 | 0.214017 | NO | NO | NA |
| Tmprss6 | ENSMUSG( | ENSMUSG( | -0.00684 | 0.986386 | NO | NO | NA |
| Paqr9 | ENSMUSG( | ENSMUSG( | 0.843275 | 0.065015 | NO | NO | NA |
| Ndufaf6 | ENSMUSG( | ENSMUSG( | 0.455092 | 0.024088 | YES | NO | NA |
| D830050J1 | ENSMUSG( | ENSMUSG( | 0.245762 | 0.369818 | NO | NO | NA |
| Map7 | ENSMUSG( | ENSMUSG( | -0.49523 | 0.01379 | YES | NO | NA |
| Efcab7 | ENSMUSG( | ENSMUSG( | 0.259288 | 0.422267 | NO | NO | NA |
| Cd46 | ENSMUSG( | ENSMUSG( | -0.18889 | 0.377938 | NO | NO | NA |
| Scd1 | ENSMUSG( | ENSMUSG( | 1.037137 | 0.051006 | NO | NO | NA |
| Ier5l | ENSMUSG( | ENSMUSG( | 0.176273 | 0.602854 | NO | NO | NA |
| Trav9-1 | ENSMUSG( | ENSMUSG( | 0.214749 | 0.669545 | NO | NO | NA |
| Zfp623 | ENSMUSG( | ENSMUSG( | 0.552073 | 0.000442 | YES | NO | NA |

| | | | | | | | |
|---|---|---|---|---|---|---|---|
| Fam221a | ENSMUSG( | ENSMUSG( | 0.62271 | 0.002118 | YES | NO | NA |
| Gm9844 | ENSMUSG( | ENSMUSG( | 0.411564 | 0.050239 | NO | NO | NA |
| Gm13436 | ENSMUSG( | ENSMUSG( | 0.381832 | 0.236486 | NO | NO | NA |
| Gm37261 | ENSMUSG( | ENSMUSG( | -0.63675 | 0.012503 | YES | NO | NA |
| Pdzk1 | ENSMUSG( | ENSMUSG( | 0.711325 | 0.000322 | YES | NO | NA |
| Fndc3b | ENSMUSG( | ENSMUSG( | 0.042804 | 0.847958 | NO | NO | NA |
| Ryr2 | ENSMUSG( | ENSMUSG( | 0.555569 | 0.571212 | NO | NO | NA |
| Gm15947 | ENSMUSG( | ENSMUSG( | 0.87991 | 0.419644 | NO | NO | NA |
| Gk5 | ENSMUSG( | ENSMUSG( | 0.726823 | 0.001618 | YES | NO | NA |
| Tagln | ENSMUSG( | ENSMUSG( | -0.21863 | 0.374035 | NO | NO | NA |
| A230020J2 | ENSMUSG( | ENSMUSG( | 0.20517 | 0.4007 | NO | NO | NA |
| Rflnb | ENSMUSG( | ENSMUSG( | -0.03659 | 0.89964 | NO | NO | NA |
| Avpi1 | ENSMUSG( | ENSMUSG( | 0.926978 | 0.000542 | YES | NO | NA |
| Rps27a-ps | ENSMUSG( | ENSMUSG( | 0.58303 | 0.017432 | YES | NO | NA |
| Csrp2 | ENSMUSG( | ENSMUSG( | 0.062094 | 0.731835 | NO | NO | NA |
| Il1r2 | ENSMUSG( | ENSMUSG( | 2.153089 | 3.34E-07 | YES | YES | Upregulated |
| Gm49378 | ENSMUSG( | ENSMUSG( | 0.437056 | 0.4904 | NO | NO | NA |
| Gm8066 | ENSMUSG( | ENSMUSG( | -0.12915 | 0.653854 | NO | NO | NA |
| Hdac9 | ENSMUSG( | ENSMUSG( | 0.947083 | 0.274949 | NO | NO | NA |
| Fmo5 | ENSMUSG( | ENSMUSG( | -0.60552 | 0.00478 | YES | NO | NA |
| Apol10b | ENSMUSG( | ENSMUSG( | -0.01184 | 0.967667 | NO | NO | NA |
| Zfp985 | ENSMUSG( | ENSMUSG( | 2.805441 | 6.64E-19 | YES | YES | Upregulated |
| Cfap74 | ENSMUSG( | ENSMUSG( | -5.78511 | 4.37E-22 | YES | YES | Downregulated |
| ENSMUSG | ENSMUSG( | ENSMUSG( | -0.12565 | 0.704289 | NO | NO | NA |
| Trav14n-2 | ENSMUSG( | ENSMUSG( | 0.060043 | 0.978282 | NO | NO | NA |
| Lrrc75a | ENSMUSG( | ENSMUSG( | -0.31326 | 0.076112 | NO | NO | NA |
| H2ac22 | ENSMUSG( | ENSMUSG( | 0.626188 | 0.029779 | YES | NO | NA |
| Trpm3 | ENSMUSG( | ENSMUSG( | 0.700466 | 0.457317 | NO | NO | NA |
| ENSMUSG( | ENSMUSG( | ENSMUSG( | -0.24411 | 0.195346 | NO | NO | NA |
| Fn1 | ENSMUSG( | ENSMUSG( | 0.163944 | 0.78461 | NO | NO | NA |
| Dab2ip | ENSMUSG( | ENSMUSG( | -0.09116 | 0.786071 | NO | NO | NA |
| Gm43654 | ENSMUSG( | ENSMUSG( | 0.108406 | 0.727799 | NO | NO | NA |
| Olfm1 | ENSMUSG( | ENSMUSG( | 0.509164 | 0.147871 | NO | NO | NA |
| Gm43890 | ENSMUSG( | ENSMUSG( | 0.539675 | 0.280628 | NO | NO | NA |
| Stag3 | ENSMUSG( | ENSMUSG( | 0.628289 | 0.071094 | NO | NO | NA |
| 2310058D | ENSMUSG( | ENSMUSG( | -0.50316 | 0.005832 | YES | NO | NA |
| Ighv9-3 | ENSMUSG( | ENSMUSG( | -1.59783 | 0.361227 | NO | NO | NA |
| Ccl27a | ENSMUSG( | ENSMUSG( | 0.094882 | 0.57285 | NO | NO | NA |
| Gm48871 | ENSMUSG( | ENSMUSG( | 1.06736 | 0.301461 | NO | NO | NA |
| Ighv1-59 | ENSMUSG( | ENSMUSG( | 0.061764 | 0.984129 | NO | NO | NA |
| Pard3b | ENSMUSG( | ENSMUSG( | 0.518917 | 0.003442 | YES | NO | NA |
| Gm3550 | ENSMUSG( | ENSMUSG( | 0.505897 | 0.034527 | YES | NO | NA |
| Efna5 | ENSMUSG( | ENSMUSG( | 0.760633 | 0.460605 | NO | NO | NA |
| Insyn2b | ENSMUSG( | ENSMUSG( | -0.85585 | 0.007195 | YES | NO | NA |
| Gm11425 | ENSMUSG( | ENSMUSG( | 0.30496 | 0.182521 | NO | NO | NA |
| Gm16185 | ENSMUSG( | ENSMUSG( | -0.58757 | 0.013718 | YES | NO | NA |
| Vmn2r17 | ENSMUSG( | ENSMUSG( | 0.715299 | 0.513234 | NO | NO | NA |

| | | | | | | | |
|---|---|---|---|---|---|---|---|
| Igkv10-94 | ENSMUSG( | ENSMUSG( | -3.93044 | 0.007565 | YES | YES | Downregulated |
| Chst14 | ENSMUSG( | ENSMUSG( | 0.209947 | 0.420191 | NO | NO | NA |
| Gm10461 | ENSMUSG( | ENSMUSG( | 0.70386 | 0.44012 | NO | NO | NA |
| Nrg1 | ENSMUSG( | ENSMUSG( | 0.615164 | 0.389469 | NO | NO | NA |
| Rad51b | ENSMUSG( | ENSMUSG( | 0.378267 | 0.113903 | NO | NO | NA |
| Gm20939 | ENSMUSG( | ENSMUSG( | -0.02406 | 0.9084 | NO | NO | NA |
| Lrp11 | ENSMUSG( | ENSMUSG( | 0.563351 | 0.097158 | NO | NO | NA |
| ENSMUSG | ENSMUSG( | ENSMUSG( | 0.326551 | 0.111027 | NO | NO | NA |
| Lhx9 | ENSMUSG( | ENSMUSG( | 1.172143 | 0.23457 | NO | NO | NA |
| Dnai4 | ENSMUSG( | ENSMUSG( | -0.25035 | 0.246816 | NO | NO | NA |
| Amz1 | ENSMUSG( | ENSMUSG( | 0.345212 | 0.576362 | NO | NO | NA |
| Pcdh17 | ENSMUSG( | ENSMUSG( | 0.684376 | 0.464038 | NO | NO | NA |
| Rbbp9 | ENSMUSG( | ENSMUSG( | 0.26283 | 0.285586 | NO | NO | NA |
| AA474408 | ENSMUSG( | ENSMUSG( | 0.515867 | 0.276993 | NO | NO | NA |
| ENSMUSG( | ENSMUSG( | ENSMUSG( | -0.33832 | 0.067478 | NO | NO | NA |
| Lif | ENSMUSG( | ENSMUSG( | 0.809035 | 0.139155 | NO | NO | NA |
| Plekhh3 | ENSMUSG( | ENSMUSG( | -1.09167 | 0.002289 | YES | YES | Downregulated |
| Rsph1 | ENSMUSG( | ENSMUSG( | 0.846106 | 8.09E-05 | YES | NO | NA |
| D130017N | ENSMUSG( | ENSMUSG( | 0.444567 | 0.084942 | NO | NO | NA |
| Eml1 | ENSMUSG( | ENSMUSG( | 0.524973 | 0.052571 | NO | NO | NA |
| Zfp273 | ENSMUSG( | ENSMUSG( | -0.09838 | 0.719466 | NO | NO | NA |
| Tatdn3 | ENSMUSG( | ENSMUSG( | 0.345591 | 0.215273 | NO | NO | NA |
| ENSMUSG | ENSMUSG( | ENSMUSG( | -0.23964 | 0.223675 | NO | NO | NA |
| Neil2 | ENSMUSG( | ENSMUSG( | 0.039019 | 0.845218 | NO | NO | NA |
| Serac1 | ENSMUSG( | ENSMUSG( | 0.066548 | 0.738942 | NO | NO | NA |
| Kbtbd3 | ENSMUSG( | ENSMUSG( | -0.18217 | 0.470684 | NO | NO | NA |
| Pou6f2 | ENSMUSG( | ENSMUSG( | 0.485983 | 0.631498 | NO | NO | NA |
| Gm14276 | ENSMUSG( | ENSMUSG( | 0.051502 | 0.858682 | NO | NO | NA |
| Map7d3 | ENSMUSG( | ENSMUSG( | 0.631926 | 0.005553 | YES | NO | NA |
| Rab31 | ENSMUSG( | ENSMUSG( | -0.85254 | 0.000804 | YES | NO | NA |
| Them4 | ENSMUSG( | ENSMUSG( | 0.62771 | 0.000387 | YES | NO | NA |
| Igkv6-17 | ENSMUSG( | ENSMUSG( | -0.62984 | 0.745768 | NO | NO | NA |
| Gm50069 | ENSMUSG( | ENSMUSG( | 0.401269 | 0.097014 | NO | NO | NA |
| Nphp4 | ENSMUSG( | ENSMUSG( | -0.67779 | 0.001474 | YES | NO | NA |
| Gm28052 | ENSMUSG( | ENSMUSG( | -1.08525 | 0.795984 | NO | NO | NA |
| Gm16153 | ENSMUSG( | ENSMUSG( | 0.096189 | 0.723831 | NO | NO | NA |
| Cnrip1 | ENSMUSG( | ENSMUSG( | 0.073562 | 0.7084 | NO | NO | NA |
| Pls3 | ENSMUSG( | ENSMUSG( | 0.628713 | 0.019869 | YES | NO | NA |
| Gm8399 | ENSMUSG( | ENSMUSG( | 0.122356 | 0.688954 | NO | NO | NA |
| Atg9b | ENSMUSG( | ENSMUSG( | 0.773262 | 0.00423 | YES | NO | NA |
| ENSMUSG | ENSMUSG( | ENSMUSG( | -0.02915 | 0.900712 | NO | NO | NA |
| Tmie | ENSMUSG( | ENSMUSG( | 0.167331 | 0.767768 | NO | NO | NA |
| Synb | ENSMUSG( | ENSMUSG( | 1.259082 | 0.112826 | NO | NO | NA |
| Aldh1a1 | ENSMUSG( | ENSMUSG( | 1.00087 | 0.048335 | YES | YES | Upregulated |
| Tspan17 | ENSMUSG( | ENSMUSG( | 1.037199 | 0.001713 | YES | YES | Upregulated |
| Gm37531 | ENSMUSG( | ENSMUSG( | 0.669004 | 0.338824 | NO | NO | NA |
| Trip10 | ENSMUSG( | ENSMUSG( | 0.008385 | 0.975111 | NO | NO | NA |

| | | | | | | | |
|---|---|---|---|---|---|---|---|
| ENSMUSG | ENSMUSG( | ENSMUSG( | -0.69587 | 0.0005 | YES | NO | NA |
| Gm5835 | ENSMUSG( | ENSMUSG( | 0.575772 | 0.09841 | NO | NO | NA |
| Gm20663 | ENSMUSG( | ENSMUSG( | -0.7494 | 0.049954 | YES | NO | NA |
| Sox5 | ENSMUSG( | ENSMUSG( | 0.491406 | 0.476489 | NO | NO | NA |
| Gm43857 | ENSMUSG( | ENSMUSG( | -0.02032 | 0.939403 | NO | NO | NA |
| Mir99ahg | ENSMUSG( | ENSMUSG( | 0.792114 | 0.391549 | NO | NO | NA |
| H2bc24 | ENSMUSG( | ENSMUSG( | 0.497355 | 0.532011 | NO | NO | NA |
| Gm29966 | ENSMUSG( | ENSMUSG( | 0.903855 | 0.384622 | NO | NO | NA |
| Gm43984 | ENSMUSG( | ENSMUSG( | 0.760829 | 0.340615 | NO | NO | NA |
| Plxna4 | ENSMUSG( | ENSMUSG( | 0.858864 | 0.369772 | NO | NO | NA |
| B4galt4 | ENSMUSG( | ENSMUSG( | -0.24415 | 0.304683 | NO | NO | NA |
| Rarres2 | ENSMUSG( | ENSMUSG( | -0.47268 | 0.060666 | NO | NO | NA |
| 6030443J( | ENSMUSG( | ENSMUSG( | -0.17933 | 0.631802 | NO | NO | NA |
| Slc25a27 | ENSMUSG( | ENSMUSG( | 0.180265 | 0.564925 | NO | NO | NA |
| Gm44888 | ENSMUSG( | ENSMUSG( | 0.519286 | 0.025988 | YES | NO | NA |
| Tmem37 | ENSMUSG( | ENSMUSG( | -1.63562 | 1.33E-05 | YES | YES | Downregulated |
| Tlr6 | ENSMUSG( | ENSMUSG( | -0.49605 | 0.044701 | YES | NO | NA |
| Tctn2 | ENSMUSG( | ENSMUSG( | 0.580127 | 0.283032 | NO | NO | NA |
| Efcab11 | ENSMUSG( | ENSMUSG( | 0.147291 | 0.413118 | NO | NO | NA |
| Epcam | ENSMUSG( | ENSMUSG( | -0.12823 | 0.751869 | NO | NO | NA |
| Snx29 | ENSMUSG( | ENSMUSG( | -0.3862 | 0.034857 | YES | NO | NA |
| Il6ra | ENSMUSG( | ENSMUSG( | 0.102119 | 0.866209 | NO | NO | NA |
| B230217O | ENSMUSG( | ENSMUSG( | -0.03303 | 0.888657 | NO | NO | NA |
| Gm36486 | ENSMUSG( | ENSMUSG( | 0.445399 | 0.5778 | NO | NO | NA |
| 1700034P: | ENSMUSG( | ENSMUSG( | -0.1481 | 0.774128 | NO | NO | NA |
| Snhg11 | ENSMUSG( | ENSMUSG( | 0.578028 | 0.596032 | NO | NO | NA |
| Ckap4 | ENSMUSG( | ENSMUSG( | 0.516731 | 0.294113 | NO | NO | NA |
| Gm47467 | ENSMUSG( | ENSMUSG( | -0.17564 | 0.454093 | NO | NO | NA |
| Ftl1-ps1 | ENSMUSG( | ENSMUSG( | 0.398088 | 0.259199 | NO | NO | NA |
| Epha5 | ENSMUSG( | ENSMUSG( | 0.789706 | 0.310173 | NO | NO | NA |
| Gm36948 | ENSMUSG( | ENSMUSG( | 0.281219 | 0.287642 | NO | NO | NA |
| 4933438K: | ENSMUSG( | ENSMUSG( | 1.20674 | 0.11648 | NO | NO | NA |
| Camkmt | ENSMUSG( | ENSMUSG( | 0.140859 | 0.587657 | NO | NO | NA |
| Gm36266 | ENSMUSG( | ENSMUSG( | -0.36356 | 0.070622 | NO | NO | NA |
| Zfp438 | ENSMUSG( | ENSMUSG( | -0.02199 | 0.926387 | NO | NO | NA |
| Gm48838 | ENSMUSG( | ENSMUSG( | -0.11944 | 0.601687 | NO | NO | NA |
| Gm42798 | ENSMUSG( | ENSMUSG( | 0.233219 | 0.430364 | NO | NO | NA |
| Bmyc | ENSMUSG( | ENSMUSG( | 1.042809 | 0.000142 | YES | YES | Upregulated |
| Tnfrsf25 | ENSMUSG( | ENSMUSG( | -1.30811 | 0.000707 | YES | YES | Downregulated |
| Lmo2 | ENSMUSG( | ENSMUSG( | 0.550989 | 0.22187 | NO | NO | NA |
| Nudt8 | ENSMUSG( | ENSMUSG( | 0.074935 | 0.708633 | NO | NO | NA |
| Gm16845 | ENSMUSG( | ENSMUSG( | 0.537381 | 0.034552 | YES | NO | NA |
| Fstl1 | ENSMUSG( | ENSMUSG( | 0.867233 | 0.215086 | NO | NO | NA |
| F13a1 | ENSMUSG( | ENSMUSG( | 0.000902 | 0.998967 | NO | NO | NA |
| Hexim2 | ENSMUSG( | ENSMUSG( | 0.416537 | 0.141138 | NO | NO | NA |
| ENSMUSG( | ENSMUSG( | ENSMUSG( | 0.693527 | 0.013765 | YES | NO | NA |
| Usp35 | ENSMUSG( | ENSMUSG( | -0.5524 | 0.001219 | YES | NO | NA |

| | | | | | | | |
|---|---|---|---|---|---|---|---|
| Ttyh2 | ENSMUSG( | ENSMUSG( | -0.50017 | 0.11693 | NO | NO | NA |
| Mcf2l | ENSMUSG( | ENSMUSG( | 0.031637 | 0.970155 | NO | NO | NA |
| Lrfn4 | ENSMUSG( | ENSMUSG( | 0.938394 | 5.64E-05 | YES | NO | NA |
| D030055H | ENSMUSG( | ENSMUSG( | -0.07471 | 0.854284 | NO | NO | NA |
| 2010110K: | ENSMUSG( | ENSMUSG( | 0.419939 | 0.087387 | NO | NO | NA |
| Tom1l1 | ENSMUSG( | ENSMUSG( | 0.222592 | 0.308887 | NO | NO | NA |
| Kcnn3 | ENSMUSG( | ENSMUSG( | 1.995202 | 0.055475 | NO | NO | NA |
| Gm47345 | ENSMUSG( | ENSMUSG( | 0.075414 | 0.816091 | NO | NO | NA |
| Tvp23a | ENSMUSG( | ENSMUSG( | -0.05963 | 0.808851 | NO | NO | NA |
| Marcks | ENSMUSG( | ENSMUSG( | 0.195213 | 0.660302 | NO | NO | NA |
| Cacnb3 | ENSMUSG( | ENSMUSG( | 0.324521 | 0.291361 | NO | NO | NA |
| Trav8d-1 | ENSMUSG( | ENSMUSG( | -0.57756 | 0.607669 | NO | NO | NA |
| Ager | ENSMUSG( | ENSMUSG( | 0.361606 | 0.122859 | NO | NO | NA |
| Gm44672 | ENSMUSG( | ENSMUSG( | -0.02487 | 0.915968 | NO | NO | NA |
| Zfp971 | ENSMUSG( | ENSMUSG( | -0.60347 | 0.000424 | YES | NO | NA |
| A430072P | ENSMUSG( | ENSMUSG( | 0.671869 | 2.52E-05 | YES | NO | NA |
| Gm15821 | ENSMUSG( | ENSMUSG( | -0.38414 | 0.085295 | NO | NO | NA |
| 2210418O | ENSMUSG( | ENSMUSG( | 1.0862 | 0.656466 | NO | NO | NA |
| H2-Eb1 | ENSMUSG( | ENSMUSG( | -0.38442 | 0.384677 | NO | NO | NA |
| 1700113A | ENSMUSG( | ENSMUSG( | 0.050447 | 0.843786 | NO | NO | NA |
| Trav14d-3 | ENSMUSG( | ENSMUSG( | -0.35355 | 0.707721 | NO | NO | NA |
| Lhpp | ENSMUSG( | ENSMUSG( | 1.167009 | 6.43E-06 | YES | YES | Upregulated |
| Gm50371 | ENSMUSG( | ENSMUSG( | 2.031209 | 0.44719 | NO | NO | NA |
| Hk1os | ENSMUSG( | ENSMUSG( | 0.723519 | 0.032169 | YES | NO | NA |
| ENSMUSG | ENSMUSG( | ENSMUSG( | -0.30367 | 0.063891 | NO | NO | NA |
| D130040H | ENSMUSG( | ENSMUSG( | -0.56605 | 0.025472 | YES | NO | NA |
| Piezo2 | ENSMUSG( | ENSMUSG( | 0.684893 | 0.518069 | NO | NO | NA |
| Rassf6 | ENSMUSG( | ENSMUSG( | -0.41635 | 0.075892 | NO | NO | NA |
| Klra9 | ENSMUSG( | ENSMUSG( | -1.43728 | 0.009317 | YES | YES | Downregulated |
| Echdc2 | ENSMUSG( | ENSMUSG( | 0.176698 | 0.354493 | NO | NO | NA |
| Mfsd3 | ENSMUSG( | ENSMUSG( | 0.393676 | 0.008966 | YES | NO | NA |
| Tefm | ENSMUSG( | ENSMUSG( | 0.117135 | 0.595915 | NO | NO | NA |
| Mapre3 | ENSMUSG( | ENSMUSG( | -0.35708 | 0.211315 | NO | NO | NA |
| Mgst3 | ENSMUSG( | ENSMUSG( | 0.720222 | 0.062254 | NO | NO | NA |
| Grhl1 | ENSMUSG( | ENSMUSG( | -0.40226 | 0.104554 | NO | NO | NA |
| B930086L( | ENSMUSG( | ENSMUSG( | 0.871203 | 0.404866 | NO | NO | NA |
| Tle2 | ENSMUSG( | ENSMUSG( | 0.110996 | 0.846519 | NO | NO | NA |
| Gm44321 | ENSMUSG( | ENSMUSG( | -0.63713 | 0.019867 | YES | NO | NA |
| Cacna1s | ENSMUSG( | ENSMUSG( | -0.05615 | 0.855673 | NO | NO | NA |
| Evx2 | ENSMUSG( | ENSMUSG( | 4.567302 | 5.31E-40 | YES | YES | Upregulated |
| 4930562F( | ENSMUSG( | ENSMUSG( | 0.022679 | 0.943383 | NO | NO | NA |
| 2010320M | ENSMUSG( | ENSMUSG( | 0.573663 | 0.001032 | YES | NO | NA |
| Gm44502 | ENSMUSG( | ENSMUSG( | -0.06171 | 0.863447 | NO | NO | NA |
| Fam171b | ENSMUSG( | ENSMUSG( | 0.010978 | 0.968345 | NO | NO | NA |
| Gm42546 | ENSMUSG( | ENSMUSG( | 0.145265 | 0.806175 | NO | NO | NA |
| Pam | ENSMUSG( | ENSMUSG( | 0.82219 | 0.005388 | YES | NO | NA |
| H3c1 | ENSMUSG( | ENSMUSG( | 0.036745 | 0.877471 | NO | NO | NA |

| | | | | | | | |
|---|---|---|---|---|---|---|---|
| Pla2g4c | ENSMUSG( | ENSMUSG( | 0.125417 | 0.585511 | NO | NO | NA |
| Bnip5 | ENSMUSG( | ENSMUSG( | -0.02548 | 0.909884 | NO | NO | NA |
| Vwa7 | ENSMUSG( | ENSMUSG( | 0.178594 | 0.647827 | NO | NO | NA |
| 4921531C: | ENSMUSG( | ENSMUSG( | 0.042674 | 0.849426 | NO | NO | NA |
| Gm38160 | ENSMUSG( | ENSMUSG( | -0.33449 | 0.257374 | NO | NO | NA |
| BC051226 | ENSMUSG( | ENSMUSG( | -0.32101 | 0.110451 | NO | NO | NA |
| Pdcd1lg2 | ENSMUSG( | ENSMUSG( | 0.966294 | 2.90E-06 | YES | NO | NA |
| Gm5905 | ENSMUSG( | ENSMUSG( | 0.433047 | 0.102769 | NO | NO | NA |
| Trav12-1 | ENSMUSG( | ENSMUSG( | -0.58849 | 0.439849 | NO | NO | NA |
| Elovl7 | ENSMUSG( | ENSMUSG( | -1.13041 | 5.37E-05 | YES | YES | Downregulated |
| ENSMUSG | ENSMUSG( | ENSMUSG( | 0.286974 | 0.275501 | NO | NO | NA |
| Ms4a4a | ENSMUSG( | ENSMUSG( | 0.307451 | 0.231678 | NO | NO | NA |
| Cadps | ENSMUSG( | ENSMUSG( | 0.650772 | 0.494121 | NO | NO | NA |
| D430041D | ENSMUSG( | ENSMUSG( | 0.474693 | 0.490375 | NO | NO | NA |
| Sox8 | ENSMUSG( | ENSMUSG( | 0.597749 | 0.395157 | NO | NO | NA |
| 3830408C: | ENSMUSG( | ENSMUSG( | 0.330869 | 0.320371 | NO | NO | NA |
| Nr2f2 | ENSMUSG( | ENSMUSG( | 0.750712 | 0.44719 | NO | NO | NA |
| Zfp354c | ENSMUSG( | ENSMUSG( | 0.419624 | 0.088193 | NO | NO | NA |
| ENSMUSG | ENSMUSG( | ENSMUSG( | -0.67501 | 4.47E-05 | YES | NO | NA |
| Dyrk3 | ENSMUSG( | ENSMUSG( | 1.016642 | 0.001968 | YES | YES | Upregulated |
| ENSMUSG | ENSMUSG( | ENSMUSG( | 0.766724 | 0.010891 | YES | NO | NA |
| Gm25679 | ENSMUSG( | ENSMUSG( | 5.165399 | NA | NO | NO | NA |
| Aldh2 | ENSMUSG( | ENSMUSG( | 0.231768 | 0.515405 | NO | NO | NA |
| Vcam1 | ENSMUSG( | ENSMUSG( | 0.152516 | 0.769418 | NO | NO | NA |
| Kif19a | ENSMUSG( | ENSMUSG( | -0.29499 | 0.207201 | NO | NO | NA |
| Slc40a1 | ENSMUSG( | ENSMUSG( | -1.35933 | 1.64E-06 | YES | YES | Downregulated |
| Fsbp | ENSMUSG( | ENSMUSG( | -0.25305 | 0.365952 | NO | NO | NA |
| ENSMUSG | ENSMUSG( | ENSMUSG( | 0.370331 | 0.132409 | NO | NO | NA |
| Pgk1-rs7 | ENSMUSG( | ENSMUSG( | 22.89022 | 1.97E-12 | YES | YES | Upregulated |
| Gm47486 | ENSMUSG( | ENSMUSG( | -0.16562 | 0.575942 | NO | NO | NA |
| Podxl | ENSMUSG( | ENSMUSG( | 1.452631 | NA | NO | NO | NA |
| Pde11a | ENSMUSG( | ENSMUSG( | 0.884028 | 0.006615 | YES | NO | NA |
| Trav14d-2 | ENSMUSG( | ENSMUSG( | 1.644812 | 0.691734 | NO | NO | NA |
| Gm17139 | ENSMUSG( | ENSMUSG( | -0.10129 | 0.722881 | NO | NO | NA |
| Zfp335os | ENSMUSG( | ENSMUSG( | 0.294467 | 0.354911 | NO | NO | NA |
| Gm48768 | ENSMUSG( | ENSMUSG( | -0.13293 | 0.55607 | NO | NO | NA |
| Igf2bp2 | ENSMUSG( | ENSMUSG( | 0.452065 | 0.393076 | NO | NO | NA |
| ENSMUSG | ENSMUSG( | ENSMUSG( | -0.45176 | 0.027124 | YES | NO | NA |
| Fam83h | ENSMUSG( | ENSMUSG( | -0.38979 | 0.059548 | NO | NO | NA |
| Gm45718 | ENSMUSG( | ENSMUSG( | -0.28694 | 0.1937 | NO | NO | NA |
| Pank1 | ENSMUSG( | ENSMUSG( | 0.016286 | 0.970343 | NO | NO | NA |
| Gm16740 | ENSMUSG( | ENSMUSG( | -0.25344 | 0.123354 | NO | NO | NA |
| AI849053 | ENSMUSG( | ENSMUSG( | 0.596042 | 0.547015 | NO | NO | NA |
| C130051F( | ENSMUSG( | ENSMUSG( | 0.406318 | 0.502358 | NO | NO | NA |
| Gm12999 | ENSMUSG( | ENSMUSG( | -0.05117 | 0.854414 | NO | NO | NA |
| Gm42846 | ENSMUSG( | ENSMUSG( | -0.68368 | 0.012344 | YES | NO | NA |
| Plscr4 | ENSMUSG( | ENSMUSG( | -0.00825 | 0.974234 | NO | NO | NA |

| | | | | | | | |
|---|---|---|---|---|---|---|---|
| Fcgr2b | ENSMUSG( | ENSMUSG( | -0.6234 | 0.017459 | YES | NO | NA |
| Sfmbt2 | ENSMUSG( | ENSMUSG( | 1.091893 | 0.004863 | YES | YES | Upregulated |
| Asb2 | ENSMUSG( | ENSMUSG( | -0.37347 | 0.386225 | NO | NO | NA |
| Epb42 | ENSMUSG( | ENSMUSG( | 0.583569 | 0.408435 | NO | NO | NA |
| Gm16201 | ENSMUSG( | ENSMUSG( | 0.149864 | 0.646383 | NO | NO | NA |
| Cux2 | ENSMUSG( | ENSMUSG( | 0.57539 | 0.521422 | NO | NO | NA |
| Tmem35b | ENSMUSG( | ENSMUSG( | -0.42339 | 0.007052 | YES | NO | NA |
| ENSMUSG | ENSMUSG( | ENSMUSG( | 1.217825 | 2.66E-09 | YES | YES | Upregulated |
| Gm9776 | ENSMUSG( | ENSMUSG( | 0.247868 | 0.272908 | NO | NO | NA |
| Selenom | ENSMUSG( | ENSMUSG( | -0.27299 | 0.334128 | NO | NO | NA |
| Sgce | ENSMUSG( | ENSMUSG( | 0.805342 | 0.439849 | NO | NO | NA |
| Ighv14-2 | ENSMUSG( | ENSMUSG( | 0.743584 | 0.729762 | NO | NO | NA |
| Tmem273 | ENSMUSG( | ENSMUSG( | -0.66186 | 0.34275 | NO | NO | NA |
| Gm26852 | ENSMUSG( | ENSMUSG( | 0.151208 | 0.46221 | NO | NO | NA |
| ENSMUSG( | ENSMUSG( | ENSMUSG( | 0.400272 | 0.029667 | YES | NO | NA |
| Casc1 | ENSMUSG( | ENSMUSG( | 0.590864 | 0.30414 | NO | NO | NA |
| Ankrd55 | ENSMUSG( | ENSMUSG( | 0.633506 | 0.031839 | YES | NO | NA |
| Trim6 | ENSMUSG( | ENSMUSG( | 0.644618 | 0.504087 | NO | NO | NA |
| Gm3636 | ENSMUSG( | ENSMUSG( | -0.46563 | 0.008979 | YES | NO | NA |
| Gm38228 | ENSMUSG( | ENSMUSG( | -0.21541 | 0.46054 | NO | NO | NA |
| Gm15893 | ENSMUSG( | ENSMUSG( | -0.62012 | 0.001876 | YES | NO | NA |
| Gm42732 | ENSMUSG( | ENSMUSG( | -0.53394 | 0.003302 | YES | NO | NA |
| Kcnmb4 | ENSMUSG( | ENSMUSG( | 0.329282 | 0.230686 | NO | NO | NA |
| Gm28417 | ENSMUSG( | ENSMUSG( | -0.00076 | 0.997273 | NO | NO | NA |
| Golga7b | ENSMUSG( | ENSMUSG( | 0.757477 | 0.291361 | NO | NO | NA |
| Ntrk2 | ENSMUSG( | ENSMUSG( | 0.872998 | 0.420513 | NO | NO | NA |
| Gm16033 | ENSMUSG( | ENSMUSG( | -0.94431 | 0.111612 | NO | NO | NA |
| ENSMUSG( | ENSMUSG( | ENSMUSG( | -0.43772 | 0.159326 | NO | NO | NA |
| Gm45220 | ENSMUSG( | ENSMUSG( | -0.20258 | 0.34275 | NO | NO | NA |
| Thap6 | ENSMUSG( | ENSMUSG( | -0.74641 | 0.859542 | NO | NO | NA |
| Mapk11 | ENSMUSG( | ENSMUSG( | -0.22028 | 0.497741 | NO | NO | NA |
| Vmn2r93 | ENSMUSG( | ENSMUSG( | 0.527179 | 0.608826 | NO | NO | NA |
| Prkcz2 | ENSMUSG( | ENSMUSG( | -0.65727 | 0.001643 | YES | NO | NA |
| Egf | ENSMUSG( | ENSMUSG( | 1.020372 | 0.00029 | YES | YES | Upregulated |
| Vmn1r173 | ENSMUSG( | ENSMUSG( | 1.029799 | 0.358441 | NO | NO | NA |
| Mroh6 | ENSMUSG( | ENSMUSG( | 0.924759 | 0.109305 | NO | NO | NA |
| Prnp | ENSMUSG( | ENSMUSG( | 0.697769 | 0.004993 | YES | NO | NA |
| Mdk | ENSMUSG( | ENSMUSG( | -0.41972 | 0.044201 | YES | NO | NA |
| Wnt3 | ENSMUSG( | ENSMUSG( | 1.112969 | 0.213096 | NO | NO | NA |
| Wwc2 | ENSMUSG( | ENSMUSG( | 0.73435 | 0.207434 | NO | NO | NA |
| Gm43412 | ENSMUSG( | ENSMUSG( | 0.526268 | 0.310617 | NO | NO | NA |
| Gm37844 | ENSMUSG( | ENSMUSG( | 0.215819 | 0.437984 | NO | NO | NA |
| Cdkl5 | ENSMUSG( | ENSMUSG( | 0.409246 | 0.5075 | NO | NO | NA |
| Gm20458 | ENSMUSG( | ENSMUSG( | -0.52413 | 0.557697 | NO | NO | NA |
| Cntln | ENSMUSG( | ENSMUSG( | 0.356239 | 0.32708 | NO | NO | NA |
| ENSMUSG | ENSMUSG( | ENSMUSG( | -0.11949 | 0.69132 | NO | NO | NA |
| Fut4 | ENSMUSG( | ENSMUSG( | 2.117913 | 1.24E-08 | YES | YES | Upregulated |

| | | | | | | | |
|---|---|---|---|---|---|---|---|
| Gm4332 | ENSMUSG( | ENSMUSG( | 0.598663 | 0.043116 | YES | NO | NA |
| Spon1 | ENSMUSG( | ENSMUSG( | 0.391194 | 0.614582 | NO | NO | NA |
| Gm45477 | ENSMUSG( | ENSMUSG( | 0.409745 | 0.574926 | NO | NO | NA |
| C330011M | ENSMUSG( | ENSMUSG( | 0.505308 | 0.013146 | YES | NO | NA |
| Gm45854 | ENSMUSG( | ENSMUSG( | -0.50619 | 0.002503 | YES | NO | NA |
| Phldb1 | ENSMUSG( | ENSMUSG( | 0.971097 | 3.68E-05 | YES | NO | NA |
| Gm49668 | ENSMUSG( | ENSMUSG( | -0.30264 | 0.296812 | NO | NO | NA |
| Eef1akmt4 | ENSMUSG( | ENSMUSG( | 0.181566 | 0.473939 | NO | NO | NA |
| BC049715 | ENSMUSG( | ENSMUSG( | 0.233194 | 0.420972 | NO | NO | NA |
| Gm37968 | ENSMUSG( | ENSMUSG( | 0.317832 | 0.442282 | NO | NO | NA |
| Gm43372 | ENSMUSG( | ENSMUSG( | 0.339143 | 0.062031 | NO | NO | NA |
| Spata1 | ENSMUSG( | ENSMUSG( | -0.26825 | 0.124372 | NO | NO | NA |
| Tssk6 | ENSMUSG( | ENSMUSG( | 0.323406 | 0.212957 | NO | NO | NA |
| Lyrm7 | ENSMUSG( | ENSMUSG( | 0.070026 | 0.750793 | NO | NO | NA |
| Retsat | ENSMUSG( | ENSMUSG( | 0.588492 | 0.009148 | YES | NO | NA |
| Pou3f3 | ENSMUSG( | ENSMUSG( | 0.812479 | 0.308977 | NO | NO | NA |
| H2bc6 | ENSMUSG( | ENSMUSG( | 0.328271 | 0.047929 | YES | NO | NA |
| Amotl1 | ENSMUSG( | ENSMUSG( | 0.469322 | 0.143251 | NO | NO | NA |
| Lyz2 | ENSMUSG( | ENSMUSG( | -1.17939 | 0.179798 | NO | NO | NA |
| Misp3 | ENSMUSG( | ENSMUSG( | 0.788303 | 0.030238 | YES | NO | NA |
| Scn2a | ENSMUSG( | ENSMUSG( | 0.89327 | 0.242832 | NO | NO | NA |
| Gm5049 | ENSMUSG( | ENSMUSG( | -0.16909 | 0.581253 | NO | NO | NA |
| Cp | ENSMUSG( | ENSMUSG( | 0.573236 | 0.374884 | NO | NO | NA |
| H2ac18 | ENSMUSG( | ENSMUSG( | -1.94214 | 0.639774 | NO | NO | NA |
| Gm16069 | ENSMUSG( | ENSMUSG( | -0.37858 | 0.173088 | NO | NO | NA |
| Slc6a11 | ENSMUSG( | ENSMUSG( | 0.452539 | 0.593294 | NO | NO | NA |
| Gm43361 | ENSMUSG( | ENSMUSG( | 0.347979 | 0.440714 | NO | NO | NA |
| Cd320 | ENSMUSG( | ENSMUSG( | 0.253649 | 0.268971 | NO | NO | NA |
| ENSMUSG | ENSMUSG( | ENSMUSG( | 0.273137 | 0.289575 | NO | NO | NA |
| Trim2 | ENSMUSG( | ENSMUSG( | 0.464397 | 0.054662 | NO | NO | NA |
| Mcee | ENSMUSG( | ENSMUSG( | 0.193993 | 0.406566 | NO | NO | NA |
| Gm27162 | ENSMUSG( | ENSMUSG( | -0.27548 | 0.307158 | NO | NO | NA |
| Aqp3 | ENSMUSG( | ENSMUSG( | 0.048218 | 0.896584 | NO | NO | NA |
| Gm20069 | ENSMUSG( | ENSMUSG( | -1.15885 | 6.17E-06 | YES | YES | Downregulated |
| Gm44667 | ENSMUSG( | ENSMUSG( | -0.59512 | 0.007353 | YES | NO | NA |
| Zfp300 | ENSMUSG( | ENSMUSG( | -0.59399 | 0.025221 | YES | NO | NA |
| Slfn10-ps | ENSMUSG( | ENSMUSG( | 1.062075 | 0.80032 | NO | NO | NA |
| Gm43769 | ENSMUSG( | ENSMUSG( | 0.690383 | 0.001196 | YES | NO | NA |
| B130021K: | ENSMUSG( | ENSMUSG( | 0.148432 | 0.786278 | NO | NO | NA |
| Gnb5 | ENSMUSG( | ENSMUSG( | 0.170187 | 0.49586 | NO | NO | NA |
| Nlgn3 | ENSMUSG( | ENSMUSG( | 0.395486 | 0.600992 | NO | NO | NA |
| Gm8355 | ENSMUSG( | ENSMUSG( | -0.3112 | 0.16372 | NO | NO | NA |
| Gm28529 | ENSMUSG( | ENSMUSG( | -0.37567 | 0.252843 | NO | NO | NA |
| 5830487J( | ENSMUSG( | ENSMUSG( | 0.875499 | 0.201773 | NO | NO | NA |
| Nphs1 | ENSMUSG( | ENSMUSG( | 0.609905 | 0.003498 | YES | NO | NA |
| Nr2c1 | ENSMUSG( | ENSMUSG( | 0.468063 | 0.006963 | YES | NO | NA |
| Trav7d-3 | ENSMUSG( | ENSMUSG( | -0.47232 | 0.623013 | NO | NO | NA |

| | | | | | | | |
|---|---|---|---|---|---|---|---|
| Gm47567 | ENSMUSG( | ENSMUSG( | 0.38294 | 0.120917 | NO | NO | NA |
| H2ac13 | ENSMUSG( | ENSMUSG( | 0.268815 | 0.409549 | NO | NO | NA |
| Tspan15 | ENSMUSG( | ENSMUSG( | 0.846297 | 0.256896 | NO | NO | NA |
| Trav14-2 | ENSMUSG( | ENSMUSG( | -0.87922 | 0.128905 | NO | NO | NA |
| Gm47218 | ENSMUSG( | ENSMUSG( | -0.17521 | 0.476831 | NO | NO | NA |
| 9930014A | ENSMUSG( | ENSMUSG( | 0.265509 | 0.158073 | NO | NO | NA |
| Sag | ENSMUSG( | ENSMUSG( | 0.168784 | 0.757584 | NO | NO | NA |
| Cdc42ep4 | ENSMUSG( | ENSMUSG( | -0.43287 | 0.032907 | YES | NO | NA |
| Dmxl2 | ENSMUSG( | ENSMUSG( | 0.826838 | 0.002656 | YES | NO | NA |
| C230096K | ENSMUSG( | ENSMUSG( | -1.30551 | 6.31E-07 | YES | YES | Downregulated |
| Igf1 | ENSMUSG( | ENSMUSG( | 0.205008 | 0.816271 | NO | NO | NA |
| Gm38257 | ENSMUSG( | ENSMUSG( | 0.2029 | 0.616932 | NO | NO | NA |
| Gm14464 | ENSMUSG( | ENSMUSG( | 0.596876 | 0.558429 | NO | NO | NA |
| Enho | ENSMUSG( | ENSMUSG( | -1.43418 | 5.28E-05 | YES | YES | Downregulated |
| Mccc1os | ENSMUSG( | ENSMUSG( | 0.615064 | 0.459367 | NO | NO | NA |
| Trio | ENSMUSG( | ENSMUSG( | 0.109718 | 0.710681 | NO | NO | NA |
| Zfp619 | ENSMUSG( | ENSMUSG( | -0.14266 | 0.653047 | NO | NO | NA |
| Arfgef3 | ENSMUSG( | ENSMUSG( | 0.858173 | 0.093701 | NO | NO | NA |
| N4bp2os | ENSMUSG( | ENSMUSG( | 0.610423 | 0.009389 | YES | NO | NA |
| Gm15751 | ENSMUSG( | ENSMUSG( | -0.10943 | 0.667108 | NO | NO | NA |
| ENSMUSG | ENSMUSG( | ENSMUSG( | 0.606333 | 0.010427 | YES | NO | NA |
| Gfod2 | ENSMUSG( | ENSMUSG( | 0.435376 | 0.054601 | NO | NO | NA |
| Masp2 | ENSMUSG( | ENSMUSG( | -0.01594 | 0.956063 | NO | NO | NA |
| Slc25a23 | ENSMUSG( | ENSMUSG( | 0.100512 | 0.797665 | NO | NO | NA |
| B020010K | ENSMUSG( | ENSMUSG( | 0.428885 | 0.147833 | NO | NO | NA |
| Drc1 | ENSMUSG( | ENSMUSG( | -0.31868 | 0.558702 | NO | NO | NA |
| Dnah17 | ENSMUSG( | ENSMUSG( | -0.38374 | 0.022265 | YES | NO | NA |
| Gatm | ENSMUSG( | ENSMUSG( | 0.565728 | 0.142719 | NO | NO | NA |
| Rab3a | ENSMUSG( | ENSMUSG( | 0.013365 | 0.963549 | NO | NO | NA |
| Usp2 | ENSMUSG( | ENSMUSG( | 0.681989 | 0.219159 | NO | NO | NA |
| Ctsl | ENSMUSG( | ENSMUSG( | 0.693853 | 0.021551 | YES | NO | NA |
| Gm4285 | ENSMUSG( | ENSMUSG( | 0.442332 | 0.002509 | YES | NO | NA |
| Ssc4d | ENSMUSG( | ENSMUSG( | 0.500622 | 0.030021 | YES | NO | NA |
| Gm50455 | ENSMUSG( | ENSMUSG( | 0.001738 | 0.99688 | NO | NO | NA |
| Gm15964 | ENSMUSG( | ENSMUSG( | 0.664621 | 0.008797 | YES | NO | NA |
| Gm32444 | ENSMUSG( | ENSMUSG( | 0.992702 | 0.251982 | NO | NO | NA |
| Lmln | ENSMUSG( | ENSMUSG( | 0.590752 | 0.003991 | YES | NO | NA |
| Ighv14-4 | ENSMUSG( | ENSMUSG( | -0.23762 | 0.906423 | NO | NO | NA |
| Septin5 | ENSMUSG( | ENSMUSG( | -0.88503 | 7.95E-05 | YES | NO | NA |
| H4c3 | ENSMUSG( | ENSMUSG( | 0.092672 | 0.72037 | NO | NO | NA |
| Pdk2 | ENSMUSG( | ENSMUSG( | 0.311642 | 0.34315 | NO | NO | NA |
| ENSMUSG | ENSMUSG( | ENSMUSG( | -0.02134 | 0.957331 | NO | NO | NA |
| Tbx6 | ENSMUSG( | ENSMUSG( | -0.92161 | 0.000922 | YES | NO | NA |
| Gm43071 | ENSMUSG( | ENSMUSG( | -0.5272 | 0.012557 | YES | NO | NA |
| Gm20045 | ENSMUSG( | ENSMUSG( | 0.395373 | 0.446301 | NO | NO | NA |
| Gm15503 | ENSMUSG( | ENSMUSG( | 1.086893 | 0.159078 | NO | NO | NA |
| Gm15542 | ENSMUSG( | ENSMUSG( | -0.24604 | 0.242752 | NO | NO | NA |

| | | | | | | | |
|---|---|---|---|---|---|---|---|
| Igkv3-2 | ENSMUSG( | ENSMUSG( | -1.11815 | 0.579864 | NO | NO | NA |
| 2610037D | ENSMUSG( | ENSMUSG( | -0.06777 | 0.82508 | NO | NO | NA |
| Sp3os | ENSMUSG( | ENSMUSG( | 0.085428 | 0.644112 | NO | NO | NA |
| Gata1 | ENSMUSG( | ENSMUSG( | 0.856334 | 0.10766 | NO | NO | NA |
| ENSMUSG | ENSMUSG( | ENSMUSG( | -0.1502 | 0.589716 | NO | NO | NA |
| Rpl31-ps8 | ENSMUSG( | ENSMUSG( | 0.469904 | 0.04931 | YES | NO | NA |
| Cask | ENSMUSG( | ENSMUSG( | 0.261601 | 0.379251 | NO | NO | NA |
| Rps6kl1 | ENSMUSG( | ENSMUSG( | -0.07456 | 0.712884 | NO | NO | NA |
| Gm15655 | ENSMUSG( | ENSMUSG( | -0.84629 | 0.006187 | YES | NO | NA |
| Cacng8 | ENSMUSG( | ENSMUSG( | -0.39694 | 0.076324 | NO | NO | NA |
| Fbxo17 | ENSMUSG( | ENSMUSG( | 0.405095 | 0.093322 | NO | NO | NA |
| Cdc42bpa | ENSMUSG( | ENSMUSG( | 1.05678 | 0.000117 | YES | YES | Upregulated |
| Afp | ENSMUSG( | ENSMUSG( | 0.258635 | 0.661527 | NO | NO | NA |
| Fam135b | ENSMUSG( | ENSMUSG( | 0.455315 | 0.552816 | NO | NO | NA |
| Dusp3 | ENSMUSG( | ENSMUSG( | 0.417896 | 0.278855 | NO | NO | NA |
| Gm614 | ENSMUSG( | ENSMUSG( | 0.443724 | 0.159503 | NO | NO | NA |
| ENSMUSG | ENSMUSG( | ENSMUSG( | -0.20925 | 0.340385 | NO | NO | NA |
| Gfi1b | ENSMUSG( | ENSMUSG( | 1.230984 | 0.09368 | NO | NO | NA |
| ENSMUSG | ENSMUSG( | ENSMUSG( | -0.03079 | 0.944326 | NO | NO | NA |
| Tmem158 | ENSMUSG( | ENSMUSG( | 0.789005 | 6.19E-05 | YES | NO | NA |
| Btnl10 | ENSMUSG( | ENSMUSG( | 0.90754 | 0.145686 | NO | NO | NA |
| Ppargc1a | ENSMUSG( | ENSMUSG( | 0.769884 | 0.460915 | NO | NO | NA |
| Rpl17-ps1( | ENSMUSG( | ENSMUSG( | 0.126263 | 0.557442 | NO | NO | NA |
| Gm4767 | ENSMUSG( | ENSMUSG( | -0.29501 | 0.921433 | NO | NO | NA |
| Fat4 | ENSMUSG( | ENSMUSG( | 1.134413 | 0.000244 | YES | YES | Upregulated |
| Map9 | ENSMUSG( | ENSMUSG( | 1.624433 | 0.0002 | YES | YES | Upregulated |
| Ighv1-80 | ENSMUSG( | ENSMUSG( | 0.593662 | 0.749795 | NO | NO | NA |
| Syt2 | ENSMUSG( | ENSMUSG( | 0.893175 | 0.244982 | NO | NO | NA |
| Creb3l1 | ENSMUSG( | ENSMUSG( | -0.48345 | 0.042189 | YES | NO | NA |
| Ighv1-75 | ENSMUSG( | ENSMUSG( | -0.34635 | 0.838588 | NO | NO | NA |
| Gm16794 | ENSMUSG( | ENSMUSG( | -0.62761 | 0.004972 | YES | NO | NA |
| Gm43660 | ENSMUSG( | ENSMUSG( | 0.028724 | 0.903031 | NO | NO | NA |
| Gm47113 | ENSMUSG( | ENSMUSG( | 0.698212 | 0.325009 | NO | NO | NA |
| Cyp2s1 | ENSMUSG( | ENSMUSG( | 0.395987 | 0.358592 | NO | NO | NA |
| Gm24305 | ENSMUSG( | ENSMUSG( | 0.044151 | NA | NO | NO | NA |
| 4732440D | ENSMUSG( | ENSMUSG( | -0.39793 | 0.028299 | YES | NO | NA |
| A930006K( | ENSMUSG( | ENSMUSG( | -0.26352 | 0.267916 | NO | NO | NA |
| Maged2 | ENSMUSG( | ENSMUSG( | -0.30254 | 0.183838 | NO | NO | NA |
| Kcnk10 | ENSMUSG( | ENSMUSG( | 0.911205 | 0.272935 | NO | NO | NA |
| Lrp4 | ENSMUSG( | ENSMUSG( | 0.623526 | 0.567548 | NO | NO | NA |
| Kel | ENSMUSG( | ENSMUSG( | 0.781453 | 0.197429 | NO | NO | NA |
| 1700010I1 | ENSMUSG( | ENSMUSG( | 0.342662 | 0.116478 | NO | NO | NA |
| Gm44775 | ENSMUSG( | ENSMUSG( | -0.25131 | 0.391117 | NO | NO | NA |
| Rhbdl1 | ENSMUSG( | ENSMUSG( | -0.1466 | 0.555025 | NO | NO | NA |
| Tead1 | ENSMUSG( | ENSMUSG( | 0.547276 | 0.153528 | NO | NO | NA |
| Gm43668 | ENSMUSG( | ENSMUSG( | 0.145846 | 0.778761 | NO | NO | NA |
| Ksr2 | ENSMUSG( | ENSMUSG( | 0.620914 | 0.487109 | NO | NO | NA |

| | | | | | | | |
|---|---|---|---|---|---|---|---|
| Gm6863 | ENSMUSG( | ENSMUSG( | 0.399412 | 0.011297 | YES | NO | NA |
| Vmn1r174 | ENSMUSG( | ENSMUSG( | 0.099703 | 0.926943 | NO | NO | NA |
| Gm26789 | ENSMUSG( | ENSMUSG( | -0.67975 | 0.000329 | YES | NO | NA |
| Gabra2 | ENSMUSG( | ENSMUSG( | 0.734328 | 0.475214 | NO | NO | NA |
| 1700021A( | ENSMUSG( | ENSMUSG( | 0.299282 | 0.08674 | NO | NO | NA |
| Npdc1 | ENSMUSG( | ENSMUSG( | 0.265738 | 0.420985 | NO | NO | NA |
| Gm6815 | ENSMUSG( | ENSMUSG( | 1.078195 | 0.366356 | NO | NO | NA |
| Dnm3os | ENSMUSG( | ENSMUSG( | 0.984491 | 0.38636 | NO | NO | NA |
| Zfp9 | ENSMUSG( | ENSMUSG( | 0.520037 | 0.451447 | NO | NO | NA |
| Ckb | ENSMUSG( | ENSMUSG( | 0.379051 | 0.129145 | NO | NO | NA |
| Gm43375 | ENSMUSG( | ENSMUSG( | -0.12273 | 0.567401 | NO | NO | NA |
| Fzd6 | ENSMUSG( | ENSMUSG( | 1.345126 | 0.000635 | YES | YES | Upregulated |
| Olfr15 | ENSMUSG( | ENSMUSG( | 0.880686 | 0.401302 | NO | NO | NA |
| Rasl12 | ENSMUSG( | ENSMUSG( | -0.73922 | 0.009186 | YES | NO | NA |
| Upk1a | ENSMUSG( | ENSMUSG( | 0.17093 | 0.60205 | NO | NO | NA |
| Ift122 | ENSMUSG( | ENSMUSG( | 1.114665 | 2.24E-06 | YES | YES | Upregulated |
| Gm37270 | ENSMUSG( | ENSMUSG( | -0.51109 | 0.008007 | YES | NO | NA |
| Gm50012 | ENSMUSG( | ENSMUSG( | -0.25184 | 0.430818 | NO | NO | NA |
| Lurap1 | ENSMUSG( | ENSMUSG( | 0.58978 | 0.043441 | YES | NO | NA |
| Gm48796 | ENSMUSG( | ENSMUSG( | -0.09621 | 0.665813 | NO | NO | NA |
| Pde1c | ENSMUSG( | ENSMUSG( | 0.493004 | 0.470839 | NO | NO | NA |
| 5330438D | ENSMUSG( | ENSMUSG( | 0.042082 | 0.888905 | NO | NO | NA |
| Gm2296 | ENSMUSG( | ENSMUSG( | 0.12379 | 0.648535 | NO | NO | NA |
| Nutf2-ps1 | ENSMUSG( | ENSMUSG( | 0.530626 | 0.178072 | NO | NO | NA |
| Arnt2 | ENSMUSG( | ENSMUSG( | 1.609439 | 0.00023 | YES | YES | Upregulated |
| A530058O | ENSMUSG( | ENSMUSG( | 0.742687 | 0.500137 | NO | NO | NA |
| Akap5 | ENSMUSG( | ENSMUSG( | 0.270662 | 0.54169 | NO | NO | NA |
| ENSMUSG | ENSMUSG( | ENSMUSG( | 5.623622 | 0.022582 | YES | YES | Upregulated |
| Nyap2 | ENSMUSG( | ENSMUSG( | 2.872742 | 0.004002 | YES | YES | Upregulated |
| Gm19557 | ENSMUSG( | ENSMUSG( | -0.22114 | 0.32436 | NO | NO | NA |
| Olfr1175 | ENSMUSG( | ENSMUSG( | 0.875647 | 0.300869 | NO | NO | NA |
| Gm17354 | ENSMUSG( | ENSMUSG( | 0.184428 | 0.581594 | NO | NO | NA |
| Lifr | ENSMUSG( | ENSMUSG( | 0.526257 | 0.564209 | NO | NO | NA |
| Vmn1r88 | ENSMUSG( | ENSMUSG( | 0.825835 | 0.442778 | NO | NO | NA |
| ENSMUSG( | ENSMUSG( | ENSMUSG( | 0.176196 | 0.490816 | NO | NO | NA |
| Gm45235 | ENSMUSG( | ENSMUSG( | -0.3171 | 0.040257 | YES | NO | NA |
| Gm15265 | ENSMUSG( | ENSMUSG( | 0.171415 | 0.499294 | NO | NO | NA |
| 9330159M | ENSMUSG( | ENSMUSG( | 0.576277 | 0.300881 | NO | NO | NA |
| Cfp | ENSMUSG( | ENSMUSG( | -0.37536 | 0.136104 | NO | NO | NA |
| H4c2 | ENSMUSG( | ENSMUSG( | 0.353271 | 0.204983 | NO | NO | NA |
| Tusc1 | ENSMUSG( | ENSMUSG( | 1.404651 | 0.000214 | YES | YES | Upregulated |
| Gm38037 | ENSMUSG( | ENSMUSG( | 0.554714 | 0.074219 | NO | NO | NA |
| Gm10605 | ENSMUSG( | ENSMUSG( | 0.782645 | 0.377352 | NO | NO | NA |
| Slc36a3os | ENSMUSG( | ENSMUSG( | 0.531501 | 0.012712 | YES | NO | NA |
| Gm38126 | ENSMUSG( | ENSMUSG( | 0.309572 | 0.683443 | NO | NO | NA |
| Gm36423 | ENSMUSG( | ENSMUSG( | 0.25044 | 0.554406 | NO | NO | NA |
| Gm50240 | ENSMUSG( | ENSMUSG( | 0.252663 | 0.132447 | NO | NO | NA |

| | | | | | | | |
|---|---|---|---|---|---|---|---|
| Klhdc8b | ENSMUSG( | ENSMUSG( | 0.285039 | 0.216855 | NO | NO | NA |
| Gm43682 | ENSMUSG( | ENSMUSG( | -0.25239 | 0.260577 | NO | NO | NA |
| Chil3 | ENSMUSG( | ENSMUSG( | 1.312004 | 0.164596 | NO | NO | NA |
| Cd79b | ENSMUSG( | ENSMUSG( | -0.5564 | 0.00916 | YES | NO | NA |
| Grm1 | ENSMUSG( | ENSMUSG( | 0.737243 | 0.460937 | NO | NO | NA |
| Tulp1 | ENSMUSG( | ENSMUSG( | 0.486799 | 0.460444 | NO | NO | NA |
| Gm18911 | ENSMUSG( | ENSMUSG( | -0.66977 | 0.001469 | YES | NO | NA |
| Rell2 | ENSMUSG( | ENSMUSG( | 0.3684 | 0.020108 | YES | NO | NA |
| Thrb | ENSMUSG( | ENSMUSG( | 0.572577 | 0.209625 | NO | NO | NA |
| Ovgp1 | ENSMUSG( | ENSMUSG( | 0.769027 | 0.014987 | YES | NO | NA |
| Gm44103 | ENSMUSG( | ENSMUSG( | -0.08998 | 0.78063 | NO | NO | NA |
| Gm38099 | ENSMUSG( | ENSMUSG( | 0.271827 | 0.664907 | NO | NO | NA |
| Zfp442 | ENSMUSG( | ENSMUSG( | -0.79147 | 0.006598 | YES | NO | NA |
| Gm21269 | ENSMUSG( | ENSMUSG( | 0.659198 | 0.540391 | NO | NO | NA |
| Gm15725 | ENSMUSG( | ENSMUSG( | -0.45899 | 0.773862 | NO | NO | NA |
| Ndrg2 | ENSMUSG( | ENSMUSG( | -0.1108 | 0.663349 | NO | NO | NA |
| Dusp9 | ENSMUSG( | ENSMUSG( | 0.741133 | 0.046642 | YES | NO | NA |
| Nova2 | ENSMUSG( | ENSMUSG( | 0.805564 | 0.368228 | NO | NO | NA |
| Gsta4 | ENSMUSG( | ENSMUSG( | 0.282569 | 0.380231 | NO | NO | NA |
| Tulp3 | ENSMUSG( | ENSMUSG( | -0.05491 | 0.885181 | NO | NO | NA |
| Dennd5b | ENSMUSG( | ENSMUSG( | 0.204615 | 0.803676 | NO | NO | NA |
| Treml2 | ENSMUSG( | ENSMUSG( | 0.109728 | 0.827588 | NO | NO | NA |
| Acad10 | ENSMUSG( | ENSMUSG( | 0.075071 | 0.747282 | NO | NO | NA |
| Gm12346 | ENSMUSG( | ENSMUSG( | 0.185352 | 0.385031 | NO | NO | NA |
| Gm17484 | ENSMUSG( | ENSMUSG( | 0.354801 | 0.522159 | NO | NO | NA |
| Cdr2l | ENSMUSG( | ENSMUSG( | 0.338991 | 0.545923 | NO | NO | NA |
| Spin2c | ENSMUSG( | ENSMUSG( | 0.117324 | 0.586824 | NO | NO | NA |
| Kit | ENSMUSG( | ENSMUSG( | 0.72498 | 0.022949 | YES | NO | NA |
| Ccdc80 | ENSMUSG( | ENSMUSG( | -0.06356 | 0.935909 | NO | NO | NA |
| Ttll11 | ENSMUSG( | ENSMUSG( | 0.80433 | 0.012661 | YES | NO | NA |
| Slc12a3 | ENSMUSG( | ENSMUSG( | 0.256111 | 0.741133 | NO | NO | NA |
| Gdap1 | ENSMUSG( | ENSMUSG( | 1.108679 | 0.201683 | NO | NO | NA |
| F420014N | ENSMUSG( | ENSMUSG( | 0.114876 | 0.660508 | NO | NO | NA |
| A530040E | ENSMUSG( | ENSMUSG( | 0.856847 | 0.004369 | YES | NO | NA |
| Srrm4 | ENSMUSG( | ENSMUSG( | 0.699052 | 0.454063 | NO | NO | NA |
| Mss51 | ENSMUSG( | ENSMUSG( | 0.246192 | 0.216593 | NO | NO | NA |
| Cers3 | ENSMUSG( | ENSMUSG( | 0.829147 | 0.295802 | NO | NO | NA |
| Actr3-ps | ENSMUSG( | ENSMUSG( | -0.07225 | 0.750708 | NO | NO | NA |
| Begain | ENSMUSG( | ENSMUSG( | 5.53649 | 2.12E-24 | YES | YES | Upregulated |
| AA465934 | ENSMUSG( | ENSMUSG( | 0.386758 | 0.091563 | NO | NO | NA |
| Gm49864 | ENSMUSG( | ENSMUSG( | 0.533818 | 0.012201 | YES | NO | NA |
| Gstt3 | ENSMUSG( | ENSMUSG( | 0.397312 | 0.045763 | YES | NO | NA |
| Gata4 | ENSMUSG( | ENSMUSG( | 1.033468 | 0.25515 | NO | NO | NA |
| Pi16 | ENSMUSG( | ENSMUSG( | 0.586442 | 0.495589 | NO | NO | NA |
| Dbp | ENSMUSG( | ENSMUSG( | -0.49976 | 0.08069 | NO | NO | NA |
| Cd200r2 | ENSMUSG( | ENSMUSG( | -0.76672 | 0.443806 | NO | NO | NA |
| 1700016P( | ENSMUSG( | ENSMUSG( | -0.48841 | 0.040697 | YES | NO | NA |

| | | | | | | | |
|---|---|---|---|---|---|---|---|
| Gm10095 | ENSMUSG( | ENSMUSG( | 0.582334 | 0.005243 | YES | NO | NA |
| Hs1bp3 | ENSMUSG( | ENSMUSG( | -0.23455 | 0.294975 | NO | NO | NA |
| Zbtb46 | ENSMUSG( | ENSMUSG( | 0.276941 | 0.624748 | NO | NO | NA |
| Emilin2 | ENSMUSG( | ENSMUSG( | 0.268884 | 0.550555 | NO | NO | NA |
| Dzank1 | ENSMUSG( | ENSMUSG( | -0.70203 | 0.001514 | YES | NO | NA |
| Homer2 | ENSMUSG( | ENSMUSG( | 0.920029 | 0.212043 | NO | NO | NA |
| 1700001L( | ENSMUSG( | ENSMUSG( | 0.052061 | 0.851529 | NO | NO | NA |
| C5ar2 | ENSMUSG( | ENSMUSG( | 0.781514 | 0.35827 | NO | NO | NA |
| ENSMUSG | ENSMUSG( | ENSMUSG( | 2.590927 | 0.021712 | YES | YES | Upregulated |
| Trav13-1 | ENSMUSG( | ENSMUSG( | 0.275055 | 0.523035 | NO | NO | NA |
| ENSMUSG | ENSMUSG( | ENSMUSG( | 1.965462 | 0.005006 | YES | YES | Upregulated |
| Ypel4 | ENSMUSG( | ENSMUSG( | 0.653794 | 0.216002 | NO | NO | NA |
| Trav5n-4 | ENSMUSG( | ENSMUSG( | 1.223097 | 0.769876 | NO | NO | NA |
| Gm38345 | ENSMUSG( | ENSMUSG( | 0.724921 | 0.002572 | YES | NO | NA |
| Wnt5a | ENSMUSG( | ENSMUSG( | 0.588289 | 0.574532 | NO | NO | NA |
| Espn | ENSMUSG( | ENSMUSG( | 0.370051 | 0.493227 | NO | NO | NA |
| ENSMUSG | ENSMUSG( | ENSMUSG( | 22.72612 | NA | NO | NO | NA |
| Gm47772 | ENSMUSG( | ENSMUSG( | 0.348005 | 0.291658 | NO | NO | NA |
| Gm9625 | ENSMUSG( | ENSMUSG( | 0.591135 | 0.008417 | YES | NO | NA |
| Shtn1 | ENSMUSG( | ENSMUSG( | 1.098968 | 0.005167 | YES | YES | Upregulated |
| Efcab2 | ENSMUSG( | ENSMUSG( | -0.08877 | 0.76261 | NO | NO | NA |
| Maml3 | ENSMUSG( | ENSMUSG( | 0.923476 | 0.187742 | NO | NO | NA |
| Nfasc | ENSMUSG( | ENSMUSG( | 0.508659 | 0.594123 | NO | NO | NA |
| Gm20633 | ENSMUSG( | ENSMUSG( | 0.387038 | 0.250293 | NO | NO | NA |
| Magi2 | ENSMUSG( | ENSMUSG( | 0.285495 | 0.191677 | NO | NO | NA |
| Tbc1d24 | ENSMUSG( | ENSMUSG( | 0.322921 | 0.184715 | NO | NO | NA |
| Aipl1 | ENSMUSG( | ENSMUSG( | 0.758769 | 0.001355 | YES | NO | NA |
| Lpl | ENSMUSG( | ENSMUSG( | 0.578839 | 0.405994 | NO | NO | NA |
| Tbc1d16 | ENSMUSG( | ENSMUSG( | 0.103803 | 0.660228 | NO | NO | NA |
| Dusp23 | ENSMUSG( | ENSMUSG( | 0.570776 | 0.007407 | YES | NO | NA |
| Prickle1 | ENSMUSG( | ENSMUSG( | -0.74985 | 0.006027 | YES | NO | NA |
| Oscp1 | ENSMUSG( | ENSMUSG( | 1.038405 | 0.025348 | YES | YES | Upregulated |
| Rasd2 | ENSMUSG( | ENSMUSG( | 1.281111 | 0.001649 | YES | YES | Upregulated |
| E130218I0 | ENSMUSG( | ENSMUSG( | 0.092925 | 0.698028 | NO | NO | NA |
| 4930520O | ENSMUSG( | ENSMUSG( | 0.945987 | 0.003741 | YES | NO | NA |
| Klhl31 | ENSMUSG( | ENSMUSG( | 0.746436 | 0.475278 | NO | NO | NA |
| ENSMUSG | ENSMUSG( | ENSMUSG( | -0.57298 | 0.037692 | YES | NO | NA |
| Eef2-ps2 | ENSMUSG( | ENSMUSG( | 0.275171 | 0.392349 | NO | NO | NA |
| Mmp14 | ENSMUSG( | ENSMUSG( | -0.79754 | 0.35166 | NO | NO | NA |
| Mreg | ENSMUSG( | ENSMUSG( | 0.601929 | 0.421905 | NO | NO | NA |
| Gvin-ps3 | ENSMUSG( | ENSMUSG( | 0.523786 | 0.250576 | NO | NO | NA |
| ENSMUSG( | ENSMUSG( | ENSMUSG( | 3.371996 | 0.000625 | YES | YES | Upregulated |
| 2810454H | ENSMUSG( | ENSMUSG( | 0.75504 | 0.032078 | YES | NO | NA |
| Oaz1-ps | ENSMUSG( | ENSMUSG( | 0.185641 | 0.50674 | NO | NO | NA |
| 4930594C: | ENSMUSG( | ENSMUSG( | 0.099386 | 0.896294 | NO | NO | NA |
| Cacnb1 | ENSMUSG( | ENSMUSG( | -1.06846 | 0.000978 | YES | YES | Downregulated |
| Gm26992 | ENSMUSG( | ENSMUSG( | -0.22268 | 0.566439 | NO | NO | NA |

| | | | | | | | |
|---|---|---|---|---|---|---|---|
| Tafa2 | ENSMUSG( | ENSMUSG( | 0.661276 | 0.495445 | NO | NO | NA |
| Dnaaf4 | ENSMUSG( | ENSMUSG( | 0.24686 | 0.407702 | NO | NO | NA |
| ENSMUSG | ENSMUSG( | ENSMUSG( | -0.0445 | 0.859288 | NO | NO | NA |
| Lrrc29 | ENSMUSG( | ENSMUSG( | -0.00048 | 0.998292 | NO | NO | NA |
| Gm47594 | ENSMUSG( | ENSMUSG( | 0.583145 | 0.508487 | NO | NO | NA |
| Wnt4 | ENSMUSG( | ENSMUSG( | 0.906325 | 0.342526 | NO | NO | NA |
| Cadm3 | ENSMUSG( | ENSMUSG( | 0.640426 | 0.095254 | NO | NO | NA |
| Gm17084 | ENSMUSG( | ENSMUSG( | 0.087016 | 0.927775 | NO | NO | NA |
| Gda | ENSMUSG( | ENSMUSG( | 0.914906 | 0.05291 | NO | NO | NA |
| Gm50335 | ENSMUSG( | ENSMUSG( | -0.9004 | 9.52E-05 | YES | NO | NA |
| Cyp2u1 | ENSMUSG( | ENSMUSG( | 0.465818 | 0.534341 | NO | NO | NA |
| Abhd14b | ENSMUSG( | ENSMUSG( | -0.26355 | 0.337555 | NO | NO | NA |
| Adam22 | ENSMUSG( | ENSMUSG( | 0.383842 | 0.354347 | NO | NO | NA |
| Slc25a29 | ENSMUSG( | ENSMUSG( | -0.19771 | 0.499325 | NO | NO | NA |
| Wfikkn1 | ENSMUSG( | ENSMUSG( | 0.251255 | 0.349342 | NO | NO | NA |
| Gm47813 | ENSMUSG( | ENSMUSG( | 0.549217 | 0.01983 | YES | NO | NA |
| Chic1 | ENSMUSG( | ENSMUSG( | -0.38103 | 0.132226 | NO | NO | NA |
| Acvr2a | ENSMUSG( | ENSMUSG( | 0.258124 | 0.353567 | NO | NO | NA |
| Npas3 | ENSMUSG( | ENSMUSG( | 0.680802 | 0.491132 | NO | NO | NA |
| Tlr7 | ENSMUSG( | ENSMUSG( | -0.70137 | 0.041332 | YES | NO | NA |
| Gm47992 | ENSMUSG( | ENSMUSG( | 0.086532 | 0.657136 | NO | NO | NA |
| Gm36527 | ENSMUSG( | ENSMUSG( | -0.62606 | 0.146854 | NO | NO | NA |
| Disc1 | ENSMUSG( | ENSMUSG( | -0.88709 | 0.000487 | YES | NO | NA |
| Gm16576 | ENSMUSG( | ENSMUSG( | -0.0882 | 0.728736 | NO | NO | NA |
| Tlr1 | ENSMUSG( | ENSMUSG( | -0.28184 | 0.217409 | NO | NO | NA |
| Myo1d | ENSMUSG( | ENSMUSG( | 0.894814 | 0.128671 | NO | NO | NA |
| Gm49284 | ENSMUSG( | ENSMUSG( | -0.13928 | 0.561653 | NO | NO | NA |
| Zfp114 | ENSMUSG( | ENSMUSG( | 0.866408 | 0.105883 | NO | NO | NA |
| Igkv1-133 | ENSMUSG( | ENSMUSG( | 1.088758 | NA | NO | NO | NA |
| Farp2 | ENSMUSG( | ENSMUSG( | -0.07917 | 0.769596 | NO | NO | NA |
| Zfp618 | ENSMUSG( | ENSMUSG( | 1.129833 | 0.083305 | NO | NO | NA |
| Cdnf | ENSMUSG( | ENSMUSG( | 0.619494 | 0.02654 | YES | NO | NA |
| Gal3st2c | ENSMUSG( | ENSMUSG( | 0.449432 | 0.672924 | NO | NO | NA |
| H4c4 | ENSMUSG( | ENSMUSG( | 0.185261 | 0.40581 | NO | NO | NA |
| Eif4ebp3 | ENSMUSG( | ENSMUSG( | -0.3183 | 0.57329 | NO | NO | NA |
| Gm15853 | ENSMUSG( | ENSMUSG( | 0.116071 | 0.618922 | NO | NO | NA |
| Gm19951 | ENSMUSG( | ENSMUSG( | -0.83559 | 0.001186 | YES | NO | NA |
| Hecw1 | ENSMUSG( | ENSMUSG( | 0.420484 | 0.670522 | NO | NO | NA |
| Gm48086 | ENSMUSG( | ENSMUSG( | -0.08891 | 0.74816 | NO | NO | NA |
| Ovol2 | ENSMUSG( | ENSMUSG( | -0.49297 | 0.081922 | NO | NO | NA |
| Ube2cbp | ENSMUSG( | ENSMUSG( | 0.797031 | 0.000436 | YES | NO | NA |
| C430042M | ENSMUSG( | ENSMUSG( | -0.00301 | 0.990073 | NO | NO | NA |
| Slc43a1 | ENSMUSG( | ENSMUSG( | 0.726576 | 0.148823 | NO | NO | NA |
| ENSMUSG( | ENSMUSG( | ENSMUSG( | 0.832878 | 0.156638 | NO | NO | NA |
| Gm5914 | ENSMUSG( | ENSMUSG( | 0.864183 | 0.003302 | YES | NO | NA |
| Nalf1 | ENSMUSG( | ENSMUSG( | 0.609746 | 0.565343 | NO | NO | NA |
| Gm53028 | ENSMUSG( | ENSMUSG( | 0.623535 | 0.495961 | NO | NO | NA |

| | | | | | | | |
|---|---|---|---|---|---|---|---|
| Zfp112 | ENSMUSG( | ENSMUSG( | 0.733401 | 0.000726 | YES | NO | NA |
| Gm7666 | ENSMUSG( | ENSMUSG( | 0.302882 | 0.335751 | NO | NO | NA |
| Map4k3 | ENSMUSG( | ENSMUSG( | 0.104365 | 0.656561 | NO | NO | NA |
| Semp2l1 | ENSMUSG( | ENSMUSG( | 0.751226 | 0.487561 | NO | NO | NA |
| Sall4 | ENSMUSG( | ENSMUSG( | 0.677732 | 0.454144 | NO | NO | NA |
| Lrrtm3 | ENSMUSG( | ENSMUSG( | 0.986858 | 0.331288 | NO | NO | NA |
| Gm43362 | ENSMUSG( | ENSMUSG( | -0.3432 | 0.210228 | NO | NO | NA |
| Anks1b | ENSMUSG( | ENSMUSG( | 0.840971 | 0.28332 | NO | NO | NA |
| Mcrip2 | ENSMUSG( | ENSMUSG( | 1.357737 | 0.000283 | YES | YES | Upregulated |
| Trim10 | ENSMUSG( | ENSMUSG( | 0.759199 | 0.188844 | NO | NO | NA |
| Ppp1r9a | ENSMUSG( | ENSMUSG( | 0.506019 | 0.552587 | NO | NO | NA |
| A430103D | ENSMUSG( | ENSMUSG( | 0.198539 | 0.364915 | NO | NO | NA |
| Gm43622 | ENSMUSG( | ENSMUSG( | -0.23802 | 0.664803 | NO | NO | NA |
| Bst1 | ENSMUSG( | ENSMUSG( | 1.916022 | 3.88E-12 | YES | YES | Upregulated |
| Pla2g2d | ENSMUSG( | ENSMUSG( | 0.696561 | 0.415065 | NO | NO | NA |
| Snhg10 | ENSMUSG( | ENSMUSG( | 0.914963 | 0.001923 | YES | NO | NA |
| Trav7d-6 | ENSMUSG( | ENSMUSG( | -0.35007 | 0.818783 | NO | NO | NA |
| Gm11998 | ENSMUSG( | ENSMUSG( | 0.058452 | 0.775658 | NO | NO | NA |
| Otub2 | ENSMUSG( | ENSMUSG( | -0.44944 | 0.006691 | YES | NO | NA |
| Gja1 | ENSMUSG( | ENSMUSG( | 0.346506 | 0.640471 | NO | NO | NA |
| Plxnb1 | ENSMUSG( | ENSMUSG( | -0.13488 | 0.605421 | NO | NO | NA |
| Mcmdc2 | ENSMUSG( | ENSMUSG( | 0.648918 | 0.464432 | NO | NO | NA |
| Gm37899 | ENSMUSG( | ENSMUSG( | 0.557903 | 0.560577 | NO | NO | NA |
| ENSMUSG( | ENSMUSG( | ENSMUSG( | 1.307877 | 0.000162 | YES | YES | Upregulated |
| Gm43079 | ENSMUSG( | ENSMUSG( | 1.221069 | 0.315031 | NO | NO | NA |
| Six3 | ENSMUSG( | ENSMUSG( | 0.838621 | 0.281181 | NO | NO | NA |
| Rps19-ps6 | ENSMUSG( | ENSMUSG( | 0.559366 | 0.113438 | NO | NO | NA |
| Tnfrsf10b | ENSMUSG( | ENSMUSG( | 0.559701 | 0.031343 | YES | NO | NA |
| Etv1 | ENSMUSG( | ENSMUSG( | 0.653259 | 0.481811 | NO | NO | NA |
| Snord3b4 | ENSMUSG( | ENSMUSG( | 0.846899 | 0.840839 | NO | NO | NA |
| B4galt2 | ENSMUSG( | ENSMUSG( | 1.50128 | 8.88E-08 | YES | YES | Upregulated |
| Garnl3 | ENSMUSG( | ENSMUSG( | 1.159483 | 0.067478 | NO | NO | NA |
| Morn4 | ENSMUSG( | ENSMUSG( | 1.843068 | 3.05E-05 | YES | YES | Upregulated |
| Gab2 | ENSMUSG( | ENSMUSG( | 0.519072 | 0.01538 | YES | NO | NA |
| Gabrg3 | ENSMUSG( | ENSMUSG( | 0.658598 | 0.539282 | NO | NO | NA |
| Ly6m | ENSMUSG( | ENSMUSG( | 0.947575 | 0.161839 | NO | NO | NA |
| Sema3a | ENSMUSG( | ENSMUSG( | 0.727156 | 0.486893 | NO | NO | NA |
| Col7a1 | ENSMUSG( | ENSMUSG( | -0.04665 | 0.861496 | NO | NO | NA |
| E030037K( | ENSMUSG( | ENSMUSG( | 0.065157 | 0.866399 | NO | NO | NA |
| Fuom | ENSMUSG( | ENSMUSG( | 0.580067 | 0.102635 | NO | NO | NA |
| Olfr1489 | ENSMUSG( | ENSMUSG( | 0.83692 | 0.417283 | NO | NO | NA |
| Ighv1-69 | ENSMUSG( | ENSMUSG( | -5.75286 | 0.000364 | YES | YES | Downregulated |
| Pcolce2 | ENSMUSG( | ENSMUSG( | 0.978302 | 0.075796 | NO | NO | NA |
| Tenm2 | ENSMUSG( | ENSMUSG( | 0.696961 | 0.504893 | NO | NO | NA |
| Trav5d-4 | ENSMUSG( | ENSMUSG( | -0.32092 | 0.894638 | NO | NO | NA |
| Trav6n-5 | ENSMUSG( | ENSMUSG( | -1.18378 | 0.473151 | NO | NO | NA |
| Gm43511 | ENSMUSG( | ENSMUSG( | -0.221 | 0.539282 | NO | NO | NA |

| | | | | | | | |
|---|---|---|---|---|---|---|---|
| Vkorc1 | ENSMUSG( | ENSMUSG( | 0.040896 | 0.879904 | NO | NO | NA |
| Egfl8 | ENSMUSG( | ENSMUSG( | 0.153417 | 0.552587 | NO | NO | NA |
| Rab26 | ENSMUSG( | ENSMUSG( | -0.17232 | 0.453495 | NO | NO | NA |
| Gm37758 | ENSMUSG( | ENSMUSG( | -0.84496 | 0.03185 | YES | NO | NA |
| Rps13-ps1 | ENSMUSG( | ENSMUSG( | 0.253419 | 0.239491 | NO | NO | NA |
| Cep112 | ENSMUSG( | ENSMUSG( | 0.846606 | 0.050664 | NO | NO | NA |
| Sqor | ENSMUSG( | ENSMUSG( | 0.625614 | 0.463063 | NO | NO | NA |
| Ighv1-81 | ENSMUSG( | ENSMUSG( | -0.05572 | 0.98 | NO | NO | NA |
| Unc5b | ENSMUSG( | ENSMUSG( | -0.08458 | 0.794703 | NO | NO | NA |
| 5730471H | ENSMUSG( | ENSMUSG( | 0.324667 | 0.168709 | NO | NO | NA |
| Zfp867 | ENSMUSG( | ENSMUSG( | 0.021293 | 0.953526 | NO | NO | NA |
| Eepd1 | ENSMUSG( | ENSMUSG( | 0.356258 | 0.204861 | NO | NO | NA |
| Apol11a | ENSMUSG( | ENSMUSG( | 0.700777 | 0.279846 | NO | NO | NA |
| Rgcc | ENSMUSG( | ENSMUSG( | 0.216312 | 0.538693 | NO | NO | NA |
| Igfbp4 | ENSMUSG( | ENSMUSG( | 0.503685 | 0.421499 | NO | NO | NA |
| H2ac20 | ENSMUSG( | ENSMUSG( | 0.240332 | 0.469209 | NO | NO | NA |
| Cldn13 | ENSMUSG( | ENSMUSG( | 1.158251 | 0.12706 | NO | NO | NA |
| Gm14681 | ENSMUSG( | ENSMUSG( | 0.8701 | 2.69E-05 | YES | NO | NA |
| Gm42572 | ENSMUSG( | ENSMUSG( | -0.40939 | 0.042886 | YES | NO | NA |
| 9230114K: | ENSMUSG( | ENSMUSG( | 0.408532 | 0.209647 | NO | NO | NA |
| Slco2a1 | ENSMUSG( | ENSMUSG( | 0.368288 | 0.586288 | NO | NO | NA |
| Tenm4 | ENSMUSG( | ENSMUSG( | 0.675662 | 0.484233 | NO | NO | NA |
| Grik4 | ENSMUSG( | ENSMUSG( | 0.492684 | 0.498479 | NO | NO | NA |
| Smim27 | ENSMUSG( | ENSMUSG( | 0.086125 | 0.694605 | NO | NO | NA |
| Bphl | ENSMUSG( | ENSMUSG( | 0.710263 | 0.001206 | YES | NO | NA |
| Tincr | ENSMUSG( | ENSMUSG( | 0.478276 | 0.184787 | NO | NO | NA |
| Gm20496 | ENSMUSG( | ENSMUSG( | -0.30542 | 0.352217 | NO | NO | NA |
| Fgd4 | ENSMUSG( | ENSMUSG( | 0.47823 | 0.5778 | NO | NO | NA |
| Gm38056 | ENSMUSG( | ENSMUSG( | 0.53328 | 0.184323 | NO | NO | NA |
| 2810403D | ENSMUSG( | ENSMUSG( | 0.429156 | 0.056628 | NO | NO | NA |
| Ptpn14 | ENSMUSG( | ENSMUSG( | 0.732151 | 0.135477 | NO | NO | NA |
| ENSMUSG | ENSMUSG( | ENSMUSG( | 0.546697 | 0.056102 | NO | NO | NA |
| Gm37123 | ENSMUSG( | ENSMUSG( | -0.15964 | 0.495195 | NO | NO | NA |
| Tmem120l | ENSMUSG( | ENSMUSG( | -0.30284 | 0.300735 | NO | NO | NA |
| Hebp1 | ENSMUSG( | ENSMUSG( | 0.523195 | 0.271037 | NO | NO | NA |
| 1700088E( | ENSMUSG( | ENSMUSG( | -0.16527 | 0.511078 | NO | NO | NA |
| Gm48374 | ENSMUSG( | ENSMUSG( | -0.02117 | 0.951406 | NO | NO | NA |
| Gm6710 | ENSMUSG( | ENSMUSG( | 0.270384 | 0.432969 | NO | NO | NA |
| Zfp931 | ENSMUSG( | ENSMUSG( | -0.46409 | 0.014396 | YES | NO | NA |
| Gm10277 | ENSMUSG( | ENSMUSG( | 0.978248 | 0.330466 | NO | NO | NA |
| Igkv13-85 | ENSMUSG( | ENSMUSG( | -3.7263 | 0.088785 | NO | NO | NA |
| ENSMUSG( | ENSMUSG( | ENSMUSG( | 0.360014 | 0.257374 | NO | NO | NA |
| Zfp94 | ENSMUSG( | ENSMUSG( | -0.14935 | 0.481807 | NO | NO | NA |
| Gm6649 | ENSMUSG( | ENSMUSG( | -0.4031 | 0.047584 | YES | NO | NA |
| Epb41l4a | ENSMUSG( | ENSMUSG( | 0.396763 | 0.562672 | NO | NO | NA |
| Trank1 | ENSMUSG( | ENSMUSG( | 0.589094 | 0.502427 | NO | NO | NA |
| Gm43775 | ENSMUSG( | ENSMUSG( | 0.622969 | 0.050894 | NO | NO | NA |

| | | | | | | | |
|---|---|---|---|---|---|---|---|
| Gm29170 | ENSMUSG( | ENSMUSG( | 0.360682 | 0.160976 | NO | NO | NA |
| 9230111E( | ENSMUSG( | ENSMUSG( | 0.337624 | 0.310965 | NO | NO | NA |
| Sox12 | ENSMUSG( | ENSMUSG( | 0.76692 | 0.002621 | YES | NO | NA |
| Lgi2 | ENSMUSG( | ENSMUSG( | -0.44744 | 0.3274 | NO | NO | NA |
| 9330020H | ENSMUSG( | ENSMUSG( | 0.440572 | 0.288228 | NO | NO | NA |
| Serpinf1 | ENSMUSG( | ENSMUSG( | 0.86008 | 0.018365 | YES | NO | NA |
| Gm43915 | ENSMUSG( | ENSMUSG( | 0.538854 | 0.594481 | NO | NO | NA |
| G0s2 | ENSMUSG( | ENSMUSG( | -1.3065 | 0.002627 | YES | YES | Downregulated |
| Gm47817 | ENSMUSG( | ENSMUSG( | 0.412857 | 0.251969 | NO | NO | NA |
| Plce1 | ENSMUSG( | ENSMUSG( | -0.62788 | 0.048465 | YES | NO | NA |
| Tlcd2 | ENSMUSG( | ENSMUSG( | 1.069919 | 5.06E-09 | YES | YES | Upregulated |
| ENSMUSG | ENSMUSG( | ENSMUSG( | 0.34687 | 0.484435 | NO | NO | NA |
| 2610035D | ENSMUSG( | ENSMUSG( | -1.34621 | 0.000101 | YES | YES | Downregulated |
| Gm9828 | ENSMUSG( | ENSMUSG( | 0.249122 | 0.217082 | NO | NO | NA |
| Senp8 | ENSMUSG( | ENSMUSG( | 0.037295 | 0.863262 | NO | NO | NA |
| Gm12610 | ENSMUSG( | ENSMUSG( | 0.513409 | 0.032164 | YES | NO | NA |
| Ophn1 | ENSMUSG( | ENSMUSG( | 0.747559 | 0.481147 | NO | NO | NA |
| BC049352 | ENSMUSG( | ENSMUSG( | -0.77921 | 0.000323 | YES | NO | NA |
| Gm42548 | ENSMUSG( | ENSMUSG( | 0.355509 | 0.495261 | NO | NO | NA |
| Zscan4e | ENSMUSG( | ENSMUSG( | 0.553744 | 0.500146 | NO | NO | NA |
| Map3k13 | ENSMUSG( | ENSMUSG( | 0.086222 | 0.850516 | NO | NO | NA |
| Igkv12-41 | ENSMUSG( | ENSMUSG( | -1.8522 | 0.640013 | NO | NO | NA |
| ENSMUSG | ENSMUSG( | ENSMUSG( | 1.014104 | 0.336704 | NO | NO | NA |
| Gm13421 | ENSMUSG( | ENSMUSG( | -0.10104 | 0.722873 | NO | NO | NA |
| Lrrk2 | ENSMUSG( | ENSMUSG( | 0.180794 | 0.63183 | NO | NO | NA |
| Gm49839 | ENSMUSG( | ENSMUSG( | 0.059133 | 0.902365 | NO | NO | NA |
| Gm20512 | ENSMUSG( | ENSMUSG( | 0.837666 | 0.335851 | NO | NO | NA |
| Gm42566 | ENSMUSG( | ENSMUSG( | -0.71332 | 0.16217 | NO | NO | NA |
| Dab1 | ENSMUSG( | ENSMUSG( | 0.858677 | 0.401556 | NO | NO | NA |
| Mgll | ENSMUSG( | ENSMUSG( | 0.712821 | 0.193762 | NO | NO | NA |
| Olfr52 | ENSMUSG( | ENSMUSG( | 0.94559 | 0.379022 | NO | NO | NA |
| Trpt1 | ENSMUSG( | ENSMUSG( | -0.40671 | 0.022724 | YES | NO | NA |
| E430014B( | ENSMUSG( | ENSMUSG( | 0.962807 | 0.009519 | YES | NO | NA |
| Dpf1 | ENSMUSG( | ENSMUSG( | 0.878583 | 0.122348 | NO | NO | NA |
| Fbn2 | ENSMUSG( | ENSMUSG( | 0.570493 | 0.513234 | NO | NO | NA |
| Klrb1f | ENSMUSG( | ENSMUSG( | 0.072234 | 0.843977 | NO | NO | NA |
| Olfr157 | ENSMUSG( | ENSMUSG( | 0.531549 | 0.60627 | NO | NO | NA |
| Guca1b | ENSMUSG( | ENSMUSG( | 0.723036 | 0.250719 | NO | NO | NA |
| Vmn1r188 | ENSMUSG( | ENSMUSG( | 0.835818 | 0.419135 | NO | NO | NA |
| ENSMUSG | ENSMUSG( | ENSMUSG( | 0.456369 | 0.220769 | NO | NO | NA |
| Gm9870 | ENSMUSG( | ENSMUSG( | 0.333469 | 0.226023 | NO | NO | NA |
| Gm48998 | ENSMUSG( | ENSMUSG( | 0.642812 | 0.252906 | NO | NO | NA |
| Setmar | ENSMUSG( | ENSMUSG( | 0.229152 | 0.496381 | NO | NO | NA |
| Pheta2 | ENSMUSG( | ENSMUSG( | -0.31871 | 0.178666 | NO | NO | NA |
| Dusp8 | ENSMUSG( | ENSMUSG( | 0.676872 | 0.272908 | NO | NO | NA |
| ENSMUSG( | ENSMUSG( | ENSMUSG( | 0.343387 | 0.391549 | NO | NO | NA |
| Gm43042 | ENSMUSG( | ENSMUSG( | 0.062345 | 0.808208 | NO | NO | NA |

| | | | | | | | |
|---|---|---|---|---|---|---|---|
| Gm45706 | ENSMUSG( | ENSMUSG( | 0.045538 | 0.906948 | NO | NO | NA |
| Prrg1 | ENSMUSG( | ENSMUSG( | 0.543129 | 0.009378 | YES | NO | NA |
| Lzts3 | ENSMUSG( | ENSMUSG( | -0.17801 | 0.537294 | NO | NO | NA |
| ENSMUSG | ENSMUSG( | ENSMUSG( | -0.41658 | 0.224118 | NO | NO | NA |
| Gm9866 | ENSMUSG( | ENSMUSG( | 0.706842 | 0.530438 | NO | NO | NA |
| Olfr373 | ENSMUSG( | ENSMUSG( | 0.603778 | 0.57351 | NO | NO | NA |
| S100a9 | ENSMUSG( | ENSMUSG( | 2.239724 | 0.058901 | NO | NO | NA |
| Tgfbr3l | ENSMUSG( | ENSMUSG( | -0.30521 | 0.342553 | NO | NO | NA |
| Nav3 | ENSMUSG( | ENSMUSG( | 0.518167 | 0.610434 | NO | NO | NA |
| Inpp5a | ENSMUSG( | ENSMUSG( | 0.274506 | 0.334843 | NO | NO | NA |
| 5830468F( | ENSMUSG( | ENSMUSG( | 0.515955 | 0.056332 | NO | NO | NA |
| Insm1 | ENSMUSG( | ENSMUSG( | 1.542898 | 0.016372 | YES | YES | Upregulated |
| Clmn | ENSMUSG( | ENSMUSG( | 0.812393 | 0.373338 | NO | NO | NA |
| Myo1b | ENSMUSG( | ENSMUSG( | 1.493025 | 5.99E-08 | YES | YES | Upregulated |
| Trav14-3 | ENSMUSG( | ENSMUSG( | 0.887744 | 0.23533 | NO | NO | NA |
| Alox8 | ENSMUSG( | ENSMUSG( | -0.29521 | 0.320058 | NO | NO | NA |
| Ifi204 | ENSMUSG( | ENSMUSG( | -0.33248 | 0.26286 | NO | NO | NA |
| Ccdc184 | ENSMUSG( | ENSMUSG( | 0.916416 | 0.164525 | NO | NO | NA |
| Aldh1b1 | ENSMUSG( | ENSMUSG( | 0.328696 | 0.124413 | NO | NO | NA |
| Gm36037 | ENSMUSG( | ENSMUSG( | 0.612306 | 0.5876 | NO | NO | NA |
| Astn1 | ENSMUSG( | ENSMUSG( | 0.55533 | 0.548545 | NO | NO | NA |
| Pakap ENS | ENSMUSG( | ENSMUSG( | 0.101951 | 0.849759 | NO | NO | NA |
| ENSMUSG | ENSMUSG( | ENSMUSG( | 0.739435 | 0.050894 | NO | NO | NA |
| Fstl4 | ENSMUSG( | ENSMUSG( | 0.838622 | 0.309058 | NO | NO | NA |
| Gm6934 | ENSMUSG( | ENSMUSG( | 0.641491 | 0.243018 | NO | NO | NA |
| Mir5108 | ENSMUSG( | ENSMUSG( | 3.951856 | NA | NO | NO | NA |
| Klra7 | ENSMUSG( | ENSMUSG( | -0.16087 | 0.61867 | NO | NO | NA |
| C3ar1 | ENSMUSG( | ENSMUSG( | 0.756152 | 0.422216 | NO | NO | NA |
| Tbc1d32 | ENSMUSG( | ENSMUSG( | 0.443145 | 0.027503 | YES | NO | NA |
| Prkra | ENSMUSG( | ENSMUSG( | -0.43313 | 0.106118 | NO | NO | NA |
| Gm28874 | ENSMUSG( | ENSMUSG( | 0.336144 | 0.609227 | NO | NO | NA |
| Ccl6 | ENSMUSG( | ENSMUSG( | -0.44106 | 0.203731 | NO | NO | NA |
| Gm43039 | ENSMUSG( | ENSMUSG( | -0.48794 | 0.129146 | NO | NO | NA |
| Dpep2 | ENSMUSG( | ENSMUSG( | 0.406886 | 0.518236 | NO | NO | NA |
| A930012O | ENSMUSG( | ENSMUSG( | 0.376868 | 0.018888 | YES | NO | NA |
| Gm43281 | ENSMUSG( | ENSMUSG( | 1.088604 | 0.281886 | NO | NO | NA |
| Cib2 | ENSMUSG( | ENSMUSG( | -0.33626 | 0.391595 | NO | NO | NA |
| B230334C( | ENSMUSG( | ENSMUSG( | 0.881638 | 0.062984 | NO | NO | NA |
| Rdh16 | ENSMUSG( | ENSMUSG( | 0.770824 | 0.456638 | NO | NO | NA |
| Galnt6os | ENSMUSG( | ENSMUSG( | -0.32503 | 0.211291 | NO | NO | NA |
| Irag1 | ENSMUSG( | ENSMUSG( | -0.19921 | 0.480943 | NO | NO | NA |
| Igkv17-127 | ENSMUSG( | ENSMUSG( | -0.41484 | 0.854408 | NO | NO | NA |
| Foxp3 | ENSMUSG( | ENSMUSG( | 1.169457 | 0.000409 | YES | YES | Upregulated |
| Trav15n-2 | ENSMUSG( | ENSMUSG( | 3.895606 | 0.214688 | NO | NO | NA |
| Itpa-ps1 | ENSMUSG( | ENSMUSG( | 0.403779 | 0.066335 | NO | NO | NA |
| Zfp93 | ENSMUSG( | ENSMUSG( | -0.19591 | 0.404432 | NO | NO | NA |
| Gm10163 | ENSMUSG( | ENSMUSG( | 0.198415 | 0.868434 | NO | NO | NA |

| | | | | | | | |
|---|---|---|---|---|---|---|---|
| Icam5 | ENSMUSG( | ENSMUSG( | 0.613585 | 0.056766 | NO | NO | NA |
| Gm45178 | ENSMUSG( | ENSMUSG( | 0.886161 | 0.412461 | NO | NO | NA |
| Gm6789 | ENSMUSG( | ENSMUSG( | 0.1483 | 0.609081 | NO | NO | NA |
| C730027H | ENSMUSG( | ENSMUSG( | 0.502554 | 0.638292 | NO | NO | NA |
| Ccdc122 | ENSMUSG( | ENSMUSG( | 1.679915 | 7.12E-06 | YES | YES | Upregulated |
| Wt1 | ENSMUSG( | ENSMUSG( | 0.699391 | 0.467432 | NO | NO | NA |
| Gm27221 | ENSMUSG( | ENSMUSG( | 8.141602 | 0.001267 | YES | YES | Upregulated |
| Cox6b2 | ENSMUSG( | ENSMUSG( | 0.442857 | 0.241243 | NO | NO | NA |
| Sec16b | ENSMUSG( | ENSMUSG( | 1.408818 | 0.009536 | YES | YES | Upregulated |
| Ifngas1 | ENSMUSG( | ENSMUSG( | -0.77707 | 0.041941 | YES | NO | NA |
| Dipk1c | ENSMUSG( | ENSMUSG( | 0.760803 | 0.466633 | NO | NO | NA |
| Ighv1-15 | ENSMUSG( | ENSMUSG( | 0.656677 | 0.730299 | NO | NO | NA |
| Foxc1 | ENSMUSG( | ENSMUSG( | 0.515785 | 0.498286 | NO | NO | NA |
| Rhbdl3 | ENSMUSG( | ENSMUSG( | 0.657406 | 0.302628 | NO | NO | NA |
| Pcsk4 | ENSMUSG( | ENSMUSG( | 0.991821 | 0.000973 | YES | NO | NA |
| Gm17655 | ENSMUSG( | ENSMUSG( | -0.32813 | 0.213096 | NO | NO | NA |
| Tdh | ENSMUSG( | ENSMUSG( | 4.470509 | 2.29E-12 | YES | YES | Upregulated |
| Rpl18-ps2 | ENSMUSG( | ENSMUSG( | 0.45147 | 0.152519 | NO | NO | NA |
| 3110045C: | ENSMUSG( | ENSMUSG( | 1.005193 | 0.131965 | NO | NO | NA |
| Stau2 | ENSMUSG( | ENSMUSG( | 1.195635 | 2.32E-07 | YES | YES | Upregulated |
| Gm37472 | ENSMUSG( | ENSMUSG( | -0.08506 | 0.784462 | NO | NO | NA |
| Akr1b8 | ENSMUSG( | ENSMUSG( | 0.523831 | 0.051278 | NO | NO | NA |
| Gm36989 | ENSMUSG( | ENSMUSG( | -0.01543 | 0.961918 | NO | NO | NA |
| Capn5 | ENSMUSG( | ENSMUSG( | 0.244782 | 0.30875 | NO | NO | NA |
| Gpat3 | ENSMUSG( | ENSMUSG( | -0.44332 | 0.318298 | NO | NO | NA |
| Igkv6-32 | ENSMUSG( | ENSMUSG( | 1.487537 | 0.32874 | NO | NO | NA |
| Celf5 | ENSMUSG( | ENSMUSG( | -0.75251 | 0.015641 | YES | NO | NA |
| Phyhd1 | ENSMUSG( | ENSMUSG( | -0.8214 | 0.000507 | YES | NO | NA |
| Gm20236 | ENSMUSG( | ENSMUSG( | 0.543168 | 0.845218 | NO | NO | NA |
| Mid1-ps1 | ENSMUSG( | ENSMUSG( | 0.368949 | 0.831172 | NO | NO | NA |
| ENSMUSG | ENSMUSG( | ENSMUSG( | 8.099465 | 0.02487 | YES | YES | Upregulated |
| Trav12d-3 | ENSMUSG( | ENSMUSG( | 0.371338 | 0.878987 | NO | NO | NA |
| Gm15472 | ENSMUSG( | ENSMUSG( | -0.45214 | 0.029598 | YES | NO | NA |
| Odad4 | ENSMUSG( | ENSMUSG( | 0.766532 | 0.436803 | NO | NO | NA |
| Robo2 | ENSMUSG( | ENSMUSG( | 0.648935 | 0.542733 | NO | NO | NA |
| Gm9958 | ENSMUSG( | ENSMUSG( | 0.530425 | 0.051554 | NO | NO | NA |
| Btnl5-ps | ENSMUSG( | ENSMUSG( | -1.58461 | 2.18E-07 | YES | YES | Downregulated |
| Pnp2 | ENSMUSG( | ENSMUSG( | 1.055615 | 0.001376 | YES | YES | Upregulated |
| Elovl4 | ENSMUSG( | ENSMUSG( | 0.724696 | 0.157795 | NO | NO | NA |
| Gm5617 | ENSMUSG( | ENSMUSG( | 0.277977 | 0.289299 | NO | NO | NA |
| A630066F: | ENSMUSG( | ENSMUSG( | 0.560328 | 0.160294 | NO | NO | NA |
| Pitx1 | ENSMUSG( | ENSMUSG( | 0.631317 | 0.512637 | NO | NO | NA |
| Mob3b | ENSMUSG( | ENSMUSG( | 0.877886 | 0.122348 | NO | NO | NA |
| Gm20768 | ENSMUSG( | ENSMUSG( | 0.722709 | 0.307944 | NO | NO | NA |
| Gm42890 | ENSMUSG( | ENSMUSG( | 0.34986 | 0.440252 | NO | NO | NA |
| Gm11769 | ENSMUSG( | ENSMUSG( | -0.07929 | 0.854072 | NO | NO | NA |
| Tbx20 | ENSMUSG( | ENSMUSG( | 0.673933 | 0.527225 | NO | NO | NA |

| | | | | | | | |
|---|---|---|---|---|---|---|---|
| Adgrb2 | ENSMUSG( | ENSMUSG( | -1.08786 | 5.93E-05 | YES | YES | Downregulated |
| Ppm1l | ENSMUSG( | ENSMUSG( | -0.19622 | 0.351904 | NO | NO | NA |
| Sv2a | ENSMUSG( | ENSMUSG( | 0.1743 | 0.635716 | NO | NO | NA |
| Rasgrf2 | ENSMUSG( | ENSMUSG( | -0.81305 | 0.001192 | YES | NO | NA |
| Bmpr1a | ENSMUSG( | ENSMUSG( | 1.134711 | 3.56E-05 | YES | YES | Upregulated |
| Rbm3-ps | ENSMUSG( | ENSMUSG( | 1.286663 | 7.47E-05 | YES | YES | Upregulated |
| Igkv1-135 | ENSMUSG( | ENSMUSG( | 3.445992 | 0.002131 | YES | YES | Upregulated |
| Acot1 | ENSMUSG( | ENSMUSG( | 0.705462 | 0.422361 | NO | NO | NA |
| Gm44510 | ENSMUSG( | ENSMUSG( | 0.685095 | 0.050567 | NO | NO | NA |
| Gm49405 | ENSMUSG( | ENSMUSG( | 0.115402 | 0.667108 | NO | NO | NA |
| Eya4 | ENSMUSG( | ENSMUSG( | 0.661077 | 0.528139 | NO | NO | NA |
| Shc3 | ENSMUSG( | ENSMUSG( | 0.662041 | 0.391249 | NO | NO | NA |
| Slc6a8 | ENSMUSG( | ENSMUSG( | -0.83902 | 2.11E-05 | YES | NO | NA |
| Sobp | ENSMUSG( | ENSMUSG( | 0.874961 | 0.259005 | NO | NO | NA |
| A630081D | ENSMUSG( | ENSMUSG( | -0.38053 | 0.286196 | NO | NO | NA |
| B530045E: | ENSMUSG( | ENSMUSG( | 0.869537 | 0.356412 | NO | NO | NA |
| Gm42878 | ENSMUSG( | ENSMUSG( | 0.188425 | 0.588195 | NO | NO | NA |
| Zbtb16 | ENSMUSG( | ENSMUSG( | -0.66217 | 0.353769 | NO | NO | NA |
| Atcayos | ENSMUSG( | ENSMUSG( | 0.66857 | 0.075498 | NO | NO | NA |
| Gm6451 | ENSMUSG( | ENSMUSG( | -0.04351 | 0.908766 | NO | NO | NA |
| Kcna2 | ENSMUSG( | ENSMUSG( | 0.180131 | 0.779287 | NO | NO | NA |
| Cracd | ENSMUSG( | ENSMUSG( | 0.614634 | 0.113012 | NO | NO | NA |
| Trav6n-6 | ENSMUSG( | ENSMUSG( | 0.420611 | 0.628903 | NO | NO | NA |
| Itga7 | ENSMUSG( | ENSMUSG( | 0.359944 | 0.074842 | NO | NO | NA |
| Gm37019 | ENSMUSG( | ENSMUSG( | -0.46457 | 0.023084 | YES | NO | NA |
| Pfn2 | ENSMUSG( | ENSMUSG( | 0.099531 | 0.756906 | NO | NO | NA |
| Slc49a4 | ENSMUSG( | ENSMUSG( | -0.05811 | 0.838685 | NO | NO | NA |
| Rassf9 | ENSMUSG( | ENSMUSG( | 0.724683 | 0.488252 | NO | NO | NA |
| Ankrd24 | ENSMUSG( | ENSMUSG( | -0.02697 | 0.902128 | NO | NO | NA |
| Gm37204 | ENSMUSG( | ENSMUSG( | 0.414259 | 0.611216 | NO | NO | NA |
| Ldlrad3 | ENSMUSG( | ENSMUSG( | 0.842232 | 0.210872 | NO | NO | NA |
| Zfp383 | ENSMUSG( | ENSMUSG( | -0.37657 | 0.226163 | NO | NO | NA |
| Wdr54 | ENSMUSG( | ENSMUSG( | 0.501638 | 0.17547 | NO | NO | NA |
| H4f16 | ENSMUSG( | ENSMUSG( | 0.260831 | 0.60722 | NO | NO | NA |
| Tmc4 | ENSMUSG( | ENSMUSG( | 0.145359 | 0.520755 | NO | NO | NA |
| Pard6b | ENSMUSG( | ENSMUSG( | 0.226431 | 0.24806 | NO | NO | NA |
| Cdh23 | ENSMUSG( | ENSMUSG( | 0.822094 | 0.002216 | YES | NO | NA |
| Edn1 | ENSMUSG( | ENSMUSG( | 1.067732 | 0.307102 | NO | NO | NA |
| Rmst | ENSMUSG( | ENSMUSG( | 0.64381 | 0.513424 | NO | NO | NA |
| Gm45501 | ENSMUSG( | ENSMUSG( | 0.148374 | 0.72037 | NO | NO | NA |
| Acaa1b | ENSMUSG( | ENSMUSG( | 1.361681 | 0.002057 | YES | YES | Upregulated |
| Sox9 | ENSMUSG( | ENSMUSG( | 0.606115 | 0.574263 | NO | NO | NA |
| 5830462I1 | ENSMUSG( | ENSMUSG( | 0.189005 | 0.45373 | NO | NO | NA |
| Akap6 | ENSMUSG( | ENSMUSG( | 0.700965 | 0.412453 | NO | NO | NA |
| A530072M | ENSMUSG( | ENSMUSG( | -0.32513 | 0.053255 | NO | NO | NA |
| Gm34730 | ENSMUSG( | ENSMUSG( | 0.647123 | 0.528481 | NO | NO | NA |
| Trim7 | ENSMUSG( | ENSMUSG( | 0.70127 | 0.213759 | NO | NO | NA |

| | | | | | | | |
|---|---|---|---|---|---|---|---|
| Gm14419 | ENSMUSG( | ENSMUSG( | -0.51834 | 0.764367 | NO | NO | NA |
| Fgd6 | ENSMUSG( | ENSMUSG( | -0.25753 | 0.229506 | NO | NO | NA |
| Fahd1 | ENSMUSG( | ENSMUSG( | 0.663126 | 0.071745 | NO | NO | NA |
| Efr3b | ENSMUSG( | ENSMUSG( | 0.923883 | 0.02414 | YES | NO | NA |
| Tmem132 | ENSMUSG( | ENSMUSG( | 0.43062 | 0.671503 | NO | NO | NA |
| Meis2 | ENSMUSG( | ENSMUSG( | 0.9198 | 0.127757 | NO | NO | NA |
| Gm29642 | ENSMUSG( | ENSMUSG( | 0.38477 | 0.624377 | NO | NO | NA |
| Synpo | ENSMUSG( | ENSMUSG( | 0.056321 | 0.840661 | NO | NO | NA |
| Zfp28 | ENSMUSG( | ENSMUSG( | 0.000338 | 0.99919 | NO | NO | NA |
| Rftn2 | ENSMUSG( | ENSMUSG( | -0.44171 | 0.02892 | YES | NO | NA |
| Fcgbp | ENSMUSG( | ENSMUSG( | 0.492235 | 0.639774 | NO | NO | NA |
| Calcb | ENSMUSG( | ENSMUSG( | 0.414579 | 0.47552 | NO | NO | NA |
| Neurl2 | ENSMUSG( | ENSMUSG( | 0.061065 | 0.84935 | NO | NO | NA |
| Gm47512 | ENSMUSG( | ENSMUSG( | -0.09479 | 0.818756 | NO | NO | NA |
| Ccdc177 | ENSMUSG( | ENSMUSG( | 0.743471 | 0.480181 | NO | NO | NA |
| Olfr718-ps | ENSMUSG( | ENSMUSG( | 0.78767 | 0.447247 | NO | NO | NA |
| Slc29a2 | ENSMUSG( | ENSMUSG( | 1.032507 | 0.000129 | YES | YES | Upregulated |
| Col23a1 | ENSMUSG( | ENSMUSG( | -0.4955 | 0.018778 | YES | NO | NA |
| ENSMUSG( | ENSMUSG( | ENSMUSG( | 0.529824 | 0.033589 | YES | NO | NA |
| Pomc | ENSMUSG( | ENSMUSG( | 0.589934 | 0.037711 | YES | NO | NA |
| Apol8 | ENSMUSG( | ENSMUSG( | 0.956074 | 0.055887 | NO | NO | NA |
| Gm17477 | ENSMUSG( | ENSMUSG( | 0.467333 | 0.439807 | NO | NO | NA |
| 0610009L1 | ENSMUSG( | ENSMUSG( | 0.53213 | 0.212989 | NO | NO | NA |
| Gm42887 | ENSMUSG( | ENSMUSG( | -0.24273 | 0.345824 | NO | NO | NA |
| Spry1 | ENSMUSG( | ENSMUSG( | 0.163195 | 0.545331 | NO | NO | NA |
| Gm11423 | ENSMUSG( | ENSMUSG( | 0.614681 | 0.016288 | YES | NO | NA |
| Arhgap20 | ENSMUSG( | ENSMUSG( | 0.773252 | 0.292571 | NO | NO | NA |
| Gm43473 | ENSMUSG( | ENSMUSG( | 0.909951 | 0.398638 | NO | NO | NA |
| Fgfr1 | ENSMUSG( | ENSMUSG( | 0.365746 | 0.644442 | NO | NO | NA |
| Tmsb15b1 | ENSMUSG( | ENSMUSG( | 0.609964 | 0.014119 | YES | NO | NA |
| Mc5r | ENSMUSG( | ENSMUSG( | 0.786842 | 0.476823 | NO | NO | NA |
| Gm37110 | ENSMUSG( | ENSMUSG( | 1.124882 | 0.741426 | NO | NO | NA |
| ENSMUSG | ENSMUSG( | ENSMUSG( | 0.462068 | 0.236434 | NO | NO | NA |
| Osr2 | ENSMUSG( | ENSMUSG( | 0.210401 | 0.654812 | NO | NO | NA |
| Daam2 | ENSMUSG( | ENSMUSG( | 1.649549 | 2.30E-07 | YES | YES | Upregulated |
| A230103L1 | ENSMUSG( | ENSMUSG( | 0.758797 | 0.342553 | NO | NO | NA |
| Gm45051 | ENSMUSG( | ENSMUSG( | 0.201739 | 0.612737 | NO | NO | NA |
| Gm10602 | ENSMUSG( | ENSMUSG( | 0.148102 | 0.559933 | NO | NO | NA |
| C030037D | ENSMUSG( | ENSMUSG( | -0.02883 | 0.949522 | NO | NO | NA |
| C86187 | ENSMUSG( | ENSMUSG( | 0.250422 | 0.741154 | NO | NO | NA |
| Rgs17 | ENSMUSG( | ENSMUSG( | 0.603665 | 0.545451 | NO | NO | NA |
| Nptxr | ENSMUSG( | ENSMUSG( | 0.532159 | 0.130616 | NO | NO | NA |
| Mboat2 | ENSMUSG( | ENSMUSG( | 0.600651 | 0.404865 | NO | NO | NA |
| E430018J2 | ENSMUSG( | ENSMUSG( | -0.00852 | 0.968621 | NO | NO | NA |
| Tfap2b | ENSMUSG( | ENSMUSG( | 0.580524 | 0.571517 | NO | NO | NA |
| Arl10 | ENSMUSG( | ENSMUSG( | -0.23847 | 0.394758 | NO | NO | NA |
| H2-Aa | ENSMUSG( | ENSMUSG( | -0.74184 | 0.11664 | NO | NO | NA |

| | | | | | | | |
|---|---|---|---|---|---|---|---|
| 1700018L( | ENSMUSG( | ENSMUSG( | 0.041231 | 0.91234 | NO | NO | NA |
| Thsd4 | ENSMUSG( | ENSMUSG( | 0.601365 | 0.512362 | NO | NO | NA |
| Hoxc9 | ENSMUSG( | ENSMUSG( | 0.80507 | 0.321549 | NO | NO | NA |
| Fcrl1 | ENSMUSG( | ENSMUSG( | -0.11359 | 0.600814 | NO | NO | NA |
| Gm12596 | ENSMUSG( | ENSMUSG( | -0.35519 | 0.170033 | NO | NO | NA |
| Dennd2c | ENSMUSG( | ENSMUSG( | 0.260886 | 0.431435 | NO | NO | NA |
| Aldh6a1 | ENSMUSG( | ENSMUSG( | -0.07181 | 0.810767 | NO | NO | NA |
| Rasd1 | ENSMUSG( | ENSMUSG( | 0.61235 | 0.048809 | YES | NO | NA |
| Gabrb1 | ENSMUSG( | ENSMUSG( | 0.713196 | 0.373981 | NO | NO | NA |
| C530043K: | ENSMUSG( | ENSMUSG( | 0.273539 | 0.797406 | NO | NO | NA |
| Vcp-rs | ENSMUSG( | ENSMUSG( | 0.374073 | 0.213047 | NO | NO | NA |
| Zfp119a | ENSMUSG( | ENSMUSG( | 0.406675 | 0.160171 | NO | NO | NA |
| Gm8750 | ENSMUSG( | ENSMUSG( | -1.25979 | 0.569739 | NO | NO | NA |
| Gm37510 | ENSMUSG( | ENSMUSG( | 0.649801 | 0.424383 | NO | NO | NA |
| Trim66 | ENSMUSG( | ENSMUSG( | 1.2974 | 0.001306 | YES | YES | Upregulated |
| Zfp334 | ENSMUSG( | ENSMUSG( | 1.046173 | 0.254516 | NO | NO | NA |
| Samd4 | ENSMUSG( | ENSMUSG( | 0.887101 | 0.338396 | NO | NO | NA |
| Cldn34c1 | ENSMUSG( | ENSMUSG( | -0.48298 | 0.345916 | NO | NO | NA |
| Amer2 | ENSMUSG( | ENSMUSG( | 0.869947 | 0.429253 | NO | NO | NA |
| Gm13477 | ENSMUSG( | ENSMUSG( | -1.29692 | 4.43E-06 | YES | YES | Downregulated |
| A930007A | ENSMUSG( | ENSMUSG( | 0.267957 | 0.35979 | NO | NO | NA |
| Gm10101 | ENSMUSG( | ENSMUSG( | 0.632267 | 0.20132 | NO | NO | NA |
| Gm37589 | ENSMUSG( | ENSMUSG( | -0.90431 | 6.34E-05 | YES | NO | NA |
| Fam124b | ENSMUSG( | ENSMUSG( | -0.48898 | 0.008366 | YES | NO | NA |
| Dclk1 | ENSMUSG( | ENSMUSG( | 1.372678 | 1.13E-05 | YES | YES | Upregulated |
| Gmpr | ENSMUSG( | ENSMUSG( | -0.41571 | 0.126934 | NO | NO | NA |
| Dpp10 | ENSMUSG( | ENSMUSG( | 0.626936 | 0.558196 | NO | NO | NA |
| Gm47078 | ENSMUSG( | ENSMUSG( | 0.738428 | 0.526752 | NO | NO | NA |
| Lonrf2 | ENSMUSG( | ENSMUSG( | 0.736085 | 0.509479 | NO | NO | NA |
| Gm10184 | ENSMUSG( | ENSMUSG( | -0.01246 | 0.970155 | NO | NO | NA |
| Tubb2b | ENSMUSG( | ENSMUSG( | 0.728235 | 0.05206 | NO | NO | NA |
| Ceacam1 | ENSMUSG( | ENSMUSG( | 0.910588 | 0.258597 | NO | NO | NA |
| Gm10874 | ENSMUSG( | ENSMUSG( | -0.46073 | 0.06689 | NO | NO | NA |
| Gm20043 | ENSMUSG( | ENSMUSG( | 0.870301 | 0.403362 | NO | NO | NA |
| Rab4a | ENSMUSG( | ENSMUSG( | 0.265865 | 0.522983 | NO | NO | NA |
| Zfp128 | ENSMUSG( | ENSMUSG( | -0.08857 | 0.799139 | NO | NO | NA |
| Ajm1 | ENSMUSG( | ENSMUSG( | -0.75003 | 0.010844 | YES | NO | NA |
| Arhgef5 | ENSMUSG( | ENSMUSG( | 0.005296 | 0.984434 | NO | NO | NA |
| Irak1bp1 | ENSMUSG( | ENSMUSG( | 1.359308 | 0.005509 | YES | YES | Upregulated |
| Zpbp2 | ENSMUSG( | ENSMUSG( | -0.01278 | 0.96972 | NO | NO | NA |
| Gm16278 | ENSMUSG( | ENSMUSG( | -0.09541 | 0.718217 | NO | NO | NA |
| Gnai1 | ENSMUSG( | ENSMUSG( | 2.9891 | 5.79E-06 | YES | YES | Upregulated |
| Ica1l | ENSMUSG( | ENSMUSG( | 0.040005 | 0.908138 | NO | NO | NA |
| Nhsl1 | ENSMUSG( | ENSMUSG( | 0.468215 | 0.593014 | NO | NO | NA |
| Gm37127 | ENSMUSG( | ENSMUSG( | -0.17565 | 0.573921 | NO | NO | NA |
| Gm13340 | ENSMUSG( | ENSMUSG( | 0.456845 | 0.081928 | NO | NO | NA |
| 2510046G | ENSMUSG( | ENSMUSG( | 0.71576 | 0.050617 | NO | NO | NA |

| | | | | | | | |
|---|---|---|---|---|---|---|---|
| ENSMUSG | ENSMUSG( | ENSMUSG( | 0.128158 | 0.649552 | NO | NO | NA |
| Plcd1 | ENSMUSG( | ENSMUSG( | 0.375675 | 0.247984 | NO | NO | NA |
| Tssk4 | ENSMUSG( | ENSMUSG( | -0.35739 | 0.277656 | NO | NO | NA |
| Gm31614 | ENSMUSG( | ENSMUSG( | 0.710373 | 0.503828 | NO | NO | NA |
| ENSMUSG | ENSMUSG( | ENSMUSG( | 0.912006 | 0.346834 | NO | NO | NA |
| Slc7a11 | ENSMUSG( | ENSMUSG( | 0.846633 | 0.264824 | NO | NO | NA |
| Rabl2 | ENSMUSG( | ENSMUSG( | 1.330145 | 0.000193 | YES | YES | Upregulated |
| Gm37274 | ENSMUSG( | ENSMUSG( | -0.21406 | 0.496053 | NO | NO | NA |
| Klhl29 | ENSMUSG( | ENSMUSG( | 0.789129 | 0.464359 | NO | NO | NA |
| Scn4b | ENSMUSG( | ENSMUSG( | 0.958604 | 0.000566 | YES | NO | NA |
| AU041133 | ENSMUSG( | ENSMUSG( | -0.20635 | 0.516419 | NO | NO | NA |
| Dmac2l | ENSMUSG( | ENSMUSG( | 0.119718 | 0.616442 | NO | NO | NA |
| Cd59a | ENSMUSG( | ENSMUSG( | 1.184402 | 0.058363 | NO | NO | NA |
| ENSMUSG( | ENSMUSG( | ENSMUSG( | -0.03837 | 0.91237 | NO | NO | NA |
| Olfr920 | ENSMUSG( | ENSMUSG( | 0.725001 | 0.389245 | NO | NO | NA |
| Gipc3 | ENSMUSG( | ENSMUSG( | 0.802495 | 0.002815 | YES | NO | NA |
| Npas4 | ENSMUSG( | ENSMUSG( | -0.61755 | 0.012072 | YES | NO | NA |
| Usp44 | ENSMUSG( | ENSMUSG( | 1.251881 | 6.73E-05 | YES | YES | Upregulated |
| Trav13n-3 | ENSMUSG( | ENSMUSG( | -1.47164 | 0.461948 | NO | NO | NA |
| Zfp987 | ENSMUSG( | ENSMUSG( | -3.28232 | 8.62E-13 | YES | YES | Downregulated |
| A330035P | ENSMUSG( | ENSMUSG( | 0.581285 | 0.115093 | NO | NO | NA |
| Elane | ENSMUSG( | ENSMUSG( | 1.212786 | 0.184315 | NO | NO | NA |
| Gm44751 | ENSMUSG( | ENSMUSG( | -0.09623 | 0.691822 | NO | NO | NA |
| Dlc1 | ENSMUSG( | ENSMUSG( | 0.103416 | 0.834139 | NO | NO | NA |
| Cdc42ep2 | ENSMUSG( | ENSMUSG( | 0.625237 | 0.376668 | NO | NO | NA |
| Gfra2 | ENSMUSG( | ENSMUSG( | 0.546088 | 0.574644 | NO | NO | NA |
| Zbtb18 | ENSMUSG( | ENSMUSG( | 0.467976 | 0.036943 | YES | NO | NA |
| Rps18-ps6 | ENSMUSG( | ENSMUSG( | 0.635199 | 0.002354 | YES | NO | NA |
| Adam23 | ENSMUSG( | ENSMUSG( | 0.859572 | 0.422267 | NO | NO | NA |
| Scn3a | ENSMUSG( | ENSMUSG( | 0.593566 | 0.58339 | NO | NO | NA |
| Rimkla | ENSMUSG( | ENSMUSG( | -0.93413 | 0.000123 | YES | NO | NA |
| Prtn3 | ENSMUSG( | ENSMUSG( | 0.568731 | 0.522159 | NO | NO | NA |
| Mmp16 | ENSMUSG( | ENSMUSG( | 0.591273 | 0.572582 | NO | NO | NA |
| Gm9762 | ENSMUSG( | ENSMUSG( | -0.07115 | 0.826387 | NO | NO | NA |
| Gm50010 | ENSMUSG( | ENSMUSG( | -0.2254 | 0.236987 | NO | NO | NA |
| B630019A | ENSMUSG( | ENSMUSG( | 0.501818 | 0.618922 | NO | NO | NA |
| Gm47770 | ENSMUSG( | ENSMUSG( | 1.145512 | 0.01635 | YES | YES | Upregulated |
| Gm45853 | ENSMUSG( | ENSMUSG( | 0.040067 | 0.899688 | NO | NO | NA |
| Slc6a1 | ENSMUSG( | ENSMUSG( | 0.548931 | 0.596766 | NO | NO | NA |
| Lrp1 | ENSMUSG( | ENSMUSG( | -0.3318 | 0.51786 | NO | NO | NA |
| Ighv1-19 | ENSMUSG( | ENSMUSG( | 1.305541 | 0.604949 | NO | NO | NA |
| Gm26614 | ENSMUSG( | ENSMUSG( | -0.72463 | 0.032784 | YES | NO | NA |
| Uckl1os | ENSMUSG( | ENSMUSG( | -0.07922 | 0.778559 | NO | NO | NA |
| ENSMUSG | ENSMUSG( | ENSMUSG( | -0.34311 | 0.23041 | NO | NO | NA |
| Lrrtm4 | ENSMUSG( | ENSMUSG( | 0.29711 | 0.777582 | NO | NO | NA |
| Hgfac | ENSMUSG( | ENSMUSG( | -0.46619 | 0.086731 | NO | NO | NA |
| Mylk | ENSMUSG( | ENSMUSG( | 0.422599 | 0.677387 | NO | NO | NA |

| | | | | | | | |
|---|---|---|---|---|---|---|---|
| Adra1a | ENSMUSG( | ENSMUSG( | 0.363021 | 0.721225 | NO | NO | NA |
| 9230116N | ENSMUSG( | ENSMUSG( | 0.590363 | 0.021795 | YES | NO | NA |
| Gm13012 | ENSMUSG( | ENSMUSG( | -0.18351 | 0.460561 | NO | NO | NA |
| Wdr66 | ENSMUSG( | ENSMUSG( | 0.575718 | 0.378023 | NO | NO | NA |
| Rhpn2 | ENSMUSG( | ENSMUSG( | 0.54622 | 0.443505 | NO | NO | NA |
| Pnck | ENSMUSG( | ENSMUSG( | 1.173696 | 0.04245 | YES | YES | Upregulated |
| Dtx4 | ENSMUSG( | ENSMUSG( | 0.181251 | 0.63602 | NO | NO | NA |
| Gm17200 | ENSMUSG( | ENSMUSG( | 1.227004 | 0.00856 | YES | YES | Upregulated |
| ENSMUSG( | ENSMUSG( | ENSMUSG( | -0.52708 | 0.367986 | NO | NO | NA |
| Gm6180 | ENSMUSG( | ENSMUSG( | 0.271322 | 0.451766 | NO | NO | NA |
| Pfn4 | ENSMUSG( | ENSMUSG( | 0.704674 | 0.391427 | NO | NO | NA |
| ENSMUSG | ENSMUSG( | ENSMUSG( | -0.86708 | 0.02564 | YES | NO | NA |
| Gm34354 | ENSMUSG( | ENSMUSG( | 0.875993 | 0.434211 | NO | NO | NA |
| Gm2245 | ENSMUSG( | ENSMUSG( | 0.087535 | 0.847094 | NO | NO | NA |
| P3h4 | ENSMUSG( | ENSMUSG( | -3.43122 | 2.95E-13 | YES | YES | Downregulated |
| B230322F( | ENSMUSG( | ENSMUSG( | 0.635105 | 0.248585 | NO | NO | NA |
| Gm45110 | ENSMUSG( | ENSMUSG( | -0.50029 | 0.027328 | YES | NO | NA |
| Nxpe2 | ENSMUSG( | ENSMUSG( | 0.774379 | 0.231925 | NO | NO | NA |
| 4930518I1 | ENSMUSG( | ENSMUSG( | -0.084 | 0.825207 | NO | NO | NA |
| Mmp17 | ENSMUSG( | ENSMUSG( | 0.331842 | 0.368327 | NO | NO | NA |
| Sox6 | ENSMUSG( | ENSMUSG( | 0.455307 | 0.365952 | NO | NO | NA |
| Bco1 | ENSMUSG( | ENSMUSG( | 1.418579 | 0.000395 | YES | YES | Upregulated |
| Gm43099 | ENSMUSG( | ENSMUSG( | 0.88034 | 0.295853 | NO | NO | NA |
| C130071C( | ENSMUSG( | ENSMUSG( | 0.708682 | 0.353974 | NO | NO | NA |
| 4932442E( | ENSMUSG( | ENSMUSG( | -0.37135 | 0.092061 | NO | NO | NA |
| Trav2 | ENSMUSG( | ENSMUSG( | -1.74606 | 0.222025 | NO | NO | NA |
| Gm45819 | ENSMUSG( | ENSMUSG( | 0.320778 | 0.488115 | NO | NO | NA |
| Grin2d | ENSMUSG( | ENSMUSG( | -0.16732 | 0.526974 | NO | NO | NA |
| Igkv6-23 | ENSMUSG( | ENSMUSG( | 0.475444 | 0.838685 | NO | NO | NA |
| Gm2237 | ENSMUSG( | ENSMUSG( | -0.2338 | 0.244414 | NO | NO | NA |
| Tex50 | ENSMUSG( | ENSMUSG( | 1.335043 | 0.166103 | NO | NO | NA |
| Gm16059 | ENSMUSG( | ENSMUSG( | 0.341115 | 0.676705 | NO | NO | NA |
| B830012L1 | ENSMUSG( | ENSMUSG( | 0.367832 | 0.689161 | NO | NO | NA |
| Apba1 | ENSMUSG( | ENSMUSG( | 0.67498 | 0.396668 | NO | NO | NA |
| Unc5d | ENSMUSG( | ENSMUSG( | 0.880666 | 0.402404 | NO | NO | NA |
| Trpm5 | ENSMUSG( | ENSMUSG( | -2.34472 | NA | NO | NO | NA |
| Ighv5-9-1 | ENSMUSG( | ENSMUSG( | -0.07572 | NA | NO | NO | NA |
| Gm28778 | ENSMUSG( | ENSMUSG( | 1.062995 | 0.576702 | NO | NO | NA |
| Gm37563 | ENSMUSG( | ENSMUSG( | 0.396826 | 0.61095 | NO | NO | NA |
| 2700078F( | ENSMUSG( | ENSMUSG( | 0.733507 | 0.496053 | NO | NO | NA |
| Frmd5 | ENSMUSG( | ENSMUSG( | 0.61204 | 0.175959 | NO | NO | NA |
| Gm50107 | ENSMUSG( | ENSMUSG( | -0.60808 | 0.003211 | YES | NO | NA |
| Hhex | ENSMUSG( | ENSMUSG( | 0.526368 | 0.16358 | NO | NO | NA |
| Scmh1 | ENSMUSG( | ENSMUSG( | 0.286831 | 0.516667 | NO | NO | NA |
| Lima1 | ENSMUSG( | ENSMUSG( | 0.281044 | 0.584094 | NO | NO | NA |
| Lipt1 | ENSMUSG( | ENSMUSG( | 0.248712 | 0.362077 | NO | NO | NA |
| Olfr354 | ENSMUSG( | ENSMUSG( | 1.036884 | 0.334319 | NO | NO | NA |

| | | | | | | | |
|---|---|---|---|---|---|---|---|
| ENSMUSG( | ENSMUSG( | ENSMUSG( | 0.286665 | 0.181221 | NO | NO | NA |
| 4933406I1 | ENSMUSG( | ENSMUSG( | -0.07281 | 0.827284 | NO | NO | NA |
| 2810455O | ENSMUSG( | ENSMUSG( | -0.07286 | 0.819148 | NO | NO | NA |
| Ccl9 | ENSMUSG( | ENSMUSG( | -0.1301 | 0.778389 | NO | NO | NA |
| Cdh3 | ENSMUSG( | ENSMUSG( | 0.668772 | 0.493946 | NO | NO | NA |
| 1700105P( | ENSMUSG( | ENSMUSG( | -0.17372 | 0.617169 | NO | NO | NA |
| Xkr6 | ENSMUSG( | ENSMUSG( | -0.33478 | 0.223269 | NO | NO | NA |
| Ggt5 | ENSMUSG( | ENSMUSG( | 0.062677 | 0.908038 | NO | NO | NA |
| Zfp661 | ENSMUSG( | ENSMUSG( | 0.315947 | 0.244364 | NO | NO | NA |
| Rbpms | ENSMUSG( | ENSMUSG( | 0.940341 | 0.30111 | NO | NO | NA |
| Rps18-ps5 | ENSMUSG( | ENSMUSG( | 0.71892 | 0.006598 | YES | NO | NA |
| Gm47050 | ENSMUSG( | ENSMUSG( | 0.585404 | 0.428302 | NO | NO | NA |
| ENSMUSG | ENSMUSG( | ENSMUSG( | 0.0981 | 0.795982 | NO | NO | NA |
| 5330409N | ENSMUSG( | ENSMUSG( | 0.653907 | 0.350628 | NO | NO | NA |
| Gm37372 | ENSMUSG( | ENSMUSG( | 0.584584 | 0.575669 | NO | NO | NA |
| P3h1 | ENSMUSG( | ENSMUSG( | 0.293992 | 0.258267 | NO | NO | NA |
| Cryzl2 | ENSMUSG( | ENSMUSG( | 0.65847 | 0.015214 | YES | NO | NA |
| Gm16279 | ENSMUSG( | ENSMUSG( | 0.498843 | 0.015366 | YES | NO | NA |
| Tnfrsf26 | ENSMUSG( | ENSMUSG( | -0.82012 | 0.023809 | YES | NO | NA |
| Gm20632 | ENSMUSG( | ENSMUSG( | 0.424688 | 0.087308 | NO | NO | NA |
| Gm26707 | ENSMUSG( | ENSMUSG( | 1.325221 | 0.100971 | NO | NO | NA |
| Pde4c | ENSMUSG( | ENSMUSG( | 0.352301 | 0.165778 | NO | NO | NA |
| Gm5526 | ENSMUSG( | ENSMUSG( | 0.270856 | 0.566439 | NO | NO | NA |
| Il11ra1 | ENSMUSG( | ENSMUSG( | -0.65237 | 0.001577 | YES | NO | NA |
| Ccnjl | ENSMUSG( | ENSMUSG( | 1.023682 | 0.132842 | NO | NO | NA |
| ENSMUSG | ENSMUSG( | ENSMUSG( | -1.74428 | 1.31E-07 | YES | YES | Downregulated |
| Sdc4 | ENSMUSG( | ENSMUSG( | 0.640558 | 0.386239 | NO | NO | NA |
| Kcnc3 | ENSMUSG( | ENSMUSG( | 0.564441 | 0.135298 | NO | NO | NA |
| Tmem88b | ENSMUSG( | ENSMUSG( | 0.518421 | 0.574104 | NO | NO | NA |
| Opcml | ENSMUSG( | ENSMUSG( | 0.357782 | 0.733607 | NO | NO | NA |
| Gm17055 | ENSMUSG( | ENSMUSG( | 0.540559 | 0.054216 | NO | NO | NA |
| Gm37469 | ENSMUSG( | ENSMUSG( | -0.59436 | 0.057654 | NO | NO | NA |
| Gm32401 | ENSMUSG( | ENSMUSG( | -0.20159 | 0.301956 | NO | NO | NA |
| Gm43681 | ENSMUSG( | ENSMUSG( | -0.39228 | 0.2123 | NO | NO | NA |
| Gm49894 | ENSMUSG( | ENSMUSG( | -0.223 | 0.416423 | NO | NO | NA |
| Maff | ENSMUSG( | ENSMUSG( | 1.486233 | 0.000219 | YES | YES | Upregulated |
| Pou3f1 | ENSMUSG( | ENSMUSG( | 0.728516 | 0.358421 | NO | NO | NA |
| Gm15747 | ENSMUSG( | ENSMUSG( | -0.16513 | 0.5075 | NO | NO | NA |
| Gm20463 | ENSMUSG( | ENSMUSG( | -0.30954 | 0.221618 | NO | NO | NA |
| Gm11400 | ENSMUSG( | ENSMUSG( | -0.01291 | 0.964701 | NO | NO | NA |
| 4930556N | ENSMUSG( | ENSMUSG( | 0.559006 | 0.515217 | NO | NO | NA |
| Efnb2 | ENSMUSG( | ENSMUSG( | 0.54647 | 0.305474 | NO | NO | NA |
| Scn2b | ENSMUSG( | ENSMUSG( | 0.486961 | 0.621132 | NO | NO | NA |
| Gm40841 | ENSMUSG( | ENSMUSG( | 0.602317 | 0.519025 | NO | NO | NA |
| Gm42899 | ENSMUSG( | ENSMUSG( | 0.678585 | 0.522159 | NO | NO | NA |
| Zfp202 | ENSMUSG( | ENSMUSG( | 0.838971 | 0.000837 | YES | NO | NA |
| Cdh4 | ENSMUSG( | ENSMUSG( | 0.34953 | 0.725061 | NO | NO | NA |

| | | | | | | | |
|---|---|---|---|---|---|---|---|
| Magi1 | ENSMUSG( | ENSMUSG( | 0.473437 | 0.513868 | NO | NO | NA |
| Gm11696 | ENSMUSG( | ENSMUSG( | -0.09346 | 0.669989 | NO | NO | NA |
| L3mbtl4 | ENSMUSG( | ENSMUSG( | 0.302737 | 0.777275 | NO | NO | NA |
| Sh3d19 | ENSMUSG( | ENSMUSG( | -0.32807 | 0.380062 | NO | NO | NA |
| Gm28536 | ENSMUSG( | ENSMUSG( | 0.332913 | 0.537294 | NO | NO | NA |
| Gm37733 | ENSMUSG( | ENSMUSG( | 0.612952 | 0.385181 | NO | NO | NA |
| Efemp2 | ENSMUSG( | ENSMUSG( | -0.08664 | 0.796109 | NO | NO | NA |
| Igkv4-57 | ENSMUSG( | ENSMUSG( | 1.074852 | 0.762649 | NO | NO | NA |
| Gm23971 | ENSMUSG( | ENSMUSG( | 4.080885 | 0.320047 | NO | NO | NA |
| Gm47205 | ENSMUSG( | ENSMUSG( | -0.13992 | 0.714323 | NO | NO | NA |
| Dgkg | ENSMUSG( | ENSMUSG( | -0.44393 | 0.286304 | NO | NO | NA |
| Mdga2 | ENSMUSG( | ENSMUSG( | 0.625136 | 0.55607 | NO | NO | NA |
| Lhfpl4 | ENSMUSG( | ENSMUSG( | 0.650061 | 0.425871 | NO | NO | NA |
| Fbxl22 | ENSMUSG( | ENSMUSG( | -0.0304 | 0.932144 | NO | NO | NA |
| 4933431K: | ENSMUSG( | ENSMUSG( | 0.130551 | 0.635606 | NO | NO | NA |
| Sema6a | ENSMUSG( | ENSMUSG( | 0.773892 | 0.460305 | NO | NO | NA |
| E330013P( | ENSMUSG( | ENSMUSG( | 0.823291 | 0.308518 | NO | NO | NA |
| Abhd15 | ENSMUSG( | ENSMUSG( | 0.441705 | 0.283811 | NO | NO | NA |
| ENSMUSG( | ENSMUSG( | ENSMUSG( | -0.24202 | 0.37966 | NO | NO | NA |
| Crnde | ENSMUSG( | ENSMUSG( | 0.554759 | 0.1418 | NO | NO | NA |
| Gm13375 | ENSMUSG( | ENSMUSG( | 1.079008 | 5.02E-06 | YES | YES | Upregulated |
| Csnk2a1-p | ENSMUSG( | ENSMUSG( | 0.127258 | 0.779029 | NO | NO | NA |
| Angptl2 | ENSMUSG( | ENSMUSG( | -0.3423 | 0.220257 | NO | NO | NA |
| Klhl8 | ENSMUSG( | ENSMUSG( | -0.04283 | 0.890997 | NO | NO | NA |
| Pdzd2 | ENSMUSG( | ENSMUSG( | 0.285388 | 0.425956 | NO | NO | NA |
| Gm12500 | ENSMUSG( | ENSMUSG( | 0.668641 | 0.065091 | NO | NO | NA |
| ENSMUSG( | ENSMUSG( | ENSMUSG( | 0.483929 | 0.10461 | NO | NO | NA |
| A130014A | ENSMUSG( | ENSMUSG( | -0.00784 | 0.979731 | NO | NO | NA |
| Slain1os | ENSMUSG( | ENSMUSG( | 0.089278 | 0.733152 | NO | NO | NA |
| Ttc30a1 | ENSMUSG( | ENSMUSG( | 0.230185 | 0.318262 | NO | NO | NA |
| A830021F: | ENSMUSG( | ENSMUSG( | 0.606715 | 0.566253 | NO | NO | NA |
| Plag1 | ENSMUSG( | ENSMUSG( | 0.357302 | 0.172886 | NO | NO | NA |
| Gm44292 | ENSMUSG( | ENSMUSG( | -0.64534 | 0.00471 | YES | NO | NA |
| Rita1 | ENSMUSG( | ENSMUSG( | -0.0683 | 0.783586 | NO | NO | NA |
| Gm7353 | ENSMUSG( | ENSMUSG( | -0.08162 | 0.808331 | NO | NO | NA |
| Ncf2 | ENSMUSG( | ENSMUSG( | -0.14671 | 0.757619 | NO | NO | NA |
| Plau | ENSMUSG( | ENSMUSG( | 0.070415 | 0.842035 | NO | NO | NA |
| BC024063 | ENSMUSG( | ENSMUSG( | -0.28985 | 0.343183 | NO | NO | NA |
| Gm4707 | ENSMUSG( | ENSMUSG( | 0.128628 | 0.783951 | NO | NO | NA |
| Gm11767 | ENSMUSG( | ENSMUSG( | -0.0892 | 0.758216 | NO | NO | NA |
| Dnd1 | ENSMUSG( | ENSMUSG( | 0.302708 | 0.330198 | NO | NO | NA |
| Olfr1440 | ENSMUSG( | ENSMUSG( | 1.007434 | 0.312376 | NO | NO | NA |
| Gm43435 | ENSMUSG( | ENSMUSG( | 0.330392 | 0.465993 | NO | NO | NA |
| Tpmt | ENSMUSG( | ENSMUSG( | 0.31646 | 0.351405 | NO | NO | NA |
| Gm38393 | ENSMUSG( | ENSMUSG( | 0.301329 | 0.671878 | NO | NO | NA |
| Vcan | ENSMUSG( | ENSMUSG( | 0.239743 | 0.43166 | NO | NO | NA |
| A930004D | ENSMUSG( | ENSMUSG( | -0.36057 | 0.203617 | NO | NO | NA |

| | | | | | | | |
|---|---|---|---|---|---|---|---|
| Gm19721 | ENSMUSG( | ENSMUSG( | -0.22691 | 0.475161 | NO | NO | NA |
| Gm10053 | ENSMUSG( | ENSMUSG( | 0.472938 | 0.359082 | NO | NO | NA |
| Zfp937 | ENSMUSG( | ENSMUSG( | -0.63943 | 0.07659 | NO | NO | NA |
| Gm44280 | ENSMUSG( | ENSMUSG( | -0.17082 | 0.518069 | NO | NO | NA |
| ENSMUSG( | ENSMUSG( | ENSMUSG( | 0.951881 | 0.199708 | NO | NO | NA |
| Nup210l | ENSMUSG( | ENSMUSG( | 0.308007 | 0.273857 | NO | NO | NA |
| ENSMUSG | ENSMUSG( | ENSMUSG( | 0.689568 | 0.372394 | NO | NO | NA |
| Rnf152 | ENSMUSG( | ENSMUSG( | 0.181553 | 0.75961 | NO | NO | NA |
| Gm15728 | ENSMUSG( | ENSMUSG( | 0.855584 | 0.237115 | NO | NO | NA |
| Trav13d-1 | ENSMUSG( | ENSMUSG( | -1.21273 | 0.206369 | NO | NO | NA |
| A930002I2 | ENSMUSG( | ENSMUSG( | -1.38824 | 0.000135 | YES | YES | Downregulated |
| Gm10419 | ENSMUSG( | ENSMUSG( | 0.543568 | 0.518236 | NO | NO | NA |
| Npffr1 | ENSMUSG( | ENSMUSG( | 0.465274 | 0.636419 | NO | NO | NA |
| Grhl2 | ENSMUSG( | ENSMUSG( | 0.408162 | 0.536077 | NO | NO | NA |
| C230085N | ENSMUSG( | ENSMUSG( | 1.07408 | 0.092957 | NO | NO | NA |
| Rilpl1 | ENSMUSG( | ENSMUSG( | 0.549103 | 0.063113 | NO | NO | NA |
| Cacna1e | ENSMUSG( | ENSMUSG( | -0.10559 | 0.901248 | NO | NO | NA |
| Gm43457 | ENSMUSG( | ENSMUSG( | -0.11981 | 0.736541 | NO | NO | NA |
| Limch1 | ENSMUSG( | ENSMUSG( | 0.552911 | 0.462442 | NO | NO | NA |
| Gm37415 | ENSMUSG( | ENSMUSG( | 0.018254 | 0.952401 | NO | NO | NA |
| Vmn2r20 | ENSMUSG( | ENSMUSG( | 0.537435 | 0.595771 | NO | NO | NA |
| Gm42829 | ENSMUSG( | ENSMUSG( | 0.040267 | 0.864262 | NO | NO | NA |
| Thnsl1 | ENSMUSG( | ENSMUSG( | 0.195355 | 0.584776 | NO | NO | NA |
| Gm45112 | ENSMUSG( | ENSMUSG( | 0.814879 | 0.402332 | NO | NO | NA |
| Gm47550 | ENSMUSG( | ENSMUSG( | -0.23864 | 0.611409 | NO | NO | NA |
| Osbpl1a | ENSMUSG( | ENSMUSG( | 0.577812 | 0.024693 | YES | NO | NA |
| B930036N | ENSMUSG( | ENSMUSG( | 0.020649 | 0.963475 | NO | NO | NA |
| Chl1 | ENSMUSG( | ENSMUSG( | 1.120032 | 0.05347 | NO | NO | NA |
| Gm10825 | ENSMUSG( | ENSMUSG( | 0.373666 | 0.742075 | NO | NO | NA |
| Garem1 | ENSMUSG( | ENSMUSG( | 0.683087 | 0.262365 | NO | NO | NA |
| Cdr1os | ENSMUSG( | ENSMUSG( | 0.642964 | 0.53935 | NO | NO | NA |
| Gm26660 | ENSMUSG( | ENSMUSG( | -0.27004 | 0.402567 | NO | NO | NA |
| Trav21-dv: | ENSMUSG( | ENSMUSG( | -0.92715 | 0.221472 | NO | NO | NA |
| Gm32031 | ENSMUSG( | ENSMUSG( | 0.011606 | 0.969697 | NO | NO | NA |
| Zfp418 | ENSMUSG( | ENSMUSG( | -0.34064 | 0.194456 | NO | NO | NA |
| Slc41a3 | ENSMUSG( | ENSMUSG( | -0.32524 | 0.201631 | NO | NO | NA |
| Gm42670 | ENSMUSG( | ENSMUSG( | -0.4054 | 0.099567 | NO | NO | NA |
| Gm9616 | ENSMUSG( | ENSMUSG( | 0.389609 | 0.160903 | NO | NO | NA |
| Noxo1 | ENSMUSG( | ENSMUSG( | 0.228026 | 0.513234 | NO | NO | NA |
| Fst | ENSMUSG( | ENSMUSG( | 0.543531 | 0.595771 | NO | NO | NA |
| Cacna2d4 | ENSMUSG( | ENSMUSG( | -2.33309 | 2.94E-05 | YES | YES | Downregulated |
| Chadl | ENSMUSG( | ENSMUSG( | -0.06322 | 0.781878 | NO | NO | NA |
| Gm16580 | ENSMUSG( | ENSMUSG( | 0.335188 | 0.220868 | NO | NO | NA |
| Zfp941 | ENSMUSG( | ENSMUSG( | 1.300731 | 1.81E-05 | YES | YES | Upregulated |
| Cacna2d2 | ENSMUSG( | ENSMUSG( | 0.033159 | 0.910266 | NO | NO | NA |
| Gm37856 | ENSMUSG( | ENSMUSG( | -0.32846 | 0.121079 | NO | NO | NA |
| Mitf | ENSMUSG( | ENSMUSG( | 0.716625 | 0.116234 | NO | NO | NA |

| | | | | | | | |
|---|---|---|---|---|---|---|---|
| 6530409C: | ENSMUSG( | ENSMUSG( | -0.39986 | 0.121641 | NO | NO | NA |
| 3110006O | ENSMUSG( | ENSMUSG( | 0.914414 | 0.276223 | NO | NO | NA |
| D630023O | ENSMUSG( | ENSMUSG( | 0.746817 | 0.503771 | NO | NO | NA |
| Tnnt2 | ENSMUSG( | ENSMUSG( | 0.761077 | 0.43892 | NO | NO | NA |
| Gria2 | ENSMUSG( | ENSMUSG( | 0.540301 | 0.603823 | NO | NO | NA |
| Gstm1 | ENSMUSG( | ENSMUSG( | 1.344737 | 0.000134 | YES | YES | Upregulated |
| Flt3 | ENSMUSG( | ENSMUSG( | 0.674383 | 0.518553 | NO | NO | NA |
| 1700007L: | ENSMUSG( | ENSMUSG( | 0.551043 | 0.015241 | YES | NO | NA |
| Kcnt1 | ENSMUSG( | ENSMUSG( | 0.345266 | 0.653047 | NO | NO | NA |
| Gm5475 | ENSMUSG( | ENSMUSG( | 0.522373 | 0.535469 | NO | NO | NA |
| 1810059H | ENSMUSG( | ENSMUSG( | 0.991906 | 0.293239 | NO | NO | NA |
| Gm11100 | ENSMUSG( | ENSMUSG( | 1.02283 | 1.72E-05 | YES | YES | Upregulated |
| A930032L( | ENSMUSG( | ENSMUSG( | 1.250571 | 0.154131 | NO | NO | NA |
| A230056P | ENSMUSG( | ENSMUSG( | 0.003411 | 0.992393 | NO | NO | NA |
| H4c8 | ENSMUSG( | ENSMUSG( | 0.612489 | 0.047347 | YES | NO | NA |
| Vmn1r179 | ENSMUSG( | ENSMUSG( | 0.393732 | 0.721244 | NO | NO | NA |
| Homer3 | ENSMUSG( | ENSMUSG( | -0.5235 | 0.023729 | YES | NO | NA |
| Gm12183 | ENSMUSG( | ENSMUSG( | -0.07558 | 0.788009 | NO | NO | NA |
| B4galnt4 | ENSMUSG( | ENSMUSG( | 1.153698 | 0.000269 | YES | YES | Upregulated |
| Gm6344 | ENSMUSG( | ENSMUSG( | 0.538638 | 0.152477 | NO | NO | NA |
| Rps10-ps2 | ENSMUSG( | ENSMUSG( | 0.343694 | 0.123354 | NO | NO | NA |
| Atp8a2 | ENSMUSG( | ENSMUSG( | 0.119664 | 0.817741 | NO | NO | NA |
| Mapk10 | ENSMUSG( | ENSMUSG( | 1.012755 | 0.286596 | NO | NO | NA |
| Ctif | ENSMUSG( | ENSMUSG( | -0.09756 | 0.69006 | NO | NO | NA |
| Ighv1-26 | ENSMUSG( | ENSMUSG( | -2.08713 | 0.105675 | NO | NO | NA |
| Gm50478 | ENSMUSG( | ENSMUSG( | -1.18529 | 0.667285 | NO | NO | NA |
| Gm11537 | ENSMUSG( | ENSMUSG( | 0.275407 | 0.34766 | NO | NO | NA |
| Faim2 | ENSMUSG( | ENSMUSG( | 0.459041 | 0.668578 | NO | NO | NA |
| Gm9008 | ENSMUSG( | ENSMUSG( | 0.532898 | 0.596227 | NO | NO | NA |
| Lgi1 | ENSMUSG( | ENSMUSG( | 0.790521 | 0.459274 | NO | NO | NA |
| Nlrc4 | ENSMUSG( | ENSMUSG( | 0.698419 | 0.349893 | NO | NO | NA |
| 5830416I1 | ENSMUSG( | ENSMUSG( | -0.22217 | 0.773862 | NO | NO | NA |
| Afap1 | ENSMUSG( | ENSMUSG( | 0.594624 | 0.216218 | NO | NO | NA |
| Fam169a | ENSMUSG( | ENSMUSG( | 0.894822 | 0.019231 | YES | NO | NA |
| Cxxc4 | ENSMUSG( | ENSMUSG( | 0.511092 | 0.63616 | NO | NO | NA |
| Sox2ot | ENSMUSG( | ENSMUSG( | 0.933393 | 0.353871 | NO | NO | NA |
| Gm12070 | ENSMUSG( | ENSMUSG( | 0.607502 | 0.317403 | NO | NO | NA |
| Trav6-4 | ENSMUSG( | ENSMUSG( | -0.11123 | 0.918145 | NO | NO | NA |
| Rnf150 | ENSMUSG( | ENSMUSG( | 1.110072 | 0.254912 | NO | NO | NA |
| Gm43963 | ENSMUSG( | ENSMUSG( | -0.09581 | 0.722171 | NO | NO | NA |
| Gm38065 | ENSMUSG( | ENSMUSG( | 0.766788 | 0.473588 | NO | NO | NA |
| Chrnb1 | ENSMUSG( | ENSMUSG( | -0.61076 | 0.016997 | YES | NO | NA |
| Hyal1 | ENSMUSG( | ENSMUSG( | -0.07109 | 0.838522 | NO | NO | NA |
| Gm10286 | ENSMUSG( | ENSMUSG( | -0.06539 | 0.857655 | NO | NO | NA |
| Zfp455 | ENSMUSG( | ENSMUSG( | -0.46029 | 0.165575 | NO | NO | NA |
| Amd2 | ENSMUSG( | ENSMUSG( | 0.147449 | 0.80545 | NO | NO | NA |
| Gm15326 | ENSMUSG( | ENSMUSG( | -0.53498 | 0.01439 | YES | NO | NA |

| | | | | | | | |
|---|---|---|---|---|---|---|---|
| Ifitm2 | ENSMUSG( | ENSMUSG( | 1.354724 | 0.00555 | YES | YES | Upregulated |
| B130034C | ENSMUSG( | ENSMUSG( | 0.788476 | 0.044865 | YES | NO | NA |
| ENSMUSG | ENSMUSG( | ENSMUSG( | -0.5249 | 0.054642 | NO | NO | NA |
| Gm30025 | ENSMUSG( | ENSMUSG( | -0.11243 | 0.653625 | NO | NO | NA |
| Shf | ENSMUSG( | ENSMUSG( | -0.40809 | 0.194086 | NO | NO | NA |
| A230103J1 | ENSMUSG( | ENSMUSG( | -0.18006 | 0.429519 | NO | NO | NA |
| Tshz2 | ENSMUSG( | ENSMUSG( | 0.813137 | 0.120489 | NO | NO | NA |
| Mpv17l | ENSMUSG( | ENSMUSG( | 0.706015 | 0.022793 | YES | NO | NA |
| Lix1l | ENSMUSG( | ENSMUSG( | 0.211448 | 0.409568 | NO | NO | NA |
| Trav6-5 | ENSMUSG( | ENSMUSG( | -0.60649 | 0.120881 | NO | NO | NA |
| Ntf5 | ENSMUSG( | ENSMUSG( | -0.08615 | 0.914876 | NO | NO | NA |
| Gm37637 | ENSMUSG( | ENSMUSG( | 0.158938 | 0.617531 | NO | NO | NA |
| Hoxb2 | ENSMUSG( | ENSMUSG( | 0.520685 | 0.611479 | NO | NO | NA |
| Lratd1 | ENSMUSG( | ENSMUSG( | 0.24991 | 0.761446 | NO | NO | NA |
| Tnfrsf22 | ENSMUSG( | ENSMUSG( | 0.286278 | 0.349342 | NO | NO | NA |
| Chaer1 | ENSMUSG( | ENSMUSG( | 0.594574 | 0.079022 | NO | NO | NA |
| Gm44209 | ENSMUSG( | ENSMUSG( | 0.621162 | 0.543583 | NO | NO | NA |
| Ap1m2 | ENSMUSG( | ENSMUSG( | 0.064135 | 0.881357 | NO | NO | NA |
| Irs2 | ENSMUSG( | ENSMUSG( | 0.684803 | 0.074371 | NO | NO | NA |
| Slc1a4 | ENSMUSG( | ENSMUSG( | 1.358381 | 1.16E-05 | YES | YES | Upregulated |
| Gm27176 | ENSMUSG( | ENSMUSG( | -0.23863 | 0.923891 | NO | NO | NA |
| Gm28513 | ENSMUSG( | ENSMUSG( | 0.37372 | 0.427511 | NO | NO | NA |
| Gm43323 | ENSMUSG( | ENSMUSG( | 0.716927 | 0.171547 | NO | NO | NA |
| Zfp667 | ENSMUSG( | ENSMUSG( | -0.15546 | 0.540932 | NO | NO | NA |
| Gm37625 | ENSMUSG( | ENSMUSG( | 0.380126 | 0.660613 | NO | NO | NA |
| Gm43876 | ENSMUSG( | ENSMUSG( | 0.715626 | 0.513114 | NO | NO | NA |
| A230072E: | ENSMUSG( | ENSMUSG( | 0.195648 | 0.315344 | NO | NO | NA |
| Gm42783 | ENSMUSG( | ENSMUSG( | -0.26798 | 0.344431 | NO | NO | NA |
| Sh3yl1 | ENSMUSG( | ENSMUSG( | -0.57893 | 0.05711 | NO | NO | NA |
| 9530018F( | ENSMUSG( | ENSMUSG( | 0.189964 | 0.560577 | NO | NO | NA |
| Fras1 | ENSMUSG( | ENSMUSG( | 0.424147 | 0.69475 | NO | NO | NA |
| Sphk1 | ENSMUSG( | ENSMUSG( | 0.399448 | 0.565868 | NO | NO | NA |
| Arpp21 | ENSMUSG( | ENSMUSG( | 0.521304 | 0.59374 | NO | NO | NA |
| ENSMUSG | ENSMUSG( | ENSMUSG( | 1.050142 | 0.169199 | NO | NO | NA |
| Dusp18 | ENSMUSG( | ENSMUSG( | 0.622013 | 0.151471 | NO | NO | NA |
| Nkd2 | ENSMUSG( | ENSMUSG( | 0.696387 | 0.048201 | YES | NO | NA |
| Gm20324 | ENSMUSG( | ENSMUSG( | 0.282597 | 0.166837 | NO | NO | NA |
| Tfap2a | ENSMUSG( | ENSMUSG( | 0.612387 | 0.211633 | NO | NO | NA |
| Camk2n2 | ENSMUSG( | ENSMUSG( | 0.62962 | 0.017066 | YES | NO | NA |
| Gm33111 | ENSMUSG( | ENSMUSG( | 0.312758 | 0.772033 | NO | NO | NA |
| Trav16d-d | ENSMUSG( | ENSMUSG( | -0.43125 | 0.51087 | NO | NO | NA |
| Pon3 | ENSMUSG( | ENSMUSG( | -0.5102 | 0.136481 | NO | NO | NA |
| Rpsa-ps2 | ENSMUSG( | ENSMUSG( | 0.485615 | 0.152586 | NO | NO | NA |
| Ctsg | ENSMUSG( | ENSMUSG( | 1.922482 | 0.032112 | YES | YES | Upregulated |
| Zfp759 | ENSMUSG( | ENSMUSG( | -0.30645 | 0.253253 | NO | NO | NA |
| 4933431E: | ENSMUSG( | ENSMUSG( | 0.961202 | 0.031357 | YES | NO | NA |
| Qser1 | ENSMUSG( | ENSMUSG( | 1.499121 | 1.22E-06 | YES | YES | Upregulated |

| | | | | | | | |
|---|---|---|---|---|---|---|---|
| Gm29438 | ENSMUSG( | ENSMUSG( | 0.882203 | 0.468739 | NO | NO | NA |
| 4930565N | ENSMUSG( | ENSMUSG( | -0.37915 | 0.18881 | NO | NO | NA |
| 9130230L2 | ENSMUSG( | ENSMUSG( | 0.287917 | 0.620133 | NO | NO | NA |
| Gm49172 | ENSMUSG( | ENSMUSG( | -0.17117 | 0.485084 | NO | NO | NA |
| Cnih2 | ENSMUSG( | ENSMUSG( | 0.200909 | 0.387383 | NO | NO | NA |
| Ttc41 | ENSMUSG( | ENSMUSG( | 0.093443 | 0.713191 | NO | NO | NA |
| Rgs6 | ENSMUSG( | ENSMUSG( | 0.517823 | 0.626606 | NO | NO | NA |
| Vwa5b2 | ENSMUSG( | ENSMUSG( | 0.839086 | 0.28599 | NO | NO | NA |
| Capn11 | ENSMUSG( | ENSMUSG( | -0.43213 | 0.803915 | NO | NO | NA |
| Nicn1 | ENSMUSG( | ENSMUSG( | 0.559083 | 0.116016 | NO | NO | NA |
| Spry4 | ENSMUSG( | ENSMUSG( | 0.457998 | 0.625041 | NO | NO | NA |
| Ighv5-4 | ENSMUSG( | ENSMUSG( | -7.83797 | 4.93E-07 | YES | YES | Downregulated |
| Gm16034 | ENSMUSG( | ENSMUSG( | 0.517141 | 0.259206 | NO | NO | NA |
| Robo1 | ENSMUSG( | ENSMUSG( | 0.398946 | 0.718279 | NO | NO | NA |
| C1ql3 | ENSMUSG( | ENSMUSG( | 0.818943 | 0.462135 | NO | NO | NA |
| 1700021J( | ENSMUSG( | ENSMUSG( | 0.072832 | 0.812244 | NO | NO | NA |
| Flrt1 | ENSMUSG( | ENSMUSG( | 0.719707 | 0.343016 | NO | NO | NA |
| Ltb4r2 | ENSMUSG( | ENSMUSG( | 0.505902 | 0.596362 | NO | NO | NA |
| Gm26569 | ENSMUSG( | ENSMUSG( | 0.716868 | 0.474914 | NO | NO | NA |
| Gm38384 | ENSMUSG( | ENSMUSG( | -0.32533 | 0.211846 | NO | NO | NA |
| Gm43549 | ENSMUSG( | ENSMUSG( | 0.527174 | 0.474192 | NO | NO | NA |
| Kpna2-ps | ENSMUSG( | ENSMUSG( | -0.1522 | 0.493081 | NO | NO | NA |
| Dbpht2 | ENSMUSG( | ENSMUSG( | 0.617454 | 0.556665 | NO | NO | NA |
| Hspa1l | ENSMUSG( | ENSMUSG( | 0.857386 | 0.000342 | YES | NO | NA |
| Renbp | ENSMUSG( | ENSMUSG( | -0.01734 | 0.937388 | NO | NO | NA |
| Zbed5 | ENSMUSG( | ENSMUSG( | 0.500099 | 0.254495 | NO | NO | NA |
| Gzmc | ENSMUSG( | ENSMUSG( | 0.926903 | 0.039299 | YES | NO | NA |
| Hpgds | ENSMUSG( | ENSMUSG( | 0.332282 | 0.576558 | NO | NO | NA |
| Gm47259 | ENSMUSG( | ENSMUSG( | 0.624143 | 0.556504 | NO | NO | NA |
| Gm9385 | ENSMUSG( | ENSMUSG( | 0.351573 | 0.145358 | NO | NO | NA |
| Pwwp2b | ENSMUSG( | ENSMUSG( | -0.8481 | 0.001958 | YES | NO | NA |
| Slc17a6 | ENSMUSG( | ENSMUSG( | 2.008453 | 1.17E-05 | YES | YES | Upregulated |
| Gm20628 | ENSMUSG( | ENSMUSG( | 0.387272 | 0.291954 | NO | NO | NA |
| Zfp651 | ENSMUSG( | ENSMUSG( | 0.280924 | 0.640929 | NO | NO | NA |
| Zbed4-ps1 | ENSMUSG( | ENSMUSG( | 0.777905 | 0.471322 | NO | NO | NA |
| Gria3 | ENSMUSG( | ENSMUSG( | 0.55335 | 0.540236 | NO | NO | NA |
| ENSMUSG | ENSMUSG( | ENSMUSG( | -0.14154 | 0.685406 | NO | NO | NA |
| Gm12472 | ENSMUSG( | ENSMUSG( | 0.811381 | 0.446439 | NO | NO | NA |
| Gm44183 | ENSMUSG( | ENSMUSG( | -0.66196 | 0.842522 | NO | NO | NA |
| Cacnb4 | ENSMUSG( | ENSMUSG( | -0.34141 | 0.235859 | NO | NO | NA |
| Paqr8 | ENSMUSG( | ENSMUSG( | 0.115405 | 0.720005 | NO | NO | NA |
| Mpc1-ps | ENSMUSG( | ENSMUSG( | 0.642276 | 0.053389 | NO | NO | NA |
| Gm14696 | ENSMUSG( | ENSMUSG( | 0.651292 | 0.569784 | NO | NO | NA |
| Tmem147( | ENSMUSG( | ENSMUSG( | -0.12307 | 0.74256 | NO | NO | NA |
| Srd5a1 | ENSMUSG( | ENSMUSG( | 0.288192 | 0.241715 | NO | NO | NA |
| Cebpd | ENSMUSG( | ENSMUSG( | 0.148616 | 0.516453 | NO | NO | NA |
| Grip1 | ENSMUSG( | ENSMUSG( | 0.419098 | 0.582357 | NO | NO | NA |

| | | | | | | | |
|---|---|---|---|---|---|---|---|
| Adamts5 | ENSMUSG( | ENSMUSG( | 0.800575 | 0.446295 | NO | NO | NA |
| Pbx1 | ENSMUSG( | ENSMUSG( | 0.410898 | 0.606581 | NO | NO | NA |
| Tymp | ENSMUSG( | ENSMUSG( | 0.442324 | 0.288828 | NO | NO | NA |
| ENSMUSG | ENSMUSG( | ENSMUSG( | -0.01505 | 0.965074 | NO | NO | NA |
| Basp1 | ENSMUSG( | ENSMUSG( | -0.01428 | 0.983122 | NO | NO | NA |
| Utf1 | ENSMUSG( | ENSMUSG( | 0.753474 | 0.006837 | YES | NO | NA |
| Gm2004 | ENSMUSG( | ENSMUSG( | 1.799955 | 0.205329 | NO | NO | NA |
| 1700102H | ENSMUSG( | ENSMUSG( | 0.591536 | 0.105599 | NO | NO | NA |
| Abhd1 | ENSMUSG( | ENSMUSG( | -0.33813 | 0.154667 | NO | NO | NA |
| Ampd3 | ENSMUSG( | ENSMUSG( | 0.812164 | 0.05174 | NO | NO | NA |
| Asic1 | ENSMUSG( | ENSMUSG( | -0.34049 | 0.147273 | NO | NO | NA |
| Elavl2 | ENSMUSG( | ENSMUSG( | 0.884193 | 0.175694 | NO | NO | NA |
| Gm37499 | ENSMUSG( | ENSMUSG( | 0.667919 | 0.416429 | NO | NO | NA |
| Zfp109 | ENSMUSG( | ENSMUSG( | 0.080766 | 0.710991 | NO | NO | NA |
| Gm3764 | ENSMUSG( | ENSMUSG( | 0.49238 | 0.639774 | NO | NO | NA |
| Neurod2 | ENSMUSG( | ENSMUSG( | 0.317514 | 0.736093 | NO | NO | NA |
| B4galt6 | ENSMUSG( | ENSMUSG( | 0.522648 | 0.217409 | NO | NO | NA |
| ENSMUSG( | ENSMUSG( | ENSMUSG( | 0.121617 | 0.893666 | NO | NO | NA |
| Gm43773 | ENSMUSG( | ENSMUSG( | 0.523813 | 0.599736 | NO | NO | NA |
| Gm38346 | ENSMUSG( | ENSMUSG( | -0.18047 | 0.572839 | NO | NO | NA |
| Gm26642 | ENSMUSG( | ENSMUSG( | 0.271848 | 0.446235 | NO | NO | NA |
| Pdgfa | ENSMUSG( | ENSMUSG( | 0.659101 | 0.531368 | NO | NO | NA |
| Utp14b | ENSMUSG( | ENSMUSG( | 0.835922 | 0.167246 | NO | NO | NA |
| Polr1has | ENSMUSG( | ENSMUSG( | 0.471445 | 0.045793 | YES | NO | NA |
| ENSMUSG( | ENSMUSG( | ENSMUSG( | 0.065129 | 0.837594 | NO | NO | NA |
| Caln1 | ENSMUSG( | ENSMUSG( | 0.894357 | 0.369 | NO | NO | NA |
| Trpm2 | ENSMUSG( | ENSMUSG( | 0.196316 | 0.789871 | NO | NO | NA |
| Foxb1 | ENSMUSG( | ENSMUSG( | 0.18451 | 0.87728 | NO | NO | NA |
| Arsg | ENSMUSG( | ENSMUSG( | 0.489579 | 0.046792 | YES | NO | NA |
| Ttc23 | ENSMUSG( | ENSMUSG( | 0.36781 | 0.431957 | NO | NO | NA |
| Rps18-ps4 | ENSMUSG( | ENSMUSG( | 1.004765 | 0.090892 | NO | NO | NA |
| Dnase1l2 | ENSMUSG( | ENSMUSG( | -0.19308 | 0.440501 | NO | NO | NA |
| Cttn | ENSMUSG( | ENSMUSG( | 0.747586 | 0.186334 | NO | NO | NA |
| Gm43667 | ENSMUSG( | ENSMUSG( | -0.01638 | 0.965301 | NO | NO | NA |
| Stc1 | ENSMUSG( | ENSMUSG( | 0.479011 | 0.646357 | NO | NO | NA |
| Nos1ap | ENSMUSG( | ENSMUSG( | 0.607833 | 0.440958 | NO | NO | NA |
| Gm36371 | ENSMUSG( | ENSMUSG( | 0.599475 | 0.262228 | NO | NO | NA |
| Nol4l | ENSMUSG( | ENSMUSG( | -0.21246 | 0.709803 | NO | NO | NA |
| Gm44005 | ENSMUSG( | ENSMUSG( | 0.870521 | 0.336832 | NO | NO | NA |
| Dmrt2 | ENSMUSG( | ENSMUSG( | 0.850986 | 0.090892 | NO | NO | NA |
| Tmem181 | ENSMUSG( | ENSMUSG( | 2.093878 | 6.43E-08 | YES | YES | Upregulated |
| Shroom3 | ENSMUSG( | ENSMUSG( | 0.482227 | 0.645205 | NO | NO | NA |
| B3galt5 | ENSMUSG( | ENSMUSG( | 0.475299 | 0.080621 | NO | NO | NA |
| Gm43126 | ENSMUSG( | ENSMUSG( | -0.51811 | 0.084679 | NO | NO | NA |
| Tmem240 | ENSMUSG( | ENSMUSG( | 0.041985 | 0.885636 | NO | NO | NA |
| Gm10036 | ENSMUSG( | ENSMUSG( | 0.155667 | 0.540236 | NO | NO | NA |
| Gm26532 | ENSMUSG( | ENSMUSG( | 0.472989 | 0.189919 | NO | NO | NA |

| | | | | | | | |
|---|---|---|---|---|---|---|---|
| C230037L1 | ENSMUSG0 | ENSMUSG0 | -0.29347 | 0.281142 | NO | NO | NA |
| Gm43127 | ENSMUSG0 | ENSMUSG0 | -0.4836 | 0.179544 | NO | NO | NA |
| Gm15559 | ENSMUSG0 | ENSMUSG0 | 0.286954 | 0.361047 | NO | NO | NA |
| Gm15710 | ENSMUSG0 | ENSMUSG0 | 0.612466 | 0.071602 | NO | NO | NA |
| Cd300a | ENSMUSG0 | ENSMUSG0 | 0.12863 | 0.865286 | NO | NO | NA |
| ENSMUSG | ENSMUSG0 | ENSMUSG0 | 0.197683 | 0.565732 | NO | NO | NA |
| Loxl2 | ENSMUSG0 | ENSMUSG0 | 0.818685 | 0.147429 | NO | NO | NA |
| C230053D | ENSMUSG0 | ENSMUSG0 | 0.413784 | 0.546654 | NO | NO | NA |
| Neu4 | ENSMUSG0 | ENSMUSG0 | 0.480312 | 0.476831 | NO | NO | NA |
| Prr36 | ENSMUSG0 | ENSMUSG0 | 0.302172 | 0.234862 | NO | NO | NA |
| Rtl10 | ENSMUSG0 | ENSMUSG0 | -0.23315 | 0.518405 | NO | NO | NA |
| 1810062O | ENSMUSG0 | ENSMUSG0 | 0.002032 | 0.993761 | NO | NO | NA |
| Syna | ENSMUSG0 | ENSMUSG0 | 1.947426 | 0.003734 | YES | YES | Upregulated |
| H2bc14 | ENSMUSG0 | ENSMUSG0 | 0.358044 | 0.218173 | NO | NO | NA |
| Prss34 | ENSMUSG0 | ENSMUSG0 | -0.63264 | 0.74342 | NO | NO | NA |
| Nphp1 | ENSMUSG0 | ENSMUSG0 | 2.088708 | 0.001041 | YES | YES | Upregulated |
| Wdr31 | ENSMUSG0 | ENSMUSG0 | -0.1722 | 0.534088 | NO | NO | NA |
| ENSMUSG | ENSMUSG0 | ENSMUSG0 | 0.32667 | 0.35551 | NO | NO | NA |
| Gm48478 | ENSMUSG0 | ENSMUSG0 | -0.23015 | 0.361309 | NO | NO | NA |
| Gm19705 | ENSMUSG0 | ENSMUSG0 | -0.37109 | 0.141618 | NO | NO | NA |
| Trav13n-1 | ENSMUSG0 | ENSMUSG0 | -1.6108 | 0.012764 | YES | YES | Downregulated |
| Lgr4 | ENSMUSG0 | ENSMUSG0 | 0.963163 | 0.119971 | NO | NO | NA |
| Tiam2 | ENSMUSG0 | ENSMUSG0 | -0.45665 | 0.177177 | NO | NO | NA |
| Gm16534 | ENSMUSG0 | ENSMUSG0 | -0.26565 | 0.625598 | NO | NO | NA |
| Acot6 | ENSMUSG0 | ENSMUSG0 | 0.914856 | 0.003136 | YES | NO | NA |
| Gm21981 | ENSMUSG0 | ENSMUSG0 | 0.270161 | 0.590346 | NO | NO | NA |
| 9930017N | ENSMUSG0 | ENSMUSG0 | -0.06969 | 0.796122 | NO | NO | NA |
| Gm42979 | ENSMUSG0 | ENSMUSG0 | -0.38658 | 0.034574 | YES | NO | NA |
| Trav9n-2 | ENSMUSG0 | ENSMUSG0 | 3.081138 | 0.031974 | YES | YES | Upregulated |
| 9430034N | ENSMUSG0 | ENSMUSG0 | -0.33575 | 0.232842 | NO | NO | NA |
| Ptk2 | ENSMUSG0 | ENSMUSG0 | 0.33455 | 0.642044 | NO | NO | NA |
| Coch | ENSMUSG0 | ENSMUSG0 | -0.5638 | 0.390459 | NO | NO | NA |
| Etv4 | ENSMUSG0 | ENSMUSG0 | 1.332429 | 0.000524 | YES | YES | Upregulated |
| Olfr344 | ENSMUSG0 | ENSMUSG0 | 0.513613 | 0.627668 | NO | NO | NA |
| 4930578M | ENSMUSG0 | ENSMUSG0 | 0.686863 | 0.336369 | NO | NO | NA |
| Lrrc27 | ENSMUSG0 | ENSMUSG0 | 0.577343 | 0.005838 | YES | NO | NA |
| Gm36298 | ENSMUSG0 | ENSMUSG0 | 0.06556 | 0.811161 | NO | NO | NA |
| Ighv1-64 | ENSMUSG0 | ENSMUSG0 | 0.166776 | 0.942217 | NO | NO | NA |
| Gm50253 | ENSMUSG0 | ENSMUSG0 | 0.408696 | 0.492958 | NO | NO | NA |
| Gm9493 | ENSMUSG0 | ENSMUSG0 | 0.245531 | 0.349385 | NO | NO | NA |
| Catsper2 | ENSMUSG0 | ENSMUSG0 | 0.222896 | 0.556504 | NO | NO | NA |
| Pgbd5 | ENSMUSG0 | ENSMUSG0 | 0.631113 | 0.562185 | NO | NO | NA |
| Crispld2 | ENSMUSG0 | ENSMUSG0 | 0.467039 | 0.619035 | NO | NO | NA |
| Etnk2 | ENSMUSG0 | ENSMUSG0 | 0.104987 | 0.924979 | NO | NO | NA |
| Gm20708 | ENSMUSG0 | ENSMUSG0 | -0.25355 | 0.62138 | NO | NO | NA |
| Gm42885 | ENSMUSG0 | ENSMUSG0 | 0.825785 | 0.449679 | NO | NO | NA |
| Rnf207 | ENSMUSG0 | ENSMUSG0 | 0.320289 | 0.726189 | NO | NO | NA |

| | | | | | | | |
|---|---|---|---|---|---|---|---|
| Mia | ENSMUSG( | ENSMUSG( | 0.876778 | 0.376339 | NO | NO | NA |
| Vmn1r76 | ENSMUSG( | ENSMUSG( | 0.465298 | 0.6715 | NO | NO | NA |
| Gm43910 | ENSMUSG( | ENSMUSG( | -0.281 | 0.384629 | NO | NO | NA |
| ENSMUSG( | ENSMUSG( | ENSMUSG( | 0.581659 | 0.136481 | NO | NO | NA |
| Hpse2 | ENSMUSG( | ENSMUSG( | 0.487774 | 0.635603 | NO | NO | NA |
| Epn2 | ENSMUSG( | ENSMUSG( | 0.824161 | 0.006182 | YES | NO | NA |
| Gm26873 | ENSMUSG( | ENSMUSG( | -0.02185 | 0.95455 | NO | NO | NA |
| Gm20621 | ENSMUSG( | ENSMUSG( | -0.29557 | 0.43691 | NO | NO | NA |
| Gm20695 | ENSMUSG( | ENSMUSG( | 0.084761 | 0.781352 | NO | NO | NA |
| Plekha7 | ENSMUSG( | ENSMUSG( | -0.27683 | 0.31733 | NO | NO | NA |
| Slc46a2 | ENSMUSG( | ENSMUSG( | 0.556409 | 0.604389 | NO | NO | NA |
| Gm7027 | ENSMUSG( | ENSMUSG( | -0.33132 | 0.088306 | NO | NO | NA |
| Slc25a47 | ENSMUSG( | ENSMUSG( | 0.483948 | 0.112117 | NO | NO | NA |
| Sdk2 | ENSMUSG( | ENSMUSG( | 0.654333 | 0.473973 | NO | NO | NA |
| C130080G | ENSMUSG( | ENSMUSG( | 0.851158 | 0.447567 | NO | NO | NA |
| Itgb4 | ENSMUSG( | ENSMUSG( | -0.74894 | 0.012214 | YES | NO | NA |
| Dnajc6 | ENSMUSG( | ENSMUSG( | 0.957332 | 0.003475 | YES | NO | NA |
| Gm37738 | ENSMUSG( | ENSMUSG( | 0.425028 | 0.521485 | NO | NO | NA |
| A930029G | ENSMUSG( | ENSMUSG( | -0.23075 | 0.580075 | NO | NO | NA |
| Gm37183 | ENSMUSG( | ENSMUSG( | 0.864775 | 0.312343 | NO | NO | NA |
| ENSMUSG | ENSMUSG( | ENSMUSG( | 0.589791 | 0.578898 | NO | NO | NA |
| Fgf22 | ENSMUSG( | ENSMUSG( | 0.65546 | 0.382461 | NO | NO | NA |
| ENSMUSG | ENSMUSG( | ENSMUSG( | -0.89983 | 2.56E-05 | YES | NO | NA |
| Ugt1a7c | ENSMUSG( | ENSMUSG( | 0.358603 | 0.249328 | NO | NO | NA |
| Gm19345 | ENSMUSG( | ENSMUSG( | 0.646344 | 0.075631 | NO | NO | NA |
| A230072C | ENSMUSG( | ENSMUSG( | -0.49563 | 0.19358 | NO | NO | NA |
| ENSMUSG | ENSMUSG( | ENSMUSG( | 0.861134 | 0.143357 | NO | NO | NA |
| Tmem255a | ENSMUSG( | ENSMUSG( | -0.30303 | 0.361782 | NO | NO | NA |
| Arhgap6 | ENSMUSG( | ENSMUSG( | 0.597099 | 0.41242 | NO | NO | NA |
| Napb | ENSMUSG( | ENSMUSG( | 0.294142 | 0.72652 | NO | NO | NA |
| Lrrc39 | ENSMUSG( | ENSMUSG( | 0.230178 | 0.451447 | NO | NO | NA |
| Lhx1 | ENSMUSG( | ENSMUSG( | 1.141865 | 0.176201 | NO | NO | NA |
| Ncam2 | ENSMUSG( | ENSMUSG( | 0.909713 | 0.283699 | NO | NO | NA |
| Syt7 | ENSMUSG( | ENSMUSG( | 0.646256 | 0.479548 | NO | NO | NA |
| Gspt2 | ENSMUSG( | ENSMUSG( | 1.727631 | 3.60E-06 | YES | YES | Upregulated |
| Vmn1r86 | ENSMUSG( | ENSMUSG( | 0.804104 | 0.426541 | NO | NO | NA |
| Clec10a | ENSMUSG( | ENSMUSG( | 0.945735 | 0.049242 | YES | NO | NA |
| Tmem14a | ENSMUSG( | ENSMUSG( | -0.13576 | 0.561259 | NO | NO | NA |
| Igkv15-103 | ENSMUSG( | ENSMUSG( | -0.39021 | 0.904271 | NO | NO | NA |
| 2610306M | ENSMUSG( | ENSMUSG( | -0.04099 | 0.894226 | NO | NO | NA |
| 9430064I2 | ENSMUSG( | ENSMUSG( | 0.460209 | 0.349254 | NO | NO | NA |
| Igkv3-4 | ENSMUSG( | ENSMUSG( | -3.42173 | 0.003275 | YES | YES | Downregulated |
| Rpl36a-ps2 | ENSMUSG( | ENSMUSG( | 0.734234 | 0.001172 | YES | NO | NA |
| Gm45191 | ENSMUSG( | ENSMUSG( | 0.079307 | 0.889049 | NO | NO | NA |
| Zfyve21 | ENSMUSG( | ENSMUSG( | 0.196315 | 0.527811 | NO | NO | NA |
| Gm28557 | ENSMUSG( | ENSMUSG( | -1.62971 | 5.06E-09 | YES | YES | Downregulated |
| Gm48502 | ENSMUSG( | ENSMUSG( | 0.821934 | 0.424153 | NO | NO | NA |

| | | | | | | | |
|---|---|---|---|---|---|---|---|
| Gm17638 | ENSMUSG( | ENSMUSG( | -0.11584 | 0.643741 | NO | NO | NA |
| Ppp1r13l | ENSMUSG( | ENSMUSG( | 0.432341 | 0.052329 | NO | NO | NA |
| 4930528J1 | ENSMUSG( | ENSMUSG( | 0.759168 | 0.477403 | NO | NO | NA |
| Gm11373 | ENSMUSG( | ENSMUSG( | 1.141529 | 0.152477 | NO | NO | NA |
| Barhl2 | ENSMUSG( | ENSMUSG( | 0.626421 | 0.533864 | NO | NO | NA |
| A330023F: | ENSMUSG( | ENSMUSG( | -0.34446 | 0.271378 | NO | NO | NA |
| Satb2 | ENSMUSG( | ENSMUSG( | 0.645101 | 0.25341 | NO | NO | NA |
| Pcdh10 | ENSMUSG( | ENSMUSG( | 0.818923 | 0.387383 | NO | NO | NA |
| Vmn2r26 | ENSMUSG( | ENSMUSG( | 0.562554 | 0.600691 | NO | NO | NA |
| Bhlhe41 | ENSMUSG( | ENSMUSG( | -0.04003 | 0.941527 | NO | NO | NA |
| Gm43029 | ENSMUSG( | ENSMUSG( | -0.25604 | 0.338282 | NO | NO | NA |
| Vmn1r80 | ENSMUSG( | ENSMUSG( | 0.702707 | 0.521665 | NO | NO | NA |
| Fkbpl | ENSMUSG( | ENSMUSG( | -0.08826 | 0.764367 | NO | NO | NA |
| ENSMUSG | ENSMUSG( | ENSMUSG( | -0.38578 | 0.142045 | NO | NO | NA |
| Gm13724 | ENSMUSG( | ENSMUSG( | 0.851945 | 0.422508 | NO | NO | NA |
| Vmn2r124 | ENSMUSG( | ENSMUSG( | 0.434237 | 0.618191 | NO | NO | NA |
| Inhba | ENSMUSG( | ENSMUSG( | 0.489106 | 0.5876 | NO | NO | NA |
| Igkv1-110 | ENSMUSG( | ENSMUSG( | -1.50845 | 0.335786 | NO | NO | NA |
| Cdh8 | ENSMUSG( | ENSMUSG( | 0.740097 | 0.527765 | NO | NO | NA |
| Gm13961 | ENSMUSG( | ENSMUSG( | 0.725051 | 0.454181 | NO | NO | NA |
| Tmem72 | ENSMUSG( | ENSMUSG( | -0.11403 | 0.73647 | NO | NO | NA |
| Tmem176a | ENSMUSG( | ENSMUSG( | 0.728754 | 0.027034 | YES | NO | NA |
| Gnaz | ENSMUSG( | ENSMUSG( | 0.587903 | 0.553596 | NO | NO | NA |
| Sirpa | ENSMUSG( | ENSMUSG( | -0.55265 | 0.260702 | NO | NO | NA |
| Fstl5 | ENSMUSG( | ENSMUSG( | 0.707814 | 0.516457 | NO | NO | NA |
| Gm16638 | ENSMUSG( | ENSMUSG( | -0.10333 | 0.793457 | NO | NO | NA |
| Vmn2r105 | ENSMUSG( | ENSMUSG( | 0.739054 | 0.480215 | NO | NO | NA |
| Gm10073 | ENSMUSG( | ENSMUSG( | 0.545845 | 0.035877 | YES | NO | NA |
| Adamts20 | ENSMUSG( | ENSMUSG( | 1.310324 | 2.45E-05 | YES | YES | Upregulated |
| Lmo7 | ENSMUSG( | ENSMUSG( | 0.754797 | 0.091625 | NO | NO | NA |
| Gm12000 | ENSMUSG( | ENSMUSG( | -0.32683 | 0.121576 | NO | NO | NA |
| Evpl | ENSMUSG( | ENSMUSG( | 0.069271 | 0.859261 | NO | NO | NA |
| Gm4149 | ENSMUSG( | ENSMUSG( | 0.197216 | 0.43991 | NO | NO | NA |
| ENSMUSG | ENSMUSG( | ENSMUSG( | 0.683058 | 0.54328 | NO | NO | NA |
| Rplp2-ps1 | ENSMUSG( | ENSMUSG( | 0.216594 | 0.810661 | NO | NO | NA |
| Gm41077 | ENSMUSG( | ENSMUSG( | 0.29513 | 0.166501 | NO | NO | NA |
| Grm5 | ENSMUSG( | ENSMUSG( | 1.026547 | 0.316968 | NO | NO | NA |
| Tjp1 | ENSMUSG( | ENSMUSG( | 0.55517 | 0.323977 | NO | NO | NA |
| Ppp1r14a | ENSMUSG( | ENSMUSG( | 1.169964 | 0.135227 | NO | NO | NA |
| Gm4166 | ENSMUSG( | ENSMUSG( | 0.977781 | 0.322605 | NO | NO | NA |
| Htr1d | ENSMUSG( | ENSMUSG( | 0.503081 | 0.567836 | NO | NO | NA |
| D330050I1 | ENSMUSG( | ENSMUSG( | 0.204656 | 0.32323 | NO | NO | NA |
| Gm42893 | ENSMUSG( | ENSMUSG( | -0.26872 | 0.39556 | NO | NO | NA |
| Epb41l1 | ENSMUSG( | ENSMUSG( | -0.89497 | 0.017199 | YES | NO | NA |
| Zfp704 | ENSMUSG( | ENSMUSG( | 0.84569 | 0.233929 | NO | NO | NA |
| Cntnap1 | ENSMUSG( | ENSMUSG( | -0.45353 | 0.233677 | NO | NO | NA |
| Krt10 | ENSMUSG( | ENSMUSG( | 1.137196 | 6.14E-05 | YES | YES | Upregulated |

| | | | | | | | |
|---|---|---|---|---|---|---|---|
| Gm17029 | ENSMUSG( | ENSMUSG( | 0.179913 | 0.523069 | NO | NO | NA |
| Gm42595 | ENSMUSG( | ENSMUSG( | 0.513522 | 0.077645 | NO | NO | NA |
| Dhh | ENSMUSG( | ENSMUSG( | 0.591532 | 0.422267 | NO | NO | NA |
| Gm33838 | ENSMUSG( | ENSMUSG( | 0.574256 | 0.579443 | NO | NO | NA |
| Msmp | ENSMUSG( | ENSMUSG( | -0.23783 | 0.527955 | NO | NO | NA |
| Myrf | ENSMUSG( | ENSMUSG( | 0.554644 | 0.554406 | NO | NO | NA |
| Teddm2 | ENSMUSG( | ENSMUSG( | 0.014961 | 0.978105 | NO | NO | NA |
| ENSMUSG | ENSMUSG( | ENSMUSG( | 0.811862 | 0.34315 | NO | NO | NA |
| Gm42600 | ENSMUSG( | ENSMUSG( | 0.138332 | 0.601079 | NO | NO | NA |
| 5730507A | ENSMUSG( | ENSMUSG( | 1.049398 | 0.344361 | NO | NO | NA |
| Olfr1178 | ENSMUSG( | ENSMUSG( | 0.737484 | 0.446552 | NO | NO | NA |
| Tlcd4 | ENSMUSG( | ENSMUSG( | 0.766908 | 0.317794 | NO | NO | NA |
| Gm11963 | ENSMUSG( | ENSMUSG( | 0.341498 | 0.616676 | NO | NO | NA |
| ENSMUSG | ENSMUSG( | ENSMUSG( | -0.62833 | 0.013783 | YES | NO | NA |
| 4833445I0 | ENSMUSG( | ENSMUSG( | 0.754774 | 0.018601 | YES | NO | NA |
| Grid2 | ENSMUSG( | ENSMUSG( | 0.45281 | 0.653625 | NO | NO | NA |
| Wdpcp | ENSMUSG( | ENSMUSG( | 0.997975 | 0.012467 | YES | NO | NA |
| Timp3 | ENSMUSG( | ENSMUSG( | 0.918717 | 0.37124 | NO | NO | NA |
| Lpar4 | ENSMUSG( | ENSMUSG( | 0.66796 | 0.513901 | NO | NO | NA |
| Gm15706 | ENSMUSG( | ENSMUSG( | 0.513427 | 0.220868 | NO | NO | NA |
| Rab26os | ENSMUSG( | ENSMUSG( | 0.825555 | 0.021356 | YES | NO | NA |
| Tnfsf15 | ENSMUSG( | ENSMUSG( | 0.736562 | 0.423913 | NO | NO | NA |
| Ankrd61 | ENSMUSG( | ENSMUSG( | -0.08333 | 0.836426 | NO | NO | NA |
| Trav13-4-c | ENSMUSG( | ENSMUSG( | 0.670096 | 0.591909 | NO | NO | NA |
| ENSMUSG | ENSMUSG( | ENSMUSG( | 0.097225 | 0.862299 | NO | NO | NA |
| Trav13n-2 | ENSMUSG( | ENSMUSG( | -0.8687 | 0.748964 | NO | NO | NA |
| Gm47540 | ENSMUSG( | ENSMUSG( | 0.715877 | 0.535806 | NO | NO | NA |
| Efcab8 | ENSMUSG( | ENSMUSG( | 0.361211 | 0.426432 | NO | NO | NA |
| Rpl17-ps3 | ENSMUSG( | ENSMUSG( | -0.02264 | 0.946174 | NO | NO | NA |
| Vmn2r99 | ENSMUSG( | ENSMUSG( | 0.514712 | 0.579864 | NO | NO | NA |
| Pcdhgc5 | ENSMUSG( | ENSMUSG( | -0.16828 | 0.588955 | NO | NO | NA |
| Gm48632 | ENSMUSG( | ENSMUSG( | 0.547665 | 0.444307 | NO | NO | NA |
| Gm43339 | ENSMUSG( | ENSMUSG( | -0.21081 | 0.515022 | NO | NO | NA |
| Aldh3b3 | ENSMUSG( | ENSMUSG( | 0.831719 | 0.410705 | NO | NO | NA |
| H2bc22 | ENSMUSG( | ENSMUSG( | 1.548347 | 5.29E-07 | YES | YES | Upregulated |
| Fbxo41 | ENSMUSG( | ENSMUSG( | 0.351941 | 0.254124 | NO | NO | NA |
| C3 | ENSMUSG( | ENSMUSG( | 0.010746 | 0.987052 | NO | NO | NA |
| Slc6a9 | ENSMUSG( | ENSMUSG( | 1.412351 | 0.027551 | YES | YES | Upregulated |
| Trav6-2 | ENSMUSG( | ENSMUSG( | -2.13837 | 0.064161 | NO | NO | NA |
| Gm9226 | ENSMUSG( | ENSMUSG( | 0.543152 | 0.698039 | NO | NO | NA |
| Ncan | ENSMUSG( | ENSMUSG( | 0.451768 | 0.668742 | NO | NO | NA |
| Ankrd33b | ENSMUSG( | ENSMUSG( | 0.808777 | 0.336678 | NO | NO | NA |
| Ly6g5b | ENSMUSG( | ENSMUSG( | 0.338547 | 0.189579 | NO | NO | NA |
| Gm45894 | ENSMUSG( | ENSMUSG( | -0.29228 | 0.193269 | NO | NO | NA |
| Gm37959 | ENSMUSG( | ENSMUSG( | 0.852725 | 0.449679 | NO | NO | NA |
| 6330537M | ENSMUSG( | ENSMUSG( | 0.491008 | 0.03814 | YES | NO | NA |
| Zfp975 | ENSMUSG( | ENSMUSG( | 0.393755 | 0.322365 | NO | NO | NA |

| | | | | | | | |
|---|---|---|---|---|---|---|---|
| Gm44860 | ENSMUSG( | ENSMUSG( | 0.206228 | 0.818006 | NO | NO | NA |
| Flicr | ENSMUSG( | ENSMUSG( | 0.602474 | 0.254516 | NO | NO | NA |
| Ninl | ENSMUSG( | ENSMUSG( | 0.653978 | 0.011513 | YES | NO | NA |
| Tmem176l | ENSMUSG( | ENSMUSG( | 0.868066 | 0.000729 | YES | NO | NA |
| Gm4890 | ENSMUSG( | ENSMUSG( | 0.615018 | 0.029346 | YES | NO | NA |
| Dtna | ENSMUSG( | ENSMUSG( | 0.728001 | 0.455036 | NO | NO | NA |
| Cacna2d1 | ENSMUSG( | ENSMUSG( | 0.525518 | 0.62014 | NO | NO | NA |
| Grin2a | ENSMUSG( | ENSMUSG( | 0.564371 | 0.56174 | NO | NO | NA |
| C9orf72 | ENSMUSG( | ENSMUSG( | 0.516808 | 0.030435 | YES | NO | NA |
| Olfr531 | ENSMUSG( | ENSMUSG( | 1.16863 | 0.259206 | NO | NO | NA |
| Myzap | ENSMUSG( | ENSMUSG( | 0.264445 | 0.680822 | NO | NO | NA |
| Gm37084 | ENSMUSG( | ENSMUSG( | 0.267291 | 0.455807 | NO | NO | NA |
| Zc4h2 | ENSMUSG( | ENSMUSG( | 0.400003 | 0.193221 | NO | NO | NA |
| Rwdd3 | ENSMUSG( | ENSMUSG( | 0.955097 | 0.018417 | YES | NO | NA |
| Tmem86a | ENSMUSG( | ENSMUSG( | 0.122043 | 0.648182 | NO | NO | NA |
| 9030407P: | ENSMUSG( | ENSMUSG( | 0.565797 | 0.027503 | YES | NO | NA |
| 9130017K: | ENSMUSG( | ENSMUSG( | 0.812962 | 0.356725 | NO | NO | NA |
| Gm44694 | ENSMUSG( | ENSMUSG( | 0.004462 | 0.989667 | NO | NO | NA |
| Cyp4f16 | ENSMUSG( | ENSMUSG( | -0.35437 | 0.066783 | NO | NO | NA |
| Sec14l2 | ENSMUSG( | ENSMUSG( | 0.413768 | 0.231678 | NO | NO | NA |
| Stox2 | ENSMUSG( | ENSMUSG( | 0.311605 | 0.734045 | NO | NO | NA |
| Slc26a1 | ENSMUSG( | ENSMUSG( | 0.393081 | 0.455036 | NO | NO | NA |
| Gm42531 | ENSMUSG( | ENSMUSG( | 0.755184 | 0.096274 | NO | NO | NA |
| Kcnk9 | ENSMUSG( | ENSMUSG( | 0.2542 | 0.80969 | NO | NO | NA |
| Srcin1 | ENSMUSG( | ENSMUSG( | 0.396821 | 0.636498 | NO | NO | NA |
| Rbm24 | ENSMUSG( | ENSMUSG( | 0.984164 | 0.301323 | NO | NO | NA |
| Cldnd2 | ENSMUSG( | ENSMUSG( | -1.21301 | 7.58E-06 | YES | YES | Downregulated |
| Rgs9 | ENSMUSG( | ENSMUSG( | 0.228977 | 0.617076 | NO | NO | NA |
| Gm14308 | ENSMUSG( | ENSMUSG( | 2.270979 | 0.001914 | YES | YES | Upregulated |
| Dbndd2 | ENSMUSG( | ENSMUSG( | 0.251842 | 0.648085 | NO | NO | NA |
| Vmn2r79 | ENSMUSG( | ENSMUSG( | 0.456031 | 0.658068 | NO | NO | NA |
| Gm21152 | ENSMUSG( | ENSMUSG( | 0.322378 | 0.778639 | NO | NO | NA |
| Vmn1r193 | ENSMUSG( | ENSMUSG( | 0.986639 | 0.308915 | NO | NO | NA |
| Vmn2r120 | ENSMUSG( | ENSMUSG( | 0.249659 | 0.772214 | NO | NO | NA |
| Sorbs3 | ENSMUSG( | ENSMUSG( | -1.04186 | 8.73E-05 | YES | YES | Downregulated |
| Gm17690 | ENSMUSG( | ENSMUSG( | 0.001207 | 0.996869 | NO | NO | NA |
| Nkx6-1 | ENSMUSG( | ENSMUSG( | 0.539027 | 0.572582 | NO | NO | NA |
| Gm12689 | ENSMUSG( | ENSMUSG( | 0.817583 | 0.42607 | NO | NO | NA |
| Ddn | ENSMUSG( | ENSMUSG( | 1.325989 | 0.010505 | YES | YES | Upregulated |
| Fnbp1l | ENSMUSG( | ENSMUSG( | 0.84156 | 0.041588 | YES | NO | NA |
| Gm43461 | ENSMUSG( | ENSMUSG( | 0.102946 | 0.714424 | NO | NO | NA |
| Spats1 | ENSMUSG( | ENSMUSG( | 0.543905 | 0.093051 | NO | NO | NA |
| Gm47230 | ENSMUSG( | ENSMUSG( | 0.48321 | 0.554406 | NO | NO | NA |
| Dmp1 | ENSMUSG( | ENSMUSG( | 0.539998 | 0.573325 | NO | NO | NA |
| Intu | ENSMUSG( | ENSMUSG( | 0.43008 | 0.139701 | NO | NO | NA |
| 4930481A | ENSMUSG( | ENSMUSG( | -1.37626 | 7.97E-06 | YES | YES | Downregulated |
| Klk9 | ENSMUSG( | ENSMUSG( | 0.802418 | 0.282081 | NO | NO | NA |

| | | | | | | | |
|---|---|---|---|---|---|---|---|
| Gm5464 | ENSMUSG( | ENSMUSG( | 0.72431 | 0.213504 | NO | NO | NA |
| Myocd | ENSMUSG( | ENSMUSG( | 0.360466 | 0.746988 | NO | NO | NA |
| Trim16 | ENSMUSG( | ENSMUSG( | 0.615131 | 0.113846 | NO | NO | NA |
| Tmod2 | ENSMUSG( | ENSMUSG( | 0.687141 | 0.355207 | NO | NO | NA |
| Ighv5-9 | ENSMUSG( | ENSMUSG( | -1.15371 | 0.782324 | NO | NO | NA |
| Gm20749 | ENSMUSG( | ENSMUSG( | 1.39084 | 0.179936 | NO | NO | NA |
| Olfr99 | ENSMUSG( | ENSMUSG( | -1.17879 | 0.000669 | YES | YES | Downregulated |
| 4921539H | ENSMUSG( | ENSMUSG( | 0.440487 | 0.69132 | NO | NO | NA |
| Gm42733 | ENSMUSG( | ENSMUSG( | -0.65157 | 0.012054 | YES | NO | NA |
| Egfl7 | ENSMUSG( | ENSMUSG( | 1.559541 | 1.01E-06 | YES | YES | Upregulated |
| Tst | ENSMUSG( | ENSMUSG( | -0.25892 | 0.481958 | NO | NO | NA |
| B430218F: | ENSMUSG( | ENSMUSG( | -0.01442 | 0.983853 | NO | NO | NA |
| Acvr1 | ENSMUSG( | ENSMUSG( | 0.803243 | 0.051776 | NO | NO | NA |
| Vmn1r184 | ENSMUSG( | ENSMUSG( | 0.832385 | 0.440714 | NO | NO | NA |
| Dlx1as | ENSMUSG( | ENSMUSG( | 0.874564 | 0.000115 | YES | NO | NA |
| Samd11 | ENSMUSG( | ENSMUSG( | -0.02473 | 0.927457 | NO | NO | NA |
| Zfhx4 | ENSMUSG( | ENSMUSG( | 0.694857 | 0.508329 | NO | NO | NA |
| Cpxm1 | ENSMUSG( | ENSMUSG( | 1.451257 | 0.000298 | YES | YES | Upregulated |
| H1f1 | ENSMUSG( | ENSMUSG( | 0.182133 | 0.446235 | NO | NO | NA |
| ENSMUSG | ENSMUSG( | ENSMUSG( | 0.053328 | 0.836132 | NO | NO | NA |
| Sgpp2 | ENSMUSG( | ENSMUSG( | -0.90076 | 0.006088 | YES | NO | NA |
| Lrp12 | ENSMUSG( | ENSMUSG( | 1.499497 | 1.98E-05 | YES | YES | Upregulated |
| Rin2 | ENSMUSG( | ENSMUSG( | 0.147048 | 0.83258 | NO | NO | NA |
| Lmbr1 | ENSMUSG( | ENSMUSG( | 0.80522 | 0.029179 | YES | NO | NA |
| Gm9926 | ENSMUSG( | ENSMUSG( | 1.054004 | 0.301956 | NO | NO | NA |
| Gm37718 | ENSMUSG( | ENSMUSG( | -0.39572 | 0.171936 | NO | NO | NA |
| Gm37283 | ENSMUSG( | ENSMUSG( | 1.018264 | 0.226737 | NO | NO | NA |
| Vmn1r68 | ENSMUSG( | ENSMUSG( | 0.650581 | 0.53864 | NO | NO | NA |
| Gm38120 | ENSMUSG( | ENSMUSG( | 0.232321 | 0.679922 | NO | NO | NA |
| Gm37151 | ENSMUSG( | ENSMUSG( | 0.890118 | 0.387383 | NO | NO | NA |
| Gm38169 | ENSMUSG( | ENSMUSG( | 0.980286 | 0.338874 | NO | NO | NA |
| Gm16299 | ENSMUSG( | ENSMUSG( | -1.47892 | 0.000558 | YES | YES | Downregulated |
| Proser2 | ENSMUSG( | ENSMUSG( | 0.590389 | 0.596194 | NO | NO | NA |
| Gm43292 | ENSMUSG( | ENSMUSG( | 0.278902 | 0.304074 | NO | NO | NA |
| Gm35363 | ENSMUSG( | ENSMUSG( | -0.80773 | 0.01782 | YES | NO | NA |
| ENSMUSG( | ENSMUSG( | ENSMUSG( | 0.171137 | 0.609694 | NO | NO | NA |
| Mgmt | ENSMUSG( | ENSMUSG( | 0.613215 | 0.42158 | NO | NO | NA |
| 2410003I1 | ENSMUSG( | ENSMUSG( | 1.107429 | 0.209547 | NO | NO | NA |
| Kirrel2 | ENSMUSG( | ENSMUSG( | 0.841841 | 0.401612 | NO | NO | NA |
| 6330418K( | ENSMUSG( | ENSMUSG( | -0.02593 | 0.929502 | NO | NO | NA |
| Gm6598 | ENSMUSG( | ENSMUSG( | 0.442229 | 0.051682 | NO | NO | NA |
| Gm43696 | ENSMUSG( | ENSMUSG( | 0.031474 | 0.938942 | NO | NO | NA |
| Rbm3os | ENSMUSG( | ENSMUSG( | -0.14101 | 0.702923 | NO | NO | NA |
| Igkv3-12 | ENSMUSG( | ENSMUSG( | -2.19724 | 0.389574 | NO | NO | NA |
| Gm10518 | ENSMUSG( | ENSMUSG( | -0.39243 | 0.272677 | NO | NO | NA |
| Aebp1 | ENSMUSG( | ENSMUSG( | 0.230645 | 0.480181 | NO | NO | NA |
| Adam1a | ENSMUSG( | ENSMUSG( | -0.17936 | 0.455689 | NO | NO | NA |

| | | | | | | | |
|---|---|---|---|---|---|---|---|
| Cpne9 | ENSMUSG( | ENSMUSG( | 0.83604 | 0.281448 | NO | NO | NA |
| Gm5676 | ENSMUSG( | ENSMUSG( | 0.45234 | 0.668469 | NO | NO | NA |
| Vmn2r63 | ENSMUSG( | ENSMUSG( | 0.736305 | 0.516667 | NO | NO | NA |
| Csmd2 | ENSMUSG( | ENSMUSG( | 0.615963 | 0.540617 | NO | NO | NA |
| 1110032F( | ENSMUSG( | ENSMUSG( | 2.171434 | 4.73E-07 | YES | YES | Upregulated |
| Kcnq3 | ENSMUSG( | ENSMUSG( | 0.612886 | 0.375718 | NO | NO | NA |
| Atp1a3 | ENSMUSG( | ENSMUSG( | 0.350708 | 0.544325 | NO | NO | NA |
| Gm43653 | ENSMUSG( | ENSMUSG( | 0.925146 | 0.110761 | NO | NO | NA |
| Nol3 | ENSMUSG( | ENSMUSG( | -0.1875 | 0.541419 | NO | NO | NA |
| Fam184b | ENSMUSG( | ENSMUSG( | 0.506999 | 0.452297 | NO | NO | NA |
| Grin3a | ENSMUSG( | ENSMUSG( | 0.542944 | 0.274152 | NO | NO | NA |
| Thsd7b | ENSMUSG( | ENSMUSG( | 0.879588 | 0.017853 | YES | NO | NA |
| Dnase1l3 | ENSMUSG( | ENSMUSG( | 0.290645 | 0.391427 | NO | NO | NA |
| Galr3 | ENSMUSG( | ENSMUSG( | 1.454539 | 5.67E-05 | YES | YES | Upregulated |
| Csprs | ENSMUSG( | ENSMUSG( | -0.0313 | 0.929007 | NO | NO | NA |
| Mycl | ENSMUSG( | ENSMUSG( | 1.040135 | 0.050132 | NO | NO | NA |
| Zic1 | ENSMUSG( | ENSMUSG( | 0.584221 | 0.562185 | NO | NO | NA |
| Spink14 | ENSMUSG( | ENSMUSG( | 1.144218 | 0.281184 | NO | NO | NA |
| Zc2hc1a | ENSMUSG( | ENSMUSG( | 0.506001 | 0.084106 | NO | NO | NA |
| Atp8b5 | ENSMUSG( | ENSMUSG( | 0.816297 | 0.459865 | NO | NO | NA |
| Gm52993 | ENSMUSG( | ENSMUSG( | -2.29597 | 0.000443 | YES | YES | Downregulated |
| Gm47237 | ENSMUSG( | ENSMUSG( | 0.809376 | 0.438286 | NO | NO | NA |
| Gm38187 | ENSMUSG( | ENSMUSG( | 0.717159 | 0.501497 | NO | NO | NA |
| Gm43162 | ENSMUSG( | ENSMUSG( | 0.024781 | 0.943682 | NO | NO | NA |
| Gm37606 | ENSMUSG( | ENSMUSG( | 0.614178 | 0.548516 | NO | NO | NA |
| Zscan18 | ENSMUSG( | ENSMUSG( | 0.929228 | 0.004602 | YES | NO | NA |
| S100a8 | ENSMUSG( | ENSMUSG( | 2.573436 | 0.004972 | YES | YES | Upregulated |
| Ltbp1 | ENSMUSG( | ENSMUSG( | 0.742662 | 0.467463 | NO | NO | NA |
| Trav15d-2- | ENSMUSG( | ENSMUSG( | 2.403407 | 0.192792 | NO | NO | NA |
| Kcnh2 | ENSMUSG( | ENSMUSG( | -0.78488 | 0.023847 | YES | NO | NA |
| Gm43290 | ENSMUSG( | ENSMUSG( | 0.802918 | 0.301956 | NO | NO | NA |
| Palld | ENSMUSG( | ENSMUSG( | 0.272717 | 0.688983 | NO | NO | NA |
| ENSMUSG( | ENSMUSG( | ENSMUSG( | 0.795231 | 0.453225 | NO | NO | NA |
| Gm11707 | ENSMUSG( | ENSMUSG( | -0.3406 | 0.246758 | NO | NO | NA |
| Gm20634 | ENSMUSG( | ENSMUSG( | 0.8087 | 0.003703 | YES | NO | NA |
| Zfp119b | ENSMUSG( | ENSMUSG( | 0.486227 | 0.036383 | YES | NO | NA |
| Eps8 | ENSMUSG( | ENSMUSG( | 0.66295 | 0.127147 | NO | NO | NA |
| Dcc | ENSMUSG( | ENSMUSG( | 0.614711 | 0.559175 | NO | NO | NA |
| Gm9929 | ENSMUSG( | ENSMUSG( | 0.113517 | 0.659188 | NO | NO | NA |
| Nqo1 | ENSMUSG( | ENSMUSG( | 0.436173 | 0.347605 | NO | NO | NA |
| Trav3-1 | ENSMUSG( | ENSMUSG( | -1.80519 | 0.041666 | YES | YES | Downregulated |
| ENSMUSG( | ENSMUSG( | ENSMUSG( | -0.50828 | 0.818468 | NO | NO | NA |
| Gm12388 | ENSMUSG( | ENSMUSG( | 0.506179 | 0.632892 | NO | NO | NA |
| Card14 | ENSMUSG( | ENSMUSG( | 0.148185 | 0.872834 | NO | NO | NA |
| Gzmm | ENSMUSG( | ENSMUSG( | -0.20854 | 0.620582 | NO | NO | NA |
| Cpne5 | ENSMUSG( | ENSMUSG( | 0.588862 | 0.402216 | NO | NO | NA |
| Cdh5 | ENSMUSG( | ENSMUSG( | -0.53015 | 0.291953 | NO | NO | NA |

| | | | | | | | |
|---|---|---|---|---|---|---|---|
| C030017B( | ENSMUSG( | ENSMUSG( | 0.705024 | 0.522939 | NO | NO | NA |
| Cspg4 | ENSMUSG( | ENSMUSG( | 0.235378 | 0.809764 | NO | NO | NA |
| Sema4g | ENSMUSG( | ENSMUSG( | 0.21814 | 0.711222 | NO | NO | NA |
| Gm28187 | ENSMUSG( | ENSMUSG( | 0.018977 | 0.958234 | NO | NO | NA |
| Gm37639 | ENSMUSG( | ENSMUSG( | -0.15047 | 0.657527 | NO | NO | NA |
| Fut2 | ENSMUSG( | ENSMUSG( | 0.813011 | 0.488783 | NO | NO | NA |
| Gm36401 | ENSMUSG( | ENSMUSG( | -0.04486 | 0.860913 | NO | NO | NA |
| Gm10443 | ENSMUSG( | ENSMUSG( | 0.623617 | 0.028758 | YES | NO | NA |
| Gm9889 | ENSMUSG( | ENSMUSG( | 0.42062 | 0.140589 | NO | NO | NA |
| Gm38043 | ENSMUSG( | ENSMUSG( | -0.16195 | 0.760343 | NO | NO | NA |
| Gm8817 | ENSMUSG( | ENSMUSG( | -0.21263 | 0.544716 | NO | NO | NA |
| Gm3365 | ENSMUSG( | ENSMUSG( | 0.004564 | 0.989102 | NO | NO | NA |
| Slc41a2 | ENSMUSG( | ENSMUSG( | 0.553549 | 0.038316 | YES | NO | NA |
| Gm14091 | ENSMUSG( | ENSMUSG( | 0.467747 | 0.109241 | NO | NO | NA |
| Catsperd | ENSMUSG( | ENSMUSG( | 0.858571 | 0.303161 | NO | NO | NA |
| Nectin3 | ENSMUSG( | ENSMUSG( | 0.840725 | 0.062188 | NO | NO | NA |
| Ccdc88a | ENSMUSG( | ENSMUSG( | 1.036142 | 4.37E-06 | YES | YES | Upregulated |
| Pcna-ps2 | ENSMUSG( | ENSMUSG( | 0.318428 | 0.221468 | NO | NO | NA |
| Gm33424 | ENSMUSG( | ENSMUSG( | 1.151577 | 0.00123 | YES | YES | Upregulated |
| Gvin-ps1 | ENSMUSG( | ENSMUSG( | 0.680986 | 0.062481 | NO | NO | NA |
| Ppm1e | ENSMUSG( | ENSMUSG( | 0.497022 | 0.053783 | NO | NO | NA |
| ENSMUSG | ENSMUSG( | ENSMUSG( | 0.072931 | 0.850294 | NO | NO | NA |
| Fignl2 | ENSMUSG( | ENSMUSG( | 0.690008 | 0.522939 | NO | NO | NA |
| Gm41442 | ENSMUSG( | ENSMUSG( | 0.784621 | 0.007567 | YES | NO | NA |
| Bik | ENSMUSG( | ENSMUSG( | -0.05676 | 0.895052 | NO | NO | NA |
| 4833413G | ENSMUSG( | ENSMUSG( | 0.152613 | 0.583116 | NO | NO | NA |
| Esr1 | ENSMUSG( | ENSMUSG( | 0.07399 | 0.913188 | NO | NO | NA |
| ENSMUSG | ENSMUSG( | ENSMUSG( | 0.678833 | 0.545865 | NO | NO | NA |
| Sfxn4 | ENSMUSG( | ENSMUSG( | -0.2424 | 0.41215 | NO | NO | NA |
| 4930404I0 | ENSMUSG( | ENSMUSG( | -0.19842 | 0.586533 | NO | NO | NA |
| Ntm | ENSMUSG( | ENSMUSG( | 1.249679 | 0.216209 | NO | NO | NA |
| Zfp14 | ENSMUSG( | ENSMUSG( | -0.4132 | 0.211315 | NO | NO | NA |
| Ak9 | ENSMUSG( | ENSMUSG( | 0.525913 | 0.575485 | NO | NO | NA |
| 4930455J1 | ENSMUSG( | ENSMUSG( | 0.675154 | 0.49586 | NO | NO | NA |
| Gm19129 | ENSMUSG( | ENSMUSG( | 0.4698 | 0.656466 | NO | NO | NA |
| Gm26704 | ENSMUSG( | ENSMUSG( | 0.948889 | 0.321336 | NO | NO | NA |
| Gm37460 | ENSMUSG( | ENSMUSG( | 0.833799 | 0.426754 | NO | NO | NA |
| Rbpjl | ENSMUSG( | ENSMUSG( | 0.863616 | 0.411823 | NO | NO | NA |
| Tmem117 | ENSMUSG( | ENSMUSG( | 0.67259 | 0.454461 | NO | NO | NA |
| Gm43254 | ENSMUSG( | ENSMUSG( | 0.034554 | 0.942632 | NO | NO | NA |
| Gm18283 | ENSMUSG( | ENSMUSG( | 1.000946 | 0.327368 | NO | NO | NA |
| Phf11 | ENSMUSG( | ENSMUSG( | -0.59066 | 0.000586 | YES | NO | NA |
| Gm28192 | ENSMUSG( | ENSMUSG( | -0.08528 | 0.771401 | NO | NO | NA |
| Pcsk5 | ENSMUSG( | ENSMUSG( | 0.629225 | 0.547307 | NO | NO | NA |
| Ndrg4 | ENSMUSG( | ENSMUSG( | 0.245068 | 0.422568 | NO | NO | NA |
| Gm10785 | ENSMUSG( | ENSMUSG( | 0.280323 | 0.367293 | NO | NO | NA |
| Hdc | ENSMUSG( | ENSMUSG( | 0.169129 | 0.645414 | NO | NO | NA |

| | | | | | | | |
|---|---|---|---|---|---|---|---|
| Ighv1-9 | ENSMUSG( | ENSMUSG( | -0.37029 | 0.87054 | NO | NO | NA |
| Gm45322 | ENSMUSG( | ENSMUSG( | 0.647748 | 0.2063 | NO | NO | NA |
| Rpl6l | ENSMUSG( | ENSMUSG( | 0.727371 | 0.003379 | YES | NO | NA |
| Adgrg3 | ENSMUSG( | ENSMUSG( | 1.070141 | 0.006264 | YES | YES | Upregulated |
| Slc18a2 | ENSMUSG( | ENSMUSG( | -0.12084 | 0.696002 | NO | NO | NA |
| Gm5786 | ENSMUSG( | ENSMUSG( | 0.161924 | 0.793457 | NO | NO | NA |
| ccdc198 | ENSMUSG( | ENSMUSG( | 0.741823 | 0.481807 | NO | NO | NA |
| ENSMUSG | ENSMUSG( | ENSMUSG( | 0.155872 | 0.563484 | NO | NO | NA |
| Gm15501 | ENSMUSG( | ENSMUSG( | 1.169474 | 0.539424 | NO | NO | NA |
| Sgip1 | ENSMUSG( | ENSMUSG( | 0.14417 | 0.721998 | NO | NO | NA |
| Hrk | ENSMUSG( | ENSMUSG( | 1.076768 | 0.289893 | NO | NO | NA |
| Zcchc14 | ENSMUSG( | ENSMUSG( | 0.480119 | 0.11748 | NO | NO | NA |
| Prlr | ENSMUSG( | ENSMUSG( | 0.59084 | 0.555693 | NO | NO | NA |
| Rnf225 | ENSMUSG( | ENSMUSG( | 0.99342 | 0.002228 | YES | NO | NA |
| Igdcc3 | ENSMUSG( | ENSMUSG( | 0.974023 | 0.349371 | NO | NO | NA |
| Gm37975 | ENSMUSG( | ENSMUSG( | 0.176268 | 0.808024 | NO | NO | NA |
| Rps13-ps2 | ENSMUSG( | ENSMUSG( | 0.219768 | 0.500516 | NO | NO | NA |
| Gm4876 | ENSMUSG( | ENSMUSG( | 0.013652 | 0.962355 | NO | NO | NA |
| Gm43411 | ENSMUSG( | ENSMUSG( | 0.956814 | 0.211315 | NO | NO | NA |
| Gm30211 | ENSMUSG( | ENSMUSG( | 0.047463 | 0.900065 | NO | NO | NA |
| ENSMUSG | ENSMUSG( | ENSMUSG( | 0.487516 | 0.055447 | NO | NO | NA |
| 9130024F: | ENSMUSG( | ENSMUSG( | 0.899143 | 0.222278 | NO | NO | NA |
| Gm20546 | ENSMUSG( | ENSMUSG( | 0.911799 | 0.37843 | NO | NO | NA |
| 2310010J1 | ENSMUSG( | ENSMUSG( | 0.468584 | 0.019845 | YES | NO | NA |
| ENSMUSG | ENSMUSG( | ENSMUSG( | 0.360204 | 0.247168 | NO | NO | NA |
| A630035G | ENSMUSG( | ENSMUSG( | 0.2112 | 0.602998 | NO | NO | NA |
| Gm37939 | ENSMUSG( | ENSMUSG( | -0.53501 | 0.402238 | NO | NO | NA |
| Brinp2 | ENSMUSG( | ENSMUSG( | 1.014218 | 0.302959 | NO | NO | NA |
| ENSMUSG( | ENSMUSG( | ENSMUSG( | 0.899322 | 0.260798 | NO | NO | NA |
| Zswim7 | ENSMUSG( | ENSMUSG( | 0.460861 | 0.051251 | NO | NO | NA |
| Gm16630 | ENSMUSG( | ENSMUSG( | 0.724213 | 0.483913 | NO | NO | NA |
| C530005A | ENSMUSG( | ENSMUSG( | 0.322366 | 0.406768 | NO | NO | NA |
| D730003I1 | ENSMUSG( | ENSMUSG( | 0.346826 | 0.297659 | NO | NO | NA |
| Gm37176 | ENSMUSG( | ENSMUSG( | 1.040465 | 0.071142 | NO | NO | NA |
| Tmprss3 | ENSMUSG( | ENSMUSG( | 0.560832 | 0.18725 | NO | NO | NA |
| 2310068J1 | ENSMUSG( | ENSMUSG( | 0.368725 | 0.502992 | NO | NO | NA |
| Gm11914 | ENSMUSG( | ENSMUSG( | 0.374605 | 0.102335 | NO | NO | NA |
| Abcg4 | ENSMUSG( | ENSMUSG( | 0.76794 | 0.06073 | NO | NO | NA |
| 1700086O | ENSMUSG( | ENSMUSG( | 0.92209 | 0.001948 | YES | NO | NA |
| AW04620( | ENSMUSG( | ENSMUSG( | -0.40975 | 0.177787 | NO | NO | NA |
| Asxl3 | ENSMUSG( | ENSMUSG( | 1.095424 | 0.239269 | NO | NO | NA |
| 1700034H | ENSMUSG( | ENSMUSG( | 0.483819 | 0.36682 | NO | NO | NA |
| Aqp7 | ENSMUSG( | ENSMUSG( | 0.761573 | 0.340615 | NO | NO | NA |
| Ffar2 | ENSMUSG( | ENSMUSG( | 1.123043 | 0.122348 | NO | NO | NA |
| Gnao1 | ENSMUSG( | ENSMUSG( | -0.09932 | 0.863593 | NO | NO | NA |
| Wfs1 | ENSMUSG( | ENSMUSG( | -0.04693 | 0.869147 | NO | NO | NA |
| Gm44260 | ENSMUSG( | ENSMUSG( | -0.18678 | 0.514619 | NO | NO | NA |

| | | | | | | | |
|---|---|---|---|---|---|---|---|
| Gm49092 | ENSMUSG( | ENSMUSG( | 0.642567 | 0.55384 | NO | NO | NA |
| Sap18b | ENSMUSG( | ENSMUSG( | 0.353588 | 0.324771 | NO | NO | NA |
| 6430571L1 | ENSMUSG( | ENSMUSG( | 0.799069 | 0.251251 | NO | NO | NA |
| ENSMUSG( | ENSMUSG( | ENSMUSG( | 0.523157 | 0.618911 | NO | NO | NA |
| Gm44560 | ENSMUSG( | ENSMUSG( | 0.462736 | 0.389771 | NO | NO | NA |
| Gm21122 | ENSMUSG( | ENSMUSG( | 0.752097 | 0.491865 | NO | NO | NA |
| Igkv8-19 | ENSMUSG( | ENSMUSG( | 0.498209 | 0.858664 | NO | NO | NA |
| Cdh1 | ENSMUSG( | ENSMUSG( | 0.411213 | 0.615791 | NO | NO | NA |
| D730044K | ENSMUSG( | ENSMUSG( | 0.335128 | 0.538299 | NO | NO | NA |
| Tmem8b | ENSMUSG( | ENSMUSG( | 0.728191 | 0.082069 | NO | NO | NA |
| Ripply3 | ENSMUSG( | ENSMUSG( | -0.50438 | 0.09836 | NO | NO | NA |
| 5930435M | ENSMUSG( | ENSMUSG( | 1.078942 | 0.294284 | NO | NO | NA |
| Gm36738 | ENSMUSG( | ENSMUSG( | -0.22159 | 0.465753 | NO | NO | NA |
| Olfr787 | ENSMUSG( | ENSMUSG( | 0.796526 | 0.481759 | NO | NO | NA |
| Gm19194 | ENSMUSG( | ENSMUSG( | 1.020462 | 0.32635 | NO | NO | NA |
| 1110035H | ENSMUSG( | ENSMUSG( | 0.369411 | 0.254189 | NO | NO | NA |
| Erfe | ENSMUSG( | ENSMUSG( | -0.01876 | 0.965837 | NO | NO | NA |
| Ogdhl | ENSMUSG( | ENSMUSG( | 1.08391 | 0.003232 | YES | YES | Upregulated |
| Snx22 | ENSMUSG( | ENSMUSG( | 0.043441 | 0.864928 | NO | NO | NA |
| ENSMUSG( | ENSMUSG( | ENSMUSG( | 1.627645 | 0.005846 | YES | YES | Upregulated |
| Atxn7l1os1 | ENSMUSG( | ENSMUSG( | 0.630493 | 0.410155 | NO | NO | NA |
| Sema5a | ENSMUSG( | ENSMUSG( | 0.729737 | 0.51542 | NO | NO | NA |
| ENSMUSG | ENSMUSG( | ENSMUSG( | 0.154829 | 0.577779 | NO | NO | NA |
| Gm18997 | ENSMUSG( | ENSMUSG( | 1.393223 | 0.167972 | NO | NO | NA |
| Abi3bp | ENSMUSG( | ENSMUSG( | 0.271751 | 0.645795 | NO | NO | NA |
| ENSMUSG( | ENSMUSG( | ENSMUSG( | 0.282295 | 0.716891 | NO | NO | NA |
| Gstt1 | ENSMUSG( | ENSMUSG( | -0.05972 | 0.875286 | NO | NO | NA |
| K230015D | ENSMUSG( | ENSMUSG( | -0.52822 | 0.087695 | NO | NO | NA |
| Tmem108 | ENSMUSG( | ENSMUSG( | 0.936546 | 0.249941 | NO | NO | NA |
| Gm43059 | ENSMUSG( | ENSMUSG( | -0.07183 | 0.811085 | NO | NO | NA |
| 2600006K( | ENSMUSG( | ENSMUSG( | 0.223021 | 0.531711 | NO | NO | NA |
| Pde5a | ENSMUSG( | ENSMUSG( | 0.817323 | 0.018983 | YES | NO | NA |
| Fgfr3-ps | ENSMUSG( | ENSMUSG( | -0.00696 | 0.984402 | NO | NO | NA |
| Gm42559 | ENSMUSG( | ENSMUSG( | -0.32567 | 0.51982 | NO | NO | NA |
| Syt13 | ENSMUSG( | ENSMUSG( | 0.893162 | 0.382317 | NO | NO | NA |
| Gm44198 | ENSMUSG( | ENSMUSG( | 0.45131 | 0.172162 | NO | NO | NA |
| A430027H | ENSMUSG( | ENSMUSG( | 0.111822 | 0.931315 | NO | NO | NA |
| Gm49395 | ENSMUSG( | ENSMUSG( | -0.25849 | 0.724845 | NO | NO | NA |
| BC065397 | ENSMUSG( | ENSMUSG( | 0.271693 | 0.370629 | NO | NO | NA |
| 1810053B2 | ENSMUSG( | ENSMUSG( | 0.825892 | 0.456151 | NO | NO | NA |
| Gng12 | ENSMUSG( | ENSMUSG( | 0.304445 | 0.170248 | NO | NO | NA |
| Sorbs2 | ENSMUSG( | ENSMUSG( | 0.228552 | 0.787106 | NO | NO | NA |
| Oxld1 | ENSMUSG( | ENSMUSG( | -0.08693 | 0.774294 | NO | NO | NA |
| Lncbate10 | ENSMUSG( | ENSMUSG( | 0.816238 | 0.457735 | NO | NO | NA |
| 4732471J( | ENSMUSG( | ENSMUSG( | 0.752877 | 0.146922 | NO | NO | NA |
| Pde6h | ENSMUSG( | ENSMUSG( | 0.909891 | 0.291361 | NO | NO | NA |
| Gm13563 | ENSMUSG( | ENSMUSG( | -0.44888 | 0.037435 | YES | NO | NA |

| | | | | | | | |
|---|---|---|---|---|---|---|---|
| Lbhd2 | ENSMUSG( | ENSMUSG( | 0.069541 | 0.828478 | NO | NO | NA |
| Cxcr5 | ENSMUSG( | ENSMUSG( | -0.32044 | 0.554715 | NO | NO | NA |
| Ubtd2 | ENSMUSG( | ENSMUSG( | -0.34134 | 0.13143 | NO | NO | NA |
| Ccn4 | ENSMUSG( | ENSMUSG( | 0.531548 | 0.53935 | NO | NO | NA |
| Trgv3 | ENSMUSG( | ENSMUSG( | -1.18076 | 0.082075 | NO | NO | NA |
| Gm44937 | ENSMUSG( | ENSMUSG( | 0.562833 | 0.492717 | NO | NO | NA |
| Gm37820 | ENSMUSG( | ENSMUSG( | 0.504243 | 0.396346 | NO | NO | NA |
| Sec14l5 | ENSMUSG( | ENSMUSG( | 1.128739 | 0.273375 | NO | NO | NA |
| Mzf1 | ENSMUSG( | ENSMUSG( | 0.777581 | 0.01523 | YES | NO | NA |
| Denn2b | ENSMUSG( | ENSMUSG( | -0.14322 | 0.818084 | NO | NO | NA |
| Grm6 | ENSMUSG( | ENSMUSG( | 0.551374 | 0.552341 | NO | NO | NA |
| Fat1 | ENSMUSG( | ENSMUSG( | 0.136204 | 0.790471 | NO | NO | NA |
| Slc6a13 | ENSMUSG( | ENSMUSG( | 1.324645 | 0.001049 | YES | YES | Upregulated |
| Doc2g | ENSMUSG( | ENSMUSG( | -0.16483 | 0.664907 | NO | NO | NA |
| Tspan6 | ENSMUSG( | ENSMUSG( | 0.664984 | 0.021627 | YES | NO | NA |
| Gm45324 | ENSMUSG( | ENSMUSG( | 0.560283 | 0.592341 | NO | NO | NA |
| Vmn2r77 | ENSMUSG( | ENSMUSG( | 0.709561 | 0.519345 | NO | NO | NA |
| Gm35315 | ENSMUSG( | ENSMUSG( | 0.606975 | 0.021963 | YES | NO | NA |
| Gm44369 | ENSMUSG( | ENSMUSG( | 0.828298 | 0.153819 | NO | NO | NA |
| Gm29325 | ENSMUSG( | ENSMUSG( | 1.069562 | 0.304251 | NO | NO | NA |
| H3c6 | ENSMUSG( | ENSMUSG( | 0.497322 | 0.038837 | YES | NO | NA |
| Gm13127 | ENSMUSG( | ENSMUSG( | 0.239027 | 0.453678 | NO | NO | NA |
| Sh3rf2 | ENSMUSG( | ENSMUSG( | 0.999122 | 0.300864 | NO | NO | NA |
| Sdc1 | ENSMUSG( | ENSMUSG( | 0.833813 | 0.081382 | NO | NO | NA |
| Gm26964 | ENSMUSG( | ENSMUSG( | -0.08011 | 0.802728 | NO | NO | NA |
| Gm10658 | ENSMUSG( | ENSMUSG( | -0.13497 | 0.758098 | NO | NO | NA |
| Trim72 | ENSMUSG( | ENSMUSG( | -0.97188 | 0.001549 | YES | NO | NA |
| Zfp647 | ENSMUSG( | ENSMUSG( | 0.572542 | 0.011526 | YES | NO | NA |
| Gm44080 | ENSMUSG( | ENSMUSG( | 0.46059 | 0.641748 | NO | NO | NA |
| ENSMUSG | ENSMUSG( | ENSMUSG( | 0.867156 | 0.178359 | NO | NO | NA |
| Dapk1 | ENSMUSG( | ENSMUSG( | 0.919864 | 0.135411 | NO | NO | NA |
| Gm29054 | ENSMUSG( | ENSMUSG( | 0.827302 | 0.456867 | NO | NO | NA |
| Gm42585 | ENSMUSG( | ENSMUSG( | 0.349002 | 0.198096 | NO | NO | NA |
| Gm4202 | ENSMUSG( | ENSMUSG( | 0.200356 | 0.43323 | NO | NO | NA |
| Rpl37rt | ENSMUSG( | ENSMUSG( | 0.264541 | 0.78063 | NO | NO | NA |
| Olfr481 | ENSMUSG( | ENSMUSG( | 0.42097 | 0.672606 | NO | NO | NA |
| Gm17435 | ENSMUSG( | ENSMUSG( | -0.15581 | 0.57298 | NO | NO | NA |
| Hoxa9 | ENSMUSG( | ENSMUSG( | 0.803177 | 0.406768 | NO | NO | NA |
| Gpi-ps | ENSMUSG( | ENSMUSG( | 0.270011 | 0.416281 | NO | NO | NA |
| Tmprss11a | ENSMUSG( | ENSMUSG( | 0.797509 | 0.455036 | NO | NO | NA |
| Gm49971 | ENSMUSG( | ENSMUSG( | 0.117598 | 0.869991 | NO | NO | NA |
| 4732416N | ENSMUSG( | ENSMUSG( | -0.1481 | 0.595836 | NO | NO | NA |
| Gm12743 | ENSMUSG( | ENSMUSG( | 0.369157 | 0.2567 | NO | NO | NA |
| Slc26a3 | ENSMUSG( | ENSMUSG( | 0.617006 | 0.581481 | NO | NO | NA |
| Gm9922 | ENSMUSG( | ENSMUSG( | 0.716592 | 0.490298 | NO | NO | NA |
| Gm8251 | ENSMUSG( | ENSMUSG( | -0.06536 | 0.904797 | NO | NO | NA |
| Gm43657 | ENSMUSG( | ENSMUSG( | 0.867636 | 0.38877 | NO | NO | NA |

| | | | | | | | |
|---|---|---|---|---|---|---|---|
| En2 | ENSMUSG( | ENSMUSG( | 0.514873 | 0.589466 | NO | NO | NA |
| Clec12a | ENSMUSG( | ENSMUSG( | 0.951533 | 0.042308 | YES | NO | NA |
| Gm16214 | ENSMUSG( | ENSMUSG( | -0.01217 | 0.970155 | NO | NO | NA |
| Sarm1 | ENSMUSG( | ENSMUSG( | -0.1861 | 0.533915 | NO | NO | NA |
| Gm10827 | ENSMUSG( | ENSMUSG( | 0.845983 | 0.100129 | NO | NO | NA |
| ENSMUSG | ENSMUSG( | ENSMUSG( | -0.39908 | 0.175984 | NO | NO | NA |
| Unc93a | ENSMUSG( | ENSMUSG( | 0.99674 | 0.33423 | NO | NO | NA |
| 2310039L1 | ENSMUSG( | ENSMUSG( | 0.59904 | 0.56241 | NO | NO | NA |
| Gm6627 | ENSMUSG( | ENSMUSG( | 0.12259 | 0.567382 | NO | NO | NA |
| Gm44154 | ENSMUSG( | ENSMUSG( | 1.073797 | 0.315975 | NO | NO | NA |
| Gm45767 | ENSMUSG( | ENSMUSG( | 0.350209 | 0.570097 | NO | NO | NA |
| Gm38375 | ENSMUSG( | ENSMUSG( | -0.51378 | 0.096441 | NO | NO | NA |
| 2900041M | ENSMUSG( | ENSMUSG( | -0.36246 | 0.156646 | NO | NO | NA |
| Cited4 | ENSMUSG( | ENSMUSG( | 0.062796 | 0.84782 | NO | NO | NA |
| Gm5898 | ENSMUSG( | ENSMUSG( | 0.187426 | 0.461665 | NO | NO | NA |
| Gm14586 | ENSMUSG( | ENSMUSG( | 0.637243 | 0.046546 | YES | NO | NA |
| Tas1r3 | ENSMUSG( | ENSMUSG( | -0.09157 | 0.787243 | NO | NO | NA |
| Tgfb1i1 | ENSMUSG( | ENSMUSG( | -0.65572 | 0.007851 | YES | NO | NA |
| Prss2 | ENSMUSG( | ENSMUSG( | 1.016589 | 0.250252 | NO | NO | NA |
| H3c8 | ENSMUSG( | ENSMUSG( | 0.585599 | 0.121613 | NO | NO | NA |
| Igkv8-30 | ENSMUSG( | ENSMUSG( | -0.23298 | 0.903282 | NO | NO | NA |
| Gm37137 | ENSMUSG( | ENSMUSG( | 0.485498 | 0.601687 | NO | NO | NA |
| ENSMUSG | ENSMUSG( | ENSMUSG( | 0.66867 | 0.431235 | NO | NO | NA |
| Trabd2b | ENSMUSG( | ENSMUSG( | 0.686785 | 0.411184 | NO | NO | NA |
| Gm48114 | ENSMUSG( | ENSMUSG( | -0.49625 | 0.272054 | NO | NO | NA |
| Gm16023 | ENSMUSG( | ENSMUSG( | -0.27987 | 0.416291 | NO | NO | NA |
| Odf3b | ENSMUSG( | ENSMUSG( | 0.919984 | 0.231297 | NO | NO | NA |
| Angptl3 | ENSMUSG( | ENSMUSG( | 1.241167 | 0.212884 | NO | NO | NA |
| E230025N | ENSMUSG( | ENSMUSG( | 0.166692 | 0.872834 | NO | NO | NA |
| Txlnb | ENSMUSG( | ENSMUSG( | -0.02889 | 0.925564 | NO | NO | NA |
| Trav12n-3 | ENSMUSG( | ENSMUSG( | -1.25571 | 0.685592 | NO | NO | NA |
| Gm19918 | ENSMUSG( | ENSMUSG( | -0.19343 | 0.450798 | NO | NO | NA |
| Styk1 | ENSMUSG( | ENSMUSG( | 0.370722 | 0.468243 | NO | NO | NA |
| Gm12174 | ENSMUSG( | ENSMUSG( | 0.422696 | 0.040051 | YES | NO | NA |
| Gm38192 | ENSMUSG( | ENSMUSG( | 0.290569 | 0.510362 | NO | NO | NA |
| Tfap2c | ENSMUSG( | ENSMUSG( | 0.741525 | 0.480639 | NO | NO | NA |
| Chrne | ENSMUSG( | ENSMUSG( | -0.45618 | 0.119138 | NO | NO | NA |
| 4921515L2 | ENSMUSG( | ENSMUSG( | 0.205197 | 0.638931 | NO | NO | NA |
| Pappa | ENSMUSG( | ENSMUSG( | 0.871986 | 0.086241 | NO | NO | NA |
| Gm11518 | ENSMUSG( | ENSMUSG( | 0.243551 | 0.606953 | NO | NO | NA |
| Gm47479 | ENSMUSG( | ENSMUSG( | 0.338371 | 0.705541 | NO | NO | NA |
| Gm12764 | ENSMUSG( | ENSMUSG( | -0.7922 | 0.007361 | YES | NO | NA |
| Mn1 | ENSMUSG( | ENSMUSG( | 0.730094 | 0.450394 | NO | NO | NA |
| F830115B( | ENSMUSG( | ENSMUSG( | 0.566345 | 0.486893 | NO | NO | NA |
| Gm37520 | ENSMUSG( | ENSMUSG( | -0.02006 | 0.962056 | NO | NO | NA |
| Bex4 | ENSMUSG( | ENSMUSG( | 1.381971 | 0.102149 | NO | NO | NA |
| Gjc1 | ENSMUSG( | ENSMUSG( | 0.587251 | 0.481958 | NO | NO | NA |

| | | | | | | | |
|---|---|---|---|---|---|---|---|
| Stc2 | ENSMUSG( | ENSMUSG( | 0.5521 | 0.411282 | NO | NO | NA |
| Mmp19 | ENSMUSG( | ENSMUSG( | 0.547711 | 0.103161 | NO | NO | NA |
| Lrrc4c | ENSMUSG( | ENSMUSG( | 0.633826 | 0.489506 | NO | NO | NA |
| Gm19261 | ENSMUSG( | ENSMUSG( | -0.44807 | 0.019908 | YES | NO | NA |
| Septin3 | ENSMUSG( | ENSMUSG( | 0.328447 | 0.728296 | NO | NO | NA |
| Klkb1 | ENSMUSG( | ENSMUSG( | 0.496911 | 0.617169 | NO | NO | NA |
| Gm45746 | ENSMUSG( | ENSMUSG( | 0.660823 | 0.411282 | NO | NO | NA |
| Gm36933 | ENSMUSG( | ENSMUSG( | -0.1181 | 0.809302 | NO | NO | NA |
| Pax9 | ENSMUSG( | ENSMUSG( | -0.06219 | 0.880057 | NO | NO | NA |
| Gpr176 | ENSMUSG( | ENSMUSG( | 1.067034 | 0.128729 | NO | NO | NA |
| ENSMUSG | ENSMUSG( | ENSMUSG( | 0.825309 | 0.188954 | NO | NO | NA |
| Gm37553 | ENSMUSG( | ENSMUSG( | -0.16444 | 0.711222 | NO | NO | NA |
| Ano5 | ENSMUSG( | ENSMUSG( | 0.616873 | 0.501479 | NO | NO | NA |
| Rab3b | ENSMUSG( | ENSMUSG( | 0.605495 | 0.443115 | NO | NO | NA |
| Vmn1r61 | ENSMUSG( | ENSMUSG( | 0.912625 | 0.332415 | NO | NO | NA |
| Gm45163 | ENSMUSG( | ENSMUSG( | 0.483452 | 0.63616 | NO | NO | NA |
| Prune2 | ENSMUSG( | ENSMUSG( | 0.282703 | 0.664907 | NO | NO | NA |
| Olfr160 | ENSMUSG( | ENSMUSG( | 0.653972 | 0.532011 | NO | NO | NA |
| Unc93a2 | ENSMUSG( | ENSMUSG( | 0.849155 | 0.411282 | NO | NO | NA |
| Atp6v0a1 | ENSMUSG( | ENSMUSG( | 0.597228 | 0.279784 | NO | NO | NA |
| Hoxd9 | ENSMUSG( | ENSMUSG( | 1.2831 | 0.192067 | NO | NO | NA |
| Gm50287 | ENSMUSG( | ENSMUSG( | -0.5985 | 0.040296 | YES | NO | NA |
| Lix1 | ENSMUSG( | ENSMUSG( | 0.89785 | 0.389049 | NO | NO | NA |
| Syt1 | ENSMUSG( | ENSMUSG( | 0.798631 | 0.444105 | NO | NO | NA |
| A430035B | ENSMUSG( | ENSMUSG( | 0.094882 | 0.707421 | NO | NO | NA |
| Olfr155 | ENSMUSG( | ENSMUSG( | 1.105241 | 0.358484 | NO | NO | NA |
| Gm10652 | ENSMUSG( | ENSMUSG( | 1.065296 | 0.201886 | NO | NO | NA |
| ENSMUSG | ENSMUSG( | ENSMUSG( | -20.7179 | NA | NO | NO | NA |
| C78283 | ENSMUSG( | ENSMUSG( | 0.585557 | 0.655162 | NO | NO | NA |
| Drd5 | ENSMUSG( | ENSMUSG( | 0.523279 | 0.566954 | NO | NO | NA |
| Gm6741 | ENSMUSG( | ENSMUSG( | -0.50747 | 0.617412 | NO | NO | NA |
| Pecr | ENSMUSG( | ENSMUSG( | 0.713535 | 0.033924 | YES | NO | NA |
| ENSMUSG | ENSMUSG( | ENSMUSG( | 0.576463 | 0.528272 | NO | NO | NA |
| Rps6ka6 | ENSMUSG( | ENSMUSG( | 0.99508 | 0.000199 | YES | NO | NA |
| Irx3os | ENSMUSG( | ENSMUSG( | 0.412065 | 0.723744 | NO | NO | NA |
| Wdfy4 | ENSMUSG( | ENSMUSG( | -1.2342 | 0.022739 | YES | YES | Downregulated |
| Olfr63 | ENSMUSG( | ENSMUSG( | 0.671718 | 0.544336 | NO | NO | NA |
| Ighv9-1 | ENSMUSG( | ENSMUSG( | 2.088473 | 0.551248 | NO | NO | NA |
| Gzme | ENSMUSG( | ENSMUSG( | 7.041663 | 4.63E-08 | YES | YES | Upregulated |
| Fam13a | ENSMUSG( | ENSMUSG( | 0.659214 | 0.542045 | NO | NO | NA |
| Kcnip2 | ENSMUSG( | ENSMUSG( | -0.12431 | 0.631695 | NO | NO | NA |
| Greb1 | ENSMUSG( | ENSMUSG( | 0.757219 | 0.491461 | NO | NO | NA |
| Vmn1r219 | ENSMUSG( | ENSMUSG( | 0.831434 | 0.398689 | NO | NO | NA |
| Gm5148 | ENSMUSG( | ENSMUSG( | 0.246565 | 0.29379 | NO | NO | NA |
| Ltbp3 | ENSMUSG( | ENSMUSG( | 0.63126 | 0.085955 | NO | NO | NA |
| Gm37370 | ENSMUSG( | ENSMUSG( | -0.30784 | 0.552362 | NO | NO | NA |
| Kcnc2 | ENSMUSG( | ENSMUSG( | 0.646014 | 0.593909 | NO | NO | NA |

| | | | | | | | |
|---|---|---|---|---|---|---|---|
| Tfr2 | ENSMUSG( | ENSMUSG( | 0.127491 | 0.860932 | NO | NO | NA |
| Mboat4 | ENSMUSG( | ENSMUSG( | -0.02041 | 0.960275 | NO | NO | NA |
| Mcidas | ENSMUSG( | ENSMUSG( | 0.611135 | 0.566439 | NO | NO | NA |
| Prrg2 | ENSMUSG( | ENSMUSG( | 0.221895 | 0.385786 | NO | NO | NA |
| Gm47652 | ENSMUSG( | ENSMUSG( | 0.138681 | 0.738771 | NO | NO | NA |
| Hif3a | ENSMUSG( | ENSMUSG( | 0.355678 | 0.49812 | NO | NO | NA |
| Gm36041 | ENSMUSG( | ENSMUSG( | 0.358922 | 0.729736 | NO | NO | NA |
| Clic3 | ENSMUSG( | ENSMUSG( | -0.05702 | 0.843184 | NO | NO | NA |
| Pou4f1 | ENSMUSG( | ENSMUSG( | 0.84396 | 0.001296 | YES | NO | NA |
| F630040K( | ENSMUSG( | ENSMUSG( | 0.767439 | 0.48915 | NO | NO | NA |
| Gm37886 | ENSMUSG( | ENSMUSG( | 0.085579 | 0.896229 | NO | NO | NA |
| Igkv6-15 | ENSMUSG( | ENSMUSG( | -0.83348 | 0.652117 | NO | NO | NA |
| Gm47017 | ENSMUSG( | ENSMUSG( | 0.97366 | 0.354347 | NO | NO | NA |
| 4931431B: | ENSMUSG( | ENSMUSG( | -1.61207 | 5.78E-06 | YES | YES | Downregulated |
| Wdr87-ps | ENSMUSG( | ENSMUSG( | 0.617908 | 0.584439 | NO | NO | NA |
| AI593442 | ENSMUSG( | ENSMUSG( | 0.72576 | 0.495046 | NO | NO | NA |
| ENSMUSG | ENSMUSG( | ENSMUSG( | 0.788998 | 0.202772 | NO | NO | NA |
| Trp63 | ENSMUSG( | ENSMUSG( | 0.85763 | 0.190858 | NO | NO | NA |
| Snap25 | ENSMUSG( | ENSMUSG( | 0.956347 | 0.314547 | NO | NO | NA |
| D430001F | ENSMUSG( | ENSMUSG( | 0.581659 | 0.495513 | NO | NO | NA |
| Gm17167 | ENSMUSG( | ENSMUSG( | 0.23614 | 0.653824 | NO | NO | NA |
| A430110C | ENSMUSG( | ENSMUSG( | 0.539841 | 0.547701 | NO | NO | NA |
| Gm45250 | ENSMUSG( | ENSMUSG( | 0.156175 | 0.691135 | NO | NO | NA |
| Dynlt2b | ENSMUSG( | ENSMUSG( | 0.920958 | 0.05289 | NO | NO | NA |
| Vmn2r59 | ENSMUSG( | ENSMUSG( | 0.492391 | 0.62793 | NO | NO | NA |
| Gm38312 | ENSMUSG( | ENSMUSG( | 0.707657 | 0.021963 | YES | NO | NA |
| Gm48719 | ENSMUSG( | ENSMUSG( | -0.23935 | 0.334634 | NO | NO | NA |
| Kcnb2 | ENSMUSG( | ENSMUSG( | 0.484635 | 0.64009 | NO | NO | NA |
| Gm35035 | ENSMUSG( | ENSMUSG( | -0.36514 | 0.517147 | NO | NO | NA |
| Dync2i1 | ENSMUSG( | ENSMUSG( | 1.043616 | 0.005914 | YES | YES | Upregulated |
| Trav3-4 | ENSMUSG( | ENSMUSG( | -1.53259 | 0.176201 | NO | NO | NA |
| Gm17083 | ENSMUSG( | ENSMUSG( | 0.156766 | 0.775195 | NO | NO | NA |
| Ctbp2 | ENSMUSG( | ENSMUSG( | 1.284845 | 0.000107 | YES | YES | Upregulated |
| Ccdc17 | ENSMUSG( | ENSMUSG( | -0.10672 | 0.79767 | NO | NO | NA |
| ENSMUSG( | ENSMUSG( | ENSMUSG( | -0.07581 | 0.793495 | NO | NO | NA |
| Gm43579 | ENSMUSG( | ENSMUSG( | -0.29621 | 0.259206 | NO | NO | NA |
| Bbs10 | ENSMUSG( | ENSMUSG( | -0.22974 | 0.43299 | NO | NO | NA |
| Ighv1-76 | ENSMUSG( | ENSMUSG( | -4.39606 | 0.00312 | YES | YES | Downregulated |
| Fgf9 | ENSMUSG( | ENSMUSG( | 0.844996 | 0.362077 | NO | NO | NA |
| Vmn2r69 | ENSMUSG( | ENSMUSG( | 0.555602 | 0.590649 | NO | NO | NA |
| Trim47 | ENSMUSG( | ENSMUSG( | -0.4191 | 0.334234 | NO | NO | NA |
| Gm33782 | ENSMUSG( | ENSMUSG( | -1.5002 | 8.74E-10 | YES | YES | Downregulated |
| Alpk2 | ENSMUSG( | ENSMUSG( | 0.577139 | 0.039169 | YES | NO | NA |
| Vmn1r185 | ENSMUSG( | ENSMUSG( | 0.357729 | 0.727144 | NO | NO | NA |
| Gm6085 | ENSMUSG( | ENSMUSG( | 0.505265 | 0.479375 | NO | NO | NA |
| Gm13264 | ENSMUSG( | ENSMUSG( | 1.159486 | 0.232128 | NO | NO | NA |
| Gm42909 | ENSMUSG( | ENSMUSG( | 0.933011 | 0.363612 | NO | NO | NA |

| | | | | | | | |
|---|---|---|---|---|---|---|---|
| Gm37250 | ENSMUSG( | ENSMUSG( | 0.643329 | 0.520168 | NO | NO | NA |
| Galr2 | ENSMUSG( | ENSMUSG( | 0.285297 | 0.693783 | NO | NO | NA |
| Col12a1 | ENSMUSG( | ENSMUSG( | 0.630853 | 0.484899 | NO | NO | NA |
| Gm43009 | ENSMUSG( | ENSMUSG( | 0.661454 | 0.499353 | NO | NO | NA |
| ENSMUSG( | ENSMUSG( | ENSMUSG( | -0.29957 | 0.580206 | NO | NO | NA |
| Jakmip2 | ENSMUSG( | ENSMUSG( | 0.593689 | 0.247115 | NO | NO | NA |
| Gm43080 | ENSMUSG( | ENSMUSG( | 0.268315 | 0.478075 | NO | NO | NA |
| Minar2 | ENSMUSG( | ENSMUSG( | 0.586767 | 0.595777 | NO | NO | NA |
| Fndc5 | ENSMUSG( | ENSMUSG( | 0.371693 | 0.693537 | NO | NO | NA |
| Vmn2r67 | ENSMUSG( | ENSMUSG( | 0.457642 | 0.653603 | NO | NO | NA |
| Vmn1r172 | ENSMUSG( | ENSMUSG( | 0.599003 | 0.568758 | NO | NO | NA |
| Amacr | ENSMUSG( | ENSMUSG( | 0.33987 | 0.216926 | NO | NO | NA |
| Spsb4 | ENSMUSG( | ENSMUSG( | 0.839711 | 0.414065 | NO | NO | NA |
| 5830448L( | ENSMUSG( | ENSMUSG( | -0.01085 | 0.976023 | NO | NO | NA |
| Gnrh1 | ENSMUSG( | ENSMUSG( | 0.242941 | 0.504794 | NO | NO | NA |
| Dpysl4 | ENSMUSG( | ENSMUSG( | 0.279821 | 0.673618 | NO | NO | NA |
| Rtn1 | ENSMUSG( | ENSMUSG( | 0.786298 | 0.306075 | NO | NO | NA |
| Prg4 | ENSMUSG( | ENSMUSG( | -0.5621 | 0.01184 | YES | NO | NA |
| Igfbp7 | ENSMUSG( | ENSMUSG( | 3.008038 | 2.47E-07 | YES | YES | Upregulated |
| ENSMUSG | ENSMUSG( | ENSMUSG( | 0.538456 | 0.372606 | NO | NO | NA |
| Dmd | ENSMUSG( | ENSMUSG( | 0.948112 | 0.070215 | NO | NO | NA |
| Fsd2 | ENSMUSG( | ENSMUSG( | -0.49762 | 0.149234 | NO | NO | NA |
| ENSMUSG( | ENSMUSG( | ENSMUSG( | -0.49853 | 0.061422 | NO | NO | NA |
| Gm37018 | ENSMUSG( | ENSMUSG( | 0.642253 | 0.561489 | NO | NO | NA |
| Gm37745 | ENSMUSG( | ENSMUSG( | 0.736072 | 0.338636 | NO | NO | NA |
| Olfr550 | ENSMUSG( | ENSMUSG( | 0.853298 | 0.49684 | NO | NO | NA |
| Msi1 | ENSMUSG( | ENSMUSG( | 0.47484 | 0.656697 | NO | NO | NA |
| Gm8744 | ENSMUSG( | ENSMUSG( | 0.514302 | 0.600373 | NO | NO | NA |
| Gm43684 | ENSMUSG( | ENSMUSG( | 0.205128 | 0.644112 | NO | NO | NA |
| Fstl3 | ENSMUSG( | ENSMUSG( | 0.7971 | 0.002349 | YES | NO | NA |
| 2810404F: | ENSMUSG( | ENSMUSG( | 1.023578 | 0.297023 | NO | NO | NA |
| Col11a1 | ENSMUSG( | ENSMUSG( | 0.240429 | 0.798402 | NO | NO | NA |
| Epha3 | ENSMUSG( | ENSMUSG( | 0.19774 | 0.682598 | NO | NO | NA |
| Mipol1 | ENSMUSG( | ENSMUSG( | 0.140023 | 0.589938 | NO | NO | NA |
| Tbx18 | ENSMUSG( | ENSMUSG( | 0.973291 | 0.324857 | NO | NO | NA |
| E030026E: | ENSMUSG( | ENSMUSG( | 0.446388 | 0.687679 | NO | NO | NA |
| Nuak2 | ENSMUSG( | ENSMUSG( | 0.742378 | 0.16428 | NO | NO | NA |
| Pde3a | ENSMUSG( | ENSMUSG( | 0.673368 | 0.507007 | NO | NO | NA |
| Pappa2 | ENSMUSG( | ENSMUSG( | 0.781293 | 0.500772 | NO | NO | NA |
| Aff3 | ENSMUSG( | ENSMUSG( | 0.303479 | 0.559849 | NO | NO | NA |
| Gm4419 | ENSMUSG( | ENSMUSG( | -0.24052 | 0.401746 | NO | NO | NA |
| 1110006O | ENSMUSG( | ENSMUSG( | 0.588309 | 0.093322 | NO | NO | NA |
| Sez6l2 | ENSMUSG( | ENSMUSG( | -0.08744 | 0.819148 | NO | NO | NA |
| Olfr398 | ENSMUSG( | ENSMUSG( | 0.693956 | 0.527278 | NO | NO | NA |
| Eng | ENSMUSG( | ENSMUSG( | 0.743542 | 0.100599 | NO | NO | NA |
| Ecm1 | ENSMUSG( | ENSMUSG( | 0.141616 | 0.56241 | NO | NO | NA |
| Fam161a | ENSMUSG( | ENSMUSG( | -0.04064 | 0.900065 | NO | NO | NA |

| | | | | | | | |
|---|---|---|---|---|---|---|---|
| Hoxb1 | ENSMUSG( | ENSMUSG( | 0.468605 | 0.651774 | NO | NO | NA |
| Gm2093 | ENSMUSG( | ENSMUSG( | 1.089723 | 0.2625 | NO | NO | NA |
| 4833422C: | ENSMUSG( | ENSMUSG( | 0.735772 | 0.351132 | NO | NO | NA |
| Tmem116 | ENSMUSG( | ENSMUSG( | -0.16684 | 0.491865 | NO | NO | NA |
| Ccr1 | ENSMUSG( | ENSMUSG( | 2.024899 | 0.000409 | YES | YES | Upregulated |
| Gm46516 | ENSMUSG( | ENSMUSG( | 0.905187 | 0.375594 | NO | NO | NA |
| Ak1 | ENSMUSG( | ENSMUSG( | 0.747568 | 0.369297 | NO | NO | NA |
| Ldhb | ENSMUSG( | ENSMUSG( | 0.18559 | 0.616442 | NO | NO | NA |
| Rbfaos | ENSMUSG( | ENSMUSG( | 0.387355 | 0.325684 | NO | NO | NA |
| Gm9531 | ENSMUSG( | ENSMUSG( | 0.140162 | 0.670951 | NO | NO | NA |
| H3c11 | ENSMUSG( | ENSMUSG( | 0.172268 | 0.618809 | NO | NO | NA |
| Olfr95 | ENSMUSG( | ENSMUSG( | 0.931256 | 0.332122 | NO | NO | NA |
| Pmel | ENSMUSG( | ENSMUSG( | -0.11472 | 0.701327 | NO | NO | NA |
| St6galnac5 | ENSMUSG( | ENSMUSG( | 0.515441 | 0.547701 | NO | NO | NA |
| Lmod1 | ENSMUSG( | ENSMUSG( | 0.778428 | 0.403558 | NO | NO | NA |
| Il1rapl1 | ENSMUSG( | ENSMUSG( | 0.721304 | 0.458252 | NO | NO | NA |
| 4930529C( | ENSMUSG( | ENSMUSG( | 0.952165 | 0.378491 | NO | NO | NA |
| Gm11223 | ENSMUSG( | ENSMUSG( | 0.268177 | 0.441555 | NO | NO | NA |
| 4930484H | ENSMUSG( | ENSMUSG( | 0.949315 | 0.332526 | NO | NO | NA |
| Slitrk5 | ENSMUSG( | ENSMUSG( | 0.190551 | 0.871077 | NO | NO | NA |
| Kcnma1 | ENSMUSG( | ENSMUSG( | 0.365326 | 0.719948 | NO | NO | NA |
| 4930412F: | ENSMUSG( | ENSMUSG( | 0.364138 | 0.372816 | NO | NO | NA |
| Gm48210 | ENSMUSG( | ENSMUSG( | 0.123399 | 0.823916 | NO | NO | NA |
| Notch3 | ENSMUSG( | ENSMUSG( | -1.42449 | 0.001334 | YES | YES | Downregulated |
| 2900060N | ENSMUSG( | ENSMUSG( | 0.797425 | 0.373022 | NO | NO | NA |
| Cxcl9 | ENSMUSG( | ENSMUSG( | 0.515423 | 0.583116 | NO | NO | NA |
| Glis2 | ENSMUSG( | ENSMUSG( | 0.366641 | 0.23029 | NO | NO | NA |
| 1110018N | ENSMUSG( | ENSMUSG( | 1.154885 | 0.22713 | NO | NO | NA |
| Tmem151a | ENSMUSG( | ENSMUSG( | -0.84677 | 0.009147 | YES | NO | NA |
| Prg2 | ENSMUSG( | ENSMUSG( | -3.96605 | 0.000339 | YES | YES | Downregulated |
| Gm11532 | ENSMUSG( | ENSMUSG( | 0.455461 | 0.412416 | NO | NO | NA |
| Gm37039 | ENSMUSG( | ENSMUSG( | -0.16695 | 0.738568 | NO | NO | NA |
| ENSMUSG | ENSMUSG( | ENSMUSG( | -0.02232 | 0.961546 | NO | NO | NA |
| Col1a1 | ENSMUSG( | ENSMUSG( | 0.248673 | 0.586288 | NO | NO | NA |
| Gm28884 | ENSMUSG( | ENSMUSG( | -1.06898 | 0.000142 | YES | YES | Downregulated |
| Gm42729 | ENSMUSG( | ENSMUSG( | 0.452331 | 0.463646 | NO | NO | NA |
| Nod2 | ENSMUSG( | ENSMUSG( | -0.67664 | 0.069104 | NO | NO | NA |
| Rnd1 | ENSMUSG( | ENSMUSG( | 0.913807 | 0.05121 | NO | NO | NA |
| Gm49320 | ENSMUSG( | ENSMUSG( | -0.17032 | 0.736727 | NO | NO | NA |
| Actr3b | ENSMUSG( | ENSMUSG( | 0.573043 | 0.057432 | NO | NO | NA |
| Trav7n-5 | ENSMUSG( | ENSMUSG( | 0.181544 | 0.937234 | NO | NO | NA |
| Olfr1416 | ENSMUSG( | ENSMUSG( | 0.777834 | 0.476775 | NO | NO | NA |
| ENSMUSG | ENSMUSG( | ENSMUSG( | 0.762778 | 0.51585 | NO | NO | NA |
| 0610038B: | ENSMUSG( | ENSMUSG( | -0.93516 | 0.003654 | YES | NO | NA |
| ENSMUSG | ENSMUSG( | ENSMUSG( | 0.842078 | 0.258558 | NO | NO | NA |
| Gm14327 | ENSMUSG( | ENSMUSG( | 0.38916 | 0.194534 | NO | NO | NA |
| Gm9246 | ENSMUSG( | ENSMUSG( | 0.494968 | 0.65479 | NO | NO | NA |

| | | | | | | | |
|---|---|---|---|---|---|---|---|
| Psd2 | ENSMUSG( | ENSMUSG( | 0.446005 | 0.138256 | NO | NO | NA |
| 4930549P: | ENSMUSG( | ENSMUSG( | 0.546334 | 0.446966 | NO | NO | NA |
| Cdkn2b | ENSMUSG( | ENSMUSG( | -0.11929 | 0.674516 | NO | NO | NA |
| Gp1ba | ENSMUSG( | ENSMUSG( | 0.010101 | 0.980563 | NO | NO | NA |
| Cdc42bpb | ENSMUSG( | ENSMUSG( | 1.54588 | 0.001373 | YES | YES | Upregulated |
| Ccdc57 | ENSMUSG( | ENSMUSG( | 1.664707 | 4.51E-06 | YES | YES | Upregulated |
| Vmn1r16 | ENSMUSG( | ENSMUSG( | 0.838339 | 0.41768 | NO | NO | NA |
| Gm37653 | ENSMUSG( | ENSMUSG( | 0.178936 | 0.709655 | NO | NO | NA |
| P4ha2 | ENSMUSG( | ENSMUSG( | 0.779039 | 0.406149 | NO | NO | NA |
| Bace2 | ENSMUSG( | ENSMUSG( | 0.760329 | 0.165658 | NO | NO | NA |
| Etnppl | ENSMUSG( | ENSMUSG( | 0.850733 | 0.335786 | NO | NO | NA |
| Ighv1-50 | ENSMUSG( | ENSMUSG( | -6.34138 | 6.40E-05 | YES | YES | Downregulated |
| Ccdc148 | ENSMUSG( | ENSMUSG( | 0.543419 | 0.163485 | NO | NO | NA |
| Trav6d-4 | ENSMUSG( | ENSMUSG( | -1.50146 | 0.131748 | NO | NO | NA |
| Gm19026 | ENSMUSG( | ENSMUSG( | 0.507084 | 0.622634 | NO | NO | NA |
| Cacna1c | ENSMUSG( | ENSMUSG( | 0.419576 | 0.367654 | NO | NO | NA |
| 2610316D | ENSMUSG( | ENSMUSG( | 1.058185 | 0.391308 | NO | NO | NA |
| Gm34220 | ENSMUSG( | ENSMUSG( | 0.048302 | 0.862699 | NO | NO | NA |
| Trav13n-4 | ENSMUSG( | ENSMUSG( | 0.552168 | 0.656466 | NO | NO | NA |
| Gm46432 | ENSMUSG( | ENSMUSG( | -0.30873 | 0.422302 | NO | NO | NA |
| Gm47370 | ENSMUSG( | ENSMUSG( | 1.045778 | 0.27646 | NO | NO | NA |
| Spef2 | ENSMUSG( | ENSMUSG( | 1.501905 | 1.85E-07 | YES | YES | Upregulated |
| Col8a2 | ENSMUSG( | ENSMUSG( | -1.19774 | 1.94E-05 | YES | YES | Downregulated |
| Hdgfl1 | ENSMUSG( | ENSMUSG( | 1.236291 | 0.19845 | NO | NO | NA |
| Gm29243 | ENSMUSG( | ENSMUSG( | -0.17368 | 0.624649 | NO | NO | NA |
| ENSMUSG | ENSMUSG( | ENSMUSG( | 1.34919 | 0.000234 | YES | YES | Upregulated |
| Klf5 | ENSMUSG( | ENSMUSG( | 0.705022 | 0.40737 | NO | NO | NA |
| Mir100hg | ENSMUSG( | ENSMUSG( | 0.459017 | 0.663346 | NO | NO | NA |
| Gm43064 | ENSMUSG( | ENSMUSG( | 0.270982 | 0.640484 | NO | NO | NA |
| Gm48678 | ENSMUSG( | ENSMUSG( | -0.66653 | 0.052075 | NO | NO | NA |
| 4930428O | ENSMUSG( | ENSMUSG( | -0.04318 | 0.910431 | NO | NO | NA |
| ENSMUSG | ENSMUSG( | ENSMUSG( | 1.016568 | 0.180785 | NO | NO | NA |
| Vmn1r43 | ENSMUSG( | ENSMUSG( | 0.236539 | 0.801651 | NO | NO | NA |
| Cntnap2 | ENSMUSG( | ENSMUSG( | 0.467135 | 0.664907 | NO | NO | NA |
| Purg | ENSMUSG( | ENSMUSG( | -0.44026 | 0.14303 | NO | NO | NA |
| 1810041H | ENSMUSG( | ENSMUSG( | 0.404594 | 0.186863 | NO | NO | NA |
| Gm47572 | ENSMUSG( | ENSMUSG( | 0.587209 | 0.042059 | YES | NO | NA |
| A430010J1 | ENSMUSG( | ENSMUSG( | 0.631544 | 0.553686 | NO | NO | NA |
| Gm9947 | ENSMUSG( | ENSMUSG( | 1.01177 | 0.076122 | NO | NO | NA |
| Gm44913 | ENSMUSG( | ENSMUSG( | 0.818911 | 0.440997 | NO | NO | NA |
| Nat8f1 | ENSMUSG( | ENSMUSG( | -0.41425 | 0.102335 | NO | NO | NA |
| Drd1 | ENSMUSG( | ENSMUSG( | 0.160752 | 0.881144 | NO | NO | NA |
| Rps6ka2 | ENSMUSG( | ENSMUSG( | 0.203506 | 0.576644 | NO | NO | NA |
| Olfr699 | ENSMUSG( | ENSMUSG( | 0.875743 | 0.389693 | NO | NO | NA |
| Col9a3 | ENSMUSG( | ENSMUSG( | -0.05991 | 0.833646 | NO | NO | NA |
| Tmem51os | ENSMUSG( | ENSMUSG( | 0.354264 | 0.763499 | NO | NO | NA |
| Gm6793 | ENSMUSG( | ENSMUSG( | -0.08099 | 0.829186 | NO | NO | NA |

| | | | | | | | |
|---|---|---|---|---|---|---|---|
| Gm28111 | ENSMUSG( | ENSMUSG( | 0.603138 | 0.568419 | NO | NO | NA |
| Gli1 | ENSMUSG( | ENSMUSG( | 0.457703 | 0.292442 | NO | NO | NA |
| Zbtb7c | ENSMUSG( | ENSMUSG( | 0.697816 | 0.513791 | NO | NO | NA |
| Olfr307 | ENSMUSG( | ENSMUSG( | 0.797431 | 0.430295 | NO | NO | NA |
| Itgb8 | ENSMUSG( | ENSMUSG( | -0.56718 | 0.326854 | NO | NO | NA |
| Gm37464 | ENSMUSG( | ENSMUSG( | 0.801699 | 0.531552 | NO | NO | NA |
| Gm7609 | ENSMUSG( | ENSMUSG( | -0.48473 | 0.351641 | NO | NO | NA |
| Hunk | ENSMUSG( | ENSMUSG( | 0.405773 | 0.513084 | NO | NO | NA |
| Gm10710 | ENSMUSG( | ENSMUSG( | 1.091294 | 0.308893 | NO | NO | NA |
| Gm36279 | ENSMUSG( | ENSMUSG( | -0.82012 | 0.009991 | YES | NO | NA |
| Dock1 | ENSMUSG( | ENSMUSG( | 0.4119 | 0.638244 | NO | NO | NA |
| Slc12a8 | ENSMUSG( | ENSMUSG( | 0.223462 | 0.489697 | NO | NO | NA |
| Oprm1 | ENSMUSG( | ENSMUSG( | 0.706424 | 0.406566 | NO | NO | NA |
| Gm19680 | ENSMUSG( | ENSMUSG( | 0.778585 | 0.047832 | YES | NO | NA |
| Cpt1c | ENSMUSG( | ENSMUSG( | 0.49921 | 0.528062 | NO | NO | NA |
| Gm48865 | ENSMUSG( | ENSMUSG( | 0.53783 | 0.115502 | NO | NO | NA |
| Gm37094 | ENSMUSG( | ENSMUSG( | 1.194477 | 0.77071 | NO | NO | NA |
| Cep112it | ENSMUSG( | ENSMUSG( | 0.797373 | 0.135872 | NO | NO | NA |
| Des | ENSMUSG( | ENSMUSG( | 1.410481 | 0.236771 | NO | NO | NA |
| Gm38155 | ENSMUSG( | ENSMUSG( | -0.17908 | 0.624649 | NO | NO | NA |
| Gm45117 | ENSMUSG( | ENSMUSG( | 1.012607 | 0.331544 | NO | NO | NA |
| Uroc1 | ENSMUSG( | ENSMUSG( | 0.870688 | 0.453836 | NO | NO | NA |
| Nkx2-2 | ENSMUSG( | ENSMUSG( | 1.06069 | 0.288257 | NO | NO | NA |
| Gm49032 | ENSMUSG( | ENSMUSG( | 0.430065 | 0.363089 | NO | NO | NA |
| Scn1a | ENSMUSG( | ENSMUSG( | 0.604505 | 0.483112 | NO | NO | NA |
| Gm38386 | ENSMUSG( | ENSMUSG( | 0.165476 | 0.873041 | NO | NO | NA |
| C130040N | ENSMUSG( | ENSMUSG( | -0.02373 | 0.939775 | NO | NO | NA |
| Gm26549 | ENSMUSG( | ENSMUSG( | -0.15214 | 0.650266 | NO | NO | NA |
| Shisal1 | ENSMUSG( | ENSMUSG( | 0.950338 | 0.353524 | NO | NO | NA |
| Cyyr1 | ENSMUSG( | ENSMUSG( | 0.572098 | 0.518236 | NO | NO | NA |
| Nudt11 | ENSMUSG( | ENSMUSG( | 0.716113 | 0.025811 | YES | NO | NA |
| Gm42992 | ENSMUSG( | ENSMUSG( | 0.497957 | 0.639774 | NO | NO | NA |
| Pard3 | ENSMUSG( | ENSMUSG( | 0.75696 | 0.349464 | NO | NO | NA |
| Ephb2 | ENSMUSG( | ENSMUSG( | 0.211069 | 0.831934 | NO | NO | NA |
| Gm44835 | ENSMUSG( | ENSMUSG( | 0.64475 | 0.446453 | NO | NO | NA |
| ENSMUSG | ENSMUSG( | ENSMUSG( | 0.72005 | 0.355589 | NO | NO | NA |
| Arhgap44 | ENSMUSG( | ENSMUSG( | 0.295058 | 0.692623 | NO | NO | NA |
| Acvr1c | ENSMUSG( | ENSMUSG( | 0.83098 | 0.409568 | NO | NO | NA |
| Gm10310 | ENSMUSG( | ENSMUSG( | 0.77012 | 0.477095 | NO | NO | NA |
| ENSMUSG | ENSMUSG( | ENSMUSG( | -0.15649 | 0.553057 | NO | NO | NA |
| 2010001A | ENSMUSG( | ENSMUSG( | 0.113888 | 0.748574 | NO | NO | NA |
| Egfr | ENSMUSG( | ENSMUSG( | 0.636612 | 0.557183 | NO | NO | NA |
| Pcdhgb6 | ENSMUSG( | ENSMUSG( | -0.5608 | 0.162434 | NO | NO | NA |
| Vmn2r115 | ENSMUSG( | ENSMUSG( | 0.393623 | 0.734336 | NO | NO | NA |
| 4632428C( | ENSMUSG( | ENSMUSG( | -1.14719 | 0.000142 | YES | YES | Downregulated |
| Gm44022 | ENSMUSG( | ENSMUSG( | 0.465518 | 0.652876 | NO | NO | NA |
| Otx2 | ENSMUSG( | ENSMUSG( | 1.33285 | 0.226254 | NO | NO | NA |

| | | | | | | | |
|---|---|---|---|---|---|---|---|
| Lrfn1 | ENSMUSG( | ENSMUSG( | 0.76157 | 0.00649 | YES | NO | NA |
| Gm16206 | ENSMUSG( | ENSMUSG( | -0.68336 | 0.01726 | YES | NO | NA |
| Gm44065 | ENSMUSG( | ENSMUSG( | 0.69448 | 0.302959 | NO | NO | NA |
| Gm42611 | ENSMUSG( | ENSMUSG( | 0.086916 | 0.863014 | NO | NO | NA |
| Zmynd10 | ENSMUSG( | ENSMUSG( | -0.77562 | 0.002146 | YES | NO | NA |
| Snca | ENSMUSG( | ENSMUSG( | -0.21906 | 0.792476 | NO | NO | NA |
| Gm14399 | ENSMUSG( | ENSMUSG( | -0.29002 | 0.444878 | NO | NO | NA |
| BC051142 | ENSMUSG( | ENSMUSG( | 0.868396 | 0.333662 | NO | NO | NA |
| Gm33460 | ENSMUSG( | ENSMUSG( | -0.24736 | 0.442886 | NO | NO | NA |
| ENSMUSG( | ENSMUSG( | ENSMUSG( | -0.57428 | 0.041449 | YES | NO | NA |
| C920021L: | ENSMUSG( | ENSMUSG( | 0.137952 | 0.646606 | NO | NO | NA |
| Fads6 | ENSMUSG( | ENSMUSG( | 0.061135 | 0.859836 | NO | NO | NA |
| Gm15832 | ENSMUSG( | ENSMUSG( | -1.00833 | 0.000601 | YES | YES | Downregulated |
| Igkv6-14 | ENSMUSG( | ENSMUSG( | -0.22382 | 0.937181 | NO | NO | NA |
| Mtcl1 | ENSMUSG( | ENSMUSG( | -0.07088 | 0.946061 | NO | NO | NA |
| Zic5 | ENSMUSG( | ENSMUSG( | 0.770794 | 0.178256 | NO | NO | NA |
| Klhl33 | ENSMUSG( | ENSMUSG( | 0.482402 | 0.581457 | NO | NO | NA |
| Lurap1l | ENSMUSG( | ENSMUSG( | 0.527262 | 0.649329 | NO | NO | NA |
| Gm8250 | ENSMUSG( | ENSMUSG( | 0.413839 | 0.177261 | NO | NO | NA |
| Gm44752 | ENSMUSG( | ENSMUSG( | -0.08658 | 0.803948 | NO | NO | NA |
| Bean1 | ENSMUSG( | ENSMUSG( | 0.239308 | 0.794099 | NO | NO | NA |
| Perm1 | ENSMUSG( | ENSMUSG( | -0.63753 | 0.062394 | NO | NO | NA |
| Grk1 | ENSMUSG( | ENSMUSG( | 0.602145 | 0.43299 | NO | NO | NA |
| Vwf | ENSMUSG( | ENSMUSG( | 0.209794 | 0.748515 | NO | NO | NA |
| Atf7ip2 | ENSMUSG( | ENSMUSG( | 0.875249 | 0.044045 | YES | NO | NA |
| Dbx2 | ENSMUSG( | ENSMUSG( | 0.71796 | 0.476823 | NO | NO | NA |
| Gm48231 | ENSMUSG( | ENSMUSG( | 1.060876 | 0.241144 | NO | NO | NA |
| Rax | ENSMUSG( | ENSMUSG( | 1.118915 | 0.220868 | NO | NO | NA |
| Redrum | ENSMUSG( | ENSMUSG( | 0.727489 | 0.359969 | NO | NO | NA |
| Acvr2b | ENSMUSG( | ENSMUSG( | 0.269413 | 0.691861 | NO | NO | NA |
| Jag1 | ENSMUSG( | ENSMUSG( | 0.760826 | 0.24806 | NO | NO | NA |
| Gpr34 | ENSMUSG( | ENSMUSG( | 1.849636 | 3.51E-05 | YES | YES | Upregulated |
| Gm43633 | ENSMUSG( | ENSMUSG( | 0.197339 | 0.642321 | NO | NO | NA |
| 4930432E: | ENSMUSG( | ENSMUSG( | 0.461484 | 0.645355 | NO | NO | NA |
| Gm37314 | ENSMUSG( | ENSMUSG( | 0.224658 | 0.799508 | NO | NO | NA |
| Gm49510 | ENSMUSG( | ENSMUSG( | 0.801294 | 0.050269 | NO | NO | NA |
| H2ac7 | ENSMUSG( | ENSMUSG( | 0.450417 | 0.364588 | NO | NO | NA |
| Gm48391 | ENSMUSG( | ENSMUSG( | 0.925421 | 0.392463 | NO | NO | NA |
| Gm20033 | ENSMUSG( | ENSMUSG( | -0.85968 | 0.008732 | YES | NO | NA |
| Gm43462 | ENSMUSG( | ENSMUSG( | 0.148391 | 0.74256 | NO | NO | NA |
| H2bc9 | ENSMUSG( | ENSMUSG( | 0.372855 | 0.260117 | NO | NO | NA |
| Gm13237 | ENSMUSG( | ENSMUSG( | 0.252446 | 0.352248 | NO | NO | NA |
| Camkk1 | ENSMUSG( | ENSMUSG( | -0.38793 | 0.218945 | NO | NO | NA |
| Gm47963 | ENSMUSG( | ENSMUSG( | -0.50033 | 0.182707 | NO | NO | NA |
| Olfr23 | ENSMUSG( | ENSMUSG( | 1.0586 | 0.270441 | NO | NO | NA |
| 94300530 | ENSMUSG( | ENSMUSG( | 0.809545 | 0.468284 | NO | NO | NA |
| Lrrtm1 | ENSMUSG( | ENSMUSG( | 0.679019 | 0.518553 | NO | NO | NA |

| | | | | | | | |
|---|---|---|---|---|---|---|---|
| Igkv12-46 | ENSMUSG( | ENSMUSG( | -0.46877 | 0.735713 | NO | NO | NA |
| Txnrd3 | ENSMUSG( | ENSMUSG( | 1.334186 | 0.002004 | YES | YES | Upregulated |
| Gm49371 | ENSMUSG( | ENSMUSG( | 0.757075 | 0.4831 | NO | NO | NA |
| Gm37530 | ENSMUSG( | ENSMUSG( | 0.949525 | 0.167626 | NO | NO | NA |
| Gjb2 | ENSMUSG( | ENSMUSG( | -0.39726 | 0.151822 | NO | NO | NA |
| Ina | ENSMUSG( | ENSMUSG( | 0.377713 | 0.499353 | NO | NO | NA |
| Ednrb | ENSMUSG( | ENSMUSG( | 0.963539 | 0.374168 | NO | NO | NA |
| Slc18a3 | ENSMUSG( | ENSMUSG( | 0.771045 | 0.398656 | NO | NO | NA |
| Cntn1 | ENSMUSG( | ENSMUSG( | 0.510982 | 0.575942 | NO | NO | NA |
| Gm45675 | ENSMUSG( | ENSMUSG( | 0.703295 | 0.536092 | NO | NO | NA |
| Plcd4 | ENSMUSG( | ENSMUSG( | -0.18406 | 0.518102 | NO | NO | NA |
| Gm5315 | ENSMUSG( | ENSMUSG( | 0.64951 | 0.566336 | NO | NO | NA |
| 9230104M | ENSMUSG( | ENSMUSG( | -0.1356 | 0.782297 | NO | NO | NA |
| Lcn2 | ENSMUSG( | ENSMUSG( | 2.891081 | 0.239055 | NO | NO | NA |
| Mdm4-ps | ENSMUSG( | ENSMUSG( | 1.026022 | 0.274415 | NO | NO | NA |
| Ebf2 | ENSMUSG( | ENSMUSG( | 0.658791 | 0.492717 | NO | NO | NA |
| Gm19696 | ENSMUSG( | ENSMUSG( | 0.750853 | 0.417592 | NO | NO | NA |
| Lmx1a | ENSMUSG( | ENSMUSG( | 0.570907 | 0.599029 | NO | NO | NA |
| Trav11 | ENSMUSG( | ENSMUSG( | -5.12094 | 0.008443 | YES | YES | Downregulated |
| Gm43430 | ENSMUSG( | ENSMUSG( | 0.764404 | 0.147988 | NO | NO | NA |
| St6galnac2 | ENSMUSG( | ENSMUSG( | 0.68832 | 0.009078 | YES | NO | NA |
| Mansc1 | ENSMUSG( | ENSMUSG( | -0.10315 | 0.727144 | NO | NO | NA |
| Ptgs1 | ENSMUSG( | ENSMUSG( | 0.080254 | 0.831994 | NO | NO | NA |
| Gm43336 | ENSMUSG( | ENSMUSG( | -0.35606 | 0.552765 | NO | NO | NA |
| Syp | ENSMUSG( | ENSMUSG( | 0.187246 | 0.73993 | NO | NO | NA |
| Tbx5 | ENSMUSG( | ENSMUSG( | 0.588115 | 0.575669 | NO | NO | NA |
| Thra | ENSMUSG( | ENSMUSG( | 0.184125 | 0.55607 | NO | NO | NA |
| Tenm1 | ENSMUSG( | ENSMUSG( | 0.517421 | 0.51145 | NO | NO | NA |
| Man2c1os | ENSMUSG( | ENSMUSG( | -0.38287 | 0.211661 | NO | NO | NA |
| Vmn1r29 | ENSMUSG( | ENSMUSG( | 0.696421 | 0.456867 | NO | NO | NA |
| Rpl39l | ENSMUSG( | ENSMUSG( | 2.284057 | 0.005673 | YES | YES | Upregulated |
| Olfr283 | ENSMUSG( | ENSMUSG( | 0.63634 | 0.572224 | NO | NO | NA |
| Gm45652 | ENSMUSG( | ENSMUSG( | 0.721787 | 0.507122 | NO | NO | NA |
| Gm5831 | ENSMUSG( | ENSMUSG( | 0.06632 | 0.843187 | NO | NO | NA |
| Peg3 | ENSMUSG( | ENSMUSG( | 1.451189 | 0.000393 | YES | YES | Upregulated |
| Nrep | ENSMUSG( | ENSMUSG( | 0.512261 | 0.625621 | NO | NO | NA |
| Zfp428 | ENSMUSG( | ENSMUSG( | 0.662507 | 0.042016 | YES | NO | NA |
| Bicc1 | ENSMUSG( | ENSMUSG( | 0.793863 | 0.449888 | NO | NO | NA |
| Adgra3 | ENSMUSG( | ENSMUSG( | 0.68272 | 0.3754 | NO | NO | NA |
| Gm10132 | ENSMUSG( | ENSMUSG( | 0.19367 | 0.652117 | NO | NO | NA |
| Tspo2 | ENSMUSG( | ENSMUSG( | 1.01656 | 0.138116 | NO | NO | NA |
| Gzmd | ENSMUSG( | ENSMUSG( | 7.093553 | 3.38E-06 | YES | YES | Upregulated |
| Gm38211 | ENSMUSG( | ENSMUSG( | 1.021463 | 0.334253 | NO | NO | NA |
| Hmga1b | ENSMUSG( | ENSMUSG( | 0.224595 | 0.455 | NO | NO | NA |
| Susd4 | ENSMUSG( | ENSMUSG( | 0.949443 | 0.168915 | NO | NO | NA |
| Cfap126 | ENSMUSG( | ENSMUSG( | 0.131988 | 0.591301 | NO | NO | NA |
| Cnpy1 | ENSMUSG( | ENSMUSG( | 0.815609 | 0.422996 | NO | NO | NA |

| | | | | | | | |
|---|---|---|---|---|---|---|---|
| Six1 | ENSMUSG( | ENSMUSG( | 0.690402 | 0.338874 | NO | NO | NA |
| Gm34455 | ENSMUSG( | ENSMUSG( | 1.431285 | 0.085841 | NO | NO | NA |
| Acyp2 | ENSMUSG( | ENSMUSG( | 0.227185 | 0.403874 | NO | NO | NA |
| Gm7285 | ENSMUSG( | ENSMUSG( | -0.01112 | 0.976422 | NO | NO | NA |
| Gm5511 | ENSMUSG( | ENSMUSG( | -0.70602 | 0.000913 | YES | NO | NA |
| 2200002D | ENSMUSG( | ENSMUSG( | 0.947583 | 0.00026 | YES | NO | NA |
| Zfp493 | ENSMUSG( | ENSMUSG( | -0.3273 | 0.405222 | NO | NO | NA |
| A130048G | ENSMUSG( | ENSMUSG( | -0.20595 | 0.78461 | NO | NO | NA |
| Phf11d | ENSMUSG( | ENSMUSG( | -0.31729 | 0.425849 | NO | NO | NA |
| Nsg2 | ENSMUSG( | ENSMUSG( | -0.11059 | 0.855874 | NO | NO | NA |
| ENSMUSG( | ENSMUSG( | ENSMUSG( | 0.528394 | 0.474914 | NO | NO | NA |
| Gm45751 | ENSMUSG( | ENSMUSG( | 0.866227 | 0.492443 | NO | NO | NA |
| Gja5 | ENSMUSG( | ENSMUSG( | 0.794689 | 0.396816 | NO | NO | NA |
| A330076C | ENSMUSG( | ENSMUSG( | 0.714325 | 0.491395 | NO | NO | NA |
| ENSMUSG | ENSMUSG( | ENSMUSG( | -0.19918 | 0.522159 | NO | NO | NA |
| Kcnj2 | ENSMUSG( | ENSMUSG( | 0.723882 | 0.449679 | NO | NO | NA |
| Cygb | ENSMUSG( | ENSMUSG( | 0.922753 | 0.356564 | NO | NO | NA |
| A430019L( | ENSMUSG( | ENSMUSG( | 0.883594 | 0.232063 | NO | NO | NA |
| Gm31763 | ENSMUSG( | ENSMUSG( | -0.09223 | 0.756346 | NO | NO | NA |
| Gm38380 | ENSMUSG( | ENSMUSG( | 0.368955 | 0.555214 | NO | NO | NA |
| Unc5a | ENSMUSG( | ENSMUSG( | 0.136842 | 0.723669 | NO | NO | NA |
| Eid3 | ENSMUSG( | ENSMUSG( | -0.0436 | 0.907269 | NO | NO | NA |
| Colec12 | ENSMUSG( | ENSMUSG( | 0.655769 | 0.435651 | NO | NO | NA |
| Ccdc38 | ENSMUSG( | ENSMUSG( | -0.25482 | 0.552241 | NO | NO | NA |
| Gm43061 | ENSMUSG( | ENSMUSG( | 0.746629 | 0.345985 | NO | NO | NA |
| Gm37082 | ENSMUSG( | ENSMUSG( | -0.79825 | 0.197041 | NO | NO | NA |
| Gm37787 | ENSMUSG( | ENSMUSG( | -0.49546 | 0.252323 | NO | NO | NA |
| Gm48430 | ENSMUSG( | ENSMUSG( | 1.10946 | 0.282727 | NO | NO | NA |
| Atp7b | ENSMUSG( | ENSMUSG( | 0.765348 | 0.128996 | NO | NO | NA |
| Fam163a | ENSMUSG( | ENSMUSG( | 0.760931 | 0.49586 | NO | NO | NA |
| 2900005J1 | ENSMUSG( | ENSMUSG( | 0.70724 | 0.321293 | NO | NO | NA |
| Trav13d-2 | ENSMUSG( | ENSMUSG( | 0.531531 | 0.855635 | NO | NO | NA |
| Tgm2 | ENSMUSG( | ENSMUSG( | -0.2262 | 0.585511 | NO | NO | NA |
| Svep1 | ENSMUSG( | ENSMUSG( | 0.505002 | 0.576644 | NO | NO | NA |
| Gm8203 | ENSMUSG( | ENSMUSG( | 0.46319 | 0.118986 | NO | NO | NA |
| Gm10388 | ENSMUSG( | ENSMUSG( | 0.530918 | 0.250124 | NO | NO | NA |
| Gm5441 | ENSMUSG( | ENSMUSG( | 0.206671 | 0.854754 | NO | NO | NA |
| 1810044D | ENSMUSG( | ENSMUSG( | 0.916975 | 0.012915 | YES | NO | NA |
| Ccdc173 | ENSMUSG( | ENSMUSG( | 0.28774 | 0.30402 | NO | NO | NA |
| Olfr251 | ENSMUSG( | ENSMUSG( | 0.47777 | 0.653824 | NO | NO | NA |
| H3c7 | ENSMUSG( | ENSMUSG( | 0.473935 | 0.209909 | NO | NO | NA |
| Gm42777 | ENSMUSG( | ENSMUSG( | 0.715974 | 0.507436 | NO | NO | NA |
| Sptbn2 | ENSMUSG( | ENSMUSG( | 0.88941 | 0.002619 | YES | NO | NA |
| Peg12 | ENSMUSG( | ENSMUSG( | 0.381982 | 0.627394 | NO | NO | NA |
| Gm37072 | ENSMUSG( | ENSMUSG( | 1.004668 | 0.345824 | NO | NO | NA |
| Sfrp4 | ENSMUSG( | ENSMUSG( | 0.685782 | 0.391427 | NO | NO | NA |
| A430028G | ENSMUSG( | ENSMUSG( | 0.391515 | 0.115584 | NO | NO | NA |

| | | | | | | | |
|---|---|---|---|---|---|---|---|
| Gm6552 | ENSMUSG( | ENSMUSG( | 0.197653 | 0.441069 | NO | NO | NA |
| Lrrc73 | ENSMUSG( | ENSMUSG( | 0.47452 | 0.060958 | NO | NO | NA |
| Gm48027 | ENSMUSG( | ENSMUSG( | 0.39525 | 0.434361 | NO | NO | NA |
| Zfp423 | ENSMUSG( | ENSMUSG( | 0.717295 | 0.552728 | NO | NO | NA |
| Gm50139 | ENSMUSG( | ENSMUSG( | 0.54459 | 0.893108 | NO | NO | NA |
| Lhfp | ENSMUSG( | ENSMUSG( | 0.437087 | 0.669364 | NO | NO | NA |
| Gm41764 | ENSMUSG( | ENSMUSG( | 0.492158 | 0.659188 | NO | NO | NA |
| Peg10 | ENSMUSG( | ENSMUSG( | 0.636843 | 0.524076 | NO | NO | NA |
| Gm10767 | ENSMUSG( | ENSMUSG( | -0.32444 | 0.158327 | NO | NO | NA |
| Ninj2 | ENSMUSG( | ENSMUSG( | 0.364451 | 0.373338 | NO | NO | NA |
| Gm26737 | ENSMUSG( | ENSMUSG( | 0.527492 | 0.471208 | NO | NO | NA |
| Trav13-2 | ENSMUSG( | ENSMUSG( | 0.477266 | 0.600191 | NO | NO | NA |
| H2bc18 | ENSMUSG( | ENSMUSG( | 0.924555 | 0.239551 | NO | NO | NA |
| Gm35040 | ENSMUSG( | ENSMUSG( | 0.640582 | 0.534005 | NO | NO | NA |
| Pou2af1 | ENSMUSG( | ENSMUSG( | 0.333842 | 0.497276 | NO | NO | NA |
| Gm34678 | ENSMUSG( | ENSMUSG( | 0.236938 | 0.584776 | NO | NO | NA |
| Gm47939 | ENSMUSG( | ENSMUSG( | 0.322458 | 0.734016 | NO | NO | NA |
| Ctdspl | ENSMUSG( | ENSMUSG( | 0.337716 | 0.55058 | NO | NO | NA |
| Atl1 | ENSMUSG( | ENSMUSG( | 0.622538 | 0.155052 | NO | NO | NA |
| 6430710M | ENSMUSG( | ENSMUSG( | 0.468771 | 0.540196 | NO | NO | NA |
| Gm47472 | ENSMUSG( | ENSMUSG( | 0.629793 | 0.57346 | NO | NO | NA |
| Tmem132a | ENSMUSG( | ENSMUSG( | 0.917762 | 0.008844 | YES | NO | NA |
| Cxxc5 | ENSMUSG( | ENSMUSG( | 0.257126 | 0.449445 | NO | NO | NA |
| Igkv5-43 | ENSMUSG( | ENSMUSG( | 1.920411 | NA | NO | NO | NA |
| Sult6b1 | ENSMUSG( | ENSMUSG( | -0.95857 | 0.00037 | YES | NO | NA |
| Nckap5 | ENSMUSG( | ENSMUSG( | 0.172267 | 0.862699 | NO | NO | NA |
| Gm44567 | ENSMUSG( | ENSMUSG( | 0.07369 | 0.886027 | NO | NO | NA |
| Ccr8 | ENSMUSG( | ENSMUSG( | 0.74551 | 0.071745 | NO | NO | NA |
| Gm2366 | ENSMUSG( | ENSMUSG( | 1.070595 | 0.29526 | NO | NO | NA |
| Dok4 | ENSMUSG( | ENSMUSG( | -0.16339 | 0.617905 | NO | NO | NA |
| A830018L1 | ENSMUSG( | ENSMUSG( | 0.718667 | 0.506637 | NO | NO | NA |
| ENSMUSG | ENSMUSG( | ENSMUSG( | -0.36835 | 0.289688 | NO | NO | NA |
| Adora1 | ENSMUSG( | ENSMUSG( | 0.595411 | 0.569142 | NO | NO | NA |
| ENSMUSG( | ENSMUSG( | ENSMUSG( | -0.30128 | 0.810036 | NO | NO | NA |
| Gm38070 | ENSMUSG( | ENSMUSG( | 0.897508 | 0.289767 | NO | NO | NA |
| Tlr2 | ENSMUSG( | ENSMUSG( | 0.043922 | 0.932228 | NO | NO | NA |
| Car7 | ENSMUSG( | ENSMUSG( | -0.10491 | 0.73361 | NO | NO | NA |
| Cfap65 | ENSMUSG( | ENSMUSG( | 0.655237 | 0.536182 | NO | NO | NA |
| 6030442K2 | ENSMUSG( | ENSMUSG( | 0.462497 | 0.371084 | NO | NO | NA |
| Mapk8ip1 | ENSMUSG( | ENSMUSG( | -0.56681 | 0.085382 | NO | NO | NA |
| Ptges3-ps | ENSMUSG( | ENSMUSG( | 0.189722 | 0.46955 | NO | NO | NA |
| Slc38a5 | ENSMUSG( | ENSMUSG( | 1.193891 | 0.032408 | YES | YES | Upregulated |
| ENSMUSG | ENSMUSG( | ENSMUSG( | -0.0879 | 0.878431 | NO | NO | NA |
| ENSMUSG | ENSMUSG( | ENSMUSG( | 0.420958 | 0.26451 | NO | NO | NA |
| ENSMUSG | ENSMUSG( | ENSMUSG( | -0.10537 | 0.800749 | NO | NO | NA |
| Gldn | ENSMUSG( | ENSMUSG( | 0.407836 | 0.706703 | NO | NO | NA |
| 1700012D | ENSMUSG( | ENSMUSG( | 0.255842 | 0.499987 | NO | NO | NA |

| | | | | | | | |
|---|---|---|---|---|---|---|---|
| Rps24-ps3 | ENSMUSG( | ENSMUSG( | 0.135721 | 0.694528 | NO | NO | NA |
| Gm35439 | ENSMUSG( | ENSMUSG( | 0.64286 | 0.483029 | NO | NO | NA |
| Cutal | ENSMUSG( | ENSMUSG( | 0.405953 | 0.686701 | NO | NO | NA |
| H4c6 | ENSMUSG( | ENSMUSG( | 0.387379 | 0.358335 | NO | NO | NA |
| A330102I1 | ENSMUSG( | ENSMUSG( | 1.560077 | 0.093973 | NO | NO | NA |
| 1700018A( | ENSMUSG( | ENSMUSG( | 0.54393 | 0.610556 | NO | NO | NA |
| Gm4724 | ENSMUSG( | ENSMUSG( | 0.421482 | 0.455036 | NO | NO | NA |
| Six5 | ENSMUSG( | ENSMUSG( | 0.191824 | 0.653047 | NO | NO | NA |
| E230029C( | ENSMUSG( | ENSMUSG( | 0.450507 | 0.280164 | NO | NO | NA |
| Slc22a3 | ENSMUSG( | ENSMUSG( | 0.952419 | 0.285299 | NO | NO | NA |
| Gm44645 | ENSMUSG( | ENSMUSG( | 0.697299 | 0.516453 | NO | NO | NA |
| Gm43652 | ENSMUSG( | ENSMUSG( | 1.072872 | 0.216209 | NO | NO | NA |
| Gm38157 | ENSMUSG( | ENSMUSG( | 0.803346 | 0.388746 | NO | NO | NA |
| Prrxl1 | ENSMUSG( | ENSMUSG( | 0.570287 | 0.584053 | NO | NO | NA |
| D830024N | ENSMUSG( | ENSMUSG( | 0.003061 | 0.995252 | NO | NO | NA |
| Zfp467 | ENSMUSG( | ENSMUSG( | -1.41014 | 0.002492 | YES | YES | Downregulated |
| Bbof1 | ENSMUSG( | ENSMUSG( | 1.127573 | 0.00106 | YES | YES | Upregulated |
| Plekha6 | ENSMUSG( | ENSMUSG( | -1.46552 | 1.77E-05 | YES | YES | Downregulated |
| Gm48904 | ENSMUSG( | ENSMUSG( | 0.264432 | 0.440587 | NO | NO | NA |
| Cacna2d3 | ENSMUSG( | ENSMUSG( | 0.618718 | 0.579864 | NO | NO | NA |
| Gm16464 | ENSMUSG( | ENSMUSG( | -1.64014 | 9.05E-11 | YES | YES | Downregulated |
| 6720427I0 | ENSMUSG( | ENSMUSG( | -0.49063 | 0.039167 | YES | NO | NA |
| Irs1 | ENSMUSG( | ENSMUSG( | 0.262502 | 0.792398 | NO | NO | NA |
| Gm44975 | ENSMUSG( | ENSMUSG( | 0.464138 | 0.660436 | NO | NO | NA |
| Fzd10 | ENSMUSG( | ENSMUSG( | 0.646855 | 0.34315 | NO | NO | NA |
| Vmn1r209 | ENSMUSG( | ENSMUSG( | 0.47853 | 0.643061 | NO | NO | NA |
| Gm45546 | ENSMUSG( | ENSMUSG( | 0.94587 | 0.009438 | YES | NO | NA |
| Olfr64 | ENSMUSG( | ENSMUSG( | 0.628303 | 0.577302 | NO | NO | NA |
| Strc | ENSMUSG( | ENSMUSG( | -1.05181 | 0.132345 | NO | NO | NA |
| Gm15792 | ENSMUSG( | ENSMUSG( | 0.571884 | 0.022537 | YES | NO | NA |
| Adap2os | ENSMUSG( | ENSMUSG( | 0.862674 | 0.402024 | NO | NO | NA |
| Tes3-ps | ENSMUSG( | ENSMUSG( | 0.271773 | 0.616448 | NO | NO | NA |
| Map1b | ENSMUSG( | ENSMUSG( | 0.244502 | 0.711964 | NO | NO | NA |
| Sall3 | ENSMUSG( | ENSMUSG( | 0.434404 | 0.62909 | NO | NO | NA |
| Fbxo43 | ENSMUSG( | ENSMUSG( | 0.887926 | 0.425871 | NO | NO | NA |
| E330032C: | ENSMUSG( | ENSMUSG( | 0.900137 | 0.173362 | NO | NO | NA |
| Dnaaf3 | ENSMUSG( | ENSMUSG( | 0.861303 | 0.041541 | YES | NO | NA |
| Gm37488 | ENSMUSG( | ENSMUSG( | -0.62754 | 0.019652 | YES | NO | NA |
| Gm37522 | ENSMUSG( | ENSMUSG( | 0.553729 | 0.552744 | NO | NO | NA |
| D430013B | ENSMUSG( | ENSMUSG( | -0.00386 | 0.989667 | NO | NO | NA |
| Gm43421 | ENSMUSG( | ENSMUSG( | 0.463837 | 0.517295 | NO | NO | NA |
| Gpr179 | ENSMUSG( | ENSMUSG( | 0.119904 | 0.836189 | NO | NO | NA |
| Bloc1s6os | ENSMUSG( | ENSMUSG( | 0.466516 | 0.428394 | NO | NO | NA |
| Lmo1 | ENSMUSG( | ENSMUSG( | -0.10919 | 0.847167 | NO | NO | NA |
| Sash1 | ENSMUSG( | ENSMUSG( | 0.019709 | 0.978064 | NO | NO | NA |
| Npl | ENSMUSG( | ENSMUSG( | 0.957011 | 0.064123 | NO | NO | NA |
| Gm43288 | ENSMUSG( | ENSMUSG( | 0.875806 | 0.271956 | NO | NO | NA |

| | | | | | | | |
|---|---|---|---|---|---|---|---|
| Gm43848 | ENSMUSG( | ENSMUSG( | 0.215562 | 0.517147 | NO | NO | NA |
| Fhdc1 | ENSMUSG( | ENSMUSG( | 0.470498 | 0.305459 | NO | NO | NA |
| Samd5 | ENSMUSG( | ENSMUSG( | 0.596662 | 0.581222 | NO | NO | NA |
| Pou3f2 | ENSMUSG( | ENSMUSG( | 0.74577 | 0.498019 | NO | NO | NA |
| Vmn1r81 | ENSMUSG( | ENSMUSG( | 0.783925 | 0.467168 | NO | NO | NA |
| Pax5 | ENSMUSG( | ENSMUSG( | 0.487562 | 0.631498 | NO | NO | NA |
| A530083I2 | ENSMUSG( | ENSMUSG( | 0.598762 | 0.379041 | NO | NO | NA |
| Pkhd1 | ENSMUSG( | ENSMUSG( | 0.85221 | 0.423238 | NO | NO | NA |
| Gm9913 | ENSMUSG( | ENSMUSG( | 0.851201 | 0.397233 | NO | NO | NA |
| Cacna1b | ENSMUSG( | ENSMUSG( | 0.381195 | 0.14602 | NO | NO | NA |
| Ctxn1 | ENSMUSG( | ENSMUSG( | 0.505525 | 0.109506 | NO | NO | NA |
| Lgr6 | ENSMUSG( | ENSMUSG( | 0.371068 | 0.504561 | NO | NO | NA |
| Cped1 | ENSMUSG( | ENSMUSG( | 1.072132 | 0.290434 | NO | NO | NA |
| Gm49504 | ENSMUSG( | ENSMUSG( | 0.573984 | 0.579864 | NO | NO | NA |
| ENSMUSG( | ENSMUSG( | ENSMUSG( | 0.969832 | 0.383975 | NO | NO | NA |
| Trim67 | ENSMUSG( | ENSMUSG( | 0.516196 | 0.582542 | NO | NO | NA |
| Akap7 | ENSMUSG( | ENSMUSG( | 0.495874 | 0.155265 | NO | NO | NA |
| Gm48369 | ENSMUSG( | ENSMUSG( | 0.254299 | 0.763451 | NO | NO | NA |
| Adarb2 | ENSMUSG( | ENSMUSG( | 0.389932 | 0.553609 | NO | NO | NA |
| Gm8663 | ENSMUSG( | ENSMUSG( | 0.075075 | 0.822616 | NO | NO | NA |
| Gm49979 | ENSMUSG( | ENSMUSG( | 0.648415 | 0.081891 | NO | NO | NA |
| 2810428J( | ENSMUSG( | ENSMUSG( | 0.135204 | 0.753767 | NO | NO | NA |
| Rab3il1 | ENSMUSG( | ENSMUSG( | 0.470221 | 0.385584 | NO | NO | NA |
| Gm38252 | ENSMUSG( | ENSMUSG( | 0.468608 | 0.663341 | NO | NO | NA |
| Pkd2 | ENSMUSG( | ENSMUSG( | 0.527802 | 0.099661 | NO | NO | NA |
| Ighv6-6 | ENSMUSG( | ENSMUSG( | -1.27102 | 0.69431 | NO | NO | NA |
| Gm43703 | ENSMUSG( | ENSMUSG( | 0.035586 | 0.935652 | NO | NO | NA |
| Kiss1r | ENSMUSG( | ENSMUSG( | -0.02192 | 0.956252 | NO | NO | NA |
| H2-M5 | ENSMUSG( | ENSMUSG( | 0.649068 | 0.438227 | NO | NO | NA |
| Vmn2r100 | ENSMUSG( | ENSMUSG( | 0.091043 | 0.93076 | NO | NO | NA |
| Ptges3l | ENSMUSG( | ENSMUSG( | 0.073389 | 0.807942 | NO | NO | NA |
| Trav12d-1 | ENSMUSG( | ENSMUSG( | -1.36821 | 0.665935 | NO | NO | NA |
| Gm9443 | ENSMUSG( | ENSMUSG( | 0.979664 | 0.336863 | NO | NO | NA |
| Gm11007 | ENSMUSG( | ENSMUSG( | -6.77662 | 0.070955 | NO | NO | NA |
| Nwd1 | ENSMUSG( | ENSMUSG( | 0.154265 | 0.893921 | NO | NO | NA |
| Tcp11 | ENSMUSG( | ENSMUSG( | -0.69292 | 0.020112 | YES | NO | NA |
| Gm49137 | ENSMUSG( | ENSMUSG( | 1.038978 | 0.324751 | NO | NO | NA |
| Elapor1 | ENSMUSG( | ENSMUSG( | -0.13457 | 0.743501 | NO | NO | NA |
| Cdx2 | ENSMUSG( | ENSMUSG( | 0.550911 | 0.60191 | NO | NO | NA |
| H2ac15 | ENSMUSG( | ENSMUSG( | 0.373033 | 0.144025 | NO | NO | NA |
| Snx7 | ENSMUSG( | ENSMUSG( | 0.813863 | 0.156284 | NO | NO | NA |
| Gm34885 | ENSMUSG( | ENSMUSG( | 0.643864 | 0.375234 | NO | NO | NA |
| Ighv5-16 | ENSMUSG( | ENSMUSG( | -5.02905 | 0.008723 | YES | YES | Downregulated |
| Vmn2r60 | ENSMUSG( | ENSMUSG( | 0.824272 | 0.288287 | NO | NO | NA |
| Gm5421 | ENSMUSG( | ENSMUSG( | 0.370775 | 0.200949 | NO | NO | NA |
| Trav4n-3 | ENSMUSG( | ENSMUSG( | -0.17731 | 0.89964 | NO | NO | NA |
| Gm29094 | ENSMUSG( | ENSMUSG( | -0.16107 | 0.634312 | NO | NO | NA |

| | | | | | | | |
|---|---|---|---|---|---|---|---|
| A230060F: | ENSMUSG( | ENSMUSG( | 0.028596 | 0.927457 | NO | NO | NA |
| Olfr1271 | ENSMUSG( | ENSMUSG( | 0.640421 | 0.547577 | NO | NO | NA |
| Mlip | ENSMUSG( | ENSMUSG( | 0.808003 | 0.468862 | NO | NO | NA |
| Gm45342 | ENSMUSG( | ENSMUSG( | -0.51454 | 0.259199 | NO | NO | NA |
| Gm37423 | ENSMUSG( | ENSMUSG( | 0.41456 | 0.349957 | NO | NO | NA |
| Gm45737 | ENSMUSG( | ENSMUSG( | 0.239597 | 0.508487 | NO | NO | NA |
| Sh3pxd2b | ENSMUSG( | ENSMUSG( | 0.716331 | 0.142842 | NO | NO | NA |
| Gm44435 | ENSMUSG( | ENSMUSG( | 0.379387 | 0.697879 | NO | NO | NA |
| 9530086O | ENSMUSG( | ENSMUSG( | 0.674674 | 0.513101 | NO | NO | NA |
| Gm37255 | ENSMUSG( | ENSMUSG( | -0.25148 | 0.596783 | NO | NO | NA |
| Gm37302 | ENSMUSG( | ENSMUSG( | 0.666528 | 0.537294 | NO | NO | NA |
| Ngfr | ENSMUSG( | ENSMUSG( | -1.91199 | 1.16E-06 | YES | YES | Downregulated |
| Itga10 | ENSMUSG( | ENSMUSG( | -0.9274 | 0.031151 | YES | NO | NA |
| Gm45844 | ENSMUSG( | ENSMUSG( | 0.703055 | 0.358045 | NO | NO | NA |
| Gm48713 | ENSMUSG( | ENSMUSG( | 0.816728 | 0.480606 | NO | NO | NA |
| Lsamp | ENSMUSG( | ENSMUSG( | 0.776081 | 0.446235 | NO | NO | NA |
| Siglecf | ENSMUSG( | ENSMUSG( | 0.839887 | 0.198254 | NO | NO | NA |
| Lgmn | ENSMUSG( | ENSMUSG( | 0.196739 | 0.546202 | NO | NO | NA |
| ENSMUSG( | ENSMUSG( | ENSMUSG( | 0.876038 | 0.422508 | NO | NO | NA |
| Gm15903 | ENSMUSG( | ENSMUSG( | 0.434448 | 0.476831 | NO | NO | NA |
| Ranbp3l | ENSMUSG( | ENSMUSG( | 0.808983 | 0.25243 | NO | NO | NA |
| Vmn1r82 | ENSMUSG( | ENSMUSG( | 0.700461 | 0.531711 | NO | NO | NA |
| Ahcyl | ENSMUSG( | ENSMUSG( | 0.403971 | 0.158522 | NO | NO | NA |
| Hoxc8 | ENSMUSG( | ENSMUSG( | 1.136495 | 0.130858 | NO | NO | NA |
| Tnfsf18 | ENSMUSG( | ENSMUSG( | 1.012751 | 0.199474 | NO | NO | NA |
| Gm45640 | ENSMUSG( | ENSMUSG( | -0.384 | 0.320055 | NO | NO | NA |
| Gm43621 | ENSMUSG( | ENSMUSG( | 0.778248 | 0.496358 | NO | NO | NA |
| Gm47857 | ENSMUSG( | ENSMUSG( | 0.553137 | 0.512821 | NO | NO | NA |
| Gm26724 | ENSMUSG( | ENSMUSG( | 0.054771 | 0.904283 | NO | NO | NA |
| Gm20655 | ENSMUSG( | ENSMUSG( | 0.497927 | 0.602228 | NO | NO | NA |
| Psmb11 | ENSMUSG( | ENSMUSG( | 0.903839 | 0.377957 | NO | NO | NA |
| Fam83g | ENSMUSG( | ENSMUSG( | 0.162629 | 0.609962 | NO | NO | NA |
| Gm45153 | ENSMUSG( | ENSMUSG( | 0.936867 | 0.301897 | NO | NO | NA |
| Arhgap5 | ENSMUSG( | ENSMUSG( | 0.440371 | 0.203782 | NO | NO | NA |
| Gstm5 | ENSMUSG( | ENSMUSG( | 1.284878 | 0.009771 | YES | YES | Upregulated |
| Gm50388 | ENSMUSG( | ENSMUSG( | -0.17554 | 0.897447 | NO | NO | NA |
| 5730409E( | ENSMUSG( | ENSMUSG( | 0.5884 | 0.155527 | NO | NO | NA |
| 6330409D | ENSMUSG( | ENSMUSG( | -0.25449 | 0.444105 | NO | NO | NA |
| D030040B | ENSMUSG( | ENSMUSG( | 0.883745 | 0.401746 | NO | NO | NA |
| Gm44901 | ENSMUSG( | ENSMUSG( | 0.387618 | 0.460937 | NO | NO | NA |
| ENSMUSG | ENSMUSG( | ENSMUSG( | 0.288618 | 0.756346 | NO | NO | NA |
| A930001M | ENSMUSG( | ENSMUSG( | 0.165069 | 0.746663 | NO | NO | NA |
| 4930470G | ENSMUSG( | ENSMUSG( | 0.769434 | 0.305908 | NO | NO | NA |
| Gm49838 | ENSMUSG( | ENSMUSG( | -0.309 | 0.458257 | NO | NO | NA |
| D630030B | ENSMUSG( | ENSMUSG( | 0.691602 | 0.484611 | NO | NO | NA |
| Gm9761 | ENSMUSG( | ENSMUSG( | 0.43338 | 0.199612 | NO | NO | NA |
| Ccr7 | ENSMUSG( | ENSMUSG( | 0.296539 | 0.480404 | NO | NO | NA |

| | | | | | | | |
|---|---|---|---|---|---|---|---|
| Vmn1r218 | ENSMUSG( | ENSMUSG( | 0.733307 | 0.368388 | NO | NO | NA |
| ENSMUSG | ENSMUSG( | ENSMUSG( | 0.795806 | 0.500155 | NO | NO | NA |
| Gm44168 | ENSMUSG( | ENSMUSG( | 0.615717 | 0.496053 | NO | NO | NA |
| Rgs7 | ENSMUSG( | ENSMUSG( | 0.66284 | 0.536077 | NO | NO | NA |
| Gm45528 | ENSMUSG( | ENSMUSG( | 0.542358 | 0.561653 | NO | NO | NA |
| Gm7993 | ENSMUSG( | ENSMUSG( | 0.873949 | 0.129078 | NO | NO | NA |
| Gm37465 | ENSMUSG( | ENSMUSG( | 0.735826 | 0.48915 | NO | NO | NA |
| Gm37558 | ENSMUSG( | ENSMUSG( | 0.006806 | 0.989896 | NO | NO | NA |
| Myh3 | ENSMUSG( | ENSMUSG( | 2.324369 | 2.38E-09 | YES | YES | Upregulated |
| Gm42731 | ENSMUSG( | ENSMUSG( | -0.77268 | 0.029851 | YES | NO | NA |
| ENSMUSG( | ENSMUSG( | ENSMUSG( | 0.709583 | 0.49586 | NO | NO | NA |
| Gm44066 | ENSMUSG( | ENSMUSG( | 1.29427 | 0.000595 | YES | YES | Upregulated |
| ENSMUSG | ENSMUSG( | ENSMUSG( | 0.373278 | 0.206019 | NO | NO | NA |
| Zdhhc22 | ENSMUSG( | ENSMUSG( | 0.147698 | 0.873814 | NO | NO | NA |
| Trav17 | ENSMUSG( | ENSMUSG( | -0.78343 | 0.501503 | NO | NO | NA |
| Gm19236 | ENSMUSG( | ENSMUSG( | 0.582424 | 0.542573 | NO | NO | NA |
| Shank2 | ENSMUSG( | ENSMUSG( | 0.627707 | 0.516718 | NO | NO | NA |
| Gm44067 | ENSMUSG( | ENSMUSG( | 0.351843 | 0.540236 | NO | NO | NA |
| Gm44812 | ENSMUSG( | ENSMUSG( | 0.96873 | 0.429222 | NO | NO | NA |
| Gm44187 | ENSMUSG( | ENSMUSG( | -0.31203 | 0.475711 | NO | NO | NA |
| Zfp711 | ENSMUSG( | ENSMUSG( | 0.978489 | 0.338282 | NO | NO | NA |
| Ap2m1-ps | ENSMUSG( | ENSMUSG( | -0.05601 | 0.824786 | NO | NO | NA |
| Gm38254 | ENSMUSG( | ENSMUSG( | 1.330762 | 0.202994 | NO | NO | NA |
| Gm44829 | ENSMUSG( | ENSMUSG( | 0.150782 | 0.610591 | NO | NO | NA |
| Gm47690 | ENSMUSG( | ENSMUSG( | 0.300172 | 0.613661 | NO | NO | NA |
| Gm12043 | ENSMUSG( | ENSMUSG( | 0.877735 | 0.354411 | NO | NO | NA |
| Vmn2r74 | ENSMUSG( | ENSMUSG( | 1.10014 | 0.307772 | NO | NO | NA |
| Gm37716 | ENSMUSG( | ENSMUSG( | 0.340631 | 0.344904 | NO | NO | NA |
| Fgd2 | ENSMUSG( | ENSMUSG( | 0.624655 | 0.129601 | NO | NO | NA |
| Nsg1 | ENSMUSG( | ENSMUSG( | 0.872851 | 0.10947 | NO | NO | NA |
| Gm38336 | ENSMUSG( | ENSMUSG( | -0.24312 | 0.435434 | NO | NO | NA |
| Gm37303 | ENSMUSG( | ENSMUSG( | -0.21975 | 0.432198 | NO | NO | NA |
| Unc13b | ENSMUSG( | ENSMUSG( | 0.438138 | 0.136624 | NO | NO | NA |
| D130058E | ENSMUSG( | ENSMUSG( | 0.643435 | 0.520926 | NO | NO | NA |
| Gm42462 | ENSMUSG( | ENSMUSG( | 0.425409 | 0.205557 | NO | NO | NA |
| Gm43196 | ENSMUSG( | ENSMUSG( | -1.00282 | 0.001924 | YES | YES | Downregulated |
| Ptx3 | ENSMUSG( | ENSMUSG( | 0.521878 | 0.631052 | NO | NO | NA |
| Gm48314 | ENSMUSG( | ENSMUSG( | 0.011725 | 0.979835 | NO | NO | NA |
| Acsl3 | ENSMUSG( | ENSMUSG( | 2.110998 | 8.04E-06 | YES | YES | Upregulated |
| Gm37443 | ENSMUSG( | ENSMUSG( | 1.208285 | 0.270662 | NO | NO | NA |
| Gm9895 | ENSMUSG( | ENSMUSG( | 0.315615 | 0.593982 | NO | NO | NA |
| Abca1 | ENSMUSG( | ENSMUSG( | 0.728055 | 0.347441 | NO | NO | NA |
| Nap1l3 | ENSMUSG( | ENSMUSG( | 0.060667 | 0.89543 | NO | NO | NA |
| Vmn2r65 | ENSMUSG( | ENSMUSG( | 0.449 | 0.653576 | NO | NO | NA |
| Fbxl7 | ENSMUSG( | ENSMUSG( | 0.544574 | 0.586396 | NO | NO | NA |
| ENSMUSG( | ENSMUSG( | ENSMUSG( | 0.894118 | 0.380749 | NO | NO | NA |
| Gm26815 | ENSMUSG( | ENSMUSG( | 0.843657 | 0.475251 | NO | NO | NA |

| | | | | | | | |
|---|---|---|---|---|---|---|---|
| Fgf6 | ENSMUSG( | ENSMUSG( | 1.046247 | 0.395256 | NO | NO | NA |
| Gm48532 | ENSMUSG( | ENSMUSG( | 0.765346 | 0.476775 | NO | NO | NA |
| 4933435G | ENSMUSG( | ENSMUSG( | 0.739667 | 0.466448 | NO | NO | NA |
| Gm20422 | ENSMUSG( | ENSMUSG( | 2.37778 | 0.028687 | YES | YES | Upregulated |
| Gm36186 | ENSMUSG( | ENSMUSG( | 0.74911 | 0.473283 | NO | NO | NA |
| Gm30400 | ENSMUSG( | ENSMUSG( | 0.882887 | 0.444352 | NO | NO | NA |
| Olfr1259 | ENSMUSG( | ENSMUSG( | 1.027783 | 0.280151 | NO | NO | NA |
| Maneal | ENSMUSG( | ENSMUSG( | 0.025886 | 0.952842 | NO | NO | NA |
| Gm35215 | ENSMUSG( | ENSMUSG( | 0.840936 | 0.441381 | NO | NO | NA |
| Slc38a11 | ENSMUSG( | ENSMUSG( | 0.185868 | 0.847974 | NO | NO | NA |
| Gm47997 | ENSMUSG( | ENSMUSG( | 0.762775 | 0.493791 | NO | NO | NA |
| Gm37829 | ENSMUSG( | ENSMUSG( | -0.05592 | 0.890997 | NO | NO | NA |
| Rwdd2a | ENSMUSG( | ENSMUSG( | 0.234989 | 0.423173 | NO | NO | NA |
| Col25a1 | ENSMUSG( | ENSMUSG( | 0.684362 | 0.474914 | NO | NO | NA |
| Fam243 | ENSMUSG( | ENSMUSG( | 0.679289 | 0.517559 | NO | NO | NA |
| Mfsd7a | ENSMUSG( | ENSMUSG( | 0.43458 | 0.688983 | NO | NO | NA |
| 2310002F( | ENSMUSG( | ENSMUSG( | 0.034432 | 0.936437 | NO | NO | NA |
| Gm16029 | ENSMUSG( | ENSMUSG( | 0.623123 | 0.557764 | NO | NO | NA |
| C1qb | ENSMUSG( | ENSMUSG( | -1.22725 | 0.000848 | YES | YES | Downregulated |
| Eml6 | ENSMUSG( | ENSMUSG( | 1.318988 | 0.020008 | YES | YES | Upregulated |
| Hpn | ENSMUSG( | ENSMUSG( | 1.082551 | 0.130971 | NO | NO | NA |
| Kctd14 | ENSMUSG( | ENSMUSG( | 0.905117 | 0.256954 | NO | NO | NA |
| Gm37248 | ENSMUSG( | ENSMUSG( | 0.464694 | 0.398888 | NO | NO | NA |
| Gm37795 | ENSMUSG( | ENSMUSG( | 0.58354 | 0.419854 | NO | NO | NA |
| Naa11 | ENSMUSG( | ENSMUSG( | 0.627778 | 0.546403 | NO | NO | NA |
| Gm28551 | ENSMUSG( | ENSMUSG( | 0.447375 | 0.607829 | NO | NO | NA |
| Vmn1r14 | ENSMUSG( | ENSMUSG( | 0.582423 | 0.580095 | NO | NO | NA |
| Gm26801 | ENSMUSG( | ENSMUSG( | 0.899785 | 0.426541 | NO | NO | NA |
| Gm45266 | ENSMUSG( | ENSMUSG( | 0.807679 | 0.192466 | NO | NO | NA |
| Gm30551 | ENSMUSG( | ENSMUSG( | 0.679735 | 0.519155 | NO | NO | NA |
| Il17rd | ENSMUSG( | ENSMUSG( | 0.271212 | 0.514562 | NO | NO | NA |
| Gm20234 | ENSMUSG( | ENSMUSG( | 0.041545 | 0.891163 | NO | NO | NA |
| Nkapl | ENSMUSG( | ENSMUSG( | 0.869229 | 0.224654 | NO | NO | NA |
| Gm19221 | ENSMUSG( | ENSMUSG( | -0.09332 | 0.756362 | NO | NO | NA |
| Adgrb3 | ENSMUSG( | ENSMUSG( | 0.203424 | 0.815336 | NO | NO | NA |
| Wipi1 | ENSMUSG( | ENSMUSG( | 0.58025 | 0.318119 | NO | NO | NA |
| Cd55os | ENSMUSG( | ENSMUSG( | -0.96593 | 0.000384 | YES | NO | NA |
| ENSMUSG( | ENSMUSG( | ENSMUSG( | 0.143717 | 0.756401 | NO | NO | NA |
| Zfp612 | ENSMUSG( | ENSMUSG( | 0.414484 | 0.522983 | NO | NO | NA |
| Gm16731 | ENSMUSG( | ENSMUSG( | 0.295725 | 0.374748 | NO | NO | NA |
| Trav4d-3 | ENSMUSG( | ENSMUSG( | -1.05069 | 0.787692 | NO | NO | NA |
| Gm3892 | ENSMUSG( | ENSMUSG( | 6.316229 | 0.062824 | NO | NO | NA |
| Gm527 | ENSMUSG( | ENSMUSG( | -0.8003 | 0.020536 | YES | NO | NA |
| Gm37742 | ENSMUSG( | ENSMUSG( | 0.960819 | 0.386988 | NO | NO | NA |
| 9530085L: | ENSMUSG( | ENSMUSG( | -0.27965 | 0.412416 | NO | NO | NA |
| Gm48583 | ENSMUSG( | ENSMUSG( | 0.837395 | 0.022031 | YES | NO | NA |
| Btbd16 | ENSMUSG( | ENSMUSG( | 0.116067 | 0.891166 | NO | NO | NA |

| | | | | | | | |
|---|---|---|---|---|---|---|---|
| Lhfpl3 | ENSMUSG( | ENSMUSG( | 0.751882 | 0.3959 | NO | NO | NA |
| Gm37309 | ENSMUSG( | ENSMUSG( | 0.168801 | 0.527649 | NO | NO | NA |
| Vmn2r57 | ENSMUSG( | ENSMUSG( | 1.02185 | 0.129948 | NO | NO | NA |
| Gm32552 | ENSMUSG( | ENSMUSG( | 0.772205 | 0.402496 | NO | NO | NA |
| 4732419C: | ENSMUSG( | ENSMUSG( | 0.786925 | 0.400828 | NO | NO | NA |
| E330035G | ENSMUSG( | ENSMUSG( | 0.482772 | 0.640484 | NO | NO | NA |
| Gm12435 | ENSMUSG( | ENSMUSG( | 0.157889 | 0.726265 | NO | NO | NA |
| E330034L1 | ENSMUSG( | ENSMUSG( | 0.125709 | 0.681501 | NO | NO | NA |
| Hsd3b7 | ENSMUSG( | ENSMUSG( | -0.73265 | 0.263075 | NO | NO | NA |
| Gm35037 | ENSMUSG( | ENSMUSG( | -0.02376 | 0.930642 | NO | NO | NA |
| Glis3 | ENSMUSG( | ENSMUSG( | 0.666319 | 0.30206 | NO | NO | NA |
| Gm38377 | ENSMUSG( | ENSMUSG( | 0.663111 | 0.32874 | NO | NO | NA |
| Ighv1-77 | ENSMUSG( | ENSMUSG( | 21.62298 | NA | NO | NO | NA |
| Plagl1 | ENSMUSG( | ENSMUSG( | 1.180206 | 0.037864 | YES | YES | Upregulated |
| Stard13 | ENSMUSG( | ENSMUSG( | 1.220645 | 0.001207 | YES | YES | Upregulated |
| Ldlrad2 | ENSMUSG( | ENSMUSG( | 0.942914 | 0.406542 | NO | NO | NA |
| Gm44172 | ENSMUSG( | ENSMUSG( | 1.253571 | 0.214499 | NO | NO | NA |
| Igkv5-48 | ENSMUSG( | ENSMUSG( | -3.96445 | 0.034681 | YES | YES | Downregulated |
| Trp53cor1 | ENSMUSG( | ENSMUSG( | 1.083592 | 0.037868 | YES | YES | Upregulated |
| Gm27010 | ENSMUSG( | ENSMUSG( | -0.28931 | 0.57 | NO | NO | NA |
| B230398E( | ENSMUSG( | ENSMUSG( | -0.01874 | 0.962708 | NO | NO | NA |
| Nxpe5 | ENSMUSG( | ENSMUSG( | 0.555223 | 0.55607 | NO | NO | NA |
| Gm26670 | ENSMUSG( | ENSMUSG( | 0.673958 | 0.402464 | NO | NO | NA |
| ENSMUSG( | ENSMUSG( | ENSMUSG( | 0.724085 | 0.49586 | NO | NO | NA |
| Smad6 | ENSMUSG( | ENSMUSG( | 0.806642 | 0.440958 | NO | NO | NA |
| Gm42651 | ENSMUSG( | ENSMUSG( | 0.527445 | 0.477439 | NO | NO | NA |
| Gm20507 | ENSMUSG( | ENSMUSG( | -0.61238 | 0.639778 | NO | NO | NA |
| Rps10-ps1 | ENSMUSG( | ENSMUSG( | 0.256763 | 0.613152 | NO | NO | NA |
| Gm45732 | ENSMUSG( | ENSMUSG( | 0.531754 | 0.201571 | NO | NO | NA |
| 1500002C: | ENSMUSG( | ENSMUSG( | -0.26733 | 0.258455 | NO | NO | NA |
| Ighv9-4 | ENSMUSG( | ENSMUSG( | -4.55912 | 0.058516 | NO | NO | NA |
| Gm13074 | ENSMUSG( | ENSMUSG( | 0.62508 | 0.560577 | NO | NO | NA |
| Vipr1 | ENSMUSG( | ENSMUSG( | -0.26791 | 0.520134 | NO | NO | NA |
| Tbx15 | ENSMUSG( | ENSMUSG( | 1.052181 | 0.182591 | NO | NO | NA |
| Gm44712 | ENSMUSG( | ENSMUSG( | 0.572717 | 0.537456 | NO | NO | NA |
| ENSMUSG | ENSMUSG( | ENSMUSG( | -0.16334 | 0.712875 | NO | NO | NA |
| Ehf | ENSMUSG( | ENSMUSG( | 0.393906 | 0.701752 | NO | NO | NA |
| Gm9892 | ENSMUSG( | ENSMUSG( | 0.661066 | 0.031185 | YES | NO | NA |
| 9330160F: | ENSMUSG( | ENSMUSG( | 0.205373 | 0.571212 | NO | NO | NA |
| Hapln1 | ENSMUSG( | ENSMUSG( | 0.682706 | 0.350224 | NO | NO | NA |
| Gm43413 | ENSMUSG( | ENSMUSG( | -0.50471 | 0.202772 | NO | NO | NA |
| Gm43403 | ENSMUSG( | ENSMUSG( | 0.847691 | 0.030902 | YES | NO | NA |
| Grb10 | ENSMUSG( | ENSMUSG( | 0.591204 | 0.519316 | NO | NO | NA |
| Fut9 | ENSMUSG( | ENSMUSG( | 0.515078 | 0.66406 | NO | NO | NA |
| Hbegf | ENSMUSG( | ENSMUSG( | 1.145202 | 0.02773 | YES | YES | Upregulated |
| ENSMUSG( | ENSMUSG( | ENSMUSG( | 0.980552 | 0.128843 | NO | NO | NA |
| Chst8 | ENSMUSG( | ENSMUSG( | 0.558673 | 0.648137 | NO | NO | NA |

| | | | | | | | |
|---|---|---|---|---|---|---|---|
| Rhoj | ENSMUSG( | ENSMUSG( | 0.578215 | 0.555025 | NO | NO | NA |
| Gm38042 | ENSMUSG( | ENSMUSG( | 0.630593 | 0.523069 | NO | NO | NA |
| Got2-ps1 | ENSMUSG( | ENSMUSG( | -0.48834 | 0.22829 | NO | NO | NA |
| Hmx2 | ENSMUSG( | ENSMUSG( | 0.900275 | 0.04363 | YES | NO | NA |
| 4930404N | ENSMUSG( | ENSMUSG( | 0.227187 | 0.584439 | NO | NO | NA |
| Gpr137c | ENSMUSG( | ENSMUSG( | -0.49647 | 0.071331 | NO | NO | NA |
| Ubtfl1 | ENSMUSG( | ENSMUSG( | 0.869516 | 0.103811 | NO | NO | NA |
| Gm37909 | ENSMUSG( | ENSMUSG( | 0.791577 | 0.34477 | NO | NO | NA |
| Gm20735 | ENSMUSG( | ENSMUSG( | -0.15351 | 0.623689 | NO | NO | NA |
| Gm47950 | ENSMUSG( | ENSMUSG( | -0.3491 | 0.184777 | NO | NO | NA |
| Gm20518 | ENSMUSG( | ENSMUSG( | -1.2261 | 0.043988 | YES | YES | Downregulated |
| 6720464F. | ENSMUSG( | ENSMUSG( | 0.11213 | 0.781352 | NO | NO | NA |
| Gm43428 | ENSMUSG( | ENSMUSG( | 0.581158 | 0.572509 | NO | NO | NA |
| Gm10030 | ENSMUSG( | ENSMUSG( | 0.217672 | 0.455255 | NO | NO | NA |
| D030068K | ENSMUSG( | ENSMUSG( | 0.527464 | 0.600045 | NO | NO | NA |
| Gm45244 | ENSMUSG( | ENSMUSG( | 0.959016 | 0.013159 | YES | NO | NA |
| Per2 | ENSMUSG( | ENSMUSG( | 0.540969 | 0.018028 | YES | NO | NA |
| Zfp551 | ENSMUSG( | ENSMUSG( | -0.47537 | 0.198254 | NO | NO | NA |
| 9030622O | ENSMUSG( | ENSMUSG( | 0.623332 | 0.553726 | NO | NO | NA |
| B230217C | ENSMUSG( | ENSMUSG( | 0.553429 | 0.245855 | NO | NO | NA |
| Gm44238 | ENSMUSG( | ENSMUSG( | -0.43727 | 0.471513 | NO | NO | NA |
| Igkv2-109 | ENSMUSG( | ENSMUSG( | -6.4063 | 0.000224 | YES | YES | Downregulated |
| Igkv17-121 | ENSMUSG( | ENSMUSG( | -2.17093 | 0.135252 | NO | NO | NA |
| ENSMUSG | ENSMUSG( | ENSMUSG( | 0.556426 | 0.164842 | NO | NO | NA |
| Gm37900 | ENSMUSG( | ENSMUSG( | 0.286512 | 0.609105 | NO | NO | NA |
| Gm43532 | ENSMUSG( | ENSMUSG( | 1.022848 | 0.338874 | NO | NO | NA |
| Lypd1 | ENSMUSG( | ENSMUSG( | -0.06715 | 0.942238 | NO | NO | NA |
| Galnt9 | ENSMUSG( | ENSMUSG( | 0.625105 | 0.289996 | NO | NO | NA |
| Gm13387 | ENSMUSG( | ENSMUSG( | -0.93699 | 0.000234 | YES | NO | NA |
| Gm20796 | ENSMUSG( | ENSMUSG( | 0.756394 | 0.440958 | NO | NO | NA |
| Gm50340 | ENSMUSG( | ENSMUSG( | -0.24963 | 0.437342 | NO | NO | NA |
| Cdh11 | ENSMUSG( | ENSMUSG( | 0.28361 | 0.569072 | NO | NO | NA |
| Gm5357 | ENSMUSG( | ENSMUSG( | 1.158099 | 0.222722 | NO | NO | NA |
| Foxn1 | ENSMUSG( | ENSMUSG( | 0.896032 | 0.326371 | NO | NO | NA |
| Gm37514 | ENSMUSG( | ENSMUSG( | 0.786848 | 0.454484 | NO | NO | NA |
| ENSMUSG | ENSMUSG( | ENSMUSG( | 1.178435 | 0.16122 | NO | NO | NA |
| ENSMUSG | ENSMUSG( | ENSMUSG( | -0.17115 | 0.968665 | NO | NO | NA |
| Gm42544 | ENSMUSG( | ENSMUSG( | 0.727067 | 0.556162 | NO | NO | NA |
| Nsun7 | ENSMUSG( | ENSMUSG( | 0.834068 | 0.036805 | YES | NO | NA |
| Gm44709 | ENSMUSG( | ENSMUSG( | 0.391185 | 0.34766 | NO | NO | NA |
| Gm15462 | ENSMUSG( | ENSMUSG( | 1.089007 | 0.29513 | NO | NO | NA |
| Gm49767 | ENSMUSG( | ENSMUSG( | 0.270009 | 0.808338 | NO | NO | NA |
| Gm38302 | ENSMUSG( | ENSMUSG( | 0.142733 | 0.8114 | NO | NO | NA |
| Gm10863 | ENSMUSG( | ENSMUSG( | -0.37826 | 0.162161 | NO | NO | NA |
| Mblac1 | ENSMUSG( | ENSMUSG( | 0.987014 | 0.016803 | YES | NO | NA |
| 3830403N | ENSMUSG( | ENSMUSG( | -0.12594 | 0.902864 | NO | NO | NA |
| Gm45343 | ENSMUSG( | ENSMUSG( | 0.158404 | 0.728114 | NO | NO | NA |

| | | | | | | | |
|---|---|---|---|---|---|---|---|
| Adamts2 | ENSMUSG( | ENSMUSG( | 1.16579 | 0.255924 | NO | NO | NA |
| BB365896 | ENSMUSG( | ENSMUSG( | 0.790429 | 0.444371 | NO | NO | NA |
| Gm10643 | ENSMUSG( | ENSMUSG( | 0.372024 | 0.436017 | NO | NO | NA |
| Ighv1-52 | ENSMUSG( | ENSMUSG( | -2.1952 | 0.309644 | NO | NO | NA |
| BC018473 | ENSMUSG( | ENSMUSG( | 0.569797 | 0.515887 | NO | NO | NA |
| Wnt7b | ENSMUSG( | ENSMUSG( | 0.578198 | 0.485027 | NO | NO | NA |
| Gm45605 | ENSMUSG( | ENSMUSG( | 0.730484 | 0.255598 | NO | NO | NA |
| C79130 | ENSMUSG( | ENSMUSG( | 0.691203 | 0.417592 | NO | NO | NA |
| Olfr1307 | ENSMUSG( | ENSMUSG( | 0.262097 | 0.751928 | NO | NO | NA |
| Mc1r | ENSMUSG( | ENSMUSG( | 0.283584 | 0.76755 | NO | NO | NA |
| Gm42955 | ENSMUSG( | ENSMUSG( | 0.609555 | 0.596194 | NO | NO | NA |
| Gm45745 | ENSMUSG( | ENSMUSG( | -0.42894 | 0.149234 | NO | NO | NA |
| Gm17322 | ENSMUSG( | ENSMUSG( | 0.455405 | 0.248985 | NO | NO | NA |
| Me3 | ENSMUSG( | ENSMUSG( | 0.30781 | 0.756944 | NO | NO | NA |
| Trbv30 | ENSMUSG( | ENSMUSG( | 0.641461 | 0.326371 | NO | NO | NA |
| D930030I( | ENSMUSG( | ENSMUSG( | 0.245131 | 0.573325 | NO | NO | NA |
| Olfml3 | ENSMUSG( | ENSMUSG( | 0.217237 | 0.808338 | NO | NO | NA |
| Tex13d | ENSMUSG( | ENSMUSG( | 0.147117 | 0.684467 | NO | NO | NA |
| Gm47493 | ENSMUSG( | ENSMUSG( | 0.987828 | 0.328452 | NO | NO | NA |
| Slc13a4 | ENSMUSG( | ENSMUSG( | 0.972968 | 0.345697 | NO | NO | NA |
| Gm44193 | ENSMUSG( | ENSMUSG( | 0.738223 | 0.472399 | NO | NO | NA |
| Gm12295 | ENSMUSG( | ENSMUSG( | 0.688175 | 0.519025 | NO | NO | NA |
| Gm48062 | ENSMUSG( | ENSMUSG( | -0.23017 | 0.546443 | NO | NO | NA |
| Sult5a1 | ENSMUSG( | ENSMUSG( | 0.716217 | 0.56174 | NO | NO | NA |
| Trim9 | ENSMUSG( | ENSMUSG( | 0.425982 | 0.692577 | NO | NO | NA |
| Kcnab1 | ENSMUSG( | ENSMUSG( | 0.365546 | 0.512289 | NO | NO | NA |
| Lzts2 | ENSMUSG( | ENSMUSG( | -0.50951 | 0.109571 | NO | NO | NA |
| Gm44101 | ENSMUSG( | ENSMUSG( | 0.783384 | 0.360943 | NO | NO | NA |
| Gm14461 | ENSMUSG( | ENSMUSG( | 0.670323 | 0.529241 | NO | NO | NA |
| Gm5637 | ENSMUSG( | ENSMUSG( | 0.105547 | 0.795358 | NO | NO | NA |
| Itgb5 | ENSMUSG( | ENSMUSG( | 0.123066 | 0.732349 | NO | NO | NA |
| Gm49484 | ENSMUSG( | ENSMUSG( | 0.91201 | 0.201715 | NO | NO | NA |
| A830082N | ENSMUSG( | ENSMUSG( | 0.351894 | 0.526045 | NO | NO | NA |
| 1700001G | ENSMUSG( | ENSMUSG( | 0.676476 | 0.03045 | YES | NO | NA |
| Zfp534 | ENSMUSG( | ENSMUSG( | 4.616651 | 1.27E-16 | YES | YES | Upregulated |
| 4930426L( | ENSMUSG( | ENSMUSG( | 0.380354 | 0.321249 | NO | NO | NA |
| Trpv3 | ENSMUSG( | ENSMUSG( | 0.948799 | 0.307125 | NO | NO | NA |
| Gm49499 | ENSMUSG( | ENSMUSG( | 0.923918 | 0.361411 | NO | NO | NA |
| Lekr1 | ENSMUSG( | ENSMUSG( | 0.75798 | 0.041393 | YES | NO | NA |
| Olfr597 | ENSMUSG( | ENSMUSG( | 0.713145 | 0.563439 | NO | NO | NA |
| Mgarp | ENSMUSG( | ENSMUSG( | 1.152576 | 0.058181 | NO | NO | NA |
| Gata2 | ENSMUSG( | ENSMUSG( | 0.627846 | 0.38454 | NO | NO | NA |
| Gm10243 | ENSMUSG( | ENSMUSG( | 0.911637 | 0.193429 | NO | NO | NA |
| 6330562C: | ENSMUSG( | ENSMUSG( | 0.672195 | 0.111881 | NO | NO | NA |
| Nlgn1 | ENSMUSG( | ENSMUSG( | 0.564023 | 0.597056 | NO | NO | NA |
| Cd200r3 | ENSMUSG( | ENSMUSG( | 0.793974 | 0.427109 | NO | NO | NA |
| H2ac4 | ENSMUSG( | ENSMUSG( | 0.344557 | 0.38917 | NO | NO | NA |

| | | | | | | | |
|---|---|---|---|---|---|---|---|
| 4930589L2 | ENSMUSG( | ENSMUSG( | 0.75944 | 0.489587 | NO | NO | NA |
| Gpr27 | ENSMUSG( | ENSMUSG( | 0.953049 | 0.301897 | NO | NO | NA |
| 4921507G | ENSMUSG( | ENSMUSG( | 0.709488 | 0.40295 | NO | NO | NA |
| Gm50462 | ENSMUSG( | ENSMUSG( | 0.382128 | 0.754539 | NO | NO | NA |
| Gm47601 | ENSMUSG( | ENSMUSG( | 1.051902 | 0.297175 | NO | NO | NA |
| 1700056N | ENSMUSG( | ENSMUSG( | -0.87169 | 0.002385 | YES | NO | NA |
| Gsta3 | ENSMUSG( | ENSMUSG( | 0.625602 | 0.473501 | NO | NO | NA |
| Spred3 | ENSMUSG( | ENSMUSG( | 0.46102 | 0.295277 | NO | NO | NA |
| Gm36932 | ENSMUSG( | ENSMUSG( | -0.74125 | 0.032352 | YES | NO | NA |
| Zeb2os | ENSMUSG( | ENSMUSG( | -0.85929 | 0.017346 | YES | NO | NA |
| C1rl | ENSMUSG( | ENSMUSG( | 0.305908 | 0.764536 | NO | NO | NA |
| Cacng4 | ENSMUSG( | ENSMUSG( | 1.069759 | 0.16498 | NO | NO | NA |
| Gm48835 | ENSMUSG( | ENSMUSG( | 1.446753 | 0.150807 | NO | NO | NA |
| Stard6 | ENSMUSG( | ENSMUSG( | 0.647099 | 0.0073 | YES | NO | NA |
| Gm6653 | ENSMUSG( | ENSMUSG( | 0.281974 | 0.411917 | NO | NO | NA |
| Gm42503 | ENSMUSG( | ENSMUSG( | 1.090742 | 0.149839 | NO | NO | NA |
| Gm44241 | ENSMUSG( | ENSMUSG( | 1.162054 | 0.280224 | NO | NO | NA |
| Olfr225 | ENSMUSG( | ENSMUSG( | 0.994402 | 0.31393 | NO | NO | NA |
| Gm47882 | ENSMUSG( | ENSMUSG( | 0.162964 | 0.537122 | NO | NO | NA |
| Rgs7bp | ENSMUSG( | ENSMUSG( | 1.128372 | 0.346331 | NO | NO | NA |
| Nacc2 | ENSMUSG( | ENSMUSG( | 0.421929 | 0.338636 | NO | NO | NA |
| Gm45159 | ENSMUSG( | ENSMUSG( | 0.664307 | 0.500435 | NO | NO | NA |
| ENSMUSG( | ENSMUSG( | ENSMUSG( | -0.03186 | 0.908629 | NO | NO | NA |
| Trdc | ENSMUSG( | ENSMUSG( | 0.222041 | 0.714767 | NO | NO | NA |
| Rab36 | ENSMUSG( | ENSMUSG( | 1.207696 | 0.023362 | YES | YES | Upregulated |
| Ribc1 | ENSMUSG( | ENSMUSG( | 0.68493 | 0.061444 | NO | NO | NA |
| Gm15894 | ENSMUSG( | ENSMUSG( | 0.809978 | 0.429315 | NO | NO | NA |
| Hoxc13 | ENSMUSG( | ENSMUSG( | 0.350359 | 0.729086 | NO | NO | NA |
| Ttc30b | ENSMUSG( | ENSMUSG( | 0.329147 | 0.486835 | NO | NO | NA |
| Gm32200 | ENSMUSG( | ENSMUSG( | 0.498987 | 0.44835 | NO | NO | NA |
| Gm44822 | ENSMUSG( | ENSMUSG( | 0.05293 | 0.91234 | NO | NO | NA |
| Dynlt2a2 | ENSMUSG( | ENSMUSG( | 0.822929 | 0.006725 | YES | NO | NA |
| Rpl28-ps2 | ENSMUSG( | ENSMUSG( | 0.716228 | 0.505024 | NO | NO | NA |
| Slc35f5 | ENSMUSG( | ENSMUSG( | 1.161629 | 0.012605 | YES | YES | Upregulated |
| Rpp38 | ENSMUSG( | ENSMUSG( | 0.232441 | 0.403558 | NO | NO | NA |
| Nkd1 | ENSMUSG( | ENSMUSG( | 0.927758 | 0.072835 | NO | NO | NA |
| Gm13204 | ENSMUSG( | ENSMUSG( | 0.450222 | 0.113445 | NO | NO | NA |
| Nacad | ENSMUSG( | ENSMUSG( | -1.10611 | 0.001106 | YES | YES | Downregulated |
| Cxcl12 | ENSMUSG( | ENSMUSG( | 0.252565 | 0.819148 | NO | NO | NA |
| Gm42989 | ENSMUSG( | ENSMUSG( | 0.393971 | 0.700651 | NO | NO | NA |
| Gm37529 | ENSMUSG( | ENSMUSG( | 0.296947 | 0.567857 | NO | NO | NA |
| Vmn1r216 | ENSMUSG( | ENSMUSG( | 0.390908 | 0.696173 | NO | NO | NA |
| Bex2 | ENSMUSG( | ENSMUSG( | 0.564218 | 0.272959 | NO | NO | NA |
| 6030400A | ENSMUSG( | ENSMUSG( | -1.35175 | 0.002826 | YES | YES | Downregulated |
| Gm37926 | ENSMUSG( | ENSMUSG( | 0.657167 | 0.506617 | NO | NO | NA |
| Lrrn4 | ENSMUSG( | ENSMUSG( | -1.29067 | 0.002693 | YES | YES | Downregulated |
| ENSMUSG | ENSMUSG( | ENSMUSG( | -0.85719 | 0.029002 | YES | NO | NA |

| | | | | | | | |
|---|---|---|---|---|---|---|---|
| Tmem151l | ENSMUSG( | ENSMUSG( | 0.101824 | 0.889961 | NO | NO | NA |
| C430002N | ENSMUSG( | ENSMUSG( | -1.00604 | 0.000271 | YES | YES | Downregulated |
| ENSMUSG | ENSMUSG( | ENSMUSG( | 0.114641 | 0.852063 | NO | NO | NA |
| Crppa | ENSMUSG( | ENSMUSG( | 0.510851 | 0.129434 | NO | NO | NA |
| ENSMUSG | ENSMUSG( | ENSMUSG( | -0.28339 | 0.424425 | NO | NO | NA |
| Gm37706 | ENSMUSG( | ENSMUSG( | 0.701073 | 0.478576 | NO | NO | NA |
| Rsph9 | ENSMUSG( | ENSMUSG( | 0.956515 | 0.164596 | NO | NO | NA |
| Gm5112 | ENSMUSG( | ENSMUSG( | 0.51532 | 0.05218 | NO | NO | NA |
| ENSMUSG( | ENSMUSG( | ENSMUSG( | 1.054809 | 0.310909 | NO | NO | NA |
| 4833417C: | ENSMUSG( | ENSMUSG( | 0.006396 | 0.983196 | NO | NO | NA |
| Magef1 | ENSMUSG( | ENSMUSG( | -0.4932 | 0.827719 | NO | NO | NA |
| Mylk3 | ENSMUSG( | ENSMUSG( | 0.895748 | 0.267362 | NO | NO | NA |
| Megf11 | ENSMUSG( | ENSMUSG( | 0.141383 | 0.783951 | NO | NO | NA |
| Rasl11a | ENSMUSG( | ENSMUSG( | -0.87654 | 0.000204 | YES | NO | NA |
| Gm17249 | ENSMUSG( | ENSMUSG( | 0.690185 | 0.389574 | NO | NO | NA |
| Marchf3 | ENSMUSG( | ENSMUSG( | 0.154135 | 0.64918 | NO | NO | NA |
| Olfr740 | ENSMUSG( | ENSMUSG( | 0.423839 | 0.686461 | NO | NO | NA |
| Gm43961 | ENSMUSG( | ENSMUSG( | -0.51804 | 0.080192 | NO | NO | NA |
| Olfr527 | ENSMUSG( | ENSMUSG( | 1.800681 | 0.00703 | YES | YES | Upregulated |
| Gm14760 | ENSMUSG( | ENSMUSG( | 0.614005 | 0.116413 | NO | NO | NA |
| Gm28320 | ENSMUSG( | ENSMUSG( | 0.878098 | 0.425416 | NO | NO | NA |
| Gm4981 | ENSMUSG( | ENSMUSG( | 1.785583 | 0.002291 | YES | YES | Upregulated |
| Vmn1r18 | ENSMUSG( | ENSMUSG( | 0.681443 | 0.503887 | NO | NO | NA |
| Prss43 | ENSMUSG( | ENSMUSG( | 0.976702 | 0.413936 | NO | NO | NA |
| C030015A | ENSMUSG( | ENSMUSG( | -0.18755 | 0.663452 | NO | NO | NA |
| Gm805 | ENSMUSG( | ENSMUSG( | 0.755106 | 0.457413 | NO | NO | NA |
| Syngap1 | ENSMUSG( | ENSMUSG( | 0.20787 | 0.579976 | NO | NO | NA |
| Pde7b | ENSMUSG( | ENSMUSG( | 0.565494 | 0.542061 | NO | NO | NA |
| Gm38021 | ENSMUSG( | ENSMUSG( | 0.671089 | 0.306823 | NO | NO | NA |
| Gm26620 | ENSMUSG( | ENSMUSG( | 0.331056 | 0.464893 | NO | NO | NA |
| Olfr839-ps | ENSMUSG( | ENSMUSG( | 0.964327 | 0.260691 | NO | NO | NA |
| ENSMUSG | ENSMUSG( | ENSMUSG( | 0.48657 | 0.351949 | NO | NO | NA |
| Mpp3 | ENSMUSG( | ENSMUSG( | 0.731828 | 0.458761 | NO | NO | NA |
| Fbln1 | ENSMUSG( | ENSMUSG( | 0.349106 | 0.603586 | NO | NO | NA |
| Zfp781 | ENSMUSG( | ENSMUSG( | -0.24053 | 0.661518 | NO | NO | NA |
| ENSMUSG | ENSMUSG( | ENSMUSG( | 0.117107 | 0.78842 | NO | NO | NA |
| 4930542C: | ENSMUSG( | ENSMUSG( | 0.124627 | 0.878742 | NO | NO | NA |
| Gm5053 | ENSMUSG( | ENSMUSG( | 0.70289 | 0.496618 | NO | NO | NA |
| 4732414G | ENSMUSG( | ENSMUSG( | 1.077005 | 0.346985 | NO | NO | NA |
| Grk3 | ENSMUSG( | ENSMUSG( | 0.344949 | 0.617106 | NO | NO | NA |
| Parvb | ENSMUSG( | ENSMUSG( | 0.411568 | 0.395256 | NO | NO | NA |
| Olfr220 | ENSMUSG( | ENSMUSG( | 1.247248 | 0.19074 | NO | NO | NA |
| Chsy3 | ENSMUSG( | ENSMUSG( | 0.419458 | 0.643515 | NO | NO | NA |
| Gm10840 | ENSMUSG( | ENSMUSG( | 0.043493 | 0.911939 | NO | NO | NA |
| 4930524O | ENSMUSG( | ENSMUSG( | -0.00661 | 0.984739 | NO | NO | NA |
| Cphx1 | ENSMUSG( | ENSMUSG( | 0.834798 | 0.261636 | NO | NO | NA |
| 4930515G | ENSMUSG( | ENSMUSG( | -0.33082 | 0.424668 | NO | NO | NA |

| | | | | | | | |
|---|---|---|---|---|---|---|---|
| BC022960 | ENSMUSG( | ENSMUSG( | 1.603284 | 0.004646 | YES | YES | Upregulated |
| Gm12191 | ENSMUSG( | ENSMUSG( | 0.080959 | 0.946066 | NO | NO | NA |
| Gm47175 | ENSMUSG( | ENSMUSG( | 0.723317 | 0.044902 | YES | NO | NA |
| Rpl36-ps1: | ENSMUSG( | ENSMUSG( | 0.392919 | 0.14327 | NO | NO | NA |
| Gm42681 | ENSMUSG( | ENSMUSG( | 0.804191 | 0.450101 | NO | NO | NA |
| Gm17825 | ENSMUSG( | ENSMUSG( | -0.00745 | 0.985008 | NO | NO | NA |
| Mast1 | ENSMUSG( | ENSMUSG( | -0.10851 | 0.736727 | NO | NO | NA |
| Gm47438 | ENSMUSG( | ENSMUSG( | 0.547835 | 0.325398 | NO | NO | NA |
| A530017D | ENSMUSG( | ENSMUSG( | -0.13088 | 0.671111 | NO | NO | NA |
| Arntl2 | ENSMUSG( | ENSMUSG( | 0.134676 | 0.866061 | NO | NO | NA |
| H4c1 | ENSMUSG( | ENSMUSG( | 0.717759 | 0.085007 | NO | NO | NA |
| AU018091 | ENSMUSG( | ENSMUSG( | 0.738883 | 0.488115 | NO | NO | NA |
| Epor | ENSMUSG( | ENSMUSG( | 0.664569 | 0.204 | NO | NO | NA |
| Ccdc142os | ENSMUSG( | ENSMUSG( | -0.24003 | 0.427653 | NO | NO | NA |
| Gm48978 | ENSMUSG( | ENSMUSG( | 0.604648 | 0.422267 | NO | NO | NA |
| ENSMUSG | ENSMUSG( | ENSMUSG( | -0.12614 | 0.691059 | NO | NO | NA |
| Sec31b | ENSMUSG( | ENSMUSG( | -0.06121 | 0.878987 | NO | NO | NA |
| Gm36931 | ENSMUSG( | ENSMUSG( | 0.42924 | 0.497169 | NO | NO | NA |
| Gm19327 | ENSMUSG( | ENSMUSG( | 0.058342 | 0.946459 | NO | NO | NA |
| Phactr1 | ENSMUSG( | ENSMUSG( | 0.487254 | 0.612379 | NO | NO | NA |
| Gm37902 | ENSMUSG( | ENSMUSG( | 0.082379 | 0.783353 | NO | NO | NA |
| Gm36329 | ENSMUSG( | ENSMUSG( | -0.36805 | 0.417925 | NO | NO | NA |
| Gcnt3 | ENSMUSG( | ENSMUSG( | 0.659932 | 0.538693 | NO | NO | NA |
| 5730488B( | ENSMUSG( | ENSMUSG( | 0.986663 | 0.360054 | NO | NO | NA |
| E130008D | ENSMUSG( | ENSMUSG( | 0.752038 | 0.463257 | NO | NO | NA |
| Gm15622 | ENSMUSG( | ENSMUSG( | 0.930875 | 0.350951 | NO | NO | NA |
| Gm43908 | ENSMUSG( | ENSMUSG( | 0.639829 | 0.543023 | NO | NO | NA |
| Gm36287 | ENSMUSG( | ENSMUSG( | 0.644222 | 0.310173 | NO | NO | NA |
| Tmem51 | ENSMUSG( | ENSMUSG( | -1.7069 | 3.00E-06 | YES | YES | Downregulated |
| Olfr484 | ENSMUSG( | ENSMUSG( | 1.258776 | 0.224698 | NO | NO | NA |
| Gm12751 | ENSMUSG( | ENSMUSG( | 0.890441 | 0.024342 | YES | NO | NA |
| Klhl1 | ENSMUSG( | ENSMUSG( | 0.33862 | 0.758402 | NO | NO | NA |
| Zbtb3 | ENSMUSG( | ENSMUSG( | -0.0788 | 0.825563 | NO | NO | NA |
| Clec1a | ENSMUSG( | ENSMUSG( | 1.157011 | 0.310319 | NO | NO | NA |
| Pou4f2 | ENSMUSG( | ENSMUSG( | 0.945002 | 0.317098 | NO | NO | NA |
| Ppm1n | ENSMUSG( | ENSMUSG( | -0.50614 | 0.126934 | NO | NO | NA |
| Gm36940 | ENSMUSG( | ENSMUSG( | 0.745577 | 0.426029 | NO | NO | NA |
| Trp53-ps | ENSMUSG( | ENSMUSG( | 0.625128 | 0.413484 | NO | NO | NA |
| 1700084E: | ENSMUSG( | ENSMUSG( | -0.03483 | 0.912407 | NO | NO | NA |
| Ighv8-5 | ENSMUSG( | ENSMUSG( | 6.727368 | NA | NO | NO | NA |
| Gm48673 | ENSMUSG( | ENSMUSG( | 0.436174 | 0.538134 | NO | NO | NA |
| Gm37780 | ENSMUSG( | ENSMUSG( | -0.08436 | 0.827325 | NO | NO | NA |
| Gm43581 | ENSMUSG( | ENSMUSG( | 0.411382 | 0.504794 | NO | NO | NA |
| 1810008I1 | ENSMUSG( | ENSMUSG( | 0.086417 | 0.942177 | NO | NO | NA |
| Gm49361 | ENSMUSG( | ENSMUSG( | -5.9688 | NA | NO | NO | NA |
| Tmem30c | ENSMUSG( | ENSMUSG( | 0.691708 | 0.440116 | NO | NO | NA |
| Gm12537 | ENSMUSG( | ENSMUSG( | 0.782354 | 0.015637 | YES | NO | NA |

| | | | | | | | |
|---|---|---|---|---|---|---|---|
| Rfpl4 | ENSMUSG( | ENSMUSG( | 1.112467 | 0.232063 | NO | NO | NA |
| 4933421A( | ENSMUSG( | ENSMUSG( | 0.439256 | 0.426541 | NO | NO | NA |
| Hoxd8 | ENSMUSG( | ENSMUSG( | 0.676624 | 0.444377 | NO | NO | NA |
| Gm27048 | ENSMUSG( | ENSMUSG( | 0.46417 | 0.69475 | NO | NO | NA |
| Gm20163 | ENSMUSG( | ENSMUSG( | -0.28838 | 0.417957 | NO | NO | NA |
| Myt1 | ENSMUSG( | ENSMUSG( | 0.626349 | 0.432246 | NO | NO | NA |
| Chrm4 | ENSMUSG( | ENSMUSG( | -0.65248 | 0.03685 | YES | NO | NA |
| Gm47585 | ENSMUSG( | ENSMUSG( | 0.145404 | 0.796234 | NO | NO | NA |
| H2-DMb2 | ENSMUSG( | ENSMUSG( | 1.224111 | 0.151822 | NO | NO | NA |
| Chil5 | ENSMUSG( | ENSMUSG( | -1.71008 | 1.76E-05 | YES | YES | Downregulated |
| Gm45555 | ENSMUSG( | ENSMUSG( | 0.123158 | 0.853129 | NO | NO | NA |
| Lbx1 | ENSMUSG( | ENSMUSG( | 0.711356 | 0.458226 | NO | NO | NA |
| Igkv4-55 | ENSMUSG( | ENSMUSG( | -7.70515 | 2.29E-08 | YES | YES | Downregulated |
| Vmn1r85 | ENSMUSG( | ENSMUSG( | 0.568314 | 0.587391 | NO | NO | NA |
| Tesmin | ENSMUSG( | ENSMUSG( | 1.499411 | 0.011466 | YES | YES | Upregulated |
| Gm44997 | ENSMUSG( | ENSMUSG( | 0.564331 | 0.581127 | NO | NO | NA |
| Opn3 | ENSMUSG( | ENSMUSG( | 0.679743 | 0.022697 | YES | NO | NA |
| Pdk4 | ENSMUSG( | ENSMUSG( | 0.840317 | 0.335852 | NO | NO | NA |
| Dnhd1 | ENSMUSG( | ENSMUSG( | -0.10301 | 0.890997 | NO | NO | NA |
| Amt | ENSMUSG( | ENSMUSG( | 0.414732 | 0.427345 | NO | NO | NA |
| Gm5124 | ENSMUSG( | ENSMUSG( | -0.55248 | 0.866209 | NO | NO | NA |
| D330037F( | ENSMUSG( | ENSMUSG( | 1.426248 | 0.210776 | NO | NO | NA |
| 4921516A( | ENSMUSG( | ENSMUSG( | 0.263312 | 0.667635 | NO | NO | NA |
| 0610039K: | ENSMUSG( | ENSMUSG( | -0.3316 | 0.508004 | NO | NO | NA |
| Cdk20 | ENSMUSG( | ENSMUSG( | 0.519416 | 0.148506 | NO | NO | NA |
| Gm49937 | ENSMUSG( | ENSMUSG( | 0.487827 | 0.63395 | NO | NO | NA |
| Gm37474 | ENSMUSG( | ENSMUSG( | -0.3889 | 0.481807 | NO | NO | NA |
| Batf3 | ENSMUSG( | ENSMUSG( | 3.102499 | 6.64E-05 | YES | YES | Upregulated |
| Sbf2 | ENSMUSG( | ENSMUSG( | 0.440879 | 0.39517 | NO | NO | NA |
| 4933437G | ENSMUSG( | ENSMUSG( | 0.975429 | 0.395828 | NO | NO | NA |
| Mmp11 | ENSMUSG( | ENSMUSG( | 0.167769 | 0.700632 | NO | NO | NA |
| 9430065F: | ENSMUSG( | ENSMUSG( | 0.63066 | 0.39575 | NO | NO | NA |
| Sh2d1b1 | ENSMUSG( | ENSMUSG( | 0.753985 | 0.405253 | NO | NO | NA |
| Dok6 | ENSMUSG( | ENSMUSG( | 0.636424 | 0.430963 | NO | NO | NA |
| Zfp993 | ENSMUSG( | ENSMUSG( | -1.1728 | 0.02564 | YES | YES | Downregulated |
| Tktl1 | ENSMUSG( | ENSMUSG( | -1.92424 | 0.000114 | YES | YES | Downregulated |
| Zc3h12b | ENSMUSG( | ENSMUSG( | 0.339414 | 0.529433 | NO | NO | NA |
| Gm42496 | ENSMUSG( | ENSMUSG( | 0.806125 | 0.479756 | NO | NO | NA |
| Gm48611 | ENSMUSG( | ENSMUSG( | -0.83685 | 0.00146 | YES | NO | NA |
| Gm45413 | ENSMUSG( | ENSMUSG( | 0.317364 | 0.716868 | NO | NO | NA |
| H3c15 | ENSMUSG( | ENSMUSG( | 0.366229 | 0.570813 | NO | NO | NA |
| 2210408I2 | ENSMUSG( | ENSMUSG( | 0.566158 | 0.04111 | YES | NO | NA |
| Ggnbp1 | ENSMUSG( | ENSMUSG( | 0.113506 | 0.805787 | NO | NO | NA |
| Calb1 | ENSMUSG( | ENSMUSG( | 1.002737 | 0.222402 | NO | NO | NA |
| Gm49616 | ENSMUSG( | ENSMUSG( | -0.19695 | 0.514894 | NO | NO | NA |
| Fam151b | ENSMUSG( | ENSMUSG( | 0.092254 | 0.777257 | NO | NO | NA |
| Sfrp1 | ENSMUSG( | ENSMUSG( | 0.937771 | 0.331156 | NO | NO | NA |

| | | | | | | | |
|---|---|---|---|---|---|---|---|
| Ncs1 | ENSMUSG( | ENSMUSG( | 0.766422 | 0.275655 | NO | NO | NA |
| Gm46597 | ENSMUSG( | ENSMUSG( | 0.04646 | 0.925531 | NO | NO | NA |
| Olfr922 | ENSMUSG( | ENSMUSG( | 0.693121 | 0.512637 | NO | NO | NA |
| Gm45838 | ENSMUSG( | ENSMUSG( | 0.361054 | 0.333517 | NO | NO | NA |
| Gm44659 | ENSMUSG( | ENSMUSG( | 0.876716 | 0.049325 | YES | NO | NA |
| Gm48735 | ENSMUSG( | ENSMUSG( | -0.03426 | 0.930442 | NO | NO | NA |
| Oaz3 | ENSMUSG( | ENSMUSG( | -0.06493 | 0.888811 | NO | NO | NA |
| Zmat4 | ENSMUSG( | ENSMUSG( | 0.507613 | 0.644893 | NO | NO | NA |
| Gm13986 | ENSMUSG( | ENSMUSG( | 1.145962 | 0.258156 | NO | NO | NA |
| Gm48346 | ENSMUSG( | ENSMUSG( | 1.00379 | 0.338282 | NO | NO | NA |
| Apba2 | ENSMUSG( | ENSMUSG( | 0.64475 | 0.466943 | NO | NO | NA |
| Mfsd9 | ENSMUSG( | ENSMUSG( | -0.31573 | 0.422217 | NO | NO | NA |
| Pld1 | ENSMUSG( | ENSMUSG( | 0.705228 | 0.292847 | NO | NO | NA |
| Prrt3 | ENSMUSG( | ENSMUSG( | 0.272639 | 0.677554 | NO | NO | NA |
| Prdm5 | ENSMUSG( | ENSMUSG( | 0.885321 | 0.000738 | YES | NO | NA |
| Apol10a | ENSMUSG( | ENSMUSG( | 0.946206 | 0.109799 | NO | NO | NA |
| Adamtsl1 | ENSMUSG( | ENSMUSG( | 0.482544 | 0.651522 | NO | NO | NA |
| Selp | ENSMUSG( | ENSMUSG( | 1.066916 | 0.2898 | NO | NO | NA |
| Gabbr2 | ENSMUSG( | ENSMUSG( | 0.659966 | 0.514619 | NO | NO | NA |
| Gm26631 | ENSMUSG( | ENSMUSG( | 0.255262 | 0.562501 | NO | NO | NA |
| Gm44907 | ENSMUSG( | ENSMUSG( | 0.916245 | 0.250293 | NO | NO | NA |
| Gm38020 | ENSMUSG( | ENSMUSG( | -0.22976 | 0.584439 | NO | NO | NA |
| Gm39556 | ENSMUSG( | ENSMUSG( | 0.425122 | 0.399941 | NO | NO | NA |
| Tmem178l | ENSMUSG( | ENSMUSG( | 1.0557 | 0.324401 | NO | NO | NA |
| Hoxc11 | ENSMUSG( | ENSMUSG( | 1.05609 | 0.320533 | NO | NO | NA |
| Mir124-2h | ENSMUSG( | ENSMUSG( | 0.666257 | 0.496053 | NO | NO | NA |
| Hoxa7 | ENSMUSG( | ENSMUSG( | 0.589984 | 0.49711 | NO | NO | NA |
| Cav2 | ENSMUSG( | ENSMUSG( | 0.96764 | 0.010207 | YES | NO | NA |
| Nwd2 | ENSMUSG( | ENSMUSG( | 0.702454 | 0.513715 | NO | NO | NA |
| Cavin1 | ENSMUSG( | ENSMUSG( | 0.4786 | 0.553112 | NO | NO | NA |
| Cmya5 | ENSMUSG( | ENSMUSG( | -0.15366 | 0.626728 | NO | NO | NA |
| Gm45633 | ENSMUSG( | ENSMUSG( | 0.446089 | 0.695107 | NO | NO | NA |
| Tmc1 | ENSMUSG( | ENSMUSG( | 0.505797 | 0.614225 | NO | NO | NA |
| Rtn2 | ENSMUSG( | ENSMUSG( | 0.574351 | 0.178773 | NO | NO | NA |
| A630072L: | ENSMUSG( | ENSMUSG( | -0.47961 | 0.24161 | NO | NO | NA |
| ENSMUSG | ENSMUSG( | ENSMUSG( | -0.40801 | NA | NO | NO | NA |
| Gm37570 | ENSMUSG( | ENSMUSG( | 0.268786 | 0.515371 | NO | NO | NA |
| Clca3a1 | ENSMUSG( | ENSMUSG( | 1.164978 | 0.153118 | NO | NO | NA |
| Gm48745 | ENSMUSG( | ENSMUSG( | -0.5071 | 0.171648 | NO | NO | NA |
| 9430091E: | ENSMUSG( | ENSMUSG( | -0.33707 | 0.296789 | NO | NO | NA |
| Sbsn | ENSMUSG( | ENSMUSG( | 0.226823 | 0.672518 | NO | NO | NA |
| Tmem132l | ENSMUSG( | ENSMUSG( | 0.399296 | 0.701752 | NO | NO | NA |
| Rmrp | ENSMUSG( | ENSMUSG( | 1.456306 | 0.089784 | NO | NO | NA |
| 1810013D | ENSMUSG( | ENSMUSG( | 0.512783 | 0.596711 | NO | NO | NA |
| Gm43858 | ENSMUSG( | ENSMUSG( | -0.23722 | 0.521168 | NO | NO | NA |
| Foxf1 | ENSMUSG( | ENSMUSG( | 0.706264 | 0.469178 | NO | NO | NA |
| Olfr898 | ENSMUSG( | ENSMUSG( | 0.63521 | 0.549519 | NO | NO | NA |

| | | | | | | | |
|---|---|---|---|---|---|---|---|
| Gm37525 | ENSMUSG( | ENSMUSG( | 0.726504 | 0.526568 | NO | NO | NA |
| Gm43143 | ENSMUSG( | ENSMUSG( | 0.426113 | 0.241308 | NO | NO | NA |
| Gm6204 | ENSMUSG( | ENSMUSG( | -0.06963 | 0.860746 | NO | NO | NA |
| Gm4430 | ENSMUSG( | ENSMUSG( | 0.112625 | 0.868739 | NO | NO | NA |
| Gm5605 | ENSMUSG( | ENSMUSG( | 0.274788 | 0.755526 | NO | NO | NA |
| Pitpnm2os | ENSMUSG( | ENSMUSG( | 0.259068 | 0.656473 | NO | NO | NA |
| Gm4279 | ENSMUSG( | ENSMUSG( | 0.979571 | 0.353977 | NO | NO | NA |
| Zic4 | ENSMUSG( | ENSMUSG( | 0.423707 | 0.683724 | NO | NO | NA |
| Slc15a1 | ENSMUSG( | ENSMUSG( | 0.956168 | 0.32203 | NO | NO | NA |
| Gm42734 | ENSMUSG( | ENSMUSG( | 1.151461 | 0.151887 | NO | NO | NA |
| Gm37677 | ENSMUSG( | ENSMUSG( | -0.10862 | 0.746718 | NO | NO | NA |
| Gm37144 | ENSMUSG( | ENSMUSG( | 0.508804 | 0.296702 | NO | NO | NA |
| Efnb3 | ENSMUSG( | ENSMUSG( | -0.36008 | 0.357098 | NO | NO | NA |
| A730063M | ENSMUSG( | ENSMUSG( | 1.056365 | 0.003348 | YES | YES | Upregulated |
| A130012E | ENSMUSG( | ENSMUSG( | 0.210082 | 0.76281 | NO | NO | NA |
| Gm19085 | ENSMUSG( | ENSMUSG( | 0.370754 | 0.757619 | NO | NO | NA |
| Gm45553 | ENSMUSG( | ENSMUSG( | -0.34213 | 0.198269 | NO | NO | NA |
| Gm49463 | ENSMUSG( | ENSMUSG( | 0.172814 | 0.663248 | NO | NO | NA |
| Agmo | ENSMUSG( | ENSMUSG( | 0.725907 | 0.48793 | NO | NO | NA |
| Gm30934 | ENSMUSG( | ENSMUSG( | -0.35417 | 0.233626 | NO | NO | NA |
| Gm49492 | ENSMUSG( | ENSMUSG( | -0.25247 | 0.589848 | NO | NO | NA |
| Selenbp1 | ENSMUSG( | ENSMUSG( | 0.348621 | 0.478566 | NO | NO | NA |
| ENSMUSG | ENSMUSG( | ENSMUSG( | 0.92056 | 0.115017 | NO | NO | NA |
| Trav9d-4 | ENSMUSG( | ENSMUSG( | 1.196159 | 0.51563 | NO | NO | NA |
| 1700066B | ENSMUSG( | ENSMUSG( | 0.706294 | 0.005399 | YES | NO | NA |
| Gdpd1 | ENSMUSG( | ENSMUSG( | 0.701089 | 0.217858 | NO | NO | NA |
| 2410004P( | ENSMUSG( | ENSMUSG( | -0.13548 | 0.736794 | NO | NO | NA |
| Col27a1 | ENSMUSG( | ENSMUSG( | 0.841516 | 0.292026 | NO | NO | NA |
| Gm12254 | ENSMUSG( | ENSMUSG( | 0.445412 | 0.10574 | NO | NO | NA |
| Gm26632 | ENSMUSG( | ENSMUSG( | 1.155614 | 0.320062 | NO | NO | NA |
| Masp1 | ENSMUSG( | ENSMUSG( | 0.683967 | 0.500772 | NO | NO | NA |
| Hddc3 | ENSMUSG( | ENSMUSG( | 0.280304 | 0.51866 | NO | NO | NA |
| Gm9214 | ENSMUSG( | ENSMUSG( | 1.070345 | 0.190304 | NO | NO | NA |
| Gm26814 | ENSMUSG( | ENSMUSG( | 1.245232 | 0.238332 | NO | NO | NA |
| Gm44130 | ENSMUSG( | ENSMUSG( | 0.385267 | 0.614734 | NO | NO | NA |
| H2-Oa | ENSMUSG( | ENSMUSG( | 0.365545 | 0.154171 | NO | NO | NA |
| Scamp1 | ENSMUSG( | ENSMUSG( | 1.299457 | 0.018803 | YES | YES | Upregulated |
| Mmrn2 | ENSMUSG( | ENSMUSG( | 0.58883 | 0.491395 | NO | NO | NA |
| Tnnt3 | ENSMUSG( | ENSMUSG( | -0.39767 | 0.398122 | NO | NO | NA |
| Gm42616 | ENSMUSG( | ENSMUSG( | 0.679537 | 0.495445 | NO | NO | NA |
| Gm48700 | ENSMUSG( | ENSMUSG( | 0.653299 | 0.52852 | NO | NO | NA |
| 9430081H | ENSMUSG( | ENSMUSG( | 0.385971 | 0.289017 | NO | NO | NA |
| Gm37730 | ENSMUSG( | ENSMUSG( | 1.111822 | 0.25243 | NO | NO | NA |
| Olfr466 | ENSMUSG( | ENSMUSG( | 0.757776 | 0.527421 | NO | NO | NA |
| Gm15753 | ENSMUSG( | ENSMUSG( | -0.06367 | 0.873059 | NO | NO | NA |
| Gm37621 | ENSMUSG( | ENSMUSG( | 0.552235 | 0.160056 | NO | NO | NA |
| Gm44250 | ENSMUSG( | ENSMUSG( | -0.2199 | 0.50674 | NO | NO | NA |

| | | | | | | | |
|---|---|---|---|---|---|---|---|
| Adamts18 | ENSMUSG( | ENSMUSG( | 0.530206 | 0.597896 | NO | NO | NA |
| A830005F: | ENSMUSG( | ENSMUSG( | -1.29012 | 0.000269 | YES | YES | Downregulated |
| Gm38575 | ENSMUSG( | ENSMUSG( | 1.053362 | 0.354826 | NO | NO | NA |
| Pax2 | ENSMUSG( | ENSMUSG( | 1.122326 | 0.320101 | NO | NO | NA |
| Unc79 | ENSMUSG( | ENSMUSG( | 0.404763 | 0.351852 | NO | NO | NA |
| Pabpc2 | ENSMUSG( | ENSMUSG( | 0.813129 | 0.461948 | NO | NO | NA |
| S100a1 | ENSMUSG( | ENSMUSG( | 0.365953 | 0.267161 | NO | NO | NA |
| Hsf2bp | ENSMUSG( | ENSMUSG( | 0.748924 | 0.402033 | NO | NO | NA |
| Prickle2 | ENSMUSG( | ENSMUSG( | 0.491777 | 0.463112 | NO | NO | NA |
| Igkv16-104 | ENSMUSG( | ENSMUSG( | -1.0402 | 0.567401 | NO | NO | NA |
| Gm43460 | ENSMUSG( | ENSMUSG( | 0.069048 | 0.847579 | NO | NO | NA |
| Prok2 | ENSMUSG( | ENSMUSG( | 0.799495 | 0.445944 | NO | NO | NA |
| 0610040B: | ENSMUSG( | ENSMUSG( | 0.347249 | 0.306823 | NO | NO | NA |
| Papss2 | ENSMUSG( | ENSMUSG( | 0.810777 | 0.263917 | NO | NO | NA |
| Camk2a | ENSMUSG( | ENSMUSG( | 0.475947 | 0.581457 | NO | NO | NA |
| ENSMUSG | ENSMUSG( | ENSMUSG( | 0.225537 | 0.488115 | NO | NO | NA |
| Rfx4 | ENSMUSG( | ENSMUSG( | 2.232669 | 1.79E-05 | YES | YES | Upregulated |
| Gm21986 | ENSMUSG( | ENSMUSG( | 0.231925 | 0.467168 | NO | NO | NA |
| D930020B | ENSMUSG( | ENSMUSG( | 0.700823 | 0.542061 | NO | NO | NA |
| Gm8233 | ENSMUSG( | ENSMUSG( | -0.75603 | 0.021828 | YES | NO | NA |
| C130089K( | ENSMUSG( | ENSMUSG( | 0.995246 | 0.361191 | NO | NO | NA |
| Gm15541 | ENSMUSG( | ENSMUSG( | 0.157376 | 0.809603 | NO | NO | NA |
| 4930532G | ENSMUSG( | ENSMUSG( | 0.421711 | 0.500155 | NO | NO | NA |
| ENSMUSG | ENSMUSG( | ENSMUSG( | 0.280952 | 0.677897 | NO | NO | NA |
| Rpl27-ps3 | ENSMUSG( | ENSMUSG( | 0.227666 | 0.746387 | NO | NO | NA |
| Gm21050 | ENSMUSG( | ENSMUSG( | 1.092664 | 0.363089 | NO | NO | NA |
| Prkn | ENSMUSG( | ENSMUSG( | 0.468048 | 0.669998 | NO | NO | NA |
| 4930572G | ENSMUSG( | ENSMUSG( | 0.698861 | 0.479548 | NO | NO | NA |
| Jcad | ENSMUSG( | ENSMUSG( | 0.945227 | 0.098347 | NO | NO | NA |
| Nudt10 | ENSMUSG( | ENSMUSG( | 0.270934 | 0.474914 | NO | NO | NA |
| 2210408F: | ENSMUSG( | ENSMUSG( | 0.359661 | 0.629263 | NO | NO | NA |
| Pcdh11x | ENSMUSG( | ENSMUSG( | 0.429402 | 0.64009 | NO | NO | NA |
| Olfr190 | ENSMUSG( | ENSMUSG( | 0.496623 | 0.661555 | NO | NO | NA |
| Pbld1 | ENSMUSG( | ENSMUSG( | 0.575299 | 0.58825 | NO | NO | NA |
| Shcbp1l | ENSMUSG( | ENSMUSG( | 1.620697 | 1.83E-05 | YES | YES | Upregulated |
| Akr1c19 | ENSMUSG( | ENSMUSG( | 0.896557 | 0.422361 | NO | NO | NA |
| 5031425F: | ENSMUSG( | ENSMUSG( | 0.274014 | 0.474856 | NO | NO | NA |
| Hoxc10 | ENSMUSG( | ENSMUSG( | 0.769499 | 0.461948 | NO | NO | NA |
| Vmn2r25 | ENSMUSG( | ENSMUSG( | 0.306942 | 0.775138 | NO | NO | NA |
| ENSMUSG( | ENSMUSG( | ENSMUSG( | 0.536089 | 0.607285 | NO | NO | NA |
| B230377A | ENSMUSG( | ENSMUSG( | 0.9794 | 0.000237 | YES | NO | NA |
| 4930444F( | ENSMUSG( | ENSMUSG( | 0.966656 | 0.32205 | NO | NO | NA |
| Gm13807 | ENSMUSG( | ENSMUSG( | 0.312433 | 0.330612 | NO | NO | NA |
| Cplx2 | ENSMUSG( | ENSMUSG( | 0.630989 | 0.263297 | NO | NO | NA |
| Ifitm7 | ENSMUSG( | ENSMUSG( | 1.325435 | 0.250296 | NO | NO | NA |
| Ccdc39 | ENSMUSG( | ENSMUSG( | 0.515388 | 0.476823 | NO | NO | NA |
| Trav5-1 | ENSMUSG( | ENSMUSG( | 0.047153 | 0.953489 | NO | NO | NA |

| | | | | | | | |
|---|---|---|---|---|---|---|---|
| Gm13568 | ENSMUSG( | ENSMUSG( | -0.26709 | 0.558676 | NO | NO | NA |
| 4933423P: | ENSMUSG( | ENSMUSG( | 0.463775 | 0.465889 | NO | NO | NA |
| Olfr709-ps | ENSMUSG( | ENSMUSG( | 0.564983 | 0.617144 | NO | NO | NA |
| Fat3 | ENSMUSG( | ENSMUSG( | 0.707516 | 0.486632 | NO | NO | NA |
| Nscme3l | ENSMUSG( | ENSMUSG( | -0.46162 | 0.28536 | NO | NO | NA |
| Gm37091 | ENSMUSG( | ENSMUSG( | 1.2367 | 0.157497 | NO | NO | NA |
| Gm50398 | ENSMUSG( | ENSMUSG( | 0.655625 | 0.448912 | NO | NO | NA |
| Gm17816 | ENSMUSG( | ENSMUSG( | -0.50039 | 0.10947 | NO | NO | NA |
| ENSMUSG | ENSMUSG( | ENSMUSG( | 0.925126 | 0.039547 | YES | NO | NA |
| Gm37333 | ENSMUSG( | ENSMUSG( | 0.661404 | 0.507007 | NO | NO | NA |
| Gm44424 | ENSMUSG( | ENSMUSG( | -0.23306 | 0.518069 | NO | NO | NA |
| Olfr341 | ENSMUSG( | ENSMUSG( | 0.735673 | 0.508226 | NO | NO | NA |
| Olfr591 | ENSMUSG( | ENSMUSG( | 1.310994 | 0.175573 | NO | NO | NA |
| Klf14 | ENSMUSG( | ENSMUSG( | 0.939134 | 0.29245 | NO | NO | NA |
| Efnb1 | ENSMUSG( | ENSMUSG( | 0.628118 | 0.074422 | NO | NO | NA |
| Gm42501 | ENSMUSG( | ENSMUSG( | 0.386317 | 0.186443 | NO | NO | NA |
| Ighv14-1 | ENSMUSG( | ENSMUSG( | -3.89161 | 0.172742 | NO | NO | NA |
| 3222401L: | ENSMUSG( | ENSMUSG( | 0.659878 | 0.477841 | NO | NO | NA |
| Nova1 | ENSMUSG( | ENSMUSG( | 0.507328 | 0.647956 | NO | NO | NA |
| Kcnrg | ENSMUSG( | ENSMUSG( | -1.04064 | 0.012239 | YES | YES | Downregulated |
| Plekhg4 | ENSMUSG( | ENSMUSG( | 2.065808 | 1.06E-07 | YES | YES | Upregulated |
| A530041M | ENSMUSG( | ENSMUSG( | 0.235348 | 0.56965 | NO | NO | NA |
| Wnt8b | ENSMUSG( | ENSMUSG( | 0.136184 | 0.90857 | NO | NO | NA |
| Gm14634 | ENSMUSG( | ENSMUSG( | 1.385966 | 0.010721 | YES | YES | Upregulated |
| Csrnp3 | ENSMUSG( | ENSMUSG( | 0.478674 | 0.669989 | NO | NO | NA |
| Gm49356 | ENSMUSG( | ENSMUSG( | 0.328354 | 0.483164 | NO | NO | NA |
| Dagla | ENSMUSG( | ENSMUSG( | -0.64927 | 0.04607 | YES | NO | NA |
| Serinc4 | ENSMUSG( | ENSMUSG( | 0.048202 | 0.948339 | NO | NO | NA |
| Pard6g | ENSMUSG( | ENSMUSG( | 0.806859 | 0.01346 | YES | NO | NA |
| Gm37485 | ENSMUSG( | ENSMUSG( | 0.9072 | 0.390199 | NO | NO | NA |
| Mtus1 | ENSMUSG( | ENSMUSG( | -0.05479 | 0.92302 | NO | NO | NA |
| Olfr332 | ENSMUSG( | ENSMUSG( | 0.546213 | 0.62031 | NO | NO | NA |
| 4930589O | ENSMUSG( | ENSMUSG( | 0.158588 | 0.832943 | NO | NO | NA |
| Gm44141 | ENSMUSG( | ENSMUSG( | 0.08407 | 0.89391 | NO | NO | NA |
| Gm10723 | ENSMUSG( | ENSMUSG( | 0.911774 | 0.428235 | NO | NO | NA |
| Tdrp | ENSMUSG( | ENSMUSG( | 0.516456 | 0.588955 | NO | NO | NA |
| Gm10482 | ENSMUSG( | ENSMUSG( | 0.878855 | 0.422105 | NO | NO | NA |
| Gm37289 | ENSMUSG( | ENSMUSG( | -0.12309 | 0.746663 | NO | NO | NA |
| C230035I1 | ENSMUSG( | ENSMUSG( | 1.344147 | 0.003503 | YES | YES | Upregulated |
| Gm44967 | ENSMUSG( | ENSMUSG( | 0.811292 | 0.416815 | NO | NO | NA |
| Gm11973 | ENSMUSG( | ENSMUSG( | 0.145167 | 0.786674 | NO | NO | NA |
| Plp1 | ENSMUSG( | ENSMUSG( | 0.760766 | 0.016104 | YES | NO | NA |
| Gm37366 | ENSMUSG( | ENSMUSG( | 0.787542 | 0.132004 | NO | NO | NA |
| Gm32390 | ENSMUSG( | ENSMUSG( | 0.863557 | 0.45895 | NO | NO | NA |
| Nthl1 | ENSMUSG( | ENSMUSG( | -0.08556 | 0.813733 | NO | NO | NA |
| Gm37581 | ENSMUSG( | ENSMUSG( | 0.087783 | 0.792881 | NO | NO | NA |
| Stmn2 | ENSMUSG( | ENSMUSG( | 0.919155 | 0.021104 | YES | NO | NA |

| | | | | | | | |
|---|---|---|---|---|---|---|---|
| Syn2 | ENSMUSG( | ENSMUSG( | 0.581351 | 0.571086 | NO | NO | NA |
| Insm2 | ENSMUSG( | ENSMUSG( | 0.510704 | 0.640929 | NO | NO | NA |
| Gm16169 | ENSMUSG( | ENSMUSG( | 0.645699 | 0.569763 | NO | NO | NA |
| Lrig1 | ENSMUSG( | ENSMUSG( | 0.811515 | 0.105756 | NO | NO | NA |
| Gm43313 | ENSMUSG( | ENSMUSG( | -0.25583 | 0.502106 | NO | NO | NA |
| Sfrp2 | ENSMUSG( | ENSMUSG( | 1.112937 | 0.21991 | NO | NO | NA |
| B430010I2 | ENSMUSG( | ENSMUSG( | 0.392688 | 0.709866 | NO | NO | NA |
| A230051N | ENSMUSG( | ENSMUSG( | 0.905288 | 0.337533 | NO | NO | NA |
| Vmn1r203 | ENSMUSG( | ENSMUSG( | 0.895548 | 0.395366 | NO | NO | NA |
| 2900093K: | ENSMUSG( | ENSMUSG( | 0.277356 | 0.301328 | NO | NO | NA |
| Cnga3 | ENSMUSG( | ENSMUSG( | 1.31638 | 0.088295 | NO | NO | NA |
| Olfr1388 | ENSMUSG( | ENSMUSG( | 0.844332 | 0.025035 | YES | NO | NA |
| Gm47093 | ENSMUSG( | ENSMUSG( | 0.712064 | 0.475134 | NO | NO | NA |
| Hdac1-ps | ENSMUSG( | ENSMUSG( | 0.180689 | 0.592341 | NO | NO | NA |
| Gm7340 | ENSMUSG( | ENSMUSG( | 0.942934 | 0.150817 | NO | NO | NA |
| ENSMUSG | ENSMUSG( | ENSMUSG( | 0.315207 | 0.630689 | NO | NO | NA |
| Gm43496 | ENSMUSG( | ENSMUSG( | 0.625432 | 0.5592 | NO | NO | NA |
| Gm44663 | ENSMUSG( | ENSMUSG( | -0.57151 | 0.410282 | NO | NO | NA |
| Slc28a3 | ENSMUSG( | ENSMUSG( | 0.582382 | 0.534573 | NO | NO | NA |
| 5430434F( | ENSMUSG( | ENSMUSG( | 0.39257 | 0.562892 | NO | NO | NA |
| Hoxa3 | ENSMUSG( | ENSMUSG( | 0.915208 | 0.415524 | NO | NO | NA |
| Zfp385b | ENSMUSG( | ENSMUSG( | 0.752158 | 0.481239 | NO | NO | NA |
| Fgfr2 | ENSMUSG( | ENSMUSG( | 0.72105 | 0.531711 | NO | NO | NA |
| Lonrf1 | ENSMUSG( | ENSMUSG( | 0.569497 | 0.073342 | NO | NO | NA |
| Gm16170 | ENSMUSG( | ENSMUSG( | 0.873763 | 0.257847 | NO | NO | NA |
| Gm26520 | ENSMUSG( | ENSMUSG( | -0.58606 | 0.2531 | NO | NO | NA |
| Trarg1 | ENSMUSG( | ENSMUSG( | 0.425377 | 0.659678 | NO | NO | NA |
| Gm45211 | ENSMUSG( | ENSMUSG( | 0.952204 | 0.358384 | NO | NO | NA |
| Fbxo10 | ENSMUSG( | ENSMUSG( | -0.01139 | 0.981881 | NO | NO | NA |
| Gm38200 | ENSMUSG( | ENSMUSG( | 0.739819 | 0.380479 | NO | NO | NA |
| Gm16124 | ENSMUSG( | ENSMUSG( | 0.472202 | 0.131748 | NO | NO | NA |
| Fam222a | ENSMUSG( | ENSMUSG( | 0.203561 | 0.600449 | NO | NO | NA |
| 5730585A | ENSMUSG( | ENSMUSG( | 0.98168 | 0.398195 | NO | NO | NA |
| Lmx1b | ENSMUSG( | ENSMUSG( | 0.654547 | 0.522926 | NO | NO | NA |
| Obscn | ENSMUSG( | ENSMUSG( | -0.83563 | 0.034919 | YES | NO | NA |
| ENSMUSG | ENSMUSG( | ENSMUSG( | -0.0752 | 0.92214 | NO | NO | NA |
| Vmn2r3 | ENSMUSG( | ENSMUSG( | 0.313202 | 0.764213 | NO | NO | NA |
| 1700023H | ENSMUSG( | ENSMUSG( | -0.55292 | 0.102184 | NO | NO | NA |
| Cst6 | ENSMUSG( | ENSMUSG( | 0.78855 | 0.188868 | NO | NO | NA |
| Gm38329 | ENSMUSG( | ENSMUSG( | -0.54183 | 0.217409 | NO | NO | NA |
| Evi5 | ENSMUSG( | ENSMUSG( | 0.373329 | 0.420436 | NO | NO | NA |
| Plppr4 | ENSMUSG( | ENSMUSG( | 0.752035 | 0.473151 | NO | NO | NA |
| Dlk1 | ENSMUSG( | ENSMUSG( | 0.587148 | 0.464134 | NO | NO | NA |
| ENSMUSG | ENSMUSG( | ENSMUSG( | 0.269342 | 0.642536 | NO | NO | NA |
| Nme4 | ENSMUSG( | ENSMUSG( | 0.858226 | 0.100739 | NO | NO | NA |
| Gm14324 | ENSMUSG( | ENSMUSG( | 0.48771 | 0.658376 | NO | NO | NA |
| Thnsl2 | ENSMUSG( | ENSMUSG( | 0.442341 | 0.264045 | NO | NO | NA |

| | | | | | | | |
|---|---|---|---|---|---|---|---|
| Pfkfb4 | ENSMUSG( | ENSMUSG( | -0.03836 | 0.92032 | NO | NO | NA |
| Gm14438 | ENSMUSG( | ENSMUSG( | 0.656094 | 0.451796 | NO | NO | NA |
| A330069E | ENSMUSG( | ENSMUSG( | 0.183359 | 0.600373 | NO | NO | NA |
| Gm37045 | ENSMUSG( | ENSMUSG( | 0.797987 | 0.159563 | NO | NO | NA |
| Gm10353 | ENSMUSG( | ENSMUSG( | -0.16843 | 0.580155 | NO | NO | NA |
| Gm11802 | ENSMUSG( | ENSMUSG( | -0.08724 | 0.917351 | NO | NO | NA |
| Svopl | ENSMUSG( | ENSMUSG( | 1.168748 | 0.264147 | NO | NO | NA |
| Gm48714 | ENSMUSG( | ENSMUSG( | 0.40825 | 0.102635 | NO | NO | NA |
| 4930558J1 | ENSMUSG( | ENSMUSG( | 0.142424 | 0.601079 | NO | NO | NA |
| Sostdc1 | ENSMUSG( | ENSMUSG( | 0.998086 | 0.331327 | NO | NO | NA |
| Gm44771 | ENSMUSG( | ENSMUSG( | 0.379046 | 0.560529 | NO | NO | NA |
| Gm6899 | ENSMUSG( | ENSMUSG( | 0.715005 | 0.338943 | NO | NO | NA |
| Tinagl1 | ENSMUSG( | ENSMUSG( | 0.767282 | 0.11648 | NO | NO | NA |
| Ehbp1 | ENSMUSG( | ENSMUSG( | 0.522935 | 0.236987 | NO | NO | NA |
| Pmepa1 | ENSMUSG( | ENSMUSG( | -0.40089 | 0.455732 | NO | NO | NA |
| Snrpn | ENSMUSG( | ENSMUSG( | -1.57443 | 0.001584 | YES | YES | Downregulated |
| 6430550D | ENSMUSG( | ENSMUSG( | -0.22271 | 0.692687 | NO | NO | NA |
| Sorcs2 | ENSMUSG( | ENSMUSG( | 0.968815 | 0.287978 | NO | NO | NA |
| Mapk4 | ENSMUSG( | ENSMUSG( | 1.168889 | 0.257725 | NO | NO | NA |
| Asb4 | ENSMUSG( | ENSMUSG( | 0.990175 | 0.348614 | NO | NO | NA |
| Gm42870 | ENSMUSG( | ENSMUSG( | 0.532756 | 0.613276 | NO | NO | NA |
| Bank1 | ENSMUSG( | ENSMUSG( | -0.38415 | 0.700937 | NO | NO | NA |
| Gm47265 | ENSMUSG( | ENSMUSG( | 1.076052 | 0.285303 | NO | NO | NA |
| 1700055D | ENSMUSG( | ENSMUSG( | 0.473374 | 0.287275 | NO | NO | NA |
| Gm19111 | ENSMUSG( | ENSMUSG( | 0.912891 | 0.377249 | NO | NO | NA |
| Gm16062 | ENSMUSG( | ENSMUSG( | -0.23725 | 0.46303 | NO | NO | NA |
| She | ENSMUSG( | ENSMUSG( | 0.962716 | 0.34766 | NO | NO | NA |
| Cpeb3 | ENSMUSG( | ENSMUSG( | 0.541431 | 0.263075 | NO | NO | NA |
| Elfn2 | ENSMUSG( | ENSMUSG( | 0.345315 | 0.672948 | NO | NO | NA |
| Lgr5 | ENSMUSG( | ENSMUSG( | 0.904248 | 0.399515 | NO | NO | NA |
| Gm13629 | ENSMUSG( | ENSMUSG( | 0.812306 | 0.471631 | NO | NO | NA |
| Siglecg | ENSMUSG( | ENSMUSG( | -0.30557 | 0.634312 | NO | NO | NA |
| Gm37766 | ENSMUSG( | ENSMUSG( | 0.307205 | 0.688954 | NO | NO | NA |
| Gm49125 | ENSMUSG( | ENSMUSG( | -0.51204 | 0.069754 | NO | NO | NA |
| Igkv9-120 | ENSMUSG( | ENSMUSG( | 0.278241 | 0.901707 | NO | NO | NA |
| Runx2os1 | ENSMUSG( | ENSMUSG( | 0.152474 | 0.729086 | NO | NO | NA |
| Fam161b | ENSMUSG( | ENSMUSG( | -0.51309 | 0.062408 | NO | NO | NA |
| Gm44704 | ENSMUSG( | ENSMUSG( | 0.449746 | 0.607785 | NO | NO | NA |
| ENSMUSG | ENSMUSG( | ENSMUSG( | 0.716628 | 0.244982 | NO | NO | NA |
| Slc22a23 | ENSMUSG( | ENSMUSG( | 0.344639 | 0.392392 | NO | NO | NA |
| Gm43605 | ENSMUSG( | ENSMUSG( | 0.389203 | 0.729989 | NO | NO | NA |
| Gm43609 | ENSMUSG( | ENSMUSG( | 0.145367 | 0.754398 | NO | NO | NA |
| Steap2 | ENSMUSG( | ENSMUSG( | 1.1834 | 0.267074 | NO | NO | NA |
| Cyp39a1 | ENSMUSG( | ENSMUSG( | 0.639123 | 0.09316 | NO | NO | NA |
| Slco5a1 | ENSMUSG( | ENSMUSG( | 0.130184 | 0.784641 | NO | NO | NA |
| A030001D | ENSMUSG( | ENSMUSG( | 0.907807 | 0.286304 | NO | NO | NA |
| Dhrs13os | ENSMUSG( | ENSMUSG( | 0.588491 | 0.374884 | NO | NO | NA |

| | | | | | | | |
|---|---|---|---|---|---|---|---|
| Src | ENSMUSG( | ENSMUSG( | 0.404741 | 0.527649 | NO | NO | NA |
| Rps19-ps3 | ENSMUSG( | ENSMUSG( | 2.187928 | 1.68E-09 | YES | YES | Upregulated |
| Gm6061 | ENSMUSG( | ENSMUSG( | 0.798642 | 0.706703 | NO | NO | NA |
| Gm45629 | ENSMUSG( | ENSMUSG( | 0.063985 | 0.892502 | NO | NO | NA |
| Gm38412 | ENSMUSG( | ENSMUSG( | 0.316049 | 0.804577 | NO | NO | NA |
| Gm30015 | ENSMUSG( | ENSMUSG( | 0.412881 | 0.704074 | NO | NO | NA |
| Gm45855 | ENSMUSG( | ENSMUSG( | 0.425122 | 0.265263 | NO | NO | NA |
| 4930429F: | ENSMUSG( | ENSMUSG( | 0.791662 | 0.22671 | NO | NO | NA |
| Gm9118 | ENSMUSG( | ENSMUSG( | 0.727595 | 0.017776 | YES | NO | NA |
| Gm48230 | ENSMUSG( | ENSMUSG( | 0.131034 | 0.911269 | NO | NO | NA |
| Pak3 | ENSMUSG( | ENSMUSG( | 0.626942 | 0.539391 | NO | NO | NA |
| Gm29253 | ENSMUSG( | ENSMUSG( | -1.15053 | 0.480378 | NO | NO | NA |
| Prkg2 | ENSMUSG( | ENSMUSG( | 0.769391 | 0.496414 | NO | NO | NA |
| Gm37607 | ENSMUSG( | ENSMUSG( | -0.06596 | 0.897813 | NO | NO | NA |
| Trbv24 | ENSMUSG( | ENSMUSG( | -0.81816 | 0.342668 | NO | NO | NA |
| Gm6754 | ENSMUSG( | ENSMUSG( | 0.609275 | 0.5716 | NO | NO | NA |
| Gm45444 | ENSMUSG( | ENSMUSG( | 0.028091 | 0.932161 | NO | NO | NA |
| 9530056E: | ENSMUSG( | ENSMUSG( | 0.55512 | 0.35827 | NO | NO | NA |
| Gm35909 | ENSMUSG( | ENSMUSG( | 1.005188 | 0.324384 | NO | NO | NA |
| Gm49673 | ENSMUSG( | ENSMUSG( | 1.242117 | 0.447795 | NO | NO | NA |
| Gm50393 | ENSMUSG( | ENSMUSG( | 0.950582 | 0.146868 | NO | NO | NA |
| Ccdc63 | ENSMUSG( | ENSMUSG( | 0.054619 | 0.891292 | NO | NO | NA |
| Btnl9 | ENSMUSG( | ENSMUSG( | 1.278373 | 0.011082 | YES | YES | Upregulated |
| Thsd7a | ENSMUSG( | ENSMUSG( | 0.547216 | 0.584776 | NO | NO | NA |
| Impdh2-ps | ENSMUSG( | ENSMUSG( | 0.931492 | 0.006386 | YES | NO | NA |
| Dcdc2a | ENSMUSG( | ENSMUSG( | -0.00693 | 0.994565 | NO | NO | NA |
| Btf3-ps1 | ENSMUSG( | ENSMUSG( | 0.379935 | 0.350951 | NO | NO | NA |
| Kcnk4 | ENSMUSG( | ENSMUSG( | 0.242635 | 0.758671 | NO | NO | NA |
| Gm47234 | ENSMUSG( | ENSMUSG( | 0.118736 | 0.680419 | NO | NO | NA |
| Atp2a1 | ENSMUSG( | ENSMUSG( | -0.97684 | 0.004235 | YES | NO | NA |
| Zfp72 | ENSMUSG( | ENSMUSG( | 0.344079 | 0.468029 | NO | NO | NA |
| ENSMUSG( | ENSMUSG( | ENSMUSG( | 0.914429 | 0.412463 | NO | NO | NA |
| Gm49494 | ENSMUSG( | ENSMUSG( | 0.638498 | 0.546166 | NO | NO | NA |
| Maoa | ENSMUSG( | ENSMUSG( | 1.20434 | 0.01365 | YES | YES | Upregulated |
| Gm47166 | ENSMUSG( | ENSMUSG( | 0.615683 | 0.473914 | NO | NO | NA |
| Carmil1 | ENSMUSG( | ENSMUSG( | 0.396835 | 0.57383 | NO | NO | NA |
| Chmp4c | ENSMUSG( | ENSMUSG( | -0.12311 | 0.768227 | NO | NO | NA |
| Schip1 | ENSMUSG( | ENSMUSG( | 0.699103 | 0.411823 | NO | NO | NA |
| Gm17968 | ENSMUSG( | ENSMUSG( | 1.044588 | 0.224015 | NO | NO | NA |
| Gm37195 | ENSMUSG( | ENSMUSG( | 0.898996 | 0.413355 | NO | NO | NA |
| H3c10 | ENSMUSG( | ENSMUSG( | 0.382051 | 0.561489 | NO | NO | NA |
| Olfr545 | ENSMUSG( | ENSMUSG( | 0.524917 | 0.57478 | NO | NO | NA |
| Gm48699 | ENSMUSG( | ENSMUSG( | 0.864214 | 0.376154 | NO | NO | NA |
| Gm38910 | ENSMUSG( | ENSMUSG( | 0.480491 | 0.576362 | NO | NO | NA |
| Nr1h3 | ENSMUSG( | ENSMUSG( | -1.02344 | 0.004367 | YES | YES | Downregulated |
| ENSMUSG( | ENSMUSG( | ENSMUSG( | -0.32484 | 0.507429 | NO | NO | NA |
| ENSMUSG | ENSMUSG( | ENSMUSG( | -0.28563 | 0.504097 | NO | NO | NA |

| | | | | | | | |
|---|---|---|---|---|---|---|---|
| Tspan8 | ENSMUSG( | ENSMUSG( | 0.771132 | 0.398144 | NO | NO | NA |
| Gm2115 | ENSMUSG( | ENSMUSG( | 0.583449 | 0.516196 | NO | NO | NA |
| Gm5869 | ENSMUSG( | ENSMUSG( | 0.604274 | 0.100456 | NO | NO | NA |
| Lpin3 | ENSMUSG( | ENSMUSG( | -0.64212 | 0.068542 | NO | NO | NA |
| Arhgdig | ENSMUSG( | ENSMUSG( | 0.928428 | 0.02921 | YES | NO | NA |
| Gm48488 | ENSMUSG( | ENSMUSG( | 0.831166 | 0.471485 | NO | NO | NA |
| Gm10644 | ENSMUSG( | ENSMUSG( | 0.102952 | 0.731535 | NO | NO | NA |
| Tmem91 | ENSMUSG( | ENSMUSG( | 0.465669 | 0.409613 | NO | NO | NA |
| Bcl2l15 | ENSMUSG( | ENSMUSG( | 0.279816 | 0.362869 | NO | NO | NA |
| Lrrc9 | ENSMUSG( | ENSMUSG( | 0.434775 | 0.483444 | NO | NO | NA |
| Spag4 | ENSMUSG( | ENSMUSG( | -0.42527 | 0.142217 | NO | NO | NA |
| Gm38999 | ENSMUSG( | ENSMUSG( | -3.45863 | 0.034857 | YES | YES | Downregulated |
| Gm50103 | ENSMUSG( | ENSMUSG( | 0.223646 | 0.568908 | NO | NO | NA |
| Gm36638 | ENSMUSG( | ENSMUSG( | 0.293212 | 0.525866 | NO | NO | NA |
| Rims1 | ENSMUSG( | ENSMUSG( | 1.10357 | 0.037676 | YES | YES | Upregulated |
| Vmn2r22 | ENSMUSG( | ENSMUSG( | 0.583199 | 0.616416 | NO | NO | NA |
| Pgam2 | ENSMUSG( | ENSMUSG( | 0.392401 | 0.45456 | NO | NO | NA |
| Gm47389 | ENSMUSG( | ENSMUSG( | 1.061662 | 0.090101 | NO | NO | NA |
| Prex2 | ENSMUSG( | ENSMUSG( | 0.316041 | 0.763066 | NO | NO | NA |
| Gm7292 | ENSMUSG( | ENSMUSG( | -0.06969 | 0.916397 | NO | NO | NA |
| Hoxd4 | ENSMUSG( | ENSMUSG( | 0.505166 | 0.641749 | NO | NO | NA |
| Gm42471 | ENSMUSG( | ENSMUSG( | -0.13195 | 0.870425 | NO | NO | NA |
| Gm37928 | ENSMUSG( | ENSMUSG( | 0.716494 | 0.515674 | NO | NO | NA |
| AU015791 | ENSMUSG( | ENSMUSG( | 0.555894 | 0.628823 | NO | NO | NA |
| ENSMUSG | ENSMUSG( | ENSMUSG( | 0.340925 | 0.463646 | NO | NO | NA |
| 6330411D | ENSMUSG( | ENSMUSG( | 1.479467 | 0.117297 | NO | NO | NA |
| Gm8126 | ENSMUSG( | ENSMUSG( | 0.596346 | 0.547351 | NO | NO | NA |
| Kif5a | ENSMUSG( | ENSMUSG( | -0.19477 | 0.668903 | NO | NO | NA |
| ENSMUSG | ENSMUSG( | ENSMUSG( | -0.67073 | 0.141943 | NO | NO | NA |
| Eppk1 | ENSMUSG( | ENSMUSG( | -0.0871 | 0.894145 | NO | NO | NA |
| Rhbdl2 | ENSMUSG( | ENSMUSG( | 0.566924 | 0.594479 | NO | NO | NA |
| Abcc2 | ENSMUSG( | ENSMUSG( | 0.610704 | 0.56241 | NO | NO | NA |
| Zic2 | ENSMUSG( | ENSMUSG( | 0.71207 | 0.556665 | NO | NO | NA |
| Gm48766 | ENSMUSG( | ENSMUSG( | 0.451529 | 0.536077 | NO | NO | NA |
| Rflna | ENSMUSG( | ENSMUSG( | 0.832771 | 0.349117 | NO | NO | NA |
| ENSMUSG | ENSMUSG( | ENSMUSG( | -0.21485 | 0.606938 | NO | NO | NA |
| Kctd21 | ENSMUSG( | ENSMUSG( | -1.18135 | 0.000775 | YES | YES | Downregulated |
| Olfr166 | ENSMUSG( | ENSMUSG( | 1.21782 | 0.025966 | YES | YES | Upregulated |
| Srrm4os | ENSMUSG( | ENSMUSG( | 0.598218 | 0.562937 | NO | NO | NA |
| Gm43429 | ENSMUSG( | ENSMUSG( | 0.997465 | 0.211951 | NO | NO | NA |
| ENSMUSG | ENSMUSG( | ENSMUSG( | 0.704168 | 0.357367 | NO | NO | NA |
| Gm28229 | ENSMUSG( | ENSMUSG( | 0.165926 | 0.574263 | NO | NO | NA |
| Gm36879 | ENSMUSG( | ENSMUSG( | 0.636656 | 0.531711 | NO | NO | NA |
| Tgfbi | ENSMUSG( | ENSMUSG( | -0.36923 | 0.53766 | NO | NO | NA |
| Slc17a7 | ENSMUSG( | ENSMUSG( | -0.2892 | 0.478362 | NO | NO | NA |
| Olfr1183 | ENSMUSG( | ENSMUSG( | 0.966725 | 0.359436 | NO | NO | NA |
| Gm20763 | ENSMUSG( | ENSMUSG( | 0.735906 | 0.502427 | NO | NO | NA |

| | | | | | | | |
|---|---|---|---|---|---|---|---|
| Syt12 | ENSMUSG( | ENSMUSG( | 0.797913 | 0.46303 | NO | NO | NA |
| Opn1sw | ENSMUSG( | ENSMUSG( | 0.665573 | 0.544325 | NO | NO | NA |
| Gm48662 | ENSMUSG( | ENSMUSG( | 0.764136 | 0.501525 | NO | NO | NA |
| Gm21284 | ENSMUSG( | ENSMUSG( | 0.694162 | 0.219882 | NO | NO | NA |
| Gm37368 | ENSMUSG( | ENSMUSG( | 0.372776 | 0.587049 | NO | NO | NA |
| Mras | ENSMUSG( | ENSMUSG( | 0.315065 | 0.595155 | NO | NO | NA |
| Gm10118 | ENSMUSG( | ENSMUSG( | 1.046577 | 0.33074 | NO | NO | NA |
| Gm47542 | ENSMUSG( | ENSMUSG( | 0.593618 | 0.550609 | NO | NO | NA |
| Sned1 | ENSMUSG( | ENSMUSG( | 0.247686 | 0.639331 | NO | NO | NA |
| ENSMUSG | ENSMUSG( | ENSMUSG( | 0.008951 | 0.977906 | NO | NO | NA |
| Nrip2 | ENSMUSG( | ENSMUSG( | 0.222371 | 0.497766 | NO | NO | NA |
| Gm48146 | ENSMUSG( | ENSMUSG( | 0.199222 | 0.733193 | NO | NO | NA |
| Gm42502 | ENSMUSG( | ENSMUSG( | -0.23409 | 0.465889 | NO | NO | NA |
| ENSMUSG | ENSMUSG( | ENSMUSG( | 0.44887 | 0.482105 | NO | NO | NA |
| Erbb2 | ENSMUSG( | ENSMUSG( | 0.190875 | 0.80634 | NO | NO | NA |
| Pcdh15 | ENSMUSG( | ENSMUSG( | 0.491149 | 0.575623 | NO | NO | NA |
| Gm47980 | ENSMUSG( | ENSMUSG( | 0.52389 | 0.576079 | NO | NO | NA |
| Gm15911 | ENSMUSG( | ENSMUSG( | -0.36389 | 0.298258 | NO | NO | NA |
| Npr3 | ENSMUSG( | ENSMUSG( | 0.396531 | 0.712916 | NO | NO | NA |
| Rgmb | ENSMUSG( | ENSMUSG( | -0.117 | 0.84861 | NO | NO | NA |
| Gm15927 | ENSMUSG( | ENSMUSG( | 0.042028 | 0.914446 | NO | NO | NA |
| Scx | ENSMUSG( | ENSMUSG( | -0.13237 | 0.78842 | NO | NO | NA |
| Hormad2 | ENSMUSG( | ENSMUSG( | 3.364932 | 5.68E-11 | YES | YES | Upregulated |
| Gm43747 | ENSMUSG( | ENSMUSG( | -0.21213 | 0.613051 | NO | NO | NA |
| Gm11225 | ENSMUSG( | ENSMUSG( | 1.027405 | 0.301461 | NO | NO | NA |
| ENSMUSG | ENSMUSG( | ENSMUSG( | -0.06917 | 0.865515 | NO | NO | NA |
| Pclo | ENSMUSG( | ENSMUSG( | 0.754447 | 0.454063 | NO | NO | NA |
| Olfr1205 | ENSMUSG( | ENSMUSG( | 1.202761 | 0.21818 | NO | NO | NA |
| Zfp469 | ENSMUSG( | ENSMUSG( | 1.123518 | 0.388544 | NO | NO | NA |
| Trav7n-4 | ENSMUSG( | ENSMUSG( | 0.333542 | 0.805716 | NO | NO | NA |
| Gm3625 | ENSMUSG( | ENSMUSG( | 0.187067 | 0.533073 | NO | NO | NA |
| Gm37737 | ENSMUSG( | ENSMUSG( | 0.602768 | 0.563166 | NO | NO | NA |
| 4930570G | ENSMUSG( | ENSMUSG( | 0.403186 | 0.708667 | NO | NO | NA |
| Rpl38-ps2 | ENSMUSG( | ENSMUSG( | 0.22998 | 0.567401 | NO | NO | NA |
| Gm49695 | ENSMUSG( | ENSMUSG( | 0.689611 | 0.53285 | NO | NO | NA |
| Gm13398 | ENSMUSG( | ENSMUSG( | 0.593965 | 0.518007 | NO | NO | NA |
| Gm43598 | ENSMUSG( | ENSMUSG( | 0.97851 | 0.167959 | NO | NO | NA |
| Proscos | ENSMUSG( | ENSMUSG( | 0.007624 | 0.985536 | NO | NO | NA |
| Gm30363 | ENSMUSG( | ENSMUSG( | 0.438681 | 0.576744 | NO | NO | NA |
| Podxl2 | ENSMUSG( | ENSMUSG( | 0.610916 | 0.538082 | NO | NO | NA |
| Hk3 | ENSMUSG( | ENSMUSG( | -0.01824 | 0.96061 | NO | NO | NA |
| Dok3 | ENSMUSG( | ENSMUSG( | 0.218207 | 0.677675 | NO | NO | NA |
| Gm12905 | ENSMUSG( | ENSMUSG( | -0.18964 | 0.586905 | NO | NO | NA |
| ENSMUSG | ENSMUSG( | ENSMUSG( | -0.18357 | 0.892753 | NO | NO | NA |
| Rdh16f1 | ENSMUSG( | ENSMUSG( | 1.643071 | 0.618667 | NO | NO | NA |
| Efna2 | ENSMUSG( | ENSMUSG( | 0.796015 | 0.357038 | NO | NO | NA |
| A630031M | ENSMUSG( | ENSMUSG( | 0.61214 | 0.56336 | NO | NO | NA |

| | | | | | | | |
|---|---|---|---|---|---|---|---|
| Efna4 | ENSMUSG( | ENSMUSG( | 0.36098 | 0.541669 | NO | NO | NA |
| Gm9754 | ENSMUSG( | ENSMUSG( | -0.2885 | 0.42652 | NO | NO | NA |
| Fam131b | ENSMUSG( | ENSMUSG( | -0.65821 | 0.07154 | NO | NO | NA |
| Ido2 | ENSMUSG( | ENSMUSG( | 0.798614 | 0.517784 | NO | NO | NA |
| Gm37345 | ENSMUSG( | ENSMUSG( | 0.902884 | 0.407641 | NO | NO | NA |
| Gm15824 | ENSMUSG( | ENSMUSG( | 0.44222 | 0.575107 | NO | NO | NA |
| Fbln5 | ENSMUSG( | ENSMUSG( | 1.271843 | 0.174565 | NO | NO | NA |
| Pate1 | ENSMUSG( | ENSMUSG( | 0.992035 | 0.389738 | NO | NO | NA |
| BC060293 | ENSMUSG( | ENSMUSG( | 0.753582 | 0.099161 | NO | NO | NA |
| Arhgef40 | ENSMUSG( | ENSMUSG( | 0.697355 | 0.160056 | NO | NO | NA |
| Gm37069 | ENSMUSG( | ENSMUSG( | -0.6035 | 0.093892 | NO | NO | NA |
| Vmn1r87 | ENSMUSG( | ENSMUSG( | 0.338821 | 0.730279 | NO | NO | NA |
| Dzip1l | ENSMUSG( | ENSMUSG( | 0.363829 | 0.726189 | NO | NO | NA |
| ENSMUSG | ENSMUSG( | ENSMUSG( | 0.919523 | 0.456646 | NO | NO | NA |
| Epx | ENSMUSG( | ENSMUSG( | -4.68593 | 0.003273 | YES | YES | Downregulated |
| Ptk7 | ENSMUSG( | ENSMUSG( | 0.2482 | 0.803272 | NO | NO | NA |
| Gm37125 | ENSMUSG( | ENSMUSG( | 0.937876 | 0.007852 | YES | NO | NA |
| ENSMUSG | ENSMUSG( | ENSMUSG( | -0.1802 | 0.521932 | NO | NO | NA |
| 4833418N | ENSMUSG( | ENSMUSG( | 0.112158 | 0.764246 | NO | NO | NA |
| Fscn1 | ENSMUSG( | ENSMUSG( | 0.066534 | 0.859911 | NO | NO | NA |
| Trav9d-1 | ENSMUSG( | ENSMUSG( | 0.716034 | 0.617724 | NO | NO | NA |
| Pik3r6 | ENSMUSG( | ENSMUSG( | 0.136564 | 0.878139 | NO | NO | NA |
| C630004M | ENSMUSG( | ENSMUSG( | 0.414076 | 0.334983 | NO | NO | NA |
| ENSMUSG( | ENSMUSG( | ENSMUSG( | 0.293273 | 0.398319 | NO | NO | NA |
| Cxcl11 | ENSMUSG( | ENSMUSG( | 0.962528 | 0.363811 | NO | NO | NA |
| Olfr556 | ENSMUSG( | ENSMUSG( | 1.061916 | 0.192562 | NO | NO | NA |
| E430002N | ENSMUSG( | ENSMUSG( | -1.14511 | 0.000511 | YES | YES | Downregulated |
| Gm12592 | ENSMUSG( | ENSMUSG( | -0.10427 | 0.798535 | NO | NO | NA |
| Glb1l2 | ENSMUSG( | ENSMUSG( | 0.171542 | 0.881883 | NO | NO | NA |
| Gpx5 | ENSMUSG( | ENSMUSG( | 0.479166 | 0.71346 | NO | NO | NA |
| Kcnmb4os | ENSMUSG( | ENSMUSG( | 0.410598 | 0.330787 | NO | NO | NA |
| Rbp1 | ENSMUSG( | ENSMUSG( | 0.890532 | 0.126934 | NO | NO | NA |
| Psg16 | ENSMUSG( | ENSMUSG( | 0.555801 | 0.487541 | NO | NO | NA |
| Dlx3 | ENSMUSG( | ENSMUSG( | 0.68666 | 0.57298 | NO | NO | NA |
| Bdnf | ENSMUSG( | ENSMUSG( | 0.889248 | 0.370733 | NO | NO | NA |
| Gm10240 | ENSMUSG( | ENSMUSG( | -20.7778 | NA | NO | NO | NA |
| ENSMUSG | ENSMUSG( | ENSMUSG( | 22.03897 | NA | NO | NO | NA |
| Gm21011 | ENSMUSG( | ENSMUSG( | 0.117428 | 0.811231 | NO | NO | NA |
| Gm43063 | ENSMUSG( | ENSMUSG( | 1.011983 | 0.298484 | NO | NO | NA |
| Emx2os | ENSMUSG( | ENSMUSG( | 0.738856 | 0.467168 | NO | NO | NA |
| Padi3 | ENSMUSG( | ENSMUSG( | 0.453377 | 0.087918 | NO | NO | NA |
| Plpp1 | ENSMUSG( | ENSMUSG( | 1.070237 | 0.05409 | NO | NO | NA |
| Gm32261 | ENSMUSG( | ENSMUSG( | 0.905809 | 0.217816 | NO | NO | NA |
| Slc44a5 | ENSMUSG( | ENSMUSG( | 0.91074 | 0.393025 | NO | NO | NA |
| Cd5l | ENSMUSG( | ENSMUSG( | -1.2277 | 0.013193 | YES | YES | Downregulated |
| Gm45577 | ENSMUSG( | ENSMUSG( | 0.465751 | 0.667955 | NO | NO | NA |
| 4930594M | ENSMUSG( | ENSMUSG( | -0.41645 | 0.256641 | NO | NO | NA |

| | | | | | | | |
|---|---|---|---|---|---|---|---|
| Gm9403 | ENSMUSG( | ENSMUSG( | -0.95838 | 0.001883 | YES | NO | NA |
| ENSMUSG | ENSMUSG( | ENSMUSG( | 0.333464 | 0.496838 | NO | NO | NA |
| Gm27209 | ENSMUSG( | ENSMUSG( | -1.24279 | 0.005487 | YES | YES | Downregulated |
| Gm49932 | ENSMUSG( | ENSMUSG( | 1.036937 | 0.298188 | NO | NO | NA |
| Fam71e1 | ENSMUSG( | ENSMUSG( | 0.559723 | 0.064878 | NO | NO | NA |
| Slc16a14 | ENSMUSG( | ENSMUSG( | 1.065271 | 0.330198 | NO | NO | NA |
| 4931439C: | ENSMUSG( | ENSMUSG( | 0.487787 | 0.647539 | NO | NO | NA |
| 1700030N | ENSMUSG( | ENSMUSG( | 0.665025 | 0.567007 | NO | NO | NA |
| ENSMUSG( | ENSMUSG( | ENSMUSG( | -0.03352 | 0.950039 | NO | NO | NA |
| Prkcg | ENSMUSG( | ENSMUSG( | -0.10084 | 0.756401 | NO | NO | NA |
| Kcnk3 | ENSMUSG( | ENSMUSG( | 0.882858 | 0.114465 | NO | NO | NA |
| Pate13 | ENSMUSG( | ENSMUSG( | 0.966802 | 0.394041 | NO | NO | NA |
| Gm43658 | ENSMUSG( | ENSMUSG( | -0.52336 | 0.096376 | NO | NO | NA |
| D930007J( | ENSMUSG( | ENSMUSG( | -0.23187 | 0.63616 | NO | NO | NA |
| Gm49260 | ENSMUSG( | ENSMUSG( | 0.639034 | 0.534067 | NO | NO | NA |
| Gm44891 | ENSMUSG( | ENSMUSG( | 0.631068 | 0.57964 | NO | NO | NA |
| Slc16a4 | ENSMUSG( | ENSMUSG( | -0.45259 | 0.255552 | NO | NO | NA |
| H4c11 | ENSMUSG( | ENSMUSG( | 0.149408 | 0.693448 | NO | NO | NA |
| 1700029J( | ENSMUSG( | ENSMUSG( | 0.492342 | 0.087639 | NO | NO | NA |
| Gm5620 | ENSMUSG( | ENSMUSG( | 0.405445 | 0.500004 | NO | NO | NA |
| Tnk2os | ENSMUSG( | ENSMUSG( | -0.15913 | 0.680822 | NO | NO | NA |
| Gm7331 | ENSMUSG( | ENSMUSG( | 0.617467 | 0.076567 | NO | NO | NA |
| Ighv3-6 | ENSMUSG( | ENSMUSG( | -2.15139 | 0.277267 | NO | NO | NA |
| Gm44013 | ENSMUSG( | ENSMUSG( | 0.796303 | 0.048708 | YES | NO | NA |
| Gm45160 | ENSMUSG( | ENSMUSG( | 0.924381 | 0.377668 | NO | NO | NA |
| V1ra8 | ENSMUSG( | ENSMUSG( | 1.005411 | 0.324777 | NO | NO | NA |
| Neurl1a | ENSMUSG( | ENSMUSG( | 0.626112 | 0.531711 | NO | NO | NA |
| Pkp2 | ENSMUSG( | ENSMUSG( | 1.234854 | 0.010924 | YES | YES | Upregulated |
| Olfr943 | ENSMUSG( | ENSMUSG( | 0.90171 | 0.393956 | NO | NO | NA |
| Zfp148-ps: | ENSMUSG( | ENSMUSG( | 0.574563 | 0.59456 | NO | NO | NA |
| ENSMUSG( | ENSMUSG( | ENSMUSG( | 0.187043 | 0.494121 | NO | NO | NA |
| Gm46123 | ENSMUSG( | ENSMUSG( | 0.480377 | 0.659678 | NO | NO | NA |
| Irf6 | ENSMUSG( | ENSMUSG( | -0.39002 | 0.250934 | NO | NO | NA |
| 2810408A | ENSMUSG( | ENSMUSG( | 0.389437 | 0.146854 | NO | NO | NA |
| Gipc2 | ENSMUSG( | ENSMUSG( | 0.210358 | 0.585728 | NO | NO | NA |
| Msantd3 | ENSMUSG( | ENSMUSG( | 0.74701 | 0.413599 | NO | NO | NA |
| Gm38359 | ENSMUSG( | ENSMUSG( | 0.384769 | 0.710991 | NO | NO | NA |
| Hspd1-ps3 | ENSMUSG( | ENSMUSG( | 0.169113 | 0.63758 | NO | NO | NA |
| Gm43692 | ENSMUSG( | ENSMUSG( | -0.31202 | 0.267916 | NO | NO | NA |
| Gm9884 | ENSMUSG( | ENSMUSG( | 1.28952 | 0.15771 | NO | NO | NA |
| Gm46440 | ENSMUSG( | ENSMUSG( | 0.818479 | 0.301687 | NO | NO | NA |
| Gm37073 | ENSMUSG( | ENSMUSG( | 0.873989 | 0.391088 | NO | NO | NA |
| Gata6os | ENSMUSG( | ENSMUSG( | 1.044222 | 0.297828 | NO | NO | NA |
| Gm45631 | ENSMUSG( | ENSMUSG( | -0.035 | 0.933725 | NO | NO | NA |
| Igfbp5 | ENSMUSG( | ENSMUSG( | 0.804168 | 0.483913 | NO | NO | NA |
| Prss41 | ENSMUSG( | ENSMUSG( | -0.23464 | 0.491395 | NO | NO | NA |
| Clic6 | ENSMUSG( | ENSMUSG( | 0.77846 | 0.434006 | NO | NO | NA |

| | | | | | | | |
|---|---|---|---|---|---|---|---|
| Dsg2 | ENSMUSG( | ENSMUSG( | -0.05607 | 0.941129 | NO | NO | NA |
| Cyp2r1 | ENSMUSG( | ENSMUSG( | -1.36218 | 2.03E-05 | YES | YES | Downregulated |
| Olfr1062 | ENSMUSG( | ENSMUSG( | 0.919276 | 0.412068 | NO | NO | NA |
| Cxadr | ENSMUSG( | ENSMUSG( | -0.04578 | 0.906987 | NO | NO | NA |
| Gm38366 | ENSMUSG( | ENSMUSG( | 0.126141 | 0.770569 | NO | NO | NA |
| Gm5111 | ENSMUSG( | ENSMUSG( | 2.410273 | 2.77E-05 | YES | YES | Upregulated |
| Gm36346 | ENSMUSG( | ENSMUSG( | 0.494469 | 0.661555 | NO | NO | NA |
| Klhl23 | ENSMUSG( | ENSMUSG( | 1.43372 | 0.002069 | YES | YES | Upregulated |
| Ube2e2 | ENSMUSG( | ENSMUSG( | 0.621604 | 0.454601 | NO | NO | NA |
| Gm35963 | ENSMUSG( | ENSMUSG( | 0.615492 | 0.583624 | NO | NO | NA |
| Gm17809 | ENSMUSG( | ENSMUSG( | -0.03581 | 0.924326 | NO | NO | NA |
| Gm15133 | ENSMUSG( | ENSMUSG( | -0.11246 | 0.82465 | NO | NO | NA |
| Pcdhb13 | ENSMUSG( | ENSMUSG( | 0.797397 | 0.434626 | NO | NO | NA |
| ENSMUSG | ENSMUSG( | ENSMUSG( | 0.427894 | 0.23298 | NO | NO | NA |
| Gm18588 | ENSMUSG( | ENSMUSG( | 0.065772 | 0.87252 | NO | NO | NA |
| Rmdn1 | ENSMUSG( | ENSMUSG( | 0.113373 | 0.778769 | NO | NO | NA |
| Gm37420 | ENSMUSG( | ENSMUSG( | -0.41669 | 0.316122 | NO | NO | NA |
| Gm48065 | ENSMUSG( | ENSMUSG( | 0.823719 | 0.422568 | NO | NO | NA |
| Cubn | ENSMUSG( | ENSMUSG( | 2.275333 | 0.072887 | NO | NO | NA |
| Cfap45 | ENSMUSG( | ENSMUSG( | 0.697656 | 0.209768 | NO | NO | NA |
| Olfr338 | ENSMUSG( | ENSMUSG( | 0.504884 | 0.560369 | NO | NO | NA |
| Gm42759 | ENSMUSG( | ENSMUSG( | 1.103532 | 0.143128 | NO | NO | NA |
| 1810024B( | ENSMUSG( | ENSMUSG( | 0.85718 | 0.025699 | YES | NO | NA |
| 4930458D | ENSMUSG( | ENSMUSG( | -0.22791 | 0.783097 | NO | NO | NA |
| Rnu2-10 | ENSMUSG( | ENSMUSG( | -0.1447 | 0.95056 | NO | NO | NA |
| Gm30371 | ENSMUSG( | ENSMUSG( | 0.626498 | 0.614864 | NO | NO | NA |
| Rps27a-ps | ENSMUSG( | ENSMUSG( | 0.137627 | 0.739353 | NO | NO | NA |
| Ywhaq-ps3 | ENSMUSG( | ENSMUSG( | 0.001278 | 0.998635 | NO | NO | NA |
| Gdpd3 | ENSMUSG( | ENSMUSG( | 0.168266 | 0.616241 | NO | NO | NA |
| Gm43112 | ENSMUSG( | ENSMUSG( | 0.567809 | 0.587391 | NO | NO | NA |
| Gm49897 | ENSMUSG( | ENSMUSG( | 0.866597 | 0.428173 | NO | NO | NA |
| Gm45399 | ENSMUSG( | ENSMUSG( | 0.233758 | 0.504016 | NO | NO | NA |
| Gm43312 | ENSMUSG( | ENSMUSG( | 0.020429 | 0.97125 | NO | NO | NA |
| Gm39244 | ENSMUSG( | ENSMUSG( | 1.006926 | 0.318088 | NO | NO | NA |
| E030042O | ENSMUSG( | ENSMUSG( | 0.107521 | 0.700801 | NO | NO | NA |
| Gm37859 | ENSMUSG( | ENSMUSG( | 0.896869 | 0.352563 | NO | NO | NA |
| Gm43106 | ENSMUSG( | ENSMUSG( | -0.97224 | 0.001036 | YES | NO | NA |
| Gm36993 | ENSMUSG( | ENSMUSG( | 0.896437 | 0.384021 | NO | NO | NA |
| Gm47615 | ENSMUSG( | ENSMUSG( | 0.727979 | 0.439353 | NO | NO | NA |
| Gm19178 | ENSMUSG( | ENSMUSG( | -0.58827 | 0.019978 | YES | NO | NA |
| 4931413K: | ENSMUSG( | ENSMUSG( | -0.12457 | 0.804695 | NO | NO | NA |
| Gm44789 | ENSMUSG( | ENSMUSG( | 1.058387 | 0.002487 | YES | YES | Upregulated |
| ENSMUSG | ENSMUSG( | ENSMUSG( | 1.232887 | 0.009564 | YES | YES | Upregulated |
| ENSMUSG( | ENSMUSG( | ENSMUSG( | 0.540688 | 0.604389 | NO | NO | NA |
| Gm37307 | ENSMUSG( | ENSMUSG( | 0.560769 | 0.585715 | NO | NO | NA |
| ENSMUSG( | ENSMUSG( | ENSMUSG( | 0.658622 | 0.354579 | NO | NO | NA |
| Gm44848 | ENSMUSG( | ENSMUSG( | 0.206446 | 0.808208 | NO | NO | NA |

| | | | | | | | |
|---|---|---|---|---|---|---|---|
| 2600014E2 | ENSMUSG( | ENSMUSG( | 0.645102 | 0.539797 | NO | NO | NA |
| Epha4 | ENSMUSG( | ENSMUSG( | 0.711341 | 0.555257 | NO | NO | NA |
| Gm19265 | ENSMUSG( | ENSMUSG( | -0.58733 | 0.099987 | NO | NO | NA |
| Prox1 | ENSMUSG( | ENSMUSG( | 0.287918 | 0.787135 | NO | NO | NA |
| Gm11747 | ENSMUSG( | ENSMUSG( | 0.733503 | 0.493946 | NO | NO | NA |
| Pcdhgb7 | ENSMUSG( | ENSMUSG( | -0.55862 | 0.233451 | NO | NO | NA |
| Gm43756 | ENSMUSG( | ENSMUSG( | 0.443059 | 0.625461 | NO | NO | NA |
| Rab30 | ENSMUSG( | ENSMUSG( | 0.472557 | 0.57298 | NO | NO | NA |
| Khdrbs2 | ENSMUSG( | ENSMUSG( | 0.512796 | 0.627247 | NO | NO | NA |
| Gm10190 | ENSMUSG( | ENSMUSG( | 0.768015 | 0.33645 | NO | NO | NA |
| 1700028K( | ENSMUSG( | ENSMUSG( | -0.37215 | 0.815297 | NO | NO | NA |
| Gm36991 | ENSMUSG( | ENSMUSG( | 0.865441 | 0.392528 | NO | NO | NA |
| Gm47073 | ENSMUSG( | ENSMUSG( | 0.496915 | 0.606874 | NO | NO | NA |
| Gm15602 | ENSMUSG( | ENSMUSG( | 0.203266 | 0.653348 | NO | NO | NA |
| Gm19202 | ENSMUSG( | ENSMUSG( | 0.785017 | 0.426562 | NO | NO | NA |
| Mamdc4 | ENSMUSG( | ENSMUSG( | -0.08879 | 0.797391 | NO | NO | NA |
| Tshz3 | ENSMUSG( | ENSMUSG( | 0.188334 | 0.820924 | NO | NO | NA |
| Slc35g2 | ENSMUSG( | ENSMUSG( | -0.97502 | 0.000688 | YES | NO | NA |
| Gm37234 | ENSMUSG( | ENSMUSG( | 0.831607 | 0.41526 | NO | NO | NA |
| Olfr1124 | ENSMUSG( | ENSMUSG( | 1.1673 | 0.264824 | NO | NO | NA |
| E430021H | ENSMUSG( | ENSMUSG( | -0.05049 | 0.931656 | NO | NO | NA |
| Vmn1r46 | ENSMUSG( | ENSMUSG( | 0.269463 | 0.759918 | NO | NO | NA |
| Gm16796 | ENSMUSG( | ENSMUSG( | -0.1401 | 0.829442 | NO | NO | NA |
| Gm44518 | ENSMUSG( | ENSMUSG( | 0.572841 | 0.613051 | NO | NO | NA |
| 4930544L( | ENSMUSG( | ENSMUSG( | 0.530854 | 0.658027 | NO | NO | NA |
| Col4a1 | ENSMUSG( | ENSMUSG( | 0.583025 | 0.528021 | NO | NO | NA |
| Fndc7 | ENSMUSG( | ENSMUSG( | 0.72734 | 0.173088 | NO | NO | NA |
| 9330188P( | ENSMUSG( | ENSMUSG( | 0.57765 | 0.58481 | NO | NO | NA |
| Gm42981 | ENSMUSG( | ENSMUSG( | 0.574252 | 0.592341 | NO | NO | NA |
| ENSMUSG | ENSMUSG( | ENSMUSG( | 0.492601 | 0.283652 | NO | NO | NA |
| Gm43545 | ENSMUSG( | ENSMUSG( | 0.188716 | 0.808024 | NO | NO | NA |
| Gm31616 | ENSMUSG( | ENSMUSG( | 0.013263 | 0.966013 | NO | NO | NA |
| Gm13091 | ENSMUSG( | ENSMUSG( | -1.30915 | 0.002481 | YES | YES | Downregulated |
| Zfp607b | ENSMUSG( | ENSMUSG( | 0.336335 | 0.371729 | NO | NO | NA |
| Gm44264 | ENSMUSG( | ENSMUSG( | 1.197502 | 0.018074 | YES | YES | Upregulated |
| 9130213A | ENSMUSG( | ENSMUSG( | 0.212495 | 0.813047 | NO | NO | NA |
| Slc7a14 | ENSMUSG( | ENSMUSG( | 0.559688 | 0.554852 | NO | NO | NA |
| Gm22317 | ENSMUSG( | ENSMUSG( | 0.690276 | 0.860844 | NO | NO | NA |
| Gm9027 | ENSMUSG( | ENSMUSG( | -0.08998 | 0.822027 | NO | NO | NA |
| Olfr39 | ENSMUSG( | ENSMUSG( | 0.954166 | 0.350151 | NO | NO | NA |
| Timp4 | ENSMUSG( | ENSMUSG( | 0.651191 | 0.561251 | NO | NO | NA |
| Apc2 | ENSMUSG( | ENSMUSG( | -0.99514 | 0.013211 | YES | NO | NA |
| Ccnd1 | ENSMUSG( | ENSMUSG( | -0.1115 | 0.791049 | NO | NO | NA |
| Ighv1-85 | ENSMUSG( | ENSMUSG( | -4.60533 | 0.079114 | NO | NO | NA |
| Gm37527 | ENSMUSG( | ENSMUSG( | 0.446716 | 0.582357 | NO | NO | NA |
| Gm31749 | ENSMUSG( | ENSMUSG( | -0.0859 | 0.788344 | NO | NO | NA |
| Gm28112 | ENSMUSG( | ENSMUSG( | 0.90146 | 0.224995 | NO | NO | NA |

| | | | | | | | |
|---|---|---|---|---|---|---|---|
| Pip5k1b | ENSMUSG( | ENSMUSG( | 0.686952 | 0.139367 | NO | NO | NA |
| Olfr607 | ENSMUSG( | ENSMUSG( | 0.24267 | 0.825884 | NO | NO | NA |
| ENSMUSG | ENSMUSG( | ENSMUSG( | -0.16462 | 0.638601 | NO | NO | NA |
| Gpr63 | ENSMUSG( | ENSMUSG( | 0.910576 | 0.306811 | NO | NO | NA |
| Flywch2 | ENSMUSG( | ENSMUSG( | 1.44513 | 0.006969 | YES | YES | Upregulated |
| Itgb6 | ENSMUSG( | ENSMUSG( | 1.008854 | 0.306212 | NO | NO | NA |
| Gm15823 | ENSMUSG( | ENSMUSG( | 0.644364 | 0.576072 | NO | NO | NA |
| Gm38082 | ENSMUSG( | ENSMUSG( | 0.482378 | 0.557764 | NO | NO | NA |
| Gm43924 | ENSMUSG( | ENSMUSG( | 1.253595 | 0.115291 | NO | NO | NA |
| Gm44708 | ENSMUSG( | ENSMUSG( | 0.788276 | 0.465617 | NO | NO | NA |
| Adam4 | ENSMUSG( | ENSMUSG( | -0.85938 | 0.017768 | YES | NO | NA |
| Sypl2 | ENSMUSG( | ENSMUSG( | -0.1687 | 0.831994 | NO | NO | NA |
| A430057M | ENSMUSG( | ENSMUSG( | 0.399932 | 0.138256 | NO | NO | NA |
| Sspn | ENSMUSG( | ENSMUSG( | 0.547726 | 0.364915 | NO | NO | NA |
| ENSMUSG | ENSMUSG( | ENSMUSG( | 0.838508 | 0.265987 | NO | NO | NA |
| Gm6525 | ENSMUSG( | ENSMUSG( | 0.996111 | 0.109136 | NO | NO | NA |
| Gm4994 | ENSMUSG( | ENSMUSG( | 0.746084 | 0.502435 | NO | NO | NA |
| Gm15910 | ENSMUSG( | ENSMUSG( | 1.33674 | 0.10941 | NO | NO | NA |
| Kcnh3 | ENSMUSG( | ENSMUSG( | -0.14639 | 0.8114 | NO | NO | NA |
| Clic5 | ENSMUSG( | ENSMUSG( | 0.321096 | 0.389738 | NO | NO | NA |
| Slc4a4 | ENSMUSG( | ENSMUSG( | 1.266801 | 0.070591 | NO | NO | NA |
| Col1a2 | ENSMUSG( | ENSMUSG( | 0.511749 | 0.642471 | NO | NO | NA |
| Gm43011 | ENSMUSG( | ENSMUSG( | -0.19887 | 0.739792 | NO | NO | NA |
| 1700123M | ENSMUSG( | ENSMUSG( | 0.28968 | 0.339868 | NO | NO | NA |
| Ankrd34b | ENSMUSG( | ENSMUSG( | 0.879455 | 0.22471 | NO | NO | NA |
| ENSMUSG( | ENSMUSG( | ENSMUSG( | 0.57361 | 0.595555 | NO | NO | NA |
| Ppp2r2c | ENSMUSG( | ENSMUSG( | -0.71998 | 0.155997 | NO | NO | NA |
| ENSMUSG | ENSMUSG( | ENSMUSG( | 0.344483 | 0.34547 | NO | NO | NA |
| A930033H | ENSMUSG( | ENSMUSG( | 0.639706 | 0.545216 | NO | NO | NA |
| Pi15 | ENSMUSG( | ENSMUSG( | 0.745797 | 0.19619 | NO | NO | NA |
| Gm15470 | ENSMUSG( | ENSMUSG( | -0.53147 | 0.132922 | NO | NO | NA |
| Gm26597 | ENSMUSG( | ENSMUSG( | 0.789176 | 0.494513 | NO | NO | NA |
| C530043A | ENSMUSG( | ENSMUSG( | 0.796229 | 0.440435 | NO | NO | NA |
| Hs3st4 | ENSMUSG( | ENSMUSG( | 0.970814 | 0.342553 | NO | NO | NA |
| Amh | ENSMUSG( | ENSMUSG( | 0.026742 | 0.939775 | NO | NO | NA |
| H4c12 | ENSMUSG( | ENSMUSG( | 0.399442 | 0.275983 | NO | NO | NA |
| Gata5 | ENSMUSG( | ENSMUSG( | 1.325737 | 0.150817 | NO | NO | NA |
| Vsig2 | ENSMUSG( | ENSMUSG( | 2.912795 | 3.08E-05 | YES | YES | Upregulated |
| Gm49413 | ENSMUSG( | ENSMUSG( | 0.375241 | 0.609694 | NO | NO | NA |
| Gm9630 | ENSMUSG( | ENSMUSG( | 1.258531 | 0.20339 | NO | NO | NA |
| Bdh2 | ENSMUSG( | ENSMUSG( | 0.798876 | 0.489365 | NO | NO | NA |
| Ighv5-17 | ENSMUSG( | ENSMUSG( | -1.10082 | 0.584776 | NO | NO | NA |
| Car13 | ENSMUSG( | ENSMUSG( | -0.18014 | 0.661527 | NO | NO | NA |
| Tbxas1 | ENSMUSG( | ENSMUSG( | 0.090379 | 0.889991 | NO | NO | NA |
| Bend5 | ENSMUSG( | ENSMUSG( | 2.329125 | 3.68E-06 | YES | YES | Upregulated |
| Erp27 | ENSMUSG( | ENSMUSG( | 0.463394 | 0.176859 | NO | NO | NA |
| Prdx6-ps2 | ENSMUSG( | ENSMUSG( | 0.784888 | 0.496053 | NO | NO | NA |

| | | | | | | | |
|---|---|---|---|---|---|---|---|
| Gm50403 | ENSMUSG( | ENSMUSG( | 0.221774 | 0.810996 | NO | NO | NA |
| Gm47445 | ENSMUSG( | ENSMUSG( | 0.390143 | 0.713817 | NO | NO | NA |
| Esrrb | ENSMUSG( | ENSMUSG( | 0.863786 | 0.416186 | NO | NO | NA |
| Gm49308 | ENSMUSG( | ENSMUSG( | 0.155755 | 0.868157 | NO | NO | NA |
| Gm48048 | ENSMUSG( | ENSMUSG( | -1.65538 | 0.417592 | NO | NO | NA |
| 9330154J( | ENSMUSG( | ENSMUSG( | 0.64144 | 0.516916 | NO | NO | NA |
| Gm4356 | ENSMUSG( | ENSMUSG( | -0.18453 | 0.680824 | NO | NO | NA |
| ENSMUSG | ENSMUSG( | ENSMUSG( | 0.471249 | 0.451408 | NO | NO | NA |
| Vmn1r49 | ENSMUSG( | ENSMUSG( | 0.662992 | 0.545057 | NO | NO | NA |
| Gm15963 | ENSMUSG( | ENSMUSG( | 0.839207 | 0.423238 | NO | NO | NA |
| Poln | ENSMUSG( | ENSMUSG( | 1.271379 | 0.002635 | YES | YES | Upregulated |
| Adamts19 | ENSMUSG( | ENSMUSG( | 0.855879 | 0.44958 | NO | NO | NA |
| Lama4 | ENSMUSG( | ENSMUSG( | 0.947236 | 0.374884 | NO | NO | NA |
| Adcy1 | ENSMUSG( | ENSMUSG( | 0.200533 | 0.702923 | NO | NO | NA |
| 4930507D | ENSMUSG( | ENSMUSG( | 0.785212 | 0.003238 | YES | NO | NA |
| Gm38374 | ENSMUSG( | ENSMUSG( | 0.712848 | 0.392392 | NO | NO | NA |
| Mkln1os | ENSMUSG( | ENSMUSG( | 0.645687 | 0.340117 | NO | NO | NA |
| Pkd1l2 | ENSMUSG( | ENSMUSG( | 1.056329 | 0.009848 | YES | YES | Upregulated |
| Gm26826 | ENSMUSG( | ENSMUSG( | 0.650936 | 0.434333 | NO | NO | NA |
| Hoxd3os1 | ENSMUSG( | ENSMUSG( | 0.75759 | 0.475251 | NO | NO | NA |
| D7Ertd443 | ENSMUSG( | ENSMUSG( | 0.440645 | 0.698039 | NO | NO | NA |
| Gm11952 | ENSMUSG( | ENSMUSG( | 1.126123 | 0.235628 | NO | NO | NA |
| Slc22a19 | ENSMUSG( | ENSMUSG( | 0.691764 | 0.486542 | NO | NO | NA |
| Gm2451 | ENSMUSG( | ENSMUSG( | 0.794844 | 0.154014 | NO | NO | NA |
| Gm17767 | ENSMUSG( | ENSMUSG( | 0.479362 | 0.503551 | NO | NO | NA |
| Trf | ENSMUSG( | ENSMUSG( | -0.48699 | 0.335244 | NO | NO | NA |
| Gm43274 | ENSMUSG( | ENSMUSG( | 0.172903 | 0.821271 | NO | NO | NA |
| Gm11264 | ENSMUSG( | ENSMUSG( | 0.597173 | 0.592479 | NO | NO | NA |
| Gm48363 | ENSMUSG( | ENSMUSG( | 0.387222 | 0.73647 | NO | NO | NA |
| Gm47705 | ENSMUSG( | ENSMUSG( | -0.01593 | 0.963738 | NO | NO | NA |
| Uox | ENSMUSG( | ENSMUSG( | 0.89308 | 0.232837 | NO | NO | NA |
| Gm13684 | ENSMUSG( | ENSMUSG( | 0.105555 | 0.728803 | NO | NO | NA |
| Jph4 | ENSMUSG( | ENSMUSG( | 0.284606 | 0.796249 | NO | NO | NA |
| Zfyve9 | ENSMUSG( | ENSMUSG( | 0.717018 | 0.029816 | YES | NO | NA |
| Pcp4l1 | ENSMUSG( | ENSMUSG( | 0.337425 | 0.765058 | NO | NO | NA |
| Gm5487 | ENSMUSG( | ENSMUSG( | 0.603482 | 0.065232 | NO | NO | NA |
| Gm26555 | ENSMUSG( | ENSMUSG( | 0.650367 | 0.587391 | NO | NO | NA |
| Morn2 | ENSMUSG( | ENSMUSG( | 0.820958 | 0.075319 | NO | NO | NA |
| Mamdc2 | ENSMUSG( | ENSMUSG( | 3.229348 | 4.29E-07 | YES | YES | Upregulated |
| ENSMUSG | ENSMUSG( | ENSMUSG( | 0.971435 | 0.307552 | NO | NO | NA |
| Trdv1 | ENSMUSG( | ENSMUSG( | -2.29937 | 0.022651 | YES | YES | Downregulated |
| Tub | ENSMUSG( | ENSMUSG( | -0.31891 | 0.351026 | NO | NO | NA |
| C730034F( | ENSMUSG( | ENSMUSG( | -0.53996 | 0.073828 | NO | NO | NA |
| Gm47608 | ENSMUSG( | ENSMUSG( | 0.298242 | 0.791836 | NO | NO | NA |
| Micall2 | ENSMUSG( | ENSMUSG( | 0.452632 | 0.580811 | NO | NO | NA |
| Gm47428 | ENSMUSG( | ENSMUSG( | 0.720024 | 0.043827 | YES | NO | NA |
| ENSMUSG | ENSMUSG( | ENSMUSG( | 0.135085 | 0.784801 | NO | NO | NA |

| | | | | | | | |
|---|---|---|---|---|---|---|---|
| Tpbg | ENSMUSG( | ENSMUSG( | 0.413592 | 0.614593 | NO | NO | NA |
| Gm43844 | ENSMUSG( | ENSMUSG( | 0.459321 | 0.208913 | NO | NO | NA |
| Gm7882 | ENSMUSG( | ENSMUSG( | 0.694495 | 0.501525 | NO | NO | NA |
| Gm26628 | ENSMUSG( | ENSMUSG( | -0.16119 | 0.699835 | NO | NO | NA |
| ENSMUSG | ENSMUSG( | ENSMUSG( | 0.003816 | 0.991804 | NO | NO | NA |
| Gm11251 | ENSMUSG( | ENSMUSG( | 1.120795 | 0.349528 | NO | NO | NA |
| Sema6d | ENSMUSG( | ENSMUSG( | 0.784462 | 0.085518 | NO | NO | NA |
| Timd4 | ENSMUSG( | ENSMUSG( | 1.153926 | 0.315559 | NO | NO | NA |
| 5730405O | ENSMUSG( | ENSMUSG( | -0.22959 | 0.638201 | NO | NO | NA |
| Pax3 | ENSMUSG( | ENSMUSG( | 0.482114 | 0.457003 | NO | NO | NA |
| Gm49132 | ENSMUSG( | ENSMUSG( | 0.705614 | 0.511078 | NO | NO | NA |
| Fmo1 | ENSMUSG( | ENSMUSG( | -0.08329 | 0.814825 | NO | NO | NA |
| A330084C | ENSMUSG( | ENSMUSG( | 1.047466 | 0.185922 | NO | NO | NA |
| ENSMUSG | ENSMUSG( | ENSMUSG( | -0.23184 | 0.579976 | NO | NO | NA |
| Gm38036 | ENSMUSG( | ENSMUSG( | 0.56698 | 0.60567 | NO | NO | NA |
| Rimklb | ENSMUSG( | ENSMUSG( | 0.613106 | 0.571509 | NO | NO | NA |
| Rps3a3 | ENSMUSG( | ENSMUSG( | 0.910559 | 0.015016 | YES | NO | NA |
| Gm42507 | ENSMUSG( | ENSMUSG( | 0.50216 | 0.492717 | NO | NO | NA |
| Htra3 | ENSMUSG( | ENSMUSG( | -0.843 | 0.020041 | YES | NO | NA |
| Lrrc14b | ENSMUSG( | ENSMUSG( | 1.227887 | 0.330198 | NO | NO | NA |
| Prrt2 | ENSMUSG( | ENSMUSG( | -1.16892 | 0.02305 | YES | YES | Downregulated |
| 4930579G | ENSMUSG( | ENSMUSG( | -0.14681 | 0.778681 | NO | NO | NA |
| Ttc8 | ENSMUSG( | ENSMUSG( | 0.676017 | 0.142842 | NO | NO | NA |
| Igkv4-63 | ENSMUSG( | ENSMUSG( | 5.308452 | NA | NO | NO | NA |
| Gm42067 | ENSMUSG( | ENSMUSG( | 1.004861 | 0.022132 | YES | YES | Upregulated |
| ENSMUSG | ENSMUSG( | ENSMUSG( | -0.51374 | 0.099161 | NO | NO | NA |
| Rab20 | ENSMUSG( | ENSMUSG( | -0.29632 | 0.466276 | NO | NO | NA |
| Gm42636 | ENSMUSG( | ENSMUSG( | 0.181749 | 0.695793 | NO | NO | NA |
| Fam135a | ENSMUSG( | ENSMUSG( | 0.318147 | 0.55572 | NO | NO | NA |
| Atp6v0e2 | ENSMUSG( | ENSMUSG( | 0.781028 | 0.377352 | NO | NO | NA |
| Hipk4 | ENSMUSG( | ENSMUSG( | 0.395404 | 0.216209 | NO | NO | NA |
| Mxra8 | ENSMUSG( | ENSMUSG( | -0.05463 | 0.910431 | NO | NO | NA |
| Aph1b | ENSMUSG( | ENSMUSG( | 0.042711 | 0.887324 | NO | NO | NA |
| Mtss2 | ENSMUSG( | ENSMUSG( | -0.27356 | 0.658117 | NO | NO | NA |
| B230369F: | ENSMUSG( | ENSMUSG( | 0.221972 | 0.661555 | NO | NO | NA |
| Rps15a-ps | ENSMUSG( | ENSMUSG( | 0.340681 | 0.328598 | NO | NO | NA |
| ENSMUSG( | ENSMUSG( | ENSMUSG( | 0.368601 | 0.422568 | NO | NO | NA |
| Cd209g | ENSMUSG( | ENSMUSG( | 0.223243 | 0.818021 | NO | NO | NA |
| ENSMUSG | ENSMUSG( | ENSMUSG( | -0.04712 | 0.892196 | NO | NO | NA |
| Olfr1366 | ENSMUSG( | ENSMUSG( | 0.832779 | 0.423328 | NO | NO | NA |
| Alpk3 | ENSMUSG( | ENSMUSG( | 0.137671 | 0.82693 | NO | NO | NA |
| Kctd4 | ENSMUSG( | ENSMUSG( | 0.435696 | 0.549432 | NO | NO | NA |
| Gm37648 | ENSMUSG( | ENSMUSG( | 0.763545 | 0.364886 | NO | NO | NA |
| Rhobtb3 | ENSMUSG( | ENSMUSG( | 0.408257 | 0.661511 | NO | NO | NA |
| Gm15638 | ENSMUSG( | ENSMUSG( | 0.360308 | 0.332236 | NO | NO | NA |
| Gm31812 | ENSMUSG( | ENSMUSG( | -0.5431 | 0.057464 | NO | NO | NA |
| ENSMUSG | ENSMUSG( | ENSMUSG( | -0.44373 | 0.33387 | NO | NO | NA |

| | | | | | | | |
|---|---|---|---|---|---|---|---|
| Gm17122 | ENSMUSG( | ENSMUSG( | -0.27861 | 0.520051 | NO | NO | NA |
| Mafa | ENSMUSG( | ENSMUSG( | -0.22506 | 0.530804 | NO | NO | NA |
| Gm10578 | ENSMUSG( | ENSMUSG( | 0.860232 | 0.501066 | NO | NO | NA |
| Nim1k | ENSMUSG( | ENSMUSG( | 0.346727 | 0.630462 | NO | NO | NA |
| H4c14 | ENSMUSG( | ENSMUSG( | 0.468945 | 0.270824 | NO | NO | NA |
| Gm46565 | ENSMUSG( | ENSMUSG( | 0.146726 | 0.881357 | NO | NO | NA |
| Gm47204 | ENSMUSG( | ENSMUSG( | -0.27543 | 0.50996 | NO | NO | NA |
| Gm16072 | ENSMUSG( | ENSMUSG( | 0.305177 | 0.756654 | NO | NO | NA |
| ENSMUSG | ENSMUSG( | ENSMUSG( | 0.691851 | 0.060375 | NO | NO | NA |
| Caskin1 | ENSMUSG( | ENSMUSG( | 0.279861 | 0.583985 | NO | NO | NA |
| Tgfb2 | ENSMUSG( | ENSMUSG( | 0.566741 | 0.502908 | NO | NO | NA |
| 4933416M | ENSMUSG( | ENSMUSG( | 0.318697 | 0.631599 | NO | NO | NA |
| Lhfpl2 | ENSMUSG( | ENSMUSG( | 1.20521 | 2.00E-05 | YES | YES | Upregulated |
| Gm6092 | ENSMUSG( | ENSMUSG( | 0.432399 | 0.690197 | NO | NO | NA |
| Ighv6-3 | ENSMUSG( | ENSMUSG( | -1.7999 | 0.178411 | NO | NO | NA |
| Gm16575 | ENSMUSG( | ENSMUSG( | -0.16699 | 0.710384 | NO | NO | NA |
| Gm50205 | ENSMUSG( | ENSMUSG( | 0.547983 | 0.202823 | NO | NO | NA |
| ENSMUSG | ENSMUSG( | ENSMUSG( | 0.602671 | 0.326107 | NO | NO | NA |
| Stamos | ENSMUSG( | ENSMUSG( | 0.842069 | 0.175573 | NO | NO | NA |
| Nnat | ENSMUSG( | ENSMUSG( | -0.53059 | 0.165406 | NO | NO | NA |
| Gm9013 | ENSMUSG( | ENSMUSG( | 0.316661 | 0.747006 | NO | NO | NA |
| Tmem231 | ENSMUSG( | ENSMUSG( | 0.606656 | 0.071111 | NO | NO | NA |
| Dab2 | ENSMUSG( | ENSMUSG( | 0.433951 | 0.633815 | NO | NO | NA |
| Gm42303 | ENSMUSG( | ENSMUSG( | 0.62847 | 0.572778 | NO | NO | NA |
| Gm48427 | ENSMUSG( | ENSMUSG( | 1.235777 | 0.254949 | NO | NO | NA |
| Gm49246 | ENSMUSG( | ENSMUSG( | 0.933126 | 0.455874 | NO | NO | NA |
| 1700086P( | ENSMUSG( | ENSMUSG( | 0.909474 | 0.326763 | NO | NO | NA |
| Gm14636 | ENSMUSG( | ENSMUSG( | 0.97368 | 0.00153 | YES | NO | NA |
| Gp6 | ENSMUSG( | ENSMUSG( | 0.497967 | 0.645996 | NO | NO | NA |
| Rims4 | ENSMUSG( | ENSMUSG( | 0.623533 | 0.51556 | NO | NO | NA |
| Slc4a3 | ENSMUSG( | ENSMUSG( | 0.709262 | 0.408588 | NO | NO | NA |
| Fgf14 | ENSMUSG( | ENSMUSG( | 0.83259 | 0.463126 | NO | NO | NA |
| Zfp697 | ENSMUSG( | ENSMUSG( | 0.290858 | 0.784462 | NO | NO | NA |
| A4galt | ENSMUSG( | ENSMUSG( | 0.317283 | 0.815645 | NO | NO | NA |
| Gm15743 | ENSMUSG( | ENSMUSG( | 1.374194 | NA | NO | NO | NA |
| Gm17108 | ENSMUSG( | ENSMUSG( | 0.807725 | 0.050492 | NO | NO | NA |
| Gm10032 | ENSMUSG( | ENSMUSG( | 0.606156 | 0.560715 | NO | NO | NA |
| ENSMUSG | ENSMUSG( | ENSMUSG( | 0.286698 | 0.585023 | NO | NO | NA |
| Gm48631 | ENSMUSG( | ENSMUSG( | 0.992387 | 0.389693 | NO | NO | NA |
| Cpa6 | ENSMUSG( | ENSMUSG( | 0.39541 | 0.725258 | NO | NO | NA |
| Piwil2 | ENSMUSG( | ENSMUSG( | 1.276312 | 0.005241 | YES | YES | Upregulated |
| Gm47896 | ENSMUSG( | ENSMUSG( | 1.317491 | 0.221021 | NO | NO | NA |
| ENSMUSG( | ENSMUSG( | ENSMUSG( | 0.086774 | 0.939243 | NO | NO | NA |
| Dio2 | ENSMUSG( | ENSMUSG( | 0.936717 | 0.021449 | YES | NO | NA |
| Gm43174 | ENSMUSG( | ENSMUSG( | 0.714905 | 0.522753 | NO | NO | NA |
| Gm37884 | ENSMUSG( | ENSMUSG( | 1.038441 | 0.406433 | NO | NO | NA |
| Gm15663 | ENSMUSG( | ENSMUSG( | 0.939884 | 0.384056 | NO | NO | NA |

| | | | | | | | |
|---|---|---|---|---|---|---|---|
| Ptprf | ENSMUSG( | ENSMUSG( | 0.620614 | 0.58825 | NO | NO | NA |
| Gm45354 | ENSMUSG( | ENSMUSG( | 0.13784 | 0.851755 | NO | NO | NA |
| Pld4 | ENSMUSG( | ENSMUSG( | -0.83154 | 0.034211 | YES | NO | NA |
| Tmem121 | ENSMUSG( | ENSMUSG( | 0.70092 | 0.185991 | NO | NO | NA |
| Gm15775 | ENSMUSG( | ENSMUSG( | -0.58323 | 0.038729 | YES | NO | NA |
| Gm37583 | ENSMUSG( | ENSMUSG( | 0.419053 | 0.629421 | NO | NO | NA |
| Zfp111 | ENSMUSG( | ENSMUSG( | 0.799718 | 0.304683 | NO | NO | NA |
| G430095P | ENSMUSG( | ENSMUSG( | -0.25183 | 0.493791 | NO | NO | NA |
| Krt222 | ENSMUSG( | ENSMUSG( | 0.610852 | 0.141098 | NO | NO | NA |
| Gm45351 | ENSMUSG( | ENSMUSG( | -0.48403 | 0.561782 | NO | NO | NA |
| Nat14 | ENSMUSG( | ENSMUSG( | -0.51917 | 0.126746 | NO | NO | NA |
| ENSMUSG( | ENSMUSG( | ENSMUSG( | 0.851379 | 0.010505 | YES | NO | NA |
| Gm4806 | ENSMUSG( | ENSMUSG( | 0.617108 | 0.556172 | NO | NO | NA |
| Upk3b | ENSMUSG( | ENSMUSG( | 1.035116 | 0.305266 | NO | NO | NA |
| Gm48582 | ENSMUSG( | ENSMUSG( | 0.846386 | 0.454019 | NO | NO | NA |
| Gm5900 | ENSMUSG( | ENSMUSG( | 2.420378 | 1.36E-10 | YES | YES | Upregulated |
| Shisa4 | ENSMUSG( | ENSMUSG( | 0.206623 | 0.841312 | NO | NO | NA |
| Gm48303 | ENSMUSG( | ENSMUSG( | 0.75074 | 0.471735 | NO | NO | NA |
| Gm10000 | ENSMUSG( | ENSMUSG( | 1.005975 | 0.328658 | NO | NO | NA |
| Hoxaas3 | ENSMUSG( | ENSMUSG( | 0.65252 | 0.34547 | NO | NO | NA |
| Gm29773 | ENSMUSG( | ENSMUSG( | 0.496707 | 0.397116 | NO | NO | NA |
| 2810032G | ENSMUSG( | ENSMUSG( | 0.815875 | 0.41347 | NO | NO | NA |
| Gm14335 | ENSMUSG( | ENSMUSG( | 0.696427 | 0.361782 | NO | NO | NA |
| Col19a1 | ENSMUSG( | ENSMUSG( | 0.617529 | 0.544688 | NO | NO | NA |
| Amy2a2 | ENSMUSG( | ENSMUSG( | -0.86662 | 0.837594 | NO | NO | NA |
| Gm42467 | ENSMUSG( | ENSMUSG( | 0.127899 | 0.729948 | NO | NO | NA |
| Gm43389 | ENSMUSG( | ENSMUSG( | 0.843938 | 0.09796 | NO | NO | NA |
| Slc5a7 | ENSMUSG( | ENSMUSG( | 0.562101 | 0.626174 | NO | NO | NA |
| Gm31583 | ENSMUSG( | ENSMUSG( | 0.899366 | 0.377772 | NO | NO | NA |
| Tmprss11f | ENSMUSG( | ENSMUSG( | 1.514774 | 0.170189 | NO | NO | NA |
| ENSMUSG( | ENSMUSG( | ENSMUSG( | -0.35609 | 0.424857 | NO | NO | NA |
| Cadm2 | ENSMUSG( | ENSMUSG( | 0.589568 | 0.563166 | NO | NO | NA |
| Tsku | ENSMUSG( | ENSMUSG( | 0.755847 | 0.373143 | NO | NO | NA |
| Olfr656 | ENSMUSG( | ENSMUSG( | 0.924491 | 0.4634 | NO | NO | NA |
| H2bc15 | ENSMUSG( | ENSMUSG( | 0.238137 | 0.628512 | NO | NO | NA |
| Ppfia3 | ENSMUSG( | ENSMUSG( | 0.538607 | 0.306823 | NO | NO | NA |
| 1700120C: | ENSMUSG( | ENSMUSG( | -0.18258 | 0.584439 | NO | NO | NA |
| Gm45251 | ENSMUSG( | ENSMUSG( | 0.654262 | 0.300467 | NO | NO | NA |
| ENSMUSG | ENSMUSG( | ENSMUSG( | -0.17971 | 0.522804 | NO | NO | NA |
| Gm29113 | ENSMUSG( | ENSMUSG( | 1.096956 | 0.250201 | NO | NO | NA |
| Ccdc189 | ENSMUSG( | ENSMUSG( | -0.2201 | 0.684648 | NO | NO | NA |
| C1qa | ENSMUSG( | ENSMUSG( | -1.06407 | 0.01094 | YES | YES | Downregulated |
| Gm37678 | ENSMUSG( | ENSMUSG( | 0.13464 | 0.851529 | NO | NO | NA |
| ENSMUSG( | ENSMUSG( | ENSMUSG( | 0.021491 | 0.976454 | NO | NO | NA |
| Gm15829 | ENSMUSG( | ENSMUSG( | -0.12797 | 0.77121 | NO | NO | NA |
| 4933413J( | ENSMUSG( | ENSMUSG( | 0.824271 | 0.245716 | NO | NO | NA |
| Gm5184 | ENSMUSG( | ENSMUSG( | 0.595441 | 0.545662 | NO | NO | NA |

| | | | | | | | |
|---|---|---|---|---|---|---|---|
| Gm609 | ENSMUSG( | ENSMUSG( | 0.852156 | 0.25757 | NO | NO | NA |
| 5930430L( | ENSMUSG( | ENSMUSG( | 0.260817 | 0.8114 | NO | NO | NA |
| Vmn1r208 | ENSMUSG( | ENSMUSG( | 0.603224 | 0.664181 | NO | NO | NA |
| Kdm4d | ENSMUSG( | ENSMUSG( | 0.511151 | 0.146903 | NO | NO | NA |
| Gm13312 | ENSMUSG( | ENSMUSG( | 1.173621 | 0.290005 | NO | NO | NA |
| Gm47411 | ENSMUSG( | ENSMUSG( | 1.423653 | 0.166132 | NO | NO | NA |
| Gm16759 | ENSMUSG( | ENSMUSG( | 0.338977 | 0.416713 | NO | NO | NA |
| Gm42686 | ENSMUSG( | ENSMUSG( | -0.5815 | 0.164928 | NO | NO | NA |
| 9330104G | ENSMUSG( | ENSMUSG( | -0.02737 | 0.962355 | NO | NO | NA |
| Gm13394 | ENSMUSG( | ENSMUSG( | -0.04307 | 0.960639 | NO | NO | NA |
| Pemt | ENSMUSG( | ENSMUSG( | 1.869724 | 0.000726 | YES | YES | Upregulated |
| Tmed2b | ENSMUSG( | ENSMUSG( | 0.169509 | 0.672518 | NO | NO | NA |
| Gm11242 | ENSMUSG( | ENSMUSG( | 1.251276 | 0.313754 | NO | NO | NA |
| Fkbp11 | ENSMUSG( | ENSMUSG( | 0.110141 | 0.841823 | NO | NO | NA |
| B9d1 | ENSMUSG( | ENSMUSG( | 1.502982 | 0.005847 | YES | YES | Upregulated |
| Gm536 | ENSMUSG( | ENSMUSG( | 0.800114 | 0.461948 | NO | NO | NA |
| Gm6278 | ENSMUSG( | ENSMUSG( | 0.371033 | 0.481807 | NO | NO | NA |
| Sparc | ENSMUSG( | ENSMUSG( | 0.663813 | 0.477841 | NO | NO | NA |
| C1qc | ENSMUSG( | ENSMUSG( | -1.06647 | 0.002213 | YES | YES | Downregulated |
| Gm6254 | ENSMUSG( | ENSMUSG( | -0.19381 | 0.798212 | NO | NO | NA |
| ENSMUSG | ENSMUSG( | ENSMUSG( | 0.659761 | 0.391249 | NO | NO | NA |
| Antxr1 | ENSMUSG( | ENSMUSG( | 0.188735 | 0.833139 | NO | NO | NA |
| Gm44553 | ENSMUSG( | ENSMUSG( | -0.38251 | 0.291961 | NO | NO | NA |
| Gm47111 | ENSMUSG( | ENSMUSG( | 0.42612 | 0.497111 | NO | NO | NA |
| ENSMUSG | ENSMUSG( | ENSMUSG( | 0.354269 | 0.614406 | NO | NO | NA |
| Gm10961 | ENSMUSG( | ENSMUSG( | 1.065714 | 0.281128 | NO | NO | NA |
| Gm37761 | ENSMUSG( | ENSMUSG( | 1.173359 | 0.25122 | NO | NO | NA |
| Gpt2 | ENSMUSG( | ENSMUSG( | 0.76903 | 0.074688 | NO | NO | NA |
| Gm41506 | ENSMUSG( | ENSMUSG( | 0.743384 | 0.369 | NO | NO | NA |
| Vmn1r28 | ENSMUSG( | ENSMUSG( | 0.560057 | 0.614838 | NO | NO | NA |
| Chrm1 | ENSMUSG( | ENSMUSG( | 0.661124 | 0.435436 | NO | NO | NA |
| Slc12a5 | ENSMUSG( | ENSMUSG( | -0.16507 | 0.725834 | NO | NO | NA |
| Gm34921 | ENSMUSG( | ENSMUSG( | 0.37237 | 0.581127 | NO | NO | NA |
| Gm16565 | ENSMUSG( | ENSMUSG( | -0.1637 | 0.650855 | NO | NO | NA |
| Gm16538 | ENSMUSG( | ENSMUSG( | -0.63812 | 0.05131 | NO | NO | NA |
| Xcr1 | ENSMUSG( | ENSMUSG( | -0.1185 | 0.840018 | NO | NO | NA |
| Ddah1 | ENSMUSG( | ENSMUSG( | 0.450771 | 0.664907 | NO | NO | NA |
| Syne4 | ENSMUSG( | ENSMUSG( | 0.558305 | 0.20912 | NO | NO | NA |
| Igkv3-10 | ENSMUSG( | ENSMUSG( | 1.627991 | 0.571926 | NO | NO | NA |
| Gm47056 | ENSMUSG( | ENSMUSG( | 0.430598 | 0.670985 | NO | NO | NA |
| Gm50470 | ENSMUSG( | ENSMUSG( | 0.315397 | 0.686039 | NO | NO | NA |
| Efcab5 | ENSMUSG( | ENSMUSG( | 0.43564 | 0.240327 | NO | NO | NA |
| Gm15283 | ENSMUSG( | ENSMUSG( | 0.84982 | 0.023799 | YES | NO | NA |
| Cntnap5a | ENSMUSG( | ENSMUSG( | 0.589639 | 0.555485 | NO | NO | NA |
| Gm42970 | ENSMUSG( | ENSMUSG( | 0.623973 | 0.562509 | NO | NO | NA |
| ENSMUSG | ENSMUSG( | ENSMUSG( | 0.283848 | 0.620441 | NO | NO | NA |
| Igkv14-12( | ENSMUSG( | ENSMUSG( | -4.20087 | 0.031111 | YES | YES | Downregulated |

| | | | | | | | |
|---|---|---|---|---|---|---|---|
| Zmiz1os1 | ENSMUSG( | ENSMUSG( | -0.75325 | 0.019562 | YES | NO | NA |
| 2810405F: | ENSMUSG( | ENSMUSG( | 0.232689 | 0.571903 | NO | NO | NA |
| Gm10392 | ENSMUSG( | ENSMUSG( | 0.833022 | 0.184404 | NO | NO | NA |
| Dcx | ENSMUSG( | ENSMUSG( | 0.380607 | 0.691861 | NO | NO | NA |
| Slc17a4 | ENSMUSG( | ENSMUSG( | 0.700223 | 0.567401 | NO | NO | NA |
| Gm35048 | ENSMUSG( | ENSMUSG( | 0.169863 | 0.881723 | NO | NO | NA |
| Gm3200 | ENSMUSG( | ENSMUSG( | 0.392031 | 0.462684 | NO | NO | NA |
| 9630001P: | ENSMUSG( | ENSMUSG( | 1.111761 | 0.33043 | NO | NO | NA |
| Gm20692 | ENSMUSG( | ENSMUSG( | -0.641 | 0.154494 | NO | NO | NA |
| Gm10638 | ENSMUSG( | ENSMUSG( | 0.111663 | 0.817607 | NO | NO | NA |
| 64305620 | ENSMUSG( | ENSMUSG( | 0.2166 | 0.647707 | NO | NO | NA |
| 4933401H | ENSMUSG( | ENSMUSG( | 0.798286 | 0.502435 | NO | NO | NA |
| Ddo | ENSMUSG( | ENSMUSG( | 1.254735 | 0.289256 | NO | NO | NA |
| Gm44271 | ENSMUSG( | ENSMUSG( | 0.145565 | 0.893947 | NO | NO | NA |
| 4732490B: | ENSMUSG( | ENSMUSG( | 0.366838 | 0.683247 | NO | NO | NA |
| Eef1akmt3 | ENSMUSG( | ENSMUSG( | 0.529989 | 0.451447 | NO | NO | NA |
| Gm13285 | ENSMUSG( | ENSMUSG( | 0.630404 | 0.03492 | YES | NO | NA |
| Tns3 | ENSMUSG( | ENSMUSG( | -0.38506 | 0.377957 | NO | NO | NA |
| Lca5l | ENSMUSG( | ENSMUSG( | 0.459121 | 0.490577 | NO | NO | NA |
| Ramp1 | ENSMUSG( | ENSMUSG( | 0.835874 | 0.061006 | NO | NO | NA |
| Gm10408 | ENSMUSG( | ENSMUSG( | 1.218355 | 0.244411 | NO | NO | NA |
| Gm43166 | ENSMUSG( | ENSMUSG( | 0.809645 | 0.499614 | NO | NO | NA |
| 4921514A | ENSMUSG( | ENSMUSG( | 0.145856 | 0.747848 | NO | NO | NA |
| Vmn2r84 | ENSMUSG( | ENSMUSG( | 0.391991 | 0.40369 | NO | NO | NA |
| Npff | ENSMUSG( | ENSMUSG( | 0.237496 | 0.471513 | NO | NO | NA |
| Gpr52 | ENSMUSG( | ENSMUSG( | -0.14884 | 0.741426 | NO | NO | NA |
| Cfap54 | ENSMUSG( | ENSMUSG( | 0.440511 | 0.135549 | NO | NO | NA |
| Acrv1 | ENSMUSG( | ENSMUSG( | 0.832575 | 0.445095 | NO | NO | NA |
| Tigd5 | ENSMUSG( | ENSMUSG( | 0.216585 | 0.530632 | NO | NO | NA |
| Adamtsl3 | ENSMUSG( | ENSMUSG( | 0.945105 | 0.384021 | NO | NO | NA |
| Yap1 | ENSMUSG( | ENSMUSG( | 0.743989 | 0.079237 | NO | NO | NA |
| Wdfy3 | ENSMUSG( | ENSMUSG( | 0.142518 | 0.714548 | NO | NO | NA |
| Camk1 | ENSMUSG( | ENSMUSG( | 0.125284 | 0.80634 | NO | NO | NA |
| Gm18828 | ENSMUSG( | ENSMUSG( | -0.11609 | 0.736727 | NO | NO | NA |
| Gm44441 | ENSMUSG( | ENSMUSG( | 0.89049 | 0.430963 | NO | NO | NA |
| Olfml2a | ENSMUSG( | ENSMUSG( | 1.31895 | 0.013274 | YES | YES | Upregulated |
| ENSMUSG( | ENSMUSG( | ENSMUSG( | 1.203895 | 0.625888 | NO | NO | NA |
| Tbx3os1 | ENSMUSG( | ENSMUSG( | 0.577017 | 0.566439 | NO | NO | NA |
| Mageb16- | ENSMUSG( | ENSMUSG( | 1.336855 | 0.232063 | NO | NO | NA |
| Tomm6os | ENSMUSG( | ENSMUSG( | 0.327461 | 0.330408 | NO | NO | NA |
| Scn9a | ENSMUSG( | ENSMUSG( | 0.57031 | 0.569096 | NO | NO | NA |
| Trav10d | ENSMUSG( | ENSMUSG( | 1.406882 | 0.04915 | YES | YES | Upregulated |
| Ighv5-6 | ENSMUSG( | ENSMUSG( | -3.03467 | NA | NO | NO | NA |
| Gm37154 | ENSMUSG( | ENSMUSG( | -0.1344 | 0.747282 | NO | NO | NA |
| Slc25a21 | ENSMUSG( | ENSMUSG( | 0.758757 | 0.247729 | NO | NO | NA |
| ENSMUSG | ENSMUSG( | ENSMUSG( | 0.158676 | 0.806261 | NO | NO | NA |
| Semp2l2a | ENSMUSG( | ENSMUSG( | 0.137449 | 0.890158 | NO | NO | NA |

| | | | | | | | |
|---|---|---|---|---|---|---|---|
| Dhrs9 | ENSMUSG( | ENSMUSG( | 0.577995 | 0.325929 | NO | NO | NA |
| Gm9621 | ENSMUSG( | ENSMUSG( | 0.287118 | 0.394559 | NO | NO | NA |
| Gm26879 | ENSMUSG( | ENSMUSG( | 0.399939 | 0.343744 | NO | NO | NA |
| Gm37105 | ENSMUSG( | ENSMUSG( | -0.13262 | 0.755934 | NO | NO | NA |
| Gm45053 | ENSMUSG( | ENSMUSG( | 0.613026 | 0.12879 | NO | NO | NA |
| Serf1 | ENSMUSG( | ENSMUSG( | 0.366967 | 0.330612 | NO | NO | NA |
| Gm15787 | ENSMUSG( | ENSMUSG( | -0.22042 | 0.569784 | NO | NO | NA |
| Gm48382 | ENSMUSG( | ENSMUSG( | 0.407203 | 0.457405 | NO | NO | NA |
| Angptl6 | ENSMUSG( | ENSMUSG( | 0.265694 | 0.491395 | NO | NO | NA |
| Gm9396 | ENSMUSG( | ENSMUSG( | 0.382939 | 0.503872 | NO | NO | NA |
| Gm36936 | ENSMUSG( | ENSMUSG( | -0.25265 | 0.626786 | NO | NO | NA |
| Slc7a7 | ENSMUSG( | ENSMUSG( | 0.121541 | 0.855452 | NO | NO | NA |
| Reep6 | ENSMUSG( | ENSMUSG( | 0.663168 | 0.173128 | NO | NO | NA |
| Slc7a3 | ENSMUSG( | ENSMUSG( | 1.004531 | 0.030061 | YES | YES | Upregulated |
| Gm44070 | ENSMUSG( | ENSMUSG( | -0.18475 | 0.611216 | NO | NO | NA |
| Gm47138 | ENSMUSG( | ENSMUSG( | 0.423856 | 0.539424 | NO | NO | NA |
| Gm37940 | ENSMUSG( | ENSMUSG( | 0.099607 | 0.868434 | NO | NO | NA |
| Gm20696 | ENSMUSG( | ENSMUSG( | 0.521665 | 0.232423 | NO | NO | NA |
| Gnasas1 | ENSMUSG( | ENSMUSG( | -0.16218 | 0.671315 | NO | NO | NA |
| Gm17590 | ENSMUSG( | ENSMUSG( | -0.08828 | 0.863183 | NO | NO | NA |
| Anxa11os | ENSMUSG( | ENSMUSG( | -0.25543 | 0.548174 | NO | NO | NA |
| Adgrg6 | ENSMUSG( | ENSMUSG( | 0.948608 | 0.309752 | NO | NO | NA |
| Gm37914 | ENSMUSG( | ENSMUSG( | 0.422179 | 0.594479 | NO | NO | NA |
| 4933430H | ENSMUSG( | ENSMUSG( | 0.07612 | 0.948585 | NO | NO | NA |
| Ednra | ENSMUSG( | ENSMUSG( | 1.150539 | 0.301322 | NO | NO | NA |
| 3300005D | ENSMUSG( | ENSMUSG( | 0.762227 | 0.10601 | NO | NO | NA |
| Gm15481 | ENSMUSG( | ENSMUSG( | 0.451307 | 0.567231 | NO | NO | NA |
| Armc2 | ENSMUSG( | ENSMUSG( | -0.56637 | 0.363052 | NO | NO | NA |
| Gm7901 | ENSMUSG( | ENSMUSG( | 0.826841 | 0.006257 | YES | NO | NA |
| Pcdhb14 | ENSMUSG( | ENSMUSG( | 0.567658 | 0.590281 | NO | NO | NA |
| 2310043M | ENSMUSG( | ENSMUSG( | -0.22351 | 0.474914 | NO | NO | NA |
| Sema3c | ENSMUSG( | ENSMUSG( | 0.565559 | 0.616503 | NO | NO | NA |
| 9530077C( | ENSMUSG( | ENSMUSG( | 0.340854 | 0.33517 | NO | NO | NA |
| Gm33994 | ENSMUSG( | ENSMUSG( | 0.311983 | 0.738822 | NO | NO | NA |
| Gm13680 | ENSMUSG( | ENSMUSG( | 0.48265 | 0.118902 | NO | NO | NA |
| Rapsn | ENSMUSG( | ENSMUSG( | 1.080898 | 0.00375 | YES | YES | Upregulated |
| Gm3325 | ENSMUSG( | ENSMUSG( | -0.07141 | 0.979817 | NO | NO | NA |
| Gm5094 | ENSMUSG( | ENSMUSG( | 0.457062 | 0.661684 | NO | NO | NA |
| Cdh2 | ENSMUSG( | ENSMUSG( | 0.667204 | 0.476775 | NO | NO | NA |
| Gm8878 | ENSMUSG( | ENSMUSG( | 0.645619 | 0.550678 | NO | NO | NA |
| Gm43705 | ENSMUSG( | ENSMUSG( | 0.705193 | 0.512637 | NO | NO | NA |
| Bend6 | ENSMUSG( | ENSMUSG( | -0.51182 | 0.27531 | NO | NO | NA |
| Clmp | ENSMUSG( | ENSMUSG( | 0.264548 | 0.82162 | NO | NO | NA |
| Vmn1r55 | ENSMUSG( | ENSMUSG( | 0.774511 | 0.420191 | NO | NO | NA |
| Tifa | ENSMUSG( | ENSMUSG( | 0.678286 | 0.329755 | NO | NO | NA |
| Gm37090 | ENSMUSG( | ENSMUSG( | 0.689875 | 0.047903 | YES | NO | NA |
| Gm13341 | ENSMUSG( | ENSMUSG( | 0.410534 | 0.415514 | NO | NO | NA |

| | | | | | | | |
|---|---|---|---|---|---|---|---|
| Gm44716 | ENSMUSG( | ENSMUSG( | 1.237807 | 0.245114 | NO | NO | NA |
| Car15 | ENSMUSG( | ENSMUSG( | -0.50745 | 0.208913 | NO | NO | NA |
| Gm12655 | ENSMUSG( | ENSMUSG( | -0.21478 | 0.518523 | NO | NO | NA |
| Cyp3a11 | ENSMUSG( | ENSMUSG( | 0.625537 | 0.579897 | NO | NO | NA |
| Clip4 | ENSMUSG( | ENSMUSG( | 0.539383 | 0.543911 | NO | NO | NA |
| Pcdhb22 | ENSMUSG( | ENSMUSG( | 0.868464 | 0.402946 | NO | NO | NA |
| Isl1 | ENSMUSG( | ENSMUSG( | 0.534309 | 0.486835 | NO | NO | NA |
| Klhl35 | ENSMUSG( | ENSMUSG( | 0.423251 | 0.185193 | NO | NO | NA |
| Gm43075 | ENSMUSG( | ENSMUSG( | 0.171451 | 0.822962 | NO | NO | NA |
| ENSMUSG( | ENSMUSG( | ENSMUSG( | 0.040964 | 0.905492 | NO | NO | NA |
| Gm20161 | ENSMUSG( | ENSMUSG( | 0.984095 | 0.052799 | NO | NO | NA |
| Gm10814 | ENSMUSG( | ENSMUSG( | -0.31782 | 0.436803 | NO | NO | NA |
| Gm45185 | ENSMUSG( | ENSMUSG( | 1.427876 | 0.1754 | NO | NO | NA |
| Gm41118 | ENSMUSG( | ENSMUSG( | 0.562305 | 0.592211 | NO | NO | NA |
| 1700122C: | ENSMUSG( | ENSMUSG( | 0.366964 | 0.691427 | NO | NO | NA |
| Abra | ENSMUSG( | ENSMUSG( | 0.845076 | 0.389754 | NO | NO | NA |
| 5330434G | ENSMUSG( | ENSMUSG( | 0.362392 | 0.74256 | NO | NO | NA |
| Dkk3 | ENSMUSG( | ENSMUSG( | 0.022259 | 0.963299 | NO | NO | NA |
| Olfr532 | ENSMUSG( | ENSMUSG( | 1.116843 | 0.263165 | NO | NO | NA |
| Itgae | ENSMUSG( | ENSMUSG( | -0.42753 | 0.118757 | NO | NO | NA |
| Asb18 | ENSMUSG( | ENSMUSG( | 0.45538 | 0.310173 | NO | NO | NA |
| Gm13856 | ENSMUSG( | ENSMUSG( | 0.109293 | 0.803364 | NO | NO | NA |
| 9630014M | ENSMUSG( | ENSMUSG( | 0.786858 | 0.433675 | NO | NO | NA |
| Gm15122 | ENSMUSG( | ENSMUSG( | -0.86767 | 0.002998 | YES | NO | NA |
| Cntf | ENSMUSG( | ENSMUSG( | -0.16303 | 0.643741 | NO | NO | NA |
| Id4 | ENSMUSG( | ENSMUSG( | 0.549535 | 0.559018 | NO | NO | NA |
| Gm47583 | ENSMUSG( | ENSMUSG( | 0.310564 | 0.453153 | NO | NO | NA |
| Gm39469 | ENSMUSG( | ENSMUSG( | -0.22592 | 0.567401 | NO | NO | NA |
| Npas1 | ENSMUSG( | ENSMUSG( | 0.528125 | 0.229729 | NO | NO | NA |
| Emx2 | ENSMUSG( | ENSMUSG( | 0.717103 | 0.505631 | NO | NO | NA |
| ENSMUSG | ENSMUSG( | ENSMUSG( | 0.011004 | 0.982039 | NO | NO | NA |
| Fsd1l | ENSMUSG( | ENSMUSG( | 0.67495 | 0.133425 | NO | NO | NA |
| Lrrc15 | ENSMUSG( | ENSMUSG( | 0.554261 | 0.572839 | NO | NO | NA |
| Trav12n-1 | ENSMUSG( | ENSMUSG( | -0.23695 | 0.915786 | NO | NO | NA |
| Slc16a2 | ENSMUSG( | ENSMUSG( | 0.971069 | 0.319978 | NO | NO | NA |
| Muc1 | ENSMUSG( | ENSMUSG( | 0.503586 | 0.230898 | NO | NO | NA |
| Gm7392 | ENSMUSG( | ENSMUSG( | 0.743241 | 0.010909 | YES | NO | NA |
| Tcaf1 | ENSMUSG( | ENSMUSG( | 0.841454 | 0.264108 | NO | NO | NA |
| Ccdc24 | ENSMUSG( | ENSMUSG( | 1.303119 | 0.028026 | YES | YES | Upregulated |
| Pakap ENS | ENSMUSG( | ENSMUSG( | 0.967876 | 0.738913 | NO | NO | NA |
| C430014B | ENSMUSG( | ENSMUSG( | 0.014323 | 0.976913 | NO | NO | NA |
| ENSMUSG | ENSMUSG( | ENSMUSG( | 0.473104 | 0.468146 | NO | NO | NA |
| Cpa4 | ENSMUSG( | ENSMUSG( | 0.646486 | 0.538735 | NO | NO | NA |
| Gm47423 | ENSMUSG( | ENSMUSG( | 0.54651 | 0.631598 | NO | NO | NA |
| Gm11579 | ENSMUSG( | ENSMUSG( | 0.772905 | 0.112968 | NO | NO | NA |
| Gm43808 | ENSMUSG( | ENSMUSG( | -0.23765 | 0.527421 | NO | NO | NA |
| Olfr735 | ENSMUSG( | ENSMUSG( | 1.402156 | 0.204686 | NO | NO | NA |

| | | | | | | | |
|---|---|---|---|---|---|---|---|
| H2bc13 | ENSMUSG( | ENSMUSG( | 0.652619 | 0.066735 | NO | NO | NA |
| Gm42577 | ENSMUSG( | ENSMUSG( | 0.54576 | 0.494358 | NO | NO | NA |
| ENSMUSG( | ENSMUSG( | ENSMUSG( | -0.53174 | 0.09051 | NO | NO | NA |
| 2010204K: | ENSMUSG( | ENSMUSG( | 1.443109 | 0.000315 | YES | YES | Upregulated |
| Spats2l | ENSMUSG( | ENSMUSG( | 0.047 | 0.953489 | NO | NO | NA |
| Gm38218 | ENSMUSG( | ENSMUSG( | 0.032324 | 0.954178 | NO | NO | NA |
| 5330431K( | ENSMUSG( | ENSMUSG( | 1.013519 | 0.343744 | NO | NO | NA |
| Zfp30 | ENSMUSG( | ENSMUSG( | 0.711749 | 0.342632 | NO | NO | NA |
| 4632418H | ENSMUSG( | ENSMUSG( | -0.24042 | 0.555322 | NO | NO | NA |
| Olfr1348 | ENSMUSG( | ENSMUSG( | 0.356014 | 0.751869 | NO | NO | NA |
| Vmn2r24 | ENSMUSG( | ENSMUSG( | 0.649662 | 0.542475 | NO | NO | NA |
| Gm10069 | ENSMUSG( | ENSMUSG( | -0.30091 | 0.627656 | NO | NO | NA |
| Gm42463 | ENSMUSG( | ENSMUSG( | 0.511097 | 0.571948 | NO | NO | NA |
| Gapt | ENSMUSG( | ENSMUSG( | 0.673406 | 0.499294 | NO | NO | NA |
| Foxa3 | ENSMUSG( | ENSMUSG( | 0.779877 | 0.516584 | NO | NO | NA |
| Olfr727 | ENSMUSG( | ENSMUSG( | 0.728001 | 0.514559 | NO | NO | NA |
| Fem1al | ENSMUSG( | ENSMUSG( | 0.802297 | 0.386056 | NO | NO | NA |
| Aqp11 | ENSMUSG( | ENSMUSG( | 0.614093 | 0.372606 | NO | NO | NA |
| Clvs2 | ENSMUSG( | ENSMUSG( | 0.676871 | 0.561489 | NO | NO | NA |
| Ms4a4d | ENSMUSG( | ENSMUSG( | -0.31937 | 0.434448 | NO | NO | NA |
| H2-Ob | ENSMUSG( | ENSMUSG( | -0.89803 | 0.015868 | YES | NO | NA |
| 1700001G | ENSMUSG( | ENSMUSG( | 0.141128 | 0.820924 | NO | NO | NA |
| Dao | ENSMUSG( | ENSMUSG( | 0.321169 | 0.642983 | NO | NO | NA |
| 2310014F( | ENSMUSG( | ENSMUSG( | 0.78938 | 0.462743 | NO | NO | NA |
| Gm44230 | ENSMUSG( | ENSMUSG( | 0.435644 | 0.701118 | NO | NO | NA |
| 5430421F: | ENSMUSG( | ENSMUSG( | 0.18252 | 0.858246 | NO | NO | NA |
| Hpdl | ENSMUSG( | ENSMUSG( | 0.873532 | 0.003601 | YES | NO | NA |
| Igkv6-25 | ENSMUSG( | ENSMUSG( | 0.984277 | NA | NO | NO | NA |
| 1700112J1 | ENSMUSG( | ENSMUSG( | 0.990318 | 0.283865 | NO | NO | NA |
| Gm37121 | ENSMUSG( | ENSMUSG( | 0.2275 | 0.789447 | NO | NO | NA |
| 9430037G | ENSMUSG( | ENSMUSG( | 0.533417 | 0.459146 | NO | NO | NA |
| Gm37551 | ENSMUSG( | ENSMUSG( | 0.948005 | 0.400316 | NO | NO | NA |
| Wt1os | ENSMUSG( | ENSMUSG( | 0.292856 | 0.760343 | NO | NO | NA |
| Vldlr | ENSMUSG( | ENSMUSG( | 1.213077 | 0.205952 | NO | NO | NA |
| Gm49588 | ENSMUSG( | ENSMUSG( | 0.506863 | 0.56965 | NO | NO | NA |
| Gm46404 | ENSMUSG( | ENSMUSG( | 1.054111 | 0.000124 | YES | YES | Upregulated |
| Gm6483 | ENSMUSG( | ENSMUSG( | 0.093792 | 0.779557 | NO | NO | NA |
| Sez6 | ENSMUSG( | ENSMUSG( | 0.525888 | 0.609539 | NO | NO | NA |
| Gm8894 | ENSMUSG( | ENSMUSG( | -0.00387 | 0.995212 | NO | NO | NA |
| Dmbx1 | ENSMUSG( | ENSMUSG( | 0.58514 | 0.560002 | NO | NO | NA |
| ENSMUSG( | ENSMUSG( | ENSMUSG( | 0.387505 | 0.66524 | NO | NO | NA |
| Rnf222 | ENSMUSG( | ENSMUSG( | 0.453275 | 0.665775 | NO | NO | NA |
| Derl3 | ENSMUSG( | ENSMUSG( | 0.439239 | 0.430963 | NO | NO | NA |
| Gm11591 | ENSMUSG( | ENSMUSG( | 0.466429 | 0.669702 | NO | NO | NA |
| Gm45833 | ENSMUSG( | ENSMUSG( | 0.512272 | 0.633247 | NO | NO | NA |
| Gm44249 | ENSMUSG( | ENSMUSG( | -0.20084 | 0.756401 | NO | NO | NA |
| Bmp4 | ENSMUSG( | ENSMUSG( | 0.480012 | 0.633815 | NO | NO | NA |

| | | | | | | | |
|---|---|---|---|---|---|---|---|
| Gm7730 | ENSMUSG( | ENSMUSG( | 0.288671 | 0.525251 | NO | NO | NA |
| 5430420F( | ENSMUSG( | ENSMUSG( | 0.828646 | 0.366592 | NO | NO | NA |
| Gm10544 | ENSMUSG( | ENSMUSG( | 1.115473 | 0.274147 | NO | NO | NA |
| Gm49197 | ENSMUSG( | ENSMUSG( | 1.227276 | 0.10893 | NO | NO | NA |
| B3galt9 | ENSMUSG( | ENSMUSG( | 0.992864 | 0.267007 | NO | NO | NA |
| Gm43533 | ENSMUSG( | ENSMUSG( | -0.0813 | 0.942764 | NO | NO | NA |
| Gm3362 | ENSMUSG( | ENSMUSG( | 0.248766 | 0.556072 | NO | NO | NA |
| Vmn2r23 | ENSMUSG( | ENSMUSG( | 0.081036 | 0.937234 | NO | NO | NA |
| Aox3 | ENSMUSG( | ENSMUSG( | 1.014836 | 0.367697 | NO | NO | NA |
| Dact3 | ENSMUSG( | ENSMUSG( | 0.503071 | 0.438825 | NO | NO | NA |
| E030032P: | ENSMUSG( | ENSMUSG( | 0.830251 | 0.467463 | NO | NO | NA |
| Itga2b | ENSMUSG( | ENSMUSG( | -0.2502 | 0.381182 | NO | NO | NA |
| Ccno | ENSMUSG( | ENSMUSG( | 0.535704 | 0.220117 | NO | NO | NA |
| Olfr394 | ENSMUSG( | ENSMUSG( | 0.665306 | 0.517147 | NO | NO | NA |
| Gm30159 | ENSMUSG( | ENSMUSG( | 0.803162 | 0.514914 | NO | NO | NA |
| Art3 | ENSMUSG( | ENSMUSG( | 0.340195 | 0.630359 | NO | NO | NA |
| ENSMUSG | ENSMUSG( | ENSMUSG( | 1.073542 | 0.314567 | NO | NO | NA |
| Gm47346 | ENSMUSG( | ENSMUSG( | 0.30424 | 0.765058 | NO | NO | NA |
| Gm43860 | ENSMUSG( | ENSMUSG( | -0.08179 | 0.797703 | NO | NO | NA |
| Otoa | ENSMUSG( | ENSMUSG( | 0.739944 | 0.41758 | NO | NO | NA |
| Gm8276 | ENSMUSG( | ENSMUSG( | 0.065622 | 0.884793 | NO | NO | NA |
| ENSMUSG | ENSMUSG( | ENSMUSG( | 0.251582 | 0.680463 | NO | NO | NA |
| Tmem215 | ENSMUSG( | ENSMUSG( | 0.98772 | 0.342342 | NO | NO | NA |
| ENSMUSG | ENSMUSG( | ENSMUSG( | -0.27978 | 0.534088 | NO | NO | NA |
| Gm5608 | ENSMUSG( | ENSMUSG( | 0.661877 | 0.406581 | NO | NO | NA |
| Gm48208 | ENSMUSG( | ENSMUSG( | 1.377196 | 0.234325 | NO | NO | NA |
| Gm50164 | ENSMUSG( | ENSMUSG( | 0.699004 | 0.529609 | NO | NO | NA |
| Olfr312 | ENSMUSG( | ENSMUSG( | 0.506238 | 0.616503 | NO | NO | NA |
| 2310015D | ENSMUSG( | ENSMUSG( | 0.876822 | 0.454144 | NO | NO | NA |
| Defb42 | ENSMUSG( | ENSMUSG( | 0.877619 | 0.422826 | NO | NO | NA |
| Zfp629 | ENSMUSG( | ENSMUSG( | 0.379605 | 0.405248 | NO | NO | NA |
| Gm38014 | ENSMUSG( | ENSMUSG( | 0.217532 | 0.74274 | NO | NO | NA |
| Rassf4 | ENSMUSG( | ENSMUSG( | -0.74178 | 0.022856 | YES | NO | NA |
| Eln | ENSMUSG( | ENSMUSG( | 0.853039 | 0.535871 | NO | NO | NA |
| Ift81 | ENSMUSG( | ENSMUSG( | 0.692316 | 0.354641 | NO | NO | NA |
| Gm6249 | ENSMUSG( | ENSMUSG( | 0.704034 | 0.516631 | NO | NO | NA |
| Gm43050 | ENSMUSG( | ENSMUSG( | 0.71896 | 0.486322 | NO | NO | NA |
| Gm50334 | ENSMUSG( | ENSMUSG( | 0.562868 | 0.493793 | NO | NO | NA |
| Wwc1 | ENSMUSG( | ENSMUSG( | 0.533114 | 0.38535 | NO | NO | NA |
| Macrod2 | ENSMUSG( | ENSMUSG( | 0.922135 | 0.16122 | NO | NO | NA |
| Obox3 | ENSMUSG( | ENSMUSG( | 0.939018 | 0.368672 | NO | NO | NA |
| B3gnt8 | ENSMUSG( | ENSMUSG( | 0.13323 | 0.773449 | NO | NO | NA |
| Gm30938 | ENSMUSG( | ENSMUSG( | 0.933913 | 0.438825 | NO | NO | NA |
| Fam217b | ENSMUSG( | ENSMUSG( | 0.109177 | 0.884746 | NO | NO | NA |
| Aqp4 | ENSMUSG( | ENSMUSG( | 0.644104 | 0.547964 | NO | NO | NA |
| Fcgr4 | ENSMUSG( | ENSMUSG( | 0.606914 | 0.317776 | NO | NO | NA |
| Tshr | ENSMUSG( | ENSMUSG( | 0.678883 | 0.476112 | NO | NO | NA |

| | | | | | | | |
|---|---|---|---|---|---|---|---|
| Tmem205 | ENSMUSG( | ENSMUSG( | 0.586544 | 0.09389 | NO | NO | NA |
| Olfr77 | ENSMUSG( | ENSMUSG( | 0.269205 | 0.560577 | NO | NO | NA |
| Nr2f1 | ENSMUSG( | ENSMUSG( | 0.933439 | 0.353555 | NO | NO | NA |
| Gm17244 | ENSMUSG( | ENSMUSG( | 0.850243 | 0.329639 | NO | NO | NA |
| Gm37367 | ENSMUSG( | ENSMUSG( | 0.558208 | 0.56488 | NO | NO | NA |
| D030025P | ENSMUSG( | ENSMUSG( | 0.851435 | 0.416164 | NO | NO | NA |
| 6430511E: | ENSMUSG( | ENSMUSG( | -0.20237 | 0.580825 | NO | NO | NA |
| Nxpe4 | ENSMUSG( | ENSMUSG( | 1.149847 | 0.085396 | NO | NO | NA |
| Gm42842 | ENSMUSG( | ENSMUSG( | 1.015079 | 0.366356 | NO | NO | NA |
| Gm50087 | ENSMUSG( | ENSMUSG( | 0.328761 | 0.46362 | NO | NO | NA |
| Ffar1 | ENSMUSG( | ENSMUSG( | 0.330598 | 0.723233 | NO | NO | NA |
| Rslcan18 | ENSMUSG( | ENSMUSG( | 0.474508 | 0.457305 | NO | NO | NA |
| Pcdhga12 | ENSMUSG( | ENSMUSG( | -0.81688 | 0.157988 | NO | NO | NA |
| Igf2 | ENSMUSG( | ENSMUSG( | 0.781438 | 0.191212 | NO | NO | NA |
| Chd5 | ENSMUSG( | ENSMUSG( | 0.696715 | 0.123478 | NO | NO | NA |
| Dlx6 | ENSMUSG( | ENSMUSG( | 0.337131 | 0.732349 | NO | NO | NA |
| Olfr161 | ENSMUSG( | ENSMUSG( | 0.642369 | 0.531114 | NO | NO | NA |
| 5430414B: | ENSMUSG( | ENSMUSG( | 0.015618 | 0.964479 | NO | NO | NA |
| Lrfn5 | ENSMUSG( | ENSMUSG( | 0.116348 | 0.91013 | NO | NO | NA |
| Pcdh19 | ENSMUSG( | ENSMUSG( | 0.543571 | 0.607785 | NO | NO | NA |
| Gm17259 | ENSMUSG( | ENSMUSG( | -0.00173 | 0.995684 | NO | NO | NA |
| Yes1 | ENSMUSG( | ENSMUSG( | 1.204045 | 0.009916 | YES | YES | Upregulated |
| Gm50100 | ENSMUSG( | ENSMUSG( | 0.315346 | 0.733927 | NO | NO | NA |
| ENSMUSG( | ENSMUSG( | ENSMUSG( | -0.64028 | 0.127985 | NO | NO | NA |
| Rubie | ENSMUSG( | ENSMUSG( | 0.749303 | 0.478634 | NO | NO | NA |
| Tectb | ENSMUSG( | ENSMUSG( | 0.731135 | 0.493099 | NO | NO | NA |
| Gm12279 | ENSMUSG( | ENSMUSG( | 0.287615 | 0.404157 | NO | NO | NA |
| Sap30bpos | ENSMUSG( | ENSMUSG( | 0.441123 | 0.391447 | NO | NO | NA |
| Akap14 | ENSMUSG( | ENSMUSG( | 1.368723 | NA | NO | NO | NA |
| Olfr1438-p | ENSMUSG( | ENSMUSG( | 0.777756 | 0.450484 | NO | NO | NA |
| ENSMUSG | ENSMUSG( | ENSMUSG( | 0.491495 | 0.322972 | NO | NO | NA |
| A830012C | ENSMUSG( | ENSMUSG( | 0.736631 | 0.283254 | NO | NO | NA |
| Ctcflos | ENSMUSG( | ENSMUSG( | 0.328871 | 0.736093 | NO | NO | NA |
| Gm19514 | ENSMUSG( | ENSMUSG( | 1.248895 | 0.182925 | NO | NO | NA |
| Igkv5-39 | ENSMUSG( | ENSMUSG( | -0.96949 | 0.602223 | NO | NO | NA |
| Gm45236 | ENSMUSG( | ENSMUSG( | 0.583645 | 0.081553 | NO | NO | NA |
| Sp8 | ENSMUSG( | ENSMUSG( | 0.627837 | 0.50518 | NO | NO | NA |
| Upb1 | ENSMUSG( | ENSMUSG( | 0.878283 | 0.219914 | NO | NO | NA |
| ENSMUSG | ENSMUSG( | ENSMUSG( | 0.329771 | 0.444307 | NO | NO | NA |
| E330037G | ENSMUSG( | ENSMUSG( | -0.34003 | 0.561517 | NO | NO | NA |
| Gm18910 | ENSMUSG( | ENSMUSG( | 0.666472 | 0.633232 | NO | NO | NA |
| Fzd4 | ENSMUSG( | ENSMUSG( | -0.28945 | 0.620176 | NO | NO | NA |
| Gm42447 | ENSMUSG( | ENSMUSG( | 0.454684 | 0.653269 | NO | NO | NA |
| Gm43267 | ENSMUSG( | ENSMUSG( | -0.08922 | 0.854133 | NO | NO | NA |
| Pter | ENSMUSG( | ENSMUSG( | 0.842463 | 0.020008 | YES | NO | NA |
| Gm42448 | ENSMUSG( | ENSMUSG( | 0.728385 | 0.463414 | NO | NO | NA |
| Gm8449 | ENSMUSG( | ENSMUSG( | 0.963473 | 0.375241 | NO | NO | NA |

| | | | | | | | |
|---|---|---|---|---|---|---|---|
| Olfr642 | ENSMUSG( | ENSMUSG( | 0.419241 | 0.727144 | NO | NO | NA |
| Gm37205 | ENSMUSG( | ENSMUSG( | 0.871056 | 0.392834 | NO | NO | NA |
| Coprs | ENSMUSG( | ENSMUSG( | 0.540613 | 0.181801 | NO | NO | NA |
| Gm40309 | ENSMUSG( | ENSMUSG( | -0.59268 | 0.146931 | NO | NO | NA |
| ENSMUSG( | ENSMUSG( | ENSMUSG( | 0.701885 | 0.386884 | NO | NO | NA |
| Gm48734 | ENSMUSG( | ENSMUSG( | -0.6789 | 0.036002 | YES | NO | NA |
| Gm21046 | ENSMUSG( | ENSMUSG( | 0.663149 | 0.490375 | NO | NO | NA |
| Flrt2 | ENSMUSG( | ENSMUSG( | 1.037869 | 0.406071 | NO | NO | NA |
| Ctrl | ENSMUSG( | ENSMUSG( | 0.111306 | 0.759918 | NO | NO | NA |
| Nrl | ENSMUSG( | ENSMUSG( | 0.720403 | 0.475513 | NO | NO | NA |
| Ccdc50-ps | ENSMUSG( | ENSMUSG( | 0.393027 | 0.645355 | NO | NO | NA |
| Ptk6 | ENSMUSG( | ENSMUSG( | 0.622534 | 0.612737 | NO | NO | NA |
| Mafb | ENSMUSG( | ENSMUSG( | -0.52468 | 0.111065 | NO | NO | NA |
| 1810006J( | ENSMUSG( | ENSMUSG( | -0.21331 | 0.724922 | NO | NO | NA |
| Gm45388 | ENSMUSG( | ENSMUSG( | 1.250997 | 0.268553 | NO | NO | NA |
| Ighd | ENSMUSG( | ENSMUSG( | 0.924973 | 0.093463 | NO | NO | NA |
| Gm38071 | ENSMUSG( | ENSMUSG( | 0.316047 | 0.784854 | NO | NO | NA |
| Cdkn1c | ENSMUSG( | ENSMUSG( | 1.046255 | 0.00118 | YES | YES | Upregulated |
| Gm16300 | ENSMUSG( | ENSMUSG( | 1.152249 | 0.274435 | NO | NO | NA |
| Gm2436 | ENSMUSG( | ENSMUSG( | -5.52856 | NA | NO | NO | NA |
| Gm12966 | ENSMUSG( | ENSMUSG( | 0.314026 | 0.355467 | NO | NO | NA |
| Ankrd60 | ENSMUSG( | ENSMUSG( | 0.803288 | 0.333591 | NO | NO | NA |
| Mzb1 | ENSMUSG( | ENSMUSG( | -1.00375 | 0.244982 | NO | NO | NA |
| Lonrf3 | ENSMUSG( | ENSMUSG( | 0.278358 | 0.637997 | NO | NO | NA |
| Gpat2 | ENSMUSG( | ENSMUSG( | 0.902955 | 0.601962 | NO | NO | NA |
| Gm40155 | ENSMUSG( | ENSMUSG( | 1.403537 | 0.200927 | NO | NO | NA |
| Apol7b | ENSMUSG( | ENSMUSG( | -0.37982 | 0.438398 | NO | NO | NA |
| Lrrc8e | ENSMUSG( | ENSMUSG( | 0.637707 | 0.524027 | NO | NO | NA |
| Vmn1r58 | ENSMUSG( | ENSMUSG( | 0.481327 | 0.636349 | NO | NO | NA |
| D530018E | ENSMUSG( | ENSMUSG( | -0.49549 | 0.229373 | NO | NO | NA |
| Zfp174 | ENSMUSG( | ENSMUSG( | -0.60018 | 0.07109 | NO | NO | NA |
| Gm44001 | ENSMUSG( | ENSMUSG( | 0.489828 | 0.642754 | NO | NO | NA |
| Olfr1096 | ENSMUSG( | ENSMUSG( | 0.796677 | 0.433539 | NO | NO | NA |
| Gm5403 | ENSMUSG( | ENSMUSG( | 0.568863 | 0.596711 | NO | NO | NA |
| Gm44724 | ENSMUSG( | ENSMUSG( | 0.905637 | 0.422508 | NO | NO | NA |
| Disp2 | ENSMUSG( | ENSMUSG( | 0.540543 | 0.603838 | NO | NO | NA |
| Gm48855 | ENSMUSG( | ENSMUSG( | 0.181013 | 0.787593 | NO | NO | NA |
| Gm12960 | ENSMUSG( | ENSMUSG( | 0.76074 | 0.031638 | YES | NO | NA |
| C430019N | ENSMUSG( | ENSMUSG( | 0.833897 | 0.43299 | NO | NO | NA |
| Grid1 | ENSMUSG( | ENSMUSG( | 0.658341 | 0.531711 | NO | NO | NA |
| Gm50078 | ENSMUSG( | ENSMUSG( | 1.233529 | 0.2583 | NO | NO | NA |
| A930016O | ENSMUSG( | ENSMUSG( | 0.828332 | 0.055568 | NO | NO | NA |
| ENSMUSG( | ENSMUSG( | ENSMUSG( | -0.43666 | 0.162143 | NO | NO | NA |
| Sgtb | ENSMUSG( | ENSMUSG( | 0.184769 | 0.816509 | NO | NO | NA |
| Foxq1 | ENSMUSG( | ENSMUSG( | 0.579047 | 0.513739 | NO | NO | NA |
| Cage1 | ENSMUSG( | ENSMUSG( | 0.912012 | 0.00363 | YES | NO | NA |
| Trav7d-5 | ENSMUSG( | ENSMUSG( | -1.89738 | 0.443772 | NO | NO | NA |

| | | | | | | | |
|---|---|---|---|---|---|---|---|
| Adgrl2 | ENSMUSG( | ENSMUSG( | 0.44409 | 0.512637 | NO | NO | NA |
| Gm9210 | ENSMUSG( | ENSMUSG( | 0.581065 | 0.638353 | NO | NO | NA |
| Lyl1 | ENSMUSG( | ENSMUSG( | 0.371729 | 0.500647 | NO | NO | NA |
| Plekha4 | ENSMUSG( | ENSMUSG( | 0.39066 | 0.446966 | NO | NO | NA |
| 2410003L1 | ENSMUSG( | ENSMUSG( | 0.358718 | 0.655714 | NO | NO | NA |
| Prrg4 | ENSMUSG( | ENSMUSG( | 0.459035 | 0.242449 | NO | NO | NA |
| Gm48822 | ENSMUSG( | ENSMUSG( | 0.447267 | 0.684434 | NO | NO | NA |
| Gm14057 | ENSMUSG( | ENSMUSG( | -0.17488 | 0.658039 | NO | NO | NA |
| Vmn1r230 | ENSMUSG( | ENSMUSG( | 0.370789 | 0.72686 | NO | NO | NA |
| Gm43422 | ENSMUSG( | ENSMUSG( | 1.199909 | 0.224252 | NO | NO | NA |
| Camp | ENSMUSG( | ENSMUSG( | 3.169528 | 0.049263 | YES | YES | Upregulated |
| C030029H | ENSMUSG( | ENSMUSG( | 1.305084 | 0.182375 | NO | NO | NA |
| Rpl17-ps5 | ENSMUSG( | ENSMUSG( | 0.482436 | 0.573931 | NO | NO | NA |
| Frg2f1 | ENSMUSG( | ENSMUSG( | -0.05821 | 0.87566 | NO | NO | NA |
| Scn7a | ENSMUSG( | ENSMUSG( | 0.72928 | 0.499353 | NO | NO | NA |
| Tmem25 | ENSMUSG( | ENSMUSG( | 0.29459 | 0.454063 | NO | NO | NA |
| Gm9911 | ENSMUSG( | ENSMUSG( | 0.956115 | 0.350224 | NO | NO | NA |
| Gm37906 | ENSMUSG( | ENSMUSG( | -0.17104 | 0.712916 | NO | NO | NA |
| Gm50196 | ENSMUSG( | ENSMUSG( | 0.296509 | 0.733718 | NO | NO | NA |
| Gm43566 | ENSMUSG( | ENSMUSG( | 0.573587 | 0.230313 | NO | NO | NA |
| Gm47923 | ENSMUSG( | ENSMUSG( | 0.779172 | 0.463661 | NO | NO | NA |
| Ddc | ENSMUSG( | ENSMUSG( | 0.73565 | 0.36873 | NO | NO | NA |
| Gm8098 | ENSMUSG( | ENSMUSG( | 0.863164 | 0.316648 | NO | NO | NA |
| Ano2 | ENSMUSG( | ENSMUSG( | 0.656882 | 0.574263 | NO | NO | NA |
| Gm20731 | ENSMUSG( | ENSMUSG( | 0.321214 | 0.322847 | NO | NO | NA |
| Gm11560 | ENSMUSG( | ENSMUSG( | 0.487906 | 0.151819 | NO | NO | NA |
| Olfr983 | ENSMUSG( | ENSMUSG( | 0.749636 | 0.497075 | NO | NO | NA |
| Cdo1 | ENSMUSG( | ENSMUSG( | -0.45793 | 0.349183 | NO | NO | NA |
| Insyn2a | ENSMUSG( | ENSMUSG( | 0.402703 | 0.616932 | NO | NO | NA |
| Zfyve28 | ENSMUSG( | ENSMUSG( | -0.1786 | 0.581457 | NO | NO | NA |
| Fabp4 | ENSMUSG( | ENSMUSG( | 0.770101 | 0.315494 | NO | NO | NA |
| Gm39041 | ENSMUSG( | ENSMUSG( | 0.362169 | 0.612413 | NO | NO | NA |
| Pla2g4d | ENSMUSG( | ENSMUSG( | 0.865644 | 0.35992 | NO | NO | NA |
| Bricd5 | ENSMUSG( | ENSMUSG( | 1.055944 | 0.002528 | YES | YES | Upregulated |
| 5430431A | ENSMUSG( | ENSMUSG( | -0.14087 | 0.792881 | NO | NO | NA |
| Rpl15-ps6 | ENSMUSG( | ENSMUSG( | 0.367871 | 0.403874 | NO | NO | NA |
| Gm21188 | ENSMUSG( | ENSMUSG( | 0.118416 | 0.857585 | NO | NO | NA |
| Gm53040 | ENSMUSG( | ENSMUSG( | 0.506948 | 0.619675 | NO | NO | NA |
| Smim40 | ENSMUSG( | ENSMUSG( | 0.050916 | 0.932228 | NO | NO | NA |
| Gm6767 | ENSMUSG( | ENSMUSG( | 0.601283 | 0.168698 | NO | NO | NA |
| Gng11 | ENSMUSG( | ENSMUSG( | 1.699809 | 0.000661 | YES | YES | Upregulated |
| ENSMUSG( | ENSMUSG( | ENSMUSG( | 0.765904 | 0.452717 | NO | NO | NA |
| Gm47625 | ENSMUSG( | ENSMUSG( | 0.971711 | 0.321265 | NO | NO | NA |
| Zcchc24 | ENSMUSG( | ENSMUSG( | -0.08751 | 0.811685 | NO | NO | NA |
| ENSMUSG | ENSMUSG( | ENSMUSG( | 0.188644 | 0.659838 | NO | NO | NA |
| Tshb | ENSMUSG( | ENSMUSG( | 0.672016 | 0.186078 | NO | NO | NA |
| Gm10686 | ENSMUSG( | ENSMUSG( | 0.010136 | 0.984173 | NO | NO | NA |

| | | | | | | | |
|---|---|---|---|---|---|---|---|
| Scimp | ENSMUSG( | ENSMUSG( | 0.266153 | 0.563091 | NO | NO | NA |
| 4921536K: | ENSMUSG( | ENSMUSG( | 0.275425 | 0.764257 | NO | NO | NA |
| Hoxb9 | ENSMUSG( | ENSMUSG( | 0.914479 | 0.430372 | NO | NO | NA |
| Gm44190 | ENSMUSG( | ENSMUSG( | 0.814392 | 0.388314 | NO | NO | NA |
| Gm31600 | ENSMUSG( | ENSMUSG( | 0.416105 | 0.701752 | NO | NO | NA |
| Csmd3 | ENSMUSG( | ENSMUSG( | -0.0813 | 0.938589 | NO | NO | NA |
| Igsf5 | ENSMUSG( | ENSMUSG( | 1.843578 | 0.009486 | YES | YES | Upregulated |
| Gm38152 | ENSMUSG( | ENSMUSG( | 1.065152 | 0.342685 | NO | NO | NA |
| Gm10537 | ENSMUSG( | ENSMUSG( | 0.149951 | 0.879282 | NO | NO | NA |
| Trpv4 | ENSMUSG( | ENSMUSG( | 0.703267 | 0.155363 | NO | NO | NA |
| Gm49493 | ENSMUSG( | ENSMUSG( | 0.366524 | 0.432602 | NO | NO | NA |
| Gm44873 | ENSMUSG( | ENSMUSG( | 0.629676 | 0.425984 | NO | NO | NA |
| Gm48206 | ENSMUSG( | ENSMUSG( | 0.336561 | 0.752616 | NO | NO | NA |
| Phyhip | ENSMUSG( | ENSMUSG( | 0.408554 | 0.639774 | NO | NO | NA |
| Gm19040 | ENSMUSG( | ENSMUSG( | 0.392092 | 0.538242 | NO | NO | NA |
| Gm36963 | ENSMUSG( | ENSMUSG( | 0.888983 | 0.498876 | NO | NO | NA |
| Gm43335 | ENSMUSG( | ENSMUSG( | 0.091542 | 0.895378 | NO | NO | NA |
| 1110019D | ENSMUSG( | ENSMUSG( | -0.86367 | 0.026644 | YES | NO | NA |
| Gm15581 | ENSMUSG( | ENSMUSG( | 1.25599 | 0.291309 | NO | NO | NA |
| Sema6c | ENSMUSG( | ENSMUSG( | -0.82386 | 0.014634 | YES | NO | NA |
| Gm14508 | ENSMUSG( | ENSMUSG( | 0.146623 | 0.660784 | NO | NO | NA |
| A530065N | ENSMUSG( | ENSMUSG( | 0.415053 | 0.720059 | NO | NO | NA |
| Myo16 | ENSMUSG( | ENSMUSG( | -0.83367 | 0.072402 | NO | NO | NA |
| Zfp488 | ENSMUSG( | ENSMUSG( | 0.62393 | 0.563174 | NO | NO | NA |
| ENSMUSG( | ENSMUSG( | ENSMUSG( | 0.552461 | 0.200627 | NO | NO | NA |
| Car8 | ENSMUSG( | ENSMUSG( | 1.100002 | 0.336143 | NO | NO | NA |
| Hmgn3 | ENSMUSG( | ENSMUSG( | 0.261656 | 0.530158 | NO | NO | NA |
| Gm2225 | ENSMUSG( | ENSMUSG( | 0.704238 | 0.028549 | YES | NO | NA |
| Snord3b2 | ENSMUSG( | ENSMUSG( | -6.22345 | NA | NO | NO | NA |
| ENSMUSG | ENSMUSG( | ENSMUSG( | -0.58699 | 0.022658 | YES | NO | NA |
| Fgfrl1 | ENSMUSG( | ENSMUSG( | 0.182035 | 0.667108 | NO | NO | NA |
| Comp | ENSMUSG( | ENSMUSG( | -0.69637 | 0.03861 | YES | NO | NA |
| Gm47948 | ENSMUSG( | ENSMUSG( | 0.743704 | 0.512691 | NO | NO | NA |
| Gm26698 | ENSMUSG( | ENSMUSG( | -0.18084 | 0.683506 | NO | NO | NA |
| Gm19096 | ENSMUSG( | ENSMUSG( | 0.676577 | 0.535846 | NO | NO | NA |
| Spint1 | ENSMUSG( | ENSMUSG( | -0.19703 | 0.680196 | NO | NO | NA |
| Gm12903 | ENSMUSG( | ENSMUSG( | 0.785013 | 0.417226 | NO | NO | NA |
| Gm15672 | ENSMUSG( | ENSMUSG( | -0.07421 | 0.811904 | NO | NO | NA |
| Gm19569 | ENSMUSG( | ENSMUSG( | 0.277656 | 0.651851 | NO | NO | NA |
| Maged1 | ENSMUSG( | ENSMUSG( | 0.84167 | 0.020981 | YES | NO | NA |
| Whrn | ENSMUSG( | ENSMUSG( | 0.701956 | 0.083555 | NO | NO | NA |
| 1110002J( | ENSMUSG( | ENSMUSG( | 1.014927 | 0.349957 | NO | NO | NA |
| Gm26910 | ENSMUSG( | ENSMUSG( | -0.79715 | 0.002262 | YES | NO | NA |
| Cacnb2 | ENSMUSG( | ENSMUSG( | 0.662302 | 0.435116 | NO | NO | NA |
| Zfp708 | ENSMUSG( | ENSMUSG( | 1.152703 | 0.421177 | NO | NO | NA |
| Olfr109 | ENSMUSG( | ENSMUSG( | 0.737405 | 0.483813 | NO | NO | NA |
| Gm45277 | ENSMUSG( | ENSMUSG( | 1.349385 | 0.28101 | NO | NO | NA |

| | | | | | | | |
|---|---|---|---|---|---|---|---|
| Atad3aos | ENSMUSG( | ENSMUSG( | 0.333571 | 0.562501 | NO | NO | NA |
| Zfp575 | ENSMUSG( | ENSMUSG( | 0.052057 | 0.87566 | NO | NO | NA |
| Elavl4 | ENSMUSG( | ENSMUSG( | 0.700917 | 0.491395 | NO | NO | NA |
| Gm13594 | ENSMUSG( | ENSMUSG( | 0.999027 | 0.037355 | YES | NO | NA |
| Gpc6 | ENSMUSG( | ENSMUSG( | 1.015219 | 0.223177 | NO | NO | NA |
| Dsg1b | ENSMUSG( | ENSMUSG( | 0.55612 | 0.635461 | NO | NO | NA |
| Syt4 | ENSMUSG( | ENSMUSG( | 0.795952 | 0.432797 | NO | NO | NA |
| Gm43371 | ENSMUSG( | ENSMUSG( | 0.64052 | 0.566196 | NO | NO | NA |
| Gm38162 | ENSMUSG( | ENSMUSG( | 0.41082 | 0.330787 | NO | NO | NA |
| Gm15772 | ENSMUSG( | ENSMUSG( | 0.533459 | 0.250089 | NO | NO | NA |
| BC048644 | ENSMUSG( | ENSMUSG( | 0.401291 | 0.610434 | NO | NO | NA |
| Gm14858 | ENSMUSG( | ENSMUSG( | 0.629113 | 0.603284 | NO | NO | NA |
| Vash2 | ENSMUSG( | ENSMUSG( | 0.865098 | 0.171937 | NO | NO | NA |
| Cfap300 | ENSMUSG( | ENSMUSG( | 0.824816 | 0.174694 | NO | NO | NA |
| Gm38299 | ENSMUSG( | ENSMUSG( | 0.206426 | 0.835561 | NO | NO | NA |
| ENSMUSG( | ENSMUSG( | ENSMUSG( | 0.582027 | 0.588052 | NO | NO | NA |
| Gm17096 | ENSMUSG( | ENSMUSG( | -0.05716 | 0.864014 | NO | NO | NA |
| Ighv2-2 | ENSMUSG( | ENSMUSG( | 1.166564 | 0.572582 | NO | NO | NA |
| Gm38115 | ENSMUSG( | ENSMUSG( | 0.50031 | 0.575381 | NO | NO | NA |
| Gm14496 | ENSMUSG( | ENSMUSG( | 0.351516 | 0.709237 | NO | NO | NA |
| Gm17971 | ENSMUSG( | ENSMUSG( | 0.373418 | 0.490566 | NO | NO | NA |
| Gm36756 | ENSMUSG( | ENSMUSG( | -0.55287 | 0.41964 | NO | NO | NA |
| Gm12953 | ENSMUSG( | ENSMUSG( | 0.049168 | 0.906287 | NO | NO | NA |
| Dmrta2 | ENSMUSG( | ENSMUSG( | 0.453794 | 0.311779 | NO | NO | NA |
| Samd15 | ENSMUSG( | ENSMUSG( | 0.825466 | 0.131806 | NO | NO | NA |
| F830112A: | ENSMUSG( | ENSMUSG( | -0.13123 | 0.869914 | NO | NO | NA |
| Gm10845 | ENSMUSG( | ENSMUSG( | 0.338649 | 0.581594 | NO | NO | NA |
| BC037039 | ENSMUSG( | ENSMUSG( | -0.43241 | 0.281848 | NO | NO | NA |
| Gm48308 | ENSMUSG( | ENSMUSG( | -0.62487 | 0.422508 | NO | NO | NA |
| Gm18881 | ENSMUSG( | ENSMUSG( | 1.070208 | 0.422503 | NO | NO | NA |
| Gm48284 | ENSMUSG( | ENSMUSG( | 0.917778 | 0.370135 | NO | NO | NA |
| Mustn1 | ENSMUSG( | ENSMUSG( | 0.676721 | 0.060626 | NO | NO | NA |
| Rab44 | ENSMUSG( | ENSMUSG( | 0.61664 | 0.534088 | NO | NO | NA |
| AV039307 | ENSMUSG( | ENSMUSG( | 0.704712 | 0.150856 | NO | NO | NA |
| Negr1 | ENSMUSG( | ENSMUSG( | 0.361409 | 0.728736 | NO | NO | NA |
| Gm26684 | ENSMUSG( | ENSMUSG( | 0.393684 | 0.691059 | NO | NO | NA |
| Gm19684 | ENSMUSG( | ENSMUSG( | -0.29678 | 0.454019 | NO | NO | NA |
| Gm37785 | ENSMUSG( | ENSMUSG( | 0.619973 | 0.377911 | NO | NO | NA |
| Lncenc1 | ENSMUSG( | ENSMUSG( | 0.819398 | 0.398656 | NO | NO | NA |
| Gm2606 | ENSMUSG( | ENSMUSG( | 0.026831 | 0.973265 | NO | NO | NA |
| ENSMUSG | ENSMUSG( | ENSMUSG( | 0.218197 | 0.619675 | NO | NO | NA |
| Rnf144b | ENSMUSG( | ENSMUSG( | -0.07814 | 0.880534 | NO | NO | NA |
| Gm43023 | ENSMUSG( | ENSMUSG( | -0.06962 | 0.956052 | NO | NO | NA |
| ENSMUSG | ENSMUSG( | ENSMUSG( | 0.559978 | 0.401589 | NO | NO | NA |
| Gm37628 | ENSMUSG( | ENSMUSG( | 1.260454 | 0.295261 | NO | NO | NA |
| Gm7278 | ENSMUSG( | ENSMUSG( | 0.817218 | 0.318864 | NO | NO | NA |
| Triml1 | ENSMUSG( | ENSMUSG( | 0.57999 | 0.575878 | NO | NO | NA |

| | | | | | | | |
|---|---|---|---|---|---|---|---|
| Ighv1-54 | ENSMUSG( | ENSMUSG( | 0.495941 | NA | NO | NO | NA |
| Prr18 | ENSMUSG( | ENSMUSG( | 0.091427 | 0.900004 | NO | NO | NA |
| Gm43527 | ENSMUSG( | ENSMUSG( | 0.34561 | 0.737669 | NO | NO | NA |
| Gm5649 | ENSMUSG( | ENSMUSG( | 0.545674 | 0.610533 | NO | NO | NA |
| Gm48482 | ENSMUSG( | ENSMUSG( | 0.590711 | 0.630467 | NO | NO | NA |
| Ptpru | ENSMUSG( | ENSMUSG( | 0.743873 | 0.2478 | NO | NO | NA |
| Gm35097 | ENSMUSG( | ENSMUSG( | 0.648619 | 0.563894 | NO | NO | NA |
| ENSMUSG | ENSMUSG( | ENSMUSG( | 0.713681 | 0.519031 | NO | NO | NA |
| Gm37652 | ENSMUSG( | ENSMUSG( | 1.033637 | 0.342628 | NO | NO | NA |
| B930025P( | ENSMUSG( | ENSMUSG( | 0.629301 | 0.615791 | NO | NO | NA |
| Gm49719 | ENSMUSG( | ENSMUSG( | 2.15066 | 0.006427 | YES | YES | Upregulated |
| Gm43960 | ENSMUSG( | ENSMUSG( | -0.22012 | 0.821695 | NO | NO | NA |
| Piwil4 | ENSMUSG( | ENSMUSG( | 1.158111 | 0.05289 | NO | NO | NA |
| ENSMUSG | ENSMUSG( | ENSMUSG( | 0.840696 | 0.078381 | NO | NO | NA |
| Gm37198 | ENSMUSG( | ENSMUSG( | 0.365894 | 0.267056 | NO | NO | NA |
| F630042J( | ENSMUSG( | ENSMUSG( | 0.878845 | 0.454144 | NO | NO | NA |
| 1700120B: | ENSMUSG( | ENSMUSG( | -0.39448 | 0.422267 | NO | NO | NA |
| ENSMUSG | ENSMUSG( | ENSMUSG( | 0.741314 | 0.046135 | YES | NO | NA |
| Gm30108 | ENSMUSG( | ENSMUSG( | 1.125694 | 0.005171 | YES | YES | Upregulated |
| 5930420M | ENSMUSG( | ENSMUSG( | -0.06651 | 0.845782 | NO | NO | NA |
| Gm14862 | ENSMUSG( | ENSMUSG( | 1.209253 | 0.208686 | NO | NO | NA |
| Gm37645 | ENSMUSG( | ENSMUSG( | 0.112875 | 0.844539 | NO | NO | NA |
| 5330432J1 | ENSMUSG( | ENSMUSG( | -0.81394 | 0.025232 | YES | NO | NA |
| Tmem229a | ENSMUSG( | ENSMUSG( | 0.215814 | 0.837333 | NO | NO | NA |
| Gm13169 | ENSMUSG( | ENSMUSG( | 2.546709 | 1.69E-05 | YES | YES | Upregulated |
| Gm15834 | ENSMUSG( | ENSMUSG( | -0.13479 | 0.789059 | NO | NO | NA |
| Gm44371 | ENSMUSG( | ENSMUSG( | -0.16016 | 0.863345 | NO | NO | NA |
| Sncb | ENSMUSG( | ENSMUSG( | -0.37555 | 0.499301 | NO | NO | NA |
| Carmil3 | ENSMUSG( | ENSMUSG( | -1.15009 | 0.005028 | YES | YES | Downregulated |
| B230216N | ENSMUSG( | ENSMUSG( | 0.47503 | 0.483913 | NO | NO | NA |
| Bbs7 | ENSMUSG( | ENSMUSG( | 0.928653 | 0.014477 | YES | NO | NA |
| Trbj2-7 | ENSMUSG( | ENSMUSG( | -0.40167 | 0.245986 | NO | NO | NA |
| Gm28221 | ENSMUSG( | ENSMUSG( | -0.22914 | 0.800263 | NO | NO | NA |
| Gm38142 | ENSMUSG( | ENSMUSG( | 0.559705 | 0.593605 | NO | NO | NA |
| Rgl1 | ENSMUSG( | ENSMUSG( | -0.05063 | 0.920459 | NO | NO | NA |
| Adgrv1 | ENSMUSG( | ENSMUSG( | 0.275251 | 0.779954 | NO | NO | NA |
| Sorcs3 | ENSMUSG( | ENSMUSG( | 0.845952 | 0.423382 | NO | NO | NA |
| Gm7162 | ENSMUSG( | ENSMUSG( | -1.05852 | 0.0002 | YES | YES | Downregulated |
| Radx | ENSMUSG( | ENSMUSG( | 0.608108 | 0.471567 | NO | NO | NA |
| Ppifos | ENSMUSG( | ENSMUSG( | -0.27786 | 0.887833 | NO | NO | NA |
| ENSMUSG( | ENSMUSG( | ENSMUSG( | -0.21893 | 0.518069 | NO | NO | NA |
| Gm47588 | ENSMUSG( | ENSMUSG( | 0.467573 | 0.454181 | NO | NO | NA |
| Gm30655 | ENSMUSG( | ENSMUSG( | 0.945454 | 0.261847 | NO | NO | NA |
| BC037032 | ENSMUSG( | ENSMUSG( | 0.132752 | 0.820002 | NO | NO | NA |
| Tdrd5 | ENSMUSG( | ENSMUSG( | 0.623714 | 0.561748 | NO | NO | NA |
| Nudt17 | ENSMUSG( | ENSMUSG( | -0.51599 | 0.315568 | NO | NO | NA |
| D130062J: | ENSMUSG( | ENSMUSG( | 0.037927 | 0.951017 | NO | NO | NA |

| | | | | | | | |
|---|---|---|---|---|---|---|---|
| 9430062P( | ENSMUSG( | ENSMUSG( | 0.527622 | 0.609621 | NO | NO | NA |
| Olfr390 | ENSMUSG( | ENSMUSG( | 0.837481 | 0.444895 | NO | NO | NA |
| Gm29597 | ENSMUSG( | ENSMUSG( | 0.945921 | 0.344041 | NO | NO | NA |
| ENSMUSG | ENSMUSG( | ENSMUSG( | 0.620584 | 0.623372 | NO | NO | NA |
| Prnd | ENSMUSG( | ENSMUSG( | 0.957082 | 0.374168 | NO | NO | NA |
| Adam28 | ENSMUSG( | ENSMUSG( | 0.791333 | 0.483813 | NO | NO | NA |
| Kcnj8 | ENSMUSG( | ENSMUSG( | -1.08763 | 0.039859 | YES | YES | Downregulated |
| Gm29257 | ENSMUSG( | ENSMUSG( | 0.503721 | 0.123254 | NO | NO | NA |
| E230020A( | ENSMUSG( | ENSMUSG( | 1.501647 | 0.062565 | NO | NO | NA |
| F8 | ENSMUSG( | ENSMUSG( | 0.284764 | 0.478628 | NO | NO | NA |
| Gm43260 | ENSMUSG( | ENSMUSG( | 0.382672 | 0.706403 | NO | NO | NA |
| Ighv1-74 | ENSMUSG( | ENSMUSG( | -6.64398 | 3.02E-05 | YES | YES | Downregulated |
| Foxl2 | ENSMUSG( | ENSMUSG( | 1.066064 | 0.358484 | NO | NO | NA |
| Ighv7-3 | ENSMUSG( | ENSMUSG( | -0.95558 | NA | NO | NO | NA |
| Frk | ENSMUSG( | ENSMUSG( | 0.934037 | 0.436467 | NO | NO | NA |
| Etl4 | ENSMUSG( | ENSMUSG( | 0.592449 | 0.465973 | NO | NO | NA |
| Lpar1 | ENSMUSG( | ENSMUSG( | 0.297207 | 0.75766 | NO | NO | NA |
| Gm13461 | ENSMUSG( | ENSMUSG( | 0.476848 | 0.080453 | NO | NO | NA |
| Spink10 | ENSMUSG( | ENSMUSG( | 0.499558 | 0.62562 | NO | NO | NA |
| Tmem26 | ENSMUSG( | ENSMUSG( | 0.596267 | 0.531552 | NO | NO | NA |
| Gm15987 | ENSMUSG( | ENSMUSG( | -0.5531 | 0.026365 | YES | NO | NA |
| Gm35572 | ENSMUSG( | ENSMUSG( | 0.810182 | 0.423134 | NO | NO | NA |
| Lhx2 | ENSMUSG( | ENSMUSG( | 0.167437 | 0.680639 | NO | NO | NA |
| ENSMUSG | ENSMUSG( | ENSMUSG( | 0.331535 | 0.45456 | NO | NO | NA |
| Otos | ENSMUSG( | ENSMUSG( | 1.456064 | 0.000181 | YES | YES | Upregulated |
| Gm19692 | ENSMUSG( | ENSMUSG( | 0.669478 | 0.578309 | NO | NO | NA |
| Sel1l3 | ENSMUSG( | ENSMUSG( | 0.699773 | 0.515887 | NO | NO | NA |
| Gm28942 | ENSMUSG( | ENSMUSG( | -0.77997 | 0.103349 | NO | NO | NA |
| Nr1i3 | ENSMUSG( | ENSMUSG( | 0.484598 | 0.410705 | NO | NO | NA |
| Col18a1 | ENSMUSG( | ENSMUSG( | -0.13524 | 0.868358 | NO | NO | NA |
| Ttyh1 | ENSMUSG( | ENSMUSG( | 0.177907 | 0.743904 | NO | NO | NA |
| Gm49721 | ENSMUSG( | ENSMUSG( | -0.35315 | 0.874161 | NO | NO | NA |
| Gm13535 | ENSMUSG( | ENSMUSG( | 0.207748 | 0.84853 | NO | NO | NA |
| Vmn2r70 | ENSMUSG( | ENSMUSG( | 0.455745 | 0.658376 | NO | NO | NA |
| Gm45210 | ENSMUSG( | ENSMUSG( | 0.777726 | 0.49586 | NO | NO | NA |
| Gm19391 | ENSMUSG( | ENSMUSG( | -0.50726 | 0.111855 | NO | NO | NA |
| Gm26901 | ENSMUSG( | ENSMUSG( | 0.294836 | 0.772141 | NO | NO | NA |
| Col5a1 | ENSMUSG( | ENSMUSG( | 1.210479 | 0.027371 | YES | YES | Upregulated |
| C130083A | ENSMUSG( | ENSMUSG( | 0.201432 | 0.787833 | NO | NO | NA |
| Olfr912 | ENSMUSG( | ENSMUSG( | 0.431146 | 0.67483 | NO | NO | NA |
| Vmn1r12 | ENSMUSG( | ENSMUSG( | 0.570695 | 0.652335 | NO | NO | NA |
| Gm47372 | ENSMUSG( | ENSMUSG( | 0.544499 | 0.446235 | NO | NO | NA |
| Gm27008 | ENSMUSG( | ENSMUSG( | -0.59047 | 0.095703 | NO | NO | NA |
| Gm43072 | ENSMUSG( | ENSMUSG( | 0.3516 | 0.747285 | NO | NO | NA |
| B930092H | ENSMUSG( | ENSMUSG( | 0.441333 | 0.667184 | NO | NO | NA |
| Gm3720 | ENSMUSG( | ENSMUSG( | 0.639194 | 0.456867 | NO | NO | NA |
| Wsb2-ps | ENSMUSG( | ENSMUSG( | -0.23405 | 0.499353 | NO | NO | NA |

| | | | | | | | |
|---|---|---|---|---|---|---|---|
| Pagr1b | ENSMUSG( | ENSMUSG( | -0.37816 | 0.841948 | NO | NO | NA |
| Arhgef26 | ENSMUSG( | ENSMUSG( | 0.589186 | 0.266887 | NO | NO | NA |
| Gm5466 | ENSMUSG( | ENSMUSG( | 0.461415 | 0.349957 | NO | NO | NA |
| Nlrp1b | ENSMUSG( | ENSMUSG( | -0.85025 | 0.012217 | YES | NO | NA |
| Gm37429 | ENSMUSG( | ENSMUSG( | 0.290585 | 0.8114 | NO | NO | NA |
| Gm43965 | ENSMUSG( | ENSMUSG( | -1.02176 | 0.018872 | YES | YES | Downregulated |
| ENSMUSG( | ENSMUSG( | ENSMUSG( | -2.08657 | 1.65E-07 | YES | YES | Downregulated |
| Uggt2 | ENSMUSG( | ENSMUSG( | 0.891555 | 0.056881 | NO | NO | NA |
| Gm44027 | ENSMUSG( | ENSMUSG( | 0.222323 | 0.792641 | NO | NO | NA |
| Gm6611 | ENSMUSG( | ENSMUSG( | -0.68785 | 0.097769 | NO | NO | NA |
| Gm50221 | ENSMUSG( | ENSMUSG( | 0.780698 | 0.227626 | NO | NO | NA |
| Gm5216 | ENSMUSG( | ENSMUSG( | 1.421048 | 0.180283 | NO | NO | NA |
| Dsc3 | ENSMUSG( | ENSMUSG( | 0.314819 | 0.756401 | NO | NO | NA |
| Gm49784 | ENSMUSG( | ENSMUSG( | 0.556347 | 0.615925 | NO | NO | NA |
| Gm45001 | ENSMUSG( | ENSMUSG( | -0.23473 | 0.479548 | NO | NO | NA |
| Gm43198 | ENSMUSG( | ENSMUSG( | 0.381241 | 0.762497 | NO | NO | NA |
| Hcn3 | ENSMUSG( | ENSMUSG( | 0.28557 | 0.49117 | NO | NO | NA |
| Galnt14 | ENSMUSG( | ENSMUSG( | -0.21813 | 0.67172 | NO | NO | NA |
| Gm28800 | ENSMUSG( | ENSMUSG( | 0.179215 | 0.66428 | NO | NO | NA |
| A830082K | ENSMUSG( | ENSMUSG( | 0.132097 | 0.896053 | NO | NO | NA |
| Gm38236 | ENSMUSG( | ENSMUSG( | 0.986959 | 0.246432 | NO | NO | NA |
| Marchf1 | ENSMUSG( | ENSMUSG( | -0.12857 | 0.857884 | NO | NO | NA |
| Spata48 | ENSMUSG( | ENSMUSG( | 0.613816 | 0.156857 | NO | NO | NA |
| Gm47586 | ENSMUSG( | ENSMUSG( | -0.78129 | 0.02349 | YES | NO | NA |
| Olfr877 | ENSMUSG( | ENSMUSG( | 0.575816 | 0.611854 | NO | NO | NA |
| Gm10635 | ENSMUSG( | ENSMUSG( | 0.383383 | 0.728271 | NO | NO | NA |
| Gm17022 | ENSMUSG( | ENSMUSG( | -0.02352 | 0.942764 | NO | NO | NA |
| Gm30346 | ENSMUSG( | ENSMUSG( | 0.261955 | 0.784011 | NO | NO | NA |
| Gm43523 | ENSMUSG( | ENSMUSG( | 0.68439 | 0.547932 | NO | NO | NA |
| Ly86 | ENSMUSG( | ENSMUSG( | -0.11399 | 0.896055 | NO | NO | NA |
| Gm9894 | ENSMUSG( | ENSMUSG( | 0.739732 | 0.428872 | NO | NO | NA |
| Gm37660 | ENSMUSG( | ENSMUSG( | 0.028153 | 0.947496 | NO | NO | NA |
| D630003M | ENSMUSG( | ENSMUSG( | 0.459245 | 0.696361 | NO | NO | NA |
| Ccr3 | ENSMUSG( | ENSMUSG( | 0.309526 | 0.514562 | NO | NO | NA |
| Gm18035 | ENSMUSG( | ENSMUSG( | 0.533969 | 0.656466 | NO | NO | NA |
| Gm37482 | ENSMUSG( | ENSMUSG( | 1.167183 | 0.25122 | NO | NO | NA |
| Rpap3-ps2 | ENSMUSG( | ENSMUSG( | 0.638959 | 0.553848 | NO | NO | NA |
| Gm43343 | ENSMUSG( | ENSMUSG( | 0.817677 | 0.471485 | NO | NO | NA |
| Gm32036 | ENSMUSG( | ENSMUSG( | 0.88041 | 0.362616 | NO | NO | NA |
| Sycp1-ps1 | ENSMUSG( | ENSMUSG( | 0.596221 | 0.596546 | NO | NO | NA |
| Gad2 | ENSMUSG( | ENSMUSG( | 1.074309 | 0.283023 | NO | NO | NA |
| Gpr165 | ENSMUSG( | ENSMUSG( | 0.782385 | 0.47774 | NO | NO | NA |
| Hmga2-ps: | ENSMUSG( | ENSMUSG( | 0.767686 | 0.028057 | YES | NO | NA |
| Olfr433 | ENSMUSG( | ENSMUSG( | 1.080344 | 0.394559 | NO | NO | NA |
| Fkbp7 | ENSMUSG( | ENSMUSG( | 0.543401 | 0.191644 | NO | NO | NA |
| 4930483P: | ENSMUSG( | ENSMUSG( | 0.862331 | 0.226504 | NO | NO | NA |
| Adamts9 | ENSMUSG( | ENSMUSG( | 0.7814 | 0.376604 | NO | NO | NA |

| | | | | | | | |
|---|---|---|---|---|---|---|---|
| Gm10734 | ENSMUSG( | ENSMUSG( | 0.039035 | 0.949522 | NO | NO | NA |
| Gm15314 | ENSMUSG( | ENSMUSG( | 1.705379 | 0.138812 | NO | NO | NA |
| Gm37100 | ENSMUSG( | ENSMUSG( | 1.130656 | 0.316924 | NO | NO | NA |
| Gm31804 | ENSMUSG( | ENSMUSG( | 0.747949 | 0.459033 | NO | NO | NA |
| Gm26781 | ENSMUSG( | ENSMUSG( | 1.342331 | 0.03799 | YES | YES | Upregulated |
| Gm4841 | ENSMUSG( | ENSMUSG( | 0.591459 | 0.166736 | NO | NO | NA |
| Slc24a3 | ENSMUSG( | ENSMUSG( | 0.551669 | 0.544264 | NO | NO | NA |
| Gm17080 | ENSMUSG( | ENSMUSG( | 0.282816 | 0.439353 | NO | NO | NA |
| Ralyl | ENSMUSG( | ENSMUSG( | 0.319741 | 0.712283 | NO | NO | NA |
| Gm973 | ENSMUSG( | ENSMUSG( | 0.743587 | 0.406768 | NO | NO | NA |
| Gm16551 | ENSMUSG( | ENSMUSG( | 0.741823 | 0.483736 | NO | NO | NA |
| Olfr1254 | ENSMUSG( | ENSMUSG( | 0.772998 | 0.470684 | NO | NO | NA |
| Gm47731 | ENSMUSG( | ENSMUSG( | 1.449905 | 0.168588 | NO | NO | NA |
| Gm7488 | ENSMUSG( | ENSMUSG( | 0.076895 | 0.858518 | NO | NO | NA |
| Esrp2 | ENSMUSG( | ENSMUSG( | 0.291003 | 0.628805 | NO | NO | NA |
| Fbxo36 | ENSMUSG( | ENSMUSG( | 0.962902 | 0.096441 | NO | NO | NA |
| Gm26604 | ENSMUSG( | ENSMUSG( | 0.128731 | 0.915328 | NO | NO | NA |
| Ifit1bl2 | ENSMUSG( | ENSMUSG( | -0.71013 | 0.110155 | NO | NO | NA |
| Gm37584 | ENSMUSG( | ENSMUSG( | -0.00079 | 0.998967 | NO | NO | NA |
| 4921527H | ENSMUSG( | ENSMUSG( | 0.956257 | 0.424923 | NO | NO | NA |
| Gm48133 | ENSMUSG( | ENSMUSG( | -0.0097 | 0.981145 | NO | NO | NA |
| Gm2223 | ENSMUSG( | ENSMUSG( | -0.07805 | 0.868297 | NO | NO | NA |
| 6430548M | ENSMUSG( | ENSMUSG( | 0.201159 | 0.592286 | NO | NO | NA |
| Olfr1423 | ENSMUSG( | ENSMUSG( | 0.68447 | 0.561491 | NO | NO | NA |
| Slc10a4-ps | ENSMUSG( | ENSMUSG( | 0.934604 | 0.327251 | NO | NO | NA |
| 9230116L( | ENSMUSG( | ENSMUSG( | -0.01696 | 0.98075 | NO | NO | NA |
| Piwil1 | ENSMUSG( | ENSMUSG( | 0.725653 | 0.493174 | NO | NO | NA |
| ENSMUSG | ENSMUSG( | ENSMUSG( | 0.412812 | 0.422267 | NO | NO | NA |
| Gm37736 | ENSMUSG( | ENSMUSG( | -0.01512 | 0.988409 | NO | NO | NA |
| 2510016D | ENSMUSG( | ENSMUSG( | -0.08197 | 0.834422 | NO | NO | NA |
| Gm43727 | ENSMUSG( | ENSMUSG( | 0.164774 | 0.865515 | NO | NO | NA |
| Vdr | ENSMUSG( | ENSMUSG( | 0.298072 | 0.665813 | NO | NO | NA |
| Olfr1277 | ENSMUSG( | ENSMUSG( | 0.531731 | 0.598076 | NO | NO | NA |
| Gm7332 | ENSMUSG( | ENSMUSG( | 0.327848 | 0.413943 | NO | NO | NA |
| Gm13036 | ENSMUSG( | ENSMUSG( | 1.621301 | 0.118408 | NO | NO | NA |
| Gm2446 | ENSMUSG( | ENSMUSG( | 0.369312 | 0.678821 | NO | NO | NA |
| Gm16419 | ENSMUSG( | ENSMUSG( | 1.299903 | 0.166283 | NO | NO | NA |
| Gm42941 | ENSMUSG( | ENSMUSG( | -0.77007 | 0.029213 | YES | NO | NA |
| Ltb4r1 | ENSMUSG( | ENSMUSG( | 0.999445 | 0.041635 | YES | NO | NA |
| C5ar1 | ENSMUSG( | ENSMUSG( | 0.125065 | 0.906059 | NO | NO | NA |
| Nphp3 | ENSMUSG( | ENSMUSG( | 1.451447 | 0.082081 | NO | NO | NA |
| Trav7-1 | ENSMUSG( | ENSMUSG( | -0.44974 | 0.624063 | NO | NO | NA |
| Ifngr2 | ENSMUSG( | ENSMUSG( | 0.613794 | 0.379041 | NO | NO | NA |
| Tnni3 | ENSMUSG( | ENSMUSG( | 1.731456 | 0.000904 | YES | YES | Upregulated |
| Gm10752 | ENSMUSG( | ENSMUSG( | 0.678899 | 0.163292 | NO | NO | NA |
| Gm9722 | ENSMUSG( | ENSMUSG( | 0.576161 | 0.560369 | NO | NO | NA |
| Gm30541 | ENSMUSG( | ENSMUSG( | 0.328947 | 0.592908 | NO | NO | NA |

| | | | | | | | |
|---|---|---|---|---|---|---|---|
| Gm49412 | ENSMUSG( | ENSMUSG( | 0.922219 | 0.3959 | NO | NO | NA |
| Gm42514 | ENSMUSG( | ENSMUSG( | 0.505923 | 0.574532 | NO | NO | NA |
| Vmn1r202 | ENSMUSG( | ENSMUSG( | 0.942444 | 0.283669 | NO | NO | NA |
| Acer2 | ENSMUSG( | ENSMUSG( | -0.52147 | 0.152958 | NO | NO | NA |
| Plk-ps1 | ENSMUSG( | ENSMUSG( | -0.25515 | 0.462015 | NO | NO | NA |
| Halr1 | ENSMUSG( | ENSMUSG( | 0.881225 | 0.354923 | NO | NO | NA |
| Slc1a1 | ENSMUSG( | ENSMUSG( | 0.580666 | 0.605484 | NO | NO | NA |
| Fbln7 | ENSMUSG( | ENSMUSG( | 0.36086 | 0.728296 | NO | NO | NA |
| Gm47096 | ENSMUSG( | ENSMUSG( | 0.526559 | 0.48821 | NO | NO | NA |
| E130119H | ENSMUSG( | ENSMUSG( | 0.65278 | 0.550072 | NO | NO | NA |
| Slc4a11 | ENSMUSG( | ENSMUSG( | 0.505315 | 0.493793 | NO | NO | NA |
| Gm49677 | ENSMUSG( | ENSMUSG( | 0.715925 | 0.541669 | NO | NO | NA |
| Vmn1r24 | ENSMUSG( | ENSMUSG( | 0.621472 | 0.560577 | NO | NO | NA |
| Rragd | ENSMUSG( | ENSMUSG( | 1.075176 | 0.076741 | NO | NO | NA |
| Gm6116 | ENSMUSG( | ENSMUSG( | 1.011147 | 0.324777 | NO | NO | NA |
| Gm20682 | ENSMUSG( | ENSMUSG( | 0.83561 | 0.319818 | NO | NO | NA |
| 2610028D | ENSMUSG( | ENSMUSG( | 0.717458 | 0.117426 | NO | NO | NA |
| Parm1 | ENSMUSG( | ENSMUSG( | 0.543423 | 0.567401 | NO | NO | NA |
| Gm47911 | ENSMUSG( | ENSMUSG( | 0.783807 | 0.471567 | NO | NO | NA |
| Padi1 | ENSMUSG( | ENSMUSG( | -0.15149 | 0.699975 | NO | NO | NA |
| Gm42519 | ENSMUSG( | ENSMUSG( | 0.187776 | 0.851529 | NO | NO | NA |
| Gm21691 | ENSMUSG( | ENSMUSG( | -0.23546 | 0.805944 | NO | NO | NA |
| Olfr458 | ENSMUSG( | ENSMUSG( | 1.073487 | 0.389754 | NO | NO | NA |
| Tas1r1 | ENSMUSG( | ENSMUSG( | 1.267876 | 0.016121 | YES | YES | Upregulated |
| Rxfp1 | ENSMUSG( | ENSMUSG( | 0.679569 | 0.512662 | NO | NO | NA |
| Il7 | ENSMUSG( | ENSMUSG( | 0.060567 | 0.902993 | NO | NO | NA |
| Olfr1508 | ENSMUSG( | ENSMUSG( | 0.371041 | 0.734336 | NO | NO | NA |
| Ccdc73 | ENSMUSG( | ENSMUSG( | -0.27671 | 0.404243 | NO | NO | NA |
| 5330439K( | ENSMUSG( | ENSMUSG( | 0.70849 | 0.423913 | NO | NO | NA |
| Lin28a | ENSMUSG( | ENSMUSG( | 0.458515 | 0.668604 | NO | NO | NA |
| Gm43648 | ENSMUSG( | ENSMUSG( | 0.724427 | 0.537219 | NO | NO | NA |
| Gm4668 | ENSMUSG( | ENSMUSG( | -0.83418 | 0.034445 | YES | NO | NA |
| 5830454E( | ENSMUSG( | ENSMUSG( | 0.223231 | 0.564931 | NO | NO | NA |
| Acot4 | ENSMUSG( | ENSMUSG( | 0.508435 | 0.545839 | NO | NO | NA |
| Ifitm1 | ENSMUSG( | ENSMUSG( | 0.953993 | 0.234786 | NO | NO | NA |
| D230049E | ENSMUSG( | ENSMUSG( | -0.12696 | 0.831255 | NO | NO | NA |
| Ctsh | ENSMUSG( | ENSMUSG( | 0.243299 | 0.675542 | NO | NO | NA |
| A530064D | ENSMUSG( | ENSMUSG( | 0.367787 | 0.717725 | NO | NO | NA |
| ENSMUSG( | ENSMUSG( | ENSMUSG( | 0.656611 | 0.530589 | NO | NO | NA |
| Gpc2 | ENSMUSG( | ENSMUSG( | -0.13612 | 0.696353 | NO | NO | NA |
| ENSMUSG | ENSMUSG( | ENSMUSG( | 0.213398 | 0.537302 | NO | NO | NA |
| Tnfrsf12a | ENSMUSG( | ENSMUSG( | -0.09883 | 0.801651 | NO | NO | NA |
| Gm20953 | ENSMUSG( | ENSMUSG( | 0.452725 | 0.659678 | NO | NO | NA |
| Tmem98 | ENSMUSG( | ENSMUSG( | 0.150238 | 0.754448 | NO | NO | NA |
| Rdh9 | ENSMUSG( | ENSMUSG( | -0.05976 | 0.971982 | NO | NO | NA |
| Gm6093 | ENSMUSG( | ENSMUSG( | 2.036105 | 0.01523 | YES | YES | Upregulated |
| Dok5 | ENSMUSG( | ENSMUSG( | 0.866823 | 0.444105 | NO | NO | NA |

| | | | | | | | |
|---|---|---|---|---|---|---|---|
| Gm40332 | ENSMUSG( | ENSMUSG( | 0.721956 | 0.533092 | NO | NO | NA |
| Gm47418 | ENSMUSG( | ENSMUSG( | -0.32564 | 0.56435 | NO | NO | NA |
| Klra5 | ENSMUSG( | ENSMUSG( | 1.113491 | NA | NO | NO | NA |
| Gm6133 | ENSMUSG( | ENSMUSG( | 0.489277 | 0.566439 | NO | NO | NA |
| Glra1 | ENSMUSG( | ENSMUSG( | 0.500185 | 0.633949 | NO | NO | NA |
| Ache | ENSMUSG( | ENSMUSG( | 0.983409 | 0.10893 | NO | NO | NA |
| Fmnl2 | ENSMUSG( | ENSMUSG( | -0.35016 | 0.507429 | NO | NO | NA |
| 9430037O | ENSMUSG( | ENSMUSG( | -0.06723 | 0.899627 | NO | NO | NA |
| ENSMUSG | ENSMUSG( | ENSMUSG( | -0.32062 | 0.507304 | NO | NO | NA |
| Ccpg1os | ENSMUSG( | ENSMUSG( | -0.53873 | 0.185287 | NO | NO | NA |
| ENSMUSG | ENSMUSG( | ENSMUSG( | 0.65334 | 0.318625 | NO | NO | NA |
| Gm14965 | ENSMUSG( | ENSMUSG( | 0.102968 | 0.939243 | NO | NO | NA |
| Gm47147 | ENSMUSG( | ENSMUSG( | -0.31057 | 0.460937 | NO | NO | NA |
| 0610009E( | ENSMUSG( | ENSMUSG( | -0.5197 | 0.167266 | NO | NO | NA |
| Olfr1316 | ENSMUSG( | ENSMUSG( | 0.284726 | 0.78906 | NO | NO | NA |
| Lum | ENSMUSG( | ENSMUSG( | 0.899261 | 0.411282 | NO | NO | NA |
| Kcnj16 | ENSMUSG( | ENSMUSG( | 1.158087 | 0.221861 | NO | NO | NA |
| Gm14321 | ENSMUSG( | ENSMUSG( | 0.404279 | 0.50265 | NO | NO | NA |
| Igkv8-27 | ENSMUSG( | ENSMUSG( | -1.28008 | 0.650641 | NO | NO | NA |
| Rnft2 | ENSMUSG( | ENSMUSG( | 1.831571 | 0.006963 | YES | YES | Upregulated |
| Slc18a1 | ENSMUSG( | ENSMUSG( | 0.506584 | 0.592991 | NO | NO | NA |
| Gm38297 | ENSMUSG( | ENSMUSG( | 0.691413 | 0.500681 | NO | NO | NA |
| ENSMUSG( | ENSMUSG( | ENSMUSG( | 0.265854 | 0.587117 | NO | NO | NA |
| Gm45141 | ENSMUSG( | ENSMUSG( | -0.35289 | 0.516629 | NO | NO | NA |
| Spic | ENSMUSG( | ENSMUSG( | -0.9349 | 0.054903 | NO | NO | NA |
| Gm43486 | ENSMUSG( | ENSMUSG( | 1.084702 | 0.281082 | NO | NO | NA |
| Tlr4 | ENSMUSG( | ENSMUSG( | 0.281924 | 0.568223 | NO | NO | NA |
| Gm47208 | ENSMUSG( | ENSMUSG( | 0.271795 | 0.785998 | NO | NO | NA |
| Muc6 | ENSMUSG( | ENSMUSG( | -0.14001 | 0.834422 | NO | NO | NA |
| Large1 | ENSMUSG( | ENSMUSG( | 0.948139 | 0.064685 | NO | NO | NA |
| Gm7389 | ENSMUSG( | ENSMUSG( | 0.571062 | 0.49519 | NO | NO | NA |
| Gm48285 | ENSMUSG( | ENSMUSG( | 0.162917 | 0.881884 | NO | NO | NA |
| Gm42879 | ENSMUSG( | ENSMUSG( | 0.97295 | 0.291045 | NO | NO | NA |
| Nrcam | ENSMUSG( | ENSMUSG( | 0.975423 | 0.247637 | NO | NO | NA |
| Capn12 | ENSMUSG( | ENSMUSG( | -0.12821 | 0.795374 | NO | NO | NA |
| Gm12696 | ENSMUSG( | ENSMUSG( | 0.036923 | 0.929734 | NO | NO | NA |
| Rbm11 | ENSMUSG( | ENSMUSG( | 0.259327 | 0.640236 | NO | NO | NA |
| Gm38124 | ENSMUSG( | ENSMUSG( | 0.956473 | 0.382461 | NO | NO | NA |
| Gm41492 | ENSMUSG( | ENSMUSG( | -0.12456 | 0.9084 | NO | NO | NA |
| Zfp964 | ENSMUSG( | ENSMUSG( | 0.295635 | 0.662264 | NO | NO | NA |
| Btk | ENSMUSG( | ENSMUSG( | 0.05324 | 0.946243 | NO | NO | NA |
| Gm37755 | ENSMUSG( | ENSMUSG( | 0.619249 | 0.396841 | NO | NO | NA |
| Gm10522 | ENSMUSG( | ENSMUSG( | -0.72577 | 0.072544 | NO | NO | NA |
| Gm48639 | ENSMUSG( | ENSMUSG( | 0.422353 | 0.644809 | NO | NO | NA |
| 4930573C( | ENSMUSG( | ENSMUSG( | 1.100454 | 0.312909 | NO | NO | NA |
| Slc18b1 | ENSMUSG( | ENSMUSG( | 0.857552 | 0.465973 | NO | NO | NA |
| Klhl32 | ENSMUSG( | ENSMUSG( | 0.605368 | 0.331659 | NO | NO | NA |

| | | | | | | | |
|---|---|---|---|---|---|---|---|
| Ptprn2 | ENSMUSG( | ENSMUSG( | 0.09499 | 0.92738 | NO | NO | NA |
| Gm5524 | ENSMUSG( | ENSMUSG( | 1.289804 | 0.228198 | NO | NO | NA |
| Gm42930 | ENSMUSG( | ENSMUSG( | 0.234925 | 0.722714 | NO | NO | NA |
| Gm43360 | ENSMUSG( | ENSMUSG( | 0.153452 | 0.681123 | NO | NO | NA |
| C4a | ENSMUSG( | ENSMUSG( | 0.246442 | 0.489576 | NO | NO | NA |
| Gm48798 | ENSMUSG( | ENSMUSG( | 0.125572 | 0.91234 | NO | NO | NA |
| Ankrd6 | ENSMUSG( | ENSMUSG( | 0.384916 | 0.408636 | NO | NO | NA |
| Gm29151 | ENSMUSG( | ENSMUSG( | 0.450252 | 0.506969 | NO | NO | NA |
| Abca5 | ENSMUSG( | ENSMUSG( | 1.026446 | 0.02687 | YES | YES | Upregulated |
| Gm11839 | ENSMUSG( | ENSMUSG( | 0.050176 | 0.948899 | NO | NO | NA |
| 6430590A( | ENSMUSG( | ENSMUSG( | -0.34201 | 0.332979 | NO | NO | NA |
| Olfr750 | ENSMUSG( | ENSMUSG( | 0.552613 | 0.650295 | NO | NO | NA |
| Gm37022 | ENSMUSG( | ENSMUSG( | 1.013014 | 0.370733 | NO | NO | NA |
| Eva1c | ENSMUSG( | ENSMUSG( | 1.371007 | 0.100639 | NO | NO | NA |
| Gm50336 | ENSMUSG( | ENSMUSG( | 0.222542 | 0.633247 | NO | NO | NA |
| Gm8036 | ENSMUSG( | ENSMUSG( | 1.069425 | 0.346493 | NO | NO | NA |
| Gm18273 | ENSMUSG( | ENSMUSG( | 0.89449 | 0.354826 | NO | NO | NA |
| Gm6883 | ENSMUSG( | ENSMUSG( | 0.515701 | 0.472908 | NO | NO | NA |
| Adamts3 | ENSMUSG( | ENSMUSG( | 1.004203 | 0.203268 | NO | NO | NA |
| Gm37537 | ENSMUSG( | ENSMUSG( | 0.175594 | 0.838522 | NO | NO | NA |
| Gm26590 | ENSMUSG( | ENSMUSG( | 0.071421 | 0.899918 | NO | NO | NA |
| Gm4013 | ENSMUSG( | ENSMUSG( | 0.755643 | 0.047878 | YES | NO | NA |
| Ptn | ENSMUSG( | ENSMUSG( | 0.504143 | 0.650468 | NO | NO | NA |
| Gm28068 | ENSMUSG( | ENSMUSG( | 0.606497 | 0.466943 | NO | NO | NA |
| Gm37566 | ENSMUSG( | ENSMUSG( | 0.561243 | 0.453271 | NO | NO | NA |
| Pdcd5-ps | ENSMUSG( | ENSMUSG( | 0.343358 | 0.30875 | NO | NO | NA |
| Pygl | ENSMUSG( | ENSMUSG( | 0.181785 | 0.699423 | NO | NO | NA |
| Gm47041 | ENSMUSG( | ENSMUSG( | 0.950155 | 0.372745 | NO | NO | NA |
| Gm37249 | ENSMUSG( | ENSMUSG( | 0.831206 | 0.178773 | NO | NO | NA |
| D430018E( | ENSMUSG( | ENSMUSG( | -0.34794 | 0.560734 | NO | NO | NA |
| Rimbp2 | ENSMUSG( | ENSMUSG( | 0.222829 | 0.819022 | NO | NO | NA |
| Gm2244 | ENSMUSG( | ENSMUSG( | 0.409156 | 0.653625 | NO | NO | NA |
| Gm45890 | ENSMUSG( | ENSMUSG( | 0.425724 | 0.513739 | NO | NO | NA |
| A930027P( | ENSMUSG( | ENSMUSG( | 0.591985 | 0.578811 | NO | NO | NA |
| Olfr5 | ENSMUSG( | ENSMUSG( | 0.832079 | 0.479158 | NO | NO | NA |
| Scube3 | ENSMUSG( | ENSMUSG( | 0.686813 | 0.503339 | NO | NO | NA |
| Gm10359 | ENSMUSG( | ENSMUSG( | 5.64426 | 0.124094 | NO | NO | NA |
| Gm42841 | ENSMUSG( | ENSMUSG( | 0.806023 | 0.462756 | NO | NO | NA |
| Gm17024 | ENSMUSG( | ENSMUSG( | 0.466596 | 0.267916 | NO | NO | NA |
| Zfr2 | ENSMUSG( | ENSMUSG( | -0.55373 | 0.171822 | NO | NO | NA |
| Gm45807 | ENSMUSG( | ENSMUSG( | 0.253346 | 0.745326 | NO | NO | NA |
| Gm44616 | ENSMUSG( | ENSMUSG( | 0.011914 | 0.975259 | NO | NO | NA |
| Tns4 | ENSMUSG( | ENSMUSG( | 0.175373 | 0.874765 | NO | NO | NA |
| AI463229 | ENSMUSG( | ENSMUSG( | 0.299742 | 0.736727 | NO | NO | NA |
| Krt77 | ENSMUSG( | ENSMUSG( | 1.11561 | 0.37594 | NO | NO | NA |
| Gm42576 | ENSMUSG( | ENSMUSG( | -0.88414 | 0.079105 | NO | NO | NA |
| Gm38125 | ENSMUSG( | ENSMUSG( | 0.398106 | 0.742703 | NO | NO | NA |

| | | | | | | | |
|---|---|---|---|---|---|---|---|
| Sh3tc2 | ENSMUSG( | ENSMUSG( | 0.510224 | 0.481811 | NO | NO | NA |
| Gm47861 | ENSMUSG( | ENSMUSG( | -0.02435 | 0.976023 | NO | NO | NA |
| Gm47806 | ENSMUSG( | ENSMUSG( | -0.20053 | 0.755585 | NO | NO | NA |
| Gm13546 | ENSMUSG( | ENSMUSG( | 1.742343 | 0.000593 | YES | YES | Upregulated |
| St8sia1 | ENSMUSG( | ENSMUSG( | 0.661805 | 0.256641 | NO | NO | NA |
| Trbj2-1 | ENSMUSG( | ENSMUSG( | 0.285831 | 0.728369 | NO | NO | NA |
| D830013O | ENSMUSG( | ENSMUSG( | 0.234827 | 0.819148 | NO | NO | NA |
| Gm16083 | ENSMUSG( | ENSMUSG( | 0.267613 | 0.713191 | NO | NO | NA |
| Gm20371 | ENSMUSG( | ENSMUSG( | 0.326998 | 0.785421 | NO | NO | NA |
| Tcim | ENSMUSG( | ENSMUSG( | 0.31797 | 0.748184 | NO | NO | NA |
| Lmo3 | ENSMUSG( | ENSMUSG( | 0.241325 | 0.835562 | NO | NO | NA |
| Ces5a | ENSMUSG( | ENSMUSG( | 0.340943 | 0.789276 | NO | NO | NA |
| BB187690 | ENSMUSG( | ENSMUSG( | 0.951469 | 0.383051 | NO | NO | NA |
| Il19 | ENSMUSG( | ENSMUSG( | 0.648376 | 0.541473 | NO | NO | NA |
| 1810014P( | ENSMUSG( | ENSMUSG( | 0.730705 | 0.558429 | NO | NO | NA |
| Mcam | ENSMUSG( | ENSMUSG( | -0.74244 | 0.032189 | YES | NO | NA |
| 4732463B( | ENSMUSG( | ENSMUSG( | 1.167899 | 0.227425 | NO | NO | NA |
| Rbfox2 | ENSMUSG( | ENSMUSG( | 0.588811 | 0.446441 | NO | NO | NA |
| Gm44237 | ENSMUSG( | ENSMUSG( | 0.827609 | 0.385467 | NO | NO | NA |
| Atp10b | ENSMUSG( | ENSMUSG( | 0.610644 | 0.495589 | NO | NO | NA |
| Nipa1 | ENSMUSG( | ENSMUSG( | -0.09742 | 0.892935 | NO | NO | NA |
| Wnt10a | ENSMUSG( | ENSMUSG( | 2.362469 | 0.003913 | YES | YES | Upregulated |
| Dbndd1 | ENSMUSG( | ENSMUSG( | -0.02849 | 0.944123 | NO | NO | NA |
| Lrrc4b | ENSMUSG( | ENSMUSG( | 0.64708 | 0.555025 | NO | NO | NA |
| Lingo4 | ENSMUSG( | ENSMUSG( | 0.267928 | 0.80634 | NO | NO | NA |
| Tmem141 | ENSMUSG( | ENSMUSG( | 0.821105 | 0.319978 | NO | NO | NA |
| 5830444F: | ENSMUSG( | ENSMUSG( | 0.813208 | 0.247168 | NO | NO | NA |
| Gm50206 | ENSMUSG( | ENSMUSG( | 0.958338 | 0.450894 | NO | NO | NA |
| Gm37593 | ENSMUSG( | ENSMUSG( | 0.31809 | 0.601687 | NO | NO | NA |
| Fmn1 | ENSMUSG( | ENSMUSG( | 0.420461 | 0.682397 | NO | NO | NA |
| Igkv4-68 | ENSMUSG( | ENSMUSG( | -2.54334 | NA | NO | NO | NA |
| Vax2os | ENSMUSG( | ENSMUSG( | 0.340898 | 0.523069 | NO | NO | NA |
| Gm29707 | ENSMUSG( | ENSMUSG( | 1.26162 | 0.273337 | NO | NO | NA |
| Gm15433 | ENSMUSG( | ENSMUSG( | -0.55413 | 0.447795 | NO | NO | NA |
| Fam43a | ENSMUSG( | ENSMUSG( | -0.1183 | 0.788575 | NO | NO | NA |
| Gm45792 | ENSMUSG( | ENSMUSG( | 0.300202 | 0.768587 | NO | NO | NA |
| Gm9577 | ENSMUSG( | ENSMUSG( | 0.780681 | 0.475513 | NO | NO | NA |
| Rgn | ENSMUSG( | ENSMUSG( | 0.956843 | 0.304251 | NO | NO | NA |
| Olfr1109 | ENSMUSG( | ENSMUSG( | 0.262656 | 0.811732 | NO | NO | NA |
| Gm43275 | ENSMUSG( | ENSMUSG( | 0.441937 | 0.675824 | NO | NO | NA |
| Lipg | ENSMUSG( | ENSMUSG( | 0.775611 | 0.41724 | NO | NO | NA |
| Zswim5 | ENSMUSG( | ENSMUSG( | 0.346863 | 0.46306 | NO | NO | NA |
| Gm45540 | ENSMUSG( | ENSMUSG( | 0.910689 | 0.346493 | NO | NO | NA |
| Gm48706 | ENSMUSG( | ENSMUSG( | 0.207586 | 0.834874 | NO | NO | NA |
| Grm2 | ENSMUSG( | ENSMUSG( | 0.468532 | 0.654136 | NO | NO | NA |
| Il23a | ENSMUSG( | ENSMUSG( | 0.73775 | 0.442003 | NO | NO | NA |
| Gm47197 | ENSMUSG( | ENSMUSG( | 0.807477 | 0.421196 | NO | NO | NA |

| | | | | | | | |
|---|---|---|---|---|---|---|---|
| Gm46209 | ENSMUSG( | ENSMUSG( | -0.06916 | 0.899627 | NO | NO | NA |
| Gm37199 | ENSMUSG( | ENSMUSG( | 0.576415 | 0.611373 | NO | NO | NA |
| Olfr518 | ENSMUSG( | ENSMUSG( | 0.764531 | 0.433772 | NO | NO | NA |
| Gm5362 | ENSMUSG( | ENSMUSG( | 0.414223 | 0.667108 | NO | NO | NA |
| Olfr1226 | ENSMUSG( | ENSMUSG( | 0.632905 | 0.557591 | NO | NO | NA |
| Gm35256 | ENSMUSG( | ENSMUSG( | 0.643999 | 0.601687 | NO | NO | NA |
| Cldn11 | ENSMUSG( | ENSMUSG( | 0.943739 | 0.406407 | NO | NO | NA |
| H3c2 | ENSMUSG( | ENSMUSG( | 0.430094 | 0.366033 | NO | NO | NA |
| Gm47863 | ENSMUSG( | ENSMUSG( | -0.19075 | 0.60683 | NO | NO | NA |
| A130051J( | ENSMUSG( | ENSMUSG( | 0.450809 | 0.282727 | NO | NO | NA |
| Gm10048 | ENSMUSG( | ENSMUSG( | 0.367623 | 0.724236 | NO | NO | NA |
| ENSMUSG | ENSMUSG( | ENSMUSG( | -0.03248 | 0.936219 | NO | NO | NA |
| Unc13c | ENSMUSG( | ENSMUSG( | 0.481607 | 0.69132 | NO | NO | NA |
| Csn1s2a | ENSMUSG( | ENSMUSG( | 1.057052 | 0.341233 | NO | NO | NA |
| 6330415G | ENSMUSG( | ENSMUSG( | 0.29764 | 0.407211 | NO | NO | NA |
| Gm37790 | ENSMUSG( | ENSMUSG( | 0.7111 | 0.104976 | NO | NO | NA |
| Cdh6 | ENSMUSG( | ENSMUSG( | 0.453475 | 0.664659 | NO | NO | NA |
| Gm31834 | ENSMUSG( | ENSMUSG( | -0.17314 | 0.597056 | NO | NO | NA |
| Syn3 | ENSMUSG( | ENSMUSG( | 0.410555 | 0.665321 | NO | NO | NA |
| Gstk1 | ENSMUSG( | ENSMUSG( | 0.172741 | 0.730991 | NO | NO | NA |
| Pm20d1 | ENSMUSG( | ENSMUSG( | 0.336704 | 0.767661 | NO | NO | NA |
| Olfr299 | ENSMUSG( | ENSMUSG( | 0.400254 | 0.761353 | NO | NO | NA |
| ENSMUSG( | ENSMUSG( | ENSMUSG( | -0.38783 | 0.686444 | NO | NO | NA |
| ENSMUSG | ENSMUSG( | ENSMUSG( | -0.08662 | 0.813465 | NO | NO | NA |
| Gm34121 | ENSMUSG( | ENSMUSG( | 0.138466 | 0.809815 | NO | NO | NA |
| Gm15764 | ENSMUSG( | ENSMUSG( | -0.25789 | 0.63616 | NO | NO | NA |
| ENSMUSG( | ENSMUSG( | ENSMUSG( | -0.44677 | 0.185221 | NO | NO | NA |
| Gm18719 | ENSMUSG( | ENSMUSG( | 0.456752 | 0.696361 | NO | NO | NA |
| Vmn1r229 | ENSMUSG( | ENSMUSG( | 0.34753 | 0.754962 | NO | NO | NA |
| 4933433G | ENSMUSG( | ENSMUSG( | 0.14369 | 0.727799 | NO | NO | NA |
| E230016M | ENSMUSG( | ENSMUSG( | 0.499707 | 0.168709 | NO | NO | NA |
| Vmn1r60 | ENSMUSG( | ENSMUSG( | 0.837316 | 0.414058 | NO | NO | NA |
| ENSMUSG( | ENSMUSG( | ENSMUSG( | -0.01921 | 0.970821 | NO | NO | NA |
| Ighv1-72 | ENSMUSG( | ENSMUSG( | -4.77278 | 0.036644 | YES | YES | Downregulated |
| Crb1 | ENSMUSG( | ENSMUSG( | 0.462759 | 0.670522 | NO | NO | NA |
| Gm17021 | ENSMUSG( | ENSMUSG( | 0.14664 | 0.827588 | NO | NO | NA |
| ENSMUSG | ENSMUSG( | ENSMUSG( | 0.255072 | 0.705422 | NO | NO | NA |
| Npas2 | ENSMUSG( | ENSMUSG( | 0.088416 | 0.925531 | NO | NO | NA |
| Lipt2 | ENSMUSG( | ENSMUSG( | 1.21306 | 0.00709 | YES | YES | Upregulated |
| Ccdc78 | ENSMUSG( | ENSMUSG( | 0.42386 | 0.423382 | NO | NO | NA |
| Gm38292 | ENSMUSG( | ENSMUSG( | -0.03576 | 0.956989 | NO | NO | NA |
| Hp | ENSMUSG( | ENSMUSG( | 0.55224 | 0.559175 | NO | NO | NA |
| Atp13a4 | ENSMUSG( | ENSMUSG( | 0.933149 | 0.400018 | NO | NO | NA |
| Gm47643 | ENSMUSG( | ENSMUSG( | 0.310895 | 0.672948 | NO | NO | NA |
| Gm49207 | ENSMUSG( | ENSMUSG( | -0.46559 | 0.48269 | NO | NO | NA |
| Gm9774 | ENSMUSG( | ENSMUSG( | 0.132602 | 0.670251 | NO | NO | NA |
| Calhm4 | ENSMUSG( | ENSMUSG( | 0.758144 | 0.461948 | NO | NO | NA |

| | | | | | | | |
|---|---|---|---|---|---|---|---|
| Gm45205 | ENSMUSG( | ENSMUSG( | -0.27763 | 0.600373 | NO | NO | NA |
| Spa17 | ENSMUSG( | ENSMUSG( | 1.349382 | 0.005822 | YES | YES | Upregulated |
| Gm32742 | ENSMUSG( | ENSMUSG( | 0.760132 | 0.500632 | NO | NO | NA |
| Aoah | ENSMUSG( | ENSMUSG( | 1.225212 | 0.030763 | YES | YES | Upregulated |
| Gm45709 | ENSMUSG( | ENSMUSG( | -0.107 | 0.763528 | NO | NO | NA |
| Gm27042 | ENSMUSG( | ENSMUSG( | 0.063802 | 0.863987 | NO | NO | NA |
| ENSMUSG | ENSMUSG( | ENSMUSG( | 0.666098 | 0.256547 | NO | NO | NA |
| ENSMUSG | ENSMUSG( | ENSMUSG( | -0.36288 | 0.52822 | NO | NO | NA |
| Gm45234 | ENSMUSG( | ENSMUSG( | 5.356162 | NA | NO | NO | NA |
| Cc2d2a | ENSMUSG( | ENSMUSG( | -0.53371 | 0.476311 | NO | NO | NA |
| Pla2r1 | ENSMUSG( | ENSMUSG( | 0.998252 | 0.333525 | NO | NO | NA |
| Ighv1-39 | ENSMUSG( | ENSMUSG( | 0.066884 | 0.978039 | NO | NO | NA |
| Cbs | ENSMUSG( | ENSMUSG( | 0.31547 | 0.682983 | NO | NO | NA |
| Brinp3 | ENSMUSG( | ENSMUSG( | 0.819554 | 0.37692 | NO | NO | NA |
| 3300002I0 | ENSMUSG( | ENSMUSG( | -0.25015 | 0.516667 | NO | NO | NA |
| Gm43027 | ENSMUSG( | ENSMUSG( | 1.298514 | 0.254949 | NO | NO | NA |
| Gm28651 | ENSMUSG( | ENSMUSG( | 0.497261 | 0.653348 | NO | NO | NA |
| Rem2 | ENSMUSG( | ENSMUSG( | 0.746441 | 0.020508 | YES | NO | NA |
| Gm32027 | ENSMUSG( | ENSMUSG( | 0.439323 | 0.558429 | NO | NO | NA |
| Usp50 | ENSMUSG( | ENSMUSG( | 0.42561 | 0.520079 | NO | NO | NA |
| Wasf3 | ENSMUSG( | ENSMUSG( | 0.72206 | 0.580206 | NO | NO | NA |
| Gm16081 | ENSMUSG( | ENSMUSG( | 1.129485 | 0.283669 | NO | NO | NA |
| ENSMUSG | ENSMUSG( | ENSMUSG( | 1.009418 | 0.324455 | NO | NO | NA |
| B230303O | ENSMUSG( | ENSMUSG( | 0.249267 | 0.716426 | NO | NO | NA |
| AA386476 | ENSMUSG( | ENSMUSG( | 0.310953 | 0.468284 | NO | NO | NA |
| 2410018L1 | ENSMUSG( | ENSMUSG( | -0.01102 | 0.984354 | NO | NO | NA |
| Olfr1490 | ENSMUSG( | ENSMUSG( | 0.756885 | 0.472117 | NO | NO | NA |
| Alx1 | ENSMUSG( | ENSMUSG( | 1.039827 | 0.327368 | NO | NO | NA |
| Gm44639 | ENSMUSG( | ENSMUSG( | 0.093934 | 0.859542 | NO | NO | NA |
| Ppara | ENSMUSG( | ENSMUSG( | 0.882009 | 0.374884 | NO | NO | NA |
| Gm43629 | ENSMUSG( | ENSMUSG( | 1.704996 | 0.077828 | NO | NO | NA |
| Gm29455 | ENSMUSG( | ENSMUSG( | 0.685014 | 0.490843 | NO | NO | NA |
| Wasf1 | ENSMUSG( | ENSMUSG( | 0.6755 | 0.416217 | NO | NO | NA |
| Gm43032 | ENSMUSG( | ENSMUSG( | 1.404572 | 0.216209 | NO | NO | NA |
| Gm47838 | ENSMUSG( | ENSMUSG( | 0.451713 | 0.681623 | NO | NO | NA |
| Pate2 | ENSMUSG( | ENSMUSG( | 0.712621 | 0.440958 | NO | NO | NA |
| Klra6 | ENSMUSG( | ENSMUSG( | 0.303979 | 0.626178 | NO | NO | NA |
| H3c4 | ENSMUSG( | ENSMUSG( | 0.296623 | 0.571421 | NO | NO | NA |
| Dlx5 | ENSMUSG( | ENSMUSG( | 1.059642 | 0.339968 | NO | NO | NA |
| Gm50402 | ENSMUSG( | ENSMUSG( | 0.130236 | 0.84357 | NO | NO | NA |
| Olfr959 | ENSMUSG( | ENSMUSG( | 0.832829 | 0.458194 | NO | NO | NA |
| Gm33027 | ENSMUSG( | ENSMUSG( | 0.879762 | 0.441418 | NO | NO | NA |
| Gm20712 | ENSMUSG( | ENSMUSG( | 0.372519 | 0.484883 | NO | NO | NA |
| Plbd1 | ENSMUSG( | ENSMUSG( | -1.42078 | 0.007975 | YES | YES | Downregulated |
| Gm42699 | ENSMUSG( | ENSMUSG( | 1.079484 | 0.263914 | NO | NO | NA |
| Chrm3 | ENSMUSG( | ENSMUSG( | 0.87772 | 0.392394 | NO | NO | NA |
| Tmc7 | ENSMUSG( | ENSMUSG( | 0.377346 | 0.729736 | NO | NO | NA |

| | | | | | | | |
|---|---|---|---|---|---|---|---|
| Aloxe3 | ENSMUSG( | ENSMUSG( | 0.473602 | 0.622267 | NO | NO | NA |
| 1700097N | ENSMUSG( | ENSMUSG( | 0.768406 | 0.435313 | NO | NO | NA |
| Mycn | ENSMUSG( | ENSMUSG( | 1.244828 | 0.015051 | YES | YES | Upregulated |
| Ppp1r18os | ENSMUSG( | ENSMUSG( | -0.10687 | 0.83236 | NO | NO | NA |
| Spata6l | ENSMUSG( | ENSMUSG( | 0.049953 | 0.929502 | NO | NO | NA |
| Cbx3-ps6 | ENSMUSG( | ENSMUSG( | 0.137924 | 0.706876 | NO | NO | NA |
| Gm42743 | ENSMUSG( | ENSMUSG( | 0.298135 | 0.439849 | NO | NO | NA |
| Aox1 | ENSMUSG( | ENSMUSG( | 0.641568 | 0.491395 | NO | NO | NA |
| Gm13293 | ENSMUSG( | ENSMUSG( | -0.27527 | 0.51162 | NO | NO | NA |
| Gm47513 | ENSMUSG( | ENSMUSG( | -0.32042 | 0.577972 | NO | NO | NA |
| ENSMUSG | ENSMUSG( | ENSMUSG( | 0.543552 | 0.200743 | NO | NO | NA |
| Gm17791 | ENSMUSG( | ENSMUSG( | 0.887543 | 0.480291 | NO | NO | NA |
| Saxo1 | ENSMUSG( | ENSMUSG( | 0.818267 | 0.458372 | NO | NO | NA |
| Gm26761 | ENSMUSG( | ENSMUSG( | 0.834587 | 0.461523 | NO | NO | NA |
| 2810030D | ENSMUSG( | ENSMUSG( | 0.612673 | 0.556286 | NO | NO | NA |
| Gm13378 | ENSMUSG( | ENSMUSG( | 0.18785 | 0.910588 | NO | NO | NA |
| Frs3 | ENSMUSG( | ENSMUSG( | 0.210039 | 0.668205 | NO | NO | NA |
| Olfr1279 | ENSMUSG( | ENSMUSG( | 0.616927 | 0.562892 | NO | NO | NA |
| Gm38376 | ENSMUSG( | ENSMUSG( | 0.323067 | 0.539237 | NO | NO | NA |
| Snn | ENSMUSG( | ENSMUSG( | 0.033516 | 0.934291 | NO | NO | NA |
| Gm28041 | ENSMUSG( | ENSMUSG( | -0.87137 | 0.168307 | NO | NO | NA |
| Skint3 | ENSMUSG( | ENSMUSG( | 0.780412 | 0.460383 | NO | NO | NA |
| Gm6012 | ENSMUSG( | ENSMUSG( | -0.22175 | 0.604949 | NO | NO | NA |
| ENSMUSG | ENSMUSG( | ENSMUSG( | 0.575097 | 0.616746 | NO | NO | NA |
| Tceal8 | ENSMUSG( | ENSMUSG( | 0.623661 | 0.293345 | NO | NO | NA |
| Gm48878 | ENSMUSG( | ENSMUSG( | 0.401172 | 0.741426 | NO | NO | NA |
| Gm42940 | ENSMUSG( | ENSMUSG( | 0.102639 | 0.797665 | NO | NO | NA |
| Gm43469 | ENSMUSG( | ENSMUSG( | 0.553077 | 0.593014 | NO | NO | NA |
| Gpr35 | ENSMUSG( | ENSMUSG( | 0.033449 | 0.973297 | NO | NO | NA |
| Olfr1385 | ENSMUSG( | ENSMUSG( | 0.500025 | 0.6524 | NO | NO | NA |
| Gm12020 | ENSMUSG( | ENSMUSG( | 0.787729 | 0.254604 | NO | NO | NA |
| Gm42913 | ENSMUSG( | ENSMUSG( | -0.10957 | 0.7985 | NO | NO | NA |
| Olfr715 | ENSMUSG( | ENSMUSG( | 1.01758 | 0.215384 | NO | NO | NA |
| Ccdc136 | ENSMUSG( | ENSMUSG( | -0.26135 | 0.635315 | NO | NO | NA |
| Gm41495 | ENSMUSG( | ENSMUSG( | 0.38691 | 0.724323 | NO | NO | NA |
| ENSMUSG( | ENSMUSG( | ENSMUSG( | 0.655555 | 0.514562 | NO | NO | NA |
| Celf3 | ENSMUSG( | ENSMUSG( | 0.605509 | 0.251044 | NO | NO | NA |
| ENSMUSG | ENSMUSG( | ENSMUSG( | 0.020408 | 0.961475 | NO | NO | NA |
| Map3k9 | ENSMUSG( | ENSMUSG( | -0.08689 | 0.856906 | NO | NO | NA |
| Sec14l3 | ENSMUSG( | ENSMUSG( | 0.445484 | 0.689779 | NO | NO | NA |
| Dct | ENSMUSG( | ENSMUSG( | 0.68973 | 0.512771 | NO | NO | NA |
| Myom1 | ENSMUSG( | ENSMUSG( | 0.625207 | 0.349342 | NO | NO | NA |
| Ndor1 ENS | ENSMUSG( | ENSMUSG( | 0.530572 | 0.692221 | NO | NO | NA |
| Olfr1322 | ENSMUSG( | ENSMUSG( | 0.734977 | 0.560755 | NO | NO | NA |
| ENSMUSG( | ENSMUSG( | ENSMUSG( | -0.14114 | 0.793457 | NO | NO | NA |
| Atxn7l1os2 | ENSMUSG( | ENSMUSG( | -0.5665 | 0.047798 | YES | NO | NA |
| C230024C | ENSMUSG( | ENSMUSG( | 0.156976 | 0.779503 | NO | NO | NA |

| | | | | | | | |
|---|---|---|---|---|---|---|---|
| A930030B | ENSMUSG( | ENSMUSG( | 0.950396 | 0.395256 | NO | NO | NA |
| Gm37846 | ENSMUSG( | ENSMUSG( | 0.521996 | 0.619199 | NO | NO | NA |
| 4931406B | ENSMUSG( | ENSMUSG( | 1.221113 | 0.136084 | NO | NO | NA |
| Rimbp3 | ENSMUSG( | ENSMUSG( | -0.26482 | 0.614855 | NO | NO | NA |
| Tchh | ENSMUSG( | ENSMUSG( | 0.589396 | 0.645714 | NO | NO | NA |
| Thsd1 | ENSMUSG( | ENSMUSG( | 0.62568 | 0.264251 | NO | NO | NA |
| Tdrkh | ENSMUSG( | ENSMUSG( | 0.686013 | 0.227437 | NO | NO | NA |
| Gm47508 | ENSMUSG( | ENSMUSG( | 0.259306 | 0.596549 | NO | NO | NA |
| Gm12243 | ENSMUSG( | ENSMUSG( | 1.150014 | 0.305056 | NO | NO | NA |
| Cbln2 | ENSMUSG( | ENSMUSG( | 0.794999 | 0.478041 | NO | NO | NA |
| 4930563J1 | ENSMUSG( | ENSMUSG( | 0.776533 | 0.42289 | NO | NO | NA |
| 4930539J( | ENSMUSG( | ENSMUSG( | 0.17355 | 0.623455 | NO | NO | NA |
| Card9 | ENSMUSG( | ENSMUSG( | -0.17399 | 0.825444 | NO | NO | NA |
| 9330118I2 | ENSMUSG( | ENSMUSG( | 0.641832 | 0.565993 | NO | NO | NA |
| Asic2 | ENSMUSG( | ENSMUSG( | 0.524574 | 0.562892 | NO | NO | NA |
| Vmn1r13 | ENSMUSG( | ENSMUSG( | 0.577597 | 0.616116 | NO | NO | NA |
| Gm10657 | ENSMUSG( | ENSMUSG( | 0.186597 | 0.887324 | NO | NO | NA |
| Igkv3-7 | ENSMUSG( | ENSMUSG( | 6.215313 | NA | NO | NO | NA |
| Gm10441 | ENSMUSG( | ENSMUSG( | 0.808244 | 0.443017 | NO | NO | NA |
| Onecut3 | ENSMUSG( | ENSMUSG( | 0.440789 | 0.650505 | NO | NO | NA |
| Gm28323 | ENSMUSG( | ENSMUSG( | 0.933968 | 0.366869 | NO | NO | NA |
| Stac3 | ENSMUSG( | ENSMUSG( | -0.2363 | 0.547053 | NO | NO | NA |
| Lrriq1 | ENSMUSG( | ENSMUSG( | 0.339952 | 0.762738 | NO | NO | NA |
| Nap1l5 | ENSMUSG( | ENSMUSG( | -0.44966 | 0.560577 | NO | NO | NA |
| Olfr485 | ENSMUSG( | ENSMUSG( | 0.796566 | 0.567401 | NO | NO | NA |
| Immp2l | ENSMUSG( | ENSMUSG( | 0.627228 | 0.125323 | NO | NO | NA |
| 1300014J1 | ENSMUSG( | ENSMUSG( | 0.343945 | 0.688954 | NO | NO | NA |
| Capsl | ENSMUSG( | ENSMUSG( | 0.636017 | 0.206884 | NO | NO | NA |
| Gm43359 | ENSMUSG( | ENSMUSG( | -0.22467 | 0.626245 | NO | NO | NA |
| Zfp974 | ENSMUSG( | ENSMUSG( | -0.25066 | 0.505843 | NO | NO | NA |
| Gcm2 | ENSMUSG( | ENSMUSG( | 0.717867 | 0.086626 | NO | NO | NA |
| Pid1 | ENSMUSG( | ENSMUSG( | -0.04126 | 0.9633 | NO | NO | NA |
| Gm12454 | ENSMUSG( | ENSMUSG( | 0.878971 | 0.02577 | YES | NO | NA |
| Gm42907 | ENSMUSG( | ENSMUSG( | 0.067803 | 0.894929 | NO | NO | NA |
| Pkhd1l1 | ENSMUSG( | ENSMUSG( | 0.16812 | 0.826347 | NO | NO | NA |
| Zfp583 | ENSMUSG( | ENSMUSG( | -0.27587 | 0.423273 | NO | NO | NA |
| Gm7108 | ENSMUSG( | ENSMUSG( | 0.228952 | 0.792396 | NO | NO | NA |
| Gm9563 | ENSMUSG( | ENSMUSG( | 1.02796 | 0.338938 | NO | NO | NA |
| Gm43203 | ENSMUSG( | ENSMUSG( | -0.52108 | 0.136826 | NO | NO | NA |
| Gm6455 | ENSMUSG( | ENSMUSG( | 1.083767 | 0.295853 | NO | NO | NA |
| ENSMUSG | ENSMUSG( | ENSMUSG( | -0.32224 | 0.462015 | NO | NO | NA |
| Gm38249 | ENSMUSG( | ENSMUSG( | 0.213757 | 0.855194 | NO | NO | NA |
| Slc23a1 | ENSMUSG( | ENSMUSG( | 0.098598 | 0.829889 | NO | NO | NA |
| Gm3235 | ENSMUSG( | ENSMUSG( | -0.16513 | 0.800693 | NO | NO | NA |
| Zfp385a | ENSMUSG( | ENSMUSG( | 0.139944 | 0.778153 | NO | NO | NA |
| Nalcn | ENSMUSG( | ENSMUSG( | 1.368801 | 0.151427 | NO | NO | NA |
| Gm14466 | ENSMUSG( | ENSMUSG( | -1.33254 | 0.006187 | YES | YES | Downregulated |

| | | | | | | | |
|---|---|---|---|---|---|---|---|
| Gm6201 | ENSMUSG( | ENSMUSG( | -0.19615 | 0.700004 | NO | NO | NA |
| Gm9544 | ENSMUSG( | ENSMUSG( | 0.236004 | 0.814938 | NO | NO | NA |
| Olfr767 | ENSMUSG( | ENSMUSG( | 0.78111 | 0.471631 | NO | NO | NA |
| Tafa1 | ENSMUSG( | ENSMUSG( | 1.107542 | 0.275475 | NO | NO | NA |
| Gm37618 | ENSMUSG( | ENSMUSG( | 0.131378 | 0.875112 | NO | NO | NA |
| Mfsd4b1 | ENSMUSG( | ENSMUSG( | 0.297568 | 0.767783 | NO | NO | NA |
| 1700041G | ENSMUSG( | ENSMUSG( | -0.49511 | 0.23712 | NO | NO | NA |
| Gm10254 | ENSMUSG( | ENSMUSG( | -0.13363 | 0.711742 | NO | NO | NA |
| Gm336 | ENSMUSG( | ENSMUSG( | 1.085318 | 0.160044 | NO | NO | NA |
| Ccn3 | ENSMUSG( | ENSMUSG( | 0.746999 | 0.453757 | NO | NO | NA |
| H2-Q2 | ENSMUSG( | ENSMUSG( | 0.380869 | 0.175573 | NO | NO | NA |
| Lin28b | ENSMUSG( | ENSMUSG( | 0.532495 | 0.387563 | NO | NO | NA |
| Gpr157 | ENSMUSG( | ENSMUSG( | 0.018153 | 0.979703 | NO | NO | NA |
| Gm37077 | ENSMUSG( | ENSMUSG( | 0.781394 | 0.482762 | NO | NO | NA |
| D330025C | ENSMUSG( | ENSMUSG( | 0.942746 | 0.401158 | NO | NO | NA |
| B230208H | ENSMUSG( | ENSMUSG( | 0.227385 | 0.556665 | NO | NO | NA |
| Olfr794 | ENSMUSG( | ENSMUSG( | 1.584211 | 0.138812 | NO | NO | NA |
| Lrat | ENSMUSG( | ENSMUSG( | 1.137277 | 0.281012 | NO | NO | NA |
| Gm31024 | ENSMUSG( | ENSMUSG( | 0.054906 | 0.868932 | NO | NO | NA |
| Gm17709 | ENSMUSG( | ENSMUSG( | -0.97412 | 0.01068 | YES | NO | NA |
| Gm43175 | ENSMUSG( | ENSMUSG( | 0.856902 | 0.432225 | NO | NO | NA |
| Gm47603 | ENSMUSG( | ENSMUSG( | -0.99501 | 0.027112 | YES | NO | NA |
| Gm16008 | ENSMUSG( | ENSMUSG( | 0.312855 | 0.533864 | NO | NO | NA |
| Gm51802 | ENSMUSG( | ENSMUSG( | 1.639615 | 0.001919 | YES | YES | Upregulated |
| ENSMUSG | ENSMUSG( | ENSMUSG( | 0.311306 | 0.761814 | NO | NO | NA |
| Gm26831 | ENSMUSG( | ENSMUSG( | 0.745591 | 0.548345 | NO | NO | NA |
| Gm14137 | ENSMUSG( | ENSMUSG( | 0.225561 | 0.542249 | NO | NO | NA |
| Gm15845 | ENSMUSG( | ENSMUSG( | 1.003585 | 0.346539 | NO | NO | NA |
| A730020E( | ENSMUSG( | ENSMUSG( | 0.926713 | 0.438825 | NO | NO | NA |
| D830039M | ENSMUSG( | ENSMUSG( | 0.890493 | 0.263012 | NO | NO | NA |
| Phex | ENSMUSG( | ENSMUSG( | 0.640763 | 0.350038 | NO | NO | NA |
| Col8a1 | ENSMUSG( | ENSMUSG( | 0.479428 | 0.659517 | NO | NO | NA |
| Gm47085 | ENSMUSG( | ENSMUSG( | 0.581766 | 0.586771 | NO | NO | NA |
| Gm31805 | ENSMUSG( | ENSMUSG( | 0.327082 | 0.749553 | NO | NO | NA |
| C1ql4 | ENSMUSG( | ENSMUSG( | 0.079391 | 0.942764 | NO | NO | NA |
| ENSMUSG | ENSMUSG( | ENSMUSG( | 0.382245 | 0.382461 | NO | NO | NA |
| Macroh2a | ENSMUSG( | ENSMUSG( | -1.7275 | 1.64E-05 | YES | YES | Downregulated |
| Rorc | ENSMUSG( | ENSMUSG( | -0.20329 | 0.748964 | NO | NO | NA |
| Rtl8c | ENSMUSG( | ENSMUSG( | -0.63725 | 0.162461 | NO | NO | NA |
| Cxcl3 | ENSMUSG( | ENSMUSG( | 0.957325 | 0.438698 | NO | NO | NA |
| 1700026D | ENSMUSG( | ENSMUSG( | 0.287021 | 0.632853 | NO | NO | NA |
| Gm8899 | ENSMUSG( | ENSMUSG( | 0.063791 | 0.919867 | NO | NO | NA |
| Gm3183 | ENSMUSG( | ENSMUSG( | -0.15429 | 0.884488 | NO | NO | NA |
| Gpr162 | ENSMUSG( | ENSMUSG( | 0.368679 | 0.402409 | NO | NO | NA |
| ENSMUSG | ENSMUSG( | ENSMUSG( | 0.438529 | 0.275319 | NO | NO | NA |
| Gm5891 | ENSMUSG( | ENSMUSG( | -0.34625 | NA | NO | NO | NA |
| Slc5a9 | ENSMUSG( | ENSMUSG( | 0.920704 | 0.381602 | NO | NO | NA |

| | | | | | | | |
|---|---|---|---|---|---|---|---|
| Gm43843 | ENSMUSG( | ENSMUSG( | 0.487712 | 0.700372 | NO | NO | NA |
| Gm26967 | ENSMUSG( | ENSMUSG( | 0.454428 | 0.74816 | NO | NO | NA |
| Olfr554 | ENSMUSG( | ENSMUSG( | 0.541856 | 0.600518 | NO | NO | NA |
| Alpk1 | ENSMUSG( | ENSMUSG( | 0.970701 | 0.011575 | YES | NO | NA |
| Gnrhr | ENSMUSG( | ENSMUSG( | 1.080509 | 0.247351 | NO | NO | NA |
| 4930528G | ENSMUSG( | ENSMUSG( | 1.16049 | 0.304481 | NO | NO | NA |
| ENSMUSG | ENSMUSG( | ENSMUSG( | -0.89713 | 0.008976 | YES | NO | NA |
| Gm49474 | ENSMUSG( | ENSMUSG( | 0.33976 | 0.613549 | NO | NO | NA |
| Myh7 | ENSMUSG( | ENSMUSG( | 0.230987 | 0.821805 | NO | NO | NA |
| Slfn14 | ENSMUSG( | ENSMUSG( | 1.010148 | 0.138778 | NO | NO | NA |
| Gm26691 | ENSMUSG( | ENSMUSG( | 0.608968 | 0.579782 | NO | NO | NA |
| ENSMUSG | ENSMUSG( | ENSMUSG( | 0.492276 | 0.314992 | NO | NO | NA |
| Gm35842 | ENSMUSG( | ENSMUSG( | 0.253506 | 0.815712 | NO | NO | NA |
| Lsmem1 | ENSMUSG( | ENSMUSG( | 1.123657 | 0.331156 | NO | NO | NA |
| Gm37724 | ENSMUSG( | ENSMUSG( | -0.19038 | 0.623382 | NO | NO | NA |
| Gm9930 | ENSMUSG( | ENSMUSG( | 0.746974 | 0.580095 | NO | NO | NA |
| Zfp345 | ENSMUSG( | ENSMUSG( | 0.634179 | 0.465973 | NO | NO | NA |
| Gm38067 | ENSMUSG( | ENSMUSG( | 0.180388 | 0.680393 | NO | NO | NA |
| Olfr1330 | ENSMUSG( | ENSMUSG( | 0.707996 | 0.567382 | NO | NO | NA |
| 4930588J1 | ENSMUSG( | ENSMUSG( | 0.910105 | 0.475513 | NO | NO | NA |
| Gm7177 | ENSMUSG( | ENSMUSG( | 0.325517 | 0.741426 | NO | NO | NA |
| Gm7449 | ENSMUSG( | ENSMUSG( | 0.71239 | 0.500028 | NO | NO | NA |
| Crybb3 | ENSMUSG( | ENSMUSG( | 0.248698 | 0.750323 | NO | NO | NA |
| Olfr691 | ENSMUSG( | ENSMUSG( | 1.048602 | 0.342997 | NO | NO | NA |
| Vmn2r95 | ENSMUSG( | ENSMUSG( | 0.593758 | 0.467681 | NO | NO | NA |
| ENSMUSG | ENSMUSG( | ENSMUSG( | 0.497054 | 0.288249 | NO | NO | NA |
| 9330161L( | ENSMUSG( | ENSMUSG( | 0.202729 | 0.689779 | NO | NO | NA |
| Gm10773 | ENSMUSG( | ENSMUSG( | -0.8435 | 0.075866 | NO | NO | NA |
| Cngb3 | ENSMUSG( | ENSMUSG( | 0.606041 | 0.580095 | NO | NO | NA |
| Igkv13-84 | ENSMUSG( | ENSMUSG( | -2.19778 | 0.376668 | NO | NO | NA |
| ENSMUSG | ENSMUSG( | ENSMUSG( | 0.549266 | 0.187531 | NO | NO | NA |
| Gucy1a2 | ENSMUSG( | ENSMUSG( | 0.227777 | 0.841956 | NO | NO | NA |
| G630064G | ENSMUSG( | ENSMUSG( | 1.220568 | 0.34493 | NO | NO | NA |
| Gm43417 | ENSMUSG( | ENSMUSG( | 0.138868 | 0.872677 | NO | NO | NA |
| Gm47246 | ENSMUSG( | ENSMUSG( | -0.09307 | 0.851096 | NO | NO | NA |
| Rin1 | ENSMUSG( | ENSMUSG( | 0.103226 | 0.749566 | NO | NO | NA |
| Arl11 | ENSMUSG( | ENSMUSG( | -0.09842 | 0.808851 | NO | NO | NA |
| Gm29017 | ENSMUSG( | ENSMUSG( | -0.24722 | 0.556665 | NO | NO | NA |
| 4931422A( | ENSMUSG( | ENSMUSG( | -0.39625 | 0.318104 | NO | NO | NA |
| Gm37781 | ENSMUSG( | ENSMUSG( | 0.655336 | 0.536401 | NO | NO | NA |
| Ackr3 | ENSMUSG( | ENSMUSG( | 0.553222 | 0.590589 | NO | NO | NA |
| Cdcp1 | ENSMUSG( | ENSMUSG( | 0.541117 | 0.429674 | NO | NO | NA |
| Olfr867 | ENSMUSG( | ENSMUSG( | 0.910192 | 0.468146 | NO | NO | NA |
| Tifab | ENSMUSG( | ENSMUSG( | -0.5147 | 0.417592 | NO | NO | NA |
| Phox2b | ENSMUSG( | ENSMUSG( | 0.345107 | 0.725203 | NO | NO | NA |
| Gm5805 | ENSMUSG( | ENSMUSG( | 0.451988 | 0.352248 | NO | NO | NA |
| ENSMUSG | ENSMUSG( | ENSMUSG( | 0.487532 | 0.612838 | NO | NO | NA |

| | | | | | | | |
|---|---|---|---|---|---|---|---|
| 4930579K1 | ENSMUSG( | ENSMUSG( | 0.848918 | 0.076531 | NO | NO | NA |
| Hmga2 | ENSMUSG( | ENSMUSG( | 0.314621 | 0.757268 | NO | NO | NA |
| Olfr352 | ENSMUSG( | ENSMUSG( | 0.263149 | 0.810521 | NO | NO | NA |
| Gm13423 | ENSMUSG( | ENSMUSG( | 0.325722 | 0.74568 | NO | NO | NA |
| Gm15401 | ENSMUSG( | ENSMUSG( | 0.028762 | 0.955338 | NO | NO | NA |
| Gm47503 | ENSMUSG( | ENSMUSG( | 0.834763 | 0.162461 | NO | NO | NA |
| Gm9817 | ENSMUSG( | ENSMUSG( | 0.593553 | 0.593125 | NO | NO | NA |
| Gm7600 | ENSMUSG( | ENSMUSG( | 1.030018 | 0.001048 | YES | YES | Upregulated |
| Zfp78 | ENSMUSG( | ENSMUSG( | -0.14464 | 0.731835 | NO | NO | NA |
| Gm49902 | ENSMUSG( | ENSMUSG( | 0.466601 | 0.342258 | NO | NO | NA |
| Gm10847 | ENSMUSG( | ENSMUSG( | 0.48577 | 0.510513 | NO | NO | NA |
| Gm48624 | ENSMUSG( | ENSMUSG( | -0.39143 | 0.466943 | NO | NO | NA |
| Foxl1 | ENSMUSG( | ENSMUSG( | 0.462613 | 0.679095 | NO | NO | NA |
| Unc5c | ENSMUSG( | ENSMUSG( | 0.412302 | 0.6888 | NO | NO | NA |
| Gm19656 | ENSMUSG( | ENSMUSG( | 0.599965 | 0.560948 | NO | NO | NA |
| Gm44224 | ENSMUSG( | ENSMUSG( | 0.747139 | 0.471567 | NO | NO | NA |
| Gm49027 | ENSMUSG( | ENSMUSG( | -0.41621 | 0.558121 | NO | NO | NA |
| Inka1 | ENSMUSG( | ENSMUSG( | 0.325016 | 0.430364 | NO | NO | NA |
| Gm4951 | ENSMUSG( | ENSMUSG( | 0.144108 | 0.824088 | NO | NO | NA |
| ENSMUSG | ENSMUSG( | ENSMUSG( | -0.00569 | 0.990219 | NO | NO | NA |
| Olfr851 | ENSMUSG( | ENSMUSG( | 0.551008 | 0.611216 | NO | NO | NA |
| Chst3 | ENSMUSG( | ENSMUSG( | 0.917456 | 0.083486 | NO | NO | NA |
| Gm43503 | ENSMUSG( | ENSMUSG( | 0.41041 | 0.682283 | NO | NO | NA |
| Olfr1170 | ENSMUSG( | ENSMUSG( | 1.020399 | 0.369772 | NO | NO | NA |
| Pcdh7 | ENSMUSG( | ENSMUSG( | 0.566326 | 0.529877 | NO | NO | NA |
| Impg2 | ENSMUSG( | ENSMUSG( | 0.217812 | 0.771535 | NO | NO | NA |
| Slc7a8 | ENSMUSG( | ENSMUSG( | 0.115523 | 0.868434 | NO | NO | NA |
| Gm42690 | ENSMUSG( | ENSMUSG( | -0.1933 | 0.684189 | NO | NO | NA |
| Gm48298 | ENSMUSG( | ENSMUSG( | 0.643143 | 0.55053 | NO | NO | NA |
| Gm44699 | ENSMUSG( | ENSMUSG( | -0.07854 | 0.913341 | NO | NO | NA |
| Gm49705 | ENSMUSG( | ENSMUSG( | 0.300325 | 0.724236 | NO | NO | NA |
| Gm26703 | ENSMUSG( | ENSMUSG( | 0.39863 | 0.749795 | NO | NO | NA |
| Tyrobp | ENSMUSG( | ENSMUSG( | -0.44017 | 0.413936 | NO | NO | NA |
| ENSMUSG | ENSMUSG( | ENSMUSG( | 1.129662 | 0.420216 | NO | NO | NA |
| Jam3 | ENSMUSG( | ENSMUSG( | 0.393953 | 0.403242 | NO | NO | NA |
| Gm45630 | ENSMUSG( | ENSMUSG( | -0.2003 | 0.766729 | NO | NO | NA |
| Gm20631 | ENSMUSG( | ENSMUSG( | 0.746023 | 0.518069 | NO | NO | NA |
| Dkk2 | ENSMUSG( | ENSMUSG( | 0.72352 | 0.518523 | NO | NO | NA |
| Fermt2 | ENSMUSG( | ENSMUSG( | 0.884322 | 0.452821 | NO | NO | NA |
| Prkag2os2 | ENSMUSG( | ENSMUSG( | -0.28934 | 0.72269 | NO | NO | NA |
| D030025E | ENSMUSG( | ENSMUSG( | 0.846141 | 0.440997 | NO | NO | NA |
| Dnmt3aos | ENSMUSG( | ENSMUSG( | 0.708576 | 0.49285 | NO | NO | NA |
| Gm43338 | ENSMUSG( | ENSMUSG( | -0.01652 | 0.968705 | NO | NO | NA |
| Gprin2 | ENSMUSG( | ENSMUSG( | 0.951221 | 0.250293 | NO | NO | NA |
| ENSMUSG | ENSMUSG( | ENSMUSG( | 0.392181 | 0.748139 | NO | NO | NA |
| Gm44985 | ENSMUSG( | ENSMUSG( | 0.087421 | 0.861004 | NO | NO | NA |
| Fcgr3 | ENSMUSG( | ENSMUSG( | -0.37271 | 0.337629 | NO | NO | NA |

| | | | | | | | |
|---|---|---|---|---|---|---|---|
| A430050A | ENSMUSG( | ENSMUSG( | 1.033967 | 0.342353 | NO | NO | NA |
| P2ry1 | ENSMUSG( | ENSMUSG( | 0.408328 | 0.613953 | NO | NO | NA |
| Gm48653 | ENSMUSG( | ENSMUSG( | 0.059213 | 0.896053 | NO | NO | NA |
| G630030J( | ENSMUSG( | ENSMUSG( | 0.142462 | 0.764246 | NO | NO | NA |
| E230032D | ENSMUSG( | ENSMUSG( | -0.61635 | 0.1307 | NO | NO | NA |
| ENSMUSG | ENSMUSG( | ENSMUSG( | 0.865577 | 0.007594 | YES | NO | NA |
| Olfr1425 | ENSMUSG( | ENSMUSG( | 1.000224 | 0.360013 | NO | NO | NA |
| Kdm6bos | ENSMUSG( | ENSMUSG( | 0.588782 | 0.402098 | NO | NO | NA |
| ENSMUSG | ENSMUSG( | ENSMUSG( | 0.562985 | 0.062199 | NO | NO | NA |
| Hhip | ENSMUSG( | ENSMUSG( | 0.304498 | 0.771858 | NO | NO | NA |
| Olfr308 | ENSMUSG( | ENSMUSG( | 0.669603 | 0.544325 | NO | NO | NA |
| Arhgef15 | ENSMUSG( | ENSMUSG( | 0.430821 | 0.672577 | NO | NO | NA |
| Ceacam3 | ENSMUSG( | ENSMUSG( | 0.811561 | 0.391427 | NO | NO | NA |
| Dppa2 | ENSMUSG( | ENSMUSG( | 0.696287 | 0.492185 | NO | NO | NA |
| Kcnk2 | ENSMUSG( | ENSMUSG( | 0.195582 | 0.88103 | NO | NO | NA |
| Ago4 | ENSMUSG( | ENSMUSG( | 0.515537 | 0.334781 | NO | NO | NA |
| Gm43859 | ENSMUSG( | ENSMUSG( | -0.14664 | 0.707721 | NO | NO | NA |
| Gm15697 | ENSMUSG( | ENSMUSG( | 0.393539 | 0.573126 | NO | NO | NA |
| Gm3953 | ENSMUSG( | ENSMUSG( | 0.914698 | 0.465084 | NO | NO | NA |
| Gm10369 | ENSMUSG( | ENSMUSG( | 0.317118 | 0.435027 | NO | NO | NA |
| Slc27a3 | ENSMUSG( | ENSMUSG( | -0.13083 | 0.729391 | NO | NO | NA |
| Gm38340 | ENSMUSG( | ENSMUSG( | 0.439175 | 0.592686 | NO | NO | NA |
| Cabcoco1 | ENSMUSG( | ENSMUSG( | 1.018211 | 0.089143 | NO | NO | NA |
| Slc6a19 | ENSMUSG( | ENSMUSG( | 0.505686 | 0.568701 | NO | NO | NA |
| Trav13d-3 | ENSMUSG( | ENSMUSG( | 0.162092 | 0.915566 | NO | NO | NA |
| Cpxm2 | ENSMUSG( | ENSMUSG( | 0.762375 | 0.48065 | NO | NO | NA |
| Platr6 | ENSMUSG( | ENSMUSG( | 0.672596 | 0.535131 | NO | NO | NA |
| Olfr1340 | ENSMUSG( | ENSMUSG( | 0.793461 | 0.433108 | NO | NO | NA |
| Gm867 | ENSMUSG( | ENSMUSG( | -0.32664 | 0.62014 | NO | NO | NA |
| Gm48961 | ENSMUSG( | ENSMUSG( | -0.61217 | 0.081799 | NO | NO | NA |
| Vmn2r31 | ENSMUSG( | ENSMUSG( | 0.810259 | 0.42412 | NO | NO | NA |
| Colq | ENSMUSG( | ENSMUSG( | 0.628219 | 0.156934 | NO | NO | NA |
| Gm3379 | ENSMUSG( | ENSMUSG( | 0.709551 | 0.288954 | NO | NO | NA |
| Gm47671 | ENSMUSG( | ENSMUSG( | 0.894234 | 0.424091 | NO | NO | NA |
| Slc16a9 | ENSMUSG( | ENSMUSG( | 0.915855 | 0.426541 | NO | NO | NA |
| Ufsp1 | ENSMUSG( | ENSMUSG( | 0.136017 | 0.756401 | NO | NO | NA |
| Khdc1a | ENSMUSG( | ENSMUSG( | 1.273357 | 0.168698 | NO | NO | NA |
| Rab7b | ENSMUSG( | ENSMUSG( | 0.105268 | 0.879395 | NO | NO | NA |
| Fbxl12os | ENSMUSG( | ENSMUSG( | -0.1254 | 0.783948 | NO | NO | NA |
| Fam71a | ENSMUSG( | ENSMUSG( | 0.821164 | 0.424153 | NO | NO | NA |
| Ccdc187 | ENSMUSG( | ENSMUSG( | 0.330379 | 0.762454 | NO | NO | NA |
| Mkx | ENSMUSG( | ENSMUSG( | 0.437774 | 0.668604 | NO | NO | NA |
| Gm7274 | ENSMUSG( | ENSMUSG( | 0.969622 | 0.409568 | NO | NO | NA |
| Lim2 | ENSMUSG( | ENSMUSG( | -1.6658 | 0.000407 | YES | YES | Downregulated |
| 4933405D | ENSMUSG( | ENSMUSG( | 1.175018 | 0.306917 | NO | NO | NA |
| Sall1 | ENSMUSG( | ENSMUSG( | 1.139798 | 0.209547 | NO | NO | NA |
| Gm47704 | ENSMUSG( | ENSMUSG( | 1.052429 | 0.440587 | NO | NO | NA |

| | | | | | | | |
|---|---|---|---|---|---|---|---|
| Gm5452 | ENSMUSG( | ENSMUSG( | 0.964524 | 0.062254 | NO | NO | NA |
| Gm43482 | ENSMUSG( | ENSMUSG( | 0.165577 | 0.770492 | NO | NO | NA |
| ENSMUSG | ENSMUSG( | ENSMUSG( | -0.07407 | 0.872232 | NO | NO | NA |
| Gm42949 | ENSMUSG( | ENSMUSG( | 0.76274 | 0.483755 | NO | NO | NA |
| Vmn1r75 | ENSMUSG( | ENSMUSG( | 0.750315 | 0.485845 | NO | NO | NA |
| Tmem150a | ENSMUSG( | ENSMUSG( | 0.26264 | 0.452821 | NO | NO | NA |
| ENSMUSG | ENSMUSG( | ENSMUSG( | 0.849961 | 0.152477 | NO | NO | NA |
| Gm13270 | ENSMUSG( | ENSMUSG( | -0.20069 | 0.796234 | NO | NO | NA |
| Gm18597 | ENSMUSG( | ENSMUSG( | 0.755995 | 0.49117 | NO | NO | NA |
| Ugt8a | ENSMUSG( | ENSMUSG( | 0.691578 | 0.377415 | NO | NO | NA |
| Gm48119 | ENSMUSG( | ENSMUSG( | -0.18857 | 0.844233 | NO | NO | NA |
| Gm12576 | ENSMUSG( | ENSMUSG( | 0.688423 | 0.525251 | NO | NO | NA |
| Gm5913 | ENSMUSG( | ENSMUSG( | 0.339025 | 0.352768 | NO | NO | NA |
| Sh3d21 | ENSMUSG( | ENSMUSG( | -0.1785 | 0.705305 | NO | NO | NA |
| Gm29571 | ENSMUSG( | ENSMUSG( | 0.788932 | 0.496838 | NO | NO | NA |
| Gm40968 | ENSMUSG( | ENSMUSG( | 0.539368 | 0.621768 | NO | NO | NA |
| Gm45601 | ENSMUSG( | ENSMUSG( | 0.694303 | 0.56241 | NO | NO | NA |
| Gm5063 | ENSMUSG( | ENSMUSG( | -0.24331 | 0.704289 | NO | NO | NA |
| Slit2 | ENSMUSG( | ENSMUSG( | 0.2966 | 0.733562 | NO | NO | NA |
| Slc35f4 | ENSMUSG( | ENSMUSG( | 0.768276 | 0.493119 | NO | NO | NA |
| Gm13561 | ENSMUSG( | ENSMUSG( | -0.20272 | 0.669294 | NO | NO | NA |
| Gm44711 | ENSMUSG( | ENSMUSG( | 0.697853 | 0.513389 | NO | NO | NA |
| Gm32051 | ENSMUSG( | ENSMUSG( | 0.709702 | 0.190962 | NO | NO | NA |
| Gdf3 | ENSMUSG( | ENSMUSG( | 1.196101 | 0.394514 | NO | NO | NA |
| Gm48139 | ENSMUSG( | ENSMUSG( | -0.8136 | 0.275571 | NO | NO | NA |
| Gm42417 | ENSMUSG( | ENSMUSG( | 0.46315 | 0.794205 | NO | NO | NA |
| Gm47339 | ENSMUSG( | ENSMUSG( | 0.581657 | 0.572582 | NO | NO | NA |
| Gm7167 | ENSMUSG( | ENSMUSG( | 0.918327 | 0.340105 | NO | NO | NA |
| Eaf2 | ENSMUSG( | ENSMUSG( | 0.178814 | 0.712916 | NO | NO | NA |
| P2ry13 | ENSMUSG( | ENSMUSG( | 0.673657 | 0.060254 | NO | NO | NA |
| Vmn1r11 | ENSMUSG( | ENSMUSG( | 0.826328 | 0.49711 | NO | NO | NA |
| Gm38250 | ENSMUSG( | ENSMUSG( | -0.28318 | 0.417268 | NO | NO | NA |
| Gm28512 | ENSMUSG( | ENSMUSG( | -0.73597 | 0.201552 | NO | NO | NA |
| Gm10232 | ENSMUSG( | ENSMUSG( | 0.401491 | 0.165085 | NO | NO | NA |
| Gm36981 | ENSMUSG( | ENSMUSG( | -0.79797 | 0.126359 | NO | NO | NA |
| Gm44220 | ENSMUSG( | ENSMUSG( | 0.694105 | 0.544325 | NO | NO | NA |
| ENSMUSG( | ENSMUSG( | ENSMUSG( | 1.025921 | 0.346493 | NO | NO | NA |
| Trgc3 | ENSMUSG( | ENSMUSG( | -0.3636 | 0.661527 | NO | NO | NA |
| C78859 | ENSMUSG( | ENSMUSG( | 0.479005 | 0.48793 | NO | NO | NA |
| Pdzd9 | ENSMUSG( | ENSMUSG( | 0.111667 | 0.816297 | NO | NO | NA |
| Smim10l2a | ENSMUSG( | ENSMUSG( | 0.300531 | 0.478694 | NO | NO | NA |
| Galntl6 | ENSMUSG( | ENSMUSG( | 0.711541 | 0.336832 | NO | NO | NA |
| Gm42986 | ENSMUSG( | ENSMUSG( | -0.19121 | 0.682983 | NO | NO | NA |
| Gm50228 | ENSMUSG( | ENSMUSG( | -0.06558 | 0.923116 | NO | NO | NA |
| ENSMUSG | ENSMUSG( | ENSMUSG( | -0.77537 | 0.0431 | YES | NO | NA |
| Trgv6 | ENSMUSG( | ENSMUSG( | -2.21826 | 0.039785 | YES | YES | Downregulated |
| Gm49024 | ENSMUSG( | ENSMUSG( | 0.813035 | 0.504241 | NO | NO | NA |

| | | | | | | | |
|---|---|---|---|---|---|---|---|
| Tmem144 | ENSMUSG( | ENSMUSG( | 0.440777 | 0.384684 | NO | NO | NA |
| Cebpa | ENSMUSG( | ENSMUSG( | 0.6927 | 0.262135 | NO | NO | NA |
| Rps6-ps1 | ENSMUSG( | ENSMUSG( | 0.481971 | 0.231678 | NO | NO | NA |
| Gm43237 | ENSMUSG( | ENSMUSG( | 0.469033 | 0.226904 | NO | NO | NA |
| Prr15 | ENSMUSG( | ENSMUSG( | 0.049796 | 0.95848 | NO | NO | NA |
| Gm37726 | ENSMUSG( | ENSMUSG( | -0.40033 | 0.473914 | NO | NO | NA |
| Smim41 | ENSMUSG( | ENSMUSG( | 0.727685 | 0.448596 | NO | NO | NA |
| Gm26621 | ENSMUSG( | ENSMUSG( | -0.35769 | 0.384382 | NO | NO | NA |
| Gm44039 | ENSMUSG( | ENSMUSG( | 1.220458 | 0.31087 | NO | NO | NA |
| Tmem150 | ENSMUSG( | ENSMUSG( | 0.679604 | 0.549903 | NO | NO | NA |
| Gm9115 | ENSMUSG( | ENSMUSG( | 0.382913 | 0.611454 | NO | NO | NA |
| Arx | ENSMUSG( | ENSMUSG( | 0.68129 | 0.434361 | NO | NO | NA |
| Dmc1 | ENSMUSG( | ENSMUSG( | 1.924357 | 0.019176 | YES | YES | Upregulated |
| ENSMUSG | ENSMUSG( | ENSMUSG( | -0.04868 | 0.92464 | NO | NO | NA |
| ENSMUSG | ENSMUSG( | ENSMUSG( | 0.215048 | 0.620133 | NO | NO | NA |
| Adtrp | ENSMUSG( | ENSMUSG( | 1.101101 | 0.30792 | NO | NO | NA |
| Gm10748 | ENSMUSG( | ENSMUSG( | 0.531971 | 0.626786 | NO | NO | NA |
| Rps6-ps3 | ENSMUSG( | ENSMUSG( | 0.365376 | 0.566439 | NO | NO | NA |
| Gm37913 | ENSMUSG( | ENSMUSG( | 0.458676 | 0.685265 | NO | NO | NA |
| ENSMUSG | ENSMUSG( | ENSMUSG( | 0.666229 | 0.347976 | NO | NO | NA |
| Cntfr | ENSMUSG( | ENSMUSG( | -1.25014 | 0.093724 | NO | NO | NA |
| Gprc5a | ENSMUSG( | ENSMUSG( | -0.4547 | 0.229095 | NO | NO | NA |
| Gm11870 | ENSMUSG( | ENSMUSG( | -0.22874 | 0.650267 | NO | NO | NA |
| Serpinb13 | ENSMUSG( | ENSMUSG( | 0.89628 | 0.39814 | NO | NO | NA |
| Pnoc | ENSMUSG( | ENSMUSG( | 0.99571 | 0.347072 | NO | NO | NA |
| ENSMUSG( | ENSMUSG( | ENSMUSG( | 1.136358 | 0.325929 | NO | NO | NA |
| Gm42683 | ENSMUSG( | ENSMUSG( | 0.344986 | 0.60697 | NO | NO | NA |
| Gm48619 | ENSMUSG( | ENSMUSG( | 1.310849 | 0.168709 | NO | NO | NA |
| 1810012K( | ENSMUSG( | ENSMUSG( | 0.057037 | 0.944999 | NO | NO | NA |
| ENSMUSG | ENSMUSG( | ENSMUSG( | 0.097437 | 0.84067 | NO | NO | NA |
| Arsj | ENSMUSG( | ENSMUSG( | 0.741272 | 0.534341 | NO | NO | NA |
| Bmp8a | ENSMUSG( | ENSMUSG( | -0.5926 | 0.372027 | NO | NO | NA |
| Gm37436 | ENSMUSG( | ENSMUSG( | 0.537631 | 0.407702 | NO | NO | NA |
| A930005G | ENSMUSG( | ENSMUSG( | 1.212843 | 0.323577 | NO | NO | NA |
| ENSMUSG | ENSMUSG( | ENSMUSG( | -0.24316 | 0.616241 | NO | NO | NA |
| Lrrc7 | ENSMUSG( | ENSMUSG( | 0.830512 | 0.411763 | NO | NO | NA |
| Gm42860 | ENSMUSG( | ENSMUSG( | 0.30895 | 0.488531 | NO | NO | NA |
| Gm50268 | ENSMUSG( | ENSMUSG( | 1.090221 | 0.389049 | NO | NO | NA |
| Pafah2 | ENSMUSG( | ENSMUSG( | 0.000322 | 0.999012 | NO | NO | NA |
| Gm44816 | ENSMUSG( | ENSMUSG( | -0.66421 | 0.130055 | NO | NO | NA |
| Dpf3 | ENSMUSG( | ENSMUSG( | 0.860986 | 0.117079 | NO | NO | NA |
| Gm10804 | ENSMUSG( | ENSMUSG( | 0.479281 | 0.365594 | NO | NO | NA |
| Gm37518 | ENSMUSG( | ENSMUSG( | 0.602662 | 0.610533 | NO | NO | NA |
| Slc2a13 | ENSMUSG( | ENSMUSG( | 0.051871 | 0.960855 | NO | NO | NA |
| Nlrp10 | ENSMUSG( | ENSMUSG( | 1.059176 | 0.192701 | NO | NO | NA |
| Tacstd2 | ENSMUSG( | ENSMUSG( | 4.931913 | 1.66E-10 | YES | YES | Upregulated |
| Gm45286 | ENSMUSG( | ENSMUSG( | -0.3292 | 0.392394 | NO | NO | NA |

| | | | | | | | |
|---|---|---|---|---|---|---|---|
| Vmn2r86 | ENSMUSG( | ENSMUSG( | 1.08367 | 0.349038 | NO | NO | NA |
| Sort1 | ENSMUSG( | ENSMUSG( | 0.05805 | 0.909855 | NO | NO | NA |
| Gm10447 | ENSMUSG( | ENSMUSG( | 0.754835 | 0.499327 | NO | NO | NA |
| ENSMUSG | ENSMUSG( | ENSMUSG( | 0.625203 | 0.662467 | NO | NO | NA |
| Pax8 | ENSMUSG( | ENSMUSG( | 0.586476 | 0.418656 | NO | NO | NA |
| Gm10263 | ENSMUSG( | ENSMUSG( | 0.601131 | 0.238548 | NO | NO | NA |
| Gm36930 | ENSMUSG( | ENSMUSG( | 0.874596 | 0.361782 | NO | NO | NA |
| Shc4 | ENSMUSG( | ENSMUSG( | 0.180589 | 0.658781 | NO | NO | NA |
| Angpt2 | ENSMUSG( | ENSMUSG( | -0.81001 | 0.160171 | NO | NO | NA |
| Gm6170 | ENSMUSG( | ENSMUSG( | 0.324744 | 0.568223 | NO | NO | NA |
| Pcdhb16 | ENSMUSG( | ENSMUSG( | 0.914756 | 0.439832 | NO | NO | NA |
| Gm42702 | ENSMUSG( | ENSMUSG( | 0.407248 | 0.70225 | NO | NO | NA |
| Gm50232 | ENSMUSG( | ENSMUSG( | -0.3006 | 0.466606 | NO | NO | NA |
| Gm26674 | ENSMUSG( | ENSMUSG( | 0.491485 | 0.698856 | NO | NO | NA |
| Gm27166 | ENSMUSG( | ENSMUSG( | -0.54583 | 0.203066 | NO | NO | NA |
| Ptafr | ENSMUSG( | ENSMUSG( | 0.494517 | 0.660508 | NO | NO | NA |
| Gm35835 | ENSMUSG( | ENSMUSG( | 1.066156 | 0.344361 | NO | NO | NA |
| 4930455G | ENSMUSG( | ENSMUSG( | 0.543079 | 0.60683 | NO | NO | NA |
| Rasgrp4 | ENSMUSG( | ENSMUSG( | -0.11252 | 0.844822 | NO | NO | NA |
| Gm20465 | ENSMUSG( | ENSMUSG( | 0.682126 | 0.579976 | NO | NO | NA |
| Tex13b | ENSMUSG( | ENSMUSG( | 0.428945 | 0.689596 | NO | NO | NA |
| Cd36 | ENSMUSG( | ENSMUSG( | 0.336457 | 0.636057 | NO | NO | NA |
| Lncbate6 | ENSMUSG( | ENSMUSG( | 0.790898 | 0.248611 | NO | NO | NA |
| Lamb1 | ENSMUSG( | ENSMUSG( | 0.126321 | 0.91877 | NO | NO | NA |
| Gm43004 | ENSMUSG( | ENSMUSG( | 0.705108 | 0.168976 | NO | NO | NA |
| ENSMUSG | ENSMUSG( | ENSMUSG( | 0.94075 | 0.377259 | NO | NO | NA |
| Dgkk | ENSMUSG( | ENSMUSG( | 0.44325 | 0.761868 | NO | NO | NA |
| Gm15889 | ENSMUSG( | ENSMUSG( | 0.478754 | 0.720059 | NO | NO | NA |
| Gm43522 | ENSMUSG( | ENSMUSG( | 0.619745 | 0.569018 | NO | NO | NA |
| ENSMUSG( | ENSMUSG( | ENSMUSG( | 0.1693 | 0.736093 | NO | NO | NA |
| BC051077 | ENSMUSG( | ENSMUSG( | 0.227555 | 0.805716 | NO | NO | NA |
| Dcbld1 | ENSMUSG( | ENSMUSG( | 0.669765 | 0.371448 | NO | NO | NA |
| Wbscr25 | ENSMUSG( | ENSMUSG( | 0.279107 | 0.808208 | NO | NO | NA |
| A230057D | ENSMUSG( | ENSMUSG( | 0.778349 | 0.399515 | NO | NO | NA |
| Tmem38a | ENSMUSG( | ENSMUSG( | 0.470693 | 0.219122 | NO | NO | NA |
| Gm47558 | ENSMUSG( | ENSMUSG( | 0.303793 | 0.668541 | NO | NO | NA |
| 9130008F: | ENSMUSG( | ENSMUSG( | -0.24942 | 0.48949 | NO | NO | NA |
| Gm5089 | ENSMUSG( | ENSMUSG( | 0.885778 | 0.400823 | NO | NO | NA |
| ENSMUSG | ENSMUSG( | ENSMUSG( | 0.007821 | 0.982021 | NO | NO | NA |
| BC051408 | ENSMUSG( | ENSMUSG( | 0.903469 | 0.399944 | NO | NO | NA |
| Gm3294 | ENSMUSG( | ENSMUSG( | 0.510142 | 0.644476 | NO | NO | NA |
| Olfr1310 | ENSMUSG( | ENSMUSG( | 0.360089 | 0.755908 | NO | NO | NA |
| B230112I2 | ENSMUSG( | ENSMUSG( | 1.153034 | 0.320047 | NO | NO | NA |
| Gm45264 | ENSMUSG( | ENSMUSG( | 1.378343 | 0.284435 | NO | NO | NA |
| Gm45495 | ENSMUSG( | ENSMUSG( | -0.08628 | 0.872516 | NO | NO | NA |
| D730045B | ENSMUSG( | ENSMUSG( | 0.856624 | 0.11648 | NO | NO | NA |
| ENSMUSG | ENSMUSG( | ENSMUSG( | 1.053819 | 0.283212 | NO | NO | NA |

| | | | | | | | |
|---|---|---|---|---|---|---|---|
| Gm43567 | ENSMUSG( | ENSMUSG( | 1.374806 | 0.346147 | NO | NO | NA |
| Insc | ENSMUSG( | ENSMUSG( | 1.147117 | 0.255296 | NO | NO | NA |
| Enpp3 | ENSMUSG( | ENSMUSG( | 0.353128 | 0.722881 | NO | NO | NA |
| 93301620: | ENSMUSG( | ENSMUSG( | 0.004584 | 0.995981 | NO | NO | NA |
| Actl6b | ENSMUSG( | ENSMUSG( | 0.477919 | 0.672193 | NO | NO | NA |
| Gm7450 | ENSMUSG( | ENSMUSG( | 0.734294 | 0.494121 | NO | NO | NA |
| Gm43719 | ENSMUSG( | ENSMUSG( | 1.434272 | 0.173088 | NO | NO | NA |
| Gm20383 | ENSMUSG( | ENSMUSG( | 0.294586 | 0.787014 | NO | NO | NA |
| Gm34776 | ENSMUSG( | ENSMUSG( | 1.006563 | 0.436798 | NO | NO | NA |
| Cmtm8 | ENSMUSG( | ENSMUSG( | 1.210822 | 0.272959 | NO | NO | NA |
| Rapgef4 | ENSMUSG( | ENSMUSG( | 0.192233 | 0.774294 | NO | NO | NA |
| Platr8 | ENSMUSG( | ENSMUSG( | 0.681455 | 0.529352 | NO | NO | NA |
| Cox7a1 | ENSMUSG( | ENSMUSG( | -0.18396 | 0.729543 | NO | NO | NA |
| Gm27029 | ENSMUSG( | ENSMUSG( | -0.15944 | 0.749922 | NO | NO | NA |
| Hes1 | ENSMUSG( | ENSMUSG( | 0.877093 | 0.053551 | NO | NO | NA |
| Gm14421 | ENSMUSG( | ENSMUSG( | -0.04707 | 0.92243 | NO | NO | NA |
| Gm45843 | ENSMUSG( | ENSMUSG( | 0.726086 | 0.500028 | NO | NO | NA |
| ENSMUSG( | ENSMUSG( | ENSMUSG( | 1.989964 | 0.001711 | YES | YES | Upregulated |
| Olfr1122 | ENSMUSG( | ENSMUSG( | 0.309963 | 0.758216 | NO | NO | NA |
| Olfr806 | ENSMUSG( | ENSMUSG( | 0.993789 | 0.351171 | NO | NO | NA |
| Mmd2 | ENSMUSG( | ENSMUSG( | 0.334183 | 0.798465 | NO | NO | NA |
| Gm47580 | ENSMUSG( | ENSMUSG( | 0.521311 | 0.311025 | NO | NO | NA |
| Olfr1500 | ENSMUSG( | ENSMUSG( | 0.51459 | 0.63481 | NO | NO | NA |
| Nxnl2 | ENSMUSG( | ENSMUSG( | 0.15106 | 0.798585 | NO | NO | NA |
| Peli2 | ENSMUSG( | ENSMUSG( | 0.112341 | 0.912705 | NO | NO | NA |
| Olfr1264 | ENSMUSG( | ENSMUSG( | 0.552899 | 0.617076 | NO | NO | NA |
| Oaf | ENSMUSG( | ENSMUSG( | 1.013238 | 0.27261 | NO | NO | NA |
| Gm48565 | ENSMUSG( | ENSMUSG( | 0.548186 | 0.590342 | NO | NO | NA |
| Gm13904 | ENSMUSG( | ENSMUSG( | 2.433919 | 0.001827 | YES | YES | Upregulated |
| Gm3289 | ENSMUSG( | ENSMUSG( | 1.010001 | 0.391549 | NO | NO | NA |
| Gm29590 | ENSMUSG( | ENSMUSG( | 0.658367 | 0.587391 | NO | NO | NA |
| Usp13 | ENSMUSG( | ENSMUSG( | 0.724921 | 0.550149 | NO | NO | NA |
| Il31ra | ENSMUSG( | ENSMUSG( | 1.876531 | 0.000115 | YES | YES | Upregulated |
| ENSMUSG | ENSMUSG( | ENSMUSG( | 0.199535 | 0.582357 | NO | NO | NA |
| 1700069B( | ENSMUSG( | ENSMUSG( | 0.05591 | 0.914344 | NO | NO | NA |
| Gm22513 | ENSMUSG( | ENSMUSG( | 1.615717 | 0.106295 | NO | NO | NA |
| Gm28731 | ENSMUSG( | ENSMUSG( | 0.184046 | 0.597141 | NO | NO | NA |
| Hoxb5 | ENSMUSG( | ENSMUSG( | 0.768467 | 0.466943 | NO | NO | NA |
| Gm13912 | ENSMUSG( | ENSMUSG( | 1.291742 | 0.169963 | NO | NO | NA |
| Mir124a-1 | ENSMUSG( | ENSMUSG( | 0.180711 | 0.87593 | NO | NO | NA |
| Gm38263 | ENSMUSG( | ENSMUSG( | 0.174871 | 0.889049 | NO | NO | NA |
| Obsl1 | ENSMUSG( | ENSMUSG( | 0.341091 | 0.651851 | NO | NO | NA |
| Gm10327 | ENSMUSG( | ENSMUSG( | 0.694334 | 0.238877 | NO | NO | NA |
| Nsa2-ps1 | ENSMUSG( | ENSMUSG( | 0.278966 | 0.51563 | NO | NO | NA |
| Gm43085 | ENSMUSG( | ENSMUSG( | 0.932746 | 0.374884 | NO | NO | NA |
| Trbv23 | ENSMUSG( | ENSMUSG( | -1.19179 | 0.349561 | NO | NO | NA |
| Grik2 | ENSMUSG( | ENSMUSG( | 0.696285 | 0.504319 | NO | NO | NA |

| | | | | | | | |
|---|---|---|---|---|---|---|---|
| Gm44090 | ENSMUSG( | ENSMUSG( | 0.545031 | 0.607428 | NO | NO | NA |
| Gm37363 | ENSMUSG( | ENSMUSG( | -0.02796 | 0.980563 | NO | NO | NA |
| ENSMUSG | ENSMUSG( | ENSMUSG( | 0.763845 | 0.502577 | NO | NO | NA |
| Tmem200 | ENSMUSG( | ENSMUSG( | 0.573515 | 0.568223 | NO | NO | NA |
| Gm30214 | ENSMUSG( | ENSMUSG( | 0.210587 | 0.660848 | NO | NO | NA |
| Gm7645 | ENSMUSG( | ENSMUSG( | -0.75645 | 0.105858 | NO | NO | NA |
| Exd1 | ENSMUSG( | ENSMUSG( | 0.570994 | 0.327392 | NO | NO | NA |
| 1110046J( | ENSMUSG( | ENSMUSG( | 0.58119 | 0.370733 | NO | NO | NA |
| Gm42535 | ENSMUSG( | ENSMUSG( | 0.467323 | 0.658738 | NO | NO | NA |
| Gm17982 | ENSMUSG( | ENSMUSG( | 1.523434 | 0.120789 | NO | NO | NA |
| Dmrt1 | ENSMUSG( | ENSMUSG( | 1.019461 | 0.317944 | NO | NO | NA |
| Cylc2 | ENSMUSG( | ENSMUSG( | 0.534782 | 0.574769 | NO | NO | NA |
| Pram1 | ENSMUSG( | ENSMUSG( | 0.954637 | 0.668205 | NO | NO | NA |
| Ppfia2 | ENSMUSG( | ENSMUSG( | 0.651991 | 0.539934 | NO | NO | NA |
| Cfap43 | ENSMUSG( | ENSMUSG( | 0.400072 | 0.285761 | NO | NO | NA |
| ENSMUSG | ENSMUSG( | ENSMUSG( | -0.49427 | 0.469238 | NO | NO | NA |
| Aqp9 | ENSMUSG( | ENSMUSG( | 0.345444 | 0.609219 | NO | NO | NA |
| Gm21182 | ENSMUSG( | ENSMUSG( | 1.787319 | 0.059102 | NO | NO | NA |
| Abcc3 | ENSMUSG( | ENSMUSG( | 0.328308 | 0.698075 | NO | NO | NA |
| Zfp608 | ENSMUSG( | ENSMUSG( | 0.366684 | 0.685613 | NO | NO | NA |
| A330009N | ENSMUSG( | ENSMUSG( | -0.52093 | 0.117208 | NO | NO | NA |
| C430049B( | ENSMUSG( | ENSMUSG( | -0.0067 | 0.991949 | NO | NO | NA |
| Cer1 | ENSMUSG( | ENSMUSG( | 1.106785 | 0.502435 | NO | NO | NA |
| Ighv1-36 | ENSMUSG( | ENSMUSG( | 4.849797 | NA | NO | NO | NA |
| Olfr50 | ENSMUSG( | ENSMUSG( | 0.093331 | 0.928982 | NO | NO | NA |
| ENSMUSG | ENSMUSG( | ENSMUSG( | -0.05175 | 0.92464 | NO | NO | NA |
| Ankmy1 | ENSMUSG( | ENSMUSG( | 0.544249 | 0.342268 | NO | NO | NA |
| Gm42508 | ENSMUSG( | ENSMUSG( | 0.579723 | 0.604254 | NO | NO | NA |
| Gm30124 | ENSMUSG( | ENSMUSG( | 0.500871 | 0.322298 | NO | NO | NA |
| Gm49949 | ENSMUSG( | ENSMUSG( | 0.709646 | 0.494674 | NO | NO | NA |
| Psapl1 | ENSMUSG( | ENSMUSG( | 0.58257 | 0.631598 | NO | NO | NA |
| Igsf11 | ENSMUSG( | ENSMUSG( | 0.424827 | 0.641748 | NO | NO | NA |
| Olfr1211 | ENSMUSG( | ENSMUSG( | 0.609427 | 0.573488 | NO | NO | NA |
| Zfp516 | ENSMUSG( | ENSMUSG( | 0.398126 | 0.307151 | NO | NO | NA |
| Gm10484 | ENSMUSG( | ENSMUSG( | 0.293103 | 0.789558 | NO | NO | NA |
| Gm6643 | ENSMUSG( | ENSMUSG( | 0.441503 | 0.719186 | NO | NO | NA |
| Ybx1-ps2 | ENSMUSG( | ENSMUSG( | 0.298442 | 0.492894 | NO | NO | NA |
| Gm44986 | ENSMUSG( | ENSMUSG( | -0.01705 | 0.972626 | NO | NO | NA |
| Gm11342 | ENSMUSG( | ENSMUSG( | 1.132406 | 0.317406 | NO | NO | NA |
| Spire2 | ENSMUSG( | ENSMUSG( | 0.261879 | 0.63481 | NO | NO | NA |
| Gm48181 | ENSMUSG( | ENSMUSG( | 0.346632 | 0.727703 | NO | NO | NA |
| Gm44949 | ENSMUSG( | ENSMUSG( | 0.733894 | 0.196454 | NO | NO | NA |
| Hoxb8 | ENSMUSG( | ENSMUSG( | 1.264786 | 0.254124 | NO | NO | NA |
| Olfr348 | ENSMUSG( | ENSMUSG( | 0.568883 | 0.593913 | NO | NO | NA |
| 1500012K( | ENSMUSG( | ENSMUSG( | 0.347111 | 0.749795 | NO | NO | NA |
| Gpr25 | ENSMUSG( | ENSMUSG( | 1.02163 | 0.221618 | NO | NO | NA |
| Rab38 | ENSMUSG( | ENSMUSG( | 1.036529 | 0.140741 | NO | NO | NA |

| | | | | | | | |
|---|---|---|---|---|---|---|---|
| Rln1 | ENSMUSG( | ENSMUSG( | 0.147273 | 0.898348 | NO | NO | NA |
| Gm3222 | ENSMUSG( | ENSMUSG( | 0.401377 | 0.474914 | NO | NO | NA |
| Gm7891 | ENSMUSG( | ENSMUSG( | 0.908428 | 0.421499 | NO | NO | NA |
| Gm12854 | ENSMUSG( | ENSMUSG( | -0.00646 | 0.989896 | NO | NO | NA |
| Galnt16 | ENSMUSG( | ENSMUSG( | 1.162686 | 0.218747 | NO | NO | NA |
| Gm43588 | ENSMUSG( | ENSMUSG( | 0.358236 | 0.569784 | NO | NO | NA |
| Atp6v1b1 | ENSMUSG( | ENSMUSG( | -0.04235 | 0.938509 | NO | NO | NA |
| Olfr204 | ENSMUSG( | ENSMUSG( | 0.346581 | 0.755183 | NO | NO | NA |
| Syngr4 | ENSMUSG( | ENSMUSG( | 0.3194 | 0.576131 | NO | NO | NA |
| Gm10802 | ENSMUSG( | ENSMUSG( | 0.085109 | 0.905316 | NO | NO | NA |
| Doc2a | ENSMUSG( | ENSMUSG( | 0.593445 | 0.630748 | NO | NO | NA |
| 1500002F: | ENSMUSG( | ENSMUSG( | 0.449256 | 0.378491 | NO | NO | NA |
| Vmn2r28 | ENSMUSG( | ENSMUSG( | 1.198775 | 0.236514 | NO | NO | NA |
| Vmn1r57 | ENSMUSG( | ENSMUSG( | 0.311491 | 0.769876 | NO | NO | NA |
| Gm40773 | ENSMUSG( | ENSMUSG( | -0.09614 | 0.850268 | NO | NO | NA |
| Adgrd1 | ENSMUSG( | ENSMUSG( | -0.14119 | 0.720821 | NO | NO | NA |
| ENSMUSG | ENSMUSG( | ENSMUSG( | -0.81874 | 0.013444 | YES | NO | NA |
| Tunar | ENSMUSG( | ENSMUSG( | 0.254178 | 0.8014 | NO | NO | NA |
| B430319G | ENSMUSG( | ENSMUSG( | 0.084824 | 0.895467 | NO | NO | NA |
| Atg4c | ENSMUSG( | ENSMUSG( | 1.165628 | 0.028847 | YES | YES | Upregulated |
| ENSMUSG( | ENSMUSG( | ENSMUSG( | 0.951017 | 0.50674 | NO | NO | NA |
| Ly6k | ENSMUSG( | ENSMUSG( | 1.305403 | 0.00384 | YES | YES | Upregulated |
| Gm45779 | ENSMUSG( | ENSMUSG( | -0.27268 | 0.607428 | NO | NO | NA |
| ENSMUSG | ENSMUSG( | ENSMUSG( | -0.0348 | 0.930548 | NO | NO | NA |
| Gm36660 | ENSMUSG( | ENSMUSG( | 0.121133 | 0.913916 | NO | NO | NA |
| Gm42726 | ENSMUSG( | ENSMUSG( | 0.074759 | 0.880165 | NO | NO | NA |
| Gucy2f | ENSMUSG( | ENSMUSG( | 0.588036 | 0.544632 | NO | NO | NA |
| Gm13219 | ENSMUSG( | ENSMUSG( | 1.180655 | 0.327607 | NO | NO | NA |
| ENSMUSG( | ENSMUSG( | ENSMUSG( | 0.249111 | 0.501922 | NO | NO | NA |
| AY512931 | ENSMUSG( | ENSMUSG( | 0.757466 | 0.484647 | NO | NO | NA |
| Gm4654 | ENSMUSG( | ENSMUSG( | 0.345559 | 0.60722 | NO | NO | NA |
| Gm43870 | ENSMUSG( | ENSMUSG( | 0.904581 | 0.42412 | NO | NO | NA |
| Gm42494 | ENSMUSG( | ENSMUSG( | 1.105003 | 0.327267 | NO | NO | NA |
| Gng4 | ENSMUSG( | ENSMUSG( | 0.644062 | 0.55012 | NO | NO | NA |
| Gm9521 | ENSMUSG( | ENSMUSG( | 0.590434 | 0.655685 | NO | NO | NA |
| Gm17189 | ENSMUSG( | ENSMUSG( | 0.581389 | 0.593442 | NO | NO | NA |
| 9930024M | ENSMUSG( | ENSMUSG( | -0.12291 | 0.826879 | NO | NO | NA |
| Gm45161 | ENSMUSG( | ENSMUSG( | 1.135967 | 0.327825 | NO | NO | NA |
| Hlx | ENSMUSG( | ENSMUSG( | -0.23279 | 0.643027 | NO | NO | NA |
| C1ra | ENSMUSG( | ENSMUSG( | -0.00017 | 0.999756 | NO | NO | NA |
| Tspyl5 | ENSMUSG( | ENSMUSG( | 0.554888 | 0.584088 | NO | NO | NA |
| Gm38265 | ENSMUSG( | ENSMUSG( | 0.826169 | 0.504087 | NO | NO | NA |
| Veph1 | ENSMUSG( | ENSMUSG( | 0.624196 | 0.616598 | NO | NO | NA |
| Cgn | ENSMUSG( | ENSMUSG( | 0.215998 | 0.776875 | NO | NO | NA |
| Gm12033 | ENSMUSG( | ENSMUSG( | 0.105051 | 0.912224 | NO | NO | NA |
| Gm22146 | ENSMUSG( | ENSMUSG( | 0.540792 | 0.437355 | NO | NO | NA |
| Myh8 | ENSMUSG( | ENSMUSG( | 0.889883 | 0.443505 | NO | NO | NA |

| | | | | | | | |
|---|---|---|---|---|---|---|---|
| Mexis | ENSMUSG( | ENSMUSG( | 0.930132 | 0.230307 | NO | NO | NA |
| Gm3839 | ENSMUSG( | ENSMUSG( | 0.331646 | 0.527031 | NO | NO | NA |
| Gm48880 | ENSMUSG( | ENSMUSG( | 0.26207 | 0.698511 | NO | NO | NA |
| Gm26787 | ENSMUSG( | ENSMUSG( | 0.288782 | 0.62406 | NO | NO | NA |
| 5430425E: | ENSMUSG( | ENSMUSG( | 1.277894 | 0.185594 | NO | NO | NA |
| Olfr279 | ENSMUSG( | ENSMUSG( | 0.367229 | 0.725594 | NO | NO | NA |
| Fam241b | ENSMUSG( | ENSMUSG( | 0.133854 | 0.752157 | NO | NO | NA |
| 4933417C: | ENSMUSG( | ENSMUSG( | 0.981823 | 0.092061 | NO | NO | NA |
| Nrk | ENSMUSG( | ENSMUSG( | 0.981212 | 0.283254 | NO | NO | NA |
| Gm48438 | ENSMUSG( | ENSMUSG( | 0.971469 | 0.354123 | NO | NO | NA |
| Gm36500 | ENSMUSG( | ENSMUSG( | 1.397341 | 0.186976 | NO | NO | NA |
| Pantr1 | ENSMUSG( | ENSMUSG( | 0.474156 | 0.668523 | NO | NO | NA |
| Olfr501-ps | ENSMUSG( | ENSMUSG( | 0.856725 | 0.431414 | NO | NO | NA |
| Gm12474 | ENSMUSG( | ENSMUSG( | -0.03016 | 0.958708 | NO | NO | NA |
| Gm44443 | ENSMUSG( | ENSMUSG( | 0.459427 | 0.517331 | NO | NO | NA |
| AA986860 | ENSMUSG( | ENSMUSG( | 0.089592 | 0.876094 | NO | NO | NA |
| Thbs1 | ENSMUSG( | ENSMUSG( | 0.344444 | 0.658279 | NO | NO | NA |
| Zfp970 | ENSMUSG( | ENSMUSG( | 0.248402 | 0.538519 | NO | NO | NA |
| Iglc2 | ENSMUSG( | ENSMUSG( | -1.6106 | 0.050582 | NO | NO | NA |
| Zfp768 | ENSMUSG( | ENSMUSG( | 1.972434 | 0.000805 | YES | YES | Upregulated |
| Gm12802 | ENSMUSG( | ENSMUSG( | 0.009365 | 0.994611 | NO | NO | NA |
| Fggy | ENSMUSG( | ENSMUSG( | 1.857361 | 0.003446 | YES | YES | Upregulated |
| Gm49087 | ENSMUSG( | ENSMUSG( | 0.842101 | 0.439849 | NO | NO | NA |
| Olfr728 | ENSMUSG( | ENSMUSG( | 0.647937 | 0.566217 | NO | NO | NA |
| Gm14176 | ENSMUSG( | ENSMUSG( | -0.52266 | 0.338094 | NO | NO | NA |
| Flrt3 | ENSMUSG( | ENSMUSG( | 1.199532 | 0.2471 | NO | NO | NA |
| Gm37376 | ENSMUSG( | ENSMUSG( | -0.98243 | 0.048196 | YES | NO | NA |
| Fam83e | ENSMUSG( | ENSMUSG( | 0.725781 | 0.202806 | NO | NO | NA |
| Gm20429 | ENSMUSG( | ENSMUSG( | 0.72299 | 0.527649 | NO | NO | NA |
| Ighv1-22 | ENSMUSG( | ENSMUSG( | -0.71691 | 0.719066 | NO | NO | NA |
| Sptbn5 | ENSMUSG( | ENSMUSG( | -0.81335 | 0.053915 | NO | NO | NA |
| Gm47574 | ENSMUSG( | ENSMUSG( | 1.208454 | 0.01444 | YES | YES | Upregulated |
| 4930519G | ENSMUSG( | ENSMUSG( | 0.712851 | 0.52971 | NO | NO | NA |
| Aldh1a3 | ENSMUSG( | ENSMUSG( | 0.121389 | 0.775138 | NO | NO | NA |
| Gm37266 | ENSMUSG( | ENSMUSG( | 0.678321 | 0.32874 | NO | NO | NA |
| Olfr31 | ENSMUSG( | ENSMUSG( | 0.348607 | 0.737991 | NO | NO | NA |
| Gm37847 | ENSMUSG( | ENSMUSG( | 1.953759 | 0.211846 | NO | NO | NA |
| Gm10115 | ENSMUSG( | ENSMUSG( | 1.432544 | 0.224204 | NO | NO | NA |
| Gm43173 | ENSMUSG( | ENSMUSG( | -0.02429 | 0.962213 | NO | NO | NA |
| R74862 | ENSMUSG( | ENSMUSG( | 0.069311 | 0.85784 | NO | NO | NA |
| Gm37806 | ENSMUSG( | ENSMUSG( | 0.648284 | 0.595012 | NO | NO | NA |
| Tmem79 | ENSMUSG( | ENSMUSG( | 0.240191 | 0.47574 | NO | NO | NA |
| Brme1 | ENSMUSG( | ENSMUSG( | -0.03856 | 0.926387 | NO | NO | NA |
| 5730521K( | ENSMUSG( | ENSMUSG( | 0.45075 | 0.164842 | NO | NO | NA |
| Ctnnd1 | ENSMUSG( | ENSMUSG( | -0.2207 | 0.514914 | NO | NO | NA |
| Gm12870 | ENSMUSG( | ENSMUSG( | 0.214822 | 0.606154 | NO | NO | NA |
| Olfr1293-p | ENSMUSG( | ENSMUSG( | 1.022143 | 0.363159 | NO | NO | NA |

| | | | | | | | |
|---|---|---|---|---|---|---|---|
| Gm48194 | ENSMUSG( | ENSMUSG( | -0.28847 | 0.414327 | NO | NO | NA |
| Olfr1454 | ENSMUSG( | ENSMUSG( | 0.525767 | 0.679576 | NO | NO | NA |
| Paqr5 | ENSMUSG( | ENSMUSG( | 0.301009 | 0.636205 | NO | NO | NA |
| Olfr1042 | ENSMUSG( | ENSMUSG( | 1.538549 | 0.110439 | NO | NO | NA |
| Gm6566 | ENSMUSG( | ENSMUSG( | 0.44484 | 0.368501 | NO | NO | NA |
| Gm43807 | ENSMUSG( | ENSMUSG( | 0.536382 | 0.612377 | NO | NO | NA |
| Gm10432 | ENSMUSG( | ENSMUSG( | -0.80272 | 0.026754 | YES | NO | NA |
| Trav13-3 | ENSMUSG( | ENSMUSG( | 0.799899 | 0.551005 | NO | NO | NA |
| Gm12493 | ENSMUSG( | ENSMUSG( | 0.71289 | 0.492717 | NO | NO | NA |
| Adamts7 | ENSMUSG( | ENSMUSG( | -0.36383 | 0.432483 | NO | NO | NA |
| Gm34297 | ENSMUSG( | ENSMUSG( | 0.812139 | 0.476489 | NO | NO | NA |
| Dynlt2a1 | ENSMUSG( | ENSMUSG( | 2.342978 | 0.053783 | NO | NO | NA |
| Fam114a1 | ENSMUSG( | ENSMUSG( | -0.63784 | 0.066122 | NO | NO | NA |
| Gm32999 | ENSMUSG( | ENSMUSG( | -0.29424 | 0.522162 | NO | NO | NA |
| H2-T3 | ENSMUSG( | ENSMUSG( | 0.611371 | 0.570158 | NO | NO | NA |
| Iglc3 | ENSMUSG( | ENSMUSG( | -1.28956 | 0.306823 | NO | NO | NA |
| Prss27 | ENSMUSG( | ENSMUSG( | 0.116985 | 0.932144 | NO | NO | NA |
| Gm50215 | ENSMUSG( | ENSMUSG( | 0.640477 | 0.547478 | NO | NO | NA |
| Acsm2 | ENSMUSG( | ENSMUSG( | 0.804212 | 0.438742 | NO | NO | NA |
| Dlx6os1 | ENSMUSG( | ENSMUSG( | 0.446901 | 0.677675 | NO | NO | NA |
| Gm11231 | ENSMUSG( | ENSMUSG( | 0.676036 | 0.523861 | NO | NO | NA |
| Ptprz1 | ENSMUSG( | ENSMUSG( | 1.271115 | 0.249389 | NO | NO | NA |
| 4933436H | ENSMUSG( | ENSMUSG( | 0.652676 | 0.557764 | NO | NO | NA |
| Gm13722 | ENSMUSG( | ENSMUSG( | 0.551608 | 0.613549 | NO | NO | NA |
| Celf6 | ENSMUSG( | ENSMUSG( | 0.619735 | 0.505715 | NO | NO | NA |
| Olfr224 | ENSMUSG( | ENSMUSG( | 0.693515 | 0.590752 | NO | NO | NA |
| Clec11a | ENSMUSG( | ENSMUSG( | 1.865512 | 8.31E-05 | YES | YES | Upregulated |
| Fam131a | ENSMUSG( | ENSMUSG( | -0.72624 | 0.062635 | NO | NO | NA |
| Vmn1r234 | ENSMUSG( | ENSMUSG( | 0.397923 | 0.70101 | NO | NO | NA |
| Gm37811 | ENSMUSG( | ENSMUSG( | 0.694005 | 0.567382 | NO | NO | NA |
| Gm16010 | ENSMUSG( | ENSMUSG( | 0.436714 | 0.701109 | NO | NO | NA |
| Gm43713 | ENSMUSG( | ENSMUSG( | 0.046043 | 0.932228 | NO | NO | NA |
| 4833422M | ENSMUSG( | ENSMUSG( | 0.188706 | 0.86613 | NO | NO | NA |
| G730003C | ENSMUSG( | ENSMUSG( | 0.990867 | 0.07724 | NO | NO | NA |
| Gm49201 | ENSMUSG( | ENSMUSG( | 0.094107 | 0.89162 | NO | NO | NA |
| Gm43748 | ENSMUSG( | ENSMUSG( | 0.874371 | 0.349712 | NO | NO | NA |
| Scarna2 | ENSMUSG( | ENSMUSG( | 0.161643 | 0.753653 | NO | NO | NA |
| 2510017J1 | ENSMUSG( | ENSMUSG( | 0.549045 | 0.185248 | NO | NO | NA |
| Gm16244 | ENSMUSG( | ENSMUSG( | -0.35035 | 0.43299 | NO | NO | NA |
| Gm15631 | ENSMUSG( | ENSMUSG( | 0.463416 | 0.707721 | NO | NO | NA |
| Gckr | ENSMUSG( | ENSMUSG( | 0.293057 | 0.789276 | NO | NO | NA |
| Zfp820 | ENSMUSG( | ENSMUSG( | -0.37817 | 0.491417 | NO | NO | NA |
| Gm6117 | ENSMUSG( | ENSMUSG( | 1.353906 | 0.113006 | NO | NO | NA |
| Dcst1 | ENSMUSG( | ENSMUSG( | 0.562274 | 0.26752 | NO | NO | NA |
| Gm13166 | ENSMUSG( | ENSMUSG( | 1.410029 | 0.008858 | YES | YES | Upregulated |
| Gm47968 | ENSMUSG( | ENSMUSG( | 0.650182 | 0.549318 | NO | NO | NA |
| Gm45719 | ENSMUSG( | ENSMUSG( | 0.504638 | 0.220371 | NO | NO | NA |

| | | | | | | | |
|---|---|---|---|---|---|---|---|
| ENSMUSG | ENSMUSG( | ENSMUSG( | 0.380173 | 0.496489 | NO | NO | NA |
| Id3 | ENSMUSG( | ENSMUSG( | 0.11506 | 0.87566 | NO | NO | NA |
| Luzp2 | ENSMUSG( | ENSMUSG( | 0.94309 | 0.347889 | NO | NO | NA |
| Pik3r3 | ENSMUSG( | ENSMUSG( | -0.13741 | 0.794925 | NO | NO | NA |
| Gmfg-ps | ENSMUSG( | ENSMUSG( | -0.01457 | 0.966629 | NO | NO | NA |
| Gm42640 | ENSMUSG( | ENSMUSG( | 0.805141 | 0.477689 | NO | NO | NA |
| Gm10472 | ENSMUSG( | ENSMUSG( | 0.624738 | 0.548283 | NO | NO | NA |
| Cxcl16 | ENSMUSG( | ENSMUSG( | 0.284614 | 0.594481 | NO | NO | NA |
| Pou1f1 | ENSMUSG( | ENSMUSG( | 0.567036 | 0.557578 | NO | NO | NA |
| Scn11a | ENSMUSG( | ENSMUSG( | -0.4106 | 0.426029 | NO | NO | NA |
| Col4a4 | ENSMUSG( | ENSMUSG( | 0.573025 | 0.476823 | NO | NO | NA |
| C230071H | ENSMUSG( | ENSMUSG( | 0.461118 | 0.653824 | NO | NO | NA |
| Trav8-2 | ENSMUSG( | ENSMUSG( | -0.10919 | 0.841744 | NO | NO | NA |
| Enox1 | ENSMUSG( | ENSMUSG( | 0.975922 | 0.283158 | NO | NO | NA |
| S100b | ENSMUSG( | ENSMUSG( | 0.927327 | 0.481807 | NO | NO | NA |
| ENSMUSG | ENSMUSG( | ENSMUSG( | -0.12955 | 0.759918 | NO | NO | NA |
| Vmn2r11 | ENSMUSG( | ENSMUSG( | 0.948922 | 0.370227 | NO | NO | NA |
| Zfp972 | ENSMUSG( | ENSMUSG( | 0.219956 | 0.642396 | NO | NO | NA |
| A730089K | ENSMUSG( | ENSMUSG( | 1.21211 | 0.064485 | NO | NO | NA |
| Gm7754 | ENSMUSG( | ENSMUSG( | 1.046174 | 0.346493 | NO | NO | NA |
| Gm43341 | ENSMUSG( | ENSMUSG( | 1.505175 | 0.161751 | NO | NO | NA |
| Ttc26 | ENSMUSG( | ENSMUSG( | 1.104967 | 0.197011 | NO | NO | NA |
| ENSMUSG( | ENSMUSG( | ENSMUSG( | -0.26509 | 0.572582 | NO | NO | NA |
| 4930551O | ENSMUSG( | ENSMUSG( | 0.516549 | 0.591796 | NO | NO | NA |
| Lrp1b | ENSMUSG( | ENSMUSG( | 0.775054 | 0.488965 | NO | NO | NA |
| Gm14812 | ENSMUSG( | ENSMUSG( | 0.742828 | 0.526045 | NO | NO | NA |
| Gpr150 | ENSMUSG( | ENSMUSG( | 0.780608 | 0.319391 | NO | NO | NA |
| Gm14549 | ENSMUSG( | ENSMUSG( | 0.73147 | 0.146688 | NO | NO | NA |
| 4933427D | ENSMUSG( | ENSMUSG( | 0.604194 | 0.526688 | NO | NO | NA |
| Jph1 | ENSMUSG( | ENSMUSG( | 0.340556 | 0.722873 | NO | NO | NA |
| Phf2os1 | ENSMUSG( | ENSMUSG( | 0.873773 | 0.11386 | NO | NO | NA |
| Gm42583 | ENSMUSG( | ENSMUSG( | 0.747798 | 0.478508 | NO | NO | NA |
| ENSMUSG( | ENSMUSG( | ENSMUSG( | -0.05593 | 0.935425 | NO | NO | NA |
| Kif9 | ENSMUSG( | ENSMUSG( | 0.396563 | 0.385467 | NO | NO | NA |
| Ctcfl | ENSMUSG( | ENSMUSG( | 0.898799 | 0.370368 | NO | NO | NA |
| 9130002K: | ENSMUSG( | ENSMUSG( | -0.57399 | 0.071956 | NO | NO | NA |
| Chdh | ENSMUSG( | ENSMUSG( | 0.049273 | 0.962708 | NO | NO | NA |
| Hyal3 | ENSMUSG( | ENSMUSG( | 0.571544 | 0.35751 | NO | NO | NA |
| Gm42449 | ENSMUSG( | ENSMUSG( | -0.08756 | 0.868932 | NO | NO | NA |
| Gm36594 | ENSMUSG( | ENSMUSG( | 0.808212 | 0.492717 | NO | NO | NA |
| Spock1 | ENSMUSG( | ENSMUSG( | 0.555033 | 0.618809 | NO | NO | NA |
| Fam216b | ENSMUSG( | ENSMUSG( | 0.594354 | 0.61095 | NO | NO | NA |
| Prss54 | ENSMUSG( | ENSMUSG( | 1.03946 | 0.341055 | NO | NO | NA |
| Pax7 | ENSMUSG( | ENSMUSG( | 0.578617 | 0.60648 | NO | NO | NA |
| Bambi | ENSMUSG( | ENSMUSG( | 0.574251 | 0.272102 | NO | NO | NA |
| Jag2 | ENSMUSG( | ENSMUSG( | -0.59413 | 0.174565 | NO | NO | NA |
| Gm15793 | ENSMUSG( | ENSMUSG( | 0.858758 | 0.155831 | NO | NO | NA |

| | | | | | | | |
|---|---|---|---|---|---|---|---|
| Gm28047 | ENSMUSG( | ENSMUSG( | 0.932198 | 0.744133 | NO | NO | NA |
| 4933406B: | ENSMUSG( | ENSMUSG( | 0.89649 | 0.439349 | NO | NO | NA |
| Gm12582 | ENSMUSG( | ENSMUSG( | 1.238484 | 0.249052 | NO | NO | NA |
| Olfr784 | ENSMUSG( | ENSMUSG( | 0.071225 | 0.922644 | NO | NO | NA |
| 9330179D | ENSMUSG( | ENSMUSG( | 0.288191 | 0.672142 | NO | NO | NA |
| Plpp3 | ENSMUSG( | ENSMUSG( | 0.687983 | 0.519459 | NO | NO | NA |
| Tnfaip8l3 | ENSMUSG( | ENSMUSG( | 0.992851 | 0.389594 | NO | NO | NA |
| a | ENSMUSG( | ENSMUSG( | 1.574556 | 0.013193 | YES | YES | Upregulated |
| Parva | ENSMUSG( | ENSMUSG( | 0.238956 | 0.843786 | NO | NO | NA |
| Pcdhga6 | ENSMUSG( | ENSMUSG( | 1.591063 | 0.166583 | NO | NO | NA |
| Fer | ENSMUSG( | ENSMUSG( | 0.851077 | 0.118086 | NO | NO | NA |
| Gm10156 | ENSMUSG( | ENSMUSG( | 1.329919 | 0.000363 | YES | YES | Upregulated |
| Gm26646 | ENSMUSG( | ENSMUSG( | 0.765505 | 0.345714 | NO | NO | NA |
| Gm38284 | ENSMUSG( | ENSMUSG( | 0.844656 | 0.450569 | NO | NO | NA |
| Kcnip4 | ENSMUSG( | ENSMUSG( | 0.179352 | 0.683308 | NO | NO | NA |
| Morc2b | ENSMUSG( | ENSMUSG( | 0.465284 | 0.639774 | NO | NO | NA |
| Gm9502 | ENSMUSG( | ENSMUSG( | -0.33191 | 0.648619 | NO | NO | NA |
| Rims2 | ENSMUSG( | ENSMUSG( | 1.8254 | 0.004979 | YES | YES | Upregulated |
| Ugt3a2 | ENSMUSG( | ENSMUSG( | 0.631298 | 0.544325 | NO | NO | NA |
| Gm44045 | ENSMUSG( | ENSMUSG( | 0.383871 | 0.712266 | NO | NO | NA |
| Ddi1 | ENSMUSG( | ENSMUSG( | 0.367312 | 0.721414 | NO | NO | NA |
| Gm37041 | ENSMUSG( | ENSMUSG( | 0.496665 | 0.649471 | NO | NO | NA |
| Gm45066 | ENSMUSG( | ENSMUSG( | 0.187595 | 0.852829 | NO | NO | NA |
| Gm10131 | ENSMUSG( | ENSMUSG( | 0.409016 | 0.268553 | NO | NO | NA |
| 4833408A | ENSMUSG( | ENSMUSG( | 0.076022 | 0.885271 | NO | NO | NA |
| Clec3b | ENSMUSG( | ENSMUSG( | -0.32084 | 0.439849 | NO | NO | NA |
| Gm9978 | ENSMUSG( | ENSMUSG( | 0.765116 | 0.540891 | NO | NO | NA |
| F11r | ENSMUSG( | ENSMUSG( | 0.458433 | 0.430226 | NO | NO | NA |
| Gm16209 | ENSMUSG( | ENSMUSG( | 0.021177 | 0.970413 | NO | NO | NA |
| Gm49714 | ENSMUSG( | ENSMUSG( | 0.963319 | 0.410374 | NO | NO | NA |
| Gm15041 | ENSMUSG( | ENSMUSG( | 1.07033 | 0.452821 | NO | NO | NA |
| Gm8991 | ENSMUSG( | ENSMUSG( | 0.993154 | 0.121489 | NO | NO | NA |
| Gm34481 | ENSMUSG( | ENSMUSG( | 0.695266 | 0.551462 | NO | NO | NA |
| Gm7962 | ENSMUSG( | ENSMUSG( | 0.45673 | 0.52971 | NO | NO | NA |
| Ighv10-1 | ENSMUSG( | ENSMUSG( | -1.15654 | 0.73647 | NO | NO | NA |
| Rgs5 | ENSMUSG( | ENSMUSG( | 0.679218 | 0.604896 | NO | NO | NA |
| Gm38170 | ENSMUSG( | ENSMUSG( | 0.173813 | 0.869991 | NO | NO | NA |
| Btf3l4b | ENSMUSG( | ENSMUSG( | 0.180163 | 0.839614 | NO | NO | NA |
| 4931415C: | ENSMUSG( | ENSMUSG( | 0.113509 | 0.83253 | NO | NO | NA |
| Car10 | ENSMUSG( | ENSMUSG( | 0.387645 | 0.713817 | NO | NO | NA |
| ENSMUSG( | ENSMUSG( | ENSMUSG( | 0.532598 | 0.554294 | NO | NO | NA |
| Gm18649 | ENSMUSG( | ENSMUSG( | 0.102965 | 0.868704 | NO | NO | NA |
| Khdrbs3 | ENSMUSG( | ENSMUSG( | 0.301583 | 0.731835 | NO | NO | NA |
| Osr1 | ENSMUSG( | ENSMUSG( | 1.054305 | 0.325898 | NO | NO | NA |
| Kl | ENSMUSG( | ENSMUSG( | 0.25017 | 0.802185 | NO | NO | NA |
| Gm19972 | ENSMUSG( | ENSMUSG( | 0.128316 | 0.900004 | NO | NO | NA |
| 4930452N | ENSMUSG( | ENSMUSG( | 1.157979 | 0.272959 | NO | NO | NA |

| | | | | | | | |
|---|---|---|---|---|---|---|---|
| Gm45527 | ENSMUSG( | ENSMUSG( | -0.5732 | 0.330466 | NO | NO | NA |
| Slc5a8 | ENSMUSG( | ENSMUSG( | 0.708003 | 0.534341 | NO | NO | NA |
| T2 | ENSMUSG( | ENSMUSG( | -0.93917 | 0.095933 | NO | NO | NA |
| Gm17969 | ENSMUSG( | ENSMUSG( | 0.621713 | 0.5474 | NO | NO | NA |
| Gm50080 | ENSMUSG( | ENSMUSG( | 0.699315 | 0.49711 | NO | NO | NA |
| Rtl4 | ENSMUSG( | ENSMUSG( | 0.78239 | 0.482948 | NO | NO | NA |
| Gm48967 | ENSMUSG( | ENSMUSG( | 0.373635 | 0.722171 | NO | NO | NA |
| Gm38355 | ENSMUSG( | ENSMUSG( | 0.624751 | 0.319978 | NO | NO | NA |
| Gm26811 | ENSMUSG( | ENSMUSG( | 0.556914 | 0.589513 | NO | NO | NA |
| Lincmd1 | ENSMUSG( | ENSMUSG( | 0.531069 | 0.615925 | NO | NO | NA |
| 1700023B: | ENSMUSG( | ENSMUSG( | 1.219652 | 0.129928 | NO | NO | NA |
| Prr22 | ENSMUSG( | ENSMUSG( | 0.341348 | 0.446235 | NO | NO | NA |
| Trav15d-1- | ENSMUSG( | ENSMUSG( | 0.224559 | 0.878139 | NO | NO | NA |
| Map10 | ENSMUSG( | ENSMUSG( | 0.178207 | 0.760732 | NO | NO | NA |
| Gm4294 | ENSMUSG( | ENSMUSG( | -0.42942 | 0.518523 | NO | NO | NA |
| Vmn1r70 | ENSMUSG( | ENSMUSG( | 0.63487 | 0.588985 | NO | NO | NA |
| Gm15408 | ENSMUSG( | ENSMUSG( | 0.505491 | 0.614624 | NO | NO | NA |
| Vmn1r211 | ENSMUSG( | ENSMUSG( | 1.075236 | 0.301393 | NO | NO | NA |
| Inava | ENSMUSG( | ENSMUSG( | 0.11139 | 0.878462 | NO | NO | NA |
| Gm43045 | ENSMUSG( | ENSMUSG( | 1.123833 | 0.30414 | NO | NO | NA |
| A730061H | ENSMUSG( | ENSMUSG( | -0.54059 | 0.207477 | NO | NO | NA |
| Gm43715 | ENSMUSG( | ENSMUSG( | 0.99274 | 0.402024 | NO | NO | NA |
| Olfr1497 | ENSMUSG( | ENSMUSG( | 0.855724 | 0.471303 | NO | NO | NA |
| Slc11a1 | ENSMUSG( | ENSMUSG( | -0.13261 | 0.775918 | NO | NO | NA |
| Dnajc28 | ENSMUSG( | ENSMUSG( | -1.33576 | 0.000206 | YES | YES | Downregulated |
| Gm49975 | ENSMUSG( | ENSMUSG( | 0.494785 | 0.558028 | NO | NO | NA |
| Gm15533 | ENSMUSG( | ENSMUSG( | -1.69489 | 1.80E-06 | YES | YES | Downregulated |
| Shank3 | ENSMUSG( | ENSMUSG( | -0.37636 | 0.569018 | NO | NO | NA |
| Pld6 | ENSMUSG( | ENSMUSG( | 0.106017 | 0.901599 | NO | NO | NA |
| Gm48551 | ENSMUSG( | ENSMUSG( | 0.064376 | 0.873681 | NO | NO | NA |
| Prob1 | ENSMUSG( | ENSMUSG( | 0.212161 | 0.664907 | NO | NO | NA |
| Gm37477 | ENSMUSG( | ENSMUSG( | 0.932324 | 0.457961 | NO | NO | NA |
| Gm29689 | ENSMUSG( | ENSMUSG( | 0.901934 | 0.494203 | NO | NO | NA |
| Draxin | ENSMUSG( | ENSMUSG( | 0.999619 | 0.280151 | NO | NO | NA |
| Oit3 | ENSMUSG( | ENSMUSG( | 0.790583 | 0.064038 | NO | NO | NA |
| Gm28177 | ENSMUSG( | ENSMUSG( | -0.44925 | 0.460605 | NO | NO | NA |
| Slitrk3 | ENSMUSG( | ENSMUSG( | 1.46418 | 0.183598 | NO | NO | NA |
| Sv2b | ENSMUSG( | ENSMUSG( | 0.738214 | 0.430897 | NO | NO | NA |
| Gm42551 | ENSMUSG( | ENSMUSG( | 0.520118 | 0.585511 | NO | NO | NA |
| Lyve1 | ENSMUSG( | ENSMUSG( | 0.144545 | 0.90316 | NO | NO | NA |
| Gm15777 | ENSMUSG( | ENSMUSG( | 0.61693 | 0.270275 | NO | NO | NA |
| Vmn2r114 | ENSMUSG( | ENSMUSG( | -0.3009 | 0.769716 | NO | NO | NA |
| Slc10a4 | ENSMUSG( | ENSMUSG( | 0.696827 | 0.528583 | NO | NO | NA |
| Gm48272 | ENSMUSG( | ENSMUSG( | -0.00403 | 0.994565 | NO | NO | NA |
| 9930120I1 | ENSMUSG( | ENSMUSG( | -0.03168 | 0.929773 | NO | NO | NA |
| Nuak1 | ENSMUSG( | ENSMUSG( | 0.713928 | 0.357038 | NO | NO | NA |
| 4921517D | ENSMUSG( | ENSMUSG( | 1.212903 | 0.259779 | NO | NO | NA |

| | | | | | | | |
|---|---|---|---|---|---|---|---|
| Gm38069 | ENSMUSG( | ENSMUSG( | 0.606672 | 0.561653 | NO | NO | NA |
| Gm31458 | ENSMUSG( | ENSMUSG( | 1.594856 | 0.137394 | NO | NO | NA |
| Trbj1-4 | ENSMUSG( | ENSMUSG( | 0.547557 | 0.523035 | NO | NO | NA |
| Gm5742 | ENSMUSG( | ENSMUSG( | 0.847843 | 0.493363 | NO | NO | NA |
| Gm2814 | ENSMUSG( | ENSMUSG( | 0.599511 | 0.645714 | NO | NO | NA |
| Mpl | ENSMUSG( | ENSMUSG( | -0.16543 | 0.884768 | NO | NO | NA |
| Mcpt8 | ENSMUSG( | ENSMUSG( | -1.27962 | 0.54328 | NO | NO | NA |
| B930018H | ENSMUSG( | ENSMUSG( | -0.0262 | 0.95056 | NO | NO | NA |
| Olfr457 | ENSMUSG( | ENSMUSG( | 0.448853 | 0.676598 | NO | NO | NA |
| Zfp976 | ENSMUSG( | ENSMUSG( | 0.156954 | 0.805447 | NO | NO | NA |
| Gm37145 | ENSMUSG( | ENSMUSG( | -0.02526 | 0.958772 | NO | NO | NA |
| Gm47403 | ENSMUSG( | ENSMUSG( | 1.101739 | 0.32708 | NO | NO | NA |
| Gm30340 | ENSMUSG( | ENSMUSG( | 1.203939 | 0.271593 | NO | NO | NA |
| Vmn1r198 | ENSMUSG( | ENSMUSG( | 0.650671 | 0.551035 | NO | NO | NA |
| Gm38111 | ENSMUSG( | ENSMUSG( | -0.09683 | 0.92985 | NO | NO | NA |
| Muc4 | ENSMUSG( | ENSMUSG( | 0.23119 | 0.753468 | NO | NO | NA |
| Gm14408 | ENSMUSG( | ENSMUSG( | 0.677997 | 0.476831 | NO | NO | NA |
| Gm13032 | ENSMUSG( | ENSMUSG( | -0.2396 | 0.693448 | NO | NO | NA |
| Gm6169 | ENSMUSG( | ENSMUSG( | 0.162183 | 0.784601 | NO | NO | NA |
| Gm21860 | ENSMUSG( | ENSMUSG( | -1.395 | 0.078313 | NO | NO | NA |
| Gm49527 | ENSMUSG( | ENSMUSG( | 0.30336 | 0.861589 | NO | NO | NA |
| Gm20379 | ENSMUSG( | ENSMUSG( | 0.65952 | 0.401233 | NO | NO | NA |
| Anks6 | ENSMUSG( | ENSMUSG( | 0.5106 | 0.309469 | NO | NO | NA |
| 4930505N | ENSMUSG( | ENSMUSG( | -0.72609 | 0.283594 | NO | NO | NA |
| Gm12091 | ENSMUSG( | ENSMUSG( | 0.274359 | 0.764532 | NO | NO | NA |
| Cobl | ENSMUSG( | ENSMUSG( | 1.047525 | 0.211136 | NO | NO | NA |
| Rph3a | ENSMUSG( | ENSMUSG( | 1.129615 | 0.259206 | NO | NO | NA |
| Gm7143 | ENSMUSG( | ENSMUSG( | 0.613739 | 0.132657 | NO | NO | NA |
| Gm49660 | ENSMUSG( | ENSMUSG( | 0.878499 | 0.391869 | NO | NO | NA |
| Gm37931 | ENSMUSG( | ENSMUSG( | 0.776732 | 0.377352 | NO | NO | NA |
| Olfr846 | ENSMUSG( | ENSMUSG( | 0.565318 | 0.561491 | NO | NO | NA |
| Degs2 | ENSMUSG( | ENSMUSG( | 0.243714 | 0.688344 | NO | NO | NA |
| Mapk8ip2 | ENSMUSG( | ENSMUSG( | -0.04806 | 0.92627 | NO | NO | NA |
| A030012G | ENSMUSG( | ENSMUSG( | 0.281024 | 0.82922 | NO | NO | NA |
| Gm4691 | ENSMUSG( | ENSMUSG( | 0.198527 | 0.874602 | NO | NO | NA |
| Gm37633 | ENSMUSG( | ENSMUSG( | 0.701044 | 0.524654 | NO | NO | NA |
| Gm37061 | ENSMUSG( | ENSMUSG( | 0.62533 | 0.560002 | NO | NO | NA |
| Gm44167 | ENSMUSG( | ENSMUSG( | 0.820886 | 0.074674 | NO | NO | NA |
| Gm18909 | ENSMUSG( | ENSMUSG( | 1.289559 | 0.096007 | NO | NO | NA |
| ENSMUSG | ENSMUSG( | ENSMUSG( | 0.95952 | 0.065334 | NO | NO | NA |
| Tspan9 | ENSMUSG( | ENSMUSG( | 0.159976 | 0.852826 | NO | NO | NA |
| Gm4208 | ENSMUSG( | ENSMUSG( | -2.13945 | 8.07E-05 | YES | YES | Downregulated |
| Gm45493 | ENSMUSG( | ENSMUSG( | 0.344175 | 0.370629 | NO | NO | NA |
| Gm43031 | ENSMUSG( | ENSMUSG( | 0.001798 | 0.99716 | NO | NO | NA |
| 4931428L1 | ENSMUSG( | ENSMUSG( | 0.858647 | 0.397233 | NO | NO | NA |
| Gm5930 | ENSMUSG( | ENSMUSG( | 0.855914 | 0.442078 | NO | NO | NA |
| Gm43322 | ENSMUSG( | ENSMUSG( | 0.603251 | 0.578811 | NO | NO | NA |

| | | | | | | | |
|---|---|---|---|---|---|---|---|
| St8sia3 | ENSMUSG( | ENSMUSG( | 0.587125 | 0.581023 | NO | NO | NA |
| Gm12669 | ENSMUSG( | ENSMUSG( | 0.53077 | 0.356689 | NO | NO | NA |
| Pla1a | ENSMUSG( | ENSMUSG( | 0.901762 | 0.059775 | NO | NO | NA |
| Mrgprx2 | ENSMUSG( | ENSMUSG( | 1.04683 | 0.347976 | NO | NO | NA |
| Gm26571 | ENSMUSG( | ENSMUSG( | 0.029664 | 0.971145 | NO | NO | NA |
| Ston1 | ENSMUSG( | ENSMUSG( | 0.282101 | 0.74914 | NO | NO | NA |
| Gm32679 | ENSMUSG( | ENSMUSG( | -0.36934 | 0.456496 | NO | NO | NA |
| Ubap1l | ENSMUSG( | ENSMUSG( | 0.726657 | 0.209573 | NO | NO | NA |
| Gm17529 | ENSMUSG( | ENSMUSG( | 0.474207 | 0.454063 | NO | NO | NA |
| Ptprm | ENSMUSG( | ENSMUSG( | 0.644402 | 0.505653 | NO | NO | NA |
| Gm47518 | ENSMUSG( | ENSMUSG( | 1.008764 | 0.376757 | NO | NO | NA |
| Ybx2 | ENSMUSG( | ENSMUSG( | 0.528847 | 0.244982 | NO | NO | NA |
| Dennd2a | ENSMUSG( | ENSMUSG( | 0.407454 | 0.684836 | NO | NO | NA |
| Lrrn4cl | ENSMUSG( | ENSMUSG( | 0.558657 | 0.10277 | NO | NO | NA |
| Gm43584 | ENSMUSG( | ENSMUSG( | -0.26595 | 0.624353 | NO | NO | NA |
| Cxcr2 | ENSMUSG( | ENSMUSG( | 0.331375 | 0.72481 | NO | NO | NA |
| ENSMUSG | ENSMUSG( | ENSMUSG( | 1.467753 | 0.163953 | NO | NO | NA |
| Kif13a | ENSMUSG( | ENSMUSG( | 0.559476 | 0.43712 | NO | NO | NA |
| 9530053A( | ENSMUSG( | ENSMUSG( | 0.184729 | 0.858246 | NO | NO | NA |
| Vmn2r52 | ENSMUSG( | ENSMUSG( | 0.614327 | 0.540391 | NO | NO | NA |
| Igkv14-10( | ENSMUSG( | ENSMUSG( | -3.24145 | NA | NO | NO | NA |
| Gm20121 | ENSMUSG( | ENSMUSG( | 0.473411 | 0.716426 | NO | NO | NA |
| 1700029I1 | ENSMUSG( | ENSMUSG( | 0.330516 | 0.301508 | NO | NO | NA |
| Fcrls | ENSMUSG( | ENSMUSG( | 0.211405 | 0.660808 | NO | NO | NA |
| Gm45033 | ENSMUSG( | ENSMUSG( | 0.115376 | 0.890997 | NO | NO | NA |
| Sh3bp4 | ENSMUSG( | ENSMUSG( | 0.031297 | 0.954267 | NO | NO | NA |
| Slc24a2 | ENSMUSG( | ENSMUSG( | 0.062535 | 0.96163 | NO | NO | NA |
| 4921523L( | ENSMUSG( | ENSMUSG( | 0.976859 | 0.297659 | NO | NO | NA |
| Thbd | ENSMUSG( | ENSMUSG( | 1.061971 | 0.094834 | NO | NO | NA |
| Gm48225 | ENSMUSG( | ENSMUSG( | 0.503631 | 0.618093 | NO | NO | NA |
| Mip | ENSMUSG( | ENSMUSG( | 0.85444 | 0.056195 | NO | NO | NA |
| Gm43820 | ENSMUSG( | ENSMUSG( | 1.251095 | 0.193149 | NO | NO | NA |
| Gm30978 | ENSMUSG( | ENSMUSG( | 1.387498 | 0.244217 | NO | NO | NA |
| Gm10358 | ENSMUSG( | ENSMUSG( | -0.63893 | 0.577302 | NO | NO | NA |
| Arhgef17 | ENSMUSG( | ENSMUSG( | 0.447943 | 0.510813 | NO | NO | NA |
| Rps2-ps13 | ENSMUSG( | ENSMUSG( | 0.731154 | 0.231009 | NO | NO | NA |
| Grin3b | ENSMUSG( | ENSMUSG( | -0.08213 | 0.870889 | NO | NO | NA |
| Gm20481 | ENSMUSG( | ENSMUSG( | -2.60626 | 0.232343 | NO | NO | NA |
| 1700029E( | ENSMUSG( | ENSMUSG( | 0.881038 | 0.410913 | NO | NO | NA |
| Gm2415 | ENSMUSG( | ENSMUSG( | 0.31744 | 0.639774 | NO | NO | NA |
| Gm38227 | ENSMUSG( | ENSMUSG( | 0.46145 | 0.683941 | NO | NO | NA |
| Olfr1116 | ENSMUSG( | ENSMUSG( | -0.30813 | 0.81143 | NO | NO | NA |
| Gm49331 | ENSMUSG( | ENSMUSG( | -0.70708 | 0.043827 | YES | NO | NA |
| 4933406P( | ENSMUSG( | ENSMUSG( | 0.070078 | 0.868434 | NO | NO | NA |
| Gm38404 | ENSMUSG( | ENSMUSG( | 0.536775 | 0.658025 | NO | NO | NA |
| Gm13362 | ENSMUSG( | ENSMUSG( | 0.727245 | 0.594479 | NO | NO | NA |
| Hm629797 | ENSMUSG( | ENSMUSG( | 1.08607 | 0.274317 | NO | NO | NA |

| | | | | | | | |
|---|---|---|---|---|---|---|---|
| Gm42897 | ENSMUSG( | ENSMUSG( | 0.287824 | 0.786288 | NO | NO | NA |
| Kifc3 | ENSMUSG( | ENSMUSG( | 0.295233 | 0.604949 | NO | NO | NA |
| Gm37963 | ENSMUSG( | ENSMUSG( | -0.25829 | 0.495503 | NO | NO | NA |
| Sprr2a2 | ENSMUSG( | ENSMUSG( | 0.094926 | 0.950135 | NO | NO | NA |
| Wdr72 | ENSMUSG( | ENSMUSG( | 0.767086 | 0.433721 | NO | NO | NA |
| Gm45509 | ENSMUSG( | ENSMUSG( | -0.93195 | 0.028739 | YES | NO | NA |
| Ildr2 | ENSMUSG( | ENSMUSG( | 0.523526 | 0.604949 | NO | NO | NA |
| 4930478M | ENSMUSG( | ENSMUSG( | 0.974914 | 0.429674 | NO | NO | NA |
| Trgv7 | ENSMUSG( | ENSMUSG( | -1.09168 | 0.292632 | NO | NO | NA |
| 4930568A | ENSMUSG( | ENSMUSG( | -0.18015 | 0.645636 | NO | NO | NA |
| Epm2a | ENSMUSG( | ENSMUSG( | 0.677602 | 0.422361 | NO | NO | NA |
| ENSMUSG | ENSMUSG( | ENSMUSG( | -0.02055 | 0.960855 | NO | NO | NA |
| Rasgrp3 | ENSMUSG( | ENSMUSG( | 0.063745 | 0.943327 | NO | NO | NA |
| Gm5903 | ENSMUSG( | ENSMUSG( | 0.47743 | 0.257684 | NO | NO | NA |
| Nostrin | ENSMUSG( | ENSMUSG( | 0.922965 | 0.026368 | YES | NO | NA |
| Cylc1 | ENSMUSG( | ENSMUSG( | 0.411285 | 0.724236 | NO | NO | NA |
| Gm13199 | ENSMUSG( | ENSMUSG( | -0.08882 | 0.820316 | NO | NO | NA |
| Rpsa-ps7 | ENSMUSG( | ENSMUSG( | 0.753207 | 0.115904 | NO | NO | NA |
| Ly6g2 | ENSMUSG( | ENSMUSG( | 1.139681 | 0.308111 | NO | NO | NA |
| Gm6736 | ENSMUSG( | ENSMUSG( | -0.12958 | 0.830207 | NO | NO | NA |
| Gm6076 | ENSMUSG( | ENSMUSG( | 1.389737 | 0.159685 | NO | NO | NA |
| 1700112D | ENSMUSG( | ENSMUSG( | 0.886441 | 0.403874 | NO | NO | NA |
| Gm49507 | ENSMUSG( | ENSMUSG( | 0.715082 | 0.48942 | NO | NO | NA |
| Gm17753 | ENSMUSG( | ENSMUSG( | 0.032029 | 0.979835 | NO | NO | NA |
| Gm12411 | ENSMUSG( | ENSMUSG( | 1.267845 | 0.157988 | NO | NO | NA |
| Gm6209 | ENSMUSG( | ENSMUSG( | 0.773711 | 0.466491 | NO | NO | NA |
| Msrb3 | ENSMUSG( | ENSMUSG( | 0.426217 | 0.553609 | NO | NO | NA |
| Catsperg1 | ENSMUSG( | ENSMUSG( | 1.063267 | 0.006366 | YES | YES | Upregulated |
| Smtnl2 | ENSMUSG( | ENSMUSG( | 0.902855 | 0.354911 | NO | NO | NA |
| Heyl | ENSMUSG( | ENSMUSG( | 0.355969 | 0.658092 | NO | NO | NA |
| Gm45297 | ENSMUSG( | ENSMUSG( | 0.581985 | 0.601436 | NO | NO | NA |
| Wipf3 | ENSMUSG( | ENSMUSG( | -0.73697 | 0.274282 | NO | NO | NA |
| 2810414N | ENSMUSG( | ENSMUSG( | 0.299362 | 0.508113 | NO | NO | NA |
| ENSMUSG | ENSMUSG( | ENSMUSG( | 1.120843 | 0.330738 | NO | NO | NA |
| Hspb1 | ENSMUSG( | ENSMUSG( | -0.59413 | 0.58481 | NO | NO | NA |
| Olfr684 | ENSMUSG( | ENSMUSG( | 0.867586 | 0.431414 | NO | NO | NA |
| Klhl10 | ENSMUSG( | ENSMUSG( | -0.83575 | 0.03906 | YES | NO | NA |
| Chrdl1 | ENSMUSG( | ENSMUSG( | 0.987352 | 0.413484 | NO | NO | NA |
| Actn3 | ENSMUSG( | ENSMUSG( | 1.31613 | 0.000192 | YES | YES | Upregulated |
| Ddr2 | ENSMUSG( | ENSMUSG( | 0.421206 | 0.673641 | NO | NO | NA |
| Gm45760 | ENSMUSG( | ENSMUSG( | 0.45433 | 0.452744 | NO | NO | NA |
| Gm15473 | ENSMUSG( | ENSMUSG( | -0.22108 | 0.619706 | NO | NO | NA |
| Igkv8-21 | ENSMUSG( | ENSMUSG( | -2.16221 | 0.325896 | NO | NO | NA |
| Gm38319 | ENSMUSG( | ENSMUSG( | -0.65425 | 0.05728 | NO | NO | NA |
| Gm43585 | ENSMUSG( | ENSMUSG( | 0.665414 | 0.412416 | NO | NO | NA |
| Gm10743 | ENSMUSG( | ENSMUSG( | 1.035138 | 0.301461 | NO | NO | NA |
| Gm5532 | ENSMUSG( | ENSMUSG( | 0.634771 | 0.360007 | NO | NO | NA |

| | | | | | | | |
|---|---|---|---|---|---|---|---|
| Epha7 | ENSMUSG( | ENSMUSG( | 0.518477 | 0.624377 | NO | NO | NA |
| 1190005I0 | ENSMUSG( | ENSMUSG( | 0.642531 | 0.178926 | NO | NO | NA |
| Gm13031 | ENSMUSG( | ENSMUSG( | -0.38249 | 0.449199 | NO | NO | NA |
| Col6a3 | ENSMUSG( | ENSMUSG( | 0.431775 | 0.599902 | NO | NO | NA |
| Dnal1 | ENSMUSG( | ENSMUSG( | 0.566272 | 0.249717 | NO | NO | NA |
| Gm44087 | ENSMUSG( | ENSMUSG( | 0.42414 | 0.26751 | NO | NO | NA |
| Gm37930 | ENSMUSG( | ENSMUSG( | 0.737019 | 0.526351 | NO | NO | NA |
| Fblim1 | ENSMUSG( | ENSMUSG( | 0.263286 | 0.784321 | NO | NO | NA |
| Gm47976 | ENSMUSG( | ENSMUSG( | 0.390581 | 0.436468 | NO | NO | NA |
| Thbs3 | ENSMUSG( | ENSMUSG( | 0.012265 | 0.984129 | NO | NO | NA |
| Trim17 | ENSMUSG( | ENSMUSG( | -0.75813 | 0.159503 | NO | NO | NA |
| Klhl40 | ENSMUSG( | ENSMUSG( | 0.374896 | 0.707721 | NO | NO | NA |
| Gm5953 | ENSMUSG( | ENSMUSG( | 0.054895 | 0.94245 | NO | NO | NA |
| Gm29601 | ENSMUSG( | ENSMUSG( | 0.912195 | 0.494203 | NO | NO | NA |
| Gm20644 | ENSMUSG( | ENSMUSG( | 0.627721 | 0.579782 | NO | NO | NA |
| Fam186b | ENSMUSG( | ENSMUSG( | -0.86985 | 0.020803 | YES | NO | NA |
| Sh3tc1 | ENSMUSG( | ENSMUSG( | -0.86433 | 0.044069 | YES | NO | NA |
| Lrtm1 | ENSMUSG( | ENSMUSG( | 0.758721 | 0.546627 | NO | NO | NA |
| ENSMUSG | ENSMUSG( | ENSMUSG( | 0.841484 | 0.216297 | NO | NO | NA |
| Elf5 | ENSMUSG( | ENSMUSG( | 1.044342 | 0.205659 | NO | NO | NA |
| Gm47590 | ENSMUSG( | ENSMUSG( | 0.384986 | 0.730991 | NO | NO | NA |
| Gm43209 | ENSMUSG( | ENSMUSG( | 1.106554 | 0.401972 | NO | NO | NA |
| Smoc1 | ENSMUSG( | ENSMUSG( | 1.035952 | 0.191461 | NO | NO | NA |
| Olfr652 | ENSMUSG( | ENSMUSG( | 0.348394 | 0.756401 | NO | NO | NA |
| Olfr876 | ENSMUSG( | ENSMUSG( | 0.210361 | 0.796234 | NO | NO | NA |
| Gm9320 | ENSMUSG( | ENSMUSG( | -0.21591 | 0.648535 | NO | NO | NA |
| Gm17917 | ENSMUSG( | ENSMUSG( | 0.558281 | 0.608323 | NO | NO | NA |
| Pipox | ENSMUSG( | ENSMUSG( | 1.318913 | 0.005628 | YES | YES | Upregulated |
| Gm43952 | ENSMUSG( | ENSMUSG( | 0.378939 | 0.729762 | NO | NO | NA |
| Gm16433 | ENSMUSG( | ENSMUSG( | 0.644675 | 0.072402 | NO | NO | NA |
| Gm2163 | ENSMUSG( | ENSMUSG( | 1.751308 | 0.644476 | NO | NO | NA |
| Lrrc31 | ENSMUSG( | ENSMUSG( | 1.624053 | 0.189768 | NO | NO | NA |
| Gm47215 | ENSMUSG( | ENSMUSG( | 0.487993 | 0.56648 | NO | NO | NA |
| Gm42946 | ENSMUSG( | ENSMUSG( | 0.607215 | 0.543878 | NO | NO | NA |
| Vmn2r13 | ENSMUSG( | ENSMUSG( | 0.799146 | 0.553738 | NO | NO | NA |
| Gm47148 | ENSMUSG( | ENSMUSG( | 0.268582 | 0.590158 | NO | NO | NA |
| Cyp4x1 | ENSMUSG( | ENSMUSG( | 0.598892 | 0.599811 | NO | NO | NA |
| Fancd2os | ENSMUSG( | ENSMUSG( | 0.372418 | 0.514255 | NO | NO | NA |
| Gm10136 | ENSMUSG( | ENSMUSG( | 0.381031 | 0.464432 | NO | NO | NA |
| Trav13d-4 | ENSMUSG( | ENSMUSG( | 0.283234 | 0.856588 | NO | NO | NA |
| Rec8 | ENSMUSG( | ENSMUSG( | 1.058949 | 0.078893 | NO | NO | NA |
| Rab3c | ENSMUSG( | ENSMUSG( | 0.630479 | 0.5369 | NO | NO | NA |
| Myt1l | ENSMUSG( | ENSMUSG( | 0.9001 | 0.404157 | NO | NO | NA |
| Gm13381 | ENSMUSG( | ENSMUSG( | 0.633833 | 0.526045 | NO | NO | NA |
| Gm45378 | ENSMUSG( | ENSMUSG( | 0.493566 | 0.576079 | NO | NO | NA |
| Rdh5 | ENSMUSG( | ENSMUSG( | 0.457662 | 0.456867 | NO | NO | NA |
| Stxbp6 | ENSMUSG( | ENSMUSG( | 1.11315 | 0.182865 | NO | NO | NA |

| | | | | | | | |
|---|---|---|---|---|---|---|---|
| Zfp968-ps | ENSMUSG( | ENSMUSG( | -0.06217 | 0.964642 | NO | NO | NA |
| Ly96 | ENSMUSG( | ENSMUSG( | -0.3196 | 0.547577 | NO | NO | NA |
| Loxl3 | ENSMUSG( | ENSMUSG( | 0.161901 | 0.765489 | NO | NO | NA |
| Gm43887 | ENSMUSG( | ENSMUSG( | 0.869304 | 0.531106 | NO | NO | NA |
| Smarca5-p | ENSMUSG( | ENSMUSG( | -0.54273 | 0.474422 | NO | NO | NA |
| Pcdhb18 | ENSMUSG( | ENSMUSG( | 0.489337 | 0.672577 | NO | NO | NA |
| G630018N | ENSMUSG( | ENSMUSG( | 0.567371 | 0.600536 | NO | NO | NA |
| Ppfibp2 | ENSMUSG( | ENSMUSG( | 0.124295 | 0.881796 | NO | NO | NA |
| Gm13135 | ENSMUSG( | ENSMUSG( | 0.718042 | 0.197491 | NO | NO | NA |
| Gm8141 | ENSMUSG( | ENSMUSG( | 0.435063 | 0.648619 | NO | NO | NA |
| Gm10060 | ENSMUSG( | ENSMUSG( | -1.07513 | 0.206258 | NO | NO | NA |
| Olfr295 | ENSMUSG( | ENSMUSG( | 0.343719 | 0.743846 | NO | NO | NA |
| Sult4a1 | ENSMUSG( | ENSMUSG( | -0.48981 | 0.330466 | NO | NO | NA |
| Gm19277 | ENSMUSG( | ENSMUSG( | 1.325042 | 0.218524 | NO | NO | NA |
| Gm45188 | ENSMUSG( | ENSMUSG( | -0.51586 | 0.250124 | NO | NO | NA |
| Cd70 | ENSMUSG( | ENSMUSG( | 2.258639 | 0.000643 | YES | YES | Upregulated |
| Gm33332 | ENSMUSG( | ENSMUSG( | 1.61214 | 0.044993 | YES | YES | Upregulated |
| Gm47879 | ENSMUSG( | ENSMUSG( | 0.522172 | 0.331562 | NO | NO | NA |
| D030051J2 | ENSMUSG( | ENSMUSG( | 0.667699 | 0.317704 | NO | NO | NA |
| Vmn1r183 | ENSMUSG( | ENSMUSG( | 0.70856 | 0.45456 | NO | NO | NA |
| Gm5398 | ENSMUSG( | ENSMUSG( | 0.481578 | 0.656785 | NO | NO | NA |
| Wnt1 | ENSMUSG( | ENSMUSG( | 1.612755 | 0.081781 | NO | NO | NA |
| Ghr | ENSMUSG( | ENSMUSG( | -0.0282 | 0.978611 | NO | NO | NA |
| Gm15228 | ENSMUSG( | ENSMUSG( | 0.672974 | 0.489113 | NO | NO | NA |
| Gm49351 | ENSMUSG( | ENSMUSG( | 0.711897 | 0.572839 | NO | NO | NA |
| Gm37885 | ENSMUSG( | ENSMUSG( | -0.61557 | 0.618922 | NO | NO | NA |
| Gm18234 | ENSMUSG( | ENSMUSG( | 0.645068 | 0.553112 | NO | NO | NA |
| 4930434F2 | ENSMUSG( | ENSMUSG( | 0.791164 | 0.491999 | NO | NO | NA |
| Gm14377 | ENSMUSG( | ENSMUSG( | -0.43947 | 0.398656 | NO | NO | NA |
| Dytn | ENSMUSG( | ENSMUSG( | 1.002129 | 0.438599 | NO | NO | NA |
| Gm10614 | ENSMUSG( | ENSMUSG( | 0.300059 | 0.768418 | NO | NO | NA |
| Gm38062 | ENSMUSG( | ENSMUSG( | -0.32589 | 0.658915 | NO | NO | NA |
| Gm26920 | ENSMUSG( | ENSMUSG( | 1.826413 | 0.425749 | NO | NO | NA |
| Gm2308 | ENSMUSG( | ENSMUSG( | 0.525988 | 0.655006 | NO | NO | NA |
| Gm17978 | ENSMUSG( | ENSMUSG( | 0.675024 | 0.561653 | NO | NO | NA |
| Igsf9 | ENSMUSG( | ENSMUSG( | 0.372635 | 0.757235 | NO | NO | NA |
| Spdya | ENSMUSG( | ENSMUSG( | 0.302876 | 0.604896 | NO | NO | NA |
| Gm48307 | ENSMUSG( | ENSMUSG( | 0.878396 | 0.530994 | NO | NO | NA |
| Gm26728 | ENSMUSG( | ENSMUSG( | -0.15878 | 0.871269 | NO | NO | NA |
| Gm34106 | ENSMUSG( | ENSMUSG( | -0.9762 | 0.040233 | YES | NO | NA |
| Naaladl1 | ENSMUSG( | ENSMUSG( | 0.389583 | 0.463063 | NO | NO | NA |
| D630044L1 | ENSMUSG( | ENSMUSG( | 0.216655 | 0.648701 | NO | NO | NA |
| Gm38335 | ENSMUSG( | ENSMUSG( | 0.584086 | 0.667108 | NO | NO | NA |
| AW04773( | ENSMUSG( | ENSMUSG( | 0.674555 | 0.502364 | NO | NO | NA |
| D830030K | ENSMUSG( | ENSMUSG( | 0.868023 | 0.595769 | NO | NO | NA |
| Rpl9-ps4 | ENSMUSG( | ENSMUSG( | 0.685196 | 0.115387 | NO | NO | NA |
| Gm44950 | ENSMUSG( | ENSMUSG( | -0.32522 | 0.692623 | NO | NO | NA |

| | | | | | | | |
|---|---|---|---|---|---|---|---|
| E230013L2 | ENSMUSG0 | ENSMUSG0 | 0.326315 | 0.566439 | NO | NO | NA |
| Gm43768 | ENSMUSG0 | ENSMUSG0 | 0.22402 | 0.759406 | NO | NO | NA |
| Rhobtb1 | ENSMUSG0 | ENSMUSG0 | 0.085125 | 0.919733 | NO | NO | NA |
| Cct6b | ENSMUSG0 | ENSMUSG0 | 1.119557 | 0.044572 | YES | YES | Upregulated |
| Gm12416 | ENSMUSG0 | ENSMUSG0 | 0.636837 | 0.223305 | NO | NO | NA |
| Ighv1-53 | ENSMUSG0 | ENSMUSG0 | 0.162729 | 0.929293 | NO | NO | NA |
| Platr1 | ENSMUSG0 | ENSMUSG0 | 0.349389 | 0.698535 | NO | NO | NA |
| Cdh10 | ENSMUSG0 | ENSMUSG0 | 0.802867 | 0.422508 | NO | NO | NA |
| Gm38352 | ENSMUSG0 | ENSMUSG0 | 0.113563 | 0.897549 | NO | NO | NA |
| Gm45060 | ENSMUSG0 | ENSMUSG0 | 0.73618 | 0.489342 | NO | NO | NA |
| Gm14886 | ENSMUSG0 | ENSMUSG0 | 0.622962 | 0.572582 | NO | NO | NA |
| Trav13-5 | ENSMUSG0 | ENSMUSG0 | 2.346795 | 0.063704 | NO | NO | NA |
| Gm10250 | ENSMUSG0 | ENSMUSG0 | 0.065048 | 0.897357 | NO | NO | NA |
| 9130019P1 | ENSMUSG0 | ENSMUSG0 | -0.38447 | 0.419179 | NO | NO | NA |
| Bhmt | ENSMUSG0 | ENSMUSG0 | 0.56653 | 0.5889 | NO | NO | NA |
| Rnls | ENSMUSG0 | ENSMUSG0 | 0.854519 | 0.359465 | NO | NO | NA |
| 6430573P0 | ENSMUSG0 | ENSMUSG0 | 0.437247 | 0.553347 | NO | NO | NA |
| Igkv9-124 | ENSMUSG0 | ENSMUSG0 | -5.39663 | 0.034857 | YES | YES | Downregulated |
| Gm20098 | ENSMUSG0 | ENSMUSG0 | 0.299868 | 0.787833 | NO | NO | NA |
| A230052G | ENSMUSG0 | ENSMUSG0 | 0.375795 | 0.644229 | NO | NO | NA |
| Ttc9 | ENSMUSG0 | ENSMUSG0 | 0.611573 | 0.596936 | NO | NO | NA |
| 7SK ENSM | ENSMUSG0 | ENSMUSG0 | 0.338926 | 0.424383 | NO | NO | NA |
| Gm15185 | ENSMUSG0 | ENSMUSG0 | 0.98785 | 0.484233 | NO | NO | NA |
| Gm9333 | ENSMUSG0 | ENSMUSG0 | 0.486057 | 0.300594 | NO | NO | NA |
| Gm16098 | ENSMUSG0 | ENSMUSG0 | -0.58939 | 0.348486 | NO | NO | NA |
| Vmn2r85 | ENSMUSG0 | ENSMUSG0 | 1.012357 | 0.388508 | NO | NO | NA |
| Dact1 | ENSMUSG0 | ENSMUSG0 | 0.496382 | 0.255372 | NO | NO | NA |
| Gm37435 | ENSMUSG0 | ENSMUSG0 | 0.035496 | 0.976346 | NO | NO | NA |
| Fbl-ps2 | ENSMUSG0 | ENSMUSG0 | 0.300646 | 0.445223 | NO | NO | NA |
| Olfr769 | ENSMUSG0 | ENSMUSG0 | 0.734684 | 0.483972 | NO | NO | NA |
| Gm47669 | ENSMUSG0 | ENSMUSG0 | 0.568295 | 0.609483 | NO | NO | NA |
| A430034D | ENSMUSG0 | ENSMUSG0 | 0.754935 | 0.499353 | NO | NO | NA |
| Klrc3 | ENSMUSG0 | ENSMUSG0 | -0.49744 | 0.478132 | NO | NO | NA |
| Eda2r | ENSMUSG0 | ENSMUSG0 | 0.836598 | 0.433696 | NO | NO | NA |
| Igkv4-80 | ENSMUSG0 | ENSMUSG0 | -5.28447 | 0.013246 | YES | YES | Downregulated |
| Usp27x | ENSMUSG0 | ENSMUSG0 | 0.581309 | 0.188868 | NO | NO | NA |
| Gm29295 | ENSMUSG0 | ENSMUSG0 | 4.621404 | NA | NO | NO | NA |
| Gm49179 | ENSMUSG0 | ENSMUSG0 | 0.498588 | 0.392082 | NO | NO | NA |
| Gm47447 | ENSMUSG0 | ENSMUSG0 | -0.90859 | 0.005884 | YES | NO | NA |
| Gm8806 | ENSMUSG0 | ENSMUSG0 | 0.244112 | 0.576927 | NO | NO | NA |
| Cldn7 | ENSMUSG0 | ENSMUSG0 | 1.115333 | 0.264275 | NO | NO | NA |
| Pitx2 | ENSMUSG0 | ENSMUSG0 | 0.93265 | 0.08558 | NO | NO | NA |
| 4930455M | ENSMUSG0 | ENSMUSG0 | 0.657499 | 0.60556 | NO | NO | NA |
| 2900022M | ENSMUSG0 | ENSMUSG0 | 0.143135 | 0.892196 | NO | NO | NA |
| Gm49477 | ENSMUSG0 | ENSMUSG0 | 1.010016 | 0.396716 | NO | NO | NA |
| Trim34b | ENSMUSG0 | ENSMUSG0 | -0.82426 | 0.03541 | YES | NO | NA |
| Gm31748 | ENSMUSG0 | ENSMUSG0 | -1.13246 | 0.734711 | NO | NO | NA |

| | | | | | | | |
|---|---|---|---|---|---|---|---|
| Cxcl1 | ENSMUSG( | ENSMUSG( | 0.376395 | 0.731835 | NO | NO | NA |
| Liph | ENSMUSG( | ENSMUSG( | 0.76304 | 0.586177 | NO | NO | NA |
| ENSMUSG | ENSMUSG( | ENSMUSG( | 0.155029 | 0.776828 | NO | NO | NA |
| H2-M2 | ENSMUSG( | ENSMUSG( | 0.80014 | 0.566954 | NO | NO | NA |
| A430018G | ENSMUSG( | ENSMUSG( | 0.39305 | 0.482105 | NO | NO | NA |
| ENSMUSG | ENSMUSG( | ENSMUSG( | 0.629396 | 0.455151 | NO | NO | NA |
| Pde6a | ENSMUSG( | ENSMUSG( | 1.197535 | 0.345215 | NO | NO | NA |
| Azin2 | ENSMUSG( | ENSMUSG( | 0.328335 | 0.728011 | NO | NO | NA |
| Gm43519 | ENSMUSG( | ENSMUSG( | 0.645733 | 0.531106 | NO | NO | NA |
| Pld5 | ENSMUSG( | ENSMUSG( | 0.757388 | 0.527421 | NO | NO | NA |
| Gm10654 | ENSMUSG( | ENSMUSG( | 1.760047 | 0.043955 | YES | YES | Upregulated |
| Olfr1336 | ENSMUSG( | ENSMUSG( | 0.478566 | 0.639774 | NO | NO | NA |
| Rps2-ps8 | ENSMUSG( | ENSMUSG( | 0.887299 | 0.132698 | NO | NO | NA |
| Rtl5 | ENSMUSG( | ENSMUSG( | -1.8706 | 0.00045 | YES | YES | Downregulated |
| Gm20337 | ENSMUSG( | ENSMUSG( | 0.534432 | 0.312445 | NO | NO | NA |
| Flnc | ENSMUSG( | ENSMUSG( | 0.808253 | 0.436798 | NO | NO | NA |
| 1700062C | ENSMUSG( | ENSMUSG( | 0.635625 | 0.552976 | NO | NO | NA |
| Ifi207 | ENSMUSG( | ENSMUSG( | -0.92036 | 0.101015 | NO | NO | NA |
| Gm9208 | ENSMUSG( | ENSMUSG( | -0.23277 | 0.66374 | NO | NO | NA |
| Gm29803 | ENSMUSG( | ENSMUSG( | 1.264118 | 0.356725 | NO | NO | NA |
| 1700101I1 | ENSMUSG( | ENSMUSG( | -0.16422 | 0.741402 | NO | NO | NA |
| Olfr1234 | ENSMUSG( | ENSMUSG( | 1.829615 | 0.05629 | NO | NO | NA |
| Gm26766 | ENSMUSG( | ENSMUSG( | 0.512826 | 0.535389 | NO | NO | NA |
| Adamts17 | ENSMUSG( | ENSMUSG( | 0.847593 | 0.526339 | NO | NO | NA |
| Gm13323 | ENSMUSG( | ENSMUSG( | 1.291768 | 0.247168 | NO | NO | NA |
| Prss56 | ENSMUSG( | ENSMUSG( | 1.071129 | 0.069426 | NO | NO | NA |
| Gm2810 | ENSMUSG( | ENSMUSG( | 0.151471 | 0.841956 | NO | NO | NA |
| Gm48882 | ENSMUSG( | ENSMUSG( | 0.374244 | 0.556502 | NO | NO | NA |
| Gabrq | ENSMUSG( | ENSMUSG( | -0.1878 | 0.868434 | NO | NO | NA |
| 4930471E | ENSMUSG( | ENSMUSG( | 0.922926 | 0.402567 | NO | NO | NA |
| Vmn1r71 | ENSMUSG( | ENSMUSG( | 0.289802 | 0.786202 | NO | NO | NA |
| Pgam1-ps | ENSMUSG( | ENSMUSG( | 0.419834 | 0.432225 | NO | NO | NA |
| Gm42735 | ENSMUSG( | ENSMUSG( | 0.278825 | 0.540932 | NO | NO | NA |
| Gm29434 | ENSMUSG( | ENSMUSG( | 0.106667 | 0.835562 | NO | NO | NA |
| Gm13615 | ENSMUSG( | ENSMUSG( | 0.732227 | 0.466606 | NO | NO | NA |
| Gm43513 | ENSMUSG( | ENSMUSG( | -0.0458 | 0.900999 | NO | NO | NA |
| Tmem204 | ENSMUSG( | ENSMUSG( | -0.06338 | 0.888811 | NO | NO | NA |
| Gas6 | ENSMUSG( | ENSMUSG( | 0.342426 | 0.532314 | NO | NO | NA |
| Gm37500 | ENSMUSG( | ENSMUSG( | -0.94837 | 0.048231 | YES | NO | NA |
| Slc5a1 | ENSMUSG( | ENSMUSG( | 1.01238 | 0.38137 | NO | NO | NA |
| Cplane2 | ENSMUSG( | ENSMUSG( | 0.046868 | 0.916181 | NO | NO | NA |
| Ptprr | ENSMUSG( | ENSMUSG( | 0.674025 | 0.318048 | NO | NO | NA |
| Gm42866 | ENSMUSG( | ENSMUSG( | 0.62 | 0.492913 | NO | NO | NA |
| Gm9964 | ENSMUSG( | ENSMUSG( | 0.965227 | 0.324184 | NO | NO | NA |
| Gm42725 | ENSMUSG( | ENSMUSG( | 0.40333 | 0.669022 | NO | NO | NA |
| ENSMUSG( | ENSMUSG( | ENSMUSG( | -0.73024 | 0.476823 | NO | NO | NA |
| Gm29394 | ENSMUSG( | ENSMUSG( | 0.454962 | 0.512662 | NO | NO | NA |

| | | | | | | | |
|---|---|---|---|---|---|---|---|
| Gm8873 | ENSMUSG( | ENSMUSG( | 0.907436 | 0.027231 | YES | NO | NA |
| ENSMUSG | ENSMUSG( | ENSMUSG( | 0.491176 | 0.417268 | NO | NO | NA |
| Gm13611 | ENSMUSG( | ENSMUSG( | 0.901187 | 0.04038 | YES | NO | NA |
| Cass4 | ENSMUSG( | ENSMUSG( | 0.846483 | 0.207883 | NO | NO | NA |
| Olfr845 | ENSMUSG( | ENSMUSG( | 0.712409 | 0.524123 | NO | NO | NA |
| Gm13185 | ENSMUSG( | ENSMUSG( | -0.29898 | 0.574837 | NO | NO | NA |
| 4930533I2 | ENSMUSG( | ENSMUSG( | 0.828933 | 0.429159 | NO | NO | NA |
| Ccdc74a | ENSMUSG( | ENSMUSG( | -0.46973 | 0.313838 | NO | NO | NA |
| Gsdmc4 | ENSMUSG( | ENSMUSG( | 1.233387 | 0.034718 | YES | YES | Upregulated |
| Gm11739 | ENSMUSG( | ENSMUSG( | 0.089184 | 0.860992 | NO | NO | NA |
| Rnd3 | ENSMUSG( | ENSMUSG( | 0.686222 | 0.439807 | NO | NO | NA |
| Gm49727 | ENSMUSG( | ENSMUSG( | 1.243247 | 0.154052 | NO | NO | NA |
| Gm47826 | ENSMUSG( | ENSMUSG( | -0.15134 | 0.796122 | NO | NO | NA |
| Gm8458 | ENSMUSG( | ENSMUSG( | 0.770865 | 0.498086 | NO | NO | NA |
| ENSMUSG | ENSMUSG( | ENSMUSG( | 0.26074 | 0.470363 | NO | NO | NA |
| Frmpd3 | ENSMUSG( | ENSMUSG( | 0.147564 | 0.890997 | NO | NO | NA |
| Psd3 | ENSMUSG( | ENSMUSG( | -0.4357 | 0.489733 | NO | NO | NA |
| 4930578E1 | ENSMUSG( | ENSMUSG( | 1.308668 | 0.18881 | NO | NO | NA |
| 5930409G | ENSMUSG( | ENSMUSG( | 0.450252 | 0.656971 | NO | NO | NA |
| Gm37357 | ENSMUSG( | ENSMUSG( | -0.10927 | 0.85712 | NO | NO | NA |
| ENSMUSG | ENSMUSG( | ENSMUSG( | -0.23017 | 0.483451 | NO | NO | NA |
| Gm45650 | ENSMUSG( | ENSMUSG( | 0.613912 | 0.557842 | NO | NO | NA |
| Gm10558 | ENSMUSG( | ENSMUSG( | 0.920263 | 0.32874 | NO | NO | NA |
| Gm32122 | ENSMUSG( | ENSMUSG( | 0.969296 | 0.188406 | NO | NO | NA |
| Gm4900 | ENSMUSG( | ENSMUSG( | 0.563273 | 0.145818 | NO | NO | NA |
| Gm37342 | ENSMUSG( | ENSMUSG( | 0.802036 | 0.450589 | NO | NO | NA |
| 4930519A | ENSMUSG( | ENSMUSG( | 0.118246 | 0.930577 | NO | NO | NA |
| Gm44587 | ENSMUSG( | ENSMUSG( | 1.479027 | 0.081048 | NO | NO | NA |
| Gm44799 | ENSMUSG( | ENSMUSG( | 0.322676 | 0.802784 | NO | NO | NA |
| Slc5a2 | ENSMUSG( | ENSMUSG( | 1.00654 | 0.432526 | NO | NO | NA |
| Gm9767 | ENSMUSG( | ENSMUSG( | 0.471166 | 0.656466 | NO | NO | NA |
| Olfr982 | ENSMUSG( | ENSMUSG( | 0.527331 | 0.63678 | NO | NO | NA |
| Gm29430 | ENSMUSG( | ENSMUSG( | 0.742966 | 0.519417 | NO | NO | NA |
| Gm45449 | ENSMUSG( | ENSMUSG( | 0.966362 | 0.391427 | NO | NO | NA |
| Rpl21-ps6 | ENSMUSG( | ENSMUSG( | 0.848243 | 0.349726 | NO | NO | NA |
| Gm38947 | ENSMUSG( | ENSMUSG( | -0.78829 | 0.047925 | YES | NO | NA |
| Gm49700 | ENSMUSG( | ENSMUSG( | 0.496456 | 0.361782 | NO | NO | NA |
| Gm37005 | ENSMUSG( | ENSMUSG( | 0.412919 | 0.754479 | NO | NO | NA |
| Syt14 | ENSMUSG( | ENSMUSG( | 0.882449 | 0.270362 | NO | NO | NA |
| Scrt1 | ENSMUSG( | ENSMUSG( | -0.73782 | 0.172523 | NO | NO | NA |
| Gm16579 | ENSMUSG( | ENSMUSG( | 0.32834 | 0.765791 | NO | NO | NA |
| Gm31356 | ENSMUSG( | ENSMUSG( | 0.357641 | 0.669318 | NO | NO | NA |
| F730035M | ENSMUSG( | ENSMUSG( | 1.038284 | 0.229957 | NO | NO | NA |
| Il33 | ENSMUSG( | ENSMUSG( | 1.736492 | 0.000766 | YES | YES | Upregulated |
| Amy1 | ENSMUSG( | ENSMUSG( | 0.492005 | 0.609944 | NO | NO | NA |
| Trbj1-1 | ENSMUSG( | ENSMUSG( | -0.48626 | 0.484229 | NO | NO | NA |
| Gm6808 | ENSMUSG( | ENSMUSG( | 0.280971 | 0.808364 | NO | NO | NA |

| | | | | | | | |
|---|---|---|---|---|---|---|---|
| Olfr54 | ENSMUSG( | ENSMUSG( | 0.418844 | 0.729783 | NO | NO | NA |
| Dbn1 | ENSMUSG( | ENSMUSG( | 0.373994 | 0.450672 | NO | NO | NA |
| Igkv4-59 | ENSMUSG( | ENSMUSG( | -7.0598 | 2.22E-06 | YES | YES | Downregulated |
| ENSMUSG( | ENSMUSG( | ENSMUSG( | 0.089935 | 0.810621 | NO | NO | NA |
| Mgat4c | ENSMUSG( | ENSMUSG( | 0.170603 | 0.888832 | NO | NO | NA |
| Inhca | ENSMUSG( | ENSMUSG( | 0.530151 | 0.452026 | NO | NO | NA |
| Agbl1 | ENSMUSG( | ENSMUSG( | 1.392296 | 0.282261 | NO | NO | NA |
| Tmem233 | ENSMUSG( | ENSMUSG( | 0.921332 | 0.461948 | NO | NO | NA |
| Gm28289 | ENSMUSG( | ENSMUSG( | -0.3032 | 0.820976 | NO | NO | NA |
| Abat | ENSMUSG( | ENSMUSG( | 0.785231 | 0.18343 | NO | NO | NA |
| Gm44068 | ENSMUSG( | ENSMUSG( | 1.186929 | 0.347218 | NO | NO | NA |
| Prss57 | ENSMUSG( | ENSMUSG( | 0.559435 | 0.597856 | NO | NO | NA |
| 4932412D | ENSMUSG( | ENSMUSG( | 0.816755 | 0.476489 | NO | NO | NA |
| Gm48542 | ENSMUSG( | ENSMUSG( | 1.074486 | 0.364181 | NO | NO | NA |
| Gm34743 | ENSMUSG( | ENSMUSG( | 0.584355 | 0.586288 | NO | NO | NA |
| Gm5234 | ENSMUSG( | ENSMUSG( | 0.180339 | 0.86879 | NO | NO | NA |
| Nsa2-ps2 | ENSMUSG( | ENSMUSG( | 0.362864 | 0.766332 | NO | NO | NA |
| Gm47884 | ENSMUSG( | ENSMUSG( | 1.457406 | 0.15771 | NO | NO | NA |
| Rps25-ps1 | ENSMUSG( | ENSMUSG( | 0.406581 | 0.476823 | NO | NO | NA |
| Gm43906 | ENSMUSG( | ENSMUSG( | 0.716773 | 0.495773 | NO | NO | NA |
| Gm48512 | ENSMUSG( | ENSMUSG( | -0.30425 | 0.562892 | NO | NO | NA |
| Gm50105 | ENSMUSG( | ENSMUSG( | -0.1934 | 0.727964 | NO | NO | NA |
| Rac3 | ENSMUSG( | ENSMUSG( | -0.08963 | 0.837721 | NO | NO | NA |
| Plekhg1 | ENSMUSG( | ENSMUSG( | 0.553776 | 0.335005 | NO | NO | NA |
| Ushbp1 | ENSMUSG( | ENSMUSG( | -0.18069 | 0.831172 | NO | NO | NA |
| 8430423G | ENSMUSG( | ENSMUSG( | 1.044206 | 0.304919 | NO | NO | NA |
| Gm17130 | ENSMUSG( | ENSMUSG( | 0.832701 | 0.427511 | NO | NO | NA |
| Gm26974 | ENSMUSG( | ENSMUSG( | 0.34266 | 0.756594 | NO | NO | NA |
| Myf5 | ENSMUSG( | ENSMUSG( | 0.28341 | 0.816371 | NO | NO | NA |
| ENSMUSG | ENSMUSG( | ENSMUSG( | 0.309217 | 0.581594 | NO | NO | NA |
| E330011M | ENSMUSG( | ENSMUSG( | -0.72975 | 0.259199 | NO | NO | NA |
| Gm32061 | ENSMUSG( | ENSMUSG( | 0.150459 | 0.894579 | NO | NO | NA |
| 4933431K: | ENSMUSG( | ENSMUSG( | 0.175761 | 0.880052 | NO | NO | NA |
| Baiap2l2 | ENSMUSG( | ENSMUSG( | 1.153519 | 0.020162 | YES | YES | Upregulated |
| A730006G | ENSMUSG( | ENSMUSG( | 0.643147 | 0.570158 | NO | NO | NA |
| Gm45698 | ENSMUSG( | ENSMUSG( | -0.06375 | 0.919297 | NO | NO | NA |
| Slitrk2 | ENSMUSG( | ENSMUSG( | 0.296023 | 0.786484 | NO | NO | NA |
| 5031415H | ENSMUSG( | ENSMUSG( | 0.32156 | 0.792398 | NO | NO | NA |
| Trav15-1-c | ENSMUSG( | ENSMUSG( | -0.54992 | 0.596711 | NO | NO | NA |
| Gcsam | ENSMUSG( | ENSMUSG( | -1.09745 | 0.196909 | NO | NO | NA |
| ENSMUSG | ENSMUSG( | ENSMUSG( | 0.435151 | 0.681191 | NO | NO | NA |
| Gm14167 | ENSMUSG( | ENSMUSG( | 0.314726 | 0.531711 | NO | NO | NA |
| Dynlt2a3 | ENSMUSG( | ENSMUSG( | 0.964737 | 0.404243 | NO | NO | NA |
| B230323A | ENSMUSG( | ENSMUSG( | 0.68911 | 0.548091 | NO | NO | NA |
| Gm13523 | ENSMUSG( | ENSMUSG( | 1.152629 | 0.047144 | YES | YES | Upregulated |
| ENSMUSG | ENSMUSG( | ENSMUSG( | 0.498818 | 0.621132 | NO | NO | NA |
| Bcar3 | ENSMUSG( | ENSMUSG( | 0.569575 | 0.562282 | NO | NO | NA |

| | | | | | | | |
|---|---|---|---|---|---|---|---|
| Gm17976 | ENSMUSG( | ENSMUSG( | 0.357983 | 0.791755 | NO | NO | NA |
| 4930579C: | ENSMUSG( | ENSMUSG( | 0.53174 | 0.287642 | NO | NO | NA |
| Cbln4 | ENSMUSG( | ENSMUSG( | 1.987587 | 0.087982 | NO | NO | NA |
| Gm13156 | ENSMUSG( | ENSMUSG( | 0.606671 | 0.54862 | NO | NO | NA |
| Gm34923 | ENSMUSG( | ENSMUSG( | 1.00944 | 0.418305 | NO | NO | NA |
| Bmp7 | ENSMUSG( | ENSMUSG( | 1.288825 | 0.003203 | YES | YES | Upregulated |
| Rpl9-ps7 | ENSMUSG( | ENSMUSG( | 0.239964 | 0.551631 | NO | NO | NA |
| Olfr145 | ENSMUSG( | ENSMUSG( | 1.138879 | 0.244364 | NO | NO | NA |
| Ccbe1 | ENSMUSG( | ENSMUSG( | 0.479625 | 0.614865 | NO | NO | NA |
| Gm17324 | ENSMUSG( | ENSMUSG( | 0.662376 | 0.132865 | NO | NO | NA |
| Gm37320 | ENSMUSG( | ENSMUSG( | 0.186179 | 0.82838 | NO | NO | NA |
| Tbc1d8 | ENSMUSG( | ENSMUSG( | 0.455387 | 0.43086 | NO | NO | NA |
| Pde8b | ENSMUSG( | ENSMUSG( | -0.0347 | 0.967852 | NO | NO | NA |
| Kcnip3 | ENSMUSG( | ENSMUSG( | 0.689214 | 0.123792 | NO | NO | NA |
| Gm29927 | ENSMUSG( | ENSMUSG( | 0.354507 | 0.764889 | NO | NO | NA |
| Ncmap | ENSMUSG( | ENSMUSG( | 0.517964 | 0.391944 | NO | NO | NA |
| 4632404M | ENSMUSG( | ENSMUSG( | 0.5431 | 0.689161 | NO | NO | NA |
| Gm37663 | ENSMUSG( | ENSMUSG( | 1.00684 | 0.311936 | NO | NO | NA |
| E130215H | ENSMUSG( | ENSMUSG( | -0.13314 | 0.735932 | NO | NO | NA |
| Gstp2 | ENSMUSG( | ENSMUSG( | -0.0333 | 0.953889 | NO | NO | NA |
| A2ml1 | ENSMUSG( | ENSMUSG( | 0.582657 | 0.582357 | NO | NO | NA |
| H2-K2 | ENSMUSG( | ENSMUSG( | -0.09591 | 0.856749 | NO | NO | NA |
| Prpf40b | ENSMUSG( | ENSMUSG( | -0.05862 | 0.897349 | NO | NO | NA |
| Olfr646 | ENSMUSG( | ENSMUSG( | 0.686712 | 0.482306 | NO | NO | NA |
| Rpl30-ps9 | ENSMUSG( | ENSMUSG( | 0.836098 | 0.843134 | NO | NO | NA |
| Stbd1 | ENSMUSG( | ENSMUSG( | -0.11057 | 0.860871 | NO | NO | NA |
| Fgfbp1 | ENSMUSG( | ENSMUSG( | 0.531356 | 0.643304 | NO | NO | NA |
| Prox2os | ENSMUSG( | ENSMUSG( | -0.0546 | 0.907201 | NO | NO | NA |
| Bves | ENSMUSG( | ENSMUSG( | 0.76401 | 0.545216 | NO | NO | NA |
| Fezf2 | ENSMUSG( | ENSMUSG( | 1.199236 | 0.202909 | NO | NO | NA |
| Edar | ENSMUSG( | ENSMUSG( | 0.358404 | 0.75729 | NO | NO | NA |
| Blnk | ENSMUSG( | ENSMUSG( | -0.59386 | 0.496053 | NO | NO | NA |
| Cryba4 | ENSMUSG( | ENSMUSG( | -0.35447 | 0.35462 | NO | NO | NA |
| Igkv3-9 | ENSMUSG( | ENSMUSG( | -5.51243 | NA | NO | NO | NA |
| Lrrn3 | ENSMUSG( | ENSMUSG( | -0.04354 | 0.929293 | NO | NO | NA |
| Aif1l | ENSMUSG( | ENSMUSG( | 0.663966 | 0.331932 | NO | NO | NA |
| Kctd16 | ENSMUSG( | ENSMUSG( | 0.934698 | 0.472888 | NO | NO | NA |
| Olfr1185-p | ENSMUSG( | ENSMUSG( | 0.663654 | 0.612152 | NO | NO | NA |
| Gm10550 | ENSMUSG( | ENSMUSG( | 1.646083 | 0.213403 | NO | NO | NA |
| Pmis2 | ENSMUSG( | ENSMUSG( | 1.061169 | 0.298524 | NO | NO | NA |
| Gm36634 | ENSMUSG( | ENSMUSG( | 0.243837 | 0.810036 | NO | NO | NA |
| Gm10340 | ENSMUSG( | ENSMUSG( | -0.40749 | 0.521432 | NO | NO | NA |
| Gm18996 | ENSMUSG( | ENSMUSG( | 0.638779 | 0.561209 | NO | NO | NA |
| 4933431C: | ENSMUSG( | ENSMUSG( | -0.3895 | 0.560317 | NO | NO | NA |
| Nron | ENSMUSG( | ENSMUSG( | 0.476696 | 0.516168 | NO | NO | NA |
| Cdh13 | ENSMUSG( | ENSMUSG( | 0.407424 | 0.694528 | NO | NO | NA |
| ENSMUSG | ENSMUSG( | ENSMUSG( | -0.12807 | 0.785602 | NO | NO | NA |

| | | | | | | | |
|---|---|---|---|---|---|---|---|
| Gm48082 | ENSMUSG( | ENSMUSG( | -0.29111 | 0.678687 | NO | NO | NA |
| Cnn3 | ENSMUSG( | ENSMUSG( | 0.469908 | 0.445328 | NO | NO | NA |
| Gm10284 | ENSMUSG( | ENSMUSG( | -0.26283 | 0.708667 | NO | NO | NA |
| Olfr1265 | ENSMUSG( | ENSMUSG( | 0.908587 | 0.398115 | NO | NO | NA |
| Gm44947 | ENSMUSG( | ENSMUSG( | 0.822025 | 0.49528 | NO | NO | NA |
| Ccr10 | ENSMUSG( | ENSMUSG( | -0.68492 | 0.127189 | NO | NO | NA |
| Gm45769 | ENSMUSG( | ENSMUSG( | 0.211138 | 0.614864 | NO | NO | NA |
| Gm6491 | ENSMUSG( | ENSMUSG( | 1.692795 | 0.158327 | NO | NO | NA |
| Etaa1os | ENSMUSG( | ENSMUSG( | -0.53357 | 0.182146 | NO | NO | NA |
| Gm48600 | ENSMUSG( | ENSMUSG( | 0.260873 | 0.805041 | NO | NO | NA |
| Adamtsl4 | ENSMUSG( | ENSMUSG( | -1.01986 | 0.207264 | NO | NO | NA |
| Olfr460 | ENSMUSG( | ENSMUSG( | 0.27322 | 0.801288 | NO | NO | NA |
| Stra6l | ENSMUSG( | ENSMUSG( | -0.11277 | 0.906496 | NO | NO | NA |
| ENSMUSG | ENSMUSG( | ENSMUSG( | -0.37428 | 0.471033 | NO | NO | NA |
| Gabrg1 | ENSMUSG( | ENSMUSG( | 1.000242 | 0.42466 | NO | NO | NA |
| Gm45137 | ENSMUSG( | ENSMUSG( | 0.160817 | 0.666332 | NO | NO | NA |
| Gm49741 | ENSMUSG( | ENSMUSG( | 0.320849 | 0.568664 | NO | NO | NA |
| Gm38137 | ENSMUSG( | ENSMUSG( | 1.126649 | 0.362672 | NO | NO | NA |
| Tdg-ps2 | ENSMUSG( | ENSMUSG( | 0.325123 | 0.491395 | NO | NO | NA |
| Gm37080 | ENSMUSG( | ENSMUSG( | 0.454434 | 0.672577 | NO | NO | NA |
| Gm42794 | ENSMUSG( | ENSMUSG( | 1.008518 | 0.359328 | NO | NO | NA |
| Mlxipl | ENSMUSG( | ENSMUSG( | 1.040119 | 0.080464 | NO | NO | NA |
| ENSMUSG( | ENSMUSG( | ENSMUSG( | -0.24159 | 0.697385 | NO | NO | NA |
| B3galt1 | ENSMUSG( | ENSMUSG( | 1.188226 | 0.095947 | NO | NO | NA |
| Gm12188 | ENSMUSG( | ENSMUSG( | -1.14224 | 0.015641 | YES | YES | Downregulated |
| Gm6682 | ENSMUSG( | ENSMUSG( | 0.733875 | 0.26864 | NO | NO | NA |
| Gm28262 | ENSMUSG( | ENSMUSG( | 0.610038 | 0.593442 | NO | NO | NA |
| Gm17039 | ENSMUSG( | ENSMUSG( | -1.02492 | 0.084276 | NO | NO | NA |
| 9430060I0 | ENSMUSG( | ENSMUSG( | -0.60547 | 0.253203 | NO | NO | NA |
| Gm38351 | ENSMUSG( | ENSMUSG( | 1.367405 | 0.328293 | NO | NO | NA |
| Gm10293 | ENSMUSG( | ENSMUSG( | 0.593262 | 0.33909 | NO | NO | NA |
| Amy2a4 | ENSMUSG( | ENSMUSG( | 1.755841 | 0.488062 | NO | NO | NA |
| Gm43912 | ENSMUSG( | ENSMUSG( | -0.10592 | 0.785998 | NO | NO | NA |
| Olfr1191-p | ENSMUSG( | ENSMUSG( | 0.230977 | 0.831994 | NO | NO | NA |
| Ankub1 | ENSMUSG( | ENSMUSG( | 0.678326 | 0.500026 | NO | NO | NA |
| Slc28a2b | ENSMUSG( | ENSMUSG( | -0.16967 | 0.818367 | NO | NO | NA |
| Trhde | ENSMUSG( | ENSMUSG( | 0.289179 | 0.800263 | NO | NO | NA |
| 2010007H | ENSMUSG( | ENSMUSG( | 0.541436 | 0.600536 | NO | NO | NA |
| Olfr1238 | ENSMUSG( | ENSMUSG( | 0.677772 | 0.541473 | NO | NO | NA |
| Gm16172 | ENSMUSG( | ENSMUSG( | -0.40264 | 0.545229 | NO | NO | NA |
| Gm14011 | ENSMUSG( | ENSMUSG( | 1.398114 | 0.216816 | NO | NO | NA |
| Gm12263 | ENSMUSG( | ENSMUSG( | 0.096516 | 0.836456 | NO | NO | NA |
| Sost | ENSMUSG( | ENSMUSG( | 0.586129 | 0.56965 | NO | NO | NA |
| Mcpt9 | ENSMUSG( | ENSMUSG( | 0.736371 | 0.528563 | NO | NO | NA |
| ENSMUSG | ENSMUSG( | ENSMUSG( | -0.51672 | 0.354818 | NO | NO | NA |
| Gm10709 | ENSMUSG( | ENSMUSG( | 1.019153 | 0.343765 | NO | NO | NA |
| Ctnna3 | ENSMUSG( | ENSMUSG( | 0.593146 | 0.561491 | NO | NO | NA |

| | | | | | | | |
|---|---|---|---|---|---|---|---|
| Lrrc70 | ENSMUSG( | ENSMUSG( | 0.165854 | 0.703822 | NO | NO | NA |
| Efcab1 | ENSMUSG( | ENSMUSG( | 0.517129 | 0.564209 | NO | NO | NA |
| Gm7599 | ENSMUSG( | ENSMUSG( | 0.593164 | 0.535508 | NO | NO | NA |
| Uchl4 | ENSMUSG( | ENSMUSG( | 0.371867 | 0.627826 | NO | NO | NA |
| Hsf3 | ENSMUSG( | ENSMUSG( | 0.841052 | 0.430364 | NO | NO | NA |
| Gm11263 | ENSMUSG( | ENSMUSG( | 0.734695 | 0.040573 | YES | NO | NA |
| Rasgef1c | ENSMUSG( | ENSMUSG( | 0.131658 | 0.797391 | NO | NO | NA |
| Prkaa2 | ENSMUSG( | ENSMUSG( | 0.64539 | 0.523478 | NO | NO | NA |
| Gm14021 | ENSMUSG( | ENSMUSG( | 1.037575 | 0.315743 | NO | NO | NA |
| Sema3e | ENSMUSG( | ENSMUSG( | 0.607164 | 0.523478 | NO | NO | NA |
| Trbj2-3 | ENSMUSG( | ENSMUSG( | -0.51712 | 0.323055 | NO | NO | NA |
| Gm17201 | ENSMUSG( | ENSMUSG( | 0.125665 | 0.82465 | NO | NO | NA |
| Mertk | ENSMUSG( | ENSMUSG( | 0.173226 | 0.826879 | NO | NO | NA |
| Rapgef3 | ENSMUSG( | ENSMUSG( | 0.33479 | 0.69132 | NO | NO | NA |
| Ephb1 | ENSMUSG( | ENSMUSG( | 0.704006 | 0.548854 | NO | NO | NA |
| Gm11353 | ENSMUSG( | ENSMUSG( | 0.610567 | 0.278187 | NO | NO | NA |
| Rpl7-ps7 | ENSMUSG( | ENSMUSG( | 0.50452 | 0.272404 | NO | NO | NA |
| Gm43081 | ENSMUSG( | ENSMUSG( | 0.655606 | 0.39729 | NO | NO | NA |
| Fbxo47 | ENSMUSG( | ENSMUSG( | 0.555687 | 0.177065 | NO | NO | NA |
| Brsk2 | ENSMUSG( | ENSMUSG( | 0.465274 | 0.561209 | NO | NO | NA |
| C330002G | ENSMUSG( | ENSMUSG( | 0.440804 | 0.682979 | NO | NO | NA |
| Ccr4 | ENSMUSG( | ENSMUSG( | 0.635686 | 0.493702 | NO | NO | NA |
| 4633401B( | ENSMUSG( | ENSMUSG( | 0.419116 | 0.453169 | NO | NO | NA |
| Adgre1 | ENSMUSG( | ENSMUSG( | -1.22915 | 0.037456 | YES | YES | Downregulated |
| 2900064K( | ENSMUSG( | ENSMUSG( | 0.840353 | 0.439278 | NO | NO | NA |
| Plek2 | ENSMUSG( | ENSMUSG( | 1.219844 | 0.038837 | YES | YES | Upregulated |
| Gm45084 | ENSMUSG( | ENSMUSG( | -0.12288 | 0.847329 | NO | NO | NA |
| A930002C( | ENSMUSG( | ENSMUSG( | 0.412789 | 0.706841 | NO | NO | NA |
| ENSMUSG( | ENSMUSG( | ENSMUSG( | 1.041058 | 0.347354 | NO | NO | NA |
| Gm35066 | ENSMUSG( | ENSMUSG( | 0.462701 | 0.68326 | NO | NO | NA |
| Gpx7 | ENSMUSG( | ENSMUSG( | 1.212253 | 0.012054 | YES | YES | Upregulated |
| Ict1os | ENSMUSG( | ENSMUSG( | 0.260928 | 0.635155 | NO | NO | NA |
| Fbxo24 | ENSMUSG( | ENSMUSG( | -0.38427 | 0.479548 | NO | NO | NA |
| Gm37285 | ENSMUSG( | ENSMUSG( | -0.56153 | 0.148127 | NO | NO | NA |
| Olfr620 | ENSMUSG( | ENSMUSG( | -0.00048 | 0.999598 | NO | NO | NA |
| 4933428G | ENSMUSG( | ENSMUSG( | 0.254302 | 0.829174 | NO | NO | NA |
| Sprn | ENSMUSG( | ENSMUSG( | 1.259118 | 0.03563 | YES | YES | Upregulated |
| Lst1 | ENSMUSG( | ENSMUSG( | 0.159467 | 0.735803 | NO | NO | NA |
| Gm8454 | ENSMUSG( | ENSMUSG( | 0.319322 | 0.764312 | NO | NO | NA |
| Smim5 | ENSMUSG( | ENSMUSG( | 0.1445 | 0.785803 | NO | NO | NA |
| Gm18988 | ENSMUSG( | ENSMUSG( | -1.55322 | 0.000932 | YES | YES | Downregulated |
| Nrip3 | ENSMUSG( | ENSMUSG( | 0.461346 | 0.454063 | NO | NO | NA |
| Sh2d4b | ENSMUSG( | ENSMUSG( | 0.324404 | 0.578846 | NO | NO | NA |
| Xlr3a | ENSMUSG( | ENSMUSG( | 0.641313 | 0.534088 | NO | NO | NA |
| Rep15 | ENSMUSG( | ENSMUSG( | 0.547538 | 0.484833 | NO | NO | NA |
| Tgm7 | ENSMUSG( | ENSMUSG( | -1.0194 | 0.183598 | NO | NO | NA |
| Adcy8 | ENSMUSG( | ENSMUSG( | 0.725389 | 0.443505 | NO | NO | NA |

| | | | | | | | |
|---|---|---|---|---|---|---|---|
| Gm16348 | ENSMUSG( | ENSMUSG( | 1.171006 | 0.301706 | NO | NO | NA |
| Erc2 | ENSMUSG( | ENSMUSG( | 0.278147 | 0.786739 | NO | NO | NA |
| Lrrc23 | ENSMUSG( | ENSMUSG( | -0.91968 | 0.125145 | NO | NO | NA |
| Gm21955 | ENSMUSG( | ENSMUSG( | 0.673065 | 0.522022 | NO | NO | NA |
| Rnf182 | ENSMUSG( | ENSMUSG( | 0.941459 | 0.416366 | NO | NO | NA |
| A530010L: | ENSMUSG( | ENSMUSG( | -0.36199 | 0.590865 | NO | NO | NA |
| Sez6l | ENSMUSG( | ENSMUSG( | 0.67835 | 0.546654 | NO | NO | NA |
| Gm18752 | ENSMUSG( | ENSMUSG( | -0.90869 | 0.197242 | NO | NO | NA |
| Gm30725 | ENSMUSG( | ENSMUSG( | 0.386394 | 0.74256 | NO | NO | NA |
| Wdr46-ps | ENSMUSG( | ENSMUSG( | 0.144348 | 0.857133 | NO | NO | NA |
| Gm6994 | ENSMUSG( | ENSMUSG( | 0.985628 | 0.401233 | NO | NO | NA |
| Gm12286 | ENSMUSG( | ENSMUSG( | 0.482748 | 0.419502 | NO | NO | NA |
| Tril | ENSMUSG( | ENSMUSG( | 0.14069 | 0.911186 | NO | NO | NA |
| Tdo2 | ENSMUSG( | ENSMUSG( | 0.310835 | 0.406115 | NO | NO | NA |
| Gm49502 | ENSMUSG( | ENSMUSG( | -0.38142 | 0.463888 | NO | NO | NA |
| Gm12968 | ENSMUSG( | ENSMUSG( | 0.169361 | 0.893133 | NO | NO | NA |
| Gm10684 | ENSMUSG( | ENSMUSG( | 1.123988 | 0.149026 | NO | NO | NA |
| ENSMUSG | ENSMUSG( | ENSMUSG( | 0.620024 | 0.412264 | NO | NO | NA |
| Gm48145 | ENSMUSG( | ENSMUSG( | -0.04818 | 0.947272 | NO | NO | NA |
| Olfr411 | ENSMUSG( | ENSMUSG( | 0.553207 | 0.653824 | NO | NO | NA |
| Gm13136 | ENSMUSG( | ENSMUSG( | 0.290602 | 0.543583 | NO | NO | NA |
| Gm37470 | ENSMUSG( | ENSMUSG( | -0.46911 | 0.475441 | NO | NO | NA |
| Cabyr | ENSMUSG( | ENSMUSG( | 0.367011 | 0.664907 | NO | NO | NA |
| Gm43088 | ENSMUSG( | ENSMUSG( | 0.386654 | 0.578309 | NO | NO | NA |
| Gm26871 | ENSMUSG( | ENSMUSG( | 0.891763 | 0.401835 | NO | NO | NA |
| Hectd2 | ENSMUSG( | ENSMUSG( | 0.305764 | 0.518069 | NO | NO | NA |
| ENSMUSG | ENSMUSG( | ENSMUSG( | 0.439957 | 0.362625 | NO | NO | NA |
| Plin1 | ENSMUSG( | ENSMUSG( | 0.540752 | 0.538621 | NO | NO | NA |
| Gm19540 | ENSMUSG( | ENSMUSG( | 1.015411 | 0.324401 | NO | NO | NA |
| Gpr62 | ENSMUSG( | ENSMUSG( | 0.786222 | 0.152477 | NO | NO | NA |
| Cpeb1 | ENSMUSG( | ENSMUSG( | 0.532028 | 0.578024 | NO | NO | NA |
| Lrrc2 | ENSMUSG( | ENSMUSG( | 0.64798 | 0.482762 | NO | NO | NA |
| Gm43990 | ENSMUSG( | ENSMUSG( | 0.622266 | 0.482448 | NO | NO | NA |
| Gm43169 | ENSMUSG( | ENSMUSG( | 0.841846 | 0.516034 | NO | NO | NA |
| Gm43941 | ENSMUSG( | ENSMUSG( | 0.931499 | 0.394006 | NO | NO | NA |
| Gm10688 | ENSMUSG( | ENSMUSG( | 0.930702 | 0.027372 | YES | NO | NA |
| Gm31897 | ENSMUSG( | ENSMUSG( | 0.133564 | 0.722714 | NO | NO | NA |
| 4930524O | ENSMUSG( | ENSMUSG( | 1.246486 | 0.13672 | NO | NO | NA |
| Gsdma | ENSMUSG( | ENSMUSG( | 0.688226 | 0.531711 | NO | NO | NA |
| Ereg | ENSMUSG( | ENSMUSG( | 0.822589 | 0.467463 | NO | NO | NA |
| Gm43102 | ENSMUSG( | ENSMUSG( | 0.584357 | 0.601436 | NO | NO | NA |
| Gm43830 | ENSMUSG( | ENSMUSG( | 1.219992 | 0.158086 | NO | NO | NA |
| Gm43294 | ENSMUSG( | ENSMUSG( | 0.322372 | 0.507358 | NO | NO | NA |
| Gm15953 | ENSMUSG( | ENSMUSG( | 0.178902 | 0.750326 | NO | NO | NA |
| Gm27011 | ENSMUSG( | ENSMUSG( | 0.154833 | 0.732776 | NO | NO | NA |
| Gm19276 | ENSMUSG( | ENSMUSG( | 0.517749 | 0.652996 | NO | NO | NA |
| Rpl19-ps1 | ENSMUSG( | ENSMUSG( | 0.580499 | 0.44899 | NO | NO | NA |

| | | | | | | | |
|---|---|---|---|---|---|---|---|
| Lncppara | ENSMUSG( | ENSMUSG( | 0.869997 | 0.073475 | NO | NO | NA |
| Gm7809 | ENSMUSG( | ENSMUSG( | -0.32414 | 0.390592 | NO | NO | NA |
| Gm5491 | ENSMUSG( | ENSMUSG( | -0.30322 | 0.487278 | NO | NO | NA |
| Gm14083 | ENSMUSG( | ENSMUSG( | 1.814842 | 0.162256 | NO | NO | NA |
| Gm28037 | ENSMUSG( | ENSMUSG( | 0.715363 | 0.810621 | NO | NO | NA |
| Gm48876 | ENSMUSG( | ENSMUSG( | -0.28699 | 0.565197 | NO | NO | NA |
| Gm37638 | ENSMUSG( | ENSMUSG( | 0.923202 | 0.450729 | NO | NO | NA |
| 4930477G | ENSMUSG( | ENSMUSG( | 0.741848 | 0.584439 | NO | NO | NA |
| Cabp4 | ENSMUSG( | ENSMUSG( | 0.164406 | 0.758117 | NO | NO | NA |
| Gm37513 | ENSMUSG( | ENSMUSG( | 1.374194 | 0.265347 | NO | NO | NA |
| Gm32479 | ENSMUSG( | ENSMUSG( | 0.550654 | 0.427576 | NO | NO | NA |
| Olfr1291-p | ENSMUSG( | ENSMUSG( | 0.562457 | 0.658027 | NO | NO | NA |
| Tubg2 | ENSMUSG( | ENSMUSG( | 0.405462 | 0.537489 | NO | NO | NA |
| A230085B | ENSMUSG( | ENSMUSG( | 0.349183 | 0.766456 | NO | NO | NA |
| Vxn | ENSMUSG( | ENSMUSG( | 0.985139 | 0.408569 | NO | NO | NA |
| Fbxo39 | ENSMUSG( | ENSMUSG( | -0.09512 | 0.888832 | NO | NO | NA |
| 1700058P: | ENSMUSG( | ENSMUSG( | -0.16555 | 0.789719 | NO | NO | NA |
| Gm45253 | ENSMUSG( | ENSMUSG( | 0.810762 | 0.448722 | NO | NO | NA |
| Gm44199 | ENSMUSG( | ENSMUSG( | -0.70477 | 0.063821 | NO | NO | NA |
| Gm43186 | ENSMUSG( | ENSMUSG( | 0.766392 | 0.512821 | NO | NO | NA |
| Gm6475 | ENSMUSG( | ENSMUSG( | 0.409798 | 0.708667 | NO | NO | NA |
| Shroom1 | ENSMUSG( | ENSMUSG( | 0.359497 | 0.722873 | NO | NO | NA |
| Zfhx2os | ENSMUSG( | ENSMUSG( | -0.2442 | 0.745283 | NO | NO | NA |
| Zfp947 | ENSMUSG( | ENSMUSG( | 0.186382 | 0.821005 | NO | NO | NA |
| Bcas3os1 | ENSMUSG( | ENSMUSG( | 0.722992 | 0.50674 | NO | NO | NA |
| Gm10323 | ENSMUSG( | ENSMUSG( | 0.525447 | 0.711331 | NO | NO | NA |
| Olfr873 | ENSMUSG( | ENSMUSG( | 0.509954 | 0.698039 | NO | NO | NA |
| Gm43437 | ENSMUSG( | ENSMUSG( | 1.093315 | 0.06479 | NO | NO | NA |
| Fhl4 | ENSMUSG( | ENSMUSG( | 0.210591 | 0.671315 | NO | NO | NA |
| ENSMUSG | ENSMUSG( | ENSMUSG( | 0.784113 | 0.588311 | NO | NO | NA |
| Kcnh7 | ENSMUSG( | ENSMUSG( | 0.398144 | 0.486973 | NO | NO | NA |
| Csf2rb2 | ENSMUSG( | ENSMUSG( | 1.340157 | 0.007102 | YES | YES | Upregulated |
| Gm12932 | ENSMUSG( | ENSMUSG( | 0.294982 | 0.477841 | NO | NO | NA |
| Ighv1-12 | ENSMUSG( | ENSMUSG( | 1.342627 | NA | NO | NO | NA |
| Gm40655 | ENSMUSG( | ENSMUSG( | 1.166455 | 0.298937 | NO | NO | NA |
| Trim58 | ENSMUSG( | ENSMUSG( | 1.254677 | 0.110237 | NO | NO | NA |
| Btg4 | ENSMUSG( | ENSMUSG( | 0.824417 | 0.438742 | NO | NO | NA |
| Gm42972 | ENSMUSG( | ENSMUSG( | 0.97743 | 0.401739 | NO | NO | NA |
| Eno2 | ENSMUSG( | ENSMUSG( | 0.149272 | 0.841321 | NO | NO | NA |
| Gm49730 | ENSMUSG( | ENSMUSG( | -0.15308 | 0.791049 | NO | NO | NA |
| Gm13830 | ENSMUSG( | ENSMUSG( | -0.02896 | 0.963858 | NO | NO | NA |
| Gm14453 | ENSMUSG( | ENSMUSG( | 0.237223 | 0.57351 | NO | NO | NA |
| Gm18716 | ENSMUSG( | ENSMUSG( | 0.419455 | 0.693448 | NO | NO | NA |
| Olfr472 | ENSMUSG( | ENSMUSG( | 1.329569 | 0.182528 | NO | NO | NA |
| Usp17la | ENSMUSG( | ENSMUSG( | 0.377621 | 0.695832 | NO | NO | NA |
| Gm10634 | ENSMUSG( | ENSMUSG( | 0.105359 | 0.882088 | NO | NO | NA |
| Gpr182 | ENSMUSG( | ENSMUSG( | 0.712285 | 0.2893 | NO | NO | NA |

| | | | | | | | |
|---|---|---|---|---|---|---|---|
| Afm | ENSMUSG( | ENSMUSG( | 0.504382 | 0.609502 | NO | NO | NA |
| ENSMUSG | ENSMUSG( | ENSMUSG( | -0.10114 | 0.835562 | NO | NO | NA |
| Olfr1195 | ENSMUSG( | ENSMUSG( | 1.351322 | 0.203472 | NO | NO | NA |
| Gm9392 | ENSMUSG( | ENSMUSG( | -0.12786 | 0.769628 | NO | NO | NA |
| Grin1 | ENSMUSG( | ENSMUSG( | 0.298122 | 0.704074 | NO | NO | NA |
| Msantd1 | ENSMUSG( | ENSMUSG( | 0.094187 | 0.856588 | NO | NO | NA |
| Gm44769 | ENSMUSG( | ENSMUSG( | 0.7674 | 0.501066 | NO | NO | NA |
| Hapln4 | ENSMUSG( | ENSMUSG( | 0.370548 | 0.774632 | NO | NO | NA |
| Vrtn | ENSMUSG( | ENSMUSG( | -0.52261 | 0.600536 | NO | NO | NA |
| Gm46447 | ENSMUSG( | ENSMUSG( | 0.420222 | 0.310195 | NO | NO | NA |
| Sbk2 | ENSMUSG( | ENSMUSG( | -1.16246 | 0.007302 | YES | YES | Downregulated |
| Platr20 | ENSMUSG( | ENSMUSG( | 0.047562 | 0.967852 | NO | NO | NA |
| Hoxd11 | ENSMUSG( | ENSMUSG( | 0.389276 | 0.70225 | NO | NO | NA |
| Anp32b-ps | ENSMUSG( | ENSMUSG( | 0.034507 | 0.942764 | NO | NO | NA |
| AI839979 | ENSMUSG( | ENSMUSG( | -0.04099 | 0.953757 | NO | NO | NA |
| Gm12735 | ENSMUSG( | ENSMUSG( | -0.66369 | 0.409325 | NO | NO | NA |
| ENSMUSG | ENSMUSG( | ENSMUSG( | -0.52273 | 0.302051 | NO | NO | NA |
| Cldn1 | ENSMUSG( | ENSMUSG( | 0.824328 | 0.482229 | NO | NO | NA |
| Gm11434 | ENSMUSG( | ENSMUSG( | 0.681022 | 0.479375 | NO | NO | NA |
| Gm45407 | ENSMUSG( | ENSMUSG( | -0.6813 | 0.187518 | NO | NO | NA |
| AI182371 | ENSMUSG( | ENSMUSG( | 0.194168 | 0.613704 | NO | NO | NA |
| Rps18-ps3 | ENSMUSG( | ENSMUSG( | 0.5444 | 0.455814 | NO | NO | NA |
| Plch1 | ENSMUSG( | ENSMUSG( | 1.220052 | 0.013228 | YES | YES | Upregulated |
| Astn2 | ENSMUSG( | ENSMUSG( | 1.41331 | 0.00404 | YES | YES | Upregulated |
| Gm42692 | ENSMUSG( | ENSMUSG( | 0.824586 | 0.487265 | NO | NO | NA |
| Gm37326 | ENSMUSG( | ENSMUSG( | 0.824704 | 0.63616 | NO | NO | NA |
| Gm10862 | ENSMUSG( | ENSMUSG( | 0.749629 | 0.490475 | NO | NO | NA |
| Gm42455 | ENSMUSG( | ENSMUSG( | 1.103652 | 0.446235 | NO | NO | NA |
| Gm37535 | ENSMUSG( | ENSMUSG( | 0.796635 | 0.34493 | NO | NO | NA |
| Gm30285 | ENSMUSG( | ENSMUSG( | 0.537594 | 0.614865 | NO | NO | NA |
| Hoxa5 | ENSMUSG( | ENSMUSG( | 0.491993 | 0.645387 | NO | NO | NA |
| A930038B | ENSMUSG( | ENSMUSG( | 0.857915 | 0.297135 | NO | NO | NA |
| Gm49625 | ENSMUSG( | ENSMUSG( | 0.202434 | 0.67207 | NO | NO | NA |
| Olfr90 | ENSMUSG( | ENSMUSG( | 0.263568 | 0.808313 | NO | NO | NA |
| Gm37457 | ENSMUSG( | ENSMUSG( | 0.674704 | 0.524192 | NO | NO | NA |
| Gm49085 | ENSMUSG( | ENSMUSG( | -0.84216 | 0.07139 | NO | NO | NA |
| Gm19087 | ENSMUSG( | ENSMUSG( | 0.498618 | 0.646614 | NO | NO | NA |
| 1110025M | ENSMUSG( | ENSMUSG( | -0.51304 | 0.449198 | NO | NO | NA |
| Olfr843 | ENSMUSG( | ENSMUSG( | 0.416087 | 0.674136 | NO | NO | NA |
| A830008E | ENSMUSG( | ENSMUSG( | 0.025357 | 0.980566 | NO | NO | NA |
| Gm6161 | ENSMUSG( | ENSMUSG( | -0.07363 | 0.938123 | NO | NO | NA |
| Gm26640 | ENSMUSG( | ENSMUSG( | 0.151843 | 0.818259 | NO | NO | NA |
| Dcaf12l2 | ENSMUSG( | ENSMUSG( | 0.340283 | 0.716591 | NO | NO | NA |
| Gm4189 | ENSMUSG( | ENSMUSG( | 0.552547 | 0.236987 | NO | NO | NA |
| 4930481B( | ENSMUSG( | ENSMUSG( | -0.40533 | 0.28 | NO | NO | NA |
| D930019O | ENSMUSG( | ENSMUSG( | 0.310588 | 0.761374 | NO | NO | NA |
| Mtmr11 | ENSMUSG( | ENSMUSG( | -0.40013 | 0.476491 | NO | NO | NA |

| | | | | | | | |
|---|---|---|---|---|---|---|---|
| Gsdmc | ENSMUSG( | ENSMUSG( | 0.816019 | 0.538708 | NO | NO | NA |
| Zcwpw2 | ENSMUSG( | ENSMUSG( | -0.07893 | 0.919746 | NO | NO | NA |
| Gm37866 | ENSMUSG( | ENSMUSG( | 1.283388 | 0.283254 | NO | NO | NA |
| Gm49398 | ENSMUSG( | ENSMUSG( | -0.26722 | 0.918995 | NO | NO | NA |
| Krt24 | ENSMUSG( | ENSMUSG( | 1.147179 | 0.284733 | NO | NO | NA |
| ENSMUSG | ENSMUSG( | ENSMUSG( | -0.0693 | 0.89857 | NO | NO | NA |
| Gm45311 | ENSMUSG( | ENSMUSG( | -0.65337 | 0.134274 | NO | NO | NA |
| Arrdc4 | ENSMUSG( | ENSMUSG( | 0.587062 | 0.361887 | NO | NO | NA |
| Olfr543 | ENSMUSG( | ENSMUSG( | 1.03955 | 0.069327 | NO | NO | NA |
| Lgi4 | ENSMUSG( | ENSMUSG( | 0.405544 | 0.61548 | NO | NO | NA |
| Olfr741 | ENSMUSG( | ENSMUSG( | 0.423528 | 0.711222 | NO | NO | NA |
| Olfr1384 | ENSMUSG( | ENSMUSG( | 1.204017 | 0.390053 | NO | NO | NA |
| A430078I( | ENSMUSG( | ENSMUSG( | 0.702325 | 0.360488 | NO | NO | NA |
| Gm43920 | ENSMUSG( | ENSMUSG( | 0.877436 | 0.187923 | NO | NO | NA |
| Gm39214 | ENSMUSG( | ENSMUSG( | 0.450847 | 0.590627 | NO | NO | NA |
| Olfr528-ps | ENSMUSG( | ENSMUSG( | 1.721061 | 0.096441 | NO | NO | NA |
| Pramel42 | ENSMUSG( | ENSMUSG( | 0.189859 | 0.908661 | NO | NO | NA |
| Spag16 | ENSMUSG( | ENSMUSG( | 0.57022 | 0.494121 | NO | NO | NA |
| 4933408J1 | ENSMUSG( | ENSMUSG( | -0.04976 | 0.934182 | NO | NO | NA |
| Gm43499 | ENSMUSG( | ENSMUSG( | 0.390425 | 0.669294 | NO | NO | NA |
| Nkx3-1 | ENSMUSG( | ENSMUSG( | 0.845615 | 0.426754 | NO | NO | NA |
| Gm38563 | ENSMUSG( | ENSMUSG( | 0.650927 | 0.304251 | NO | NO | NA |
| Rpl10-ps6 | ENSMUSG( | ENSMUSG( | -0.0955 | 0.867268 | NO | NO | NA |
| Gm42995 | ENSMUSG( | ENSMUSG( | 1.018633 | 0.377421 | NO | NO | NA |
| Ptgir | ENSMUSG( | ENSMUSG( | 0.486313 | 0.336818 | NO | NO | NA |
| Gm28720 | ENSMUSG( | ENSMUSG( | -0.09363 | 0.930577 | NO | NO | NA |
| Klk7 | ENSMUSG( | ENSMUSG( | 0.709979 | 0.499353 | NO | NO | NA |
| Adgrf2 | ENSMUSG( | ENSMUSG( | 0.646407 | 0.600373 | NO | NO | NA |
| Olfr1152 | ENSMUSG( | ENSMUSG( | 0.86074 | 0.505677 | NO | NO | NA |
| BB218582 | ENSMUSG( | ENSMUSG( | 1.211228 | 0.212043 | NO | NO | NA |
| Mal2 | ENSMUSG( | ENSMUSG( | 0.309243 | 0.822566 | NO | NO | NA |
| Adgrg2 | ENSMUSG( | ENSMUSG( | 0.651247 | 0.532666 | NO | NO | NA |
| Gm43395 | ENSMUSG( | ENSMUSG( | 1.140389 | 0.333497 | NO | NO | NA |
| Gm43483 | ENSMUSG( | ENSMUSG( | 0.431836 | 0.706185 | NO | NO | NA |
| Sh2b2 | ENSMUSG( | ENSMUSG( | 1.134125 | 0.055972 | NO | NO | NA |
| Cdkl4 | ENSMUSG( | ENSMUSG( | 0.117008 | 0.773862 | NO | NO | NA |
| Olfr889 | ENSMUSG( | ENSMUSG( | 0.658266 | 0.534393 | NO | NO | NA |
| Csnk2a1-p | ENSMUSG( | ENSMUSG( | 0.944386 | 0.420436 | NO | NO | NA |
| 4930509H | ENSMUSG( | ENSMUSG( | 0.971423 | 0.176802 | NO | NO | NA |
| Cox5b-ps | ENSMUSG( | ENSMUSG( | 0.101353 | 0.76667 | NO | NO | NA |
| Gm44625 | ENSMUSG( | ENSMUSG( | 0.716806 | 0.452297 | NO | NO | NA |
| Slc16a7 | ENSMUSG( | ENSMUSG( | 0.628498 | 0.518553 | NO | NO | NA |
| En1 | ENSMUSG( | ENSMUSG( | 1.208791 | 0.278631 | NO | NO | NA |
| Gm15872 | ENSMUSG( | ENSMUSG( | -0.77266 | 0.205874 | NO | NO | NA |
| Ephb3 | ENSMUSG( | ENSMUSG( | -1.12814 | 0.083234 | NO | NO | NA |
| Frrs1l | ENSMUSG( | ENSMUSG( | 0.584851 | 0.580206 | NO | NO | NA |
| Sms-ps | ENSMUSG( | ENSMUSG( | 0.024088 | 0.958234 | NO | NO | NA |

| | | | | | | | |
|---|---|---|---|---|---|---|---|
| Apol6 | ENSMUSG( | ENSMUSG( | 0.585669 | 0.657527 | NO | NO | NA |
| ENSMUSG | ENSMUSG( | ENSMUSG( | 0.430822 | 0.446439 | NO | NO | NA |
| Ptprn | ENSMUSG( | ENSMUSG( | 0.823642 | 0.38777 | NO | NO | NA |
| Gm15200 | ENSMUSG( | ENSMUSG( | 0.243484 | 0.658162 | NO | NO | NA |
| Gm2412 | ENSMUSG( | ENSMUSG( | -0.54191 | NA | NO | NO | NA |
| Gm9970 | ENSMUSG( | ENSMUSG( | -0.17128 | 0.729543 | NO | NO | NA |
| 4930583K( | ENSMUSG( | ENSMUSG( | 0.841816 | 0.333264 | NO | NO | NA |
| Grem1 | ENSMUSG( | ENSMUSG( | 0.511058 | 0.623052 | NO | NO | NA |
| Gm4045 | ENSMUSG( | ENSMUSG( | 0.351445 | 0.477193 | NO | NO | NA |
| ENSMUSG | ENSMUSG( | ENSMUSG( | -1.22618 | 0.004972 | YES | YES | Downregulated |
| Gm14303 | ENSMUSG( | ENSMUSG( | 0.51848 | 0.153819 | NO | NO | NA |
| Gm45481 | ENSMUSG( | ENSMUSG( | -0.22377 | 0.645795 | NO | NO | NA |
| Vmn1r51 | ENSMUSG( | ENSMUSG( | 0.47097 | 0.716822 | NO | NO | NA |
| Gm38114 | ENSMUSG( | ENSMUSG( | 1.202815 | 0.223204 | NO | NO | NA |
| Gm28941 | ENSMUSG( | ENSMUSG( | -0.76392 | 0.141138 | NO | NO | NA |
| Vwa3a | ENSMUSG( | ENSMUSG( | 0.722538 | 0.40581 | NO | NO | NA |
| Gm37756 | ENSMUSG( | ENSMUSG( | 0.435005 | 0.518438 | NO | NO | NA |
| Gm14204 | ENSMUSG( | ENSMUSG( | 0.961958 | 0.397264 | NO | NO | NA |
| Gm5197 | ENSMUSG( | ENSMUSG( | 0.947539 | 0.387092 | NO | NO | NA |
| Bambi-ps1 | ENSMUSG( | ENSMUSG( | 0.682332 | 0.122502 | NO | NO | NA |
| Gm48924 | ENSMUSG( | ENSMUSG( | 0.665064 | 0.560755 | NO | NO | NA |
| ENSMUSG | ENSMUSG( | ENSMUSG( | -0.13155 | 0.839537 | NO | NO | NA |
| ENSMUSG( | ENSMUSG( | ENSMUSG( | 0.187931 | 0.803915 | NO | NO | NA |
| Gm13408 | ENSMUSG( | ENSMUSG( | 0.131348 | 0.81153 | NO | NO | NA |
| Gm43942 | ENSMUSG( | ENSMUSG( | 0.640459 | 0.58825 | NO | NO | NA |
| ENSMUSG | ENSMUSG( | ENSMUSG( | 0.199954 | 0.86031 | NO | NO | NA |
| Hpgd | ENSMUSG( | ENSMUSG( | -1.41197 | 0.061736 | NO | NO | NA |
| Acad12 | ENSMUSG( | ENSMUSG( | -0.11191 | 0.810767 | NO | NO | NA |
| Hoxa10 | ENSMUSG( | ENSMUSG( | 0.931679 | 0.238316 | NO | NO | NA |
| Tmprss11c | ENSMUSG( | ENSMUSG( | 0.738812 | 0.517788 | NO | NO | NA |
| Gm47258 | ENSMUSG( | ENSMUSG( | -0.31494 | 0.590846 | NO | NO | NA |
| Slc6a20a | ENSMUSG( | ENSMUSG( | 0.570914 | 0.429185 | NO | NO | NA |
| Gm8566 | ENSMUSG( | ENSMUSG( | 0.402391 | 0.426432 | NO | NO | NA |
| ENSMUSG | ENSMUSG( | ENSMUSG( | 0.431408 | 0.432246 | NO | NO | NA |
| Ppp1r3g | ENSMUSG( | ENSMUSG( | -0.28491 | 0.65264 | NO | NO | NA |
| Gm43720 | ENSMUSG( | ENSMUSG( | 0.000802 | 0.999756 | NO | NO | NA |
| Gm19187 | ENSMUSG( | ENSMUSG( | -0.20134 | 0.650523 | NO | NO | NA |
| 6430500D | ENSMUSG( | ENSMUSG( | 1.242536 | 0.295022 | NO | NO | NA |
| Dach1 | ENSMUSG( | ENSMUSG( | 0.785389 | 0.418047 | NO | NO | NA |
| Gm7496 | ENSMUSG( | ENSMUSG( | -0.53017 | 0.155632 | NO | NO | NA |
| Cldn19 | ENSMUSG( | ENSMUSG( | 0.565795 | 0.593125 | NO | NO | NA |
| Tex11 | ENSMUSG( | ENSMUSG( | 3.19021 | 0.055447 | NO | NO | NA |
| Marco | ENSMUSG( | ENSMUSG( | 1.10544 | 0.010788 | YES | YES | Upregulated |
| Gm45697 | ENSMUSG( | ENSMUSG( | 1.044618 | 0.432526 | NO | NO | NA |
| 4932441J( | ENSMUSG( | ENSMUSG( | 1.477945 | 0.099117 | NO | NO | NA |
| Hck | ENSMUSG( | ENSMUSG( | -0.92506 | 0.04196 | YES | NO | NA |
| Gm44745 | ENSMUSG( | ENSMUSG( | 0.062563 | 0.927457 | NO | NO | NA |

| | | | | | | | |
|---|---|---|---|---|---|---|---|
| Cd63 | ENSMUSG( | ENSMUSG( | 1.001562 | 0.266242 | NO | NO | NA |
| Gm47720 | ENSMUSG( | ENSMUSG( | 0.588969 | 0.582405 | NO | NO | NA |
| Hdhd3 | ENSMUSG( | ENSMUSG( | -0.20687 | 0.628876 | NO | NO | NA |
| Krt1 | ENSMUSG( | ENSMUSG( | 0.874279 | 0.418303 | NO | NO | NA |
| Olfr384 | ENSMUSG( | ENSMUSG( | 1.136146 | 0.275399 | NO | NO | NA |
| Gpc3 | ENSMUSG( | ENSMUSG( | 0.908225 | 0.397319 | NO | NO | NA |
| Gm10209 | ENSMUSG( | ENSMUSG( | 0.057754 | 0.889318 | NO | NO | NA |
| Gm9874 | ENSMUSG( | ENSMUSG( | 0.004997 | 0.994565 | NO | NO | NA |
| Cyp19a1 | ENSMUSG( | ENSMUSG( | 0.928209 | 0.35495 | NO | NO | NA |
| Tlcd5 | ENSMUSG( | ENSMUSG( | 0.878364 | 0.232205 | NO | NO | NA |
| Gapdh-ps1 | ENSMUSG( | ENSMUSG( | -0.8786 | 0.123512 | NO | NO | NA |
| Spi1 | ENSMUSG( | ENSMUSG( | -0.58953 | 0.164596 | NO | NO | NA |
| Gm15880 | ENSMUSG( | ENSMUSG( | 0.576977 | 0.372606 | NO | NO | NA |
| Bmp15 | ENSMUSG( | ENSMUSG( | 0.832472 | 0.457098 | NO | NO | NA |
| Cldn34c3 | ENSMUSG( | ENSMUSG( | 1.11132 | 0.319978 | NO | NO | NA |
| Gm10829 | ENSMUSG( | ENSMUSG( | 0.666013 | 0.580075 | NO | NO | NA |
| Gm48569 | ENSMUSG( | ENSMUSG( | -0.92519 | 0.260999 | NO | NO | NA |
| Olfr285 | ENSMUSG( | ENSMUSG( | -0.23401 | 0.80634 | NO | NO | NA |
| Cstad | ENSMUSG( | ENSMUSG( | -0.39942 | 0.430963 | NO | NO | NA |
| Gm1110 | ENSMUSG( | ENSMUSG( | 0.659485 | 0.511586 | NO | NO | NA |
| Gm42685 | ENSMUSG( | ENSMUSG( | 0.390376 | 0.58825 | NO | NO | NA |
| Barhl1 | ENSMUSG( | ENSMUSG( | 0.642271 | 0.423173 | NO | NO | NA |
| Gm42632 | ENSMUSG( | ENSMUSG( | -0.25882 | 0.691734 | NO | NO | NA |
| Fzd3 | ENSMUSG( | ENSMUSG( | 0.849272 | 0.5075 | NO | NO | NA |
| Gm36070 | ENSMUSG( | ENSMUSG( | 0.336216 | 0.746696 | NO | NO | NA |
| ENSMUSG | ENSMUSG( | ENSMUSG( | 0.54135 | 0.276786 | NO | NO | NA |
| Gm43485 | ENSMUSG( | ENSMUSG( | 0.495739 | 0.672193 | NO | NO | NA |
| ENSMUSG | ENSMUSG( | ENSMUSG( | 0.231295 | 0.669318 | NO | NO | NA |
| Gm43604 | ENSMUSG( | ENSMUSG( | -0.32358 | 0.413484 | NO | NO | NA |
| Gm20511 | ENSMUSG( | ENSMUSG( | -0.28466 | 0.690467 | NO | NO | NA |
| ENSMUSG | ENSMUSG( | ENSMUSG( | -0.00379 | 0.994565 | NO | NO | NA |
| Gm42863 | ENSMUSG( | ENSMUSG( | 0.490098 | 0.581702 | NO | NO | NA |
| Gm38034 | ENSMUSG( | ENSMUSG( | -0.25424 | 0.682983 | NO | NO | NA |
| Gm43846 | ENSMUSG( | ENSMUSG( | 1.010952 | 0.306075 | NO | NO | NA |
| Fgf15 | ENSMUSG( | ENSMUSG( | 1.320372 | 0.23756 | NO | NO | NA |
| Syt15 | ENSMUSG( | ENSMUSG( | 0.773623 | 0.433052 | NO | NO | NA |
| Gm44668 | ENSMUSG( | ENSMUSG( | 0.253696 | 0.736093 | NO | NO | NA |
| 1600012P: | ENSMUSG( | ENSMUSG( | 0.160342 | 0.901184 | NO | NO | NA |
| Gm44049 | ENSMUSG( | ENSMUSG( | 0.925898 | 0.386988 | NO | NO | NA |
| Trnp1 | ENSMUSG( | ENSMUSG( | 2.163965 | 0.042725 | YES | YES | Upregulated |
| Gask1b | ENSMUSG( | ENSMUSG( | 0.617108 | 0.48389 | NO | NO | NA |
| Dazl | ENSMUSG( | ENSMUSG( | 0.330343 | 0.717542 | NO | NO | NA |
| Gm28198 | ENSMUSG( | ENSMUSG( | 1.464874 | 0.542733 | NO | NO | NA |
| Greb1l | ENSMUSG( | ENSMUSG( | -0.14218 | 0.837983 | NO | NO | NA |
| Gm38073 | ENSMUSG( | ENSMUSG( | 0.845344 | 0.454144 | NO | NO | NA |
| 4933412L: | ENSMUSG( | ENSMUSG( | 1.018861 | 0.422508 | NO | NO | NA |
| Olfr814 | ENSMUSG( | ENSMUSG( | 0.732102 | 0.557591 | NO | NO | NA |

| | | | | | | | |
|---|---|---|---|---|---|---|---|
| Prrt4 | ENSMUSG( | ENSMUSG( | 0.802187 | 0.156284 | NO | NO | NA |
| Six3os1 | ENSMUSG( | ENSMUSG( | 0.200754 | 0.85174 | NO | NO | NA |
| Gm34785 | ENSMUSG( | ENSMUSG( | 0.131695 | 0.896865 | NO | NO | NA |
| Enthd1 | ENSMUSG( | ENSMUSG( | 0.451682 | 0.42756 | NO | NO | NA |
| Gm15879 | ENSMUSG( | ENSMUSG( | 0.156932 | 0.731835 | NO | NO | NA |
| B230214G | ENSMUSG( | ENSMUSG( | 0.601691 | 0.571384 | NO | NO | NA |
| Gm14023 | ENSMUSG( | ENSMUSG( | 0.214195 | 0.654781 | NO | NO | NA |
| Rpl18-ps1 | ENSMUSG( | ENSMUSG( | 0.693241 | 0.263743 | NO | NO | NA |
| Gm45289 | ENSMUSG( | ENSMUSG( | 0.481993 | 0.508487 | NO | NO | NA |
| 5830418P: | ENSMUSG( | ENSMUSG( | 2.132488 | 0.00382 | YES | YES | Upregulated |
| Olfr1107 | ENSMUSG( | ENSMUSG( | 0.874986 | 0.481147 | NO | NO | NA |
| Gm28959 | ENSMUSG( | ENSMUSG( | 0.014286 | 0.990073 | NO | NO | NA |
| Dll1 | ENSMUSG( | ENSMUSG( | -0.10577 | 0.828882 | NO | NO | NA |
| Gm42736 | ENSMUSG( | ENSMUSG( | -0.23001 | 0.627244 | NO | NO | NA |
| Emcn | ENSMUSG( | ENSMUSG( | 0.877112 | 0.431344 | NO | NO | NA |
| 1700049E: | ENSMUSG( | ENSMUSG( | 0.627495 | 0.561209 | NO | NO | NA |
| 4930509G | ENSMUSG( | ENSMUSG( | 0.642869 | 0.268921 | NO | NO | NA |
| Arhgap29 | ENSMUSG( | ENSMUSG( | 0.621511 | 0.410712 | NO | NO | NA |
| Matn1 | ENSMUSG( | ENSMUSG( | 0.46113 | 0.583984 | NO | NO | NA |
| Prdx6b | ENSMUSG( | ENSMUSG( | 0.798779 | 0.50712 | NO | NO | NA |
| Gm50245 | ENSMUSG( | ENSMUSG( | 1.130279 | 0.291361 | NO | NO | NA |
| Ighv1-42 | ENSMUSG( | ENSMUSG( | 0.850258 | NA | NO | NO | NA |
| Gm43570 | ENSMUSG( | ENSMUSG( | 0.924694 | 0.400572 | NO | NO | NA |
| Gm19145 | ENSMUSG( | ENSMUSG( | 1.203784 | 0.309494 | NO | NO | NA |
| ENSMUSG( | ENSMUSG( | ENSMUSG( | -0.68607 | 0.359644 | NO | NO | NA |
| Gm10817 | ENSMUSG( | ENSMUSG( | -0.3042 | 0.561491 | NO | NO | NA |
| ENSMUSG | ENSMUSG( | ENSMUSG( | 0.271014 | 0.72269 | NO | NO | NA |
| Gm37109 | ENSMUSG( | ENSMUSG( | 0.393674 | 0.748139 | NO | NO | NA |
| Fbxo40 | ENSMUSG( | ENSMUSG( | -0.22078 | 0.729736 | NO | NO | NA |
| Gm37519 | ENSMUSG( | ENSMUSG( | 0.356375 | 0.645714 | NO | NO | NA |
| 9630050E: | ENSMUSG( | ENSMUSG( | 1.051725 | 0.290871 | NO | NO | NA |
| Gm29562 | ENSMUSG( | ENSMUSG( | -0.62818 | 0.562748 | NO | NO | NA |
| Col6a2 | ENSMUSG( | ENSMUSG( | 0.61989 | 0.133175 | NO | NO | NA |
| Gm31138 | ENSMUSG( | ENSMUSG( | 0.667617 | 0.529352 | NO | NO | NA |
| Gm48515 | ENSMUSG( | ENSMUSG( | 0.370941 | 0.762728 | NO | NO | NA |
| Lilra6 | ENSMUSG( | ENSMUSG( | 0.913877 | 0.383864 | NO | NO | NA |
| Gm53 | ENSMUSG( | ENSMUSG( | 0.675227 | 0.545216 | NO | NO | NA |
| Gm11574 | ENSMUSG( | ENSMUSG( | 0.313457 | 0.704289 | NO | NO | NA |
| Igsf6 | ENSMUSG( | ENSMUSG( | -0.05721 | 0.90247 | NO | NO | NA |
| ENSMUSG | ENSMUSG( | ENSMUSG( | 0.107166 | 0.818756 | NO | NO | NA |
| Esrrg | ENSMUSG( | ENSMUSG( | 0.702112 | 0.465753 | NO | NO | NA |
| Trpc3 | ENSMUSG( | ENSMUSG( | 0.401373 | 0.722103 | NO | NO | NA |
| Gm20457 | ENSMUSG( | ENSMUSG( | 0.341151 | 0.430908 | NO | NO | NA |
| Olfr462 | ENSMUSG( | ENSMUSG( | 1.394862 | 0.208913 | NO | NO | NA |
| Col4a3 | ENSMUSG( | ENSMUSG( | 0.701951 | 0.564209 | NO | NO | NA |
| Gm37447 | ENSMUSG( | ENSMUSG( | -0.41445 | 0.522926 | NO | NO | NA |
| Iglon5 | ENSMUSG( | ENSMUSG( | 0.355078 | 0.718783 | NO | NO | NA |

| | | | | | | | |
|---|---|---|---|---|---|---|---|
| Gm42820 | ENSMUSG( | ENSMUSG( | 0.088027 | 0.888656 | NO | NO | NA |
| Tstd1 | ENSMUSG( | ENSMUSG( | -0.91682 | 0.047925 | YES | NO | NA |
| Rtbdn | ENSMUSG( | ENSMUSG( | 0.34521 | 0.596936 | NO | NO | NA |
| Klrb1 | ENSMUSG( | ENSMUSG( | 0.109228 | 0.906059 | NO | NO | NA |
| ENSMUSG | ENSMUSG( | ENSMUSG( | 0.949505 | 0.241308 | NO | NO | NA |
| Mif-ps4 | ENSMUSG( | ENSMUSG( | 1.219339 | 0.072929 | NO | NO | NA |
| Gm11539 | ENSMUSG( | ENSMUSG( | 0.66269 | 0.298258 | NO | NO | NA |
| Gm44229 | ENSMUSG( | ENSMUSG( | 1.128092 | 0.315849 | NO | NO | NA |
| Pde1a | ENSMUSG( | ENSMUSG( | 0.657058 | 0.552759 | NO | NO | NA |
| Gm52989 | ENSMUSG( | ENSMUSG( | 1.811586 | 0.087876 | NO | NO | NA |
| Gm14036 | ENSMUSG( | ENSMUSG( | 0.788947 | 0.579498 | NO | NO | NA |
| Gm42873 | ENSMUSG( | ENSMUSG( | 0.099078 | 0.850944 | NO | NO | NA |
| Ccdc60 | ENSMUSG( | ENSMUSG( | 0.521383 | 0.677025 | NO | NO | NA |
| A330076H | ENSMUSG( | ENSMUSG( | 1.441363 | 0.214168 | NO | NO | NA |
| Olfr600 | ENSMUSG( | ENSMUSG( | 1.113293 | 0.395365 | NO | NO | NA |
| Tbc1d9 | ENSMUSG( | ENSMUSG( | -1.42063 | 0.004168 | YES | YES | Downregulated |
| Gm42868 | ENSMUSG( | ENSMUSG( | -0.01318 | 0.981038 | NO | NO | NA |
| ENSMUSG | ENSMUSG( | ENSMUSG( | -0.0863 | 0.863994 | NO | NO | NA |
| Gm29154 | ENSMUSG( | ENSMUSG( | 0.780316 | 0.508487 | NO | NO | NA |
| Gpr83 | ENSMUSG( | ENSMUSG( | 1.369173 | 0.207717 | NO | NO | NA |
| Gm19792 | ENSMUSG( | ENSMUSG( | 0.726585 | 0.558679 | NO | NO | NA |
| 4930474N | ENSMUSG( | ENSMUSG( | 0.365096 | 0.751103 | NO | NO | NA |
| Gm12511 | ENSMUSG( | ENSMUSG( | 0.9272 | 0.417592 | NO | NO | NA |
| Gm27198 | ENSMUSG( | ENSMUSG( | 0.599849 | 0.596711 | NO | NO | NA |
| Gm26749 | ENSMUSG( | ENSMUSG( | -0.50364 | 0.288325 | NO | NO | NA |
| Gm10233 | ENSMUSG( | ENSMUSG( | 0.346378 | 0.814223 | NO | NO | NA |
| Gm12116 | ENSMUSG( | ENSMUSG( | 0.401395 | 0.515981 | NO | NO | NA |
| Gm13999 | ENSMUSG( | ENSMUSG( | -0.07851 | 0.950605 | NO | NO | NA |
| Gnat2 | ENSMUSG( | ENSMUSG( | 1.010243 | 0.376739 | NO | NO | NA |
| Gm15455 | ENSMUSG( | ENSMUSG( | 0.120852 | 0.893108 | NO | NO | NA |
| Gadl1 | ENSMUSG( | ENSMUSG( | 0.685089 | 0.062097 | NO | NO | NA |
| Gm37962 | ENSMUSG( | ENSMUSG( | -0.37011 | 0.596134 | NO | NO | NA |
| 0610005C: | ENSMUSG( | ENSMUSG( | 0.308305 | 0.656664 | NO | NO | NA |
| 4930421C: | ENSMUSG( | ENSMUSG( | 0.689136 | 0.59053 | NO | NO | NA |
| Pou5f1 | ENSMUSG( | ENSMUSG( | 1.549671 | 0.007181 | YES | YES | Upregulated |
| Rnf208 | ENSMUSG( | ENSMUSG( | 0.265265 | 0.614855 | NO | NO | NA |
| Gm31121 | ENSMUSG( | ENSMUSG( | 0.436448 | 0.670459 | NO | NO | NA |
| Spns3 | ENSMUSG( | ENSMUSG( | 0.652931 | 0.062188 | NO | NO | NA |
| Hdx | ENSMUSG( | ENSMUSG( | -0.61541 | 0.673364 | NO | NO | NA |
| Alx4 | ENSMUSG( | ENSMUSG( | 0.301005 | 0.767433 | NO | NO | NA |
| ENSMUSG | ENSMUSG( | ENSMUSG( | 0.273148 | 0.72833 | NO | NO | NA |
| Fhitos | ENSMUSG( | ENSMUSG( | 0.813187 | 0.518236 | NO | NO | NA |
| Gm2155 | ENSMUSG( | ENSMUSG( | 0.693124 | 0.513389 | NO | NO | NA |
| ENSMUSG( | ENSMUSG( | ENSMUSG( | 0.677148 | 0.527026 | NO | NO | NA |
| Olfr1225 | ENSMUSG( | ENSMUSG( | 0.498946 | 0.635203 | NO | NO | NA |
| Gm42473 | ENSMUSG( | ENSMUSG( | -0.28754 | 0.80012 | NO | NO | NA |
| A530058N | ENSMUSG( | ENSMUSG( | 0.360459 | 0.759918 | NO | NO | NA |

| | | | | | | | |
|---|---|---|---|---|---|---|---|
| ENSMUSG( | ENSMUSG( | ENSMUSG( | 0.653906 | 0.609087 | NO | NO | NA |
| Tex14 | ENSMUSG( | ENSMUSG( | -0.03235 | 0.936489 | NO | NO | NA |
| Rps13-ps4 | ENSMUSG( | ENSMUSG( | 0.248859 | 0.636965 | NO | NO | NA |
| Gm7700 | ENSMUSG( | ENSMUSG( | 1.229293 | 0.299905 | NO | NO | NA |
| Dapl1 | ENSMUSG( | ENSMUSG( | 0.125762 | 0.895165 | NO | NO | NA |
| 1700065D | ENSMUSG( | ENSMUSG( | 0.228067 | 0.66539 | NO | NO | NA |
| Gm43650 | ENSMUSG( | ENSMUSG( | -0.50421 | 0.644476 | NO | NO | NA |
| Gm20594 | ENSMUSG( | ENSMUSG( | 0.88137 | 0.470957 | NO | NO | NA |
| Gm43445 | ENSMUSG( | ENSMUSG( | -0.83103 | 0.254696 | NO | NO | NA |
| Gm45179 | ENSMUSG( | ENSMUSG( | 0.445417 | 0.733562 | NO | NO | NA |
| Olfr1346 | ENSMUSG( | ENSMUSG( | 1.230013 | 0.369406 | NO | NO | NA |
| Gm2396 | ENSMUSG( | ENSMUSG( | 0.779229 | 0.432065 | NO | NO | NA |
| Vill | ENSMUSG( | ENSMUSG( | 0.052078 | 0.937576 | NO | NO | NA |
| Atp8b1 | ENSMUSG( | ENSMUSG( | 0.250954 | 0.793871 | NO | NO | NA |
| Gm3226 | ENSMUSG( | ENSMUSG( | 0.140472 | 0.71024 | NO | NO | NA |
| ENSMUSG( | ENSMUSG( | ENSMUSG( | 1.12288 | 0.26752 | NO | NO | NA |
| Prelid3a | ENSMUSG( | ENSMUSG( | 0.825108 | 0.055642 | NO | NO | NA |
| Gm42819 | ENSMUSG( | ENSMUSG( | 0.663259 | 0.388271 | NO | NO | NA |
| Tma7-ps | ENSMUSG( | ENSMUSG( | -0.16553 | 0.721604 | NO | NO | NA |
| Gm39090 | ENSMUSG( | ENSMUSG( | 0.36863 | 0.68683 | NO | NO | NA |
| Gm36199 | ENSMUSG( | ENSMUSG( | 0.796769 | 0.528041 | NO | NO | NA |
| ENSMUSG | ENSMUSG( | ENSMUSG( | -0.53177 | 0.171618 | NO | NO | NA |
| Vmn2r112 | ENSMUSG( | ENSMUSG( | 0.54045 | 0.609081 | NO | NO | NA |
| Gm16540 | ENSMUSG( | ENSMUSG( | 0.836422 | 0.343905 | NO | NO | NA |
| Olfr887 | ENSMUSG( | ENSMUSG( | 1.220348 | 0.365952 | NO | NO | NA |
| 6330408M | ENSMUSG( | ENSMUSG( | 1.118142 | 0.03717 | YES | YES | Upregulated |
| Gal3st4 | ENSMUSG( | ENSMUSG( | -0.1624 | 0.920601 | NO | NO | NA |
| Gm43670 | ENSMUSG( | ENSMUSG( | 0.88798 | 0.432526 | NO | NO | NA |
| Gm13577 | ENSMUSG( | ENSMUSG( | 0.049371 | 0.939373 | NO | NO | NA |
| Gm16574 | ENSMUSG( | ENSMUSG( | -0.39596 | 0.383051 | NO | NO | NA |
| Fgf10 | ENSMUSG( | ENSMUSG( | 0.632243 | 0.523906 | NO | NO | NA |
| Gm37008 | ENSMUSG( | ENSMUSG( | -0.29309 | 0.605695 | NO | NO | NA |
| Chrna1os | ENSMUSG( | ENSMUSG( | 0.838304 | 0.257847 | NO | NO | NA |
| Proca1 | ENSMUSG( | ENSMUSG( | 0.387735 | 0.630359 | NO | NO | NA |
| Wdr38 | ENSMUSG( | ENSMUSG( | -0.06371 | 0.90825 | NO | NO | NA |
| Pzp | ENSMUSG( | ENSMUSG( | 0.707845 | 0.544716 | NO | NO | NA |
| Gm7045 | ENSMUSG( | ENSMUSG( | 0.366001 | 0.442677 | NO | NO | NA |
| ENSMUSG( | ENSMUSG( | ENSMUSG( | 0.387477 | 0.545774 | NO | NO | NA |
| Gm33280 | ENSMUSG( | ENSMUSG( | -0.37159 | 0.480832 | NO | NO | NA |
| Gm49886 | ENSMUSG( | ENSMUSG( | -0.14311 | 0.747571 | NO | NO | NA |
| Adamtsl2 | ENSMUSG( | ENSMUSG( | 1.314709 | 0.33979 | NO | NO | NA |
| Evc | ENSMUSG( | ENSMUSG( | 0.229049 | 0.540709 | NO | NO | NA |
| Pianp | ENSMUSG( | ENSMUSG( | -0.3747 | 0.495121 | NO | NO | NA |
| Tceal1 | ENSMUSG( | ENSMUSG( | 0.653934 | 0.553739 | NO | NO | NA |
| 2900089D | ENSMUSG( | ENSMUSG( | -0.01151 | 0.987528 | NO | NO | NA |
| Chrna2 | ENSMUSG( | ENSMUSG( | -0.10613 | 0.757236 | NO | NO | NA |
| Prss36 | ENSMUSG( | ENSMUSG( | 0.155599 | 0.800263 | NO | NO | NA |

| | | | | | | | |
|---|---|---|---|---|---|---|---|
| Gm17764 | ENSMUSG( | ENSMUSG( | -0.97127 | 0.035784 | YES | NO | NA |
| Vmn2r73 | ENSMUSG( | ENSMUSG( | 0.347706 | 0.731835 | NO | NO | NA |
| Gm43868 | ENSMUSG( | ENSMUSG( | 0.815123 | 0.380024 | NO | NO | NA |
| Gm37848 | ENSMUSG( | ENSMUSG( | 0.494361 | 0.654684 | NO | NO | NA |
| Sspnos | ENSMUSG( | ENSMUSG( | 1.261604 | 0.277347 | NO | NO | NA |
| Gm49390 | ENSMUSG( | ENSMUSG( | 0.624019 | 0.558673 | NO | NO | NA |
| Rnf32 | ENSMUSG( | ENSMUSG( | 0.832182 | 0.397767 | NO | NO | NA |
| Myoz3 | ENSMUSG( | ENSMUSG( | 0.950264 | 0.221997 | NO | NO | NA |
| Gm9993 | ENSMUSG( | ENSMUSG( | -0.31466 | 0.53935 | NO | NO | NA |
| C130083N | ENSMUSG( | ENSMUSG( | 0.157951 | 0.741182 | NO | NO | NA |
| B430219N | ENSMUSG( | ENSMUSG( | -0.06309 | 0.916645 | NO | NO | NA |
| 4930590J( | ENSMUSG( | ENSMUSG( | 0.044551 | 0.935826 | NO | NO | NA |
| Icam4 | ENSMUSG( | ENSMUSG( | 1.406867 | 0.0812 | NO | NO | NA |
| Gm44043 | ENSMUSG( | ENSMUSG( | 0.824844 | 0.463661 | NO | NO | NA |
| Gm16892 | ENSMUSG( | ENSMUSG( | -0.57193 | 0.182925 | NO | NO | NA |
| Phactr3 | ENSMUSG( | ENSMUSG( | -1.49551 | 0.246275 | NO | NO | NA |
| Plxnb2 | ENSMUSG( | ENSMUSG( | 0.288713 | 0.496564 | NO | NO | NA |
| Gm17879 | ENSMUSG( | ENSMUSG( | 0.234855 | 0.660808 | NO | NO | NA |
| Gm5560 | ENSMUSG( | ENSMUSG( | 0.466554 | 0.389897 | NO | NO | NA |
| ENSMUSG | ENSMUSG( | ENSMUSG( | 0.684981 | 0.494121 | NO | NO | NA |
| Gm17841 | ENSMUSG( | ENSMUSG( | 0.52467 | 0.652228 | NO | NO | NA |
| Sntg1 | ENSMUSG( | ENSMUSG( | 0.977578 | 0.358705 | NO | NO | NA |
| Muc3a | ENSMUSG( | ENSMUSG( | -1.00708 | 0.044942 | YES | YES | Downregulated |
| Gm44951 | ENSMUSG( | ENSMUSG( | 0.508219 | 0.609667 | NO | NO | NA |
| Bpifb4 | ENSMUSG( | ENSMUSG( | 0.648225 | 0.197319 | NO | NO | NA |
| Ceacam20 | ENSMUSG( | ENSMUSG( | 1.207187 | 0.321293 | NO | NO | NA |
| Gm44152 | ENSMUSG( | ENSMUSG( | 0.097005 | 0.901916 | NO | NO | NA |
| Pmch | ENSMUSG( | ENSMUSG( | -0.12892 | 0.816509 | NO | NO | NA |
| Sox17 | ENSMUSG( | ENSMUSG( | 1.025429 | 0.36325 | NO | NO | NA |
| Igkv6-13 | ENSMUSG( | ENSMUSG( | -4.26023 | NA | NO | NO | NA |
| Gm12792 | ENSMUSG( | ENSMUSG( | 0.231209 | 0.827325 | NO | NO | NA |
| C8g | ENSMUSG( | ENSMUSG( | 0.631456 | 0.161847 | NO | NO | NA |
| Gm38085 | ENSMUSG( | ENSMUSG( | 0.843774 | 0.422361 | NO | NO | NA |
| Sh3bgrl2 | ENSMUSG( | ENSMUSG( | 0.246113 | 0.766456 | NO | NO | NA |
| Olfr461 | ENSMUSG( | ENSMUSG( | 1.627636 | 0.16516 | NO | NO | NA |
| Gm18558 | ENSMUSG( | ENSMUSG( | 1.12191 | 0.484233 | NO | NO | NA |
| Sp9 | ENSMUSG( | ENSMUSG( | 0.062875 | 0.92309 | NO | NO | NA |
| Gm44644 | ENSMUSG( | ENSMUSG( | 0.67481 | 0.542733 | NO | NO | NA |
| Gm3509 | ENSMUSG( | ENSMUSG( | 0.562479 | 0.666819 | NO | NO | NA |
| Gm45259 | ENSMUSG( | ENSMUSG( | 0.93608 | 0.372481 | NO | NO | NA |
| Gm12994 | ENSMUSG( | ENSMUSG( | 2.96521 | 0.03684 | YES | YES | Upregulated |
| Gm7324 | ENSMUSG( | ENSMUSG( | 0.212842 | 0.681405 | NO | NO | NA |
| Sh3pxd2a | ENSMUSG( | ENSMUSG( | -0.30392 | 0.627656 | NO | NO | NA |
| Gm48236 | ENSMUSG( | ENSMUSG( | 0.699621 | 0.518553 | NO | NO | NA |
| Syngr1 | ENSMUSG( | ENSMUSG( | 1.31047 | 0.034552 | YES | YES | Upregulated |
| Dynlt4 | ENSMUSG( | ENSMUSG( | 0.751407 | 0.171008 | NO | NO | NA |
| Gm47794 | ENSMUSG( | ENSMUSG( | -0.55722 | 0.313759 | NO | NO | NA |

| | | | | | | | |
|---|---|---|---|---|---|---|---|
| A3galt2 | ENSMUSG( | ENSMUSG( | 0.673222 | 0.377555 | NO | NO | NA |
| Rnf180 | ENSMUSG( | ENSMUSG( | 0.372711 | 0.685034 | NO | NO | NA |
| Map2k3os | ENSMUSG( | ENSMUSG( | 0.159351 | 0.736727 | NO | NO | NA |
| Gm9755 | ENSMUSG( | ENSMUSG( | 0.234218 | 0.656466 | NO | NO | NA |
| Apof | ENSMUSG( | ENSMUSG( | 0.785234 | 0.474071 | NO | NO | NA |
| Ighv1-34 | ENSMUSG( | ENSMUSG( | -6.51682 | 0.000314 | YES | YES | Downregulated |
| ENSMUSG | ENSMUSG( | ENSMUSG( | -0.18046 | 0.694528 | NO | NO | NA |
| Il17re | ENSMUSG( | ENSMUSG( | 0.334953 | 0.765086 | NO | NO | NA |
| Alkbh3os1 | ENSMUSG( | ENSMUSG( | -0.64552 | 0.394648 | NO | NO | NA |
| ENSMUSG( | ENSMUSG( | ENSMUSG( | -0.39184 | 0.445814 | NO | NO | NA |
| Gm43332 | ENSMUSG( | ENSMUSG( | 0.292605 | 0.776769 | NO | NO | NA |
| Gm15302 | ENSMUSG( | ENSMUSG( | 0.165064 | 0.888443 | NO | NO | NA |
| Sdsl | ENSMUSG( | ENSMUSG( | 1.304363 | 0.132579 | NO | NO | NA |
| Gm19000 | ENSMUSG( | ENSMUSG( | 0.589372 | 0.546654 | NO | NO | NA |
| Gm2961 | ENSMUSG( | ENSMUSG( | 1.042011 | 0.417592 | NO | NO | NA |
| Ddit4l | ENSMUSG( | ENSMUSG( | -3.04174 | 0.004962 | YES | YES | Downregulated |
| ENSMUSG | ENSMUSG( | ENSMUSG( | -0.08696 | 0.831728 | NO | NO | NA |
| Gm48939 | ENSMUSG( | ENSMUSG( | 0.171001 | 0.846149 | NO | NO | NA |
| Fam20c | ENSMUSG( | ENSMUSG( | 0.887174 | 0.356412 | NO | NO | NA |
| Kcna7 | ENSMUSG( | ENSMUSG( | 0.454734 | 0.684648 | NO | NO | NA |
| Etos1 | ENSMUSG( | ENSMUSG( | 0.2045 | 0.840237 | NO | NO | NA |
| Gm16133 | ENSMUSG( | ENSMUSG( | 0.802597 | 0.238548 | NO | NO | NA |
| Gal3st1 | ENSMUSG( | ENSMUSG( | 0.332544 | 0.642321 | NO | NO | NA |
| 1700126G | ENSMUSG( | ENSMUSG( | -0.14572 | 0.777011 | NO | NO | NA |
| Gm11573 | ENSMUSG( | ENSMUSG( | -0.45831 | 0.472454 | NO | NO | NA |
| ENSMUSG( | ENSMUSG( | ENSMUSG( | 0.546494 | 0.593014 | NO | NO | NA |
| Mecom | ENSMUSG( | ENSMUSG( | 0.397112 | 0.639073 | NO | NO | NA |
| Gm37647 | ENSMUSG( | ENSMUSG( | 1.249335 | 0.365768 | NO | NO | NA |
| Mmp8 | ENSMUSG( | ENSMUSG( | 1.105191 | 0.268296 | NO | NO | NA |
| Gm49373 | ENSMUSG( | ENSMUSG( | 0.441133 | 0.629334 | NO | NO | NA |
| Vmn1r204 | ENSMUSG( | ENSMUSG( | 0.887604 | 0.513234 | NO | NO | NA |
| ENSMUSG | ENSMUSG( | ENSMUSG( | 0.769389 | 0.325929 | NO | NO | NA |
| Gm33378 | ENSMUSG( | ENSMUSG( | -0.26771 | 0.690986 | NO | NO | NA |
| Gm5648 | ENSMUSG( | ENSMUSG( | 0.671652 | 0.183032 | NO | NO | NA |
| ENSMUSG | ENSMUSG( | ENSMUSG( | 1.052586 | 0.058486 | NO | NO | NA |
| Gm14150 | ENSMUSG( | ENSMUSG( | -0.01732 | 0.972271 | NO | NO | NA |
| Olfr1033 | ENSMUSG( | ENSMUSG( | 0.253573 | 0.80032 | NO | NO | NA |
| Gm10268 | ENSMUSG( | ENSMUSG( | 1.345478 | 0.110439 | NO | NO | NA |
| Slc9a3r2 | ENSMUSG( | ENSMUSG( | 0.772834 | 0.252227 | NO | NO | NA |
| Gm16477 | ENSMUSG( | ENSMUSG( | 0.886485 | 0.046642 | YES | NO | NA |
| Gm42966 | ENSMUSG( | ENSMUSG( | 0.757338 | 0.474914 | NO | NO | NA |
| Pilra | ENSMUSG( | ENSMUSG( | 0.41915 | 0.623206 | NO | NO | NA |
| Gm42711 | ENSMUSG( | ENSMUSG( | 0.145011 | 0.883697 | NO | NO | NA |
| Hsf5 | ENSMUSG( | ENSMUSG( | 0.0792 | 0.940462 | NO | NO | NA |
| Ugt1a10 | ENSMUSG( | ENSMUSG( | 2.120206 | 0.344431 | NO | NO | NA |
| Gm49871 | ENSMUSG( | ENSMUSG( | 0.393479 | 0.535657 | NO | NO | NA |
| Jam2 | ENSMUSG( | ENSMUSG( | -0.297 | 0.549104 | NO | NO | NA |

| | | | | | | | |
|---|---|---|---|---|---|---|---|
| Gm38309 | ENSMUSG( | ENSMUSG( | 0.192928 | 0.74816 | NO | NO | NA |
| Gpr85 | ENSMUSG( | ENSMUSG( | -0.44817 | 0.691734 | NO | NO | NA |
| Cstdc4 | ENSMUSG( | ENSMUSG( | 0.700244 | 0.46417 | NO | NO | NA |
| Il22ra2 | ENSMUSG( | ENSMUSG( | 0.277079 | 0.829293 | NO | NO | NA |
| Vmn2r-ps1 | ENSMUSG( | ENSMUSG( | 1.558033 | 0.242187 | NO | NO | NA |
| Adcyap1 | ENSMUSG( | ENSMUSG( | 1.047382 | 0.438698 | NO | NO | NA |
| Gm11788 | ENSMUSG( | ENSMUSG( | 0.455047 | 0.449417 | NO | NO | NA |
| Gm37759 | ENSMUSG( | ENSMUSG( | 0.037107 | 0.91639 | NO | NO | NA |
| Gm48936 | ENSMUSG( | ENSMUSG( | 0.671683 | 0.420191 | NO | NO | NA |
| Gm6252 | ENSMUSG( | ENSMUSG( | 0.37999 | 0.474373 | NO | NO | NA |
| Krt82 | ENSMUSG( | ENSMUSG( | 0.438866 | 0.73408 | NO | NO | NA |
| Rcan2 | ENSMUSG( | ENSMUSG( | 0.450981 | 0.677728 | NO | NO | NA |
| Gm12009 | ENSMUSG( | ENSMUSG( | 0.052869 | 0.950071 | NO | NO | NA |
| Cd79a | ENSMUSG( | ENSMUSG( | -0.30693 | 0.583116 | NO | NO | NA |
| 9330159F1 | ENSMUSG( | ENSMUSG( | 0.902209 | 0.424893 | NO | NO | NA |
| Alpl | ENSMUSG( | ENSMUSG( | 0.062749 | 0.879511 | NO | NO | NA |
| Necab3 | ENSMUSG( | ENSMUSG( | -0.30397 | 0.60683 | NO | NO | NA |
| Gm40518 | ENSMUSG( | ENSMUSG( | 0.025586 | 0.985317 | NO | NO | NA |
| Gm42413 | ENSMUSG( | ENSMUSG( | -0.04452 | 0.977984 | NO | NO | NA |
| Asphd1 | ENSMUSG( | ENSMUSG( | 0.066604 | 0.926428 | NO | NO | NA |
| 6030440G | ENSMUSG( | ENSMUSG( | 0.73648 | 0.50518 | NO | NO | NA |
| Gm15518 | ENSMUSG( | ENSMUSG( | 1.818343 | 0.019009 | YES | YES | Upregulated |
| Gpx4-ps2 | ENSMUSG( | ENSMUSG( | 2.105048 | NA | NO | NO | NA |
| Olfr446 | ENSMUSG( | ENSMUSG( | 0.768476 | 0.502992 | NO | NO | NA |
| Gm12479 | ENSMUSG( | ENSMUSG( | 1.250438 | 0.372473 | NO | NO | NA |
| Cntn2 | ENSMUSG( | ENSMUSG( | 0.774748 | 0.479784 | NO | NO | NA |
| Saa3 | ENSMUSG( | ENSMUSG( | 0.548268 | 0.581594 | NO | NO | NA |
| Nkain3 | ENSMUSG( | ENSMUSG( | 0.881285 | 0.449488 | NO | NO | NA |
| Gm40932 | ENSMUSG( | ENSMUSG( | 1.110178 | 0.278359 | NO | NO | NA |
| Gm18878 | ENSMUSG( | ENSMUSG( | 0.995777 | 0.412416 | NO | NO | NA |
| ENSMUSG | ENSMUSG( | ENSMUSG( | 0.155304 | 0.7454 | NO | NO | NA |
| Gm45797 | ENSMUSG( | ENSMUSG( | -0.73859 | 0.134692 | NO | NO | NA |
| ENSMUSG | ENSMUSG( | ENSMUSG( | 0.503221 | 0.592341 | NO | NO | NA |
| Gm14546 | ENSMUSG( | ENSMUSG( | 0.708289 | 0.541019 | NO | NO | NA |
| Pcdhb6 | ENSMUSG( | ENSMUSG( | 0.258184 | 0.830186 | NO | NO | NA |
| Slc5a11 | ENSMUSG( | ENSMUSG( | 0.105025 | 0.841227 | NO | NO | NA |
| ENSMUSG( | ENSMUSG( | ENSMUSG( | -1.03445 | 0.055229 | NO | NO | NA |
| Gm18166 | ENSMUSG( | ENSMUSG( | -0.42046 | 0.88733 | NO | NO | NA |
| Gm16275 | ENSMUSG( | ENSMUSG( | 0.419941 | 0.732315 | NO | NO | NA |
| Gm48743 | ENSMUSG( | ENSMUSG( | 0.572966 | 0.613046 | NO | NO | NA |
| ENSMUSG( | ENSMUSG( | ENSMUSG( | -0.09742 | 0.812357 | NO | NO | NA |
| Gm43363 | ENSMUSG( | ENSMUSG( | 1.057252 | 0.331156 | NO | NO | NA |
| Gm8489 | ENSMUSG( | ENSMUSG( | 0.969328 | 0.428679 | NO | NO | NA |
| Slc22a21 | ENSMUSG( | ENSMUSG( | 0.834918 | 0.128905 | NO | NO | NA |
| Phxr4 | ENSMUSG( | ENSMUSG( | -0.0913 | 0.909884 | NO | NO | NA |
| Cnr2 | ENSMUSG( | ENSMUSG( | 0.328309 | 0.677728 | NO | NO | NA |
| Cfap97d2 | ENSMUSG( | ENSMUSG( | 0.773022 | 0.453495 | NO | NO | NA |

| | | | | | | | |
|---|---|---|---|---|---|---|---|
| Mpped2 | ENSMUSG( | ENSMUSG( | 0.406753 | 0.558407 | NO | NO | NA |
| ENSMUSG | ENSMUSG( | ENSMUSG( | 0.597569 | 0.254038 | NO | NO | NA |
| Rbm20 | ENSMUSG( | ENSMUSG( | -0.48006 | 0.346871 | NO | NO | NA |
| Rps15-ps2 | ENSMUSG( | ENSMUSG( | 0.353502 | 0.501916 | NO | NO | NA |
| Prlhr | ENSMUSG( | ENSMUSG( | 2.345508 | 0.065767 | NO | NO | NA |
| Gm49785 | ENSMUSG( | ENSMUSG( | 0.667207 | 0.154375 | NO | NO | NA |
| Gm16364 | ENSMUSG( | ENSMUSG( | -0.87387 | 0.055365 | NO | NO | NA |
| Gm49164 | ENSMUSG( | ENSMUSG( | 0.081269 | 0.867506 | NO | NO | NA |
| Col22a1 | ENSMUSG( | ENSMUSG( | 0.735207 | 0.576025 | NO | NO | NA |
| Gm44707 | ENSMUSG( | ENSMUSG( | 0.357146 | 0.745859 | NO | NO | NA |
| Gm38156 | ENSMUSG( | ENSMUSG( | 0.449504 | 0.678481 | NO | NO | NA |
| Gm21388 | ENSMUSG( | ENSMUSG( | 0.787055 | 0.560369 | NO | NO | NA |
| Shox2 | ENSMUSG( | ENSMUSG( | 0.421604 | 0.710086 | NO | NO | NA |
| Gm49795 | ENSMUSG( | ENSMUSG( | 0.818388 | 0.476271 | NO | NO | NA |
| Gm43285 | ENSMUSG( | ENSMUSG( | 0.616038 | 0.57329 | NO | NO | NA |
| Slc22a8 | ENSMUSG( | ENSMUSG( | 0.210777 | 0.859925 | NO | NO | NA |
| Gm7266 | ENSMUSG( | ENSMUSG( | 0.644537 | 0.272626 | NO | NO | NA |
| Dsg1a | ENSMUSG( | ENSMUSG( | 0.627206 | 0.63825 | NO | NO | NA |
| Gm38116 | ENSMUSG( | ENSMUSG( | 0.590928 | 0.599709 | NO | NO | NA |
| 2810471M | ENSMUSG( | ENSMUSG( | 0.414515 | 0.683119 | NO | NO | NA |
| 1110020A | ENSMUSG( | ENSMUSG( | 0.044116 | 0.927405 | NO | NO | NA |
| Gm48137 | ENSMUSG( | ENSMUSG( | 0.185173 | 0.754639 | NO | NO | NA |
| Olfr1281 | ENSMUSG( | ENSMUSG( | 0.527241 | 0.647804 | NO | NO | NA |
| Gm37174 | ENSMUSG( | ENSMUSG( | -0.01253 | 0.980729 | NO | NO | NA |
| Zbtb10 | ENSMUSG( | ENSMUSG( | 0.654237 | 0.41927 | NO | NO | NA |
| Gm42537 | ENSMUSG( | ENSMUSG( | -0.06851 | 0.954204 | NO | NO | NA |
| Gm29072 | ENSMUSG( | ENSMUSG( | 0.72913 | 0.475084 | NO | NO | NA |
| Gm48045 | ENSMUSG( | ENSMUSG( | -0.30591 | 0.422983 | NO | NO | NA |
| Cdk3 | ENSMUSG( | ENSMUSG( | 0.439614 | 0.415514 | NO | NO | NA |
| Cd177 | ENSMUSG( | ENSMUSG( | 2.059237 | 0.164596 | NO | NO | NA |
| Gm47664 | ENSMUSG( | ENSMUSG( | 0.016401 | 0.985161 | NO | NO | NA |
| Gpr45 | ENSMUSG( | ENSMUSG( | 1.44232 | 0.048707 | YES | YES | Upregulated |
| Gm16259 | ENSMUSG( | ENSMUSG( | -0.22232 | 0.650865 | NO | NO | NA |
| Dgat2l6 | ENSMUSG( | ENSMUSG( | 0.988159 | 0.286304 | NO | NO | NA |
| Gm4258 | ENSMUSG( | ENSMUSG( | 0.108303 | 0.909337 | NO | NO | NA |
| Gm42436 | ENSMUSG( | ENSMUSG( | 0.697068 | 0.265897 | NO | NO | NA |
| ENSMUSG | ENSMUSG( | ENSMUSG( | -0.13943 | 0.817522 | NO | NO | NA |
| Gm48914 | ENSMUSG( | ENSMUSG( | -0.01347 | 0.983122 | NO | NO | NA |
| Gm3045 | ENSMUSG( | ENSMUSG( | 0.506604 | 0.690197 | NO | NO | NA |
| Gm48265 | ENSMUSG( | ENSMUSG( | 0.668137 | 0.544332 | NO | NO | NA |
| Igdcc4 | ENSMUSG( | ENSMUSG( | 1.012539 | 0.166659 | NO | NO | NA |
| Gm20387 | ENSMUSG( | ENSMUSG( | 0.711945 | 0.516268 | NO | NO | NA |
| Gm30262 | ENSMUSG( | ENSMUSG( | 0.196491 | 0.629727 | NO | NO | NA |
| Klri2 | ENSMUSG( | ENSMUSG( | 0.341701 | 0.579897 | NO | NO | NA |
| 5530401A | ENSMUSG( | ENSMUSG( | 0.450867 | 0.684078 | NO | NO | NA |
| Siah3 | ENSMUSG( | ENSMUSG( | 0.452548 | 0.424383 | NO | NO | NA |
| Frmd3 | ENSMUSG( | ENSMUSG( | 0.327565 | 0.755541 | NO | NO | NA |

| | | | | | | | |
|---|---|---|---|---|---|---|---|
| Ooep | ENSMUSG( | ENSMUSG( | 0.677203 | 0.525434 | NO | NO | NA |
| Notch4 | ENSMUSG( | ENSMUSG( | 0.425917 | 0.356094 | NO | NO | NA |
| Gm16216 | ENSMUSG( | ENSMUSG( | 0.489554 | 0.269346 | NO | NO | NA |
| Mrgprc4-p | ENSMUSG( | ENSMUSG( | 1.275431 | 0.288478 | NO | NO | NA |
| Olfr644 | ENSMUSG( | ENSMUSG( | 1.036128 | 0.337914 | NO | NO | NA |
| Noto | ENSMUSG( | ENSMUSG( | 0.293514 | 0.787833 | NO | NO | NA |
| Gm12333 | ENSMUSG( | ENSMUSG( | 0.499781 | 0.449166 | NO | NO | NA |
| Gm37052 | ENSMUSG( | ENSMUSG( | -0.34596 | 0.606364 | NO | NO | NA |
| Gm9974 | ENSMUSG( | ENSMUSG( | -0.12907 | 0.812414 | NO | NO | NA |
| Gm43788 | ENSMUSG( | ENSMUSG( | 0.211561 | 0.793559 | NO | NO | NA |
| Gm37805 | ENSMUSG( | ENSMUSG( | 0.470068 | 0.351132 | NO | NO | NA |
| Mrc1 | ENSMUSG( | ENSMUSG( | -0.62681 | 0.283594 | NO | NO | NA |
| Lrp2bp | ENSMUSG( | ENSMUSG( | 0.629401 | 0.28125 | NO | NO | NA |
| Npm3-ps1 | ENSMUSG( | ENSMUSG( | 0.975493 | 0.027634 | YES | NO | NA |
| Gm47687 | ENSMUSG( | ENSMUSG( | 0.79256 | 0.512771 | NO | NO | NA |
| Gm39213 | ENSMUSG( | ENSMUSG( | -0.59349 | 0.34315 | NO | NO | NA |
| Cldn34c2 | ENSMUSG( | ENSMUSG( | 0.467176 | 0.713313 | NO | NO | NA |
| Col20a1 | ENSMUSG( | ENSMUSG( | -0.3298 | 0.549104 | NO | NO | NA |
| Fam151a | ENSMUSG( | ENSMUSG( | 0.064198 | 0.970821 | NO | NO | NA |
| Gm42967 | ENSMUSG( | ENSMUSG( | 0.302387 | 0.679095 | NO | NO | NA |
| Gm43891 | ENSMUSG( | ENSMUSG( | 0.037938 | 0.97207 | NO | NO | NA |
| Tppp | ENSMUSG( | ENSMUSG( | -0.72847 | 0.035038 | YES | NO | NA |
| ENSMUSG( | ENSMUSG( | ENSMUSG( | 0.579373 | 0.229256 | NO | NO | NA |
| Gm13370 | ENSMUSG( | ENSMUSG( | 6.586047 | NA | NO | NO | NA |
| Gm19084 | ENSMUSG( | ENSMUSG( | 0.919197 | 0.388497 | NO | NO | NA |
| Gm15344 | ENSMUSG( | ENSMUSG( | -0.22119 | 0.587391 | NO | NO | NA |
| Foxd1 | ENSMUSG( | ENSMUSG( | -0.13211 | 0.893365 | NO | NO | NA |
| Lrig3 | ENSMUSG( | ENSMUSG( | 0.653352 | 0.339365 | NO | NO | NA |
| Edil3 | ENSMUSG( | ENSMUSG( | 0.173152 | 0.872055 | NO | NO | NA |
| Gm48799 | ENSMUSG( | ENSMUSG( | -0.23233 | 0.733563 | NO | NO | NA |
| Gm43981 | ENSMUSG( | ENSMUSG( | 0.953299 | 0.382706 | NO | NO | NA |
| Telomeras | ENSMUSG( | ENSMUSG( | 0.864831 | 0.6204 | NO | NO | NA |
| Ppbp | ENSMUSG( | ENSMUSG( | -2.53088 | 0.000907 | YES | YES | Downregulated |
| Gm18075 | ENSMUSG( | ENSMUSG( | 0.767306 | 0.517828 | NO | NO | NA |
| Gm43637 | ENSMUSG( | ENSMUSG( | 0.955015 | 0.257374 | NO | NO | NA |
| Gm2238 | ENSMUSG( | ENSMUSG( | -0.29895 | 0.734496 | NO | NO | NA |
| Gm38300 | ENSMUSG( | ENSMUSG( | -0.83313 | 0.096123 | NO | NO | NA |
| Gm15527 | ENSMUSG( | ENSMUSG( | -1.70969 | 0.000309 | YES | YES | Downregulated |
| Gm45700 | ENSMUSG( | ENSMUSG( | -2.18478 | 0.435889 | NO | NO | NA |
| Gm30329 | ENSMUSG( | ENSMUSG( | 0.639986 | 0.535806 | NO | NO | NA |
| Gm8784 | ENSMUSG( | ENSMUSG( | 0.253913 | 0.764246 | NO | NO | NA |
| Gm4335 | ENSMUSG( | ENSMUSG( | 0.14909 | 0.861112 | NO | NO | NA |
| Vmn1r181 | ENSMUSG( | ENSMUSG( | 0.182771 | 0.878742 | NO | NO | NA |
| Tlx2 | ENSMUSG( | ENSMUSG( | 1.655067 | 0.107665 | NO | NO | NA |
| Gm42998 | ENSMUSG( | ENSMUSG( | 0.498692 | 0.466606 | NO | NO | NA |
| Zfp286 | ENSMUSG( | ENSMUSG( | 0.323248 | 0.417592 | NO | NO | NA |
| Gm10826 | ENSMUSG( | ENSMUSG( | 0.654157 | 0.477659 | NO | NO | NA |

| | | | | | | | |
|---|---|---|---|---|---|---|---|
| Dgat2 | ENSMUSG( | ENSMUSG( | 0.846573 | 0.16428 | NO | NO | NA |
| Plxna2 | ENSMUSG( | ENSMUSG( | 0.608258 | 0.459198 | NO | NO | NA |
| Slc25a31 | ENSMUSG( | ENSMUSG( | 4.091702 | 0.256595 | NO | NO | NA |
| Gm3696 | ENSMUSG( | ENSMUSG( | 2.017412 | 0.097851 | NO | NO | NA |
| Lrrc51 | ENSMUSG( | ENSMUSG( | 0.113142 | 0.871524 | NO | NO | NA |
| Fgf23 | ENSMUSG( | ENSMUSG( | -0.17104 | 0.725185 | NO | NO | NA |
| Matn2 | ENSMUSG( | ENSMUSG( | 0.025602 | 0.976785 | NO | NO | NA |
| Lrrtm2 | ENSMUSG( | ENSMUSG( | 0.558914 | 0.520226 | NO | NO | NA |
| Gm36888 | ENSMUSG( | ENSMUSG( | 0.149736 | 0.899918 | NO | NO | NA |
| Olfr1120 | ENSMUSG( | ENSMUSG( | 1.044267 | 0.324751 | NO | NO | NA |
| Bhlha15 | ENSMUSG( | ENSMUSG( | -0.59155 | 0.267916 | NO | NO | NA |
| Krt18 | ENSMUSG( | ENSMUSG( | 1.198424 | 0.035902 | YES | YES | Upregulated |
| Gm10080 | ENSMUSG( | ENSMUSG( | -0.20378 | 0.648535 | NO | NO | NA |
| Gm44624 | ENSMUSG( | ENSMUSG( | 0.004182 | 0.994581 | NO | NO | NA |
| Gm47530 | ENSMUSG( | ENSMUSG( | 0.725206 | 0.471631 | NO | NO | NA |
| Scn1b | ENSMUSG( | ENSMUSG( | -0.15947 | 0.789216 | NO | NO | NA |
| Kcnv1 | ENSMUSG( | ENSMUSG( | 0.69569 | 0.555644 | NO | NO | NA |
| Gm43623 | ENSMUSG( | ENSMUSG( | 1.08719 | 0.294876 | NO | NO | NA |
| Vipr2 | ENSMUSG( | ENSMUSG( | 1.787242 | 0.030815 | YES | YES | Upregulated |
| Ntng1 | ENSMUSG( | ENSMUSG( | 0.155416 | 0.913536 | NO | NO | NA |
| Gm6145 | ENSMUSG( | ENSMUSG( | 0.659295 | 0.645129 | NO | NO | NA |
| Gm42769 | ENSMUSG( | ENSMUSG( | 0.395007 | 0.748417 | NO | NO | NA |
| Olfr670 | ENSMUSG( | ENSMUSG( | 1.399858 | 0.308111 | NO | NO | NA |
| Gm9860 | ENSMUSG( | ENSMUSG( | 0.436401 | 0.618922 | NO | NO | NA |
| ENSMUSG | ENSMUSG( | ENSMUSG( | 1.028007 | 0.021356 | YES | YES | Upregulated |
| Gm7308 | ENSMUSG( | ENSMUSG( | 0.365479 | 0.631048 | NO | NO | NA |
| Grb14 | ENSMUSG( | ENSMUSG( | 0.585256 | 0.327368 | NO | NO | NA |
| Gm2582 | ENSMUSG( | ENSMUSG( | 0.248453 | 0.821783 | NO | NO | NA |
| Olfr259 | ENSMUSG( | ENSMUSG( | 0.930137 | 0.427576 | NO | NO | NA |
| Gm14125 | ENSMUSG( | ENSMUSG( | -2.33974 | 0.000503 | YES | YES | Downregulated |
| Pygo1 | ENSMUSG( | ENSMUSG( | 0.348058 | 0.745508 | NO | NO | NA |
| Gm6176 | ENSMUSG( | ENSMUSG( | -0.53259 | 0.786395 | NO | NO | NA |
| Gm43702 | ENSMUSG( | ENSMUSG( | -0.61316 | 0.315819 | NO | NO | NA |
| Cdc20b | ENSMUSG( | ENSMUSG( | -0.78217 | 0.127557 | NO | NO | NA |
| Gtsf1 | ENSMUSG( | ENSMUSG( | 0.621038 | 0.326678 | NO | NO | NA |
| Gm5171 | ENSMUSG( | ENSMUSG( | 0.140464 | 0.715963 | NO | NO | NA |
| Gm37954 | ENSMUSG( | ENSMUSG( | -0.32398 | 0.71682 | NO | NO | NA |
| ENSMUSG( | ENSMUSG( | ENSMUSG( | 0.318323 | 0.76755 | NO | NO | NA |
| Alox5ap | ENSMUSG( | ENSMUSG( | -0.19026 | 0.779546 | NO | NO | NA |
| Gm34471 | ENSMUSG( | ENSMUSG( | 0.734382 | 0.544725 | NO | NO | NA |
| H2ac5-ps | ENSMUSG( | ENSMUSG( | 0.165551 | 0.786484 | NO | NO | NA |
| Gm15270 | ENSMUSG( | ENSMUSG( | 0.417527 | 0.673641 | NO | NO | NA |
| Pacsin3 | ENSMUSG( | ENSMUSG( | 0.377914 | 0.315494 | NO | NO | NA |
| Svop | ENSMUSG( | ENSMUSG( | 0.270607 | 0.567401 | NO | NO | NA |
| Gm48262 | ENSMUSG( | ENSMUSG( | 0.446301 | 0.685613 | NO | NO | NA |
| Olfr1349 | ENSMUSG( | ENSMUSG( | 0.780708 | 0.544848 | NO | NO | NA |
| Gm36976 | ENSMUSG( | ENSMUSG( | -0.43034 | 0.338389 | NO | NO | NA |

| | | | | | | | |
|---|---|---|---|---|---|---|---|
| Muc13 | ENSMUSG( | ENSMUSG( | 1.045834 | 0.215946 | NO | NO | NA |
| Rxfp2 | ENSMUSG( | ENSMUSG( | 0.264481 | 0.814028 | NO | NO | NA |
| Gm37843 | ENSMUSG( | ENSMUSG( | 0.278007 | 0.797406 | NO | NO | NA |
| Gm49417 | ENSMUSG( | ENSMUSG( | -0.76517 | 0.09552 | NO | NO | NA |
| Gm42872 | ENSMUSG( | ENSMUSG( | -1.08787 | 0.004288 | YES | YES | Downregulated |
| Ddx25 | ENSMUSG( | ENSMUSG( | 0.815278 | 0.114903 | NO | NO | NA |
| Lce1i | ENSMUSG( | ENSMUSG( | 0.229588 | 0.835599 | NO | NO | NA |
| A530030E: | ENSMUSG( | ENSMUSG( | 1.299349 | 0.254163 | NO | NO | NA |
| Gm42892 | ENSMUSG( | ENSMUSG( | 0.345303 | 0.737206 | NO | NO | NA |
| Tmem102 | ENSMUSG( | ENSMUSG( | 0.363843 | 0.732954 | NO | NO | NA |
| Marchf10 | ENSMUSG( | ENSMUSG( | 1.593821 | 0.137858 | NO | NO | NA |
| Gm49692 | ENSMUSG( | ENSMUSG( | 0.947592 | 0.389693 | NO | NO | NA |
| ENSMUSG | ENSMUSG( | ENSMUSG( | 0.280848 | 0.74342 | NO | NO | NA |
| ENSMUSG | ENSMUSG( | ENSMUSG( | 0.787874 | 0.114822 | NO | NO | NA |
| Srgap1 | ENSMUSG( | ENSMUSG( | 0.499234 | 0.618635 | NO | NO | NA |
| Zfp558 | ENSMUSG( | ENSMUSG( | -0.28547 | 0.523632 | NO | NO | NA |
| Osmr | ENSMUSG( | ENSMUSG( | 0.425074 | 0.528002 | NO | NO | NA |
| Fam83f | ENSMUSG( | ENSMUSG( | -0.32286 | 0.756952 | NO | NO | NA |
| 8430406P: | ENSMUSG( | ENSMUSG( | -0.21473 | 0.679591 | NO | NO | NA |
| Gm36501 | ENSMUSG( | ENSMUSG( | -1.30343 | 0.00058 | YES | YES | Downregulated |
| 3010001F: | ENSMUSG( | ENSMUSG( | 0.712299 | 0.475513 | NO | NO | NA |
| Dkk4 | ENSMUSG( | ENSMUSG( | 0.557512 | 0.643278 | NO | NO | NA |
| Gm20274 | ENSMUSG( | ENSMUSG( | -0.16834 | 0.732625 | NO | NO | NA |
| St8sia2 | ENSMUSG( | ENSMUSG( | 0.713267 | 0.51145 | NO | NO | NA |
| Gm44740 | ENSMUSG( | ENSMUSG( | 0.337726 | 0.765903 | NO | NO | NA |
| Gm50353 | ENSMUSG( | ENSMUSG( | 0.576377 | 0.664834 | NO | NO | NA |
| Hoxb3os | ENSMUSG( | ENSMUSG( | 0.593672 | 0.621561 | NO | NO | NA |
| Vdac3-ps1 | ENSMUSG( | ENSMUSG( | 0.18793 | 0.722103 | NO | NO | NA |
| Olfr103 | ENSMUSG( | ENSMUSG( | 1.343142 | 0.216767 | NO | NO | NA |
| Olfr45 | ENSMUSG( | ENSMUSG( | 0.920563 | 0.278087 | NO | NO | NA |
| Olfr102 | ENSMUSG( | ENSMUSG( | 0.384197 | 0.680133 | NO | NO | NA |
| ENSMUSG | ENSMUSG( | ENSMUSG( | 0.517251 | 0.579897 | NO | NO | NA |
| Amotl2 | ENSMUSG( | ENSMUSG( | 0.845557 | 0.124515 | NO | NO | NA |
| A330015K( | ENSMUSG( | ENSMUSG( | 1.464866 | 0.121941 | NO | NO | NA |
| Gm36827 | ENSMUSG( | ENSMUSG( | 0.450944 | 0.695793 | NO | NO | NA |
| Gm6316 | ENSMUSG( | ENSMUSG( | 0.449677 | 0.610654 | NO | NO | NA |
| 5430430K: | ENSMUSG( | ENSMUSG( | 0.376343 | 0.705305 | NO | NO | NA |
| Prxl2b | ENSMUSG( | ENSMUSG( | -0.13276 | 0.734883 | NO | NO | NA |
| Gm49559 | ENSMUSG( | ENSMUSG( | 0.893597 | 0.437329 | NO | NO | NA |
| Rhou | ENSMUSG( | ENSMUSG( | 0.790722 | 0.371199 | NO | NO | NA |
| Gm44697 | ENSMUSG( | ENSMUSG( | 0.16056 | 0.869306 | NO | NO | NA |
| Gm50270 | ENSMUSG( | ENSMUSG( | -0.21063 | 0.744963 | NO | NO | NA |
| Gm37864 | ENSMUSG( | ENSMUSG( | 0.636082 | 0.529014 | NO | NO | NA |
| Gm5106 | ENSMUSG( | ENSMUSG( | 0.344936 | 0.397767 | NO | NO | NA |
| Gm37789 | ENSMUSG( | ENSMUSG( | 0.56654 | 0.548121 | NO | NO | NA |
| Gls2 | ENSMUSG( | ENSMUSG( | 1.174345 | 0.048698 | YES | YES | Upregulated |
| Olfr22-ps1 | ENSMUSG( | ENSMUSG( | 0.65095 | 0.538424 | NO | NO | NA |

| | | | | | | | |
|---|---|---|---|---|---|---|---|
| 3110070M | ENSMUSG( | ENSMUSG( | 0.2309 | 0.586824 | NO | NO | NA |
| Gm13487 | ENSMUSG( | ENSMUSG( | 0.250872 | 0.784646 | NO | NO | NA |
| Gm44284 | ENSMUSG( | ENSMUSG( | -0.01816 | 0.980821 | NO | NO | NA |
| Rnf183 | ENSMUSG( | ENSMUSG( | 0.207291 | 0.792835 | NO | NO | NA |
| 4930526F1 | ENSMUSG( | ENSMUSG( | 0.38361 | 0.731148 | NO | NO | NA |
| Stk36 | ENSMUSG( | ENSMUSG( | -0.42263 | 0.548707 | NO | NO | NA |
| Dusp15 | ENSMUSG( | ENSMUSG( | 0.67383 | 0.548283 | NO | NO | NA |
| Gm44584 | ENSMUSG( | ENSMUSG( | 0.725247 | 0.574591 | NO | NO | NA |
| Ccdc162 | ENSMUSG( | ENSMUSG( | -0.13361 | 0.799673 | NO | NO | NA |
| Gm36839 | ENSMUSG( | ENSMUSG( | 0.376877 | 0.638353 | NO | NO | NA |
| Scg3 | ENSMUSG( | ENSMUSG( | 0.706318 | 0.496053 | NO | NO | NA |
| Syndig1l | ENSMUSG( | ENSMUSG( | 0.006193 | 0.995759 | NO | NO | NA |
| Gm8670 | ENSMUSG( | ENSMUSG( | 1.085495 | 0.096382 | NO | NO | NA |
| Gm32885 | ENSMUSG( | ENSMUSG( | 0.583519 | 0.502992 | NO | NO | NA |
| Irak3 | ENSMUSG( | ENSMUSG( | 0.081157 | 0.878139 | NO | NO | NA |
| Olfm3 | ENSMUSG( | ENSMUSG( | 1.856349 | 0.174565 | NO | NO | NA |
| Gm16385 | ENSMUSG( | ENSMUSG( | 0.183125 | 0.722171 | NO | NO | NA |
| Gm12341 | ENSMUSG( | ENSMUSG( | 0.382044 | 0.677181 | NO | NO | NA |
| Gm12167 | ENSMUSG( | ENSMUSG( | 0.194372 | 0.822413 | NO | NO | NA |
| Gm45486 | ENSMUSG( | ENSMUSG( | 1.510464 | 0.096274 | NO | NO | NA |
| Igkv4-72 | ENSMUSG( | ENSMUSG( | 1.036961 | 0.676301 | NO | NO | NA |
| Unc80 | ENSMUSG( | ENSMUSG( | 0.535502 | 0.559139 | NO | NO | NA |
| 4732465J0 | ENSMUSG( | ENSMUSG( | 0.997127 | 0.394006 | NO | NO | NA |
| Adamtsl5 | ENSMUSG( | ENSMUSG( | 1.395089 | 0.01124 | YES | YES | Upregulated |
| Gm26839 | ENSMUSG( | ENSMUSG( | -0.11903 | 0.75647 | NO | NO | NA |
| C230066G | ENSMUSG( | ENSMUSG( | -0.50489 | 0.261815 | NO | NO | NA |
| Olfr1247 | ENSMUSG( | ENSMUSG( | 0.846805 | 0.359472 | NO | NO | NA |
| Zfp872 | ENSMUSG( | ENSMUSG( | 1.184585 | 0.291309 | NO | NO | NA |
| Gm8494 | ENSMUSG( | ENSMUSG( | 0.298232 | 0.572981 | NO | NO | NA |
| Slc25a34 | ENSMUSG( | ENSMUSG( | -0.92853 | 0.280151 | NO | NO | NA |
| Gm26558 | ENSMUSG( | ENSMUSG( | 0.601891 | 0.203991 | NO | NO | NA |
| Gm31182 | ENSMUSG( | ENSMUSG( | 1.006596 | 0.407858 | NO | NO | NA |
| Gm13055 | ENSMUSG( | ENSMUSG( | 0.188354 | 0.890997 | NO | NO | NA |
| Phospho1 | ENSMUSG( | ENSMUSG( | 0.982348 | 0.37124 | NO | NO | NA |
| Pirb | ENSMUSG( | ENSMUSG( | -1.02762 | 0.105772 | NO | NO | NA |
| Gm49503 | ENSMUSG( | ENSMUSG( | 0.769682 | 0.208423 | NO | NO | NA |
| 1700034K( | ENSMUSG( | ENSMUSG( | 1.158669 | 0.448011 | NO | NO | NA |
| Gm47016 | ENSMUSG( | ENSMUSG( | 0.717913 | 0.582375 | NO | NO | NA |
| Gm37720 | ENSMUSG( | ENSMUSG( | 0.920807 | 0.373789 | NO | NO | NA |
| C030017G | ENSMUSG( | ENSMUSG( | -0.24704 | 0.623747 | NO | NO | NA |
| Gm2885 | ENSMUSG( | ENSMUSG( | 0.302034 | 0.596711 | NO | NO | NA |
| 1700047G | ENSMUSG( | ENSMUSG( | 0.457094 | 0.485081 | NO | NO | NA |
| Gm50142 | ENSMUSG( | ENSMUSG( | 0.548202 | 0.646764 | NO | NO | NA |
| Gm11525 | ENSMUSG( | ENSMUSG( | 0.594561 | 0.348108 | NO | NO | NA |
| Gm10305 | ENSMUSG( | ENSMUSG( | 1.065939 | 0.321399 | NO | NO | NA |
| Gm44283 | ENSMUSG( | ENSMUSG( | 0.799449 | 0.416423 | NO | NO | NA |
| Gm19142 | ENSMUSG( | ENSMUSG( | -0.28433 | 0.665838 | NO | NO | NA |

| | | | | | | | |
|---|---|---|---|---|---|---|---|
| ENSMUSG | ENSMUSG( | ENSMUSG( | 0.703507 | 0.581594 | NO | NO | NA |
| F830210D( | ENSMUSG( | ENSMUSG( | 0.886195 | 0.401746 | NO | NO | NA |
| Fzd1 | ENSMUSG( | ENSMUSG( | 1.396641 | 0.117245 | NO | NO | NA |
| Gm36745 | ENSMUSG( | ENSMUSG( | 0.440332 | 0.673911 | NO | NO | NA |
| Gm28877 | ENSMUSG( | ENSMUSG( | 0.975745 | 0.474814 | NO | NO | NA |
| Msx1 | ENSMUSG( | ENSMUSG( | 0.989428 | 0.211633 | NO | NO | NA |
| Gm50256 | ENSMUSG( | ENSMUSG( | -0.37176 | 0.455689 | NO | NO | NA |
| Gm5822 | ENSMUSG( | ENSMUSG( | 0.219299 | 0.645355 | NO | NO | NA |
| Gm44790 | ENSMUSG( | ENSMUSG( | 0.694153 | 0.545957 | NO | NO | NA |
| Gm35507 | ENSMUSG( | ENSMUSG( | 0.223995 | 0.836247 | NO | NO | NA |
| Gm47573 | ENSMUSG( | ENSMUSG( | 0.481065 | 0.610654 | NO | NO | NA |
| D830035M | ENSMUSG( | ENSMUSG( | 0.455894 | 0.391549 | NO | NO | NA |
| Gm38188 | ENSMUSG( | ENSMUSG( | 0.227502 | 0.624516 | NO | NO | NA |
| Olfr318 | ENSMUSG( | ENSMUSG( | 0.143527 | 0.92009 | NO | NO | NA |
| Gm49273 | ENSMUSG( | ENSMUSG( | -0.92134 | 0.113545 | NO | NO | NA |
| Gm48405 | ENSMUSG( | ENSMUSG( | -0.01529 | 0.981351 | NO | NO | NA |
| Hdac11 | ENSMUSG( | ENSMUSG( | -0.61294 | 0.232443 | NO | NO | NA |
| Gm42515 | ENSMUSG( | ENSMUSG( | 0.600925 | 0.516453 | NO | NO | NA |
| Tmprss11g | ENSMUSG( | ENSMUSG( | 0.691785 | 0.520755 | NO | NO | NA |
| Col26a1 | ENSMUSG( | ENSMUSG( | 0.259196 | 0.786071 | NO | NO | NA |
| Gata6 | ENSMUSG( | ENSMUSG( | 0.749363 | 0.420525 | NO | NO | NA |
| Gm14667 | ENSMUSG( | ENSMUSG( | 1.000698 | 0.380197 | NO | NO | NA |
| BC055308 | ENSMUSG( | ENSMUSG( | -0.22422 | 0.672948 | NO | NO | NA |
| Tcam1 | ENSMUSG( | ENSMUSG( | 0.257248 | 0.665838 | NO | NO | NA |
| Ggn | ENSMUSG( | ENSMUSG( | 0.385413 | 0.522983 | NO | NO | NA |
| Cbln3 | ENSMUSG( | ENSMUSG( | 0.087846 | 0.828824 | NO | NO | NA |
| Gm19500 | ENSMUSG( | ENSMUSG( | 0.524838 | 0.623864 | NO | NO | NA |
| Slc27a2 | ENSMUSG( | ENSMUSG( | 0.450819 | 0.692551 | NO | NO | NA |
| Aif1 | ENSMUSG( | ENSMUSG( | -0.55777 | 0.199049 | NO | NO | NA |
| Gm15515 | ENSMUSG( | ENSMUSG( | 0.779171 | 0.473973 | NO | NO | NA |
| Kcna1 | ENSMUSG( | ENSMUSG( | 0.3284 | 0.750846 | NO | NO | NA |
| Axdnd1 | ENSMUSG( | ENSMUSG( | -0.14578 | 0.748417 | NO | NO | NA |
| Gm43289 | ENSMUSG( | ENSMUSG( | 1.029005 | 0.329411 | NO | NO | NA |
| 2810029C( | ENSMUSG( | ENSMUSG( | 0.572849 | 0.486893 | NO | NO | NA |
| A530076I1 | ENSMUSG( | ENSMUSG( | 0.572497 | 0.609502 | NO | NO | NA |
| Gm42858 | ENSMUSG( | ENSMUSG( | 0.400833 | 0.554806 | NO | NO | NA |
| ENSMUSG | ENSMUSG( | ENSMUSG( | 1.606235 | 0.054068 | NO | NO | NA |
| Siglech | ENSMUSG( | ENSMUSG( | 1.368752 | 0.198188 | NO | NO | NA |
| Gm7895 | ENSMUSG( | ENSMUSG( | -0.58252 | 0.251928 | NO | NO | NA |
| Gm10322 | ENSMUSG( | ENSMUSG( | -0.59931 | 0.544325 | NO | NO | NA |
| Clec2l | ENSMUSG( | ENSMUSG( | 0.013355 | 0.980226 | NO | NO | NA |
| Gm4513 | ENSMUSG( | ENSMUSG( | -1.34802 | 0.729736 | NO | NO | NA |
| Ighv1-18 | ENSMUSG( | ENSMUSG( | -3.13971 | 0.117484 | NO | NO | NA |
| Kmo | ENSMUSG( | ENSMUSG( | -0.34373 | 0.487953 | NO | NO | NA |
| Slc24a4 | ENSMUSG( | ENSMUSG( | 0.516952 | 0.680822 | NO | NO | NA |
| Gm37215 | ENSMUSG( | ENSMUSG( | 0.608081 | 0.454063 | NO | NO | NA |
| Gm47031 | ENSMUSG( | ENSMUSG( | 0.70063 | 0.555644 | NO | NO | NA |

| | | | | | | | |
|---|---|---|---|---|---|---|---|
| Tcea1-ps1 | ENSMUSG( | ENSMUSG( | 0.045885 | 0.91013 | NO | NO | NA |
| Cd163 | ENSMUSG( | ENSMUSG( | 0.368379 | 0.495374 | NO | NO | NA |
| Wnt3a | ENSMUSG( | ENSMUSG( | 0.911694 | 0.370629 | NO | NO | NA |
| Gng7 | ENSMUSG( | ENSMUSG( | 1.386265 | 0.301062 | NO | NO | NA |
| Gm11411 | ENSMUSG( | ENSMUSG( | 0.882082 | 0.459936 | NO | NO | NA |
| Gm13292 | ENSMUSG( | ENSMUSG( | 2.061399 | 0.035396 | YES | YES | Upregulated |
| Myo15b | ENSMUSG( | ENSMUSG( | 0.124841 | 0.847174 | NO | NO | NA |
| Gm10501 | ENSMUSG( | ENSMUSG( | 0.379551 | 0.601693 | NO | NO | NA |
| A330032P. | ENSMUSG( | ENSMUSG( | 1.909566 | 0.07332 | NO | NO | NA |
| Morc1 | ENSMUSG( | ENSMUSG( | -2.04693 | 0.001836 | YES | YES | Downregulated |
| Gm48393 | ENSMUSG( | ENSMUSG( | 1.138142 | 0.360125 | NO | NO | NA |
| Gm48562 | ENSMUSG( | ENSMUSG( | 0.155085 | 0.767349 | NO | NO | NA |
| Gm43054 | ENSMUSG( | ENSMUSG( | 0.028959 | 0.956656 | NO | NO | NA |
| Olfr270 | ENSMUSG( | ENSMUSG( | 0.702001 | 0.591085 | NO | NO | NA |
| Gm37242 | ENSMUSG( | ENSMUSG( | 0.58823 | 0.260276 | NO | NO | NA |
| Gm48125 | ENSMUSG( | ENSMUSG( | 0.860614 | 0.541419 | NO | NO | NA |
| Olfr748 | ENSMUSG( | ENSMUSG( | 0.545861 | 0.609095 | NO | NO | NA |
| Olfr904 | ENSMUSG( | ENSMUSG( | 0.60849 | 0.538948 | NO | NO | NA |
| Gm14797 | ENSMUSG( | ENSMUSG( | 1.278721 | 0.306821 | NO | NO | NA |
| Tead2 | ENSMUSG( | ENSMUSG( | -0.22 | 0.544012 | NO | NO | NA |
| Ms4a8a | ENSMUSG( | ENSMUSG( | 0.501387 | 0.576362 | NO | NO | NA |
| Gm16105 | ENSMUSG( | ENSMUSG( | 0.750876 | 0.444377 | NO | NO | NA |
| Gm44695 | ENSMUSG( | ENSMUSG( | 0.608307 | 0.540187 | NO | NO | NA |
| Gm44770 | ENSMUSG( | ENSMUSG( | 1.100937 | 0.369 | NO | NO | NA |
| Gm16174 | ENSMUSG( | ENSMUSG( | 0.409427 | 0.408656 | NO | NO | NA |
| ENSMUSG | ENSMUSG( | ENSMUSG( | 1.959563 | 0.09551 | NO | NO | NA |
| Gm47522 | ENSMUSG( | ENSMUSG( | 0.832429 | 0.490176 | NO | NO | NA |
| Gm12002 | ENSMUSG( | ENSMUSG( | 0.841145 | 0.465954 | NO | NO | NA |
| Gm21043 | ENSMUSG( | ENSMUSG( | 1.13763 | 0.356245 | NO | NO | NA |
| Gm15387 | ENSMUSG( | ENSMUSG( | -5.59285 | 0.127842 | NO | NO | NA |
| Dpp6 | ENSMUSG( | ENSMUSG( | 0.214988 | 0.850641 | NO | NO | NA |
| 4921513H | ENSMUSG( | ENSMUSG( | 0.374012 | 0.756346 | NO | NO | NA |
| Syde1 | ENSMUSG( | ENSMUSG( | 1.020802 | 0.015555 | YES | YES | Upregulated |
| Gm42869 | ENSMUSG( | ENSMUSG( | -0.20502 | 0.65276 | NO | NO | NA |
| Gm13479 | ENSMUSG( | ENSMUSG( | 0.46201 | 0.757003 | NO | NO | NA |
| Iglv2 | ENSMUSG( | ENSMUSG( | -3.92028 | 0.109974 | NO | NO | NA |
| ENSMUSG( | ENSMUSG( | ENSMUSG( | 0.05756 | 0.959325 | NO | NO | NA |
| 4833412C: | ENSMUSG( | ENSMUSG( | 1.003307 | 0.354826 | NO | NO | NA |
| ENSMUSG( | ENSMUSG( | ENSMUSG( | 0.817139 | 0.582781 | NO | NO | NA |
| Habp2 | ENSMUSG( | ENSMUSG( | 0.483605 | 0.693448 | NO | NO | NA |
| Brd8dc | ENSMUSG( | ENSMUSG( | 0.217454 | 0.740776 | NO | NO | NA |
| Hao1 | ENSMUSG( | ENSMUSG( | 0.885266 | 0.411247 | NO | NO | NA |
| Sema3d | ENSMUSG( | ENSMUSG( | 1.476315 | 0.198985 | NO | NO | NA |
| Gm42772 | ENSMUSG( | ENSMUSG( | 1.111695 | 0.411011 | NO | NO | NA |
| Gm14679 | ENSMUSG( | ENSMUSG( | 0.939265 | 0.392349 | NO | NO | NA |
| ENSMUSG( | ENSMUSG( | ENSMUSG( | 1.202669 | 0.244364 | NO | NO | NA |
| Gm10662 | ENSMUSG( | ENSMUSG( | 1.301449 | 0.613046 | NO | NO | NA |

| | | | | | | | |
|---|---|---|---|---|---|---|---|
| Gm6192 | ENSMUSG( | ENSMUSG( | 0.208235 | 0.729544 | NO | NO | NA |
| Gm20400 | ENSMUSG( | ENSMUSG( | 0.310521 | 0.511409 | NO | NO | NA |
| Gm16741 | ENSMUSG( | ENSMUSG( | 0.535131 | 0.381703 | NO | NO | NA |
| Gm10705 | ENSMUSG( | ENSMUSG( | 0.371575 | 0.606212 | NO | NO | NA |
| Gm11948 | ENSMUSG( | ENSMUSG( | -0.42075 | 0.319978 | NO | NO | NA |
| F930017D: | ENSMUSG( | ENSMUSG( | 0.851805 | 0.244364 | NO | NO | NA |
| 8030445P: | ENSMUSG( | ENSMUSG( | 0.965135 | 0.417268 | NO | NO | NA |
| Gm6198 | ENSMUSG( | ENSMUSG( | 1.734891 | 0.040221 | YES | YES | Upregulated |
| Gm49004 | ENSMUSG( | ENSMUSG( | -0.22837 | 0.731673 | NO | NO | NA |
| Olfr753-ps | ENSMUSG( | ENSMUSG( | 0.69767 | 0.600072 | NO | NO | NA |
| Gm27017 | ENSMUSG( | ENSMUSG( | 0.242402 | 0.729187 | NO | NO | NA |
| ENSMUSG | ENSMUSG( | ENSMUSG( | 0.613769 | 0.626357 | NO | NO | NA |
| Siglec1 | ENSMUSG( | ENSMUSG( | -0.47105 | 0.375234 | NO | NO | NA |
| Neurod1 | ENSMUSG( | ENSMUSG( | 1.064895 | 0.107184 | NO | NO | NA |
| Platr16 | ENSMUSG( | ENSMUSG( | 2.189526 | 0.126951 | NO | NO | NA |
| A430027C | ENSMUSG( | ENSMUSG( | 0.215215 | 0.661476 | NO | NO | NA |
| Bmp3 | ENSMUSG( | ENSMUSG( | -0.2195 | 0.841744 | NO | NO | NA |
| Gm44957 | ENSMUSG( | ENSMUSG( | 1.009273 | 0.347649 | NO | NO | NA |
| ENSMUSG | ENSMUSG( | ENSMUSG( | 0.412816 | 0.711169 | NO | NO | NA |
| Gm43721 | ENSMUSG( | ENSMUSG( | -0.35393 | 0.593768 | NO | NO | NA |
| Gm49863 | ENSMUSG( | ENSMUSG( | -0.02399 | 0.956989 | NO | NO | NA |
| Ptprb | ENSMUSG( | ENSMUSG( | 0.851838 | 0.398656 | NO | NO | NA |
| Vmn1r6 | ENSMUSG( | ENSMUSG( | 0.603978 | 0.547964 | NO | NO | NA |
| Gm12366 | ENSMUSG( | ENSMUSG( | -0.25894 | 0.736727 | NO | NO | NA |
| ENSMUSG | ENSMUSG( | ENSMUSG( | 0.133522 | 0.911939 | NO | NO | NA |
| Tpt1-ps6 | ENSMUSG( | ENSMUSG( | 0.164365 | 0.808851 | NO | NO | NA |
| Gm11743 | ENSMUSG( | ENSMUSG( | -0.85482 | 0.294095 | NO | NO | NA |
| Gm47982 | ENSMUSG( | ENSMUSG( | 0.770054 | 0.532755 | NO | NO | NA |
| Gm16222 | ENSMUSG( | ENSMUSG( | 1.169255 | 0.2874 | NO | NO | NA |
| Rab27b | ENSMUSG( | ENSMUSG( | 1.035614 | 0.306811 | NO | NO | NA |
| Spx | ENSMUSG( | ENSMUSG( | -0.37893 | 0.460439 | NO | NO | NA |
| Hoxc6 | ENSMUSG( | ENSMUSG( | 0.688384 | 0.518069 | NO | NO | NA |
| Gm43314 | ENSMUSG( | ENSMUSG( | 0.124879 | 0.911939 | NO | NO | NA |
| 2610017A | ENSMUSG( | ENSMUSG( | -0.64188 | 0.196143 | NO | NO | NA |
| Gm49194 | ENSMUSG( | ENSMUSG( | -0.21634 | 0.849434 | NO | NO | NA |
| Slc6a15 | ENSMUSG( | ENSMUSG( | 0.43547 | 0.703614 | NO | NO | NA |
| 4932438H | ENSMUSG( | ENSMUSG( | 0.561974 | 0.626728 | NO | NO | NA |
| Gm37120 | ENSMUSG( | ENSMUSG( | 0.412989 | 0.730207 | NO | NO | NA |
| Gm46218 | ENSMUSG( | ENSMUSG( | -0.35671 | 0.62299 | NO | NO | NA |
| Gm26600 | ENSMUSG( | ENSMUSG( | 0.245967 | 0.829321 | NO | NO | NA |
| Gm18515 | ENSMUSG( | ENSMUSG( | 1.034333 | 0.389245 | NO | NO | NA |
| Gm48559 | ENSMUSG( | ENSMUSG( | 0.006052 | 0.990219 | NO | NO | NA |
| Olfr459 | ENSMUSG( | ENSMUSG( | 1.50951 | 0.170586 | NO | NO | NA |
| Gm10575 | ENSMUSG( | ENSMUSG( | -0.36852 | 0.529107 | NO | NO | NA |
| Olfr862 | ENSMUSG( | ENSMUSG( | 0.560731 | 0.651735 | NO | NO | NA |
| Il4i1 | ENSMUSG( | ENSMUSG( | 0.006457 | 0.993356 | NO | NO | NA |
| 2210039B | ENSMUSG( | ENSMUSG( | 0.33022 | 0.764031 | NO | NO | NA |

| | | | | | | | |
|---|---|---|---|---|---|---|---|
| Fgf8 | ENSMUSG( | ENSMUSG( | 1.202697 | 0.300618 | NO | NO | NA |
| ENSMUSG | ENSMUSG( | ENSMUSG( | 1.326442 | 0.242752 | NO | NO | NA |
| Foxg1 | ENSMUSG( | ENSMUSG( | 1.116653 | 0.298442 | NO | NO | NA |
| Gm28424 | ENSMUSG( | ENSMUSG( | 0.464556 | 0.275655 | NO | NO | NA |
| Csgalnact1 | ENSMUSG( | ENSMUSG( | 0.137494 | 0.85316 | NO | NO | NA |
| ENSMUSG | ENSMUSG( | ENSMUSG( | 0.304769 | 0.540932 | NO | NO | NA |
| Gm37509 | ENSMUSG( | ENSMUSG( | -0.55593 | 0.198996 | NO | NO | NA |
| Gm10516 | ENSMUSG( | ENSMUSG( | 0.533757 | 0.404239 | NO | NO | NA |
| Pdzd7 | ENSMUSG( | ENSMUSG( | 0.421129 | 0.569784 | NO | NO | NA |
| Gm37226 | ENSMUSG( | ENSMUSG( | -1.21246 | 0.030409 | YES | YES | Downregulated |
| Gm36529 | ENSMUSG( | ENSMUSG( | 0.523363 | 0.656736 | NO | NO | NA |
| Iqck | ENSMUSG( | ENSMUSG( | 0.918145 | 0.218214 | NO | NO | NA |
| Gm14705 | ENSMUSG( | ENSMUSG( | 1.620262 | 0.164408 | NO | NO | NA |
| Gm14648 | ENSMUSG( | ENSMUSG( | -0.03413 | 0.968865 | NO | NO | NA |
| Gm10241 | ENSMUSG( | ENSMUSG( | -0.16386 | 0.772675 | NO | NO | NA |
| Gm18094 | ENSMUSG( | ENSMUSG( | 0.987673 | 0.455732 | NO | NO | NA |
| Gm4779 | ENSMUSG( | ENSMUSG( | 5.329314 | NA | NO | NO | NA |
| Tdgf1 | ENSMUSG( | ENSMUSG( | 0.356123 | 0.425984 | NO | NO | NA |
| Olfr355 | ENSMUSG( | ENSMUSG( | 0.518494 | 0.662767 | NO | NO | NA |
| Gm43024 | ENSMUSG( | ENSMUSG( | 0.279333 | 0.661527 | NO | NO | NA |
| Art4 | ENSMUSG( | ENSMUSG( | 0.758859 | 0.397767 | NO | NO | NA |
| ENSMUSG( | ENSMUSG( | ENSMUSG( | 0.902305 | 0.326281 | NO | NO | NA |
| ENSMUSG | ENSMUSG( | ENSMUSG( | 1.032258 | 0.243074 | NO | NO | NA |
| Gm28044 | ENSMUSG( | ENSMUSG( | -1.31931 | 0.223803 | NO | NO | NA |
| E130310I0 | ENSMUSG( | ENSMUSG( | 0.669792 | 0.572224 | NO | NO | NA |
| Ldb3 | ENSMUSG( | ENSMUSG( | 0.564222 | 0.589914 | NO | NO | NA |
| Gm19113 | ENSMUSG( | ENSMUSG( | 0.940267 | 0.426939 | NO | NO | NA |
| Gm32171 | ENSMUSG( | ENSMUSG( | 1.215977 | 0.280897 | NO | NO | NA |
| Kctd8 | ENSMUSG( | ENSMUSG( | 1.934325 | 0.132447 | NO | NO | NA |
| Gm48771 | ENSMUSG( | ENSMUSG( | 0.295559 | 0.535272 | NO | NO | NA |
| Gm6027 | ENSMUSG( | ENSMUSG( | 0.699595 | 0.548076 | NO | NO | NA |
| Cpa2 | ENSMUSG( | ENSMUSG( | 1.591469 | 0.013972 | YES | YES | Upregulated |
| Gm20501 | ENSMUSG( | ENSMUSG( | 0.266643 | 0.791514 | NO | NO | NA |
| Olfr551 | ENSMUSG( | ENSMUSG( | 1.614271 | 0.116191 | NO | NO | NA |
| Ccdc85a | ENSMUSG( | ENSMUSG( | 0.555837 | 0.587507 | NO | NO | NA |
| Olfr1262 | ENSMUSG( | ENSMUSG( | 0.141311 | 0.893131 | NO | NO | NA |
| Gm48720 | ENSMUSG( | ENSMUSG( | -0.04969 | 0.90247 | NO | NO | NA |
| Rgr | ENSMUSG( | ENSMUSG( | 0.078283 | 0.948074 | NO | NO | NA |
| ENSMUSG( | ENSMUSG( | ENSMUSG( | -1.02137 | 0.388427 | NO | NO | NA |
| Ppp1r3b | ENSMUSG( | ENSMUSG( | -0.35569 | 0.471129 | NO | NO | NA |
| Gm42602 | ENSMUSG( | ENSMUSG( | 0.309649 | 0.74816 | NO | NO | NA |
| Gm43210 | ENSMUSG( | ENSMUSG( | 0.442473 | 0.736696 | NO | NO | NA |
| Gm48159 | ENSMUSG( | ENSMUSG( | 0.240589 | 0.814086 | NO | NO | NA |
| Stox1 | ENSMUSG( | ENSMUSG( | 0.48934 | 0.382461 | NO | NO | NA |
| Megf8 | ENSMUSG( | ENSMUSG( | 0.159386 | 0.83236 | NO | NO | NA |
| Gm37323 | ENSMUSG( | ENSMUSG( | 0.369159 | 0.729989 | NO | NO | NA |
| Gm45758 | ENSMUSG( | ENSMUSG( | 0.434227 | 0.374748 | NO | NO | NA |

| | | | | | | | |
|---|---|---|---|---|---|---|---|
| Gm29560 | ENSMUSG( | ENSMUSG( | 0.789373 | 0.461523 | NO | NO | NA |
| Olfr1009 | ENSMUSG( | ENSMUSG( | 0.831226 | 0.560273 | NO | NO | NA |
| Gm12811 | ENSMUSG( | ENSMUSG( | 0.553526 | 0.732555 | NO | NO | NA |
| Gm15968 | ENSMUSG( | ENSMUSG( | 0.724739 | 0.508487 | NO | NO | NA |
| D330023K | ENSMUSG( | ENSMUSG( | 0.392345 | 0.440916 | NO | NO | NA |
| Ephx3 | ENSMUSG( | ENSMUSG( | -2.09569 | 0.00196 | YES | YES | Downregulated |
| S1pr3 | ENSMUSG( | ENSMUSG( | 0.331074 | 0.738629 | NO | NO | NA |
| Gm38025 | ENSMUSG( | ENSMUSG( | 0.780662 | 0.555058 | NO | NO | NA |
| Gm44235 | ENSMUSG( | ENSMUSG( | 0.929308 | 0.434524 | NO | NO | NA |
| Cend1 | ENSMUSG( | ENSMUSG( | -0.12773 | 0.871269 | NO | NO | NA |
| Gm42826 | ENSMUSG( | ENSMUSG( | 0.534387 | 0.653576 | NO | NO | NA |
| Rpl7-ps9 | ENSMUSG( | ENSMUSG( | 0.690895 | 0.153294 | NO | NO | NA |
| Gm36128 | ENSMUSG( | ENSMUSG( | 1.045611 | 0.412453 | NO | NO | NA |
| Gm47571 | ENSMUSG( | ENSMUSG( | 1.341423 | 0.242187 | NO | NO | NA |
| Olfr916 | ENSMUSG( | ENSMUSG( | 0.568757 | 0.569763 | NO | NO | NA |
| Gm44200 | ENSMUSG( | ENSMUSG( | -0.36786 | 0.410248 | NO | NO | NA |
| Gm37504 | ENSMUSG( | ENSMUSG( | 1.115609 | 0.361191 | NO | NO | NA |
| Olfr1326-p | ENSMUSG( | ENSMUSG( | 0.843633 | 0.432212 | NO | NO | NA |
| Pdzrn4 | ENSMUSG( | ENSMUSG( | 0.531736 | 0.575279 | NO | NO | NA |
| 1110003F1 | ENSMUSG( | ENSMUSG( | 0.274504 | 0.674285 | NO | NO | NA |
| Gm48375 | ENSMUSG( | ENSMUSG( | -0.23769 | 0.716638 | NO | NO | NA |
| ENSMUSG( | ENSMUSG( | ENSMUSG( | 0.068811 | 0.94245 | NO | NO | NA |
| Cysltr1 | ENSMUSG( | ENSMUSG( | 0.42782 | 0.665917 | NO | NO | NA |
| Gm43103 | ENSMUSG( | ENSMUSG( | 1.172083 | 0.265846 | NO | NO | NA |
| Olfr1157 | ENSMUSG( | ENSMUSG( | 0.862818 | 0.495503 | NO | NO | NA |
| Gpr12 | ENSMUSG( | ENSMUSG( | 1.09847 | 0.311264 | NO | NO | NA |
| ENSMUSG | ENSMUSG( | ENSMUSG( | -0.2049 | 0.718587 | NO | NO | NA |
| Gm49534 | ENSMUSG( | ENSMUSG( | 0.205486 | 0.844515 | NO | NO | NA |
| 3000002C1 | ENSMUSG( | ENSMUSG( | 6.097255 | 0.021647 | YES | YES | Upregulated |
| Gm26583 | ENSMUSG( | ENSMUSG( | -1.44495 | 0.07241 | NO | NO | NA |
| Gm38348 | ENSMUSG( | ENSMUSG( | 0.213935 | 0.835562 | NO | NO | NA |
| Rps2-ps6 | ENSMUSG( | ENSMUSG( | 0.654872 | 0.108783 | NO | NO | NA |
| Adamts4 | ENSMUSG( | ENSMUSG( | 0.252672 | 0.772586 | NO | NO | NA |
| Cd14 | ENSMUSG( | ENSMUSG( | 0.804184 | 0.283258 | NO | NO | NA |
| H2-DMb1 | ENSMUSG( | ENSMUSG( | 0.193988 | 0.74256 | NO | NO | NA |
| Lin7a | ENSMUSG( | ENSMUSG( | 0.752852 | 0.527421 | NO | NO | NA |
| Gm7446 | ENSMUSG( | ENSMUSG( | 0.04147 | 0.96183 | NO | NO | NA |
| ENSMUSG( | ENSMUSG( | ENSMUSG( | 0.2617 | 0.794188 | NO | NO | NA |
| ENSMUSG | ENSMUSG( | ENSMUSG( | -0.5387 | 0.299432 | NO | NO | NA |
| ENSMUSG | ENSMUSG( | ENSMUSG( | -0.25066 | 0.666189 | NO | NO | NA |
| Prom2 | ENSMUSG( | ENSMUSG( | 0.16915 | 0.820802 | NO | NO | NA |
| Nr2e1 | ENSMUSG( | ENSMUSG( | 0.169645 | 0.880153 | NO | NO | NA |
| Gm10390 | ENSMUSG( | ENSMUSG( | -0.42183 | 0.391427 | NO | NO | NA |
| Hr | ENSMUSG( | ENSMUSG( | 0.62429 | 0.108882 | NO | NO | NA |
| Gm10300 | ENSMUSG( | ENSMUSG( | -0.25774 | 0.719466 | NO | NO | NA |
| Gm14427 | ENSMUSG( | ENSMUSG( | 0.30869 | 0.778248 | NO | NO | NA |
| C330024D | ENSMUSG( | ENSMUSG( | 0.9315 | 0.498019 | NO | NO | NA |

| | | | | | | | |
|---|---|---|---|---|---|---|---|
| Gm44698 | ENSMUSG( | ENSMUSG( | -0.29165 | 0.627751 | NO | NO | NA |
| A730091E: | ENSMUSG( | ENSMUSG( | 0.049008 | 0.942177 | NO | NO | NA |
| 4930528D | ENSMUSG( | ENSMUSG( | 1.208146 | 0.177644 | NO | NO | NA |
| Strip2 | ENSMUSG( | ENSMUSG( | 0.808526 | 0.130256 | NO | NO | NA |
| Drc3 | ENSMUSG( | ENSMUSG( | 1.187059 | 0.073847 | NO | NO | NA |
| Gm15983 | ENSMUSG( | ENSMUSG( | -0.2512 | 0.665005 | NO | NO | NA |
| Gm40617 | ENSMUSG( | ENSMUSG( | 0.644647 | 0.57964 | NO | NO | NA |
| P3h3 | ENSMUSG( | ENSMUSG( | 0.342301 | 0.664323 | NO | NO | NA |
| Gm14414 | ENSMUSG( | ENSMUSG( | -0.393 | 0.919171 | NO | NO | NA |
| Gm42843 | ENSMUSG( | ENSMUSG( | 0.850328 | 0.498883 | NO | NO | NA |
| Gm47920 | ENSMUSG( | ENSMUSG( | -0.45631 | 0.271539 | NO | NO | NA |
| Gm19018 | ENSMUSG( | ENSMUSG( | 0.602182 | 0.563484 | NO | NO | NA |
| Calr4 | ENSMUSG( | ENSMUSG( | 0.904846 | 0.429294 | NO | NO | NA |
| Gm48737 | ENSMUSG( | ENSMUSG( | 0.686257 | 0.528109 | NO | NO | NA |
| Gm32913 | ENSMUSG( | ENSMUSG( | 0.67131 | 0.583554 | NO | NO | NA |
| Gm5035 | ENSMUSG( | ENSMUSG( | -0.02942 | 0.993351 | NO | NO | NA |
| Klhl41 | ENSMUSG( | ENSMUSG( | 0.978475 | 0.347317 | NO | NO | NA |
| Vmn1r197 | ENSMUSG( | ENSMUSG( | 0.806094 | 0.460439 | NO | NO | NA |
| Gm19817 | ENSMUSG( | ENSMUSG( | 0.650471 | 0.645911 | NO | NO | NA |
| Gm7432 | ENSMUSG( | ENSMUSG( | 0.747233 | 0.305474 | NO | NO | NA |
| Olfr1173 | ENSMUSG( | ENSMUSG( | 0.284281 | 0.797665 | NO | NO | NA |
| A930012L: | ENSMUSG( | ENSMUSG( | 0.903829 | 0.431293 | NO | NO | NA |
| 9530059O | ENSMUSG( | ENSMUSG( | 0.04446 | 0.961918 | NO | NO | NA |
| Gm9944 | ENSMUSG( | ENSMUSG( | 0.048776 | 0.963858 | NO | NO | NA |
| Gm19510 | ENSMUSG( | ENSMUSG( | -0.20234 | 0.652373 | NO | NO | NA |
| Gm46649 | ENSMUSG( | ENSMUSG( | 0.06767 | 0.904371 | NO | NO | NA |
| Vmn2r27 | ENSMUSG( | ENSMUSG( | 0.825621 | 0.527421 | NO | NO | NA |
| Plppr2 | ENSMUSG( | ENSMUSG( | 0.638362 | 0.197539 | NO | NO | NA |
| Pheta1 | ENSMUSG( | ENSMUSG( | 0.105018 | 0.819022 | NO | NO | NA |
| Gm37163 | ENSMUSG( | ENSMUSG( | -0.05779 | 0.939403 | NO | NO | NA |
| Lingo1 | ENSMUSG( | ENSMUSG( | 1.079202 | 0.187274 | NO | NO | NA |
| Vmn2r111 | ENSMUSG( | ENSMUSG( | 1.45414 | 0.214327 | NO | NO | NA |
| 8030456M | ENSMUSG( | ENSMUSG( | 0.276142 | 0.583984 | NO | NO | NA |
| Gm37777 | ENSMUSG( | ENSMUSG( | 0.259061 | 0.840825 | NO | NO | NA |
| Gm33370 | ENSMUSG( | ENSMUSG( | 0.786187 | 0.304061 | NO | NO | NA |
| Slc2a6 | ENSMUSG( | ENSMUSG( | -0.04139 | 0.933874 | NO | NO | NA |
| Gm45242 | ENSMUSG( | ENSMUSG( | 0.681466 | 0.12198 | NO | NO | NA |
| Ascl4 | ENSMUSG( | ENSMUSG( | 0.64228 | 0.480046 | NO | NO | NA |
| Gm47739 | ENSMUSG( | ENSMUSG( | -0.5124 | 0.595207 | NO | NO | NA |
| F12 | ENSMUSG( | ENSMUSG( | 0.365362 | 0.626728 | NO | NO | NA |
| Vmn2r8 | ENSMUSG( | ENSMUSG( | 0.745471 | 0.563717 | NO | NO | NA |
| Gm40377 | ENSMUSG( | ENSMUSG( | 0.627877 | 0.535762 | NO | NO | NA |
| ENSMUSG | ENSMUSG( | ENSMUSG( | 1.223762 | 0.303721 | NO | NO | NA |
| 1810010H | ENSMUSG( | ENSMUSG( | 0.917389 | 0.078381 | NO | NO | NA |
| Myom2 | ENSMUSG( | ENSMUSG( | -0.59333 | 0.328452 | NO | NO | NA |
| Olfr1010 | ENSMUSG( | ENSMUSG( | 0.525729 | 0.639774 | NO | NO | NA |
| Gm48888 | ENSMUSG( | ENSMUSG( | 0.831765 | 0.423479 | NO | NO | NA |

| | | | | | | | |
|---|---|---|---|---|---|---|---|
| Gm37410 | ENSMUSG( | ENSMUSG( | 1.11592 | 0.299276 | NO | NO | NA |
| ENSMUSG | ENSMUSG( | ENSMUSG( | 0.492386 | 0.29736 | NO | NO | NA |
| Gm4544 | ENSMUSG( | ENSMUSG( | 1.165223 | 0.31092 | NO | NO | NA |
| Gm10941 | ENSMUSG( | ENSMUSG( | 1.779366 | 0.002009 | YES | YES | Upregulated |
| Col5a3 | ENSMUSG( | ENSMUSG( | -0.18932 | 0.80602 | NO | NO | NA |
| Vmn2r50 | ENSMUSG( | ENSMUSG( | 0.621979 | 0.63321 | NO | NO | NA |
| Gm18188 | ENSMUSG( | ENSMUSG( | 1.444318 | 0.192053 | NO | NO | NA |
| 1700013F( | ENSMUSG( | ENSMUSG( | 0.656321 | 0.590936 | NO | NO | NA |
| Gm10357 | ENSMUSG( | ENSMUSG( | 0.68267 | 0.223572 | NO | NO | NA |
| Gm26971 | ENSMUSG( | ENSMUSG( | -2.58968 | 0.001955 | YES | YES | Downregulated |
| F530104D | ENSMUSG( | ENSMUSG( | -0.0962 | 0.839261 | NO | NO | NA |
| Olfr921 | ENSMUSG( | ENSMUSG( | 0.435535 | 0.698039 | NO | NO | NA |
| P3h2 | ENSMUSG( | ENSMUSG( | 2.596898 | 0.017488 | YES | YES | Upregulated |
| Bco2 | ENSMUSG( | ENSMUSG( | 0.770291 | 0.167887 | NO | NO | NA |
| Gm41611 | ENSMUSG( | ENSMUSG( | 0.671814 | 0.48065 | NO | NO | NA |
| Vmn1r233 | ENSMUSG( | ENSMUSG( | 0.440294 | 0.673065 | NO | NO | NA |
| Gm19098 | ENSMUSG( | ENSMUSG( | 0.414311 | 0.706403 | NO | NO | NA |
| Stap2 | ENSMUSG( | ENSMUSG( | -0.03228 | 0.962598 | NO | NO | NA |
| Pou2f3 | ENSMUSG( | ENSMUSG( | 0.384217 | 0.739587 | NO | NO | NA |
| Gm38001 | ENSMUSG( | ENSMUSG( | 0.419357 | 0.674809 | NO | NO | NA |
| Gm42900 | ENSMUSG( | ENSMUSG( | -0.57344 | 0.198985 | NO | NO | NA |
| Olfr659 | ENSMUSG( | ENSMUSG( | 0.506173 | 0.633232 | NO | NO | NA |
| Lrrc3 | ENSMUSG( | ENSMUSG( | 0.471966 | 0.64988 | NO | NO | NA |
| Gm14849 | ENSMUSG( | ENSMUSG( | 0.30538 | 0.822027 | NO | NO | NA |
| Gm42697 | ENSMUSG( | ENSMUSG( | 0.596402 | 0.556504 | NO | NO | NA |
| Olfr641 | ENSMUSG( | ENSMUSG( | 1.041103 | 0.334413 | NO | NO | NA |
| Olfr1477 | ENSMUSG( | ENSMUSG( | 1.018423 | 0.40431 | NO | NO | NA |
| Sugct | ENSMUSG( | ENSMUSG( | 0.330614 | 0.455954 | NO | NO | NA |
| Gm17193 | ENSMUSG( | ENSMUSG( | 0.428481 | 0.269406 | NO | NO | NA |
| Gm26903 | ENSMUSG( | ENSMUSG( | 0.987562 | 0.438698 | NO | NO | NA |
| Tc2n | ENSMUSG( | ENSMUSG( | 0.753081 | 0.44331 | NO | NO | NA |
| Gm6855 | ENSMUSG( | ENSMUSG( | -0.37079 | 0.35827 | NO | NO | NA |
| Gm12424 | ENSMUSG( | ENSMUSG( | -0.37673 | 0.35329 | NO | NO | NA |
| Spef1l | ENSMUSG( | ENSMUSG( | 0.767076 | 0.346331 | NO | NO | NA |
| Gm9634 | ENSMUSG( | ENSMUSG( | 0.213004 | 0.844131 | NO | NO | NA |
| Gm7862 | ENSMUSG( | ENSMUSG( | 0.773579 | 0.195917 | NO | NO | NA |
| Olfr1487 | ENSMUSG( | ENSMUSG( | 0.444128 | 0.677884 | NO | NO | NA |
| Gm45861 | ENSMUSG( | ENSMUSG( | 0.232513 | 0.83009 | NO | NO | NA |
| Olfr1167 | ENSMUSG( | ENSMUSG( | 0.073778 | 0.948899 | NO | NO | NA |
| Gm7334 | ENSMUSG( | ENSMUSG( | -1.42808 | 0.00349 | YES | YES | Downregulated |
| Dnajb3 | ENSMUSG( | ENSMUSG( | 0.057099 | 0.934384 | NO | NO | NA |
| Pantr2 | ENSMUSG( | ENSMUSG( | 0.643173 | 0.6204 | NO | NO | NA |
| Gm48854 | ENSMUSG( | ENSMUSG( | 0.220406 | 0.861256 | NO | NO | NA |
| Gm49479 | ENSMUSG( | ENSMUSG( | 0.805686 | 0.475168 | NO | NO | NA |
| Gm5855 | ENSMUSG( | ENSMUSG( | 0.716958 | 0.546443 | NO | NO | NA |
| 5430401H | ENSMUSG( | ENSMUSG( | 0.810193 | 0.242237 | NO | NO | NA |
| Ctf2 | ENSMUSG( | ENSMUSG( | 0.71367 | 0.530632 | NO | NO | NA |

| | | | | | | | |
|---|---|---|---|---|---|---|---|
| Rgs9bp | ENSMUSG( | ENSMUSG( | -0.59247 | 0.209122 | NO | NO | NA |
| 1700063D | ENSMUSG( | ENSMUSG( | 0.383747 | 0.738182 | NO | NO | NA |
| 1700011L( | ENSMUSG( | ENSMUSG( | -0.53839 | 0.313759 | NO | NO | NA |
| Lct | ENSMUSG( | ENSMUSG( | -0.17011 | 0.692012 | NO | NO | NA |
| ENSMUSG | ENSMUSG( | ENSMUSG( | 1.089399 | 0.382686 | NO | NO | NA |
| Snora73b | ENSMUSG( | ENSMUSG( | 0.950251 | 0.524405 | NO | NO | NA |
| Gm34549 | ENSMUSG( | ENSMUSG( | 0.894715 | 0.402464 | NO | NO | NA |
| Klk13 | ENSMUSG( | ENSMUSG( | 0.456369 | 0.584344 | NO | NO | NA |
| Gm37776 | ENSMUSG( | ENSMUSG( | 0.580413 | 0.602523 | NO | NO | NA |
| Olfr1408 | ENSMUSG( | ENSMUSG( | 1.587415 | 0.151822 | NO | NO | NA |
| Gm34667 | ENSMUSG( | ENSMUSG( | 0.094086 | 0.935818 | NO | NO | NA |
| Gm43258 | ENSMUSG( | ENSMUSG( | 0.693515 | 0.497159 | NO | NO | NA |
| Ccin | ENSMUSG( | ENSMUSG( | 0.266682 | 0.664907 | NO | NO | NA |
| D130009I1 | ENSMUSG( | ENSMUSG( | 0.870414 | 0.456373 | NO | NO | NA |
| Aspn | ENSMUSG( | ENSMUSG( | 1.09613 | 0.137993 | NO | NO | NA |
| Gm14703 | ENSMUSG( | ENSMUSG( | 0.413123 | 0.57346 | NO | NO | NA |
| 4930412C: | ENSMUSG( | ENSMUSG( | 1.094489 | 0.284435 | NO | NO | NA |
| Gm47606 | ENSMUSG( | ENSMUSG( | 1.33633 | 0.262115 | NO | NO | NA |
| Akt2-ps | ENSMUSG( | ENSMUSG( | -0.24661 | 0.579397 | NO | NO | NA |
| Gm42910 | ENSMUSG( | ENSMUSG( | 0.576245 | 0.583554 | NO | NO | NA |
| Gm48701 | ENSMUSG( | ENSMUSG( | -0.10592 | 0.771401 | NO | NO | NA |
| Vmn2r88 | ENSMUSG( | ENSMUSG( | 0.713598 | 0.501292 | NO | NO | NA |
| Zfp229 | ENSMUSG( | ENSMUSG( | 0.578551 | 0.334843 | NO | NO | NA |
| Gm48840 | ENSMUSG( | ENSMUSG( | 0.520444 | 0.334627 | NO | NO | NA |
| Gm48541 | ENSMUSG( | ENSMUSG( | 0.210319 | 0.803999 | NO | NO | NA |
| Olfr414 | ENSMUSG( | ENSMUSG( | 1.355956 | 0.24636 | NO | NO | NA |
| Itgb1bp2 | ENSMUSG( | ENSMUSG( | 0.071643 | 0.926005 | NO | NO | NA |
| ENSMUSG | ENSMUSG( | ENSMUSG( | -0.14885 | 0.756906 | NO | NO | NA |
| Gm43611 | ENSMUSG( | ENSMUSG( | 0.304871 | 0.816371 | NO | NO | NA |
| Olfr1219 | ENSMUSG( | ENSMUSG( | 0.587643 | 0.665321 | NO | NO | NA |
| Gm4961 | ENSMUSG( | ENSMUSG( | 0.372879 | 0.59852 | NO | NO | NA |
| Lrp2 | ENSMUSG( | ENSMUSG( | 1.10026 | 0.391549 | NO | NO | NA |
| Soga1 | ENSMUSG( | ENSMUSG( | 0.609422 | 0.280224 | NO | NO | NA |
| H3f3aos | ENSMUSG( | ENSMUSG( | 0.178476 | 0.727799 | NO | NO | NA |
| ENSMUSG | ENSMUSG( | ENSMUSG( | 0.782181 | 0.476215 | NO | NO | NA |
| Gm14682 | ENSMUSG( | ENSMUSG( | 1.571183 | 0.297828 | NO | NO | NA |
| Gm16236 | ENSMUSG( | ENSMUSG( | -0.05477 | 0.964427 | NO | NO | NA |
| Gm49291 | ENSMUSG( | ENSMUSG( | 0.89561 | 0.505844 | NO | NO | NA |
| 3110067C( | ENSMUSG( | ENSMUSG( | 0.211282 | 0.783902 | NO | NO | NA |
| Gm36602 | ENSMUSG( | ENSMUSG( | -1.02559 | 0.078883 | NO | NO | NA |
| Cald1 | ENSMUSG( | ENSMUSG( | -0.23405 | 0.779749 | NO | NO | NA |
| Cdcp2 | ENSMUSG( | ENSMUSG( | -0.098 | 0.895628 | NO | NO | NA |
| ENSMUSG | ENSMUSG( | ENSMUSG( | -0.82298 | 0.104616 | NO | NO | NA |
| Ighv1-47 | ENSMUSG( | ENSMUSG( | -4.92564 | NA | NO | NO | NA |
| Tmem88 | ENSMUSG( | ENSMUSG( | 0.507193 | 0.474362 | NO | NO | NA |
| 4930517E: | ENSMUSG( | ENSMUSG( | -0.22006 | 0.725572 | NO | NO | NA |
| Gm40999 | ENSMUSG( | ENSMUSG( | 0.529643 | 0.616442 | NO | NO | NA |

| | | | | | | | |
|---|---|---|---|---|---|---|---|
| Map3k19 | ENSMUSG( | ENSMUSG( | 0.679234 | 0.294909 | NO | NO | NA |
| Ms4a3 | ENSMUSG( | ENSMUSG( | 1.358983 | 0.19074 | NO | NO | NA |
| Adgre4 | ENSMUSG( | ENSMUSG( | -0.05231 | 0.958772 | NO | NO | NA |
| Tm6sf2 | ENSMUSG( | ENSMUSG( | 0.149686 | 0.722714 | NO | NO | NA |
| Gm10400 | ENSMUSG( | ENSMUSG( | 0.23198 | 0.836181 | NO | NO | NA |
| Gm20784 | ENSMUSG( | ENSMUSG( | -0.00222 | 0.99896 | NO | NO | NA |
| Sorcs1 | ENSMUSG( | ENSMUSG( | 0.785756 | 0.481958 | NO | NO | NA |
| Gm5422 | ENSMUSG( | ENSMUSG( | -0.27891 | 0.775658 | NO | NO | NA |
| Gm10503 | ENSMUSG( | ENSMUSG( | 0.080889 | 0.945917 | NO | NO | NA |
| Rsph14 | ENSMUSG( | ENSMUSG( | -0.33821 | 0.762738 | NO | NO | NA |
| Gm49167 | ENSMUSG( | ENSMUSG( | 0.531492 | 0.723831 | NO | NO | NA |
| ENSMUSG | ENSMUSG( | ENSMUSG( | 0.828681 | 0.153294 | NO | NO | NA |
| Tspan12 | ENSMUSG( | ENSMUSG( | 0.710854 | 0.55484 | NO | NO | NA |
| Dgkb | ENSMUSG( | ENSMUSG( | 0.859091 | 0.344361 | NO | NO | NA |
| 4933435F: | ENSMUSG( | ENSMUSG( | 0.34699 | 0.735201 | NO | NO | NA |
| Gpr135 | ENSMUSG( | ENSMUSG( | 1.001922 | 0.178926 | NO | NO | NA |
| 4930428E( | ENSMUSG( | ENSMUSG( | 0.761456 | 0.534067 | NO | NO | NA |
| Gm4940 | ENSMUSG( | ENSMUSG( | 0.404644 | 0.733315 | NO | NO | NA |
| Pcsk9 | ENSMUSG( | ENSMUSG( | 1.224139 | 0.183655 | NO | NO | NA |
| Gm20745 | ENSMUSG( | ENSMUSG( | 0.931467 | 0.486916 | NO | NO | NA |
| Gm26692 | ENSMUSG( | ENSMUSG( | -0.07045 | 0.938653 | NO | NO | NA |
| Gm5525 | ENSMUSG( | ENSMUSG( | 0.356398 | 0.408597 | NO | NO | NA |
| S100a3 | ENSMUSG( | ENSMUSG( | 0.52461 | 0.2254 | NO | NO | NA |
| Gm12799 | ENSMUSG( | ENSMUSG( | -0.11213 | 0.814825 | NO | NO | NA |
| Gm4130 | ENSMUSG( | ENSMUSG( | 0.012556 | 0.983447 | NO | NO | NA |
| Fhl1 | ENSMUSG( | ENSMUSG( | 0.918805 | 0.325219 | NO | NO | NA |
| Speer4f2 | ENSMUSG( | ENSMUSG( | 1.015536 | 0.399941 | NO | NO | NA |
| Gm4928 | ENSMUSG( | ENSMUSG( | 0.428214 | 0.446781 | NO | NO | NA |
| Gm11766 | ENSMUSG( | ENSMUSG( | -1.61763 | 0.000729 | YES | YES | Downregulated |
| Rex2 | ENSMUSG( | ENSMUSG( | 3.869628 | 1.02E-05 | YES | YES | Upregulated |
| Gm49214 | ENSMUSG( | ENSMUSG( | 0.691402 | 0.455954 | NO | NO | NA |
| ENSMUSG | ENSMUSG( | ENSMUSG( | 0.11702 | 0.852301 | NO | NO | NA |
| Grifin | ENSMUSG( | ENSMUSG( | 1.285016 | 0.028374 | YES | YES | Upregulated |
| Gm5940 | ENSMUSG( | ENSMUSG( | 1.102602 | 0.609081 | NO | NO | NA |
| Gm43930 | ENSMUSG( | ENSMUSG( | 0.219175 | 0.849581 | NO | NO | NA |
| Rpl31-ps9 | ENSMUSG( | ENSMUSG( | 0.023586 | 0.980452 | NO | NO | NA |
| Gm8274 | ENSMUSG( | ENSMUSG( | -0.37111 | 0.826387 | NO | NO | NA |
| Slc39a5 | ENSMUSG( | ENSMUSG( | -0.10958 | 0.905937 | NO | NO | NA |
| Fam229b | ENSMUSG( | ENSMUSG( | 1.067157 | 0.149429 | NO | NO | NA |
| Ackr1 | ENSMUSG( | ENSMUSG( | 1.351026 | 0.043827 | YES | YES | Upregulated |
| Hadhb-ps | ENSMUSG( | ENSMUSG( | -0.05983 | 0.912074 | NO | NO | NA |
| Gm7935 | ENSMUSG( | ENSMUSG( | 0.381665 | 0.595771 | NO | NO | NA |
| Vmn2r40 | ENSMUSG( | ENSMUSG( | 1.240529 | 0.342628 | NO | NO | NA |
| Asb10 | ENSMUSG( | ENSMUSG( | 1.302103 | 0.201637 | NO | NO | NA |
| ENSMUSG | ENSMUSG( | ENSMUSG( | 0.244641 | 0.8114 | NO | NO | NA |
| Lnx1 | ENSMUSG( | ENSMUSG( | 0.207747 | 0.711222 | NO | NO | NA |
| Olfr1212 | ENSMUSG( | ENSMUSG( | 1.077144 | 0.354826 | NO | NO | NA |

| | | | | | | | |
|---|---|---|---|---|---|---|---|
| Gm4609 | ENSMUSG( | ENSMUSG( | -0.15875 | 0.874941 | NO | NO | NA |
| E230020D | ENSMUSG( | ENSMUSG( | 0.82772 | 0.465525 | NO | NO | NA |
| Olfr370 | ENSMUSG( | ENSMUSG( | 0.500539 | 0.661985 | NO | NO | NA |
| Gm45208 | ENSMUSG( | ENSMUSG( | 1.809449 | 0.42412 | NO | NO | NA |
| Pycr1 | ENSMUSG( | ENSMUSG( | -0.15954 | 0.746839 | NO | NO | NA |
| Olfr300-ps | ENSMUSG( | ENSMUSG( | 0.627826 | 0.572005 | NO | NO | NA |
| Gm14168 | ENSMUSG( | ENSMUSG( | 0.229917 | 0.620289 | NO | NO | NA |
| Flg | ENSMUSG( | ENSMUSG( | 0.179233 | 0.881723 | NO | NO | NA |
| Gm44078 | ENSMUSG( | ENSMUSG( | 1.103315 | 0.027197 | YES | YES | Upregulated |
| Olfr24 | ENSMUSG( | ENSMUSG( | 0.621852 | 0.559018 | NO | NO | NA |
| Pgbd1 | ENSMUSG( | ENSMUSG( | 0.789164 | 0.123816 | NO | NO | NA |
| Chad | ENSMUSG( | ENSMUSG( | -0.69253 | 0.154439 | NO | NO | NA |
| Krt2 | ENSMUSG( | ENSMUSG( | 0.492299 | 0.673578 | NO | NO | NA |
| Gm6882 | ENSMUSG( | ENSMUSG( | -0.76171 | 0.547864 | NO | NO | NA |
| Gm10291 | ENSMUSG( | ENSMUSG( | 0.923314 | 0.137529 | NO | NO | NA |
| BC024386 | ENSMUSG( | ENSMUSG( | 0.720349 | 0.476481 | NO | NO | NA |
| 4931408D | ENSMUSG( | ENSMUSG( | -0.86945 | 0.447868 | NO | NO | NA |
| C330013E: | ENSMUSG( | ENSMUSG( | 0.183681 | 0.712797 | NO | NO | NA |
| Gm6368 | ENSMUSG( | ENSMUSG( | 0.242571 | 0.611216 | NO | NO | NA |
| Gm50008 | ENSMUSG( | ENSMUSG( | 0.796301 | 0.460998 | NO | NO | NA |
| Gm45059 | ENSMUSG( | ENSMUSG( | -0.24954 | 0.710987 | NO | NO | NA |
| Gm37708 | ENSMUSG( | ENSMUSG( | 1.050991 | 0.39701 | NO | NO | NA |
| Gm38307 | ENSMUSG( | ENSMUSG( | 0.49319 | 0.690757 | NO | NO | NA |
| Asb16 | ENSMUSG( | ENSMUSG( | 1.409371 | 0.036538 | YES | YES | Upregulated |
| Gm14048 | ENSMUSG( | ENSMUSG( | -0.09036 | 0.857524 | NO | NO | NA |
| ENSMUSG( | ENSMUSG( | ENSMUSG( | -0.50045 | 0.475272 | NO | NO | NA |
| Tle6 | ENSMUSG( | ENSMUSG( | 0.743609 | 0.268859 | NO | NO | NA |
| Ptpdc1 | ENSMUSG( | ENSMUSG( | 0.838 | 0.10595 | NO | NO | NA |
| Tmem40 | ENSMUSG( | ENSMUSG( | 0.870217 | 0.351951 | NO | NO | NA |
| Gm38259 | ENSMUSG( | ENSMUSG( | 1.035593 | 0.514255 | NO | NO | NA |
| Gm7887 | ENSMUSG( | ENSMUSG( | -0.12901 | 0.826539 | NO | NO | NA |
| 4933438A | ENSMUSG( | ENSMUSG( | 0.21486 | 0.847974 | NO | NO | NA |
| Gm15657 | ENSMUSG( | ENSMUSG( | 0.057331 | 0.956941 | NO | NO | NA |
| Pih1d2 | ENSMUSG( | ENSMUSG( | 0.541014 | 0.243419 | NO | NO | NA |
| Cadps2 | ENSMUSG( | ENSMUSG( | 0.842145 | 0.491395 | NO | NO | NA |
| Gm3807 | ENSMUSG( | ENSMUSG( | 0.425186 | 0.5201 | NO | NO | NA |
| Gm10655 | ENSMUSG( | ENSMUSG( | -0.3092 | 0.703822 | NO | NO | NA |
| Gm45834 | ENSMUSG( | ENSMUSG( | 0.601197 | 0.565679 | NO | NO | NA |
| ENSMUSG | ENSMUSG( | ENSMUSG( | -0.08111 | 0.894999 | NO | NO | NA |
| Gm13231 | ENSMUSG( | ENSMUSG( | -2.12085 | 0.010709 | YES | YES | Downregulated |
| Olfr1105 | ENSMUSG( | ENSMUSG( | 0.738765 | 0.568223 | NO | NO | NA |
| Gm20136 | ENSMUSG( | ENSMUSG( | 0.737172 | 0.595947 | NO | NO | NA |
| Olfr1 | ENSMUSG( | ENSMUSG( | 0.95211 | 0.454653 | NO | NO | NA |
| Il11 | ENSMUSG( | ENSMUSG( | 0.475703 | 0.564493 | NO | NO | NA |
| Gm42876 | ENSMUSG( | ENSMUSG( | 0.104605 | 0.858109 | NO | NO | NA |
| Rnf223 | ENSMUSG( | ENSMUSG( | 0.376592 | 0.585486 | NO | NO | NA |
| ENSMUSG | ENSMUSG( | ENSMUSG( | 0.850297 | 0.246037 | NO | NO | NA |

| | | | | | | | |
|---|---|---|---|---|---|---|---|
| ENSMUSG( | ENSMUSG( | ENSMUSG( | 0.31095 | 0.501197 | NO | NO | NA |
| Islr | ENSMUSG( | ENSMUSG( | -0.52993 | 0.41347 | NO | NO | NA |
| 4933431J2 | ENSMUSG( | ENSMUSG( | 0.914868 | 0.429286 | NO | NO | NA |
| Olfr1118 | ENSMUSG( | ENSMUSG( | 0.331586 | 0.763396 | NO | NO | NA |
| Lactbl1 | ENSMUSG( | ENSMUSG( | -0.20308 | 0.658781 | NO | NO | NA |
| Slc38a4 | ENSMUSG( | ENSMUSG( | 0.750201 | 0.495773 | NO | NO | NA |
| Gm11455 | ENSMUSG( | ENSMUSG( | 0.187181 | 0.779114 | NO | NO | NA |
| ENSMUSG | ENSMUSG( | ENSMUSG( | 0.898563 | 0.38657 | NO | NO | NA |
| Gm10269 | ENSMUSG( | ENSMUSG( | 0.922548 | 0.018616 | YES | NO | NA |
| Fbll1 | ENSMUSG( | ENSMUSG( | -0.20905 | 0.868358 | NO | NO | NA |
| Kif26a | ENSMUSG( | ENSMUSG( | 0.431402 | 0.707602 | NO | NO | NA |
| Extl1 | ENSMUSG( | ENSMUSG( | 0.861254 | 0.342553 | NO | NO | NA |
| ENSMUSG | ENSMUSG( | ENSMUSG( | 0.818497 | 0.458202 | NO | NO | NA |
| A930006L( | ENSMUSG( | ENSMUSG( | -0.22561 | 0.637771 | NO | NO | NA |
| Thbs2 | ENSMUSG( | ENSMUSG( | 0.314497 | 0.772658 | NO | NO | NA |
| Ighv1-5 | ENSMUSG( | ENSMUSG( | -0.30855 | 0.939729 | NO | NO | NA |
| Bsx | ENSMUSG( | ENSMUSG( | 0.913507 | 0.417582 | NO | NO | NA |
| Olfr911-ps | ENSMUSG( | ENSMUSG( | 0.873761 | 0.495392 | NO | NO | NA |
| Gm11298 | ENSMUSG( | ENSMUSG( | 1.334842 | 0.306821 | NO | NO | NA |
| Ece2 | ENSMUSG( | ENSMUSG( | 0.547478 | 0.543488 | NO | NO | NA |
| Slc35f1 | ENSMUSG( | ENSMUSG( | 0.383989 | 0.743114 | NO | NO | NA |
| Bcar1 | ENSMUSG( | ENSMUSG( | 0.281311 | 0.595965 | NO | NO | NA |
| Mst1 | ENSMUSG( | ENSMUSG( | -0.06783 | 0.878431 | NO | NO | NA |
| Gm37065 | ENSMUSG( | ENSMUSG( | -0.16593 | 0.702237 | NO | NO | NA |
| 1700102P( | ENSMUSG( | ENSMUSG( | -0.48788 | 0.254124 | NO | NO | NA |
| Gm10481 | ENSMUSG( | ENSMUSG( | -0.07765 | 0.929889 | NO | NO | NA |
| Mest | ENSMUSG( | ENSMUSG( | -0.3404 | 0.503776 | NO | NO | NA |
| Acnat1 | ENSMUSG( | ENSMUSG( | 0.247809 | 0.802041 | NO | NO | NA |
| Gm6485 | ENSMUSG( | ENSMUSG( | -0.07334 | 0.899717 | NO | NO | NA |
| Prokr2 | ENSMUSG( | ENSMUSG( | -0.34599 | 0.827561 | NO | NO | NA |
| Bahcc1 | ENSMUSG( | ENSMUSG( | -0.0192 | 0.965837 | NO | NO | NA |
| Zkscan2 | ENSMUSG( | ENSMUSG( | 0.091124 | 0.932161 | NO | NO | NA |
| Gm42862 | ENSMUSG( | ENSMUSG( | 1.270105 | 0.191906 | NO | NO | NA |
| Rab34 | ENSMUSG( | ENSMUSG( | 0.50128 | 0.495961 | NO | NO | NA |
| Gm49450 | ENSMUSG( | ENSMUSG( | -0.00808 | 0.997769 | NO | NO | NA |
| Cfap157 | ENSMUSG( | ENSMUSG( | 1.114623 | 0.115391 | NO | NO | NA |
| Gm7276 | ENSMUSG( | ENSMUSG( | 1.070118 | 0.296203 | NO | NO | NA |
| Gm43311 | ENSMUSG( | ENSMUSG( | 0.723177 | 0.507817 | NO | NO | NA |
| Gm44165 | ENSMUSG( | ENSMUSG( | 0.895365 | 0.440714 | NO | NO | NA |
| 7SK ENSM | ENSMUSG( | ENSMUSG( | 0.650504 | 0.750846 | NO | NO | NA |
| Csf2 | ENSMUSG( | ENSMUSG( | 0.316671 | 0.462135 | NO | NO | NA |
| Gm43508 | ENSMUSG( | ENSMUSG( | -0.12094 | 0.922935 | NO | NO | NA |
| Tmem45a2 | ENSMUSG( | ENSMUSG( | 0.920848 | 0.416514 | NO | NO | NA |
| Olfr772 | ENSMUSG( | ENSMUSG( | 0.194039 | 0.860489 | NO | NO | NA |
| Gm37092 | ENSMUSG( | ENSMUSG( | 0.860525 | 0.505024 | NO | NO | NA |
| Gabra4 | ENSMUSG( | ENSMUSG( | 0.326191 | 0.766456 | NO | NO | NA |
| Gm28590 | ENSMUSG( | ENSMUSG( | 0.669651 | 0.576707 | NO | NO | NA |

| | | | | | | | |
|---|---|---|---|---|---|---|---|
| Wnk4 | ENSMUSG( | ENSMUSG( | 0.265839 | 0.664323 | NO | NO | NA |
| Gm11599 | ENSMUSG( | ENSMUSG( | 0.349909 | 0.748417 | NO | NO | NA |
| Wee2 | ENSMUSG( | ENSMUSG( | 0.773624 | 0.572582 | NO | NO | NA |
| Prss39 | ENSMUSG( | ENSMUSG( | 0.853877 | 0.451408 | NO | NO | NA |
| Trav7n-6 | ENSMUSG( | ENSMUSG( | 0.333066 | NA | NO | NO | NA |
| Gm38255 | ENSMUSG( | ENSMUSG( | 0.675411 | 0.529241 | NO | NO | NA |
| Gm37562 | ENSMUSG( | ENSMUSG( | 1.051017 | 0.416121 | NO | NO | NA |
| Pdzd4 | ENSMUSG( | ENSMUSG( | 0.126634 | 0.794149 | NO | NO | NA |
| Gm4129 | ENSMUSG( | ENSMUSG( | 0.10009 | 0.828338 | NO | NO | NA |
| ENSMUSG | ENSMUSG( | ENSMUSG( | 0.737956 | 0.507429 | NO | NO | NA |
| Gm22567 | ENSMUSG( | ENSMUSG( | 1.842947 | 0.03589 | YES | YES | Upregulated |
| Vmn1r72 | ENSMUSG( | ENSMUSG( | 0.462155 | 0.697974 | NO | NO | NA |
| Ntn4 | ENSMUSG( | ENSMUSG( | 0.267 | 0.781352 | NO | NO | NA |
| Fam13c | ENSMUSG( | ENSMUSG( | -0.19787 | 0.843643 | NO | NO | NA |
| Avpr1b | ENSMUSG( | ENSMUSG( | 0.806119 | 0.46441 | NO | NO | NA |
| Gm12599 | ENSMUSG( | ENSMUSG( | 0.470993 | 0.636333 | NO | NO | NA |
| Rpl36a-ps1 | ENSMUSG( | ENSMUSG( | 0.643619 | 0.308328 | NO | NO | NA |
| Srsf3-ps | ENSMUSG( | ENSMUSG( | 0.612328 | 0.220452 | NO | NO | NA |
| Galnt13 | ENSMUSG( | ENSMUSG( | 0.602706 | 0.55059 | NO | NO | NA |
| Gm4565 | ENSMUSG( | ENSMUSG( | 0.281349 | 0.902186 | NO | NO | NA |
| Gm48182 | ENSMUSG( | ENSMUSG( | 0.779821 | 0.526309 | NO | NO | NA |
| Gm48566 | ENSMUSG( | ENSMUSG( | 0.751432 | 0.588985 | NO | NO | NA |
| Gm36189 | ENSMUSG( | ENSMUSG( | -0.44814 | 0.336818 | NO | NO | NA |
| Olfr535 | ENSMUSG( | ENSMUSG( | -0.40708 | 0.749282 | NO | NO | NA |
| Rpl18a-ps1 | ENSMUSG( | ENSMUSG( | 0.139952 | 0.888811 | NO | NO | NA |
| Pcdhgb8 | ENSMUSG( | ENSMUSG( | -0.58318 | 0.624845 | NO | NO | NA |
| Klra2 | ENSMUSG( | ENSMUSG( | 1.119439 | 0.306811 | NO | NO | NA |
| Vmn2r30 | ENSMUSG( | ENSMUSG( | 0.993387 | 0.384859 | NO | NO | NA |
| Gm38387 | ENSMUSG( | ENSMUSG( | -0.21108 | 0.734149 | NO | NO | NA |
| Gm37396 | ENSMUSG( | ENSMUSG( | 0.313107 | 0.781484 | NO | NO | NA |
| 2610203C2 | ENSMUSG( | ENSMUSG( | 0.428044 | 0.732447 | NO | NO | NA |
| Gm44008 | ENSMUSG( | ENSMUSG( | -2.50974 | 0.000566 | YES | YES | Downregulated |
| Gm6644 | ENSMUSG( | ENSMUSG( | 0.145957 | 0.812696 | NO | NO | NA |
| Glycam1 | ENSMUSG( | ENSMUSG( | 0.324775 | 0.806236 | NO | NO | NA |
| Gm36033 | ENSMUSG( | ENSMUSG( | 0.94071 | 0.438742 | NO | NO | NA |
| ENSMUSG | ENSMUSG( | ENSMUSG( | 0.605741 | 0.355689 | NO | NO | NA |
| Gm13034 | ENSMUSG( | ENSMUSG( | 0.969431 | 0.103007 | NO | NO | NA |
| Kdelr3 | ENSMUSG( | ENSMUSG( | 0.443244 | 0.525251 | NO | NO | NA |
| Gm16152 | ENSMUSG( | ENSMUSG( | 1.246015 | 0.30414 | NO | NO | NA |
| Dhx58os | ENSMUSG( | ENSMUSG( | -0.60084 | 0.218427 | NO | NO | NA |
| Cep112os1 | ENSMUSG( | ENSMUSG( | 0.329333 | 0.760745 | NO | NO | NA |
| Olfr681 | ENSMUSG( | ENSMUSG( | 0.991159 | 0.327932 | NO | NO | NA |
| Gm44053 | ENSMUSG( | ENSMUSG( | -0.29089 | 0.524628 | NO | NO | NA |
| ENSMUSG | ENSMUSG( | ENSMUSG( | -0.2553 | 0.635155 | NO | NO | NA |
| Bmp2 | ENSMUSG( | ENSMUSG( | 0.860153 | 0.441308 | NO | NO | NA |
| Neo1 | ENSMUSG( | ENSMUSG( | 0.870011 | 0.2761 | NO | NO | NA |
| Olfr731 | ENSMUSG( | ENSMUSG( | 0.685588 | 0.509107 | NO | NO | NA |

| | | | | | | | |
|---|---|---|---|---|---|---|---|
| Gm17814 | ENSMUSG( | ENSMUSG( | 0.540268 | 0.655215 | NO | NO | NA |
| Gm11290 | ENSMUSG( | ENSMUSG( | 0.827946 | 0.366632 | NO | NO | NA |
| Olfr553 | ENSMUSG( | ENSMUSG( | 0.573938 | 0.62014 | NO | NO | NA |
| Gm19144 | ENSMUSG( | ENSMUSG( | -0.04019 | 0.932022 | NO | NO | NA |
| Gm5879 | ENSMUSG( | ENSMUSG( | 0.4043 | 0.566179 | NO | NO | NA |
| Catsperg2 | ENSMUSG( | ENSMUSG( | 1.024345 | 0.319978 | NO | NO | NA |
| Gm47004 | ENSMUSG( | ENSMUSG( | 0.11626 | 0.880844 | NO | NO | NA |
| Gm8926 | ENSMUSG( | ENSMUSG( | 1.515029 | 0.353769 | NO | NO | NA |
| Gm34961 | ENSMUSG( | ENSMUSG( | 0.657997 | 0.46419 | NO | NO | NA |
| Ms4a7 | ENSMUSG( | ENSMUSG( | 0.492408 | 0.479158 | NO | NO | NA |
| Gm12382 | ENSMUSG( | ENSMUSG( | 0.136534 | 0.920093 | NO | NO | NA |
| Gm43131 | ENSMUSG( | ENSMUSG( | 1.585096 | 0.181931 | NO | NO | NA |
| BC006965 | ENSMUSG( | ENSMUSG( | 0.873528 | 0.496053 | NO | NO | NA |
| ENSMUSG | ENSMUSG( | ENSMUSG( | 0.910964 | 0.439278 | NO | NO | NA |
| Svet1 | ENSMUSG( | ENSMUSG( | 1.281621 | 0.292887 | NO | NO | NA |
| Gm26774 | ENSMUSG( | ENSMUSG( | 1.159459 | 0.309679 | NO | NO | NA |
| 4930455C: | ENSMUSG( | ENSMUSG( | 0.139069 | 0.797008 | NO | NO | NA |
| Gm18003 | ENSMUSG( | ENSMUSG( | 0.441981 | 0.693347 | NO | NO | NA |
| Gm12844 | ENSMUSG( | ENSMUSG( | 0.12101 | 0.797055 | NO | NO | NA |
| Gm34006 | ENSMUSG( | ENSMUSG( | -1.12471 | 0.008235 | YES | YES | Downregulated |
| Atf3 | ENSMUSG( | ENSMUSG( | 0.375588 | 0.711222 | NO | NO | NA |
| Sapcd1 | ENSMUSG( | ENSMUSG( | 0.574764 | 0.25241 | NO | NO | NA |
| Mfap3l | ENSMUSG( | ENSMUSG( | 0.713054 | 0.322609 | NO | NO | NA |
| Prodh | ENSMUSG( | ENSMUSG( | 1.299266 | 0.114342 | NO | NO | NA |
| Hacd4 | ENSMUSG( | ENSMUSG( | -0.04546 | 0.941752 | NO | NO | NA |
| Gm12276 | ENSMUSG( | ENSMUSG( | -0.50313 | 0.179788 | NO | NO | NA |
| Gpc4 | ENSMUSG( | ENSMUSG( | -0.04661 | 0.975184 | NO | NO | NA |
| Zfp804a | ENSMUSG( | ENSMUSG( | 0.69767 | 0.563166 | NO | NO | NA |
| Gm44941 | ENSMUSG( | ENSMUSG( | 0.197516 | 0.703822 | NO | NO | NA |
| Gm34016 | ENSMUSG( | ENSMUSG( | 0.485329 | 0.641488 | NO | NO | NA |
| Hepacam | ENSMUSG( | ENSMUSG( | 0.900585 | 0.288272 | NO | NO | NA |
| Gm44886 | ENSMUSG( | ENSMUSG( | 0.604806 | 0.648171 | NO | NO | NA |
| Gm19409 | ENSMUSG( | ENSMUSG( | -0.2343 | 0.765414 | NO | NO | NA |
| Gm9937 | ENSMUSG( | ENSMUSG( | -0.86902 | 0.175573 | NO | NO | NA |
| Olfr1501 | ENSMUSG( | ENSMUSG( | 0.56668 | 0.583602 | NO | NO | NA |
| Gm39327 | ENSMUSG( | ENSMUSG( | 0.720049 | 0.553957 | NO | NO | NA |
| Rps23-ps2 | ENSMUSG( | ENSMUSG( | 0.218631 | 0.675677 | NO | NO | NA |
| AI314278 | ENSMUSG( | ENSMUSG( | 0.948509 | 0.491395 | NO | NO | NA |
| Gm37040 | ENSMUSG( | ENSMUSG( | 0.196328 | 0.849635 | NO | NO | NA |
| Kcnd1 | ENSMUSG( | ENSMUSG( | 0.555194 | 0.359187 | NO | NO | NA |
| Akr1cl | ENSMUSG( | ENSMUSG( | 1.131331 | 0.334128 | NO | NO | NA |
| 4930444P: | ENSMUSG( | ENSMUSG( | -0.047 | 0.970343 | NO | NO | NA |
| ENSMUSG | ENSMUSG( | ENSMUSG( | 0.387796 | 0.75435 | NO | NO | NA |
| Gm7890 | ENSMUSG( | ENSMUSG( | -0.20063 | 0.648535 | NO | NO | NA |
| Gm16229 | ENSMUSG( | ENSMUSG( | 0.631782 | 0.406768 | NO | NO | NA |
| Gm44916 | ENSMUSG( | ENSMUSG( | 0.48307 | 0.42929 | NO | NO | NA |
| Rbpms2 | ENSMUSG( | ENSMUSG( | 0.379972 | 0.748836 | NO | NO | NA |

| | | | | | | | |
|---|---|---|---|---|---|---|---|
| Gm6733 | ENSMUSG( | ENSMUSG( | -0.17476 | 0.737905 | NO | NO | NA |
| P2ry4 | ENSMUSG( | ENSMUSG( | 0.393177 | 0.698787 | NO | NO | NA |
| Gm9247 | ENSMUSG( | ENSMUSG( | 0.825813 | 0.230497 | NO | NO | NA |
| Gm33847 | ENSMUSG( | ENSMUSG( | 0.259762 | 0.873602 | NO | NO | NA |
| 1190001M | ENSMUSG( | ENSMUSG( | 0.456197 | 0.515887 | NO | NO | NA |
| Vmn2r-ps4 | ENSMUSG( | ENSMUSG( | 0.900039 | 0.493225 | NO | NO | NA |
| C630044B | ENSMUSG( | ENSMUSG( | 0.25509 | 0.810758 | NO | NO | NA |
| Igkv1-122 | ENSMUSG( | ENSMUSG( | -3.97085 | 0.271018 | NO | NO | NA |
| Pth1r | ENSMUSG( | ENSMUSG( | 0.849416 | 0.49193 | NO | NO | NA |
| Gm49380 | ENSMUSG( | ENSMUSG( | -0.27806 | 0.874197 | NO | NO | NA |
| Olfr395 | ENSMUSG( | ENSMUSG( | 0.738234 | 0.537122 | NO | NO | NA |
| Myoz1 | ENSMUSG( | ENSMUSG( | 1.091556 | 0.344374 | NO | NO | NA |
| Gm47578 | ENSMUSG( | ENSMUSG( | -0.40547 | 0.631695 | NO | NO | NA |
| BB557941 | ENSMUSG( | ENSMUSG( | -0.88138 | 0.257899 | NO | NO | NA |
| Gm20522 | ENSMUSG( | ENSMUSG( | 0.098679 | 0.816045 | NO | NO | NA |
| Gm42997 | ENSMUSG( | ENSMUSG( | -1.68983 | 0.010623 | YES | YES | Downregulated |
| Frzb | ENSMUSG( | ENSMUSG( | 1.347725 | 0.32874 | NO | NO | NA |
| Actg2 | ENSMUSG( | ENSMUSG( | -0.67044 | 0.202665 | NO | NO | NA |
| Gm31645 | ENSMUSG( | ENSMUSG( | 0.908248 | 0.188076 | NO | NO | NA |
| ENSMUSG( | ENSMUSG( | ENSMUSG( | 0.300763 | 0.80634 | NO | NO | NA |
| 2610042L( | ENSMUSG( | ENSMUSG( | 0.937776 | 0.335381 | NO | NO | NA |
| Slitrk4 | ENSMUSG( | ENSMUSG( | 0.594453 | 0.612241 | NO | NO | NA |
| Wnt2b | ENSMUSG( | ENSMUSG( | -0.24707 | 0.714349 | NO | NO | NA |
| Gm40153 | ENSMUSG( | ENSMUSG( | 0.986547 | 0.422229 | NO | NO | NA |
| Gm18164 | ENSMUSG( | ENSMUSG( | 5.662214 | 0.043248 | YES | YES | Upregulated |
| Tmbim7 | ENSMUSG( | ENSMUSG( | 0.929128 | 0.449488 | NO | NO | NA |
| Tgm1 | ENSMUSG( | ENSMUSG( | -0.01538 | 0.976978 | NO | NO | NA |
| Gpr3 | ENSMUSG( | ENSMUSG( | 0.688097 | 0.351445 | NO | NO | NA |
| Olfr156 | ENSMUSG( | ENSMUSG( | 0.810252 | 0.480378 | NO | NO | NA |
| Topaz1 | ENSMUSG( | ENSMUSG( | 1.212165 | 0.160073 | NO | NO | NA |
| Gm21378 | ENSMUSG( | ENSMUSG( | 0.310894 | 0.827284 | NO | NO | NA |
| Alox12 | ENSMUSG( | ENSMUSG( | 0.259959 | 0.717192 | NO | NO | NA |
| Zic3 | ENSMUSG( | ENSMUSG( | 0.618548 | 0.569018 | NO | NO | NA |
| Vmn2r101 | ENSMUSG( | ENSMUSG( | 0.173141 | 0.897257 | NO | NO | NA |
| Gm50165 | ENSMUSG( | ENSMUSG( | 0.138094 | 0.908102 | NO | NO | NA |
| Pak5 | ENSMUSG( | ENSMUSG( | 0.216097 | 0.879096 | NO | NO | NA |
| Gm34267 | ENSMUSG( | ENSMUSG( | 0.775984 | 0.500004 | NO | NO | NA |
| Gm13464 | ENSMUSG( | ENSMUSG( | 0.630603 | 0.523035 | NO | NO | NA |
| Rps15-ps3 | ENSMUSG( | ENSMUSG( | 0.643956 | 0.40479 | NO | NO | NA |
| Gm52946 | ENSMUSG( | ENSMUSG( | 1.13291 | 0.364181 | NO | NO | NA |
| Hsd3b3 | ENSMUSG( | ENSMUSG( | 0.32754 | 0.770725 | NO | NO | NA |
| Gm35766 | ENSMUSG( | ENSMUSG( | 1.492464 | 0.198876 | NO | NO | NA |
| Syn1 | ENSMUSG( | ENSMUSG( | 0.055909 | 0.928272 | NO | NO | NA |
| Rln3 | ENSMUSG( | ENSMUSG( | -0.72615 | 0.231297 | NO | NO | NA |
| Gm37138 | ENSMUSG( | ENSMUSG( | 0.845145 | 0.305121 | NO | NO | NA |
| Gm42556 | ENSMUSG( | ENSMUSG( | 0.42093 | 0.726838 | NO | NO | NA |
| Nr5a2 | ENSMUSG( | ENSMUSG( | 0.738522 | 0.627163 | NO | NO | NA |

| | | | | | | | |
|---|---|---|---|---|---|---|---|
| Gm22009 | ENSMUSG( | ENSMUSG( | -0.56306 | 0.534972 | NO | NO | NA |
| Morc4 | ENSMUSG( | ENSMUSG( | 0.374263 | 0.728803 | NO | NO | NA |
| Trim69 | ENSMUSG( | ENSMUSG( | -0.63558 | 0.256616 | NO | NO | NA |
| Cdh12 | ENSMUSG( | ENSMUSG( | 1.325711 | 0.262692 | NO | NO | NA |
| Gm45338 | ENSMUSG( | ENSMUSG( | 0.241201 | 0.866416 | NO | NO | NA |
| 2510009E( | ENSMUSG( | ENSMUSG( | -0.13086 | 0.887324 | NO | NO | NA |
| 1520401A( | ENSMUSG( | ENSMUSG( | 0.609634 | 0.449198 | NO | NO | NA |
| Gm4598 | ENSMUSG( | ENSMUSG( | 1.944216 | 0.154778 | NO | NO | NA |
| Gm44562 | ENSMUSG( | ENSMUSG( | 0.248565 | 0.826657 | NO | NO | NA |
| Paupar | ENSMUSG( | ENSMUSG( | 0.345135 | 0.747848 | NO | NO | NA |
| Tie1 | ENSMUSG( | ENSMUSG( | 0.723437 | 0.572005 | NO | NO | NA |
| Gm31793 | ENSMUSG( | ENSMUSG( | 0.058923 | 0.897193 | NO | NO | NA |
| Olfr1318 | ENSMUSG( | ENSMUSG( | 0.874873 | 0.475513 | NO | NO | NA |
| Gm6296 | ENSMUSG( | ENSMUSG( | -0.43592 | 0.454063 | NO | NO | NA |
| Gm35850 | ENSMUSG( | ENSMUSG( | 0.672007 | 0.519025 | NO | NO | NA |
| Rps2-ps11 | ENSMUSG( | ENSMUSG( | -0.08407 | 0.907906 | NO | NO | NA |
| Crym | ENSMUSG( | ENSMUSG( | 0.953873 | 0.478415 | NO | NO | NA |
| Gm15637 | ENSMUSG( | ENSMUSG( | 1.06813 | 0.338636 | NO | NO | NA |
| Gm47676 | ENSMUSG( | ENSMUSG( | 0.622512 | 0.569784 | NO | NO | NA |
| Slfn4 | ENSMUSG( | ENSMUSG( | -0.02218 | 0.968726 | NO | NO | NA |
| Dlgap2 | ENSMUSG( | ENSMUSG( | 0.243703 | 0.829174 | NO | NO | NA |
| Gm42675 | ENSMUSG( | ENSMUSG( | 0.373873 | 0.746839 | NO | NO | NA |
| AA388235 | ENSMUSG( | ENSMUSG( | 0.061634 | 0.894638 | NO | NO | NA |
| Serinc2 | ENSMUSG( | ENSMUSG( | 3.551436 | NA | NO | NO | NA |
| Gm16144 | ENSMUSG( | ENSMUSG( | 0.171546 | 0.747769 | NO | NO | NA |
| Gm13841 | ENSMUSG( | ENSMUSG( | 0.399082 | 0.547825 | NO | NO | NA |
| Gm8249 | ENSMUSG( | ENSMUSG( | 0.755344 | 0.713303 | NO | NO | NA |
| Snai1 | ENSMUSG( | ENSMUSG( | 0.80087 | 0.187802 | NO | NO | NA |
| Gm48291 | ENSMUSG( | ENSMUSG( | 0.907589 | 0.334781 | NO | NO | NA |
| Cbln1 | ENSMUSG( | ENSMUSG( | 0.153454 | 0.888811 | NO | NO | NA |
| ENSMUSG | ENSMUSG( | ENSMUSG( | 0.522426 | 0.522159 | NO | NO | NA |
| Gm20275 | ENSMUSG( | ENSMUSG( | 0.254013 | 0.607595 | NO | NO | NA |
| Gm45360 | ENSMUSG( | ENSMUSG( | -0.99281 | 0.137848 | NO | NO | NA |
| B930059L( | ENSMUSG( | ENSMUSG( | -0.42605 | 0.471513 | NO | NO | NA |
| ENSMUSG | ENSMUSG( | ENSMUSG( | 0.364554 | 0.56241 | NO | NO | NA |
| Col5a2 | ENSMUSG( | ENSMUSG( | 0.053935 | 0.958234 | NO | NO | NA |
| Aox2 | ENSMUSG( | ENSMUSG( | 0.878316 | 0.156638 | NO | NO | NA |
| Gm45515 | ENSMUSG( | ENSMUSG( | -0.28821 | 0.47581 | NO | NO | NA |
| Bbs1 | ENSMUSG( | ENSMUSG( | -0.15614 | 0.759918 | NO | NO | NA |
| Gm29675 | ENSMUSG( | ENSMUSG( | -0.17406 | 0.722714 | NO | NO | NA |
| Ppp1r1c | ENSMUSG( | ENSMUSG( | 1.189478 | 0.304663 | NO | NO | NA |
| Fcna | ENSMUSG( | ENSMUSG( | -1.75591 | 0.014572 | YES | YES | Downregulated |
| Gm42648 | ENSMUSG( | ENSMUSG( | 0.049738 | 0.933701 | NO | NO | NA |
| Gm11510 | ENSMUSG( | ENSMUSG( | 0.488627 | 0.622431 | NO | NO | NA |
| Aire | ENSMUSG( | ENSMUSG( | -0.75186 | 0.452595 | NO | NO | NA |
| Gm49066 | ENSMUSG( | ENSMUSG( | 0.891391 | 0.397627 | NO | NO | NA |
| Mir5110 | ENSMUSG( | ENSMUSG( | 3.048153 | NA | NO | NO | NA |

| | | | | | | | |
|---|---|---|---|---|---|---|---|
| Col15a1 | ENSMUSG( | ENSMUSG( | 0.596986 | 0.436008 | NO | NO | NA |
| Gm45321 | ENSMUSG( | ENSMUSG( | 1.222521 | 0.344307 | NO | NO | NA |
| Ptgs2os | ENSMUSG( | ENSMUSG( | 0.650262 | 0.608703 | NO | NO | NA |
| Gpr158 | ENSMUSG( | ENSMUSG( | 0.997261 | 0.310463 | NO | NO | NA |
| Gm11942 | ENSMUSG( | ENSMUSG( | 0.683526 | 0.22233 | NO | NO | NA |
| Rskr | ENSMUSG( | ENSMUSG( | 0.091896 | 0.874972 | NO | NO | NA |
| Rpl31-ps1: | ENSMUSG( | ENSMUSG( | 0.037387 | 0.966706 | NO | NO | NA |
| Gm43592 | ENSMUSG( | ENSMUSG( | -0.15161 | 0.889961 | NO | NO | NA |
| Tead4 | ENSMUSG( | ENSMUSG( | -0.44403 | 0.638353 | NO | NO | NA |
| Dlgap3 | ENSMUSG( | ENSMUSG( | 0.55044 | 0.39815 | NO | NO | NA |
| Gm2848 | ENSMUSG( | ENSMUSG( | 1.244503 | 0.2583 | NO | NO | NA |
| Gm8190 | ENSMUSG( | ENSMUSG( | 1.119495 | 0.263914 | NO | NO | NA |
| Btnl6 | ENSMUSG( | ENSMUSG( | -0.24595 | 0.72269 | NO | NO | NA |
| Iqch | ENSMUSG( | ENSMUSG( | -0.37125 | 0.757619 | NO | NO | NA |
| Brdt | ENSMUSG( | ENSMUSG( | 0.220269 | 0.662185 | NO | NO | NA |
| G630022F: | ENSMUSG( | ENSMUSG( | 0.369923 | 0.689625 | NO | NO | NA |
| Gm9200 | ENSMUSG( | ENSMUSG( | 1.391406 | 0.191184 | NO | NO | NA |
| Gm37393 | ENSMUSG( | ENSMUSG( | 0.33378 | 0.769935 | NO | NO | NA |
| Gm43387 | ENSMUSG( | ENSMUSG( | 0.719598 | 0.457562 | NO | NO | NA |
| Pcsk1 | ENSMUSG( | ENSMUSG( | 0.71939 | 0.586184 | NO | NO | NA |
| Gm15708 | ENSMUSG( | ENSMUSG( | 0.469376 | 0.419605 | NO | NO | NA |
| Gm15510 | ENSMUSG( | ENSMUSG( | 1.210652 | 0.460937 | NO | NO | NA |
| Ighv10-3 | ENSMUSG( | ENSMUSG( | 4.576437 | NA | NO | NO | NA |
| Gm26861 | ENSMUSG( | ENSMUSG( | 0.42498 | 0.683941 | NO | NO | NA |
| Gm48190 | ENSMUSG( | ENSMUSG( | 0.944779 | 0.440252 | NO | NO | NA |
| Gm4247 | ENSMUSG( | ENSMUSG( | 0.452402 | 0.649957 | NO | NO | NA |
| Vmn1r200 | ENSMUSG( | ENSMUSG( | 0.520101 | 0.629327 | NO | NO | NA |
| Gm19212 | ENSMUSG( | ENSMUSG( | 0.29961 | 0.765499 | NO | NO | NA |
| Asprv1 | ENSMUSG( | ENSMUSG( | 0.72814 | 0.112413 | NO | NO | NA |
| Gm7370 | ENSMUSG( | ENSMUSG( | -0.58837 | 0.681348 | NO | NO | NA |
| Hus1b | ENSMUSG( | ENSMUSG( | -0.18648 | 0.800263 | NO | NO | NA |
| Dlx2 | ENSMUSG( | ENSMUSG( | 1.10958 | 0.339866 | NO | NO | NA |
| Gm50331 | ENSMUSG( | ENSMUSG( | 0.082682 | 0.914981 | NO | NO | NA |
| 4833412K: | ENSMUSG( | ENSMUSG( | 0.059082 | 0.943364 | NO | NO | NA |
| Gm47342 | ENSMUSG( | ENSMUSG( | 0.625412 | 0.672697 | NO | NO | NA |
| Gm43443 | ENSMUSG( | ENSMUSG( | 0.087718 | 0.940957 | NO | NO | NA |
| Gm39139 | ENSMUSG( | ENSMUSG( | 0.617709 | 0.561489 | NO | NO | NA |
| Heph | ENSMUSG( | ENSMUSG( | 0.458077 | 0.688848 | NO | NO | NA |
| Gm7816 | ENSMUSG( | ENSMUSG( | -0.41428 | 0.547864 | NO | NO | NA |
| Gm50358 | ENSMUSG( | ENSMUSG( | 0.463583 | 0.293237 | NO | NO | NA |
| Gm42421 | ENSMUSG( | ENSMUSG( | -0.15284 | 0.895376 | NO | NO | NA |
| A530083M | ENSMUSG( | ENSMUSG( | 0.523604 | 0.53462 | NO | NO | NA |
| Gm26588 | ENSMUSG( | ENSMUSG( | 0.21236 | 0.859542 | NO | NO | NA |
| Gm6974 | ENSMUSG( | ENSMUSG( | 0.168845 | 0.733562 | NO | NO | NA |
| Gm11491 | ENSMUSG( | ENSMUSG( | 0.278925 | 0.702923 | NO | NO | NA |
| Gm28988 | ENSMUSG( | ENSMUSG( | -0.42124 | 0.489778 | NO | NO | NA |
| 1700022I1 | ENSMUSG( | ENSMUSG( | 0.35049 | 0.607228 | NO | NO | NA |

| | | | | | | | |
|---|---|---|---|---|---|---|---|
| Gm16582 | ENSMUSG( | ENSMUSG( | 1.491747 | 0.099096 | NO | NO | NA |
| Gm47624 | ENSMUSG( | ENSMUSG( | -0.26197 | 0.861045 | NO | NO | NA |
| 5033417F: | ENSMUSG( | ENSMUSG( | -0.52258 | 0.430999 | NO | NO | NA |
| Gm47161 | ENSMUSG( | ENSMUSG( | 0.689573 | 0.563055 | NO | NO | NA |
| Gm14673 | ENSMUSG( | ENSMUSG( | 1.198195 | 0.309494 | NO | NO | NA |
| Gm44417 | ENSMUSG( | ENSMUSG( | 0.660286 | 0.544716 | NO | NO | NA |
| Gm35290 | ENSMUSG( | ENSMUSG( | -0.64042 | 0.295433 | NO | NO | NA |
| Armcx6 | ENSMUSG( | ENSMUSG( | -0.01996 | 0.96972 | NO | NO | NA |
| Vmn1r67 | ENSMUSG( | ENSMUSG( | 0.513229 | 0.693448 | NO | NO | NA |
| Gm12350 | ENSMUSG( | ENSMUSG( | -0.19442 | 0.800263 | NO | NO | NA |
| Gm48751 | ENSMUSG( | ENSMUSG( | 0.492961 | 0.679095 | NO | NO | NA |
| Gm10420 | ENSMUSG( | ENSMUSG( | -0.09663 | 0.917421 | NO | NO | NA |
| Zdhhc14 | ENSMUSG( | ENSMUSG( | 0.402159 | 0.635203 | NO | NO | NA |
| Gm13042 | ENSMUSG( | ENSMUSG( | -0.57699 | 0.56435 | NO | NO | NA |
| Gm48976 | ENSMUSG( | ENSMUSG( | 1.024471 | 0.369406 | NO | NO | NA |
| Ndst4 | ENSMUSG( | ENSMUSG( | 0.286357 | 0.77811 | NO | NO | NA |
| Omg | ENSMUSG( | ENSMUSG( | -0.10363 | 0.877421 | NO | NO | NA |
| Gm7598 | ENSMUSG( | ENSMUSG( | 0.020252 | 0.977199 | NO | NO | NA |
| Gm2395 | ENSMUSG( | ENSMUSG( | -0.54598 | 0.680543 | NO | NO | NA |
| Mmp28 | ENSMUSG( | ENSMUSG( | 0.256378 | 0.718217 | NO | NO | NA |
| Gm47641 | ENSMUSG( | ENSMUSG( | 1.192911 | 0.259143 | NO | NO | NA |
| Gm48742 | ENSMUSG( | ENSMUSG( | 0.302055 | 0.778559 | NO | NO | NA |
| Gm2594 | ENSMUSG( | ENSMUSG( | 0.257431 | 0.83101 | NO | NO | NA |
| Stard8 | ENSMUSG( | ENSMUSG( | 1.239501 | 0.030397 | YES | YES | Upregulated |
| ENSMUSG | ENSMUSG( | ENSMUSG( | 0.073231 | 0.901859 | NO | NO | NA |
| 4930546K( | ENSMUSG( | ENSMUSG( | 0.139878 | 0.888811 | NO | NO | NA |
| Gm12253 | ENSMUSG( | ENSMUSG( | -1.80692 | 0.053519 | NO | NO | NA |
| Gm28791 | ENSMUSG( | ENSMUSG( | -0.70524 | 0.302448 | NO | NO | NA |
| Gm42790 | ENSMUSG( | ENSMUSG( | 0.501978 | 0.645171 | NO | NO | NA |
| Aox4 | ENSMUSG( | ENSMUSG( | 0.309898 | 0.746663 | NO | NO | NA |
| Gm8539 | ENSMUSG( | ENSMUSG( | -0.25161 | 0.618809 | NO | NO | NA |
| Gm5444 | ENSMUSG( | ENSMUSG( | 0.68676 | 0.519484 | NO | NO | NA |
| Gm37729 | ENSMUSG( | ENSMUSG( | -0.16602 | 0.881144 | NO | NO | NA |
| Gm29237 | ENSMUSG( | ENSMUSG( | 0.376694 | 0.470839 | NO | NO | NA |
| Gm12059 | ENSMUSG( | ENSMUSG( | 0.493346 | 0.597141 | NO | NO | NA |
| Rps4x-ps | ENSMUSG( | ENSMUSG( | 0.322246 | 0.647827 | NO | NO | NA |
| Slc25a2 | ENSMUSG( | ENSMUSG( | -0.77119 | 0.048523 | YES | NO | NA |
| Gm14494 | ENSMUSG( | ENSMUSG( | 0.220021 | 0.740111 | NO | NO | NA |
| Btbd18 | ENSMUSG( | ENSMUSG( | 0.501296 | 0.396623 | NO | NO | NA |
| Enam | ENSMUSG( | ENSMUSG( | 0.859123 | 0.422059 | NO | NO | NA |
| Ebf3 | ENSMUSG( | ENSMUSG( | 0.501007 | 0.658393 | NO | NO | NA |
| Lep | ENSMUSG( | ENSMUSG( | 0.251591 | 0.832943 | NO | NO | NA |
| Gm13625 | ENSMUSG( | ENSMUSG( | -0.68442 | 0.796234 | NO | NO | NA |
| Igkv12-98 | ENSMUSG( | ENSMUSG( | 1.737566 | NA | NO | NO | NA |
| B430306N | ENSMUSG( | ENSMUSG( | -0.37446 | 0.640929 | NO | NO | NA |
| Il18 | ENSMUSG( | ENSMUSG( | 0.182954 | 0.683724 | NO | NO | NA |
| Gm11715 | ENSMUSG( | ENSMUSG( | 0.114797 | 0.930577 | NO | NO | NA |

| | | | | | | | |
|---|---|---|---|---|---|---|---|
| Muc2 | ENSMUSG( | ENSMUSG( | 0.130944 | 0.892196 | NO | NO | NA |
| Gm44432 | ENSMUSG( | ENSMUSG( | -0.9379 | 0.170848 | NO | NO | NA |
| Mkrn2os | ENSMUSG( | ENSMUSG( | 0.509675 | 0.518236 | NO | NO | NA |
| Gm8176 | ENSMUSG( | ENSMUSG( | 0.215723 | 0.847785 | NO | NO | NA |
| Cnnm1 | ENSMUSG( | ENSMUSG( | 0.420325 | 0.590846 | NO | NO | NA |
| 2700080J2 | ENSMUSG( | ENSMUSG( | 1.036781 | 0.367024 | NO | NO | NA |
| Gm17746 | ENSMUSG( | ENSMUSG( | 0.723599 | 0.48801 | NO | NO | NA |
| Gm37941 | ENSMUSG( | ENSMUSG( | 0.432533 | 0.70225 | NO | NO | NA |
| Otp | ENSMUSG( | ENSMUSG( | 0.051241 | 0.96972 | NO | NO | NA |
| Lcat | ENSMUSG( | ENSMUSG( | -0.12415 | 0.827824 | NO | NO | NA |
| Eif5al3-ps | ENSMUSG( | ENSMUSG( | 0.601107 | 0.363739 | NO | NO | NA |
| ENSMUSG | ENSMUSG( | ENSMUSG( | -0.32863 | 0.521197 | NO | NO | NA |
| Gm8520 | ENSMUSG( | ENSMUSG( | -0.0095 | 0.993785 | NO | NO | NA |
| Sim1 | ENSMUSG( | ENSMUSG( | 0.805424 | 0.515543 | NO | NO | NA |
| Serpina12 | ENSMUSG( | ENSMUSG( | 0.887955 | 0.347354 | NO | NO | NA |
| Igsf10 | ENSMUSG( | ENSMUSG( | 0.982856 | 0.130425 | NO | NO | NA |
| Pcsk2 | ENSMUSG( | ENSMUSG( | 0.578604 | 0.655776 | NO | NO | NA |
| 5730408A | ENSMUSG( | ENSMUSG( | 1.482312 | 0.194787 | NO | NO | NA |
| Cfap161 | ENSMUSG( | ENSMUSG( | 0.529063 | 0.63616 | NO | NO | NA |
| Gm50070 | ENSMUSG( | ENSMUSG( | -0.20809 | 0.81077 | NO | NO | NA |
| Gm9294 | ENSMUSG( | ENSMUSG( | 0.382413 | 0.461948 | NO | NO | NA |
| Gm21399 | ENSMUSG( | ENSMUSG( | 0.27531 | 0.616241 | NO | NO | NA |
| Gm26797 | ENSMUSG( | ENSMUSG( | 0.109733 | 0.919733 | NO | NO | NA |
| Gm43689 | ENSMUSG( | ENSMUSG( | 3.282272 | 0.017067 | YES | YES | Upregulated |
| ENSMUSG | ENSMUSG( | ENSMUSG( | 0.135672 | 0.778722 | NO | NO | NA |
| Gm42984 | ENSMUSG( | ENSMUSG( | -0.30771 | 0.550072 | NO | NO | NA |
| Gm4545 | ENSMUSG( | ENSMUSG( | 0.089584 | 0.972271 | NO | NO | NA |
| ENSMUSG | ENSMUSG( | ENSMUSG( | 0.433565 | 0.731835 | NO | NO | NA |
| Kcnab3 | ENSMUSG( | ENSMUSG( | -0.63774 | 0.236987 | NO | NO | NA |
| Chat | ENSMUSG( | ENSMUSG( | 0.129773 | 0.926387 | NO | NO | NA |
| Slc17a3 | ENSMUSG( | ENSMUSG( | 0.268944 | 0.819978 | NO | NO | NA |
| Gm9316 | ENSMUSG( | ENSMUSG( | -0.46167 | 0.480526 | NO | NO | NA |
| Skint4 | ENSMUSG( | ENSMUSG( | 0.768891 | 0.529107 | NO | NO | NA |
| ENSMUSG( | ENSMUSG( | ENSMUSG( | 1.199917 | 0.207706 | NO | NO | NA |
| Olfr859 | ENSMUSG( | ENSMUSG( | -0.07168 | 0.963848 | NO | NO | NA |
| Gm45591 | ENSMUSG( | ENSMUSG( | -0.75619 | 0.351004 | NO | NO | NA |
| Gm13899 | ENSMUSG( | ENSMUSG( | 0.683206 | 0.548323 | NO | NO | NA |
| ENSMUSG | ENSMUSG( | ENSMUSG( | -0.83911 | 0.065886 | NO | NO | NA |
| Gm15726 | ENSMUSG( | ENSMUSG( | 1.210519 | 0.016464 | YES | YES | Upregulated |
| Gm42751 | ENSMUSG( | ENSMUSG( | 0.879859 | 0.521422 | NO | NO | NA |
| ENSMUSG( | ENSMUSG( | ENSMUSG( | 0.63111 | 0.627 | NO | NO | NA |
| Smok3c | ENSMUSG( | ENSMUSG( | 0.076203 | 0.958652 | NO | NO | NA |
| Gm8797 | ENSMUSG( | ENSMUSG( | -0.33172 | 0.520881 | NO | NO | NA |
| Gm20589 | ENSMUSG( | ENSMUSG( | 1.109528 | 0.289575 | NO | NO | NA |
| Rpl21-ps8 | ENSMUSG( | ENSMUSG( | 0.404665 | 0.474503 | NO | NO | NA |
| Adora3 | ENSMUSG( | ENSMUSG( | 0.299383 | 0.553137 | NO | NO | NA |
| Gm5844 | ENSMUSG( | ENSMUSG( | -0.09324 | 0.853406 | NO | NO | NA |

| | | | | | | | |
|---|---|---|---|---|---|---|---|
| ENSMUSG( | ENSMUSG( | ENSMUSG( | 0.839063 | 0.423578 | NO | NO | NA |
| Vmn2r-ps1 | ENSMUSG( | ENSMUSG( | 0.224868 | 0.840863 | NO | NO | NA |
| ENSMUSG | ENSMUSG( | ENSMUSG( | 0.407228 | 0.481636 | NO | NO | NA |
| Gm37868 | ENSMUSG( | ENSMUSG( | -0.13141 | 0.889961 | NO | NO | NA |
| Gm42717 | ENSMUSG( | ENSMUSG( | 0.151136 | 0.814718 | NO | NO | NA |
| Grm8 | ENSMUSG( | ENSMUSG( | 0.180758 | 0.867057 | NO | NO | NA |
| Gm15411 | ENSMUSG( | ENSMUSG( | 0.530067 | 0.394041 | NO | NO | NA |
| Vmn1r226 | ENSMUSG( | ENSMUSG( | 0.220718 | 0.85863 | NO | NO | NA |
| Platr27 | ENSMUSG( | ENSMUSG( | 0.476732 | 0.467096 | NO | NO | NA |
| Gm6546 | ENSMUSG( | ENSMUSG( | 0.585646 | 0.63885 | NO | NO | NA |
| Gm18284 | ENSMUSG( | ENSMUSG( | 0.563211 | 0.439866 | NO | NO | NA |
| Gm47905 | ENSMUSG( | ENSMUSG( | 0.805841 | 0.571967 | NO | NO | NA |
| Olfr629 | ENSMUSG( | ENSMUSG( | 0.719515 | 0.527421 | NO | NO | NA |
| Gm48966 | ENSMUSG( | ENSMUSG( | 0.585603 | 0.513234 | NO | NO | NA |
| Chrnb3 | ENSMUSG( | ENSMUSG( | 0.528837 | 0.655215 | NO | NO | NA |
| Dok7 | ENSMUSG( | ENSMUSG( | -0.37928 | 0.573931 | NO | NO | NA |
| Gm9006 | ENSMUSG( | ENSMUSG( | 0.90922 | 0.539797 | NO | NO | NA |
| Fbn1 | ENSMUSG( | ENSMUSG( | 0.491741 | 0.607454 | NO | NO | NA |
| 7SK ENSM | ENSMUSG( | ENSMUSG( | -0.51903 | 0.396816 | NO | NO | NA |
| Stab2 | ENSMUSG( | ENSMUSG( | -0.39252 | 0.601687 | NO | NO | NA |
| Gm43466 | ENSMUSG( | ENSMUSG( | -0.18523 | 0.766791 | NO | NO | NA |
| Gm14418 | ENSMUSG( | ENSMUSG( | 0.935782 | 0.025468 | YES | NO | NA |
| F730311O | ENSMUSG( | ENSMUSG( | -0.40898 | 0.462511 | NO | NO | NA |
| ENSMUSG | ENSMUSG( | ENSMUSG( | 0.939863 | 0.272085 | NO | NO | NA |
| Gm11650 | ENSMUSG( | ENSMUSG( | 0.625466 | 0.596956 | NO | NO | NA |
| Flt4 | ENSMUSG( | ENSMUSG( | 0.647317 | 0.557764 | NO | NO | NA |
| Gm19938 | ENSMUSG( | ENSMUSG( | 0.629086 | 0.561491 | NO | NO | NA |
| ENSMUSG | ENSMUSG( | ENSMUSG( | 1.308774 | 0.239999 | NO | NO | NA |
| Nid1 | ENSMUSG( | ENSMUSG( | 0.632399 | 0.345714 | NO | NO | NA |
| Gm42680 | ENSMUSG( | ENSMUSG( | 0.958292 | 0.466523 | NO | NO | NA |
| ENSMUSG | ENSMUSG( | ENSMUSG( | 0.481911 | 0.430489 | NO | NO | NA |
| Mpzl2 | ENSMUSG( | ENSMUSG( | 0.481032 | 0.640892 | NO | NO | NA |
| Alk | ENSMUSG( | ENSMUSG( | 0.476979 | 0.654862 | NO | NO | NA |
| Gm33699 | ENSMUSG( | ENSMUSG( | 0.963698 | 0.491865 | NO | NO | NA |
| ENSMUSG | ENSMUSG( | ENSMUSG( | -0.44603 | 0.41964 | NO | NO | NA |
| Gm7418 | ENSMUSG( | ENSMUSG( | 0.282944 | 0.579897 | NO | NO | NA |
| Vmn2r45 | ENSMUSG( | ENSMUSG( | 0.876951 | 0.402464 | NO | NO | NA |
| Gm29488 | ENSMUSG( | ENSMUSG( | -0.1524 | 0.765455 | NO | NO | NA |
| Samd12 | ENSMUSG( | ENSMUSG( | 0.29232 | 0.758216 | NO | NO | NA |
| Tbc1d22bc | ENSMUSG( | ENSMUSG( | -0.2579 | 0.789235 | NO | NO | NA |
| 5330426L2 | ENSMUSG( | ENSMUSG( | 0.015385 | 0.982039 | NO | NO | NA |
| Fibcd1 | ENSMUSG( | ENSMUSG( | 0.687618 | 0.347354 | NO | NO | NA |
| Gm16224 | ENSMUSG( | ENSMUSG( | -0.301 | 0.636419 | NO | NO | NA |
| Gm42446 | ENSMUSG( | ENSMUSG( | 0.659311 | 0.610381 | NO | NO | NA |
| Csdc2 | ENSMUSG( | ENSMUSG( | 0.105619 | 0.839038 | NO | NO | NA |
| Gm11634 | ENSMUSG( | ENSMUSG( | 0.403762 | 0.692687 | NO | NO | NA |
| Gm12781 | ENSMUSG( | ENSMUSG( | 0.482281 | 0.406631 | NO | NO | NA |

| | | | | | | | |
|---|---|---|---|---|---|---|---|
| Cr2 | ENSMUSG( | ENSMUSG( | 0.407655 | 0.615733 | NO | NO | NA |
| Pcsk2os1 | ENSMUSG( | ENSMUSG( | 1.106859 | 0.293315 | NO | NO | NA |
| Gm15976 | ENSMUSG( | ENSMUSG( | 1.757231 | 0.19877 | NO | NO | NA |
| Gm49732 | ENSMUSG( | ENSMUSG( | 0.813496 | 0.473525 | NO | NO | NA |
| Gm15489 | ENSMUSG( | ENSMUSG( | -0.2554 | 0.637771 | NO | NO | NA |
| Cnksr2 | ENSMUSG( | ENSMUSG( | 0.703828 | 0.601079 | NO | NO | NA |
| Gm15594 | ENSMUSG( | ENSMUSG( | 0.260443 | 0.80397 | NO | NO | NA |
| Neto1 | ENSMUSG( | ENSMUSG( | 0.748442 | 0.586177 | NO | NO | NA |
| Gm10128 | ENSMUSG( | ENSMUSG( | 0.523085 | 0.722103 | NO | NO | NA |
| Gm27240 | ENSMUSG( | ENSMUSG( | -0.43838 | 0.508112 | NO | NO | NA |
| ENSMUSG | ENSMUSG( | ENSMUSG( | -0.1511 | 0.77861 | NO | NO | NA |
| Rhbdf1 | ENSMUSG( | ENSMUSG( | 0.302888 | 0.672786 | NO | NO | NA |
| Gm2832 | ENSMUSG( | ENSMUSG( | 0.638218 | 0.615733 | NO | NO | NA |
| Gm13126 | ENSMUSG( | ENSMUSG( | 0.743149 | 0.558429 | NO | NO | NA |
| ENSMUSG( | ENSMUSG( | ENSMUSG( | 0.257365 | 0.664861 | NO | NO | NA |
| Pdpn | ENSMUSG( | ENSMUSG( | 0.099523 | 0.940462 | NO | NO | NA |
| Gm37749 | ENSMUSG( | ENSMUSG( | 0.095378 | 0.942605 | NO | NO | NA |
| Met | ENSMUSG( | ENSMUSG( | 0.43107 | 0.568223 | NO | NO | NA |
| Gm9949 | ENSMUSG( | ENSMUSG( | -0.45716 | 0.438825 | NO | NO | NA |
| Spanxn4 | ENSMUSG( | ENSMUSG( | 1.015346 | 0.396346 | NO | NO | NA |
| Gm37877 | ENSMUSG( | ENSMUSG( | 0.85205 | 0.520905 | NO | NO | NA |
| ENSMUSG | ENSMUSG( | ENSMUSG( | 0.988383 | 0.430091 | NO | NO | NA |
| Gm42856 | ENSMUSG( | ENSMUSG( | 0.094436 | 0.872207 | NO | NO | NA |
| 6330420H | ENSMUSG( | ENSMUSG( | 0.687372 | 0.609714 | NO | NO | NA |
| ENSMUSG | ENSMUSG( | ENSMUSG( | 0.730822 | 0.279164 | NO | NO | NA |
| Nmbr | ENSMUSG( | ENSMUSG( | 0.360806 | 0.787948 | NO | NO | NA |
| Itga9 | ENSMUSG( | ENSMUSG( | 0.785474 | 0.205619 | NO | NO | NA |
| Gm14912 | ENSMUSG( | ENSMUSG( | 0.921552 | 0.530329 | NO | NO | NA |
| Vmn1r221 | ENSMUSG( | ENSMUSG( | 0.921157 | 0.47696 | NO | NO | NA |
| 1700067K( | ENSMUSG( | ENSMUSG( | -0.0814 | 0.889318 | NO | NO | NA |
| Cd59b | ENSMUSG( | ENSMUSG( | 0.424936 | 0.610556 | NO | NO | NA |
| Boc | ENSMUSG( | ENSMUSG( | 0.014339 | 0.981038 | NO | NO | NA |
| Gm27241 | ENSMUSG( | ENSMUSG( | 0.046444 | 0.964701 | NO | NO | NA |
| Nodal | ENSMUSG( | ENSMUSG( | -1.01835 | 0.063378 | NO | NO | NA |
| 5033425B( | ENSMUSG( | ENSMUSG( | 0.677434 | 0.515032 | NO | NO | NA |
| Gm49620 | ENSMUSG( | ENSMUSG( | -0.13008 | 0.916437 | NO | NO | NA |
| Gm49801 | ENSMUSG( | ENSMUSG( | 0.906778 | 0.4904 | NO | NO | NA |
| ENSMUSG | ENSMUSG( | ENSMUSG( | 1.196464 | 0.113663 | NO | NO | NA |
| Zfp957 | ENSMUSG( | ENSMUSG( | 0.434145 | 0.684648 | NO | NO | NA |
| Gm48754 | ENSMUSG( | ENSMUSG( | 0.06127 | 0.931063 | NO | NO | NA |
| Gm6210 | ENSMUSG( | ENSMUSG( | -0.05896 | 0.901638 | NO | NO | NA |
| B3galt2 | ENSMUSG( | ENSMUSG( | 0.078072 | 0.900004 | NO | NO | NA |
| Gm44037 | ENSMUSG( | ENSMUSG( | 0.273796 | 0.638583 | NO | NO | NA |
| Gm9311 | ENSMUSG( | ENSMUSG( | -0.75147 | 0.1307 | NO | NO | NA |
| Gm31219 | ENSMUSG( | ENSMUSG( | 0.539455 | 0.600536 | NO | NO | NA |
| Olfr1418 | ENSMUSG( | ENSMUSG( | 0.104004 | 0.918895 | NO | NO | NA |
| Gm38104 | ENSMUSG( | ENSMUSG( | 0.376726 | 0.583984 | NO | NO | NA |

| | | | | | | | |
|---|---|---|---|---|---|---|---|
| Lta | ENSMUSG( | ENSMUSG( | 0.396972 | 0.545553 | NO | NO | NA |
| Gm28447 | ENSMUSG( | ENSMUSG( | 0.594557 | 0.644442 | NO | NO | NA |
| Ms4a1 | ENSMUSG( | ENSMUSG( | 0.598893 | 0.547317 | NO | NO | NA |
| Gm26644 | ENSMUSG( | ENSMUSG( | 0.706331 | 0.528583 | NO | NO | NA |
| Zfp449 | ENSMUSG( | ENSMUSG( | 0.394671 | 0.49287 | NO | NO | NA |
| Pcdhga11 | ENSMUSG( | ENSMUSG( | -0.91142 | 0.099093 | NO | NO | NA |
| Kcnh1 | ENSMUSG( | ENSMUSG( | 0.860835 | 0.45817 | NO | NO | NA |
| ENSMUSG | ENSMUSG( | ENSMUSG( | 0.335661 | 0.449488 | NO | NO | NA |
| Jph3 | ENSMUSG( | ENSMUSG( | -0.20077 | 0.751869 | NO | NO | NA |
| Gm10668 | ENSMUSG( | ENSMUSG( | 0.053978 | 0.982586 | NO | NO | NA |
| Gm43712 | ENSMUSG( | ENSMUSG( | 0.713732 | 0.261819 | NO | NO | NA |
| 1110028F: | ENSMUSG( | ENSMUSG( | 0.632425 | 0.508329 | NO | NO | NA |
| Gm45762 | ENSMUSG( | ENSMUSG( | 0.225405 | 0.725782 | NO | NO | NA |
| BC051537 | ENSMUSG( | ENSMUSG( | 0.392434 | 0.724365 | NO | NO | NA |
| Tbx2 | ENSMUSG( | ENSMUSG( | 0.885949 | 0.064038 | NO | NO | NA |
| Cyp1b1 | ENSMUSG( | ENSMUSG( | 0.36281 | 0.735748 | NO | NO | NA |
| Vmn1r38 | ENSMUSG( | ENSMUSG( | 0.851509 | 0.397165 | NO | NO | NA |
| Gm19016 | ENSMUSG( | ENSMUSG( | 0.617833 | 0.564995 | NO | NO | NA |
| Pramel4 | ENSMUSG( | ENSMUSG( | -0.19217 | 0.838501 | NO | NO | NA |
| Gm8518 | ENSMUSG( | ENSMUSG( | 0.351581 | 0.470978 | NO | NO | NA |
| Gm37115 | ENSMUSG( | ENSMUSG( | 0.556004 | 0.604262 | NO | NO | NA |
| Igkv9-129 | ENSMUSG( | ENSMUSG( | -4.96841 | NA | NO | NO | NA |
| Tubb4b-ps | ENSMUSG( | ENSMUSG( | 0.526903 | 0.280093 | NO | NO | NA |
| Btc | ENSMUSG( | ENSMUSG( | 0.760602 | 0.469728 | NO | NO | NA |
| Gm32647 | ENSMUSG( | ENSMUSG( | 0.9928 | 0.346054 | NO | NO | NA |
| Gm19689 | ENSMUSG( | ENSMUSG( | 0.433437 | 0.647539 | NO | NO | NA |
| Olfr720 | ENSMUSG( | ENSMUSG( | 0.827596 | 0.543583 | NO | NO | NA |
| Gm50218 | ENSMUSG( | ENSMUSG( | -1.10536 | 0.025442 | YES | YES | Downregulated |
| Sphkap | ENSMUSG( | ENSMUSG( | 0.858592 | 0.561491 | NO | NO | NA |
| Gm46409 | ENSMUSG( | ENSMUSG( | 0.034685 | 0.977005 | NO | NO | NA |
| Gm13497 | ENSMUSG( | ENSMUSG( | 0.608798 | 0.600045 | NO | NO | NA |
| 5330406N | ENSMUSG( | ENSMUSG( | 0.180606 | 0.741426 | NO | NO | NA |
| Tmem100 | ENSMUSG( | ENSMUSG( | 0.651104 | 0.522576 | NO | NO | NA |
| ENSMUSG( | ENSMUSG( | ENSMUSG( | -1.14167 | 0.06537 | NO | NO | NA |
| Gm12176 | ENSMUSG( | ENSMUSG( | 0.655393 | 0.515346 | NO | NO | NA |
| Gm49594 | ENSMUSG( | ENSMUSG( | 0.302202 | 0.888811 | NO | NO | NA |
| Mxra7 | ENSMUSG( | ENSMUSG( | 0.862481 | 0.351949 | NO | NO | NA |
| Ighv1-63 | ENSMUSG( | ENSMUSG( | -5.33834 | 0.080192 | NO | NO | NA |
| ENSMUSG | ENSMUSG( | ENSMUSG( | -0.1333 | 0.771401 | NO | NO | NA |
| Gm45684 | ENSMUSG( | ENSMUSG( | -0.08386 | 0.902186 | NO | NO | NA |
| Gli2 | ENSMUSG( | ENSMUSG( | 0.635292 | 0.61797 | NO | NO | NA |
| Gm37979 | ENSMUSG( | ENSMUSG( | -0.29525 | 0.633271 | NO | NO | NA |
| Gm7594 | ENSMUSG( | ENSMUSG( | 0.707389 | 0.517274 | NO | NO | NA |
| Gm9060 | ENSMUSG( | ENSMUSG( | 0.534896 | 0.616523 | NO | NO | NA |
| Slco4c1 | ENSMUSG( | ENSMUSG( | 0.41311 | 0.566439 | NO | NO | NA |
| Gm16589 | ENSMUSG( | ENSMUSG( | -0.31107 | 0.626178 | NO | NO | NA |
| Gm43183 | ENSMUSG( | ENSMUSG( | 0.524948 | 0.715024 | NO | NO | NA |

| | | | | | | | |
|---|---|---|---|---|---|---|---|
| Rpsa-ps9 | ENSMUSG( | ENSMUSG( | 0.057948 | 0.921903 | NO | NO | NA |
| Gm48837 | ENSMUSG( | ENSMUSG( | 0.609534 | 0.375678 | NO | NO | NA |
| Gm47248 | ENSMUSG( | ENSMUSG( | -1.14432 | 0.1153 | NO | NO | NA |
| Gm18560 | ENSMUSG( | ENSMUSG( | 1.134115 | 0.32348 | NO | NO | NA |
| Rpl13a-ps: | ENSMUSG( | ENSMUSG( | 0.33055 | 0.52971 | NO | NO | NA |
| 4930594A( | ENSMUSG( | ENSMUSG( | 1.55254 | 0.280628 | NO | NO | NA |
| 9130604C: | ENSMUSG( | ENSMUSG( | 0.47561 | 0.318492 | NO | NO | NA |
| Ighv1-4 | ENSMUSG( | ENSMUSG( | -3.92303 | NA | NO | NO | NA |
| Tm4sf5 | ENSMUSG( | ENSMUSG( | 0.086156 | 0.894959 | NO | NO | NA |
| Gm50210 | ENSMUSG( | ENSMUSG( | -0.1641 | 0.790471 | NO | NO | NA |
| Gm45643 | ENSMUSG( | ENSMUSG( | 0.713494 | 0.589301 | NO | NO | NA |
| Nrap | ENSMUSG( | ENSMUSG( | 0.591406 | 0.410148 | NO | NO | NA |
| Klra1 | ENSMUSG( | ENSMUSG( | 0.227759 | 0.813047 | NO | NO | NA |
| Gm8024 | ENSMUSG( | ENSMUSG( | 0.842143 | 0.526869 | NO | NO | NA |
| Ighv1-58 | ENSMUSG( | ENSMUSG( | 0.113396 | 0.966814 | NO | NO | NA |
| Olfr830 | ENSMUSG( | ENSMUSG( | 0.465204 | 0.736358 | NO | NO | NA |
| Ighv1-78 | ENSMUSG( | ENSMUSG( | -5.82754 | 0.020717 | YES | YES | Downregulated |
| Trpc4 | ENSMUSG( | ENSMUSG( | 0.295435 | 0.847701 | NO | NO | NA |
| Cacna1h | ENSMUSG( | ENSMUSG( | -0.38522 | 0.691861 | NO | NO | NA |
| Gm3234 | ENSMUSG( | ENSMUSG( | 0.577239 | 0.655006 | NO | NO | NA |
| ENSMUSG | ENSMUSG( | ENSMUSG( | 0.946716 | 0.456373 | NO | NO | NA |
| L3hypdh | ENSMUSG( | ENSMUSG( | 0.628923 | 0.468146 | NO | NO | NA |
| Gm7879 | ENSMUSG( | ENSMUSG( | -0.09661 | 0.885836 | NO | NO | NA |
| Vmn1r227 | ENSMUSG( | ENSMUSG( | 0.706531 | 0.520825 | NO | NO | NA |
| Ehd2 | ENSMUSG( | ENSMUSG( | 0.412712 | 0.53278 | NO | NO | NA |
| Gm49844 | ENSMUSG( | ENSMUSG( | 1.46858 | 0.178773 | NO | NO | NA |
| Pcdhb19 | ENSMUSG( | ENSMUSG( | 0.775111 | 0.469209 | NO | NO | NA |
| Gas1 | ENSMUSG( | ENSMUSG( | 0.483519 | 0.612079 | NO | NO | NA |
| Gm35569 | ENSMUSG( | ENSMUSG( | 0.354534 | 0.758872 | NO | NO | NA |
| Pisd-ps2 | ENSMUSG( | ENSMUSG( | -0.42971 | 0.442629 | NO | NO | NA |
| Gm16141 | ENSMUSG( | ENSMUSG( | 0.574639 | 0.631515 | NO | NO | NA |
| Cyp2d22 | ENSMUSG( | ENSMUSG( | -0.28775 | 0.802278 | NO | NO | NA |
| Ptger3 | ENSMUSG( | ENSMUSG( | 0.997593 | 0.247517 | NO | NO | NA |
| Ift43 | ENSMUSG( | ENSMUSG( | 0.106655 | 0.843187 | NO | NO | NA |
| Gm35371 | ENSMUSG( | ENSMUSG( | 0.72241 | 0.664907 | NO | NO | NA |
| Gm8153 | ENSMUSG( | ENSMUSG( | -0.36263 | 0.680393 | NO | NO | NA |
| Car3 | ENSMUSG( | ENSMUSG( | 0.007217 | 0.992393 | NO | NO | NA |
| Prom1 | ENSMUSG( | ENSMUSG( | 1.139721 | 0.093668 | NO | NO | NA |
| Fhad1 | ENSMUSG( | ENSMUSG( | 0.067913 | 0.946702 | NO | NO | NA |
| Bcas1 | ENSMUSG( | ENSMUSG( | 0.525229 | 0.408636 | NO | NO | NA |
| Cyp26b1 | ENSMUSG( | ENSMUSG( | 0.712836 | 0.542287 | NO | NO | NA |
| Ssc5d | ENSMUSG( | ENSMUSG( | -0.03577 | 0.955431 | NO | NO | NA |
| Calml4 | ENSMUSG( | ENSMUSG( | 0.174943 | 0.887324 | NO | NO | NA |
| Gm14900 | ENSMUSG( | ENSMUSG( | 0.556427 | 0.607228 | NO | NO | NA |
| Zfp457 | ENSMUSG( | ENSMUSG( | 0.686551 | 0.280014 | NO | NO | NA |
| Gm15421 | ENSMUSG( | ENSMUSG( | 0.110478 | 0.872677 | NO | NO | NA |
| Gm43200 | ENSMUSG( | ENSMUSG( | -0.72862 | 0.198705 | NO | NO | NA |

| | | | | | | | |
|---|---|---|---|---|---|---|---|
| Gm42596 | ENSMUSG( | ENSMUSG( | -0.52927 | 0.586288 | NO | NO | NA |
| Gm44680 | ENSMUSG( | ENSMUSG( | 0.486346 | 0.653409 | NO | NO | NA |
| Gm48353 | ENSMUSG( | ENSMUSG( | 1.019132 | 0.404157 | NO | NO | NA |
| Epha8 | ENSMUSG( | ENSMUSG( | 1.426229 | 0.074389 | NO | NO | NA |
| Gm42937 | ENSMUSG( | ENSMUSG( | 0.005578 | 0.990716 | NO | NO | NA |
| Igkv4-61 | ENSMUSG( | ENSMUSG( | 3.599692 | NA | NO | NO | NA |
| Gm42560 | ENSMUSG( | ENSMUSG( | 0.76659 | 0.513114 | NO | NO | NA |
| Nkx2-3 | ENSMUSG( | ENSMUSG( | 1.140804 | 0.313303 | NO | NO | NA |
| Gm49486 | ENSMUSG( | ENSMUSG( | 0.621563 | 0.5325 | NO | NO | NA |
| Slc7a2 | ENSMUSG( | ENSMUSG( | 0.452473 | 0.63616 | NO | NO | NA |
| Gm43759 | ENSMUSG( | ENSMUSG( | -0.38369 | 0.502364 | NO | NO | NA |
| Gm6430 | ENSMUSG( | ENSMUSG( | 0.336846 | 0.595991 | NO | NO | NA |
| Adamts15 | ENSMUSG( | ENSMUSG( | -0.05035 | 0.964479 | NO | NO | NA |
| Olfr1565 | ENSMUSG( | ENSMUSG( | 0.103573 | 0.923116 | NO | NO | NA |
| ENSMUSG | ENSMUSG( | ENSMUSG( | -0.60789 | 0.493007 | NO | NO | NA |
| Gm5468 | ENSMUSG( | ENSMUSG( | 0.614593 | 0.552734 | NO | NO | NA |
| Gm45440 | ENSMUSG( | ENSMUSG( | 0.726077 | 0.537122 | NO | NO | NA |
| Gstp-ps | ENSMUSG( | ENSMUSG( | -0.14306 | 0.794371 | NO | NO | NA |
| Rnd2 | ENSMUSG( | ENSMUSG( | 0.387633 | 0.534067 | NO | NO | NA |
| Col2a1 | ENSMUSG( | ENSMUSG( | 0.564081 | 0.495601 | NO | NO | NA |
| Gm4833 | ENSMUSG( | ENSMUSG( | 0.240125 | 0.706898 | NO | NO | NA |
| Chn1 | ENSMUSG( | ENSMUSG( | 0.320117 | 0.601687 | NO | NO | NA |
| Gm13274 | ENSMUSG( | ENSMUSG( | 0.460983 | 0.611373 | NO | NO | NA |
| Tnfsf11 | ENSMUSG( | ENSMUSG( | 1.323581 | 0.05387 | NO | NO | NA |
| Gm42676 | ENSMUSG( | ENSMUSG( | 0.915708 | 0.518697 | NO | NO | NA |
| Klk11 | ENSMUSG( | ENSMUSG( | 1.438022 | 0.10351 | NO | NO | NA |
| Gm8172 | ENSMUSG( | ENSMUSG( | 0.982522 | 0.117937 | NO | NO | NA |
| ENSMUSG | ENSMUSG( | ENSMUSG( | 1.063 | 0.354296 | NO | NO | NA |
| Baiap2l1 | ENSMUSG( | ENSMUSG( | -0.40632 | 0.611454 | NO | NO | NA |
| Gm15471 | ENSMUSG( | ENSMUSG( | 1.474651 | 0.222366 | NO | NO | NA |
| Gm5424 | ENSMUSG( | ENSMUSG( | 0.268341 | 0.57346 | NO | NO | NA |
| Yjefn3 | ENSMUSG( | ENSMUSG( | 0.150698 | 0.829991 | NO | NO | NA |
| A230083G | ENSMUSG( | ENSMUSG( | 0.286113 | 0.600449 | NO | NO | NA |
| Gm9194 | ENSMUSG( | ENSMUSG( | 0.728152 | 0.554054 | NO | NO | NA |
| Triqk | ENSMUSG( | ENSMUSG( | 0.617769 | 0.56336 | NO | NO | NA |
| Gm15892 | ENSMUSG( | ENSMUSG( | 0.61278 | 0.449198 | NO | NO | NA |
| Fbxw19 | ENSMUSG( | ENSMUSG( | 0.560361 | 0.689584 | NO | NO | NA |
| 1700120G | ENSMUSG( | ENSMUSG( | 1.045464 | 0.289398 | NO | NO | NA |
| Spaca9 | ENSMUSG( | ENSMUSG( | -0.94684 | 0.129141 | NO | NO | NA |
| Dnah10 | ENSMUSG( | ENSMUSG( | -0.32379 | 0.618809 | NO | NO | NA |
| Matr3-ps2 | ENSMUSG( | ENSMUSG( | 0.728536 | 0.532211 | NO | NO | NA |
| Gm32063 | ENSMUSG( | ENSMUSG( | 1.698264 | 0.177506 | NO | NO | NA |
| Wfdc13 | ENSMUSG( | ENSMUSG( | 1.703428 | 0.247737 | NO | NO | NA |
| Vmn1r89 | ENSMUSG( | ENSMUSG( | 0.861705 | 0.505024 | NO | NO | NA |
| Gm43761 | ENSMUSG( | ENSMUSG( | 0.911619 | 0.324777 | NO | NO | NA |
| Gm9207 | ENSMUSG( | ENSMUSG( | -0.79227 | 0.159538 | NO | NO | NA |
| Glt8d2 | ENSMUSG( | ENSMUSG( | -0.32539 | 0.690509 | NO | NO | NA |

| | | | | | | | |
|---|---|---|---|---|---|---|---|
| Gm43571 | ENSMUSG( | ENSMUSG( | -0.17214 | 0.851225 | NO | NO | NA |
| Gm19522 | ENSMUSG( | ENSMUSG( | 0.104672 | 0.919248 | NO | NO | NA |
| Gm18717 | ENSMUSG( | ENSMUSG( | 0.425922 | 0.677554 | NO | NO | NA |
| Gm12846 | ENSMUSG( | ENSMUSG( | 0.366076 | 0.755226 | NO | NO | NA |
| 9430007M | ENSMUSG( | ENSMUSG( | 1.459889 | 0.223204 | NO | NO | NA |
| Ell3 | ENSMUSG( | ENSMUSG( | -0.42842 | 0.435459 | NO | NO | NA |
| Gm45073 | ENSMUSG( | ENSMUSG( | 0.925982 | 0.306823 | NO | NO | NA |
| Gm7061 | ENSMUSG( | ENSMUSG( | -0.79562 | 0.080785 | NO | NO | NA |
| Gm43062 | ENSMUSG( | ENSMUSG( | -0.3045 | 0.639778 | NO | NO | NA |
| Gm43831 | ENSMUSG( | ENSMUSG( | 0.669376 | 0.561782 | NO | NO | NA |
| Rps2-ps9 | ENSMUSG( | ENSMUSG( | -0.06894 | 0.941616 | NO | NO | NA |
| 4930444A | ENSMUSG( | ENSMUSG( | -1.22197 | 0.010211 | YES | YES | Downregulated |
| Nr0b2 | ENSMUSG( | ENSMUSG( | 0.110081 | 0.946872 | NO | NO | NA |
| Zfp521 | ENSMUSG( | ENSMUSG( | 0.735908 | 0.522182 | NO | NO | NA |
| Gm11889 | ENSMUSG( | ENSMUSG( | 0.033469 | 0.954655 | NO | NO | NA |
| Gvin-ps5 | ENSMUSG( | ENSMUSG( | -1.06401 | 0.150037 | NO | NO | NA |
| Gucy2d | ENSMUSG( | ENSMUSG( | 3.572511 | NA | NO | NO | NA |
| Gm43366 | ENSMUSG( | ENSMUSG( | 0.430114 | 0.77617 | NO | NO | NA |
| Gm37973 | ENSMUSG( | ENSMUSG( | -0.37523 | 0.590342 | NO | NO | NA |
| Plppr1 | ENSMUSG( | ENSMUSG( | 0.867124 | 0.452821 | NO | NO | NA |
| Gm32391 | ENSMUSG( | ENSMUSG( | 0.368223 | 0.77071 | NO | NO | NA |
| Gm43882 | ENSMUSG( | ENSMUSG( | 0.755384 | 0.474914 | NO | NO | NA |
| Ecm2 | ENSMUSG( | ENSMUSG( | -0.46535 | 0.424153 | NO | NO | NA |
| Gm15693 | ENSMUSG( | ENSMUSG( | 0.808243 | 0.517719 | NO | NO | NA |
| Trdn | ENSMUSG( | ENSMUSG( | 0.94052 | 0.442282 | NO | NO | NA |
| ENSMUSG | ENSMUSG( | ENSMUSG( | -0.7136 | 0.10385 | NO | NO | NA |
| 9930022D | ENSMUSG( | ENSMUSG( | 0.936186 | 0.050344 | NO | NO | NA |
| Gm44056 | ENSMUSG( | ENSMUSG( | -0.08589 | 0.955945 | NO | NO | NA |
| EU599041 | ENSMUSG( | ENSMUSG( | 0.405879 | 0.726446 | NO | NO | NA |
| Gm48755 | ENSMUSG( | ENSMUSG( | -0.24379 | 0.855572 | NO | NO | NA |
| Itga5 | ENSMUSG( | ENSMUSG( | 0.127216 | 0.826879 | NO | NO | NA |
| Gm44940 | ENSMUSG( | ENSMUSG( | 0.609447 | 0.574263 | NO | NO | NA |
| Gm18329 | ENSMUSG( | ENSMUSG( | 0.355551 | 0.512102 | NO | NO | NA |
| Slc39a2 | ENSMUSG( | ENSMUSG( | 0.269185 | 0.565775 | NO | NO | NA |
| Rps15a-ps | ENSMUSG( | ENSMUSG( | 1.002767 | 0.126454 | NO | NO | NA |
| Gm37217 | ENSMUSG( | ENSMUSG( | 0.769907 | 0.478338 | NO | NO | NA |
| 1810062G | ENSMUSG( | ENSMUSG( | 0.898019 | 0.235038 | NO | NO | NA |
| Gm43923 | ENSMUSG( | ENSMUSG( | 0.780929 | 0.604904 | NO | NO | NA |
| Gm44288 | ENSMUSG( | ENSMUSG( | 0.113074 | 0.930307 | NO | NO | NA |
| Gm5134 | ENSMUSG( | ENSMUSG( | 0.164024 | 0.773408 | NO | NO | NA |
| Gm5619 | ENSMUSG( | ENSMUSG( | 0.307938 | 0.645355 | NO | NO | NA |
| Cnr1 | ENSMUSG( | ENSMUSG( | 0.846616 | 0.462756 | NO | NO | NA |
| ENSMUSG | ENSMUSG( | ENSMUSG( | -0.09394 | 0.877471 | NO | NO | NA |
| Gm10729 | ENSMUSG( | ENSMUSG( | 0.081254 | 0.946237 | NO | NO | NA |
| Bfsp1 | ENSMUSG( | ENSMUSG( | -0.84453 | 0.209133 | NO | NO | NA |
| Wdr93 | ENSMUSG( | ENSMUSG( | 0.178612 | 0.745208 | NO | NO | NA |
| Gm6311 | ENSMUSG( | ENSMUSG( | 0.204415 | 0.769416 | NO | NO | NA |

| | | | | | | | |
|---|---|---|---|---|---|---|---|
| Gm45560 | ENSMUSG( | ENSMUSG( | 0.9921 | 0.475218 | NO | NO | NA |
| Gm47569 | ENSMUSG( | ENSMUSG( | 1.119603 | 0.408456 | NO | NO | NA |
| Gm8871 | ENSMUSG( | ENSMUSG( | 1.167107 | 0.552241 | NO | NO | NA |
| Olfr1391 | ENSMUSG( | ENSMUSG( | 0.884444 | 0.401447 | NO | NO | NA |
| Zfp57 | ENSMUSG( | ENSMUSG( | -0.04065 | 0.968633 | NO | NO | NA |
| Gm47875 | ENSMUSG( | ENSMUSG( | -0.12765 | 0.929003 | NO | NO | NA |
| Gm8924 | ENSMUSG( | ENSMUSG( | 1.05186 | 0.373608 | NO | NO | NA |
| Zfp184 | ENSMUSG( | ENSMUSG( | 0.99411 | 0.252174 | NO | NO | NA |
| Gm15991 | ENSMUSG( | ENSMUSG( | 0.671846 | 0.542494 | NO | NO | NA |
| Trav6d-5 | ENSMUSG( | ENSMUSG( | -0.00331 | 0.997273 | NO | NO | NA |
| Gm15608 | ENSMUSG( | ENSMUSG( | 0.249301 | 0.72046 | NO | NO | NA |
| Gm47003 | ENSMUSG( | ENSMUSG( | -0.44384 | 0.495445 | NO | NO | NA |
| C730045M | ENSMUSG( | ENSMUSG( | 0.314572 | 0.653824 | NO | NO | NA |
| Gm17641 | ENSMUSG( | ENSMUSG( | 2.524825 | 0.002377 | YES | YES | Upregulated |
| Gm44487 | ENSMUSG( | ENSMUSG( | 0.858087 | 0.44719 | NO | NO | NA |
| Gm48764 | ENSMUSG( | ENSMUSG( | -0.785 | 0.113644 | NO | NO | NA |
| Gm14165 | ENSMUSG( | ENSMUSG( | 0.156981 | 0.850268 | NO | NO | NA |
| Mhrt | ENSMUSG( | ENSMUSG( | 0.419637 | 0.725572 | NO | NO | NA |
| Gm44185 | ENSMUSG( | ENSMUSG( | 0.471779 | 0.4007 | NO | NO | NA |
| ENSMUSG | ENSMUSG( | ENSMUSG( | 0.268418 | 0.65669 | NO | NO | NA |
| Gm13333 | ENSMUSG( | ENSMUSG( | -0.08973 | 0.938419 | NO | NO | NA |
| Gm37186 | ENSMUSG( | ENSMUSG( | 0.479558 | 0.492185 | NO | NO | NA |
| Tmem17 | ENSMUSG( | ENSMUSG( | 1.830302 | 0.010944 | YES | YES | Upregulated |
| A530064N | ENSMUSG( | ENSMUSG( | -0.77181 | 0.333497 | NO | NO | NA |
| Tnfsfm13 | ENSMUSG( | ENSMUSG( | -0.09829 | 0.914579 | NO | NO | NA |
| Slc34a1 | ENSMUSG( | ENSMUSG( | 0.440546 | 0.654043 | NO | NO | NA |
| Gm13226 | ENSMUSG( | ENSMUSG( | 1.090646 | 0.034137 | YES | YES | Upregulated |
| Gm3854 | ENSMUSG( | ENSMUSG( | 0.346017 | 0.656466 | NO | NO | NA |
| Mrgprc2-p | ENSMUSG( | ENSMUSG( | 0.822515 | 0.504438 | NO | NO | NA |
| Shisa2 | ENSMUSG( | ENSMUSG( | -0.08226 | 0.946459 | NO | NO | NA |
| Tceal3 | ENSMUSG( | ENSMUSG( | 0.532589 | 0.599416 | NO | NO | NA |
| Gm5628 | ENSMUSG( | ENSMUSG( | -0.07756 | 0.953489 | NO | NO | NA |
| Cd180 | ENSMUSG( | ENSMUSG( | -0.3354 | 0.727799 | NO | NO | NA |
| Olfr1192-p | ENSMUSG( | ENSMUSG( | 0.266138 | 0.846339 | NO | NO | NA |
| Gm19169 | ENSMUSG( | ENSMUSG( | 0.790214 | 0.471513 | NO | NO | NA |
| ENSMUSG | ENSMUSG( | ENSMUSG( | 0.589998 | 0.326489 | NO | NO | NA |
| ENSMUSG | ENSMUSG( | ENSMUSG( | -0.65875 | 0.24754 | NO | NO | NA |
| Gm42684 | ENSMUSG( | ENSMUSG( | -1.48231 | 0.020204 | YES | YES | Downregulated |
| Nat8f3 | ENSMUSG( | ENSMUSG( | 1.557523 | 0.109241 | NO | NO | NA |
| Gm10916 | ENSMUSG( | ENSMUSG( | -0.11407 | 0.818006 | NO | NO | NA |
| Gm48585 | ENSMUSG( | ENSMUSG( | 0.334442 | 0.630021 | NO | NO | NA |
| Gm43781 | ENSMUSG( | ENSMUSG( | 0.955197 | 0.432522 | NO | NO | NA |
| Igkv2-112 | ENSMUSG( | ENSMUSG( | -2.43782 | 0.47047 | NO | NO | NA |
| Gm16701 | ENSMUSG( | ENSMUSG( | -1.46288 | 0.002465 | YES | YES | Downregulated |
| Ephb4 | ENSMUSG( | ENSMUSG( | 0.468074 | 0.552591 | NO | NO | NA |
| Gm43708 | ENSMUSG( | ENSMUSG( | 0.917615 | 0.493174 | NO | NO | NA |
| Olfr729 | ENSMUSG( | ENSMUSG( | 0.810235 | 0.433696 | NO | NO | NA |

| | | | | | | | |
|---|---|---|---|---|---|---|---|
| Gm27249 | ENSMUSG( | ENSMUSG( | 0.900939 | 0.531667 | NO | NO | NA |
| Gm27184 | ENSMUSG( | ENSMUSG( | 0.209537 | 0.706226 | NO | NO | NA |
| ENSMUSG( | ENSMUSG( | ENSMUSG( | -1.68169 | 0.21605 | NO | NO | NA |
| Ifi205 | ENSMUSG( | ENSMUSG( | -0.76641 | 0.491858 | NO | NO | NA |
| C630043F( | ENSMUSG( | ENSMUSG( | 0.185321 | 0.823474 | NO | NO | NA |
| Ppp4r4 | ENSMUSG( | ENSMUSG( | 0.674406 | 0.198615 | NO | NO | NA |
| Gm13230 | ENSMUSG( | ENSMUSG( | 1.57691 | 0.083352 | NO | NO | NA |
| Tgm5 | ENSMUSG( | ENSMUSG( | 0.866553 | 0.537302 | NO | NO | NA |
| Nmrk2 | ENSMUSG( | ENSMUSG( | 0.359436 | 0.734433 | NO | NO | NA |
| Il1r1 | ENSMUSG( | ENSMUSG( | 0.738308 | 0.401589 | NO | NO | NA |
| Gm16283 | ENSMUSG( | ENSMUSG( | -0.16059 | 0.798402 | NO | NO | NA |
| Gm7557 | ENSMUSG( | ENSMUSG( | 0.46843 | 0.748181 | NO | NO | NA |
| Gm44509 | ENSMUSG( | ENSMUSG( | 0.343598 | 0.722714 | NO | NO | NA |
| ENSMUSG | ENSMUSG( | ENSMUSG( | 0.696551 | 0.519638 | NO | NO | NA |
| Zscan4-ps3 | ENSMUSG( | ENSMUSG( | 0.643476 | 0.557697 | NO | NO | NA |
| Gm31392 | ENSMUSG( | ENSMUSG( | -0.08296 | 0.924369 | NO | NO | NA |
| Tnr | ENSMUSG( | ENSMUSG( | 0.587681 | 0.576938 | NO | NO | NA |
| Nhs | ENSMUSG( | ENSMUSG( | 0.333854 | 0.753653 | NO | NO | NA |
| Vmn1r217 | ENSMUSG( | ENSMUSG( | 0.500757 | 0.700004 | NO | NO | NA |
| Xpnpep2 | ENSMUSG( | ENSMUSG( | 0.618906 | 0.545331 | NO | NO | NA |
| Rapgefl1 | ENSMUSG( | ENSMUSG( | -0.15399 | 0.85211 | NO | NO | NA |
| Gm38178 | ENSMUSG( | ENSMUSG( | 1.293009 | 0.288828 | NO | NO | NA |
| Prss35 | ENSMUSG( | ENSMUSG( | 1.288956 | 0.103506 | NO | NO | NA |
| Apoa2 | ENSMUSG( | ENSMUSG( | 0.601738 | 0.43991 | NO | NO | NA |
| Gm3830 | ENSMUSG( | ENSMUSG( | -0.15564 | 0.846519 | NO | NO | NA |
| Gm48309 | ENSMUSG( | ENSMUSG( | -1.42124 | 0.123597 | NO | NO | NA |
| Gm43329 | ENSMUSG( | ENSMUSG( | -0.63446 | 0.196645 | NO | NO | NA |
| Gm37595 | ENSMUSG( | ENSMUSG( | 0.568344 | 0.659678 | NO | NO | NA |
| Gm48682 | ENSMUSG( | ENSMUSG( | 0.55174 | 0.720556 | NO | NO | NA |
| D030024E( | ENSMUSG( | ENSMUSG( | 0.165248 | 0.910266 | NO | NO | NA |
| Ldb2 | ENSMUSG( | ENSMUSG( | 0.286462 | 0.806441 | NO | NO | NA |
| Snhg9 | ENSMUSG( | ENSMUSG( | 0.428038 | 0.526045 | NO | NO | NA |
| Gm48360 | ENSMUSG( | ENSMUSG( | 1.34707 | 0.354347 | NO | NO | NA |
| Gm45094 | ENSMUSG( | ENSMUSG( | -0.30984 | 0.596711 | NO | NO | NA |
| Cyp11b1 | ENSMUSG( | ENSMUSG( | 1.032136 | 0.244485 | NO | NO | NA |
| ENSMUSG | ENSMUSG( | ENSMUSG( | -0.08842 | 0.909378 | NO | NO | NA |
| Gm20900 | ENSMUSG( | ENSMUSG( | -0.17082 | 0.741308 | NO | NO | NA |
| Gm47251 | ENSMUSG( | ENSMUSG( | 0.809364 | 0.237758 | NO | NO | NA |
| Traj21 | ENSMUSG( | ENSMUSG( | 0.204884 | 0.752952 | NO | NO | NA |
| Gm16287 | ENSMUSG( | ENSMUSG( | 0.165368 | 0.815645 | NO | NO | NA |
| Speg | ENSMUSG( | ENSMUSG( | 0.045578 | 0.955293 | NO | NO | NA |
| Zfp811 | ENSMUSG( | ENSMUSG( | -0.00548 | 0.993519 | NO | NO | NA |
| ENSMUSG | ENSMUSG( | ENSMUSG( | -0.35019 | 0.573192 | NO | NO | NA |
| Mep1a | ENSMUSG( | ENSMUSG( | 0.733405 | 0.627656 | NO | NO | NA |
| Smim6 | ENSMUSG( | ENSMUSG( | -0.09288 | 0.875834 | NO | NO | NA |
| Gm49300 | ENSMUSG( | ENSMUSG( | 0.479838 | 0.70225 | NO | NO | NA |
| BC048559 | ENSMUSG( | ENSMUSG( | -0.61258 | 0.295575 | NO | NO | NA |

| | | | | | | | |
|---|---|---|---|---|---|---|---|
| Tcf23 | ENSMUSG( | ENSMUSG( | 1.233568 | 0.278918 | NO | NO | NA |
| Gm6501 | ENSMUSG( | ENSMUSG( | 0.13902 | 0.764232 | NO | NO | NA |
| Gm42656 | ENSMUSG( | ENSMUSG( | 0.25864 | 0.86261 | NO | NO | NA |
| Gm42894 | ENSMUSG( | ENSMUSG( | 0.733526 | 0.539734 | NO | NO | NA |
| Gm37238 | ENSMUSG( | ENSMUSG( | -0.01263 | 0.979976 | NO | NO | NA |
| Slc8a3 | ENSMUSG( | ENSMUSG( | 0.494564 | 0.596956 | NO | NO | NA |
| Osbp2 | ENSMUSG( | ENSMUSG( | 0.058731 | 0.963858 | NO | NO | NA |
| Gm11551 | ENSMUSG( | ENSMUSG( | 1.541726 | 0.28741 | NO | NO | NA |
| Ndst3 | ENSMUSG( | ENSMUSG( | 0.459781 | 0.544017 | NO | NO | NA |
| Brinp1 | ENSMUSG( | ENSMUSG( | 0.300534 | 0.814825 | NO | NO | NA |
| Epha10 | ENSMUSG( | ENSMUSG( | -0.42405 | 0.63183 | NO | NO | NA |
| Epha1 | ENSMUSG( | ENSMUSG( | -0.71579 | 0.16738 | NO | NO | NA |
| Gm47400 | ENSMUSG( | ENSMUSG( | 0.880777 | 0.477371 | NO | NO | NA |
| Bcl6b | ENSMUSG( | ENSMUSG( | -0.39456 | 0.545509 | NO | NO | NA |
| Gm2707 | ENSMUSG( | ENSMUSG( | 0.816226 | 0.500026 | NO | NO | NA |
| Gm26736 | ENSMUSG( | ENSMUSG( | 0.723212 | 0.552496 | NO | NO | NA |
| Gm10687 | ENSMUSG( | ENSMUSG( | 0.22386 | 0.819148 | NO | NO | NA |
| Themis3 | ENSMUSG( | ENSMUSG( | -0.28443 | 0.835599 | NO | NO | NA |
| Pcolce | ENSMUSG( | ENSMUSG( | -0.09541 | 0.838501 | NO | NO | NA |
| Gm38140 | ENSMUSG( | ENSMUSG( | 0.278361 | 0.665539 | NO | NO | NA |
| Htr1a | ENSMUSG( | ENSMUSG( | -0.63567 | 0.629039 | NO | NO | NA |
| Gm13864 | ENSMUSG( | ENSMUSG( | 0.90785 | 0.397767 | NO | NO | NA |
| Aldh3b1 | ENSMUSG( | ENSMUSG( | -1.17027 | 0.081331 | NO | NO | NA |
| Gm48657 | ENSMUSG( | ENSMUSG( | 0.492295 | 0.534573 | NO | NO | NA |
| Gm15558 | ENSMUSG( | ENSMUSG( | 1.066305 | 0.249328 | NO | NO | NA |
| Gm43301 | ENSMUSG( | ENSMUSG( | 1.151472 | 0.350484 | NO | NO | NA |
| Gm44550 | ENSMUSG( | ENSMUSG( | -0.32968 | 0.547137 | NO | NO | NA |
| Acsm3 | ENSMUSG( | ENSMUSG( | 1.05698 | 0.182925 | NO | NO | NA |
| Sod3 | ENSMUSG( | ENSMUSG( | 1.133346 | 0.295967 | NO | NO | NA |
| Gm11956 | ENSMUSG( | ENSMUSG( | 0.257291 | 0.872677 | NO | NO | NA |
| Gm7535 | ENSMUSG( | ENSMUSG( | -0.30439 | 0.71691 | NO | NO | NA |
| ENSMUSG | ENSMUSG( | ENSMUSG( | 0.211134 | 0.765362 | NO | NO | NA |
| Vmn1r79 | ENSMUSG( | ENSMUSG( | 0.368531 | 0.732409 | NO | NO | NA |
| Gm48054 | ENSMUSG( | ENSMUSG( | -0.01357 | 0.990103 | NO | NO | NA |
| Gm128 | ENSMUSG( | ENSMUSG( | -0.31046 | 0.674542 | NO | NO | NA |
| Tnfsf12 | ENSMUSG( | ENSMUSG( | -0.06997 | 0.926943 | NO | NO | NA |
| ENSMUSG( | ENSMUSG( | ENSMUSG( | 0.21416 | 0.821556 | NO | NO | NA |
| Vmn1r84 | ENSMUSG( | ENSMUSG( | 0.456089 | 0.656466 | NO | NO | NA |
| ENSMUSG | ENSMUSG( | ENSMUSG( | -0.46964 | 0.371845 | NO | NO | NA |
| Olfr2 | ENSMUSG( | ENSMUSG( | 0.513277 | 0.519155 | NO | NO | NA |
| Vmn1r-ps1 | ENSMUSG( | ENSMUSG( | 1.408402 | 0.239491 | NO | NO | NA |
| 4930512H | ENSMUSG( | ENSMUSG( | -1.38007 | 0.066828 | NO | NO | NA |
| Nox1 | ENSMUSG( | ENSMUSG( | -0.12878 | 0.842612 | NO | NO | NA |
| Dlg5 | ENSMUSG( | ENSMUSG( | 0.071999 | 0.932228 | NO | NO | NA |
| Gm14659 | ENSMUSG( | ENSMUSG( | 0.98433 | 0.481807 | NO | NO | NA |
| Gm14800 | ENSMUSG( | ENSMUSG( | 0.732905 | 0.544498 | NO | NO | NA |
| Gm2800 | ENSMUSG( | ENSMUSG( | 0.245552 | 0.739749 | NO | NO | NA |

| | | | | | | | |
|---|---|---|---|---|---|---|---|
| Avpr1a | ENSMUSG( | ENSMUSG( | 0.435064 | 0.683101 | NO | NO | NA |
| Olfr480 | ENSMUSG( | ENSMUSG( | 1.419836 | 0.224654 | NO | NO | NA |
| Gm29417 | ENSMUSG( | ENSMUSG( | 0.58405 | 0.589158 | NO | NO | NA |
| Gm45358 | ENSMUSG( | ENSMUSG( | -0.40213 | 0.610637 | NO | NO | NA |
| Foxl2os | ENSMUSG( | ENSMUSG( | 0.027564 | 0.986345 | NO | NO | NA |
| Gm7367 | ENSMUSG( | ENSMUSG( | 0.581449 | 0.422508 | NO | NO | NA |
| Tnfrsf23 | ENSMUSG( | ENSMUSG( | 0.044425 | 0.943313 | NO | NO | NA |
| Tmprss11l | ENSMUSG( | ENSMUSG( | 0.750774 | 0.472355 | NO | NO | NA |
| Olfr1202 | ENSMUSG( | ENSMUSG( | 0.47592 | 0.684265 | NO | NO | NA |
| ENSMUSG | ENSMUSG( | ENSMUSG( | 0.208936 | 0.703822 | NO | NO | NA |
| Gm44682 | ENSMUSG( | ENSMUSG( | 0.373819 | 0.729187 | NO | NO | NA |
| Vegfa | ENSMUSG( | ENSMUSG( | 1.343383 | 0.126477 | NO | NO | NA |
| Gm7658 | ENSMUSG( | ENSMUSG( | 0.068836 | 0.90316 | NO | NO | NA |
| Rps15a-ps | ENSMUSG( | ENSMUSG( | 0.033651 | 0.949621 | NO | NO | NA |
| Mansc4 | ENSMUSG( | ENSMUSG( | -0.65789 | 0.423063 | NO | NO | NA |
| Kctd15 | ENSMUSG( | ENSMUSG( | -0.13706 | 0.884166 | NO | NO | NA |
| Frem2 | ENSMUSG( | ENSMUSG( | 2.431928 | 0.006727 | YES | YES | Upregulated |
| Rit2 | ENSMUSG( | ENSMUSG( | 0.876053 | 0.456978 | NO | NO | NA |
| Gm10553 | ENSMUSG( | ENSMUSG( | 0.352023 | 0.526045 | NO | NO | NA |
| Pcdhga2 | ENSMUSG( | ENSMUSG( | 0.655451 | 0.665687 | NO | NO | NA |
| 1700100I1 | ENSMUSG( | ENSMUSG( | 0.334261 | 0.755183 | NO | NO | NA |
| Inhbe | ENSMUSG( | ENSMUSG( | 0.540935 | 0.552744 | NO | NO | NA |
| Gm26542 | ENSMUSG( | ENSMUSG( | -0.34073 | 0.564931 | NO | NO | NA |
| Gm44186 | ENSMUSG( | ENSMUSG( | 1.053778 | 0.334781 | NO | NO | NA |
| Gm44095 | ENSMUSG( | ENSMUSG( | 1.159986 | 0.422361 | NO | NO | NA |
| Gm37478 | ENSMUSG( | ENSMUSG( | 0.963717 | 0.393942 | NO | NO | NA |
| Gm3739 | ENSMUSG( | ENSMUSG( | 0.288553 | 0.70225 | NO | NO | NA |
| Zfp986 | ENSMUSG( | ENSMUSG( | 5.66477 | 0.008085 | YES | YES | Upregulated |
| Olfr825 | ENSMUSG( | ENSMUSG( | 0.194581 | 0.857884 | NO | NO | NA |
| Tmem220 | ENSMUSG( | ENSMUSG( | -0.12396 | 0.807141 | NO | NO | NA |
| Gm45366 | ENSMUSG( | ENSMUSG( | 0.857951 | 0.552744 | NO | NO | NA |
| Olfr878 | ENSMUSG( | ENSMUSG( | 0.731137 | 0.548545 | NO | NO | NA |
| Gm2553 | ENSMUSG( | ENSMUSG( | 0.569 | 0.678924 | NO | NO | NA |
| A430110L: | ENSMUSG( | ENSMUSG( | -0.94094 | 0.219358 | NO | NO | NA |
| 1700001G | ENSMUSG( | ENSMUSG( | 1.564857 | 0.334781 | NO | NO | NA |
| Eif1ad12 | ENSMUSG( | ENSMUSG( | 1.358387 | 0.505843 | NO | NO | NA |
| Olfr671 | ENSMUSG( | ENSMUSG( | 0.463839 | 0.735159 | NO | NO | NA |
| Gm42936 | ENSMUSG( | ENSMUSG( | 4.900334 | NA | NO | NO | NA |
| ENSMUSG( | ENSMUSG( | ENSMUSG( | -4.70701 | NA | NO | NO | NA |
| D130004A | ENSMUSG( | ENSMUSG( | 0.799003 | 0.495195 | NO | NO | NA |
| Gm14635 | ENSMUSG( | ENSMUSG( | 1.054491 | 0.43682 | NO | NO | NA |
| Flvcr2 | ENSMUSG( | ENSMUSG( | -0.08876 | 0.879627 | NO | NO | NA |
| Gm7461 | ENSMUSG( | ENSMUSG( | 0.017696 | 0.982586 | NO | NO | NA |
| Cfb | ENSMUSG( | ENSMUSG( | 0.150445 | 0.824613 | NO | NO | NA |
| Gm37849 | ENSMUSG( | ENSMUSG( | 1.352283 | 0.273857 | NO | NO | NA |
| Gm18142 | ENSMUSG( | ENSMUSG( | 1.408105 | 0.278918 | NO | NO | NA |
| Gm16235 | ENSMUSG( | ENSMUSG( | -0.22205 | 0.740776 | NO | NO | NA |

| | | | | | | | |
|---|---|---|---|---|---|---|---|
| Gm6652 | ENSMUSG( | ENSMUSG( | 0.125438 | 0.827515 | NO | NO | NA |
| 9630010A | ENSMUSG( | ENSMUSG( | 0.142378 | 0.836679 | NO | NO | NA |
| ENSMUSG( | ENSMUSG( | ENSMUSG( | -0.37797 | 0.504319 | NO | NO | NA |
| Gm15886 | ENSMUSG( | ENSMUSG( | 0.351218 | 0.624879 | NO | NO | NA |
| Nox4 | ENSMUSG( | ENSMUSG( | -0.09197 | 0.936437 | NO | NO | NA |
| Cdh7 | ENSMUSG( | ENSMUSG( | 0.222142 | 0.857884 | NO | NO | NA |
| Vit | ENSMUSG( | ENSMUSG( | 0.711865 | 0.545662 | NO | NO | NA |
| Gm12488 | ENSMUSG( | ENSMUSG( | -0.12159 | 0.914757 | NO | NO | NA |
| Btf3-ps15 | ENSMUSG( | ENSMUSG( | 1.346287 | 0.303179 | NO | NO | NA |
| Gm49905 | ENSMUSG( | ENSMUSG( | 1.298359 | 0.350706 | NO | NO | NA |
| Gm36264 | ENSMUSG( | ENSMUSG( | 0.750888 | 0.532905 | NO | NO | NA |
| ENSMUSG( | ENSMUSG( | ENSMUSG( | -0.0468 | 0.966629 | NO | NO | NA |
| Nppc | ENSMUSG( | ENSMUSG( | 0.736364 | 0.551031 | NO | NO | NA |
| Gm20537 | ENSMUSG( | ENSMUSG( | 0.011906 | 0.987624 | NO | NO | NA |
| Gm19001 | ENSMUSG( | ENSMUSG( | 0.733828 | 0.544308 | NO | NO | NA |
| Acsl6 | ENSMUSG( | ENSMUSG( | 0.416167 | 0.570158 | NO | NO | NA |
| A230006K | ENSMUSG( | ENSMUSG( | 0.439537 | 0.735159 | NO | NO | NA |
| Gm12874 | ENSMUSG( | ENSMUSG( | 0.373311 | 0.816153 | NO | NO | NA |
| Gm36949 | ENSMUSG( | ENSMUSG( | 1.226984 | 0.129646 | NO | NO | NA |
| Gm43113 | ENSMUSG( | ENSMUSG( | 0.634921 | 0.612079 | NO | NO | NA |
| Gm45098 | ENSMUSG( | ENSMUSG( | 0.065964 | 0.968633 | NO | NO | NA |
| 4930570N | ENSMUSG( | ENSMUSG( | 0.07904 | 0.863904 | NO | NO | NA |
| Gm9512 | ENSMUSG( | ENSMUSG( | -0.1899 | 0.797665 | NO | NO | NA |
| Gm27252 | ENSMUSG( | ENSMUSG( | 0.184414 | 0.707365 | NO | NO | NA |
| Noxred1 | ENSMUSG( | ENSMUSG( | -0.73744 | 0.206197 | NO | NO | NA |
| Csf1r-ps | ENSMUSG( | ENSMUSG( | 0.739468 | 0.492239 | NO | NO | NA |
| Gm42480 | ENSMUSG( | ENSMUSG( | 0.786877 | 0.594479 | NO | NO | NA |
| Cyp2c29 | ENSMUSG( | ENSMUSG( | 0.714651 | 0.614256 | NO | NO | NA |
| Gm15941 | ENSMUSG( | ENSMUSG( | -0.69433 | 0.31724 | NO | NO | NA |
| Gm43647 | ENSMUSG( | ENSMUSG( | 0.165067 | 0.836198 | NO | NO | NA |
| Gm2614 | ENSMUSG( | ENSMUSG( | 0.907359 | 0.132965 | NO | NO | NA |
| Gm13161 | ENSMUSG( | ENSMUSG( | 0.71757 | 0.542061 | NO | NO | NA |
| Gm47359 | ENSMUSG( | ENSMUSG( | 0.476963 | 0.726412 | NO | NO | NA |
| Vmn2r106 | ENSMUSG( | ENSMUSG( | 1.280086 | 0.328402 | NO | NO | NA |
| Gm5763 | ENSMUSG( | ENSMUSG( | -0.1912 | 0.709842 | NO | NO | NA |
| Gm18860 | ENSMUSG( | ENSMUSG( | -0.04502 | 0.930577 | NO | NO | NA |
| Hoxc4 | ENSMUSG( | ENSMUSG( | 0.845955 | 0.154019 | NO | NO | NA |
| Gm37078 | ENSMUSG( | ENSMUSG( | -0.4461 | 0.63996 | NO | NO | NA |
| 8430422M | ENSMUSG( | ENSMUSG( | 0.394872 | 0.461176 | NO | NO | NA |
| Gm43334 | ENSMUSG( | ENSMUSG( | 0.843508 | 0.575669 | NO | NO | NA |
| Gm44270 | ENSMUSG( | ENSMUSG( | 0.829712 | 0.13394 | NO | NO | NA |
| Dhtkd1 | ENSMUSG( | ENSMUSG( | 0.172106 | 0.81886 | NO | NO | NA |
| Gm43138 | ENSMUSG( | ENSMUSG( | 1.265348 | 0.104188 | NO | NO | NA |
| Gm15328 | ENSMUSG( | ENSMUSG( | 0.501683 | 0.692686 | NO | NO | NA |
| 5033406O | ENSMUSG( | ENSMUSG( | 0.656307 | 0.36861 | NO | NO | NA |
| Albfm1 | ENSMUSG( | ENSMUSG( | -1.12925 | 0.093463 | NO | NO | NA |
| Gm45677 | ENSMUSG( | ENSMUSG( | 0.413537 | 0.736236 | NO | NO | NA |

| | | | | | | | |
|---|---|---|---|---|---|---|---|
| Dmpk | ENSMUSG( | ENSMUSG( | -0.16486 | 0.791755 | NO | NO | NA |
| Sertad4 | ENSMUSG( | ENSMUSG( | 1.111164 | 0.285761 | NO | NO | NA |
| Gm44664 | ENSMUSG( | ENSMUSG( | -1.66853 | 0.032953 | YES | YES | Downregulated |
| Kcnj10 | ENSMUSG( | ENSMUSG( | -0.90551 | 0.171966 | NO | NO | NA |
| 4930599N | ENSMUSG( | ENSMUSG( | -0.35237 | 0.580811 | NO | NO | NA |
| Gm45903 | ENSMUSG( | ENSMUSG( | -0.03941 | 0.964642 | NO | NO | NA |
| Gm47037 | ENSMUSG( | ENSMUSG( | 0.786738 | 0.481044 | NO | NO | NA |
| Gm6235 | ENSMUSG( | ENSMUSG( | 0.655196 | 0.557016 | NO | NO | NA |
| Gm11249 | ENSMUSG( | ENSMUSG( | 0.79013 | 0.211222 | NO | NO | NA |
| 2310020H | ENSMUSG( | ENSMUSG( | 1.248043 | 0.304251 | NO | NO | NA |
| ENSMUSG | ENSMUSG( | ENSMUSG( | -0.2627 | 0.683247 | NO | NO | NA |
| AI661453 | ENSMUSG( | ENSMUSG( | 0.528392 | 0.474503 | NO | NO | NA |
| Gm45015 | ENSMUSG( | ENSMUSG( | 0.892341 | 0.236061 | NO | NO | NA |
| Gm48621 | ENSMUSG( | ENSMUSG( | -0.5362 | 0.482172 | NO | NO | NA |
| Olfr704 | ENSMUSG( | ENSMUSG( | 0.437479 | 0.705078 | NO | NO | NA |
| Gm42513 | ENSMUSG( | ENSMUSG( | 0.494338 | 0.62909 | NO | NO | NA |
| Gm4575 | ENSMUSG( | ENSMUSG( | 0.762852 | 0.527649 | NO | NO | NA |
| 1700016L2 | ENSMUSG( | ENSMUSG( | 0.593937 | 0.626685 | NO | NO | NA |
| Gm37296 | ENSMUSG( | ENSMUSG( | 0.572978 | 0.552134 | NO | NO | NA |
| Gm47969 | ENSMUSG( | ENSMUSG( | 0.495193 | 0.707602 | NO | NO | NA |
| Cib3 | ENSMUSG( | ENSMUSG( | -0.19304 | 0.792444 | NO | NO | NA |
| Gm15950 | ENSMUSG( | ENSMUSG( | -0.23762 | 0.757727 | NO | NO | NA |
| Fgf12 | ENSMUSG( | ENSMUSG( | 1.045731 | 0.395366 | NO | NO | NA |
| Asap3 | ENSMUSG( | ENSMUSG( | -0.26536 | 0.776428 | NO | NO | NA |
| Gm16409 | ENSMUSG( | ENSMUSG( | 1.534136 | 0.163035 | NO | NO | NA |
| Gm38235 | ENSMUSG( | ENSMUSG( | 0.284344 | 0.770926 | NO | NO | NA |
| Zfp366 | ENSMUSG( | ENSMUSG( | -1.88368 | 0.014804 | YES | YES | Downregulated |
| Zp3r | ENSMUSG( | ENSMUSG( | 1.108988 | 0.336657 | NO | NO | NA |
| Olfr381 | ENSMUSG( | ENSMUSG( | 1.071292 | 0.42523 | NO | NO | NA |
| Mmp9 | ENSMUSG( | ENSMUSG( | 0.510143 | 0.657253 | NO | NO | NA |
| 1700028J1 | ENSMUSG( | ENSMUSG( | 0.313385 | 0.683666 | NO | NO | NA |
| Gm2956 | ENSMUSG( | ENSMUSG( | 0.586027 | 0.731346 | NO | NO | NA |
| ENSMUSG | ENSMUSG( | ENSMUSG( | 1.351528 | 0.254258 | NO | NO | NA |
| Gm5385 | ENSMUSG( | ENSMUSG( | -0.26769 | 0.857884 | NO | NO | NA |
| Gm31048 | ENSMUSG( | ENSMUSG( | 2.033286 | 0.067152 | NO | NO | NA |
| ENSMUSG | ENSMUSG( | ENSMUSG( | 0.424021 | 0.74274 | NO | NO | NA |
| Igfn1 | ENSMUSG( | ENSMUSG( | 1.396706 | 0.349653 | NO | NO | NA |
| Gm26559 | ENSMUSG( | ENSMUSG( | -0.17143 | 0.879404 | NO | NO | NA |
| Vmn1r26 | ENSMUSG( | ENSMUSG( | 0.185847 | 0.890997 | NO | NO | NA |
| Astl | ENSMUSG( | ENSMUSG( | 0.221937 | 0.78842 | NO | NO | NA |
| Fau-ps2 | ENSMUSG( | ENSMUSG( | -0.29553 | 0.824386 | NO | NO | NA |
| Gm19710 | ENSMUSG( | ENSMUSG( | 0.354444 | 0.656122 | NO | NO | NA |
| Hey2 | ENSMUSG( | ENSMUSG( | 0.446082 | 0.717106 | NO | NO | NA |
| Mmp25 | ENSMUSG( | ENSMUSG( | 0.666032 | 0.538508 | NO | NO | NA |
| Gm43344 | ENSMUSG( | ENSMUSG( | 0.249277 | 0.840113 | NO | NO | NA |
| Hspe1-rs1 | ENSMUSG( | ENSMUSG( | 0.268758 | 0.750432 | NO | NO | NA |
| Gm2423 | ENSMUSG( | ENSMUSG( | -0.2218 | 0.806319 | NO | NO | NA |

| | | | | | | | |
|---|---|---|---|---|---|---|---|
| Gm11350 | ENSMUSG( | ENSMUSG( | 1.82001 | 0.126022 | NO | NO | NA |
| Gm49239 | ENSMUSG( | ENSMUSG( | 1.200341 | 0.405412 | NO | NO | NA |
| Gm28085 | ENSMUSG( | ENSMUSG( | -0.82006 | 0.232059 | NO | NO | NA |
| Gm11832 | ENSMUSG( | ENSMUSG( | 0.501442 | 0.641488 | NO | NO | NA |
| Gm48965 | ENSMUSG( | ENSMUSG( | 0.743811 | 0.459618 | NO | NO | NA |
| Myo1a | ENSMUSG( | ENSMUSG( | -0.42984 | 0.608452 | NO | NO | NA |
| Apon | ENSMUSG( | ENSMUSG( | 0.425801 | 0.469517 | NO | NO | NA |
| Olfr777 | ENSMUSG( | ENSMUSG( | 0.290242 | 0.813794 | NO | NO | NA |
| Gm44758 | ENSMUSG( | ENSMUSG( | -0.83428 | 0.613704 | NO | NO | NA |
| Akr1d1 | ENSMUSG( | ENSMUSG( | 1.107432 | 0.474914 | NO | NO | NA |
| Gm37814 | ENSMUSG( | ENSMUSG( | 0.764757 | 0.600373 | NO | NO | NA |
| Gm37696 | ENSMUSG( | ENSMUSG( | -0.82075 | 0.06282 | NO | NO | NA |
| Gm3619 | ENSMUSG( | ENSMUSG( | 1.139462 | 0.406671 | NO | NO | NA |
| Gm37349 | ENSMUSG( | ENSMUSG( | 0.995814 | 0.065369 | NO | NO | NA |
| ENSMUSG | ENSMUSG( | ENSMUSG( | 0.642639 | 0.470978 | NO | NO | NA |
| Gm20878 | ENSMUSG( | ENSMUSG( | 0.222468 | 0.664861 | NO | NO | NA |
| Gm37099 | ENSMUSG( | ENSMUSG( | 0.384902 | 0.713641 | NO | NO | NA |
| Gm48744 | ENSMUSG( | ENSMUSG( | 0.756749 | 0.546443 | NO | NO | NA |
| Trib3 | ENSMUSG( | ENSMUSG( | -0.60686 | 0.389693 | NO | NO | NA |
| Gm38090 | ENSMUSG( | ENSMUSG( | 0.713321 | 0.517274 | NO | NO | NA |
| Gm7589 | ENSMUSG( | ENSMUSG( | 0.054409 | 0.936782 | NO | NO | NA |
| Gm4968 | ENSMUSG( | ENSMUSG( | 0.341824 | 0.756346 | NO | NO | NA |
| Iqsec3 | ENSMUSG( | ENSMUSG( | -0.19684 | 0.868688 | NO | NO | NA |
| Gm40264 | ENSMUSG( | ENSMUSG( | 0.449636 | 0.784849 | NO | NO | NA |
| 3110083C: | ENSMUSG( | ENSMUSG( | -0.03256 | 0.953913 | NO | NO | NA |
| Gm44091 | ENSMUSG( | ENSMUSG( | -0.42743 | 0.453381 | NO | NO | NA |
| Rps23rg1 | ENSMUSG( | ENSMUSG( | 0.417265 | 0.516631 | NO | NO | NA |
| Mageb18 | ENSMUSG( | ENSMUSG( | 0.192174 | 0.898283 | NO | NO | NA |
| Bcl2a1a | ENSMUSG( | ENSMUSG( | 0.466343 | 0.523069 | NO | NO | NA |
| Pira2 | ENSMUSG( | ENSMUSG( | -0.23922 | 0.743374 | NO | NO | NA |
| Myof | ENSMUSG( | ENSMUSG( | -0.57234 | 0.466943 | NO | NO | NA |
| Capn3 | ENSMUSG( | ENSMUSG( | 1.554995 | 0.023003 | YES | YES | Upregulated |
| Amdhd1 | ENSMUSG( | ENSMUSG( | 0.006253 | 0.99076 | NO | NO | NA |
| Gm42435 | ENSMUSG( | ENSMUSG( | -0.12154 | 0.855452 | NO | NO | NA |
| Gm49496 | ENSMUSG( | ENSMUSG( | 1.551648 | 0.443997 | NO | NO | NA |
| Mecomos | ENSMUSG( | ENSMUSG( | 0.57265 | 0.63535 | NO | NO | NA |
| Gm34589 | ENSMUSG( | ENSMUSG( | 0.647118 | 0.590649 | NO | NO | NA |
| Cd209c | ENSMUSG( | ENSMUSG( | -0.26815 | 0.639774 | NO | NO | NA |
| Olfr711 | ENSMUSG( | ENSMUSG( | 0.248153 | 0.796234 | NO | NO | NA |
| Zfp85os | ENSMUSG( | ENSMUSG( | 0.382543 | 0.465954 | NO | NO | NA |
| Olfm5 | ENSMUSG( | ENSMUSG( | 0.329796 | 0.773408 | NO | NO | NA |
| Gm12404 | ENSMUSG( | ENSMUSG( | -0.16963 | 0.750323 | NO | NO | NA |
| Gm10647 | ENSMUSG( | ENSMUSG( | 0.31288 | 0.832243 | NO | NO | NA |
| Atoh7 | ENSMUSG( | ENSMUSG( | 0.59814 | 0.562672 | NO | NO | NA |
| 6330415B: | ENSMUSG( | ENSMUSG( | 0.53568 | 0.700632 | NO | NO | NA |
| Pcdh9 | ENSMUSG( | ENSMUSG( | 0.228227 | 0.845723 | NO | NO | NA |
| Xirp1 | ENSMUSG( | ENSMUSG( | -1.56249 | 0.103013 | NO | NO | NA |

| | | | | | | | |
|---|---|---|---|---|---|---|---|
| Gm10421 | ENSMUSG( | ENSMUSG( | 0.593453 | 0.603586 | NO | NO | NA |
| B630019K( | ENSMUSG( | ENSMUSG( | 1.065304 | 0.230821 | NO | NO | NA |
| Mir703 | ENSMUSG( | ENSMUSG( | 0.008586 | 0.987891 | NO | NO | NA |
| Gm52988 | ENSMUSG( | ENSMUSG( | 0.276326 | 0.835507 | NO | NO | NA |
| ENSMUSG | ENSMUSG( | ENSMUSG( | 0.76529 | 0.449674 | NO | NO | NA |
| Gm14979 | ENSMUSG( | ENSMUSG( | 0.346956 | 0.779557 | NO | NO | NA |
| Stac2 | ENSMUSG( | ENSMUSG( | -0.67208 | 0.345824 | NO | NO | NA |
| Gm11292 | ENSMUSG( | ENSMUSG( | 0.770869 | 0.471848 | NO | NO | NA |
| Hmx1 | ENSMUSG( | ENSMUSG( | 0.829526 | 0.451342 | NO | NO | NA |
| Ighv5-15 | ENSMUSG( | ENSMUSG( | -0.43258 | 0.795809 | NO | NO | NA |
| Slc34a2 | ENSMUSG( | ENSMUSG( | 0.777251 | 0.530589 | NO | NO | NA |
| Gm15114 | ENSMUSG( | ENSMUSG( | 0.510247 | 0.703614 | NO | NO | NA |
| Pcdha9 | ENSMUSG( | ENSMUSG( | 0.494401 | 0.824342 | NO | NO | NA |
| Gap43 | ENSMUSG( | ENSMUSG( | 0.394965 | 0.763095 | NO | NO | NA |
| Gm10745 | ENSMUSG( | ENSMUSG( | 0.721377 | 0.520813 | NO | NO | NA |
| Vmn2r32 | ENSMUSG( | ENSMUSG( | 1.247834 | 0.367037 | NO | NO | NA |
| Gm10165 | ENSMUSG( | ENSMUSG( | 0.618946 | 0.615925 | NO | NO | NA |
| Rdh1 | ENSMUSG( | ENSMUSG( | 0.49809 | 0.554494 | NO | NO | NA |
| C230013L1 | ENSMUSG( | ENSMUSG( | -0.02093 | 0.978085 | NO | NO | NA |
| Il15 | ENSMUSG( | ENSMUSG( | -0.23446 | 0.69705 | NO | NO | NA |
| Olfr1065 | ENSMUSG( | ENSMUSG( | 0.763643 | 0.503551 | NO | NO | NA |
| Galnt5 | ENSMUSG( | ENSMUSG( | 1.057626 | 0.311191 | NO | NO | NA |
| Igkv2-137 | ENSMUSG( | ENSMUSG( | -1.09977 | 0.633247 | NO | NO | NA |
| Gm48617 | ENSMUSG( | ENSMUSG( | 0.611545 | 0.620382 | NO | NO | NA |
| Spdye4b | ENSMUSG( | ENSMUSG( | 0.769707 | 0.506969 | NO | NO | NA |
| Sppl2c | ENSMUSG( | ENSMUSG( | 1.628525 | 0.183655 | NO | NO | NA |
| Olfr890 | ENSMUSG( | ENSMUSG( | 1.842425 | 0.129796 | NO | NO | NA |
| Smoc2 | ENSMUSG( | ENSMUSG( | 0.234592 | 0.804298 | NO | NO | NA |
| 4930525G | ENSMUSG( | ENSMUSG( | -0.56529 | 0.592341 | NO | NO | NA |
| Gm36447 | ENSMUSG( | ENSMUSG( | 0.619634 | 0.587034 | NO | NO | NA |
| ENSMUSG | ENSMUSG( | ENSMUSG( | -0.12637 | 0.924993 | NO | NO | NA |
| Vmn2r108 | ENSMUSG( | ENSMUSG( | 0.764983 | 0.454359 | NO | NO | NA |
| 2900060B1 | ENSMUSG( | ENSMUSG( | -0.46867 | 0.454883 | NO | NO | NA |
| Gm3534 | ENSMUSG( | ENSMUSG( | -0.62906 | 0.557697 | NO | NO | NA |
| Gm10113 | ENSMUSG( | ENSMUSG( | 1.753318 | 0.175836 | NO | NO | NA |
| Gpr141 | ENSMUSG( | ENSMUSG( | 0.447949 | 0.660508 | NO | NO | NA |
| Gm37876 | ENSMUSG( | ENSMUSG( | -0.90696 | 0.093724 | NO | NO | NA |
| Postn | ENSMUSG( | ENSMUSG( | 0.201461 | 0.851412 | NO | NO | NA |
| ENSMUSG( | ENSMUSG( | ENSMUSG( | 0.612108 | 0.582375 | NO | NO | NA |
| Cnksr3 | ENSMUSG( | ENSMUSG( | 0.92534 | 0.121973 | NO | NO | NA |
| H2ac12 | ENSMUSG( | ENSMUSG( | 0.120531 | 0.844265 | NO | NO | NA |
| Gm17041 | ENSMUSG( | ENSMUSG( | -1.53891 | 0.111618 | NO | NO | NA |
| AI429214 | ENSMUSG( | ENSMUSG( | 0.657491 | 0.405003 | NO | NO | NA |
| Gm15567 | ENSMUSG( | ENSMUSG( | 0.149411 | 0.814825 | NO | NO | NA |
| 0610033M | ENSMUSG( | ENSMUSG( | 0.633948 | 0.636839 | NO | NO | NA |
| Gm9633 | ENSMUSG( | ENSMUSG( | 1.172553 | 0.280406 | NO | NO | NA |
| Vmn1r20 | ENSMUSG( | ENSMUSG( | 0.60708 | 0.65434 | NO | NO | NA |

| | | | | | | | |
|---|---|---|---|---|---|---|---|
| Gm32515 | ENSMUSG( | ENSMUSG( | -1.16663 | 0.065617 | NO | NO | NA |
| Nupr1l | ENSMUSG( | ENSMUSG( | 0.284052 | 0.664907 | NO | NO | NA |
| Odaph | ENSMUSG( | ENSMUSG( | 0.915513 | 0.358045 | NO | NO | NA |
| Gm5781 | ENSMUSG( | ENSMUSG( | 0.589606 | 0.476831 | NO | NO | NA |
| ENSMUSG | ENSMUSG( | ENSMUSG( | 1.000923 | 0.272677 | NO | NO | NA |
| Nynrin | ENSMUSG( | ENSMUSG( | -0.35517 | 0.709532 | NO | NO | NA |
| Gm13223 | ENSMUSG( | ENSMUSG( | 1.462893 | 0.349893 | NO | NO | NA |
| Gm45731 | ENSMUSG( | ENSMUSG( | 0.155801 | 0.795982 | NO | NO | NA |
| Ighv1-61 | ENSMUSG( | ENSMUSG( | -1.24446 | NA | NO | NO | NA |
| Vmn2r7 | ENSMUSG( | ENSMUSG( | -0.02082 | 0.988483 | NO | NO | NA |
| Fgf5 | ENSMUSG( | ENSMUSG( | 0.671443 | 0.586177 | NO | NO | NA |
| Gm1947 | ENSMUSG( | ENSMUSG( | 0.321285 | 0.736541 | NO | NO | NA |
| Gm44257 | ENSMUSG( | ENSMUSG( | 0.988103 | 0.390415 | NO | NO | NA |
| Tslp | ENSMUSG( | ENSMUSG( | 1.190754 | 0.322609 | NO | NO | NA |
| Gm6807 | ENSMUSG( | ENSMUSG( | 0.752564 | 0.234325 | NO | NO | NA |
| Gm5277 | ENSMUSG( | ENSMUSG( | 0.313891 | 0.768416 | NO | NO | NA |
| Gm44958 | ENSMUSG( | ENSMUSG( | 0.300518 | 0.807555 | NO | NO | NA |
| Gcnt7 | ENSMUSG( | ENSMUSG( | -0.32765 | 0.698856 | NO | NO | NA |
| Gm43201 | ENSMUSG( | ENSMUSG( | -0.28576 | 0.646219 | NO | NO | NA |
| Olfr492 | ENSMUSG( | ENSMUSG( | 0.579624 | 0.601079 | NO | NO | NA |
| Gm45483 | ENSMUSG( | ENSMUSG( | 0.647913 | 0.511174 | NO | NO | NA |
| Gm33318 | ENSMUSG( | ENSMUSG( | 0.136971 | 0.785602 | NO | NO | NA |
| ENSMUSG | ENSMUSG( | ENSMUSG( | 0.933227 | 0.212895 | NO | NO | NA |
| Nrg2 | ENSMUSG( | ENSMUSG( | 0.468156 | 0.625072 | NO | NO | NA |
| Gm9472 | ENSMUSG( | ENSMUSG( | 0.722356 | 0.536092 | NO | NO | NA |
| Gm48138 | ENSMUSG( | ENSMUSG( | -0.03492 | 0.976719 | NO | NO | NA |
| Gm15902 | ENSMUSG( | ENSMUSG( | 1.499232 | 0.263565 | NO | NO | NA |
| Olfr301 | ENSMUSG( | ENSMUSG( | 1.54138 | 0.225137 | NO | NO | NA |
| Olfr1297 | ENSMUSG( | ENSMUSG( | -0.1463 | 0.899918 | NO | NO | NA |
| Pnldc1 | ENSMUSG( | ENSMUSG( | -0.20779 | 0.693858 | NO | NO | NA |
| Gm15624 | ENSMUSG( | ENSMUSG( | 0.536915 | 0.522159 | NO | NO | NA |
| ENSMUSG | ENSMUSG( | ENSMUSG( | -0.25563 | 0.588526 | NO | NO | NA |
| Msx3 | ENSMUSG( | ENSMUSG( | 0.915452 | 0.327201 | NO | NO | NA |
| Fam110c | ENSMUSG( | ENSMUSG( | 0.272825 | 0.803364 | NO | NO | NA |
| Pcdhgb5 | ENSMUSG( | ENSMUSG( | 0.09502 | 0.899882 | NO | NO | NA |
| Trav11n | ENSMUSG( | ENSMUSG( | 0.586113 | 0.827544 | NO | NO | NA |
| Gm43792 | ENSMUSG( | ENSMUSG( | 0.543832 | 0.718103 | NO | NO | NA |
| Gm28594 | ENSMUSG( | ENSMUSG( | 0.732224 | 0.589529 | NO | NO | NA |
| Gm48207 | ENSMUSG( | ENSMUSG( | 0.891975 | 0.507778 | NO | NO | NA |
| Stk32a | ENSMUSG( | ENSMUSG( | 0.690349 | 0.604548 | NO | NO | NA |
| Rai2 | ENSMUSG( | ENSMUSG( | 0.79802 | 0.460937 | NO | NO | NA |
| Pira1 | ENSMUSG( | ENSMUSG( | 1.136907 | 0.693448 | NO | NO | NA |
| Gm14913 | ENSMUSG( | ENSMUSG( | 1.306658 | 0.260999 | NO | NO | NA |
| Dnajb6-ps | ENSMUSG( | ENSMUSG( | 0.751952 | 0.216926 | NO | NO | NA |
| Tnc | ENSMUSG( | ENSMUSG( | 0.350497 | 0.747414 | NO | NO | NA |
| Ccdc180 | ENSMUSG( | ENSMUSG( | 0.096602 | 0.886368 | NO | NO | NA |
| Gm21010 | ENSMUSG( | ENSMUSG( | 1.375776 | 0.126385 | NO | NO | NA |

| | | | | | | | |
|---|---|---|---|---|---|---|---|
| Erg | ENSMUSG( | ENSMUSG( | -0.7844 | 0.208027 | NO | NO | NA |
| Gm19046 | ENSMUSG( | ENSMUSG( | 0.092149 | 0.906948 | NO | NO | NA |
| Gm42630 | ENSMUSG( | ENSMUSG( | 0.688999 | 0.609902 | NO | NO | NA |
| Gm45172 | ENSMUSG( | ENSMUSG( | -0.30879 | 0.80545 | NO | NO | NA |
| Prkg1 | ENSMUSG( | ENSMUSG( | 0.452771 | 0.710819 | NO | NO | NA |
| Barx2 | ENSMUSG( | ENSMUSG( | 0.746671 | 0.557697 | NO | NO | NA |
| Gm43490 | ENSMUSG( | ENSMUSG( | -0.52869 | 0.664677 | NO | NO | NA |
| Cyp17a1 | ENSMUSG( | ENSMUSG( | -1.95166 | 0.058447 | NO | NO | NA |
| Ube2n-ps1 | ENSMUSG( | ENSMUSG( | 0.579398 | 0.238215 | NO | NO | NA |
| ENSMUSG | ENSMUSG( | ENSMUSG( | 0.516247 | 0.672697 | NO | NO | NA |
| Pip5kl1 | ENSMUSG( | ENSMUSG( | 0.451269 | 0.556502 | NO | NO | NA |
| Klra4 | ENSMUSG( | ENSMUSG( | 1.032532 | 0.295427 | NO | NO | NA |
| Olfr1233 | ENSMUSG( | ENSMUSG( | 0.562076 | 0.580206 | NO | NO | NA |
| Cntn3 | ENSMUSG( | ENSMUSG( | 0.780266 | 0.503171 | NO | NO | NA |
| Gm5431 | ENSMUSG( | ENSMUSG( | -0.91947 | 0.054628 | NO | NO | NA |
| Insyn1 | ENSMUSG( | ENSMUSG( | -0.01183 | 0.991619 | NO | NO | NA |
| ENSMUSG | ENSMUSG( | ENSMUSG( | 0.642192 | 0.520079 | NO | NO | NA |
| Spaar | ENSMUSG( | ENSMUSG( | 0.75183 | 0.625041 | NO | NO | NA |
| Gm26596 | ENSMUSG( | ENSMUSG( | -0.24076 | 0.812047 | NO | NO | NA |
| Zscan4-ps2 | ENSMUSG( | ENSMUSG( | 0.298188 | 0.772331 | NO | NO | NA |
| Gm10051 | ENSMUSG( | ENSMUSG( | 0.07145 | 0.938123 | NO | NO | NA |
| Gm13066 | ENSMUSG( | ENSMUSG( | -0.30081 | 0.931063 | NO | NO | NA |
| Samd13 | ENSMUSG( | ENSMUSG( | 0.691417 | 0.565522 | NO | NO | NA |
| Olfr986 | ENSMUSG( | ENSMUSG( | 0.059846 | 0.966706 | NO | NO | NA |
| Gm16120 | ENSMUSG( | ENSMUSG( | -0.7872 | 0.236978 | NO | NO | NA |
| Gm39307 | ENSMUSG( | ENSMUSG( | 1.097625 | 0.361227 | NO | NO | NA |
| Slc22a13 | ENSMUSG( | ENSMUSG( | 0.527714 | 0.722275 | NO | NO | NA |
| Gm42991 | ENSMUSG( | ENSMUSG( | 0.002008 | 0.998642 | NO | NO | NA |
| ENSMUSG | ENSMUSG( | ENSMUSG( | 0.39988 | 0.711591 | NO | NO | NA |
| Gm43677 | ENSMUSG( | ENSMUSG( | -0.0358 | 0.98075 | NO | NO | NA |
| Uncx | ENSMUSG( | ENSMUSG( | -0.29419 | 0.819342 | NO | NO | NA |
| Gngt1 | ENSMUSG( | ENSMUSG( | 0.343949 | 0.730991 | NO | NO | NA |
| D830026I1 | ENSMUSG( | ENSMUSG( | 0.613868 | 0.63602 | NO | NO | NA |
| Gm10145 | ENSMUSG( | ENSMUSG( | 0.477852 | 0.444093 | NO | NO | NA |
| Junos | ENSMUSG( | ENSMUSG( | 0.702408 | 0.170142 | NO | NO | NA |
| Gm7785 | ENSMUSG( | ENSMUSG( | -0.04272 | 0.944504 | NO | NO | NA |
| Gm15965 | ENSMUSG( | ENSMUSG( | 0.275402 | 0.75175 | NO | NO | NA |
| Gm42815 | ENSMUSG( | ENSMUSG( | 0.590164 | 0.316122 | NO | NO | NA |
| Rps11-ps1 | ENSMUSG( | ENSMUSG( | -0.07258 | 0.899457 | NO | NO | NA |
| Ckm | ENSMUSG( | ENSMUSG( | 0.663155 | 0.180398 | NO | NO | NA |
| Gm42555 | ENSMUSG( | ENSMUSG( | 0.607013 | 0.355276 | NO | NO | NA |
| ENSMUSG | ENSMUSG( | ENSMUSG( | -0.05785 | 0.899882 | NO | NO | NA |
| Cfap77 | ENSMUSG( | ENSMUSG( | 0.523939 | 0.549271 | NO | NO | NA |
| Gm36644 | ENSMUSG( | ENSMUSG( | 0.529349 | 0.711437 | NO | NO | NA |
| Htr2c | ENSMUSG( | ENSMUSG( | 1.304028 | 0.245586 | NO | NO | NA |
| Rpsa-ps4 | ENSMUSG( | ENSMUSG( | 0.523 | 0.511124 | NO | NO | NA |
| Gm37211 | ENSMUSG( | ENSMUSG( | 0.369415 | 0.783213 | NO | NO | NA |

| | | | | | | | |
|---|---|---|---|---|---|---|---|
| Gm13243 | ENSMUSG( | ENSMUSG( | 0.762334 | 0.69034 | NO | NO | NA |
| Gm43484 | ENSMUSG( | ENSMUSG( | 0.108231 | 0.844661 | NO | NO | NA |
| Gm6159 | ENSMUSG( | ENSMUSG( | 0.127931 | 0.829174 | NO | NO | NA |
| A330040F: | ENSMUSG( | ENSMUSG( | -0.45164 | 0.535468 | NO | NO | NA |
| ENSMUSG | ENSMUSG( | ENSMUSG( | 0.052917 | 0.926387 | NO | NO | NA |
| Gm45534 | ENSMUSG( | ENSMUSG( | -0.58782 | 0.323853 | NO | NO | NA |
| Kdf1 | ENSMUSG( | ENSMUSG( | -0.54962 | 0.495773 | NO | NO | NA |
| Gprasp2 | ENSMUSG( | ENSMUSG( | -0.13395 | 0.847785 | NO | NO | NA |
| Cd300ld | ENSMUSG( | ENSMUSG( | 0.011428 | 0.991511 | NO | NO | NA |
| 4930483J1 | ENSMUSG( | ENSMUSG( | 0.573041 | 0.579897 | NO | NO | NA |
| Ifnlr1 | ENSMUSG( | ENSMUSG( | 0.182984 | 0.871067 | NO | NO | NA |
| G630093K | ENSMUSG( | ENSMUSG( | 0.372404 | 0.769397 | NO | NO | NA |
| Gm16998 | ENSMUSG( | ENSMUSG( | -0.0843 | 0.860095 | NO | NO | NA |
| Dnah9 | ENSMUSG( | ENSMUSG( | 0.302117 | 0.725782 | NO | NO | NA |
| Mrgpra4 | ENSMUSG( | ENSMUSG( | 0.614479 | 0.613076 | NO | NO | NA |
| Gm15303 | ENSMUSG( | ENSMUSG( | -0.50914 | 0.746198 | NO | NO | NA |
| Gm48552 | ENSMUSG( | ENSMUSG( | 0.209582 | 0.816371 | NO | NO | NA |
| Iftap | ENSMUSG( | ENSMUSG( | 1.464421 | 0.064419 | NO | NO | NA |
| Olfr1414 | ENSMUSG( | ENSMUSG( | 0.543572 | 0.618496 | NO | NO | NA |
| Gm9816 | ENSMUSG( | ENSMUSG( | 0.070911 | 0.903228 | NO | NO | NA |
| Kbtbd12 | ENSMUSG( | ENSMUSG( | -0.14369 | 0.899918 | NO | NO | NA |
| Slc38a8 | ENSMUSG( | ENSMUSG( | -0.80763 | 0.541187 | NO | NO | NA |
| Gm2387 | ENSMUSG( | ENSMUSG( | 0.954028 | 0.265152 | NO | NO | NA |
| Gm44902 | ENSMUSG( | ENSMUSG( | 1.67563 | 0.221308 | NO | NO | NA |
| Lrguk | ENSMUSG( | ENSMUSG( | -0.04511 | 0.976236 | NO | NO | NA |
| Pabpc5 | ENSMUSG( | ENSMUSG( | -0.03244 | 0.98075 | NO | NO | NA |
| Gm5420 | ENSMUSG( | ENSMUSG( | 0.523666 | 0.63395 | NO | NO | NA |
| Gm9432 | ENSMUSG( | ENSMUSG( | 0.298583 | 0.797703 | NO | NO | NA |
| 5430400D | ENSMUSG( | ENSMUSG( | -0.16173 | 0.844265 | NO | NO | NA |
| Hey1 | ENSMUSG( | ENSMUSG( | 0.04668 | 0.938123 | NO | NO | NA |
| Sowaha | ENSMUSG( | ENSMUSG( | 0.653208 | 0.345794 | NO | NO | NA |
| Gm6286 | ENSMUSG( | ENSMUSG( | 1.160411 | 0.289136 | NO | NO | NA |
| ENSMUSG | ENSMUSG( | ENSMUSG( | 0.834389 | 0.308438 | NO | NO | NA |
| Fgf18 | ENSMUSG( | ENSMUSG( | 0.692081 | 0.361227 | NO | NO | NA |
| Gm34294 | ENSMUSG( | ENSMUSG( | 0.591434 | 0.442731 | NO | NO | NA |
| Gm18782 | ENSMUSG( | ENSMUSG( | -0.84542 | 0.274342 | NO | NO | NA |
| 1700014B( | ENSMUSG( | ENSMUSG( | -0.06936 | 0.939403 | NO | NO | NA |
| Gm45507 | ENSMUSG( | ENSMUSG( | 1.347924 | 0.315372 | NO | NO | NA |
| Gpr153 | ENSMUSG( | ENSMUSG( | -0.23251 | 0.610556 | NO | NO | NA |
| Hecw2 | ENSMUSG( | ENSMUSG( | 0.259315 | 0.654278 | NO | NO | NA |
| Gm10244 | ENSMUSG( | ENSMUSG( | 0.320171 | 0.563166 | NO | NO | NA |
| Olfr1448 | ENSMUSG( | ENSMUSG( | 0.658339 | 0.540932 | NO | NO | NA |
| Gm18981 | ENSMUSG( | ENSMUSG( | 1.012541 | 0.451058 | NO | NO | NA |
| Gm3455 | ENSMUSG( | ENSMUSG( | 0.845144 | 0.436017 | NO | NO | NA |
| Slc22a17 | ENSMUSG( | ENSMUSG( | -0.91538 | 0.137848 | NO | NO | NA |
| Gm38047 | ENSMUSG( | ENSMUSG( | 0.293481 | 0.822027 | NO | NO | NA |
| Gm39318 | ENSMUSG( | ENSMUSG( | -0.21166 | 0.715635 | NO | NO | NA |

| | | | | | | | |
|---|---|---|---|---|---|---|---|
| Hmgb1-ps! | ENSMUSG( | ENSMUSG( | -0.75329 | 0.171822 | NO | NO | NA |
| D430040D | ENSMUSG( | ENSMUSG( | 0.460672 | 0.460605 | NO | NO | NA |
| Gm37675 | ENSMUSG( | ENSMUSG( | -0.08588 | 0.94868 | NO | NO | NA |
| ENSMUSG | ENSMUSG( | ENSMUSG( | -0.38807 | 0.56304 | NO | NO | NA |
| Gm38269 | ENSMUSG( | ENSMUSG( | 1.040648 | 0.380895 | NO | NO | NA |
| Reps2 | ENSMUSG( | ENSMUSG( | 0.498934 | 0.531711 | NO | NO | NA |
| Gm4221 | ENSMUSG( | ENSMUSG( | -0.03536 | 0.978105 | NO | NO | NA |
| ENSMUSG | ENSMUSG( | ENSMUSG( | 1.110786 | 0.211065 | NO | NO | NA |
| Kcnj12 | ENSMUSG( | ENSMUSG( | 0.759558 | 0.567401 | NO | NO | NA |
| AY074887 | ENSMUSG( | ENSMUSG( | 0.507557 | 0.627656 | NO | NO | NA |
| Gm6680 | ENSMUSG( | ENSMUSG( | 0.514957 | 0.676249 | NO | NO | NA |
| Rpl19-ps7 | ENSMUSG( | ENSMUSG( | 0.383124 | 0.665401 | NO | NO | NA |
| 1810021B: | ENSMUSG( | ENSMUSG( | 0.453142 | 0.373865 | NO | NO | NA |
| Ntn1 | ENSMUSG( | ENSMUSG( | 0.003411 | 0.996448 | NO | NO | NA |
| Gm26947 | ENSMUSG( | ENSMUSG( | 0.307458 | 0.740776 | NO | NO | NA |
| ENSMUSG | ENSMUSG( | ENSMUSG( | -0.14072 | 0.864457 | NO | NO | NA |
| Gm2628 | ENSMUSG( | ENSMUSG( | 0.937143 | 0.430908 | NO | NO | NA |
| Gm45757 | ENSMUSG( | ENSMUSG( | 1.507094 | 0.199499 | NO | NO | NA |
| Gm16401 | ENSMUSG( | ENSMUSG( | 0.484933 | 0.668578 | NO | NO | NA |
| Gm50273 | ENSMUSG( | ENSMUSG( | -0.75964 | 0.381121 | NO | NO | NA |
| Olfr1085 | ENSMUSG( | ENSMUSG( | 0.179054 | 0.896294 | NO | NO | NA |
| ENSMUSG( | ENSMUSG( | ENSMUSG( | -0.20997 | 0.80634 | NO | NO | NA |
| St8sia6 | ENSMUSG( | ENSMUSG( | 0.204606 | 0.834422 | NO | NO | NA |
| Gm47289 | ENSMUSG( | ENSMUSG( | 0.486098 | 0.675021 | NO | NO | NA |
| Gm37969 | ENSMUSG( | ENSMUSG( | 0.056162 | 0.932483 | NO | NO | NA |
| Zc2hc1c | ENSMUSG( | ENSMUSG( | -0.28421 | 0.514562 | NO | NO | NA |
| Gm16617 | ENSMUSG( | ENSMUSG( | 0.791098 | 0.500683 | NO | NO | NA |
| Gm15513 | ENSMUSG( | ENSMUSG( | 0.294414 | 0.653576 | NO | NO | NA |
| Gm28913 | ENSMUSG( | ENSMUSG( | -0.42905 | 0.612413 | NO | NO | NA |
| Prph | ENSMUSG( | ENSMUSG( | 1.112043 | 0.096182 | NO | NO | NA |
| Gm37334 | ENSMUSG( | ENSMUSG( | 0.192033 | 0.729736 | NO | NO | NA |
| Scel | ENSMUSG( | ENSMUSG( | 0.632455 | 0.423134 | NO | NO | NA |
| Arhgap24 | ENSMUSG( | ENSMUSG( | -0.16016 | 0.813165 | NO | NO | NA |
| Gpx8 | ENSMUSG( | ENSMUSG( | -0.74218 | 0.426056 | NO | NO | NA |
| Tpbgl | ENSMUSG( | ENSMUSG( | -1.34576 | 0.083367 | NO | NO | NA |
| Mak | ENSMUSG( | ENSMUSG( | 0.58832 | 0.670459 | NO | NO | NA |
| Gm15071 | ENSMUSG( | ENSMUSG( | 0.522571 | 0.656933 | NO | NO | NA |
| Gm48301 | ENSMUSG( | ENSMUSG( | 0.415977 | 0.730403 | NO | NO | NA |
| Il1rn | ENSMUSG( | ENSMUSG( | 1.101148 | 0.368166 | NO | NO | NA |
| Olfr1406 | ENSMUSG( | ENSMUSG( | -0.27591 | 0.812371 | NO | NO | NA |
| Gm9614 | ENSMUSG( | ENSMUSG( | 0.215741 | 0.854072 | NO | NO | NA |
| Gm15445 | ENSMUSG( | ENSMUSG( | -0.78903 | 0.11499 | NO | NO | NA |
| Gm43880 | ENSMUSG( | ENSMUSG( | 0.879991 | 0.464937 | NO | NO | NA |
| Myo5c | ENSMUSG( | ENSMUSG( | 0.617929 | 0.611373 | NO | NO | NA |
| Pdgfra | ENSMUSG( | ENSMUSG( | 0.336808 | 0.752633 | NO | NO | NA |
| Olfr1387 | ENSMUSG( | ENSMUSG( | 0.572207 | 0.557016 | NO | NO | NA |
| Gm7720 | ENSMUSG( | ENSMUSG( | 1.046706 | 0.4523 | NO | NO | NA |

| | | | | | | | |
|---|---|---|---|---|---|---|---|
| Plb1 | ENSMUSG( | ENSMUSG( | 0.649334 | 0.636296 | NO | NO | NA |
| Diras1 | ENSMUSG( | ENSMUSG( | -1.20348 | 0.45373 | NO | NO | NA |
| Gm38224 | ENSMUSG( | ENSMUSG( | 0.633411 | 0.544237 | NO | NO | NA |
| Lrrc4 | ENSMUSG( | ENSMUSG( | 0.064336 | 0.936437 | NO | NO | NA |
| Gm10764 | ENSMUSG( | ENSMUSG( | 0.57401 | 0.590846 | NO | NO | NA |
| 5730403I0 | ENSMUSG( | ENSMUSG( | 0.69889 | 0.538242 | NO | NO | NA |
| Gm29638 | ENSMUSG( | ENSMUSG( | -0.09627 | 0.940697 | NO | NO | NA |
| Gm45120 | ENSMUSG( | ENSMUSG( | -0.00494 | 0.994353 | NO | NO | NA |
| Ripor3 | ENSMUSG( | ENSMUSG( | -0.34399 | 0.502577 | NO | NO | NA |
| Pabpc4l | ENSMUSG( | ENSMUSG( | 0.7357 | 0.557168 | NO | NO | NA |
| Gm47232 | ENSMUSG( | ENSMUSG( | -0.66249 | 0.174565 | NO | NO | NA |
| Myoz2 | ENSMUSG( | ENSMUSG( | 1.303203 | 0.233306 | NO | NO | NA |
| F5 | ENSMUSG( | ENSMUSG( | 0.190836 | 0.87566 | NO | NO | NA |
| Gm11674 | ENSMUSG( | ENSMUSG( | 0.301178 | 0.809017 | NO | NO | NA |
| Kif21a | ENSMUSG( | ENSMUSG( | 0.166098 | 0.863994 | NO | NO | NA |
| 7530414M | ENSMUSG( | ENSMUSG( | 0.57571 | 0.771995 | NO | NO | NA |
| Gm47427 | ENSMUSG( | ENSMUSG( | 0.637073 | 0.476831 | NO | NO | NA |
| Gm16045 | ENSMUSG( | ENSMUSG( | -0.11515 | 0.901935 | NO | NO | NA |
| Gm15591 | ENSMUSG( | ENSMUSG( | 0.061848 | 0.930577 | NO | NO | NA |
| Gm45880 | ENSMUSG( | ENSMUSG( | 0.761845 | 0.522159 | NO | NO | NA |
| Gm27003 | ENSMUSG( | ENSMUSG( | -0.39002 | 0.505754 | NO | NO | NA |
| B430212C | ENSMUSG( | ENSMUSG( | 0.193241 | 0.859092 | NO | NO | NA |
| Gm48335 | ENSMUSG( | ENSMUSG( | 0.926418 | 0.438599 | NO | NO | NA |
| Mixl1 | ENSMUSG( | ENSMUSG( | -0.40305 | 0.664677 | NO | NO | NA |
| Fpr2 | ENSMUSG( | ENSMUSG( | 0.945886 | 0.27044 | NO | NO | NA |
| Gm36640 | ENSMUSG( | ENSMUSG( | 0.172087 | 0.909942 | NO | NO | NA |
| Gm50431 | ENSMUSG( | ENSMUSG( | 0.053659 | 0.914981 | NO | NO | NA |
| 9830107B | ENSMUSG( | ENSMUSG( | 1.193993 | 0.242611 | NO | NO | NA |
| Gm12352 | ENSMUSG( | ENSMUSG( | -0.14149 | 0.787014 | NO | NO | NA |
| Hmgb1-ps | ENSMUSG( | ENSMUSG( | -0.31626 | 0.584983 | NO | NO | NA |
| Igll1 | ENSMUSG( | ENSMUSG( | 0.279882 | 0.798585 | NO | NO | NA |
| Traj23 | ENSMUSG( | ENSMUSG( | 0.472823 | 0.636386 | NO | NO | NA |
| Gm43182 | ENSMUSG( | ENSMUSG( | 0.403291 | 0.574593 | NO | NO | NA |
| Ocln | ENSMUSG( | ENSMUSG( | 0.650668 | 0.448115 | NO | NO | NA |
| Olfr1138 | ENSMUSG( | ENSMUSG( | 0.693137 | 0.612867 | NO | NO | NA |
| Gm14914 | ENSMUSG( | ENSMUSG( | 1.662836 | 0.171513 | NO | NO | NA |
| Gm3468 | ENSMUSG( | ENSMUSG( | 0.389029 | 0.590627 | NO | NO | NA |
| Gm6218 | ENSMUSG( | ENSMUSG( | -0.20259 | 0.833242 | NO | NO | NA |
| Gm20760 | ENSMUSG( | ENSMUSG( | -2.61677 | 0.046642 | YES | YES | Downregulated |
| Gm35769 | ENSMUSG( | ENSMUSG( | 0.362216 | 0.741308 | NO | NO | NA |
| Nr2e3 | ENSMUSG( | ENSMUSG( | 1.003772 | 0.433722 | NO | NO | NA |
| Rpl13-ps6 | ENSMUSG( | ENSMUSG( | 0.171829 | 0.887695 | NO | NO | NA |
| Ugt1a1 | ENSMUSG( | ENSMUSG( | 0.329638 | NA | NO | NO | NA |
| Platr9 | ENSMUSG( | ENSMUSG( | -0.17888 | 0.895376 | NO | NO | NA |
| Gm3558 | ENSMUSG( | ENSMUSG( | 0.689591 | 0.525866 | NO | NO | NA |
| Vmn2r47 | ENSMUSG( | ENSMUSG( | 1.211497 | 0.565362 | NO | NO | NA |
| Gm50211 | ENSMUSG( | ENSMUSG( | 0.493131 | 0.487276 | NO | NO | NA |

| | | | | | | | |
|---|---|---|---|---|---|---|---|
| Gm18761 | ENSMUSG( | ENSMUSG( | 0.449618 | 0.731535 | NO | NO | NA |
| Gm43399 | ENSMUSG( | ENSMUSG( | 1.704488 | 0.124783 | NO | NO | NA |
| 2900027N | ENSMUSG( | ENSMUSG( | 0.520131 | 0.640961 | NO | NO | NA |
| Olfr57 | ENSMUSG( | ENSMUSG( | 0.106143 | 0.941997 | NO | NO | NA |
| Platr21 | ENSMUSG( | ENSMUSG( | 0.757664 | 0.567401 | NO | NO | NA |
| A430105J( | ENSMUSG( | ENSMUSG( | 0.403509 | 0.569951 | NO | NO | NA |
| Gm13559 | ENSMUSG( | ENSMUSG( | 0.575073 | 0.689779 | NO | NO | NA |
| Gm10399 | ENSMUSG( | ENSMUSG( | -0.15988 | 0.843245 | NO | NO | NA |
| Gm31517 | ENSMUSG( | ENSMUSG( | -0.40284 | 0.711913 | NO | NO | NA |
| Gm42461 | ENSMUSG( | ENSMUSG( | 0.679643 | 0.525866 | NO | NO | NA |
| Gm21844 | ENSMUSG( | ENSMUSG( | 1.378827 | 0.296468 | NO | NO | NA |
| Ighv14-3 | ENSMUSG( | ENSMUSG( | -4.26745 | NA | NO | NO | NA |
| Gm48267 | ENSMUSG( | ENSMUSG( | 0.45646 | 0.645285 | NO | NO | NA |
| Atp5pb-ps | ENSMUSG( | ENSMUSG( | 0.13402 | 0.822281 | NO | NO | NA |
| Kctd12b | ENSMUSG( | ENSMUSG( | -0.13862 | 0.819779 | NO | NO | NA |
| ENSMUSG( | ENSMUSG( | ENSMUSG( | 0.560925 | 0.658027 | NO | NO | NA |
| Gm36669 | ENSMUSG( | ENSMUSG( | 1.180994 | 0.308111 | NO | NO | NA |
| Tenm3 | ENSMUSG( | ENSMUSG( | 0.908296 | 0.389321 | NO | NO | NA |
| Mef2b | ENSMUSG( | ENSMUSG( | -0.44999 | 0.660508 | NO | NO | NA |
| Gm45548 | ENSMUSG( | ENSMUSG( | -0.32906 | 0.592341 | NO | NO | NA |
| Gm36120 | ENSMUSG( | ENSMUSG( | 0.85292 | 0.389686 | NO | NO | NA |
| ENSMUSG | ENSMUSG( | ENSMUSG( | 0.235288 | 0.773862 | NO | NO | NA |
| Cngb1 | ENSMUSG( | ENSMUSG( | 0.553397 | 0.693798 | NO | NO | NA |
| Gm37694 | ENSMUSG( | ENSMUSG( | 0.47153 | 0.652231 | NO | NO | NA |
| Hoxb3 | ENSMUSG( | ENSMUSG( | 0.060504 | 0.908692 | NO | NO | NA |
| Igfbpl1 | ENSMUSG( | ENSMUSG( | 0.292838 | 0.821463 | NO | NO | NA |
| Gm5976 | ENSMUSG( | ENSMUSG( | -0.31097 | 0.51761 | NO | NO | NA |
| Gm45212 | ENSMUSG( | ENSMUSG( | 0.352766 | 0.633247 | NO | NO | NA |
| Gm13919 | ENSMUSG( | ENSMUSG( | 0.517721 | 0.533291 | NO | NO | NA |
| 1700054O | ENSMUSG( | ENSMUSG( | 0.742895 | 0.57298 | NO | NO | NA |
| Ppp1ccb | ENSMUSG( | ENSMUSG( | 0.515085 | 0.826387 | NO | NO | NA |
| Gm15215 | ENSMUSG( | ENSMUSG( | 0.976212 | 0.37988 | NO | NO | NA |
| Trim30b | ENSMUSG( | ENSMUSG( | 0.029391 | 0.972995 | NO | NO | NA |
| ENSMUSG | ENSMUSG( | ENSMUSG( | 0.039593 | 0.938452 | NO | NO | NA |
| Gm32633 | ENSMUSG( | ENSMUSG( | -0.20429 | 0.771506 | NO | NO | NA |
| Platr28 | ENSMUSG( | ENSMUSG( | 0.509309 | 0.7281 | NO | NO | NA |
| Gm42943 | ENSMUSG( | ENSMUSG( | 0.559735 | 0.643515 | NO | NO | NA |
| Olfr837 | ENSMUSG( | ENSMUSG( | 0.861854 | 0.652883 | NO | NO | NA |
| Gm36974 | ENSMUSG( | ENSMUSG( | 1.057034 | 0.48793 | NO | NO | NA |
| ENSMUSG | ENSMUSG( | ENSMUSG( | -0.06374 | 0.898497 | NO | NO | NA |
| Gm12657 | ENSMUSG( | ENSMUSG( | 0.227237 | 0.71346 | NO | NO | NA |
| Gm48203 | ENSMUSG( | ENSMUSG( | 1.134333 | 0.34766 | NO | NO | NA |
| Hephl1 | ENSMUSG( | ENSMUSG( | 0.796543 | 0.514713 | NO | NO | NA |
| C230038L( | ENSMUSG( | ENSMUSG( | 1.112668 | 0.298164 | NO | NO | NA |
| Gm43693 | ENSMUSG( | ENSMUSG( | -0.19904 | 0.736727 | NO | NO | NA |
| 3110080O | ENSMUSG( | ENSMUSG( | 0.611439 | 0.636129 | NO | NO | NA |
| Cgnl1 | ENSMUSG( | ENSMUSG( | 0.759226 | 0.473866 | NO | NO | NA |

| | | | | | | | |
|---|---|---|---|---|---|---|---|
| Gm5447 | ENSMUSG( | ENSMUSG( | 0.619765 | 0.343744 | NO | NO | NA |
| Mtag2 | ENSMUSG( | ENSMUSG( | 0.257714 | 0.830763 | NO | NO | NA |
| Kcna4 | ENSMUSG( | ENSMUSG( | 0.642682 | 0.413484 | NO | NO | NA |
| 4933415A( | ENSMUSG( | ENSMUSG( | 0.110022 | 0.92736 | NO | NO | NA |
| Cd302 | ENSMUSG( | ENSMUSG( | -0.45164 | 0.545839 | NO | NO | NA |
| Rpl28-ps3 | ENSMUSG( | ENSMUSG( | 0.602264 | 0.440716 | NO | NO | NA |
| Appbp2os | ENSMUSG( | ENSMUSG( | 0.107955 | 0.899319 | NO | NO | NA |
| Gm38979 | ENSMUSG( | ENSMUSG( | -0.08683 | 0.858471 | NO | NO | NA |
| Wwtr1 | ENSMUSG( | ENSMUSG( | 0.464797 | 0.628512 | NO | NO | NA |
| Ajap1 | ENSMUSG( | ENSMUSG( | 0.588467 | 0.575942 | NO | NO | NA |
| Gm16184 | ENSMUSG( | ENSMUSG( | 0.492089 | 0.405253 | NO | NO | NA |
| Gm48804 | ENSMUSG( | ENSMUSG( | 1.167743 | 0.310179 | NO | NO | NA |
| Hoxc12 | ENSMUSG( | ENSMUSG( | 0.137558 | 0.906802 | NO | NO | NA |
| P4ha3 | ENSMUSG( | ENSMUSG( | 0.237828 | 0.840514 | NO | NO | NA |
| Gm26634 | ENSMUSG( | ENSMUSG( | -0.70819 | 0.534393 | NO | NO | NA |
| ENSMUSG | ENSMUSG( | ENSMUSG( | 0.757934 | 0.444377 | NO | NO | NA |
| Coro2b | ENSMUSG( | ENSMUSG( | -1.05498 | 0.126781 | NO | NO | NA |
| Gm26653 | ENSMUSG( | ENSMUSG( | 1.080337 | 0.50674 | NO | NO | NA |
| Gm42917 | ENSMUSG( | ENSMUSG( | -0.07213 | 0.956971 | NO | NO | NA |
| Rab7-ps1 | ENSMUSG( | ENSMUSG( | -0.25498 | 0.735932 | NO | NO | NA |
| Gm37897 | ENSMUSG( | ENSMUSG( | 0.621373 | 0.647827 | NO | NO | NA |
| Gm32624 | ENSMUSG( | ENSMUSG( | 0.479442 | 0.759918 | NO | NO | NA |
| Gm44753 | ENSMUSG( | ENSMUSG( | 0.423106 | 0.679132 | NO | NO | NA |
| Olfr173 | ENSMUSG( | ENSMUSG( | -0.37184 | 0.555025 | NO | NO | NA |
| Tcf24 | ENSMUSG( | ENSMUSG( | -0.12046 | 0.929293 | NO | NO | NA |
| Gm16618 | ENSMUSG( | ENSMUSG( | 0.375536 | 0.544716 | NO | NO | NA |
| Gm11732 | ENSMUSG( | ENSMUSG( | 0.602885 | 0.632323 | NO | NO | NA |
| Plxna4os1 | ENSMUSG( | ENSMUSG( | 0.996628 | 0.374748 | NO | NO | NA |
| Gm26807 | ENSMUSG( | ENSMUSG( | 0.150889 | 0.90825 | NO | NO | NA |
| ENSMUSG( | ENSMUSG( | ENSMUSG( | 0.143526 | 0.91013 | NO | NO | NA |
| Gm42922 | ENSMUSG( | ENSMUSG( | 0.151688 | 0.840661 | NO | NO | NA |
| Gm46355 | ENSMUSG( | ENSMUSG( | 0.292577 | 0.80032 | NO | NO | NA |
| Spaca1 | ENSMUSG( | ENSMUSG( | 0.499808 | 0.509107 | NO | NO | NA |
| Rpl27a-ps1 | ENSMUSG( | ENSMUSG( | -0.6765 | 0.371982 | NO | NO | NA |
| Myct1 | ENSMUSG( | ENSMUSG( | 0.417079 | 0.667955 | NO | NO | NA |
| Gm24950 | ENSMUSG( | ENSMUSG( | 1.096425 | 0.749922 | NO | NO | NA |
| Gm6034 | ENSMUSG( | ENSMUSG( | -0.7686 | 0.174455 | NO | NO | NA |
| Gm15868 | ENSMUSG( | ENSMUSG( | -0.43881 | 0.518248 | NO | NO | NA |
| Gm38106 | ENSMUSG( | ENSMUSG( | 0.908947 | 0.353488 | NO | NO | NA |
| C1ql1 | ENSMUSG( | ENSMUSG( | 0.651527 | 0.414327 | NO | NO | NA |
| Olfr1324 | ENSMUSG( | ENSMUSG( | 0.340886 | 0.810767 | NO | NO | NA |
| Fhip1a | ENSMUSG( | ENSMUSG( | -0.70852 | 0.519563 | NO | NO | NA |
| Olfr315 | ENSMUSG( | ENSMUSG( | -0.3558 | 0.840237 | NO | NO | NA |
| Hmcn1 | ENSMUSG( | ENSMUSG( | -0.02958 | 0.978405 | NO | NO | NA |
| Gm48342 | ENSMUSG( | ENSMUSG( | 0.424038 | 0.512102 | NO | NO | NA |
| Gpr15 | ENSMUSG( | ENSMUSG( | -0.12699 | 0.917548 | NO | NO | NA |
| Abca15 | ENSMUSG( | ENSMUSG( | 1.648712 | 0.231083 | NO | NO | NA |

| | | | | | | | |
|---|---|---|---|---|---|---|---|
| Gm15759 | ENSMUSG( | ENSMUSG( | 0.600407 | 0.432246 | NO | NO | NA |
| 9530018H | ENSMUSG( | ENSMUSG( | 1.310971 | 0.365388 | NO | NO | NA |
| Tmc3 | ENSMUSG( | ENSMUSG( | 0.046497 | 0.96437 | NO | NO | NA |
| Trbj2-4 | ENSMUSG( | ENSMUSG( | -0.41296 | 0.564209 | NO | NO | NA |
| Btbd17 | ENSMUSG( | ENSMUSG( | 0.809831 | 0.356637 | NO | NO | NA |
| Grm3 | ENSMUSG( | ENSMUSG( | 1.66722 | 0.243746 | NO | NO | NA |
| Gm49921 | ENSMUSG( | ENSMUSG( | 0.646818 | 0.668767 | NO | NO | NA |
| Gm37662 | ENSMUSG( | ENSMUSG( | 1.032087 | 0.422217 | NO | NO | NA |
| Gm14448 | ENSMUSG( | ENSMUSG( | 0.4525 | 0.549432 | NO | NO | NA |
| ENSMUSG | ENSMUSG( | ENSMUSG( | 0.633365 | 0.507567 | NO | NO | NA |
| Prap1 | ENSMUSG( | ENSMUSG( | 0.406663 | 0.741426 | NO | NO | NA |
| Kif26b | ENSMUSG( | ENSMUSG( | 0.146946 | 0.868739 | NO | NO | NA |
| Nxph4 | ENSMUSG( | ENSMUSG( | 0.617432 | 0.683107 | NO | NO | NA |
| Gm43465 | ENSMUSG( | ENSMUSG( | 0.315542 | 0.814086 | NO | NO | NA |
| Gm4034 | ENSMUSG( | ENSMUSG( | 1.068413 | 0.401018 | NO | NO | NA |
| Grtp1 | ENSMUSG( | ENSMUSG( | -0.03828 | 0.956941 | NO | NO | NA |
| Gm49887 | ENSMUSG( | ENSMUSG( | -0.03313 | 0.974434 | NO | NO | NA |
| Pla2g3 | ENSMUSG( | ENSMUSG( | -0.03121 | 0.958713 | NO | NO | NA |
| Slc26a8 | ENSMUSG( | ENSMUSG( | 0.535867 | 0.358484 | NO | NO | NA |
| Gm48232 | ENSMUSG( | ENSMUSG( | 0.730598 | 0.606154 | NO | NO | NA |
| Crb2 | ENSMUSG( | ENSMUSG( | 0.044289 | 0.974234 | NO | NO | NA |
| Gm6542 | ENSMUSG( | ENSMUSG( | 0.2739 | 0.624516 | NO | NO | NA |
| Gm43884 | ENSMUSG( | ENSMUSG( | 1.005403 | 0.501622 | NO | NO | NA |
| Gm11928 | ENSMUSG( | ENSMUSG( | 0.105304 | 0.907759 | NO | NO | NA |
| Gm13056 | ENSMUSG( | ENSMUSG( | -0.02486 | 0.983174 | NO | NO | NA |
| Gm13168 | ENSMUSG( | ENSMUSG( | -1.54214 | 0.301461 | NO | NO | NA |
| Gm12324 | ENSMUSG( | ENSMUSG( | 1.229146 | 0.254604 | NO | NO | NA |
| Gm49254 | ENSMUSG( | ENSMUSG( | 0.848619 | 0.517571 | NO | NO | NA |
| 4930405A | ENSMUSG( | ENSMUSG( | 0.423431 | 0.614734 | NO | NO | NA |
| Gm2136 | ENSMUSG( | ENSMUSG( | 0.391324 | 0.729989 | NO | NO | NA |
| Gm40770 | ENSMUSG( | ENSMUSG( | 0.162223 | 0.888811 | NO | NO | NA |
| 4930546C | ENSMUSG( | ENSMUSG( | 0.98962 | 0.515032 | NO | NO | NA |
| Gm15204 | ENSMUSG( | ENSMUSG( | -0.49515 | 0.624821 | NO | NO | NA |
| Vmn2r56 | ENSMUSG( | ENSMUSG( | 0.191688 | 0.890997 | NO | NO | NA |
| Gm8267 | ENSMUSG( | ENSMUSG( | 0.43822 | 0.73702 | NO | NO | NA |
| Olfr46 | ENSMUSG( | ENSMUSG( | 0.23666 | 0.821463 | NO | NO | NA |
| Clec2g | ENSMUSG( | ENSMUSG( | 0.00199 | 0.998635 | NO | NO | NA |
| Fitm1 | ENSMUSG( | ENSMUSG( | 0.063757 | 0.956997 | NO | NO | NA |
| 2810407A | ENSMUSG( | ENSMUSG( | 0.98984 | 0.416871 | NO | NO | NA |
| Gm13303 | ENSMUSG( | ENSMUSG( | 1.409808 | NA | NO | NO | NA |
| Zfp786 | ENSMUSG( | ENSMUSG( | 0.526022 | 0.675324 | NO | NO | NA |
| Nrg3 | ENSMUSG( | ENSMUSG( | 0.39824 | 0.746663 | NO | NO | NA |
| Gm42737 | ENSMUSG( | ENSMUSG( | 0.546441 | 0.707602 | NO | NO | NA |
| Vgll3 | ENSMUSG( | ENSMUSG( | 0.714546 | 0.562508 | NO | NO | NA |
| Gm12829 | ENSMUSG( | ENSMUSG( | -0.01126 | 0.993293 | NO | NO | NA |
| Slc44a3 | ENSMUSG( | ENSMUSG( | 0.309034 | 0.785273 | NO | NO | NA |
| Gm8637 | ENSMUSG( | ENSMUSG( | 0.927015 | 0.546603 | NO | NO | NA |

| | | | | | | | |
|---|---|---|---|---|---|---|---|
| ENSMUSG | ENSMUSG( | ENSMUSG( | 0.111229 | 0.871801 | NO | NO | NA |
| Gm34780 | ENSMUSG( | ENSMUSG( | 1.129284 | 0.384649 | NO | NO | NA |
| Megf10 | ENSMUSG( | ENSMUSG( | 0.79682 | 0.548849 | NO | NO | NA |
| Gm47011 | ENSMUSG( | ENSMUSG( | 0.968731 | 0.435651 | NO | NO | NA |
| Gm4022 | ENSMUSG( | ENSMUSG( | 1.400666 | 0.15939 | NO | NO | NA |
| Wnk2 | ENSMUSG( | ENSMUSG( | 0.387447 | 0.66593 | NO | NO | NA |
| Aplnr | ENSMUSG( | ENSMUSG( | 0.805804 | 0.48915 | NO | NO | NA |
| Gm10653 | ENSMUSG( | ENSMUSG( | 4.671384 | NA | NO | NO | NA |
| Gm43562 | ENSMUSG( | ENSMUSG( | 4.671384 | NA | NO | NO | NA |
| Rnu1a1 | ENSMUSG( | ENSMUSG( | 4.790284 | NA | NO | NO | NA |
| Gm44684 | ENSMUSG( | ENSMUSG( | 1.06834 | 0.328444 | NO | NO | NA |
| Olfr1029 | ENSMUSG( | ENSMUSG( | 0.266168 | 0.802041 | NO | NO | NA |
| Vmn1r-ps1 | ENSMUSG( | ENSMUSG( | 1.567748 | 0.165825 | NO | NO | NA |
| Gm42971 | ENSMUSG( | ENSMUSG( | 0.424764 | 0.712916 | NO | NO | NA |
| A430090L1 | ENSMUSG( | ENSMUSG( | 0.081874 | 0.898004 | NO | NO | NA |
| Gm4787 | ENSMUSG( | ENSMUSG( | -0.6869 | 0.299612 | NO | NO | NA |
| Gm49890 | ENSMUSG( | ENSMUSG( | 0.832118 | 0.276671 | NO | NO | NA |
| Gng13 | ENSMUSG( | ENSMUSG( | 0.782572 | 0.356637 | NO | NO | NA |
| Olfr1181 | ENSMUSG( | ENSMUSG( | 0.521946 | 0.667069 | NO | NO | NA |
| 4930563E2 | ENSMUSG( | ENSMUSG( | 0.046232 | 0.954162 | NO | NO | NA |
| Clstn3 | ENSMUSG( | ENSMUSG( | 0.791743 | 0.544716 | NO | NO | NA |
| ENSMUSG | ENSMUSG( | ENSMUSG( | 0.184616 | 0.761874 | NO | NO | NA |
| Rps11-ps2 | ENSMUSG( | ENSMUSG( | 0.245782 | 0.692686 | NO | NO | NA |
| Gm44113 | ENSMUSG( | ENSMUSG( | 0.456142 | 0.749636 | NO | NO | NA |
| Gm44104 | ENSMUSG( | ENSMUSG( | 0.016383 | 0.986912 | NO | NO | NA |
| Onecut1 | ENSMUSG( | ENSMUSG( | 0.444666 | 0.711914 | NO | NO | NA |
| ENSMUSG | ENSMUSG( | ENSMUSG( | 0.128174 | 0.907201 | NO | NO | NA |
| Emid1 | ENSMUSG( | ENSMUSG( | 0.145113 | 0.781929 | NO | NO | NA |
| Gm44511 | ENSMUSG( | ENSMUSG( | -0.06597 | 0.982586 | NO | NO | NA |
| Gm49706 | ENSMUSG( | ENSMUSG( | 0.179529 | 0.769716 | NO | NO | NA |
| Gm10640 | ENSMUSG( | ENSMUSG( | 0.482783 | 0.544688 | NO | NO | NA |
| Dsc2 | ENSMUSG( | ENSMUSG( | 0.698684 | 0.56435 | NO | NO | NA |
| Gm5846 | ENSMUSG( | ENSMUSG( | -0.03006 | 0.961527 | NO | NO | NA |
| Gm29994 | ENSMUSG( | ENSMUSG( | 0.519775 | 0.550555 | NO | NO | NA |
| Clec4e | ENSMUSG( | ENSMUSG( | 0.132866 | 0.92032 | NO | NO | NA |
| Gm17399 | ENSMUSG( | ENSMUSG( | -0.32463 | 0.695672 | NO | NO | NA |
| ENSMUSG | ENSMUSG( | ENSMUSG( | 0.817567 | 0.22713 | NO | NO | NA |
| Fam187b | ENSMUSG( | ENSMUSG( | 0.301382 | 0.618409 | NO | NO | NA |
| Tmed10-p | ENSMUSG( | ENSMUSG( | 0.102183 | 0.865515 | NO | NO | NA |
| 4930511M | ENSMUSG( | ENSMUSG( | 1.321166 | 0.321959 | NO | NO | NA |
| Trim30c | ENSMUSG( | ENSMUSG( | -1.57115 | 0.109955 | NO | NO | NA |
| Gm43447 | ENSMUSG( | ENSMUSG( | -0.96976 | 0.541669 | NO | NO | NA |
| Gm44021 | ENSMUSG( | ENSMUSG( | 0.452556 | 0.656628 | NO | NO | NA |
| Trpc7 | ENSMUSG( | ENSMUSG( | 0.039389 | 0.974234 | NO | NO | NA |
| Hspa12b | ENSMUSG( | ENSMUSG( | 0.454294 | 0.492834 | NO | NO | NA |
| Gm5555 | ENSMUSG( | ENSMUSG( | -0.21571 | 0.699514 | NO | NO | NA |
| Gm7560 | ENSMUSG( | ENSMUSG( | 0.508775 | 0.716497 | NO | NO | NA |

| | | | | | | | |
|---|---|---|---|---|---|---|---|
| Gm37629 | ENSMUSG( | ENSMUSG( | 0.17176 | 0.889318 | NO | NO | NA |
| Adgra2 | ENSMUSG( | ENSMUSG( | -0.68468 | 0.281466 | NO | NO | NA |
| Tcfl5 | ENSMUSG( | ENSMUSG( | 0.11379 | 0.878742 | NO | NO | NA |
| Olfr755-ps | ENSMUSG( | ENSMUSG( | -1.95978 | 0.055847 | NO | NO | NA |
| ENSMUSG( | ENSMUSG( | ENSMUSG( | 0.192954 | 0.795139 | NO | NO | NA |
| 4930550C: | ENSMUSG( | ENSMUSG( | 0.583726 | 0.399515 | NO | NO | NA |
| Gm37768 | ENSMUSG( | ENSMUSG( | -0.24659 | 0.653348 | NO | NO | NA |
| Padi4 | ENSMUSG( | ENSMUSG( | 0.218371 | 0.759335 | NO | NO | NA |
| Galnt18 | ENSMUSG( | ENSMUSG( | 1.446206 | 0.235128 | NO | NO | NA |
| Efhb | ENSMUSG( | ENSMUSG( | 1.11081 | 0.335106 | NO | NO | NA |
| Mamstr | ENSMUSG( | ENSMUSG( | 0.530706 | 0.679132 | NO | NO | NA |
| Aph1c | ENSMUSG( | ENSMUSG( | 0.416993 | 0.562501 | NO | NO | NA |
| Gm48273 | ENSMUSG( | ENSMUSG( | -0.10189 | 0.914861 | NO | NO | NA |
| Kitl | ENSMUSG( | ENSMUSG( | 1.724035 | 0.001801 | YES | YES | Upregulated |
| Apol9b | ENSMUSG( | ENSMUSG( | -0.2461 | 0.696107 | NO | NO | NA |
| Gm15998 | ENSMUSG( | ENSMUSG( | -0.02571 | 0.984035 | NO | NO | NA |
| 1700016A( | ENSMUSG( | ENSMUSG( | -1.05028 | 0.117937 | NO | NO | NA |
| Gm9923 | ENSMUSG( | ENSMUSG( | 0.14857 | 0.85211 | NO | NO | NA |
| Gm36958 | ENSMUSG( | ENSMUSG( | 0.002376 | 0.997831 | NO | NO | NA |
| Esr2 | ENSMUSG( | ENSMUSG( | 0.516919 | 0.682588 | NO | NO | NA |
| Abca9 | ENSMUSG( | ENSMUSG( | 0.637482 | 0.54328 | NO | NO | NA |
| Gm7470 | ENSMUSG( | ENSMUSG( | 1.108061 | 0.412202 | NO | NO | NA |
| Clnk | ENSMUSG( | ENSMUSG( | 1.416292 | 0.028057 | YES | YES | Upregulated |
| ENSMUSG | ENSMUSG( | ENSMUSG( | 0.557387 | 0.731673 | NO | NO | NA |
| Mag | ENSMUSG( | ENSMUSG( | 1.156637 | 0.174565 | NO | NO | NA |
| Gm15749 | ENSMUSG( | ENSMUSG( | 0.897777 | 0.459033 | NO | NO | NA |
| 1700016H | ENSMUSG( | ENSMUSG( | 0.152992 | 0.907741 | NO | NO | NA |
| Gm18867 | ENSMUSG( | ENSMUSG( | 1.30672 | 0.347295 | NO | NO | NA |
| Gm42437 | ENSMUSG( | ENSMUSG( | 0.668899 | 0.505568 | NO | NO | NA |
| Gm53038 | ENSMUSG( | ENSMUSG( | -0.06262 | 0.915968 | NO | NO | NA |
| Gm5987 | ENSMUSG( | ENSMUSG( | -0.33777 | 0.773862 | NO | NO | NA |
| Gm44627 | ENSMUSG( | ENSMUSG( | 0.00837 | 0.994734 | NO | NO | NA |
| Gm49604 | ENSMUSG( | ENSMUSG( | 0.28917 | 0.74568 | NO | NO | NA |
| Pde6g | ENSMUSG( | ENSMUSG( | -0.72337 | 0.471757 | NO | NO | NA |
| Platr10 | ENSMUSG( | ENSMUSG( | 0.413246 | 0.690616 | NO | NO | NA |
| Mypn | ENSMUSG( | ENSMUSG( | 0.493831 | 0.636821 | NO | NO | NA |
| Gm42498 | ENSMUSG( | ENSMUSG( | 0.977235 | 0.427616 | NO | NO | NA |
| Gm43051 | ENSMUSG( | ENSMUSG( | -0.07306 | 0.952073 | NO | NO | NA |
| Gm5697 | ENSMUSG( | ENSMUSG( | 0.533919 | 0.5601 | NO | NO | NA |
| Olfr250 | ENSMUSG( | ENSMUSG( | 0.886251 | 0.533787 | NO | NO | NA |
| Gm15901 | ENSMUSG( | ENSMUSG( | 2.403203 | 0.06466 | NO | NO | NA |
| Arhgap27c | ENSMUSG( | ENSMUSG( | 1.069763 | 0.236728 | NO | NO | NA |
| Gm30239 | ENSMUSG( | ENSMUSG( | 0.930796 | 0.57629 | NO | NO | NA |
| Gm9570 | ENSMUSG( | ENSMUSG( | 0.108694 | 0.911394 | NO | NO | NA |
| Cachd1 | ENSMUSG( | ENSMUSG( | 0.601472 | 0.521932 | NO | NO | NA |
| Gm34590 | ENSMUSG( | ENSMUSG( | -0.0137 | 0.986185 | NO | NO | NA |
| Gm17826 | ENSMUSG( | ENSMUSG( | 0.969753 | 0.330985 | NO | NO | NA |

| | | | | | | | |
|---|---|---|---|---|---|---|---|
| Cyp2b23 | ENSMUSG( | ENSMUSG( | 0.740658 | 0.572224 | NO | NO | NA |
| Kcnh4 | ENSMUSG( | ENSMUSG( | 0.036435 | 0.973749 | NO | NO | NA |
| Gm3321 | ENSMUSG( | ENSMUSG( | 0.742 | 0.661527 | NO | NO | NA |
| Slc2a7 | ENSMUSG( | ENSMUSG( | 0.201715 | 0.878742 | NO | NO | NA |
| ENSMUSG | ENSMUSG( | ENSMUSG( | -0.56582 | 0.412993 | NO | NO | NA |
| Gm42571 | ENSMUSG( | ENSMUSG( | -0.29149 | 0.648873 | NO | NO | NA |
| Gstt4 | ENSMUSG( | ENSMUSG( | 0.619564 | 0.603586 | NO | NO | NA |
| Spaca6 | ENSMUSG( | ENSMUSG( | 0.036513 | 0.96163 | NO | NO | NA |
| Adgra1 | ENSMUSG( | ENSMUSG( | 0.663417 | 0.575418 | NO | NO | NA |
| Gm6141 | ENSMUSG( | ENSMUSG( | -0.29821 | 0.832745 | NO | NO | NA |
| Gm19131 | ENSMUSG( | ENSMUSG( | 1.210568 | 0.671934 | NO | NO | NA |
| Elfn1 | ENSMUSG( | ENSMUSG( | 0.514397 | 0.628512 | NO | NO | NA |
| Lhfpl5 | ENSMUSG( | ENSMUSG( | -0.07004 | 0.942764 | NO | NO | NA |
| Gm44678 | ENSMUSG( | ENSMUSG( | 0.869987 | 0.484806 | NO | NO | NA |
| 4930488L2 | ENSMUSG( | ENSMUSG( | 0.387868 | 0.732409 | NO | NO | NA |
| Gm42432 | ENSMUSG( | ENSMUSG( | 0.001987 | 0.996944 | NO | NO | NA |
| Gm42830 | ENSMUSG( | ENSMUSG( | 1.027357 | 0.318119 | NO | NO | NA |
| ENSMUSG( | ENSMUSG( | ENSMUSG( | 5.444808 | 0.151614 | NO | NO | NA |
| Gm18384 | ENSMUSG( | ENSMUSG( | 0.897734 | 0.504087 | NO | NO | NA |
| Grik1 | ENSMUSG( | ENSMUSG( | 0.842799 | 0.545865 | NO | NO | NA |
| Gm37315 | ENSMUSG( | ENSMUSG( | 0.494678 | 0.712298 | NO | NO | NA |
| Gm47171 | ENSMUSG( | ENSMUSG( | 0.786374 | 0.386187 | NO | NO | NA |
| Lexm | ENSMUSG( | ENSMUSG( | 1.250321 | 0.087339 | NO | NO | NA |
| Tpbpa | ENSMUSG( | ENSMUSG( | 0.945985 | 0.438442 | NO | NO | NA |
| Gm43686 | ENSMUSG( | ENSMUSG( | 0.458386 | 0.640929 | NO | NO | NA |
| Fat2 | ENSMUSG( | ENSMUSG( | 0.559524 | 0.65572 | NO | NO | NA |
| 4930502E( | ENSMUSG( | ENSMUSG( | 0.492629 | 0.723598 | NO | NO | NA |
| Adam7 | ENSMUSG( | ENSMUSG( | 0.788011 | 0.454656 | NO | NO | NA |
| Olig3 | ENSMUSG( | ENSMUSG( | 0.295847 | 0.78459 | NO | NO | NA |
| Gm37602 | ENSMUSG( | ENSMUSG( | 0.056864 | 0.962787 | NO | NO | NA |
| Gm48655 | ENSMUSG( | ENSMUSG( | -0.23105 | 0.784462 | NO | NO | NA |
| Gm43557 | ENSMUSG( | ENSMUSG( | 1.060818 | 0.429224 | NO | NO | NA |
| Gm45215 | ENSMUSG( | ENSMUSG( | 5.18102 | 1.95E-05 | YES | YES | Upregulated |
| Rpl10-ps5 | ENSMUSG( | ENSMUSG( | -0.0759 | 0.912705 | NO | NO | NA |
| 9530052E( | ENSMUSG( | ENSMUSG( | 0.646946 | 0.471631 | NO | NO | NA |
| Sptbn4 | ENSMUSG( | ENSMUSG( | -0.34567 | 0.603367 | NO | NO | NA |
| Gm3466 | ENSMUSG( | ENSMUSG( | -2.69749 | 0.157181 | NO | NO | NA |
| Prokr1 | ENSMUSG( | ENSMUSG( | 0.414659 | 0.658438 | NO | NO | NA |
| Zar1 | ENSMUSG( | ENSMUSG( | 0.093977 | 0.946702 | NO | NO | NA |
| Dchs1 | ENSMUSG( | ENSMUSG( | -0.93348 | 0.258394 | NO | NO | NA |
| Gm5373 | ENSMUSG( | ENSMUSG( | 0.653235 | 0.6204 | NO | NO | NA |
| Gm11984 | ENSMUSG( | ENSMUSG( | 0.678788 | 0.579407 | NO | NO | NA |
| Gm27021 | ENSMUSG( | ENSMUSG( | 0.441789 | 0.715041 | NO | NO | NA |
| Aatk | ENSMUSG( | ENSMUSG( | 0.211177 | 0.85302 | NO | NO | NA |
| Gm43168 | ENSMUSG( | ENSMUSG( | 0.497817 | 0.534067 | NO | NO | NA |
| Lhx5 | ENSMUSG( | ENSMUSG( | -0.08417 | 0.942899 | NO | NO | NA |
| 2210010C( | ENSMUSG( | ENSMUSG( | 0.694535 | 0.417592 | NO | NO | NA |

| | | | | | | | |
|---|---|---|---|---|---|---|---|
| Gm4978 | ENSMUSG( | ENSMUSG( | 0.112391 | 0.835507 | NO | NO | NA |
| Amy2a3 | ENSMUSG( | ENSMUSG( | -2.11249 | NA | NO | NO | NA |
| Aknad1 | ENSMUSG( | ENSMUSG( | -0.07174 | 0.941476 | NO | NO | NA |
| Pin4-ps | ENSMUSG( | ENSMUSG( | 0.422259 | 0.474362 | NO | NO | NA |
| Gm16085 | ENSMUSG( | ENSMUSG( | -0.99856 | 0.089446 | NO | NO | NA |
| Gm37166 | ENSMUSG( | ENSMUSG( | -4.43689 | 2.33E-05 | YES | YES | Downregulated |
| Gm3375 | ENSMUSG( | ENSMUSG( | 0.263093 | 0.623149 | NO | NO | NA |
| Hspb9 | ENSMUSG( | ENSMUSG( | 0.260557 | 0.650855 | NO | NO | NA |
| Vmn2r90 | ENSMUSG( | ENSMUSG( | 0.554048 | 0.637463 | NO | NO | NA |
| Gal3st3 | ENSMUSG( | ENSMUSG( | 0.243986 | 0.832943 | NO | NO | NA |
| Trpd52l3 | ENSMUSG( | ENSMUSG( | 0.726428 | 0.644413 | NO | NO | NA |
| Trgv1 | ENSMUSG( | ENSMUSG( | -0.16071 | 0.80971 | NO | NO | NA |
| Adam33 | ENSMUSG( | ENSMUSG( | 1.450552 | 0.052139 | NO | NO | NA |
| Lepr | ENSMUSG( | ENSMUSG( | 0.915395 | 0.334697 | NO | NO | NA |
| Utp14b-ps | ENSMUSG( | ENSMUSG( | 0.59763 | 0.665416 | NO | NO | NA |
| Apom | ENSMUSG( | ENSMUSG( | 0.805684 | 0.098313 | NO | NO | NA |
| Gm26608 | ENSMUSG( | ENSMUSG( | 0.62054 | 0.71644 | NO | NO | NA |
| Gm43663 | ENSMUSG( | ENSMUSG( | 0.781405 | 0.467096 | NO | NO | NA |
| BC037704 | ENSMUSG( | ENSMUSG( | 0.599444 | 0.254696 | NO | NO | NA |
| Mup9 | ENSMUSG( | ENSMUSG( | 0.457284 | 0.738771 | NO | NO | NA |
| A330008L1 | ENSMUSG( | ENSMUSG( | 0.730929 | 0.521932 | NO | NO | NA |
| Vmn2r4 | ENSMUSG( | ENSMUSG( | 0.723981 | 0.534475 | NO | NO | NA |
| Vwa2 | ENSMUSG( | ENSMUSG( | 1.742607 | 0.007941 | YES | YES | Upregulated |
| Gm43765 | ENSMUSG( | ENSMUSG( | 1.265059 | 0.378105 | NO | NO | NA |
| Olfr678 | ENSMUSG( | ENSMUSG( | 0.211877 | 0.864929 | NO | NO | NA |
| Cttnbp2 | ENSMUSG( | ENSMUSG( | -0.47883 | 0.62774 | NO | NO | NA |
| ENSMUSG | ENSMUSG( | ENSMUSG( | 0.466875 | 0.548323 | NO | NO | NA |
| Gm46378 | ENSMUSG( | ENSMUSG( | -0.6542 | 0.29433 | NO | NO | NA |
| A530016L2 | ENSMUSG( | ENSMUSG( | 1.034673 | 0.357367 | NO | NO | NA |
| Stx1b | ENSMUSG( | ENSMUSG( | -0.33916 | 0.838501 | NO | NO | NA |
| Gm30117 | ENSMUSG( | ENSMUSG( | 0.158849 | 0.873681 | NO | NO | NA |
| Gm13562 | ENSMUSG( | ENSMUSG( | -0.50022 | 0.390657 | NO | NO | NA |
| Cabp1 | ENSMUSG( | ENSMUSG( | 0.603151 | 0.286864 | NO | NO | NA |
| Rpsa-ps5 | ENSMUSG( | ENSMUSG( | 0.410319 | 0.626538 | NO | NO | NA |
| Gm38344 | ENSMUSG( | ENSMUSG( | 1.441212 | 0.359969 | NO | NO | NA |
| Lama2 | ENSMUSG( | ENSMUSG( | 0.543863 | 0.666874 | NO | NO | NA |
| Actg-ps1 | ENSMUSG( | ENSMUSG( | -0.32184 | 0.503722 | NO | NO | NA |
| Gm37122 | ENSMUSG( | ENSMUSG( | 1.110409 | 0.349957 | NO | NO | NA |
| Gm49804 | ENSMUSG( | ENSMUSG( | 1.819278 | 0.582357 | NO | NO | NA |
| Gm45130 | ENSMUSG( | ENSMUSG( | 0.558994 | 0.625372 | NO | NO | NA |
| Adap2 | ENSMUSG( | ENSMUSG( | -0.42627 | 0.543023 | NO | NO | NA |
| Emp2 | ENSMUSG( | ENSMUSG( | 0.201242 | 0.883479 | NO | NO | NA |
| Slc16a5 | ENSMUSG( | ENSMUSG( | -1.01656 | 0.244982 | NO | NO | NA |
| Gm4926 | ENSMUSG( | ENSMUSG( | 1.60917 | 0.178411 | NO | NO | NA |
| Kazald1 | ENSMUSG( | ENSMUSG( | 0.926048 | 0.118998 | NO | NO | NA |
| Ccdc92b | ENSMUSG( | ENSMUSG( | 0.400177 | 0.681566 | NO | NO | NA |
| P2rx3 | ENSMUSG( | ENSMUSG( | 0.169503 | 0.784462 | NO | NO | NA |

| | | | | | | | |
|---|---|---|---|---|---|---|---|
| H2bu1-ps | ENSMUSG( | ENSMUSG( | 0.646536 | 0.256835 | NO | NO | NA |
| Gm2962 | ENSMUSG( | ENSMUSG( | -0.34592 | 0.479553 | NO | NO | NA |
| Pros1 | ENSMUSG( | ENSMUSG( | 0.253244 | 0.721638 | NO | NO | NA |
| Gm14049 | ENSMUSG( | ENSMUSG( | 0.342667 | 0.719466 | NO | NO | NA |
| Gm37651 | ENSMUSG( | ENSMUSG( | 1.327947 | 0.110414 | NO | NO | NA |
| Gm38973 | ENSMUSG( | ENSMUSG( | 0.759048 | 0.216209 | NO | NO | NA |
| Gm20420 | ENSMUSG( | ENSMUSG( | -0.11962 | 0.899457 | NO | NO | NA |
| Reln | ENSMUSG( | ENSMUSG( | 0.154347 | 0.863022 | NO | NO | NA |
| Olfr1253 | ENSMUSG( | ENSMUSG( | 0.317611 | 0.788624 | NO | NO | NA |
| Treml4 | ENSMUSG( | ENSMUSG( | -0.88222 | 0.424867 | NO | NO | NA |
| 4732491K: | ENSMUSG( | ENSMUSG( | 0.408912 | 0.5656 | NO | NO | NA |
| Gm47801 | ENSMUSG( | ENSMUSG( | 0.402042 | 0.58564 | NO | NO | NA |
| Olfr881 | ENSMUSG( | ENSMUSG( | 0.444175 | 0.744557 | NO | NO | NA |
| BC051665 | ENSMUSG( | ENSMUSG( | 0.958942 | 0.526772 | NO | NO | NA |
| Gm48237 | ENSMUSG( | ENSMUSG( | 0.469215 | 0.746839 | NO | NO | NA |
| 4930563N | ENSMUSG( | ENSMUSG( | -0.32708 | 0.722855 | NO | NO | NA |
| Gm49946 | ENSMUSG( | ENSMUSG( | 0.150537 | 0.895322 | NO | NO | NA |
| Gm6533 | ENSMUSG( | ENSMUSG( | -0.25865 | 0.76667 | NO | NO | NA |
| Gm29500 | ENSMUSG( | ENSMUSG( | 0.435585 | 0.711964 | NO | NO | NA |
| Gm37398 | ENSMUSG( | ENSMUSG( | -0.27892 | 0.861045 | NO | NO | NA |
| Gabrg2 | ENSMUSG( | ENSMUSG( | 0.637284 | 0.631515 | NO | NO | NA |
| Gm12031 | ENSMUSG( | ENSMUSG( | 1.508939 | 0.243074 | NO | NO | NA |
| ENSMUSG | ENSMUSG( | ENSMUSG( | -0.18288 | 0.860095 | NO | NO | NA |
| Olfr237 | ENSMUSG( | ENSMUSG( | 0.529382 | 0.686352 | NO | NO | NA |
| 2010109A | ENSMUSG( | ENSMUSG( | -0.36313 | 0.604949 | NO | NO | NA |
| Gm48935 | ENSMUSG( | ENSMUSG( | 0.590275 | 0.565868 | NO | NO | NA |
| Gm8130 | ENSMUSG( | ENSMUSG( | 0.210938 | 0.763528 | NO | NO | NA |
| Gm12568 | ENSMUSG( | ENSMUSG( | 1.051646 | 0.183882 | NO | NO | NA |
| Vmn1r15 | ENSMUSG( | ENSMUSG( | 0.175063 | 0.895628 | NO | NO | NA |
| Gm11224 | ENSMUSG( | ENSMUSG( | 1.215157 | 0.257464 | NO | NO | NA |
| Pcdhga10 | ENSMUSG( | ENSMUSG( | -0.17912 | 0.928262 | NO | NO | NA |
| D030045P | ENSMUSG( | ENSMUSG( | 0.663932 | 0.600035 | NO | NO | NA |
| 4930570E( | ENSMUSG( | ENSMUSG( | 0.960426 | 0.412487 | NO | NO | NA |
| Gm11991 | ENSMUSG( | ENSMUSG( | 0.278414 | 0.663248 | NO | NO | NA |
| Gm38026 | ENSMUSG( | ENSMUSG( | -0.19582 | 0.829442 | NO | NO | NA |
| Cdcp3 | ENSMUSG( | ENSMUSG( | 0.018217 | 0.984434 | NO | NO | NA |
| Gm37671 | ENSMUSG( | ENSMUSG( | 1.770322 | 0.126244 | NO | NO | NA |
| Gm38091 | ENSMUSG( | ENSMUSG( | 0.188291 | 0.806405 | NO | NO | NA |
| Gm3893 | ENSMUSG( | ENSMUSG( | 1.544241 | 0.589529 | NO | NO | NA |
| Gm49188 | ENSMUSG( | ENSMUSG( | 1.015059 | 0.286804 | NO | NO | NA |
| Gm13233 | ENSMUSG( | ENSMUSG( | 1.83319 | 0.639729 | NO | NO | NA |
| Palmd | ENSMUSG( | ENSMUSG( | -0.40825 | 0.760531 | NO | NO | NA |
| Gm18621 | ENSMUSG( | ENSMUSG( | 0.35471 | 0.80668 | NO | NO | NA |
| Nell1 | ENSMUSG( | ENSMUSG( | 0.529808 | 0.647707 | NO | NO | NA |
| ENSMUSG | ENSMUSG( | ENSMUSG( | -0.19825 | 0.738918 | NO | NO | NA |
| ENSMUSG | ENSMUSG( | ENSMUSG( | 1.04026 | 0.493081 | NO | NO | NA |
| 4933421D | ENSMUSG( | ENSMUSG( | -0.20937 | 0.824393 | NO | NO | NA |

| | | | | | | | |
|---|---|---|---|---|---|---|---|
| Gm5207 | ENSMUSG( | ENSMUSG( | -0.36474 | 0.78906 | NO | NO | NA |
| Gm26577 | ENSMUSG( | ENSMUSG( | 0.525763 | 0.383074 | NO | NO | NA |
| Gm8093 | ENSMUSG( | ENSMUSG( | -0.69152 | 0.513114 | NO | NO | NA |
| Naaladl2 | ENSMUSG( | ENSMUSG( | 1.918949 | 0.124185 | NO | NO | NA |
| Gm38123 | ENSMUSG( | ENSMUSG( | 0.721521 | 0.505017 | NO | NO | NA |
| Gm36860 | ENSMUSG( | ENSMUSG( | 1.348255 | 0.268487 | NO | NO | NA |
| ENSMUSG | ENSMUSG( | ENSMUSG( | -0.57733 | 0.424153 | NO | NO | NA |
| Ccnb1-ps | ENSMUSG( | ENSMUSG( | 0.421879 | 0.491646 | NO | NO | NA |
| Gm49629 | ENSMUSG( | ENSMUSG( | 1.3508 | 0.26258 | NO | NO | NA |
| Efcab9 | ENSMUSG( | ENSMUSG( | 0.615238 | 0.273506 | NO | NO | NA |
| Hepacam2 | ENSMUSG( | ENSMUSG( | 0.435569 | 0.709815 | NO | NO | NA |
| 1700030J2 | ENSMUSG( | ENSMUSG( | -0.39679 | 0.505024 | NO | NO | NA |
| Gm8173 | ENSMUSG( | ENSMUSG( | -0.27872 | 0.78842 | NO | NO | NA |
| Calr3 | ENSMUSG( | ENSMUSG( | 0.402065 | 0.462692 | NO | NO | NA |
| Gm17910 | ENSMUSG( | ENSMUSG( | 0.180782 | 0.781929 | NO | NO | NA |
| Slc5a5 | ENSMUSG( | ENSMUSG( | -1.06796 | 0.159401 | NO | NO | NA |
| Gm11783 | ENSMUSG( | ENSMUSG( | -0.07855 | 0.944554 | NO | NO | NA |
| Cdiptos | ENSMUSG( | ENSMUSG( | -0.13664 | 0.851529 | NO | NO | NA |
| ENSMUSG | ENSMUSG( | ENSMUSG( | -0.06029 | 0.949367 | NO | NO | NA |
| Gm52969 | ENSMUSG( | ENSMUSG( | 1.300803 | 0.431457 | NO | NO | NA |
| Gm13623 | ENSMUSG( | ENSMUSG( | 0.284391 | 0.802906 | NO | NO | NA |
| Chst1 | ENSMUSG( | ENSMUSG( | 0.250934 | 0.796934 | NO | NO | NA |
| ENSMUSG( | ENSMUSG( | ENSMUSG( | 0.52869 | 0.638572 | NO | NO | NA |
| Pdlim3 | ENSMUSG( | ENSMUSG( | -0.01477 | 0.989896 | NO | NO | NA |
| Gm43753 | ENSMUSG( | ENSMUSG( | -0.13163 | 0.895322 | NO | NO | NA |
| Gm26673 | ENSMUSG( | ENSMUSG( | -0.49385 | 0.619675 | NO | NO | NA |
| Klhl4 | ENSMUSG( | ENSMUSG( | 0.471962 | 0.500293 | NO | NO | NA |
| Olfr142 | ENSMUSG( | ENSMUSG( | 1.532284 | 0.208343 | NO | NO | NA |
| Gm43896 | ENSMUSG( | ENSMUSG( | 0.551636 | 0.642925 | NO | NO | NA |
| Slc25a43 | ENSMUSG( | ENSMUSG( | 0.424795 | 0.497255 | NO | NO | NA |
| Pgr | ENSMUSG( | ENSMUSG( | 0.555928 | 0.702414 | NO | NO | NA |
| Hydin | ENSMUSG( | ENSMUSG( | 0.514876 | 0.53749 | NO | NO | NA |
| Gm28465 | ENSMUSG( | ENSMUSG( | 0.64203 | 0.665475 | NO | NO | NA |
| Olfr147 | ENSMUSG( | ENSMUSG( | 0.94387 | 0.414083 | NO | NO | NA |
| Gm14964 | ENSMUSG( | ENSMUSG( | 1.540665 | 0.271723 | NO | NO | NA |
| Gm14046 | ENSMUSG( | ENSMUSG( | 0.756302 | 0.165778 | NO | NO | NA |
| Pde10a | ENSMUSG( | ENSMUSG( | 0.414086 | 0.476823 | NO | NO | NA |
| Asb5 | ENSMUSG( | ENSMUSG( | 0.609579 | 0.509801 | NO | NO | NA |
| ENSMUSG( | ENSMUSG( | ENSMUSG( | 0.683416 | 0.511586 | NO | NO | NA |
| Gm7803 | ENSMUSG( | ENSMUSG( | 0.494227 | 0.624649 | NO | NO | NA |
| Gm2762 | ENSMUSG( | ENSMUSG( | 0.273254 | 0.86099 | NO | NO | NA |
| Gm26887 | ENSMUSG( | ENSMUSG( | 1.651288 | 0.027545 | YES | YES | Upregulated |
| Gm26511 | ENSMUSG( | ENSMUSG( | 0.547769 | 0.454653 | NO | NO | NA |
| Sftpd | ENSMUSG( | ENSMUSG( | 0.099409 | 0.930568 | NO | NO | NA |
| Olfr1028 | ENSMUSG( | ENSMUSG( | 2.470087 | 0.008117 | YES | YES | Upregulated |
| 4930500H | ENSMUSG( | ENSMUSG( | -0.04525 | 0.975611 | NO | NO | NA |
| 1700092M | ENSMUSG( | ENSMUSG( | 0.411722 | 0.647827 | NO | NO | NA |

| | | | | | | | |
|---|---|---|---|---|---|---|---|
| Gm37672 | ENSMUSG( | ENSMUSG( | -0.45757 | 0.515314 | NO | NO | NA |
| Gm43119 | ENSMUSG( | ENSMUSG( | 1.08437 | 0.41201 | NO | NO | NA |
| Gm45183 | ENSMUSG( | ENSMUSG( | 0.047898 | 0.970343 | NO | NO | NA |
| Gm10135 | ENSMUSG( | ENSMUSG( | 0.039654 | 0.987646 | NO | NO | NA |
| 4930412M | ENSMUSG( | ENSMUSG( | 0.125265 | 0.815418 | NO | NO | NA |
| Lrrc24 | ENSMUSG( | ENSMUSG( | -0.85483 | 0.334627 | NO | NO | NA |
| Gm4875 | ENSMUSG( | ENSMUSG( | -0.86637 | 0.19845 | NO | NO | NA |
| Gm47904 | ENSMUSG( | ENSMUSG( | -0.35673 | 0.774247 | NO | NO | NA |
| Gm43015 | ENSMUSG( | ENSMUSG( | 0.132726 | 0.917228 | NO | NO | NA |
| Usp51 | ENSMUSG( | ENSMUSG( | 0.259945 | 0.588319 | NO | NO | NA |
| ENSMUSG( | ENSMUSG( | ENSMUSG( | 0.934001 | 0.465954 | NO | NO | NA |
| Fgd5 | ENSMUSG( | ENSMUSG( | 0.965899 | 0.388618 | NO | NO | NA |
| Gm37060 | ENSMUSG( | ENSMUSG( | 0.718651 | 0.544325 | NO | NO | NA |
| Amot | ENSMUSG( | ENSMUSG( | 1.121722 | 0.177644 | NO | NO | NA |
| Rps2-ps5 | ENSMUSG( | ENSMUSG( | 0.019288 | 0.9833 | NO | NO | NA |
| Gm16156 | ENSMUSG( | ENSMUSG( | 0.296036 | 0.712916 | NO | NO | NA |
| Gm2108 | ENSMUSG( | ENSMUSG( | 1.375667 | 0.178034 | NO | NO | NA |
| Ccdc158 | ENSMUSG( | ENSMUSG( | 0.806611 | 0.473607 | NO | NO | NA |
| Pyroxd2 | ENSMUSG( | ENSMUSG( | 0.666221 | 0.41326 | NO | NO | NA |
| Olfr1231 | ENSMUSG( | ENSMUSG( | 1.792543 | 0.162161 | NO | NO | NA |
| ENSMUSG | ENSMUSG( | ENSMUSG( | 0.308984 | 0.814938 | NO | NO | NA |
| Olfr392 | ENSMUSG( | ENSMUSG( | 0.788383 | 0.539424 | NO | NO | NA |
| ENSMUSG | ENSMUSG( | ENSMUSG( | -0.34929 | 0.542573 | NO | NO | NA |
| Rpl19-ps6 | ENSMUSG( | ENSMUSG( | 0.713275 | 0.268548 | NO | NO | NA |
| Gm16310 | ENSMUSG( | ENSMUSG( | 0.23001 | 0.657997 | NO | NO | NA |
| Gm48602 | ENSMUSG( | ENSMUSG( | 0.612938 | 0.596699 | NO | NO | NA |
| Zfp981 | ENSMUSG( | ENSMUSG( | 4.905175 | 0.0002 | YES | YES | Upregulated |
| Gm7390 | ENSMUSG( | ENSMUSG( | 0.405225 | 0.485768 | NO | NO | NA |
| Ccdc83 | ENSMUSG( | ENSMUSG( | -0.38536 | 0.76761 | NO | NO | NA |
| Gm43180 | ENSMUSG( | ENSMUSG( | 0.379043 | 0.682983 | NO | NO | NA |
| Synm | ENSMUSG( | ENSMUSG( | 0.437912 | 0.66558 | NO | NO | NA |
| Epha2 | ENSMUSG( | ENSMUSG( | 0.089204 | 0.923834 | NO | NO | NA |
| Mybpc1 | ENSMUSG( | ENSMUSG( | 1.20498 | 0.047764 | YES | YES | Upregulated |
| Trim52 | ENSMUSG( | ENSMUSG( | 0.135876 | 0.908038 | NO | NO | NA |
| Gm29358 | ENSMUSG( | ENSMUSG( | 0.680593 | 0.434626 | NO | NO | NA |
| ENSMUSG | ENSMUSG( | ENSMUSG( | 1.035222 | 0.143737 | NO | NO | NA |
| Ms4a20 | ENSMUSG( | ENSMUSG( | 1.387316 | 0.29044 | NO | NO | NA |
| Gm16861 | ENSMUSG( | ENSMUSG( | 0.003449 | 0.994917 | NO | NO | NA |
| Gm15899 | ENSMUSG( | ENSMUSG( | -0.14818 | 0.897251 | NO | NO | NA |
| C2 | ENSMUSG( | ENSMUSG( | 0.258959 | 0.687358 | NO | NO | NA |
| Scamp5 | ENSMUSG( | ENSMUSG( | -0.62669 | 0.375455 | NO | NO | NA |
| Crry-ps | ENSMUSG( | ENSMUSG( | 0.366256 | 0.633647 | NO | NO | NA |
| Pak1 | ENSMUSG( | ENSMUSG( | -0.44188 | 0.576362 | NO | NO | NA |
| Gm18697 | ENSMUSG( | ENSMUSG( | 0.634226 | 0.539055 | NO | NO | NA |
| D130079A | ENSMUSG( | ENSMUSG( | -0.21485 | 0.815932 | NO | NO | NA |
| St18 | ENSMUSG( | ENSMUSG( | 0.541707 | 0.540236 | NO | NO | NA |
| Foxa1 | ENSMUSG( | ENSMUSG( | -0.07205 | 0.95455 | NO | NO | NA |

| | | | | | | | |
|---|---|---|---|---|---|---|---|
| Gm4526 | ENSMUSG( | ENSMUSG( | -0.05343 | 0.977981 | NO | NO | NA |
| Gm5611 | ENSMUSG( | ENSMUSG( | 0.76961 | 0.254155 | NO | NO | NA |
| Sox1 | ENSMUSG( | ENSMUSG( | 0.548746 | 0.674136 | NO | NO | NA |
| Gm9575 | ENSMUSG( | ENSMUSG( | 0.369039 | 0.619675 | NO | NO | NA |
| Gm49756 | ENSMUSG( | ENSMUSG( | 0.506887 | 0.658915 | NO | NO | NA |
| Gm11893 | ENSMUSG( | ENSMUSG( | 1.123731 | 0.443092 | NO | NO | NA |
| Dach2 | ENSMUSG( | ENSMUSG( | 1.199264 | 0.248944 | NO | NO | NA |
| Gm39317 | ENSMUSG( | ENSMUSG( | 1.194964 | 0.29155 | NO | NO | NA |
| ENSMUSG | ENSMUSG( | ENSMUSG( | 1.422653 | 0.042226 | YES | YES | Upregulated |
| Gm45389 | ENSMUSG( | ENSMUSG( | 0.453516 | 0.743114 | NO | NO | NA |
| Gm50019 | ENSMUSG( | ENSMUSG( | -0.33168 | 0.509441 | NO | NO | NA |
| Rem1 | ENSMUSG( | ENSMUSG( | 0.137113 | 0.898999 | NO | NO | NA |
| Gm1965 | ENSMUSG( | ENSMUSG( | -0.03108 | 0.98 | NO | NO | NA |
| Tm4sf4 | ENSMUSG( | ENSMUSG( | 0.202661 | 0.891542 | NO | NO | NA |
| ENSMUSG | ENSMUSG( | ENSMUSG( | 0.106082 | 0.916811 | NO | NO | NA |
| Gm44666 | ENSMUSG( | ENSMUSG( | 0.473742 | 0.784854 | NO | NO | NA |
| Olfr683 | ENSMUSG( | ENSMUSG( | 1.073387 | 0.451796 | NO | NO | NA |
| Cox6a2 | ENSMUSG( | ENSMUSG( | -2.19107 | 0.001438 | YES | YES | Downregulated |
| Gm19007 | ENSMUSG( | ENSMUSG( | 0.375097 | 0.741472 | NO | NO | NA |
| Nudc-ps1 | ENSMUSG( | ENSMUSG( | 0.508204 | 0.388366 | NO | NO | NA |
| Scube1 | ENSMUSG( | ENSMUSG( | 0.41554 | 0.807036 | NO | NO | NA |
| ENSMUSG | ENSMUSG( | ENSMUSG( | -1.09668 | 0.125415 | NO | NO | NA |
| Kirrel3 | ENSMUSG( | ENSMUSG( | -0.15606 | 0.860142 | NO | NO | NA |
| 4933402N | ENSMUSG( | ENSMUSG( | 1.817908 | 0.349893 | NO | NO | NA |
| Gm15773 | ENSMUSG( | ENSMUSG( | 0.433192 | 0.479941 | NO | NO | NA |
| Zfp641 | ENSMUSG( | ENSMUSG( | 1.198821 | 0.377352 | NO | NO | NA |
| Gm37353 | ENSMUSG( | ENSMUSG( | 0.78305 | 0.355386 | NO | NO | NA |
| Gm15454 | ENSMUSG( | ENSMUSG( | 0.523868 | 0.680822 | NO | NO | NA |
| Gm43190 | ENSMUSG( | ENSMUSG( | 0.184371 | 0.880844 | NO | NO | NA |
| Gm7204 | ENSMUSG( | ENSMUSG( | 0.373473 | 0.526511 | NO | NO | NA |
| Gm36161 | ENSMUSG( | ENSMUSG( | -0.42182 | 0.635606 | NO | NO | NA |
| Kcnn2 | ENSMUSG( | ENSMUSG( | -0.01897 | 0.98694 | NO | NO | NA |
| Gm14791 | ENSMUSG( | ENSMUSG( | 1.007678 | 0.637329 | NO | NO | NA |
| Gm15161 | ENSMUSG( | ENSMUSG( | 1.122724 | 0.386118 | NO | NO | NA |
| Gm15023 | ENSMUSG( | ENSMUSG( | 1.66941 | 0.491395 | NO | NO | NA |
| Gm44877 | ENSMUSG( | ENSMUSG( | 0.561299 | 0.462135 | NO | NO | NA |
| Gm45639 | ENSMUSG( | ENSMUSG( | 0.541939 | 0.668205 | NO | NO | NA |
| Phyhipl | ENSMUSG( | ENSMUSG( | 0.724218 | 0.549883 | NO | NO | NA |
| Gm15873 | ENSMUSG( | ENSMUSG( | 1.133845 | 0.107958 | NO | NO | NA |
| Vmn2r68 | ENSMUSG( | ENSMUSG( | 0.797346 | 0.581594 | NO | NO | NA |
| Gm8581 | ENSMUSG( | ENSMUSG( | 0.328644 | 0.853293 | NO | NO | NA |
| Gm36315 | ENSMUSG( | ENSMUSG( | 1.324361 | 0.13697 | NO | NO | NA |
| ENSMUSG | ENSMUSG( | ENSMUSG( | 0.056814 | 0.946459 | NO | NO | NA |
| Gm12977 | ENSMUSG( | ENSMUSG( | 0.634875 | 0.435507 | NO | NO | NA |
| Bend7 | ENSMUSG( | ENSMUSG( | 1.085577 | 0.113267 | NO | NO | NA |
| Gm37383 | ENSMUSG( | ENSMUSG( | 0.499458 | 0.580312 | NO | NO | NA |
| Gm12506 | ENSMUSG( | ENSMUSG( | 0.606508 | 0.708667 | NO | NO | NA |

| | | | | | | | |
|---|---|---|---|---|---|---|---|
| Rpl21-ps1! | ENSMUSG( | ENSMUSG( | 0.870056 | 0.503887 | NO | NO | NA |
| Hnf1b | ENSMUSG( | ENSMUSG( | 0.81655 | 0.39983 | NO | NO | NA |
| Gm13420 | ENSMUSG( | ENSMUSG( | 0.430616 | 0.560577 | NO | NO | NA |
| Olfr477 | ENSMUSG( | ENSMUSG( | 0.419745 | 0.731835 | NO | NO | NA |
| Ckmt1 | ENSMUSG( | ENSMUSG( | 0.553192 | 0.53766 | NO | NO | NA |
| Gm37397 | ENSMUSG( | ENSMUSG( | -0.57094 | 0.528583 | NO | NO | NA |
| 4932411K: | ENSMUSG( | ENSMUSG( | 0.005593 | 0.995646 | NO | NO | NA |
| Gm12383 | ENSMUSG( | ENSMUSG( | 1.511502 | 0.44354 | NO | NO | NA |
| Gm48776 | ENSMUSG( | ENSMUSG( | 2.846489 | 0.128446 | NO | NO | NA |
| Gm11725 | ENSMUSG( | ENSMUSG( | -0.46519 | 0.625719 | NO | NO | NA |
| Gm42763 | ENSMUSG( | ENSMUSG( | 0.238526 | 0.668822 | NO | NO | NA |
| Gm6418 | ENSMUSG( | ENSMUSG( | 0.941312 | 0.436468 | NO | NO | NA |
| Gm14493 | ENSMUSG( | ENSMUSG( | 0.176047 | 0.855008 | NO | NO | NA |
| Cd300lg | ENSMUSG( | ENSMUSG( | 0.191943 | 0.858532 | NO | NO | NA |
| 2210411M | ENSMUSG( | ENSMUSG( | 0.891172 | 0.480717 | NO | NO | NA |
| Rab11b-ps | ENSMUSG( | ENSMUSG( | 0.333082 | 0.521197 | NO | NO | NA |
| Gm33093 | ENSMUSG( | ENSMUSG( | 0.891467 | 0.459146 | NO | NO | NA |
| Gm15958 | ENSMUSG( | ENSMUSG( | 0.611754 | 0.58518 | NO | NO | NA |
| Iqank1 | ENSMUSG( | ENSMUSG( | 0.500405 | 0.361514 | NO | NO | NA |
| AY512915 | ENSMUSG( | ENSMUSG( | 0.060954 | 0.954092 | NO | NO | NA |
| Atp8b3 | ENSMUSG( | ENSMUSG( | 0.526905 | 0.507331 | NO | NO | NA |
| Gm45206 | ENSMUSG( | ENSMUSG( | 0.566925 | 0.642754 | NO | NO | NA |
| Slit3 | ENSMUSG( | ENSMUSG( | 3.178967 | 0.007594 | YES | YES | Upregulated |
| Vmn2r6 | ENSMUSG( | ENSMUSG( | 0.094257 | 0.947496 | NO | NO | NA |
| Syde2 | ENSMUSG( | ENSMUSG( | 0.998916 | 0.137575 | NO | NO | NA |
| 4930598N | ENSMUSG( | ENSMUSG( | 1.226957 | 0.319818 | NO | NO | NA |
| Gm47628 | ENSMUSG( | ENSMUSG( | 0.277002 | 0.822358 | NO | NO | NA |
| Gm45204 | ENSMUSG( | ENSMUSG( | 0.173028 | 0.880076 | NO | NO | NA |
| Speer3 | ENSMUSG( | ENSMUSG( | 0.475801 | 0.672658 | NO | NO | NA |
| Gm37534 | ENSMUSG( | ENSMUSG( | 0.967879 | 0.276306 | NO | NO | NA |
| Gm49972 | ENSMUSG( | ENSMUSG( | -0.3793 | 0.629989 | NO | NO | NA |
| Gm9905 | ENSMUSG( | ENSMUSG( | 0.122586 | 0.874422 | NO | NO | NA |
| Gm15535 | ENSMUSG( | ENSMUSG( | 0.503096 | 0.624452 | NO | NO | NA |
| Gm26592 | ENSMUSG( | ENSMUSG( | 0.281924 | 0.772033 | NO | NO | NA |
| Gm38260 | ENSMUSG( | ENSMUSG( | 0.528345 | 0.648053 | NO | NO | NA |
| Nutm1 | ENSMUSG( | ENSMUSG( | 0.193787 | 0.759335 | NO | NO | NA |
| Olfr822 | ENSMUSG( | ENSMUSG( | 0.485652 | 0.652127 | NO | NO | NA |
| Gm43488 | ENSMUSG( | ENSMUSG( | 0.632605 | 0.508669 | NO | NO | NA |
| Gm49041 | ENSMUSG( | ENSMUSG( | 0.130607 | 0.861547 | NO | NO | NA |
| 2310034G | ENSMUSG( | ENSMUSG( | 0.529295 | 0.478132 | NO | NO | NA |
| Gm44243 | ENSMUSG( | ENSMUSG( | 0.394353 | 0.518553 | NO | NO | NA |
| 1700054A( | ENSMUSG( | ENSMUSG( | 0.384769 | 0.68649 | NO | NO | NA |
| Gm14988 | ENSMUSG( | ENSMUSG( | -0.28223 | 0.859925 | NO | NO | NA |
| Olfr1135 | ENSMUSG( | ENSMUSG( | 0.987309 | 0.557458 | NO | NO | NA |
| Gm29016 | ENSMUSG( | ENSMUSG( | 0.694681 | 0.655215 | NO | NO | NA |
| Gm47132 | ENSMUSG( | ENSMUSG( | -0.8872 | 0.291319 | NO | NO | NA |
| Synpo2l | ENSMUSG( | ENSMUSG( | 0.940676 | 0.21335 | NO | NO | NA |

| | | | | | | | |
|---|---|---|---|---|---|---|---|
| Camsap3 | ENSMUSG( | ENSMUSG( | -0.31219 | 0.645176 | NO | NO | NA |
| Zscan4a | ENSMUSG( | ENSMUSG( | 0.476498 | 0.760533 | NO | NO | NA |
| Gm43187 | ENSMUSG( | ENSMUSG( | 0.459065 | 0.713194 | NO | NO | NA |
| ENSMUSG( | ENSMUSG( | ENSMUSG( | 0.528025 | 0.586707 | NO | NO | NA |
| Gm26686 | ENSMUSG( | ENSMUSG( | 0.174819 | 0.830848 | NO | NO | NA |
| Esrp1 | ENSMUSG( | ENSMUSG( | -0.04966 | 0.964167 | NO | NO | NA |
| Gm5080 | ENSMUSG( | ENSMUSG( | 0.479931 | 0.679095 | NO | NO | NA |
| ENSMUSG | ENSMUSG( | ENSMUSG( | 1.467759 | 0.229656 | NO | NO | NA |
| Olfr418 | ENSMUSG( | ENSMUSG( | 1.049049 | 0.461648 | NO | NO | NA |
| Gm37364 | ENSMUSG( | ENSMUSG( | -0.09068 | 0.949307 | NO | NO | NA |
| Gm8623 | ENSMUSG( | ENSMUSG( | 0.32821 | 0.631598 | NO | NO | NA |
| Gm7290 | ENSMUSG( | ENSMUSG( | 0.143205 | 0.909888 | NO | NO | NA |
| Spata31 | ENSMUSG( | ENSMUSG( | 0.067495 | 0.961918 | NO | NO | NA |
| Gm11557 | ENSMUSG( | ENSMUSG( | -0.00156 | 0.998967 | NO | NO | NA |
| Trpv6 | ENSMUSG( | ENSMUSG( | -0.11217 | 0.896584 | NO | NO | NA |
| ENSMUSG | ENSMUSG( | ENSMUSG( | 0.368532 | 0.714767 | NO | NO | NA |
| 4930413G | ENSMUSG( | ENSMUSG( | 0.053057 | 0.922644 | NO | NO | NA |
| Gm3081 | ENSMUSG( | ENSMUSG( | -0.53003 | 0.466606 | NO | NO | NA |
| Slc3a1 | ENSMUSG( | ENSMUSG( | -0.07136 | 0.946562 | NO | NO | NA |
| Gm9903 | ENSMUSG( | ENSMUSG( | -0.65568 | 0.290434 | NO | NO | NA |
| C130093G | ENSMUSG( | ENSMUSG( | 0.551067 | 0.626342 | NO | NO | NA |
| Cers1 | ENSMUSG( | ENSMUSG( | -0.73368 | 0.372481 | NO | NO | NA |
| B3gnt4 | ENSMUSG( | ENSMUSG( | -0.87604 | 0.356412 | NO | NO | NA |
| Gm12532 | ENSMUSG( | ENSMUSG( | 1.728867 | 0.197539 | NO | NO | NA |
| Gm48909 | ENSMUSG( | ENSMUSG( | 0.910146 | 0.240544 | NO | NO | NA |
| Spertl | ENSMUSG( | ENSMUSG( | 2.477566 | 0.137143 | NO | NO | NA |
| Gm7776 | ENSMUSG( | ENSMUSG( | 0.598574 | 0.480291 | NO | NO | NA |
| Gm43191 | ENSMUSG( | ENSMUSG( | 3.932073 | 0.146084 | NO | NO | NA |
| Gm7595 | ENSMUSG( | ENSMUSG( | -0.35596 | 0.85174 | NO | NO | NA |
| ENSMUSG | ENSMUSG( | ENSMUSG( | 1.014461 | 0.425871 | NO | NO | NA |
| Gm6394 | ENSMUSG( | ENSMUSG( | 0.277429 | 0.760732 | NO | NO | NA |
| 4930519E( | ENSMUSG( | ENSMUSG( | 0.743897 | 0.485363 | NO | NO | NA |
| AV320801 | ENSMUSG( | ENSMUSG( | 0.619765 | 0.805179 | NO | NO | NA |
| Gm20756 | ENSMUSG( | ENSMUSG( | 1.625867 | 0.268921 | NO | NO | NA |
| Gm7638 | ENSMUSG( | ENSMUSG( | -0.07521 | 0.923834 | NO | NO | NA |
| Gm2694 | ENSMUSG( | ENSMUSG( | -0.18092 | 0.87252 | NO | NO | NA |
| Gm31881 | ENSMUSG( | ENSMUSG( | -0.19261 | 0.888832 | NO | NO | NA |
| Gm5186 | ENSMUSG( | ENSMUSG( | 0.970148 | 0.513234 | NO | NO | NA |
| Gm8100 | ENSMUSG( | ENSMUSG( | -0.79168 | 0.452595 | NO | NO | NA |
| Gm36955 | ENSMUSG( | ENSMUSG( | -0.03342 | 0.973265 | NO | NO | NA |
| Fzd7 | ENSMUSG( | ENSMUSG( | 0.238698 | 0.751327 | NO | NO | NA |
| Gm37788 | ENSMUSG( | ENSMUSG( | 0.014268 | 0.991949 | NO | NO | NA |
| Igkv8-28 | ENSMUSG( | ENSMUSG( | -4.73463 | 0.072352 | NO | NO | NA |
| Gm19582 | ENSMUSG( | ENSMUSG( | -0.37706 | 0.544276 | NO | NO | NA |
| ENSMUSG( | ENSMUSG( | ENSMUSG( | -1.01352 | 0.589182 | NO | NO | NA |
| Gm37066 | ENSMUSG( | ENSMUSG( | 0.739556 | 0.508534 | NO | NO | NA |
| Zfp92 | ENSMUSG( | ENSMUSG( | 0.423636 | 0.472355 | NO | NO | NA |

| | | | | | | | |
|---|---|---|---|---|---|---|---|
| Gm15651 | ENSMUSG( | ENSMUSG( | 2.096053 | 0.103506 | NO | NO | NA |
| Gm26981 | ENSMUSG( | ENSMUSG( | 0.216935 | 0.721604 | NO | NO | NA |
| Gm30593 | ENSMUSG( | ENSMUSG( | -0.04168 | 0.978564 | NO | NO | NA |
| 4833411C( | ENSMUSG( | ENSMUSG( | 0.816291 | 0.492717 | NO | NO | NA |
| Gm48336 | ENSMUSG( | ENSMUSG( | 0.652441 | 0.409549 | NO | NO | NA |
| Gm4986 | ENSMUSG( | ENSMUSG( | 1.004993 | 0.561653 | NO | NO | NA |
| Olfr756-ps | ENSMUSG( | ENSMUSG( | -0.64988 | 0.457263 | NO | NO | NA |
| 9830166K( | ENSMUSG( | ENSMUSG( | 0.976154 | 0.446476 | NO | NO | NA |
| Gm6023 | ENSMUSG( | ENSMUSG( | -0.08227 | 0.908038 | NO | NO | NA |
| ENSMUSG | ENSMUSG( | ENSMUSG( | 0.465406 | 0.707965 | NO | NO | NA |
| 4930478P: | ENSMUSG( | ENSMUSG( | 0.968756 | 0.464432 | NO | NO | NA |
| Prickle4 | ENSMUSG( | ENSMUSG( | -0.05955 | 0.976913 | NO | NO | NA |
| Slc10a1 | ENSMUSG( | ENSMUSG( | 0.434024 | 0.613276 | NO | NO | NA |
| Gm16071 | ENSMUSG( | ENSMUSG( | 0.662794 | 0.301162 | NO | NO | NA |
| Gm17906 | ENSMUSG( | ENSMUSG( | 0.966734 | 0.413484 | NO | NO | NA |
| Tll1 | ENSMUSG( | ENSMUSG( | 0.338216 | 0.817522 | NO | NO | NA |
| Gm42639 | ENSMUSG( | ENSMUSG( | 0.459412 | 0.68133 | NO | NO | NA |
| Olfr6 | ENSMUSG( | ENSMUSG( | -0.03332 | 0.975877 | NO | NO | NA |
| Gys2 | ENSMUSG( | ENSMUSG( | 1.555634 | 0.25122 | NO | NO | NA |
| Pkib | ENSMUSG( | ENSMUSG( | 0.258255 | 0.787243 | NO | NO | NA |
| Pcdha5 | ENSMUSG( | ENSMUSG( | 2.055464 | 0.364595 | NO | NO | NA |
| Gm47723 | ENSMUSG( | ENSMUSG( | 1.568107 | 0.215303 | NO | NO | NA |
| Gm37850 | ENSMUSG( | ENSMUSG( | -0.30224 | 0.664907 | NO | NO | NA |
| Ugt1a5 | ENSMUSG( | ENSMUSG( | 0.240572 | 0.928272 | NO | NO | NA |
| Gm43931 | ENSMUSG( | ENSMUSG( | 0.405774 | 0.747644 | NO | NO | NA |
| Gm37502 | ENSMUSG( | ENSMUSG( | 1.412151 | 0.222025 | NO | NO | NA |
| Gm34758 | ENSMUSG( | ENSMUSG( | -0.62492 | 0.459146 | NO | NO | NA |
| 4930417O | ENSMUSG( | ENSMUSG( | 1.146181 | 0.491199 | NO | NO | NA |
| Gm7104 | ENSMUSG( | ENSMUSG( | 0.692037 | 0.626403 | NO | NO | NA |
| Ahrr | ENSMUSG( | ENSMUSG( | 0.586106 | 0.458202 | NO | NO | NA |
| Ppp1r3a | ENSMUSG( | ENSMUSG( | 0.71813 | 0.560369 | NO | NO | NA |
| Gm45873 | ENSMUSG( | ENSMUSG( | 0.017037 | 0.984258 | NO | NO | NA |
| Myo18b | ENSMUSG( | ENSMUSG( | 0.832728 | 0.330466 | NO | NO | NA |
| A930006I( | ENSMUSG( | ENSMUSG( | 1.011818 | 0.343845 | NO | NO | NA |
| Cdk5r2 | ENSMUSG( | ENSMUSG( | 0.928048 | 0.398122 | NO | NO | NA |
| Gm44735 | ENSMUSG( | ENSMUSG( | -0.94491 | 0.069591 | NO | NO | NA |
| ENSMUSG | ENSMUSG( | ENSMUSG( | 0.396473 | 0.749566 | NO | NO | NA |
| Gm4557 | ENSMUSG( | ENSMUSG( | -0.1678 | 0.942688 | NO | NO | NA |
| Sis | ENSMUSG( | ENSMUSG( | 0.449907 | 0.723449 | NO | NO | NA |
| Gm47430 | ENSMUSG( | ENSMUSG( | 0.341129 | 0.767165 | NO | NO | NA |
| Fbxl16 | ENSMUSG( | ENSMUSG( | 0.336392 | 0.513834 | NO | NO | NA |
| Gm24924 | ENSMUSG( | ENSMUSG( | 1.01757 | 0.639778 | NO | NO | NA |
| Gm37158 | ENSMUSG( | ENSMUSG( | 0.277476 | 0.757131 | NO | NO | NA |
| E330017L1 | ENSMUSG( | ENSMUSG( | 1.557482 | 0.164026 | NO | NO | NA |
| Gm50375 | ENSMUSG( | ENSMUSG( | 0.310933 | 0.758216 | NO | NO | NA |
| ENSMUSG | ENSMUSG( | ENSMUSG( | 0.129126 | 0.873805 | NO | NO | NA |
| 9330175M | ENSMUSG( | ENSMUSG( | 0.89172 | 0.508669 | NO | NO | NA |

| | | | | | | | |
|---|---|---|---|---|---|---|---|
| 2210412B: | ENSMUSG( | ENSMUSG( | 0.246472 | 0.806872 | NO | NO | NA |
| Bnc1 | ENSMUSG( | ENSMUSG( | 0.882517 | 0.452004 | NO | NO | NA |
| Olfr588-ps | ENSMUSG( | ENSMUSG( | 0.965995 | 0.373347 | NO | NO | NA |
| Gm27206 | ENSMUSG( | ENSMUSG( | -0.57461 | 0.429222 | NO | NO | NA |
| Gm37612 | ENSMUSG( | ENSMUSG( | -1.39742 | 0.023362 | YES | YES | Downregulated |
| Bpifb5 | ENSMUSG( | ENSMUSG( | 0.437479 | 0.760265 | NO | NO | NA |
| ENSMUSG( | ENSMUSG( | ENSMUSG( | 0.725642 | 0.607428 | NO | NO | NA |
| Dpep1 | ENSMUSG( | ENSMUSG( | 1.194322 | 0.349342 | NO | NO | NA |
| Cpn2 | ENSMUSG( | ENSMUSG( | 0.705433 | 0.593861 | NO | NO | NA |
| ENSMUSG( | ENSMUSG( | ENSMUSG( | 0.879265 | 0.476531 | NO | NO | NA |
| Olfr679 | ENSMUSG( | ENSMUSG( | 0.282356 | 0.807516 | NO | NO | NA |
| Gm26575 | ENSMUSG( | ENSMUSG( | 0.881745 | 0.436274 | NO | NO | NA |
| Spata25 | ENSMUSG( | ENSMUSG( | 0.761112 | 0.266753 | NO | NO | NA |
| Gm38451 | ENSMUSG( | ENSMUSG( | 1.359891 | 0.211098 | NO | NO | NA |
| ENSMUSG | ENSMUSG( | ENSMUSG( | 0.742041 | 0.08912 | NO | NO | NA |
| 4930512B( | ENSMUSG( | ENSMUSG( | 0.216157 | 0.847974 | NO | NO | NA |
| Fgf1 | ENSMUSG( | ENSMUSG( | 1.175656 | 0.269521 | NO | NO | NA |
| Gm17227 | ENSMUSG( | ENSMUSG( | 0.343714 | 0.640929 | NO | NO | NA |
| Pvalb | ENSMUSG( | ENSMUSG( | 1.856342 | 0.003977 | YES | YES | Upregulated |
| Mroh8 | ENSMUSG( | ENSMUSG( | 0.986114 | 0.058258 | NO | NO | NA |
| Ip6k3 | ENSMUSG( | ENSMUSG( | 1.492151 | 0.447868 | NO | NO | NA |
| Gm4949 | ENSMUSG( | ENSMUSG( | 0.33671 | 0.767976 | NO | NO | NA |
| Slc23a4 | ENSMUSG( | ENSMUSG( | 0.843562 | 0.338824 | NO | NO | NA |
| Gm16239 | ENSMUSG( | ENSMUSG( | -0.77526 | 0.143121 | NO | NO | NA |
| Gm14537 | ENSMUSG( | ENSMUSG( | 0.903534 | 0.458492 | NO | NO | NA |
| D6Ertd527 | ENSMUSG( | ENSMUSG( | -0.24913 | 0.688511 | NO | NO | NA |
| Gm16304 | ENSMUSG( | ENSMUSG( | 1.624111 | 0.215058 | NO | NO | NA |
| Gm41787 | ENSMUSG( | ENSMUSG( | 0.796233 | 0.5656 | NO | NO | NA |
| Prkar1b | ENSMUSG( | ENSMUSG( | -0.03312 | 0.970413 | NO | NO | NA |
| Hmgb1-ps: | ENSMUSG( | ENSMUSG( | 0.759662 | 0.317179 | NO | NO | NA |
| Gm42908 | ENSMUSG( | ENSMUSG( | -0.43443 | 0.582357 | NO | NO | NA |
| Gm12666 | ENSMUSG( | ENSMUSG( | 0.850792 | 0.475134 | NO | NO | NA |
| Gm26688 | ENSMUSG( | ENSMUSG( | 0.102761 | 0.899457 | NO | NO | NA |
| Tmem217 | ENSMUSG( | ENSMUSG( | -0.05872 | 0.932598 | NO | NO | NA |
| Gm39526 | ENSMUSG( | ENSMUSG( | -0.46504 | 0.545662 | NO | NO | NA |
| Gm12264 | ENSMUSG( | ENSMUSG( | -0.61829 | 0.518553 | NO | NO | NA |
| Gm5931 | ENSMUSG( | ENSMUSG( | 0.251379 | 0.653854 | NO | NO | NA |
| 1700010B: | ENSMUSG( | ENSMUSG( | -0.99308 | 0.457961 | NO | NO | NA |
| D630008O | ENSMUSG( | ENSMUSG( | 0.431471 | 0.444377 | NO | NO | NA |
| Olfr1314 | ENSMUSG( | ENSMUSG( | 1.471903 | 0.365156 | NO | NO | NA |
| A630023P | ENSMUSG( | ENSMUSG( | 0.04338 | 0.96662 | NO | NO | NA |
| Gm8606 | ENSMUSG( | ENSMUSG( | 0.714694 | 0.591301 | NO | NO | NA |
| Gm43188 | ENSMUSG( | ENSMUSG( | 1.206892 | 0.415327 | NO | NO | NA |
| Gm42801 | ENSMUSG( | ENSMUSG( | 0.710795 | 0.575147 | NO | NO | NA |
| ENSMUSG( | ENSMUSG( | ENSMUSG( | -0.18874 | 0.874679 | NO | NO | NA |
| Slc6a4 | ENSMUSG( | ENSMUSG( | -0.66147 | 0.500435 | NO | NO | NA |
| Gm37818 | ENSMUSG( | ENSMUSG( | 0.698635 | 0.553957 | NO | NO | NA |

| | | | | | | | |
|---|---|---|---|---|---|---|---|
| Gm19337 | ENSMUSG( | ENSMUSG( | -0.33193 | 0.480277 | NO | NO | NA |
| Gm47732 | ENSMUSG( | ENSMUSG( | 4.351566 | 0.072544 | NO | NO | NA |
| Atp6v0a4 | ENSMUSG( | ENSMUSG( | 0.849907 | 0.400018 | NO | NO | NA |
| Tmem92 | ENSMUSG( | ENSMUSG( | 0.983558 | 0.437342 | NO | NO | NA |
| Gm14938 | ENSMUSG( | ENSMUSG( | 1.557797 | 0.429629 | NO | NO | NA |
| Olfr267 | ENSMUSG( | ENSMUSG( | 0.328115 | 0.758707 | NO | NO | NA |
| Gm37541 | ENSMUSG( | ENSMUSG( | -0.19953 | 0.795942 | NO | NO | NA |
| Gm44116 | ENSMUSG( | ENSMUSG( | 0.380205 | 0.63616 | NO | NO | NA |
| Cd248 | ENSMUSG( | ENSMUSG( | 1.135841 | 0.187531 | NO | NO | NA |
| ENSMUSG | ENSMUSG( | ENSMUSG( | 0.137797 | 0.873733 | NO | NO | NA |
| 4933427J( | ENSMUSG( | ENSMUSG( | 0.074822 | 0.956604 | NO | NO | NA |
| Gm13167 | ENSMUSG( | ENSMUSG( | 1.886592 | 0.384056 | NO | NO | NA |
| Gm18193 | ENSMUSG( | ENSMUSG( | 0.837996 | 0.561237 | NO | NO | NA |
| Gm10354 | ENSMUSG( | ENSMUSG( | 1.11381 | 0.387383 | NO | NO | NA |
| Gm6981 | ENSMUSG( | ENSMUSG( | 0.016736 | 0.978039 | NO | NO | NA |
| Gm5533 | ENSMUSG( | ENSMUSG( | 0.435755 | 0.572489 | NO | NO | NA |
| ENSMUSG | ENSMUSG( | ENSMUSG( | 0.785425 | 0.158327 | NO | NO | NA |
| Gm19552 | ENSMUSG( | ENSMUSG( | 0.131194 | 0.840237 | NO | NO | NA |
| Gm43365 | ENSMUSG( | ENSMUSG( | 0.997943 | 0.334634 | NO | NO | NA |
| Siglec15 | ENSMUSG( | ENSMUSG( | 0.16378 | 0.919733 | NO | NO | NA |
| Gm26779 | ENSMUSG( | ENSMUSG( | 0.015329 | 0.980563 | NO | NO | NA |
| ENSMUSG( | ENSMUSG( | ENSMUSG( | 9.68E-05 | 0.999926 | NO | NO | NA |
| Otx2os1 | ENSMUSG( | ENSMUSG( | 1.189909 | 0.413484 | NO | NO | NA |
| Ankrd34c | ENSMUSG( | ENSMUSG( | 1.553506 | 0.345676 | NO | NO | NA |
| Tyr | ENSMUSG( | ENSMUSG( | 0.971864 | 0.574591 | NO | NO | NA |
| Nudt12 | ENSMUSG( | ENSMUSG( | 0.460739 | 0.450923 | NO | NO | NA |
| Cfhr1 | ENSMUSG( | ENSMUSG( | 2.589756 | 0.112856 | NO | NO | NA |
| Gm10566 | ENSMUSG( | ENSMUSG( | 1.761394 | 0.043233 | YES | YES | Upregulated |
| A330049N | ENSMUSG( | ENSMUSG( | 0.38391 | 0.810189 | NO | NO | NA |
| Ccdc152 | ENSMUSG( | ENSMUSG( | -0.35714 | 0.697385 | NO | NO | NA |
| Gm16166 | ENSMUSG( | ENSMUSG( | -0.49382 | 0.354579 | NO | NO | NA |
| Gm17093 | ENSMUSG( | ENSMUSG( | 0.820245 | 0.505754 | NO | NO | NA |
| Dalir | ENSMUSG( | ENSMUSG( | 0.626684 | 0.658376 | NO | NO | NA |
| Gm42701 | ENSMUSG( | ENSMUSG( | 1.793387 | 0.166736 | NO | NO | NA |
| Gm14342 | ENSMUSG( | ENSMUSG( | 0.142118 | 0.894564 | NO | NO | NA |
| Gm7502 | ENSMUSG( | ENSMUSG( | 0.384024 | 0.720058 | NO | NO | NA |
| Gm37390 | ENSMUSG( | ENSMUSG( | -0.47445 | 0.533517 | NO | NO | NA |
| Gm15580 | ENSMUSG( | ENSMUSG( | 0.210579 | 0.917064 | NO | NO | NA |
| Gm29811 | ENSMUSG( | ENSMUSG( | 0.279727 | 0.802041 | NO | NO | NA |
| Olfr1087 | ENSMUSG( | ENSMUSG( | 1.24517 | 0.394465 | NO | NO | NA |
| Gm20661 | ENSMUSG( | ENSMUSG( | 1.261197 | 0.26474 | NO | NO | NA |
| Gm11605 | ENSMUSG( | ENSMUSG( | 0.368472 | 0.535315 | NO | NO | NA |
| Gm47357 | ENSMUSG( | ENSMUSG( | -0.00244 | 0.996654 | NO | NO | NA |
| Trbj1-6 | ENSMUSG( | ENSMUSG( | 0.058064 | 0.96146 | NO | NO | NA |
| 4930466K: | ENSMUSG( | ENSMUSG( | 0.227003 | 0.803298 | NO | NO | NA |
| A130023I2 | ENSMUSG( | ENSMUSG( | 0.879883 | 0.508669 | NO | NO | NA |
| Gm11964 | ENSMUSG( | ENSMUSG( | 0.909429 | 0.239359 | NO | NO | NA |

| | | | | | | | |
|---|---|---|---|---|---|---|---|
| Rnps1-ps | ENSMUSG( | ENSMUSG( | 0.516211 | 0.328864 | NO | NO | NA |
| Trav7-2 | ENSMUSG( | ENSMUSG( | -5.8374 | NA | NO | NO | NA |
| Gm6032 | ENSMUSG( | ENSMUSG( | -0.77983 | 0.430295 | NO | NO | NA |
| ENSMUSG | ENSMUSG( | ENSMUSG( | 0.227421 | 0.725782 | NO | NO | NA |
| Gm16061 | ENSMUSG( | ENSMUSG( | 0.318959 | 0.73702 | NO | NO | NA |
| Gm5281 | ENSMUSG( | ENSMUSG( | 0.005059 | 0.997651 | NO | NO | NA |
| 4930556J2 | ENSMUSG( | ENSMUSG( | 0.635842 | 0.408636 | NO | NO | NA |
| Rpl5-ps1 | ENSMUSG( | ENSMUSG( | 0.426832 | 0.531711 | NO | NO | NA |
| Ighv4-1 | ENSMUSG( | ENSMUSG( | 0.699998 | 0.797665 | NO | NO | NA |
| Gm13092 | ENSMUSG( | ENSMUSG( | 0.381433 | 0.542733 | NO | NO | NA |
| Gm28175 | ENSMUSG( | ENSMUSG( | 0.500885 | 0.691861 | NO | NO | NA |
| Gm10517 | ENSMUSG( | ENSMUSG( | 0.538446 | 0.50818 | NO | NO | NA |
| Gm18722 | ENSMUSG( | ENSMUSG( | -0.48476 | 0.645299 | NO | NO | NA |
| Trav15n-1 | ENSMUSG( | ENSMUSG( | 2.100903 | 0.410802 | NO | NO | NA |
| Zim1 | ENSMUSG( | ENSMUSG( | 0.774372 | 0.520755 | NO | NO | NA |
| Olfr1263 | ENSMUSG( | ENSMUSG( | 0.844601 | 0.650865 | NO | NO | NA |
| Nmur1 | ENSMUSG( | ENSMUSG( | 0.628876 | 0.561209 | NO | NO | NA |
| Skint7 | ENSMUSG( | ENSMUSG( | 0.995521 | 0.461523 | NO | NO | NA |
| Gm35164 | ENSMUSG( | ENSMUSG( | 0.190588 | 0.888811 | NO | NO | NA |
| Gm49376 | ENSMUSG( | ENSMUSG( | -0.72688 | 0.522588 | NO | NO | NA |
| Fmr1nb | ENSMUSG( | ENSMUSG( | 3.341486 | NA | NO | NO | NA |
| Trbv21 | ENSMUSG( | ENSMUSG( | 0.943315 | 0.587391 | NO | NO | NA |
| Gm8108 | ENSMUSG( | ENSMUSG( | 3.544602 | 0.063705 | NO | NO | NA |
| Paqr6 | ENSMUSG( | ENSMUSG( | 0.82212 | 0.294263 | NO | NO | NA |
| Gm8104 | ENSMUSG( | ENSMUSG( | 0.754869 | 0.39556 | NO | NO | NA |
| ENSMUSG | ENSMUSG( | ENSMUSG( | 1.461517 | 0.006878 | YES | YES | Upregulated |
| Gm45644 | ENSMUSG( | ENSMUSG( | 0.03487 | 0.980908 | NO | NO | NA |
| Gm1826 | ENSMUSG( | ENSMUSG( | -0.18778 | 0.771401 | NO | NO | NA |
| Cfap20dc | ENSMUSG( | ENSMUSG( | 2.329192 | 0.005205 | YES | YES | Upregulated |
| ENSMUSG | ENSMUSG( | ENSMUSG( | -0.02485 | 0.977476 | NO | NO | NA |
| Gm13256 | ENSMUSG( | ENSMUSG( | 0.543905 | 0.472454 | NO | NO | NA |
| Gm37786 | ENSMUSG( | ENSMUSG( | 0.647877 | 0.400602 | NO | NO | NA |
| Gm19210 | ENSMUSG( | ENSMUSG( | 0.113606 | 0.945764 | NO | NO | NA |
| 1700003G | ENSMUSG( | ENSMUSG( | -0.04764 | 0.957599 | NO | NO | NA |
| Gm17473 | ENSMUSG( | ENSMUSG( | 0.583752 | 0.626073 | NO | NO | NA |
| Myh11 | ENSMUSG( | ENSMUSG( | 1.319192 | 0.138443 | NO | NO | NA |
| Olfr1258 | ENSMUSG( | ENSMUSG( | 0.226545 | 0.808567 | NO | NO | NA |
| Tas1r2 | ENSMUSG( | ENSMUSG( | 3.038447 | NA | NO | NO | NA |
| 4933402J1 | ENSMUSG( | ENSMUSG( | 0.799569 | 0.544012 | NO | NO | NA |
| Gm16315 | ENSMUSG( | ENSMUSG( | 0.466117 | 0.553133 | NO | NO | NA |
| Gm12630 | ENSMUSG( | ENSMUSG( | 0.638342 | 0.526045 | NO | NO | NA |
| A930002H | ENSMUSG( | ENSMUSG( | 0.936813 | 0.460843 | NO | NO | NA |
| Gm43450 | ENSMUSG( | ENSMUSG( | 0.021017 | 0.987528 | NO | NO | NA |
| Pknox2 | ENSMUSG( | ENSMUSG( | 0.681473 | 0.564209 | NO | NO | NA |
| Klf17 | ENSMUSG( | ENSMUSG( | 2.047241 | 0.185875 | NO | NO | NA |
| Ms4a15 | ENSMUSG( | ENSMUSG( | -0.5451 | 0.647726 | NO | NO | NA |
| Gm37053 | ENSMUSG( | ENSMUSG( | -0.3532 | 0.646271 | NO | NO | NA |

| | | | | | | | |
|---|---|---|---|---|---|---|---|
| 4930549G | ENSMUSG( | ENSMUSG( | -0.14234 | 0.851437 | NO | NO | NA |
| Gm53015 | ENSMUSG( | ENSMUSG( | -4.19327 | 0.138443 | NO | NO | NA |
| Nppb | ENSMUSG( | ENSMUSG( | 1.337256 | 0.321549 | NO | NO | NA |
| ENSMUSG | ENSMUSG( | ENSMUSG( | 0.912793 | 0.470363 | NO | NO | NA |
| Cetn1 | ENSMUSG( | ENSMUSG( | 0.527202 | 0.727144 | NO | NO | NA |
| 2210406H | ENSMUSG( | ENSMUSG( | 0.496902 | 0.476823 | NO | NO | NA |
| Gm7530 | ENSMUSG( | ENSMUSG( | 1.024245 | 0.370476 | NO | NO | NA |
| Gm16982 | ENSMUSG( | ENSMUSG( | 0.179462 | 0.738771 | NO | NO | NA |
| Gm10248 | ENSMUSG( | ENSMUSG( | -0.2396 | 0.90316 | NO | NO | NA |
| Gdf6 | ENSMUSG( | ENSMUSG( | 0.513321 | 0.732776 | NO | NO | NA |
| Olfr420 | ENSMUSG( | ENSMUSG( | 1.554013 | 0.278583 | NO | NO | NA |
| Gm17530 | ENSMUSG( | ENSMUSG( | 0.234961 | 0.726573 | NO | NO | NA |
| Gm12519 | ENSMUSG( | ENSMUSG( | 1.515988 | 0.203489 | NO | NO | NA |
| ENSMUSG | ENSMUSG( | ENSMUSG( | -0.00558 | 0.993583 | NO | NO | NA |
| Angptl7 | ENSMUSG( | ENSMUSG( | 0.551114 | 0.551583 | NO | NO | NA |
| Ces1f | ENSMUSG( | ENSMUSG( | 0.862761 | 0.513791 | NO | NO | NA |
| Gm16573 | ENSMUSG( | ENSMUSG( | 0.240414 | 0.726573 | NO | NO | NA |
| Gm20089 | ENSMUSG( | ENSMUSG( | 0.164937 | 0.919297 | NO | NO | NA |
| 4921525O | ENSMUSG( | ENSMUSG( | 0.581599 | 0.44899 | NO | NO | NA |
| Gm28826 | ENSMUSG( | ENSMUSG( | 1.849526 | 0.331789 | NO | NO | NA |
| Rarres1 | ENSMUSG( | ENSMUSG( | 0.19388 | 0.887464 | NO | NO | NA |
| Gm45231 | ENSMUSG( | ENSMUSG( | -0.06534 | 0.933155 | NO | NO | NA |
| ENSMUSG | ENSMUSG( | ENSMUSG( | 0.723299 | 0.62299 | NO | NO | NA |
| Gm19782 | ENSMUSG( | ENSMUSG( | 0.446553 | 0.737054 | NO | NO | NA |
| Gm43845 | ENSMUSG( | ENSMUSG( | 1.051116 | 0.573325 | NO | NO | NA |
| Gm19272 | ENSMUSG( | ENSMUSG( | 0.36421 | 0.778559 | NO | NO | NA |
| Gm50277 | ENSMUSG( | ENSMUSG( | 3.051054 | 0.291658 | NO | NO | NA |
| Gm48752 | ENSMUSG( | ENSMUSG( | 0.595287 | 0.671253 | NO | NO | NA |
| Gm15394 | ENSMUSG( | ENSMUSG( | 0.666895 | 0.525251 | NO | NO | NA |
| Mylk4 | ENSMUSG( | ENSMUSG( | -0.06038 | 0.961918 | NO | NO | NA |
| Gm5312 | ENSMUSG( | ENSMUSG( | 0.692107 | 0.556665 | NO | NO | NA |
| Gm43132 | ENSMUSG( | ENSMUSG( | 1.198038 | 0.349653 | NO | NO | NA |
| 4930465I2 | ENSMUSG( | ENSMUSG( | 0.86099 | 0.653801 | NO | NO | NA |
| Gm5921 | ENSMUSG( | ENSMUSG( | 0.436025 | 0.535131 | NO | NO | NA |
| C9 | ENSMUSG( | ENSMUSG( | 1.677336 | 0.130018 | NO | NO | NA |
| Gm36595 | ENSMUSG( | ENSMUSG( | 0.36938 | 0.786629 | NO | NO | NA |
| Gm12057 | ENSMUSG( | ENSMUSG( | 0.315772 | 0.701529 | NO | NO | NA |
| Gm3671 | ENSMUSG( | ENSMUSG( | 0.567547 | 0.55053 | NO | NO | NA |
| Rab9b | ENSMUSG( | ENSMUSG( | 0.74763 | 0.604949 | NO | NO | NA |
| Ogn | ENSMUSG( | ENSMUSG( | 0.242331 | 0.835045 | NO | NO | NA |
| Gm15696 | ENSMUSG( | ENSMUSG( | 0.235675 | 0.738158 | NO | NO | NA |
| 2610035F2 | ENSMUSG( | ENSMUSG( | 0.481449 | 0.724236 | NO | NO | NA |
| Tmem74b | ENSMUSG( | ENSMUSG( | 0.94999 | 0.473873 | NO | NO | NA |
| Gm8170 | ENSMUSG( | ENSMUSG( | -0.1259 | 0.901489 | NO | NO | NA |
| Gm2424 | ENSMUSG( | ENSMUSG( | 0.755435 | 0.62899 | NO | NO | NA |
| Trav4-2 | ENSMUSG( | ENSMUSG( | -0.80415 | 0.514914 | NO | NO | NA |
| Gm5523 | ENSMUSG( | ENSMUSG( | 1.597105 | 0.377713 | NO | NO | NA |

| | | | | | | | |
|---|---|---|---|---|---|---|---|
| ENSMUSG | ENSMUSG( | ENSMUSG( | 0.70626 | 0.572489 | NO | NO | NA |
| Btnl4 | ENSMUSG( | ENSMUSG( | -0.53919 | 0.510765 | NO | NO | NA |
| Olfr1251 | ENSMUSG( | ENSMUSG( | 1.690193 | 0.381121 | NO | NO | NA |
| Gm15127 | ENSMUSG( | ENSMUSG( | 1.371324 | 0.491395 | NO | NO | NA |
| Ern2 | ENSMUSG( | ENSMUSG( | 0.763826 | 0.504097 | NO | NO | NA |
| Zfp979 | ENSMUSG( | ENSMUSG( | 1.62304 | 0.111828 | NO | NO | NA |
| 9430073C: | ENSMUSG( | ENSMUSG( | 1.239539 | 0.412303 | NO | NO | NA |
| Gm6129 | ENSMUSG( | ENSMUSG( | 0.707184 | 0.377259 | NO | NO | NA |
| Vmn1r74 | ENSMUSG( | ENSMUSG( | 0.385984 | 0.741426 | NO | NO | NA |
| Olfr1131 | ENSMUSG( | ENSMUSG( | 0.599981 | 0.615791 | NO | NO | NA |
| 5430402P( | ENSMUSG( | ENSMUSG( | 1.27588 | 0.437342 | NO | NO | NA |
| Gm6443 | ENSMUSG( | ENSMUSG( | 0.6159 | 0.5876 | NO | NO | NA |
| Gm8850 | ENSMUSG( | ENSMUSG( | -1.19502 | 0.054751 | NO | NO | NA |
| Gm48670 | ENSMUSG( | ENSMUSG( | -0.05347 | 0.954162 | NO | NO | NA |
| Olfr1396 | ENSMUSG( | ENSMUSG( | -0.49475 | 0.513679 | NO | NO | NA |
| Fscn2 | ENSMUSG( | ENSMUSG( | 0.321075 | 0.647539 | NO | NO | NA |
| Nxph1 | ENSMUSG( | ENSMUSG( | 0.83953 | 0.483813 | NO | NO | NA |
| Gm37873 | ENSMUSG( | ENSMUSG( | 0.291435 | 0.767165 | NO | NO | NA |
| Gm6526 | ENSMUSG( | ENSMUSG( | 0.708006 | 0.582357 | NO | NO | NA |
| Adam12 | ENSMUSG( | ENSMUSG( | 0.34654 | 0.837865 | NO | NO | NA |
| Ano1 | ENSMUSG( | ENSMUSG( | 0.37536 | 0.699848 | NO | NO | NA |
| 6720475M | ENSMUSG( | ENSMUSG( | 0.173171 | 0.851529 | NO | NO | NA |
| D030062O | ENSMUSG( | ENSMUSG( | -0.11092 | 0.932346 | NO | NO | NA |
| Gm48008 | ENSMUSG( | ENSMUSG( | 0.394308 | 0.779954 | NO | NO | NA |
| Spag17 | ENSMUSG( | ENSMUSG( | 0.620937 | 0.688851 | NO | NO | NA |
| Gm28724 | ENSMUSG( | ENSMUSG( | 1.550924 | 0.325061 | NO | NO | NA |
| Gm31659 | ENSMUSG( | ENSMUSG( | 1.92206 | 0.224491 | NO | NO | NA |
| Fgd1 | ENSMUSG( | ENSMUSG( | -0.39983 | 0.459773 | NO | NO | NA |
| Gm40787 | ENSMUSG( | ENSMUSG( | 1.059667 | 0.44719 | NO | NO | NA |
| Gm4111 | ENSMUSG( | ENSMUSG( | 0.353349 | 0.775159 | NO | NO | NA |
| ENSMUSG | ENSMUSG( | ENSMUSG( | 0.692687 | 0.396816 | NO | NO | NA |
| Gm48313 | ENSMUSG( | ENSMUSG( | 0.858449 | 0.556901 | NO | NO | NA |
| Zfp960 | ENSMUSG( | ENSMUSG( | 0.546194 | 0.683874 | NO | NO | NA |
| Mpzl1 | ENSMUSG( | ENSMUSG( | 0.802387 | 0.269321 | NO | NO | NA |
| Trpa1 | ENSMUSG( | ENSMUSG( | 1.46496 | 0.216002 | NO | NO | NA |
| Gm16213 | ENSMUSG( | ENSMUSG( | -1.09523 | 0.046498 | YES | YES | Downregulated |
| Gm38216 | ENSMUSG( | ENSMUSG( | 0.535347 | 0.732776 | NO | NO | NA |
| Gm42721 | ENSMUSG( | ENSMUSG( | 0.134155 | 0.898999 | NO | NO | NA |
| ENSMUSG | ENSMUSG( | ENSMUSG( | 0.890013 | 0.541393 | NO | NO | NA |
| Gm42524 | ENSMUSG( | ENSMUSG( | 1.024359 | 0.50674 | NO | NO | NA |
| Apol9a | ENSMUSG( | ENSMUSG( | -0.13942 | 0.905135 | NO | NO | NA |
| Faddos | ENSMUSG( | ENSMUSG( | 0.530494 | 0.603756 | NO | NO | NA |
| 4930471G | ENSMUSG( | ENSMUSG( | 0.391102 | 0.693776 | NO | NO | NA |
| B230208B( | ENSMUSG( | ENSMUSG( | 1.611699 | 0.371752 | NO | NO | NA |
| Gm26784 | ENSMUSG( | ENSMUSG( | 0.984604 | 0.385531 | NO | NO | NA |
| Gm42895 | ENSMUSG( | ENSMUSG( | 0.432798 | 0.696353 | NO | NO | NA |
| Gm29684 | ENSMUSG( | ENSMUSG( | 1.994335 | 0.292762 | NO | NO | NA |

| | | | | | | | |
|---|---|---|---|---|---|---|---|
| Gm42497 | ENSMUSG( | ENSMUSG( | 0.00448 | 0.997702 | NO | NO | NA |
| Gm13446 | ENSMUSG( | ENSMUSG( | 1.816552 | 0.208913 | NO | NO | NA |
| Gm5733 | ENSMUSG( | ENSMUSG( | 0.227092 | 0.888811 | NO | NO | NA |
| Pdgfc | ENSMUSG( | ENSMUSG( | -0.06312 | 0.936437 | NO | NO | NA |
| Lhx3 | ENSMUSG( | ENSMUSG( | 0.700243 | 0.554054 | NO | NO | NA |
| Gm5117 | ENSMUSG( | ENSMUSG( | 0.669567 | 0.624878 | NO | NO | NA |
| Ptx4 | ENSMUSG( | ENSMUSG( | -0.90381 | 0.49246 | NO | NO | NA |
| Ldhb-ps | ENSMUSG( | ENSMUSG( | 0.725973 | 0.539391 | NO | NO | NA |
| Gdf1 | ENSMUSG( | ENSMUSG( | 1.447161 | 0.615867 | NO | NO | NA |
| Gm42490 | ENSMUSG( | ENSMUSG( | 0.684546 | 0.584042 | NO | NO | NA |
| Fcer2a | ENSMUSG( | ENSMUSG( | 0.862727 | 0.43785 | NO | NO | NA |
| Alox5 | ENSMUSG( | ENSMUSG( | 0.66073 | 0.467432 | NO | NO | NA |
| Gm34059 | ENSMUSG( | ENSMUSG( | 1.44963 | 0.36899 | NO | NO | NA |
| Gm32175 | ENSMUSG( | ENSMUSG( | 1.489604 | 0.209279 | NO | NO | NA |
| Aldh1l2 | ENSMUSG( | ENSMUSG( | 0.808846 | 0.573488 | NO | NO | NA |
| Tubb1 | ENSMUSG( | ENSMUSG( | -0.55391 | 0.696668 | NO | NO | NA |
| 8030474K( | ENSMUSG( | ENSMUSG( | 4.601114 | NA | NO | NO | NA |
| Fzd8 | ENSMUSG( | ENSMUSG( | 1.808836 | 0.020204 | YES | YES | Upregulated |
| Gm15839 | ENSMUSG( | ENSMUSG( | 1.02512 | 0.471757 | NO | NO | NA |
| ENSMUSG( | ENSMUSG( | ENSMUSG( | 1.485734 | 0.656736 | NO | NO | NA |
| Gm28140 | ENSMUSG( | ENSMUSG( | 1.119279 | 0.436017 | NO | NO | NA |
| Gm30414 | ENSMUSG( | ENSMUSG( | 0.974778 | 0.48613 | NO | NO | NA |
| Gm4799 | ENSMUSG( | ENSMUSG( | 0.339828 | 0.592899 | NO | NO | NA |
| 2810459M | ENSMUSG( | ENSMUSG( | 0.893991 | 0.471322 | NO | NO | NA |
| Pik3c2g | ENSMUSG( | ENSMUSG( | 0.941943 | 0.557591 | NO | NO | NA |
| Gm4567 | ENSMUSG( | ENSMUSG( | -0.23999 | 0.946243 | NO | NO | NA |
| Gm14258 | ENSMUSG( | ENSMUSG( | 0.842349 | 0.66539 | NO | NO | NA |
| Gm5157 | ENSMUSG( | ENSMUSG( | -0.71695 | 0.688848 | NO | NO | NA |
| Gm36899 | ENSMUSG( | ENSMUSG( | 1.395565 | 0.101829 | NO | NO | NA |
| Wnt16 | ENSMUSG( | ENSMUSG( | 0.335677 | 0.632543 | NO | NO | NA |
| Gm48079 | ENSMUSG( | ENSMUSG( | 0.341663 | 0.624649 | NO | NO | NA |
| Pcdhga8 | ENSMUSG( | ENSMUSG( | 0.575451 | 0.72738 | NO | NO | NA |
| 3110009F2 | ENSMUSG( | ENSMUSG( | 0.101431 | 0.947061 | NO | NO | NA |
| Armh2 | ENSMUSG( | ENSMUSG( | 0.176439 | 0.865309 | NO | NO | NA |
| Gm13240 | ENSMUSG( | ENSMUSG( | 0.040732 | 0.978405 | NO | NO | NA |
| Zscan10 | ENSMUSG( | ENSMUSG( | -0.52913 | 0.642085 | NO | NO | NA |
| Caly | ENSMUSG( | ENSMUSG( | 0.149471 | 0.926296 | NO | NO | NA |
| Gm8920 | ENSMUSG( | ENSMUSG( | -1.55495 | 0.338943 | NO | NO | NA |
| Gm44936 | ENSMUSG( | ENSMUSG( | 0.599788 | 0.612737 | NO | NO | NA |
| ENSMUSG | ENSMUSG( | ENSMUSG( | 0.596415 | 0.713566 | NO | NO | NA |
| Gm19967 | ENSMUSG( | ENSMUSG( | -0.12387 | 0.883252 | NO | NO | NA |
| Gm39468 | ENSMUSG( | ENSMUSG( | 0.536598 | 0.783586 | NO | NO | NA |
| Myo3a | ENSMUSG( | ENSMUSG( | 0.438328 | 0.734336 | NO | NO | NA |
| Gm48204 | ENSMUSG( | ENSMUSG( | 0.206275 | 0.841312 | NO | NO | NA |
| Gm15593 | ENSMUSG( | ENSMUSG( | 1.264249 | 0.472454 | NO | NO | NA |
| Olfr914 | ENSMUSG( | ENSMUSG( | 0.950439 | 0.502423 | NO | NO | NA |
| Gm16046 | ENSMUSG( | ENSMUSG( | -0.3808 | 0.681191 | NO | NO | NA |

| | | | | | | | |
|---|---|---|---|---|---|---|---|
| Gm16177 | ENSMUSG( | ENSMUSG( | 0.308634 | 0.794235 | NO | NO | NA |
| Sybu | ENSMUSG( | ENSMUSG( | 0.636565 | 0.614707 | NO | NO | NA |
| Slc25a18 | ENSMUSG( | ENSMUSG( | -1.25441 | 0.095703 | NO | NO | NA |
| Olfr297 | ENSMUSG( | ENSMUSG( | 1.318892 | 0.267056 | NO | NO | NA |
| Gm31045 | ENSMUSG( | ENSMUSG( | -0.0193 | 0.989102 | NO | NO | NA |
| ENSMUSG | ENSMUSG( | ENSMUSG( | 0.968186 | 0.516977 | NO | NO | NA |
| D9Wsu149 | ENSMUSG( | ENSMUSG( | 1.059779 | 0.369 | NO | NO | NA |
| Gm43707 | ENSMUSG( | ENSMUSG( | 0.642026 | 0.515526 | NO | NO | NA |
| Nek10 | ENSMUSG( | ENSMUSG( | 0.110496 | 0.886726 | NO | NO | NA |
| ENSMUSG | ENSMUSG( | ENSMUSG( | -0.85885 | 0.205774 | NO | NO | NA |
| Psg29 | ENSMUSG( | ENSMUSG( | 0.703994 | 0.519758 | NO | NO | NA |
| Gm10371 | ENSMUSG( | ENSMUSG( | 0.469392 | 0.692874 | NO | NO | NA |
| Gm42646 | ENSMUSG( | ENSMUSG( | 0.341982 | 0.772331 | NO | NO | NA |
| Cep126 | ENSMUSG( | ENSMUSG( | 1.078555 | 0.386598 | NO | NO | NA |
| Gm10416 | ENSMUSG( | ENSMUSG( | 0.086489 | 0.921653 | NO | NO | NA |
| 3110099E( | ENSMUSG( | ENSMUSG( | 0.979684 | 0.400823 | NO | NO | NA |
| Gm2546 | ENSMUSG( | ENSMUSG( | -0.03249 | 0.970155 | NO | NO | NA |
| Slc4a9 | ENSMUSG( | ENSMUSG( | -0.03282 | 0.976454 | NO | NO | NA |
| F2 | ENSMUSG( | ENSMUSG( | -0.22636 | 0.85786 | NO | NO | NA |
| ENSMUSG( | ENSMUSG( | ENSMUSG( | 0.476007 | 0.676289 | NO | NO | NA |
| Plekhb1 | ENSMUSG( | ENSMUSG( | -0.03832 | 0.968083 | NO | NO | NA |
| Spem2 | ENSMUSG( | ENSMUSG( | 0.861909 | 0.626178 | NO | NO | NA |
| 4930447F2 | ENSMUSG( | ENSMUSG( | 0.833474 | 0.244063 | NO | NO | NA |
| Gm3126 | ENSMUSG( | ENSMUSG( | -1.15822 | 0.278615 | NO | NO | NA |
| Pnpla1 | ENSMUSG( | ENSMUSG( | -0.50632 | 0.55012 | NO | NO | NA |
| Snord3b4 | ENSMUSG( | ENSMUSG( | -5.47871 | NA | NO | NO | NA |
| Gm5915 | ENSMUSG( | ENSMUSG( | 0.041375 | 0.950039 | NO | NO | NA |
| Gm42486 | ENSMUSG( | ENSMUSG( | 0.423049 | 0.744834 | NO | NO | NA |
| Gm16386 | ENSMUSG( | ENSMUSG( | -1.18743 | 0.207706 | NO | NO | NA |
| Gm5065 | ENSMUSG( | ENSMUSG( | 0.849699 | 0.437161 | NO | NO | NA |
| Gm37102 | ENSMUSG( | ENSMUSG( | -0.71508 | 0.423479 | NO | NO | NA |
| Gm30771 | ENSMUSG( | ENSMUSG( | 0.101675 | 0.898499 | NO | NO | NA |
| Olfr739 | ENSMUSG( | ENSMUSG( | 0.348243 | 0.78842 | NO | NO | NA |
| Gm18112 | ENSMUSG( | ENSMUSG( | 1.123779 | 0.478783 | NO | NO | NA |
| Gm26810 | ENSMUSG( | ENSMUSG( | 0.024784 | 0.968633 | NO | NO | NA |
| Prr23a2 | ENSMUSG( | ENSMUSG( | 0.155851 | 0.919733 | NO | NO | NA |
| Gm43268 | ENSMUSG( | ENSMUSG( | 0.907225 | 0.214073 | NO | NO | NA |
| Gm13141 | ENSMUSG( | ENSMUSG( | 1.081366 | 0.513919 | NO | NO | NA |
| Gm44530 | ENSMUSG( | ENSMUSG( | 0.18516 | 0.814334 | NO | NO | NA |
| Phldb2 | ENSMUSG( | ENSMUSG( | 0.332278 | 0.619252 | NO | NO | NA |
| Rpl31-ps18 | ENSMUSG( | ENSMUSG( | 0.173682 | 0.908038 | NO | NO | NA |
| Gm37258 | ENSMUSG( | ENSMUSG( | 0.465202 | 0.790442 | NO | NO | NA |
| Mok | ENSMUSG( | ENSMUSG( | 0.742349 | 0.574532 | NO | NO | NA |
| Gm12763 | ENSMUSG( | ENSMUSG( | 1.406456 | 0.336398 | NO | NO | NA |
| 4930558F1 | ENSMUSG( | ENSMUSG( | 1.153737 | 0.596414 | NO | NO | NA |
| Teddm3 | ENSMUSG( | ENSMUSG( | -0.25311 | 0.849635 | NO | NO | NA |
| Gm45165 | ENSMUSG( | ENSMUSG( | 0.086319 | 0.962199 | NO | NO | NA |

| | | | | | | | |
|---|---|---|---|---|---|---|---|
| Ccdc170 | ENSMUSG( | ENSMUSG( | 0.208104 | 0.835115 | NO | NO | NA |
| Slc38a3 | ENSMUSG( | ENSMUSG( | -1.82218 | 0.147915 | NO | NO | NA |
| ENSMUSG | ENSMUSG( | ENSMUSG( | -0.76183 | 0.369902 | NO | NO | NA |
| Fbln2 | ENSMUSG( | ENSMUSG( | 0.122327 | 0.89804 | NO | NO | NA |
| Gm15881 | ENSMUSG( | ENSMUSG( | 0.86755 | 0.688571 | NO | NO | NA |
| Gm18030 | ENSMUSG( | ENSMUSG( | 0.462495 | 0.664907 | NO | NO | NA |
| Gm13158 | ENSMUSG( | ENSMUSG( | -1.54453 | 0.205787 | NO | NO | NA |
| Gm4812 | ENSMUSG( | ENSMUSG( | -1.11516 | 0.455732 | NO | NO | NA |
| ENSMUSG | ENSMUSG( | ENSMUSG( | 0.029531 | 0.973749 | NO | NO | NA |
| 4933439K: | ENSMUSG( | ENSMUSG( | 0.159796 | 0.896294 | NO | NO | NA |
| Chrna1 | ENSMUSG( | ENSMUSG( | 0.455133 | 0.665321 | NO | NO | NA |
| Gm8178 | ENSMUSG( | ENSMUSG( | 1.520452 | 0.614336 | NO | NO | NA |
| Dnah3 | ENSMUSG( | ENSMUSG( | 0.630929 | 0.431414 | NO | NO | NA |
| 5730414N | ENSMUSG( | ENSMUSG( | 0.752231 | 0.245002 | NO | NO | NA |
| C4b | ENSMUSG( | ENSMUSG( | -0.46201 | 0.624706 | NO | NO | NA |
| Vmn1r39 | ENSMUSG( | ENSMUSG( | 0.716619 | 0.668541 | NO | NO | NA |
| Gm37025 | ENSMUSG( | ENSMUSG( | -0.18255 | 0.853948 | NO | NO | NA |
| Gm9256 | ENSMUSG( | ENSMUSG( | -0.707 | 0.661008 | NO | NO | NA |
| Gm18529 | ENSMUSG( | ENSMUSG( | 0.868648 | 0.576552 | NO | NO | NA |
| Tmem44 | ENSMUSG( | ENSMUSG( | 0.28897 | 0.758872 | NO | NO | NA |
| H3f4 | ENSMUSG( | ENSMUSG( | -0.73181 | 0.33196 | NO | NO | NA |
| Igkv4-86 | ENSMUSG( | ENSMUSG( | -4.4593 | 0.140549 | NO | NO | NA |
| ENSMUSG | ENSMUSG( | ENSMUSG( | 0.635125 | 0.39361 | NO | NO | NA |
| Upp2 | ENSMUSG( | ENSMUSG( | 1.69522 | 0.012458 | YES | YES | Upregulated |
| Gm33843 | ENSMUSG( | ENSMUSG( | 0.778193 | 0.490843 | NO | NO | NA |
| Ube2d2b | ENSMUSG( | ENSMUSG( | 0.545087 | 0.692623 | NO | NO | NA |
| Adam30 | ENSMUSG( | ENSMUSG( | 0.32109 | 0.773712 | NO | NO | NA |
| 4933412O | ENSMUSG( | ENSMUSG( | 2.173625 | 0.101346 | NO | NO | NA |
| Gm43330 | ENSMUSG( | ENSMUSG( | -0.7463 | 0.567382 | NO | NO | NA |
| Cemip | ENSMUSG( | ENSMUSG( | 1.37888 | 0.326393 | NO | NO | NA |
| ENSMUSG | ENSMUSG( | ENSMUSG( | 0.212919 | 0.896865 | NO | NO | NA |
| Olfr177 | ENSMUSG( | ENSMUSG( | 0.430032 | 0.762738 | NO | NO | NA |
| Gm8168 | ENSMUSG( | ENSMUSG( | -1.13939 | 0.489989 | NO | NO | NA |
| ENSMUSG( | ENSMUSG( | ENSMUSG( | 0.076219 | 0.951037 | NO | NO | NA |
| ENSMUSG | ENSMUSG( | ENSMUSG( | 0.490945 | 0.642775 | NO | NO | NA |
| Gm26705 | ENSMUSG( | ENSMUSG( | 1.447813 | 0.112548 | NO | NO | NA |
| Gm9029 | ENSMUSG( | ENSMUSG( | 0.619843 | 0.881489 | NO | NO | NA |
| Igkv4-58 | ENSMUSG( | ENSMUSG( | 4.575482 | 0.104188 | NO | NO | NA |
| Plscr2 | ENSMUSG( | ENSMUSG( | 0.629392 | 0.480639 | NO | NO | NA |
| Timd2 | ENSMUSG( | ENSMUSG( | 0.184285 | 0.889649 | NO | NO | NA |
| Chrng | ENSMUSG( | ENSMUSG( | 0.465952 | 0.699119 | NO | NO | NA |
| Gm49387 | ENSMUSG( | ENSMUSG( | -5.21103 | 0.194435 | NO | NO | NA |
| Gm37775 | ENSMUSG( | ENSMUSG( | 0.683665 | 0.621768 | NO | NO | NA |
| ENSMUSG | ENSMUSG( | ENSMUSG( | 0.790821 | 0.32874 | NO | NO | NA |
| Rps10-ps3 | ENSMUSG( | ENSMUSG( | 1.281757 | 0.408247 | NO | NO | NA |
| Gm49152 | ENSMUSG( | ENSMUSG( | 0.982275 | 0.468862 | NO | NO | NA |
| Gm1123 | ENSMUSG( | ENSMUSG( | -0.39334 | 0.681397 | NO | NO | NA |

| | | | | | | | |
|---|---|---|---|---|---|---|---|
| A630014C | ENSMUSG( | ENSMUSG( | 0.824866 | 0.544781 | NO | NO | NA |
| 2900079G | ENSMUSG( | ENSMUSG( | 0.131543 | 0.929773 | NO | NO | NA |
| Ctrb1 | ENSMUSG( | ENSMUSG( | 0.311379 | 0.888378 | NO | NO | NA |
| Gm2058 | ENSMUSG( | ENSMUSG( | -0.27765 | 0.709959 | NO | NO | NA |
| Gm33472 | ENSMUSG( | ENSMUSG( | 0.039451 | 0.983122 | NO | NO | NA |
| Gm43056 | ENSMUSG( | ENSMUSG( | -0.36364 | 0.823786 | NO | NO | NA |
| Psors1c2 | ENSMUSG( | ENSMUSG( | 0.686748 | 0.440741 | NO | NO | NA |
| Spdye4a | ENSMUSG( | ENSMUSG( | 1.702257 | 0.176203 | NO | NO | NA |
| Gm32509 | ENSMUSG( | ENSMUSG( | 1.669936 | 0.461018 | NO | NO | NA |
| Slit1 | ENSMUSG( | ENSMUSG( | 0.91738 | 0.303357 | NO | NO | NA |
| Gm7183 | ENSMUSG( | ENSMUSG( | 0.213254 | 0.894999 | NO | NO | NA |
| Gm4956 | ENSMUSG( | ENSMUSG( | -5.28943 | 0.080687 | NO | NO | NA |
| Medag | ENSMUSG( | ENSMUSG( | -0.23616 | 0.867268 | NO | NO | NA |
| Igkv4-74 | ENSMUSG( | ENSMUSG( | -2.33011 | 0.334697 | NO | NO | NA |
| F830016B( | ENSMUSG( | ENSMUSG( | -0.1352 | 0.874608 | NO | NO | NA |
| Serpinb9g | ENSMUSG( | ENSMUSG( | 1.912563 | 0.120559 | NO | NO | NA |
| Crispld1 | ENSMUSG( | ENSMUSG( | 1.507396 | 0.571509 | NO | NO | NA |
| Gm5860 | ENSMUSG( | ENSMUSG( | 0.479787 | 0.796087 | NO | NO | NA |
| 7120432I0 | ENSMUSG( | ENSMUSG( | 1.340243 | 0.460561 | NO | NO | NA |
| Gm11198 | ENSMUSG( | ENSMUSG( | 0.410797 | 0.822566 | NO | NO | NA |
| Gm26444 | ENSMUSG( | ENSMUSG( | 3.025256 | 0.482239 | NO | NO | NA |
| Gm5576 | ENSMUSG( | ENSMUSG( | 0.992612 | 0.203595 | NO | NO | NA |
| Gm47036 | ENSMUSG( | ENSMUSG( | 0.777344 | 0.558551 | NO | NO | NA |
| Gm15429 | ENSMUSG( | ENSMUSG( | -0.19163 | 0.816509 | NO | NO | NA |
| Gm30270 | ENSMUSG( | ENSMUSG( | 1.083847 | 0.638931 | NO | NO | NA |
| 1700036A | ENSMUSG( | ENSMUSG( | 0.518248 | 0.802684 | NO | NO | NA |
| Gm8146 | ENSMUSG( | ENSMUSG( | 0.017277 | 0.983245 | NO | NO | NA |
| Gm2534 | ENSMUSG( | ENSMUSG( | 1.713463 | 0.223803 | NO | NO | NA |
| ENSMUSG | ENSMUSG( | ENSMUSG( | 2.516927 | 0.13582 | NO | NO | NA |
| Ighv3-8 | ENSMUSG( | ENSMUSG( | -4.02285 | 0.205952 | NO | NO | NA |
| Gm2314 | ENSMUSG( | ENSMUSG( | 1.21983 | 0.425837 | NO | NO | NA |
| D630002J1 | ENSMUSG( | ENSMUSG( | -0.67782 | 0.661008 | NO | NO | NA |
| Gm13170 | ENSMUSG( | ENSMUSG( | 0.70301 | 0.716158 | NO | NO | NA |
| Btf3-ps2 | ENSMUSG( | ENSMUSG( | 1.138748 | 0.50831 | NO | NO | NA |
| Smok4a | ENSMUSG( | ENSMUSG( | 2.419249 | 0.571926 | NO | NO | NA |
| Gm26714 | ENSMUSG( | ENSMUSG( | -1.46921 | 0.300676 | NO | NO | NA |
| Gm3164 | ENSMUSG( | ENSMUSG( | -0.42198 | 0.717004 | NO | NO | NA |
| Igkv3-1 | ENSMUSG( | ENSMUSG( | -3.88951 | 0.166132 | NO | NO | NA |
| Gm28707 | ENSMUSG( | ENSMUSG( | 0.225624 | 0.729805 | NO | NO | NA |
| Gm32089 | ENSMUSG( | ENSMUSG( | 0.474108 | 0.748515 | NO | NO | NA |
| E230016K1 | ENSMUSG( | ENSMUSG( | -0.08025 | 0.924792 | NO | NO | NA |
| Kcnh6 | ENSMUSG( | ENSMUSG( | -1.80828 | 0.103888 | NO | NO | NA |
| Olfr1458 | ENSMUSG( | ENSMUSG( | -1.62929 | 0.461948 | NO | NO | NA |
| Gm49931 | ENSMUSG( | ENSMUSG( | 0.85581 | 0.636057 | NO | NO | NA |
| Prss32 | ENSMUSG( | ENSMUSG( | -0.94067 | 0.538735 | NO | NO | NA |
| Tsacc | ENSMUSG( | ENSMUSG( | NA | NA | NO | NO | NA |
| Clhc1 | ENSMUSG( | ENSMUSG( | NA | NA | NO | NO | NA |

| | | | | | | | |
|---|---|---|---|---|---|---|---|
| Fkbp1b | ENSMUSG( | ENSMUSG( | NA | NA | NO | NO | NA |
| Ccl1 | ENSMUSG( | ENSMUSG( | NA | NA | NO | NO | NA |
| Nid2 | ENSMUSG( | ENSMUSG( | NA | NA | NO | NO | NA |
| Dnah12 | ENSMUSG( | ENSMUSG( | NA | NA | NO | NO | NA |
| Ajuba | ENSMUSG( | ENSMUSG( | NA | NA | NO | NO | NA |
| Pdia5 | ENSMUSG( | ENSMUSG( | NA | NA | NO | NO | NA |
| Trpm6 | ENSMUSG( | ENSMUSG( | NA | NA | NO | NO | NA |
| Itpka | ENSMUSG( | ENSMUSG( | NA | NA | NO | NO | NA |
| Mall | ENSMUSG( | ENSMUSG( | NA | NA | NO | NO | NA |
| Rspo1 | ENSMUSG( | ENSMUSG( | NA | NA | NO | NO | NA |
| Dpysl5 | ENSMUSG( | ENSMUSG( | NA | NA | NO | NO | NA |
| Crybb1 | ENSMUSG( | ENSMUSG( | NA | NA | NO | NO | NA |
| Ppef2 | ENSMUSG( | ENSMUSG( | NA | NA | NO | NO | NA |
| Tmtc1 | ENSMUSG( | ENSMUSG( | NA | NA | NO | NO | NA |
| Nucb2 | ENSMUSG( | ENSMUSG( | NA | NA | NO | NO | NA |
| Rab33a | ENSMUSG( | ENSMUSG( | NA | NA | NO | NO | NA |
| Cd40lg | ENSMUSG( | ENSMUSG( | NA | NA | NO | NO | NA |
| Xkrx | ENSMUSG( | ENSMUSG( | NA | NA | NO | NO | NA |
| Me1 | ENSMUSG( | ENSMUSG( | NA | NA | NO | NO | NA |
| Ppp1r32 | ENSMUSG( | ENSMUSG( | NA | NA | NO | NO | NA |
| Ppp1r26 | ENSMUSG( | ENSMUSG( | NA | NA | NO | NO | NA |
| Clba1 | ENSMUSG( | ENSMUSG( | NA | NA | NO | NO | NA |
| Minar1 | ENSMUSG( | ENSMUSG( | NA | NA | NO | NO | NA |
| Prrx2 | ENSMUSG( | ENSMUSG( | NA | NA | NO | NO | NA |
| Dock3 | ENSMUSG( | ENSMUSG( | NA | NA | NO | NO | NA |
| Plcb4 | ENSMUSG( | ENSMUSG( | NA | NA | NO | NO | NA |
| Aicda | ENSMUSG( | ENSMUSG( | NA | NA | NO | NO | NA |
| Hesx1 | ENSMUSG( | ENSMUSG( | NA | NA | NO | NO | NA |
| C1qtnf4 | ENSMUSG( | ENSMUSG( | NA | NA | NO | NO | NA |
| Fbxo2 | ENSMUSG( | ENSMUSG( | NA | NA | NO | NO | NA |
| Khdc1c | ENSMUSG( | ENSMUSG( | NA | NA | NO | NO | NA |
| H1f8 | ENSMUSG( | ENSMUSG( | NA | NA | NO | NO | NA |
| Mgat3 | ENSMUSG( | ENSMUSG( | NA | NA | NO | NO | NA |
| Gm7729 | ENSMUSG( | ENSMUSG( | NA | NA | NO | NO | NA |
| Psma5-ps | ENSMUSG( | ENSMUSG( | NA | NA | NO | NO | NA |
| Fam171a1 | ENSMUSG( | ENSMUSG( | NA | NA | NO | NO | NA |
| Fam228b | ENSMUSG( | ENSMUSG( | NA | NA | NO | NO | NA |
| Gm6139 | ENSMUSG( | ENSMUSG( | NA | NA | NO | NO | NA |
| Mab21l2 | ENSMUSG( | ENSMUSG( | NA | NA | NO | NO | NA |
| Zfp105 | ENSMUSG( | ENSMUSG( | NA | NA | NO | NO | NA |
| 4933430I1 | ENSMUSG( | ENSMUSG( | NA | NA | NO | NO | NA |
| Gm8116 | ENSMUSG( | ENSMUSG( | NA | NA | NO | NO | NA |
| Hacd1 | ENSMUSG( | ENSMUSG( | NA | NA | NO | NO | NA |
| Ccdc146 | ENSMUSG( | ENSMUSG( | NA | NA | NO | NO | NA |
| Gm5529 | ENSMUSG( | ENSMUSG( | NA | NA | NO | NO | NA |
| Gm7931 | ENSMUSG( | ENSMUSG( | NA | NA | NO | NO | NA |
| Gm10221 | ENSMUSG( | ENSMUSG( | NA | NA | NO | NO | NA |

| | | | | | | | |
|---|---|---|---|---|---|---|---|
| Fcrl6 | ENSMUSG( | ENSMUSG( | NA | NA | NO | NO | NA |
| Amy2a5 | ENSMUSG( | ENSMUSG( | NA | NA | NO | NO | NA |
| Gm15417 | ENSMUSG( | ENSMUSG( | NA | NA | NO | NO | NA |
| Ism1 | ENSMUSG( | ENSMUSG( | NA | NA | NO | NO | NA |
| 4930558J2 | ENSMUSG( | ENSMUSG( | NA | NA | NO | NO | NA |
| Trav15-2-c | ENSMUSG( | ENSMUSG( | NA | NA | NO | NO | NA |
| Trdv2-2 | ENSMUSG( | ENSMUSG( | NA | NA | NO | NO | NA |
| Trdv5 | ENSMUSG( | ENSMUSG( | NA | NA | NO | NO | NA |
| Rps12l1 | ENSMUSG( | ENSMUSG( | NA | NA | NO | NO | NA |
| Erich3 | ENSMUSG( | ENSMUSG( | NA | NA | NO | NO | NA |
| Nkain1 | ENSMUSG( | ENSMUSG( | NA | NA | NO | NO | NA |
| Smt3h2-ps | ENSMUSG( | ENSMUSG( | NA | NA | NO | NO | NA |
| Gm14780 | ENSMUSG( | ENSMUSG( | NA | NA | NO | NO | NA |
| Rpl3-ps2 | ENSMUSG( | ENSMUSG( | NA | NA | NO | NO | NA |
| Gm16104 | ENSMUSG( | ENSMUSG( | NA | NA | NO | NO | NA |
| Platr14 | ENSMUSG( | ENSMUSG( | NA | NA | NO | NO | NA |
| Gm12534 | ENSMUSG( | ENSMUSG( | NA | NA | NO | NO | NA |
| Gm4750 | ENSMUSG( | ENSMUSG( | NA | NA | NO | NO | NA |
| Gm13493 | ENSMUSG( | ENSMUSG( | NA | NA | NO | NO | NA |
| Gm2541 | ENSMUSG( | ENSMUSG( | NA | NA | NO | NO | NA |
| Gm10171 | ENSMUSG( | ENSMUSG( | NA | NA | NO | NO | NA |
| Gm16163 | ENSMUSG( | ENSMUSG( | NA | NA | NO | NO | NA |
| Gm6158 | ENSMUSG( | ENSMUSG( | NA | NA | NO | NO | NA |
| Cdv3-ps | ENSMUSG( | ENSMUSG( | NA | NA | NO | NO | NA |
| Zfp422-ps | ENSMUSG( | ENSMUSG( | NA | NA | NO | NO | NA |
| Gm5778 | ENSMUSG( | ENSMUSG( | NA | NA | NO | NO | NA |
| Gm20479 | ENSMUSG( | ENSMUSG( | NA | NA | NO | NO | NA |
| Gm20531 | ENSMUSG( | ENSMUSG( | NA | NA | NO | NO | NA |
| Gm3788 | ENSMUSG( | ENSMUSG( | NA | NA | NO | NO | NA |
| Igkv3-5 | ENSMUSG( | ENSMUSG( | NA | NA | NO | NO | NA |
| Gm16793 | ENSMUSG( | ENSMUSG( | NA | NA | NO | NO | NA |
| Kansl2-ps | ENSMUSG( | ENSMUSG( | NA | NA | NO | NO | NA |
| Gm26636 | ENSMUSG( | ENSMUSG( | NA | NA | NO | NO | NA |
| Gm26916 | ENSMUSG( | ENSMUSG( | NA | NA | NO | NO | NA |
| Gm29123 | ENSMUSG( | ENSMUSG( | NA | NA | NO | NO | NA |
| Trpc2 | ENSMUSG( | ENSMUSG( | NA | NA | NO | NO | NA |
| Pcdhga4 | ENSMUSG( | ENSMUSG( | NA | NA | NO | NO | NA |
| 6820402A( | ENSMUSG( | ENSMUSG( | NA | NA | NO | NO | NA |
| Gm5837 | ENSMUSG( | ENSMUSG( | NA | NA | NO | NO | NA |
| D5Ertd605 | ENSMUSG( | ENSMUSG( | NA | NA | NO | NO | NA |
| Gm45155 | ENSMUSG( | ENSMUSG( | NA | NA | NO | NO | NA |
| Gm5596 | ENSMUSG( | ENSMUSG( | NA | NA | NO | NO | NA |
| Gm48646 | ENSMUSG( | ENSMUSG( | NA | NA | NO | NO | NA |
| Gm47327 | ENSMUSG( | ENSMUSG( | NA | NA | NO | NO | NA |
| Gm5425 | ENSMUSG( | ENSMUSG( | NA | NA | NO | NO | NA |
| Gm5182 | ENSMUSG( | ENSMUSG( | NA | NA | NO | NO | NA |
| Gm36099 | ENSMUSG( | ENSMUSG( | NA | NA | NO | NO | NA |

| | | | | | | | |
|---|---|---|---|---|---|---|---|
| Gm46339 | ENSMUSG( | ENSMUSG( | NA | NA | NO | NO | NA |
| Gm47870 | ENSMUSG( | ENSMUSG( | NA | NA | NO | NO | NA |
| Gm49355 | ENSMUSG( | ENSMUSG( | NA | NA | NO | NO | NA |
| Aldoa ENS | ENSMUSG( | ENSMUSG( | NA | NA | NO | NO | NA |
| Gm48933 | ENSMUSG( | ENSMUSG( | NA | NA | NO | NO | NA |
| 1700087I2 | ENSMUSG( | ENSMUSG( | NA | NA | NO | NO | NA |
| Gm50083 | ENSMUSG( | ENSMUSG( | NA | NA | NO | NO | NA |
| Gm35551 | ENSMUSG( | ENSMUSG( | NA | NA | NO | NO | NA |
| Gm25432 | ENSMUSG( | ENSMUSG( | NA | NA | NO | NO | NA |
| Gm50216 | ENSMUSG( | ENSMUSG( | NA | NA | NO | NO | NA |
| Gm22068 | ENSMUSG( | ENSMUSG( | NA | NA | NO | NO | NA |
| Snord3b1 | ENSMUSG( | ENSMUSG( | NA | NA | NO | NO | NA |
| Snord3b3 | ENSMUSG( | ENSMUSG( | NA | NA | NO | NO | NA |
| ENSMUSG | ENSMUSG( | ENSMUSG( | NA | NA | NO | NO | NA |
| ENSMUSG | ENSMUSG( | ENSMUSG( | NA | NA | NO | NO | NA |
| ENSMUSG | ENSMUSG( | ENSMUSG( | NA | NA | NO | NO | NA |
| ENSMUSG( | ENSMUSG( | ENSMUSG( | NA | NA | NO | NO | NA |
