## Supplemental Table 2 for "Upregulation of TNFR2 Precedes TOX Expression by Exhausted T cells and Restricts Antitumor and Antiviral Immunity"

**Glycolysis and glucose import**
Adh1
Aldh1a3
Aldh1b1
Aldh3a1
Aldh3b2
Aldoa
Aldob
Dld
Fbp2
Gapdhs
Gckr
Ldha
Ldhal6b
Nup54
Pck1
Pdhb
Ppp2ca
Ppp2cb
Ppp2r1a
Ppp2r1b
Slc2a12
Slc2a3
Slc2a6

**Type II Interferon**
Camk2d
Ciita
Fcgr1
Gbp2
Gbp2b
Icam1
Ifng
Irf2
Irf4
Irf8
Jak2
Oas1a
Oas2
Oas3
Pias1
Pml
Ptafr
Ptpn11
Ptpn6
Socs1
Socs3

Sp100
Sumo1
Trim2
Trim29
Trim35
Trim46
Vcam1

**IL-1 signaling**
Alox5
Btrc
Cul1
Hmgb1
Il13
Il18
Il1a
Il1b
Il1f9
Il1r1
Il1r2
Il1rapl1
Il1rl1
Il1rl2
Il1rn
Il4
Irak3
Irak4
Ptpn11
Ptpn18
Ptpn2
Ptpn6
Ptpn7
Rbx1
Tab2

**AP-1 signaling**
Batf
Cebpb
Fos
Jun
Junb
Jund
Mapk10
Mapk11
Mapk14
Mapk8
Mapk9

Nrl
Spag9

**RAR signaling**
Aldh1a1
Aldh1a3
Cyp26a1
Cyp26c1
Dhrs3
Dld
Pdhb
Pdk4
Rara
Rarb
Rxrg

**Fatty Acid metabolism**
Acaca
Acadl
Acadvl
Acat2
Acot3
Acsl3
Acsl4
Acsl6
Adh1
Aldh1b1
Cpt1c
Cyp4a12a/b
Cyp4a31/32
Echs1
Ehhadh
Elovl2
Elovl3
Elovl4
Elovl7
Hadhb
Pctp
Ppt1
Prkaa2

**Glutamine metabolism**
Gls
Gls2
Glud1
Got2
Oat

Pycr1
Pycr2
Rimkla

**PPAR signaling**
Acadl
Acsl3
Acsl4
Acsl6
Apoc3
Aqp7
Cpt1c
Cyp4a12a/b
Cyp4a31/32
Cyp7a1
Cyp8b1
Ehhadh
Fabp1
Fabp4
Gk
Lpl
Pck1
Pparg
Rxrg
Scp2
Ucp1

**IL-7 signaling**
Brwd1
Cish
Crlf2
Hgf
Il7
Il7r
Irs1
Rag1
Smarca4
Socs1
Socs2
Tslp

**cell cycle**
Anapc4
Atm
Atr
Ccna1
Ccnb1

Ccnb3
Cdc14b
Cdc16
Cdc20
Cdc25a
Cdkn1c
Cdkn2c
Chek1
Crebbp
Cul1
E2f1
E2f2
E2f5
Gadd45a
Gsk3b
Mcm5
Mcm7
Orc2
Orc4
Pkmyt1
Prkdc
Rbl2
Rbx1
Sfn
Smc3
Stag2
Ttk
Ywhaz

**T cell checkpoint signaling**
Adora2a
Btla
Cd200r1
Cd27
Cd274
Cd276
Cd28
Cd40
Cd40lg
Cd70
Cd80
Cd86
Cdh1
Ctla4
Havcr2
Icos
Icosl

Lag3
Pdcd1
Pdcd1lg2
Pecam1
Pvrig
Tigit
Tnfrsf14
Tnfrsf18
Tnfrsf4
Tnfrsf9
Tnfsf18
Tnfsf4
Tnfsf9
Vsir
Vtcn1

**NK receptors**
Cd160
Cd22
Cd226
Cd244a
Cd27
Cd33
Cd96
Ceacam1
Crtam
Fcer1g
Hcst
Kir3dl1/2
Klrb1
Klrb1a
Klrc1
Klrd1
Klrg1
Lair1
Sema4d
Siglec1
Slamf6
Tyrobp

**Chemokine signaling**
Ccl12
Ccl2
Ccl20
Ccl21a/b/c
Ccl22
Ccl3

Ccl4
Ccl5
Ccr1
Ccr2
Ccr5
Ccr7
Cmklr1
Cxcl1
Cxcl11
Cxcl9
Cxcr3
Cxcr6
Xcl1

**mTOR signaling**
Atp6v1g3
Dvl1
Eif4b
Fzd2
Fzd7
Grb10
Grb2
Gsk3b
Ins1
Irs1
Kras
Lamtor3
Lrp6
Nprl3
Ppm1a
Prkaa1
Prkaa2
Prkag3
Prkcb
Rheb
Rictor
Rragc
Sesn2
Slc7a5
Sos1
Sos2
Telo2
Tnfrsf1a
Wnt11
Wnt2
Wnt3
Wnt3a

Wnt4
Wnt6

**IL-6 signaling**
Cbl
Clcf1
Cntf
Crlf1
Ctf1
Il11
Il11ra1/2
Il31
Il31ra
Il6
Il6ra
Il6st
Jak2
Lif
Lifr
Osm
Osmr
Ptpn11
Socs3
Tyk2

**IL-2 signaling**
Csf2
Grb2
Havcr2
Il15
Il2
Il21
Il21r
Il2ra
Il3
Il5
Il9
Il9r
Inpp5d
Inppl1
Lgals9
Ptk2b
Ptpn6
Sos2

**MAPK signaling**
Angpt1

Atf4
Cacnb2
Cacnb4
Cd14
Cdc42
Csf1
Csf1r
Dusp1
Dusp9
Efna1
Egf
Egfr
Fgf10
Fgf15
Fgf18
Fgf2
Fgf22
Gadd45a
Gng12
Grb2
Hgf
Hras
Ins1
Irak4
Jund
Kras
Lamtor3
Map2k2
Map3k13
Map3k8
Mapk1
Mapk10
Mapk11
Mapk13
Mapk14
Mapk3
Mapk8
Mapk9
Mras
Nfatc1
Nfatc3
Nr4a1
Nras
Ntf3
Pdgfd
Pdgfra
Pdgfrb

Ppm1a
Ppp3ca
Ppp3cc
Ppp3r1
Prkcb
Ptpn7
Rac1
Rasgrf1
Rasgrp1
Rasgrp3
Rras
Sos1
Sos2
Tab2
Tnfrsf1a
Traf2
Vegfa

**epigenetic modification**
Atm
Bcl6
Camk2d
Ccnb1
Cdyl
Chek1
Cpa4
Crebbp
Dnmt3a
Dnmt3l
Dydc1
Eya1
Eya4
Fbll1
Foxp3
Gata3
Gnas
Gsk3b
Hdac7
Hdac9
Hnf1a
Irf4
Jak2
Jarid2
Kdm3a
Lef1
Lif
Lrrk2

Mbd3
Mysm1
Naa50
Pml
Ppargc1a
Ppm1f
Prdm16
Prkcb
Rag1
Sf3b1
Smad4
Snai2
Snw1
Taf6l
Tbl1xr1
Tdrd1
Tgfb1
Twist1
Uhrf1
Usp15
Usp16
Yy1

**TGFb signaling**
Amh
Bmp2
Bmp8b
Cbl
Ccnc
Ccnt1
Ccnt2
Cer1
Chrd
Chrdl1
Crebbp
Cul1
Dcn
Drap1
E2f5
Gdf6
Grem2
Id2
Ifng
Junb
Lefty1
Ltbp1
Pitx2

Pmepa1
Ppm1a
Ppp1cb
Prkcz
Rbx1
Rock1
Smad2
Smad3
Smad4
Smad9
Snw1
Tgfb1
Tgfb2
Trim33
Ube2d1
Usp15

**TLR signaling**
Apob
Bpi
Btk
Btrc
Casp8
Ccl3
Ccl4
Ccl5
Cd14
Cd180
Cd40
Cd80
Cd86
Cul1
Cxcl11
Cxcl9
Fadd
Fga
Hmgb1
Ifna1
Il12a
Il12b
Irak3
Irak4
Plcg2
Ptpn11
Rac1
Sftpa1
Socs1

Stat1
Tab2
Tbk1
Ticam1
Tlr1
Tlr2
Tlr7
Ube2d1

**hypoxia response**
Ak4
Aldoa
Aldob
Angpt1
Aqp1
Atp7a
Camk2d
Cflar
Clca1
Crebbp
Cul2
Dnmt3a
E2f1
Egf
Egfr
Fabp1
Foxo3
Gata6
Icam1
Ifng
Il6ra
Ins1
Kcnk2
Ldha
Ldhal6b
Nfe2l2
Nos2
Npepps
Pck1
Pdhb
Plcg1
Plcg2
Ptgs2
Rbx1
Rora
Scn2a
Sfrp1

Slc8a1
Stat3
Stc1
Stc2
Tert
Timp1
Trpc6
Twist1
Ube2d1
Vegfa

**CTLA4 signaling**
Cd80
Cd86
Ctla4
Fyn
Lck
Lyn
Ppp2ca
Ppp2cb
Ppp2r1a
Ppp2r1b
Ppp2r5a
Ppp2r5b
Ppp2r5c
Ppp2r5e
Ptpn11
Yes1

**T cell exhaustion**
Ahr
Batf
Casp3
Ccl3
Cd160
Cd244a
Cd86
Ctla4
Dgka
Dgkz
Eomes
Fasl
Havcr2
Irf4
Klrg1
Lag3
Nfil3

Pbx3
Pdcd1
Pirb
Plscr1
Ptger4
Rgs16
Tigit
Tnfrsf9
Tnfsf9
Tox2
Tox3
Tox4
Yy1

**JAK/STAT signaling**
Cish
Cntf
Crebbp
Crlf2
Csf2
Csf3
Csf3r
Ctf1
Gh
Ghr
Grb2
Ifna1
Ifnk
Jak1
Jak2
Jak3
Lepr
Lif
Lifr
Mcl1
Osm
Osmr
Pdgfra
Pdgfrb
Pias1
Pias2
Prl
Prlr
Ptpn11
Ptpn2
Ptpn6
Socs1

Socs2
Socs3
Sos1
Sos2
Stat1
Stat3
Stat4
Stat5a/b
Tslp
Tyk2

**Immune checkpoints**
Adora2a
Btla
Cd200r1
Cd27
Cd274
Cd276
Cd28
Cd3d
Cd3e
Cd3g
Cd4
Cd40
Cd40lg
Cd70
Cd80
Cd86
Cdh1
Ctla4
Fyn
H2-Aa
H2-Ab1
H2-D1
H2-DMa
H2-DMb1
H2-DMb2
H2-Eb1
H2-K1
H2-M3
H2-Ob
H2-Pa
H2-Q1
H2-Q10
H2-Q2
H2-T23
Havcr2

Icos
Icosl
Lag3
Lck
Lyn
Pdcd1
Pdcd1lg2
Pecam1
Ppp2ca
Ppp2cb
Ppp2r1a
Ppp2r1b
Ppp2r5a
Ppp2r5b
Ppp2r5c
Ppp2r5e
Ptpn11
Ptpn6
Pvrig
Tigit
Tnfrsf14
Tnfrsf18
Tnfrsf4
Tnfrsf9
Tnfsf18
Tnfsf4
Tnfsf9
Trac
Vsir
Vtcn1
Yes1

**Senescence & Quiescence**
Anapc4
Atm
Atr
Btrc
Calm1
Ccna1
Ccnb1
Ccnb3
Cdc16
Cdc25a
Cdkn2c
Cebpb
Chek1
E2f1

E2f2
E2f5
Foxo1
Foxo3
Gadd45a
Hipk1
Il1a
Itpr1
Itpr2
Klf2
Kras
Mapk10
Mapk11
Mapk13
Mapk14
Mapk9
Mras
Nfatc1
Nfatc2
Nfatc3
Nfatc4
Phc3
Ppp1cb
Ppp3ca
Ppp3cc
Ppp3r1
Rbl2
Rheb
Rras
Sesn1
Sesn2
Sesn3
Smad2
Smad3
Tert
Tgfb1
Tgfb2
Trpm7
Ube2d1
Vdac1
Zfp36l2

**PI3K-AKT**
Akt1
Akt2
Akt3
Atf4

Bcl2
Bcl2l1
Casp9
Creb3l2
Creb5
Efna1
Eif4b
Foxo3
Gh
Ghr
Gng12
Grb2
Gsk3b
Gys2
Hgf
Ifna1
Irs1
Itga10
Itga6
Itgav
Itgb3
Itgb8
Lpar1
Magi2
Mcl1
Nr4a1
Ntf3
Pck1
Pdgfd
Pdgfra
Pdgfrb
Pik3ap1
Pik3ca
Pik3cb
Pik3cd
Pik3r1
Pik3r2
Pik3r3
Pkn2
Ppp2ca
Ppp2cb
Ppp2r1a
Ppp2r1b
Ppp2r5a
Ppp2r5b
Ppp2r5c
Ppp2r5e

Prkaa1
Prkaa2
Prl
Prlr
Rac1
Rbl2
Reln
Rheb
Sos2
Syk
Tcl1
Tlr2
Vegfa
Ywhaz

**PD1 signaling**
Cd274
Cd3d
Cd3e
Cd3g
Cd4
H2-Aa
H2-Ab1
H2-D1
H2-DMa
H2-DMb1
H2-DMb2
H2-Eb1
H2-K1
H2-M3
H2-Ob
H2-Pa
H2-Q1
H2-Q10
H2-Q2
H2-T23
Lck
Pdcd1
Pdcd1lg2
Ptpn11
Ptpn6
Trac

**NK-kB signaling**
Atm
Btk
Card11

Ccl2
Ccl21a/b/c
Ccl4
Cd14
Cd40
Cd40lg
Cflar
Chuk
Cxcl1
Cxcl2
Cxcl3
Cyld
Gadd45a
Icam1
Il1r1
Irak4
Lck
Lyn
Malt1
Nfkb1
Nfkb2
Nfkbia
Plcg1
Plcg2
Prkcb
Ptgs2
Rela
Syk
Tab2
Ticam1
Tnfaip3
Tnfrsf1a
Traf2
Ube2i
Vcam1
Xiap
Zap70

**Type I interferon**
Bst2
Gbp2
Ifit1
Ifit3/3b
Irf2
Irf4
Irf8
Mx1

Mx2
Oas1a
Oas2
Oas3
Ptpn11
Ptpn6
Rsad2
Socs1
Socs3
Tyk2
Xaf1

**IL10 signaling**
Ccl12
Ccl20
Ccl22
Ccl3
Ccl4
Ccl5
Ccr1
Ccr2
Ccr5
Cd80
Cd86
Csf1
Csf2
Csf3
Cxcl2
Fcer2a
Fpr1
Icam1
Il10
Il10ra
Il10rb
Il12a
Il12b
Il18
Il1a
Il1r1
Il1r2
Il1rn
Lif
Ptafr
Ptgs2
Timp1
Tnf
Tnfrsf1a

Tnfrsf1b
Tyk2

**TNF signaling**
Atf4
Casp3
Casp8
Ccl12
Ccl20
Ccl5
Cebpb
Cflar
Creb3l2
Creb5
Csf1
Csf2
Cxcl1
Cxcl2
Cxcl3
Cyld
Dnm1l
Fadd
Fas
Icam1
Il15
Itch
Jag1
Junb
Lif
Mmp14
Mmp3
Mmp9
Ptgs2
Sele
Socs3
Tab2
Tax1bp1
Tnf
Tnfaip3
Tnfrsf1a
Tnfrsf1b
Traf2
Vcam1
Xiap
